# MoBPS - Modular Breeding Program Simulator

**DOI:** 10.1101/829333

**Authors:** T. Pook, M. Schlather, H. Simianer

## Abstract

The R-package MoBPS provides a computationally efficient and flexible framework to simulate complex breeding programs and compare their economic and genetic impact. Simulations are performed on the base of individuals and haplotypes are calculated on-the-fly by only saving founder haplotypes, points of recombination and mutations. MoBPS utilizes a highly efficient implementation with bit-wise storage of data and matrix multiplications from the associated R-package miraculix allowing to handle large scale populations. The modular structure of MoBPS allows to combine rather coarse simulations, as needed to generate founder populations, with a very detailed modeling of todays’ complex breeding programs, making use of all available biotechnologies. MoBPS provides pre-implemented functions for common breeding practices such as optimum genetic contributions and single-step GBLUP but also allows the user to replace certain steps with personalized and/or self-written solutions.

## INTRODUCTION

Breeding programs aim at improving the genetic properties of livestock and crop populations w.r.t. productivity, fitness and adaptation. Progress towards the target is limited by the available resources, but also negative effects, such as inbreeding depression or health issues, have to be avoided or at least controlled. Hence, the allocation of resources in a breeding program is a complex optimization problem. Additionally, population history, such as fluctuating population sizes and selection pressures, has an impact on the current genomic architecture and thus the potential for future improvement.

Over the years a variety of simulation tools have been developed to assist breeders to evaluate and optimize their breeding programs. A general problem of simulation studies is that the underlying genomic processes are highly complex and have to be simplified for modeling. In addition, users often have rather different objectives in mind when setting up their simulation studies. Since tools often do not provide the necessary flexibility to execute the specific breeding actions and/or it is not possible to export all necessary outputs, this commonly leads to the use of self-developed solutions that tend to be more error-prone, less sophisticated and computationally inefficient. The functionality of existing software ranges from cohort based deterministic simulation that relies on expected gains like ZPLAN+ (Täubert *et al.* 2010) to applications on the base of the stochastic simulation of single individuals such as QMSim (Sargolzaei and Schenkel 2009) and AlphaSim (Faux *et al.* 2016). The functionality of each of these tools highly depends on the intended use. ZPLAN+ (Täubert *et al.* 2010) focuses on the economic impact from a macro-perspective. Since analytic formulas for cohorts are required, it has limitations when simulating complex mating schemes or when focusing on other quantities than genetic or economic gain. QMSim (Sargolzaei and Schenkel 2009) is able to simulate each individual meiosis but is limited in the options for the design of the breeding program itself. As QMSim is mostly designed for population genetic studies, a typical application of the tool is the generation of a historical population, often followed by self-developed solution in later steps. On the contrary, AlphaSim (Faux *et al.* 2016) provides a lot of flexibility in term of the design of the breeding program, especially for plant breeding and when the number of cohorts in the breeding program is small. However, AlphaSim lacks the efficiency and flexibility to simulate complex and large scale populations.

Our goal was to develop a tool that combines the simulation of a historical population and the evaluation of a subsequent complex breeding program in a computationally efficient way. The Modular Breeding Program Simulator (MoBPS) is not only flexible in terms of parameters and design of breeding programs, but also allows the user to replace standard procedures of the package with own ones.

## METHODS

Simulations in MoBPS are ultimately based on the simulation of single individuals. In principle, this allows the user to control each singular mating and modify recombination or mutation rates for the respective meiosis. Breeding programs are constructed in a modular form as a combination of cohorts (Hill 1974), representing a group of contemporary individuals with similar characteristics, and transformations, which link one or several parent cohorts to a child cohort. Examples for such transformations are aging, selection, or reproduction, and each transformation reflects a set of rules how the characteristics of the parent cohort(s) are transformed into the characteristics of the child cohort. Cohorts and transformations are defined in a generic way and are parametrized, so that any breeding program of arbitrary complexity can be modeled as a suitable sequence of cohorts and transformations.

All data for a population is stored in a list that contains general and individual information. The general part provides information on the underlying genetics like the physical position of each marker, allelic variants or structure of the underlying genetic traits. The individual part contains information that is specific to the individual. Haplotypes are stored for founder individuals only. For all other individuals only points of recombination and mutation and their genetic origins are stored and haplotypes are derived on-the-fly. Therefore, the required memory is minimized and only increases slightly with increasing marker density. When thousands of generations are simulated it is advisable to classify additional generations as new founders to reduce the number of recombinations and mutations to be stored in subsequent generations.

Simulation of multiple correlated traits with and without underlying QTL is supported. Classical additive, dominant and epistatic or pleiotropic QTL can be defined and any effect structure of multiple interacting loci is supported. Each locus has to be assigned with a position in Morgan and different recombination rates for subgroups (e.g. males/females) are supported. Information on the number of markers can be manually entered or imported via a database (Ensemble, (Zerbino *et al.* 2017)), a map-file (Purcell *et al.* 2007) or a vcf-file (Danecek *et al.* 2011). For common species, exemplary map files are provided in the associated package MoBPSmaps (Pook 2019). Genotype data for a base population can be imported via PLINK (Purcell *et al.* 2007) and/or vcf-format (Danecek *et al.* 2011), sampled internally or generated by executing prior simulation in MoBPS and/or other tools (Chen *et al.* 2009; Sargolzaei and Schenkel 2009) to generate the required population structure. All breeding actions performed in the simulation can be tracked and assigned with costs to derive the expenses of the program. Different breeding programs can be compared in terms of their economic revenue or other target functions (e.g. development of the inbreeding rate) one is interested in.

Common methods for selection such as optimal genetic contributions (Meuwissen 1997) are implemented and a variety of different packages for breeding value estimation can be switched on. This includes BGLR (Pérez and de los Campos 2014), sommer (Covarrubias-Pazaran 2016) and rrBLUP (Endelman 2011), as well as an efficient implementation for solving the mixed model (Henderson 1975) in the traditional GBLUP model (Meuwissen *et al.* 2001; VanRaden 2008) that is assuming known heritability and is using the R-package RandomFieldsUtils (Schlather *et al.* 2019b) for the matrix inversion. Inputs for these packages such as the different pedigree and genomic relationship matrices (VanRaden 2008; Legarra *et al.* 2014; Martini *et al.* 2017) can be derived via highly efficient and fully-parallelized bit-wise matrix multiplications (R-package miraculix (Schlather *et al.* 2019a)). Non of the mentioned packages, however, is required to execute simulations in MoBPS. In particular all functionality of the MoBPS R-package is still available when miraculix is not installed, with the downside of higher computing times and memory demands.

The simulations in MoBPS are based on two main functions: *creating.diploid()* and *breeding.diploid()*. Here, *creating.diploid()* initializes the base-line population and *breeding.diploid()* performs breeding actions on an existing population list. As a simple example consider the following script:

~~~
library (MoBPS)
pop <- creating . diploid (nsnp =10000, nindi =50,
    chr. nr=5, chromosome . length =2,
    n. additive =25, n. dominant =5,
    name. cohort=“Founder”)
pop <- breeding . diploid (pop, heritability =0.5,
    new . bv. observation =“all”)
pop <- breeding . diploid (pop, bve= TRUE)
pop <- breeding . diploid (pop, breeding . size =50,
    selection . size= c(5,10),
    selection . m=“function “,
    selection . m. cohorts=“Founder_ M”,
    selection . f. cohorts=“Founder_ F”,
    name. cohort=“Offspring”)
~~~

Via this code, we first generate a base population containing 50 individuals with 10,000 markers. The underlying genome consists of 5 chromosomes with a length of 2 Morgan each and equidistant markers. Furthermore, we generated a single trait that is impacted by 25 purely additive QTLs and 5 dominant QTLs.

In the next step, we initialize a breeding action to generate phenotypes for all individuals in the population with an assumed heritability of 0.5. Next, a breeding value estimation is performed. Since no cohorts are selected, the last (and only) generation of the population list will be considered for the breeding value estimation. Lastly, we generate 50 offspring by randomly mating the top 5 male and top 10 female individuals. In principle, all three breeding actions performed via *breeding.diploid()* could have also been executed in a joint step. For a full list of all possible breeding actions and available parameters we refer to our user manual (available at https://github.com/tpook92/MoBPS).

For a quick overview of the simulated population, the function *summary()* can be used:

~~~
> summary (pop)
Population size:
Total: 100 Individuals
Of which 50 are male and 50 are female.
There are 2 generations
and 4 unique cohorts.

Genome Info :
There are 5 unique chromosomes.
In total there are 10000 SNPs.
The genome has a total length of 10 Morgan .
No physical positions are stored .

Trait Info :
There is 1 modelled trait.
The trait has underlying QTL
The trait is named : Trait 1
~~~

A variety of functions is provided to export required information such as the phenotypes (*get.pheno()*), the genotypes (*get.geno()*) and the pedigree (*get.pedigree()*) for selected individuals from the population list. These functions are thoroughly described in chapter 9 of the user manual (available at https://github.com/tpook92/MoBPS). Furthermore, functions to derive rates of inbreeding (*kinship.emp()*), development of breeding values (*bv.development()*) or changes in allele frequency over time (*analyze.population()*) are provided to further analyze the resulting population list.

## RESULTS AND DISCUSSION

The package MoBPS is completely written in R (R Core Team 2017) so that all functionalities for genetic applications are platform independent. The R-packages miraculix (Schlather *et al.* 2019a) can be activated in MoBPS and leads to more efficient data storage and shorter simulation times. In particular vector multiplications with genetics data (0,1,2) are performed via bitwise operations on a whole register (128/256 bit) using SSE2/AVX2. Computing times are similar to the ones in PLINK (Purcell *et al.* 2007) with one fourth of the memory usage.

Even though basically all information regarding each individual is stored, the required memory in MoBPS is still relatively low as a highly efficient storage structure is used. Haplotypes of founders and details on the origin of the segment between points of recombination are stored bitwise. E.g. the simulation of 20 generations with 50,000 cows with 50,000 markers and breeding value estimation via GBLUP takes 26.2 hours using 24 cores on a server cluster with Intel E5-2650 (2X12 core 2.2GHz) processors. At peak, 65 GB of memory was used. The main share of this was required for the storage of the genomic relationship matrix whereas the resulting population list, containing more than a million individuals, only had a size of about 0.44 GB. The biggest proportion of the computing time is used for breeding value estimation (25.3 hours, 96.4%). The generation of new animals took 55 minutes (3.5%, 304 animals per second using a single core). All other parts needed negligible computing time (132 seconds, 0.1%). Computing times for most parts (except breeding value estimation) increase linearly with the number of individuals. This highly efficient storage structure therefore also allows for the simulation of historical populations with thousands of generations and undergone population dynamics such as genetic bottlenecks, migration or mutational drift.

The flexible and efficient enviroment of MoBPS allows for the simulation of a variety of different and potential large-scale breeding programs. For exemplary scripts of more complex breeding programs we refer to the user manual. Exemplary simulations are given for the effect of gene editing in a cattle breeding program (Simianer *et al.* 2018), the simulation of a multi-parent advanced generation intercross in maize (Pook *et al.* 2019), an introgression scheme in chicken (Ha *et al.* 2017) and the generation of a base population with a hard sweep. A further advantage of MoBPS in comparison to other simulation tools is its flexible structure that allows the user to substitute single steps of the breeding program with a personalized and/or self-written solution. For this consider the following example to execute one owns breeding value estimation:

~~~
genos <- get. geno (pop, gen =1)
y <- get. pheno (pop, gen =1)
indi_names <- colnames(genos)
*# Execute one owns function to perform*
*# the breeding value estimation*
y_ hat <- own . method . for. bve(genos, y)

*# Enter BVEs in the population - list*
pop <- insert. bve(pop,
    bves = cbind (indi_ names, y_ hat))
~~~

Even though a simulation study can never fully reflect reality and is relying on model assumptions, the use of a simulation study comes with major benefits and still allows the user to draw important conclusions. In contrast to reality the underlying truth in a simulation study is known, and therefore new methods can be thoroughly evaluated and compared to existing ones. Furthermore, the effects of particular breeding actions on a variety of output dimension can be assessed and compared. This in turn can be used to derive an ideal resource allocation and optimize potentially highly complex breeding scenarios in a setting that can be evaluated multiple times and without constrains both in terms of money and time.

## WEB RESOURCES

An executable version of MoBPS and the associated R-packages miraculix (Schlather *et al.* 2019a), RandomFieldsUtils (Schlather *et al.* 2019b) and MoBPSmaps (Pook 2019) for Windows and Linux are freely available at https://github.com/tpook92/MoBPS. This directory also contains an comprehensive user manual explaining the functionality of all input parameters and utility functions in MoBPS. A frozen version of the R-packages MoBPS (v1.4.15), miraculix (v0.9.7), RandomFieldsUtils (v0.5.9), and our user manual at submission can be found in the Supplementary material. The MoBPS R-package can be directly installed within your R session via following commands:

~~~
install. packages(“devtools”)
devtools :: install_ github (“tpook92 / MoBPS”,
    subdir=“pkg”)
~~~

## Supporting information

User manual

MoBPS R-package

miraculix R-package

RandomFieldsUtils R-package

## ACKNOWLEDGMENTS

This package was developed in the context of the European Union’s Horizon 2020 Research and Innovation Program under grant agreement n°677353 IMAGE.

