## Supplementary material for "MoBPS - Modular Breeding Program Simulator": User manual

### MOBPS

Torsten Pook, Martin Schlather, Henner Simianer

OCTOBER 29, 2019

UNIVERSITY OF GOETTINGEN
Animal Breeding and Genetics Group

### Table of Contents

*1 General* 4

*2 Installation* 4

*3 Citation* 4

*4 Creation of the starting population (creating.diploid())* 5

**4.1 Importing/Generating of a genetic dataset** 5

**4.2 Importing a genetic map** 6

**4.3 Simulating/Generating the genetic architecture underlying each trait** 6

4.3.1 Custom-made genetic architectures 7
4.3.2 Predefined genetic architectures 8
4.3.3 Correlated Traits 8

**4.4 Position of Markers** 8

*5 Simulation of breeding processes (breeding.diploid())* 9

**5.1 General setup** 9

**5.2 Control of heritability, breeding values, genotypes and phenotypes** 10

**5.3 Breeding value estimation** 11

5.3.1 Direct approach with known heritability 12
5.3.2 Bayesian approaches (BGLR) 12
5.3.3 GBLUP (EMMREML) 13
5.3.4 GBLUP (sommer) 13
5.3.5 GBLUP (rrBLUP) 13
5.3.6 Pedigree-based (breedR) 13
5.3.7 Parent/Grandparent mean 13
5.3.8 Own function 13
5.3.9 Calculating marker effects & GWAS 14
5.3.10 Calculation of reliabilities 14

**5.4 Selection techniques & mating strategies** 14

5.4.1 Multiple traits 15
5.4.2 Higher procreation of genetically favored individuals 15
5.4.3 Maximum number of offspring per individual 16
5.4.4 Plant breeding (no-sexes & selfing & DH-production & cloning) 16
5.4.5 Generate offspring for all sire combination 16
5.4.6 Targeted/Fixed mating/Manual selection of individuals 16
5.4.7 Gene-Editing 16
5.4.8 Optimum genetic contribution 17

**5.5 Genetic architecture** 17

**5.6 Other** 17

5.6.1 Culling / Death 17
5.6.2 Parameter for target-mating with J.W.R. Martini 18
5.6.3 Allele-frequency per generation 18
5.6.4 Set a random seed 18

5.6.5 save.recombination.history 18

**5.7 Storage & computing time 18**

5.7.1 Reducing the size of the population list 19

5.7.2 Reducing memory needs in the BVE 19

5.7.3 Inverting G using miraculix 19

5.7.4 On-the-fly calculation of haplotypes 19

***6 Exporting information from the population-list (get.XXXX) 20***

**6.1 get.genos 20**

**6.2 get.haplos 20**

**6.3 get.bv / get.bve / get.pheno / get.reliability / get.selectionindex 21**

**6.4 get.recombi 21**

**6.5 get.pedigree (1/2/3) 21**

**6.6 get.cohorts 22**

**6.7 get.class 22**

**6.8 get.genotyped 23**

**6.9 get.time.point 23**

**6.10 get.creating.type 23**

**6.11 get.cullingtime 24**

**6.12 get.individual.loc 24**

**6.13 get.vcf 24**

**6.14 get.pedmap 24**

**6.15 get.database 25**

***7 Importing information to the population-list 25***

**7.1 Insert.bve 25**

***8 Data structure of the population list 26***

**8.1 $info 26**

**8.2 $breeding 28**

8.2.1 Storage per generation 28

8.2.2 Storage per individual 28

***9 Utility functions 29***

**9.1 bv.development 29**

**9.2 bv.development.box 30**

**9.3 Kinship.development 31**

**9.4 Kinship.emp / kinship.emp.fast 31**

**9.5 Kinship.exp 32**

9.6 analyze.population 32

9.7 new.base.generation 33

9.8 creating.trait 33

9.9 ensembl.map 33

9.10 compute.costs 34

9.11 compute.costs.cohorts 34

9.12 summary 35

9.13 pedmap.to.phasedbeaglevcf 35

*10 Memory and computation times 36*

*11 List of input parameters in breeding.diploid() 36*

*12 List of input parameters in creating.diploid() 43*

*13 List of datasets included in the package 46*

*14 User-interface 47*

*15 Commonly used word definitions 54*

*16 Exemplary scripts 54*

16.1 Simulation of a MAGIC population in maize 54

16.2 Simulation of Introgression on blue eggshell QTL 55

16.3 Simulation of gene editing in a cow breeding program 58

16.4 Simulation of a base population with a hard sweep 59

*References 60*

*17 Acknowledgements 62*

### 1 General

MoBPS is an R-package to simulate complex and large scale breeding programs with focus on livestock and crop populations. Simulations are performed on an individual basis. MoBPS is a versatile tool, providing standard procedures applied in animal and plant breeding like GBLUP and OGC, but also allowing to use own selection schemes while still controlling the simulation of phenotypes, meiosis and costs of the simulated scheme. The actual process of the simulation can be split up into two steps: the creation of a starting population and the simulation of breeding processes.

As it is our goal to provide a lot of flexibility while performing the simulation, there is a need of many parameters – luckily only a few of those will be needed for most simulations. For a better understanding of the workflow required to set up a simulation, we refer to section 16 for exemplary simulations. For a list of all input parameters and possible initializations, we refer to section 11.

For questions regarding the tool or how to set up your simulation feel free to contact me. We are always happy for questions as it really helps improve the tool and its documentation. For quick reply, it would help to provide a small example of our problem.

In addition, we are currently in the process of developing a graphical interface for the R-package (available at www.mobps.de). Note that this interface is still in active development and not part of the MoBPS paper. The use of the interface is still encouraged, but for major projects we highly recommend (and are looking for) close collaboration for the further development.

### 2 Installation

The current version of MoBPS requires R 3.0 or higher. We highly recommend the use of the packages RandomFieldsUtils (version 0.5.9+) and miraculix (0.9.7+) as they significantly reduce computing time when working with a high number of markers and individuals. All functionality is available without both package, but simulations can be significantly slower. For direct install via GitHub use the following line of code (this requires the R-package devtools):

```
devtools::install_github("tpook92/MoBPS", subdir="pkg")
```

This option is currently not available for miraculix and RandomFieldsUtils. RandomFieldsUtils is available on CRAN, miraculix will hopefully soon follow. As updates to CRAN are only possible every few month we highly recommend to use the version available on GitHub itself. Packages can be downloaded at https://github.com/tpook92/MoBPS and directly install via the R function *install.packages().* Usage was tested on Linux and Windows (under windows set type="source", repo=NULL). The usage on Mac OS is currently not recommended. Commonly used Ensembl-maps (Zerbino et al. 2017) are available in the associated R-package MoBPSmaps.

```
devtools::install_github("tpook92/MoBPS", subdir="pkg-maps")
```

For Windows the installation of Rtools is required. RandomFieldsUtils does require the package spam.

### 3 Citation

There is currently no paper published on our R-package. This will hopefully soon change. For so long we suggest using following to citations for the R-packages MoBPS and miraculix:

```
@Manual{,
```

```
    title = {MoBPS: Modular Breeding Program Simulator},
    author = {Torsten Pook},
    year = {2019},
    note = { Available at https://github.com/tpook92/MoBPS; R-package versi
on 1.4.15},
  }

  @Manual{,
    title = {miraculix: Statistical Functions for Animal Breeding},
    author = {Malena Erbe and Martin Schlather and Florian Skene and Alexan
der Freudenberg},
    year = {2019},
    note = { Available at https://github.com/tpook92/MoBPS; R-package versi
on 0.9.6},
  }
```

### 4 Creation of the starting population (*creating.diploid()*)

The input for the simulation of a breeding process is a population list. This list is created via *creating.diploid()*.

We provide exemplary genetic maps for some common species, which can be selected via the parameter **template.chip**. Note that primary the number of chromosomes and their genetic length is imported at this step (especially not real markers with known allele frequencies, effects or base pairs). The maps provided via **template.chip** are “cattle” (Ma et al. 2015), “pig” (Rohrer et al. 1994), “chicken” (Groenen et al. 2009), “sheep” (Prieur et al. 2017) and “maize” (Lee et al. 2002). Alternatively, a genomic map can be inserted via the parameter **map** with exemplary maps being provide in the package itself or via import from Ensembl (section 9.9).

#### 4.1 Importing/Generating of a genetic dataset

In case one has haplotype data for the founders/starting population this can be imported via the parameter **dataset** in form of a haplotype dataset:

```
> dataset
      Indi1Haplo1 Indi1Haplo2 Indi2Haplo1 Indi2Haplo2 Indi3Haplo1 Indi3Haplo2 Indi4Haplo1 Indi4Haplo2
SNP1            1           1           1           1           0           1           1           0
SNP2            0           1           1           0           1           0           1           1
SNP3            0           0           0           1           1           0           0           1
SNP4            1           1           1           1           0           0           1           1
SNP5            0           1           0           1           0           1           1           1
SNP6            0           1           0           0           0           1           1           1
SNP7            1           0           1           1           1           1           1           0
SNP8            0           0           0           1           0           1           0           1
SNP9            1           1           0           0           0           0           0           1
SNP10           0           1           0           0           1           1           1           0
```

Datasets can also be imported via entering the path of the vcf file in the parameter **vcfpath**. The R-package vcfR is needed for this. Otherwise, a dataset can be generated by setting the number of SNPs (**nsnp**) and individuals (**nindi**) – we offer four possible modes to simulate starting haplotypes ("all0", "allhetero", "random", "homorandom") leading to haplotypes (000…/000…, 000…/111…,X_1 X_2 X_3… /X_4 X_5 X_6 … with X_i~B(**freq**) X_1 X_2 X_3 … /X_1 X_2 X_3… with X_i~B(**freq**)). On default "random" is used. The allele frequencies for each marker are enter via the parameter **freq** and are sampled from a beta distribution with **beta_shape1** and **beta_shape2** (default: 1,1; leading to a uniform distribution of allele frequencies.

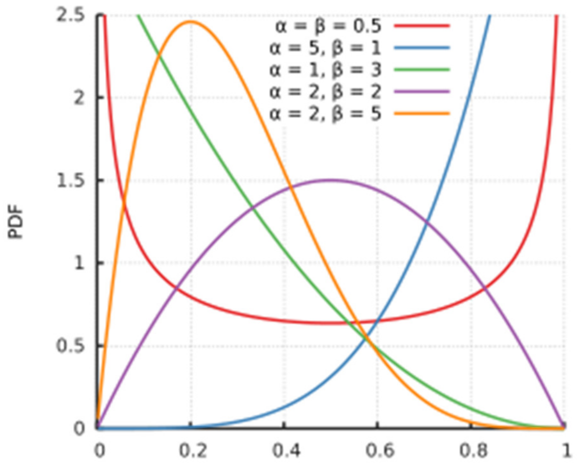

*Figure 1: https://en.wikipedia.org/wiki/Beta_distribution*

To generate an LD and haplotype structure without using real data we recommend to start with one of the simple datasets and simulate some random/non-random mating generations using *breeding.diploid()* (cf. section 5). Wrapper functions for the automatic generation of those base populations in MoBPS are planned but not yet implemented.

If a vcf file is used for data import or a map is provided, the chromosome, marker name and base pair position are automatically imported. Alternatively, those can be provided via **chr.nr**, **bp** and **snp.name**. By doing this, multiple chromosomes can be inputted jointly. If no input for **chr.nr** is provided all markers are assumed to be on the same chromosome (for more on the usage of a genetic map, we refer to section 4.2).

In case more markers are to be added to and existing population list set **add.chromosome** to TRUE and repeat the previous process with the population list as an additional input.

To specify the sex of each sample either assign each individual a probability to be female (**sex.quota**) or alternatively use a vector (**sex.s**) assigning each individual its sex (M=1, F=2).

#### 4.2 Importing a genetic map

The user can provide a genetic map of up to five colums via the parameter **map**. The first column contains the chromosome of the respective marker, the second column the name of the marker, the third column the physical position of the marker, the forth column the position in centimorgan and the last column the allele frequency in the population. All values not provided are automatically set to NA and values are used as input for **chr.nr**, **snp.name**, **bp**, **snp.position**, **freq**.

Alternatively maps can be imported via *ensembl.map()*. For more on this we refer to Chapter 9.9.

#### 4.3 Simulating/Generating the genetic architecture underlying each trait

As the manual input of effects can be tiring, we provide some automated procedures to simulate some common effect structures (additive, dominant, qualitative and quantitative epistasis) – if you do not need more, you can just skip to section 4.3.2).

Note that this is the generation of an actual genetic value that is underlying each individual of the population. In reality you cannot observe this, as traits will be caused by far more complex interactions and effects are not known. This, on the other hand, enables opportunities to evaluate a model fit given a known structure (e.g. GWAS hits can be compared to actual effect markers instead of previously identified markers or similar). Traits can be named via the parameter **trait.name**.

##### 4.3.1 Custom-made genetic architectures

To simulate a custom-made genetic architecture we allow for effects caused by one (**real.bv.add**), two (**real.bv.mult**) or more SNPs (**real.bv.dice**). These effects can either be added directly while using *creating.diploid()* or added later using *creating.trait()*. To delete previously existing effects set **replace.real.bv** to TRUE. For multiple traits use lists as inputs for all parameters in this section with each list element containing information for one trait.

Input structure for the first two is a matrix with each row coding a single effect:

```
> real.bv.add
     SNP chromosome Effect 0 effect 1 effect 2
[1,] 120          1     -1.0      0.0        1
[2,]  42          5      0.0      0.0        2
[3,]  17         22      0.1      0.1        0
```

**real.bv.add** should be able to model any additive or dominance effects of single markers.

```
> real.bv.mult
     First SNP First chromosome Second SNP Second chromosome effect 00 effect 01 effect 02 effect 10 effect 11 effect 12 effect 20 effect 21 effect 22
[1,]       144                1        145                 1      1.00      0.00      0.00      0.00      0.00      0.00      0.00      0.00      0.00
[2,]         6                3        188                 5      0.37      0.16      1.33      1.49      1.58      2.51      0.38      2.12      0.98
[3,]         5               17          1                10      1.18      2.60      0.18      1.74      0.69      1.39     -1.21      0.96      1.94
```

**real.bv.mult** should be able to model any epistatic interaction between two markers.

To simulate even more complex effect structures use the parameter **real.bv.dice** allowing the modelling of effects caused by more than two SNPs.

Input for **real.bv.dice** is a list containing a list of all locations and a list of all effects:

```
> real.bv.dice
$location
$location[[1]]
     SNP chromosome
[1,]  11          1
[2,]  12          1
[3,]  16          4

$location[[2]]
     SNP chromosome
[1,]  14          2
[2,]  77          6
[3,]  15          9

$effects
$effects[[1]]
 [1]  1.8212212  1.5939013  1.9189774  1.7821363  1.0745650 -0.9893517  1.6198257  0.9438713  0.8442045 -0.4707524  0.5218499  1.4179416  2.3586796
[14]  0.8972123  1.3876716  0.9461950 -0.3770596  0.5850054  0.6057100  0.9406866  2.1000254  1.7631757  0.8354764  0.7466383  1.6969634  1.5566632
[27]  0.3112443

$effects[[2]]
 [1]  0.29250484  1.36458196  1.76853292  0.88765379  1.88110773  1.39810588  0.38797361  1.34111969 -0.12936310  2.43302370  2.98039990  0.63277852
[13] -0.04413463  1.56971963  0.86494540  3.40161776  0.96076000  1.68973936  1.02800216  0.25672679  1.18879230 -0.80495863  2.46555486  1.15325334
[25]  3.17261167  1.47550953  0.29005357
```

Each network of interacting markers is giving in the first list (location) and their effects are given in a second list (effects). Effects are sorted in following order: 0…0, 0…01, … 2…2 – resulting in a vector with $3^n$ elements, where n is the number of markers involved in the effect.

Marker effects assign to positions that currently do not exist (e.g. SNP 100 on chromosome 1 in case chromosome 1 only contains 50 SNPs) are automatically removed from the stored effects unless **remove.invalid.qtl** is set to FALSE.

##### 4.3.2 Predefined genetic architectures

In case of a predefined genetic architecture, all markers are assigned with the same probability to be drawn as effect markers. To exclude markers use a parameter **exclude.snps** containing a vector of all numeric positions of excluded markers. Numbering is consecutively starting with chromosome 1. Note that only markers, that are already included in the dataset, can be chosen as effect markers – so in case of more than one chromosome with no generation via **chr.nr** the effects should be added using *creating.trait()* (cf. section 9.2) or in the last run of *creating.diploid()*.

The number of additive (**n.additive**) and dominate (**n.dominant**) QTL as well as effects caused by qualitative (**n.qualitative**) and quantitative (**n.quantitative**) epistasis can be included directly. To assign the variance, one can input a vector containing the variance for each effect (**var.additive, var.dominant,…).** On default, each variance is set to 1 and effects are drawn from a Gaussian distribution.

Qualitative epistasis is simulated by drawing 3 random effects for both involved markers, taking the absolute values from those, sorting them from low (0) to high (2) and multiplying those effects with each other. By this, we obtain the lowest effect for 00 and the highest for 22 with selection for alternative allele to be beneficial in all cases.

Quantitative epistasis is simulated by drawing 9 random effects and assigning the absolute values of two of those to the corner 02 and 20. All other combinations are assigned the minus absolute values of drawn random number.

To simulate more than one trait use vectors for **n.**XXX and lists for **var.**XXX instead.

##### 4.3.3 Correlated Traits

To generate correlated quantitative traits selected the traits that should be correlated via **shuffle.traits** and provide the needed correlation in **shuffle.cor**. Note that QTLs are then assigned to the same markers to get correlations independent of the underlying LD structure. To set a correlation for traits with no underlying QTL, use **new.breeding.correlation**. As simulation is done via sampling from a Gaussian distribution and genetic traits do not fulfil all requirements of a dependent multivariate Gaussian distribution (which is here used to model dependency), the obtained resulting correlations can be different to the correlation set in **new.breeding.correlation** if non-QTL traits have correlations with QTL-traits. We are currently working on alternatives for this.

#### 4.4 Position of Markers

For our simulations, the physical position in base pairs does not really matter as we are interested in a position in Morgan internally. We assume points of recombination to be distributed according to a Poisson distribution. On default, we assume markers to be equidistant with the total length of the chromosome in Morgan given by the parameter **chromosome.length** (default: 5). When performing a joint generation of multiple chromosomes with different sizes enter a vector instead. If non of the following options for the position of each marker are provides, markers are assumed to be equidistant. This will also minimize computation time and therefore should only be changed if needed.

Based on the physical positions entered in the parameter **bp** the position in Morgan can be derived by providing a conversion rate in **bpcm.conversion**. Note that tools like BEAGLE (Browning et al. 2018) assume 100.000.000 bp per Morgan. For chicken, we would recommend the use of 30.000.000 bp per Morgan.

Another way of entering the genetic position (in M) is via the parameter **snp.position** manually. Scaling can be performed internally by activating the parameter **position.scaling** and the **chromosome.length**. In addition, one should input a value for the number of base pairs before and after the last position (**length.before**, **length.behind**). Both should be chosen to be larger zero (default: 5) if scaling is performed.

For some applications, the recombination rate might not be the same for all individuals (e.g. male/female differences). To input an additional recombination map enter alternative positions in the parameter **add.architecture**. You can select which architecture is used for every parent in the actual simulation process.

### 5 Simulation of breeding processes (*breeding.diploid()*)

To perform the actual simulation of matings the function *breeding.diploid()* is used. Especially for that step, the sheer amount of different options can be deterring – in reality only a few parameters will actually be relevant. In this section, we will first discuss absolutely necessary parameters, their default options and afterwards discuss possible deviations. For exemplary simulation we refer to section 16. Note that you most likely can skip through some of the sections if you are not interested in changes in that dimension. There will be a lot of cases where there is the same parameter for the male and female part of the breeding program. We will limit ourselves here to the male parameter (parametername**.m**) – usage of the female version (parametername**.f**) will always be the same. The default setting for the female parameter is to be the same as the male parameter (<u>NOT</u> the default of the male parameter). Exception to this is course the case in when groups of individuals are selected to be used as parents and similar (e.g. **selection.f.cohorts**).

#### 5.1 General setup

The output of *breeding.diploid()* is an updated population list. All newly generated individuals are added as an additional generation – to add them to a previously existing generation set **add.gen** to the generation you want to add to.

The number of newly generated individuals can be chosen via **breeding.size**. Input for this is a numeric value and sex of each offspring is randomly determined via **breeding.sex** (probability of a male offspring). To remove randomness set **breeding.sex.random** to FALSE or input a vector containing the number of new male/female individuals in **breeding.sex** instead.

To control which individuals are used in the mating procedure use the parameter **selection.size** (vector of size 2, containing the number of used male/female individuals). By default only the individuals of the last generation and class 0 (this is usually all and you will realize when this is not the case) are used. Classes can be used to model migration in store groups of different genetic origin but same sex and generation or to just remove individuals from the pool of individuals considered for selection.

To control which individuals are used every cohort generated can be named via **name.cohort** in *breeding.diploid()* & *creating.diploid()*. The individuals used in the selection procedure can be chosen via **selection.m.gen**, **selection.m.database** and **selection.m.cohorts** (paternal side) and same syntax for the maternal side. In the old version of the code, this is equivalent to the use **of best1.from.cohorts** and **best1.from.groups** that are still alternative input parameters for **selection.m.database** and **selection.m.cohorts**.

To combine individuals to a new cohorts of individuals set **combine** to TRUE. This will generate a new cohort of all selected individuals - do not combine male and females individuals!

On default, the selection of individuals is done at random (For more on this we refer to 3.4).

To select the individuals of which class to consider in the selection procedure, use the parameter **class.m**, containing a vector of all usable classes. To control the class of the new individuals use the parameter **new.class** (default 0). Classes of all cohorts added are automatically added to the vector of considered classes (**class.m**) unless **add.class.cohorts** is set to FALSE.

To generate multiple offspring from the same dam/sire pair set **repeat.mating** to the desired number.

Both the time of the generation of new individuals (**time.point**) and the type of the mating (**creating.type**) can be stored. Both parameters are mostly used internally in the web-based application and are automatically tracked internally.

#### 5.2 Control of heritability, breeding values, genotypes and phenotypes

For each individual, an underlying true genetic value is calculated for each trait. Based on this, phenotypes can be generated. For which individuals to generate new phenotypes can be controlled via **new.bv.observation.gen**, **new.bv.observation.database** and **new.bv.observation.cohorts**. For a quick input of all individuals previously not phenotyped set **new.bv.observation** to "all", "non_obs", "non_obs_m" or "non_obs_f" for all or all previously not phenotyped individuals (potential of only one sex). On default, for each individual at most one phenotype is generate. Set **multiple.observation** to TRUE to allow for more than one observation per individual. To generate multiple observations in a single run of *breeding.diploid()* set **n.observation** to that number. Note that the number of times an observation for an individual is generated does matter since the environmental variance will be reduced with each observation as observations are assumed to be independet. To model a correlation between the environmental variances for different traits set provide the desired correlation matrix via **new.phenotyp.correlation** (this can also be done in *creating.diploid()*). For the simulation of correlated genetic values we refer to section 4.3.3.

The environmental variance $\sigma_e$ can be controlled by the usage of **sigma.e**. This can either be set to a fixed numeric value or be estimated to fulfill a target heritability. For the second possibility, the genetic variance is calculated based on the individuals specified in **sigma.e.gen**, **sigma.e.database** and **sigma.e.cohorts** and the needed environmental variance is then calculated by the usage of a predefined **heritability**. You can also use the environmental variance of the previous simulation by setting **use.last.sigma.e** to TRUE. A manual change of the genetic variance **sigma.g** is not recommended but in principle possible (this will only affect the breeding value estimation). On default it is estimated using all individuals used in the current breeding value estimation (set **forecast.sigma.g** to FALSE to deactivate). To specify which groups to use to estimate $\sigma_g$ use **sigma.g.gen**, **sigma.g.database**, **sigma.g.cohorts**.

For newly created individuals the phenotype is set to 0. Alternatively, one can it to the mean of the parents or create an observation by setting **new.bv.child** to "mean" or "obs" instead of "zero". In case of generating individuals via **copy.individual** one can also use "addobs" to import existing observations but also potential generate additional ones via **n.observation**. Estimated breeding values are also kept unless **copy.individual.keep.bve** is set to FALSE.

To select the share of individuals genotyped use the parameter **share.genotyped** or select it manually via **genotyped.s** (in concordance to **sex.s** in *creating.diploid()*). In case an individual is generated via **copy.individual** the genotyping state is keep and the share of previously not genotyped individuals that is now genotyped can be controlled via **added.genotyped**.

In some applications, the genetic value of the individual itself is not of importance – instead the performance of its offspring is of relevancy. To select for which individuals to import offspring phenotypes use **offspring.bve.parent.gen**, **offspring.bve.parent.database** and **offspring.bve.parent.cohorts**. Unless specified in **offspring.bve.offspring.gen**, **offspring.bve.offspring.database** and **offspring.bve.offspring.cohorts** all offspring are considered here.

For better comparison of and between breeding values it is possible to standardizing breeding values before the first generation by activating **standardize.bv**. By this the average breeding value is set to **standardize.bv.level** (default: 100) – for the calculation of this, the average of the individuals in generation **standardize.bv.gen** (default: 1) is used.

Scaling in case of index selection with multiple traits is performed in the selection process itself.

#### 5.3 Breeding value estimation

To perform selection one can perform breeding value estimation. To activate this set **bve** to TRUE. In the simplest case, one has to input which groups to use in the breeding value estimation via the parameters **bve.gen**, **bve.database** and **bve.cohorts**.

As there are a lot of different ways to perform breeding value estimation, we implemented multiple variants:

1. GBLUP with assumed known heritability and direct solving of the mixed model without REML variance component estimation
2. Bayesian approaches implemented in BGLR
3. GBLUP using EMMREML
4. GBLUP using sommer
5. Pedigree-based BLUP via breedR
6. Parent/Grandparent mean
7. Own function

In any case estimated breeding values are entered for all individuals unless **bve.insert.gen**, **bve.insert.database** or **bve.insert.cohorts** directly classifies for which groups breeding values are to be entered. The accuracy of the breeding value estimation is automatically reported unless **report.accuracy** is set to FALSE.

For the calculation of G we offer multiple methods with **computation.A**=”vanRaden” being the default (VanRaden 2008). Alternatives include “kinship”, “CM”, “CE” (Martini et al. 2017) and the non-Z-standardized version of the vanRaden method (“non_stand”). In case “kinship” is selected the depth of the pedigree has to be provided via the parameter **depth.pedigree** (default: 7). Note that these individuals are just used to calculate the A matrix. Individuals used in the actual breeding value estimation still need to be selected via **bve.gen**, **bve.database** and **bve.cohorts**. Individuals with no observed phenotype start with a value of 0. Internally all phenotypes that are exactly 0 are handled as an NA – suppress this by setting **bve.0isNA** to FALSE (note that only methods 1./2./4. are able to handle NAs in the data). To active the use of the single step genomic relationship matrix set **singlestep.active** to TRUE – otherwise non-genotyped individuals are not considered in the breeding value estimation unless **remove.non.genotyped** is set to FALSE (Legarra et al. 2014).

To not perform statistical breeding value estimation but instead using the phenotypes as breeding value estimates set **phenotype.bv** to TRUE.

As the presence of true effect markers in the dataset might be a strong assumption one can set **remove.effect.position** to not use markers associated with any traits in the breeding value estimation.

To only included individuals with a certain class set **bve.class** to a vector containing all classes to consider.

In case individuals were generated via **copy.individual** (this is especially relevant for the web-based application) each individuals is only used at most once. To consider the same individual multiple times set **bve.avoid.duplicates** to FALSE. Note that cloning will not lead to the same ID.

##### 5.3.1 Direct approach with known heritability

Main advantage of a direct estimation is a massive improvement in computation time as the usually necessary REML estimation takes most of the time. In practice, it might not be realistic but since genetic values are known it is possible. Note that this is still an empirical measure that can change when using different individuals in the estimation process. Especially for bigger populations heritability estimation should not be problematic and is not performed in each breeding value estimation in practice as well. To instead estimate the additive genetic variance using a parental model activate **estimate.add.gen.var**.

In case of missing phenotypes, estimates will be based on (VanRaden 2008) method 2. This will also be used in case of the use of single step. Alternatively one can use rrBLUP based estimates by setting **bve.direct.est** to FALSE. Note that this second version is slower and requires the presence of individuals that are phenotyped and genotyped.

##### 5.3.2 Bayesian approaches (BGLR)

For performing Bayesian methods we are using the R-package BGLR. To activate the usage of BGLR set **BGLR.bve** to TRUE. On default a Reproducing-kernel-hilbert-space is used – alternatively one can use BayesA, BayesB, BayesC by setting **BGLR.model** to "BayesA", "BayesB", "BayesC", "BRR", or "BL" instead of "RKHS". Other parameter values will all be chosen according to the defaults of the BGLR package. If you want to test alternative parameter settings or use other methods implemented in BGLR either do the estimation manually (Chapter 5.3.8) or contact the me to add it to the package.

To control the number of the burn-in and iterations use **BGLR.burnin** and **BGLR.iteration**. To deactivate printing of results of interim steps set **BGLR.print** to FALSE (equal to verbose=FALSE in BGLR). On default BGLR will generate some internal files in its computations. To select a path of where to store them chose it via **BGLR.save**. Especially when parallelizing thousands of simulations BGLR tents to crash when the same path is used multiple times. Activating **BGLR.save.random** will hinder this.

##### 5.3.3 GBLUP (EMMREML)

Traditional GBLUP including variance component estimation using REML is performed by using the package EMMREML. To activate the usage set **emmreml.bve** to TRUE. EMMREML does not support missing phenotypes and therefore can only be used if phenotypes for all individuals in the BVE are available (if not use the direct approach, sommer or rrBLUP).

##### 5.3.4 GBLUP (sommer)

Traditional GBLUP including variance component estimation using REML is performed by using the package sommer. To activate the usage set **sommer.bve** to TRUE. Sommer does support missing phenotypes.

To activate the use of the multi-trait model implemented in sommer use **sommer.multi.bve** to TRUE. Note that this will take substantially longer than single trait models.

##### 5.3.5 GBLUP (rrBLUP)

Traditional GBLUP including variance component estimation using REML is performed by using the package rrBLUP. To activate the usage set **rrBLUP.bve** to TRUE. rrBLUP is about 2.5 times as fast as sommer for breeding value estimation.

##### 5.3.6 Pedigree-based (breedR)

Traditional breeding value estimation using pedigree data. To activate set **breedR.bve** to TRUE – especially for bigger populations this is much faster computation wise. Overall heritabilities in breedR seem to be underestimated and accuracies slightly lower. Alternatively, the pedigree-based relationship matrix can also be used in all other methods by setting **computation.A** to “kinship”. This procedure takes about the same time as with other relationship matrices (No usage of high number of zeros in the relationship matrix or the direct inverse of the pedigree matrix).

##### 5.3.7 Parent/Grandparent mean

To use the mean performance of the parents / grandparents as the breeding value use **bve.parent.mean** / **bve.grandparent.mean**. On default breeding value estimates for the parents are used and if those are not available phenotypes. Alternatively on can select to use breeding values only (“bve”), phenotypes only (“pheno”) or genomic values (“bv”) via the parameter **bve.mean.between**.

##### 5.3.8 Own function

Instead of performing breeding value estimation inside of *breeding.diploid()* one can implement his own methodology by exporting all information needed to those computations and inserting own breeding values estimates via the function *insert.bve().*

According code could look like this:

```
# Simulate Phenotypes for generation 4 with heritability 0.4
population <- breeding.diploid(population, heritability = 0.4,
```

```
                                  sigma.e.gen = 4,
                                  new.bv.observation.gen = 4)
# Export genotypes and phenotypes for generation 4
genos <- get.geno(population, gen = 4)
phenos <- get.pheno(population, gen = 4)

# Here you perform your own method to assign breeding values to each individual
bve <- runif(ncol(genos)) # This is probably not the best technique for this =)

# Import breeding values estimated for generation 4
bves <- cbind(colnames(genos), bve)
population <- insert.bve(population, bves=bves)
```

For details on exporting functions, we refer to section 6. For details on importing function, we refer to section 7.

##### 5.3.9 Calculating marker effects & GWAS

For some applications (e.g. gene editing) it is necessary to identify causal markers. Although marker effects are known in a simulation, in practice one has to identify them. Options here are either a direct calculation of the effect size of each marker based on the computations performed in 5.3.1 (rrBLUP) or the performance of a GWAS-study without correction for population structure. Methods can be activated by setting **estimate.u** or **gwas.u** to TRUE.

In case of a GWAS study one can additionally select the groups used in the study by setting **gwas.gen**, **gwas.database** and **gwas.cohorts** (default is same as for breeding value estimation). As a value for y, one can use the phenotype ("pheno"), true breeding value ("bv") or the estimated breeding value ("bve"). Additionally it might be necessary to standardize the y value by the mean of the group by activating **gwas.group.standard**. Note that this is a basic implementation of GWAS with no correction for population structure or similar.

##### 5.3.10 Calculation of reliabilities

Reliabilities are not derived in any of the used R-packages. In the direct approach (Chapter 5.3.1), they can be derived by setting **calculate.reliability** to TRUE according to (VanRaden 2008).

#### 5.4 Selection techniques & mating strategies

Selection of the individuals for matings in the following generations is of key importance for any breeding program. Especially here, one is limited to the techniques that work in the species one wants to simulate. On default settings, the selection of the new founders is done at random. To use estimated breeding values as a selection criteria set **selection.m** to "function". To ignore the best selected individuals set **ignore.best** to that value – note that this value will be internally subtracted from **selection.size**. E.g. to simulate mating between the top 100 female individuals with the third and fourth best male individual set **selection.size** = c(4,100) and **ignore.best**=c(2,0). To exclude specific individuals from the set of individuals to select from use **reduced.selection.panel.m**. The vector should contain all individuals to use (e.g. 1:10 when selecting from the first ten individuals).

To store details on which individuals were selected, which mating were performed and the currently estimated breeding values activate **store.breeding.totals**.

Selection can be performed based on the phenotype, genetic value or the breeding value estimates. To select what to use set **selection.criteria.type** to “pheno”, “bv” or “bve” (default: “bve”).

##### 5.4.1 Multiple traits

When working with multiple traits, the selection of the best individuals is typically done by the use of a combination of those traits. All single values can either be added up directly (**multiple.bve**=”add”) or one can use a selection index just accounting for the ranking (**multiple.bve**=”ranking”). To reduce scaling problems for different traits one can use **multiple.bve.scale.m** to standardize the variance in each trait. Note that this scaling in the cohort mode is for all individuals together whereas in the old selection modes it is done per group (! – needs to be the same!!!). Additionally, each trait can be assigned a weighting via **multiple.bve.weights.m**.

To derive the ideal index based on phenotypic/genotypic variance, reliabilities and economic gains per unit according to (Miesenberger 1997) set **selection.m.miesenberger** to TRUE. Economic gains can be provided in **multiple.bve.weights.m**. On default, the gain has to be provided per standard deviation of the breeding value estimations. Alternatives can be provided in **selection.miesenberger.w** (default: “bve_sd”) and are per unit (“unit”) and per phenotypic standard deviation (“pheno_sd”).

In case reliabilities are not derived (Chapter 5.3.10), they need to be estimated. On default, this is done by dividing the standard deviation of the breeding value estimation by the standard deviation of the phenotypes. Alternatives can be entered in **selection.miesenberger.reliability.est**, as the direct use of the heritability (“heritability”) or the actual calculation according to the correlation between breeding value estimates and true underlying genomic values (“derived”) which is of course not possible in practice but should be the most accurate.

##### 5.4.2 Higher procreation of genetically favored individuals

Genetically favored individuals tend to procreate more often. To model this set a ratio between the likelihood of the best individual to mate compared to the worst individual (in the group of selected individuals) in **best.selection.ratio.m**. This ratio is the ratio between the frequency the best selected individual and the worst selected individual. All other frequencies are then calculated linearly. E.g. in a group of selected individuals with breeding values 105, 103 and 100 with a ratio of 6 the relative frequencies are of 6,4,1 (this just is a linear function – comp for individual 2: (103 – 100)/ (105-100) * (6-1) +1). Criteria behind can be either “bv”, “bve” or “pheno” and can be entered in **best.selection.criteria.m**. To manually enter the probability of each individual in the group of selected individuals input a vector with frequencies for each individual in **best.selection.manual.ratio.m**. Individuals selected are sorted with the individual with the highest estimated breeding value being the first one.

This does not require breeding value estimation and can also be used to simulate slow natural selection processes over thousands of generations.

Higher procreation is also relevant for optimum genetic contribution theory and the use of **ogc** will automatically change these parameters accordingly (section 5.4.8).

##### 5.4.3 Maximum number of offspring per individual

To control the maximum number of times each individual is used for reproduction set **max.offspring**. Either enter a numeric if that boundary is for both sexes or a vector with the first value coding the maximum for male and the second the one for female.

##### 5.4.4 Plant breeding (no-sexes & selfing & DH-production & cloning)

For some applications the sex of an individuals is not relevant (or an individual does not even have a sex). Even though a sex is still stored internal it might be neglectable for the application at hand. In this case, one can allow matings between individuals from the same sex by usage of **same.sex.activ**. The probability to select a female individuals as a parent can be set via **same.sex.sex** (default=0.5). To additionally allow for selfing set **same.sex.selfing** to TRUE. The probability for this mating is the same as any other mating combination.

To perform exclusively selfings, activate **selfing.mating** and selected the probability to use a female parent via **selfing.sex**.

To generate doubled haploid lines active **dh.mating** and selected the probability to use a female parent via **dh.sex**.

To generate and exact copy of an individual to **copy.individual** to TRUE. Instead of simulating the meiosis both chromosome sets of the selected first parent (usually father) will be copied (to copy female individuals use **same.sex.activ** and set the probability to use females to 1 (**same.sex.sex**=1).

##### 5.4.5 Generate offspring for all sire combination

To generate offspring from each possible parental combination set **breeding.all.combination** to TRUE. In case only individuals from one sex are selected sex is ignored when deriving potential matings. **Breeding.size** still has to be set.

##### 5.4.6 Targeted/Fixed mating/Manual selection of individuals

If none of the previously described methods works for your simulation, you can also manually enter a list of all matings that should be performed. For this, use the parameter **fixed.breeding**. To perform targeted mating in the group of the best individuals use **fixed.breeding.best**. Here each row just contains the sex and position in the list of selected individuals. In both cases, an additional column can be added that is coding the likelihood of the offspring being female.

##### 5.4.7 Gene-Editing

With increasing popularity of methods like CRISPR/Cas9 one might be interested in performing gene editing to increase the genetic gain. Gene editing can be activated by setting **gene.editing** to TRUE. The number of edits can be controlled via **nr.edits** and effect markers are picked via usage of the predictions via rrBLUP/GWAS in section 5.3.9. We only count actually performed edits – if an allele is already beneficial, the next best marker is edited instead. Although such a procedure is not possible in practice, the first integrated way of editing is to edit all selected individuals – this technique is also performed in the approach PAGE (Jenko et al. 2015) and our counter version (Simianer et al. 2018).

As a more realistic scenario, we also allow for the editing of offspring via **gene.editing.offspring**. To only perform editing on male or female individuals set **gene.editing.offspring.sex** / **gene.editing.best.sex** to 1 (male) or 2 (female).

Note that traditionally modelled effects often neglected strongly deleterious mutations and we are here assuming that 100% of all edits to work, all possible offspring will survive the procedure and traits are as simple as designed (usually single marker QTL).

##### 5.4.8 Optimum genetic contribution

To use optimum genetic contribution theory from the group of selected individuals set **ogc** to TRUE. The target increase of the average relationship can be selected via **ogc.cAc**. On default, the increase is minimized. MoBPS is used the traditional formula according to (Meuwissen 1997) and a pedigree based relationship (change via the parameter **computation.A.ogc**). One can also provide on weightings via **best.selection.manual.ratio.m** (Chapter 5.4.2) and/or **fixed.breeding** (Chapter 5.4.6). We are also willing to implemented alternatives if they are needed/wanted.

#### 5.5 Genetic architecture

When simulating meiosis, we are accounting for recombination and mutation. We are assuming that recombination points are Poisson distributed with one expected point of recombination per 1 Morgan. To change this, set **recombination.rate** to the needed value. To not use, a fixed value but a step-function instead use **recom.f.indicator**. Additional genetic architectures can be added the same way as in *creating.diploid()* via **add.architecture**. To select the genetic architecture of recombination for set **gen.archicture.m** to the architecture that should be used for males.

Regarding mutation rates we are assuming that each marker has the same probability for a mutation – this can be changed via **mutation.rate** (default: $10^{-5}$). A mutation back to the reference is assigned the probability of **remutation.rate** (default: $10^{-5}$). Those values tend to be a lot higher than what you would expect in nature. Depending on the base pair one would expect something around $10^{-8}$ for a random loci.

Duplications are implemented but the modelling is absolutely adhoc and probably needs refinement – talk to me if you plan to do something in that direction!

#### 5.6 Other

##### 5.6.1 Culling / Death

###### 5.6.1.1 Culling module (Web-interface)

The new culling module is currently only available for cohorts and mainly intended for interface users. Here, parameters settings will take care of itself. To manually execute this in R use **culling.cohort** to select for which cohort to execute the module. The age of the individual can be provided in **culling.time**. The name of the culling reason can be provided in **culling.name**. Additional one can provide two breeding values (**culling.bv1**, **culling.bv2**) and two culling probabilities (**culling.share1**, **culling.share2**). For all other genomic values the probability of culling this then derived with linear extension.

An index of weighting between traits can be selected via **culling.index** (similar to **multiple.bve.weights.m** – Chapter 5.4.1). On default, no genomic influence is assumed and all individuals are culled with the same probability (**culling.share1**).

###### 5.6.1.2 Old module

Especially for cost calculation it might be necessary to know the time of death for each individual. A group of individuals can be reduced to **reduce.group** with each row coding generation, sex, number of individuals to keep, class. To set the selection criteria use **reduce.group.selection** (default: "random").

##### 5.6.2 Parameter for target-mating with J.W.R. Martini

Don't think anyone needs documentation here – code is specific to the planned paper with Johannes and does not generalize (quick and dirty - https://github.com/Droogans/unmaintainable-code)

**martini.selection / Special.comp / Special.comp.add / Max_auswahl / Predict.effects / SNP.density / Use.effect.markers / Use.effect.combination**

##### 5.6.3 Allele-frequency per generation

To store the frequency of each allele per generation activate **store.effect.freq**.

##### 5.6.4 Set a random seed

For repeatability it might be helpful to set a random seed in R. This can be done via the parameter **randomSeed** or directly performed in R using *set.seed().*

##### 5.6.5 save.recombination.history

To store the time of occurance of each point of recombination activate **save.recombination.history**. This has to be done starting with the first generation and currently crashes after setting a new founder population (Currently nobody needs it! – but should be an easy fix!)

#### 5.7 Storage & computing time

Computing time for the whole simulation (population$info$comp.times), the steps of the breeding value estimation (population$info$comp.times.bve) and the generation of new individuals (population$info$comp.times.generation) are stored in the population list. If this is not required you can deactive this by setting **store.comp.times**, **store.comp.times.bve** and **store.comp.times.generation** to FALSE. Unless **verbose** is set to FALSE you should automatically receive notifications on the current step of the algorithm and computing times of individual steps. Activation of **Rprof** can provide even more information.

##### 5.7.1 Reducing the size of the population list

Especially when simulating populations with lots of markers, individuals and/or generations, data storage can become a problem. As the internal structure of a population list is complex and manually deleting of things is not recommended – use following settings in *breeding.diploid()* instead:

**delete.haplotypes**: Vector containing all generations for which haplotypes no longer need to be stored (note that only founder generations are stored anyway – everything else is calculated on-the-fly)

**delete.same.origin**: Merge two adjacent segments with the same founding haplotypes (deletion of a recombination point with no influence)

**delete.individuals**: Vector containing all generations for complete deletion– to only delete one sex use **delete.sex** (vector contain sex to delete – 1 (male), 2 (female). Especially when the number of recombinations stored per individual, becomes bigger this is of relevancy.

##### 5.7.2 Reducing memory needs in the BVE

To calculate the genomic relationship matrix, one has to perform matrix multiplication of a matrix containing n*p entries (individuals x markers). Note that for every entry only 2 bits are needed when using miraculix. Nevertheless, this can become extremely big – to reduce this, one can perform the calculation of G sequentially and only load a part of Z into memory at any time.

To activate this, set **sequenceZ** to TRUE. The number of markers in memory can be controlled via **maxZ** (default: 5000). Alternatively, one can put **maxZtotal** to control the total number of entries instead. As this is increasing the computation time, we first recommend to activation of miraculix. As long as miraculix is installed this will happen automatically unless the parameter **miraculix** in *creating.diploid()* is set to FALSE. For more on this we refer to section 9.12.

To speed up commutation one can use multiple cores by the usage of **miraculix.cores** (default: 1) or in case miraculix is not active **ncore**. The backend outside of miraculix is using doParallel and mclapply but doMPI is supported as well – we highly recommend the use of miraculix instead.

Setting **fast.compiler** to TRUE will additionally activate a just-in-time-compiler (enableJIT(3)).

To save computation speed in the GWAS, one can use **approx.residuals** – this does not influence the order of the predicted effect markers but will influence p-values slightly.

##### 5.7.3 Inverting G using miraculix

The inversion of $(G + I_n \cdot \lambda)$ can take a lot of time, is numerically unstable and might not even be possible at all if the matrix is not invertible at all. Instead of the standard cholesky procedure using *chol2inv(chol())*, the inversion can also be done in RandomFieldsUtils/miraculix by activating **chol.miraculix**. Leading to slightly reduced computing times – but also includes screening for semi definite matrices and an automatically changed algorithm, if needed, and thus proceeding without error.

##### 5.7.4 On-the-fly calculation of haplotypes

To save memory, haplotypes are calculated on-the-fly. For this, the location of each recombination point (between which markers) has to be stored. In case one is working with equidistant markers, it basically takes no time. For other cM-positions it might increase computation speed to provide a function that derives the last marker in front of a certain position in Morgan. This function can be entered via **import.position.calculation**. Only in extreme cases (lots of markers) this should even matter!

### 6 Exporting information from the population-list (get.XXXX)

Most of the data stored in a population list is highly compressed since saving haplotypes of all individuals of the dataset for all generations would often exceed most local machines or even servers. For some applications (especially if one wants to perform his own fancy simulations without contacting the author and asking him to extent the package) it might be useful to understand the data structure behind. For that, we refer to section 8. In most cases, using our predefined export functions should be enough:

In what individuals one is interested in can be controlled by usage of the parameter **gen**, **database** and/or **cohorts**. Here **cohorts** and **gen** are vectors containing all generations/cohort to include whereas **database** contains a matrix with each row coding a group to export:

```
> database
     Generation sex
[1,]          1   2
[2,]          5   1
```

This **database** will export the information for all female individuals from generation 1 and all male individuals from generation 5. If require a third and fourth column in **database** can be added to indicate the first and last individual of the group to consider. MoBPS will automatically add this to be the full group when *get.database()* is used.

#### 6.1 get.genos

This function will export genotypes. To additionally output the base pair of the minor/major allele set the parameter **export.alleles** to TRUE. Each column contains one genotype with column names indicating sex, individual number and generation.

```
> genos <- get.geno(population,gen=3)
> genos[1:5,1:10]
         M1_3 M2_3 M3_3 M4_3 M5_3 M6_3 M7_3 M8_3 M9_3 M10_3
Chr1SNP1    1    2    1    0    2    1    2    2    2     1
Chr1SNP2    1    0    1    0    2    1    1    2    1     0
Chr1SNP3    1    0    1    1    0    1    0    1    0     1
Chr1SNP4    0    0    1    1    1    1    0    0    0     1
Chr1SNP5    1    0    1    2    1    1    0    0    0     0
```

#### 6.2 get.haplos

This function will export haplotypes. To additionally output the base pair of the minor/major allele set the parameter **export.alleles** to TRUE. Each column contains one haplotype with column names indicating sex, individual number, chromosome set (currently only diploid individuals) and generation.

```
> haplos <- get.haplo(population,gen=3)
> haplos[1:5,1:10]
        M1_3_set1 M1_3_set2 M2_3_set1 M2_3_set2 M3_3_set1 M3_3_set2 M4_3_set1 M4_3_set2 M5_3_set1 M5_3_set2
Chr1SNP1        0         1         1         1         0         1         0         0         1         1
Chr1SNP2        0         1         0         0         0         1         0         0         1         1
Chr1SNP3        1         0         0         0         1         0         0         1         0         0
Chr1SNP4        0         0         0         0         1         0         1         0         1         0
Chr1SNP5        1         0         0         0         1         0         1         1         1         0
```

#### 6.3 get.bv / get.bve / get.pheno / get.reliability / get.selectionindex

These functions will export the true underlying breeding value (“bv”), the estimated breeding value (“bve”), the phenotype (“pheno”), the reliability for each breeding value estimation/trait (“reliability”) and the selection index used in the last selection procedure (“selectionindex”).

```
> bv <- get.bv(population,gen=1)
> bve <- get.bve(population,gen=1)
> pheno <- get.pheno(population,gen=1)
> bve[1:5]
[1] 45.95833 48.85317 31.53675 35.73686 49.55310
> bv[1:5]
[1] 33.35516 47.52439 31.92523 59.13089 37.96659
> pheno[1:5]
[1] -22.230518  67.241256  -9.337895 -16.056936 101.497754
```

#### 6.4 get.recombi

This function will export all points of recombination and the genetic origin of each segment. The structure here is a list of 4 elements with elements 1 (paternal) and 2 (maternal) containing recombination points and elements 3 (paternal) and 4 (maternal) containing the genetic origin.

Each row is coding the genetic origin between two points (generation, sex, individual number, chromosome set). In the example provided this would mean that the segment between 0.000 and 0.218 of the paternal chromosome originates from the second chromosome set of the 532nd female individual of the first generation.

Note that in addition to all recombination points the start and end points of chromosomes are also exported.

```
> recombi <- get.recombi(population, gen=3)
> recombi[[1]][[1]]
 [1]  0.0000000  0.2183368  0.3517032  0.6187816  1.0718667  1.3089574  1.6053402  1.6390138  1.6627053  1.9827624  2.0846120
[12]  2.1364282  2.4399836  3.0289287  3.7351480  4.0304151  4.6348112  5.5724172  5.9576469  6.5091164  6.9677451  7.4578594
[23]  8.5493007  8.8105940  8.9809813  9.2018383 10.0248883 11.4693043 11.6175350 11.6398517 12.6182508 13.4061942 13.5190658
[34] 13.8967908 14.4158473 14.4829801 14.7182578 15.1508620 15.1545097 15.5787925 15.7266202 15.8199412 16.3273204 17.1619292
[45] 17.3799130 17.7438021 18.1580695 19.0026216 19.8908821 20.1429153 21.3074407 22.2018853 22.5145334 22.8492019 22.8980965
[56] 23.2768180 23.4729906 23.8671758 24.1965660 24.2540496 24.5550846 24.9169786 25.2573038 25.3671158 25.5634302 25.7719966
[67] 25.9989692 26.2255921 26.8175851 26.8552645 27.0569816 27.3228323 27.7300196 27.7441943 28.0331752 28.0909684 28.0925820
[78] 28.4657455 28.7984996 29.0263514 29.1274251 29.2751559 29.3417943 29.5015208 29.6375127 29.8279188 30.0385607 30.1115719
[89] 30.1383062 30.2662008 30.2851334 30.4432174
> recombi[[1]][[3]][1:5,]
     [,1] [,2] [,3] [,4]
[1,]    1    2  352    2
[2,]    1    2  352    1
[3,]    1    1   78    1
[4,]    1    2  352    2
[5,]    1    2  352    1
> recombi[[1]][[5]]
[1] "M1"
```

#### 6.5 get.pedigree (1/2/3)

This function will export the pedigree. Individuals are coded by sex, individual number and generation.

```
> ped <- get.pedigree(population, gen=12)
> ped[1:10,]
     offspring father     mother
[1,] "M1_12"   "M2214_11" "W1776_11"
[2,] "M2_12"   "M2052_11" "W1125_11"
[3,] "M3_12"   "M1529_11" "W1904_11"
[4,] "M4_12"   "M1712_11" "W1256_11"
[5,] "M5_12"   "M221_11"  "W326_11"
```

Instead of character string with "M"/"W" indicting sex, one can also directly export a table with 9 colums indicating Sex (1/2), Generation(1,2,3,…) and individidal number (1,2,3,…) in a numeric format by setting **raw** to TRUE.

To export grandparents use *get.pedigree2()*, to get both *get.pedigree3()*. In *get.pedigree2()* one can additionally export the share of the genome inherited by which grandparent by setting the parameter **shares** to TRUE.

```
> ped <- get.pedigree2(population, gen=12)
> ped[1:5,]
     offspring grandfatherf grandmotherf grandfatherm grandmotherm
[1,] "M1_12"   "M734_10"    "W2135_10"   "M2342_10"   "W313_10"
[2,] "M2_12"   "M1188_10"   "W1436_10"   "M489_10"    "W539_10"
[3,] "M3_12"   "M1702_10"   "W1860_10"   "M413_10"    "W1234_10"
[4,] "M4_12"   "M1560_10"   "W1211_10"   "M1649_10"   "W254_10"
[5,] "M5_12"   "M1649_10"   "W2093_10"   "M1593_10"   "W2390_10"
> ped <- get.pedigree3(population, gen=12)
> ped[1:5,]
     offspring father     mother     grandfatherf grandmotherf grandfatherm grandmotherm
[1,] "M1_12"   "M2214_11" "W1776_11" "M734_10"    "W2135_10"   "M2342_10"   "W313_10"
[2,] "M2_12"   "M2052_11" "W1125_11" "M1188_10"   "W1436_10"   "M489_10"    "W539_10"
[3,] "M3_12"   "M1529_11" "W1904_11" "M1702_10"   "W1860_10"   "M413_10"    "W1234_10"
[4,] "M4_12"   "M1712_11" "W1256_11" "M1560_10"   "W1211_10"   "M1649_10"   "W254_10"
[5,] "M5_12"   "M221_11"  "W326_11"  "M1649_10"   "W2093_10"   "M1593_10"   "W2390_10"
```

#### 6.6 get.cohorts

This function extracts all existing cohorts from the population list. Set **extended** to TRUE to also extract further information on the cohorts:

```
> get.cohorts(population)[1:5]
[1] "Founder_M"      "Founder_W"      "Cows"           "Bulls"          "Selected_bulls"
> get.cohorts(population, extended=TRUE)[1:5,]
     name             generation male individuals female individuals class position first male position first female time point creating.type
[1,] "Founder_M"      "1"        "50"             "0"                "0"   "1"                 "0"                   "0"        "0"
[2,] "Founder_W"      "1"        "0"              "50"               "0"   "0"                 "1"                   "0"        "0"
[3,] "Cows"           "2"        "0"              "50"               "1"   "1"                 "1"                   "1"        "2"
[4,] "Bulls"          "2"        "50"             "0"                "1"   "1"                 "51"                  "1"        "2"
[5,] "Selected_bulls" "3"        "10"             "0"                "2"   "1"                 "1"                   "1"        "1"
```

#### 6.7 get.class

This function extracts the class of each individual:

```
> get.class(population, gen=1:2)
 M1_1  M2_1  M3_1  M4_1  M5_1  M6_1  M7_1  M8_1  M9_1 M10_1 M11_1 M12_1 M13_1
    0     0     0     0     0     0     0     0     0     0     0     0     0
M14_1 M15_1 M16_1 M17_1 M18_1 M19_1 M20_1 M21_1 M22_1 M23_1 M24_1 M25_1 M26_1
    0     0     0     0     0     0     0     0     0     0     0     0     0
```

…

```
W42_1 W43_1 W44_1 W45_1 W46_1 W47_1 W48_1 W49_1 W50_1  M1_2  M2_2  M3_2  M4_2
    0     0     0     0     0     0     0     0     0     1     1     1     1
 M5_2  M6_2  M7_2  M8_2  M9_2 M10_2 M11_2 M12_2 M13_2 M14_2 M15_2 M16_2 M17_2
    1     1     1     1     1     1     1     1     1     1     1     1     1
M18_2 M19_2 M20_2 M21_2 M22_2 M23_2 M24_2 M25_2 M26_2 M27_2 M28_2 M29_2 M30_2
    1     1     1     1     1     1     1     1     1     1     1     1     1
```

#### 6.8 get.genotyped

This function extracts if an individual is genotyped or not (mostly relevant for cost calculation and/or use in Single Step GBLUP (Legarra et al. 2014).

```
> get.genotyped(population, gen=6)
 M1_6  M2_6  M3_6  M4_6  M5_6  M6_6  M7_6  M8_6  M9_6 M10_6 M11_6 M12_6
FALSE  TRUE FALSE  TRUE  TRUE  TRUE  TRUE  TRUE  TRUE  TRUE  TRUE  TRUE
M13_6 M14_6 M15_6 M16_6 M17_6 M18_6 M19_6 M20_6 M21_6 M22_6 M23_6 M24_6
 TRUE FALSE  TRUE FALSE FALSE  TRUE FALSE  TRUE  TRUE  TRUE FALSE  TRUE
M25_6 M26_6 M27_6 M28_6 M29_6 M30_6 M31_6 M32_6 M33_6 M34_6 M35_6 M36_6
FALSE FALSE  TRUE  TRUE  TRUE  TRUE  TRUE  TRUE  TRUE FALSE FALSE FALSE
M37_6 M38_6 M39_6 M40_6 M41_6 M42_6 M43_6 M44_6 M45_6 M46_6 M47_6 M48_6
 TRUE FALSE  TRUE FALSE FALSE FALSE  TRUE  TRUE  TRUE FALSE  TRUE FALSE
```

#### 6.9 get.time.point

This function extract the time point of generation – this is mostly applicable when using the web-based application since there the first possible time point of generation is automatically calculated.

```
> get.time.point(population , gen=1:2)
 M1_1  M2_1  M3_1  M4_1  M5_1  M6_1  M7_1  M8_1  M9_1 M10_1 M11_1 M12_1 M13_1 M14_1 M15_1
    0     0     0     0     0     0     0     0     0     0     0     0     0     0     0
M16_1 M17_1 M18_1 M19_1 M20_1 M21_1 M22_1 M23_1 M24_1 M25_1 M26_1 M27_1 M28_1 M29_1 M30_1
    0     0     0     0     0     0     0     0     0     0     0     0     0     0     0

…

W41_1 W42_1 W43_1 W44_1 W45_1 W46_1 W47_1 W48_1 W49_1 W50_1  M1_2  M2_2  M3_2  M4_2  M5_2
    0     0     0     0     0     0     0     0     0     0     1     1     1     1     1
 M6_2  M7_2  M8_2  M9_2 M10_2 M11_2 M12_2 M13_2 M14_2 M15_2 M16_2 M17_2 M18_2 M19_2 M20_2
    1     1     1     1     1     1     1     1     1     1     1     1     1     1     1
```

#### 6.10 get.creating.type

This function extracts the creating type of each individual – this is mostly applicable when using the web-based application of the package. Here following coding is used:

0 – Founder

1 – Selection

2 – Reproduction

3 – Recombination

4 – Selfing

5 – DH-Production

6 – Cloning

7 – Combine

8 – Aging

9 – Split

```
> get.creating.type(population, gen=1:3)
  M1_1   M2_1   M3_1   M4_1   M5_1   M6_1   M7_1   M8_1   M9_1  M10_1  M11_1  M12_1  M13_1  M14_1  M15_1
     0      0      0      0      0      0      0      0      0      0      0      0      0      0      0
 M16_1  M17_1  M18_1  M19_1  M20_1  M21_1  M22_1  M23_1  M24_1  M25_1  M26_1  M27_1  M28_1  M29_1  M30_1
     0      0      0      0      0      0      0      0      0      0      0      0      0      0      0
```

…

```
W31_2 W32_2 W33_2 W34_2 W35_2 W36_2 W37_2 W38_2 W39_2 W40_2 W41_2 W42_2 W43_2 W44_2 W45_2
    2     2     2     2     2     2     2     2     2     2     2     2     2     2     2
W46_2 W47_2 W48_2 W49_2 W50_2  M1_3  M2_3  M3_3  M4_3  M5_3  M6_3  M7_3  M8_3  M9_3 M10_3
    2     2     2     2     2     1     1     1     1     1     1     1     1     1     1
```

#### 6.11 get.cullingtime

This function extracts the time of culling of each individual – this is mostly applicable when using the web-based application of the package.

#### 6.12 get.individual.loc

Function to derive the position in the stored population-list.

```
> get.individual.loc(population, gen=1)
               generation sex individual nr.
M1_1                    1   1              1
M2_1                    1   1              2
M3_1                    1   1              3
M4_1                    1   1              4
M5_1                    1   1              5
```

#### 6.13 get.vcf

Function to export genomic data in a vcf-file (currently using the synbreed-package but more efficient implementation including stored bp etc. is planned). Set **path** to the path you want to write to:

```
##fileformat=VCFv4.1
##filedate= 17980
##source="write.vcf of R-synbreed"
##FORMAT=<ID=GT,Number=1,Type=String,Description="Genotype">
#CHROM  POS ID  REF ALT QUAL    FILTER  INFO    FORMAT  M1_3_set2   M10_3_set2  M2_3_set2   M3_3_set2
chr1    .   Chr1SNP1    A   G   .   PASS    .   GT  0|0 0|1 1|0 0|0 0|0 0|0 0|0 0|0 0|0 0|0
chr1    .   Chr1SNP2    A   G   .   PASS    .   GT  1|0 0|1 1|1 0|1 0|1 0|1 0|0 0|0 1|0 0|0
chr1    .   Chr1SNP3    A   G   .   PASS    .   GT  0|1 1|0 0|0 1|0 0|0 1|0 1|1 0|1 0|1 0|0
chr1    .   Chr1SNP4    A   G   .   PASS    .   GT  1|0 0|1 1|0 1|1 0|1 1|1 0|0 1|0 0|0 0|0
chr1    .   Chr1SNP5    A   G   .   PASS    .   GT  0|0 0|1 0|0 0|0 0|0 0|0 0|0 0|0 0|0 0|0
```

#### 6.14 get.pedmap

Function to export genomic data in a ped and a map-file (PLINK format (Purcell et al. 2007)). The first of the two colums of each marker is representing the first haplotype of the individual. Set **path** to the path you want to write to:

```
1 1 0 0 0 0 C C C C C C C A A A C C C A C A C A A C C A C A
1 2 0 0 0 0 A A A A A A A A A A A A A C C A A A A A A C C C A
1 3 0 0 0 0 A C A A C C C C C C A A A A A A C A C C A A A C C
1 4 0 0 0 0 A C A A C C A C C A A A A A C C A C C C C A A A A
1 5 0 0 0 0 A A A A A A A A A A A A C A A A A A A A A C C C A
1 6 0 0 0 0 A A A A C C A A A A A A A A A C A A A A C A C C A
1 7 0 0 0 0 A A A A A C C C C C A A A A A A C C C C C C A A C
1 8 0 0 0 0 A C A C C C A C A C C C A A A C C C C A C A A C C
1 9 0 0 0 0 C C C A C C C C A A C A C A A A C C A A A A A C A
1 10 0 0 0 0 C A A A C C C A C C C C A A A A A A A C C C C A C
1 11 0 0 0 0 A C A A C C C C C C A A A A C A C C C C A C A C C
```

```
1 SNP1 0 0
1 SNP2 0 0
1 SNP3 0 0
1 SNP4 0 0
1 SNP5 0 0
1 SNP6 0 0
1 SNP7 0 0
1 SNP8 0 0
1 SNP9 0 0
```

#### 6.15 get.database

Function to merge **gen**, **database** and **cohorts** –info into a joint database. This is only needed internally – as it is the only internal *get.*X*()* function it is still mentioned here for completeness.

```
> get.database(population, gen=1, cohorts="Selected_bulls")
         start end
[1,] 1 1     1  50
[2,] 1 2     1  50
[3,] 3 1     1  10
```

### 7 Importing information to the population-list

#### 7.1 Insert.bve

To manually insert breeding values (type=”bve”), true genetic values (type=”bv”) or phenotypes (type=”pheno”) use the function insert.bve. Output is a modified population list. In case new phenotypes are observed this is counted as **count** observations. In case bve are changed it is assumed that genotyping was necessary unless **count** is set to 0. This is only relevant for economic calculations.

New observations are entered in the parameter **bves** with the first column coding the individual and the others containing values for the traits:

```
> bves
     Individual Name Trait 1 Trait 2
[1,] "M1_1"          "101.5" "104.2"
[2,] "M2_1"          "102"   "98.9"
[3,] "M3_1"          "99.7"  "98.4"
[4,] "M4_1"          "98.2"  "101.2"
[5,] "M5_1"          "103.8" "101.1"
> population <- insert.bve(population, bves)
```

Structure of individual names is the same in all export functions (sex [“M”/”F”], individual number [1,2,…], and generation [1,2,…]). It is recommend to just use the names as they are exported via *get.geno()* ect..

### 8 Data structure of the population list

All information regarding the breeding program are stored in a population list (R-object: list) which is modified by each run of *breeding.diploid()* and *creating.diploid()*. The population list contains matrices, inside of lists, inside of lists, inside of lists, inside of lists, inside of lists, inside of lists (you get the point!) – when understanding the structure behind it is actually not that bad, luckily you do not have to understand the structure behind for most applications since you can use exporting function discussed in section 6.

The list contains two major parts - $info ((or [[1]])) and $breeding ((or[[2]]):

#### 8.1 $info

$info contains all general information concerning genetic architecture, size of the program and internal information needed to perform the simulations. Each entry is names according to what it is supposed to contain.

| | |
|---|---|
| schlather.slot1 | Internal variable for miraculix (cre: M. Schlather) |
| chromosome | Number of chromosomes in the population list |
| snp | Number of SNPs per chromosome |
| position | Position (in Morgan) on the chromosome for each marker |
| snp.base | Major/Minor Allele (e.g. characters since internally 0/1 is used) |
| snp.position | Overall position in the genome (ongoing over chromosomes |
| length | Length of each chromosome |
| length.total | Cumulative length of chromosomes |
| func | It’s just FALSE – placeholder for later |
| size | Size of each group (generation/sex) |
| bve | Coding if breeding values are simulated |
| bv.calculated | Coding if breeding values are calculated for the founders (will be after first run of *breeding.diploid()*) |
| breeding.totals | Recap of each run of *breeding.diploid()* (if stored) |
| bve.data | Recap of each breeding value estimation (if stored) |
| bve.nr | Number of traits (with QTLs behind) to consider |

| | |
|---|---|
| bv.random | Coding which traits have underlying QTLs behind |
| bv.random.variance | Genetic variance for traits with no QTLs behind |
| snps.equidistant | Are SNPs equidistant on every chromosome (speed up!) |
| origin.gen | List of founding generations (with stored haplotypes) |
| cumsnp | Cumulative sum of SNP number (just to save computational time) |
| bp | Physical position of each marker (bp) |
| snp.name | Name of each marker |
| next.animal | ID of the next individual to generate |
| bve.mult.factor | (bv) * this |
| bve.poly.factor | (bv) ^ this |
| base.bv | This + QTL_effects |
| bv.calc | Number of total traits (including those with no QTL behind) |
| real.bv.length | Traits with (additive/multiplicative/dice-effects) |
| sex | Sex of the founders added in *creation.diploid* |
| real.bv.add | Lists with an overview of all single marker QTLs for each trait |
| real.bv.mult | Lists with an overview of all two marker QTLs for each trait |
| real.bv.dice | Lists with an overview of all three+ marker QTLs for each trait |
| pheno.correlation | Correlation matrix of the environmental variance between traits |
| bv.correlation | Correlation matrix of the genetic variance between traits (only for non-QTL traits) |
| miraculix | Coding if miraculix was used to generate the data – only miraculix users will be able to work with those population lists |
| cohorts | List of all cohorts with name and position in the population list |
| effect.p.add | Markers involved as QTL in the single QTL proportion |
| effect.p.mult1 | Markers involved as QTL in the two-SNP QTL proportion |
| effect.p.mult2 | Markers involved as QTL in the two-SNP QTL proportion |
| effect.p | Markers involved as QTL in any trait |
| store.effect.freq | Frequency of each marker in each generation |
| last.sigma.e.database | Database to derive the last used environmental variance |
| last.sigma.e.value | Last used environmental variance |
| last.sigma.e.heritabilty | Heritability assumed to derive last used environmental variance |
| comp.times | Computation times needed in each use of *breeding.diploid()* (if stored) |
| comp.times.bve | Computation times needed in the breeding value estimation in each use of *breeding.diploid()* (if stored) |
| comp.times.generation | Computation times needed in for the generation of new individuals in each use of *breeding.diploid()* (if stored) |
| Culling.stats | Information on the culling reason of each individual (mostly releveant for the web-interface) |

#### 8.2 $breeding

$breeding contains all relevant information concerning the individuals of the breeding scheme. For efficiency purposes a lot of this is internally coded or computed on-the-fly.

Individuals are sorted according to generation, sex and individual number. In case data has to be stored for both male and female (or father/mother) there will be two entries with the first one being the male (Have to talk with the equality commissioner about that!).

$breeding[[generation]][[sex]][[ individual nr.]] (( or [[2]][[generation]][[sex]][[ individual nr.]]))

##### 8.2.1 Storage per generation

| | |
|---|---|
| $breeding[[generation]][[3,4]] | Estimated breeding values of males (3) and females (4) |
| $breeding[[generation]][[5,6]] | Class of males (5) and females (6) |
| $breeding[[generation]][[7,8]] | Underlying "true" genetic values of males (7) and females (8) |
| $breeding[[generation]][[9,10]] | Observed phenotypes for males (9) and females (10) |
| $breeding[[generation]][[11,12]] | Time point of generation for male (11) and females (12) |
| $breeding[[generation]][[13,14]] | Creating type of generation for males(13) and females (13)<br>This is only relevant for the web-based application |
| $breeding[[generation]][[15,16]] | Individual IDs for male (15) and females (16) |
| $breeding[[generation]][[17,18]] | Time of culling for male(17) and female (18) individuals |
| $breeding[[generation]][[19,20]] | Reliability estimated for male (19) and females (20) |
| $breeding[[generation]][[21,22]] | Last applied selection index (mostly relevant for complex selection indices like (Miesenberger 1997) |

```
> str(population$breeding[[1]])
 .. [list output truncated]
$ : num [1, 1:100] 46 48.9 31.5 35.7 49.6 ...
$ : num [1, 1:1000] 67.6 57.4 68.8 53 39.6 ...
$ : int [1:100] 0 0 0 0 0 0 0 0 0 0 ...
$ : int [1:1000] 0 0 0 0 0 0 0 0 0 0 ...
$ : num [1, 1:100] 33.4 47.5 31.9 59.1 38 ...
$ : num [1, 1:1000] 77.79 54.65 63.42 54.49 7.45 ...
$ : num [1, 1:100] -22.23 67.24 -9.34 -16.06 101.5 ...
$ : num [1, 1:1000] 80.4 48.1 96.2 37.1 30.5 ...
```

##### 8.2.2 Storage per individual

$breeding[[generation]][[sex]][[ individual nr.]]…

| | |
|---|---|
| [[1,2]] | Points of recombination on the first (1) and second (2) chromosome set |
| [[3,4]] | Points of mutations |
| [[5,6]] | Efficiently stored origins of segments between two points of recombination. Decoding using *decodeOrigins()* (miraculix) / *decodeOriginsR()* (else).<br>Output in *get.recombi()* is automatically decoded |
| [[7,8]] | Father / Mother |
| [[9,10]] | Efficiently stored haplotypes (if it is a founder – else empty) |
| [[11,12]] | Storage of duplications (long not used!) |

| | |
|---|---|
| [[13,14]] | Storage of history of recombinations |
| [[15]] | How often a phenotype was generated for the individual |
| [[16]] | Is the individual genotyped |
| [[17]] | True breeding value before gene editing |
| [[18]] | Generation of death and previous class |
| [[19]] | Share of the genetic material of the grandfather of the father inherited |
| [[20]] | Share of the genetic material of the grandfather of the mother inherited |
| [[21]] | List of all individuals with the same id. |

```
> str(population$breeding[[1]][[1]][[1]])
List of 21
 $ : num [1:6] 0 0.2 0.4 0.6 0.8 1
 $ : num [1:6] 0 0.2 0.4 0.6 0.8 1
 $ : NULL
 $ : NULL
 $ : int [1:5] 0 0 0 0 0
 $ : int [1:5] 1 1 1 1 1
 $ : num [1:3] 1 1 1
 $ : num [1:3] 1 1 1
 $ :Haplotype information at 1000 loci.
Attributes are a List of 3
 $ information: int [1:22] 0 1000 1 530385344 0 530385344 0 0 0 NA ...
 $ method     : int 8
 $ class      : chr "haplomatrix"
 $ : chr "Placeholder_Pointer_Martin"
 $ : NULL
 $ : NULL
 $ : NULL
 $ : NULL
 $ : num [1:21] 0 0 0 0 0 0 0 0 0 0 ...
 $ : int 0
 $ : NULL
 $ : NULL
 $ : NULL
 $ : NULL
 $ : num [1:2, 1:3] 1 2 1 1 1 63
  ..- attr(*, "dimnames")=List of 2
  .. ..$ : NULL
  .. ..$ : chr [1:3] "generation" "sex" ""
```

### 9 Utility functions

Since the inclusion of miraculix, I did not need many utility functions any more – older version of function to show development of allele frequencies and others still exists but need modifications. Just tell me if you wish to have a certain function to help you with the package.

#### 9.1 bv.development

This function will generate a plot showing the development of breeding values and phenotypes over generations. 95% confidence bands are included in a dotted line:

Which groups to display is selected via the parameters **gen**, **database** & **cohorts**. In case the user interface was used to generate the population list set **json** to TRUE to automatically display all selected cohorts. Confidence bands are drawn for "bv" (1), "bve" (2) & "pheno"(3) – to change the quantile use the parameter **quantile** (default: 0.95), to exclude selected the for with to draw a confidence band via the parameter **confidence** (default: c(1,2,3)). Groups with only zeros are ignored on default – if you want lines to be included for all selected cohorts set **ignore.zero** to FALSE.

To display the time point, the creating type, the sex, and cohort name set **display.time.point**, **display.creating.type**, **display.cohort.name**, **display.sex** to TRUE. In case the generating interface between groups is highly heterozygous it might be useful to use **equal.spacing** between displayed cohorts. To not display the line displaying a long-term trend of the breeding values set **display.line** to FALSE.

```
population <- json.simulation(total = ex_json)
bv.development(population, json=TRUE, bvrow=1, confidence = 1, development = 1,
               display.creating.type = TRUE, display.sex = TRUE,
               display.cohort.name = TRUE)
```

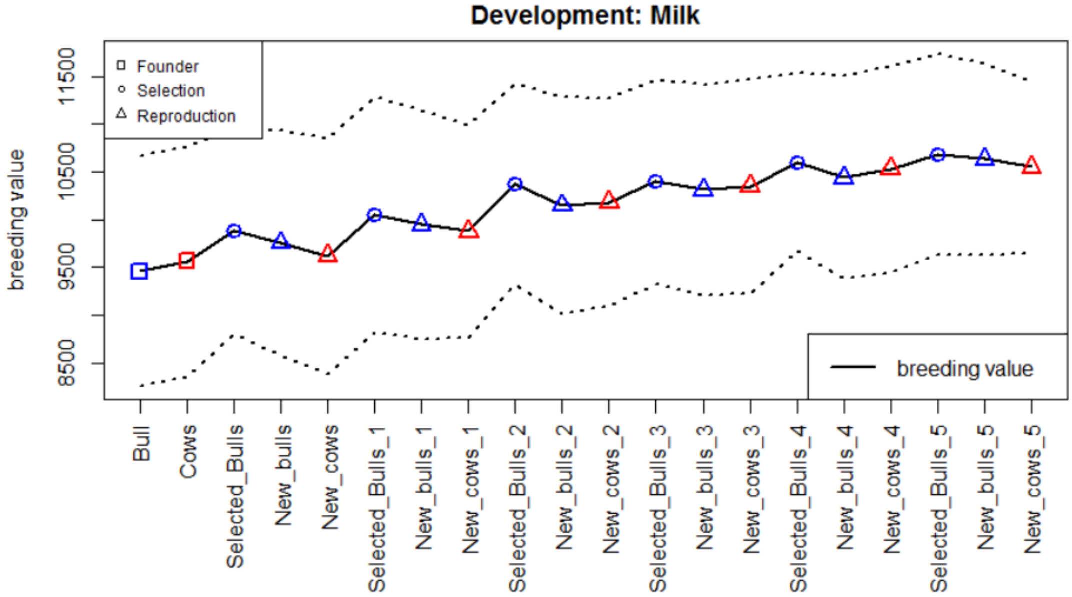

#### 9.2 bv.development.box

This function will generate a plot displaying the development of breeding values with a boxplot for each selected **gen**, **database**, **cohorts**. In case the user-interface was used to generate the population set **json** to TRUE to automatically display all selected cohorts. To only display a subset of trait set **bvrow** to those traits. To display phenotypes or breeding value estimations for the individuals instead of breeding values, set **display** to "pheno" / "bve".

In case the user interface was used to generate the population, one can display which cohorts where generated by which other cohort (via selection or reproduction) by setting **display.selection** and **display.reproduction** to TRUE.

```
population <- json.simulation(total = ex_json)
bv.development.box(population, json = TRUE, bvrow = 1)
```

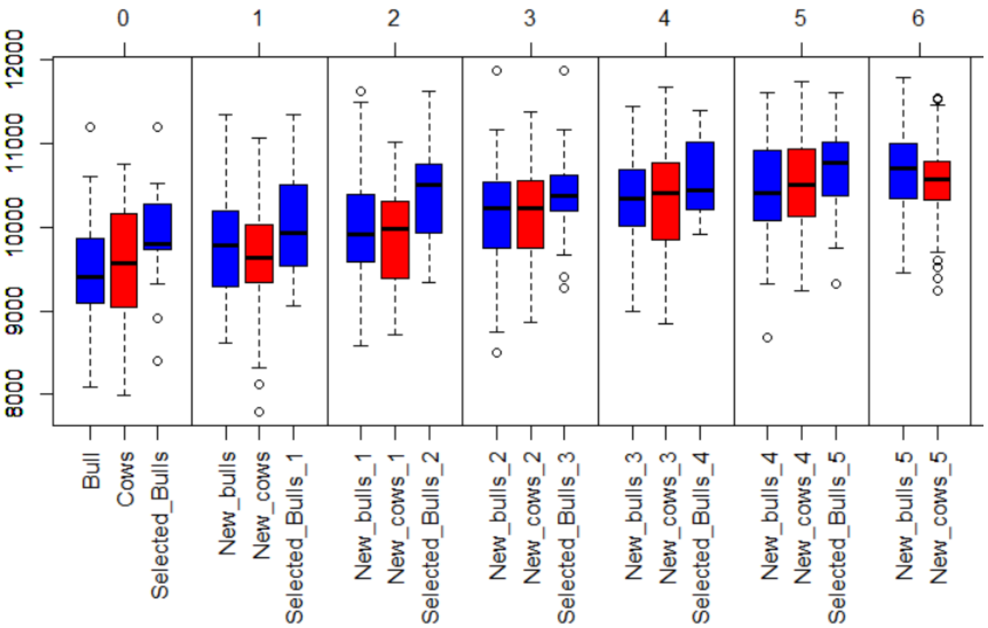

#### 9.3 Kinship.development

Function to display the development of kinship over different gen, database, cohorts. Internally kinship.emp.fast is used and same optional parameters can be used to improve computation time / accuracy.

```
population <- json.simulation(total = ex_json)
kinship.development(population, json = TRUE, display.cohort.name = TRUE)
```

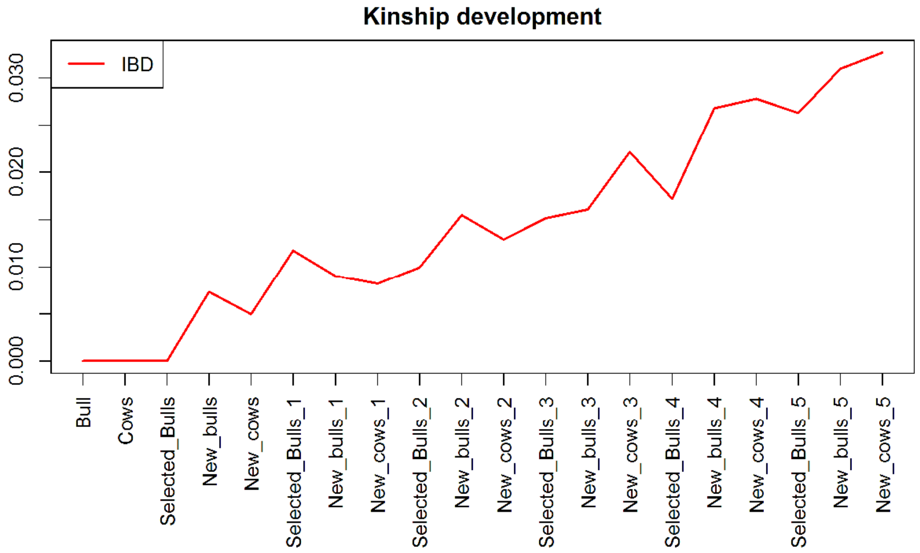

#### 9.4 Kinship.emp / kinship.emp.fast

These functions can be used to derive empirical kinship between a set of individuals. Either directly supply a list containing all stored information for the respective individuals via the parameter **animals** or selected them by usage of **gen**, **database**, **cohorts**. In that case the **population** list needs to also be provided.

On default this call lead to all pairwise relations being evaluated. For a quick evaluation use *kinship.emp.fast()* and provide the total number of pairwise relationships (**ibd.obs**) and relations with the individual with itself (**hbd.obs**).

#### 9.5 Kinship.exp

This function can be used to derive the expected kinship between individuals. To how many generations are used back use **prev.gen**. On default it’s assumed all individuals before are unrelated. Alternatively, one can provide a **kinship.matrix** the individuals of the first generation via **first.individuals**. It should be noted that there are more efficient ways to derive a pedigree matrix than this – alternatively one can export the pedigree via *get.pedigree()* and use that as input for breedR or tools outside of R.

#### 9.6 analyze.population

With this function, one can analyze the allele frequency of a specific marker over time. Select the marker to analyze via parameters **chromosome** & **snp**. To selected with generations to compare use **gen**, **database**, **cohorts**.

```
population <- json.simulation(total = ex_json)
analyze.population(population,5,2, gen=1:8)
```

```
> analyze.population(population,5,2, gen=1:8)
     [,1] [,2] [,3] [,4] [,5] [,6] [,7] [,8]
[1,]  198   27  220  200  207  151  162  143
[2,]  214   12  278  278  250  264  269  239
[3,]   88   11   52   72   93  135  119  118
```

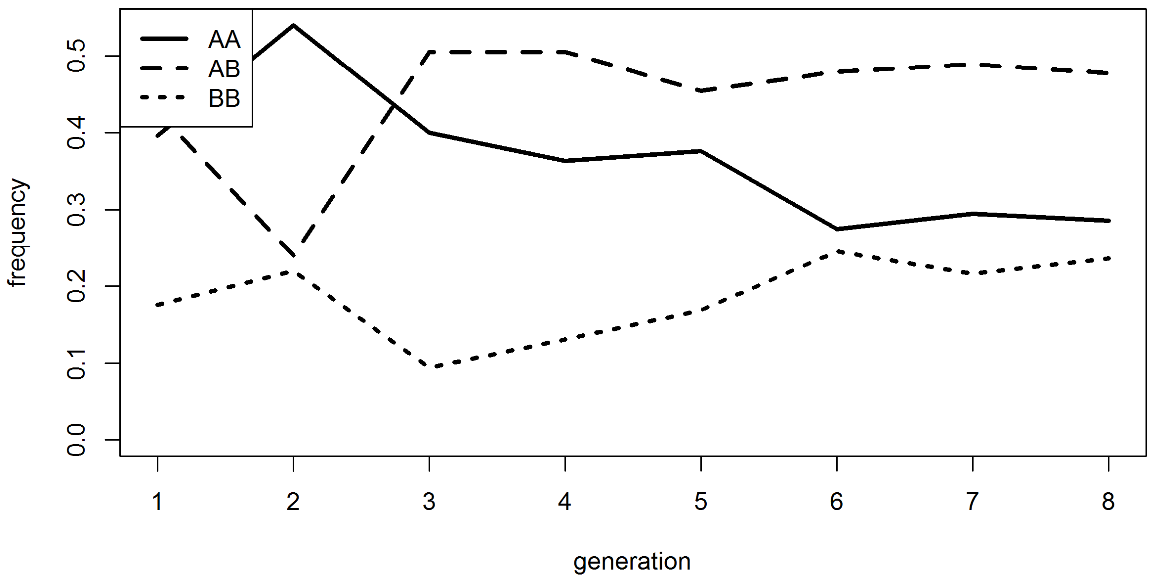

#### 9.7 new.base.generation

With rising number of generations the number of points of recombinations to store is increasing. For efficient storing it can make sense to compute and store haplotypes for a later generation and use those individuals as a new founder generation. For this use *new.base.generation()* and select the new base generation via the parameter **base.gen**. To further reduce memory needs and computation time you can additionally delete data of previous generations via **delete.previous.gen**, **delete.breeding.totals** and **delete.bve.data**.

#### 9.8 creating.trait

With this function one can generate additional traits for the base population without the need to add genetic datasets. Functionality is the same as *creating.diploid()* otherwise.

#### 9.9 ensembl.map

Via this functions genetic maps provided in Ensembl (https://www.ensembl.org/index.html & http://plants.ensembl.org/index.html) can be imported. Internally the package biomaRt is used – for guidelines on how to install this package we refer to https://bioconductor.org/packages/release/bioc/html/biomaRt.html.

Naming of parameters is orientated according to the biomaRt package. Set **dataset** to the dataset you want to access (e.g. for cattle-SNPs: "btaurus_snp") – for a list of possible datasets run this function with **export.datasets** set to TRUE.

To import a subset of all markers use the parameter filter and **filter.values**. To limited the markers to a specific SNP-chip just set **filter.values** to the name of the chip (e.g. **filter.values**="Illumina BovineSNP50 BeadChip"). Names of potential filters for a dataset can be exported by setting **export.filters** to TRUE.

For us the direct export in R via Ensembl was painfully slow and the package did not always do what we intended it to do. Therefore, we are providing exemplary map files for most common species in the associated R-package MoBPSmaps. For a list of map that are already included in MoBPS and the associated data package MoBPS_maps we refer to section 12.

```
cattle_map <- ensembl.map(dataset="btaurus_snp", filter.values="Illumina BovineSNP50 BeadChip")
```

```
> cattle_map[1:10,]
      Chromosome SNP-ID        bp        M  freq
 [1,] "1"        "rs42778024"  "435963"  NA NA
 [2,] "1"        "rs41609588"  "776231"  NA NA
 [3,] "1"        "rs108982244" "907810"  NA NA
 [4,] "1"        "rs29026917"  "1073496" NA NA
 [5,] "1"        "rs29015852"  "1150763" NA NA
 [6,] "1"        "rs108981857" "1566539" NA NA
 [7,] "1"        "rs108994381" "1695632" NA NA
 [8,] "1"        "rs41635940"  "1929664" NA NA
 [9,] "1"        "rs41255293"  "2082139" NA NA
[10,] "1"        "rs41580909"  "2235492" NA NA
```

#### 9.10 compute.costs

To calculate the costs of the currently simulated breeding program use the function *compute.costs()*. Currently implemented cost factors include the following:

| Cost factor | MoBPS -Parameter | Default |
|---|---|---|
| Phenotyping | phenotyping.costs | 10 |
| Genotyping | genotyping.costs | 100 |
| Housing/Field costs | Housing.costs | 0 |
| Fixed costs | fix.costs | 0 |
| Annual costs | fix.costs.annual | 0 |
| Profit per BV | profit.per.bv | 1 |

Note that all default settings are basically chosen at random and should be modified when analyzing a real breeding program. In case costs/gains between sexes are different, use a vector. To separate between generations use a matrix with each row coding costs/gains per generation.

To only calculate the resulting costs of some generations/cohorts use database/gen.

To model an interest rate set **interest.rate** (default: 1 – meaning i = 0%. We here assume inputs of the form 1 + i) – with costs/gain changed according to a base generation (**base.gen** – default: 1)

```
> compute.costs(population,gen=1:10)
 [1]  -63014.02 -242252.85 -274059.21 -144566.68   12944.92  132774.58  165916.47  299725.18  339646.66  510074.31
```

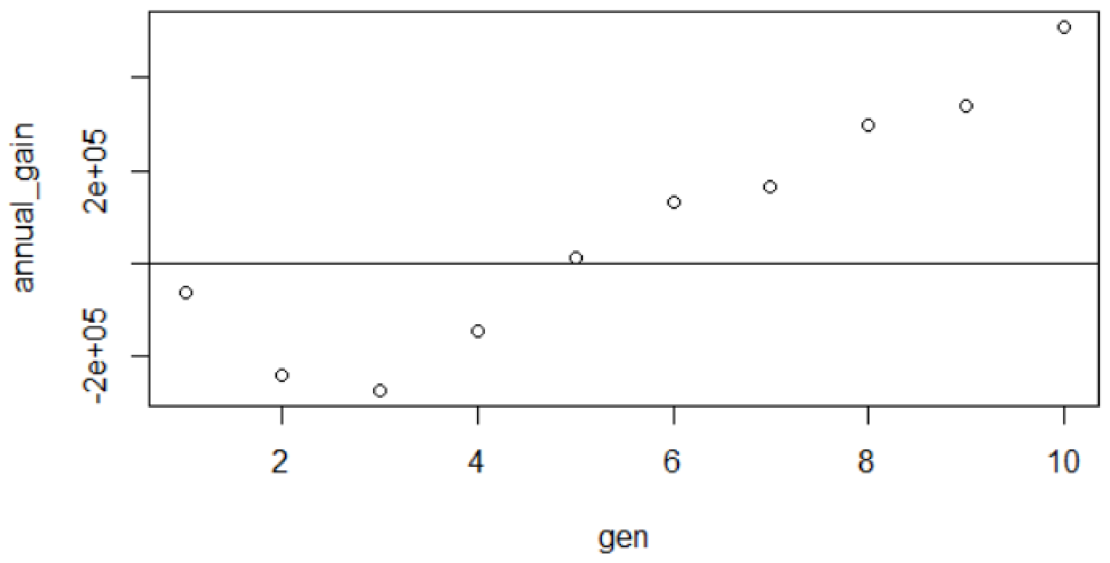

#### 9.11 compute.costs.cohorts

The functionally of *compute.costs.cohorts()* is similar to *compute.costs()* with the added benefit of usability with input parameters provided in our user-interface. Input parameters include **phenotyping.costs**, **genotyping.costs**, **housing.costs**, **fix.costs**, **fix.costs.annual**, **profit.per.bv**, **interest.rate**. In case the user interface is provided considered groups (**gen**, **database**, **cohorts**) are automatically assigned to their time point of creation for discounting.

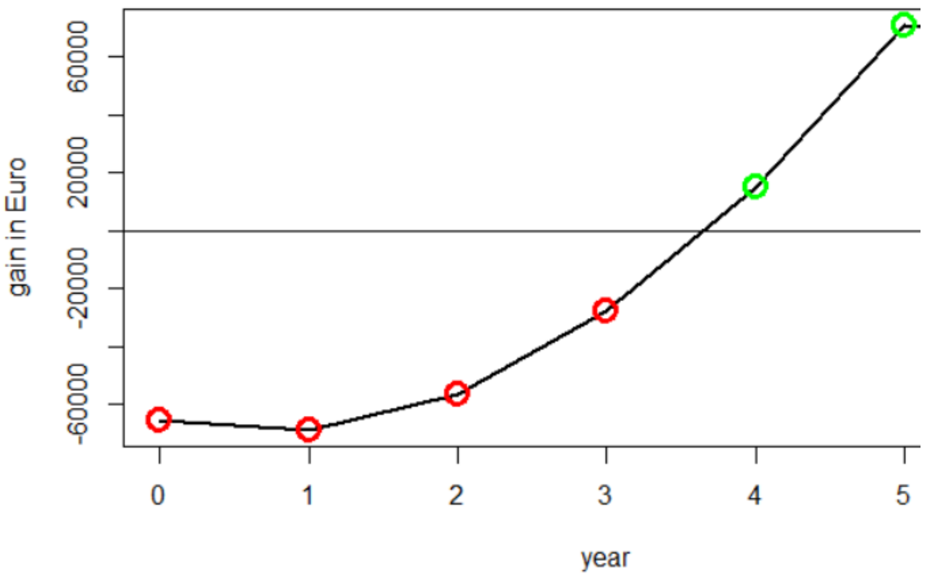

#### 9.12 summary

The population list generated via *creating.diploid()* and *breeding.diploid()* is of class "population". Application of the generic function summary leads to an overview of the population list including the number of individuals, cohorts, structure of the genome and traits:

```
> summary(pop)
Population size:
Total: 100 Individuals
Of which 50 are male and 50 are female.
There are 2 generations
and 4 unique cohorts.

Genome Info:
There are 5 unique chromosomes.
In total there are 10000 SNPs.
The genome has a total length of 10 Morgan.
No physical positions are stored.

Trait Info:
There is 1 modelled trait.
The trait has underlying QTL
The trait is named: Trait 1
```

#### 9.13 pedmap.to.phasedbeaglevcf

The standard input of MoBPS are haplotypes (not genotypes!). In case of using own genomic data it is highly encourages to perform genomic phased before using the dataset as an input. In this function a routine pipeline to generate a phased dataset is executed. In this pipeline BEAGLE 5.0 (https://faculty.washington.edu/browning/beagle/beagle.html) and PLINK 1.9 (https://www.cog-genomics.org/plink/1.9/) are used. Additional file are generated in a selected directory. Path for all three have to be provided in **beagle_jar**, **plink_dir**, **db_dir**. Input can either be a dataset in PLINK format (**ped_path**, **map_path**) or a vcf-file (**vcf_path**). Defaults are all chosen to work on the webserver for our web-interface (Chapter 14). Phasing can also be directly performed in the web-interface.

### 10 Memory and computation times

Critical parts of MoBPS concerning memory requirements and computation times can be performed using the associated R-package miraculix. By using SSE2 operations and bit-wise storing computation speed can be massively increased leading to about 10 times faster matrix multiplications than the regular R implementation while needing only 1/16 of the regularly needed memory.

To speed up commutation of the breeding value estimation one can use multiple cores by the usage of **miraculix.cores** (default: 1) or in case miraculix is not active **ncore**. To parallelize generation of new individuals set **parallel.generation** to TRUE and set the number of cores used via **ncore.generation**. This will only lead to significant improvement in computation time for the generation of a lot of individuals. Even when using a single core ~1'000 individuals are generated per second. The packages doParallel (Microsoft Corporation and Steve Weston 2018) and doRNG (Renaud Gaujoux 2018) are used for parallelization in R.

### 11 List of input parameters in breeding.diploid()

For a description of each parameter we refer to the use of the help function in R (?*breeding.diploid*) and/or other sections of this Guidelines.

| **Parameter** | **Default** | **options** |
|---|---|---|
| population | NULL | A previous population list |
| mutation.rate | 10^-5 | Value between 0 and 1 |
| remutation.rate | 10^-5 | Value between 0 and 1 |
| recombination.rate | 1 | Any positive numeric |
| selection.m | "random" | "function" |
| selection.f | NULL (selection.m) | "random", "function" |
| new.selection.calculation | TRUE | FALSE |
| selection.function.matrix | NULL | Don't touch – will be removed |
| selection.size | C(0,0) | Vector with two non-negative values |
| breeding.size | 0 | Positive number // 2 element vector (male/female) |
| breeding.sex | 0.5 | Value between 0 and 1 |
| breeding.sex.random | FALSE | TRUE |
| class.m | 0 | Vector with all classes to consider |
| class.f | NULL (class.m) | Vector with all classes to consider |
| add.gen | 0 (will lead to added generation) | Value between 1 and number of generations |

| | | |
|---|---|---|
| recom.f.indicator | NULL | Not necessary (use modified marker position instead!) |
| recom.f.polynom | NULL | Not necessary (use modified marker position instead!) |
| duplication.rate | 0 | |
| duplication.length | 0.01 | Duplication modelling needs changes! |
| duplication.recombination | 1 | |
| same.sex.active | FALSE | TRUE |
| new.class | 0 | Numeric value (ideally positive integer; -1 is reserved for dead individuals) |
| bve | FALSE | TRUE |
| bve.direct.est | TRUE | FALSE |
| bve.gen | NULL | 1:3 |
| bve.database | NULL | Generation sex<br>[1,] 1 2<br>[2,] 5 1 |
| bve.cohorts | NULL | c("Founder_M", "F1") |
| bve.avoid.duplicates | TRUE | FALSE |
| report.accuracy | TRUE | FALSE |
| sigma.e | NULL | Numeric value above 0 |
| sigma.s | 100 | Numeric value above 0 |
| new.bv.observation | NULL | "all" for all individuals<br>"non_obs" for all previously not observed<br>"non_obs_m" for all previously not observed male individuals<br>"non_obs_f" for all previously not observed female individuals |
| new.bv.observation.gen | NULL | 1:3 |
| new.bv.observation.database | NULL | Generation sex<br>[1,] 1 2<br>[2,] 5 1 |
| new.bv.observation.cohorts | NULL | c("Founder_M", "F1") |
| new.bv.child | "zero" | "mean", "obs" |
| computation.A | "vanRaden" | "kinship", "CE", "CM", "non_stand" |
| depth.pedigree | 7 | Positive Integer |
| delete.haplotypes | NULL | Vector of all generations to delete (natural number) |
| delete.individuals | NULL | Vector of all generations to delete (natural number) |

| | | |
|---|---|---|
| fixed.breeding | NULL | matrix with each row containing (gen1,sex1,nr1, gen2,sex2,nr2, sex.probability) with 1 being father, 2 being mother) |
| fixed.breeding.best | NULL | matrix with each row containing (sex1, nr1, sex2, nr2, sex.probability) chosen from the group of selected individuals |
| max.offspring | C(Inf,Inf) | vector with two natural numbers (first male, second female) |
| store.breeding.totals | FALSE | TRUE |
| forecast.sigma.s | TRUE | FALSE |
| multiple.bve | “add” | “ranking” |
| multiple.bve.weights.m | 1 | Any weights – use a vector with length equal to number of traits |
| multiple.bve.weights.f | NULL (multiple.bve.weights.m) | Any weights – use a vector with length equal to number of traits |
| store.bve.data | FALSE | TRUE |
| fixed.assignment | FALSE | “bestworst”, “worstbest” |
| reduce.group | NULL | Per row: Generation, Sex, Individuals to survive, class of individuals |
| reduce.group.selection | “random” | “function” |
| selection.criteria | c(TRUE,TRUE) | C(FALSE/TRUE,FALSE/TRUE) |
| same.sex.sex | 0.5 | Numeric value between 0 and 1 |
| same.sex.selfing | TRUE | FALSE |
| selfing.mating | FALSE | TRUE |
| selfing.sex | 0.5 | Numeric value between 0 and 1 |
| praeimplantation | NULL | No use currently recommended |
| sigma.e.gen | NULL | 1:3 |
| sigma.e.database | NULL | ` Generation sex`<br>`[1,] 1 2`<br>`[2,] 5 1` |
| sigma.e.cohorts | NULL | c(“Founder_M“, “F1“) |
| heritability | NULL | Numeric value between 0 and 1 |
| multiple.bve.scale.m | FALSE | TRUE |
| multiple.bve.scale.f | NULL (multiple.bve.scale.m) | TRUE |
| use.last.sigma.e | FALSE | TRUE |
| save.recombination.history | FALSE | TRUE |
| martini.selection | FALSE | TRUE |

| | | |
|---|---|---|
| BGLR.bve | FALSE | TRUE |
| BGLR.model | “RKHS” | “BRR”, “BL”, “BayesA”, “BayesB”, “BayesC” |
| BGLR.burnin | 500 | natural number |
| BGLR.iteration | 5000 | natural number |
| BGLR.save | “RKHS” | any path you want |
| BGLR.save.random | FALSE | TRUE |
| BGLR.print | FALSE | TRUE |
| copy.individual | FALSE | TRUE |
| copy.individual.keep.bve | TRUE | FALSE |
| dh.mating | FALSE | TRUE |
| dh.sex | 0.5 | Numeric value between 0 and 1 |
| offspring.bve.parents.gen | NULL | 1:3 |
| offspring.bve.parents.database | NULL | Generation sex<br>[1,] 1 2<br>[2,] 5 1 |
| offspring.bve.parents.cohorts | NULL | c(“Founder_M“, “F1“) |
| offspring.bve.offspring.gen | NULL | 1:3 |
| offspring.bve.offspring.database | NULL | Generation sex<br>[1,] 1 2<br>[2,] 5 1 |
| offspring.bve.offspring.cohorts | NULL | c(“Founder_M“, “F1“) |
| bve.parent.mean | FALSE | TRUE |
| bve.grandparent.mean | FALSE | TRUE |
| bve.mean.between | “bvepheno” | „bve“, „pheno“, „bv“ |
| n.observation | 1 | Natural number |
| share.genotyped | 1 | Numeric value between 0 and 1 |
| added.genotyped | 0 | Numeric value between 0 and 1 |
| remove.non.genotyped | TRUE | FALSE |
| Singlestep.active | FALSE | TRUE |
| bve.0isNA | TRUE | FALSE |
| phenotype.bv | FALSE | TRUE |
| standardize.bv | FALSE | TRUE |
| standardize.bv.level | 100 | Numeric value |
| standardize.bv.gen | 1 | Natural number <= generation number |
| delete.same.origin | FALSE | TRUE |
| remove.effect.position | FALSE | TRUE |
| estimate.u | FALSE | TRUE |
| fast.uhat | TRUE | FALSE |

| | | |
|---|---|---|
| new.phenotyp.correlation | NULL | Positive definite matrix |
| new.breeding.correlation | NULL | Positive definite matrix |
| recalculate.bv.var.correlation | FALSE | TRUE |
| new.bv.random.correlated | TRUE | FALSE |
| estimate.add.gen.var | FALSE | TRUE |
| estimate.pheno.var | FALSE | TRUE |
| selection.m.gen | NULL | 1:3 |
| selection.f.gen | NULL | 1:3 |
| selection.m.database | NULL | Generation sex<br>[1,] 1 2<br>[2,] 5 1 |
| selection.f.database | NULL | Generation sex<br>[1,] 1 2<br>[2,] 5 1 |
| selection.m.cohorts | NULL | c("Founder_M", "F1") |
| selection.f.cohorts | NULL | c("Founder_M", "F1") |
| best1.from.group | NULL | Matrix with one group per row |
| best2.from.group | NULL | Matrix with one group per row |
| best1.from.cohort | NULL | Vector containing names of cohorts |
| best2.from.cohort | NULL | Vector containing names of cohorts |
| Reduced.selection.panel.m | NULL | Vector containing numeric values |
| Reduced.selection.panel.f | NULL | Vector containing numeric values |
| store.comp.times | TRUE | FALSE |
| store.comp.times.bve | TRUE | FALSE |
| special.comb | FALSE | Part of martini selection – do not use! |
| max.auswahl | Inf | Part of martini selection – do not use! |
| predict.effects | FALSE | Part of martini selection – do not use! |
| SNP.density | 10 | Part of martini selection – do not use! |
| use.effect.markers | FALSE | Part of martini selection – do not use! |
| use.effect.combination | FALSE | Part of martini selection – do not use! |
| import.position.calculation | NULL | Function f(cm_position) = Last previous SNP |

| | | |
|---|---|---|
| special.comb.add | FALSE | Part of martini selection – dont use! |
| ogc | FALSE | TRUE |
| ogc.cAc | NA (minimal gain in inbreeding) | Numeric value between 0 and 1 |
| computation.A.ogc | “kinship” | “vanRaden” (see computation.A) |
| depth.pedigree.ogc | 7 | Positive Integer |
| emmreml.bve | FALSE | TRUE |
| nr.edits | 0 | any natural number |
| gene.editing | FALSE | TRUE |
| gene.editing.offspring | FALSE | TRUE |
| gene.editing.best | FALSE | TRUE |
| gene.editing.offspring.sex | c(TRUE,TRUE) | Vector with two boole variables |
| gene.editing.best.sex | c(TRUE,TRUE) | vector with two boole variables |
| gwas.u | FALSE | TRUE |
| approx.residuals | TRUE | FALSE |
| sequenceZ | FALSE | TRUE |
| maxZ | 5000 | Any natural number |
| maxZtotal | 0 | Any natural number |
| gwas.gen | NULL | 1:3 |
| gwas.database | NULL | `Generation sex`<br>`[1,] 1 2`<br>`[2,] 5 1` |
| gwas.cohorts | NULL | c(“Founder_M“, “F1“) |
| delete.sex | c(1,2) | 1 (male), 2 (female) |
| gwas.group.standard | FALSE | TRUE |
| y.gwas.used | “pheno” | “bv“, “bve“ |
| culling.cohort | NULL | Any cohort name |
| culling.time | Inf | Numeric value |
| culling.name | “Not_named” | Any character string |
| culling.bv1 | 100 | numeric value |
| culling.share1 | 0 | Probability between 0 and 1 |
| culling.bv2 | 110 | numeric value |
| culling.share2 | 0 | Probability between 0 and 1 |
| culling.index | 0 | Any weights – use a vector with length equal to number of traits, “lastindex” |
| gen.architecture.m | 0 | Natural number (select one of the previously stored architectures) |

| | | |
|---|---|---|
| gen.architecture.f | NULL (gen.architecture.m) | Natural number (select one of the previously stored architectures) |
| ncore | 1 | Natural number |
| Z.integer | FALSE | TRUE |
| store.effect.freq | TRUE | FALSE |
| backend | “doParallel” | „doMPI“ |
| randomSeed | NULL | natural number |
| randomSeed.generation | NULL | Natural number |
| Rprof | FALSE | TRUE |
| miraculix | FALSE | TRUE (automatically activated when miraculix is used is *creating.diploid()*) |
| miraculix.mult | NULL (leading to FALSE) | TRUE / FALSE |
| fast.compiler | 0 | 3 (For R >= 3.4 this is default in R) |
| miraculix.cores | 1 | natrual number |
| store.bve.parameter | FALSE | TRUE |
| print.error.sources | FALSE | TRUE |
| chol.miraculix | FALSE | TRUE |
| bve.insert.gen | NULL | 1:3 |
| bve.insert.database | NULL | Generation sex<br>[1,] 1 2<br>[2,] 5 1 |
| bve.insert.cohorts | NULL | c(“Founder_M“, “F1“) |
| best.selection.ratio.m | 1 | positive numeric value |
| best.selection.ratio.f | NULL (best.selection.ratio.m) | positive numeric value |
| best.selection.criteria.m | “bv” | “bve“, “pheno“ |
| best.selection.criteria.f | NULL (best.selection.criteria.m) | “bve“, “pheno“ |
| best.selection.manual.ratio.m | NULL | positive numeric value |
| best.selection.manual.ratio.f | NULL (best.selection.manual.ratio.m) | positive numeric value |
| bve.class | NULL (take all!) | vector containing numeric values |
| parallel.generation | FALSE | TRUE |
| ncore.generation | 1 | Positive numeric value |
| name.cohort | NULL | “Founders” or any other character string |
| add.class.cohorts | TRUE | FALSE |
| display.progress | TRUE | FALSE |

| | | |
|---|---|---|
| ignore.best | C(0,0) | Any two element vector (first male, second female) |
| combine | FALSE | TRUE |
| repeat.mating | 1 | Positive numeric value |
| time.point | 0 | Positive numeric value (this will be automatically processed in the web-based-application) |
| creating.type | 0 | This is automatically stored in the web-based-application<br># 0 – Founder<br># 1 – Selection<br># 2 – Reproduction<br># 3 – Recombination<br># 4 – Selfing<br># 5 – DH-Production<br># 6 – Cloning<br># 7 – Combine<br># 8 – Aging<br># 9 – Split |

### 12 List of input parameters in creating.diploid()

For a description of each parameter we refer to the use of the help function in R (?*creating.diploid*) and/or other sections of this Guidelines.

| **Parameter** | **Default** | **options** |
|---|---|---|
| population | NULL (will lead to “random”) | A previous population list |
| dataset | “random” | SNP-dataset (One haplotype per colum), “random”, “all0”, “homorandom”, “allhetero” |
| nsnp | 0 | Positive integer value |
| nindi | 0 | Positive integer value |
| vcf | NULL | Path to a vcf-file |
| map | NULL | Matrix with up to 5 colums containing (chr.nr, snp.name, bp, position in Morgan, allele freq). Rest will be set to NULL/NA.<br>For more see section 13 |
| chr.nr | NULL (all markers on the same chromosome) | Vector containing the chromosome for each generated marker, or natural number with the number of chromosomes |
| bp | NULL | Vector containing the base-pair for each generated marker |

| | | |
|---|---|---|
| snp.name | NULL | Vector containing the snp-name for each generated marker |
| bpcm.conversion | 0 | Recommendations:<br>For human: 100.000.000<br>For chicken: 30.000.000 |
| chromosome.length | NULL (will lead to 5M) | Positive numeric value |
| freq | "beta" | Numeric value or vector for each marker |
| beta.shape1 | 1 | Positive numeric value |
| beta.shape2 | 1 | Positive numeric value |
| sex.s | "fixed" | "random", vector containing the sex of each newly added individual. |
| share.genotyped | 1 | Numeric value between 0 and 1 |
| genotyped.s | NULL | vector containing the sex of each newly added individual. |
| add.chromosome | FALSE | TRUE |
| generation | 1 | Positive integer value (no empty generations inbetween!) |
| class | 0 | Numeric value (positive integer recommended) |
| sex.quota | 0.5 | Numeric value between 0 and 1 |
| snps.equidistant | NULL (will be TRUE if no other way to derive Morgan-position is provided) | FALSE/TRUE |
| change.order | FALSE | TRUE |
| position.scaling | FALSE | TRUE |
| length.before | 5 | Positive numeric value |
| length.behind | 5 | Positive numeric value |
| hom0 | NULL (automatically derived) | Vector containing major allele for each generated marker. |
| hom1 | NULL (automatically derived) | Vector containing minor allele for each generated marker. |
| miraculix | TRUE | FALSE |
| bit.storing | FALSE | TRUE (this is less efficient than miraculix but does not rely on C-code) |
| nbits | 30 | Integer value between 1 and 30 |
| bv.total | 0 (automatically set according to traits provided) | Integer value. If higher than the number of traits simulate traits based on pedigree/inbreeding rates |
| trait.name | NULL | Vector containing the names of the traits (e.g. "milk") |
| real.bv.add | NULL | List with each element containing effect matrices |

| | | |
|---|---|---|
| real.bv.mult | NULL | List with each element containing effect matrices |
| real.bv.dice | NULL | List with each element containing effect lists |
| n.additive | 0 | Positive integer value |
| n.dominant | 0 | Positive integer value |
| n.qualitative | 0 | Positive integer value |
| n.quantitative | 0 | Positive integer value |
| var.additive.l | NULL | List containing a single numeric value or vector with variances for each trait |
| var.dominant.l | NULL | List containing a single numeric value or vector with variances for each trait |
| var.qualitative.l | NULL | List containing a single numeric value or vector with variances for each trait |
| var.quantitative.l | NULL | List containing a single numeric value or vector with variances for each trait |
| exclude.snps | NULL | Vector containing marker positions with no simulated random effects |
| shuffle.traits | NULL | TRUE |
| shuffle.cor | NULL | Correlation matrix for the traits to shuffle |
| replace.real.bv | FALSE | TRUE |
| name.cohort | NULL | Character string |
| skip.rest | FALSE | TRUE (INTERNAL PARAMETER!) |
| randomSeed | NULL | Integer value |
| template.chip | NULL | "cattle", "chicken", "pig", "sheep", "maize" |
| time.point | 0 | Positive numeric value (this will be automatically processed in the web-based-application) |
| creating.type | 0 | This is automatically stored in the web-based-application<br># 0 – Founder<br># 1 – Selection<br># 2 – Reproduction<br># 3 – Recombination<br># 4 – Selfing<br># 5 – DH-Production<br># 6 – Cloning<br># 7 – Combine<br># 8 – Aging<br># 9 – Split |

| | | |
|---|---|---|
| remove.invalid.qtl | TRUE | FALSE |

### 13 List of datasets included in the package

MoBPS does contain a variety of maps that are preimported from Ensembl since the actual import takes quite long for bigger map-files. In case you feel a certain map is missing feel free to contact us to we can add it to the tool. Maps are available in the associated R-package MoBPS_maps. Only map_chicken1, map_cattle1 and map_maize1 are included in MoBPS itself. To use a specific map use it as an input for the parameter **map** in *creating.diploid()*.

In addition to all those maps an exemplary json-file (**ex_json**) generated by a recent version of our interface is included for text use in *json.simulation()* and other function that utilize datasets generated by *json.simulation()*. Note that this file is automatically generated via the user-interface and you do not have to worry about its structure.

| Dataset name | Corresponding Chip | Number of Markers | Contains:<br>1. Physical position<br>2. Morgan position<br>3. allele frequency |
|---|---|---|---|
| Map_pig1 | Axiom Genotyping Array | 590‘318 | |
| Map_pig2 | GGP Porcine HD | 63’113 | |
| Map_pig3 | GGP Porcine LD | 8’624 | |
| Map_pig4 | Illumina_PorcineSNP60 | 55‘684 | |
| Map_chicken1 | Affymetrix Chicken600K Array | 547’024 | |
| Map_chicken2 | Affymetrix Chicken600K Array (diversity subset) | 293’251 | |
| Map_chicken3 | Affymetrix Chicken600K Array (50k subset) | 50’000 | |
| Map_cattle1 | Illumina BovineSNP50 BeadChip | 45’613 | |
| Map_cattle2 | Illumina BovineHD BeadChip | 727‘605 | |
| Map_cattle3 | Illumina BovineLD BeadChip | 6’600 | |
| Map_cattle4 | Genotyping chip variations | 732‘645 | |
| Map_horse1 | Illumina EquineSNP50 BeadChip | 51’105 | |
| Map_sheep1 | IlluminaOvineHDSNP | 575‘256 | |
| Map_sheep2 | IlluminaOvineSNP50 | 46‘545 | |
| Map_sheep3 | Genotyping chip variants | 580‘661 | |
| Map_goat1 | Illumina_GoatSNP50 | 55‘050 | |
| Map_human1 | Affy GeneChip500K | 483‘418 | |
| Map_human2 | Illumina_1M-duo | 1‘122‘013 | |
| Map_human3 | Illumina_HumanHap550 | 545‘902 | |
| Map_maize1 | Affymetrix Axiom Maize Genotyping Array | 501’124 | |

| | | |
|---|---|---|
| Map_wheat1 | Subset of a 55k chip from (Liu et al. 2018) | 12’109 |
| Map_wheat2 | Subset of a 90K chip from (Wen et al. 2017) | 29’692 |
| Map_sorghum1 | Subset of a 90k chip from (Bekele et al. 2013) | 3’000 |

Ex_json – the first few rows. Do not bother trying to understand it. *Json.simulation()* will do the job for you:

```
"Nodes": [
  {
    "id": "Bull",
    "Number of Individuals": "100",
    "x": -152,
    "y": -165,
    "individualsVar": "100",
    "Founder": "Yes",
    "Path": "",
    "Proportion of Male": 1,
    "BV Plot": "Yes",
    "Sex": "Male",
    "Phenotyping Class": "Default PhenoC",
    "Housing Cost Class": "Male individuals",
    "Proportion of genotyped individuals": 1,
    "label": "Bull",
    "color": "#9acef4",
    "title": "Bull: 100 Ind"
  },
```

### 14 User-interface

The development of the user-interface of MoBPS is a joint project of Torsten Pook, Amudha Ganesa, Ngoc-Thuy Ha, Lisa Büttgen, Johannes Geibel and Henner Simianer (All: Department of Animal Sciences, Center for Integrated Breeding Research, University of Goettingen, Goettingen, 37075, Germany). The user-interface is currently in active development and available at www.mobps.de. Note that this is still a development version and is constantly undergoing change.

**There will be a separate publication for the user interface that is in preparation but this will still take a while. The user-interface will take more developing to fulfil our standards for publication!** In case you have a given simulation you want to perform feel free to contact us for possible collaboration. When looking through the openly-available code you will find code snippets that are only relevant for the interface (*json.simulation()*). At the current state, this chapter can mostly be seen as a news page on the current stage of development.

Main goal of the user-interface is the usage of the R-package without the need of programming skills in R or knowledge of the details of the package to set up your simulation. Note that the interface will not be able to grasp the full functionality/efficiency of the R-package but the goal is to get close. Input parameters can be entered in a web-based application (java-script) – especially the breeding scheme can be entered in an intuitive way via nodes (cohorts of individuals) and edges (breeding & selection processes).

Simulations can be directly started via the web-interface with a VM server hosted from Goettingen. AT the current state we can provide computational resource for smaller generations (20 CPU, 64 GB Memory), but it is also population to download the json-file containing all information of the user interface and run the simulation via *json.simulation()* in R. The global test account (EAAPguest) has no permission to use our computing resources but scripts can still be exported. Student-Users are allowed to use up to five cores and are limited to 6 hour simulations, professional users are allowed up to 10 cores with no time limit for simulations. To generate an account or upgrade your available resources contact me.

To input information in regard to the breeding program in a user friendly way we provide the following module:

1. Design you Genome
2. Design your Traits
3. Multiple Subpopulations
4. Design your Selection Index
5. Reasons for Culling
6. Economic Parameters
7. Draw your Breeding Scheme
8. Analyze your Population

For most inputs we provide implemented help buttons to briefly explain what kind of input is expected. The exemplary json-script provided in the R-package would look like this (note that all advanced parameter options are deactivate to not further complicate things):

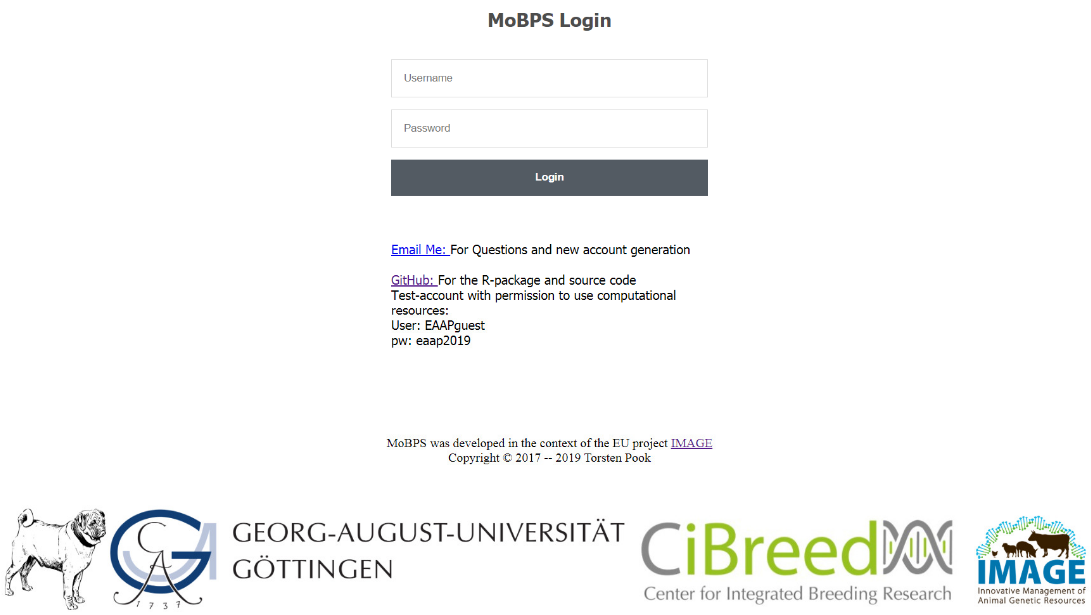

<< Ex_json >> Save Project | Save As ... | Download Project | Delete Project | Show/Hide Info | Show/Hide Warnings

### MoBPS

**Load a new/existent project from your own database:**

Project: Ex_json ▾ Version: (most recent) ▾

**Or choose a template as a starting point:**

▾

#### General Information

| | |
|---|---|
| Project Name ⓘ | Ex_json |
| Advanced settings | ☐ |
| Species ⓘ | Cattle ▾ |
| Time Unit ⓘ | Years ▾ |
| Genetic Data ⓘ | ◉ Use Ensembl Map<br>○ Upload Own Map (vcf/plink)<br>○ Create customized Map |
| Ensembl Dataset ⓘ | Illumina BovineSNP50 BeadChip ▾ |
| Max. Number of SNPs ⓘ | 10000 |

#### Phenotype Information ⓘ

Add new phenotype | Show/Hide 3 phenotypes | Show/Hide QTLs | Show/Hide phenotypic correlation | Show/Hide genetic correlation

| Phenotype ⓘ | Unit ⓘ | Pheno. Mean ⓘ | Pheno. SD ⓘ | Heritability ⓘ | # polygenic loci ⓘ | Major QTL ⓘ | Value per unit (€) ⓘ | Show Cor | |
|---|---|---|---|---|---|---|---|---|---|
| Milk | liters | 9300 | 900 | 0.35 | 1000 | 1 | 0,30 | ☑ | X |
| Fat | % | 3.9 | 0.4 | 0.4 | 1000 | 1 | 100 | ☑ | X |
| Protein | % | 3.4 | 0.3 | 0.38 | 100 | 0 | 100 | ☑ | X |

| SNP for Milk | SNP ID ⓘ | bp | chromo | Effect AA | Effect AB | Effect BB | Allele Freq. (B) ⓘ | Optional Info | |
|---|---|---|---|---|---|---|---|---|---|
| 1 | QTL DGA | 545 | 14 | 0 | -400 | -800 | 0.2 | DGAT1 | X |

| SNP for Fat | SNP ID ⓘ | bp | chromo | Effect AA | Effect AB | Effect BB | Allele Freq. (B) ⓘ | Optional Info | |
|---|---|---|---|---|---|---|---|---|---|
| 1 | QTL DGA | 545 | 14 | 0 | 0.5 | 1 | 0.2 | DGAT1 | X |

**Upload Excel file for Correlation:**

ⓘ

Datei auswählen Keine ausgewählt Save to Project

**Residual Correlation** ⓘ

| | Milk | Fat | Protein |
|---|---|---|---|
| Milk | 1 | 0.1 | 0.3 |
| Fat | 0.1 | 1 | -0.2 |
| Protein | 0.3 | -0.2 | 1 |

Enter Phenotypic correlation instead of residual correlation ☐

**Genetic Correlation** ⓘ

| | Milk | Fat | Protein |
|---|---|---|---|
| Milk | 1 | 0.3 | 0.4 |
| Fat | 0.3 | 1 | 0.1 |
| Protein | 0.4 | 0.1 | 1 |

**Creating own Selection Indexes (Optional)** ⓘ

Type SI name | Add new SI

| SI ⓘ | Milk | Fat | Protein |
|---|---|---|---|
| Default Index | 1 | 1 | 1 |
| Milk increase | 5 | 1 | 1 |

| SI ⓘ | Selection Index (Hazel and Lush 1943, Miesenberger 1997) ⓘ | Standardization ⓘ |
|---|---|---|
| Default Index | ☐ | Per Phenotypic SD |
| Milk increase | ☐ | Per Phenotypic SD |

**Creating own Phenotyping Classes (Optional)** ⓘ

Type PhenoC name | Add new PhenoC

| PhenoClass ⓘ | Phenotyping Cost | Milk | Fat | Protein | |
|---|---|---|---|---|---|
| Default PhenoC | 800 | 1 | 1 | 1 | |
| Male individuals | 0 | 0 | 0 | 0 | X |

**Breeding Scheme** ⓘ

***Important: Please AVOID node names with underscore and trailing numbers combined, e.g. ABC_1, DEF_2, ...***

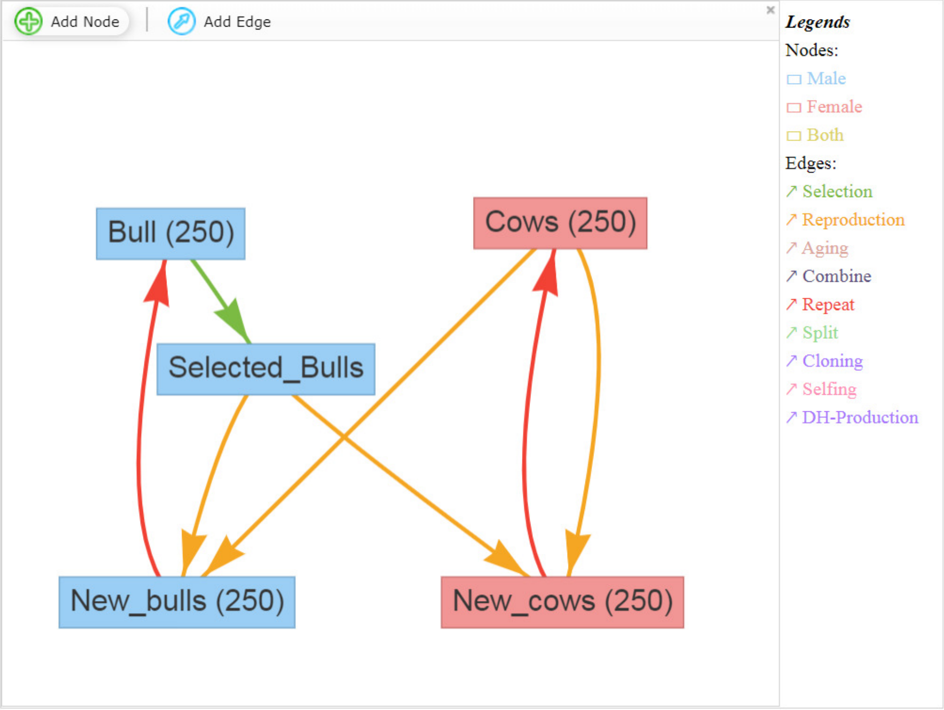

After the simulations are execute the resulting, population-list can be download in R and then be manually analyzed. Alternatively, we also provide some basic evaluation functions. In case multiple simulations are performed it is also possible to analyze average between multiple runs of the simulation:

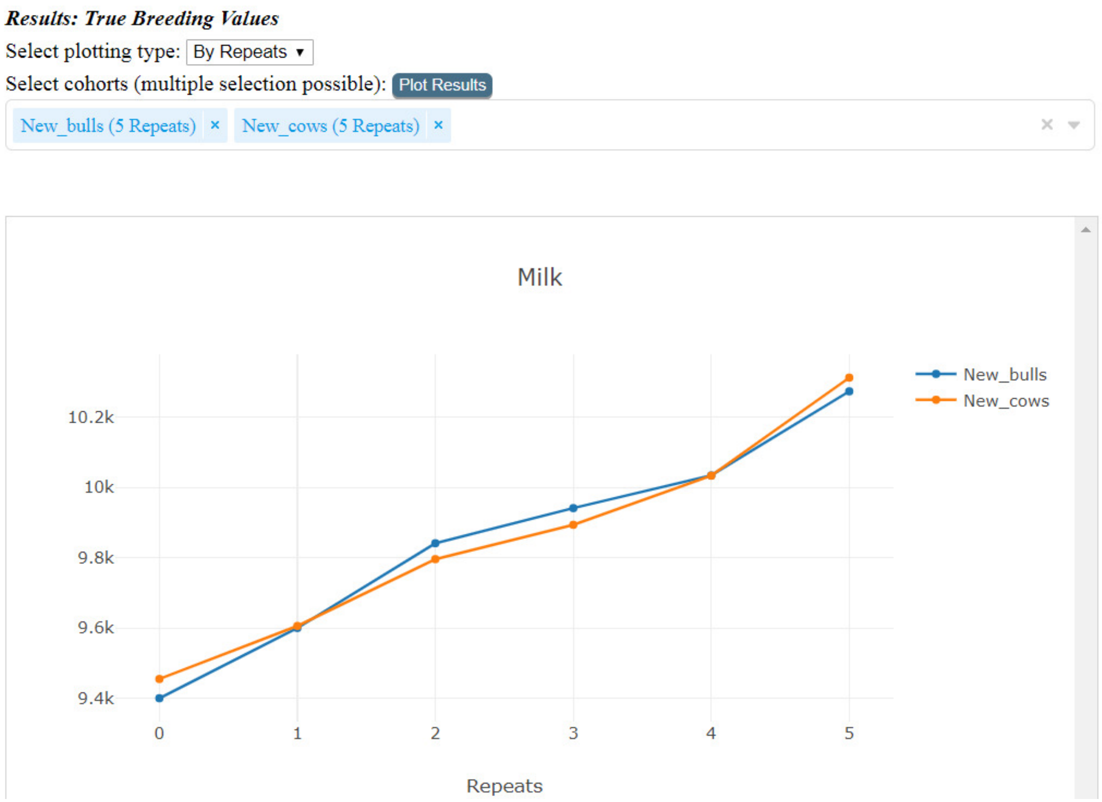

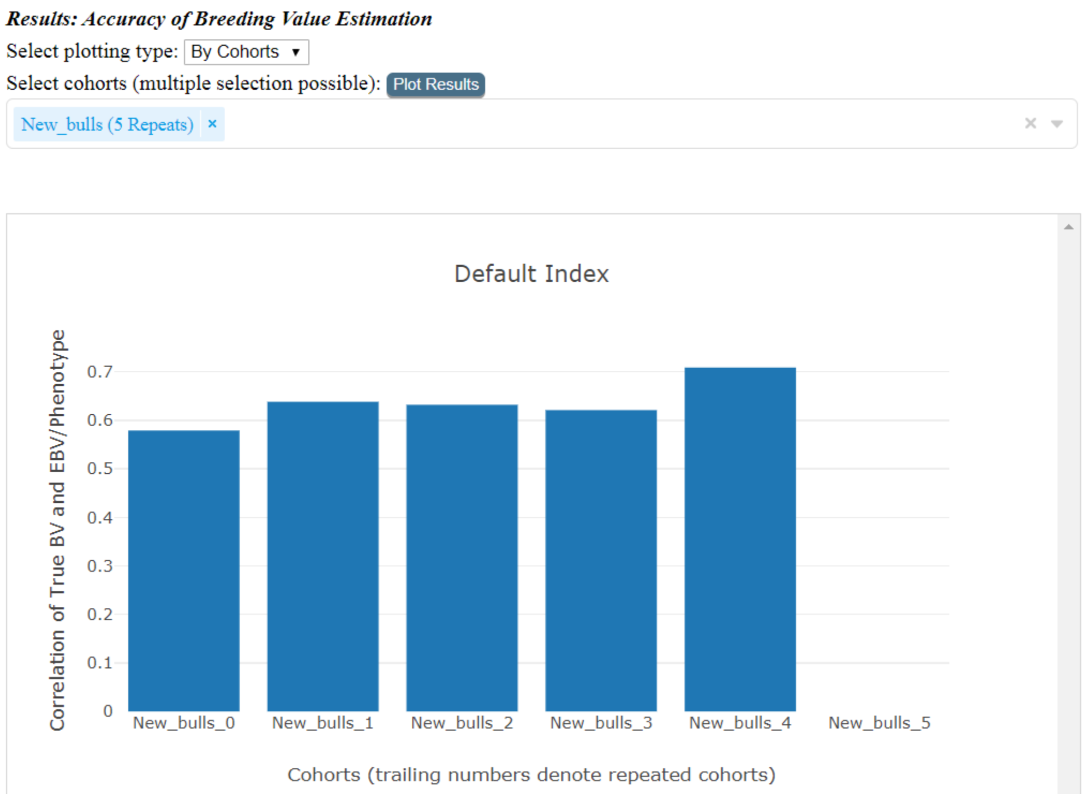

***Results: Relationship and Inbreeding within Cohorts***

Select plotting type: By Repeats ▾

Select cohorts (multiple selection possible): Plot Results

New_bulls (5 Repeats) × New_cows (5 Repeats) ×

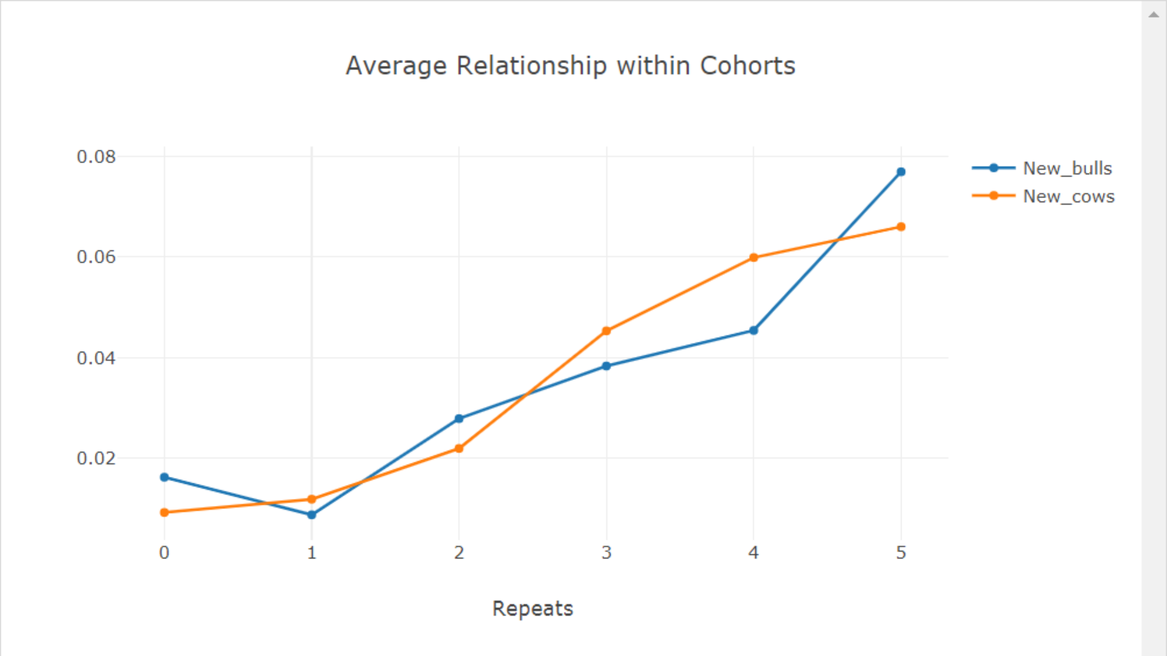

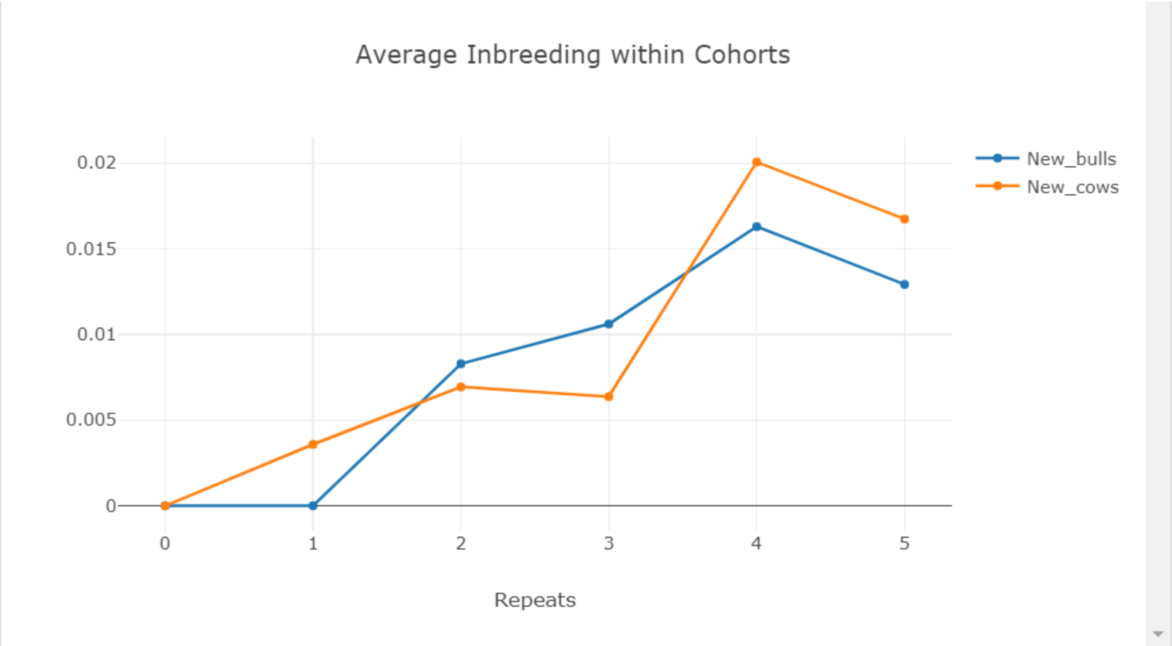

***Results: Relationship between Cohorts***

Select cohort 1 : New_bulls (5 Repeats) ▾ Select cohort 2 : New_cows (5 Repeats) ▾ Plot Results

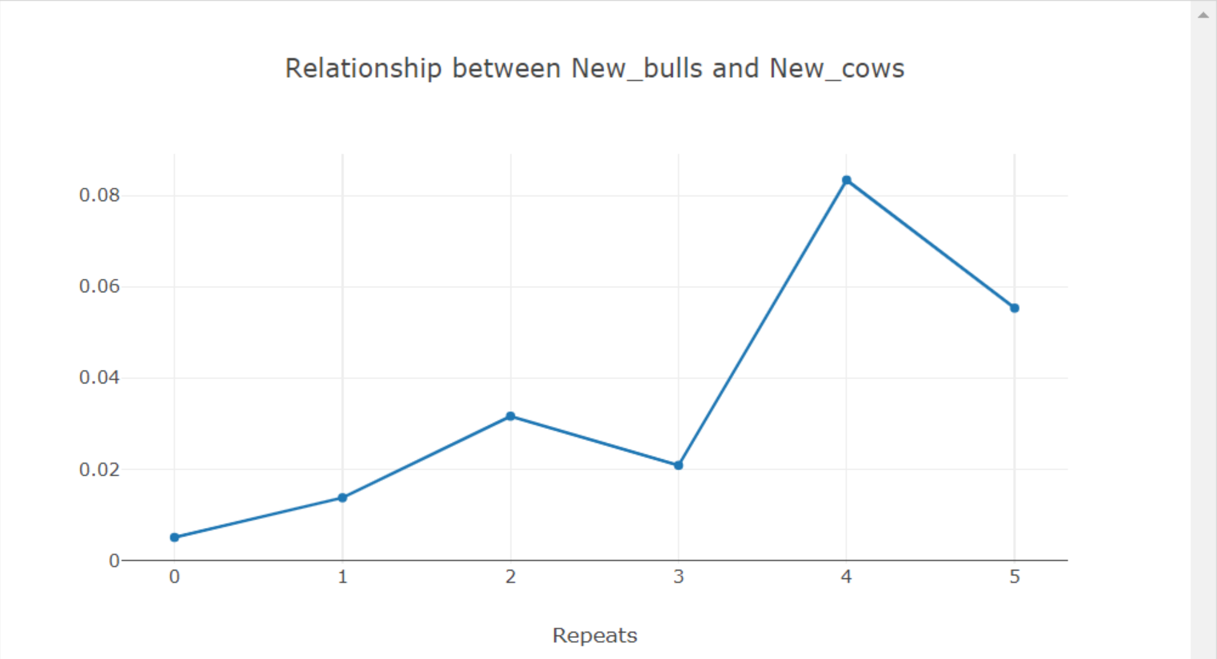

***Results: Major QTLs (Allele Frequency, exp./obs. Heterozygosity)***

Select Trait : Milk ▾ Select QTL : QTL DGAT1 ▾

Select Cohorts (multiple selection possible): Plot Results

New_bulls (5 Repeats) × New_cows (5 Repeats) ×

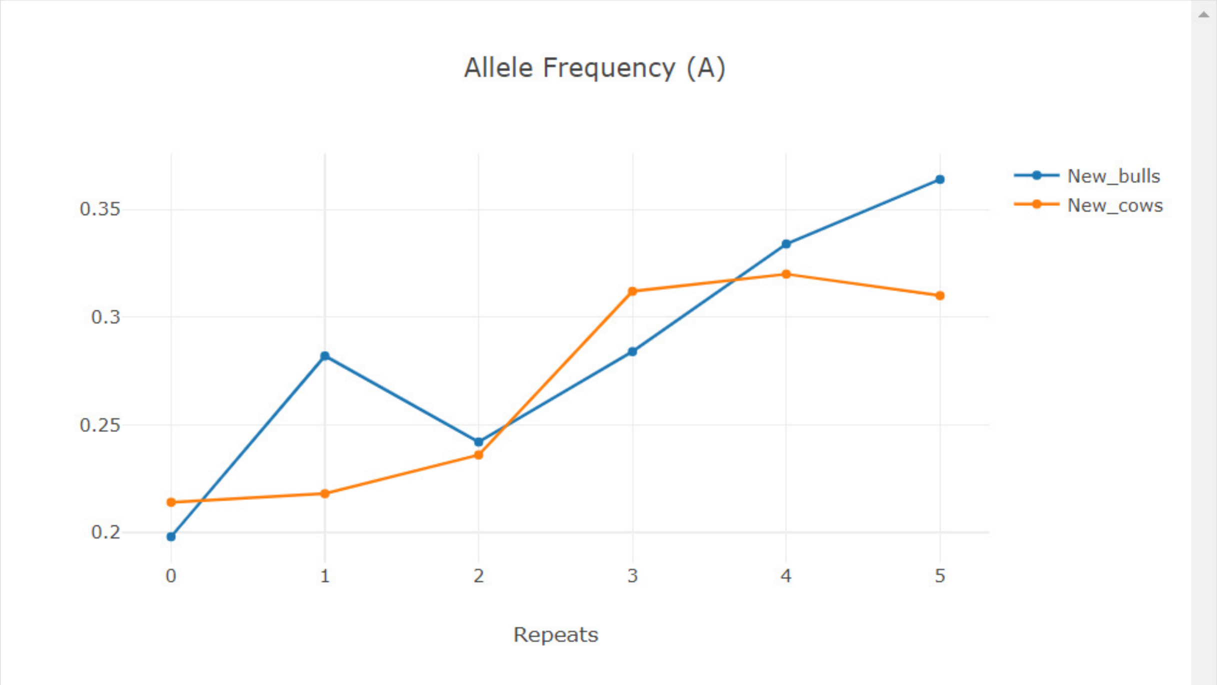

### 15 Commonly used word definitions

Group: Group of individuals with the same sex and belonging to the same generation

Cohorts: Group of individuals with the same sex generated in a single run of *breeding.diploid()*

Class: Auxiliary variable to classify individuals in an additional dimension (besides sex & generation)

Founder: Founder individuals are the start-point of a simulation and all individuals in the population can be traced back to the founders. Because of this only for those individuals genotype/haplotype data has to be saved.

### 16 Exemplary scripts

#### 16.1 Simulation of a MAGIC population in maize

We here show how to perform an exemplary simulation of a MAGIC population in maize with a mating scheme given in (Zheng et al. 2015) – cf. adjacent Figure.

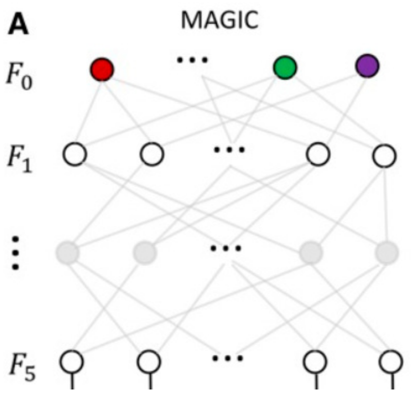

Since default settings in MoBPS are to always use the last generation anyway the needed code is quite short even without cohort mode. For the sake of completeness, we provide a cohort version for the script as well.

In term of computation time this simulation with a 15.3M genome, 31k SNPs and a total of 780 individuals took 1.6 seconds on one core of my local maschine.

<u>Non-cohort-modus:</u>

```
# Generation of 20 fully-homozygous founders lines
# All plants are stored as male individuals (sex=0)
population <- creating.diploid(nindi = 20, sex.quota = 0, template.chip = "maize",
                               dataset = "homorandom")

# Simulate matings between all founders.
# Each plan is involved in exactly 19 matings.
population <- breeding.diploid(population, breeding.size = c(190,0),
                               breeding.all.combination = TRUE,
                               selection.size = c(20,0), max.offspring = 19)

# Simulate matings between plants of the last generation.
# Each plant is involved in exactly 2 matings.

population <- breeding.diploid(population, breeding.size = c(190,0),
                               selection.size = c(190,0), same.sex.activ = TRUE,
                               same.sex.sex = 0, max.offspring = 2)
population <- breeding.diploid(population, breeding.size = c(190,0),
                               selection.size = c(190,0), same.sex.activ = TRUE,
                               same.sex.sex = 0, max.offspring = 2)
population <- breeding.diploid(population, breeding.size = c(190,0),
                               selection.size = c(190,0), same.sex.activ = TRUE,
                               same.sex.sex = 0, max.offspring = 2)
```

<u>Cohort-modus:</u>

```
# Generation of 20 fully-homozygous founders lines
# All plants are stored as male individuals (sex=0)
population <- creating.diploid(nindi = 20, sex.quota = 0, template.chip = "maize",
                               dataset = "homorandom", name.cohort = "F0")
```

```
# Simulate matings between all founders.
# Each plan is involved in exactly 19 matings.
population <- breeding.diploid(population, breeding.size = c(190,0),
                               breeding.all.combination = TRUE,
                               selection.size = c(20,0),
                               selection.m.cohort = "F0", name.cohort = "F1")

# Simulate matings between plants of the last generation.
# Each plant is involved in exactly 2 matings.

population <- breeding.diploid(population, breeding.size = c(190,0),
                               selection.size = c(190,0), same.sex.activ = TRUE,
                               same.sex.sex = 0, max.offspring = c(2,0),
                               selection.m.cohort = "F1", name.cohort = "F2")
population <- breeding.diploid(population, breeding.size = c(190,0),
                               selection.size = c(190,0), same.sex.activ = TRUE,
                               same.sex.sex = 0, max.offspring = c(2,0),
                               selection.m.cohort = "F2", name.cohort = "F3")
population <- breeding.diploid(population, breeding.size = c(190,0),
                               selection.size = c(190,0), same.sex.activ = TRUE,
                               same.sex.sex = 0, max.offspring = c(2,0),
                               selection.m.cohort = "F3", name.cohort = "F4")
```

#### 16.2 Simulation of Introgression on blue eggshell QTL

We here show how to perform an exemplary simulation of a breeding scheme to perform introgression of a single QTL. In term of computing time this simulation with a 5M genome, 5k SNPs and a total of 520 individuals took 1.2 seconds on one core of my local maschine without the usage of miraculix.

```
# Generate an input SNP-dataset

# 10 White-Layer (0) (20 haplotypes, 5'000 SNPs)
# 10 Wild population (1) (20 haplotypes, 5'000 SNPs)
dataset1 <- matrix(0, nrow = 5000, ncol = 20)
dataset2 <- matrix(1, nrow = 5000, ncol = 20)

# Generation of a trait
# Colums code: SNP, chromosome, effect 00, effect 01, effect 11
# Blue Eggshell QTL is positioned on SNP 2000, chromosome 1
major_qtl <- c(2000, 1, 0, 10000, 20000)
# In all other positions the white layer genome is assumed to be favorable
# All marker effects combiened are smaller than the blue eggshell QTL
rest <- cbind(1:5000, 1, 1, 0.5, 0)
trait <- rbind(major_qtl, rest)

# Generation of the base-population
# First 10 individuals are female (sex=2)
# Next 10 individuals are male (sex=1)
population <- creating.diploid(dataset = cbind(dataset1, dataset2),
                               real.bv.add = trait, name.cohort = "Founders",
                               sex.s = c(rep(2,10), rep(1,10)))

# Simulate random mating:
population <- breeding.diploid(population, breeding.size = c(100,100),
                               selection.size = c(10,10),
                               selection.m.cohorts = "Founders_M",
                               selection.f.cohorts = "Founders_F",
                               name.cohort = "F1")

# Simuation of matings with selection:
# Top 50 cocks are mated to the 10 founder hens
# Selection of the cocks based on their genomic value ("bv")
# Target: Increase share of white layer while preserving blue egg shell QTL
```

```
population <- breeding.diploid(population, breeding.size = c(100,100),
                               selection.size = c(50,10),
                               selection.m.cohorts = "F1_M",
                               selection.f.cohorts = "Founders_F",
                               name.cohort = "BC1", selection.m = "function",
                               selection.criteria.type = "bv")
population <- breeding.diploid(population, breeding.size = c(100,100),
                               selection.size = c(50,10),
                               selection.m.cohorts = "BC1_M",
                               selection.f.cohorts = "Founders_F",
                               name.cohort = "BC2", selection.m = "function",
                               selection.criteria.type = "bv")
population <- breeding.diploid(population, breeding.size = c(100,100),
                               selection.size = c(50,10),
                               selection.m.cohorts = "BC2_M",
                               selection.f.cohorts = "Founders_F",
                               name.cohort = "BC3", selection.m = "function",
                               selection.criteria.type = "bv")

# Mating of cocks and hens that are heterozygous in blue egg shell QTL
# 25% of resulting offspring should be homozygous in blue egg shell QTL

population <- breeding.diploid(population, breeding.size = c(100,100),
                               selection.size = c(50,50),
                               selection.m.cohorts = "BC3_M",
                               selection.f.cohorts = "BC3_F",
                               name.cohort = "IC", selection.m = "function",
                               selection.criteria.type = "bv")

# Check genomic share of wild race in the final generation
genoIC <- get.geno(population, cohorts = "IC_F")
plot(rowSums(genoIC)/200, xlab = "genome", ylab = "frequency of wild allele", type
= "l")
abline(v = 2000, lwd = 2, col = "red")
```

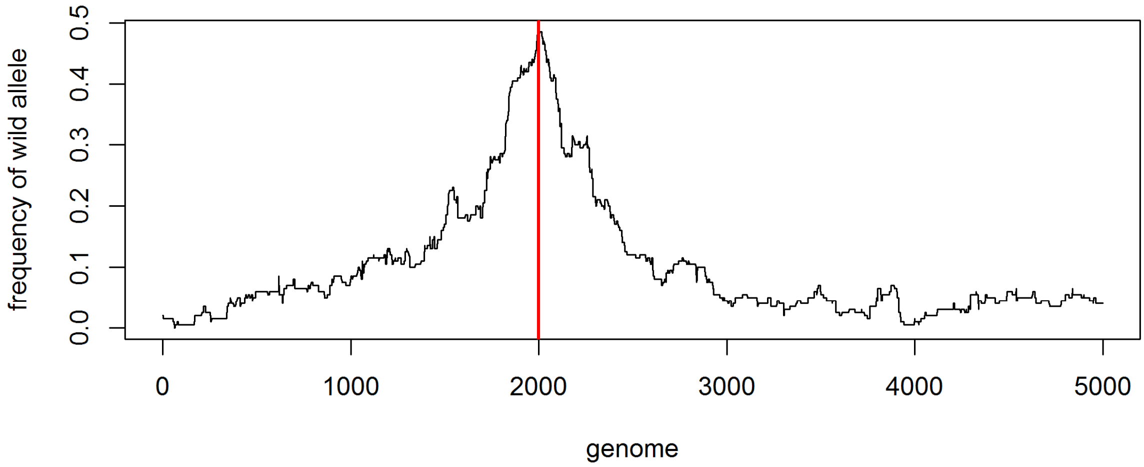

As expected, the frequency of genetic material stemming from the wild type is higher in the region of the QTL.

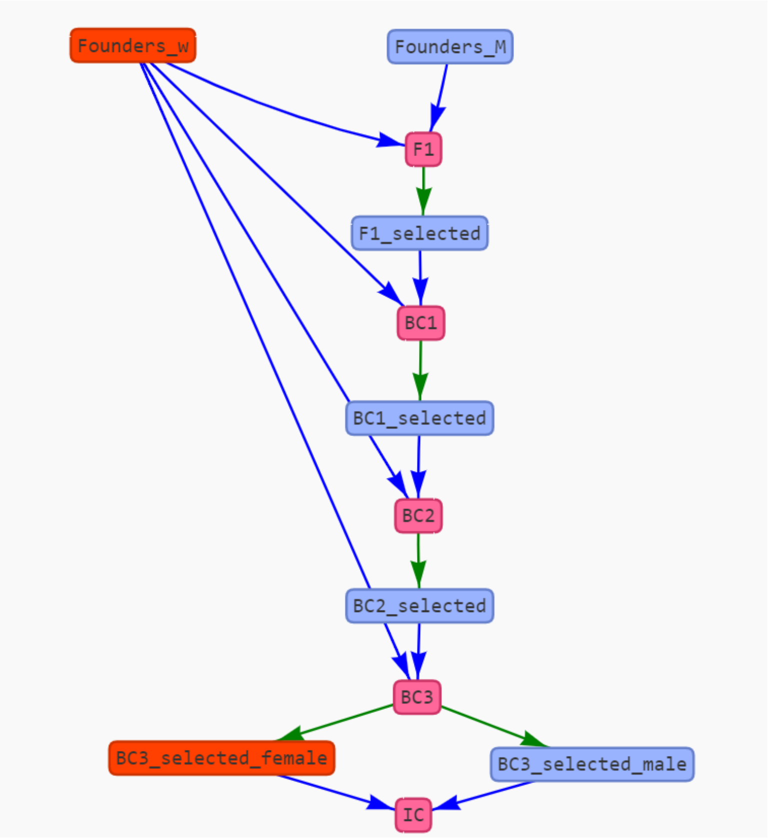

*Figure 2: Mating Scheme for Introgression of the blue-egg-shell QTL. Graph is generated via user-interface in MoBPS*

#### 16.3 Simulation of gene editing in a cow breeding program

The following script can be used to simulate a breeding program that is utilizing genome editing. Design is chosen according to (Jenko et al. 2015; Simianer et al. 2018). Note that individual numbers are much smaller than in the two references to ensure low computation times. Simulation of 20 generations with 50’000 cows per generation would take 26.3 hours using 24 cores on the gwdg-hpc (Intel E5-2650 (2X12 core 2.2GHz)). This small example with a 5 Morgan chromosome, 5k SNPs and 4300 individuals took 7.4 seconds.

```
# Generation of a base population:
# 1'000 Founder individuals
# 5'000 SNPs
# 100 additive single marker QTL
population <- creating.diploid(nindi = 1000, nsnp = 5000,
                               n.additive = 100, name.cohort = "Founders")

# Simulation of a random mating generation
# 100 bulls (sex=1), 1'000 cows (sex=2) are generated
population <- breeding.diploid(population, breeding.size = c(100,1000),
                               selection.size = c(500,500),
                               selection.m.cohorts = "Founders_M",
                               selection.f.cohorts = "Founders_F",
                               name.cohort = "Random")

# Generate 200 offspring of both from the top 5 bulls / 200 cows
# Heritability of the trait is set to 0.5
# only phenotypes previously unobserved cows are generated
population <- breeding.diploid(population, breeding.size = 200,
                               selection.size = c(5,200), bve = TRUE,
                               heritability = 0.5,
                               new.bv.observation = "non_obs_f",
                               selection.m = "function", name.cohort = "Top",
                               selection.m.cohorts = "Random_M",
                               selection.f.cohorts = "Random_F")

# Generate additional cows using all cows of the previous generation
# Cows are added to the same generation as the previous simulation
population <- breeding.diploid(population, breeding.size = c(0,900),
                               selection.size = c(5,1000),
                               selection.m = "function", name.cohort = "Sec_F",
                               selection.m.cohorts = "Random_M",
                               selection.f.cohorts = "Random_F",
                               add.gen = 3)

# Same cycle as before with additional genome editing
# Edits are chosen based on highest effects in rrBLUP
population <- breeding.diploid(population, breeding.size = c(100,100),
                               selection.size = c(5,200), bve = TRUE,
                               new.bv.observation = "non_obs_f",
                               selection.m = "function",
                               name.cohort = "Top_Edit",
                               selection.m.cohorts = "Top_M",
                               selection.f.cohorts = c("Top_F","Sec_F"),
                               nr.edits = 20, estimate.u = TRUE)

population <- breeding.diploid(population, breeding.size = c(0,900),
                               selection.size = c(5,1000),
                               selection.m = "function", name.cohort = "Sec_Edit",
                               selection.m.cohorts = "Top_M",
                               selection.f.cohorts = c("Top_F","Sec_F"),
                               add.gen = 4)

bv.development(population, cohorts = c("Founders_F", "Random_F", "Sec_F",
                                       "Top_F", "Sec_Edit", "Top_Edit_F"),
```

```
                    display.cohort.name = TRUE, display.sex = TRUE, development = 1)
```

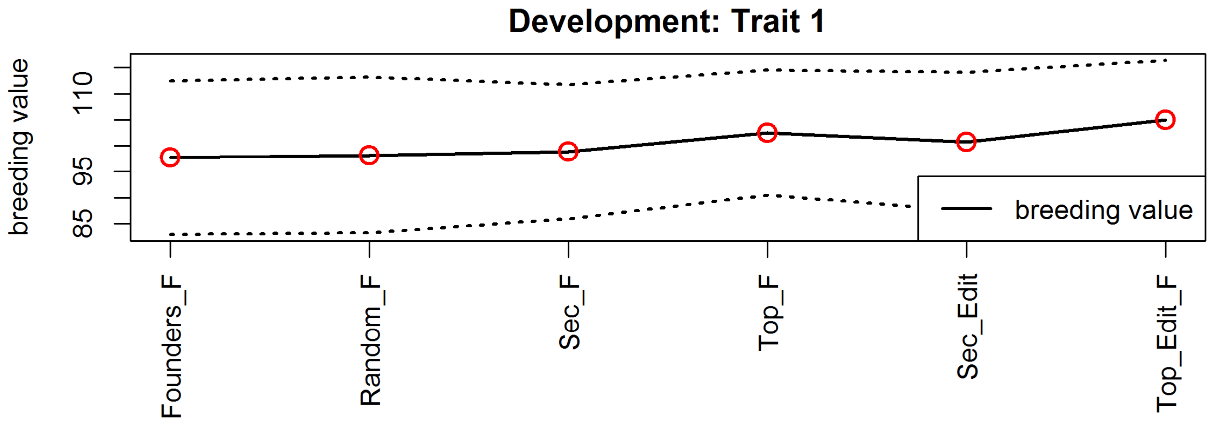

#### 16.4 Simulation of a base population with a hard sweep

We here show how to perform an exemplary simulation to generate a base population and a hard sweep.

```
# Generate a starting population with 5000 SNPs and 200 individuals
# and a single chromosome of length 2 Morgan.
population <- creating.diploid(nsnp = 5000, nindi = 200, chromosome.length = 2)

# LD build up via 100 generations of random mating
# Each generation contains 200 individuals
for(index in 1:100){
  population <- breeding.diploid(population, breeding.size = 200,
                                 selection.size = c(100,100))
}

# Derive allele frequency and check LD for the last generation:
genotype.check <- get.geno(population, gen = length(population$breeding))
p_i <- rowMeans(genotype.check)/2
ld.decay(population, genotype.dataset = genotype.check, step = 10, max = 500)
```

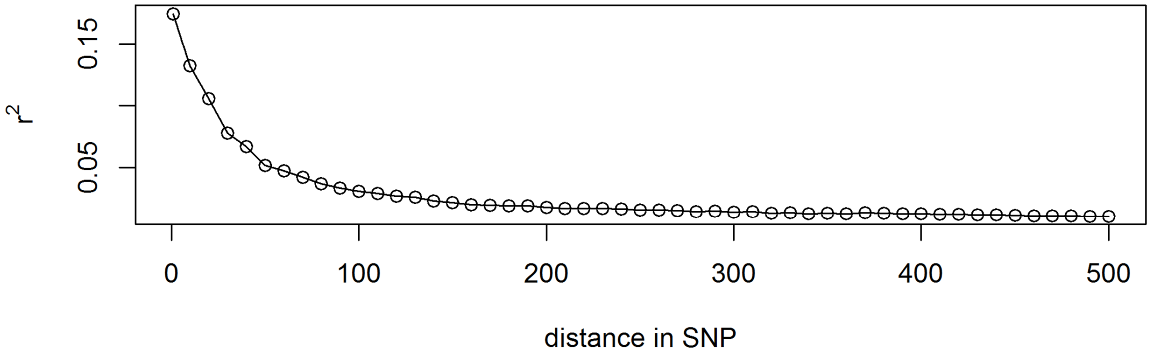

```
# Simulate a favorable mutation in a previously fixed marker
fixated_markers <- which(p_i==0) # Which markers are fixated
qtl_posi <- sample(fixated_markers, 1) # Selected a fixated marker in A
trait <- cbind(qtl_posi, 1, 0, 1, 2) # SNP, Chromosome, Effect AA, Effect AB,
Effect BB
population <- creating.trait(population, real.bv.add = trait)
```

```
# Generate a mutation in the first male individual
population <- mutation.intro(population, 101, 1, 1, qtl_posi)

# Simulate generations with selection pressure
# Individuals with the favorable SNP are picked 5 times as often
for(index in 1:25){
  population <- breeding.diploid(population, breeding.size = 200,
                                 selection.size = c(100,100),
                                 best.selection.ratio.m = 5,
                                 best.selection.ratio.f = 5)
}

analyze.population(population, gen = 98:115, chromosome = 1, snp = qtl_posi)
```

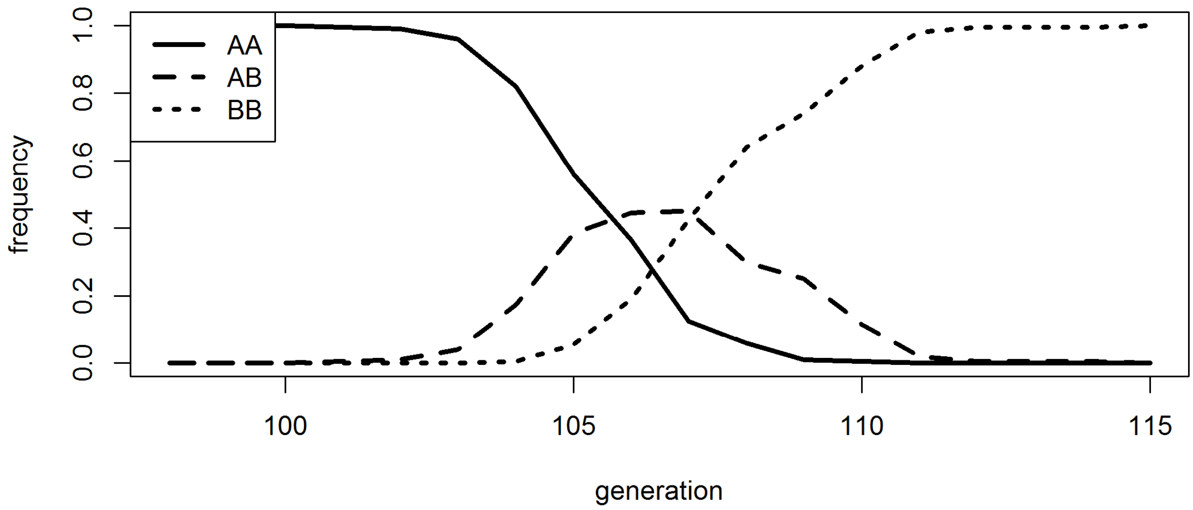

### 17 Acknowledgements

This package was developed in the context of the European Union's Horizon 2020 Research and Innovation Program under grant agreement n°677353 IMAGE

Additional thanks goes to the Research Training Group 1644 "Scaling Problems in Statistics" for financing travelling, BMBF project "MAZE – Accessing the genomic and functional diversity of maize to improve quantitative traits" (Grant ID 031B0195) and all members of the Animal Breeding and Genetics Group at the University of Goettingen for all the helpful advice to people with less genetic background and ideas of things to implement.

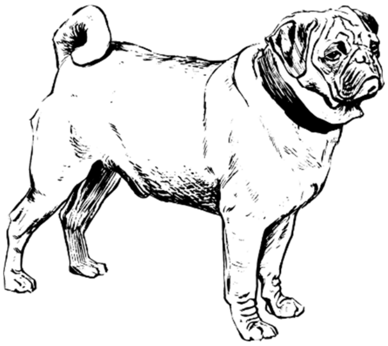

IMAGE
Innovative Management of Animal Genetic Resources

CiBreed
Center for Integrated Breeding Research

GEORG-AUGUST-UNIVERSITÄT GÖTTINGEN
