## Supplementary material for "MoBPS - Modular Breeding Program Simulator": MoBPS R-package: add.diag.R

```
'#
  Authors
Torsten Pook,

Copyright (C) 2017 -- 2018  Torsten Pook

This program is free software; you can redistribute it and/or
modify it under the terms of the GNU General Public License
as published by the Free Software Foundation; either version 3
of the License, or (at your option) any later version.

This program is distributed in the hope that it will be useful,
but WITHOUT ANY WARRANTY; without even the implied warranty of
MERCHANTABILITY or FITNESS FOR A PARTICULAR PURPOSE.  See the
GNU General Public License for more details.

You should have received a copy of the GNU General Public License
along with this program; if not, write to the Free Software
Foundation, Inc., 59 Temple Place - Suite 330, Boston, MA  02111-1307, USA.
'#

#' Add something to the diagonal
#'
#' Function to add numeric to the diagonal of a matrix
#' @param M Matrix Population list
#' @param d Vector to add to the diagonal of the matrix
#' @export

add.diag <- function(M, d){
  diag(M) <- diag(M) + d
  return(M)
}
```
