## Supplementary material for "MoBPS - Modular Breeding Program Simulator": MoBPS R-package: analyze.bv.R

```
'#
  Authors
Torsten Pook,

Copyright (C) 2017 -- 2018  Torsten Pook

This program is free software; you can redistribute it and/or
modify it under the terms of the GNU General Public License
as published by the Free Software Foundation; either version 3
of the License, or (at your option) any later version.

This program is distributed in the hope that it will be useful,
but WITHOUT ANY WARRANTY; without even the implied warranty of
MERCHANTABILITY or FITNESS FOR A PARTICULAR PURPOSE.  See the
GNU General Public License for more details.

You should have received a copy of the GNU General Public License
along with this program; if not, write to the Free Software
Foundation, Inc., 59 Temple Place - Suite 330, Boston, MA  02111-1307, USA.
'#

#' Analyze genomic values
#'
#' Function to analyze correlation between bv/bve/pheno
#' @param population Population list
#' @param database Groups of individuals to consider for the export
#' @param gen Quick-insert for database (vector of all generations to export)
#' @param cohorts Quick-insert for database (vector of names of cohorts to export)
#' @param bvrow Which traits to display (for multiple traits separte plots (par(mfrow)))
#' @export
#'

analyze.bv <- function(population, gen=NULL, database=NULL, cohorts=NULL, bvrow="all"){

  if(length(bvrow)==1 && bvrow=="all"){
    bvrow <- 1:population$info$bv.nr
  }
  bv <- get.bv(population, gen=gen, database = database, cohorts = cohorts)[bvrow,,drop=FALSE]
  bve <- get.bve(population, gen=gen, database = database, cohorts = cohorts)[bvrow,,drop=FALSE]
  pheno <-get.pheno(population, gen=gen, database = database, cohorts = cohorts)[bvrow,,drop=FALSE]


  cor_matrix <- matrix(0, nrow=3, ncol=length(bvrow))
  var_vector <- numeric(length(bvrow))
  for(index in 1:length(bvrow)){
    cor_matrix[1,index] <- stats::cor(bv[index,], bve[index,])
    cor_matrix[2,index] <- stats::cor(bv[index,], pheno[index,])
    cor_matrix[3,index] <- stats::cor(bve[index,], pheno[index,])
    var_vector[index] <- stats::var(bv[index,])
  }
  names(var_vector) <- population$info$trait.name
  colnames(cor_matrix) <- population$info$trait.name
  rownames(cor_matrix) <- c("BV / BVE", "BV / Pheno", "BVE / Pheno")
  return(list(cor_matrix, var_vector))

}
```
