## Supplementary material for "MoBPS - Modular Breeding Program Simulator": MoBPS R-package: analyze.population.R

```
'#
  Authors
Torsten Pook,

Copyright (C) 2017 -- 2018  Torsten Pook

This program is free software; you can redistribute it and/or
modify it under the terms of the GNU General Public License
as published by the Free Software Foundation; either version 3
of the License, or (at your option) any later version.

This program is distributed in the hope that it will be useful,
but WITHOUT ANY WARRANTY; without even the implied warranty of
MERCHANTABILITY or FITNESS FOR A PARTICULAR PURPOSE.  See the
GNU General Public License for more details.

You should have received a copy of the GNU General Public License
along with this program; if not, write to the Free Software
Foundation, Inc., 59 Temple Place - Suite 330, Boston, MA  02111-1307, USA.
'#

#' Analyze allele frequency of a single marker
#'
#' Analyze allele frequency of a single marker
#' @param population Population list
#' @param database Groups of individuals to consider for the export
#' @param gen Quick-insert for database (vector of all generations to export)
#' @param cohorts Quick-insert for database (vector of names of cohorts to export)
#' @param chromosome Number of the chromosome of the relevant SNP
#' @param snp Number of the relevant SNP
#' @export

analyze.population <- function(population, chromosome, snp, database=NULL, gen=NULL, cohorts=NULL){

  groups <- sum(nrow(database), length(gen), length(cohorts))
  state <- matrix(0, nrow=3, ncol = groups)
  p.snp <- sum(population$info$length[0:(chromosome-1)]) + population$info$position[[chromosome]][snp]
  n.snp <- sum(population$info$snp[0:(chromosome-1)]) + snp

  col <- 1
  if(length(gen)>0){
    for(index in 1:length(gen)){
      genos <- get.geno(population, gen = gen[index])[n.snp,]
      state[,col] <- c(sum(genos==0), sum(genos==1), sum(genos==2))
      col <- col + 1
    }
  }
  if(length(database)>0){
    for(index in 1:nrow(database)){
      genos <- get.geno(population, database = database[index,])[n.snp,]
      state[,col] <- c(sum(genos==0), sum(genos==1), sum(genos==2))
      col <- col + 1
    }
  }
  if(length(cohorts)>0){
    for(index in 1:length(cohorts)){
      genos <- get.geno(population, cohorts = cohorts[index])[n.snp,]
      state[,col] <- c(sum(genos==0), sum(genos==1), sum(genos==2))
      col <- col + 1
    }
  }
  datatime <- c(gen, database[,1], get.database(population, cohorts=cohorts)[,1])

  state.prob <- t(t(state)/colSums(state))
  maxp <- max(state.prob)
  graphics::plot(datatime ,state.prob[1,],xlim=c(min(datatime),max(datatime)),ylim=c(0,maxp),type="l",xlab="generation", ylab="frequency", lwd=3,
                 main="")
  graphics::lines(datatime ,state.prob[2,],lty=2, lwd=3)
  graphics::lines(datatime ,state.prob[3,],lty=3, lwd=3)
  graphics::legend("topleft",legend = c("AA","AB","BB"),lty=c(1,2,3), lwd=c(3,3,3))
  return(state)
}


#' Old Analyze allele frequency of a single marker
#'
#' Old Analyze allele frequency of a single marker
#' @param population Population list
#' @param chromosome Number of the chromosome of the relevant SNP
#' @param snp Number of the relevant SNP
#' @param include include male/female (1/2) individuals in computation default( c(1,2))
#' @export

analyze.population.old <- function(population,  chromosome, snp, include=c(1,2)){

  if (population$info$miraculix){
    if(requireNamespace("miraculix", quietly = TRUE)) {
      codeOriginsU <- miraculix::codeOrigins
      decodeOriginsU <- miraculix::decodeOrigins
    } else{
      cat("Package miraculix required. Original simulations used the package.")
    }
  } else{
    codeOriginsU <- codeOriginsR
    decodeOriginsU <- decodeOriginsR
  }
  generations <- length(population$breeding)
  state <- matrix(0, nrow=3, ncol = generations)
  p.snp <- sum(population$info$length[0:(chromosome-1)]) + population$info$position[[chromosome]][snp]
  n.snp <- sum(population$info$snp[0:(chromosome-1)]) + snp
  for(index in 1:generations){
    #    active.animals <- NULL
    for(sex in include){
      n.animals <- length(population$breeding[[index]][[sex]])
      if(n.animals>0){
        for(animal in 1:n.animals){

          if(population$info$miraculix){
            geno <- miraculix::computeSNPS(population, index, sex, animal)[n.snp]
          } else{
            geno <- sum(compute.snps(population, index, sex, animal)[,n.snp])
          }

          state[(geno+1), index] <- state[(geno+1), index] +1

        }
      }

    }
  }
  state.prob <- t(t(state)/colSums(state))
  maxp <- max(state.prob)
  graphics::plot(1:generations ,state.prob[1,],xlim=c(1,generations),ylim=c(0,maxp),type="l",xlab="generation", ylab="share", lwd=3,
                 main=paste("Allele frequencies in Chr", chromosome, "SNP", snp))
  graphics::lines(1:generations ,state.prob[2,],lty=2, lwd=3)
  graphics::lines(1:generations ,state.prob[3,],lty=3, lwd=3)
  graphics::legend("topleft",legend = c("hom0","hetero","hom1"),lty=c(1,2,3), lwd=c(3,3,3))
  return(state)
}
```
