## Supplementary material for "MoBPS - Modular Breeding Program Simulator": MoBPS R-package: bitwise.storing.R

```
'#
  Authors
Torsten Pook,

Copyright (C) 2017 -- 2018  Torsten Pook

This program is free software; you can redistribute it and/or
modify it under the terms of the GNU General Public License
as published by the Free Software Foundation; either version 3
of the License, or (at your option) any later version.

This program is distributed in the hope that it will be useful,
but WITHOUT ANY WARRANTY; without even the implied warranty of
MERCHANTABILITY or FITNESS FOR A PARTICULAR PURPOSE.  See the
GNU General Public License for more details.

You should have received a copy of the GNU General Public License
along with this program; if not, write to the Free Software
Foundation, Inc., 59 Temple Place - Suite 330, Boston, MA  02111-1307, USA.
'#

#' Bitwise-storing in R
#'
#' Function for bitwise-storing in R (only 30 of 32 bits are used!)
#' @param snpseq SNP sequence
#' @param nbits Number of usable bits (default: 30)
#' @export


bit.storing <- function(snpseq, nbits){
  n <- length(snpseq)
  q <- ceiling(n/nbits)
  if(n!=q*nbits){
    snpseq <- c(snpseq, rep(0, q*nbits-n))
  }
  mult <- c(1,rep(2,nbits-1))
  bit.seq <- as.integer(cumprod(mult) %*% matrix(snpseq, nrow=nbits))
}

#' Decoding of bitwise-storing in R
#'
#' Function for decoding in bitwise-storing in R (only 30 of 32 bits are used!)
#' @param bit.seq bitweise gespeicherte SNP-Sequenz
#' @param nbits Number of usable bits (default: 30)
#' @param population Population list
#' @param from.p.bit Bit to start on
#' @export

bit.snps <- function(bit.seq, nbits, population=NULL, from.p.bit=1){
  raw <- intToBits(bit.seq)
  keep <- rep(1:nbits, length(bit.seq)) + rep(32*(0:(length(bit.seq)-1)+ from.p.bit - 1), each=nbits)
  if(length(population) >0 && keep[length(keep)] > floor(sum(population$info$snp)/nbits)*32+population$info$leftover){
    keep <- keep[keep<=floor(sum(population$info$snp)/nbits)*32+population$info$leftover]
  }
  snp_seq <- as.integer(raw[keep])
}
```
