## Supplementary material for "MoBPS - Modular Breeding Program Simulator": MoBPS R-package: breeding.diploid.R

```
'#
  Authors
Torsten Pook,

Copyright (C) 2017 -- 2018  Torsten Pook

This program is free software; you can redistribute it and/or
modify it under the terms of the GNU General Public License
as published by the Free Software Foundation; either version 3
of the License, or (at your option) any later version.

This program is distributed in the hope that it will be useful,
but WITHOUT ANY WARRANTY; without even the implied warranty of
MERCHANTABILITY or FITNESS FOR A PARTICULAR PURPOSE.  See the
GNU General Public License for more details.

You should have received a copy of the GNU General Public License
along with this program; if not, write to the Free Software
Foundation, Inc., 59 Temple Place - Suite 330, Boston, MA  02111-1307, USA.
'#


#' Breeding function
#'
#' Function to simulate a step in a breeding scheme
#' @param population Population list
#' @param mutation.rate Mutation rate in each marker (default: 10^-5)
#' @param remutation.rate Remutation rate in each marker (default: 10^-5)
#' @param recombination.rate Average number of recombination per 1 length unit (default: 1M)
#' @param selection.m Selection criteria for male individuals (default: "random", alt: "function")
#' @param selection.f Selection criteria for female individuals (default: selection.m , alt: "random", function")
#' @param selection.function.matrix Manuel generation of a temporary selection function (Use BVs instead!)
#' @param selection.size Number of selected individuals for breeding (default: c(0,0) - alt: positive numbers)
#' @param breeding.size Number of individuals to generate
#' @param breeding.sex Share of female animals (if single value is used for breeding size; default: 0.5)
#' @param breeding.sex.random If TRUE randomly chose sex of new individuals (default: FALSE - use expected values)
#' @param relative.selection Use best.selection.ratio instead!
#' @param recom.f.indicator Use step function for recombination map (transform snp.positions if possible instead)
#' @param add.gen New animals are generated in the next generation (default: length(population$breeding))
#' @param recom.f.polynom Polynomical function to determine expected number of recombinations (transform snp.positions if possible instead)
#' @param duplication.rate Share of recombination points with a duplication (default: 0 - DEACTIVATED)
#' @param duplication.length Average length of a duplication (Exponentially distributed)
#' @param duplication.recombination Average number of recombinations per 1 length uit of duplication (default: 1)
#' @param new.class Migration level of newly generated individuals (default: 0)
#' @param bve If TRUE perform a breeding value estimation (default: FALSE)
#' @param bve.gen Generations of individuals to consider in breeding value estimation (default: NULL)
#' @param bve.cohorts Cohorts of individuals to consider in breeding value estimation (default: NULL)
#' @param bve.database Groups of individuals to consider in breeding value estimation (default: NULL)
#' @param bve.insert.gen Generations of individuals to compute breeding values for (default: all groups in bve.database)
#' @param bve.insert.cohorts Cohorts of individuals to compute breeding values for (default: all groups in bve.database)
#' @param bve.insert.database Groups of individuals to compute breeding values for (default: all groups in bve.database)
#' @param sigma.e Enviromental variance (default: 100)
#' @param sigma.e.gen Generations to consider when estimating sigma.e when using hertability
#' @param sigma.e.cohorts Cohorts to consider when estimating sigma.e when using hertability
#' @param sigma.e.database Groups to consider when estimating sigma.e when using hertability
#' @param forecast.sigma.g Set FALSE to not estimate sigma.g (Default: TRUE)
#' @param heritability Use sigma.e to obtain a certain heritability (default: NULL)
#' @param sigma.g Genetic variance (default: 100 - only used if not computed via estimate.sigma.g^2 in der Zuchtwertschaetzung (Default: 100)
#' @param sigma.g.gen Generations to consider when estimating sigma.g
#' @param sigma.g.cohorts Cohorts to consider when estimating sigma.g
#' @param sigma.g.database Groups to consider when estimating sigma.g
#' @param new.bv.observation Vector of all generation for which breeding values are observed (alt: "all" for all & "non_obs" for all non-observed individuals)
#' @param new.bv.child Starting phenotypes of newly generated individuals (default: "mean" of both parents, "obs" - regular observation, "zero" - 0)
#' @param computation.A Method to calculate pedigree matrix (Default: "vanRaden", alt: "kinship", "CE", "non_stand", "CE2", "CM")
#' @param computation.A.ogc Method to calculate pedigree matrix in OGC (Default: "kinship", alt: "vanRaden", "CE", "non_stand", "CE2", "CM")
#' @param delete.haplotypes Generations for with haplotypes of founders can be deleted (only use if storage problem!)
#' @param delete.individuals Generations for with individuals are completley deleted (only use if storage problem!)
#' @param praeimplantation Only use matings the lead to a specific genotype in a specific marker
#' @param new.phenotype.correlation Correlation of the simulated enviromental variance
#' @param new.breeding.correlation Correlation of the simulated genetic variance (child share! heritage is not influenced!)
#' @param best1.from.group (OLD!- use selection.m.database) Groups of individuals to consider as First Parent / Father (also female individuals are possible)
#' @param best2.from.group (OLD!- use selection.f.database) Groups of individuals to consider as Second Parent / Mother (also male individuals are possible)
#' @param best1.from.cohort (OLD!- use selection.m.cohorts) Groups of individuals to consider as First Parent / Father (also female individuals are possible)
#' @param best2.from.cohort (OLD! - use selection.f.cohorts) Groups of individuals to consider as Second Parent / Mother (also male individuals are possible)
#' @param fixed.breeding Set of targeted matings to perform
#' @param fixed.breeding.best Perform targeted matings in the group of selected individuals
#' @param max.offspring Maximum number of offspring per individual (default: c(Inf,Inf) - (m,w))
#' @param store.breeding.totals If TRUE store information on selected animals in $info$breeding.totals
#' @param multiple.bve Way to handle multiple traits in bv/selection (default: "add", alt: "ranking")
#' @param multiple.bve.weights.m Weighting between traits when using "add" (default: 1)
#' @param multiple.bve.weights.f Weighting between traits when using "add" (default: same as multiple.bve.weights.m)
#' @param store.bve.data If TRUE store information of bve in $info$bve.data
#' @param fixed.assignment Set TRUE for targeted mating of best-best individual till worst-worst (of selected). set to "bestworst" for best-worst mating
#' @param reduce.group (OLD! - use culling modules) Groups of animals for reduce to a new size (by changing class to -1)
#' @param reduce.group.selection (OLD! - use culling modules) Selection criteria for reduction of groups (cf. selection.m / selection.f - default: "random")
#' @param selection.critera If 0 individuals with lowest bve are selected as best individuals (default c(1,1) - (m,w))
#' @param same.sex.activ If TRUE allow matings of individuals of same sex
#' @param same.sex.sex Probability to use female individuals as parents (default: 0.5)
#' @param same.sex.selfing If FALSE no matings between an individual with itself
#' @param selfing.mating If TRUE generate new individuals via selfing
#' @param selfing.sex Share of female individuals used for selfing (default: 0.5)
#' @param multiple.bve.scale.m Set to "pheno_sd" when using gains per phenotypic SD, "unit" when using gains per unit, default: "bve_sd"
#' @param multiple.bve.scale.f Set to "pheno_sd" when using gains per phenotypic SD, "unit" when using gains per unit, default: "bve_sd"
#' @param use.last.sigma.e If TRUE use the sigma.e used in the previous simulation (default: FALSE)
#' @param class.m Migrationlevels of male individuals to consider for mating process (default: 0)
#' @param class.f Migrationlevels of female individuals to consider for mating process (default: 0)
#' @param save.recombination.history If TRUE store the time point of each recombination event
#' @param martini.selection If TRUE use the group of non-selected individuals as second parent
#' @param BGLR.bve If TRUE use BGLR to perform breeding value estimation
#' @param BGLR.model Select which BGLR model to use (default: "RKHS", alt: "BRR", "BL", "BayesA", "BayesB", "BayesC")
#' @param BGLR.burnin Number of burn-in steps in BGLR (default: 1000)
#' @param BGLR.iteration Number of iterations in BGLR (default: 5000)
#' @param BGLR.print If TRUE set verbose to TRUE in BGLR
#' @param BGLR.save Method to use in BGLR (default: "RKHS" - alt: NON currently)
#' @param BGLR.save.random Add random number to store location of internal BGLR computations (only needed when simulating a lot in parallel!)
#' @param copy.individual If TRUE copy the selected father for a mating
#' @param copy.individual.keep.bve Set to FALSE to not keep estimated breeding value in case of use of copy.individuals
#' @param dh.mating If TRUE generate a DH-line in mating process
#' @param dh.sex Share of DH-lines generated from selected female individuals
#' @param n.observation Number of phenotypes generated per individuals (influences enviromental variance)
#' @param bve.0isNA Individuals with phenotype 0 are used as NA in breeding value estimation
#' @param phenotype.bv If TRUE use phenotype as estimated breeding value
#' @param standardize.bv If TRUE standardize breeding value (additive transformation to mean standardize.bv.level)
#' @param standardize.bv.level Level for the standardization (default: 100)
#' @param standardize.bv.gen Generations to use in standardize.bv
#' @param delete.same.origin If TRUE delete recombination points when genetic origin of adjacent segments is the same
#' @param estimate.u If TRUE estimate u in breeding value estimation (Y = Xb + Zu + e)
#' @param gwas.u If TRUE estimate u via GWAS (relevant for gene editing)
#' @param approx.residuals If FALSE calculate the variance for each marker separatly instead of using a set variance (doesnt change order - only p-values)
#' @param gwas.gen Generations to consider in GWAS analysis
#' @param gwas.cohorts Cohorts to consider in GWAS analysis
#' @param gwas.database Groups to consider in GWAS analysis
#' @param gwas.group.standard If TRUE standardize phenotypes by group mean
#' @param y.gwas.used What y value to use in GWAS study (Default: "pheno", alt: "bv", "bve")
#' @param remove.effect.position If TRUE remove real QTLs in breeding value estimation
#' @param estimate.add.gen.var If TRUE estimate additive genetic variance and heritability based on parent model
#' @param estimate.pheno.var If TRUE estimate total variance in breeding value estimation
#' @param store.comp.times If TRUE store computation times in $info$comp.times (default: TRUE)
#' @param store.comp.times.bve If TRUE store computation times of breeding value estimation in $info$comp.times.bve (default: TRUE)
#' @param store.comp.times.generation If TRUE store computation times of mating simulations in $info$comp.times.generation (default: TRUE)
#' @param special.comb If TRUE selected individuals based on their special combining ability
#' @param predict.effects If TRUE use estimated effects instead of true values in special.comb
#' @param SNP.density Use every _ SNP in predict.effects (default: 10)
#' @param use.effect.markers If TRUE use QTL markers in special.comb
#' @param use.effect.combination If TRUE use QTL combinations (epistatic/mult) in special.comb
#' @param max.auswahl Maximum amount of times a individuals is considered in special.comb
#' @param ogc If TRUE use optimal genetic contribution theory to perform selection (Needs rework!)
#' @param ogc.cAc Increase of average relationship in ogc. Default: minimize inbreeding rate.
#' @param gene.editing.offspring If TRUE perform gene editing on newly generated individuals
#' @param gene.editing.best If TRUE perform gene editing on selected individuals
#' @param gene.editing.offspring.sex Which sex to perform editing on (Default c(TRUE,TRUE), mw)
#' @param gene.editing.best.sex Which sex to perform editing on (Default c(TRUE,TRUE), mw)
#' @param nr.edits Number of edits to perform per individual
#' @param import.position.calculation Function to calculate recombination point into adjacent/following SNP
#' @param special.comb.add If TRUE select matings based on additive combining ability
#' @param emmreml.bve If TRUE use REML estimator from R-package EMMREML in breeding value estimation
#' @param rrblup.bve If TRUE use REML estimator from R-package rrBLUP in breeding value estimation
#' @param sommer.bve If TRUE use REML estimator from R-package sommer in breeding value estimation
#' @param breedR.bve If TRUE use pedigree-based breeding value estimation implemented in the r-package breedR
#' @param bve.direct.est If TRUE predict BVEs in direct estimation according to vanRaden 2008 method 2 (default: TRUE)
#' @param breedR.groups Cohorts to consider for breedR-breeding value estimation
#' @param sequenceZ Split genomic matric into parts (relevent if high memory usage)
#' @param maxZ Number of SNPs to consider in each part of sequenceZ
#' @param maxZtotal Number of matrix entries to consider jointly (maxZ = maxZtotal/number of animals)
#' @param delete.sex Remove all individuals from these sex from generation delete.individuals (default: 1:2 ; note:delete individuals=NULL)
#' @param gen.architecture.m Genetic architecture for male animal (default: 0 - no transformation)
#' @param gen.architecture.f Genetic architecture for female animal (default: gen.architecture.m - no transformation)
#' @param add.architecture List with two vectors containing (A: length of chromosomes, B: position in cM of SNPs)
#' @param ncore Cores used for parallel computing in compute.snps
#' @param Z.integer If TRUE save Z as a integer in parallel computing
#' @param store.effect.freq If TRUE store the allele frequency of effect markers perss generation
#' @param backend Chose the used backend (default: "doParallel", alt: "doMPI")
#' @param Rprof Store computation times of each function
#' @param miraculix If TRUE use miraculix to perform computations (ideally already generate population in creating.diploid with this; default: automatic detection from population list)
#' @param miraculix.mult If TRUE use miraculix for matrix multiplications even if miraculix is not used for storage
#' @param miraculix.chol If TRUE perform Cholesky-decomposition implemented in miraculix (also works for semi-definite matrices)
#' @param miraculix.cores Number of cores used in miraculix applications (default: 1)
#' @param best.selection.ratio.m Ratio of the frequency of the selection of the best best animal and the worst best animal (default=1)
#' @param best.selection.ratio.f Ratio of the frequency of the selection of the best best animal and the worst best animal (default=1)
#' @param best.selection.criteria.m Criteria to calculate this ratio (default: "bv", alt: "bve", "pheno")
#' @param best.selection.criteria.f Criteria to calculate this ratio (default: "bv", alt: "bve", "pheno")
#' @param best.selection.manual.ratio.m vector containing probability to draw from for every individual (e.g. c(0.1,0.2,0.7))
#' @param best.selection.manual.ratio.f vector containing probability to draw from for every individual (e.g. c(0.1,0.2,0.7))
#' @param selection.criteria.type What to use in the selection proces (default: "bve", alt: "bv", "pheno")
#' @param bve.class Consider only animals of those class classes in breeding value estimation (default: NULL - use all)
#' @param parallel.generation Set TRUE to active parallel computing in animal generation
#' @param ncore.generation Number of cores to use in parallel generation
#' @param randomSeed Set random seed of the process
#' @param randomSeed.generation Set random seed for parallel generation process
#' @param new.selection.calculation If TRUE recalculate breeding values obtained by selection.function.matrix
#' @param fast.compiler Use a just-in-time compiler from the compiler package (Default: 0, alt: 1,2,3)
#' @param name.cohort Name of the newly added cohort
#' @param add.class.cohorts Migration levels of all cohorts selected for reproduction are automatically added to class.m/class.f (default: TRUE)
#' @param display.progress Set FALSE to not display progress bars
#' @param max.ticks Maximal number of ticks in a progress bar
#' @param ignore.best Not consider the top individuals of the selected individuals (e.g. to use 2-10 best individuals)
#' @param combine Copy existing individuals (e.g. to merge individuals from different groups in a joined cohort). Individuals to use are used as the first parent
#' @param repeat.mating Generate multiple mating from the same dam/sire combination
#' @param time.point Time point at which the new individuals are generated
#' @param creating.type Technique to generate new individuals (usage in web-based application)
#' @param multiple.observation Set TRUE to allow for more than one phenotype observation per individual (this will decrease enviromental variance!)
#' @param new.bv.observation.gen Vector of generation from which to generate additional phenotypes
#' @param new.bv.observation.cohorts Vector of cohorts from which to generate additional phenotype
#' @param new.bv.observation.database Matrix of groups from which to generate additional phenotypes
#' @param reduced.selection.panel.m Use only a subset of individuals of the potential selected ones ("Split in user-interface")
#' @param reduced.selection.panel.f Use only a subset of individuals of the potential selected ones ("Split in user-interface")
#' @param breeding.all.combination Set to TRUE to automatically perform each mating combination possible exactly ones.
#' @param depth.pedigree Depth of the pedigree in generations (default: 7)
#' @param depth.pedigree.ogc Depth of the pedigree in generations (default: 7)
#' @param bve.avoid.duplicates If set to FALSE multiple generatations of the same individual can be used in the bve (only possible by using copy.individual to generate individuals)
#' @param report.accuracy Report the accuracy of the breeding value estimation
#' @param share.genotyped Share of individuals genotyped in the founders
#' @param added.genotyped Share of individuals that is additionally genotyped (only for copy.individuals)
#' @param singlestep.active Set TRUE to use single step in breeding value estimation (only implemented for vanRaden- G matrix and without use sequenceZ) (Legarra 2014)
#' @param remove.non.genotyped Set to FALSE to manually include non-genotyped individuals in genetic BVE, single-step will deactive this as well
#' @param fast.uhat Set to FALSE to  derive inverse of A in rrBLUP
#' @param offspring.bve.parents.gen Generations to consider to derive phenotype from offspring phenotypes
#' @param offspring.bve.parents.database Groups to consider to derive phenotype from offspring phenotypes
#' @param offspring.bve.parents.cohorts Cohorts to consider to derive phenotype from offspring phenotypes
#' @param offspring.bve.offspring.gen Active generations for import of offspring phenotypes
#' @param offspring.bve.offspring.database Active groups for import of offspring phenotypes
#' @param offspring.bve.offspring.cohorts Active cohorts for import of offspring phenotypes
#' @param sommer.multi.bve Set TRUE to use a mulit-trait model in the R-package sommer for BVE
#' @param calculate.reliability Set TRUE to calculate a reliability when performing Direct-Mixed-Model BVE
#' @param selection.m.gen Generations available for selection of paternal parent
#' @param selection.f.gen Generations available for selection of maternal parent
#' @param selection.m.database Groups available for selection of paternal parent
#' @param selection.f.database Groups available for selection of maternal parent
#' @param selection.m.cohorts Cohorts available for selection of paternal parent
#' @param selection.f.cohorts Cohorts available for selection of maternal parent
#' @param selection.m.miesenberger Use Weighted selection index according to Miesenberger 1997 for paternal selection
#' @param selection.f.miesenberger Use Weighted selection index according to Miesenberger 1997 for maternal selection
#' @param selection.miesenberger.reliability.est If available reliability estimated are used. If not use default:"estimated" (SD BVE / SD Pheno), alt: "heritability", "derived" (cor(BVE,BV)^2) as replacement
#' @param culling.cohort Cohort to consider to culling
#' @param culling.time Age of the individuals at culling
#' @param culling.name Name of the culling action (user-interface stuff)
#' @param culling.bv1 Reference Breeding value
#' @param culling.share1 Probability of death for individuals with bv1
#' @param culling.bv2 Alternative breeding value (linear extended for other bvs)
#' @param culling.share2 Probability of death for individuals with bv2
#' @param culling.index Genomic index (default:0 - no genomic impact, use: "lastindex" to use the last selection index applied in selection)
#' @param verbose Set to FALSE to not display any prints
#' @param bve.parent.mean Set to TRUE to use the average parental performance as the breeding value estimate
#' @param bve.grandparent.mean Set to TRUE to use the average grandparental performance as the breeding value estimate
#' @param bve.mean.between Select if you want to use the "bve", "bv", "pheno" or "bvepheno" to form the mean (default: "bvepheno" - if available bve, else pheno)
#' @export


breeding.diploid <- function(population,
                             mutation.rate = 10^-5,
                             remutation.rate = 10^-5,
            recombination.rate = 1,
            selection.m = "random",
            selection.f = NULL,
            new.selection.calculation = TRUE,
            selection.function.matrix = NULL,
            selection.size = 0,
            ignore.best = 0,
            breeding.size = 0,
            breeding.sex = NULL,
            breeding.sex.random = FALSE,
            relative.selection = FALSE,
            class.m = 0,
            class.f = 0,
            add.gen = 0,
            recom.f.indicator = NULL,
            recom.f.polynom = NULL,
            duplication.rate = 0,
            duplication.length = 0.01,
            duplication.recombination = 1,
            new.class = 0L,
            bve = FALSE,
            sigma.e = NULL,
            sigma.g = 100,
            new.bv.child = "zero",
            computation.A = "vanRaden",
            computation.A.ogc = "kinship",
            delete.haplotypes = NULL,
            delete.individuals = NULL,
            fixed.breeding = NULL,
            fixed.breeding.best = NULL,
            max.offspring = Inf,
            store.breeding.totals = FALSE,
            forecast.sigma.g = TRUE,
            multiple.bve = "add",
            store.bve.data = FALSE,
            fixed.assignment = FALSE,
            reduce.group = NULL,
            reduce.group.selection = "random",
            selection.critera = c(TRUE,TRUE),
            selection.criteria.type = c("bve", "bve"),
            same.sex.activ = FALSE,
            same.sex.sex = 0.5,
            same.sex.selfing = TRUE,
            selfing.mating = FALSE,
            selfing.sex = 0.5,
            praeimplantation = NULL,
            heritability = NULL,
            use.last.sigma.e = FALSE,
            save.recombination.history = FALSE,
            martini.selection = FALSE,
            BGLR.bve = FALSE,
            BGLR.model = "RKHS",
            BGLR.burnin = 500,
            BGLR.iteration = 5000,
            BGLR.print = FALSE,
            copy.individual = FALSE,
            dh.mating = FALSE,
            dh.sex = 0.5,
            n.observation = 1,
            bve.0isNA = TRUE,
            phenotype.bv = FALSE,
            standardize.bv = FALSE,
            standardize.bv.level = 100,
            standardize.bv.gen = 1,
            delete.same.origin = FALSE,
            remove.effect.position = FALSE,
            estimate.u = FALSE,
            new.phenotype.correlation = NULL,
            new.breeding.correlation = NULL,
            estimate.add.gen.var = FALSE,
            estimate.pheno.var = FALSE,
            best1.from.group = NULL,
            best2.from.group = NULL,
            best1.from.cohort = NULL,
            best2.from.cohort = NULL,
            add.class.cohorts = TRUE,
            store.comp.times = TRUE,
            store.comp.times.bve = TRUE,
            store.comp.times.generation = TRUE,
            special.comb = FALSE,
            max.auswahl = Inf,
            predict.effects = FALSE,
            SNP.density = 10,
            use.effect.markers = FALSE,
            use.effect.combination = FALSE,
            import.position.calculation = NULL,
            special.comb.add = FALSE,
            BGLR.save = "RKHS",
            BGLR.save.random = FALSE,
            ogc = FALSE,
            ogc.cAc = NA,
            emmreml.bve = FALSE,
            rrblup.bve = FALSE,
            sommer.bve = FALSE,
            sommer.multi.bve=FALSE,
            breedR.bve = FALSE,
            breedR.groups = NULL,
            nr.edits = 0,
            gene.editing.offspring = FALSE,
            gene.editing.best = FALSE,
            gene.editing.offspring.sex = c(TRUE,TRUE),
            gene.editing.best.sex = c(TRUE,TRUE),
            gwas.u = FALSE,
            approx.residuals = TRUE,
            sequenceZ = FALSE,
            maxZ = 5000,
            maxZtotal = 0,
            delete.sex = 1:2,
            gwas.group.standard = FALSE,
            y.gwas.used = "pheno",
            gen.architecture.m = 0,
            gen.architecture.f = NULL,
            add.architecture = NULL,
            ncore = 1,
            ncore.generation = 1,
            Z.integer = FALSE,
            store.effect.freq = FALSE,
            backend = "doParallel",
            randomSeed = NULL,
            randomSeed.generation = NULL,
            Rprof = FALSE,
            miraculix = NULL,
            miraculix.cores = 1,
            miraculix.mult = NULL,
            miraculix.chol = FALSE,
            fast.compiler = 0,
            best.selection.ratio.m = 1,
            best.selection.ratio.f = NULL,
            best.selection.criteria.m = "bv",
            best.selection.criteria.f = NULL,
            best.selection.manual.ratio.m = NULL,
            best.selection.manual.ratio.f = NULL,
            bve.class = NULL,
            parallel.generation = FALSE,
            name.cohort = NULL,
            display.progress = TRUE,
            max.ticks = Inf,
            combine = FALSE,
            repeat.mating = 1,
            time.point = 0,
            creating.type = 0,
            multiple.observation = FALSE,
            new.bv.observation = NULL,
            new.bv.observation.gen = NULL,
            new.bv.observation.cohorts = NULL,
            new.bv.observation.database = NULL,
            bve.gen = NULL,
            bve.cohorts = NULL,
            bve.database = NULL,
            sigma.e.gen = NULL,
            sigma.e.cohorts = NULL,
            sigma.e.database = NULL,
            sigma.g.gen = NULL,
            sigma.g.cohorts = NULL,
            sigma.g.database = NULL,
            gwas.gen = NULL,
            gwas.cohorts = NULL,
            gwas.database = NULL,
            bve.insert.gen = NULL,
            bve.insert.cohorts = NULL,
            bve.insert.database = NULL,
            reduced.selection.panel.m = NULL,
            reduced.selection.panel.f = NULL,
            breeding.all.combination = FALSE,
            depth.pedigree = 7,
            depth.pedigree.ogc = 7,
            copy.individual.keep.bve = TRUE,
            bve.avoid.duplicates = TRUE,
            report.accuracy = TRUE,
            share.genotyped = 1,
            singlestep.active = FALSE,
            remove.non.genotyped = TRUE,
            added.genotyped = 0,
            fast.uhat = TRUE,
            offspring.bve.parents.gen = NULL,
            offspring.bve.parents.database = NULL,
            offspring.bve.parents.cohorts = NULL,
            offspring.bve.offspring.gen = NULL,
            offspring.bve.offspring.database = NULL,
            offspring.bve.offspring.cohorts = NULL,
            culling.cohort=NULL,
            culling.time = Inf,
            culling.name = "Not_named",
            culling.bv1 = 0,
            culling.share1 = 0,
            culling.bv2 = NULL,
            culling.share2 = NULL,
            culling.index = 0,
            calculate.reliability=FALSE,
            selection.m.gen = NULL,
            selection.f.gen = NULL,
            selection.m.database = NULL,
            selection.f.database = NULL,
            selection.m.cohorts=NULL,
            selection.f.cohorts=NULL,
            selection.m.miesenberger=FALSE,
            selection.f.miesenberger=NULL,
            selection.miesenberger.reliability.est="estimated",
            multiple.bve.weights.m = 1,
            multiple.bve.weights.f = NULL,
            multiple.bve.scale.m = "bve_sd",
            multiple.bve.scale.f = NULL,
            verbose=TRUE,
            bve.parent.mean=FALSE,
            bve.grandparent.mean=FALSE,
            bve.mean.between="bvepheno",
            bve.direct.est=TRUE
            ){


  #######################################################################
  ############################### To-Dos ################################
  #######################################################################
  # Ignore.best implementieren.
  # Duplication re-work - account for true genetic structure of duplications - known?
  # Duplikationen in Duplizierten Bereichen werden nicht doppelt dupliziert
  # Keine feste matrix-struktur in 11/12? - testelement 13 entfernen
  # Duplikation am Rand benötigt info ob am start oder Ende bei mehreren Chromosomen
  #
  # Matrixstrukur bei mehreren Zuchtwerten geht verloren wenn nur ein relevantes Tier enthalten
  #
  # STORE BREEDING.totals fur fixed.breeding implementieren
  #######################################################################
  # Implementiere default selection.size => alle
  #######################################################################
  ## Pre-work to allow for flexiblity when inputing parameter values ####
  # Initialisize parameters that were not initialized in early versions #
  #######################################################################
{
  reduced.selection.panel <- list(reduced.selection.panel.m, reduced.selection.panel.f)

  if(culling.share1>0 || (length(culling.share2)>0 && culling.share2>0)){
    culling <- TRUE
  } else{
    culling <- FALSE
  }

  if(length(randomSeed)>0){
    set.seed(randomSeed)
  }

  if(length(bve.database)==0 && length(bve.gen)==0 && length(bve.cohorts)==0){
    bve.gen <- nrow(population$info$size)
  }
  # Fill databases

  bve.gen.input <- bve.gen
  bve.database.input <- bve.database
  bve.cohorts.input <- bve.cohorts

  bve.database <- get.database(population, bve.gen, bve.database, bve.cohorts) # NOT DONE

  if(length(bve.insert.gen)==0 && length(bve.insert.cohorts)==0 && length(bve.insert.database)==0){
    bve.insert.gen <- bve.gen.input
    bve.insert.cohorts <- bve.cohorts.input
    bve.insert.database <- bve.database.input
  }
  bve.insert.database <- get.database(population, bve.insert.gen, bve.insert.database, bve.insert.cohorts) # NOT DONE

  # Add Individuals in bve.insert that are not in bve.database
  for(index in 1:nrow(bve.insert.database)){
    included <- (bve.insert.database[index,1] == bve.database[,1]) & (bve.insert.database[index,2] == bve.database[,2]) & (bve.insert.database[index,3] >= bve.database[,3]) & (bve.insert.database[index,4] <= bve.database[,4])
    if(sum(included)==0){
      if(verbose) cat("Not all individuals with breeding values to insert included in breeding value estimation.\n")
      if(verbose) cat("Missing individuals are automatically added in BVE.\n")
      bve.database <- rbind(bve.database, bve.insert.database[index,])
    }
  }
  if(length(sigma.e.gen)==0 && length(sigma.e.cohorts)==0 && length(sigma.e.database)==0){
    sigma.e.gen <- bve.gen.input
    sigma.e.cohorts <- bve.cohorts.input
    sigma.e.database <- bve.database.input
  }
  sigma.e.database <- get.database(population, sigma.e.gen, sigma.e.database, sigma.e.cohorts) # NOT DONE

  if(length(sigma.g.gen)==0 && length(sigma.g.cohorts)==0 && length(sigma.g.database)==0){
    sigma.g.gen <- bve.gen.input
    sigma.g.cohorts <- bve.cohorts.input
    sigma.g.database <- bve.database.input
  }
  sigma.g.database <- get.database(population, sigma.g.gen, sigma.g.database, sigma.g.cohorts) # NOT DONE

  if(length(gwas.gen)==0 && length(gwas.cohorts)==0 && length(gwas.database)==0){
    gwas.gen <- bve.gen.input
    gwas.cohorts <- bve.cohorts.input
    gwas.database <- bve.database.input
  }
  gwas.database <- get.database(population, gwas.gen, gwas.database, gwas.cohorts) # NOT DONE

  if(length(new.bv.observation)==1 && (new.bv.observation=="all" || new.bv.observation=="non_obs" )){
    new.bv.observation.gen <- 1:length(population$breeding)
    if(new.bv.observation=="non_obs"){
      multiple.observation <- FALSE
    }
  }
  if(length(new.bv.observation)==1 && new.bv.observation=="non_obs_m"){
    new.bv.observation.database <- cbind(1:length(population$breeding),1)
  }
  if(length(new.bv.observation)==1 && new.bv.observation=="non_obs_f"){
    new.bv.observation.database <- cbind(1:length(population$breeding),2)
  }

  new.bv.observation.database <- get.database(population, new.bv.observation.gen, new.bv.observation.database, new.bv.observation.cohorts)

  if(combine==TRUE){
    # combine is modelled via cloning with no recombination
    copy.individual <- TRUE
    selfing.mating <- TRUE
    selfing.sex <- 0
    class.m <- unique(c(class.m, class.f))
    best1.from.cohort <- c(best1.from.cohort, best2.from.cohort)
    best2.from.cohort <- NULL
    best1.from.group <- c(best1.from.group, best2.from.group)
    best2.from.group <- NULL
    max.offspring = c(1,1)
  }

  if(length(selection.size)==1){
    selection.size <- rep(selection.size,2)
  }

  if(length(best1.from.cohort)){
    if(length(selection.m.cohorts)==0){
      selection.m.cohorts <- best1.from.cohort
    } else{
      selection.m.cohorts <- c(selection.m.cohorts,best1.from.cohort)
    }
  }
  if(length(best2.from.cohort)){
    if(length(selection.m.cohorts)==0){
      selection.f.cohorts <- best2.from.cohort
    } else{
      selection.f.cohorts <- c(selection.f.cohorts,best2.from.cohort)
    }
  }
  if(length(best1.from.group)){
    if(length(selection.m.database)==0){
      selection.m.database <- best1.from.group
    } else{
      selection.m.database <- c(selection.m.database,best1.from.group)
    }
  }
  if(length(best2.from.group)){
    if(length(selection.f.database)==0){
      selection.f.database <- best2.from.group
    } else{
      selection.f.database <- c(selection.f.database,best2.from.group)
    }
  }

  if(length(  population$info$bv.random.activ)==0){
    population$info$bv.random.activ <- which(population$info$bv.random[1:population$info$bv.calc]==FALSE)
  }

  if(length(selection.m.gen)==0 && length(selection.m.database)==0 && length(selection.m.cohorts)==0 && selection.size[1]>0){
    if(verbose) cat("No individuals for selection provided (male side). Use last available.\n")
    selection.m.database <- cbind(max(which(population$info$size[,1]>0)),1)
  }
  if(length(selection.f.gen)==0 && length(selection.f.database)==0 && length(selection.f.cohorts)==0 && selection.size[2]>0){
    if(verbose) cat("No individuals for selection provided (female side). Use last available.\n")
    selection.f.database <- cbind(max(which(population$info$size[,2]>0)),2)
  }

  selection.m.database <- get.database(population, selection.m.gen, selection.m.database, selection.m.cohorts)
  selection.f.database <- get.database(population, selection.f.gen, selection.f.database, selection.f.cohorts)

  if(length(population$info$cumsnp)==0){
    population$info$cumsnp <- cumsum(population$info$snp)
  }
  if(length(gen.architecture.f)==0){
    gen.architecture.f=gen.architecture.m
  }
  if(length(selection.criteria.type)==1){
    selection.criteria.type <- rep(selection.criteria.type,2)
  }
  if(length(max.offspring)==1){
    max.offspring <- rep(max.offspring,2)
  }
  best.selection.ratio <- list()
  best.selection.ratio[[1]] <- best.selection.ratio.m
  if(length(best.selection.ratio.f)==0){
    best.selection.ratio.f <- best.selection.ratio.m
  }
  best.selection.ratio[[2]] <- best.selection.ratio.f

  best.selection.criteria <- list()
  best.selection.criteria[[1]] <- best.selection.criteria.m
  if(length(best.selection.criteria.f)==0){
    best.selection.criteria.f <- best.selection.criteria.m
  }
  best.selection.criteria[[2]] <- best.selection.criteria.f

  best.selection.manual.ratio <- list()
  best.selection.manual.ratio[[1]] <- best.selection.manual.ratio.m
  if(length(best.selection.manual.ratio.f)==0 && length(best.selection.manual.ratio.f)!=0 ){
    best.selection.manual.ratio.f <- best.selection.manual.ratio.m
  }
  best.selection.manual.ratio[[2]] <- best.selection.manual.ratio.f
  best.selection.manual.ratio[[3]] <- "placeholder"

  if(length(population$info$origin.gen)>0){
    population$info$origin.gen <- base::as.integer(population$info$origin.gen)
  } else{
    population$info$origin.gen <- 1:64L
  }

  if(Rprof){
    Rprof()
  }

  if(length(population$info$origin)==0){
    population$info$origin <- 1:64
  }

  if(fast.compiler && requireNamespace("compiler", quietly = TRUE)){
    compiler::enableJIT(3)
  }
  if(sum(population$info$snp)<=maxZ){
    sequenceZ <- FALSE
  }
  if(length(miraculix)==0){
    miraculix <- population$info$miraculix
    if(length(miraculix)==0){
      stop("No input on use of miraculix provided!")
    }
  }
  if(miraculix && length(miraculix.mult)==0){
    miraculix.mult <- TRUE
  } else if(length(miraculix.mult)==0){
    miraculix.mult <- FALSE
  }
  if(store.comp.times){
    comp.times <- numeric(7)
    comp.times[1] <- as.numeric(Sys.time())
  }

  if(gene.editing.offspring || gene.editing.best){
    gene.editing <- TRUE
  } else{
    gene.editing <- FALSE
  }
  if(length(population$info$bitstoring)>0){
    bit.storing <- TRUE
    nbits <- population$info$bitstoring
    if(maxZ%%nbits!=0){
      maxZ <- maxZ + nbits - maxZ%%nbits
    }
  } else{
    nbits <- 30
    bit.storing <- FALSE
  }
  # Increase maxZ to improve run-time based on bitwise computing.
  if(miraculix && maxZ%%512!=0){
    maxZ <- maxZ + 512 - maxZ%%512
  }

  if(length(add.architecture)>0){
    population$info$gen.architecture[[length(population$info$gen.architecture)+1]] <- list()
    population$info$gen.architecture[[length(population$info$gen.architecture)]]$length.total <- cumsum(c(0,add.architecture[[1]]))
    population$info$gen.architecture[[length(population$info$gen.architecture)]]$snp.position <- add.architecture[[2]]

  }
  if (requireNamespace("miraculix", quietly = TRUE)) {
    if(miraculix.cores>1){
      RandomFieldsUtils::RFoptions(cores = miraculix.cores)
    }
    codeOriginsU <- miraculix::codeOrigins
    decodeOriginsU <- miraculix::decodeOrigins
  } else{
    codeOriginsU <- codeOriginsR
    decodeOriginsU <- decodeOriginsR
  }


  if(length(new.phenotype.correlation)>0){
    population$info$pheno.correlation <- t(chol(new.phenotype.correlation))
  }
  if(length(new.breeding.correlation)>0){
    population$info$bv.correlation <- new.breeding.correlation
  }


  #standardize.bv nie betrachtete Werte nicht standardisieren.


  class <- list()
  class[[1]] <- class.m
  class[[2]] <- class.f
  if(length(sigma.e)==0 && length(population$info$last.sigma.e.value)>0){
    sigma.e <- population$info$last.sigma.e.value
  } else{
    sigma.e <- 100
  }
  if(use.last.sigma.e){
    if(length(population$info$last.sigma.e.value)){
      sigma.e <- population$info$last.sigma.e.value
    }
  }
  if(length(sigma.e)==1){
    sigma.e <- rep(sigma.e, population$info$bv.nr)
  }
  if(length(sigma.g)==1){
    sigma.g <- rep(sigma.g, population$info$bv.nr)
  }
  # Berechnung der tatsaechlichen Zuchtwerte

  if(store.comp.times){
    comp.times[2] <- as.numeric(Sys.time())
  }


  if(length(selection.function.matrix)!=0 && is.matrix(selection.function.matrix)==FALSE){
    selection.function.matrix <- matrix(selection.function.matrix,nrow=1)
  }
  if(length(selection.f)==0){
    selection.f <- selection.m
  }
  if(length(multiple.bve.weights.m)< population$info$bv.nr){
    multiple.bve.weights.m <- rep(multiple.bve.weights.m, length.out = population$info$bv.nr)
  }
  if(length(multiple.bve.weights.f)==0){
    multiple.bve.weights.f <- multiple.bve.weights.m
  }
  if(length(multiple.bve.weights.f)< population$info$bv.nr){
    multiple.bve.weights.f <- rep(multiple.bve.weights.f, length.out = population$info$bv.nr)
  }
  if(length(multiple.bve.scale.f)==0){
    multiple.bve.scale.f <- multiple.bve.scale.m
  }

  if(length(selection.f.miesenberger)==0){
    selection.f.miesenberger <- selection.m.miesenberger
  }


  if(length(ignore.best)==1){
    ignore.best <- rep(ignore.best,2)
  }
  if(length(breeding.sex)==0 && length(breeding.size)==2){
    breeding.sex <- breeding.size[2] / sum(breeding.size)
  } else if(length(breeding.sex)==0){
    breeding.sex <- 0.5
  }
  if(length(breeding.size)==1){
    breeding.size.total <- breeding.size
    breeding.size.m <- round(breeding.size*(1-breeding.sex))
    if(breeding.sex.random){
      breeding.size.m <- sum(stats::rbinom(breeding.size,1,(1-breeding.sex)))
    }
    breeding.size <- c(breeding.size.m, breeding.size - breeding.size.m)
  } else{
    breeding.size.total <- sum(breeding.size)
  }

  sex.animal <- rep(2, breeding.size.total)
  if(breeding.size[1] > 0){
    sex.animalr <- sample(1:breeding.size.total, breeding.size[1])
    sex.animal[sex.animalr] <- 1
  }


  if(add.gen==0){
    current.gen <- length(population$breeding)
  } else{
    current.gen <- add.gen - 1
  }

  if(breeding.all.combination){
    if(selection.size[1]>0 && selection.size[2]>0){
      fixed.breeding.best <- matrix(0, nrow= selection.size[1] * selection.size[2], ncol=4)
      for(index in 1:selection.size[1]){
        fixed.breeding.best[1:selection.size[2] + (index-1) * selection.size[2],] <- cbind(1,index, 2, 1:selection.size[2])
      }
    } else{
      nindi <- max(selection.size[1:2])
      sex.fixed <- 1 + as.numeric(selection.size[2]>0)
      fixed.breeding.best <- matrix(0, nrow= nindi * (nindi-1) / 2, ncol=4)
      prior <- 0
      for(index in 1:(nindi-1)){
        fixed.breeding.best[1:(nindi-index)+ prior,] <- cbind(sex.fixed,index, sex.fixed, (index+1):nindi)
        prior <- prior +nindi-index
      }
    }
  }

  if(length(fixed.breeding) >0 && ncol(fixed.breeding)==6){
    fixed.breeding <- cbind(fixed.breeding,breeding.sex)
  }
  if(length(fixed.breeding.best) >0 && ncol(fixed.breeding.best)==4){
    fixed.breeding.best <- cbind(fixed.breeding.best,breeding.sex)
  }
  if(repeat.mating>1 && length(fixed.breeding)>0){
    fixed.breeding <- matrix(rep(t(fixed.breeding), repeat.mating), ncol=7, byrow=TRUE)
  }
  if(repeat.mating>1 && length(fixed.breeding.best)>0){
    fixed.breeding <- matrix(rep(t(fixed.breeding.best), repeat.mating), ncol=5, byrow=TRUE)
  }

}
  #######################################################################
  ############## Start of the actual Simulation #########################
  #######################################################################
{
  if(standardize.bv){
    for(bven in 1:population$info$bv.nr){
      bv_previous <- c(population$breeding[[standardize.bv.gen]][[7]][bven,],population$breeding[[standardize.bv.gen]][[8]][bven,])
      needed_change <- standardize.bv.level - mean(bv_previous)
      population$info$base.bv <- population$info$base.bv + needed_change
      for(times in 1:length(population$breeding)){
        for(to_change in c(3,4,7,8,9,10)){
          population$breeding[[times]][[to_change]][bven,] <- population$breeding[[times]][[to_change]][bven,] + needed_change
        }

      }
    }
  }

  # Only first TRUE/FALSE necessary for new datasets.
  if(population$info$bv.calculated==FALSE || length(population$info$effect.p)==0 || (length(population$info$effect.p.add)==0 && length(population$info$effect.p.mult1)==0 && length(population$info$effect.p.dice)==0)){
    excludes <- NULL
    snp.before <- cumsum(c(0,population$info$snp))
    population$info$effect.p.add <- list()
    if(population$info$real.bv.length[1]>0){
      for(index in 1:population$info$real.bv.length[1]){
        if(length(population$info$real.bv.add[[index]])>0 && nrow(population$info$real.bv.add[[index]])>0){
          population$info$effect.p.add[[index]] <- population$info$real.bv.add[[index]][,1]+ snp.before[population$info$real.bv.add[[index]][,2]]
          excludes <- unique(c(excludes, population$info$effect.p.add))

        }
      }
    }

    if(population$info$real.bv.length[2]>0){
      population$info$effect.p.mult1 <- list()
      population$info$effect.p.mult2 <- list()
      for(index in 1:population$info$real.bv.length[2]){
        if(length(population$info$real.bv.mult[[index]])>0 &&nrow(population$info$real.bv.mult[[index]])>0){
          population$info$effect.p.mult1[[index]] <- population$info$real.bv.mult[[index]][,1]+ snp.before[population$info$real.bv.mult[[index]][,2]]
          population$info$effect.p.mult2[[index]] <- population$info$real.bv.mult[[index]][,3]+ snp.before[population$info$real.bv.mult[[index]][,4]]
          excludes <- unique(c(excludes, population$info$effect.p.mult1,population$info$effect.p.mult2))
        }
      }
    }
    if(population$info$real.bv.length[3]>0){

      for(index in 1:population$info$real.bv.length[3]){
        if(length(population$info$real.bv.dice[[index]])>0){
          for(index2 in 1:length(population$info$real.bv.dice[[index]][[1]])){
            population$info$effect.p.dice <- c(population$info$effect.p.dice, population$info$real.bv.dice[[index]][[1]][[index2]][,1]+ snp.before[population$info$real.bv.dice[[index]][[1]][[index2]][,2]])
          }
          excludes <- unique(c(excludes, population$info$effect.p.dice))
        }
      }
    }
    population$info$effect.p <- unlist(excludes)

  }

  temp1 <- population$info$bv.calculated
  if(population$info$bve && population$info$bv.calculated==FALSE){
    if(verbose) cat("Derive genomic values of founders. \n")
    for(index in 1:length(population$breeding)){
      for(sex in 1:2){
        nanimals <- length(population$breeding[[index]][[sex]])
        if(nanimals >0){
          for(nr.animal in 1:nanimals){
            activ_bv <- population$info$bv.random.activ
            if(length(activ_bv)>0){
              temp_out <- calculate.bv(population, index, sex, nr.animal,
                                       activ_bv, import.position.calculation=import.position.calculation,
                                       decodeOriginsU=decodeOriginsU,
                                       store.effect.freq=store.effect.freq,
                                       bit.storing=bit.storing, nbits=nbits, output_compressed=FALSE)
              population$breeding[[index]][[6+sex]][activ_bv,nr.animal] <- temp_out[[1]]
              if(store.effect.freq){
                if(length(population$info$store.effect.freq) < index || length(population$info$store.effect.freq[[index]])==0){
                  population$info$store.effect.freq[[index]] <- temp_out[[2]]
                } else{
                  population$info$store.effect.freq[[index]] <- population$info$store.effect.freq[[index]] + temp_out[[2]]
                }
              }
            }
          }
        }
      }
    }
    population$info$bv.calculated <- TRUE
  }

  # Reale Zuchtwerte fuer Nicht-SNP-Effekte
  if(population$info$bv.calc>0 && population$info$bv.random[population$info$bv.calc]){
    mu1 <- numeric((population$info$bv.calc-1))
    # Schaetzung sigma.g fuer SNP-Effekte-Effekte
    if(population$info$bv.calc>1){

      n.animals <- 0
      for(index in 1:nrow(sigma.g.database)){
        n.animals <- n.animals + diff(sigma.g.database[index,3:4]) + 1
      }
      y_real <- array(0, dim=c(n.animals,(population$info$bv.calc-1))) # schaetzung sigma.g
      cindex <- 1
      temp1 <- 1:(population$info$bv.nr)
      for(index in 1:nrow(sigma.g.database)){
        k.database <- sigma.g.database[index,]
        if(diff(k.database[3:4])>=0){
          for(kindex in k.database[3]:k.database[4]){
            y_real[cindex,temp1] <- population$breeding[[k.database[[1]]]][[6+k.database[[2]]]][bven,temp1]
            cindex <- cindex +1
          }
        }
      }
      for(bven in temp1){
        population$info$bv.random.variance[bven] <- stats::var(y_real[,bven])
        mu1[bven] <- mean(y_real[,bven])
      }

    }
    population$info$current.bv.correlation <- population$info$bv.correlation
    if(sum(is.na(population$info$bv.correlation))){
      emp_cor <- which(is.na(population$info$bv.correlation), arr.ind=TRUE)
      for(index in 1:nrow(emp_cor)){
        population$info$current.bv.correlation[emp_cor[index,1], emp_cor[index,2]] <-
          stats::cor(y_real[,emp_cor[index,1]], y_real[,emp_cor[index,2]])
      }
    }
    if(temp1==FALSE){
      population$info$current.bv.random.variance  <- population$info$bv.random.variance
      if(population$info$bv.calc==1){
        bv.var <- diag(sqrt(population$info$current.bv.random.variance)) %*%population$info$current.bv.correlation %*% diag(sqrt(population$info$current.bv.random.variance))
      } else{
        AA <- diag(sqrt(population$info$current.bv.random.variance)[1:(population$info$bv.calc-1)]) %*% population$info$current.bv.correlation[1:(population$info$bv.calc-1), 1:(population$info$bv.calc-1)]%*% diag(sqrt(population$info$current.bv.random.variance)[(1:(population$info$bv.calc-1))])
        BB <- diag(sqrt(population$info$current.bv.random.variance)[1:(population$info$bv.calc-1)]) %*%population$info$current.bv.correlation[1:(population$info$bv.calc-1), -(1:(population$info$bv.calc-1))]%*% diag(sqrt(population$info$current.bv.random.variance)[-(1:(population$info$bv.calc-1))])
        CC <- diag(sqrt(population$info$current.bv.random.variance)[-(1:(population$info$bv.calc-1))]) %*%population$info$current.bv.correlation[-(1:(population$info$bv.calc-1)), -(1:(population$info$bv.calc-1))] %*% diag(sqrt(population$info$current.bv.random.variance)[-(1:(population$info$bv.calc-1))])
        if (requireNamespace("MASS", quietly = TRUE)) {
          bv.var <- CC - t(BB) %*% MASS::ginv(AA) %*% BB
          mu_mult <- t(BB) %*% MASS::ginv(AA)
        } else{
          bv.var <- CC - t(BB) %*% solve(AA) %*% BB
          mu_mult <- t(BB) %*% solve(AA)
        }
      }
      bv.var_chol <- t(chol(bv.var))
      population$info$bv.correlation_col <- bv.var_chol
      for(index in 1:length(population$breeding)){
        for(sex in 1:2){
          nanimals <- length(population$breeding[[index]][[sex]])
          if(nanimals >0){
            for(nr.animal in 1:nanimals){
              if(population$info$bv.calc==1){
                bv.mean <- population$info$base.bv
              } else{
                bv.mean <- population$info$base.bv[-(1:(population$info$bv.calc-1))] +  mu_mult %*% (population$breeding[[index]][[sex+6]][1:(population$info$bv.calc-1), nr.animal] - mu1)
              }
              population$breeding[[index]][[6+sex]][population$info$bv.calc:population$info$bv.nr, nr.animal] <- bv.mean + bv.var_chol %*% stats::rnorm(population$info$bv.nr-population$info$bv.calc+1,0,1)
            }
          }
        }
      }
    }

  }
  if(store.comp.times){
    comp.times[3] <- as.numeric(Sys.time())
  }


  if(length(heritability)>0){

    if(length(population$info$last.sigma.e.database)==0 || !(nrow(population$info$last.sigma.e.database)==nrow(sigma.e.database) && prod(population$info$last.sigma.e.database==sigma.e.database)==1) || length(heritability)>0 && (length(population$info$last.sigma.e.heritability)==0 || prod(population$info$last.sigma.e.heritability==heritability)==0)){
      if(verbose) cat("Start deriving enviromental variance (according to given heritability).\n")
      if(length(heritability)!=population$info$bv.nr){
        heritability <- rep(heritability, population$info$bv.nr)
      }
      n.animals <- 0
      for(index in 1:nrow(sigma.e.database)){
        n.animals <- n.animals + diff(sigma.e.database[index,3:4]) + 1
      }
      y_real <- array(0, dim=c(n.animals,(population$info$bv.nr))) # schaetzung sigma.g

      cindex <- 1
      temp1 <- 1:(population$info$bv.nr)
      for(index in 1:nrow(sigma.e.database)){
        k.database <- sigma.e.database[index,]
        if(diff(k.database[3:4])>=0){
          for(kindex in k.database[3]:k.database[4]){
            y_real[cindex,temp1] <- population$breeding[[k.database[1]]][[6+k.database[2]]][temp1,kindex]
            cindex <- cindex +1
          }
        }
      }

      for(bven in 1:population$info$bv.nr){
        if(forecast.sigma.g){
          sigma.g.temp <- stats::var(y_real[,bven])
        }
        sigma.e[bven] <- ((1- heritability[bven]) * sigma.g.temp)/ heritability[bven]
      }
    } else{
      sigma.e <- population$info$last.sigma.e.value
    }

  }


  if(length(new.bv.observation.database)>0 && population$info$bve && sum(n.observation)>0){
    if(length(n.observation)<population$info$bv.nr){
      n.observation <- rep(n.observation, length.out=population$info$bv.nr)

    }
    if(verbose) cat("Start simulating phenotypes.\n")

    observation_reps <- sort(unique(c(0,n.observation)))
    for(index in 1:nrow(new.bv.observation.database)){
      gen <- new.bv.observation.database[index,1]
      sex <- new.bv.observation.database[index,2]
      if(diff(new.bv.observation.database[index,3:4])>=0){
        for(nr.animal in new.bv.observation.database[index,3]:new.bv.observation.database[index,4]){
          multi_check <- (population$breeding[[gen]][[sex]][[nr.animal]][[15]]==0)
          for(observation_rep in 2:length(observation_reps)){
            new.obs <- observation_reps[observation_rep] - observation_reps[observation_rep-1]
            temp_random <- matrix(stats::rnorm(population$info$bv.nr*new.obs,0,1), ncol=new.obs)
            active.traits <- (n.observation >=observation_reps[observation_rep])
            active.traits <- active.traits*(1:length(active.traits))
            if(!multiple.observation){
              active.traits <- active.traits * multi_check
              # No phenotypes for traits which were already observed
            }
            for(bven in (1:population$info$bv.nr)[active.traits]){
              bobs <- population$breeding[[gen]][[sex]][[nr.animal]][[15]][bven]
              fobs <- new.obs
              population$breeding[[gen]][[8+sex]][bven,nr.animal] <-  bobs/(bobs+fobs) * population$breeding[[gen]][[8+sex]][[bven,nr.animal]] + fobs/(bobs+fobs) * (rowMeans(population$info$pheno.correlation %*% temp_random)[bven] * sqrt(sigma.e[bven]) + population$breeding[[gen]][[6+sex]][bven, nr.animal])
              population$breeding[[gen]][[sex]][[nr.animal]][[15]][bven] <- population$breeding[[gen]][[sex]][[nr.animal]][[15]][bven] + new.obs
            }
          }
        }
      }
    }
  }


  # Import offspring phenotypes for parents
  offspring.bve.parents.database <- get.database(population, offspring.bve.parents.gen, offspring.bve.parents.database, offspring.bve.parents.cohorts)
  if(length(offspring.bve.parents.database)>0){
    offspring.bve <- TRUE
  } else{
    offspring.bve <- FALSE
  }

  if(length(offspring.bve.offspring.gen)>0 || length(offspring.bve.offspring.database)>0 || length(offspring.bve.offspring.cohorts)>0){
    offspring.bve.offspring.database <- get.database(population, offspring.bve.offspring.gen, offspring.bve.offspring.database, offspring.bve.offspring.cohorts)
  } else if(offspring.bve){
    if(verbose) cat("No potential offspring for phenotype import given. Consider all potential individuals")
    first_gen <- min(offspring.bve.parents.database[,1])
    list_of_copy <- list()
    for(gen_check in 1:nrow(offspring.bve.parents.database)){
      for(gen_check2 in offspring.bve.parents.database[gen_check,3]:offspring.bve.parents.database[gen_check,4]){
        first_gen <- min(first_gen, population$breeding[[offspring.bve.parents.database[gen_check,1]]][[offspring.bve.parents.database[gen_check,2]]][[gen_check2]][[21]][,1])
        list_of_copy[[length(list_of_copy)+1]] <- rbind(offspring.bve.parents.database[gen_check,1], offspring.bve.parents.database[gen_check,2], gen_check2 ,
                                                         t(population$breeding[[offspring.bve.parents.database[gen_check,1]]][[offspring.bve.parents.database[gen_check,2]]][[gen_check2]][[21]]))
      }
    }
    list_of_copy <- matrix(unlist(list_of_copy), ncol=6, byrow=TRUE)
    list_of_copy <- list_of_copy[!(list_of_copy[,1]==list_of_copy[,4] & list_of_copy[,2]==list_of_copy[,5] & list_of_copy[,3]==list_of_copy[,6]),]
    ped_off <- get.pedigree(population, gen = first_gen:nrow(population$info$size), raw=TRUE)
    candidates <- rep(TRUE,nrow(ped_off))
    for(index in 1:nrow(offspring.bve.parents.database)){
      act <- offspring.bve.parents.database[index,]
      candidates[ped_off[,1]==act[1] & ped_off[,2] == act[2] & ped_off[,3]>= act[3] & ped_off[,6]<=act[4]] <- FALSE
    }
    ped_off <- ped_off[candidates,]

    candidates <- rep(FALSE,nrow(ped_off))
    for(index in 1:nrow(offspring.bve.parents.database)){
      act <- offspring.bve.parents.database[index,]
      candidates[ped_off[,4]==act[1] & ped_off[,5] == act[2] & ped_off[,6]>= act[3] & ped_off[,6]<=act[4]] <- TRUE
      candidates[ped_off[,7]==act[1] & ped_off[,8] == act[2] & ped_off[,9]>= act[3] & ped_off[,9]<=act[4]] <- TRUE
    }
    candidates <- TRUE
    male_candidates <- (ped_off[candidates & ped_off[,2]==1 ,])
    female_candidates <- (ped_off[candidates & ped_off[,2]==2 ,])

    male_gen <- unique(male_candidates[,1])
    female_gen  <- unique(female_candidates[,1])

    offspring.bve.offspring.database <- NULL
    if(length(male_gen)>0){
      for(index in male_gen){
        offspring.bve.offspring.database <- rbind(offspring.bve.offspring.database,
                                                  c(index,1,min(male_candidates[male_candidates[,1]==index,3]),
                                                    max(male_candidates[male_candidates[,1]==index,3])))
      }
    }
    if(length(female_gen)>0){
      for(index in female_gen){
        offspring.bve.offspring.database <- rbind(offspring.bve.offspring.database,
                                                  c(index,2,min(female_candidates[female_candidates[,1]==index,3]),
                                                    max(female_candidates[female_candidates[,1]==index,3])))
      }
    }

  }

  if(offspring.bve){
    offspring_id <- get.id(population, offspring.bve.offspring.database)
    is_double <- duplicated(offspring_id[length(offspring_id):1])[length(offspring_id):1]
    if(verbose) cat("Import phenotypes of offspring.\n")
    for(index in 1:nrow(offspring.bve.parents.database)){
      activ.parents <- offspring.bve.parents.database[index,]
      new.bv <- counter <- matrix(0, nrow=population$info$bv.nr, ncol=activ.parents[4]-activ.parents[3]+1)
      next_indi <- 1
      for(index2 in 1:nrow(offspring.bve.offspring.database)){
        activ.offspring <- offspring.bve.offspring.database[index2,]
        n.animals <- activ.offspring[4]-activ.offspring[3] +1
        if(n.animals>0){
          for(index3 in (activ.offspring[3]:activ.offspring[4])[!is_double[next_indi:(next_indi+n.animals-1)]]){
            parent1 <- population$breeding[[activ.offspring[1]]][[activ.offspring[2]]][[index3]][[7]]
            parent2 <- population$breeding[[activ.offspring[1]]][[activ.offspring[2]]][[index3]][[8]]
            if(length(list_of_copy)>0){
              replace1 <- which(parent1[1]==list_of_copy[,4] & parent1[2]==list_of_copy[,5] & parent1[3]==list_of_copy[,6])
              replace2 <- which(parent2[1]==list_of_copy[,4] & parent2[2]==list_of_copy[,5] & parent2[3]==list_of_copy[,6])
              if(length(replace1)==1){
                parent1 <- list_of_copy[replace1,1:3]
              }
              if(length(replace2)==1){
                parent2 <- list_of_copy[replace2,1:3]
              }
            }
            if(parent1[1]==activ.parents[1] && parent1[2]==activ.parents[2] && parent1[3]>= activ.parents[3] && parent1[3]<= activ.parents[4]){
              new.bv[,parent1[3] - activ.parents[3]+1] <- new.bv[,parent1[3]- activ.parents[3]+1] + population$breeding[[activ.offspring[1]]][[activ.offspring[2]+8]][,index3] * population$breeding[[activ.offspring[1]]][[activ.offspring[2]]][[activ.offspring[3]]][[15]]
              counter[,parent1[3]- activ.parents[3]+1] <- counter[,parent1[3]- activ.parents[3]+1] + population$breeding[[activ.offspring[1]]][[activ.offspring[2]]][[activ.offspring[3]]][[15]]
            }
            if(parent2[1]==activ.parents[1] && parent2[2]==activ.parents[2] && parent2[3]>= activ.parents[3] && parent2[3]<= activ.parents[4]){
              new.bv[,parent2[3]- activ.parents[3]+1] <- new.bv[,parent2[3]- activ.parents[3]+1] + population$breeding[[activ.offspring[1]]][[activ.offspring[2]+8]][,index3]
              counter[,parent2[3]- activ.parents[3]+1] <- counter[,parent2[3]- activ.parents[3]+1] + population$breeding[[activ.offspring[1]]][[activ.offspring[2]]][[activ.offspring[3]]][[15]]
            }
          }
        }
        next_indi <- next_indi + n.animals

      }
      population$breeding[[activ.parents[1]]][[activ.parents[2]+2]][,activ.parents[3]:activ.parents[4]] <- new.bv / counter
      population$breeding[[activ.parents[1]]][[activ.parents[2]+8]][,activ.parents[3]:activ.parents[4]] <- new.bv / counter
      if(sum(counter==0)>0){
        if(verbose) cat(paste0(sum(counter==0), " phenotype entries without valid offspring for phenotype import from offspring! Set phenotype to 0."))
        population$breeding[[activ.parents[1]]][[activ.parents[2]+2]][counter==0] <- 0
        population$breeding[[activ.parents[1]]][[activ.parents[2]+8]][counter==0] <- 0
      }
    }

  }
}
{
  ## Culling Module
  if(culling){
    if(culling.index=="lastindex"){
      culling.index <- get.selectionindex(population, cohorts=culling.cohort)
    }
    if(population$info$bv.nr>0){
      culling.bv <- colSums(get.bv(population, cohorts = culling.cohort) * culling.index)
    } else{
      culling.bv <- numeric(length(get.id(population, cohorts= culling.cohort)))
    }
    n_animals <- length(culling.bv)

    if(sum(abs(culling.index))==0){
      culling.prob <- rep(culling.share1, n_animals)
    } else{
      culling.prob <- (culling.bv - culling.bv1) * (culling.share1 - culling.share2)/ (culling.bv1 - culling.bv2) + culling.share1
      culling.prob[culling.prob>1] <- 1
      culling.prob[culling.prob<0] <- 0
    }
    culling.action <- stats::rbinom(n_animals,1,culling.prob)==1

    culling.database <- get.database(population, cohorts=culling.cohort)
    store <- population$breeding[[culling.database[1]]][[culling.database[2]+4]][culling.database[3]:culling.database[4]]
    population$breeding[[culling.database[1]]][[culling.database[2]+4]][culling.database[3]:culling.database[4]][culling.action] <- (-1)
    new_death <- population$breeding[[culling.database[1]]][[culling.database[2]+4]][culling.database[3]:culling.database[4]] != store
    n_death <- sum(new_death)

    if(n_death>0){
      population$breeding[[culling.database[1]]][[culling.database[2]+16]][culling.database[3]:culling.database[4]][which(new_death)] <- population$breeding[[culling.database[1]]][[culling.database[2]+10]][culling.database[3]:culling.database[4]][which(new_death)] + culling.time
      active_cohort <- which(population$info$cohorts[,1]==culling.cohort)
      if(length(population$info$culling.stats)<active_cohort){
        population$info$culling.stats[[active_cohort]] <- cbind(culling.name, n_death)
      } else{
        population$info$culling.stats[[active_cohort]] <- rbind(population$info$culling.stats[[active_cohort]],
                                                                c(culling.name, n_death))
      }

    }
  }
}

{
  ## Breeding value estimation
  if(store.comp.times){
    comp.times[4] <- as.numeric(Sys.time())
  }

  ## Track computing times
  if(store.comp.times.bve){
    comp.times.bve <- numeric(5)
    z_chol <- 0
    z_uhat <- 0
    zcalc <- 0
    z_ped <- 0
    z_h <- 0
    comp.times.bve[1] <- as.numeric(Sys.time())
  }


  u_hat <- NULL
  gwas_hat <- NULL
  u_hat_possible <- FALSE

  ## BVE according to average performance of parents / grandparents
  if(bve.parent.mean || bve.grandparent.mean){
    addpheno <- FALSE
    if(bve.mean.between=="pheno"){
      addmean <- 8
    } else if(bve.mean.between=="bv"){
      addmean <- 6
    } else if(bve.mean.between=="bve"){
      addmean <- 2
    } else{
      addmean <- 2
      addpheno <- TRUE
    }
    for(index in 1:nrow(bve.insert.database)){
      activ.base <- bve.insert.database[index,]
      if(diff(activ.base[3:4])>=0){
        for(index2 in activ.base[3]:activ.base[4]){
          reference <- rbind(population$breeding[[activ.base[1]]][[activ.base[2]]][[index2]][[7]],population$breeding[[activ.base[1]]][[activ.base[2]]][[index2]][[8]])
          if(bve.grandparent.mean){
            reference <- rbind(population$breeding[[reference[1,1]]][[reference[1,2]]][[reference[1,3]]][[7]],
                              population$breeding[[reference[1,1]]][[reference[1,2]]][[reference[1,3]]][[8]],
                              population$breeding[[reference[2,1]]][[reference[2,2]]][[reference[2,3]]][[7]],
                              population$breeding[[reference[2,1]]][[reference[2,2]]][[reference[2,3]]][[8]])
          }
          bve_import <- matrix(0, nrow=nrow(reference), ncol=population$info$bv.nr)
          for(index3 in 1:nrow(reference)){
            bve_import[index3,] <- population$breeding[[reference[index3,1]]][[reference[index3,2]+addmean]][,reference[index3,3]]
          }
          if(addpheno){
            for(index3 in 1:nrow(reference)){
              bve_import[index3,bve_import[index3,]==0] <- population$breeding[[reference[index3,1]]][[reference[index3,2]+8]][bve_import[index3,]==0,reference[index3,3]]
            }
          }
          bve_import[bve_import==0] <- NA
          population$breeding[[activ.base[1]]][[activ.base[2]+2]][,index2] <- colMeans(bve_import, na.rm=TRUE)
          if(sum(is.na(population$breeding[[activ.base[1]]][[activ.base[2]+2]][,index2]))>0){
            population$breeding[[activ.base[1]]][[activ.base[2]+2]][is.na(population$breeding[[activ.base[1]]][[activ.base[2]+2]][,index2]),index2] <- 0
          }

        }
      }
      if(report.accuracy){
        y_real_report <- NULL
        y_hat_report <- NULL
        for(index in 1:nrow(bve.insert.database)){
          activ.base <- bve.insert.database[index,]
          if(diff(activ.base[3:4])>=0){
            y_real_report <- cbind(y_real_report, population$breeding[[activ.base[1]]][[activ.base[2]+6]][,activ.base[3]:activ.base[4], drop=FALSE])
            y_hat_report <- cbind(y_hat_report, population$breeding[[activ.base[1]]][[activ.base[2]+2]][,activ.base[3]:activ.base[4], drop=FALSE])
          }
        }

        y_hat_report[y_hat_report==0] <- NA
        if(verbose) cat("Correlation between genetic values and BVE (parent-mean BVE):\n")
        acc <- suppressWarnings(stats::cor(t(y_real_report), t(y_hat_report), use="pairwise.complete.obs"))


        if(sum(is.na(acc))>0){
          acc[is.na(acc)] <- 0
        }
        if(verbose) cat(diag(acc))
        if(verbose) cat("\n")
      }

    }
  } else if(phenotype.bv || bve && sum(sigma.e)==0 && BGLR.bve==FALSE ){
    ## Use Phenotypes as breeding values // You can also use phenotypes in the selection procedure - so potentially removable at some point.
    if(store.bve.data){
      y <- array(0,dim=c(n.animals,population$info$bv.nr))
      y_parent <- array(0,dim=c(n.animals,population$info$bv.nr))
      y_real <- array(0, dim=c(n.animals,population$info$bv.nr)) # schaetzung sigma.g
      y_hat <- array(0, dim=c(n.animals,population$info$bv.nr))
      cindex <- 1
      size <- cumsum(c(0,as.vector(t(population$info$size))))
      for(index in 1:nrow(bve.database)){
        k.database <- bve.database[index,]
        temp1 <- 1:population$info$bv.nr
        if(diff(k.database[3:4])>=0){
          for(kindex in k.database[3]:k.database[4]){
            # t() not needed just to be safe when using multiple individuals at once later
            y[cindex,temp1] <- y_hat[cindex,temp1] <- t(population$breeding[[k.database[[1]]]][[8+k.database[[2]]]][temp1,kindex])
            y_real[cindex,temp1] <- t(population$breeding[[k.database[[1]]]][[6+k.database[[2]]]][temp1,kindex])
            cindex <- cindex +1
          }
        }

      }
    }
    for(index in 1:nrow(bve.insert.database)){
      activ.base <- bve.insert.database[index,]
      if(diff(activ.base[3:4])>=0){
        population$breeding[[activ.base[1]]][[activ.base[2]+2]][,activ.base[3]:activ.base[4]] <- population$breeding[[activ.base[1]]][[activ.base[2]+8]][,activ.base[3]:activ.base[4]]
      }
      if(report.accuracy){
        y_real_report <- NULL
        y_hat_report <- NULL
        for(index in 1:nrow(bve.insert.database)){
          activ.base <- bve.insert.database[index,]
          if(diff(activ.base[3:4])>=0){
            y_real_report <- cbind(y_real_report, population$breeding[[activ.base[1]]][[activ.base[2]+6]][,activ.base[3]:activ.base[4], drop=FALSE])
            y_hat_report <- cbind(y_hat_report, population$breeding[[activ.base[1]]][[activ.base[2]+8]][,activ.base[3]:activ.base[4], drop=FALSE])
          }
        }

        y_hat_report[y_hat_report==0] <- NA
        if(verbose) cat("Correlation between genetic values and BVE (phenotypic BVE):\n")
        acc <- suppressWarnings(stats::cor(t(y_real_report), t(y_hat_report), use="pairwise.complete.obs"))


        if(sum(is.na(acc))>0){
          acc[is.na(acc)] <- 0
        }
        if(verbose) cat(diag(acc))
        if(verbose) cat("\n")
      }

    }
  } else if(bve && breedR.bve==TRUE){
    ## Breeding value estimation using breedR package (as this is just pedigree based all other computational steps are not needed)
    ## Add accuracy report!

    if(verbose) cat("Start pedigree-based BVE.\n")

    y <- get.pheno(population, database=breedR.groups)
    animal_list <- get.individual.loc(population, database = breedR.groups)

    ped1 <- ped <- get.pedigree(population, database=breedR.groups, founder.zero=FALSE)
    y_real <- y_hat <- array(0,dim=c(ncol(y),population$info$bv.nr))
    setna <- 1
    while(sum(setna)>0){
      setna <- !duplicated(c(ped1[,1], ped1[,2]))[-(1:nrow(ped))]
      ped1[setna,2] <- NA
      setna <- !duplicated(c(ped1[,1], ped1[,3]))[-(1:nrow(ped))]
      ped1[setna,3] <- NA
    }
    for(index in 1:nrow(ped1)){
      change <- which(ped1==ped[index,1])
      ped1[change] <- index
    }
    x <- rep(1,length(pheno[1,]))
    for(bven in 1:population$info$bv.nr){
      fit <- breedR::remlf90(fixed = pheno~1,genetic=list('add_animal', pedigree=ped1, id='self'), data=data.frame(pheno=y[bven,], x=x, self=ped1[,1], mum = ped1[,3], dad=ped1[,2]))
      y_hat[bven] <- stats::fitted(fit)
    }
    for(index in 1:nrow(animal_list)){
      population$breeding[[animal_list[index,1]]][[animal_list[index,2]+2]][, animal_list[index,3]] <- y_hat[,index]
    }

    best.selection.ratio.m

  } else if(bve){
    ## Breeding value estimation using a mixed model

    if(verbose) cat("Start genomic BVE.\n")
    ## Derive if individuals are using multiple times in the BVE
    ## Import Phenotypes from all copies of an individual (potentially use individual entry [[21]] for efficiency?)
    loop_elements_list <- derive.loop.elements(population=population, bve.database=bve.database,
                                          bve.class=bve.class, bve.avoid.duplicates=bve.avoid.duplicates,
                                          store.which.adding = TRUE, list.of.copys = TRUE)
    loop_elements <- loop_elements_list[[1]]
    loop_elements_copy <- loop_elements_list[[3]]

    n.animals <- nrow(loop_elements)
    genotyped <- numeric(n.animals)

    stay.loop.elements <- NULL
    n.rep <- 0
    if(length(loop_elements_copy)>0){
      n.rep <- nrow(loop_elements_copy)
    }
    # remove non-genotyped samples in case no pedigree-based estimation // single-step
    if(remove.non.genotyped && singlestep.active==FALSE && computation.A!="kinship"){
      for(index in 1:n.animals){

        k.database <- bve.database[loop_elements[index,3],]
        kindex <- loop_elements[index,2]
        genotyped[index] <- population$breeding[[k.database[[1]]]][[k.database[[2]]]][[kindex]][[16]]
      }


      if(n.rep>0){
        genotyped.copy <- numeric(n.rep)
        for(index in 1:n.rep){
          kindex <- loop_elements_copy[index,2]
          k.database <- bve.database[loop_elements_copy[index,3],]
          if(population$breeding[[k.database[[1]]]][[k.database[[2]]]][[kindex]][[16]]==1){
            genotyped[loop_elements_copy[index,6]] <-  genotyped.copy[index] <- population$breeding[[k.database[[1]]]][[k.database[[2]]]][[kindex]][[16]]

          }
        }
        remove.loop.elements <- which(genotyped.copy==0)

        if(length(remove.loop.elements)>0){
          loop_elements_list[[3]] <- loop_elements_list[[3]][-remove.loop.elements,]
        }
      }
      stay.loop.elements <- which(genotyped==1)
      remove.loop.elements <- which(genotyped==0)
      if(length(remove.loop.elements)>0){
        loop_elements_list[[1]] <- loop_elements_list[[1]][-remove.loop.elements,]
        loop_elements_list[[2]] <- loop_elements_list[[2]][-remove.loop.elements]
      }


      loop_elements <- loop_elements_list[[1]]
      n.animals <- nrow(loop_elements)
      loop_elements_list[[1]][,1] <- 1:n.animals
      genotyped <- numeric(n.animals)

    }
    # sequenceZ is to not process all SNPs at the same time. Especially relevant for large scale datasets!
    if(sequenceZ){
      if(maxZtotal>0){
        maxZ <- floor(maxZtotal / n.animals)
      }
      Zt <- array(0L,dim=c(maxZ,n.animals))
    } else{
      Zt <- array(0L,dim=c(sum(population$info$snp), n.animals))
    }
    y <- y_real <- y_hat <- y_reli <- array(0,dim=c(n.animals,population$info$bv.nr))
    X <- matrix(1, nrow=n.animals,ncol=1)

    grid.position <- numeric(n.animals)
    cindex <- 1
    size <- cumsum(c(0,as.vector(t(population$info$size))))
    y_obs <- matrix(0, nrow=n.animals, ncol=population$info$bv.nr)
    # Import phenotypes / genomic values
    for(index in 1:n.animals){

      k.database <- bve.database[loop_elements[index,3],]
      kindex <- loop_elements[index,2]
      y[index,] <- population$breeding[[k.database[[1]]]][[8+k.database[[2]]]][,kindex]
      y_real[index,] <- population$breeding[[k.database[[1]]]][[6+k.database[[2]]]][,kindex]
      y_obs[index,] <- population$breeding[[k.database[[1]]]][[k.database[[2]]]][[kindex]][[15]]
      grid.position[index] <- kindex + size[sum(k.database[1:2]*c(2,1))-2] # how many individuals are in earlier generations
      genotyped[index] <- population$breeding[[k.database[[1]]]][[k.database[[2]]]][[kindex]][[16]]
      if(!remove.non.genotyped && singlestep.active==FALSE){
        genotyped[index] <- 1
      }

      if(estimate.add.gen.var){
        father <- population$breeding[[k.database[1]]][[k.database[2]]][[kindex]][[7]]
        mother <- population$breeding[[k.database[1]]][[k.database[2]]][[kindex]][[8]]
        if(sum(father[1:3]==c(k.database[1:2], kindex))==3 ||sum(mother[1:3]==c(k.database[1:2], kindex))==3){
          if(verbose) cat("Schaetzung der additiv genetischen Varianz extrem problematisch. Kein Elterntier fuer jedes Tier vorhanden!\n")
        }
        y_parent[index,] <- mean(population$breeding[[father[1]]][[8+father[2]]][,father[3]],population$breeding[[mother[1]]][[8+mother[2]]][,mother[3]])
      }
    }
    if(n.rep>0){
      for(index in 1:n.rep){
        kindex <- loop_elements_copy[index,2]
        k.database <- bve.database[loop_elements_copy[index,3],]
        if(length(stay.loop.elements)>0){
          non_copy <- which(stay.loop.elements==loop_elements_copy[index,6])
        } else{
          non_copy <- loop_elements_copy[index,6]
        }

        if(length(non_copy)==1 && prod(population$breeding[[k.database[[1]]]][[k.database[[2]]]][[kindex]][[15]]>= y_obs[non_copy,])==1){
          y[non_copy,] <- population$breeding[[k.database[[1]]]][[8+k.database[[2]]]][,kindex]
        }
      }
    }

    # Check if some copy is genotyped
    if(nrow(loop_elements_list[[3]])>0){
      for(index in 1:nrow(loop_elements_list[[3]])){
        kindex <- loop_elements_list[[3]][index,2]
        k.database <- bve.database[loop_elements_list[[3]][index,3],]
        if(length(stay.loop.elements)>0){
          non_copy <- which(stay.loop.elements==loop_elements_copy[index,6])
        } else{
          non_copy <- loop_elements_copy[index,6]
        }
        genotyped[non_copy] <- population$breeding[[k.database[[1]]]][[k.database[[2]]]][[kindex]][[16]]
      }
    }
    genotype.included <- which(genotyped==1)

    batch <- ceiling(nrow(loop_elements)/ncore)
    batche <- list()
    for(index in 1:ncore){
      batche[[index]] <- (batch*(index-1)+1):min(batch*index, nrow(loop_elements))
    }


    # Import Z
    if(sequenceZ==FALSE){
      if(miraculix){

        if(store.comp.times.bve){
          before <- as.numeric(Sys.time())
        }

        if (requireNamespace("miraculix", quietly = TRUE)) {
          Z.code <- miraculix::computeSNPS(population, loop_elements[,4], loop_elements[,5], loop_elements[,2], what="geno", output_compressed=TRUE)
          if(computation.A!="vanRaden"){
            Zt <- miraculix::computeSNPS(population, loop_elements[,4], loop_elements[,5], loop_elements[,2], what="geno", output_compressed=FALSE)
          }
        }
        if(store.comp.times.bve){
          after <- as.numeric(Sys.time())
          zcalc <- zcalc + after - before
        }

      } else if(ncore>1){
        if(store.comp.times.bve){
          before <- as.numeric(Sys.time())
        }

        if(backend=="doParallel"){
          doParallel::registerDoParallel(cores=ncore)
        } else if(backend=="doMPI"){
          if (requireNamespace("doMPI", quietly = TRUE)) {
            cl <- doMPI::startMPIcluster(count=ncore)
            doMPI::registerDoMPI(cl)
          } else{
            stop("Usage of doMPI without being installed!")
          }
        } else{
          if(verbose) cat("No valid backend specified.\n")
        }

        if (requireNamespace("foreach", quietly = TRUE)) {
        } else{
          stop("Usage of foreach without being installed!")
        }
        Zt <- foreach::foreach(indexb=1:ncore, .combine = "cbind", .multicombine = TRUE,.maxcombine = 1000,
                     .packages="MoBPS") %dopar% {
          Ztpar <- array(0,dim=c(sum(population$info$snp), length(batche[[indexb]])))
          sub <- min(batche[[indexb]]) -1
          for(index in batche[[indexb]]){
            k.database <- bve.database[loop_elements[index,3],]
            cindex <- loop_elements[index,1] - sub
            kindex <- loop_elements[index,2]
            Ztpar[,cindex] <- base::as.integer(colSums(compute.snps(population, k.database[1],k.database[2],kindex, import.position.calculation=import.position.calculation,decodeOriginsU=decodeOriginsU, bit.storing=bit.storing, nbits=nbits, output_compressed=FALSE)))
          }
          if(Z.integer){
            storage.mode(Zpar) <- "integer"
          }
          Ztpar
        }
        if(backend=="doParallel"){
          doParallel::stopImplicitCluster()
        } else if(backend=="doMPI"){
          if (requireNamespace("doMPI", quietly = TRUE)) {
            doMPI::closeCluster(cl)
          } else{
            stop("Usage of doMPI without being installed!")
          }


        }
        if(store.comp.times.bve){
          after <- as.numeric(Sys.time())
          zcalc <- zcalc + after - before
        }
      } else{
        if(store.comp.times.bve){
          before <- as.numeric(Sys.time())
        }

        for(index in 1:n.animals){
          k.database <- bve.database[loop_elements[index,3],]
          cindex <- loop_elements[index,1]
          kindex <- loop_elements[index,2]
          Zt[,cindex] <- base::as.integer(colSums(compute.snps(population, k.database[1],k.database[2],kindex, import.position.calculation=import.position.calculation,decodeOriginsU=decodeOriginsU, bit.storing=bit.storing, nbits=nbits, output_compressed=FALSE)))
        }
        if(store.comp.times.bve){
          after <- as.numeric(Sys.time())
          zcalc <- zcalc + after - before
        }
      }

    }


    if(remove.effect.position && sequenceZ==FALSE){
      if(miraculix && exists("Z.code")){
        if (requireNamespace("miraculix", quietly = TRUE)) {
          Z.code <- miraculix::zeroNthGeno(Z.code, population$info$effect.p)
        }
      } else{
        Zt <- Zt[-population$info$effect.p,]
      }

    }
    if(remove.effect.position=="only_effect" && sequenceZ==FALSE){
      if(miraculix && exists("Z.code")){
        if (requireNamespace("miraculix", quietly = TRUE)) {
          Z.code <- miraculix::zeroNthGeno(Z.code, (1:sum(population$info$snp))[-population$info$effect.p])
        }
      } else{
        Zt <- Zt[population$info$effect.p,]
      }
    }
    if(bve.0isNA){
      y[y==0] <- NA
    }

    if(store.comp.times.bve){
      comp.times.bve[2] <- as.numeric(Sys.time())
    }

    # Derive relationship matrix sequenceZ and single-step only for vanRaden based genomic relationship
    if(computation.A=="kinship"){
      z_ped <- z_ped - as.numeric(Sys.time())
      A <- kinship.exp(population, database=bve.database, depth.pedigree=depth.pedigree)[loop_elements_list[[2]],loop_elements_list[[2]]] * 2
      z_ped <- z_ped + as.numeric(Sys.time())
    } else if(computation.A=="vanRaden"){
      if(sequenceZ){
        total_n <- sum(population$info$snp)
        p_i <- numeric(total_n)
        first <- 1
        last <- min(maxZ, total_n)
        A <- matrix(0, n.animals, n.animals)
        for(index3 in 1:ceiling(total_n/maxZ)){
          if(miraculix){
            if(store.comp.times.bve){
              before <- as.numeric(Sys.time())
            }

            if (requireNamespace("miraculix", quietly = TRUE)) {
              Z.code <- miraculix::computeSNPS(population, loop_elements[,4], loop_elements[,5], loop_elements[,2], from_p=first,
                                  to_p=last, what="geno", output_compressed=TRUE)
            }
            if(store.comp.times.bve){
              after <- as.numeric(Sys.time())
              zcalc <- zcalc + after - before
            }


          } else if(ncore>1){
            if(store.comp.times.bve){
              before <- as.numeric(Sys.time())
            }

            if(backend=="doParallel"){
              doParallel::registerDoParallel(cores=ncore)
            } else if(backend=="doMPI"){
              if (requireNamespace("doMPI", quietly = TRUE)) {
                cl <- doMPI::startMPIcluster(count=ncore)
                doMPI::registerDoMPI(cl)
              } else{
                stop("Usage of doMPI without being installed!")
              }
            } else{
              if(verbose) cat("No valid backend specified.\n")
            }
            if (requireNamespace("foreach", quietly = TRUE)) {
            } else{
              stop("Usage of foreach without being installed!")
            }
            Zt <- foreach::foreach(indexb=1:ncore, .combine = "rbind", .multicombine = TRUE,.maxcombine = 1000,
                         .packages="MoBPS") %dopar% {
              Ztpar <- array(0,dim=c(last-first+1, length(batche[[indexb]])))
              sub <- min(batche[[indexb]]) -1
              for(index in batche[[indexb]]){
                k.database <- bve.database[loop_elements[index,3],]
                cindex <- loop_elements[index,1] - sub
                kindex <- loop_elements[index,2]
                Zpar[,cindex] <- base::as.integer(colSums(compute.snps(population, k.database[1],k.database[2],kindex, import.position.calculation=import.position.calculation, from_p=first, to_p=last, decodeOriginsU=decodeOriginsU, bit.storing=bit.storing, nbits=nbits, output_compressed=FALSE)))
              }
              if(Z.integer){
                storage.mode(Ztpar) <- "integer"
              }
              Ztpar
            }
            if(backend=="doParallel"){
              doParallel::stopImplicitCluster()
            } else if(backend=="doMPI"){
              if (requireNamespace("doMPI", quietly = TRUE)) {
                doMPI::closeCluster(cl)
              } else{
                stop("Usage of doMPI without being installed!")
              }

            }
            if(store.comp.times.bve){
              after <- as.numeric(Sys.time())
              zcalc <- zcalc + after - before
            }
          } else{
            if(store.comp.times.bve){
              before <- as.numeric(Sys.time())
            }

            for(index in 1:n.animals){
              k.database <- bve.database[loop_elements[index,3],]
              cindex <- loop_elements[index,1]
              kindex <- loop_elements[index,2]
              Zt[1:(last-first+1), cindex] <- base::as.integer(colSums(compute.snps(population, k.database[1],k.database[2],kindex, import.position.calculation=import.position.calculation, from_p=first, to_p=last,decodeOriginsU=decodeOriginsU, bit.storing=bit.storing, nbits=nbits, output_compressed=FALSE)))
            }
            if(store.comp.times.bve){
              after <- as.numeric(Sys.time())
              zcalc <- zcalc + after - before
            }
          }

          activ_effect <- population$info$effect.p - first + 1
          activ_effect <- activ_effect[activ_effect>0]
          activ_effect <- activ_effect[activ_effect<=last]
          if(remove.effect.position==TRUE){
            if(miraculix && exists("Z.code")){
              if (requireNamespace("miraculix", quietly = TRUE)) {
                Z.code <- miraculix::zeroNthGeno(Z.code, activ_effect)
              }
            } else{
              Zt <- Zt[-activ_effect,]
            }

          } else if(remove.effect.position=="only_effect"){
            if(miraculix && exists("Z.code")){
              if (requireNamespace("miraculix", quietly = TRUE)) {
                Z.code <- miraculix::zeroNthGeno(Z.code, (1:(last-first+1))[-activ_effect])
              }
            } else{
              Zt <- Zt[activ_effect,]
            }
          }

          if(miraculix){
            if (requireNamespace("miraculix", quietly = TRUE)) {
              p_i[first:last] <- miraculix::allele_freq(Z.code)
              A <- A + miraculix::relationshipMatrix(Z.code, centered=TRUE, normalized=FALSE)
            }
          } else if(miraculix.mult){
            storage.mode(Zt) <- "integer"
            p_i[first:last] <- rowSums(Zt[1:(last-first+1),])/ncol(Zt)/2
            if (requireNamespace("miraculix", quietly = TRUE)) {
              Zt_miraculix <- miraculix::genomicmatrix(Zt[1:(last-first+1),])
              A <- A + miraculix::relationshipMatrix(Zt_miraculix, centered=TRUE, normalized=FALSE)
            }
          } else{
            p_i[first:last] <- rowSums(Zt[1:(last-first+1),])/ncol(Zt)/2
            Ztm <- Zt[1:(last-first+1),] - p_i[first:last] * 2
            A <- A + crossprod(Ztm[1:(last-first+1),])
          }

          first <- first + maxZ
          last <- min(maxZ*(index3+1), total_n)
        }
        A <- A / (2 * sum(p_i*(1-p_i)))


      } else{

        if(singlestep.active){
          n.geno <- length(genotype.included)
          n.ped <- n.animals - n.geno
          if(n.ped==0){
            if(verbose) cat("No non genotyped individuals included. Automatically switch to regular GBLUP instead of ssGBLUP\n")
            singlestep.active <- FALSE
          }
          if(n.geno==0){
            z_ped <- z_ped - as.numeric(Sys.time())
            if(verbose) cat("No genotyped individuals included. Automatically switch to pedigree BLUP instead of ssGBLUP\n")
            singlestep.active <- FALSE
            computation.A <- "kinship"
            A <- kinship.exp(population, database=bve.database, depth.pedigree=depth.pedigree)[loop_elements_list[[2]],loop_elements_list[[2]]] * 2
            z_ped <- z_ped + as.numeric(Sys.time())
          }
        }
        if(singlestep.active){
          if(verbose) cat("Start derive Single-step relationship matrix \n")
          if(verbose) cat(paste0("Construct pedigree matrix for ", length(loop_elements_list[[2]]), " individuals.\n"))
          z_ped <- z_ped - as.numeric(Sys.time())
          A_pedigree <-  kinship.exp(population, database=bve.database, depth.pedigree=depth.pedigree)[loop_elements_list[[2]],loop_elements_list[[2]]] * 2
          z_ped <- z_ped + as.numeric(Sys.time())
          if(verbose) cat(paste0("Derived pedigree matrix in  ", round(z_ped, digits=2), " seconds.\n"))
          if(miraculix){
            if (requireNamespace("miraculix", quietly = TRUE)) {
              Z.code.small <- miraculix::computeSNPS(population, loop_elements[genotype.included,4], loop_elements[genotype.included,5], loop_elements[genotype.included,2], what="geno", output_compressed=TRUE)
              p_i <- miraculix::allele_freq(Z.code.small)
              A_geno <- miraculix::relationshipMatrix(Z.code.small, centered=TRUE, normalized=TRUE)
            }
          } else if(miraculix.mult){
            if (requireNamespace("miraculix", quietly = TRUE)) {
              p_i <- rowSums(Zt[,genotype.included])/ncol(Zt[,genotype.included])/2
              Zt_miraculix <- miraculix::genomicmatrix(Zt[,genotype.included])
              A_geno <- miraculix::relationshipMatrix(Zt_miraculix, centered=TRUE, normalized=TRUE)
            }
          } else{
            p_i <- rowSums(Zt[,genotype.included])/ncol(Zt[,genotype.included])/2
            Ztm <- Zt[,genotype.included] - p_i * 2
            A_geno <- crossprod(Ztm)/ (2 * sum(p_i*(1-p_i)))
          }

          z_h <- z_h - as.numeric(Sys.time())

          reorder <- c((1:n.animals)[-genotype.included], genotype.included)
          if(verbose) cat("Start deriving of H matrix for", length(genotype.included), "genotyped and", nrow(A_pedigree)-length(genotype.included), "non-genotyped individuals.\n")
          H_order <- H <- ssGBLUP(A11= A_pedigree[-genotype.included, -genotype.included],
                                  A12 = A_pedigree[-genotype.included, genotype.included],
                                  A22 = A_pedigree[genotype.included, genotype.included], G = A_geno)

          H[genotype.included, genotype.included] <- H_order[(ncol(H_order)-length(genotype.included)+1):ncol(H_order), (ncol(H_order)-length(genotype.included)+1):ncol(H_order)]
          H[-genotype.included, -genotype.included] <- H_order[1:(ncol(H_order)-length(genotype.included)), 1:(ncol(H_order)-length(genotype.included))]
          H[-genotype.included, genotype.included] <- H_order[1:(ncol(H_order)-length(genotype.included)), (ncol(H_order)-length(genotype.included)+1):ncol(H_order)]
          H[genotype.included, -genotype.included] <- H_order[(ncol(H_order)-length(genotype.included)+1):ncol(H_order), 1:(ncol(H_order)-length(genotype.included))]

          rm(A_geno)
          rm(H_order)
          A <- H
          rm(H)
          rm(A_pedigree)

          z_h <- z_h + as.numeric(Sys.time())
          if(verbose) cat(paste0("Derived H matrix in  ", round(z_h, digits=2), " seconds.\n"))

        } else if(computation.A=="vanRaden"){
          if(miraculix){
            if (requireNamespace("miraculix", quietly = TRUE)) {
              p_i <- miraculix::allele_freq(Z.code) # Noch nicht implementiert?
              A <- miraculix::relationshipMatrix(Z.code, centered=TRUE, normalized=TRUE)
            }
          } else if(miraculix.mult){
            if (requireNamespace("miraculix", quietly = TRUE)) {
              p_i <- rowSums(Zt)/ncol(Zt)/2
              Zt_miraculix <- miraculix::genomicmatrix(Zt)
              A <- miraculix::relationshipMatrix(Zt_miraculix, centered=TRUE, normalized=TRUE)
            }
          } else{
            p_i <- rowSums(Zt)/ncol(Zt)/2
            Ztm <- Zt - p_i * 2
            A <- crossprod(Ztm)/ (2 * sum(p_i*(1-p_i)))
          }
        }


      }

      if(store.comp.times.bve){
        comp.times.bve[3] <- as.numeric(Sys.time())
      }

    } else if(computation.A=="CM"){
      #CM SCHAETZER
      Ztm <- rbind(Zt==0, Zt==1, Zt==2)
      A <- crossprod(Ztm) / nrow(Zt)
    } else if(computation.A=="CE"){

      Ztm <- rbind(Zt==0, Zt==1, Zt==2)
      A <- crossprod(Ztm)
      A <- (A^2 - 0.5*A)/(nrow(Zt)^2)
    } else if(computation.A=="CE2"){
      A2 <- crossprod(Zt)
      A <- 0.5 * A2 * A2 + 0.5 * crossprod(Zt*Zt)
      A <- A/mean(diag(A))
    } else if(computation.A=="CE3"){
      A2 <- crossprod(Zt)
      A <- 0.5 * A2 * A2 - 0.5 * crossprod(Zt*Zt)
      A <- A/mean(diag(A))
    } else if(computation.A=="non_stand"){
      A <- crossprod(Zt) / nrow(Zt)
    } else if(computation.A=="vanRaden2"){
      p_i <- rowSums(Zt)/ncol(Zt)/2
      Ztm <- Zt - p_i * 2
      A <- crossprod(Ztm)/ (2 * sum(p_i*(1-p_i)))
    }

    sigma.a.hat <- numeric(length(sigma.g))
    sigma.e.hat <- sigma.e

    beta_hat <-  colMeans(y, na.rm=TRUE) # Rest faellt weg! (X' R^-1 X)^-1 X' R^-1 y
    if(sum(is.na(beta_hat))>0){
      if(verbose) cat("No phenotypes available for traits:", population$info$trait.name[which(is.na(beta_hat))],"\n")
      if(verbose) cat("Set all breeding value estimates for these trait(s) to 0. \n")
      beta_hat[is.na(beta_hat)] <- 0
    }

    prev_rest_take <- NULL

    n.rep <- 0
    if(length(bve.database)==length(bve.insert.database) && prod(bve.database==bve.insert.database) && nrow(loop_elements_list[[3]])==0){
      bve.insert <- rep(TRUE, n.animals)
      bve.insert.copy <- NULL
    } else{
      bve.insert <- rep(FALSE, n.animals)
      before <- 0
      for(index in 1:nrow(bve.insert.database)){
        add_insert <- loop_elements[,2] <= bve.insert.database[index,4] & loop_elements[,2] >= bve.insert.database[index,3] & loop_elements[,4] == bve.insert.database[index,1] & loop_elements[,5] == bve.insert.database[index,2]
        bve.insert[add_insert] <- TRUE
      }

      loop_elements_copy <- loop_elements_list[[3]]
      n.rep <- nrow(loop_elements_copy)
      bve.insert.copy <- rep(FALSE, n.rep)
      if(n.rep>0){
        for(index in 1:nrow(bve.insert.database)){
          add_insert <- loop_elements_copy[,2] <= bve.insert.database[index,4] & loop_elements_copy[,2] >= bve.insert.database[index,3] & loop_elements_copy[,4] == bve.insert.database[index,1] & loop_elements_copy[,5] == bve.insert.database[index,2]
          bve.insert.copy[add_insert] <- TRUE
        }
      }

    }
    # Breeding value estimation (Solving of the mixed-model) - all single trait models execpt multi-sommer
    for(bven in 1:population$info$bv.nr){
      if(forecast.sigma.g){
        sigma.g[bven] <- stats::var(y_real[,bven])
      }
      if(estimate.pheno.var){
        sigma.e.hat[bven] <- max(0, stats::var(y[,bven], na.rm=TRUE) - sigma.g[bven])
      }
      sigma.a.hat[bven] <- sigma.g[bven]
      if(estimate.add.gen.var){
        sigma.a.hat[bven] <- max(min(stats::lm(y[,bven]~y_parent[,bven])$coefficients[2],1),0.001) * sigma.e.hat[bven]
      }
      if(BGLR.bve){

        if(miraculix){
          Zt <- as.matrix(Z.code)
        }
        Zt <- t(scale(t(Zt), center=TRUE, scale=FALSE))
        fixed <- which(is.na(Zt[,1]))
        if(length(fixed)>0){
          Zt <- Zt[-fixed,]
        }
        if(BGLR.model=="RKHS"){
          ETA <- list(list(K=A, model='RKHS'))
        } else if(BGLR.model=="BayesA"){
          ETA <- list(list(X=t(Zt), model='BayesA'))
        } else if(BGLR.model=="BayesB"){
          ETA <- list(list(X=t(Zt), model='BayesB'))
        } else if(BGLR.model=="BayesC"){
          ETA <- list(list(X=t(Zt), model='BayesC'))
        } else if(BGLR.model=="BL"){
          ETA <- list(list(X=t(Zt), model='BL'))
        } else if(BGLR.model=="BRR"){
          ETA <- list(list(X=t(Zt), model='BRR'))
        }

        if(min(y[!is.na(y[,bven]),bven])==max(y[!is.na(y[,bven]),bven])){
          y_hat[,bven] <- rep(max(y[!is.na(y[,bven]),bven]), length(y[,bven]))
        } else{
          if(BGLR.save.random){
            BGLR.save <- paste0(BGLR.save, sample(1:10000,1))
          }
          if(requireNamespace("BGLR", quietly = TRUE)){
            fm <- BGLR::BGLR(y=y[,bven], ETA=ETA, nIter=BGLR.iteration, burnIn=BGLR.burnin, saveAt=BGLR.save, verbose=BGLR.print)
          } else{
            stop("Use of BGLR without being installed!")
          }
          y_hat[,bven] <- fm$yHat
        }
      } else if(rrblup.bve){

        check <- sum(is.na(y[,bven]))
        if(verbose) cat(paste0(length(y[,bven]) - check, " phenotyped individuals in BVE.\n"))
        if(check == length(y[,bven])){
          if(verbose) cat(paste0("No phenotyped individuals for trait ", population$info$trait.name[bven], "\n"))
          if(verbose) cat(paste0("Skip this BVE.\n"))
          next
        }

        t1 <- as.numeric(Sys.time())
        test <- rrBLUP::mixed.solve(y[,bven], K = A, method="REML", bounds = c(1e-9,1e9))
        t2 <- as.numeric(Sys.time())

        if(verbose) cat(paste0(round(t2-t1, digits=2), " seconds for BVE.\n"))
        y_hat[,bven] <- as.numeric(test$beta) + test$u

      } else if(emmreml.bve){
        check <- sum(is.na(y[,bven]))
        if(verbose) cat(paste0(length(y[,bven]) - check, " phenotyped individuals in BVE.\n"))
        if(check == length(y[,bven])){
          if(verbose) cat(paste0("No phenotyped individuals for trait ", population$info$trait.name[bven], "\n"))
          if(verbose) cat(paste0("Skip this BVE."))
          next
        }
        if(check>0){
          if(verbose) cat(paste0("Breeding value estimation with ", check, " NA phenotypes! EMMREML does not support this!\n"))
          if(verbose) cat(paste0("No estimation is performed to NA individuals. \n"))
          take <- which(!is.na(y[,bven]))
        } else{
          take <- 1:length(y[,bven])
        }
        n <- length(y[take,bven])
        p <- nrow(Zt)
        stopifnot(n == ncol(Zt[,take]))

        if(requireNamespace("EMMREML", quietly = TRUE)){
          fm <- EMMREML::emmreml(
            y[take,bven],
            matrix(1,nrow=n),
            diag(n),
            A[take,take])
        } else{
          stop("Usage of EMMREML without being installed!")
        }


        y_hat[take,bven] <- as.numeric(fm$uhat) + as.numeric(fm$betahat)
        if(estimate.u){
          u_hat <- cbind(u_hat, alpha_to_beta(drop(fm$uhat),A[take,take],t(Zt[,take])))
        }

      } else if(sommer.bve){

        check <- sum(is.na(y[,bven]))
        if(verbose) cat(paste0(length(y[,bven]) - check, " phenotyped individuals in BVE.\n"))
        if(check == length(y[,bven])){
          if(verbose) cat(paste0("No phenotyped individuals for trait ", population$info$trait.name[bven], "\n"))
          if(verbose) cat(paste0("Skip this BVE.\n"))
          next
        }

        traitnames <- (paste0("name", 1:ncol(y)))
        traitnames[bven] <- "name"
        traitnames <- as.factor(traitnames)
        colnames(y) <- traitnames
        id <- as.factor(paste0("P", 1:nrow(y)))
        y_som <- data.frame(y, id)
        rownames(y_som) <- id
        colnames(A) <- rownames(A) <- id

        test <- sommer::mmer(name ~1, random=~sommer::vs(id, Gu=A), rcov = ~units, data=y_som)

        y_hat[sort(as.character(id), index.return=TRUE)$ix,bven] <- test$U[[1]][[1]] + as.numeric(test$Beta[3])

      } else if(sommer.multi.bve){

        check <- sum(is.na(y))

        if(verbose) cat(paste0(length(y[,bven]) - check, " phenotyped individuals in BVE.\n"))
        if(check == length(y)){
          if(verbose) cat(paste0("No phenotyped individuals for multi-trait mixed model\n"))
          if(verbose) cat(paste0("Skip this BVE.\n"))
          next
        }


        if(bven==population$info$bv.nr){

          traitnames <- paste0("name", 1:ncol(y))
          colnames(y) <- as.factor(traitnames)
          id <- as.factor(paste0("P", 1:nrow(y)))
          y_som <- data.frame(y, id)
          rownames(y_som) <- id
          colnames(A) <- rownames(A) <- id
          text <- "cbind("
          for(index in 1:length(traitnames)){
            if(index==length(traitnames)){
              text <- paste0(text, traitnames[index], ")")
            } else{
              text <- paste0(text, traitnames[index], ",")
            }

          }
          text <- paste0("sommer::mmer(",text,"~1, random=~sommer::vs(id, Gu=A, Gtc=sommer::unsm(bven)), rcov = ~sommer::vs(units, Gtc=diag(bven)), data=y_som)")
          test <- eval(parse(text=text))
          for(bven1 in 1:bven){
            y_hat[sort(as.character(id), index.return=TRUE)$ix,bven1] <- test$U[[1]][[bven1]]
          }


          if(estimate.u){
            if(verbose) cat("U estimation not available in sommer")
          }
        }
      } else if(sigma.e[bven]>0 || sum(is.na(y[,bven]>0))){
        # sigma.a.hat / sigma.e.hat are modified to avoid h=0, h=1 cases (numeric instability)
        if(sigma.a.hat[bven]==0){
          if(sigma.e.hat[bven]>0){
            sigma.a.hat[bven] <- sigma.e.hat[bven] * 0.001
          } else{
            sigma.e.hat[bven] <- 1
            sigma.a.hat[bven] <- 1000
          }
        }
        if(sigma.e.hat[bven]==0){
          sigma.e.hat[bven] <- sigma.a.hat[bven] * 0.001
        }
        bve.direct.est.now <- bve.direct.est
        check <- sum(is.na(y[,bven]))

        u_hat_possible <- TRUE
        # This will be more complicated in case other fixed-effects are added to the model !
        multi <- y[,bven] - X %*% beta_hat[bven]

        rrblup.required <- FALSE

        if(verbose) cat(paste0(length(y[,bven]) - check, " phenotyped individuals in BVE.\n"))
        if(check == length(y[,bven])){
          if(verbose) cat(paste0("No phenotyped individuals for trait ", population$info$trait.name[bven], "\n"))
          if(verbose) cat(paste0("Skip this BVE.\n"))
          next
        }

        # take Individuals used to trait the mixed model on
        # take2 individuals for which to enter a breeding value
        # take3 individuals to used for estimation of allele effects (rrBLUP)
        if(check>0){
          if(sum(genotyped==1 & is.na(y[,bven]))==sum(genotyped==1)){
            if(bve.direct.est.now==FALSE){
              if(verbose) cat("No genotyped and phenotyped individuals. Application of rrBLUP not possible!\n")
              if(verbose) cat("Assume non phenotyped individuals to have average phenotype.\n")
              if(bve.0isNA && sum(is.na(multi))>0){
                multi[is.na(multi)] <- 0
              }
              take <- take2 <- 1:length(y[,bven])
            } else{
              take <- which(!is.na(y[,bven]))
              take2 <- 1:length(y[,bven])
            }


          } else if(sum(genotyped==1 & is.na(y[,bven]))>0){
            if(verbose) cat("Some genotyped individuals without phenotype.\n")
            take <- which(!is.na(y[,bven])) # individuals for GBLUP
            if(!bve.direct.est.now){
              rrblup.required <- TRUE
              take2 <- which(!is.na(y[,bven]) & genotyped==1) # individuals for rrBLUP
            } else if(estimate.u){
              take2 <- which(!is.na(y[,bven]) & genotyped==1) # individuals for rrBLUP
            } else{
              take2 <- 1:length(y[,bven])
            }


            bve.insert.full <- bve.insert
            if(length(bve.insert.copy)>0){

              bve.insert.full[loop_elements_copy[bve.insert.copy,6]] <- TRUE
            }
            take3 <- which(is.na(y[,bven]) & genotyped==1 & bve.insert.full) # individuals to estimate via rrBLUP
            if(length(take3)>0 && !bve.direct.est.now){
              if(verbose) cat("Use rrBLUP to estimate breeding value for those individuals!\n")
            }
          } else{
            take <- take2 <- which(!is.na(y[,bven]))
            if(bve.direct.est.now){
              take2 <- 1:length(y[,bven])
            }
          }
        } else{
          take <- take2 <- 1:length(y[,bven])
        }

        if(length(take)==length(take2)){
          bve.direct.est.now <- FALSE
        }


        if(store.comp.times.bve){
          before <- as.numeric(Sys.time())
        }
        # Solve mixed model which assumed known heritability
        # This is not 100% accurate but massively reduces computing time and should be ok for large scale breeding programs
        t1 <- as.numeric(Sys.time())

        if(estimate.u || rrblup.required){
          if(calculate.reliability){
            if(verbose) cat("Reliabilities are currently only calculated for phenotyped individuals. Extension according to vanRaden 2008 planned.\n")
            if(verbose) cat("Use of Z2 Z instead of ZZ (C instead of G). \n")
            GR1 <- chol2inv(chol(add.diag(A[take,take],sigma.e.hat[bven] / sigma.a.hat[bven])))
            Rest_term <- GR1 %*% multi[take]
            y_hat[take,bven] <- A[take,take] %*% Rest_term  + beta_hat[bven]
            y_reli[take,bven] <- diag( A[take,take] %*% GR1 %*% A[take,take])
          } else if(miraculix && miraculix.chol && !bve.direct.est.now){
            if (requireNamespace("miraculix", quietly = TRUE)) {
              temp1 <- miraculix::solveRelMat(A[take,take], sigma.e.hat[bven] / sigma.a.hat[bven], multi[take], beta_hat[bven])
            }
            Rest_term <- temp1[[1]]
            y_hat[take,bven] <- temp1[[2]]
          } else{
            Rest_term <- (chol2inv(chol(add.diag(A[take,take], sigma.e.hat[bven] / sigma.a.hat[bven]))) %*% multi[take])
            y_hat[take,bven] <- A[take,take] %*% Rest_term  + beta_hat[bven]
          }
        } else{
          if(calculate.reliability){
            GR1 <- chol2inv(chol(add.diag(A[take,take], sigma.e.hat[bven] / sigma.a.hat[bven])))
            if(bve.direct.est.now){
              y_hat[take2,bven] <- A[take2,take] %*% (GR1 %*% multi[take])  + beta_hat[bven]
              y_reli[take1,bven] <- diag( A[take,take] %*% GR1 %*% A[take,take])
              y_reli[take2,bven] <- diag( A[,take2] %*% GR1 %*% A[take2,])[which(duplicated(c(take,take2))[-(1:length(take))])]
            } else{
              y_hat[take,bven] <- A[take,take] %*% (GR1 %*% multi[take])  + beta_hat[bven]
              y_reli[take,bven] <- diag( A[take,take] %*% GR1 %*% A[take,take])
            }

          } else if(miraculix && miraculix.chol && !bve.direct.est.now){
            if (requireNamespace("miraculix", quietly = TRUE)) {
              y_hat[take,bven] <- miraculix::solveRelMat(A[take,take], sigma.e.hat[bven] / sigma.a.hat[bven], multi[take],beta_hat[bven])[[2]]
            }
          } else{
            if(bve.direct.est.now){
              y_hat[take2,bven] <- A[take2,take] %*% (chol2inv(chol(add.diag(A[take,take], sigma.e.hat[bven] / sigma.a.hat[bven]))) %*% multi[take]) + beta_hat[bven]
            } else{
              y_hat[take,bven] <- A[take,take] %*% (chol2inv(chol(add.diag(A[take,take],sigma.e.hat[bven] / sigma.a.hat[bven]))) %*% multi[take]) + beta_hat[bven]
            }
          }
        }

        t2 <- as.numeric(Sys.time())
        if(verbose) cat(paste0(round(t2-t1, digits=2), " seconds for BVE.\n"))

        if(store.comp.times.bve){
          after <- as.numeric(Sys.time())
          z_chol <- after - before
        }

        # Estimation of marker effects via rrBLUP

        if(estimate.u || rrblup.required){

          while(bven>1 && (length(u_hat)==0 || ncol(u_hat)<(bven-1))){
            u_hat <- cbind(u_hat,rep(0, sum(population$info$snp)))
          }
          rest_take <- which(duplicated(c(take,take2))[-(1:length(take))])

          if(sequenceZ){
            total_n <- sum(population$info$snp)
            u_hat_new <- numeric(total_n)
            first <- 1
            last <- min(maxZ, total_n)
            for(index3 in 1:ceiling(total_n/maxZ)){
              if(miraculix){
                if(store.comp.times.bve){
                  before <- as.numeric(Sys.time())
                }
                if (requireNamespace("miraculix", quietly = TRUE)) {
                  Z.code2 <- miraculix::computeSNPS(population, loop_elements[take2,4], loop_elements[take2,5], loop_elements[take2,2],
                                      from_p=first, to_p=last, what="geno", output_compressed=TRUE
                                      )
                }

                if(store.comp.times.bve){
                  after <- as.numeric(Sys.time())
                  zcalc <- zcalc + after - before
                }
              } else if(ncore>1){
                if(store.comp.times.bve){
                  before <- as.numeric(Sys.time())
                }

                if(backend=="doParallel"){
                  doParallel::registerDoParallel(cores=ncore)
                } else if(backend=="doMPI"){
                  if (requireNamespace("doMPI", quietly = TRUE)) {
                    cl <- doMPI::startMPIcluster(count=ncore)
                    doMPI::registerDoMPI(cl)
                  } else{
                    stop("Usage of doMPI without being installed!")
                  }
                } else{
                  if(verbose) cat("No valid backend specified.\n")
                }
                if (requireNamespace("foreach", quietly = TRUE)) {
                } else{
                  stop("Usage of foreach without being installed!")
                }
                Zt <- foreach::foreach(indexb=1:ncore, .combine = "cbind", .multicombine = TRUE,.maxcombine = 1000,
                           .packages="MoBPS") %dopar% {
                  Ztpar <- array(0,dim=c(sum(population$info$snp), length(batche[[indexb]])))
                  sub <- min(batche[[indexb]]) -1
                  for(index in batche[[indexb]]){
                    k.database <- bve.database[loop_elements[index,3],]
                    cindex <- loop_elements[index,1] - sub
                    kindex <- loop_elements[index,2]
                    Ztpar[,cindex] <- base::as.integer(colSums(compute.snps(population, k.database[1],k.database[2],kindex, import.position.calculation=import.position.calculation, from_p=first, to_p=last, decodeOriginsU=decodeOriginsU, bit.storing=bit.storing, nbits=nbits, output_compressed=FALSE)))
                  }
                  if(Z.integer){
                    storage.mode(Ztpar) <- "integer"
                  }
                  Ztpar
                }
                if(backend=="doParallel"){
                  doParallel::stopImplicitCluster()
                } else if(backend=="doMPI"){
                  if (requireNamespace("doMPI", quietly = TRUE)) {
                    doMPI::closeCluster(cl)
                  } else{
                    stop("Usage of doMPI without being installed!")
                  }

                }
                if(store.comp.times.bve){
                  after <- as.numeric(Sys.time())
                  zcalc <- zcalc + after - before
                }
              } else{
                if(store.comp.times.bve){
                  before <- as.numeric(Sys.time())
                }
                for(index in 1:n.animals){
                  k.database <- bve.database[loop_elements[index,3],]
                  cindex <- loop_elements[index,1]
                  kindex <- loop_elements[index,2]
                  Zt[1:(last-first+1), cindex] <- base::as.integer(colSums(compute.snps(population, k.database[1],k.database[2],kindex, import.position.calculation=import.position.calculation, from_p=first, to_p=last,decodeOriginsU=decodeOriginsU, bit.storing=bit.storing, nbits=nbits, output_compressed=FALSE)))
                }
                if(store.comp.times.bve){
                  after <- as.numeric(Sys.time())
                  zcalc <- zcalc + after - before
                }
              }

              if(store.comp.times.bve){
                before <- as.numeric(Sys.time())
              }

              if(!(length(rest_take)==length(prev_rest_take) && prod(rest_take==prev_rest_take)==1) && !fast.uhat){
                A1 <- MASS::ginv(A[take2[rest_take], take2[rest_take]])
              }
              if(computation.A!="vanRaden"){
                if(miraculix){
                  if (requireNamespace("miraculix", quietly = TRUE)) {
                    p_i <- miraculix::allele_freq(Z.code2)
                  }
                } else{
                  p_i <- rowSums(Zt[,take2[rest_take]])/2
                }
              }
              if(miraculix){

                if (requireNamespace("miraculix", quietly = TRUE)) {
                  if(fast.uhat){
                    u_hat_new[first:last] <- 1/ 2 / sum(p_i*(1-p_i))* miraculix::genoVector(Z.code2, Rest_term[rest_take])
                  } else{
                    u_hat_new[first:last] <- 1/ 2 / sum(p_i*(1-p_i))* miraculix::genoVector(Z.code2, (A1 %*% (y_hat[take2[rest_take],bven] - beta_hat[bven])))
                  }
                }
              } else if(miraculix.mult){
                if (requireNamespace("miraculix", quietly = TRUE)) {
                  if(fast.uhat){
                    u_hat_new[first:last] <- 1/ 2 / sum(p_i*(1-p_i))* miraculix::genoVector(miraculix::genomicmatrix(Zt[,take2[rest_take]]), Rest_term[rest_take])
                  } else{
                    u_hat_new[first:last] <- 1/ 2 / sum(p_i*(1-p_i))* miraculix::genoVector(miraculix::genomicmatrix(Zt[,take2[rest_take]]), (A1 %*% (y_hat[take2[rest_take],bven] - beta_hat[bven])))
                  }
                }
              } else{
                if(fast.uhat){
                  u_hat_new[first:last] <- 1/ 2 / sum(p_i*(1-p_i))*(Zt[,rest_take] %*% Rest_term[rest_take])
                } else{
                  u_hat_new[first:last] <- 1/ 2 / sum(p_i*(1-p_i))*(Zt[,rest_take] %*% (A1 %*% (y_hat[rest_take,bven] - beta_hat[bven])))
                }
              }

              if(store.comp.times.bve){
                after <- as.numeric(Sys.time())
                z_uhat <- z_uhat + after - before
              }


              first <- first + maxZ
              last <- min(maxZ*(index3+1), total_n)
            }
            u_hat <- cbind(u_hat, u_hat_new)

          } else{

            if(store.comp.times.bve){
              before <- as.numeric(Sys.time())
            }
            rest_take <- which(duplicated(c(take,take2))[-(1:length(take))])
            if(!(length(rest_take)==length(prev_rest_take) && prod(rest_take==prev_rest_take)==1)){
              if(miraculix && length(take2)!=nrow(loop_elements)){
                if (requireNamespace("miraculix", quietly = TRUE)) {
                  Z.code2 <- miraculix::computeSNPS(population, loop_elements[take2,4], loop_elements[take2,5], loop_elements[take2,2], what="geno",
                                                  output_compressed = TRUE)
                }
              } else if(length(take2)!=nrow(loop_elements)){
                Zt2 <- Zt[,take2[rest_take]]
              } else if(miraculix){
                Z.code2 <- Z.code
              } else {
                Zt2 <- Zt
              }
              if(!fast.uhat){
                A1 <- MASS::ginv(A[take2[rest_take], take2[rest_take]])
              }

              prev_rest_take <- rest_take
            } else if(length(prev_rest_take)==0){
              if(miraculix){
                Z.code2 <- Z.code
              } else{
                Zt2 <- Zt
              }

            }

            if(computation.A!="vanRaden"){
              if(miraculix){
                if (requireNamespace("miraculix", quietly = TRUE)) {
                  p_i <- miraculix::allele_freq(Z.code2)
                }
              } else{
                p_i <- rowSums(Zt[,take2[rest_take]])/2
              }
            }

            if(miraculix){
              if (requireNamespace("miraculix", quietly = TRUE)) {
                if(fast.uhat){
                  u_hat <- cbind(u_hat, 1/ 2 / sum(p_i*(1-p_i))* miraculix::genoVector(Z.code2, Rest_term[rest_take]))
                } else{
                  u_hat <- cbind(u_hat, 1/ 2 / sum(p_i*(1-p_i)) * miraculix::genoVector(Z.code2, A1 %*% (y_hat[take2[rest_take],bven] - beta_hat[bven])))
                }
              }
            } else if(miraculix.mult){
              if (requireNamespace("miraculix", quietly = TRUE)) {
                if(fast.uhat){
                  u_hat <- cbind(u_hat, 1/ 2 / sum(p_i*(1-p_i))* miraculix::genoVector(miraculix::genomicmatrix(Zt[,take2[rest_take]]), Rest_term[rest_take]))
                } else{
                  u_hat <- cbind(u_hat, 1/ 2 / sum(p_i*(1-p_i))* miraculix::genoVector(miraculix::genomicmatrix(Zt[,take2[rest_take]]), A1 %*% (y_hat[take2[rest_take],bven] - beta_hat[bven])))
                }
              }
            } else{
              if(fast.uhat){
                u_hat <- cbind(u_hat, 1/ 2 / sum(p_i*(1-p_i))*(Zt[,take2[rest_take]] %*% Rest_term[rest_take]))
              } else{
                u_hat <- cbind(u_hat, 1/ 2 / sum(p_i*(1-p_i))*(Zt[,take2[rest_take]] %*% (A1 %*% (y_hat[take2[rest_take],bven] - beta_hat[bven]))))
              }
            }

            if(store.comp.times.bve){
              after <- as.numeric(Sys.time())
              z_uhat <- z_uhat + after - before
            }

          }

          if(rrblup.required){
            if(sequenceZ){
              stop("Not implemented!")
            } else{
              if(miraculix){
                if (requireNamespace("miraculix", quietly = TRUE)) {
                  y_hat[take3,bven] <- u_hat[,bven] %*% (as.matrix(Z.code)[,take3]-2*p_i) + beta_hat[bven]
                }
              } else {
                y_hat[take3,bven] <- u_hat[,bven] %*% Ztm[,take3] + beta_hat[bven]
              }
            }

          }
         }
      } else{
        y_hat[,bven] <- y[,bven]
      }

      if(store.comp.times.bve==TRUE){
        comp.times.bve[4] <- as.numeric(Sys.time())
      }


      if(report.accuracy && bven==population$info$bv.nr){
        if(verbose) cat("Correlation between genetic values and BVE:\n")
        if(n.rep==0){
          y_hat_temp <- y_hat
          y_hat_temp[y_hat_temp==0] <- NA
          acc <- suppressWarnings(stats::cor(y_real[bve.insert,], y_hat_temp[bve.insert,], use="pairwise.complete.obs"))
        } else{
          insert.temp <- numeric(length(bve.insert.copy))

          if(length(stay.loop.elements)>0){
            for(index in (1:nrow(loop_elements_copy))[bve.insert.copy]){
              insert.temp[index] <- which(stay.loop.elements==loop_elements_copy[index,6])
            }
          } else{
            for(index in (1:nrow(loop_elements_copy))[bve.insert.copy]){
              insert.temp[index] <- loop_elements_copy[index,6]
            }

          }
          y_hat_temp <- rbind(y_hat[bve.insert,,drop=FALSE], y_hat[insert.temp,,drop=FALSE])
          y_hat_temp[y_hat_temp==0] <- NA
          acc <- suppressWarnings(stats::cor(rbind(y_real[bve.insert,,drop=FALSE], y_real[insert.temp,, drop=FALSE]),
                                             y_hat_temp, use="pairwise.complete.obs"))
        }
        if(length(acc)==1){
          acc <- matrix(acc,nrow=1)
        }

        if(sum(is.na(acc))>0){
          acc[is.na(acc)] <- 0
        }
        if(verbose) cat(diag(acc))
        if(verbose) cat("\n")
      }
      for(index in (1:nrow(loop_elements))[bve.insert]){
        population$breeding[[loop_elements[index,4]]][[loop_elements[index,5]+2]][, loop_elements[index,2]] <- y_hat[index,]
      }
      if(calculate.reliability){
        for(index in (1:nrow(loop_elements))[bve.insert]){
          population$breeding[[loop_elements[index,4]]][[loop_elements[index,5]+18]][, loop_elements[index,2]] <- y_reli[index,]
        }
      }

      if(n.rep>0){
        for(index in (1:nrow(loop_elements_copy))[bve.insert.copy]){
          if(length(stay.loop.elements)>0){
            non_copy <- which(stay.loop.elements==loop_elements_copy[index,6])
          } else{
            non_copy <- loop_elements_copy[index,6]
          }
          if(length(non_copy)==1){
            population$breeding[[loop_elements_copy[index,4]]][[loop_elements_copy[index,5]+2]][, loop_elements_copy[index,2]] <- y_hat[non_copy,]

          }
        }
      }


      #  GWAS CODE NOT WRITEN FOR PARALLEL COMPUTING
      if(gwas.u){

        if(y.gwas.used=="pheno"){
          y_gwas <- y
        } else if(y.gwas.used=="bv"){
          y_gwas <- y_real
        } else if(y.gwas.used=="bve"){
          y_gwas <- y_hat
        }

        if(nrow(gwas.database)!=nrow(bve.database) || prod(gwas.database==bve.database)==0){

          loop_elements_gwas_list <- derive.loop.elements(population=population, bve.database=bve.database,
                                                bve.class=bve.class, bve.avoid.duplicates=bve.avoid.duplicates,
                                                store.adding=TRUE)
          loop_elements_gwas <- loop_elements_gwas_list[[1]]
          n.animals.gwas <- nrow(loop_elements_gwas)
        }
        if(gwas.group.standard){
          gwas_start <- loop_elements_gwas_list[[2]]
        }
        if(sequenceZ){
          if(nrow(gwas.database)!=nrow(bve.database) || prod(gwas.database==bve.database)==0){
            if(maxZtotal>0){
              maxZ <- floor(maxZtotal / n.animals.gwas)
            }
            if(ncore<=1 && miraculix==FALSE){
              Zt <- array(0L,dim=c(maxZ,n.animals.gwas))
            }
            y_gwas <- array(0, dim=c(n.animals.gwas, population$info$bv.nr))

          }
          total_n <- sum(population$info$snp)
          x_mean <- x2_mean <- xy_mean <- numeric(total_n)
          first <- 1
          last <- min(maxZ, total_n)
          for(index3 in 1:ceiling(total_n/maxZ)){

            cindex <- 1
            for(index in 1:n.animals.gwas){
              k.database <- gwas.database[loop_elements_gwas[index,3],]
              if(miraculix){
                if (requireNamespace("miraculix", quietly = TRUE)) {
                  Zt[1:(last-first+1), cindex] <- miraculix::computeSNPS(population, k.database[1],k.database[2],kindex, from_p=first, to_p=last, what="geno", output_compressed=FALSE)
                }
              } else{
                Zt[1:(last-first+1), cindex] <- base::as.integer(colSums(compute.snps(population, k.database[1],k.database[2],kindex, import.position.calculation=import.position.calculation, from_p=first, to_p=last, decodeOriginsU=decodeOriginsU, bit.storing=bit.storing, nbits=nbits, output_compressed=FALSE)))
              }
              population$breeding[[k.database[1]]][[k.database[2]]][[kindex]][[16]] <- 1
              for(bven in 1:population$info$bv.nr){
                # HIER EVENTUELL SNPs auslesen!
                if(y.gwas.used=="pheno"){
                  y_gwas[cindex,bven] <- population$breeding[[k.database[1]]][[8+k.database[2]]][bven,kindex]
                } else if(y.gwas.used=="bv"){
                  y_gwas[cindex,bven] <- population$breeding[[k.database[1]]][[6+k.database[2]]][bven,kindex]
                } else if(y.gwas.used=="bve"){
                  y_gwas[cindex,bven] <- population$breeding[[k.database[1]]][[2+k.database[2]]][bven,kindex]
                }

              }
              cindex <- cindex +1
            }
          }
          if(gwas.group.standard){
            for(indexg in 1:(length(gwas_start)-1)){
              if(gwas_start[indexg]<=(gwas_start[indexg+1]-1)){
                y_gwas[gwas_start[indexg]:(gwas_start[indexg+1]-1), bven] <- y_gwas[gwas_start[indexg]:(gwas_start[indexg+1]-1), bven] - mean(y_gwas[gwas_start[indexg]:(gwas_start[indexg+1]-1), bven])
              }
            }
          }

          x_mean[first:last] <- rowMeans(Zt[1:(last-first+1),])
          x2_mean[first:last] <- rowMeans(Zt[1:(last-first+1),]^2)
          xy_mean[first:last] <- rowMeans(Zt[1:(last-first+1),]*y_gwas[,bven])
          first <- first + maxZ
          last <- min(maxZ*(index3+1), total_n)
        } else{
          if(nrow(gwas.database)!=nrow(bve.database) || prod(gwas.database==bve.database)==0){
            Zt <- array(0L,dim=c(sum(population$info$snp), n.animals.gwas))
            y_gwas <- array(0, dim=c(n.animals.gwas, population$info$bv.nr))
            cindex <- 1
            for(index in 1:n.animals.gwas){
              k.database <- gwas.database[loop_elements_gwas[index,3],]
              if(miraculix){
                if (requireNamespace("miraculix", quietly = TRUE)) {
                  Zt[,cindex] <- miraculix::computeSNPS(population, k.database[1],k.database[2],kindex, what="geno", output_compressed=FALSE)
                }
              } else{
                Zt[,cindex] <- base::as.integer(colSums(compute.snps(population, k.database[1],k.database[2],kindex, import.position.calculation=import.position.calculation, decodeOriginsU=decodeOriginsU, bit.storing=bit.storing, nbits=nbits, output_compressed=FALSE)))
              }
              for(bven in 1:population$info$bv.nr){
                # HIER EVENTUELL SNPs auslesen!
                if(y.gwas.used=="pheno"){
                  y_gwas[cindex,bven] <- population$breeding[[k.database[1]]][[8+k.database[2]]][bven,kindex]
                } else if(y.gwas.used=="bv"){
                  y_gwas[cindex,bven] <- population$breeding[[k.database[1]]][[6+k.database[2]]][bven,kindex]
                } else if(y.gwas.used=="bve"){
                  y_gwas[cindex,bven] <- population$breeding[[k.database[1]]][[2+k.database[2]]][bven,kindex]
                }
              }
              cindex <- cindex +1
            }
          }

          if(gwas.group.standard){

            for(indexg in 1:(length(gwas_start)-1)){
              if(gwas_start[indexg]>=(gwas_start[indexg+1]-1)){
                y_gwas[gwas_start[indexg]:(gwas_start[indexg+1]-1), bven] <- y_gwas[gwas_start[indexg]:(gwas_start[indexg+1]-1), bven] - mean(y_gwas[gwas_start[indexg]:(gwas_start[indexg+1]-1), bven])
              }
            }
          }
          x_mean <- rowMeans(Zt)
          x2_mean <- rowMeans(Zt^2)
#          xy_mean <- colMeans(Z*y_gwas[,bven])
          xy_mean <- colMeans(t(Zt)*y_gwas[,bven])
        }

        n <- length(y_gwas[,bven])
        y_mean <- mean(y_gwas[,bven])

        b1 <- (n *xy_mean - n *x_mean * y_mean) / (x2_mean*n - n *x_mean^2)


        if(approx.residuals==FALSE){
          sigma1 <- 1/(n * (x2_mean-(x_mean)^2))
          b0 <- y_mean - b1 * x_mean
          var1 <- numeric(length(sigma1))
          for(index in 1:length(sigma1)){
            var1[index] <- sigma1[index] * stats::var(y_gwas[,bven] - b1[index] * Zt[index,] - b0[index]) * (n-1)/(n-2)
          }

        } else{
          var1 <- 1/(n * (x2_mean-(x_mean)^2)) * stats::var(y_gwas[,bven])
        }
        test <- b1/sqrt(var1)
        gwas_hat <- cbind(gwas_hat, test)
        #sorted <- sort(abs(test), index.return=TRUE)
      }

    }

  }
  if(u_hat_possible && bve && estimate.u && computation.A=="vanRaden"){
    population$info$u_hat[[length(population$info$u_hat)+1]] <- u_hat
    population$info$u_hat_single[[length(population$info$u_hat)]] <- list()
    for(bven in 1:ncol(u_hat)){
      population$info$u_hat_single[[length(population$info$u_hat)]][[bven]] <- cbind((-2*p_i) *u_hat[,bven],(-2*p_i+1) *u_hat[,bven],(-2*p_i+2) *u_hat[,bven])
    }
  } else if(u_hat_possible && bve && estimate.u && computation.A=="CM"){
    population$info$u_hat[[length(population$info$u_hat)+1]] <- u_hat
    population$info$u_hat_single[[length(population$info$u_hat)]] <- list()
    for(bven in 1:ncol(u_hat)){
      population$info$u_hat_single[[length(population$info$u_hat)]][[bven]] <- cbind(u_hat[1:nrow(Zt),bven],u_hat[1:nrow(Zt)+ nrow(Zt),bven],u_hat[1:nrow(Zt)+2*nrow(Zt),bven])
    }
  }
  if(gwas.u){
    if(sum(is.na(gwas_hat)>0)){
      gwas_hat[is.na(gwas_hat)] <- 0
    }
    population$info$gwas_hat[[length(population$info$gwas_hat)+1]] <- gwas_hat
  }

  if(store.comp.times.bve){
    comp.times.bve[5] <- as.numeric(Sys.time())
  }
}

  #############################################################################################################
  #############################################################################################################
  ########################## TEST MARTINI######################################################################
{
  if(special.comb==TRUE){
    pos_grp <- matrix(c(length(population$breeding),1), nrow=1)

    n <- sum(population$info$size[pos_grp])
    combination <- matrix(NA, n, n)

    haplo_data <- get.haplo(population, database = pos_grp, chromosomen = 1:10)
    geno_data <- get.geno(population, database = pos_grp, chromosomen = 1:10)

    if(predict.effects==TRUE){
      if(use.effect.combination==TRUE){
        markers_used <- c(population$info$real.bv.mult[[1]][,1] + 500 * population$info$real.bv.mult[[1]][,2] - 500, population$info$real.bv.mult[[1]][,3] + 500 * population$info$real.bv.mult[[1]][,4] - 500)
        X_sub <- geno_data[markers_used, -(1:2)]
        p <- (nrow(population$info$real.bv.mult[[1]]))
        temp1 <- 0
        Z <- matrix(NA, ncol=n, nrow=p*9)
        for(index in 1:(nrow(population$info$real.bv.mult[[1]]))){
          Z1 <- rbind((X_sub[index,]==0)*(X_sub[index+p,]==0), (X_sub[index,]==0)*(X_sub[index+p,]==1),(X_sub[index,]==0)*(X_sub[index+p,]==2),
                      (X_sub[index,]==1)*(X_sub[index+p,]==0), (X_sub[index,]==1)*(X_sub[index+p,]==1),(X_sub[index,]==1)*(X_sub[index+p,]==2),
                      (X_sub[index,]==2)*(X_sub[index+p,]==0), (X_sub[index,]==2)*(X_sub[index+p,]==1),(X_sub[index,]==2)*(X_sub[index+p,]==2))
          Z[1:9+9*temp1,] <- Z1
          temp1 <- temp1+1

        }
        if(mean(population$breeding[[length(population$breeding)]][[9]][1,])!=0){
          y <- population$breeding[[length(population$breeding)]][[9]][1,]
        } else{
          y <- population$breeding[[length(population$breeding)]][[7]][1,]
        }


#        A_BGLR <- tcrossprod(Z) / ncol(Z)
#        ETA <- list(list(K=A_BGLR, model='RKHS'))
#        fm_BGLR <- BGLR(y=y, ETA=ETA, verbose=BGLR.print)
#        u_hat <-  fm_BGLR$yHat - y
#        effects <- crossprod(Z,solve(A_BGLR,u_hat))

        effects <- epi(y, t(Z))
#        model <- rq.fit.lasso(Z,y, lambda = 10)
#        effects <- model$coefficients
        real.effect.mult  <- cbind(population$info$real.bv.mult[[1]][,1:4] , matrix(effects, ncol=9, byrow=TRUE))
#        cor(as.numeric(real.effect.mult[,5:13]), as.numeric(population$info$real.bv.mult[[1]][,5:13]))
        real.effect.add <- NULL
      } else{
        if(use.effect.markers==TRUE){
          markers_used <- c(population$info$real.bv.mult[[1]][,1] + 500 * population$info$real.bv.mult[[1]][,2] - 500, population$info$real.bv.mult[[1]][,3] + 500 * population$info$real.bv.mult[[1]][,4] - 500)
        } else{
          markers_used <- 1:(nrow(geno_data)/SNP.density) * SNP.density
        }

        X_sub <- geno_data[markers_used, -(1:2)]
        Z <- matrix(NA, ncol=ncol(X_sub), nrow=9*(nrow(X_sub))*(nrow(X_sub)+1)/2)
        temp1 <- 0
        for(index in 1:nrow(X_sub)){
          for(index2 in index:nrow(X_sub)){
            Z1 <- rbind((X_sub[index,]==0)*(X_sub[index2,]==0), (X_sub[index,]==0)*(X_sub[index2,]==1),(X_sub[index,]==0)*(X_sub[index2,]==2),
                        (X_sub[index,]==1)*(X_sub[index2,]==0), (X_sub[index,]==1)*(X_sub[index2,]==1),(X_sub[index,]==1)*(X_sub[index2,]==2),
                        (X_sub[index,]==2)*(X_sub[index2,]==0), (X_sub[index,]==2)*(X_sub[index2,]==1),(X_sub[index,]==2)*(X_sub[index2,]==2))
            Z[1:9+9*temp1,] <- Z1
            temp1 <- temp1+1
          }
        }
        #

        Z <- t(Z)
        if(mean(population$breeding[[length(population$breeding)]][[9]][1,])!=0){
          y <- population$breeding[[length(population$breeding)]][[9]][1,]
        } else{
          y <- population$breeding[[length(population$breeding)]][[7]][1,]
        }
        effects <- epi(y, Z)


        pos <- cbind((markers_used-1)%%500+1, (markers_used - (markers_used-1)%%500 -1)/500 +1)
        tz <- which(matrix(0, ncol=nrow(pos), nrow=nrow(pos))==0, arr.ind = TRUE)
        tz <- tz[tz[,1]<=tz[,2],]

        real.effect.mult  <- cbind(pos[tz[,1],1:2], pos[tz[,2],1:2], matrix(effects, ncol=9, byrow=TRUE))
        real.effect.add <- NULL
      }
    } else{
      real.effect.add <- population$info$real.bv.add[[1]]
      real.effect.mult <- population$info$real.bv.mult[[1]]
    }

    p <- nrow(real.effect.mult)


    position0 <- real.effect.add[,1]+ cumsum(c(0,population$info$snp))[real.effect.add[,2]]
    position1 <- real.effect.mult[,1]+ cumsum(c(0,population$info$snp))[real.effect.mult[,2]]
    position2 <- real.effect.mult[,3]+ cumsum(c(0,population$info$snp))[real.effect.mult[,4]]

    positions <- numeric(length(position1)*2)
    positions[1:length(position1)*2-1] <- position1
    positions[1:length(position2)*2] <- position2

    geno_data0 <- geno_data[position0, -(1:2)]
    haplo_data <- haplo_data[positions,-(1:2)]
    geno_data <- geno_data[positions,-(1:2)]
#    haplo_data <- haplo_data[c(position1, position2),-(1:2)]
#    geno_data <- geno_data[c(position1, position2),-(1:2)]

    storage.mode(haplo_data) <- "numeric"
    storage.mode(geno_data) <- "numeric"
    storage.mode(geno_data0) <- "numeric"

    same_chromo_effects <- which(real.effect.mult[,2]==real.effect.mult[,4])
    non_same_chromo_effects <- which(real.effect.mult[,2]!= real.effect.mult[,4])

    if(length(p)==0){
      p <- 0
    }
    effect.prob <- matrix(0, nrow= 4 * n, ncol= p)

    for(index in non_same_chromo_effects){
#      p1 <- geno_data[index*2-1,]/2
#      p2 <- geno_data[index*2,]/2
#      effect.prob[,index] <- c((1-p1)*(1-p2), (1-p1)*(p2), p1 * (1-p2), p1*p2)
      p1 <- geno_data[index*2-1,]
      p2 <- geno_data[index*2,]
      effect.prob[,index] <- c((p1==0)*(p2==0) + 0.5*(p1==1)*(p2==0)+ 0.5*(p1==0)*(p2==1)+ 0.25*(p1==1)*(p2==1),
                               (p1==0)*(p2==2) + 0.5*(p1==0)*(p2==1)+ 0.5*(p1==1)*(p2==2)+ 0.25*(p1==1)*(p2==1),
                               (p1==2)*(p2==0) + 0.5*(p1==1)*(p2==0)+ 0.5*(p1==2)*(p2==1)+ 0.25*(p1==1)*(p2==1),
                               (p1==2)*(p2==2) + 0.5*(p1==1)*(p2==2)+ 0.5*(p1==2)*(p2==1)+ 0.25*(p1==1)*(p2==1))
    }


    switch <- 1:(2*n)
    switch[1:n*2] <- switch[1:n*2]-1
    switch[1:n*2-1] <- switch[1:n*2-1]+1
    for(index in same_chromo_effects){
      distanz <- abs(population$info$snp.position[position1[index]] - population$info$snp.position[position2[index]])
      ab <- sinh(distanz) * exp(-distanz) * 0.5
      aa <- 0.5 - ab

      aa_real <- paste(haplo_data[index*2-1,], haplo_data[index*2,], sep="")
      r00 <- round(which(aa_real=="00")/2+0.01)
      r01 <- round(which(aa_real=="01")/2+0.01)
      r10 <- round(which(aa_real=="10")/2+0.01)
      r11 <- round(which(aa_real=="11")/2+0.01)
      if(length(r00)>0){
        dubs00 <- r00[duplicated(r00)]
        effect.prob[r00,index] <- effect.prob[r00,index] + aa
        effect.prob[dubs00,index] <- effect.prob[dubs00,index] + aa
      }
      if(length(r01)>0){
        dubs01 <- r01[duplicated(r01)]
        effect.prob[r01+n,index] <- effect.prob[r01+n,index] + aa
        effect.prob[dubs01+n,index] <- effect.prob[dubs01+n,index] + aa
      }
      if(length(r10)>0){
        dubs10 <- r10[duplicated(r10)]
        effect.prob[r10+2*n,index] <- effect.prob[r10+2*n,index] + aa
        effect.prob[dubs10+2*n,index] <- effect.prob[dubs10+2*n,index] + aa
      }
      if(length(r11)>0){
        dubs11 <- r11[duplicated(r11)]
        effect.prob[r11+3*n,index] <- effect.prob[r11+3*n,index] + aa
        effect.prob[dubs11+3*n,index] <- effect.prob[dubs11+3*n,index] + aa
      }


      ab_real <- paste(haplo_data[index*2-1,], haplo_data[index*2,switch], sep="")


      r00 <- round(which(ab_real=="00")/2+0.01)
      r01 <- round(which(ab_real=="01")/2+0.01)
      r10 <- round(which(ab_real=="10")/2+0.01)
      r11 <- round(which(ab_real=="11")/2+0.01)
      if(length(r00)>0){
        dubs00 <- r00[duplicated(r00)]
        effect.prob[r00,index] <- effect.prob[r00,index] + ab
        effect.prob[dubs00,index] <- effect.prob[dubs00,index] + ab
      }
      if(length(r01)>0){
        dubs01 <- r01[duplicated(r01)]
        effect.prob[r01 +n,index] <- effect.prob[r01+n,index] + ab
        effect.prob[dubs01+n,index] <- effect.prob[dubs01+n,index] + ab
      }
      if(length(r10)>0){
        dubs10 <- r10[duplicated(r10)]
        effect.prob[r10+2*n,index] <- effect.prob[r10+2*n,index] + ab
        effect.prob[dubs10+2*n,index] <- effect.prob[dubs10+2*n,index] + ab
      }
      if(length(r11)>0){
        dubs11 <- r11[duplicated(r11)]
        effect.prob[r11+3*n,index] <- effect.prob[r11+3*n,index] + ab
        effect.prob[dubs11+3*n,index] <- effect.prob[dubs11+3*n,index] + ab
      }

    }


    teffect.prob <- t(effect.prob)
    for(index2 in 1:n){
      if(index2%%25==0) print(index2)
      if(index2==1){
        combination[1:index2, index2] <- combination[index2, 1:index2] <-sum(real.effect.mult[,5]* teffect.prob[,1:index2] * teffect.prob[,index2] + # 00 Effekt
                                                                                   real.effect.mult[,6]* (teffect.prob[,1:index2+n] * teffect.prob[,index2]+teffect.prob[,1:index2] * teffect.prob[,index2+n]) + # 01 Effekt
                                                                                   real.effect.mult[,7]* (teffect.prob[,1:index2+n] * teffect.prob[,index2+n]) + # 02 Effekt
                                                                                   real.effect.mult[,8]* (teffect.prob[,1:index2+2*n] * teffect.prob[,index2]+teffect.prob[,1:index2] * teffect.prob[,index2+2*n]) + # 10 Effekt
                                                                                   real.effect.mult[,9]* (teffect.prob[,1:index2+3*n] * teffect.prob[,index2]+teffect.prob[,1:index2] * teffect.prob[,index2+3*n]+teffect.prob[,1:index2+n] * teffect.prob[,index2+2*n]+teffect.prob[,1:index2+2*n] * teffect.prob[,index2+n]) + # 11 Effekt
                                                                                   real.effect.mult[,10]* (teffect.prob[,1:index2+3*n] * teffect.prob[,index2+n]+teffect.prob[,1:index2+n] * teffect.prob[,index2+3*n]) + # 12 Effekt
                                                                                   real.effect.mult[,11]* (teffect.prob[,1:index2+2*n] * teffect.prob[,index2+2*n]) + # 20 Effekt
                                                                                   real.effect.mult[,12]* (teffect.prob[,1:index2+3*n] * teffect.prob[,index2+2*n]+teffect.prob[,1:index2+2*n] * teffect.prob[,index2+3*n]) + # 21 Effekt
                                                                                   real.effect.mult[,13]* (teffect.prob[,1:index2+3*n] * teffect.prob[,index2+3*n]))  # 22 Effekt

      } else{
        combination[1:index2, index2] <- combination[index2, 1:index2] <-colSums(real.effect.mult[,5]* teffect.prob[,1:index2] * teffect.prob[,index2] + # 00 Effekt
                                                                                   real.effect.mult[,6]* (teffect.prob[,1:index2+n] * teffect.prob[,index2]+teffect.prob[,1:index2] * teffect.prob[,index2+n]) + # 01 Effekt
                                                                                   real.effect.mult[,7]* (teffect.prob[,1:index2+n] * teffect.prob[,index2+n]) + # 02 Effekt
                                                                                   real.effect.mult[,8]* (teffect.prob[,1:index2+2*n] * teffect.prob[,index2]+teffect.prob[,1:index2] * teffect.prob[,index2+2*n]) + # 10 Effekt
                                                                                   real.effect.mult[,9]* (teffect.prob[,1:index2+3*n] * teffect.prob[,index2]+teffect.prob[,1:index2] * teffect.prob[,index2+3*n]+teffect.prob[,1:index2+n] * teffect.prob[,index2+2*n]+teffect.prob[,1:index2+2*n] * teffect.prob[,index2+n]) + # 11 Effekt
                                                                                   real.effect.mult[,10]* (teffect.prob[,1:index2+3*n] * teffect.prob[,index2+n]+teffect.prob[,1:index2+n] * teffect.prob[,index2+3*n]) + # 12 Effekt
                                                                                   real.effect.mult[,11]* (teffect.prob[,1:index2+2*n] * teffect.prob[,index2+2*n]) + # 20 Effekt
                                                                                   real.effect.mult[,12]* (teffect.prob[,1:index2+3*n] * teffect.prob[,index2+2*n]+teffect.prob[,1:index2+2*n] * teffect.prob[,index2+3*n]) + # 21 Effekt
                                                                                   real.effect.mult[,13]* (teffect.prob[,1:index2+3*n] * teffect.prob[,index2+3*n]))  # 22 Effekt

      }
    }


    #removes <- 1:500*500-499:0
    #plot(combination0[-removes], combination1[-removes])
    #cor(combination[-removes], combination2[-removes])

    if(max.auswahl<1000){
      auswahl <- which(combination>=(-999), arr.ind=TRUE)
      values <- combination[auswahl]
      removes <- (auswahl[,1]<auswahl[,2])*1:length(values)
      values <- values[removes]
      auswahl <- auswahl[removes,]
      ordering <- sort(values, index.return=TRUE, decreasing = TRUE)
      auswahl <- auswahl[ordering$ix,]
      counts <- numeric(n)
      choosen <- numeric(n)
      row <- 1
      activ <- 1
      while(sum(counts)<(2*n)){
        if(counts[auswahl[row,1]]<max.auswahl && counts[auswahl[row,2]]<max.auswahl){
          counts[auswahl[row,1]] <- counts[auswahl[row,1]] +1
          counts[auswahl[row,2]] <- counts[auswahl[row,2]] +1
          choosen[activ] <- row
          activ <- activ +1
        }
        row <- row +1
      }
      auswahl <- auswahl[choosen,]
    } else{
      cutoff <- sort(combination, decreasing = TRUE)[n*4+1]
      auswahl <- which(combination>=cutoff, arr.ind = TRUE)
      auswahl <- auswahl[auswahl[,1]<auswahl[,2],]
      values <- combination[auswahl]

      ordering <- sort(values, index.return=TRUE, decreasing = TRUE)
      auswahl <- auswahl[ordering$ix[1:n],]
    }

    animals <- c(auswahl[,1], auswahl[,2])

    animals1 <- unique(animals)
    count <- numeric(length(animals1))
    for(index in 1:length(animals1)){
      count[index] <- sum(animals==animals1[index])
    }
    fixed.breeding <- cbind(length(population$breeding), 1, auswahl[,1], length(population$breeding), 1, auswahl[,2], 0)
    population$breeding[[length(population$breeding)]][[11]] <- rbind(animals1, count)
    population$breeding[[length(population$breeding)]][[12]] <- ordering$x[1:n]
  }

  if(special.comb.add==TRUE){
    add_bv <- population$breeding[[length(population$breeding)]][[3]][1,]
    n <- length(add_bv)
    combination <- matrix(NA, n, n)
    for(index in 1:n){
      for(index2 in index:n){
        combination[index, index2]  <- add_bv[index] + add_bv[index2]
      }
    }
    if(max.auswahl<500){
      auswahl <- which(combination>=(-999), arr.ind=TRUE)
      values <- combination[auswahl]
      removes <- (auswahl[,1]<auswahl[,2])*1:length(values)
      values <- values[removes]
      auswahl <- auswahl[removes,]
      ordering <- sort(values, index.return=TRUE, decreasing = TRUE)
      auswahl <- auswahl[ordering$ix,]
      counts <- numeric(n)
      choosen <- numeric(n)
      row <- 1
      activ <- 1
      while(sum(counts)<(2*n)){
        if(counts[auswahl[row,1]]<max.auswahl && counts[auswahl[row,2]]<max.auswahl){
          counts[auswahl[row,1]] <- counts[auswahl[row,1]] +1
          counts[auswahl[row,2]] <- counts[auswahl[row,2]] +1
          choosen[activ] <- row
          activ <- activ +1
        }
        row <- row +1
      }
      auswahl <- auswahl[choosen,]
    } else{
      cutoff <- sort(combination, decreasing = TRUE)[n*4+1]
      auswahl <- which(combination>=cutoff, arr.ind = TRUE)
      auswahl <- auswahl[auswahl[,1]<auswahl[,2],]
      values <- combination[auswahl]

      ordering <- sort(values, index.return=TRUE, decreasing = TRUE)
      auswahl <- auswahl[ordering$ix[1:n],]
    }

    animals <- c(auswahl[,1], auswahl[,2])

    animals1 <- unique(animals)
    count <- numeric(length(animals1))
    for(index in 1:length(animals1)){
      count[index] <- sum(animals==animals1[index])
    }
    fixed.breeding <- cbind(length(population$breeding), 1, auswahl[,1], length(population$breeding), 1, auswahl[,2], 0)
    population$breeding[[length(population$breeding)]][[11]] <- rbind(animals1, count)
    population$breeding[[length(population$breeding)]][[12]] <- ordering$x[1:n]

  }
}
  ######################### ENDE TEST MARTINI ######################
  #############################################################################################################
  #############################################################################################################
  #############################################################################################################
{
  if(gene.editing){
    if(estimate.u){
      effect_order<- sort(abs(population$info$u_hat[[length(population$info$u_hat)]]), index.return=TRUE, decreasing=TRUE)$ix
      direction <- (population$info$u_hat[[length(population$info$u_hat)]] < 0)
      distance_actual <- integer(length(effect_order))
      for(index in 1:length(effect_order)){
        distance_actual[index] <- min(abs(effect_order[index]-population$info$effect.p))
      }
      if(length(population$info$editing_info)==0){
        population$info$editing_info <- list()
      }
      population$info$editing_info[[length(population$info$editing_info)+1]] <- cbind(effect_order, direction[effect_order], distance_actual, deparse.level = 0)
    }
    if(gwas.u){
      effect_order <- sort(abs(population$info$gwas_hat[[length(population$info$gwas_hat)]]), index.return=TRUE, decreasing=TRUE)$ix
      direction <- (population$info$gwas_hat[[length(population$info$gwas_hat)]] <0 )
      distance_actual <- integer(length(effect_order))
      for(index in 1:length(effect_order)){
        distance_actual[index] <- min(abs(effect_order[index]-population$info$effect.p))
      }
      if(length(population$info$editing_info)==0){
        population$info$editing_info <- list()
      }
      population$info$editing_info[[length(population$info$editing_info)+1]] <- cbind(effect_order, direction[effect_order], distance_actual, deparse.level = 0)
    }
  }


  if(store.comp.times){
    comp.times[5] <- as.numeric(Sys.time())
  }
}

{
    {
    best.m <- matrix(0, nrow=selection.size[1],ncol=5)
    best.f <- matrix(0, nrow=selection.size[2],ncol=5)

    addsel <- c(2,2)
    if(selection.criteria.type[1]=="bv"){
      addsel[1] = 6
    }
    if(selection.criteria.type[2]=="bv"){
      addsel[2] = 6
    }
    if(selection.criteria.type[1]=="pheno"){
      addsel[1] = 8
    }
    if(selection.criteria.type[2]=="pheno"){
      addsel[2] = 8
    }

    if(add.class.cohorts && length(selection.m.cohorts)>0){
      for(index in 1:length(selection.m.cohorts)){
        take <- which(population$info$cohorts[,1]==selection.m.cohorts[index])
        class.m <- c(class.m, as.numeric(population$info$cohorts[take,5]))
      }
      class[[1]] <- class.m
    }
    if(add.class.cohorts && length(selection.f.cohorts)>0){
      for(index in 1:length(selection.f.cohorts)){
        take <- which(population$info$cohorts[,1]==selection.f.cohorts[index])
        class.f <- c(class.f, as.numeric(population$info$cohorts[take,5]))
      }
      class[[2]] <- class.f
    }


    best <- list(best.m,best.f)

  }

  activ_groups_list <- list(selection.m.database, selection.f.database)
  chosen.animals.list <- list()
  selection.size.sex <- list(selection.size[1], selection.size[2])
  selection.sex <- c(selection.m, selection.f)
  selection.miesenberger <- c(selection.m.miesenberger, selection.f.miesenberger)
  multiple.bve.weights <- list(multiple.bve.weights.m, multiple.bve.weights.f)
  multiple.bve.scale <- c(multiple.bve.scale.m, multiple.bve.scale.f)
  sd_scaling <- rep(1, population$info$bv.nr)
  if(length(fixed.breeding)==0){
    if(sum(selection.size)>0){
      if(verbose) cat("Start selection procedure.\n")
      for(sex in (1:2)[selection.size>0]){
        possible_animals <- NULL
        activ_groups <- activ_groups_list[[sex]]
        for(index5 in 1:nrow(activ_groups)){
          possible_animals <- rbind(possible_animals, cbind(activ_groups[index5,1], activ_groups[index5,2], activ_groups[index5,3]:activ_groups[index5,4], population$breeding[[activ_groups[index5,1]]][[4+activ_groups[index5,2]]][activ_groups[index5,3]:activ_groups[index5,4]]))
        }
        relevant.animals <- rep(possible_animals[,4], length(class[[sex]])) == rep(class[[sex]],each=nrow(possible_animals))
        relevant.animals <- unique(c(0,relevant.animals * 1:nrow(possible_animals)))[-1]
        possible_animals <- rbind(NULL, possible_animals[unique(relevant.animals),])
        if(length(reduced.selection.panel[[sex]])>0 && length(possible_animals)>0){
          possible_animals <- possible_animals[reduced.selection.panel[[sex]],,drop=FALSE]
        }
        n.animals <- nrow(possible_animals)

        if(selection.sex[sex]=="random"){
          if(population$info$bv.nr==0){
            import.bv <- rep(0, n.animals)
          } else if(population$info$bv.nr==1){
            import.bv <- rep(0, n.animals)
            for(index5 in 1:nrow(possible_animals)){
              import.bv[index5] <- population$breeding[[possible_animals[index5,1]]][[possible_animals[index5,2]+addsel[possible_animals[index5,2]]]][,possible_animals[index5,3]]
            }
          } else{
            if(multiple.bve=="add"){
              breeding.values <- matrix(0, ncol=nrow(possible_animals), nrow= population$info$bv.nr)
              for(index5 in 1:nrow(possible_animals)){
                breeding.values[,index5] <- population$breeding[[possible_animals[index5,1]]][[possible_animals[index5,2]+addsel[possible_animals[index5,2]]]][,possible_animals[index5,3]]
              }
              if(multiple.bve.scale[sex]=="bve_sd" || multiple.bve.scale[sex]=="pheno_sd"){

                sd_store <- numeric(population$info$bv.nr)
                if(ncol(breeding.values)>1){
                  P <- stats::var(t(breeding.values))
                } else{
                  P <- diag(nrow(breeding.values))
                }

                if(multiple.bve.scale[sex]=="pheno_sd"){
                  pheno.values <- genomic.values <-  matrix(0, ncol=nrow(possible_animals), nrow= population$info$bv.nr)
                  for(index5 in 1:nrow(possible_animals)){
                    pheno.values[,index5] <- population$breeding[[possible_animals[index5,1]]][[possible_animals[index5,2]+8]][,possible_animals[index5,3]]
                    genomic.values[,index5] <- population$breeding[[possible_animals[index5,1]]][[possible_animals[index5,2]+6]][,possible_animals[index5,3]]
                  }
                }
                for(bven in 1:population$info$bv.nr){
                  breeding.values[bven,] <- breeding.values[bven,] - mean(breeding.values[bven,])
                  if(ncol(breeding.values)!=1){
                    if(multiple.bve.scale[sex]=="bve_sd"){
                      sd_store[bven] <- stats::sd(breeding.values[bven,])
                    } else{
                      sd_store[bven] <- stats::sd(pheno.values[bven,])
                      if(sd_store[bven]==0){
                        if(verbose) cat("No observed phenotypes in the group of selected individuals. Sure you want to scale according to phenotypes?\n")
                        if(verbose) cat("Expected phenotypic sd based on one observation was used!\n")
                        sd_store[bven] <- sqrt(stats::var(genomic.values[bven,]) + sigma.e[bven])
                      }
                    }

                  } else{
                    sd_store[bven] <- 1
                  }

                  if(sd_store[bven]>0){
                    breeding.values[bven,] <- breeding.values[bven,] / sd_store[bven]
                  }
                }
              }

              bve.sum <- colSums(breeding.values * multiple.bve.weights[[sex]])

            } else if(multiple.bve=="ranking"){
              breeding.values <- matrix(0, ncol=nrow(possible_animals), nrow= population$info$bv.nr)
              for(index5 in 1:nrow(possible_animals)){
                breeding.values[,index5] <- population$breeding[[possible_animals[index5,1]]][[possible_animals[index5,2]+addsel[possible_animals[index5,2]]]][,possible_animals[index5,3]]
              }
              ranking <- matrix(0, ncol=nrow(possible_animals), nrow= population$info$bv.nr)
              for(bven in 1:population$info$bv.nr){
                order <- sort(breeding.values[bven,], index.return=TRUE, decreasing=selection.critera[sex])$ix
                ranking[bven,order] <- length(order):1
              }
              bve.sum <- colSums(ranking*multiple.bve.weights[[sex]])
            }
            import.bv <- bve.sum
          }
          chosen.animals <- relevant.animals[sample(1:n.animals, selection.size[sex])]
          best[[sex]][,1:3] <- possible_animals[chosen.animals,1:3]
          best[[sex]][,4] <- import.bv[chosen.animals]
        } else if(selection.sex[sex]=="function"){
          if(population$info$bv.nr==1){
            import.bv <- numeric(nrow(possible_animals))
            for(index5 in 1:nrow(possible_animals)){
              import.bv[index5] <- population$breeding[[possible_animals[index5,1]]][[possible_animals[index5,2]+addsel[possible_animals[index5,2]]]][,possible_animals[index5,3]]
            }
            chosen.animals <- sort(import.bv, index.return=TRUE, decreasing=selection.critera[sex])$ix[1:sum(selection.size[[sex]])] # Diese 3er werden zu 4 in Weiblich
            best[[sex]][,1:4] <- cbind(possible_animals[chosen.animals, 1], possible_animals[chosen.animals, 2], possible_animals[chosen.animals, 3], import.bv[chosen.animals])
          } else{
            if(multiple.bve=="add"){
              breeding.values <- matrix(0, ncol=nrow(possible_animals), nrow= population$info$bv.nr)
              for(index5 in 1:nrow(possible_animals)){
                breeding.values[,index5] <- population$breeding[[possible_animals[index5,1]]][[possible_animals[index5,2]+addsel[possible_animals[index5,2]]]][,possible_animals[index5,3]]
              }
              if((multiple.bve.scale[sex]=="bve_sd" || multiple.bve.scale[sex]=="pheno_sd") && selection.miesenberger[sex]==FALSE){

                sd_scaling <- numeric(population$info$bv.nr)
                if(ncol(breeding.values)>1){
                  P <- stats::var(t(breeding.values))
                } else{
                  P <- diag(nrow(breeding.values))
                }

                if(multiple.bve.scale[sex]=="pheno_sd"){
                  pheno.values <- genomic.values <-  matrix(0, ncol=nrow(possible_animals), nrow= population$info$bv.nr)
                  for(index5 in 1:nrow(possible_animals)){
                    pheno.values[,index5] <- population$breeding[[possible_animals[index5,1]]][[possible_animals[index5,2]+8]][,possible_animals[index5,3]]
                    genomic.values[,index5] <- population$breeding[[possible_animals[index5,1]]][[possible_animals[index5,2]+6]][,possible_animals[index5,3]]
                  }
                }
                for(bven in 1:population$info$bv.nr){
                  breeding.values[bven,] <- breeding.values[bven,] - mean(breeding.values[bven,])
                  if(ncol(breeding.values)!=1){
                    if(multiple.bve.scale[sex]=="bve_sd"){
                      sd_scaling[bven] <- stats::sd(breeding.values[bven,])
                    } else{
                      sd_scaling[bven] <- stats::sd(pheno.values[bven,])
                      if(sd_scaling[bven]==0){
                        if(verbose) cat("No observed phenotypes in the group of selected individuals. Sure you want to scale according to phenotypes?\n")
                        if(verbose) cat("Expected phenotypic sd based on one observation was used!\n")
                        sd_scaling[bven] <- sqrt(stats::var(genomic.values[bven,]) + sigma.e[bven])
                      }
                    }

                  } else{
                    sd_scaling[bven] <- 1
                  }

                  if(sd_scaling[bven]>0){
                    breeding.values[bven,] <- breeding.values[bven,] / sd_scaling[bven]
                  }
                }
              }

              if(selection.miesenberger[sex]){
                stop("MIESENBERGER IS CURRENTLY NOT FUNCTIONAL!")
                genomic.values <- matrix(0, ncol=nrow(possible_animals), nrow= population$info$bv.nr)
                bve.values <- matrix(0, ncol=nrow(possible_animals), nrow= population$info$bv.nr)
                pheno.values <- matrix(0, ncol=nrow(possible_animals), nrow= population$info$bv.nr)
                reliability.values <- matrix(0, ncol=nrow(possible_animals), nrow= population$info$bv.nr)
                bve.index <- matrix(0, ncol=nrow(possible_animals), nrow= population$info$bv.nr)

                for(index5 in 1:nrow(possible_animals)){
                  genomic.values[,index5] <- population$breeding[[possible_animals[index5,1]]][[possible_animals[index5,2]+6]][,possible_animals[index5,3]]
                  bve.values[,index5] <- population$breeding[[possible_animals[index5,1]]][[possible_animals[index5,2]+2]][,possible_animals[index5,3]]
                  pheno.values[,index5] <- population$breeding[[possible_animals[index5,1]]][[possible_animals[index5,2]+8]][,possible_animals[index5,3]]
                  reliability.values[,index5] <- population$breeding[[possible_animals[index5,1]]][[possible_animals[index5,2]+18]][,possible_animals[index5,3]]
                }
                if(sum(bve.values==0)==length(bve.values)){
                  if(verbose) cat("No breeding values for selected individuals! Miesenberger selection was skip.\n")

                  index.weights <- multiple.bve.weights[[sex]]
                  for(index5 in 1:nrow(possible_animals)){
                    bve.index[,index5] <- index.weights
                    population$breeding[[possible_animals[index5,1]]][[possible_animals[index5,2]+20]][,possible_animals[index5,3]] <- bve.index[,index5]
                  }
                } else{
                  active_traits <- which(rowSums(bve.values==0)<nrow(bve.values))
                  if(length(active_traits)<population$info$bv.nr){
                    if(verbose) cat("Only use traits", population$info$trait.name[active_traits], "for Miesenberger.\n")
                    if(verbose) cat("Traits", population$info$trait.name[-active_traits], "with no BVE.\n")
                  }
                  genomic.cov <- stats::var(t(genomic.values[active_traits,,drop=FALSE]))
                  bve.cov <- stats::var(t(bve.values[active_traits,,drop=FALSE]))
                  pheno.cov <- stats::var(t(pheno.values[active_traits,,drop=FALSE]))

#                  diag(genomic.cov) <- diag(genomic.cov) + 10^-8
#                  diag(bve.cov) <- diag(bve.cov) + 10^-8
#                  diag(pheno.cov) <- diag(pheno.cov) + 10^-8

                  heri.emp <- diag(genomic.cov) / diag(pheno.cov)
                  derived.reliability <- diag(stats::cor(t(genomic.values[active_traits,,drop=FALSE]), t(bve.values[active_traits,,drop=FALSE])))^2
                  estimated.reliability <- sqrt(diag(bve.cov) / diag(pheno.cov))
                  estimated.reliability[diag(pheno.cov)==0] <- derived.reliability[diag(pheno.cov)==0]
                  estimated.reliability[estimated.reliability>1] <- 1


                  for(bven in 1:length(active_traits)){
                    trait_index <- active_traits[bven]
                    if(selection.miesenberger.reliability.est=="estimated"){
                      reliability.values[trait_index, reliability.values[trait_index,]==0] <- estimated.reliability[bven]
                    } else if(selection.miesenberger.reliability.est=="heritability"){
                      reliability.values[trait_index, reliability.values[trait_index,]==0] <- heri.emp[bven]
                    } else if(selection.miesenberger.reliability.est=="derived"){
                      reliability.values[trait_index, reliability.values[trait_index,]==0] <- derived.reliability[bven]
                    }

                  }
                  reliability.values[is.na(reliability.values)] <- 0

                  V <- bve.cov
                  V1 <- MASS::ginv(V)
                  # V1 <- chol2inv(chol(V))

                  G_cov <- genomic.cov
                  RG <- sqrt(diag(1/diag(G_cov))) %*% G_cov %*% sqrt(diag(1/diag(G_cov)))

                  if(multiple.bve.scale[sex]=="pheno_sd"){
                    index.weights <- multiple.bve.weights[[sex]][active_traits] / sqrt(diag(pheno.cov))
                    if(sum(diag(pheno.cov)==0)>0){
                      if(verbose) cat("Are you sure you want to scale Miesenberger w by phenotypic sd? No phenotypes for some traits!\n")
                      if(verbose) cat("Expected phenotypic sd was used!\n")
                      index.weights <- multiple.bve.weights[[sex]][active_traits] / sqrt(diag(genomic.cov) + sigma.e[active_traits])
                    }
                  } else if(multiple.bve.scale[sex]=="unit"){
                    index.weights <- multiple.bve.weights[[sex]][active_traits]
                  } else if(multiple.bve.scale[sex]=="bve_sd"){
                    index.weights <- multiple.bve.weights[[sex]][active_traits] / sqrt(diag(bve.cov))
                    index.weights[diag(bve.cov)==0] <- 0
                  }
                  for(index5 in 1:nrow(possible_animals)){
                    bve.index[active_traits,index5] <- miesenberger.index(V1=V1, V= V, G = G_cov, RG = RG, r = sqrt(reliability.values[active_traits,index5]), w = index.weights)
                    population$breeding[[possible_animals[index5,1]]][[possible_animals[index5,2]+20]][,possible_animals[index5,3]] <- bve.index[,index5]
                  }
                  if(verbose) cat("Average index used for in selection according to Miesenberger:\n")
                  if(multiple.bve.scale[sex]=="pheno_sd"){
                    if(sum(diag(pheno.cov)==0)>0){
                      if(verbose) cat(rowMeans(bve.index) * sqrt(diag(genomic.cov) + sigma.e[active_traits]))
                    } else{
                      if(verbose) cat(rowMeans(bve.index) * sqrt(diag(pheno.cov)))
                    }
                  } else if(multiple.bve.scale[sex]=="unit"){
                    if(verbose) cat(rowMeans(bve.index))
                  } else if(multiple.bve.scale[sex]=="bve_sd"){
                    if(verbose) cat(rowMeans(bve.index) * sqrt(diag(bve.cov)))
                  }

                  if(verbose) cat("\n")

                }

                bve.sum <- colSums(bve.index * breeding.values, na.rm=TRUE)

              } else{
                bve.sum <- colSums(breeding.values * multiple.bve.weights[[sex]], na.rm=TRUE)
              }
            } else if(multiple.bve=="ranking"){
              breeding.values <- matrix(0, ncol=nrow(possible_animals), nrow= population$info$bv.nr)
              for(index5 in 1:nrow(possible_animals)){
                breeding.values[,index5] <- population$breeding[[possible_animals[index5,1]]][[possible_animals[index5,2]+addsel[possible_animals[index5,2]]]][,possible_animals[index5,3]]
              }
              ranking <- matrix(0, ncol=nrow(possible_animals), nrow= population$info$bv.nr)
              for(bven in 1:population$info$bv.nr){
                order <- sort(breeding.values[bven,], index.return=TRUE, decreasing=selection.critera[sex])$ix
                ranking[bven,order] <- length(order):1
              }
              bve.sum <- colSums(ranking*multiple.bve.weights[[sex]], na.rm=TRUE)
            }
            chosen.animals <- sort(bve.sum, index.return=TRUE, decreasing=selection.critera[sex])$ix[1:sum(selection.size.sex[[sex]])] # Diese 3er werden zu 4 in Weiblich
            best[[sex]][,1:4] <- cbind(possible_animals[chosen.animals,1:3, drop=FALSE], bve.sum[chosen.animals])
          }

        }

        if(selection.miesenberger[sex]==FALSE){
          for(index5 in 1:nrow(possible_animals)){
            population$breeding[[possible_animals[index5,1]]][[possible_animals[index5,2]+20]][,possible_animals[index5,3]] <- multiple.bve.weights[[sex]]
          }
        }

        ## Just some follow up computation in case individuasl are not picked with equal share
        if(population$info$bv.nr==1){
          breeding.values <- numeric(length(chosen.animals))
          for(running in 1:length(chosen.animals)){
            if(best.selection.criteria[[sex]]=="bv"){
              breeding.values[running] <- population$breeding[[possible_animals[chosen.animals[running],1]]][[possible_animals[chosen.animals[running],2]+6]][1,possible_animals[chosen.animals[running],3]]
            } else if(best.selection.criteria[[sex]]=="pheno"){
              breeding.values[running] <- population$breeding[[possible_animals[chosen.animals[running],1]]][[possible_animals[chosen.animals[running],2]+8]][1,possible_animals[chosen.animals[running],3]]
            } else{
              breeding.values[running] <- population$breeding[[possible_animals[chosen.animals[running],1]]][[possible_animals[chosen.animals[running],2]+2]][1,possible_animals[chosen.animals[running],3]]
            }
          }
          best[[sex]][,5] <- breeding.values
        } else if(population$info$bv.nr>1){
          if(multiple.bve=="add"){
            breeding.values <- matrix(0, nrow=population$info$bv.nr, ncol=length(chosen.animals))
            breeding.values.scaling <- matrix(0, nrow=population$info$bv.nr, ncol=length(chosen.animals))
            for(running in 1:length(chosen.animals)){
              if(best.selection.criteria[[sex]]=="bv"){
                breeding.values[,running] <- population$breeding[[possible_animals[chosen.animals[running],1]]][[possible_animals[chosen.animals[running],2]+6]][,possible_animals[chosen.animals[running],3]]
              } else if(best.selection.criteria[[sex]]=="pheno"){
                breeding.values[,running] <- population$breeding[[possible_animals[chosen.animals[running],1]]][[possible_animals[chosen.animals[running],2]+8]][,possible_animals[chosen.animals[running],3]]
              } else{
                breeding.values[,running] <- population$breeding[[possible_animals[chosen.animals[running],1]]][[possible_animals[chosen.animals[running],2]+2]][,possible_animals[chosen.animals[running],3]]
              }
              breeding.values.scaling[,running] <- population$breeding[[possible_animals[chosen.animals[running],1]]][[possible_animals[chosen.animals[running],2]+20]][,possible_animals[chosen.animals[running],3]]

            }
            if((multiple.bve.scale[sex]=="bve_sd" || multiple.bve.scale[sex]=="pheno_sd") && selection.miesenberger[sex]==FALSE){

              sd_scaling <- numeric(population$info$bv.nr)
              if(ncol(breeding.values)>1){
                P <- stats::var(t(breeding.values))
              } else{
                P <- diag(nrow(breeding.values))
              }

              if(multiple.bve.scale[sex]=="pheno_sd"){
                pheno.values <- genomic.values <-  matrix(0, ncol=nrow(possible_animals), nrow= population$info$bv.nr)
                for(index5 in 1:nrow(possible_animals)){
                  pheno.values[,index5] <- population$breeding[[possible_animals[index5,1]]][[possible_animals[index5,2]+8]][,possible_animals[index5,3]]
                  genomic.values[,index5] <- population$breeding[[possible_animals[index5,1]]][[possible_animals[index5,2]+6]][,possible_animals[index5,3]]
                }
              }
              for(bven in 1:population$info$bv.nr){
                breeding.values[bven,] <- breeding.values[bven,] - mean(breeding.values[bven,])
                if(ncol(breeding.values)!=1){
                  if(multiple.bve.scale[sex]=="bve_sd"){
                    sd_scaling[bven] <- stats::sd(breeding.values[bven,])
                  } else{
                    sd_scaling[bven] <- stats::sd(pheno.values[bven,])
                    if(sd_scaling[bven]==0){
                      if(verbose) cat("No observed phenotypes in the group of selected individuals. Sure you want to scale according to phenotypes?\n")
                      if(verbose) cat("Expected phenotypic sd based on one observation was used!\n")
                      sd_scaling[bven] <- sqrt(stats::var(genomic.values[bven,]) + sigma.e[bven])
                    }
                  }

                } else{
                  sd_scaling[bven] <- 1
                }

                if(sd_scaling[bven]>0){
                  breeding.values[bven,] <- breeding.values[bven,] / sd_scaling[bven]
                }
              }
            }
            bve.sum <- colSums(breeding.values * breeding.values.scaling, na.rm=TRUE)
          } else if(multiple.bve=="ranking"){
            ranking <- matrix(0, nrow=population$info$bv.nr, ncol=length(chosen.animals))
            for(running in 1:length(chosen.animals)){
              if(best.selection.criteria[[sex]]=="bv"){
                ranking[,running] <- population$breeding[[possible_animals[chosen.animals[running],1]]][[possible_animals[chosen.animals[running],2]+6]][,possible_animals[chosen.animals[running],3]]
              } else if(best.selection.criteria[[sex]]=="pheno"){
                ranking[,running] <- population$breeding[[possible_animals[chosen.animals[running],1]]][[possible_animals[chosen.animals[running],2]+8]][,possible_animals[chosen.animals[running],3]]
              } else{
                ranking[,running] <- population$breeding[[possible_animals[chosen.animals[running],1]]][[possible_animals[chosen.animals[running],2]+2]][,possible_animals[chosen.animals[running],3]]
              }
            }
            for(bven in 1:population$info$bv.nr){
              order <- sort(ranking[bven,], index.return=TRUE, decreasing=selection.critera[sex])$ix
              ranking[bven,order] <- length(order):1
            }
            bve.sum <- colSums(ranking*breeding.values.scaling, na.rm=TRUE)
          }
          best[[sex]][,5] <- bve.sum
        }

        reorder <- sort(best[[sex]][,4], index.return=TRUE, decreasing = selection.critera[sex])$ix
        best[[sex]] <- best[[sex]][reorder,,drop=FALSE]
        chosen.animals <- chosen.animals[reorder]
        if(ignore.best[sex]>0){
          best[[sex]] <- best[[sex]][-(1:ignore.best[sex]),,drop=FALSE]
          chosen.animals <- chosen.animals[-(1:ignore.best[sex])]
          selection.size[sex] <- selection.size[sex] - ignore.best[sex]

        }

        chosen.animals.list[[sex]] <- chosen.animals
      }

      if(fixed.assignment!=FALSE){
        nm <- nrow(best[[1]])
        nw <- nrow(best[[2]])
        male.rounds <- breeding.size.total / nm
        female.rounds <- breeding.size.total / nw
        full.m <- nm*(male.rounds - round(male.rounds - 0.5))
        full.f <- nw*(female.rounds - round(female.rounds -0.5))
        fixed.assignment.m <- c(rep(1:full.m,each=male.rounds-full.m/nm+1), rep((full.m+1):nm, each =male.rounds-full.m/nm))
        fixed.assignment.f <- c(rep(1:full.f,each=female.rounds - full.f/nw +1), rep((full.f+1):nw, each =female.rounds-full.f/nw))
        fixed.assignment.m <- fixed.assignment.m[1:breeding.size.total]
        fixed.assignment.f <- fixed.assignment.f[1:breeding.size.total]
        if(fixed.assignment=="bestworst" || fixed.assignment=="worstbest"){
          fixed.assignment.f <- fixed.assignment.f[breeding.size.total:1]
        }
        fixed.breeding <- cbind(best[[1]][fixed.assignment.m,1:3], best[[2]][fixed.assignment.f,1:3], sex.animal)
      }
      if(ogc){
        if(verbose) cat("Inefficient implementation of OGC. Currently not available for large scale datasets.\n")

        animallist <- rbind(cbind(best[[1]],1), cbind(best[[2]],2))
        n.animals <- nrow(animallist)

        if(sum(abs(animallist[,4]))==0){
          if(verbose) cat("No breeding values stored for OGC. Use breeding value of first trait as u!\n")
          for(index in 1:nrow(animallist)){
            animallist[index,4] <- population$breeding[[animallist[index,1]]][[2+animallist[index,2]]][1,animallist[index,3]]
          }
        }

        if(sum(abs(animallist[,4]))==0){
          if(verbose) cat("No breeding value estimated available! Use genomic value of first trait as u!\n")
          for(index in 1:nrow(animallist)){
            animallist[index,4] <- population$breeding[[animallist[index,1]]][[6+animallist[index,2]]][1,animallist[index,3]]
          }
        }

        u <- animallist[,4]
        Q <- cbind((animallist[,6]==1), animallist[,6]==2)

        if(miraculix){
          Z.code <- miraculix::computeSNPS(population, animallist[,1], animallist[,2], animallist[,3], what="geno", output_compressed = TRUE)
        } else{
          Zt <- array(0,dim=c(sum(population$info$snp), n.animals))
          for(index in 1:n.animals){
            Zt[,index] <- colSums(compute.snps(population, animallist[index,1], animallist[index,2], animallist[index,3], import.position.calculation=import.position.calculation, decodeOriginsU=decodeOriginsU, bit.storing=bit.storing, nbits=nbits, output_compressed=FALSE))
          }
        }

        # Verwandtschaftsmatrix:
        if(computation.A.ogc=="kinship"){
          A <- kinship.exp(population, database = animallist[,c(1,2,3,3)], depth.pedigree = depth.pedigree.ogc)
        } else if(computation.A.ogc=="vanRaden"){
          if(miraculix){
            p_i <- miraculix::allele_freq(Z.code) # Noch nicht implementiert?
            A <- miraculix::relationshipMatrix(Z.code, centered=TRUE, normalized=TRUE)

          } else if(miraculix.mult){
            p_i <- rowSums(Zt)/ncol(Zt)/2
            Zt_miraculix <- miraculix::genomicmatrix(Zt)
            A <- miraculix::relationshipMatrix(Zt_miraculix, centered=TRUE, normalized=TRUE)
          } else{
            p_i <- rowSums(Zt)/ncol(Zt)/2
            Ztm <- Zt - p_i * 2
            A <- crossprod(Ztm)/ (2 * sum(p_i*(1-p_i)))
          }

        } else if(computation.A.ogc=="CM"){
          #CM SCHAETZER
          Ztm <- rbind(Zt==0, Zt==1, Zt==2)
          A <- crossprod(Ztm) / ncol(Zt)
        } else if(computation.A.ogc=="CE"){
          Ztm <- rbind(Zt==0, Zt==1, Zt==2)
          A <- crossprod(Ztm)
          A <- (A^2 - 0.5*A)/(nrow(Zt)^2)

        } else if(computation.A.ogc=="non_stand"){
          A <- crossprod(Zt) / nrow(Zt)
        }
        contribution <- OGC(A, u, Q, ogc.cAc, single=TRUE)
        contribution <- list(contribution$`Optimal c`[Q[,1]], contribution$`Optimal c`[Q[,2]])

      }
    }

  } else{
    breeding.size.total <- nrow(fixed.breeding)
    sex.animal <- fixed.breeding[,7] <- stats::rbinom(breeding.size.total, 1, fixed.breeding[,7]) +1
    breeding.size <- c(sum(fixed.breeding[,7]==1), sum(fixed.breeding[,7]==2))
  }

}

  if(gene.editing.best){
    for(sex in (1:2)[gene.editing.best.sex]){
      if(length(best[[sex]])>0){
        for(index in 1:nrow(best[[sex]])){
          activ <- best[[sex]][index,]
          population$breeding[[activ[1]]][[activ[2]]][[activ[3]]] <- edit_animal(population, activ[1], activ[2], activ[3], nr.edits, decodeOriginsU=decodeOriginsU, bit.storing=bit.storing, nbits=nbits)
        }
        for(index in 1:nrow(best[[sex]])){
          activ <- best[[sex]][index,]
          if(length(population$breeding[[activ[1]]][[activ[2]]][[activ[3]]])>=17){
            population$breeding[[activ[1]]][[activ[2]]][[activ[3]]][[17]] <- c(population$breeding[[activ[1]]][[activ[2]]][[activ[3]]][[17]],population$breeding[[activ[1]]][[6+activ[2]]][,activ[3]])
          } else{
            population$breeding[[activ[1]]][[activ[2]]][[activ[3]]][[17]] <- population$breeding[[activ[1]]][[6+activ[2]]][,activ[3]]
          }
          population$breeding[[activ[1]]][[activ[2]]][[activ[3]]][[15]] <- rep(0, population$info$bv.nr)
          activ_bv <- population$info$bv.random.activ
          if(length(activ_bv)>0){
            temp_out <- calculate.bv(population, activ[1], activ[2], activ[3], activ_bv, import.position.calculation=import.position.calculation, decodeOriginsU=decodeOriginsU, bit.storing=bit.storing, nbits=nbits, output_compressed=FALSE)
            population$breeding[[activ[1]]][[6+activ[2]]][activ_bv,activ[3]] <- temp_out[[1]]
          }
          population$breeding[[activ[1]]][[activ[2]]][[activ[3]]][[17]] <- c(population$breeding[[activ[1]]][[activ[2]]][[activ[3]]][[17]] ,population$breeding[[activ[1]]][[6+activ[2]]][,activ[3]])
        }
      }
    }
  }


  # Gezielte Alterung
  if(is.matrix(reduce.group)==0 && length(reduce.group)>0){
    reduce.group <- t(reduce.group)
  }
  if(length(reduce.group)>0){
    reduce.group <- reduce.group[reduce.group[,1]!=0,]
  }
  if(length(reduce.group)>0){
    for(index in 1:nrow(reduce.group)){
      activ.reduce <- reduce.group[index,]
      group.animals <- (population$breeding[[activ.reduce[1]]][[activ.reduce[2]+4]] == activ.reduce[4])
      n.animals <- sum(group.animals)
      to.kill <- n.animals - activ.reduce[3]
      animal.position <- unique(c(0,group.animals * 1:length(group.animals)))[-1]
      if(to.kill>0){
        if(reduce.group.selection=="random"){
          death <- sample(animal.position, to.kill)

        } else if(reduce.group.selection=="selection"){
          if(multiple.bve=="add"){
            print(population$breeding[[activ.reduce[1]]][[activ.reduce[2]+2]])
            bve.sum <- colSums(rbind(population$breeding[[activ.reduce[1]]][[activ.reduce[2]+2]][,animal.position]*multiple.bve.weights[[sex]],0))
          } else if(multiple.bve=="ranking"){
            ranking <- population$breeding[[index]][[sex+2]][,relevant.animals]
            for(bven in 1:population$info$bv.nr){
              order <- sort(ranking[bven,], index.return=TRUE, decreasing=selection.critera[activ.reduce[2]])$ix
              ranking[bven,order] <- length(order):1
            }
            bve.sum <- colSums(ranking*multiple.bve.weights[[sex]])
          }
          chosen.animals <- sort(bve.sum, index.return=TRUE, decreasing=selection.critera[activ.reduce[2]])$ix
          death <- chosen.animals[(length(chosen.animals)-to.kill+1):length(chosen.animals)]
        }
        for(modanimal in death){
          population$breeding[[activ.reduce[1]]][[activ.reduce[2]]][[modanimal]][[18]] <- c(population$breeding[[activ.reduce[1]]][[activ.reduce[2]+4]][modanimal], current.gen)
          population$breeding[[activ.reduce[1]]][[activ.reduce[2]+4]][modanimal] <- -1
        }
      }


    }
  }
  if(store.comp.times){
    comp.times[6] <- as.numeric(Sys.time())
  }

  sample_prob <- list()
  for(sex in 1:2){
    if(length(best[[sex]])>0){
      if(ogc){
        sample_prob[[sex]] <- contribution[[sex]]
      } else if(length(best.selection.manual.ratio[[sex]])==0){
        individual_bv <- best[[sex]][,5]
        min_bv <- min(individual_bv)
        max_bv <- max(individual_bv)
        if(min_bv==max_bv){
          sample_prob[[sex]] <- rep(1, length(individual_bv))
        } else{
          sample_prob[[sex]] <- (individual_bv- min_bv) / (max_bv-min_bv) * (best.selection.ratio[[sex]]-1) + 1
        }
      } else{
        sample_prob[[sex]] <- best.selection.manual.ratio[[sex]]
      }

      if(length(unique(sample_prob[[sex]]))==1){
        sample_prob[[sex]] <- NULL
      }
    }
  }
  sample_prob[[3]] <- "placeholder"


  if(length(fixed.breeding.best)>0){
    fixed.breeding <- matrix(0, nrow=nrow(fixed.breeding.best), ncol=7)

    for(index in 1:nrow(fixed.breeding.best)){
      fixed.breeding[index,1:3] <- best[[fixed.breeding.best[index,1]]][fixed.breeding.best[index,2], 1:3]
      fixed.breeding[index,4:6] <- best[[fixed.breeding.best[index,3]]][fixed.breeding.best[index,4], 1:3]
    }

    fixed.breeding[,7] <- fixed.breeding.best[,5]
    if(breeding.size.total==0){
      breeding.size.total <- nrow(fixed.breeding)
    }
    sex.animal <- fixed.breeding[,7] <- stats::rbinom(breeding.size.total, 1, fixed.breeding[,7]) +1
    breeding.size <- c(sum(fixed.breeding[,7]==1), sum(fixed.breeding[,7]==2))
  }


  # Rekombinationsprozess
  if(length(population$breeding)<=current.gen && sum(breeding.size)>0 ){
    population$breeding[[current.gen+1]] <- list()
    population$info$size <- rbind(population$info$size, 0)
  }

  selection.rate <- list(numeric(selection.size[1]), numeric(selection.size[2]))
  activ.selection.size <- selection.size
  availables.m <- 1:selection.size[1]
  availables.f <- 1:selection.size[2]
  availables <- list(availables.m, availables.f)

  if(relative.selection){
    if(sum(best[[1]][,4]<0)>0){
      best[[1]][(best[[1]][,4]<0),4] <- 0
    }
    if(sum(best[[2]][,4]<0)>0){
      best[[2]][(best[[2]][,4]<0),4] <- 0
    }

    cum.sum.m <- cumsum(best[[1]][,4])
    cum.sum.f <- cumsum(best[[2]][,4])
    sum.m <- sum(best[[1]][,4])
    sum.f <- sum(best[[2]][,4])
    cum.sum <- list(cum.sum.m, cum.sum.f)
    sum <- list(sum.m,sum.f)
  }
  current.size <- c(1,1)
  for(sex in 1:2){
    if(sum(breeding.size)>0){
      if(length(population$breeding[[current.gen+1]])<=2 || length(population$breeding[[current.gen+1]][[sex]])==0){
        population$breeding[[current.gen+1]][[2+sex]] <- matrix(0, nrow=population$info$bv.nr, ncol=breeding.size[sex])
        population$breeding[[current.gen+1]][[4+sex]] <- rep(new.class, breeding.size[sex])
        population$breeding[[current.gen+1]][[6+sex]] <- matrix(0, nrow=population$info$bv.nr, ncol=breeding.size[sex])
        population$breeding[[current.gen+1]][[8+sex]] <- matrix(0, nrow=population$info$bv.nr, ncol=breeding.size[sex])
        population$breeding[[current.gen+1]][[10+sex]] <- rep(time.point, breeding.size[sex])
        population$breeding[[current.gen+1]][[12+sex]] <- rep(creating.type, breeding.size[sex])

        if(copy.individual){
          population$breeding[[current.gen+1]][[14+sex]] <- rep(0, breeding.size[sex])
        } else{
          population$breeding[[current.gen+1]][[14+sex]] <- seq(population$info$next.animal, population$info$next.animal + breeding.size[sex] -1, length.out= breeding.size[sex])
          population$info$next.animal <- population$info$next.animal + breeding.size[sex]
        }
        population$breeding[[current.gen+1]][[16+sex]] <- rep(NA, breeding.size[sex])
        population$breeding[[current.gen+1]][[18+sex]] <- matrix(0, nrow=population$info$bv.nr, ncol=breeding.size[sex])
        population$breeding[[current.gen+1]][[20+sex]] <- matrix(0, nrow=population$info$bv.nr, ncol=breeding.size[sex])

        #    } else if(length(population$breeding[[current.gen+1]][[sex+2]])==0){
        #      population$breeding[[current.gen+1]][[2+sex]] <- rep(0, breeding.size[sex])
        #      population$breeding[[current.gen+1]][[4+sex]] <- rep(new.class, breeding.size[sex])
        #     population$breeding[[current.gen+1]][[6+sex]] <- new.bv[sex,]
        #      population$breeding[[current.gen+1]][[8+sex]] <- new.bv.approx[sex,]
      } else{
        current.size[sex] <- length(population$breeding[[current.gen+1]][[4+sex]]) + 1
        population$breeding[[current.gen+1]][[2+sex]] <- cbind(population$breeding[[current.gen+1]][[sex+2]], matrix(0, nrow= population$info$bv.nr, ncol=breeding.size[sex]))
        population$breeding[[current.gen+1]][[4+sex]] <- c(population$breeding[[current.gen+1]][[sex+4]], rep(new.class, breeding.size[sex]))
        population$breeding[[current.gen+1]][[6+sex]] <- cbind(population$breeding[[current.gen+1]][[6+sex]], matrix(0, nrow= population$info$bv.nr, ncol=breeding.size[sex]))
        population$breeding[[current.gen+1]][[8+sex]] <- cbind(population$breeding[[current.gen+1]][[8+sex]], matrix(0, nrow= population$info$bv.nr, ncol=breeding.size[sex]))
        population$breeding[[current.gen+1]][[10+sex]] <- c(population$breeding[[current.gen+1]][[sex+10]], rep(time.point, breeding.size[sex]))
        population$breeding[[current.gen+1]][[12+sex]] <- c(population$breeding[[current.gen+1]][[sex+12]], rep(creating.type, breeding.size[sex]))
        if(copy.individual){
          population$breeding[[current.gen+1]][[14+sex]] <- c(population$breeding[[current.gen+1]][[14+sex]], rep(0,breeding.size[sex]))
        } else{
          population$breeding[[current.gen+1]][[14+sex]] <- c(population$breeding[[current.gen+1]][[14+sex]], seq(population$info$next.animal, population$info$next.animal + breeding.size[sex] -1, length.out= breeding.size[sex]))
          population$info$next.animal <- population$info$next.animal + breeding.size[sex]
        }
        population$breeding[[current.gen+1]][[16+sex]] <- c(population$breeding[[current.gen+1]][[sex+16]], rep(NA, breeding.size[sex]))
        population$breeding[[current.gen+1]][[18+sex]] <- cbind(population$breeding[[current.gen+1]][[18+sex]], matrix(0, nrow= population$info$bv.nr, ncol=breeding.size[sex]))
        population$breeding[[current.gen+1]][[20+sex]] <- cbind(population$breeding[[current.gen+1]][[20+sex]], matrix(0, nrow= population$info$bv.nr, ncol=breeding.size[sex]))

      }
      if(length(population$breeding[[current.gen+1]][[sex]])==0){
        population$breeding[[current.gen+1]][[sex]] <- list()
      }

    }
  }
  if(length(praeimplantation)>0){
    nsnps <- c(0,population$info$cumsnp)
    praeimplantation <- cbind(praeimplantation, nsnps[praeimplantation[,1]] + praeimplantation[,2])
    praeimplantation.max <- list()
    for(sex in 1:2){
      praeimplantation.max[[sex]] <- numeric(nrow(best[[sex]]))
      for(index in 1:length(best[[sex]])){
        activ <- best[[sex]][index,]
        hap1 <- population$breeding[[activ[1]]][[activ[2]]][[activ[3]]][[9]][praeimplantation[,4]]
        hap2 <- population$breeding[[activ[1]]][[activ[2]]][[activ[3]]][[10]][praeimplantation[,4]]
        pos <- ((hap1==praeimplantation[,3]) + (hap2==praeimplantation[,3]))>0
        praeimplantation.max[[sex]][index] <- sum(pos)
      }
    }
    print("praeimplantation bisher nicht in Selektion enthalten")
  }


##  store.effect.freq and multiple correlated bvs deactivated
  if(store.comp.times.generation){
    pre_stuff <- 0
    generation_stuff <- 0
    bv_stuff <- 0
  }
  if(parallel.generation && breeding.size.total>0){

    if(requireNamespace("doParallel", quietly = TRUE)) {

      if(length(name.cohort)>0){
        if(verbose) cat(paste0("Start generation of new individuals (cohort: ", name.cohort,").\n"))
      } else{
        if(verbose) cat("Start generation of new individuals.\n")
      }


      if(store.comp.times.generation){
        tick <- as.numeric(Sys.time())
      }
      info_father_list <- info_mother_list <- matrix(0, nrow=breeding.size.total, ncol=5)


      runs <- repeat.mating
      for(animal.nr in 1:breeding.size.total){

        sex <- sex.animal[animal.nr]
        if(length(fixed.breeding)>0){
          info.father <- fixed.breeding[animal.nr,1:3]
          info.mother <- fixed.breeding[animal.nr,4:6]
        } else{
          if(runs!=repeat.mating && runs>0){
            runs <- runs - 1
          } else{
            runs <- repeat.mating - 1
            if(selfing.mating==FALSE){
              sex1 <- 1
              sex2 <- 2
              if(same.sex.activ==FALSE && relative.selection==FALSE){

                number1 <- availables[[sex1]][sample(1:activ.selection.size[sex1],1, prob=sample_prob[[sex1]][availables[[sex1]]])]
                number2 <- availables[[sex2]][sample(1:activ.selection.size[sex2],1, prob=sample_prob[[sex2]][availables[[sex2]]])]

                info.father <- best[[sex1]][number1,]
                info.mother <- best[[sex2]][number2,]
              } else if(same.sex.activ==FALSE && relative.selection){
                info.father <- best[[sex1]][sum(cum.sum[[sex1]] <stats::runif(1,0,sum[[1]])) +1 ,1:3]
                info.mother <- best[[sex2]][sum(cum.sum[[sex2]] <stats::runif(1,0,sum[[2]])) +1 ,1:3]
              } else{
                sex1 <- stats::rbinom(1,1,same.sex.sex) + 1 # ungleichviele tiere erhöht

                sex2 <- stats::rbinom(1,1,same.sex.sex) + 1

                number1 <- availables[[sex1]][sample(1:activ.selection.size[sex1],1, prob=sample_prob[[sex1]][availables[[sex1]]])]
                number2 <- availables[[sex2]][sample(1:activ.selection.size[sex2],1, prob=sample_prob[[sex2]][availables[[sex2]]])]
                test <- 1
                while(same.sex.selfing==FALSE && number1==number2 && test < 100){
                  number2 <- availables[[sex2]][sample(1:activ.selection.size[sex2],1, prob=sample_prob[[sex2]][availables[[sex2]]])]
                  test <- test+1
                  if(test==100 && number1==number2){
                    print("Selbstung durchgefuehrt da nur ein Tier in Selektionsgruppe")
                  }
                }

                if(relative.selection==FALSE){
                  info.father <- best[[sex1]][number1,]
                  # waehle fuers bveite Tier ein Tier aus der Gruppe der Nicht-besten Tiere
                  if(martini.selection){
                    options <- (1:population$info$size[best[[sex2]][1,1],sex2])[-best[[sex2]][,3]]
                    number2 <- sample(options,1)
                    info.mother <- c(best[[sex2]][1,1:2], number2)
                  } else{
                    info.mother <- best[[sex2]][number2,]
                  }
                } else{
                  info.father <- best[[sex1]][sum(cum.sum[[sex1]] <stats::runif(1,0,cum.sum[[sex1]])) +1 ,1:3]
                  info.mother <- best[[sex2]][sum(cum.sum[[sex2]] <stats::runif(1,0,cum.sum[[sex2]])) +1 ,1:3]
                }

              }
            } else{
              sex1 <- sex2 <- stats::rbinom(1,1,selfing.sex)+1
              if(length(availables[[sex1]])==0){
                sex1 <- sex2 <- 3 - sex1
              }
              number1 <- number2 <- availables[[sex1]][sample(1:activ.selection.size[sex1],1, prob=sample_prob[[sex1]][availables[[sex1]]])]
              info.father <- best[[sex1]][number1,]
              info.mother <- best[[sex2]][number2,]

            }
            if(martini.selection==FALSE){
              selection.rate[[sex1]][number1] <- selection.rate[[sex1]][number1] + repeat.mating
              selection.rate[[sex2]][number2] <- selection.rate[[sex2]][number2] + repeat.mating

              if(selection.rate[[sex1]][number1] >= max.offspring[sex1]){
                activ.selection.size[sex1] <-  activ.selection.size[sex1] - 1
                availables[[sex1]] <- availables[[sex1]][availables[[sex1]]!=number1]
              }
              if(selection.rate[[sex2]][number2] >= max.offspring[sex2]){
                if(sex1!= sex2 || number1 != number2){
                  activ.selection.size[sex2] <-  activ.selection.size[sex2] -1
                }
                availables[[sex2]] <- availables[[sex2]][availables[[sex2]]!=number2]
              }
            }
          }


        }
        info_father_list[animal.nr,] <- info.father
        info_mother_list[animal.nr,] <- info.mother
      }

      if(store.comp.times.generation){
        tock <- as.numeric(Sys.time())
        pre_stuff <- tock-tick
      }

      if(Sys.info()[['sysname']]=="Windows"){
        doParallel::registerDoParallel(cores=ncore.generation)
        if(length(randomSeed.generation)>0){
          if (requireNamespace("doRNG", quietly = TRUE)) {
            doRNG::registerDoRNG(seed=randomSeed.generation)
          } else{
            stop("Usage of doRNG without being installed!")
          }
        }
      }


      for(sex_running in 1:2){

        if(store.comp.times.generation){
          tick <- as.numeric(Sys.time())
        }

        if(Sys.info()[['sysname']]=="Windows"){
          new_animal <- foreach::foreach(indexb=(1:breeding.size.total)[sex.animal==sex_running],
                                         .packages="MoBPS") %dopar% {

                                           child <- generation.individual(indexb,
                                                                          population, info_father_list, info_mother_list,
                                                                          copy.individual, mutation.rate, remutation.rate, recombination.rate,
                                                                          recom.f.indicator, recom.f.polynom, duplication.rate, duplication.length,
                                                                          duplication.recombination, delete.same.origin,
                                                                          (gene.editing.offspring * gene.editing.offspring.sex[1]), nr.edits,
                                                                          gen.architecture.m, gen.architecture.f, decodeOriginsU,
                                                                          current.gen, save.recombination.history, new.bv.child,
                                                                          dh.mating, share.genotyped, added.genotyped,
                                                                          dh.sex, n.observation)
                                           child

          }
        } else {
          activ_stuff <- (1:breeding.size.total)[sex.animal==sex_running]
          new_animal <- parallel::mclapply(activ_stuff, function(x) generation.individual(x,
                                                                                population, info_father_list, info_mother_list,
                                                                                copy.individual, mutation.rate, remutation.rate, recombination.rate,
                                                                                recom.f.indicator, recom.f.polynom, duplication.rate, duplication.length,
                                                                                duplication.recombination, delete.same.origin,
                                                                                (gene.editing.offspring * gene.editing.offspring.sex[1]), nr.edits,
                                                                                gen.architecture.m, gen.architecture.f, decodeOriginsU,
                                                                                current.gen, save.recombination.history, new.bv.child,
                                                                                dh.mating, share.genotyped, added.genotyped,
                                                                                dh.sex, n.observation),
                                 mc.cores=ncore.generation)
        }


        prev_ani <- length(population$breeding[[current.gen+1]][[sex_running]])
        present_before <- population$info$size[current.gen+1,sex_running]
        population$breeding[[current.gen+1]][[sex_running]] <-  c(population$breeding[[current.gen+1]][[sex_running]], new_animal)
        population$info$size[current.gen+1,sex_running] <- length(population$breeding[[current.gen+1]][[sex_running]])

        if(length(new_animal)>0){
          for(index6 in 1:length(new_animal) + prev_ani){
            if(copy.individual){
              first_copy <- population$breeding[[current.gen+1]][[sex_running]][[index6]][[21]]
              new_copy <- rbind(population$breeding[[first_copy[1,1]]][[first_copy[1,2]]][[first_copy[1,3]]][[21]],
                                c(current.gen+1, sex_running, index6))
              for(index7 in 1:nrow(new_copy)){
                population$breeding[[new_copy[index7,1]]][[new_copy[index7,2]]][[new_copy[index7,3]]][[21]] <- new_copy
              }

            } else{
              population$breeding[[current.gen+1]][[sex_running]][[index6]][[21]] <- cbind(current.gen+1, sex_running, index6)
            }
          }
        }

        if(store.comp.times.generation){
          tack <- as.numeric(Sys.time())
          generation_stuff <- tack-tick + generation_stuff
        }
        if(store.effect.freq){
          store.effect.freq <- FALSE
          cat("Effect-Freq not available in parallel computing.\n")
        }

        index_loop <- NULL
        if(length(new_animal)>0) index_loop <- 1:length(new_animal)

        if(Sys.info()[['sysname']]!="Windows"){
          doParallel::registerDoParallel(cores=ncore.generation)
        }

        new.bv.list <- foreach::foreach(indexb=index_loop,
                              .packages=c("MoBPS", "miraculix")) %dopar% {

                              info.father <- info_father_list[indexb,]
                              info.mother <- info_mother_list[indexb,]
                              new.bv <- new.bv_approx <-  new.bve <- numeric(population$info$bv.nr)
                              activ_bv <- population$info$bv.random.activ

                              if(length(activ_bv)>0){
                                temp_out <- calculate.bv(population, current.gen+1, sex_running, indexb + present_before, activ_bv, import.position.calculation=import.position.calculation, decodeOriginsU=decodeOriginsU, store.effect.freq=store.effect.freq, bit.storing=bit.storing, nbits=nbits, output_compressed=FALSE)
                                new.bv[activ_bv] <- temp_out[[1]]
                                if(store.effect.freq){
                                  if(length(population$info$store.effect.freq) < (current.gen+1) || length(population$info$store.effect.freq[[current.gen+1]])==0){
                                    population$info$store.effect.freq[[current.gen+1]] <- temp_out[[2]]
                                  } else{
                                    population$info$store.effect.freq[[current.gen+1]] <- population$info$store.effect.freq[[current.gen+1]] + temp_out[[2]]
                                  }
                                }
                              }

                              if(population$info$bv.calc > 0  && population$info$bv.random[population$info$bv.calc]){
                                means <- 0.5*(population$breeding[[info.father[1]]][[6+info.father[2]]][[population$info$bv.calc:population$info$bv.nr,info.father[3]]] + population$breeding[[info.mother[1]]][[6+info.mother[2]]][[population$info$bv.calc:population$info$bv.nr,info.mother[3]]])
                                varp <- kinship.emp(list(population$breeding[[info.father[1]]][[info.father[2]]][[info.father[3]]], population$breeding[[info.mother[1]]][[info.mother[2]]][[info.mother[3]]]))
                                varp <- (2 - (2* (varp[1,1]-0.5) + 2 * (varp[2,2]- 0.5)))/4

                                if(FALSE){
                                  new.bv[population$info$bv.calc:population$info$bv.nr] <- stats::rnorm(1, mean=means, sd= sqrt(var*population$info$bv.random.variance[bven]))
                                } else{
                                  if(population$info$bv.calc==1){
                                    population$info$current.bv.random.variance <- varp * population$info$bv.random.variance
                                    bv.var <- diag(sqrt(population$info$current.bv.random.variance)) %*%population$info$current.bv.correlation %*% diag(sqrt(population$info$current.bv.random.variance))
                                    single.mean <- means
                                  } else{
                                    population$info$current.bv.random.variance <- c(population$info$bv.random.variance[1:(population$info$bv.calc-1)],varp * population$info$bv.random.variance[population$info$bv.calc:population$info$bv.nr])
                                    AA <- diag(sqrt(population$info$current.bv.random.variance)[1:(population$info$bv.calc-1)]) %*% population$info$current.bv.correlation[1:(population$info$bv.calc-1), 1:(population$info$bv.calc-1)]%*% diag(sqrt(population$info$current.bv.random.variance)[(1:(population$info$bv.calc-1))])
                                    BB <- diag(sqrt(population$info$current.bv.random.variance)[1:(population$info$bv.calc-1)]) %*%population$info$current.bv.correlation[1:(population$info$bv.calc-1), -(1:(population$info$bv.calc-1))]%*% diag(sqrt(population$info$current.bv.random.variance)[-(1:(population$info$bv.calc-1))])
                                    CC <- diag(sqrt(population$info$current.bv.random.variance)[-(1:(population$info$bv.calc-1))]) %*%population$info$current.bv.correlation[-(1:(population$info$bv.calc-1)), -(1:(population$info$bv.calc-1))] %*% diag(sqrt(population$info$current.bv.random.variance)[-(1:(population$info$bv.calc-1))])
                                    if (requireNamespace("MASS", quietly = TRUE)) {
                                      bv.var <- CC - t(BB) %*% MASS::ginv(AA) %*% BB
                                      single.mean <- means + t(BB) %*% MASS::ginv(AA) %*% ( new.bv[1:(population$info$bv.calc-1)]-mu1[1:(population$info$bv.calc-1)])
                                    } else{
                                      bv.var <- CC - t(BB) %*% solve(AA) %*% BB
                                      single.mean <- means + t(BB) %*% solve(AA) %*% ( new.bv[1:(population$info$bv.calc-1)]-mu1[1:(population$info$bv.calc-1)])
                                    }
                                  }

                                  bv.var_chol <- t(chol(bv.var))
                                  population$info$bv.correlation_col <- bv.var_chol

                                  random_part <- bv.var_chol %*% stats::rnorm(population$info$bv.nr - population$info$bv.calc +1, 0,1 )
                                  means_part <- single.mean
                                  new.bv[population$info$bv.calc:population$info$bv.nr] <-random_part + means_part
                                }

                              }

                              if(new.bv.child=="mean" || new.bv.child=="addobs"){
                                activ_father <- population$breeding[[current.gen+1]][[sex_running]][[indexb + present_before]][[7]]
                                activ_mother <- population$breeding[[current.gen+1]][[sex_running]][[indexb + present_before]][[8]]
                                if(copy.individual){
                                  new.bv_approx <- population$breeding[[info.father[1]]][[8+info.father[2]]][,info.father[3]]
                                } else{
                                  for(bven in 1:population$info$bv.nr){
                                    new.bv_approx[bven] <- mean(c(population$breeding[[activ_father[1]]][[8+activ_father[2]]][bven,activ_father[3]],population$breeding[[activ_mother[1]]][[8+activ_mother[2]]][bven,activ_mother[3]]))
                                  }
                                }

                              }
                              if(new.bv.child=="obs" || new.bv.child=="addobs"){
                                if(sum(n.observation)>0){
                                  if(new.bv.child=="addobs"){
                                    prior_obs <- population$breeding[[info.father[1]]][[info.father[2]]][[info.father[3]]][[15]]
                                    total_obs <- prior_obs + n.observation
                                    new.bv_approx <- (new.bv_approx - population$breeding[[info.father[1]]][[info.father[2]+6]][,info.father[3]]) *  prior_obs / total_obs
                                  } else{
                                    total_obs <- n.observation
                                  }
                                  observation_reps <- sort(unique(c(0,n.observation)))
                                  for(observation_rep in 2:length(observation_reps)){
                                    new.obs <- observation_reps[observation_rep] - observation_reps[observation_rep-1]
                                    temp_random <- matrix(stats::rnorm(population$info$bv.nr*new.obs,0,1), ncol=new.obs)
                                    active.traits <- (n.observation >=observation_reps[observation_rep])
                                    active.traits <- active.traits*(1:length(active.traits))
                                    for(bven in (1:population$info$bv.nr)[active.traits]){
                                      new.bv_approx[bven] <-  new.bv_approx[bven] + new.obs/(total_obs[bven]) * rowMeans(population$info$pheno.correlation %*% temp_random)[bven] * sqrt(sigma.e[bven])
                                    }

                                  }
                                  new.bv_approx <- new.bv + new.bv_approx
                                  new.bv_approx[n.observation==0] <- 0
                                }

                              }

                              new.bve <- population$breeding[[info.father[1]]][[2+info.father[2]]][,info.father[3]]
                              new.reli <- population$breeding[[info.father[1]]][[18+info.father[2]]][,info.father[3]]

                              temp1 <- c(new.bv, new.bv_approx, new.bve, new.reli)
                              temp1
                              }


        if(length(index_loop)>0){
          new.bv.list <- matrix(unlist(new.bv.list), ncol=length(new_animal))
          n_bv <- nrow(new.bv.list) / 4
          population$breeding[[current.gen+1]][[sex_running+6]][,(present_before+1):(present_before+length(new_animal))] <- new.bv.list[1:n_bv,,drop=FALSE]
          population$breeding[[current.gen+1]][[sex_running+8]][,(present_before+1):(present_before+length(new_animal))] <- new.bv.list[1:n_bv+n_bv,,drop=FALSE]
          if(copy.individual && copy.individual.keep.bve){
            population$breeding[[current.gen+1]][[sex_running+2]][,(present_before+1):(present_before+length(new_animal))] <- new.bv.list[1:n_bv+2*n_bv,,drop=FALSE]
            population$breeding[[current.gen+1]][[sex_running+18]][,(present_before+1):(present_before+length(new_animal))] <- new.bv.list[1:n_bv+3*n_bv,,drop=FALSE]
          }
        }

        if(store.comp.times.generation){
          tock <- as.numeric(Sys.time())
          bv_stuff <- tock - tack + bv_stuff
        }
      }

      if(Sys.info()[['sysname']]=="Windows" || TRUE){
        doParallel::stopImplicitCluster()
      }

      if(copy.individual){
        activ_prior <- length(population$breeding[[current.gen+1]][[sex_running]]) - length(new_animal) +1
        for(index in 1:nrow(info_father_list)){
          info.father <- info_father_list[index,]
          population$breeding[[current.gen+1]][[14+sex_running]][activ_prior] <- population$breeding[[info.father[1]]][[info.father[2]+14]][[info.father[3]]]
        }
      }
    } else{
      stop("Usage of doParallel without being installed!")
    }

  } else if(breeding.size.total>0){
    if(length(name.cohort)>0){
      if(verbose) cat(paste0("Start generation of new individuals (cohort: ", name.cohort,").\n"))
    } else{
      if(verbose) cat("Start generation of new individuals.\n")
    }
    if(display.progress){
          pb <- utils::txtProgressBar(min = 0, max = breeding.size.total, style = 3)
    }

    runs <- repeat.mating

    for(animal.nr in 1:breeding.size.total){
      if(store.comp.times.generation){
        tick <- as.numeric(Sys.time())
      }
      sex <- sex.animal[animal.nr]
      new.bv <- new.bv_approx <- new.bve <- new.reli <- individual.id <- numeric(population$info$bv.nr)
      if(length(fixed.breeding)>0){
        info.father <- fixed.breeding[animal.nr,1:3]
        info.mother <- fixed.breeding[animal.nr,4:6]
      } else{
        if(runs!=repeat.mating && runs>0){
          runs <- runs - 1
        } else{
          runs <- repeat.mating - 1
          if(selfing.mating==FALSE){
            sex1 <- 1
            sex2 <- 2
            if(same.sex.activ==FALSE && relative.selection==FALSE){
              number1 <- availables[[sex1]][sample(1:activ.selection.size[sex1],1, prob=sample_prob[[sex1]][availables[[sex1]]])]
              number2 <- availables[[sex2]][sample(1:activ.selection.size[sex2],1, prob=sample_prob[[sex2]][availables[[sex2]]])]
              info.father <- best[[sex1]][number1,]
              info.mother <- best[[sex2]][number2,]
            } else if(same.sex.activ==FALSE && relative.selection){
              info.father <- best[[sex1]][sum(cum.sum[[sex1]] <stats::runif(1,0,sum[[1]])) +1 ,1:3]
              info.mother <- best[[sex2]][sum(cum.sum[[sex2]] <stats::runif(1,0,sum[[2]])) +1 ,1:3]
            } else{
              sex1 <- stats::rbinom(1,1,same.sex.sex) + 1 # ungleichviele tiere erhöht

              sex2 <- stats::rbinom(1,1,same.sex.sex) + 1
              number1 <- availables[[sex1]][sample(1:activ.selection.size[sex1],1, prob=sample_prob[[sex1]][availables[[sex1]]])]
              number2 <- availables[[sex2]][sample(1:activ.selection.size[sex2],1, prob=sample_prob[[sex2]][availables[[sex2]]])]
              test <- 1
              while(same.sex.selfing==FALSE && number1==number2 && test < 100){
                number2 <- availables[[sex2]][sample(1:activ.selection.size[sex2],1, prob=sample_prob[[sex2]][availables[[sex2]]])]
                test <- test+1
                if(test==100 && number1==number2){
                  print("Selbstung durchgefuehrt da nur ein Tier in Selektionsgruppe")
                }
              }

              if(relative.selection==FALSE){
                info.father <- best[[sex1]][number1,]
                # waehle fuers bveite Tier ein Tier aus der Gruppe der Nicht-besten Tiere
                if(martini.selection){
                  options <- (1:population$info$size[best[[sex2]][1,1],sex2])[-best[[sex2]][,3]]
                  number2 <- sample(options,1)
                  info.mother <- c(best[[sex2]][1,1:2], number2)
                } else{
                  info.mother <- best[[sex2]][number2,]
                }
              } else{
                info.father <- best[[sex1]][sum(cum.sum[[sex1]] <stats::runif(1,0,cum.sum[[sex1]])) +1 ,1:3]
                info.mother <- best[[sex2]][sum(cum.sum[[sex2]] <stats::runif(1,0,cum.sum[[sex2]])) +1 ,1:3]
              }

            }
          } else{
            sex1 <- sex2 <- stats::rbinom(1,1,selfing.sex)+1
            if(length(availables[[sex1]])==0){
              if(verbose) cat(paste0("No ", if(sex1==1){"male"} else{"female"}, " individuals left. Use other sex individuals"))
              sex1 <- sex2 <- 3 - sex1
            }
            number1 <- number2 <- availables[[sex1]][sample(1:activ.selection.size[sex1],1, prob=sample_prob[[sex1]][availables[[sex1]]])]
            info.father <- best[[sex1]][number1,]
            info.mother <- best[[sex2]][number2,]

          }
          if(martini.selection==FALSE){
            selection.rate[[sex1]][number1] <- selection.rate[[sex1]][number1] + repeat.mating
            selection.rate[[sex2]][number2] <- selection.rate[[sex2]][number2] + repeat.mating

            if(selection.rate[[sex1]][number1] >= max.offspring[sex1]){
              activ.selection.size[sex1] <-  activ.selection.size[sex1] -1
              availables[[sex1]] <- availables[[sex1]][availables[[sex1]]!=number1]
            }
            if(selection.rate[[sex2]][number2] >= max.offspring[sex2]){
              if(sex1!= sex2 || number1 != number2){
                activ.selection.size[sex2] <-  activ.selection.size[sex2] -1
              }
              availables[[sex2]] <- availables[[sex2]][availables[[sex2]]!=number2]
            }
          }
        }


      }
      if(store.comp.times.generation){
        tack <- as.numeric(Sys.time())
        pre_stuff <- pre_stuff + tack -tick
      }

      father <- population$breeding[[info.father[1]]][[info.father[2]]][[info.father[3]]]
      mother <- population$breeding[[info.mother[1]]][[info.mother[2]]][[info.mother[3]]]
      if(copy.individual){
        info.mother <- info.father
        child1 <- list(father[[1]], father[[3]], father[[5]], father[[7]], father[[11]], 0, if(length(father)>19){father[[19]]} else{0})
        child2 <- list(father[[2]], father[[4]], father[[6]], father[[8]], father[[12]], 0, if(length(father)>19){father[[20]]} else{0})
      } else{
        child1 <- breeding.intern(info.father, father, population,
                                  mutation.rate, remutation.rate, recombination.rate,
                                  recom.f.indicator, recom.f.polynom, duplication.rate, duplication.length,
                                  duplication.recombination, delete.same.origin=delete.same.origin,
                                  gene.editing=(gene.editing.offspring*gene.editing.offspring.sex[1]), nr.edits= nr.edits,
                                  gen.architecture=gen.architecture.m,
                                  decodeOriginsU=decodeOriginsU)

        child2 <- breeding.intern(info.mother, mother, population,
                                  mutation.rate, remutation.rate, recombination.rate,
                                  recom.f.indicator, recom.f.polynom, duplication.rate, duplication.length,
                                  duplication.recombination, delete.same.origin=delete.same.origin,
                                  gene.editing=(gene.editing.offspring * gene.editing.offspring.sex[1]) , nr.edits= nr.edits,
                                  gen.architecture=gen.architecture.f,
                                  decodeOriginsU=decodeOriginsU)
      }
      if(dh.mating){
        if(stats::rbinom(1,1,dh.sex)==0){
          child2 <- child1
        } else{
          child1 <- child2
        }
      }

      # Fuer Praeimplantationsdiagnostik muesste hier die relevanten SNPs berechnet werden.

      if(length(praeimplantation)>0){
        # sex1/sex2 und number1/number2 werden von Selektion uebernommen und nicht bestimmt!
        # Praeimplantationsdiagnostik 1. Chromosom
        counter <- 1
        good1 <- 0
        n.snps <- sum(population$info$snp)
        while(good1==0){
          hap1 <- rep(0,n.snps)
          temp1 <- 0
          current.animal <- child1
          for(index2 in 1:(length(current.animal[[1]])-1)){
            relevant.snp <- (population$info$snp.position < current.animal[[1]][index2+1])*(population$info$snp.position >= current.animal[[1]][index2])*(1:n.snps)
            ursprung <- decodeOriginsU(current.animal[[3]],index2)
            ursprung[1] <- population$info$origin.gen[ursprung[1]]
            hap1[relevant.snp] <-population$breeding[[ursprung[1]]][[ursprung[2]]][[ursprung[3]]][[ursprung[4]+8]][relevant.snp]
          }
          if(length(current.animal[[2]])>0){
            for(index2 in 1:length(current.animal[[2]])){
              position <- which(population$info$snp.position==current.animal[[2]][index2])
              hap1[position] <- 1-population$breeding[[current.gen+1]][[sex]][[current.size[sex]]][[9+temp1]][position]
            }
          }
          if(hap1[pos] == praeimplantation.max[[sex1]][[number1]] || counter==25){
            good1 <- 1
            if(counter==25){print("Praeimplantation gescheitert!")}
          } else{
            child1 <- breeding.intern(info.father, father, population,
                                      mutation.rate, remutation.rate, recombination.rate,
                                      recom.f.indicator, recom.f.polynom, duplication.rate, duplication.length,
                                      duplication.recombination, delete.same.origin=delete.same.origin,
                                      gene.editing=gene.editing, nr.edits= nr.edits,gen.architecture=gen.architecture.m,
                                      decodeOriginsU=decodeOriginsU)
            counter <- counter +1
          }
        }

        # Praeimplantationsdiagnostik 2. Chromosom

        good1 <- 0
        n.snps <- sum(population$info$snp)
        while(good1==0){
          hap1 <- rep(0,n.snps)
          temp1 <- 0
          current.animal <- child2
          for(index2 in 1:(length(current.animal[[1]])-1)){
            relevant.snp <- (population$info$snp.position < current.animal[[1]][index2+1])*(population$info$snp.position >= current.animal[[1]][index2])*(1:n.snps)
            ursprung <-  decodeOriginsU(current.animal[[3]],index2)
            ursprung[1] <- population$info$origin.gen[ursprung[1]]
            hap1[relevant.snp] <-population$breeding[[ursprung[1]]][[ursprung[2]]][[ursprung[3]]][[ursprung[4]+8]][relevant.snp]
          }
          if(length(current.animal[[2]])>0){
            for(index2 in 1:length(current.animal[[2]])){
              position <- which(population$info$snp.position==current.animal[[2]][index2])
              hap1[position] <- 1-population$breeding[[current.gen+1]][[sex]][[current.size[sex]]][[9+temp1]][position]
            }
          }
          if(hap1[pos] == praeimplantation.max[[sex2]][[number2]] || counter==25){
            good1 <- 1
            if(counter==25){print("Praeimplantation gescheitert!")}
          } else{
            child2 <- breeding.intern(info.mother, mother, population,
                                      mutation.rate, remutation.rate, recombination.rate,
                                      recom.f.indicator, recom.f.polynom, duplication.rate, duplication.length,
                                      duplication.recombination, delete.same.origin=delete.same.origin,
                                      gene.editing=gene.editing, nr.edits= nr.edits,
                                      gen.architecture=gen.architecture.f, decodeOriginsU=decodeOriginsU)
            counter <- counter +1
          }
        }
      }


      population$breeding[[current.gen+1]][[sex]][[current.size[sex]]] <- list()
      population$breeding[[current.gen+1]][[sex]][[current.size[sex]]][[1]] <- child1[[1]]
      population$breeding[[current.gen+1]][[sex]][[current.size[sex]]][[2]] <- child2[[1]]
      population$breeding[[current.gen+1]][[sex]][[current.size[sex]]][[3]] <- child1[[2]]
      population$breeding[[current.gen+1]][[sex]][[current.size[sex]]][[4]] <- child2[[2]]
      population$breeding[[current.gen+1]][[sex]][[current.size[sex]]][[5]] <- child1[[3]]
      population$breeding[[current.gen+1]][[sex]][[current.size[sex]]][[6]] <- child2[[3]]
      population$breeding[[current.gen+1]][[sex]][[current.size[sex]]][[7]] <- child1[[4]]
      population$breeding[[current.gen+1]][[sex]][[current.size[sex]]][[8]] <- child2[[4]]

      population$info$size[current.gen+1 ,sex] <- population$info$size[current.gen+1,sex] + 1

      if(is.vector(child1[[5]])){
        population$breeding[[current.gen+1]][[sex]][[current.size[sex]]][[11]] <- t(as.matrix(child1[[5]]))
      } else{
        population$breeding[[current.gen+1]][[sex]][[current.size[sex]]][[11]] <- child1[[5]]
      }
      if(is.vector(child2[[5]])){
        population$breeding[[current.gen+1]][[sex]][[current.size[sex]]][[12]] <- t(as.matrix(child2[[5]]))
      } else{
        population$breeding[[current.gen+1]][[sex]][[current.size[sex]]][[12]] <- child2[[5]]
      }
      if(save.recombination.history && current.gen==1){
        if(length(child1[[6]][-c(1,length(child1[[6]]))])>0){
          population$breeding[[current.gen+1]][[sex]][[current.size[sex]]][[13]] <- cbind(current.gen, child1[[6]][-c(1,length(child1[[6]]))])
        } else{
          population$breeding[[current.gen+1]][[sex]][[current.size[sex]]][[13]] <- cbind(0,0)
        }
        if(length( child2[[6]][-c(1,length(child2[[6]]))])>0){
          population$breeding[[current.gen+1]][[sex]][[current.size[sex]]][[14]] <- cbind(current.gen, child2[[6]][-c(1,length(child2[[6]]))])
        } else{
          population$breeding[[current.gen+1]][[sex]][[current.size[sex]]][[14]] <- cbind(0,0)
        }

      } else if(save.recombination.history && current.gen>1){
        if(length(child1[[6]][-c(1,length(child1[[6]]))])>0){
          population$breeding[[current.gen+1]][[sex]][[current.size[sex]]][[13]] <- rbind(population$breeding[[info.father[1]]][[info.father[2]]][[info.father[3]]][[13]], population$breeding[[info.father[1]]][[info.father[2]]][[info.father[3]]][[14]], cbind(current.gen, child1[[6]][-c(1,length(child1[[6]]))]))
        } else{
          population$breeding[[current.gen+1]][[sex]][[current.size[sex]]][[13]] <- rbind(population$breeding[[info.father[1]]][[info.father[2]]][[info.father[3]]][[13]], population$breeding[[info.father[1]]][[info.father[2]]][[info.father[3]]][[14]])

        }
        if(length( child2[[6]][-c(1,length(child2[[6]]))])>0){
          population$breeding[[current.gen+1]][[sex]][[current.size[sex]]][[14]] <- rbind(population$breeding[[info.mother[1]]][[info.mother[2]]][[info.mother[3]]][[13]], population$breeding[[info.mother[1]]][[info.mother[2]]][[info.mother[3]]][[14]], cbind(current.gen, child2[[6]][-c(1,length(child2[[6]]))]))
        } else{
          population$breeding[[current.gen+1]][[sex]][[current.size[sex]]][[14]] <- rbind(population$breeding[[info.mother[1]]][[info.mother[2]]][[info.mother[3]]][[13]], population$breeding[[info.mother[1]]][[info.mother[2]]][[info.mother[3]]][[14]])

        }

      } else{
        #population$breeding[[current.gen+1]][[sex]][[current.size[sex]]][[13]] <- "test"
      }

      if(new.bv.child=="obs"){
        population$breeding[[current.gen+1]][[sex]][[current.size[sex]]][[15]] <- n.observation
      } else if(new.bv.child=="addobs"){
        population$breeding[[current.gen+1]][[sex]][[current.size[sex]]][[15]] <- n.observation +
          population$breeding[[info.mother[1]]][[info.mother[2]]][[info.mother[3]]][[15]]/2   +
          population$breeding[[info.father[1]]][[info.father[2]]][[info.father[3]]][[15]]/2
      } else{
        population$breeding[[current.gen+1]][[sex]][[current.size[sex]]][[15]] <- rep(0, population$info$bv.nr)
      }

      if(copy.individual){
        population$breeding[[current.gen+1]][[sex]][[current.size[sex]]][[16]] <- population$breeding[[info.father[1]]][[info.father[2]]][[info.father[3]]][[16]]
        if(added.genotyped>0 && population$breeding[[current.gen+1]][[sex]][[current.size[sex]]][[16]]==0){
          population$breeding[[current.gen+1]][[sex]][[current.size[sex]]][[16]] <- stats::rbinom(1,1,added.genotyped)
        }

        first_copy <- population$breeding[[info.father[1]]][[info.father[2]]][[info.father[3]]][[21]]
        new_copy <- rbind(population$breeding[[first_copy[1,1]]][[first_copy[1,2]]][[first_copy[1,3]]][[21]],
                          c(current.gen+1, sex, current.size[sex]))
        for(index7 in 1:nrow(new_copy)){
          population$breeding[[new_copy[index7,1]]][[new_copy[index7,2]]][[new_copy[index7,3]]][[21]] <- new_copy
        }
      } else{
        population$breeding[[current.gen+1]][[sex]][[current.size[sex]]][[16]] <- stats::rbinom(1,1,share.genotyped)
        population$breeding[[current.gen+1]][[sex]][[current.size[sex]]][[21]] <- cbind(current.gen+1, sex, current.size[sex])
      }
      if(length(child1[[7]])>0){
        population$breeding[[current.gen+1]][[sex]][[current.size[sex]]][[19]] <- child1[[7]]
      }
      if(length(child2[[7]])>0){
        population$breeding[[current.gen+1]][[sex]][[current.size[sex]]][[20]] <- child2[[7]]
      }


      if(store.comp.times.generation){
        tock <- as.numeric(Sys.time())
        generation_stuff <- generation_stuff + tock -tack
      }
      if(population$info$bve){
        activ_bv <- population$info$bv.random.activ
        if(length(activ_bv)>0){
          temp_out <- calculate.bv(population, current.gen+1, sex, current.size[sex], activ_bv, import.position.calculation=import.position.calculation, decodeOriginsU=decodeOriginsU, store.effect.freq=store.effect.freq, bit.storing=bit.storing, nbits=nbits, output_compressed=FALSE)
          new.bv[activ_bv] <- temp_out[[1]]
          if(store.effect.freq){
            if(length(population$info$store.effect.freq) < (current.gen+1) || length(population$info$store.effect.freq[[current.gen+1]])==0){
              population$info$store.effect.freq[[current.gen+1]] <- temp_out[[2]]
            } else{
              population$info$store.effect.freq[[current.gen+1]] <- population$info$store.effect.freq[[current.gen+1]] + temp_out[[2]]
            }
          }
        }

        if(population$info$bv.calc > 0  && population$info$bv.random[population$info$bv.calc]){

          #Means passt (Korrelation exakt wie gewünscht)
          means <- 0.5*(population$breeding[[info.father[1]]][[6+info.father[2]]][population$info$bv.calc:population$info$bv.nr,info.father[3]] + population$breeding[[info.mother[1]]][[6+info.mother[2]]][population$info$bv.calc:population$info$bv.nr,info.mother[3]])

          # Berechnung i (0-0.5)
          varp <- kinship.emp(list(population$breeding[[info.father[1]]][[info.father[2]]][[info.father[3]]], population$breeding[[info.mother[1]]][[info.mother[2]]][[info.mother[3]]]))
          varp <- (2 - (2* (varp[1,1]-0.5) + 2 * (varp[2,2]- 0.5)))/4

          if(FALSE){
            new.bv[population$info$bv.calc:population$info$bv.nr] <- stats::rnorm(1, mean=means, sd= sqrt(var*population$info$bv.random.variance[bven]))
          } else{
            if(population$info$bv.calc==1){
              population$info$current.bv.random.variance <- varp * population$info$bv.random.variance
              bv.var <- diag(sqrt(population$info$current.bv.random.variance)) %*%population$info$current.bv.correlation %*% diag(sqrt(population$info$current.bv.random.variance))
              single.mean <- means
            } else{
              #population$info$current.bv.random.variance <- c(population$info$bv.random.variance[1:(population$info$bv.calc-1)], population$info$bv.random.variance[population$info$bv.calc:population$info$bv.nr])
              population$info$current.bv.random.variance <- c(population$info$bv.random.variance[1:(population$info$bv.calc-1)],varp * population$info$bv.random.variance[population$info$bv.calc:population$info$bv.nr])

              AA <- diag(sqrt(population$info$current.bv.random.variance)[1:(population$info$bv.calc-1)]) %*% population$info$current.bv.correlation[1:(population$info$bv.calc-1), 1:(population$info$bv.calc-1)]%*% diag(sqrt(population$info$current.bv.random.variance)[(1:(population$info$bv.calc-1))])
              BB <- diag(sqrt(population$info$current.bv.random.variance)[1:(population$info$bv.calc-1)]) %*%population$info$current.bv.correlation[1:(population$info$bv.calc-1), -(1:(population$info$bv.calc-1))]%*% diag(sqrt(population$info$current.bv.random.variance)[-(1:(population$info$bv.calc-1))])
              CC <- diag(sqrt(population$info$current.bv.random.variance)[-(1:(population$info$bv.calc-1))]) %*%population$info$current.bv.correlation[-(1:(population$info$bv.calc-1)), -(1:(population$info$bv.calc-1))] %*% diag(sqrt(population$info$current.bv.random.variance)[-(1:(population$info$bv.calc-1))])
              if (requireNamespace("MASS", quietly = TRUE)) {
                bv.var <- CC - t(BB) %*% MASS::ginv(AA) %*% BB
                single.mean <- means + t(BB) %*% MASS::ginv(AA) %*% ( new.bv[1:(population$info$bv.calc-1)]-mu1[1:(population$info$bv.calc-1)])
              } else{
                bv.var <- CC - t(BB) %*% solve(AA) %*% BB
                single.mean <- means + t(BB) %*% solve(AA) %*% ( new.bv[1:(population$info$bv.calc-1)]-mu1[1:(population$info$bv.calc-1)])
              }


            }

            bv.var_chol <- t(chol(bv.var))
            population$info$bv.correlation_col <- bv.var_chol

#            random_part <- varp * bv.var_chol %*% stats:rnorm(population$info$bv.nr - population$info$bv.calc +1, 0,1 )
            random_part <- bv.var_chol %*% stats::rnorm(population$info$bv.nr - population$info$bv.calc +1, 0,1 )
            means_part <- single.mean


            new.bv[population$info$bv.calc:population$info$bv.nr] <-   single.mean  + random_part
            #new.bv[population$info$bv.calc:population$info$bv.nr] <- bv.var_chol %*% stats::rnorm(population$info$bv.nr - population$info$bv.calc +1, 0,1 ) + single.mean
          }


        }

        if(copy.individual && copy.individual.keep.bve){
          new.bve <- population$breeding[[info.father[1]]][[2+info.father[2]]][,info.father[3]]
          new.reli <- population$breeding[[info.father[1]]][[18+info.father[2]]][,info.father[3]]
        }
        if(copy.individual){
          individual.id <- population$breeding[[info.father[1]]][[14+info.father[2]]][info.father[3]]
        }
        if(new.bv.child=="mean" || new.bv.child=="addobs"){
          if(copy.individual){
            new.bv_approx <- population$breeding[[info.father[1]]][[8+info.father[2]]][,info.father[3]]
          } else{
            new.bv_approx <- 0.5 * (population$breeding[[info.father[1]]][[8+info.father[2]]][,info.father[3]] + population$breeding[[info.mother[1]]][[8+info.mother[2]]][,info.mother[3]])
          }

        }
        if(new.bv.child=="obs" || new.bv.child=="addobs"){
          if(sum(n.observation)>0){
            if(new.bv.child=="addobs"){
              prior_obs <- population$breeding[[info.father[1]]][[info.father[2]]][[info.father[3]]][[15]]
              total_obs <- prior_obs + n.observation
              new.bv_approx <- (new.bv_approx - population$breeding[[info.father[1]]][[info.father[2]+6]][,info.father[3]]) *  prior_obs / total_obs
            } else{
              total_obs <- n.observation
            }
            observation_reps <- sort(unique(c(0,n.observation)))
            for(observation_rep in 2:length(observation_reps)){
              new.obs <- observation_reps[observation_rep] - observation_reps[observation_rep-1]
              temp_random <- matrix(stats::rnorm(population$info$bv.nr*new.obs,0,1), ncol=new.obs)
              active.traits <- (n.observation >=observation_reps[observation_rep])
              active.traits <- active.traits*(1:length(active.traits))
              for(bven in (1:population$info$bv.nr)[active.traits]){
                new.bv_approx[bven] <-  new.bv_approx[bven] + new.obs/(total_obs[bven]) * rowMeans(population$info$pheno.correlation %*% temp_random)[bven] * sqrt(sigma.e[bven])
              }

            }
            new.bv_approx <- new.bv + new.bv_approx
            new.bv_approx[total_obs==0] <- 0
          }


        }
      }

      population$breeding[[current.gen+1]][[2+sex]][,current.size[sex]] <- new.bve
      population$breeding[[current.gen+1]][[6+sex]][,current.size[sex]] <- new.bv
      population$breeding[[current.gen+1]][[8+sex]][,current.size[sex]] <- new.bv_approx
      population$breeding[[current.gen+1]][[18+sex]][,current.size[sex]] <- new.reli
      if(copy.individual){
        population$breeding[[current.gen+1]][[14+sex]][current.size[sex]] <- individual.id
      }
      #    } else if(length(population$breeding[[current.gen+1]][[sex+2]])==0){
      #      population$breeding[[current.gen+1]][[2+sex]] <- rep(0, breeding.size[sex])
      #      population$breeding[[current.gen+1]][[4+sex]] <- rep(new.class, breeding.size[sex])
      #     population$breeding[[current.gen+1]][[6+sex]] <- new.bv[sex,]
      #      population$breeding[[current.gen+1]][[8+sex]] <- new.bv.approx[sex,]

      if(store.comp.times.generation){
        tock2 <- as.numeric(Sys.time())
        bv_stuff <- bv_stuff+tock2-tock
      }
      if(display.progress){
        utils::setTxtProgressBar(pb, animal.nr)
      }

      current.size[sex] <- current.size[sex] +1
    }
    if(display.progress){
      close(pb)
    }

  }

  delete.haplotypes <- delete.haplotypes[delete.haplotypes>1]
  if(length(delete.haplotypes)>0){
    for(gen_r in delete.haplotypes){
      for(sex in 1:2){
        n.animals <- length(population$breeding[[gen_r]][[sex]])
        if(n.animals>0){
          for(index2 in 1:n.animals){
            population$breeding[[gen_r]][[sex]][[index2]][[9]] <- "removed"
            population$breeding[[gen_r]][[sex]][[index2]][[10]] <- "removed"
          }
        }
      }
    }
  }
  delete.individuals <- delete.individuals[delete.individuals>1]
  if(length(delete.individuals)>0){
    for(gen_r in delete.individuals){
      for(sex in delete.sex){
        n.animals <- length(population$breeding[[gen_r]][[sex]])
        if(n.animals>0){
          population$breeding[[gen_r]][[sex]]<- "removed"

        }
      }
    }
  }
  if(store.breeding.totals){
    cur <- length(population$info$breeding.totals) + 1L
    population$info$breeding.totals[[cur]] <- list()
    population$info$breeding.totals[[cur]][[1]] <- current.gen +1L
    population$info$breeding.totals[[cur]][[2]] <- breeding.size
    population$info$breeding.totals[[cur]][[3]] <- best[[1]]
    population$info$breeding.totals[[cur]][[4]] <- best[[2]]
    population$info$breeding.totals[[cur]][[5]] <- selection.rate[[1]]
    population$info$breeding.totals[[cur]][[6]] <- selection.rate[[2]]
    population$info$breeding.totals[[cur]][[7]] <- chosen.animals.list
  }

  if(store.bve.data && bve){
    cur <- length(population$info$bve.data) + 1
    population$info$bve.data[[cur]] <- list()
    population$info$bve.data[[cur]][[1]] <- current.gen
    population$info$bve.data[[cur]][[2]] <- sigma.e
    population$info$bve.data[[cur]][[3]] <- sigma.e.hat
    population$info$bve.data[[cur]][[4]] <- sigma.g
    population$info$bve.data[[cur]][[5]] <- sigma.a.hat
    population$info$bve.data[[cur]][[6]] <- bve.database
    population$info$bve.data[[cur]][[7]] <- new.bv.observation
    population$info$bve.data[[cur]][[8]] <- y_real
    population$info$bve.data[[cur]][[9]] <- y_hat
    population$info$bve.data[[cur]][[10]] <- y
  }

  population$info$last.sigma.e.database <- sigma.e.database
  population$info$last.sigma.e.value <- sigma.e
  if(length(heritability)>0){
    population$info$last.sigma.e.heritability <- heritability
  }


  if(store.comp.times){
    comp.times[7] <- as.numeric(Sys.time())
    comp.times <- c(comp.times[-1] - comp.times[-length(comp.times)], comp.times[length(comp.times)]-comp.times[1])
    population$info$comp.times <- round(rbind(population$info$comp.times, comp.times, deparse.level = 0), digits=4)
    if(nrow(population$info$comp.times)==1){
      colnames(population$info$comp.times) <- c("preparation", "new real BV", "phenotypes", "BVE","selection","generate new individuals","total")
    }
  }
  if(store.comp.times.bve){
    comp.times.bve <- c(comp.times.bve[-1] - comp.times.bve[-length(comp.times.bve)], zcalc, z_chol, z_uhat, z_ped, z_h, comp.times.bve[length(comp.times.bve)]-comp.times.bve[1])
    population$info$comp.times.bve <- round(rbind(population$info$comp.times.bve, comp.times.bve, deparse.level = 0), digits=4)
    if(nrow(population$info$comp.times.bve)==1){
      colnames(population$info$comp.times.bve) <- c("y_z_import", "A genomic", "solveMixed","Gwas_stuff", "Derive Z", "A inversion", "rrBlup", "A-Pedigree","SingleStep H", "Total")
    }
  }
  if(store.comp.times.generation){
    comp.times.generation <- c(pre_stuff, generation_stuff, bv_stuff, sum(pre_stuff, generation_stuff, bv_stuff))
    population$info$comp.times.generation <- round(rbind(population$info$comp.times.generation, comp.times.generation, deparse.level = 0), digits=4)
    if(nrow(population$info$comp.times.generation)==1){
      colnames(population$info$comp.times.generation) <- c("Preparation", "Generation", "BV-Calculation", "Total")
    }
  }

  if(length(name.cohort)>0){
    if(breeding.size[1] > 0 && breeding.size[2]>0){
      population$info$cohorts <- rbind(population$info$cohorts, c(paste0(name.cohort, "_M"), current.gen+1, breeding.size[1],0, new.class, (current.size-breeding.size)[1], 0,
                                                                  time.point, creating.type),
                                       c(paste0(name.cohort, "_F"), current.gen+1, 0, breeding.size[2], new.class, 0, (current.size-breeding.size)[2],time.point, creating.type))
      print("Added _M, _F to cohort names!")
      rownames(population$info$cohorts)[(nrow(population$info$cohorts)-1):nrow(population$info$cohorts)] <- paste0(name.cohort, c("_M", c("_F")))
    } else{
      population$info$cohorts <- rbind(population$info$cohorts, c(name.cohort, current.gen+1, breeding.size[1:2], new.class, current.size-breeding.size,
                                                                  time.point, creating.type))
      rownames(population$info$cohorts)[nrow(population$info$cohorts)] <- paste0(name.cohort)
    }
    if(nrow(population$info$cohorts)<=2){
      colnames(population$info$cohorts) <- c("name","generation", "male individuals", "female individuals", "class", "position first male", "position first female",
                                             "time point", "creating.type")    }
  }
  if(Rprof){
    Rprof(NULL)
    population$info$Rprof[[length(population$info$Rprof)+1]] <- utils::summaryRprof()
  }
  return(population)
}
```
