## Supplementary material for "MoBPS - Modular Breeding Program Simulator": MoBPS R-package: breeding.intern.R

```
'#
  Authors
Torsten Pook,

Copyright (C) 2017 -- 2018  Torsten Pook

This program is free software; you can redistribute it and/or
modify it under the terms of the GNU General Public License
as published by the Free Software Foundation; either version 3
of the License, or (at your option) any later version.

This program is distributed in the hope that it will be useful,
but WITHOUT ANY WARRANTY; without even the implied warranty of
MERCHANTABILITY or FITNESS FOR A PARTICULAR PURPOSE.  See the
GNU General Public License for more details.

You should have received a copy of the GNU General Public License
along with this program; if not, write to the Free Software
Foundation, Inc., 59 Temple Place - Suite 330, Boston, MA  02111-1307, USA.
'#

#' Internal function to simulate one meiosis
#'
#' Internal function to simulate one meiosis
#' @param info.parent position of the parent in the dataset
#' @param parent list of information regarding the parent
#' @param population Population list
#' @param mutation.rate Mutation rate in each marker (default: 10^-5)
#' @param remutation.rate Remutation rate in each marker (default: 10^-5)
#' @param recombination.rate Average number of recombination per 1 length unit (default: 1M)
#' @param recom.f.indicator Use step function for recombination map (transform snp.positions if possible instead)
#' @param recom.f.polynom Polynomical function to determine expected number of recombinations (transform snp.positions if possible instead)
#' @param duplication.rate Share of recombination points with a duplication (default: 0 - DEACTIVATED)
#' @param duplication.length Average length of a duplication (Exponentially distributed)
#' @param duplication.recombination Average number of recombinations per 1 length uit of duplication (default: 1)
#' @param gene.editing If TRUE perform gene editing on newly generated individual
#' @param gen.architecture Used underlying genetic architecture (genome length in M)
#' @param nr.edits Number of edits to perform per individual
#' @param decodeOriginsU Used function for the decoding of genetic origins [[5]]/[[6]]
#' @param delete.same.origin If TRUE delete recombination points when genetic origin of adjacent segments is the same
#' @export

breeding.intern <- function(info.parent, parent,  population , mutation.rate, remutation.rate, recombination.rate,
                            recom.f.indicator, recom.f.polynom, duplication.rate, duplication.length,
                            duplication.recombination, delete.same.origin=FALSE,
                            gene.editing=gene.editing, nr.edits= nr.edits,
                            gen.architecture=0,
                            decodeOriginsU=decodeOriginsR){
  n_snps <- sum(population$info$snp)
  if(gen.architecture==0){
    length.total <- population$info$length.total
  } else{
    length.total <- population$info$gen.architecture[[gen.architecture]]$length.total

    # Transform points of recombination according to position
    for(haplo in 1:2){
      for(index in 1:length(parent[[haplo]])){
#        chromo <- find.chromo(parent[[haplo]][index], population$info$length.total)
        before <- find.snpbefore(parent[[haplo]][index], population$info$snp.position)
        if(before>0){
          p_before <- population$info$snp.position[before]
          new_p_before <- population$info$gen.architecture[[gen.architecture]]$snp.position[before]
        } else{
          p_before <- population$info$length.total[1]
          new_p_before <-length.total[1]
        }
        if(before<n_snps){
          p_after <- population$info$snp.position[before+1]
          new_p_after <- population$info$gen.architecture[[gen.architecture]]$snp.position[before+1]
        } else{
          p_after <- population$info$length.total[length(population$info$length.total)]
          new_p_after <- length.total[length(length.total)]
        }

        share <- (parent[[haplo]][index]-p_before) / (p_after-p_before)
        parent[[haplo]][index] <- new_p_before + share * (new_p_after-new_p_before)
      }
    }
  }

  n.chromosome <- length(length.total)-1

  # Außerhalb von breeding.intern berechnen
  if(length(recom.f.indicator)!=0){
    # Für Polynom Numerische Bestimmung anstrengend?
    recom.f.indicator <- rbind(recom.f.indicator, c(length.total[n.chromosome+1],0))
    indicator.vol <- sum((recom.f.indicator[-1,1] -recom.f.indicator[-nrow(recom.f.indicator),1])*recom.f.indicator[-nrow(recom.f.indicator),2])
#    polynom.vol <- cumprod(c(1,rep(length.total[n.chromosome+1], length(recom.f.polynom)))) *c(0,recom.f.polynom)
#    polynom.vol <- sum(polynom.vol * c(0,(1/(1:(length(recom.f.polynom))))))
    recom.vol <- indicator.vol #+ polynom.vol
  } else{
    recom.vol <- length.total[n.chromosome+1]*recombination.rate
  }

  noc <- stats::rpois(1, recom.vol) #Anzahl Rekombinationspunkte
  if(length(recom.f.indicator)!=0){
    porc <- stats::runif(noc,0,recom.vol)
    if(length(porc)>0){
      cums <- cumsum((recom.f.indicator[-1,1] -recom.f.indicator[-nrow(recom.f.indicator),1])*recom.f.indicator[-nrow(recom.f.indicator),2])
      for(index in 1:length(porc)){
        actuel <- porc[index]
        before <- sum(actuel < cums)
        prev1 <- recom.f.indicator[nrow(recom.f.indicator)-before,]
        next1 <- recom.f.indicator[nrow(recom.f.indicator)-before+1,]

        porc[index] <- prev1[1] + (actuel-c(0,cums)[nrow(recom.f.indicator)-before]) /prev1[2]
      }
    }
  } else{
    porc <- stats::runif(noc,0,length.total[n.chromosome+1]) #Position der Rekombinationspunkte
  }


  if(duplication.rate>0){
    porc.d <- sortd(c(porc, 0, length.total[n.chromosome+1]))
    rpod <- c(0, (stats::rbinom(noc,1,duplication.rate) * 2:(noc+1)),0)
    pod <- porc.d[rpod]
    pod2 <- rep(0,length(pod))

    count <- 1
    add.one <- rep(0,length(pod))


    for(index in unique(c(0,rpod))[-1]){
      activ.chromosome <- sum(pod[count] > length.total)
      length.d <- stats::rexp(1, rate=(1/duplication.length)) * (-1)^(stats::rbinom(1,1,0.5))
      #print(c("pod",pod, "count", count, "porc.d", porc.d, "index", index, "activ.chromosome", activ.chromosome))
      pod2[count] <- max(min(pod[count]+ length.d, porc.d[index+1], length.total[activ.chromosome+1]),porc.d[index-1], length.total[activ.chromosome])
      if(pod2[count]==(porc.d[index])|| sum(length.total[start.point]==porc.d[index])){
        add.one[count] <- 1
      }
      count <- count+1
    }
    pod.start <- pod
    pod.start[pod.start>pod2] <- pod2[pod.start>pod2]
    pod.end <- pod
    pod.end[pod.end<pod2] <- pod2[pod.end<pod2]

  } else{
    pod <- pod2 <- rpod <- pod.start <- pod.end <- numeric(0)

  }


  start.point <- c(stats::rbinom(n.chromosome,1,0.5),0) * (1:(n.chromosome+1)) #Wechsel zu Beginn des Chromosoms
  porc <- sortd((c(length.total[start.point],porc))) # Sortieren der Rekombinationspunkte
  #Füge bvei irrelevante Punkte hinzu die in jedemfall Außerhalb des Gens liegen
  porc <- c(-1,porc,length.total[n.chromosome+1]+1)


  # recombination on dupliciation sequences
  dup <- list()
    dup[[1]] <- (parent[[11]])
  dup[[2]] <- (parent[[12]])
  dup[[3]] <- "test"

  for(abc in 1:2){
    counter <- 0
    if(length(dup[[abc]])>0){
      posi <- 1:nrow(dup[[abc]])
      for(index in 1:nrow(dup[[abc]])){
        ndup <- stats::rpois(1, (dup[[abc]][index,3]- dup[[abc]][index,2])* duplication.recombination)
#        print(ndup)
        if(ndup>0){
          pdup <- sort(stats::runif(ndup, min=dup[[abc]][index,2], max= dup[[abc]][index,3]))
        } else{
          pdup <- numeric(0)
        }
        if(counter==1 && dup[[abc]][index,1] == dup[[abc]][(index-1),1]){
          pdup <- c(dup[[abc]][index,2], pdup)
        }
        counter <- 0
        if(ndup%%2 && (sum(dup[[abc]][index,1]>=porc)%%2)==(abc%%2)){ # recombination on a relevant section
          porc <- sort(c(porc, dup[[abc]][index,1]))
        }
        if(ndup>0){
          if(ndup%%2==0){ #garanty odd number of elements
            pdup <- c(pdup, dup[[abc]][index,3])
            counter <- 1
            }
          dup[[abc]][index,3] <- pdup[1]
          if(ndup>1){
            for(index2 in seq(2,ndup,by=2)){
              dup[[abc]] <- rbind(dup[[abc]], dup[[abc]][index,])
              dup[[abc]][nrow(dup[[abc]]),2:3] <- pdup[index2:(index2+1)] # adding additional duplication segment
              posi <- c(posi,index)
            }
          }
        }
      }
      posi <- sort(posi,index.return=TRUE)$ix
      dup[[abc]] <- matrix(dup[[abc]][posi,], ncol=8)
      #remove duplication segements with length 0
#      print(dup[[1]])
      remove0 <- (dup[[abc]][,2] == dup[[abc]][,3])
#      print(remove0)
      if(sum(remove0)>0){
        remove0 <- remove0 * 1:length(remove0)
        dup[[abc]] <- dup[[abc]][-remove0,]
      }
    }
  }


  #
  new.poc <- NULL
  new.mut <- NULL
  new.origin <- NULL
  new.dup <- NULL
  activ <- 1
  store_mut <- list(population$info$snp.position[parent[[3]]], population$info$snp.position[parent[[4]]])

  for(index in 1:(length(porc)-1)){
    #    print(parent[[activ+10]])
    #    if(length(parent[[activ+10]])>0) print(is.matrix[parent[[activ+10]]])

    activ.porc <- (parent[[activ]]<porc[index+1]) & (parent[[activ]]>porc[index])

    if(length(store_mut[[activ]])==0){
      activ.mut <- logical(0)
    } else{
      activ.mut <- (store_mut[[activ]]<porc[index+1]) & (store_mut[[activ]]>porc[index])
    }

    if(length(dup[[activ]])>0){
      activ.dup1 <- (dup[[activ]][,1] <= porc[index+1]) * (dup[[activ]][,1] >= porc[index]) * ((dup[[activ]][,8] == 1) + (dup[[activ]][,8] == 3))
      # duplication section on the start/end of a chromosome
      activ.dup2 <- (dup[[activ]][,1] < porc[index+1]) * (dup[[activ]][,1] > porc[index]) * (dup[[activ]][,8] == 2)
      # duplication section in the middle of a chromosome

      # if the adjacting porcs are because of a start.point recombination some duplication sections have to be excluded
      leftb <- sum(porc[index]==porc) - sum(porc[index] == length.total[start.point])
      rightb <- sum(porc[index+1]==porc) - sum(porc[index+1] == length.total[start.point])

      activ.dup3 <- (1-leftb) * (dup[[activ]][,1] <= porc[index+1]) * (dup[[activ]][,1] == porc[index]) * ((dup[[activ]][,8] == 1) + (dup[[activ]][,8] == 3))
      activ.dup4 <- (1-rightb) * (dup[[activ]][,1] == porc[index+1]) * (dup[[activ]][,1] >= porc[index]) * ((dup[[activ]][,8] == 1) + (dup[[activ]][,8] == 3))

      activ.dup <- activ.dup1 + activ.dup2 - ((activ.dup3+ activ.dup4)>0)
      new.dup <- rbind(new.dup, dup[[activ]][activ.dup * (1:length(activ.dup)),])
    } else{
      active.dup <- integer(0)
      new.dup <- NULL
    }

    save1 <- max(new.poc,-2)!=porc[index+1]

    if(save1){
      new.poc <- c(new.poc, parent[[activ]][activ.porc], porc[index+1])
    }


    if(length(activ.mut)>0){
      new.mut <- c(new.mut, parent[[activ+2]][activ.mut])
    }


    start <- max(sum(parent[[activ]]<=porc[index]),0)

    activ.origin <- start:(start+sum(activ.porc))
    if(start==0){
      activ.origin <- activ.origin[-1]
    }

    if(length(activ.origin)>0){
      while(max(activ.origin,-1)>length(parent[[activ+4]])){ # add -1 avoid max(numeric(0))
        activ.origin <- activ.origin[-length(activ.origin)]

      }
    }

    if(porc[index+1]>0 && save1){
      new.origin <- c(new.origin, parent[[activ+4]][activ.origin])
    }
    #    first.following <- porc[index+1]<=parent[[activ]]
    #   first.following[length(first.following)] <- 1
    #  activ.porc[which(first.following == 1)[1]] <- 1
    # activ.porc <- activ.porc * 1:length(activ.porc)
    #new.origin <- rbind(new.origin, parent[[activ+4]][activ.porc,])
    activ <- 3 - activ
  }
  new.poc <- new.poc[-length(new.poc)]

  # mutation process
  n_mut <- stats::rbinom(1,n_snps, mutation.rate)
  if(length(new.mut)==0){
    n_remut <- 0
  } else{
    n_remut <- stats::rbinom(1, length(new.mut), remutation.rate) ############### BOTH CHECKER
  }

  if(n_mut==0){
    mutationen <- integer(0)
  } else{
    mutationen <- sample(1:n_snps, n_mut)
  }

  if(length(new.mut)>0){
    checker <- duplicated(c(new.mut, mutationen))
    mutationen <- mutationen[(1:length(checker))[-(1:length(new.mut))]-length(new.mut)]
  }

  if(n_remut==0){
    remutationen <- integer(0)
  } else{
    remutationen <- sample(1:length(new.mut), n_remut)
  }

  if(length(remutationen)==0){
    new.mut <- sort(unique(c(mutationen, new.mut)))
  } else{
    new.mut <- c(mutationen, new.mut[-remutationen])
    if(length(new.mut)>0){
      new.mut <- sort(new.mut)
      new.mut <- new.mut[!duplicated(new.mut)]
    }

  }


#  print(pod)
  new.dups <- NULL

  if(length(pod)>0){
    new.dups <- cbind(0,pod.start, pod.end, info.parent[1],info.parent[2],info.parent[3], 0, 0)
#    print(new.dups)
    # calculation of the position on the chromosome and the origin of the sequence
    for(index in 1:length(pod)){

      position.options <- c(length.total[sum(length.total<pod.start[index])], pod.start[index], length.total[sum(length.total<pod.start[index])+ 1])#start, mid, end
      samp<- sample(1:3,1)
      new.dups[index,1] <- position.options[samp]
      activ.chromo <- (sum(porc <= pod.start[index])+1+ add.one[index])%%2 +1
      new.dups[index,7] <- activ.chromo
      new.dups[index,8] <- samp
    }
  }
  # Duplication process

  new.dup <- rbind(new.dup, new.dups)
  if(length(new.dup)>0){
    order <- sort(new.dup[,1],index.return=TRUE)$ix
    new.dup <- new.dup[order,]
  }
  new.origin_old <- new.origin
  new.poc_old <- new.poc
  if(delete.same.origin==TRUE && length(new.origin)>1){
    for(index in length(new.origin):2){
      check <- prod(new.origin[index] == new.origin[index-1])
      if(check==TRUE){
        new.origin <- new.origin[-index]
        new.poc <- new.poc[-index]
      }
    }
  }

  if(gen.architecture!=0){
    # Transform points of recombination according to position
    for(index in 1:length(new.poc)){
      #        chromo <- find.chromo(parent[[haplo]][index], population$info$length.total)
      before <- find.snpbefore(new.poc[index], population$info$gen.architecture[[gen.architecture]]$snp.position)
      if(before>0){
        new_p_before <- population$info$snp.position[before]
        p_before <- population$info$gen.architecture[[gen.architecture]]$snp.position[before]
      } else{
        new_p_before <- population$info$length.total[1]
        p_before <-length.total[1]
      }
      if(before<n_snps){
        new_p_after <- population$info$snp.position[before+1]
        p_after <- population$info$gen.architecture[[gen.architecture]]$snp.position[before+1]
      } else{
        new_p_after <- population$info$length.total[length(population$info$length.total)]
        p_after <- length.total[length(length.total)]
      }
      share <- (new.poc[index]-p_before) / (p_after-p_before)
      new.poc[index] <- new_p_before + share * (new_p_after-new_p_before)
    }
  }


  if(length(new.poc)!=(length(new.origin)+1)){
    print("Rekombination - origin Inkonsistenz!!")
      print(new.poc)
      print(new.origin)
      print(parent)
      print(porc)

  }


  if(gene.editing==TRUE){
    hap_sequence <- compute.snps_single(population, new.poc, new.mut, new.origin, decodeOriginsU=decodeOriginsU)
    ed_info <- population$info$editing_info[[length(population$info$editing_info)]]
    edits_p <- numeric(nr.edits)
    edits <- 0
    current_p <- 1
    max_p <- sum(population$info$snp)
    while(edits < nr.edits && current_p <= max_p){
      if(hap_sequence[ed_info[current_p,1]] == ed_info[current_p,2]){
        edits <- edits + 1
        edits_p[edits] <- ed_info[current_p,1]
      }
      current_p <- current_p +1
    }
    changes <- edits_p

    new.mut <- sort(c(new.mut, edits_p))
  }
  maxl <- max(length.total)
  segment_length <- diff(c(0,porc[-unique(c(1,length(porc)))], maxl))
  share_a <- sum(segment_length[(1:length(segment_length)%%2)==1])/maxl
  return(list(new.poc, new.mut, new.origin, info.parent, new.dup, porc, share_a))
}
```
