## Supplementary material for "MoBPS - Modular Breeding Program Simulator": MoBPS R-package: bv.development.box.R

```
'#
  Authors
Torsten Pook,

Copyright (C) 2017 -- 2018  Torsten Pook

This program is free software; you can redistribute it and/or
modify it under the terms of the GNU General Public License
as published by the Free Software Foundation; either version 3
of the License, or (at your option) any later version.

This program is distributed in the hope that it will be useful,
but WITHOUT ANY WARRANTY; without even the implied warranty of
MERCHANTABILITY or FITNESS FOR A PARTICULAR PURPOSE.  See the
GNU General Public License for more details.

You should have received a copy of the GNU General Public License
along with this program; if not, write to the Free Software
Foundation, Inc., 59 Temple Place - Suite 330, Boston, MA  02111-1307, USA.
'#

#' Devolopment of genetic/breeding value
#'
#' Function to plot genetic/breeding values for multiple generation/cohorts
#' @param population population list
#' @param display Choose between "bv", "pheno", "bve" (default: "bv")
#' @param display.selection Display lines between generated cohorts via selection (webinterface)
#' @param display.reproduction Display lines between generated cohorts via reproduction (webinterface)
#' @param database Groups of individuals to consider for the export
#' @param gen Quick-insert for database (vector of all generations to export)
#' @param cohorts Quick-insert for database (vector of names of cohorts to export)
#' @param bvrow Which traits to display (for multiple traits separte plots (par(mfrow)))
#' @param json If TRUE extract which cohorts to plot according to the json-file used in json.simulation
#' @param ylim Set this to fix the y-axis of the plot
#' @param fix_mfrow Set TRUE to not use mfrow - use for custom plots
#' @export

bv.development.box <- function(population, database=NULL, gen=NULL, cohorts=NULL,
                           bvrow="all", json=FALSE, display="bv",
                           display.selection=FALSE,
                           display.reproduction=FALSE,
                           ylim=NULL, fix_mfrow=FALSE){

  if(length(bvrow)==1 && bvrow=="all"){
    bvrow <- 1:population$info$bv.nr
  }

  values_total <- list()
  values <- list()
  creating.type <- list()
  time.point <- list()
  sex <- list()
  if(!fix_mfrow){
    graphics::par(mfrow=c(1,length(bvrow)))
  }

  if(json){
    ids <- to_plot <- numeric(length(population$info$json[[1]]))
    for(index in 1:length(ids)){
      ids[index] <- population$info$json[[1]][[index]]$id
      if(length(population$info$json[[1]][[index]]$'BV Plot')>0 && sum(population$info$cohorts[,1]==ids[index])>0){
        to_plot[index] <- population$info$json[[1]][[index]]$'BV Plot'
      }
      cohorts <- ids[which(to_plot=="Yes")]
      if(length(cohorts)==0){
        cohorts <- population$info$cohorts[,1]
      }
    }
  }
  if(length(gen)>0){
    for(index in gen){
      if(display=="bv"){
        values[[length(values)+1]] <- get.bv(population, gen=index)
      } else if(display=="bve"){
          values[[length(values)+1]] <- get.bve(population, gen=index)
      } else if(display=="pheno"){
        values[[length(values)+1]] <- get.pheno(population, gen=index)
      }
      time.point[[length(time.point)+1]] <- get.time.point(population, gen=index)
      creating.type[[length(creating.type)+1]] <- get.creating.type(population, gen=index)
      sex[[length(sex)+1]] <- 0
    }
  }
  if(length(database)>0){
    for(index in 1:nrow(database)){
      if(display=="bv"){
        values[[length(values)+1]] <- get.bv(population, database=database[index,,drop=FALSE])
      } else if(display=="bve"){
        values[[length(values)+1]] <- get.bve(population, database=database[index,,drop=FALSE])
      } else if(display=="pheno"){
        values[[length(values)+1]] <- get.pheno(population, database=database[index,,drop=FALSE])
      }
      time.point[[length(time.point)+1]] <- get.time.point(population, database=database[index,,drop=FALSE])
      creating.type[[length(creating.type)+1]] <- get.creating.type(population, database=database[index,,drop=FALSE])
      sex[[length(sex)+1]] <- database[index,2]
    }
  }
  if(length(cohorts)>0){
    for(index in cohorts){
      if(display=="bv"){
        values[[length(values)+1]] <- get.bv(population, cohorts=index)
      } else if(display=="bve"){
        values[[length(values)+1]] <- get.bve(population, cohorts=index)
      } else if(display=="pheno"){
        values[[length(values)+1]] <- get.pheno(population, cohorts=index)
      }
      time.point[[length(time.point)+1]] <- get.time.point(population, cohorts=index)
      creating.type[[length(creating.type)+1]] <- get.creating.type(population, cohorts=index)
      sex[[length(sex)+1]] <- 1+ (as.numeric(population$info$cohorts[population$info$cohorts[,1]==index,4])>0)
    }
  }
  sex <- unlist(sex)
  time_plot <-  1:length(values)
  type_plot <-  rep(1, length(values))
  for(index in 1:length(values)){
    time_plot[index] <- mean(time.point[[index]])
    if(length(unique(time.point[[index]]))>1){
      print("More than one time point in a plotted element")
    }
  }
  for(index in 1:length(values)){
    type_plot[index] <- stats::median(creating.type[[index]])
    if(length(unique(creating.type[[index]]))>1){
      print("More than one creating type in a plotted element")
    }
  }
  for(nr in bvrow){
    time_points <- sort(unique(time_plot))
    x_axis <- length(time_points) + length(values)
    y_min <- Inf
    y_max <- -Inf
    for(index in 1:length(values)){
      y_min <- min(y_min, values[[index]][nr,])
      y_max <- max(y_max, values[[index]][nr,])
    }

    graphics::boxplot(x=c(-10^10), xlim=c(0,x_axis), ylim=if(length(ylim)==0){c(y_min,y_max)} else{ylim})
    pos <- 0
    pref <- 0
    label_pos <- NULL
    cohort_pos <- numeric(length(values))

    for(index in time_points){
      for(activ_c in which(time_plot==index)){
        graphics::boxplot(values[[activ_c]][nr,], add=TRUE, at=pos, width=0.95,
                          col = c("black", "blue", "red")[sex[activ_c]+1])
        cohort_pos[activ_c] <- pos
        pos <- pos + 1
      }
      graphics::abline(v=pos)
      pos <- pos + 1
      label_pos <- c(label_pos, mean(c(pref, pos-2)))
      pref <- pos
    }
    display.names <- NULL
    if(length(gen)>0){
      display.names <- c(display.names, paste("Generation", gen))
    }
    if(length(database)>0){
      display.names <- c(display.names, paste0(c("M", "F")[database[,2]], "_", database[,1]))
    }
    if(length(cohorts)>0){
      display.names <- c(display.names, cohorts)
    }
    graphics::axis(1, at=cohort_pos, labels=display.names,las=2)
    graphics::axis(3, at=label_pos, label=time_points)
    edges <- population$info$json[[2]]
    for(index in 1:length(edges)){
      if(display.selection){
        if(edges[[index]]$'Breeding Type'=="Selection"){
          from <- which(edges[[index]]$from==cohorts)
          to <- which(edges[[index]]$to==cohorts)
          if(length(from)>0 && length(to)>0){
            graphics::lines( c(cohort_pos[from], cohort_pos[to]),c(stats::median(values[[from]][nr,]), stats::median(values[[to]][nr,])), col="green", lwd=2)
          }
        }
      }
      if(display.reproduction){
        if(edges[[index]]$'Breeding Type'=="Reproduction"){
          from <- which(edges[[index]]$from==cohorts)
          to <- which(edges[[index]]$to==cohorts)
          if(length(from)>0 && length(to)>0){
            graphics::lines( c(cohort_pos[from], cohort_pos[to]),c(stats::median(values[[from]][nr,]), stats::median(values[[to]][nr,])), col="orange", lwd=2)
          }
        }
      }
    }
  }
  names(values) <- display.names

  return(values)
}
```
