## Supplementary material for "MoBPS - Modular Breeding Program Simulator": MoBPS R-package: bv.development.R

```
'#
  Authors
Torsten Pook,

Copyright (C) 2017 -- 2018  Torsten Pook

This program is free software; you can redistribute it and/or
modify it under the terms of the GNU General Public License
as published by the Free Software Foundation; either version 3
of the License, or (at your option) any later version.

This program is distributed in the hope that it will be useful,
but WITHOUT ANY WARRANTY; without even the implied warranty of
MERCHANTABILITY or FITNESS FOR A PARTICULAR PURPOSE.  See the
GNU General Public License for more details.

You should have received a copy of the GNU General Public License
along with this program; if not, write to the Free Software
Foundation, Inc., 59 Temple Place - Suite 330, Boston, MA  02111-1307, USA.
'#

#' Devolopment of genetic/breeding value
#'
#' Function to plot genetic/breeding values for multiple generation/cohorts
#' @param population population list
#' @param database Groups of individuals to consider for the export
#' @param gen Quick-insert for database (vector of all generations to export)
#' @param cohorts Quick-insert for database (vector of names of cohorts to export)
#' @param confidence Draw confidence intervals for (1- bv, 2- bve, 3- pheno; default: c(1,2,3))
#' @param quantile Quantile of the confidence interval to draw (default: 0.05)
#' @param bvrow Which traits to display (for multiple traits separte plots (par(mfrow)))
#' @param ignore.zero Cohorts with only 0 individuals are not displayed (default: TRUE)
#' @param json If TRUE extract which cohorts to plot according to the json-file used in json.simulation
#' @param display.time.point Set TRUE to use time point of generated to sort groups
#' @param display.creating.type Set TRUE to show Breedingtype used in generation (web-interface)
#' @param equal.spacing Equal distance between groups (independent of time.point)
#' @param development Include development of (1- bv, 2- bve, 3- pheno; default: c(1,2,3))
#' @param display.cohort.name Set TRUE to display the name of the cohort in the x-axis
#' @param display.sex Set TRUE to display the creating.type (Shape of Points - web-based-application)
#' @param display.line Set FALSE to not display the line connecting cohorts
#' @param time_reorder Set TRUE to order cohorts according to the time point of generation
#' @param ylim Set this to fix the y-axis of the plot
#' @param fix_mfrow Set TRUE to not use mfrow - use for custom plots
#' @export

bv.development <- function(population, database=NULL, gen=NULL, cohorts=NULL,
                           confidence= c(1,2,3), development = c(1,2,3), quantile=0.95, bvrow="all", ignore.zero=TRUE,
                           json=FALSE,
                           display.time.point=FALSE,
                           display.creating.type=FALSE,
                           display.cohort.name=FALSE,
                           display.sex=FALSE,
                           equal.spacing = FALSE,
                           time_reorder=FALSE,
                           display.line=TRUE,
                           ylim=NULL,
                           fix_mfrow=FALSE){

  if(length(bvrow)==1 && bvrow=="all"){
    bvrow <- 1:population$info$bv.nr
  }
  bv <- list()
  bve <- list()
  pheno <- list()
  time.point <- list()
  creating.type <- list()
  sex <- list()
  if(!fix_mfrow){
    graphics::par(mfrow=c(1,length(bvrow)))
  }


  if(json){
    ids <- to_plot <- numeric(length(population$info$json[[1]]))
    for(index in 1:length(ids)){
      ids[index] <- population$info$json[[1]][[index]]$id
      if(length(population$info$json[[1]][[index]]$'BV Plot')>0){
        to_plot[index] <- population$info$json[[1]][[index]]$'BV Plot'
      }
      cohorts <- ids[which(to_plot=="Yes")]
      if(length(cohorts)==0){
        cohorts <- population$info$cohorts[,1]
      }
    }
  }
  if(length(gen)>0){
    for(index in gen){
      bv[[length(bv)+1]] <- get.bv(population, gen=index)
      bve[[length(bve)+1]] <- get.bve(population, gen=index)
      pheno[[length(pheno)+1]] <- get.pheno(population, gen=index)
      time.point[[length(time.point)+1]] <- get.time.point(population, gen=index)
      creating.type[[length(creating.type)+1]] <- get.creating.type(population, gen=index)
      sex[[length(sex)+1]] <- 0
    }
  }
  if(length(database)>0){
    for(index in 1:nrow(database)){
      bv[[length(bv)+1]] <- get.bv(population, database=database[index,,drop=FALSE])
      bve[[length(bve)+1]] <- get.bve(population, database=database[index,,drop=FALSE])
      pheno[[length(pheno)+1]] <- get.pheno(population, database=database[index,,drop=FALSE])
      time.point[[length(time.point)+1]] <- get.time.point(population, database=database[index,,drop=FALSE])
      creating.type[[length(creating.type)+1]] <- get.creating.type(population, database=database[index,,drop=FALSE])
      sex[[length(sex)+1]] <- database[index,2]
    }
  }
  if(length(cohorts)>0){
    for(index in cohorts){
      bv[[length(bv)+1]] <- get.bv(population, cohorts=index)
      bve[[length(bve)+1]] <- get.bve(population, cohorts=index)
      pheno[[length(pheno)+1]] <- get.pheno(population, cohorts=index)
      time.point[[length(time.point)+1]] <- get.time.point(population, cohorts=index)
      creating.type[[length(creating.type)+1]] <- get.creating.type(population, cohorts=index)
      sex[[length(sex)+1]] <- 1+ (as.numeric(population$info$cohorts[population$info$cohorts[,1]==index,4])>0)
    }
  }
  sex <- unlist(sex)
  if(!display.sex){
    sex <- rep(0, length(sex))
  }

  for(nr in bvrow){
    color <- c("black", "blue", "red")
    q <- stats::qnorm(1 - (1-quantile)/2)
    means <- sds <- all0 <- matrix(NA, nrow=3, ncol=length(bv))
    time_plot <-  1:length(bv)
    type_plot <-  rep(1, length(bv))

    if(display.time.point){
      for(index in 1:length(bv)){
        time_plot[index] <- mean(time.point[[index]])
        if(length(unique(time.point[[index]]))>1){
          print("More than one time point in a plotted element")
        }
      }
    }
    if(display.creating.type){
      for(index in 1:length(bv)){
        type_plot[index] <- stats::median(creating.type[[index]])
        if(length(unique(creating.type[[index]]))>1){
          print("More than one creating type in a plotted element")
        }
      }
    }

    for(index in 1:length(bv)){
      means[1,index] <- base::mean(bv[[index]][nr,])
      means[2,index] <- base::mean(bve[[index]][nr,])
      means[3,index] <- base::mean(pheno[[index]][nr,])
      sds[1,index] <- stats::sd(bv[[index]][nr,])
      sds[2,index] <- stats::sd(bve[[index]][nr,])
      sds[3,index] <- stats::sd(pheno[[index]][nr,])
      all0[1,index] <- base::prod(bv[[index]][nr,]==0)
      all0[2,index] <- base::prod(bve[[index]][nr,]==0)
      all0[3,index] <- base::prod(pheno[[index]][nr,]==0)
    }
    for(index in 1:3){
      if(sum(development==index)==0){
        means[index,] <- NA
        all0[index,] <-  1
      }
    }

    if(ignore.zero==TRUE){
      means[all0*(1:length(means))] <- NA
      sds[all0*(1:length(means))] <- NA
    }

    ub <- means - sds * q
    ob <- means + sds * q

    # time sort
    if(time_reorder){
      reorder <- sort(time_plot, index.return=TRUE)$ix
    } else{
      reorder <- 1:length(time_plot)
    }

    means <- means[,reorder]
    sds <- sds[,reorder]
    all0 <- all0[,reorder]
    time_plot <- time_plot[reorder]
    type_plot <- type_plot[reorder]
    sex <- sex[reorder]

    if(equal.spacing){
      xlabel <- time_plot
      time_plot <- 1:length(time_plot)
    }

    inc <- (rowSums(all0)!=ncol(all0))* (1:3)
    if(display.cohort.name){
      graphics::par(mar=c(8.1,4.1,2.1,0.1))
    } else{
      graphics::par(mar=c(4.1,4.1,2.1,0.1))
    }

    graphics::plot(time_plot, means[1,],type=if(display.line) {"l"} else {NULL},  main=paste("Development:", population$info$trait.name[nr]),
                   xlab=if(display.cohort.name){""}else{"time"}, ylab="breeding value", ylim=if(length(ylim)==0) {c(min(ub, na.rm=TRUE),max(ob, na.rm=TRUE))} else{ylim}, lwd=2,
                   xaxt = if(display.cohort.name || equal.spacing){'n'}else {NULL}, cex=1.5)
    graphics::lines(time_plot, means[2,], col="blue", lwd=2)
    graphics::lines(time_plot, means[3,], col="red", lwd=2)
    graphics::points(time_plot, means[1,], pch=type_plot, col= c("black", "blue", "red")[sex+1], cex=1.5, lwd=2)
#    graphics::points(means[2,], pch=type_plot, col="blue")
#    graphics::points(means[3,], pch=type_plot, col="red")
    for(art in confidence){
      graphics::lines(time_plot, ob[art,], col=color[art], lty=3, lwd=2)
      graphics::lines(time_plot, ub[art,], col=color[art], lty=3, lwd=2)
    }
    graphics::legend("bottomright",c("breeding value","breeding value estimate","phenotype")[inc], lty=c(1,1,1)[inc], col=color[inc], lwd=c(2,2,2)[inc])
    temp1 <- unique(type_plot)
    if(length(temp1)>1){
      graphics::legend("topleft", c("Founder", "Selection", "Reproduction", "Recombination", "Selfing", "DH-Gene", "Cloning", "Combine", "Aging", "Split")[temp1+1],
                       pch=temp1, cex=0.75)
    }
    if(equal.spacing && !display.cohort.name){
      graphics::axis(1, at=time_plot, labels=xlabel)
    }

    if(display.cohort.name){
      display.names <- NULL
      if(length(gen)>0){
        display.names <- c(display.names, paste("Generation", gen))
      }
      if(length(database)>0){
        display.names <- c(display.names, paste0(c("M", "F")[database[,2]], "_", database[,1]))
      }
      if(length(cohorts)>0){
        display.names <- c(display.names, cohorts)
      }
      display.names <- display.names[reorder]
      graphics::axis(1, at=time_plot, labels=display.names,las=2)
    }
  }


}
```
