## Supplementary material for "MoBPS - Modular Breeding Program Simulator": MoBPS R-package: calculate.bv.R

```
'#
  Authors
Torsten Pook,

Copyright (C) 2017 -- 2018  Torsten Pook

This program is free software; you can redistribute it and/or
modify it under the terms of the GNU General Public License
as published by the Free Software Foundation; either version 3
of the License, or (at your option) any later version.

This program is distributed in the hope that it will be useful,
but WITHOUT ANY WARRANTY; without even the implied warranty of
MERCHANTABILITY or FITNESS FOR A PARTICULAR PURPOSE.  See the
GNU General Public License for more details.

You should have received a copy of the GNU General Public License
along with this program; if not, write to the Free Software
Foundation, Inc., 59 Temple Place - Suite 330, Boston, MA  02111-1307, USA.
'#

#' Calculate breeding values
#'
#' Internal function to calculate the breeding value of a given individual
#' @param population Population list
#' @param gen Generation of the individual of interest
#' @param sex Sex of the individual of interest
#' @param nr Number of the individual of interest
#' @param activ_bv traits to consider
#' @param decodeOriginsU Used function for the decoding of genetic origins [[5]]/[[6]]
#' @param import.position.calculation Function to calculate recombination point into adjacent/following SNP
#' @param store.effect.freq If TRUE store the allele frequency of effect markers per generation
#' @param bit.storing Set to TRUE if the RekomBre (not-miraculix! bit-storing is used)
#' @param nbits Bits available in RekomBre-bit-storing
#' @param output_compressed Set to TRUE to get a miraculix-compressed genotype/haplotype
#' @export

calculate.bv <- function(population, gen, sex, nr, activ_bv, import.position.calculation=NULL,
                         decodeOriginsU=decodeOriginsR, store.effect.freq=FALSE,
                         bit.storing=FALSE, nbits=30, output_compressed=FALSE){

  # Falls nötig könnten Haplotypen hier erst bestimmt werden.
  if(population$info$miraculix){
    if (requireNamespace("miraculix", quietly = TRUE)) {
      geno <- miraculix::computeSNPS(population, gen, sex, nr, what="geno")
    } else{
      stop("Usage of miraculix without being installed!")
    }

  } else{
    hap <- compute.snps(population, gen, sex, nr, import.position.calculation=import.position.calculation, decodeOriginsU=decodeOriginsU, bit.storing=bit.storing, nbits=nbits, output_compressed=output_compressed)
    geno <- colSums(hap)
  }

  bv_final <- numeric(length(activ_bv))
  snp.before <- population$info$cumsnp
  cindex <- 1
  for(bven in activ_bv){
    bv <- population$info$base.bv[bven] # Mittelwert

    real.bv.adds <- population$info$real.bv.add[[bven]]
    # Additive Effekte -Vektorielle loesung
    if(length(real.bv.adds)>0){


      position <- population$info$effect.p.add[[bven]]
      neff <- nrow(real.bv.adds)
      take <- (geno[position] + 2L ) * neff + 1:neff
      ## ^ seems to be extremely inefficient!
      if(population$info$bve.poly.factor[bven]==1){
        bv <- bv + population$info$bve.mult.factor[bven] * sum((real.bv.adds[take]))
      } else{
        bv <- bv + population$info$bve.mult.factor[bven] * sum((real.bv.adds[take])^population$info$bve.poly.factor[bven])
      }

    }


    real.bv.mults <- population$info$real.bv.mult[[bven]]
    # Multiplikative Effekte - Vektorielle Version
    if(length(real.bv.mults)>0){
      position1 <- population$info$effect.p.mult1[[bven]]
      position2 <- population$info$effect.p.mult2[[bven]]
      bv <- bv + sum(real.bv.mults[cbind(1:nrow(real.bv.mults), 5L + geno[position1]*3L + geno[position2])])
    }

    # Wuerfel Effekte - Nicht vektorielle da Listenzugriffe (verbesserbar wenn viele solche Effekte betrachtet werden)
    real.bv.dices <- population$info$real.bv.dice[[bven]]
    if(length(real.bv.dices)>0 && length(real.bv.dices[[1]])>0){
      for(index in 1:length(real.bv.dices[[1]])){
        posis <- numeric(nrow(real.bv.dices[[1]][[index]]))
        for(index2 in 1:length(posis)){
          posis[index2] <- snp.before[real.bv.dices[[1]][[index]][index2,2]] + real.bv.dices[[1]][[index]][index2,1L]
        }
        activ_p <- 1L
        for(index2 in 1:length(posis)){
          activ_p <- activ_p + geno[posis[index2]] * (3L^(length(posis)-index2))
        }
        bv <- bv + real.bv.dices[[2]][[index]][activ_p]
      }
    }
    bv_final[cindex] <- bv
    cindex <- cindex + 1
  }

  if(store.effect.freq){
    freq_list <- cbind(geno[population$info$effect.p]==0, geno[population$info$effect.p]==1, geno[population$info$effect.p]==2)
    storage.mode(freq_list) <- "integer"
  } else{
    freq_list <- "notrelevant"
  }

  return(list(bv_final, freq_list))

}
```
