## Supplementary material for "MoBPS - Modular Breeding Program Simulator": MoBPS R-package: clean.up.R

```
'#
  Authors
Torsten Pook,

Copyright (C) 2017 -- 2018  Torsten Pook

This program is free software; you can redistribute it and/or
modify it under the terms of the GNU General Public License
as published by the Free Software Foundation; either version 3
of the License, or (at your option) any later version.

This program is distributed in the hope that it will be useful,
but WITHOUT ANY WARRANTY; without even the implied warranty of
MERCHANTABILITY or FITNESS FOR A PARTICULAR PURPOSE.  See the
GNU General Public License for more details.

You should have received a copy of the GNU General Public License
along with this program; if not, write to the Free Software
Foundation, Inc., 59 Temple Place - Suite 330, Boston, MA  02111-1307, USA.
'#

#' Clean-up recombination points
#'
#' Function to remove recombination points + origins with no influence on markers
#' @param population Population list
#' @param gen Generations to clean up (default: "current")
#' @param remove.no.snp.segments If TRUE remove segments which do not contain SNPs
#' @export

clean.up <- function(population, gen="current", remove.no.snp.segments=FALSE){
  #remove.no.snp.segments nur auf die Schnelle und super ineffizient!
  generations <- gen
  if(gen=="current"){
    generations <- length(population$breeding)
  }
  if(gen=="all"){
    generations <- 1:length(population$breeding)
  }

  for(current.gen in generations){
  for(sex in 1:2){
    n <- length(population$breeding[[current.gen]][[sex]])
    if(n>0){
      for(index in 1:n){
        for(row in 1:2){
          if(length(population$breeding[[current.gen]][[sex]][[index]][[4+row]])>7){
            remove.list <- rep(0,(nrow(population$breeding[[current.gen]][[sex]][[index]][[4+row]])-1))
            for(abc in 1:(nrow(population$breeding[[current.gen]][[sex]][[index]][[4+row]])-1)){

              if(prod(population$breeding[[current.gen]][[sex]][[index]][[4+row]][abc,]==population$breeding[[current.gen]][[sex]][[index]][[4+row]][abc+1,])){
                remove.list[abc] <- 1
               }
            }
            if(sum(remove.list)>0){
              remove.list <- remove.list * 1:(length(remove.list))
              population$breeding[[current.gen]][[sex]][[index]][[4+row]] <- population$breeding[[current.gen]][[sex]][[index]][[4+row]][-remove.list,]
              population$breeding[[current.gen]][[sex]][[index]][[row]] <- population$breeding[[current.gen]][[sex]][[index]][[row]][-(remove.list+1*(remove.list!=0))]

            }

          }

        }
      }
    }
  }
  }
  # SUPER INEFFZIENT!
  # ES MUESSTEN DEUTLICHER WENIGER VERGLEICHE DURCHGEFUEHRT WERDEN!
  if(remove.no.snp.segments==TRUE){
    snp.p <- population$info$snp.position
    for(current.gen in generations){
      for(sex in 1:2){
        n <- length(population$breeding[[current.gen]][[sex]])
        if(n>0){
          for(index in 1:n){
            for(row in 1:2){
              recom <- population$breeding[[current.gen]][[sex]][[index]][[row]]
              nr <- length(recom)-1
              rm <- numeric(nr)
              rm1 <- numeric(nr)
              for(index2 in 1:nr){
                prev1 <- recom[index2]
                next1 <- recom[index2+1]
                if(sum(prev1<snp.p)== sum(next1<snp.p)){
                  rm[index2] <- index2
                  rm1[index2] <- index2 +1
                }
              }
              if(rm[nr]>0){
                rm[nr] <- rm[nr] - 1
                rm1[nr] <- rm1[nr] -1
              }
              if(sum(rm)>0){
                population$breeding[[current.gen]][[sex]][[index]][[row]] <- population$breeding[[current.gen]][[sex]][[index]][[row]][-rm1]
                population$breeding[[current.gen]][[sex]][[index]][[row+4]] <- population$breeding[[current.gen]][[sex]][[index]][[row+4]][-rm,]
              }


            }
          }
        }
      }
    }
  }
  return(population)
}
```
