## Supplementary material for "MoBPS - Modular Breeding Program Simulator": MoBPS R-package: codeOriginsR.R

```
'#
  Authors
Torsten Pook,

Copyright (C) 2017 -- 2018  Torsten Pook

This program is free software; you can redistribute it and/or
modify it under the terms of the GNU General Public License
as published by the Free Software Foundation; either version 3
of the License, or (at your option) any later version.

This program is distributed in the hope that it will be useful,
but WITHOUT ANY WARRANTY; without even the implied warranty of
MERCHANTABILITY or FITNESS FOR A PARTICULAR PURPOSE.  See the
GNU General Public License for more details.

You should have received a copy of the GNU General Public License
along with this program; if not, write to the Free Software
Foundation, Inc., 59 Temple Place - Suite 330, Boston, MA  02111-1307, USA.
'#

#' Origins-coding(R)
#'
#' R-Version of the internal bitwise-coding of origins
#' @param M Origins matrix
#' @export

codeOriginsR <- function(M){
  P <- as.integer(colSums(t(M-1) * c(2^26,2^25,2^3,1)))
  return(P)
}

#' Origins-Decoding(R)
#'
#' R-Version of the internal bitwise-decoding of origins
#' @param P coded origins vector
#' @param row row
#' @export

decodeOriginsR <- function(P, row){
  activ <- P[row]
  gen <- floor(activ/(2^26))+1
  activ <- activ - (gen-1)*2^26
  sex <- floor(activ/(2^25))+1
  activ <- activ - (sex-1)*2^25
  nr <- floor(activ/(2^3))+1
  activ <- activ - (nr-1)*2^3
  chromo <- activ+1
  return(c(gen,sex,nr,chromo))
}


#' Check if package is available
#'
#' Funktion to check availablity of a package
#' @param mypkg package to test
#' @export

is.installed <- function(mypkg) is.element(mypkg, utils::installed.packages()[,1])
```
