## Supplementary material for "MoBPS - Modular Breeding Program Simulator": MoBPS R-package: compute.costs.cohorts.R

```
'#
  Authors
Torsten Pook,

Copyright (C) 2017 -- 2018  Torsten Pook

This program is free software; you can redistribute it and/or
modify it under the terms of the GNU General Public License
as published by the Free Software Foundation; either version 3
of the License, or (at your option) any later version.

This program is distributed in the hope that it will be useful,
but WITHOUT ANY WARRANTY; without even the implied warranty of
MERCHANTABILITY or FITNESS FOR A PARTICULAR PURPOSE.  See the
GNU General Public License for more details.

You should have received a copy of the GNU General Public License
along with this program; if not, write to the Free Software
Foundation, Inc., 59 Temple Place - Suite 330, Boston, MA  02111-1307, USA.
'#

#' Compute costs of a breeding program
#'
#' Function to derive the costs of a breeding program / population-list
#' @param population population-list
#' @param json If TRUE extract which cohorts to plot according to the json-file used in json.simulation
#' @param phenotyping.costs Costs for the generation of a phenotype
#' @param genotyping.costs Costs for the geneation of a genotype
#' @param housing.costs Costs for housing
#' @param fix.costs one time occuring fixed costs
#' @param fix.costs.annual annually occuring fixed costs
#' @param profit.per.bv profit generated by bv per animal
#' @param database Groups of individuals to consider
#' @param gen Quick-insert for database (vector of all generations to consider)
#' @param cohorts Quick-insert for database (vector of names of cohorts to consider)
#' @param interest.rate Applied yearly interest rate
#' @export

compute.costs.cohorts <- function(population, gen=NULL, database=NULL, cohorts=NULL, json=TRUE,
                          phenotyping.costs = NULL, genotyping.costs = 0, housing.costs=NULL, fix.costs= 0, fix.costs.annual = 0,
                          profit.per.bv = 1, interest.rate = 1){


  if(json){
    ids <- to_plot <- numeric(length(population$info$json[[1]]))
    for(index in 1:length(ids)){
      ids[index] <- population$info$json[[1]][[index]]$label
      if(length(population$info$json[[1]][[index]]$'BV Plot')>0 && sum(population$info$cohorts[,1]==ids[index])>0){
        to_plot[index] <- population$info$json[[1]][[index]]$'BV Plot'
      }
      cohorts <- ids[which(to_plot=="Yes")]
      if(length(cohorts)==0){
        cohorts <- population$info$cohorts[,1]
      }
    }
  }

  database <- get.database(population, gen=gen, database=database,cohorts=cohorts)


  cost_table <- NULL
  cost_table_interest <- NULL
  time_span <- 0
  if(json==TRUE){
    time_point <- population$info$cohorts[cohorts,8]
  } else{
    time_point <- database[,1] - 1
  }
  for(index in 1:nrow(database)){
    cost_genotypes <- 0
    cost_phenotypes <- 0
    cost_housing <- 0
    profit_bv <- 0
    if(json==TRUE){
      node <- which(population$info$json[[8]]==cohorts[index])
    }
    for(index2 in database[index,3]:database[index,4]){
      cost_genotypes <- cost_genotypes + population$breeding[[database[index,1]]][[database[index,2]]][[index2]][[16]] * genotyping.costs
      if(json==FALSE){
        cost_phenotypes <- cost_phenotypes + sum(population$breeding[[database[index,1]]][[database[index,2]]][[index2]][[15]] * phenotyping.costs)
      } else{
        cost_phenotypes <- cost_phenotypes + population$info$json[[7]][[1]][which(population$info$json[[1]][[node]]$'Phenotyping Class' == population$info$json[[7]][[2]])]
      }
      if(json==FALSE){
        cost_housing <- cost_housing + housing.costs
      } else{
        cost_housing <- cost_housing + population$info$json[[6]][[1]][which(population$info$json[[1]][[node]]$'Housing Cost Class' == population$info$json[[6]][[2]])]
      }
      profit_bv <- profit_bv + sum(profit.per.bv * population$breeding[[database[index,1]]][[6+database[index,2]]][,index2])

    }

    cost_table <- rbind(cost_table, c(cost_genotypes, cost_phenotypes, cost_housing, profit_bv, profit_bv-cost_phenotypes- cost_genotypes - cost_housing))
    if(json==TRUE){
      cost_table_interest <- rbind(cost_table_interest, c(cost_genotypes, cost_phenotypes, cost_housing, profit_bv, (profit_bv-cost_phenotypes- cost_genotypes- cost_housing) *
                                     interest.rate ^ (population$breeding[[database[index,1]]][[database[index,2]+10]][[index2]])))
    } else{
      cost_table_interest <- rbind(cost_table_interest, c(cost_genotypes, cost_phenotypes, cost_housing, profit_bv, (profit_bv-cost_phenotypes- cost_genotypes- cost_housing) *
                                     interest.rate ^ (database[index,1]-1)))
    }
    time_span <- max(time_span, (population$breeding[[database[index,1]]][[database[index,2]+10]]))

  }

  gain_total <- colSums(cost_table_interest)
  if(time_span==0){
    time_span <- nrow(population$info$size)
  }
  gain_total <- round(gain_total, digits=2)
  cat(paste0("The given programs takes ",time_span," years and results in:\n"))
  cat(paste0("Genotyping costs: ", gain_total[1], " Euro \n"))
  cat(paste0("Phenotyping costs: ", gain_total[2], " Euro \n"))
  cat(paste0("Housing costs: ", gain_total[3], " Euro \n"))
  cat(paste0("Fixed costs: ", round(fix.costs, digits=2), " Euro \n"))
  cat(paste0("Annual fixed costs: ", round(sum(fix.costs.annual * cumprod(rep(interest.rate,time_span))/interest.rate), digits=2), " Euro \n"))
  cat(paste0("Gains: ", round(gain_total[4], digits=2), " Euro \n"))
  cat(paste0("Total: ", round(gain_total[4] - gain_total[3] - gain_total[1] - gain_total[2] - fix.costs -sum(fix.costs.annual * cumprod(rep(interest.rate,time_span))/interest.rate), digits=2), " Euro \n"))
  colnames(cost_table_interest) <- c("Genotyping", "Phenotyping", "Gains", "Housing costs",  "Total")

  time_point <- as.numeric(time_point)
  plot_points <- sort(unique(time_point))
  time_eco <- numeric(length(plot_points))
  annual <- round(fix.costs.annual * cumprod(rep(interest.rate,time_span))/interest.rate, digits=2)
  for(index in 1:length(plot_points)){
    time_eco[index] <- sum(cost_table_interest[time_point==plot_points[index],5])
  }
  for(index in 1:length(annual)){
    time_eco[which(plot_points<=index)[1]] <- time_eco[which(plot_points<=index)[1]] - annual[index]
  }
  eco_total <- cumsum(time_eco) - fix.costs

  graphics::par(mfrow=c(1,1))
  graphics::plot(plot_points, eco_total, type="l", ylab="gain in Euro", xlab="year", lwd=2)
  colr <- rep("red", length(plot_points))
  colr[eco_total>0] <- "green"
  graphics::points(plot_points, eco_total, col=colr, cex=2,lwd=3)
  graphics::abline(h=0)

  cost_table_interest
}
```
