## Supplementary material for "MoBPS - Modular Breeding Program Simulator": MoBPS R-package: compute.costs.R

```
'#
  Authors
Torsten Pook,

Copyright (C) 2017 -- 2018  Torsten Pook

This program is free software; you can redistribute it and/or
modify it under the terms of the GNU General Public License
as published by the Free Software Foundation; either version 3
of the License, or (at your option) any later version.

This program is distributed in the hope that it will be useful,
but WITHOUT ANY WARRANTY; without even the implied warranty of
MERCHANTABILITY or FITNESS FOR A PARTICULAR PURPOSE.  See the
GNU General Public License for more details.

You should have received a copy of the GNU General Public License
along with this program; if not, write to the Free Software
Foundation, Inc., 59 Temple Place - Suite 330, Boston, MA  02111-1307, USA.
'#

#' Compute costs of a breeding program
#'
#' Function to derive the costs of a breeding program / population-list
#' @param population population-list
#' @param phenotyping.costs Costs for the generation of a phenotype
#' @param genotyping.costs Costs for the geneation of a genotype
#' @param fix.costs one time occuring fixed costs
#' @param fix.costs.annual annually occuring fixed costs
#' @param profit.per.bv profit generated by bv per animal
#' @param database Groups of individuals to consider
#' @param gen Quick-insert for database (vector of all generations to consider)
#' @param cohorts Quick-insert for database (vector of names of cohorts to consider)
#' @param interest.rate Applied yearly interest rate
#' @param base.gen Base generation (application of interest rate)
#' @export

compute.costs <- function(population, phenotyping.costs = 10, genotyping.costs = 100, fix.costs= 0, fix.costs.annual = 0,
                          profit.per.bv = 1, database=NULL, gen=NULL, cohorts=NULL,
                          interest.rate = 1, base.gen=1){


  if(length(gen)>0){
    database <- cbind(rep(gen,each=2), rep(1:2, length(gen)))
  }
  if(length(database)>0 && ncol(database)==2){
    start <- end <- numeric(nrow(database))
    for(index in 1:nrow(database)){
      start[index] <- 1
      end[index] <- population$info$size[database[index,1], database[index,2]]
    }
    database <- cbind(database, start, end)
  }
  if(length(cohorts)>0){
    database2 <- matrix(0L, nrow=length(cohorts), ncol=4)
    for(index in 1:length(cohorts)){
      row <- which(population$info$cohorts==cohorts[index])
      gen <- as.numeric(population$info$cohorts[row,2])
      sex <- 1 + (as.numeric(population$info$cohorts[row,4])>0)
      first <- as.numeric(population$info$cohorts[row,5 + sex])
      last <- first + as.numeric(population$info$cohorts[row,2 + sex]) - 1
      database2[index,] <- c(gen,sex,first,last)
    }
    database <- rbind(database, database2)
  }

  generation <- max(database[,1])

  t <- length(phenotyping.costs)
  if(t != (2*generation)){
    if(t == 1){
      phenotyping.costs <- matrix(phenotyping.costs, nrow=generation, ncol=2)
    }
    if(t==generation){
      phenotyping.costs <- matrix(phenotyping.costs, nrow=generation, ncol=2, byrow=FALSE)
    }
    if(t==2){
      phenotyping.costs <- matrix(phenotyping.costs, nrow=generation, ncol=2, byrow=TRUE)
      if(generation==2){
        print("Specify if phenotyping.costs are per generation or per sex. Default use: Gender")
      }
    }
  }

  t <- length(genotyping.costs)
  if(t != (2*generation)){
    if(t == 1){
      genotyping.costs <- matrix(genotyping.costs, nrow=generation, ncol=2)
    }
    if(t==generation){
      genotyping.costs <- matrix(genotyping.costs, nrow=generation, ncol=2, byrow=FALSE)
    }
    if(t==2){
      genotyping.costs <- matrix(genotyping.costs, nrow=generation, ncol=2, byrow=TRUE)
      if(generation==2){
        print("Specify if genotyping.costs are per generation or per sex. Default use: Gender")
      }
    }
  }

  t <- length(profit.per.bv)
  if(t != (2*generation)){
    if(t == 1){
      profit.per.bv <- matrix(profit.per.bv, nrow=generation, ncol=2)
    }
    if(t==generation){
      profit.per.bv <- matrix(profit.per.bv, nrow=generation, ncol=2, byrow=FALSE)
    }
    if(t==2){
      profit.per.bv <- matrix(profit.per.bv, nrow=generation, ncol=2, byrow=TRUE)
      if(generation==2){
        print("Specify if profit.per.bv are per generation or per sex. Default use: Gender")
      }
    }
  }

  if(length(fix.costs.annual)!=generation){
    fix.costs.annual <- rep(fix.costs.annual, length.out=generation)
  }

  if(interest.rate!=1){
    phenotyping.costs <- phenotyping.costs * (interest.rate ^ (1:generation-base.gen))
    genotyping.costs <- genotyping.costs * (interest.rate ^ (1:generation-base.gen))
    profit.per.bv <- profit.per.bv * (interest.rate ^ (1:generation-base.gen))
    fix.costs.annual <- fix.costs.annual * (interest.rate ^ (1:generation-base.gen))
  }

  costs_pheno <- costs_geno <- profit_bv <-numeric(nrow(database))

  for(index in 1:nrow(database)){
    activ <- database[index,]
    for(index2 in activ[3]:activ[4]){
      costs_pheno[index] <- costs_pheno[index] + population$breeding[[activ[1]]][[activ[2]]][[index]][[15]] * phenotyping.costs[activ[1], activ[2]]
      costs_geno[index] <- costs_geno[index] + population$breeding[[activ[1]]][[activ[2]]][[index]][[16]] * genotyping.costs[activ[1], activ[2]]
      profit_bv[index] <- profit_bv[index] + population$breeding[[activ[1]]][[activ[2]+6]][1,index] * profit.per.bv[activ[1], activ[2]]
    }
  }

  costs_pergroup <- costs_pheno + costs_geno

  gains_group <- (profit_bv) - (costs_pheno) - (costs_geno)

  timest <- unique(database[,1])
  gains_year <- numeric(length(timest))
  t <- 1
  for(index in timest){
    gains_year[t] <- sum(gains_group[which(database[,1]==index)]) + fix.costs.annual[index]
    t <- t + 1
  }

  graphics::par(mfrow=(c(1,2)))
  graphics::plot(timest, gains_year, main="Gains per Year", xlab="generation", ylab="gains")
  graphics::abline(h=0)
  graphics::plot(gains_group, main="Gains per Cohort", xlab="cohort", ylab="gains", xaxt="n")
  graphics::axis(1, at=1:length(gains_group), labels=cohorts)
  graphics::abline(h=0)

  total <- sum(gains_year) - fix.costs
  return(total)
}
```
