## Supplementary material for "MoBPS - Modular Breeding Program Simulator": MoBPS R-package: compute.snps.R

```
'#
  Authors
Torsten Pook,

Copyright (C) 2017 -- 2018  Torsten Pook

This program is free software; you can redistribute it and/or
modify it under the terms of the GNU General Public License
as published by the Free Software Foundation; either version 3
of the License, or (at your option) any later version.

This program is distributed in the hope that it will be useful,
but WITHOUT ANY WARRANTY; without even the implied warranty of
MERCHANTABILITY or FITNESS FOR A PARTICULAR PURPOSE.  See the
GNU General Public License for more details.

You should have received a copy of the GNU General Public License
along with this program; if not, write to the Free Software
Foundation, Inc., 59 Temple Place - Suite 330, Boston, MA  02111-1307, USA.
'#

#' Compute genotype/haplotype
#'
#' Internal function for the computation of genotypes & haplotypes
#' @param population Population list
#' @param gen Generation of the individual to compute
#' @param sex Gender of the individual to compute
#' @param nr Number of the individual to compute
#' @param faster If FALSE use slower version to compute markers between recombination points
#' @param import.position.calculation Function to calculate recombination point into adjacent/following SNP
#' @param from_p First SNP to consider
#' @param to_p Last SNP to consider
#' @param decodeOriginsU Used function for the decoding of genetic origins [[5]]/[[6]]
#' @param bit.storing Set to TRUE if the RekomBre (not-miraculix! bit-storing is used)
#' @param nbits Bits available in RekomBre-bit-storing
#' @param output_compressed Set to TRUE to get a miraculix-compressed genotype/haplotype
#' @export

compute.snps <- function(population, gen, sex, nr, faster=TRUE, import.position.calculation=NULL,
                         from_p=1, to_p=Inf, decodeOriginsU=decodeOriginsR,
                         bit.storing=FALSE, nbits=30, output_compressed=FALSE){
  n.snps <- sum(population$info$snp)
  if(bit.storing){
    hap <- matrix(0L, ncol=ceiling(n.snps/nbits), nrow=2)
  } else{
    hap <- matrix(0L, ncol=n.snps, nrow=2)
  }
  if(bit.storing==TRUE){
    from.p.bit <- (from_p-1)/nbits + 1
    to.p.bit <- ceiling(to_p/nbits)
  }

  temp1 <- 0
  for(hapnr in 1:2){
    current.recombi <- population$breeding[[gen]][[sex]][[nr]][[hapnr]]
    current.mut <- population$breeding[[gen]][[sex]][[nr]][[hapnr+2]]
    current.ursprung <- population$breeding[[gen]][[sex]][[nr]][[hapnr+4]]
    if(is.function(import.position.calculation)){
      till <- import.position.calculation(current.recombi)
    } else if(faster==TRUE && sum(population$info$snps.equidistant==TRUE)){
      till <- rep(0, length(current.recombi))
      for(index in 1:length(current.recombi)){
        current.chromo <- sum(current.recombi[index] >= population$info$length.total)
        if(current.recombi[index]>population$info$snp.position[sum(population$info$snp)]){
          till[index] <- sum(population$info$snp)
        } else{
          between2 <- population$info$position[[current.chromo]][1]
          length_chromo <- sum(population$info$length[0:(current.chromo-1)])
          position_chromo <- sum(population$info$snp[0:(current.chromo-1)])
          till[index] <- ceiling((current.recombi[index] - length_chromo - between2) / (between2*2)) + position_chromo

        }
      }
    } else if(faster==TRUE){
      till <- rep(0, length(current.recombi))
      for(index in 1:length(current.recombi)){
        till[index] <- find.snpbefore(current.recombi[index], population$info$snp.position)
#        till[index] <- sum(population$info$snp.position <= current.recombi[index])
      }

    }
    if(bit.storing){
      leftover <- NULL

      for(index2 in 1:(length(current.recombi)-1)){
        ursprung <- decodeOriginsU(current.ursprung,index2)
        ursprung[1] <- population$info$origin.gen[ursprung[1]]
        if(faster==TRUE){
          if((till[index2+1] >=(till[index2]+1))&& (till[index2]+1 <= to_p) && (till[index2+1])>= from_p){
            start <- ceiling(max(from.p.bit,(till[index2]+1)/nbits))
            start_snp <- (max(from_p,(till[index2]+1))-1)%%nbits +1
#            if(ceiling((till[index2]+1)/nbits)<start){
#              start_snp <- 1
#            }
            tail <- ceiling(min(to.p.bit,till[index2+1]/nbits))
            tail_snp <- (min(to_p,till[index2+1])-1)%%nbits + 1
#            if(tail > ceiling(till[index2+1]/nbits)){
#              tail_snp <- (to_p-1)%%nbits - (from_p-1)%%nbits + 1
#            }
            mid <- (start+1):(tail-1)

            # Wenn start außerhalb vom bereichen start_snp <- 1
            # end tail außerhalb vom bereich dann tail_snp <- nbits bbv. Ende

            if(start==tail){
              leftover <- c(leftover, bit.snps(population$breeding[[ursprung[1]]][[ursprung[2]]][[ursprung[3]]][[ursprung[4]+8]][start], nbits)[start_snp:tail_snp])
              if(length(leftover)==nbits){
                hap[hapnr,start] <- bit.storing(leftover, nbits)
                leftover <- NULL
              }
            } else{
              leftover <- c(leftover, bit.snps(population$breeding[[ursprung[1]]][[ursprung[2]]][[ursprung[3]]][[ursprung[4]+8]][start], nbits)[start_snp:nbits])
              hap[hapnr,start] <- bit.storing(leftover, nbits)
              leftover <- bit.snps(population$breeding[[ursprung[1]]][[ursprung[2]]][[ursprung[3]]][[ursprung[4]+8]][tail], nbits)[1:tail_snp]
              if(length(leftover)==nbits){
                hap[hapnr,tail] <- bit.storing(leftover, nbits)
                leftover <- NULL
              }
            }

            if(start<(tail-1)){
              hap[hapnr,mid] <- population$breeding[[ursprung[1]]][[ursprung[2]]][[ursprung[3]]][[ursprung[4]+8]][mid]
            }
          }
        }
      }
      if(length(leftover)>0){
        hap[hapnr,ncol(hap)] <- bit.storing(leftover, nbits)
        leftover <- NULL
      }
      if(length(current.mut)>0){
        for(changes in current.mut){
          variablennr <- ceiling(changes/nbits)
          bit <- changes - (variablennr-1) * nbits
          snpseq <- bit.snps(hap[hapnr,variablennr], nbits)
          snpseq[bit] <- 1L - snpseq[bit]
          hap[hapnr,variablennr] <- bit.storing(snpseq, nbits)
        }
      }
    } else{
      for(index2 in 1:(length(current.recombi)-1)){
        ursprung <- decodeOriginsU(current.ursprung,index2)
        ursprung[1] <- population$info$origin.gen[ursprung[1]]
        if(faster==TRUE){
          if((till[index2+1] >=(till[index2]+1))&& (till[index2]+1 <= to_p) && (till[index2+1])>= from_p){
            relevant.snp <- max(from_p,(till[index2]+1)):min(to_p,till[index2+1])
          } else{
            relevant.snp <- NULL
          }
          hap[hapnr,relevant.snp] <-population$breeding[[ursprung[1]]][[ursprung[2]]][[ursprung[3]]][[ursprung[4]+8]][relevant.snp]
        } else{
          relevant.snp <- (population$info$snp.position < current.recombi[index2+1])*(population$info$snp.position >= current.recombi[index2])*(1:n.snps)
          hap[hapnr,relevant.snp] <-population$breeding[[ursprung[1]]][[ursprung[2]]][[ursprung[3]]][[ursprung[4]+8]][relevant.snp]
        }
      }
      if(length(current.mut)>0){
        position <- current.mut
        hap[hapnr,position] <- 1L - hap[hapnr,position]
      }
    }

  }


  if(output_compressed==FALSE && bit.storing==TRUE){
    hap_snp <- rbind(bit.snps(hap[1, from.p.bit:min(to.p.bit, ceiling(n.snps/nbits))], nbits, population=population, from.p.bit=from.p.bit),
                     bit.snps(hap[2, from.p.bit:min(to.p.bit, ceiling(n.snps/nbits))], nbits, population=population, from.p.bit=from.p.bit))
    return(hap_snp)

  } else if(bit.storing==TRUE){
    return(hap[,from_p:min(to_p, ceiling(n.snps/nbits))])
  }

  return(hap[, from_p:min(to_p, n.snps)])
}

#' Compute genotype/haplotype in gene editing application
#'
#' Internal function for the computation of genotypes & haplotypes in gene editing application
#' @param population Population list
#' @param current.recombi vector of currently activ recombination points
#' @param current.mut vector of currently activ mutations
#' @param current.ursprung vector of currently activ origins
#' @param faster If FALSE use slower version to compute markers between recombination points
#' @param import.position.calculation Function to calculate recombination point into adjacent/following SNP
#' @param decodeOriginsU Used function for the decoding of genetic origins [[5]]/[[6]]

compute.snps_single <- function(population, current.recombi, current.mut, current.ursprung, faster=TRUE,
                                import.position.calculation=NULL, decodeOriginsU=decodeOriginsR){
  n.snps <- sum(population$info$snp)
  hap<- numeric(n.snps)
  temp1 <- 0
  hapnr <- 1

  if(is.function(import.position.calculation)){
    till <- import.position.calculation(current.recombi)
  } else if(faster==TRUE && sum(population$info$snps.equidistant==TRUE)){
    till <- rep(0, length(current.recombi))
    for(index in 1:length(current.recombi)){
      current.chromo <- sum(current.recombi[index] >= population$info$length.total)
      if(current.recombi[index]>population$info$snp.position[sum(population$info$snp)]){
        till[index] <- sum(population$info$snp)
      } else{
        between2 <- population$info$position[[current.chromo]][1]
        length_chromo <- sum(population$info$length[0:(current.chromo-1)])
        position_chromo <- sum(population$info$snp[0:(current.chromo-1)])
        till[index] <- ceiling((current.recombi[index] - length_chromo - between2) / (between2*2)) + position_chromo

      }
    }
  } else if(faster==TRUE){
    till <- rep(0, length(current.recombi))
    for(index in 1:length(current.recombi)){
      till[index] <- sum(population$info$snp.position <= current.recombi[index])
    }

  }
  for(index2 in 1:(length(current.recombi)-1)){
    ursprung <- decodeOriginsU(current.ursprung,index2)
    ursprung[1] <- population$info$origin.gen[ursprung[1]]
    if(faster==TRUE){
      if(till[index2+1] >=(till[index2]+1)){
        relevant.snp <- (till[index2]+1):till[index2+1]
      } else{
        relevant.snp <- NULL
      }

      hap[relevant.snp] <-population$breeding[[ursprung[1]]][[ursprung[2]]][[ursprung[3]]][[ursprung[4]+8]][relevant.snp]
    } else{
      relevant.snp <- (population$info$snp.position < current.recombi[index2+1])*(population$info$snp.position >= current.recombi[index2])*(1:n.snps)
      hap[relevant.snp] <-population$breeding[[ursprung[1]]][[ursprung[2]]][[ursprung[3]]][[ursprung[4]+8]][relevant.snp]
    }
  }
  if(length(current.mut)>0){
    position <- current.mut
    hap[position] <- 1-hap[position]
  }
  return(hap)
}
```
