## Supplementary material for "MoBPS - Modular Breeding Program Simulator": MoBPS R-package: creating.diploid.R

```
'#
  Authors
Torsten Pook,

Copyright (C) 2017 -- 2018  Torsten Pook

This program is free software; you can redistribute it and/or
modify it under the terms of the GNU General Public License
as published by the Free Software Foundation; either version 3
of the License, or (at your option) any later version.

This program is distributed in the hope that it will be useful,
but WITHOUT ANY WARRANTY; without even the implied warranty of
MERCHANTABILITY or FITNESS FOR A PARTICULAR PURPOSE.  See the
GNU General Public License for more details.

You should have received a copy of the GNU General Public License
along with this program; if not, write to the Free Software
Foundation, Inc., 59 Temple Place - Suite 330, Boston, MA  02111-1307, USA.
'#

#' Generation of the starting population
#'
#' Generation of the starting population
#' @param dataset SNP dataset, use "random", "allhetero" "all0" when generating a dataset via nsnp,nindi
#' @param nsnp number of markers to generate in a random dataset
#' @param nindi number of inidividuals to generate in a random dataset
#' @param freq frequency of allele 1 when randomly generating a dataset
#' @param population Population list
#' @param sex.s Specify with newly added individuals are male (1) or female (2)
#' @param sex.quota Share of newly added female individuals (deterministic if sex.s="fixed", alt: sex.s="random")
#' @param add.chromosome If TRUE add an additional chromosome to the dataset
#' @param generation Generation of the newly added individuals (default: 1)
#' @param class Migration level of the newly added individuals
#' @param chromosome.length Length of the newly added chromosome (default: 5)
#' @param length.before Length before the first SNP of the dataset (default: 5)
#' @param length.behind Length after the last SNP of the dataset (default: 5)
#' @param snps.equidistant Use equidistant markers (computationally faster! ; default: TRUE)
#' @param snp.position Location of each marker on the genetic map
#' @param change.order If TRUE sort markers according to given marker positions
#' @param bv.total Number of traits (If more than traits via real.bv.X use traits with no directly underlying QTL)
#' @param polygenic.variance Genetic variance of traits with no underlying QTL
#' @param bit.storing Set to TRUE if the RekomBre (not-miraculix! bit-storing is used)
#' @param nbits Bits available in RekomBre-bit-storing
#' @param randomSeed Set random seed of the process
#' @param miraculix If TRUE use miraculix package for data storage, computations and dataset generation
#' @param miraculix.dataset Set FALSE to deactive miraculix package for dataset generation
#' @param n.additive Number of additive QTL
#' @param n.dominant Number of dominante QTL
#' @param n.qualitative Number of qualitative epistatic QTL
#' @param n.quantitative Number of quantitative epistatic QTL
#' @param var.additive.l Variance of additive QTL
#' @param var.dominant.l Variance of dominante QTL
#' @param var.qualitative.l Variance of qualitative epistatic QTL
#' @param var.quantitative.l Variance of quantitative epistatic QTL
#' @param exclude.snps Marker were no QTL are simulated on
#' @param replace.real.bv If TRUE delete the simulated traits added before
#' @param shuffle.traits Combine different traits into a joined trait
#' @param shuffle.cor Target Correlation between shuffeled traits
#' @param real.bv.add Single Marker effects
#' @param real.bv.mult Two Marker effects
#' @param real.bv.dice Multi-marker effects
#' @param bve.mult.factor Multiplicate trait value times this
#' @param bve.poly.factor Potency trait value over this
#' @param base.bv Average genetic value of a trait
#' @param add.chromosome.ends Add chromosome ends as recombination points
#' @param new.phenotype.correlation Correlation of the simulated enviromental variance
#' @param new.breeding.correlation Correlation of the simulated genetic variance (child share! heritage is not influenced!
#' @param add.architecture Add genetic architecture (marker positions)
#' @param position.scaling Manual scaling of snp.position
#' @param name.cohort Name of the newly added cohort
#' @param template.chip Import genetic map and chip from a species ("cattle", "chicken", "pig")
#' @param vcf Path to a vcf-file used as input genotypes (correct haplotype phase is assumed!)
#' @param chr.nr Vector containing the assosiated chromosome for each marker (default: all on the same)
#' @param bp Vector containing the physical position (bp) for each marker (default: 1,2,3...)
#' @param bpcm.conversion Convert physical position (bp) into a cM position (default: 0 - not done)
#' @param snp.name Vector containing the name of each marker (default ChrXSNPY - XY chosen accordingly)
#' @param hom0 Vector containing the first allelic variant in each marker (default: 0)
#' @param hom1 Vector containing the second allelic variant in each marker (default: 1)
#' @param skip.rest Internal variable needed when adding multipe chromosomes jointly
#' @param beta.shape1 First parameter of the beta distribution for simulating allele frequencies
#' @param beta.shape2 Second parameter of the beta distribution for simulating allele frequencies
#' @param time.point Time point at which the new individuals are generated
#' @param creating.type Technique to generate new individuals (usage in web-based application)
#' @param trait.name Name of the trait generated
#' @param share.genotyped Share of individuals genotyped in the founders
#' @param genotyped.s Specify with newly added individuals are genotyped (1) or not (0)
#' @param map map-file that contains up to 5 colums (Chromsome, SNP-id, Bp-position, M-position, allele freq - Everything not provides it set to NA). A map can be imported via ensembl.map()
#' @param remove.invalid.qtl Set to FALSE to deactive the automatic removal of QTLs on markers that do not exist
#' @export


creating.diploid <- function(dataset=NULL, vcf=NULL, chr.nr=NULL, bp=NULL, snp.name=NULL, hom0=NULL, hom1=NULL,
                             bpcm.conversion=0,
                             nsnp=0, nindi=0, freq="beta", population=NULL, sex.s="fixed",
                             add.chromosome=FALSE, generation=1, class=0L,
                             sex.quota = 0.5, chromosome.length=NULL,length.before=5, length.behind=5,
                             real.bv.add=NULL, real.bv.mult=NULL, real.bv.dice=NULL, snps.equidistant=NULL,
                             change.order=FALSE, bv.total=0, polygenic.variance=100,
                             bve.mult.factor=NULL, bve.poly.factor=NULL,
                             base.bv=NULL, add.chromosome.ends=TRUE,
                             new.phenotype.correlation=NULL,
                             new.breeding.correlation=NULL,
                             add.architecture=NULL, snp.position=NULL,
                             position.scaling=FALSE,
                             bit.storing=FALSE,
                             nbits=30, randomSeed=NULL,
                             miraculix=TRUE,
                             miraculix.dataset=TRUE,
                             n.additive=0,
                             n.dominant=0,
                             n.qualitative=0,
                             n.quantitative=0,
                             var.additive.l=NULL,
                             var.dominant.l=NULL,
                             var.qualitative.l=NULL,
                             var.quantitative.l=NULL,
                             exclude.snps=NULL,
                             replace.real.bv=FALSE,
                             shuffle.traits=NULL,
                             shuffle.cor=NULL,
                             skip.rest=FALSE,
                             name.cohort=NULL,
                             template.chip=NULL,
                             beta.shape1=1,
                             beta.shape2=1,
                             time.point=0,
                             creating.type=0,
                             trait.name=NULL,
                             share.genotyped=1,
                             genotyped.s=NULL,
                             map=NULL,
                             remove.invalid.qtl=TRUE){

  if(length(randomSeed)>0){
    set.seed(randomSeed)
  }
  if(!miraculix && miraculix.dataset){
    cat("miraculix.dataset only possible when miraculix is active")
    miraculix.dataset <- FALSE
  }

  if(length(map)>0){
    while(ncol(map)<5){
      map <- cbind(map, NA)
    }
    chr.nr <- map[,1]
    snp.name <- map[,2]
    if(sum(!is.na(map[,3]))>0){
      if(length(bp)==0){
        bp <- numeric(nrow(map))
      }
      bp[!is.na(map[,3])] <- map[!is.na(map[,3]),3]
    }
    if(sum(!is.na(map[,4]))==nrow(map)){
      snp.position <- map[,4]
    } else if(sum(is.na(map[,4]))==nrow(map) || length(chromosome.length)==0){
      if(bpcm.conversion==0){
        cat("Assume 1cm per 100.000.000bp - to change use bpcm.conversion")
        bpcm.conversion <- 100000000
      }
      map[,4] <- as.numeric(bp) /  bpcm.conversion
    }
    if(sum(!is.na(map[,4]))==nrow(map) || length(chromosome.length)==0){
      chr.opt <- unique(chr.nr)
      chromosome.length <- numeric(length(chr.opt))
      for(index in 1:length(chr.opt)){
        chromosome.length[index] <- max(as.numeric(map[map[,1]==chr.opt[index],4])) + min(as.numeric(map[map[,1]==chr.opt[index],4]))
      }
    }
    if(sum(!is.na(map[,5]))==nrow(map)){
      freq <- map[,5]
    }
    if(nsnp!=0 && nsnp!=nrow(map)){
      cat("Number of SNPs not in concordance with used map!\n")

      cat("Set number of SNPs to")
      cat(nrow(map))
      cat("!\n")
    }
    nsnp <- nrow(map)

  }
  if(length(chromosome.length)==0 || (length(chromosome.length)==1 && chromosome.length==0)){
    if(length(snp.position)>0){
      chromosome.length <- max(snp.position) + min(snp.position)
      if(chromosome.length<=0){
        print("invalid setting for snp.position - Proceed with chromosome of 5M and equidistant markers")
        chromosome.length <- 5
        snps.equidistant <- TRUE
      }
    } else{
      chromosome.length <- 5
    }
  }
  if (requireNamespace("miraculix", quietly = TRUE)) {
    codeOriginsU <- miraculix::codeOrigins
    decodeOriginsU <- miraculix::decodeOrigins
  } else{
    codeOriginsU <- codeOriginsR
    decodeOriginsU <- decodeOriginsR
    miraculix <- FALSE
    miraculix.dataset <- FALSE
  }

  if(length(template.chip)==1){
    if(template.chip=="cattle"){
      target_snp <- nsnp
      chromosome.length <- MoBPS::cattle_chip[,2]
      nsnp <- round(MoBPS::cattle_chip[,3] * chromosome.length)
      if(target_snp>0){
        nsnp_temp  <- nsnp * target_snp / sum(nsnp)
        nsnp <- floor(nsnp_temp)
        if(sum(nsnp)<target_snp){
          add1 <- sort(nsnp_temp-nsnp, index.return=TRUE, decreasing = TRUE)$ix[1:(target_snp-sum(nsnp))]
          nsnp[add1] <- nsnp[add1] +1
        }
      }

      chr.nr <- numeric(sum(nsnp))
      start1 <- 1
      for(index in 1:length(nsnp)){
        if(nsnp[index]!=0){
          chr.nr[start1:(start1+nsnp[index]-1)] <- index
          start1 <- start1 + nsnp[index]
        }

      }
    } else if(template.chip=="chicken"){
      target_snp <- nsnp
      chromosome.length <- MoBPS::chicken_chip[,2]/100
      nsnp <- round(MoBPS::chicken_chip[,3] * chromosome.length)
      if(target_snp>0){
        nsnp_temp  <- nsnp * target_snp / sum(nsnp)
        nsnp <- floor(nsnp_temp)
        if(sum(nsnp)<target_snp){
          add1 <- sort(nsnp_temp-nsnp, index.return=TRUE, decreasing = TRUE)$ix[1:(target_snp-sum(nsnp))]
          nsnp[add1] <- nsnp[add1] +1
        }
      }
      chr.nr <- numeric(sum(nsnp))
      start1 <- 1
      for(index in 1:length(nsnp)){
        if(nsnp[index]!=0){
          chr.nr[start1:(start1+nsnp[index]-1)] <- index
          start1 <- start1 + nsnp[index]
        }
      }
    } else if(template.chip=="pig"){
      target_snp <- nsnp
      chromosome.length <- MoBPS::pig_chip[,2]/100
      nsnp <- round(MoBPS::pig_chip[,3] * chromosome.length)
      if(target_snp>0){
        nsnp_temp  <- nsnp * target_snp / sum(nsnp)
        nsnp <- floor(nsnp_temp)
        if(sum(nsnp)<target_snp){
          add1 <- sort(nsnp_temp-nsnp, index.return=TRUE, decreasing = TRUE)$ix[1:(target_snp-sum(nsnp))]
          nsnp[add1] <- nsnp[add1] +1
        }
      }
      chr.nr <- numeric(sum(nsnp))
      start1 <- 1
      for(index in 1:length(nsnp)){
        if(nsnp[index]!=0){
          chr.nr[start1:(start1+nsnp[index]-1)] <- index
          start1 <- start1 + nsnp[index]
        }
      }
    } else if(template.chip=="sheep"){
      target_snp <- nsnp
      chromosome.length <- MoBPS::sheep_chip[,2]
      nsnp <- round(MoBPS::sheep_chip[,3] * chromosome.length)
      if(target_snp>0){
        nsnp_temp  <- nsnp * target_snp / sum(nsnp)
        nsnp <- floor(nsnp_temp)
        if(sum(nsnp)<target_snp){
          add1 <- sort(nsnp_temp-nsnp, index.return=TRUE, decreasing = TRUE)$ix[1:(target_snp-sum(nsnp))]
          nsnp[add1] <- nsnp[add1] +1
        }
      }
      chr.nr <- numeric(sum(nsnp))
      start1 <- 1
      for(index in 1:length(nsnp)){
        if(nsnp[index]!=0){
          chr.nr[start1:(start1+nsnp[index]-1)] <- index
          start1 <- start1 + nsnp[index]
        }
      }
    } else if(template.chip=="maize"){
      target_snp <- nsnp
      chromosome.length <- MoBPS::maize_chip[,2]
      nsnp <- round(MoBPS::maize_chip[,3] * chromosome.length)
      if(target_snp>0){
        nsnp_temp  <- nsnp * target_snp / sum(nsnp)
        nsnp <- floor(nsnp_temp)
        if(sum(nsnp)<target_snp){
          add1 <- sort(nsnp_temp-nsnp, index.return=TRUE, decreasing = TRUE)$ix[1:(target_snp-sum(nsnp))]
          nsnp[add1] <- nsnp[add1] +1
        }
      }
      chr.nr <- numeric(sum(nsnp))
      start1 <- 1
      for(index in 1:length(nsnp)){
        if(nsnp[index]!=0){
          chr.nr[start1:(start1+nsnp[index]-1)] <- index
          start1 <- start1 + nsnp[index]
        }
      }
    }

  }
  if(length(dataset)>0 && class(dataset)=="haplomatrix"){
    dataset <- list(dataset)
  }
  if(length(dataset)>0 && class(dataset)=="matrix"){
    miraculix.dataset <- FALSE
  }

  if(length(chr.nr)==1 && chr.nr>1){
    if(length(nsnp)==1  && nsnp > chr.nr){
      chr.nr <- sort(rep(1:chr.nr, length.out=nsnp))
      nsnp <- numeric(length(chr.nr))
      for(index in 1:length(nsnp)){
        nsnp[index] <- sum(chr.nr==index)
      }
    } else if(class(dataset)=="matrix" && nrow(dataset)>chr.nr){
      chr.nr <- sort(rep(1:chr.nr, length.out=nrow(dataset)))
    } else if(class(dataset)=="haplomatrix" && attr(dataset[[1]], "information")[2]==1){
      cat("Are you sure you want to generate a 1 SNP chromosome via miraculix?")
    }

  }

  if(skip.rest==FALSE){
    if(length(vcf)>0 && requireNamespace("vcfR", quietly = TRUE)){
      vcf_file <- vcfR::read.vcfR(vcf)
      vcf_data <- vcf_file@gt[,-1]
      dataset <- matrix(0L, nrow=nrow(vcf_data), ncol=ncol(vcf_data)*2)
      dataset[,(1:ncol(vcf_data))*2-1] <- as.integer(substr(vcf_data, start=1,stop=1))
      dataset[,(1:ncol(vcf_data))*2-1] <- as.integer(substr(vcf_data, start=3,stop=3))

      chr.nr <- as.numeric(vcf_file@fix[,1])
      bp <- as.numeric(vcf_file@fix[,2])
      snp.name <- vcf_file@fix[,3]
      hom0 <- vcf_file@fix[,4]
      hom1 <- vcf_file@fix[,5]

    }


    if(sum(nsnp)>0 && length(freq)==1 && freq=="beta"){
      freq <- stats::rbeta(sum(nsnp), shape1=beta.shape1, shape2=beta.shape2)
    }
    if(sum(nsnp)>0 && length(freq)<sum(nsnp)){
      freq <- rep(freq, length.out=sum(nsnp))
    }
    if(length(freq)>1 && length(freq)>sum(nsnp)){
      nsnp <- length(freq)
    }
    if(length(freq)>0 && sum(is.na(freq))>0){
      replace <- which(is.na(freq))
      freq[replace] <- stats::rbeta(sum(is.na(freq)), shape1=beta.shape1, shape2=beta.shape2)
      freq <- as.numeric(freq)
    }


    if(!is.list(var.additive.l) ){
      var.additive.l <- list(var.additive.l)
    }
    if(!is.list(var.dominant.l)){
      var.dominant.l <- list(var.dominant.l)
    }
    if(!is.list(var.qualitative.l)){
      var.qualitative.l <- list(var.qualitative.l)
    }
    if(!is.list(var.quantitative.l)){
      var.quantitative.l <- list(var.quantitative.l)
    }

    trait_sum <- n.additive + n.dominant + n.qualitative + n.quantitative
    test <- list(NULL)

    if(length(var.additive.l) < length(trait_sum)){
      var.additive.l <- c(var.additive.l, rep(test,length.out=length(trait_sum)-length(var.additive.l)))
    }
    if(length(var.dominant.l) < length(trait_sum)){
      var.dominant.l <- c(var.dominant.l, rep(test,length.out=length(trait_sum)-length(var.dominant.l)))
    }
    if(length(var.qualitative.l) < length(trait_sum)){
      var.qualitative.l <- c(var.qualitative.l, rep(test,length.out=length(trait_sum)-length(var.qualitative.l)))
    }
    if(length(var.quantitative.l) < length(trait_sum)){
      var.quantitative.l <- c(var.quantitative.l, rep(test,length.out=length(trait_sum)-length(var.quantitative.l)))
    }

    ntraits <- length(trait_sum)
    n.additive <- c(n.additive, rep(0, length.out=ntraits-length(n.additive)))
    n.dominant <- c(n.dominant, rep(0, length.out=ntraits-length(n.dominant)))
    n.qualitative <- c(n.qualitative, rep(0, length.out=ntraits-length(n.qualitative)))
    n.quantitative <- c(n.quantitative, rep(0, length.out=ntraits-length(n.quantitative)))

    if(length(unlist(c(var.qualitative.l, var.quantitative.l, var.additive.l, var.dominant.l)))>0){
      ntraits <- max(length(trait_sum), length(var.additive.l),length(var.dominant.l), length(var.qualitative.l), length(var.quantitative.l) )
      n.additive <- c(n.additive, rep(0, length.out=ntraits-length(n.additive)))
      n.dominant <- c(n.dominant, rep(0, length.out=ntraits-length(n.dominant)))
      n.qualitative <- c(n.qualitative, rep(0, length.out=ntraits-length(n.qualitative)))
      n.quantitative <- c(n.quantitative, rep(0, length.out=ntraits-length(n.quantitative)))
      trait_sum <- n.additive + n.dominant + n.qualitative + n.quantitative
      if(length(var.additive.l) < length(trait_sum)){
        var.additive.l <- rep(var.additive.l, length.out=length(trait_sum))
      }
      if(length(var.dominant.l) < length(trait_sum)){
        var.dominant.l <- rep(var.dominant.l, length.out=length(trait_sum))
      }
      if(length(var.qualitative.l) < length(trait_sum)){
        var.qualitative.l <- rep(var.qualitative.l, length.out=length(trait_sum))
      }
      if(length(var.quantitative.l) < length(trait_sum)){
        var.quantitative.l <- rep(var.quantitative.l, length.out=length(trait_sum))
      }
    }


    if(length(population)>0){
      if(length(real.bv.add)==0 && replace.real.bv==FALSE){
        real.bv.add <- population$info$real.bv.add
        real.bv.add[[population$info$bv.calc+1]] <- NULL
      } else if(replace.real.bv==FALSE){
        if(!is.list(real.bv.add)){
          real.bv.add <- list(real.bv.add)
        }
        real.bv.add <- c(population$info$real.bv.add, real.bv.add)
        real.bv.add[[population$info$bv.calc+1]] <- NULL
      }
      if(length(real.bv.mult)==0 && replace.real.bv==FALSE){
        real.bv.mult <- population$info$real.bv.mult
        real.bv.mult[[population$info$bv.calc+1]] <- NULL
      } else if(replace.real.bv==FALSE){
        if(!is.list(real.bv.mult)){
          real.bv.mult <- list(real.bv.mult)
        }
        real.bv.mult <- c(population$info$real.bv.mult, real.bv.mult)
        real.bv.mult[[population$info$bv.calc+1]] <- NULL
      }
      if(length(real.bv.dice)==0 && replace.real.bv==FALSE){
        real.bv.dice <- population$info$real.bv.dice
        real.bv.dice[[population$info$bv.calc+1]] <- NULL
      } else if(replace.real.bv==FALSE){
        if(!is.list(real.bv.dice)){
          real.bv.dice <- list(real.bv.dice)
        }
        real.bv.dice <- c(population$info$real.bv.dice, real.bv.dice)
        real.bv.dice[[population$info$bv.calc+1]] <- NULL
      }

    }
    if(length(real.bv.add)>0 && !is.list(real.bv.add)){
      real.bv.add <- list(real.bv.add)
    }
    if(length(real.bv.mult)>0 && !is.list(real.bv.mult)){
      real.bv.mult <- list(real.bv.mult)
    }
    if(length(real.bv.dice)>0 && !is.list(real.bv.dice)){
      real.bv.dice <- list(real.bv.dice)
    }
    so_far <- max(length(real.bv.dice), length(real.bv.add), length(real.bv.mult))
    if(length(trait_sum)>0){
      for(index_trait in 1:length(trait_sum)){
        var_additive <- var.additive.l[[index_trait]]
        var_dominant <- var.dominant.l[[index_trait]]
        var_qualitative <- var.qualitative.l[[index_trait]]
        var_quantitative <- var.quantitative.l[[index_trait]]
        if(n.additive[index_trait]>0 && length(var_additive)<n.additive[index_trait]){
          if(length(var_additive)==0){
            var_additive <- 1
          }
          var_additive <- rep(1, length.out=n.additive[index_trait])
          var.additive.l[[index_trait]] <- var_additive
        }
        if(n.dominant[index_trait]>0 && length(var_dominant)<n.dominant[index_trait]){
          if(length(var_dominant)==0){
            var_dominant <- 1
          }
          var_dominant <- rep(1, length.out=n.dominant[index_trait])
          var.dominant.l[[index_trait]] <- var_dominant
        }
        if(n.qualitative[index_trait]>0 && length(var_qualitative)<n.qualitative[index_trait]){
          if(length(var_qualitative)==0){
            var_qualitative <- 1
          }
          var_qualitative <- rep(1, length.out=n.qualitative[index_trait])
          var.qualitative.l[[index_trait]] <- var_qualitative
        }
        if(n.quantitative[index_trait]>0 && length(var_quantitative)<n.quantitative[index_trait]){
          if(length(var_quantitative)==0){
            var_quantitative <- 1
          }
          var_quantitative <- rep(1, length.out=n.quantitative[index_trait])
          var.quantitative.l[[index_trait]] <- var_quantitative

        }

        if(length(var_additive)!= n.additive[index_trait]){
          n.additive[index_trait] <- length(var_additive)
        }
        if(length(var_dominant)!= n.dominant[index_trait]){
          n.dominant[index_trait] <- length(var_dominant)
        }
        if(length(var_qualitative)!= n.qualitative[index_trait]){
          n.qualitative[index_trait] <- length(var_qualitative)
        }
        if(length(var_quantitative)!= n.quantitative[index_trait]){
          n.quantitative[index_trait] <- length(var_quantitative)
        }


        snpdata <- population$info$snp


        #This part is only needed in creating.diploid
        if(sum(nsnp)>0){
          snpdata <- c(snpdata, nsnp)
        } else if((is.matrix(dataset) && nrow(dataset)>0 ) || is.list(dataset)){
          if(length(chr.nr)>0 && length(unique(chr.nr))>1){
            rindex <- 1
            for(chr.index in unique(chr.nr)){
              if(is.list(dataset)){
                snpdata <- c(snpdata, attr(dataset[[rindex]], "information")[2])
                rindex <- rindex + 1
              } else{
                snpdata <- c(snpdata, sum(chr.nr==chr.index))
              }
            }
          } else{
            if(class(dataset)=="haplomatrix"){
              snpdata <- c(snpdata, attr(dataset[[1]], "information")[2])
            } else{
              snpdata <- c(snpdata, nrow(dataset))
            }

          }
        }

        # Generating additive
        effect_marker <- (1:sum(snpdata))
        if(length(exclude.snps)>0){
          effect_marker <- effect_marker[-exclude.snps]
        }

        add_marker <- sample(effect_marker, n.additive[index_trait], replace=if(n.additive[index_trait]>length(effect_marker)){TRUE} else{FALSE})
        dom_marker <- sample(effect_marker, n.dominant[index_trait], replace=if(n.dominant[index_trait]>length(effect_marker)){TRUE} else{FALSE})
        epi1_marker <- sample(effect_marker, n.quantitative[index_trait]*2, replace=if(n.quantitative[index_trait]*2>length(effect_marker)){TRUE} else{FALSE})
        epi2_marker <- sample(effect_marker, n.qualitative[index_trait]*2, replace=if(n.qualitative[index_trait]*2>length(effect_marker)){TRUE} else{FALSE})


        cum_snp <- cumsum(snpdata)
        real.bv.add.new <- NULL
        real.bv.mult.new <- NULL
        if(n.additive[index_trait]>0){
          add_snp <- add_chromo <- numeric(n.additive[index_trait])
          for(index in 1:n.additive[index_trait]){
            add_chromo[index] <- sum(add_marker[index] > cum_snp) + 1
            add_snp[index] <- add_marker[index] - c(0,cum_snp)[add_chromo[index]]
          }
          add_effect <- stats::rnorm(n.additive[index_trait], 1, var_additive)
          real.bv.add.new <- cbind(add_snp, add_chromo, add_effect,0,-add_effect)
        }
        if(n.dominant[index_trait]>0){
          dom_snp <- dom_chromo <- numeric(n.dominant[index_trait])
          for(index in 1:n.dominant[index_trait]){
            dom_chromo[index] <- sum(dom_marker[index] > cum_snp) + 1
            dom_snp[index] <- dom_marker[index] - c(0,cum_snp)[dom_chromo[index]]
          }
          dom_effect <- stats::rnorm(n.dominant[index_trait], 1, var_dominant)
          real.bv.add.new <- rbind(real.bv.add.new, cbind(dom_snp, dom_chromo, 0 ,dom_effect,dom_effect))

        }

        if(n.quantitative[index_trait]){
          epi1_snp <- epi1_chromo <- numeric(n.quantitative[index_trait]*2)
          for(index in 1:(n.quantitative[index_trait]*2)){
            epi1_chromo[index] <- sum(epi1_marker[index] > cum_snp) + 1
            epi1_snp[index] <- epi1_marker[index] - c(0,cum_snp)[epi1_chromo[index]]
          }

          effect_matrix <- matrix(0,nrow=n.quantitative[index_trait], ncol=9)
          for(index in 1:n.quantitative[index_trait]){
            d1 <- sort(abs(stats::rnorm(3, 1, var_quantitative[index])))
            d2 <- sort(abs(stats::rnorm(3, 1, var_quantitative[index])))
            effect_matrix[index,] <- c(d1*d2[1], d1*d2[2], d1*d2[3])
          }
          real.bv.mult.new <- cbind(epi1_snp[1:n.quantitative[index_trait]], epi1_chromo[1:n.quantitative[index_trait]],
                                    epi1_snp[-(1:n.quantitative[index_trait])], epi1_chromo[-(1:n.quantitative[index_trait])],
                                    effect_matrix)
        }

        if(n.qualitative[index_trait]>0){
          epi2_snp <- epi2_chromo <- numeric(n.qualitative[index_trait]*2)
          for(index in 1:(n.qualitative[index_trait]*2)){
            epi2_chromo[index] <- sum(epi2_marker[index] > cum_snp) + 1
            epi2_snp[index] <- epi2_marker[index] - c(0,cum_snp)[epi2_chromo[index]]
          }

          effect_matrix <- matrix(0,nrow=n.qualitative[index_trait], ncol=9)
          for(index in 1:n.qualitative[index_trait]){

            d1 <- -abs(stats::rnorm(9, 1, var_qualitative[index]))
            d1[c(3,7)] <- -d1[c(3,7)]
            effect_matrix[index,] <- d1
          }
          real.bv.mult.new <- rbind(real.bv.mult.new, cbind(epi2_snp[1:n.qualitative[index_trait]], epi2_chromo[1:n.qualitative[index_trait]],
                                                            epi2_snp[-(1:n.qualitative[index_trait])], epi2_chromo[-(1:n.qualitative[index_trait])],
                                                            effect_matrix))
        }

        real.bv.add[[index_trait+so_far]] <- real.bv.add.new
        real.bv.mult[[index_trait+so_far]] <- real.bv.mult.new

      }
    }


    perserve_bve <- length(population)==0

    if(length(population)>0 && length(population$info$bitstoring)>0){
      nbits <- population$info$bitstoring
      leftover <- population$info$leftover
      bit.storing <- TRUE
    }
    if(length(population)>0 && length(population$info$miraculix)>0 && population$info$miraculix){
      miraculix <- TRUE
    }


    nbv <- max(if(is.list(real.bv.add)){length(real.bv.add)} else{as.numeric(length(real.bv.add)>0)},
               if(is.list(real.bv.mult)){length(real.bv.mult)} else{as.numeric(length(real.bv.mult)>0)},
               if(is.list(real.bv.dice)){length(real.bv.dice)} else{as.numeric(length(real.bv.dice)>0)},
               if(length(trait_sum)>1){length(trait_sum)} else{0})

    if(nbv >= bv.total){
      bv.total <- nbv
      bv.calc <- nbv
      bv.random <- rep(FALSE, bv.total)
      bv.random.variance <- c(rep(0, nbv))
    }
    if(bv.total > nbv){
      if(length(polygenic.variance)< (bv.total - nbv)){
        polygenic.variance <- rep(polygenic.variance, bv.total - nbv)
      }
      bv.random <- c(rep(FALSE, nbv), rep(TRUE, bv.total - nbv))

      bv.random.variance <- c(rep(0, nbv), polygenic.variance)
      bv.calc <- nbv +1 ## WARUM STEHT HIER +1 SINN ist calc nicht gerade anzahl der ZW mit berechenbaren wert? # ALLES SO RICHTIG in breeding.diploid!
    }

    if(length(chr.nr)>0){
      chr.opt <- unique(chr.nr)
      nsnp <- numeric(length(chr.opt))
      for(index in 1:length(chr.opt)){
        nsnp[index] <- sum(chr.nr==chr.opt[index])
      }
    }
    if(length(dataset)==0 || (length(dataset)==1 && !is.list(dataset) && dataset=="random")){
      if(miraculix && length(chr.nr)==0 && miraculix.dataset){
        suppressWarnings(dataset <- list(miraculix::rhaplo(freq, indiv = nindi, loci = nsnp)))
      } else if(miraculix && miraculix.dataset){
        dataset <- list()
        for(chr_index in 1:length(chr.opt)){
          suppressWarnings(dataset[[chr_index]] <- miraculix::rhaplo(freq[which(chr.nr==chr.opt[chr_index])], indiv = nindi, loci = nsnp[chr_index]))
        }
      } else{
        dataset <- matrix((c(stats::rbinom(nindi*2*sum(nsnp),1,freq))),ncol=nindi*2, nrow=sum(nsnp))
      }
    }

    if(length(dataset)==1 && !is.list(dataset) && dataset=="all0"){
      if(miraculix && length(chr.nr)==0 && miraculix.dataset){
        suppressWarnings(dataset <- list(miraculix::rhaplo(0, indiv = nindi, loci = nsnp)))
      } else if(miraculix && miraculix.dataset){
        dataset <- list()
        for(chr_index in 1:length(chr.opt)){
          suppressWarnings(dataset[[chr_index]] <- miraculix::rhaplo(0, indiv = nindi, loci = nsnp[chr_index]))
        }
      } else{
        dataset <- matrix((c(rep(0,nindi*2*sum(nsnp)))),ncol=nindi*2, nrow=sum(nsnp))
      }

    }
    if(length(dataset)==1 && !is.list(dataset) && dataset=="homorandom"){
      if(miraculix && miraculix.dataset && length(chr.nr)==0){
        suppressWarnings(dataset <- list(miraculix::rhaplo(freq, indiv = nindi, loci = nsnp, freq2="IRGENDWAS")))
      } else if(miraculix && miraculix.dataset){
        dataset <- list()
        for(chr_index in 1:length(chr.opt)){
          suppressWarnings(dataset[[chr_index]] <- miraculix::rhaplo(freq[which(chr.nr==chr.opt[chr_index])], indiv = nindi, loci = nsnp[chr_index], freq2="IRGENDWAS"))
        }
      } else{
        dataset <- matrix((c(stats::rbinom(nindi*sum(nsnp),1,freq))),ncol=nindi, nrow=sum(nsnp))
        dataset <- dataset[,rep(1:nindi, each=2)]
      }
    }
    if(length(dataset)==1 && !is.list(dataset) && dataset=="allhetero"){
      if(miraculix && miraculix.dataset && length(chr.nr)==0){
        suppressWarnings(dataset <- list(miraculix::rhaplo(0, indiv = nindi, loci = nsnp,1)))
      } else if(miraculix && miraculix.dataset){
        dataset <- list()
        for(chr_index in 1:length(chr.opt)){
          suppressWarnings(dataset[[chr_index]] <- miraculix::rhaplo(0, indiv = nindi, loci = nsnp[chr_index],1))
        }
      } else{
        dataset <- matrix((c(rep(0,nindi*2*sum(nsnp)))),ncol=nindi*2, nrow=sum(nsnp))
        dataset[,1:nindi*2] <- 1
      }
    }

    if(length(nsnp)<2){
      if(is.list(dataset)){
        nsnp <- numeric(length(dataset))
        for(index in 1:length(dataset)){
          nsnp[index] <- attr(dataset[[index]], "information")[2]
        }

      } else{
        if(length(chr.nr)>0){
          nsnp <- numeric(length(chr.opt))
          for(index in 1:length(chr.opt)){
            nsnp[index] <- sum(chr.nr==chr.opt[index])
          }
        } else{
          nsnp <- nrow(dataset)
        }
      }
    }
    if(is.list(dataset)){
      nindi <- attr(dataset[[1]], "information")[3]
    } else{
      nindi <- ncol(dataset)/2
    }


    if(change.order && length(snp.position)>0){
      if(miraculix && miraculix.dataset){
        stop("Change order has to be executed before dataset generation // is not possible for imported datasets")
      }
      if(length(chr.nr)>0 && length(unique(chr.nr))>0){
        dataset_temp <- dataset
        snp.position_temp <- snp.position
        bp_temp <- bp
        snp.name_temp <- snp.name
        chr.nr_temp <- chr.nr
        for(index in unique(chr.nr)){
          consider <- which(chr.nr==index)
          order <- sort(snp.position_temp, index.return=TRUE)$ix
          snp.position <- snp.position_temp[consider][order]
          chr.nr <- chr.nr_temp[consider][order]
          if(length(bp)==length(snp.position)){
            bp <- bp_temp[consider][order]
          }
          if(length(snp.name)==length(snp.position)){
            snp.name <- snp.name_temp[consider][order]
          }

        }
      } else{
        order <- sort(snp.position,index.return=TRUE)$ix
        snp.position <- snp.position[order]
        dataset <- dataset[order,]
        if(length(bp)==length(snp.position)){
          bp <- bp[order]
        }
        if(length(snp.name)==length(snp.position)){
          snp.name <- snp.name[order]
        }
      }

    }

    if(length(genotyped.s)==0){
      genotyped.s <- stats::rbinom(nindi, 1, share.genotyped)
    }
    if(sex.s[1]=="random"){

      sex.s <- stats::rbinom(nindi, 1,sex.quota) +1
      if(add.chromosome==TRUE){
        sex.s <- population$info$sex
      }
    }
    if(sex.s[1]=="fixed"){
      sex.s <- rep(1, nindi*(1-sex.quota))
      sex.s <- c(sex.s, rep(2, nindi - length(sex.s)))
    }


    if(length(hom0)==0 && (is.list(dataset) || is.numeric(dataset[1,1]))){
      hom0 <- integer(sum(nsnp))
      hom1 <- rep(1L,sum(nsnp))
    }

  } else{
    if(is.list(dataset)){
      nsnp <- attr(dataset[[1]], "information")[2]
      nindi <- attr(dataset[[1]], "information")[3]
    } else{
      nsnp <- nrow(dataset)
      nindi <- ncol(dataset)/2
    }

  }

  if(length(chr.nr)==0){
    chr.nr <- numeric(sum(nsnp))
    start1 <- 1
    for(index in 1:length(nsnp)){
      if(nsnp[index]!=0){
        chr.nr[start1:(start1+nsnp[index]-1)] <- index
        start1 <- start1 + nsnp[index]
      }

    }
  }
  if(length(bp)==0){
    bp <- numeric(sum(nsnp))
    start1 <- 1
    for(index in 1:length(nsnp)){
      bp[start1:(start1+nsnp[index]-1)] <- 1:nsnp[index]
      start1 <- start1 + nsnp[index]
    }
  }
  if(length(snp.name)==0){
    nsnpnr <- numeric(sum(nsnp))
    start1 <- 1
    for(index in 1:length(nsnp)){
      nsnpnr[start1:(start1+nsnp[index]-1)] <- 1:nsnp[index]
      start1 <- start1 + nsnp[index]
    }
    snp.name <- paste0("Chr", chr.nr, "SNP", nsnpnr)
  }

  chr.opt <- unique(chr.nr)

  if(length(chr.opt)==1){
    if(bpcm.conversion>0 && length(snp.position)==0){
      snp.position <- as.numeric(bp) / bpcm.conversion
      chromosome.length <- max(snp.position) - min(snp.position)
    } else if(bpcm.conversion>0 && length(snp.position)>0){
      cat("Do not use bpcm.conversion and snp.position jointly!\n")
    }
  }
  if(length(bpcm.conversion)!=length(chr.opt)){
    bpcm.conversion <- rep(bpcm.conversion, length.out=length(chr.opt))
  }


  if(is.matrix(dataset) && is.numeric(dataset[1,1])){
    if(storage.mode(dataset)!="integer"){
      storage.mode(dataset) <- "integer"
    }
  } else if(is.matrix(dataset)){
    if(length(hom0)==0){
      hom0 <- dataset[,1]
    }
    if(length(hom1)==0){
      hom1 <- dataset[,1]
    }
    dataset <- (dataset==hom1)
    storage.mode(dataset) <- "integer"
  }

  if(length(snp.position)>0 && snp.position[1]<=0 && (length(snps.equidistant)==0 || snps.equidistant!=TRUE)){
    snp.position[1] <- snp.position[2]/2
    print(paste("Illegal position for SNP 1 - changed to",snp.position[1]))
  }
  mindex <- max(which(chr.opt[1]==chr.nr))
  if(length(snp.position)>1 && snp.position[mindex]>=chromosome.length[1] && (length(snps.equidistant)==0 || snps.equidistant!=TRUE)){
    snp.position[mindex] <- mean(c(snp.position[mindex-1], chromosome.length[1]))
    print(paste("Illegal position for last SNP - changed to",snp.position[mindex]))
  }

  position <- snp.position
  if(length(snp.position)>0 && length(snps.equidistant)==0){
    snps.equidistant <- FALSE
  } else if(length(snps.equidistant)==0){
    snps.equidistant <- TRUE
  }

  if(length(chr.opt)==1 && (snps.equidistant || position.scaling || max(position) > sum(chromosome.length) )){
    if(snps.equidistant){
      position <- 0.5:(sum(nsnp))*10
    }
    min.p <- min(position)
    max.p <- max(position)
    #  position.scal <- (position - (min.p - length.before)) / (max.p-min.p + length.before+ length.behind) * chromosome.length
    position <- (position) / (max.p-min.p + length.before+ length.behind) * chromosome.length[1]

  }

  if(length(chr.opt)!=length(chromosome.length)){
    if(length(snp.position)==length(chr.nr)){
      chromosome.length <- numeric(length(chr.opt))
      for(index in 1:length(chr.opt)){
        chromosome.length[index] <- max(snp.position[chr.nr==chr.opt[index]]) + min(snp.position[chr.nr==chr.opt[index]])
      }
    } else{
      chromosome.length <- rep(chromosome.length, length.out=length(chr.opt))
    }

  }

  if(length(chr.opt)==1){
    if(length(population)==0){
      population <- list()

      ## KEINE EDITS ZWISCHEN [[1]] und [[18]] snps.equidistant ++ Miraculix aenderungen sonst erforderlich!
      population$info <- list()
      population$info$schlather.slot1 <- "miraculix_not_activated"
      population$info$chromosome <- 1L
      population$info$snp <- as.integer(nsnp)
      population$info$position <- list()
      population$info$position[[1]] <- position
      population$info$snp.base <- rbind(hom0,hom1, deparse.level = 0)
      population$info$snp.position <- position
      population$info$length <- chromosome.length
      population$info$length.total <- c(0,population$info$length)
      population$info$func <- FALSE
      population$info$size <- matrix(0L,nrow=1, ncol=2)
      population$info$bve <- FALSE
      population$info$bv.calculated <- FALSE
      population$info$breeding.totals <- list()
      population$info$bve.data <- list()
      population$info$bv.nr <- bv.total
      population$info$bv.random <- bv.random
      population$info$bv.random.variance <- bv.random.variance
      population$info$snps.equidistant <- snps.equidistant
      population$info$origin.gen <- 1L
      population$info$cumsnp <- nsnp
      population$info$bp <- bp
      population$info$snp.name <- snp.name
      population$info$next.animal <- 1
      population$info$culling.stats <- list()
      if(length(bve.mult.factor)==0){
        population$info$bve.mult.factor <- rep(1L, bv.total)
      } else{
        population$info$bve.mult.factor <- bve.mult.factor
      }
      if(length(bve.poly.factor)==0){
        population$info$bve.poly.factor <- rep(1L, bv.total)
      } else{
        population$info$bve.poly.factor <- bve.poly.factor
      }
      if(length(base.bv)==0){
        population$info$base.bv <- rep(100L, bv.total)
      } else{
        population$info$base.bv <- base.bv
      }


    } else if(add.chromosome==TRUE){
      if(length(chr.opt)>1){
        stop("You can only add a single chromosome using add.chromosome!")
      }
      population$info$chromosome <- population$info$chromosome + 1L
      population$info$snp <- c(population$info$snp, as.integer(nsnp))
      population$info$position[[length(population$info$position)+1]] <- position
      population$info$snp.position <- c(population$info$snp.position, position + max(population$info$length.total))
      population$info$length <- c(population$info$length, chromosome.length)
      population$info$length.total <- cumsum(c(0,population$info$length))
      population$info$snp.base <- cbind(population$info$snp.base , rbind(hom0,hom1, deparse.level = 0), deparse.level = 0)
      population$info$cumsnp <- c(population$info$cumsnp, sum(population$info$snp))
      population$info$bp <- c(population$info$bp, bp)
      population$info$snp.name <- c(population$info$snp.name, snp.name)
    }
    if(generation!=1){
      take <- which(population$info$origin.gen==generation)
      if(length(take)==1){
        origin_code <- population$info$origin.gen[take]
      } else{
        if(length(population$info$origin.gen)<64){
          population$info$origin.gen <- c(population$info$origin.gen, as.integer(generation))
          origin_code <- length(population$info$origin.gen)
        } else{
          print("To many origin generation!")
          print("Delete second lowest origin.gen")
          switch <- sort(population$info$origin.gen, index.return=TRUE)[[2]]
          population$info$origin.gen[switch] <- as.integer(generation)
          origin_code <- switch
        }
      }
    } else{
      origin_code <- generation
    }

    counter <- c(length(population$breeding[[generation]][[1]]),length(population$breeding[[generation]][[2]]))+1L # maennlich/weibliche Tiere bisher
    counter.start <- counter

    if(length(population)==1){
      population$breeding <- list()
      population$breeding[[1]] <- list()
    }
    if(length(population$breeding)==0 || length(population$breeding[[1]])==0){
      population$breeding[[1]][[1]] <- list()
    }
    if(length(population$breeding[[1]])==1 ){
      population$breeding[[1]][[2]] <- list()
    }

    if(generation!=1){
      if(length(population$breeding)==(generation-1) || length(population$breeding[[generation]])==0){
        population$breeding[[generation]] <- list()
        population$breeding[[generation]][[1]] <- list()
        population$info$size <- rbind(population$info$size,0L, deparse.level = 0)
      }
      if(length(population$breeding[[generation]])==1){
        population$breeding[[generation]][[2]] <- list()
      }
    }

    if(add.chromosome==FALSE){

      for(index in 1:length(sex.s)){
        sex <- sex.s[index]
        genotyped <- genotyped.s[index]

        population$breeding[[generation]][[sex]][[counter[sex]]] <- list()
        population$breeding[[generation]][[sex]][[counter[sex]]][[1]] <- c(0, sum(population$info$length))
        population$breeding[[generation]][[sex]][[counter[sex]]][[2]] <- c(0,sum(population$info$length))
        if(add.chromosome.ends==TRUE){
          population$breeding[[generation]][[sex]][[counter[sex]]][[1]] <- population$info$length.total
          population$breeding[[generation]][[sex]][[counter[sex]]][[2]] <- population$info$length.total
        }
        population$breeding[[generation]][[sex]][[counter[sex]]][[3]] <- NULL
        population$breeding[[generation]][[sex]][[counter[sex]]][[4]] <- NULL
        population$breeding[[generation]][[sex]][[counter[sex]]][[5]] <- codeOriginsU(matrix(c(origin_code, sex, counter[sex], 1),nrow=(length(population$breeding[[generation]][[sex]][[counter[sex]]][[1]])-1), ncol=4, byrow=TRUE))
        population$breeding[[generation]][[sex]][[counter[sex]]][[6]] <- codeOriginsU(matrix(c(origin_code, sex, counter[sex], 2),nrow=(length(population$breeding[[generation]][[sex]][[counter[sex]]][[2]])-1), ncol=4, byrow=TRUE))
        population$breeding[[generation]][[sex]][[counter[sex]]][[7]] <- c(generation, sex, counter[sex])
        population$breeding[[generation]][[sex]][[counter[sex]]][[8]] <- c(generation, sex, counter[sex])

        if(miraculix && miraculix.dataset){
          population$breeding[[generation]][[sex]][[counter[sex]]][[9]] <- miraculix::haplomatrix(as.matrix(dataset[[1]],indiv = index))
          population$breeding[[generation]][[sex]][[counter[sex]]][[10]] <- "Placeholder_Pointer_Martin"
        } else if(miraculix){
          population$breeding[[generation]][[sex]][[counter[sex]]][[9]] <- miraculix::haplomatrix(dataset[,(index*2-c(1,0))])
          population$breeding[[generation]][[sex]][[counter[sex]]][[10]] <- "Placeholder_Pointer_Martin"
        } else if(bit.storing){
          population$breeding[[generation]][[sex]][[counter[sex]]][[9]] <- bit.storing(dataset[,(index*2-1)], nbits)
          population$breeding[[generation]][[sex]][[counter[sex]]][[10]] <- bit.storing(dataset[,(index*2)], nbits)
        } else{
          population$breeding[[generation]][[sex]][[counter[sex]]][[9]] <- dataset[,(index*2-1)]
          population$breeding[[generation]][[sex]][[counter[sex]]][[10]] <- dataset[,(index*2)]
        }

        population$breeding[[generation]][[sex]][[counter[sex]]][[11]] <- NULL
        population$breeding[[generation]][[sex]][[counter[sex]]][[12]] <- NULL
        #      population$breeding[[generation]][[sex]][[counter[sex]]][[13]] <- "test"
        population$breeding[[generation]][[sex]][[counter[sex]]][[15]] <- rep(0, population$info$bv.nr)
        population$breeding[[generation]][[sex]][[counter[sex]]][[16]] <- genotyped
        population$breeding[[generation]][[sex]][[counter[sex]]][[21]] <- cbind(generation, sex, counter[sex])
        population$info$size[generation,sex] <- population$info$size[generation,sex] +1L
        counter[sex] <- counter[sex] + 1L
      }

      if(length(population$breeding[[generation]])==2){
        population$breeding[[generation]][[3]] <- matrix(0, nrow= population$info$bv.nr, ncol=counter[1]-1) # Selektionsfunktion
        population$breeding[[generation]][[4]] <- matrix(0, nrow= population$info$bv.nr, ncol=counter[2]-1)
        population$breeding[[generation]][[5]] <- rep(class,counter[1]-1) # Migrationslevel
        population$breeding[[generation]][[6]] <- rep(class,counter[2]-1)
        population$breeding[[generation]][[7]] <- matrix(0, nrow= population$info$bv.nr, ncol=counter[1]-1) # realer ZW
        population$breeding[[generation]][[8]] <- matrix(0, nrow= population$info$bv.nr, ncol=counter[2]-1)
        population$breeding[[generation]][[9]] <- matrix(0, nrow= population$info$bv.nr, ncol=counter[1]-1) # geschaetzer ZW
        population$breeding[[generation]][[10]] <- matrix(0, nrow= population$info$bv.nr, ncol=counter[2]-1)
        population$breeding[[generation]][[11]] <- rep(time.point,counter[1]-1) # Time point
        population$breeding[[generation]][[12]] <- rep(time.point,counter[2]-1)
        population$breeding[[generation]][[13]] <- rep(creating.type,counter[1]-1) # Creating.type
        population$breeding[[generation]][[14]] <- rep(creating.type,counter[2]-1)
        population$breeding[[generation]][[15]] <- seq(population$info$next.animal, population$info$next.animal + counter[1] -2, length.out= counter[1] -1)
        population$info$next.animal <- population$info$next.animal + counter[1] -1
        population$breeding[[generation]][[16]] <- seq(population$info$next.animal, population$info$next.animal + counter[2] -2, length.out= counter[2] -1)
        population$info$next.animal <- population$info$next.animal + counter[2] -1
        population$breeding[[generation]][[17]] <- rep(NA,counter[1]-1) # Time of death point
        population$breeding[[generation]][[18]] <- rep(NA,counter[2]-1)
        population$breeding[[generation]][[19]] <- matrix(0, nrow= population$info$bv.nr, ncol=counter[1]-1) # Reliabilities
        population$breeding[[generation]][[20]] <- matrix(0, nrow= population$info$bv.nr, ncol=counter[2]-1)
        population$breeding[[generation]][[21]] <- matrix(0, nrow= population$info$bv.nr, ncol=counter[1]-1) # Last applied selection index
        population$breeding[[generation]][[22]] <- matrix(0, nrow= population$info$bv.nr, ncol=counter[2]-1)

        # calculate Real-ZW
      } else{
        population$breeding[[generation]][[3]] <- cbind(population$breeding[[generation]][[3]], matrix(0, nrow= population$info$bv.nr, ncol=counter[1]-counter.start[1])) # Selektionsfunktion
        population$breeding[[generation]][[4]] <- cbind(population$breeding[[generation]][[4]], matrix(0, nrow= population$info$bv.nr, ncol=counter[2]-counter.start[2]))
        population$breeding[[generation]][[5]] <- c(population$breeding[[generation]][[5]], rep(class ,counter[1]-counter.start[1])) # Migrationslevel
        population$breeding[[generation]][[6]] <- c(population$breeding[[generation]][[6]], rep(class ,counter[2]-counter.start[2]))
        population$breeding[[generation]][[7]] <- cbind(population$breeding[[generation]][[7]] , matrix(0, nrow= population$info$bv.nr, ncol=counter[1]-counter.start[1])) # realer ZW
        population$breeding[[generation]][[8]] <- cbind(population$breeding[[generation]][[8]] , matrix(0, nrow= population$info$bv.nr, ncol=counter[2]-counter.start[2]))
        population$breeding[[generation]][[9]] <- cbind(population$breeding[[generation]][[9]] , matrix(0, nrow= population$info$bv.nr, ncol=counter[1]-counter.start[1])) # geschaetzer ZW
        population$breeding[[generation]][[10]] <-cbind(population$breeding[[generation]][[10]] , matrix(0, nrow= population$info$bv.nr, ncol=counter[2]-counter.start[2]))
        population$breeding[[generation]][[11]] <- c(population$breeding[[generation]][[11]], rep(time.point ,counter[1]-counter.start[1])) # Time point
        population$breeding[[generation]][[12]] <- c(population$breeding[[generation]][[12]], rep(time.point ,counter[2]-counter.start[2]))
        population$breeding[[generation]][[13]] <- c(population$breeding[[generation]][[13]], rep(time.point ,counter[1]-counter.start[1])) # Creating type
        population$breeding[[generation]][[14]] <- c(population$breeding[[generation]][[14]], rep(time.point ,counter[2]-counter.start[2]))

        population$breeding[[generation]][[15]] <- c(population$breeding[[generation]][[15]] , seq(population$info$next.animal, population$info$next.animal + counter[1] -2, length.out= counter[1] -1))
        population$info$next.animal <- population$info$next.animal + counter[1] -1
        population$breeding[[generation]][[16]] <- c(population$breeding[[generation]][[16]] , seq(population$info$next.animal, population$info$next.animal + counter[2] -2, length.out= counter[2] -1))
        population$info$next.animal <- population$info$next.animal + counter[2] -1
        population$breeding[[generation]][[17]] <- c(population$breeding[[generation]][[17]], rep(NA ,counter[1]-counter.start[1])) # Time of death
        population$breeding[[generation]][[18]] <- c(population$breeding[[generation]][[18]], rep(NA ,counter[2]-counter.start[2]))
        population$breeding[[generation]][[19]] <- cbind(population$breeding[[generation]][[19]], matrix(0, nrow= population$info$bv.nr, ncol=counter[1]-counter.start[1])) # Reliabilities
        population$breeding[[generation]][[20]] <- cbind(population$breeding[[generation]][[20]], matrix(0, nrow= population$info$bv.nr, ncol=counter[2]-counter.start[2]))
        population$breeding[[generation]][[21]] <- cbind(population$breeding[[generation]][[21]], matrix(0, nrow= population$info$bv.nr, ncol=counter[1]-counter.start[1])) # Last applied selection index
        population$breeding[[generation]][[22]] <- cbind(population$breeding[[generation]][[22]], matrix(0, nrow= population$info$bv.nr, ncol=counter[2]-counter.start[2]))


      }

      population$info$sex <- c(population$info$sex, sex.s)


    } else{
      counter <- c(1,1)
      for(index in 1:length(sex.s)){
        sex <- sex.s[index]
        genotyped <- genotyped.s[index]
        population$breeding[[generation]][[sex]][[counter[sex]]][[1]][2] <- sum(population$info$length)
        population$breeding[[generation]][[sex]][[counter[sex]]][[2]][2] <- sum(population$info$length)
        if(add.chromosome.ends==TRUE){
          population$breeding[[generation]][[sex]][[counter[sex]]][[1]] <- population$info$length.total
          population$breeding[[generation]][[sex]][[counter[sex]]][[2]] <- population$info$length.total
          population$breeding[[generation]][[sex]][[counter[sex]]][[5]] <- codeOriginsU(matrix(c(origin_code, sex, counter[sex], 1),nrow=(length(population$breeding[[generation]][[sex]][[counter[sex]]][[1]])-1), ncol=4, byrow=TRUE))
          population$breeding[[generation]][[sex]][[counter[sex]]][[6]] <- codeOriginsU(matrix(c(origin_code, sex, counter[sex], 2),nrow=(length(population$breeding[[generation]][[sex]][[counter[sex]]][[2]])-1), ncol=4, byrow=TRUE))

        }
        population$breeding[[generation]][[sex]][[counter[sex]]][[7]] <- c(generation, sex, counter[sex])
        population$breeding[[generation]][[sex]][[counter[sex]]][[8]] <- c(generation, sex, counter[sex])
        if(miraculix && miraculix.dataset){
          population$breeding[[generation]][[sex]][[counter[sex]]][[9]] <- miraculix::haplomatrix(rbind(as.matrix(population$breeding[[generation]][[sex]][[counter[sex]]][[9]]),as.matrix(dataset[[1]], index)))
          population$breeding[[generation]][[sex]][[counter[sex]]][[10]] <- "Placeholder_Pointer_Martin"
        } else if(miraculix){
          population$breeding[[generation]][[sex]][[counter[sex]]][[9]] <- miraculix::haplomatrix(rbind(as.matrix(population$breeding[[generation]][[sex]][[counter[sex]]][[9]]),dataset[,(index*2-c(1,0))]))
          population$breeding[[generation]][[sex]][[counter[sex]]][[10]] <- "Placeholder_Pointer_Martin"
        } else if(bit.storing){
          if(leftover==0){
            population$breeding[[generation]][[sex]][[counter[sex]]][[9]] <- c(population$breeding[[generation]][[sex]][[counter[sex]]][[9]], bit.storing(dataset[,(index*2-1)]),nbits)
            population$breeding[[generation]][[sex]][[counter[sex]]][[10]] <- c(population$breeding[[generation]][[sex]][[counter[sex]]][[10]], bit.storing(dataset[,(index*2)]), nbits)
          } else{
            population$breeding[[generation]][[sex]][[counter[sex]]][[9]] <- c(population$breeding[[generation]][[sex]][[counter[sex]]][[9]][-length(population$breeding[[generation]][[sex]][[counter[sex]]][[9]])],
                                                                                     bit.storing(c(bit.snps(population$breeding[[generation]][[sex]][[counter[sex]]][[9]][length(population$breeding[[generation]][[sex]][[counter[sex]]][[9]])], nbits)[(nbits-leftover+1):nbits],dataset[,(index*2-1)]),nbits))
            population$breeding[[generation]][[sex]][[counter[sex]]][[10]] <- c(population$breeding[[generation]][[sex]][[counter[sex]]][[10]][-length(population$breeding[[generation]][[sex]][[counter[sex]]][[10]])],
                                                                                      bit.storing(c(bit.snps(population$breeding[[generation]][[sex]][[counter[sex]]][[10]][length(population$breeding[[generation]][[sex]][[counter[sex]]][[10]])], nbits)[(nbits-leftover+1):nbits],dataset[,(index*2)]),nbits))

          }
        } else{
          population$breeding[[generation]][[sex]][[counter[sex]]][[9]] <- c(population$breeding[[generation]][[sex]][[counter[sex]]][[9]], dataset[,(index*2-1)])
          population$breeding[[generation]][[sex]][[counter[sex]]][[10]] <- c(population$breeding[[generation]][[sex]][[counter[sex]]][[10]], dataset[,(index*2)])

        }

        counter[sex] <- counter[sex] + 1
      }

    }


    if(skip.rest==FALSE){
      if(bv.total>0 ||length(real.bv.add)>0  || length(real.bv.mult) >0 || length(real.bv.dice)>0){
        population$info$bve <- TRUE
        if(is.list(real.bv.add)){
          population$info$real.bv.add <- real.bv.add
        } else{
          population$info$real.bv.add <- list(real.bv.add)
        }
        if(is.list(real.bv.mult)){
          population$info$real.bv.mult <- real.bv.mult
        } else{
          population$info$real.bv.mult <- list(real.bv.mult)
        }
        if(is.list(real.bv.dice)){
          population$info$real.bv.dice <- real.bv.dice
        } else{
          if(length(real.bv.dice)>0){
            print("Keine vorschriftmaessige Eingabe fuer real.bv.dice!")
          }
          population$info$real.bv.dice <- list(real.bv.dice)
        }

        population$info$bv.nr <- bv.total
        population$info$bv.calc <- bv.calc

        population$info$real.bv.length <- c(length(population$info$real.bv.add),
                                            length(population$info$real.bv.mult),
                                            length(population$info$real.bv.dice))

        population$info$real.bv.add[[nbv+1]] <- "placeholder" # Use nbv instead of bv.calc
        population$info$real.bv.mult[[nbv+1]] <- "placeholder"
        population$info$real.bv.dice[[nbv+1]] <- "placeholder"


      } else if(perserve_bve){
        population$info$bve <- FALSE
        population$info$bv.nr <- 0
        population$info$bv.calc <- 0
        population$info$real.bv.length <- c(0,0,0)
      }

      if(bv.total>0){
        population$info$pheno.correlation <- diag(1L, bv.total)
      }
      if(length(new.phenotype.correlation)>0){
        population$info$pheno.correlation <- t(chol(new.phenotype.correlation))
      }
      if(bv.total>0){
        population$info$bv.correlation <- diag(1L, bv.total)
      }
      if(length(new.breeding.correlation)>0){
        population$info$bv.correlation <- new.breeding.correlation
      }


      if(length(shuffle.traits)>0){
        if(length(shuffle.traits)==1){
          shuffle.traits <- which(population$info$bv.random==FALSE)
        }
        LT <- chol(shuffle.cor)
        if(nrow(LT)!=length(shuffle.traits)){
          stop("Dimension of shuffle correlation matrix doesnt work with traits to shuffle")
        } else{

          population$info$bv.correlation[shuffle.traits,shuffle.traits] <- t(LT) %*% LT
          if(sum(abs(population$info$bv.correlation[shuffle.traits,shuffle.traits]- shuffle.cor))>0.0001){
            print("No-covariance matrix for traits given! Values above diagonal used.")
          }

          store.add <- population$info$real.bv.add
          store.mult <- population$info$real.bv.mult
          store.dice <- population$info$real.bv.dice


          col <- 1
          for(index in shuffle.traits){
            new.add <- new.mult <- new.dice1 <- new.dice2 <- NULL
            row <- 1
            for(index2 in shuffle.traits){
              if(length(store.add[[index2]])>0){
                new.add <- rbind(new.add, store.add[[index2]] %*% diag(c(1,1,rep(LT[row,col],3))))
                zeros <- rowSums(abs(new.add[,3:5, drop=FALSE]))
                new.add <- new.add[zeros>0,,drop=FALSE]
              }
              if(length(store.mult[[index2]])>0){
                new.mult <- rbind(new.mult, store.mult[[index2]] %*% diag(c(1,1,1,1,rep(LT[row,col],9))))
                zeros <- rowSums(abs(new.mult[,5:13, drop=FALSE]))
                new.mult <- new.add[zeros>0,,drop=FALSE]
              }
              if(length(store.dice[[index2]])>0){
                before <- length(new.dice2)
                new.dice1 <- c(new.dice1,store.dice[[index2]][[1]])
                new.dice2 <- c(new.dice2,store.dice[[index2]][[2]])
                for(index3 in (before+1):length(new.dice2)){
                  new.dice2[[index3]] <- new.dice2[[index3]] * LT[row,col]
                }
              }
              row <- row +1
            }
            # DONT REMOVE NULL - MORE WORK NEEDED HERE!
            if(length(new.add)==0){

            } else{
              population$info$real.bv.add[[index]] <- new.add
            }
            if(length(new.mult)==0){

            } else{
              population$info$real.bv.mult[[index]] <- new.mult
            }
            if(length(new.add)==0){

            } else{
              population$info$real.bv.dice[[index]] <- list(new.dice1,new.dice2)
            }
            col <- col +1
          }

        }

        for(index in shuffle.traits){
          population$info$real.bv.length[1] <- max(population$info$real.bv.length[1], if(length(population$info$real.bv.add[[index]])>0){index} else{0})
          population$info$real.bv.length[2] <- max(population$info$real.bv.length[2], if(length(population$info$real.bv.mult[[index]])>0){index} else{0})
          population$info$real.bv.length[3] <- max(population$info$real.bv.length[3], if(length(population$info$real.bv.dice[[index]][[1]])>0){index} else{0})
        }
      }


      if(length(add.architecture)>0){
        population$info$gen.architecture[[length(population$info$gen.architecture)+1]] <- list()
        population$info$gen.architecture[[length(population$info$gen.architecture)]]$length.total <- cumsum(c(0,add.architecture[[1]]))
        population$info$gen.architecture[[length(population$info$gen.architecture)]]$snp.position <- add.architecture[[2]]

      }
      if(bit.storing){
        population$info$bitstoring <- nbits
        population$info$leftover <-  sum(population$info$snp)%%nbits
      }
      if(miraculix || miraculix.dataset){
        population$info$miraculix <- TRUE
      } else{
        population$info$miraculix <- FALSE
      }

    }
    if(length(name.cohort)>=1 && add.chromosome==FALSE){

      if((counter-counter.start)[1]>0 && (counter-counter.start)[2]>0){
        population$info$cohorts <- rbind(population$info$cohorts, c(paste0(name.cohort, "_M"), generation, (counter - counter.start)[1], 0, class, counter.start[1], 0,
                                                                    time.point, creating.type),
                                                                  c(paste0(name.cohort, "_F"), generation, 0, (counter - counter.start)[2], class, 0, counter.start[2],
                                                                    time.point, creating.type))

        cat("Both genders in the cohort. Added _M, _F to cohort names!\n")
      } else{
        population$info$cohorts <- rbind(population$info$cohorts, c(name.cohort, generation, counter - counter.start, class, counter.start,
                                                                    time.point, creating.type))
      }
      if(nrow(population$info$cohorts)<=2){
        colnames(population$info$cohorts) <- c("name","generation", "male individuals", "female individuals", "class", "position first male", "position first female",
                                               "time point", "creating.type")
      }
    }


  } else{
    if(length(population)==0 || add.chromosome==TRUE){
      for(chr_index in 1:length(chr.opt)){
        index <- chr.opt[chr_index]
        activ <- which(chr.nr==index)
        chr_activ <- chr.nr[activ]
        bp_activ <- bp[activ]
        snp.name_activ <- snp.name[activ]
        hom0_activ <- hom0[activ]
        hom1_activ <- hom1[activ]
        if(is.list(dataset)){
          dataset_activ <- dataset[[chr_index]]
        } else{
          dataset_activ <- dataset[activ,,drop=FALSE]
        }

        snp.position_activ <- position[activ]
        if(length(freq)>1){
          freq_activ <- freq[activ]
        } else{
          freq_activ <- freq
        }

        if(chr_index==1){
          skip.rest <- FALSE
          add.chromosome <- add.chromosome
        } else{
          skip.rest <- TRUE
          add.chromosome <- TRUE
        }
        if(add.chromosome==FALSE){
          name.cohort <- name.cohort
        } else{
          name.cohort <- NULL
        }

        population <- creating.diploid(population=population, dataset=dataset_activ,
                                       nsnp=nsnp[chr_index], nindi=nindi,
                                       add.chromosome=add.chromosome, chr.nr = chr_activ,
                                       bp= bp_activ, snp.name = snp.name_activ,
                                       hom0= hom0_activ, hom1 = hom1_activ,
                                       class = class,
                                       generation = generation,
                                       add.chromosome.ends = add.chromosome.ends,
                                       miraculix = miraculix,
                                       miraculix.dataset = miraculix.dataset,
                                       snp.position = if(bpcm.conversion[chr_index]==0){snp.position_activ} else NULL,
                                       snps.equidistant= snps.equidistant,
                                       position.scaling= position.scaling,
                                       chromosome.length= chromosome.length[chr_index],
                                       length.before = length.before,
                                       length.behind = length.behind,
                                       skip.rest = skip.rest,
                                       sex.s = sex.s,
                                       genotyped.s = genotyped.s,
                                       name.cohort = name.cohort,
                                       real.bv.add = real.bv.add,
                                       real.bv.mult = real.bv.mult,
                                       real.bv.dice = real.bv.dice,
                                       freq = freq_activ,
                                       bpcm.conversion = bpcm.conversion[chr_index],
                                       remove.invalid.qtl=FALSE)
      }
    } else{
      if(min(diff(chr.nr))<0 || !miraculix.dataset){
        dataset_temp <- dataset
        till <- 0
        for(chr_index in 1:length(chr.opt)){
          index <- chr.opt[chr_index]
          activ <- which(chr.nr==index)
          chr_activ <- chr.nr[activ]
          bp_activ <- bp[activ]
          snp.name_activ <- snp.name[activ]
          hom0_activ <- hom0[activ]
          hom1_activ <- hom1[activ]
          dataset_activ <- dataset[activ,]


          snp.position_activ <- position[activ]
          if(length(activ)>0){
            dataset[1:length(activ)+till,] <- dataset_temp[activ,]
          }
          if((dataset[1+till,1]!=hom0_activ[1] && dataset[1+till,1]!=hom1_activ[1])){
            dataset[1:length(activ)+till,][dataset[1:length(activ)+till,]==hom0_activ] <- 0
            dataset[1:length(activ)+till,][dataset[1:length(activ)+till,]==hom1_activ] <- 1
          }
          till <- till + length(activ)

        }
      } else{
        if((dataset[1,1]!=hom0[1] && dataset[1,1]!=hom1[1]) && !miraculix.dataset){
          dataset[dataset==hom0_activ] <- 0
          dataset[dataset==hom1_activ] <- 1
        }
      }
      skip.rest <- TRUE

      population <- creating.diploid(population=population, dataset=dataset,
                                       class = class,
                                       generation = generation,
                                       add.chromosome.ends = add.chromosome.ends,
                                       miraculix = miraculix,
                                       skip.rest = skip.rest,
                                       sex.s = sex.s,
                                       genotyped.s = genotyped.s,
                                       name.cohort = name.cohort,
                                       real.bv.add = real.bv.add,
                                       real.bv.mult = real.bv.mult,
                                       real.bv.dice = real.bv.dice,
                                       hom0 =population$info$snp.base[1,],
                                       hom1 =population$info$snp.base[2,],
                                       remove.invalid.qtl =FALSE)
    }

  }

  if(bv.total){
    population$info$trait.name <- trait.name
    if(length(trait.name)<bv.total){
      population$info$trait.name <- c(population$info$trait.name, paste0("Trait ", (length(trait.name)+1):bv.total))
    }
  }
  if(length(population$info$real.bv.add)==0){
    population$info$real.bv.add <- list()
    population$info$real.bv.mult <- list()
    population$info$real.bv.dice <- list()
    population$info$real.bv.add[[1]] <- "placeholder" # Use nbv instead of bv.calc
    population$info$real.bv.mult[[1]] <- "placeholder"
    population$info$real.bv.dice[[1]] <- "placeholder"
  }
  if(remove.invalid.qtl && length(population$info$real.bv.add)>1){
    for(index in 1:(length(population$info$real.bv.add)-1)){
      removes <- which(population$info$real.bv.add[[index]][,1] > population$info$snp[population$info$real.bv.add[[index]][,2]])
      if(length(removes)>0){
        population$info$real.bv.add[[index]] <- population$info$real.bv.add[[index]][-removes,,drop=FALSE]
        cat(paste0(length(removes), " QTL-effects entered on markers that do not exist for ", population$info$trait.name[index], ".\n"))
        cat(paste0(nrow(population$info$real.bv.add[[index]]), " QTL-effects remain.\n"))
      }
    }
    for(index in 1:(length(population$info$real.bv.mult)-1)){
      removes <- which(population$info$real.bv.mult[[index]][,1] > population$info$snp[population$info$real.bv.mult[[index]][,2]])
      if(length(removes)>0){
        population$info$real.bv.mult[[index]] <- population$info$real.bv.mult[[index]][-removes,,drop=FALSE]
        cat(paste0(length(removes), " QTL-effects entered on markers that do not exist for ", population$info$trait.name[index], ".\n"))
        cat(paste0(nrow(population$info$real.bv.mult[[index]]), " QTL-effects remain.\n"))
      }
    }
    for(index in 1:(length(population$info$real.bv.mult)-1)){
      removes <- which(population$info$real.bv.mult[[index]][,3] > population$info$snp[population$info$real.bv.mult[[index]][,4]])
      if(length(removes)>0){
        population$info$real.bv.mult[[index]] <- population$info$real.bv.mult[[index]][-removes,,drop=FALSE]
        cat(paste0(length(removes), " QTL-effects entered on markers that do not exist for ", population$info$trait.name[index], ".\n"))
        cat(paste0(nrow(population$info$real.bv.mult[[index]]), " QTL-effects remain.\n"))
      }
    }

  }

  class(population) <- "population"
  return(population)
}
```
