## Supplementary material for "MoBPS - Modular Breeding Program Simulator": MoBPS R-package: derive.loop.elements.R

```
'#
  Authors
Torsten Pook,

Copyright (C) 2017 -- 2018  Torsten Pook

This program is free software; you can redistribute it and/or
modify it under the terms of the GNU General Public License
as published by the Free Software Foundation; either version 3
of the License, or (at your option) any later version.

This program is distributed in the hope that it will be useful,
but WITHOUT ANY WARRANTY; without even the implied warranty of
MERCHANTABILITY or FITNESS FOR A PARTICULAR PURPOSE.  See the
GNU General Public License for more details.

You should have received a copy of the GNU General Public License
along with this program; if not, write to the Free Software
Foundation, Inc., 59 Temple Place - Suite 330, Boston, MA  02111-1307, USA.
'#

#' Derive loop elements
#'
#' Internal function to derive the position of all individuals to consider for BVE/GWAS
#' @param population Population list
#' @param bve.database Groups of individuals to consider in breeding value estimation
#' @param bve.class Consider only animals of those class classes in breeding value estimation (default: NULL - use all)
#' @param bve.avoid.duplicates If set to FALSE multiple generatations of the same individual can be used in the bve (only possible by using copy.individual to generate individuals)
#' @param store.adding Internal parameter to derive number of added individuals per database entry (only relevant internally for GWAS)
#' @param store.which.adding Internal parameter to derive which individuals are copy entries
#' @param list.of.copys Internal parameter to derive further information on the copies individuals
#' @export


derive.loop.elements <- function(population, bve.database, bve.class, bve.avoid.duplicates, store.adding=FALSE,
                                 store.which.adding=FALSE, list.of.copys=FALSE){
  max.animals <- 0
  start <- non_start <- 1
  full_adding <- NULL
  for(index in 1:nrow(bve.database)){
    if(length(bve.class)>0){
      for(mig in bve.class){
        max.animals <- max.animals + sum(population$breeding[[bve.database[index,1]]][[bve.database[index,2]+4]][bve.database[3]:bve.database[4]]==mig)
      }
    } else{
      max.animals <- max.animals + diff(bve.database[index,3:4]) +1
    }
  }
  loop_elements <- matrix(nrow=max.animals, ncol=5)
  loop_elements[,1] <- 1:max.animals
  if(list.of.copys){
    copy_elements <- cbind(loop_elements,0)
  }

  used <- NULL
  prior <- non_prior <- 0
  prior3 <- 0
  for(index in 1:nrow(bve.database)){
    k.database <- bve.database[index,]
    if(length(bve.class)==0){
      news <- population$breeding[[k.database[1]]][[k.database[2]+14]][k.database[3]:k.database[4]]
      if(length(news)>0){
        if(bve.avoid.duplicates){
          to_add <- which(!duplicated(c(news,used), fromLast=TRUE)[1:length(news)])
        } else{
          to_add <- 1:length(news)
        }

        adding <- news[to_add]


        kn <- length(adding)
        start <- c(start, max(start) + kn)
        if(kn>0){
          if(store.which.adding){
            full_adding <- c(full_adding, to_add + prior3)
          }
          used <- c(used, adding)
          loop_elements[1:kn+ prior,2] <- (k.database[3]:k.database[4])[to_add]
          loop_elements[1:kn+ prior,3] <- index
          loop_elements[1:kn+ prior,4] <- k.database[1]
          loop_elements[1:kn+ prior,5] <- k.database[2]
          prior <- prior + kn
        }

        if(list.of.copys){
          if(length(to_add)==0){
            non_adding <- news
          } else{
            non_adding <- news[-to_add]
          }

          non_kn <- length(non_adding)

          non_start <- c(non_start, max(non_start) + non_kn)
          if(non_kn>0){
            if(length(to_add)==0){
              copy_elements[1:non_kn + non_prior,2] <- (k.database[3]:k.database[4])
            } else{
              copy_elements[1:non_kn + non_prior,2] <- (k.database[3]:k.database[4])[-to_add]
            }

            copy_elements[1:non_kn + non_prior,3] <- index
            copy_elements[1:non_kn + non_prior,4] <- k.database[1]
            copy_elements[1:non_kn + non_prior,5] <- k.database[2]

            for(check_id in 1:non_kn){
              copy_elements[check_id + non_prior,6] <- which(used==non_adding[check_id])
            }
            non_prior <- non_prior + non_kn


          }


        }

      }
    } else{
      start <- c(start, max(start))
      istart <- length(start)
      for(mig in bve.class){
        news <- population$breeding[[k.database[1]]][[k.database[2]+14]][k.database[3]:k.database[4]][population$breeding[[k.database[1]]][[k.database[2]+4]][k.database[3]:k.database[4]]==mig]
        if(length(news)>0){
          if(bve.avoid.duplicates){
            to_add <- which(!duplicated(c(news,used), fromLast=TRUE)[1:length(news)])
          } else{
            to_add <- 1:length(news)
          }

          adding <- news[to_add]
          kn <- length(adding)
          start[istart] <- start[istart] + kn
          if(kn>0){
            if(store.which.adding){
              full_adding <- c(full_adding, to_add + prior3)
            }
            used <- c(used, adding)
            loop_elements[1:kn+ prior,2] <- (k.database[3]:k.database[4])[population$breeding[[k.database[1]]][[k.database[2]+4]][k.database[3]:k.database[4]]==mig][to_add]
            loop_elements[1:kn+ prior,3] <- index
            loop_elements[1:kn+ prior,4] <- k.database[1]
            loop_elements[1:kn+ prior,5] <- k.database[2]
            prior <- prior + kn
          }

          if(list.of.copys){
            if(length(to_add)==0){
              non_adding <- news
            } else{
              non_adding <- news[-to_add]
            }

            non_kn <- length(non_adding)

            non_start <- c(non_start, max(non_start) + non_kn)
            if(non_kn>0){
              if(length(to_add)==0){
                copy_elements[1:non_kn + non_prior,2] <- (k.database[3]:k.database[4])
              } else{
                copy_elements[1:non_kn + non_prior,2] <- (k.database[3]:k.database[4])[-to_add]
              }

              copy_elements[1:non_kn + non_prior,3] <- index
              copy_elements[1:non_kn + non_prior,4] <- k.database[1]
              copy_elements[1:non_kn + non_prior,5] <- k.database[2]

              for(check_id in 1:non_kn){
                copy_elements[check_id + non_prior,6] <- which(used==non_adding[check_id])
              }
              non_prior <- non_prior + non_kn


            }


          }
        }
      }
    }
    prior3 <- prior3 + k.database[4]-k.database[3] +1

  }
  loop_elements <- loop_elements[1:prior,]

  output <- list()

  output[[1]] <- loop_elements
  if(store.adding){
    output[[length(output) +1]] <- start
  }
  if(store.which.adding){
    output[[length(output) +1]] <- full_adding
  }
  if(list.of.copys){
    if(non_prior>0){
      copy_elements <- copy_elements[1:non_prior,,drop=FALSE]
    } else{
      copy_elements <- copy_elements[0,,drop=FALSE]
    }

    output[[length(output) +1]] <- copy_elements
  }
  return(output)

}
```
