## Supplementary material for "MoBPS - Modular Breeding Program Simulator": MoBPS R-package: edges.fromto.R

```
'#
  Authors
Torsten Pook,

Copyright (C) 2017 -- 2018  Torsten Pook

This program is free software; you can redistribute it and/or
modify it under the terms of the GNU General Public License
as published by the Free Software Foundation; either version 3
of the License, or (at your option) any later version.

This program is distributed in the hope that it will be useful,
but WITHOUT ANY WARRANTY; without even the implied warranty of
MERCHANTABILITY or FITNESS FOR A PARTICULAR PURPOSE.  See the
GNU General Public License for more details.

You should have received a copy of the GNU General Public License
along with this program; if not, write to the Free Software
Foundation, Inc., 59 Temple Place - Suite 330, Boston, MA  02111-1307, USA.
'#

#' gen/database/cohorts conversion
#'
#' Function to derive a database based on gen/database/cohorts
#' @param edges Edges of the json-file generated via the web-interface
#' @export


edges.fromto<- function(edges){

  edges_fromto <- matrix(0, nrow=length(edges), ncol=2)
  for(index in 1:length(edges)){
    edges_fromto[index,] <- c(edges[[index]]$from, edges[[index]]$to)
  }

  return(edges_fromto)
}
```
