## Supplementary material for "MoBPS - Modular Breeding Program Simulator": MoBPS R-package: edit.animal.R

```
'#
  Authors
Torsten Pook,

Copyright (C) 2017 -- 2018  Torsten Pook

This program is free software; you can redistribute it and/or
modify it under the terms of the GNU General Public License
as published by the Free Software Foundation; either version 3
of the License, or (at your option) any later version.

This program is distributed in the hope that it will be useful,
but WITHOUT ANY WARRANTY; without even the implied warranty of
MERCHANTABILITY or FITNESS FOR A PARTICULAR PURPOSE.  See the
GNU General Public License for more details.

You should have received a copy of the GNU General Public License
along with this program; if not, write to the Free Software
Foundation, Inc., 59 Temple Place - Suite 330, Boston, MA  02111-1307, USA.
'#

#' Internal gene editing function
#'
#' Internal function to perform gene editing
#' @param gen Generation of the individual to edit
#' @param sex Gender of the individual to edit
#' @param nr Number of the individual to edit
#' @param nr.edits Number of edits to perform
#' @param decodeOriginsU Used function for the decoding of genetic origins [[5]]/[[6]]
#' @param bit.storing Set to TRUE if the RekomBre (not-miraculix! bit-storing is used)
#' @param nbits Bits available in RekomBre-bit-storing
#' @param population Population list
#' @export


edit_animal <- function(population, gen, sex, nr, nr.edits, decodeOriginsU=decodeOriginsR,
                        bit.storing=FALSE, nbits=30){

  animal <- population$breeding[[gen]][[sex]][[nr]]
  if(length(population$info$miraculix)>0 && population$info$miraculix){
    haplo <- miraculix::computeSNPS(population, gen, sex, nr, output_compressed=FALSE, what="haplo")
  } else{
    haplo <- compute.snps(population, gen, sex, nr, decodeOriginsU=decodeOriginsU, bit.storing=bit.storing, nbits=nbits, output_compressed=FALSE)
  }
  geno <- colSums(haplo)
#  multi <- rep(2, sum(population$info$snp))
#  multi[geno==2] <- (-2)
#  multi[geno==1] <- (population$info$u_hat[[length(population$info$u_hat)]][geno==1] > 0)*2-1
#  editing_effect <- geno * population$info$u_hat[[length(population$info$u_hat)]]
  ed_info <- population$info$editing_info[[length(population$info$editing_info)]]
  edits_p <- numeric(nr.edits)
  edits_row <- numeric(nr.edits)
  edits <- 0
  current_p <- 1
  max_p <- sum(population$info$snp)
  while(edits < nr.edits && current_p <= max_p){
    if(geno[ed_info[current_p,1]] == (2*ed_info[current_p,2]) || geno[ed_info[current_p,1]]==1 ){
      edits <- edits + 1
      edits_p[edits] <- ed_info[current_p,1]
      edits_row[edits] <- current_p
    }
    current_p <- current_p +1
  }
  for(set in 1:2){
    #address position instead of location genome
    #changes <- population$info$snp.position[edits_p][(haplo[set,edits_p]==(ed_info[edits_row,2]))]
    changes <- as.integer(edits_p[(haplo[set,edits_p]==(ed_info[edits_row,2]))])
    both <- intersect(animal[[2+set]], changes)
    animal[[2+set]] <- sort(c(animal[[2+set]], changes))
    if(length(both)>0){
      removes <- numeric(length(both))
      for(index in 1:length(both)){
        removes[index] <- which(animal[[2+set]]==both[index])[[1]]
      }
      animal[[2+set]] <- animal[[2+set]][-removes]
    }
  }
  if(length(animal)>=18){
    animal[[18]] <- c(animal[[18]],length(intersect(population$info$effect.p, edits_p)))
  } else{
    animal[[18]] <- length(intersect(population$info$effect.p, edits_p))
  }

  return(animal)
}
```
