## Supplementary material for "MoBPS - Modular Breeding Program Simulator": MoBPS R-package: ensembl.map.R

```
'#
  Authors
Torsten Pook,

Copyright (C) 2017 -- 2018  Torsten Pook

This program is free software; you can redistribute it and/or
modify it under the terms of the GNU General Public License
as published by the Free Software Foundation; either version 3
of the License, or (at your option) any later version.

This program is distributed in the hope that it will be useful,
but WITHOUT ANY WARRANTY; without even the implied warranty of
MERCHANTABILITY or FITNESS FOR A PARTICULAR PURPOSE.  See the
GNU General Public License for more details.

You should have received a copy of the GNU General Public License
along with this program; if not, write to the Free Software
Foundation, Inc., 59 Temple Place - Suite 330, Boston, MA  02111-1307, USA.
'#

#' Ensemble Map
#'
#' Function to generate a ensemble map file

#' @export
#' @param host Host to use in Ensembl (default: "www.ensembl.org" , alt: "plants.ensembl.org")
#' @param dataset Dataset used in Ensembl
#' @param filter Filters to apply in Ensembl
#' @param filter.values Values for the used filters in Ensembl
#' @param nchromo Number of chromosomes to export in the map
#' @param export.filters Export possible filters for parameter filters
#' @param export.datasets Export possible datasets for usage in parameter dataset
#' @export

ensembl.map <- function(host="www.ensembl.org", dataset="btaurus_snp", export.filters=FALSE, export.datasets=FALSE,
                        filter="variation_set_name", filter.values="Illumina BovineSNP50 BeadChip",
                        nchromo=NULL){
  if(export.datasets){
    if(host!="www.ensembl.org"){
      mart <- biomaRt::useMart(biomaRt::listMarts(host=host)[1,1],host=host)
    } else{
      mart = biomaRt::useEnsembl('ENSEMBL_MART_SNP')
    }

    export1 <- biomaRt::listDatasets(mart)
    return(export1)
  }
  ensembl = biomaRt::useEnsembl(biomart="snp", dataset=dataset, host=host)

  if(export.filters){
    export1 <- biomaRt::listFilters(ensembl)
    return(export1)
  }
  # head(listAttributes(ensembl))

  snps <- biomaRt::getBM(attributes=c('chr_name','refsnp_id', 'chrom_start', 'minor_allele_freq'),
                         filters=filter, values=filter.values, mart=ensembl)

    ## Map generation
  map <- NULL
  if(length(nchromo)==0){
    nchromo <- max(as.numeric((snps[,1])), na.rm =TRUE)
  }

  for(index in 1:nchromo){
    take <- which(snps[,1]==index)
    current_chromo <- snps[take,]
    reorder <- sort(as.numeric(current_chromo[,3]), index.return=TRUE)$ix
    current_chromo <- current_chromo[reorder,]

    map <- rbind(map, cbind(current_chromo[,1], current_chromo[,2], current_chromo[,3], NA, current_chromo[,4])) # 4 should be cM position, 5 should be allele freq
  }
  colnames(map) <- c("Chromosome", "SNP-ID", "bp", "M", "freq")
  return(map)

}
```
