## Supplementary material for "MoBPS - Modular Breeding Program Simulator": MoBPS R-package: epi.R

```
'#
  Authors
Torsten Pook,

Copyright (C) 2017 -- 2018  Torsten Pook

This program is free software; you can redistribute it and/or
modify it under the terms of the GNU General Public License
as published by the Free Software Foundation; either version 3
of the License, or (at your option) any later version.

This program is distributed in the hope that it will be useful,
but WITHOUT ANY WARRANTY; without even the implied warranty of
MERCHANTABILITY or FITNESS FOR A PARTICULAR PURPOSE.  See the
GNU General Public License for more details.

You should have received a copy of the GNU General Public License
along with this program; if not, write to the Free Software
Foundation, Inc., 59 Temple Place - Suite 330, Boston, MA  02111-1307, USA.
'#

#' Moore-Penrose-Transfomration
#'
#' Internal transformation using Moore-Penrose
#' @param alpha alpha
#' @param G kinship-matrix
#' @param Z genomic information matrix
#' @export

alpha_to_beta <- function(alpha,G,Z) {
  if (requireNamespace("MASS", quietly = TRUE)) {
    crossprod(Z,crossprod(MASS::ginv(G),alpha))
  } else{
    crossprod(Z,crossprod(solve(G),alpha))
  }
}

#' Martini-Test function
#'
#' Internal function to perform martini test
#' @param y y
#' @param Z genomic information matrix
#' @param G kinship matrix

epi <- function(y,Z, G=NULL) {
  n <- length(y)
  p <- ncol(Z)
  stopifnot(n == nrow(Z))
  if(length(G)==0){
    G <- tcrossprod(Z)
  }
  if(requireNamespace("EMMREML", quietly = TRUE)){
    fm <- EMMREML::emmreml(
      y,
      matrix(1,nrow=n),
      diag(n),
      G)
  } else{
    stop("Usage of EMMREML without being installed!")
  }

  beta <- alpha_to_beta(drop(fm$uhat),G,Z)
  return(drop(beta))
}
```
