## Supplementary material for "MoBPS - Modular Breeding Program Simulator": MoBPS R-package: example.dataset.R

```
'#
  Authors
Torsten Pook,

Copyright (C) 2017 -- 2018  Torsten Pook

This program is free software; you can redistribute it and/or
modify it under the terms of the GNU General Public License
as published by the Free Software Foundation; either version 3
of the License, or (at your option) any later version.

This program is distributed in the hope that it will be useful,
but WITHOUT ANY WARRANTY; without even the implied warranty of
MERCHANTABILITY or FITNESS FOR A PARTICULAR PURPOSE.  See the
GNU General Public License for more details.

You should have received a copy of the GNU General Public License
along with this program; if not, write to the Free Software
Foundation, Inc., 59 Temple Place - Suite 330, Boston, MA  02111-1307, USA.
'#

#' @name cattle_chip
#' @title Cattle chip
#' @description Genome for cattle according to Ma et al.
#' @usage cattle_chip
#' @source Ma et al
#' @author Torsten Pook \email{}
NULL

#' @name pig_chip
#' @title pig chip
#' @description Genome for pig according to Rohrer et al.
#' @usage pig_chip
#' @source Rohrer et al
#' @author Torsten Pook \email{}
NULL

#' @name chicken_chip
#' @title chicken chip
#' @description Genome for chicken according to Groenen et al.
#' @usage chicken_chip
#' @source Groenen et al
#' @author Torsten Pook \email{}
NULL

#' @name sheep_chip
#' @title sheep chip
#' @description Genome for sheep according to Prieur et al.
#' @usage sheep_chip
#' @source Prieur et al
#' @author Torsten Pook \email{}
NULL

#' @name maize_chip
#' @title maize chip
#' @description Genome for maize according to Lee et al.
#' @usage maize_chip
#' @source Lee et al
#' @author Torsten Pook \email{}
NULL

#' @name ex_json
#' @title ex_json
#' @description Exemplary json-data
#' @usage ex_json
#' @source Web-interface
#' @author Torsten Pook \email{}
NULL
```
