## Supplementary material for "MoBPS - Modular Breeding Program Simulator": MoBPS R-package: find.chromo.R

```
'#
  Authors
Torsten Pook,

Copyright (C) 2017 -- 2018  Torsten Pook

This program is free software; you can redistribute it and/or
modify it under the terms of the GNU General Public License
as published by the Free Software Foundation; either version 3
of the License, or (at your option) any later version.

This program is distributed in the hope that it will be useful,
but WITHOUT ANY WARRANTY; without even the implied warranty of
MERCHANTABILITY or FITNESS FOR A PARTICULAR PURPOSE.  See the
GNU General Public License for more details.

You should have received a copy of the GNU General Public License
along with this program; if not, write to the Free Software
Foundation, Inc., 59 Temple Place - Suite 330, Boston, MA  02111-1307, USA.
'#

#' Position detection
#'
#' Internal function for the detection on which chromosome each marker is
#' @param position position in the genome
#' @param length.total Length of each chromosome


find.chromo <- function(position, length.total){
  chromo <- min(sum(position<=length.total), length(length.total)-1)
  return(chromo)
}
```
