## Supplementary material for "MoBPS - Modular Breeding Program Simulator": MoBPS R-package: find.snpbefore.R

```
'#
  Authors
Torsten Pook,

Copyright (C) 2017 -- 2018  Torsten Pook

This program is free software; you can redistribute it and/or
modify it under the terms of the GNU General Public License
as published by the Free Software Foundation; either version 3
of the License, or (at your option) any later version.

This program is distributed in the hope that it will be useful,
but WITHOUT ANY WARRANTY; without even the implied warranty of
MERCHANTABILITY or FITNESS FOR A PARTICULAR PURPOSE.  See the
GNU General Public License for more details.

You should have received a copy of the GNU General Public License
along with this program; if not, write to the Free Software
Foundation, Inc., 59 Temple Place - Suite 330, Boston, MA  02111-1307, USA.
'#

#' Position detection
#'
#' Internal function for the detection on which position each marker is
#' @param position Position on the genome
#' @param snp.position Position of the SNPs on the genome
#' @export


find.snpbefore <- function(position, snp.position){
  n <- length(snp.position)
  before <- ceiling(n/2)
  step_size <- ceiling(n/4)
  check0 <- checkn <- 0
  while(before!=0 && before!=n && (snp.position[before]>position || snp.position[before+1]<=position)){
    if(snp.position[before]>position){
      before <- before - step_size
    } else{
      before <- before + step_size
    }
    step_size <- ceiling(step_size/2)
    if(step_size > min(before, n-before)){
      step_size <- 1
    }
    if(before==0 && check0==0){
      before <- 1
      check0 <- 1
    }
    if(before==n && checkn==0){
      before <- n -1
      checkn <- 1
    }
  }
  return(before)
}
```
