## Supplementary material for "MoBPS - Modular Breeding Program Simulator": MoBPS R-package: generation.individual.R

```
'#
  Authors
Torsten Pook,

Copyright (C) 2017 -- 2018  Torsten Pook

This program is free software; you can redistribute it and/or
modify it under the terms of the GNU General Public License
as published by the Free Software Foundation; either version 3
of the License, or (at your option) any later version.

This program is distributed in the hope that it will be useful,
but WITHOUT ANY WARRANTY; without even the implied warranty of
MERCHANTABILITY or FITNESS FOR A PARTICULAR PURPOSE.  See the
GNU General Public License for more details.

You should have received a copy of the GNU General Public License
along with this program; if not, write to the Free Software
Foundation, Inc., 59 Temple Place - Suite 330, Boston, MA  02111-1307, USA.
'#

#' Function to generate a new individual
#'
#' Function to generate a new individual
#' @param indexb internal test
#' @param population internal test
#' @param info_father_list internal test
#' @param info_mother_list internal test
#' @param copy.individual internal test
#' @param mutation.rate internal test
#' @param remutation.rate internal test
#' @param recombination.rate internal test
#' @param recom.f.indicator internal test
#' @param recom.f.polynom internal test
#' @param duplication.rate internal test
#' @param duplication.length internal test
#' @param duplication.recombination internal test
#' @param delete.same.origin internal test
#' @param gene.editing internal test
#' @param nr.edits internal test
#' @param gen.architecture.m internal test
#' @param gen.architecture.f internal test
#' @param decodeOriginsU internal test
#' @param current.gen internal test
#' @param save.recombination.history internal test
#' @param new.bv.child internal test
#' @param dh.mating internal test
#' @param share.genotyped internal test
#' @param added.genotyped internal test
#' @param dh.sex internal test
#' @param n.observation internal test


generation.individual <- function(indexb, population, info_father_list, info_mother_list,
                                  copy.individual, mutation.rate, remutation.rate, recombination.rate,
                                  recom.f.indicator, recom.f.polynom, duplication.rate, duplication.length,
                                  duplication.recombination, delete.same.origin,
                                  gene.editing, nr.edits, gen.architecture.m, gen.architecture.f,
                                  decodeOriginsU, current.gen, save.recombination.history, new.bv.child,
                                  dh.mating, share.genotyped, added.genotyped,
                                  dh.sex, n.observation){

  info.father <- info_father_list[indexb,]
  info.mother <- info_mother_list[indexb,]

  father <- population$breeding[[info.father[1]]][[info.father[2]]][[info.father[3]]]
  mother <- population$breeding[[info.mother[1]]][[info.mother[2]]][[info.mother[3]]]


  if(copy.individual){
    info.mother <- info.father
    child1 <- list(father[[1]], father[[3]], father[[5]], father[[7]], father[[11]], 0, if(length(father)>19){father[[19]]} else{0})
    child2 <- list(father[[2]], father[[4]], father[[6]], father[[8]], father[[12]], 0, if(length(father)>19){father[[20]]} else{0})
  } else{
    child1 <- breeding.intern(info.father, father, population,
                              mutation.rate, remutation.rate, recombination.rate,
                              recom.f.indicator, recom.f.polynom, duplication.rate, duplication.length,
                              duplication.recombination, delete.same.origin=delete.same.origin,
                              gene.editing=gene.editing, nr.edits= nr.edits,
                              gen.architecture=gen.architecture.m,
                              decodeOriginsU=decodeOriginsU)

    child2 <- breeding.intern(info.mother, mother, population,
                              mutation.rate, remutation.rate, recombination.rate,
                              recom.f.indicator, recom.f.polynom, duplication.rate, duplication.length,
                              duplication.recombination, delete.same.origin=delete.same.origin,
                              gene.editing=gene.editing , nr.edits= nr.edits,
                              gen.architecture=gen.architecture.f,
                              decodeOriginsU=decodeOriginsU)
  }
  if(dh.mating){
    if(stats::rbinom(1,1,dh.sex)==0){
      child2 <- child1
    } else{
      child1 <- child2
    }
  }
  child <- list()
  child[[1]] <- child1[[1]]
  child[[2]] <- child2[[1]]
  child[[3]] <- child1[[2]]
  child[[4]] <- child2[[2]]
  child[[5]] <- child1[[3]]
  child[[6]] <- child2[[3]]
  child[[7]] <- child1[[4]]
  child[[8]] <- child2[[4]]

  if(is.vector(child1[[5]])){
    child[[11]] <- t(as.matrix(child1[[5]]))
  } else{
    child[[11]] <- child1[[5]]
  }
  if(is.vector(child2[[5]])){
    child[[12]] <- t(as.matrix(child2[[5]]))
  } else{
    child[[12]] <- child2[[5]]
  }

  if(save.recombination.history && current.gen==1){
    if(length(child1[[6]][-c(1,length(child1[[6]]))])>0){
      child[[13]] <- cbind(current.gen, child1[[6]][-c(1,length(child1[[6]]))])
    } else{
      child[[13]] <- cbind(0,0)
    }
    if(length( child2[[6]][-c(1,length(child2[[6]]))])>0){
      child[[14]] <- cbind(current.gen, child2[[6]][-c(1,length(child2[[6]]))])
    } else{
      child[[14]] <- cbind(0,0)
    }

  } else if(save.recombination.history && current.gen>1){
    if(length(child1[[6]][-c(1,length(child1[[6]]))])>0){
      child[[13]] <- rbind(population$breeding[[info.father[1]]][[info.father[2]]][[info.father[3]]][[13]], population$breeding[[info.father[1]]][[info.father[2]]][[info.father[3]]][[14]], cbind(current.gen, child1[[6]][-c(1,length(child1[[6]]))]))
    } else{
      child[[13]] <- rbind(population$breeding[[info.father[1]]][[info.father[2]]][[info.father[3]]][[13]], population$breeding[[info.father[1]]][[info.father[2]]][[info.father[3]]][[14]])

    }
    if(length( child2[[6]][-c(1,length(child2[[6]]))])>0){
      child[[14]] <- rbind(population$breeding[[info.mother[1]]][[info.mother[2]]][[info.mother[3]]][[13]], population$breeding[[info.mother[1]]][[info.mother[2]]][[info.mother[3]]][[14]], cbind(current.gen, child2[[6]][-c(1,length(child2[[6]]))]))
    } else{
      child[[14]] <- rbind(population$breeding[[info.mother[1]]][[info.mother[2]]][[info.mother[3]]][[13]], population$breeding[[info.mother[1]]][[info.mother[2]]][[info.mother[3]]][[14]])

    }

  }
  if(new.bv.child=="obs"){
    child[[15]] <- n.observation
  } else if(new.bv.child=="addobs"){
    child[[15]] <- n.observation +
      population$breeding[[info.mother[1]]][[info.mother[2]]][[info.mother[3]]][[15]]/2   +
      population$breeding[[info.father[1]]][[info.father[2]]][[info.father[3]]][[15]]/2
  } else{
    child[[15]] <- rep(0, population$info$bv.nr)
  }
  if(copy.individual){
    child[[16]] <- population$breeding[[info.father[1]]][[info.father[2]]][[info.father[3]]][[16]]
    if(added.genotyped>0 && child[[16]]==0){
      child[[16]] <- stats::rbinom(1,1,added.genotyped)
    }
  } else{
    child[[16]] <- stats::rbinom(1,1,share.genotyped)
  }

  child[[19]] <- child1[[7]]
  child[[20]] <- child2[[7]]

  if(copy.individual){
    child[[21]] <-  matrix(info.father[1:3], nrow=1)
  }

  return(child)

}
```
