## Supplementary material for "MoBPS - Modular Breeding Program Simulator": MoBPS R-package: get.bv.R

```
'#
  Authors
Torsten Pook,

Copyright (C) 2017 -- 2018  Torsten Pook

This program is free software; you can redistribute it and/or
modify it under the terms of the GNU General Public License
as published by the Free Software Foundation; either version 3
of the License, or (at your option) any later version.

This program is distributed in the hope that it will be useful,
but WITHOUT ANY WARRANTY; without even the implied warranty of
MERCHANTABILITY or FITNESS FOR A PARTICULAR PURPOSE.  See the
GNU General Public License for more details.

You should have received a copy of the GNU General Public License
along with this program; if not, write to the Free Software
Foundation, Inc., 59 Temple Place - Suite 330, Boston, MA  02111-1307, USA.
'#

#' Export underlying true breeding values
#'
#' Function to export underlying true breeding values
#' @param population Population list
#' @param database Groups of individuals to consider for the export
#' @param gen Quick-insert for database (vector of all generations to export)
#' @param cohorts Quick-insert for database (vector of names of cohorts to export)
#' @export

get.bv<- function(population, database=NULL, gen=NULL, cohorts=NULL){

  database <- get.database(population, gen, database, cohorts)

  n.animals <- sum(database[,4] - database[,3] +1)
  data <- matrix(0, ncol=n.animals, nrow=population$info$bv.nr)
  before <- 0
  names <- numeric(n.animals)
  for(row in 1:nrow(database)){
    animals <- database[row,]
    nanimals <- database[row,4] - database[row,3] +1
    if(nanimals>0){
      data[,(before+1):(before+nanimals)] <- population$breeding[[animals[1]]][[6+ animals[2]]][, animals[3]:animals[4]]
      names[(before+1):(before+nanimals)] <- paste(if(animals[2]==1) "M" else "F", animals[3]:animals[4],"_", animals[1], sep="")
      before <- before + nanimals
    }
  }
  row_names <- paste("Trait", 1:population$info$bv.nr)
  colnames(data) <- names
  rownames(data) <- row_names
  return(data)
}
```
