## Supplementary material for "MoBPS - Modular Breeding Program Simulator": MoBPS R-package: get.database.R

```
'#
  Authors
Torsten Pook,

Copyright (C) 2017 -- 2018  Torsten Pook

This program is free software; you can redistribute it and/or
modify it under the terms of the GNU General Public License
as published by the Free Software Foundation; either version 3
of the License, or (at your option) any later version.

This program is distributed in the hope that it will be useful,
but WITHOUT ANY WARRANTY; without even the implied warranty of
MERCHANTABILITY or FITNESS FOR A PARTICULAR PURPOSE.  See the
GNU General Public License for more details.

You should have received a copy of the GNU General Public License
along with this program; if not, write to the Free Software
Foundation, Inc., 59 Temple Place - Suite 330, Boston, MA  02111-1307, USA.
'#

#' gen/database/cohorts conversion
#'
#' Function to derive a database based on gen/database/cohorts
#' @param population Population list
#' @param database Groups of individuals to consider for the export
#' @param gen Quick-insert for database (vector of all generations to export)
#' @param cohorts Quick-insert for database (vector of names of cohorts to export)
#' @export


get.database<- function(population, gen=NULL, database=NULL, cohorts=NULL){

  if(length(gen)>0){
    database_gen <- cbind(rep(gen,each=2), rep(1:2, length(gen)))
    if(length(database)>0 && ncol(database)==4){
      database_gen <- cbind(database_gen,1,population$info$size[database_gen])
    }
    database <- rbind(database, database_gen)
  }
  if(length(database)>0 && ncol(database)==2){
    start <- end <- numeric(nrow(database))
    for(index in 1:nrow(database)){
      start[index] <- 1
      end[index] <- population$info$size[database[index,1], database[index,2]]
    }
    database <- cbind(database, start, end)
  }
  if(length(cohorts)>0){
    database2 <- matrix(0L, nrow=length(cohorts), ncol=4)
    for(index in 1:length(cohorts)){
      row <- which(population$info$cohorts==cohorts[index])[1]
      gen <- as.numeric(population$info$cohorts[row,2])
      sex <- 1 + (as.numeric(population$info$cohorts[row,4])>0)
      first <- as.numeric(population$info$cohorts[row,5 + sex])
      last <- first + as.numeric(population$info$cohorts[row,2 + sex]) - 1
      database2[index,] <- c(gen,sex,first,last)
    }
    database <- rbind(database, database2)
  }

  keep <- database[,3]<=database[,4]
  database <- database[keep,,drop=FALSE]

  if(length(database)>0 && nrow(database)>1){
    order <- sort(database[,1]*1e10 + database[,2]*1e5 + database[,3], index.return=TRUE)$ix
    database <- database[order,,drop=FALSE]
    for(index in 2:nrow(database)){
      checks <- (which(database[1:(index-1),1]==database[index,1] & database[1:(index-1),2] == database[index,2]))
      for(index2 in checks){
        if(database[index,3] < (database[index2,4]+1)){
          database[index2,4] <- max(database[index2,4], database[index,4])
          database[index,] <- 0
        }

      }
    }
    database <- database[database[,1]!=0,,drop=FALSE]
  }

  database <- unique(database)

  return(database)
}
```
