## Supplementary material for "MoBPS - Modular Breeding Program Simulator": MoBPS R-package: get.geno.R

```
'#
  Authors
Torsten Pook,

Copyright (C) 2017 -- 2018  Torsten Pook

This program is free software; you can redistribute it and/or
modify it under the terms of the GNU General Public License
as published by the Free Software Foundation; either version 3
of the License, or (at your option) any later version.

This program is distributed in the hope that it will be useful,
but WITHOUT ANY WARRANTY; without even the implied warranty of
MERCHANTABILITY or FITNESS FOR A PARTICULAR PURPOSE.  See the
GNU General Public License for more details.

You should have received a copy of the GNU General Public License
along with this program; if not, write to the Free Software
Foundation, Inc., 59 Temple Place - Suite 330, Boston, MA  02111-1307, USA.
'#

#' Derive genotypes of selected individuals
#'
#' Function to devide genotypes of selected individuals
#' @param population Population list
#' @param database Groups of individuals to consider for the export
#' @param gen Quick-insert for database (vector of all generations to export)
#' @param cohorts Quick-insert for database (vector of names of cohorts to export)
#' @param chromosomen Beschraenkung des Genotypen auf bestimmte Chromosomen (default: 1)
#' @param export.alleles If TRUE export underlying alleles instead of just 012
#' @export

get.geno <- function(population, database=NULL, gen=NULL, cohorts=NULL, chromosomen="all", export.alleles=FALSE){

  if(length(chromosomen)==1 && chromosomen=="all"){
    subsetting <- FALSE
    chromosomen <- 1:length(population$info$snp)
  } else{
    subsetting <- TRUE
  }

  if(population$info$miraculix){
    if (requireNamespace("miraculix", quietly = TRUE)) {
      codeOriginsU <- miraculix::codeOrigins
      decodeOriginsU <- miraculix::decodeOrigins
    } else{
      codeOriginsU <- codeOriginsR
      decodeOriginsU <- decodeOriginsR
    }
  } else{
    codeOriginsU <- codeOriginsR
    decodeOriginsU <- decodeOriginsR
  }
  if(length(population$info$origin.gen)>0){
    population$info$origin.gen <- as.integer(population$info$origin.gen)
  } else{
    population$info$origin.gen <- 1:64L
  }


  database <- get.database(population, gen, database, cohorts)

  start.chromo <- cumsum(c(1,population$info$snp)[-(length(population$info$snp)+1)])
  end.chromo <- population$info$cumsnp

  relevant.snps <- NULL
  for(index in chromosomen){
    relevant.snps <- c(relevant.snps, start.chromo[index]:end.chromo[index])
  }
  nsnp <- length(relevant.snps)

  titel <- t(population$info$snp.base[,relevant.snps])

  n.animals <- sum(database[,4] - database[,3] +1)
  data <- matrix(0, ncol=n.animals, nrow=nsnp)
  before <- 0
  names <- numeric(n.animals)
  colnames(titel) <- c("Major_Allel", "Minor_Allel")

  for(row in 1:nrow(database)){
    animals <- database[row,]
    nanimals <- database[row,4] - database[row,3] +1

    if(nanimals>0){
      if(population$info$miraculix){
        if(subsetting){
          data[, (before+1):(before+nanimals)] <- miraculix::computeSNPS(population,rep(animals[1], nanimals), rep(animals[2], nanimals) ,animals[3]:animals[4], what="geno")[relevant.snps,]

        } else{
          data[, (before+1):(before+nanimals)] <- miraculix::computeSNPS(population,rep(animals[1], nanimals), rep(animals[2], nanimals) ,animals[3]:animals[4], what="geno")
        }
      }
      names[(before+1):(before+nanimals)] <- paste(if(animals[2]==1) "M" else "F", animals[3]:animals[4], "_", animals[1], sep="")
      if(!population$info$miraculix){
        rindex <- 1
        for(index in animals[3]:animals[4]){
          data[, before + rindex] <- colSums(compute.snps(population,animals[1], animals[2],index, decodeOriginsU=decodeOriginsU))[relevant.snps]
          rindex <- rindex + 1
        }
      }
    }
    before <- before + nanimals
  }
  colnames(data) <- names
  rownames(data) <- population$info$snp.name[relevant.snps]
  if(export.alleles){
    return(list(titel,data))
  } else{
    return(data)
  }
}
```
