## Supplementary material for "MoBPS - Modular Breeding Program Simulator": MoBPS R-package: get.genotyped.R

```
'#
  Authors
Torsten Pook,

Copyright (C) 2017 -- 2018  Torsten Pook

This program is free software; you can redistribute it and/or
modify it under the terms of the GNU General Public License
as published by the Free Software Foundation; either version 3
of the License, or (at your option) any later version.

This program is distributed in the hope that it will be useful,
but WITHOUT ANY WARRANTY; without even the implied warranty of
MERCHANTABILITY or FITNESS FOR A PARTICULAR PURPOSE.  See the
GNU General Public License for more details.

You should have received a copy of the GNU General Public License
along with this program; if not, write to the Free Software
Foundation, Inc., 59 Temple Place - Suite 330, Boston, MA  02111-1307, USA.
'#

#' Derive genetic origins
#'
#' Function to derive genetic origin
#' @param population Population list
#' @param database Groups of individuals to consider for the export
#' @param gen Quick-insert for database (vector of all generations to export)
#' @param cohorts Quick-insert for database (vector of names of cohorts to export)
#' @export

get.genotyped <- function(population, database=NULL, gen=NULL, cohorts=NULL){

  database <- get.database(population, gen, database, cohorts)
  n.animals <- sum(database[,4] - database[,3] +1)
  genotyped <- colnamed <- numeric(n.animals)
  rindex <- 1

  for(row in 1:nrow(database)){
    animals <- database[row,]
    if(diff(database[row,3:4])>0){
      for(index in database[row,3]:database[row,4]){
        genotyped[rindex] <- population$breeding[[database[row,1]]][[database[row,2]]][[index]][[16]]
        colnamed[rindex] <- paste(if(animals[2]==1) "M" else "F", index,"_", animals[1],sep="")
        rindex <- rindex + 1
      }
    }

  }
  names(genotyped) <- colnamed
  genotyped <- genotyped>0
  return(genotyped)
}
```
