## Supplementary material for "MoBPS - Modular Breeding Program Simulator": MoBPS R-package: get.infos.R

```
'#
  Authors
Torsten Pook,

Copyright (C) 2017 -- 2018  Torsten Pook

This program is free software; you can redistribute it and/or
modify it under the terms of the GNU General Public License
as published by the Free Software Foundation; either version 3
of the License, or (at your option) any later version.

This program is distributed in the hope that it will be useful,
but WITHOUT ANY WARRANTY; without even the implied warranty of
MERCHANTABILITY or FITNESS FOR A PARTICULAR PURPOSE.  See the
GNU General Public License for more details.

You should have received a copy of the GNU General Public License
along with this program; if not, write to the Free Software
Foundation, Inc., 59 Temple Place - Suite 330, Boston, MA  02111-1307, USA.
'#

#' Extract bv/pheno/geno of selected individuals
#'
#' Function to extract bv/pheno/geno of selected individuals
#' @param population Population list
#' @param database Groups of individuals to consider for the export
#' @param gen Quick-insert for database (vector of all generations to export)
#' @param cohorts Quick-insert for database (vector of names of cohorts to export)


get.infos<- function(population, database=NULL, gen=NULL, cohorts=NULL){

  if(length(population$info$origin.gen)>0){
    population$info$origin.gen <- as.integer(population$info$origin.gen)
  } else{
    population$info$origin.gen <- 1:64L
  }
  database <- get.database(population, gen, database, cohorts)

  pheno <- as.numeric(get.pheno(population, database=database)[-1])
  bv <- as.numeric(get.bv(population, database=database)[-1])

  size <- database[,4]-database[,3]+1
  cindex <- 1
  loop_elements <- matrix(0,nrow=sum(size), ncol=3)
  for(index in 1:nrow(database)){
    loop_elements[cindex:(cindex+size[index]-1),] <- cbind(database[index,1], database[index,2], 1:size[index])
    cindex <- cindex + size[index]
  }

  if(population$info$miraculix){
    if (requireNamespace("miraculix", quietly = TRUE)) {
      Z <- miraculix::computeSNPS(population, loop_elements[,1], loop_elements[,2], loop_elements[,3],
                                  what="geno", output_compressed=FALSE)
    } else{
      stop("Usage of miraculix without being installed!")
    }

  } else{
    Z <- matrix(NA, nrow=sum(population$info$snp), ncol=nrow(loop_elements))
    for(index in 1:nrow(loop_elements)){
      Z[,index] <- colSums(compute.snps(population, loop_elements[index,1] , loop_elements[index,2] , loop_elements[index,3], decodeOriginsU=miraculix::decodeOrigins))

    }
  }


  add_eff <- population$info$real.bv.add
  mult_eff <- population$info$real.bv.mult
  dice_eff <- population$info$real.bv.dice

  return(list(pheno, bv, Z, add_eff, mult_eff, dice_eff))
}
```
