## Supplementary material for "MoBPS - Modular Breeding Program Simulator": MoBPS R-package: get.pedigree.R

```
'#
  Authors
Torsten Pook,

Copyright (C) 2017 -- 2018  Torsten Pook

This program is free software; you can redistribute it and/or
modify it under the terms of the GNU General Public License
as published by the Free Software Foundation; either version 3
of the License, or (at your option) any later version.

This program is distributed in the hope that it will be useful,
but WITHOUT ANY WARRANTY; without even the implied warranty of
MERCHANTABILITY or FITNESS FOR A PARTICULAR PURPOSE.  See the
GNU General Public License for more details.

You should have received a copy of the GNU General Public License
along with this program; if not, write to the Free Software
Foundation, Inc., 59 Temple Place - Suite 330, Boston, MA  02111-1307, USA.
'#

#' Derive pedigree
#'
#' Derive pedigree for selected individuals
#' @param population Population list
#' @param database Groups of individuals to consider for the export
#' @param gen Quick-insert for database (vector of all generations to export)
#' @param cohorts Quick-insert for database (vector of names of cohorts to export)
#' @param founder.zero Parents of founders are displayed as "0" (default: TRUE)
#' @param raw Set to TRUE to not convert numbers into Sex etc.
#' @export


get.pedigree <- function(population, database=NULL, gen=NULL, cohorts=NULL, founder.zero=TRUE,
                         raw=FALSE){

  database <- get.database(population, gen, database, cohorts)

  n.animals <- sum(database[,4] - database[,3] +1)

  pedigree <- matrix(0, nrow=n.animals, ncol=3 + 6 * raw)
  rindex <- 1

  if(raw){
    for(row in 1:nrow(database)){
      animals <- database[row,]
      for(index in database[row,3]:database[row,4]){
        father <- population$breeding[[database[row,1]]][[database[row,2]]][[index]][[7]][1:3]
        mother <- population$breeding[[database[row,1]]][[database[row,2]]][[index]][[8]][1:3]
        pedigree[rindex,] <- c(database[row,1:2], index, father, mother)
        rindex <- rindex + 1
      }
    }
  } else{
    for(row in 1:nrow(database)){
      animals <- database[row,]
      for(index in database[row,3]:database[row,4]){
        father <- population$breeding[[database[row,1]]][[database[row,2]]][[index]][[7]]
        mother <- population$breeding[[database[row,1]]][[database[row,2]]][[index]][[8]]
        father_t <- paste(if(father[2]==1) "M" else "F", father[3], "_", father[1], sep="")
        mother_t <- paste(if(mother[2]==1) "M" else "F", mother[3], "_", mother[1], sep="")
        child_t <- paste(if(database[row,2]==1) "M" else "F", index, "_", database[row,1], sep="")
        pedigree[rindex,] <- c(child_t, father_t, mother_t)
        rindex <- rindex + 1
      }
    }
  }

  if(!raw){
    colnames(pedigree) <- c("offspring", "father", "mother")
  }


  if(founder.zero && !raw){
    set0 <- which(pedigree[,1]==pedigree[,2])
    if(length(set0)>0){
      pedigree[set0,2] <- "0"
    }
    set0 <- which(pedigree[,1]==pedigree[,3])
    if(length(set0)>0){
      pedigree[set0,3] <- "0"
    }
  }
  return(pedigree)
}

#' Derive pedigree including grandparents
#'
#' Derive pedigree for selected individuals including grandparents
#' @param population Population list
#' @param database Groups of individuals to consider for the export
#' @param gen Quick-insert for database (vector of all generations to export)
#' @param cohorts Quick-insert for database (vector of names of cohorts to export)
#' @param shares Determine actual inherited shares of grandparents
#' @param founder.zero Parents of founders are displayed as "0" (default: TRUE)
#' @export

get.pedigree2 <- function(population, database=NULL, gen=NULL, cohorts=NULL, shares=FALSE, founder.zero=TRUE){

  database <- get.database(population, gen, database, cohorts)

  n.animals <- sum(database[,4] - database[,3] +1)


  if(shares){
    cols <- 9
  } else{
    cols <- 5
  }
  pedigree <- matrix(0, nrow=n.animals, ncol=cols)
  rindex <- 1

  for(row in 1:nrow(database)){
    animals <- database[row,]
    for(index in database[row,3]:database[row,4]){
      father <- population$breeding[[database[row,1]]][[database[row,2]]][[index]][[7]]
      mother <- population$breeding[[database[row,1]]][[database[row,2]]][[index]][[8]]
      grandf1 <- population$breeding[[father[[1]]]][[father[[2]]]][[father[[3]]]][[7]]
      grandf1share <- population$breeding[[database[row,1]]][[database[row,2]]][[index]][[17]]
      grandm1 <- population$breeding[[father[[1]]]][[father[[2]]]][[father[[3]]]][[8]]
      grandm1share <- 1 - population$breeding[[database[row,1]]][[database[row,2]]][[index]][[17]]
      grandf2 <- population$breeding[[mother[[1]]]][[mother[[2]]]][[mother[[3]]]][[7]]
      grandf2share <- population$breeding[[database[row,1]]][[database[row,2]]][[index]][[18]]
      grandm2 <- population$breeding[[mother[[1]]]][[mother[[2]]]][[mother[[3]]]][[8]]
      grandm2share <- 1 - population$breeding[[database[row,1]]][[database[row,2]]][[index]][[18]]

      grandf1_t <- paste(if(grandf1[2]==1) "M" else "F", grandf1[3], "_", grandf1[1], sep="")
      grandm1_t <- paste(if(grandm1[2]==1) "M" else "F", grandm1[3], "_", grandm1[1], sep="")
      grandf2_t <- paste(if(grandf2[2]==1) "M" else "F", grandf2[3], "_", grandf2[1], sep="")
      grandm2_t <- paste(if(grandm2[2]==1) "M" else "F", grandm2[3], "_", grandm2[1], sep="")
      child_t <- paste(if(database[row,2]==1) "M" else "F", index, "_", gen, sep="")

      father_t <- paste(if(father[2]==1) "M" else "F", father[3], "_", father[1], sep="")
      mother_t <- paste(if(mother[2]==1) "M" else "F", mother[3], "_", mother[1], sep="")
      child_t <- paste(if(database[row,2]==1) "M" else "F", index, "_", database[row,1], sep="")
      if(shares){
        pedigree[rindex,] <- c(child_t, grandf1_t, grandm1_t, grandf2_t, grandm2_t, grandf1share, grandm1share, grandf2share, grandm2share)
      } else{
        pedigree[rindex,] <- c(child_t, grandf1_t, grandm1_t, grandf2_t, grandm2_t)
      }
      rindex <- rindex + 1
    }
  }

  if(founder.zero){
    set0 <- which(pedigree[,1]==pedigree[,2])
    if(length(set0)>0){
      pedigree[set0,2] <- "0"
    }
    set0 <- which(pedigree[,1]==pedigree[,3])
    if(length(set0)>0){
      pedigree[set0,3] <- "0"
    }
  }
  if(founder.zero){
    set0 <- which(pedigree[,1]==pedigree[,4])
    if(length(set0)>0){
      pedigree[set0,4] <- "0"
    }
    set0 <- which(pedigree[,1]==pedigree[,5])
    if(length(set0)>0){
      pedigree[set0,5] <- "0"
    }
  }
  return(pedigree)
}

#' Derive pedigree parents and grandparents
#'
#' Derive pedigree for selected individuals including parents/grandparents
#' @param population Population list
#' @param database Groups of individuals to consider for the export
#' @param gen Quick-insert for database (vector of all generations to export)
#' @param cohorts Quick-insert for database (vector of names of cohorts to export)
#' @param founder.zero Parents of founders are displayed as "0" (default: TRUE)
#' @export
#'
get.pedigree3 <- function(population, database=NULL, gen=NULL, cohorts=NULL, founder.zero=TRUE){

  database <- get.database(population, gen, database, cohorts)

  n.animals <- sum(database[,4] - database[,3] +1)
  pedigree <- matrix(0, nrow=n.animals, ncol=7)
  rindex <- 1

  for(row in 1:nrow(database)){
    animals <- database[row,]
    for(index in database[row,3]:database[row,4]){
      father <- population$breeding[[database[row,1]]][[database[row,2]]][[index]][[7]]
      mother <- population$breeding[[database[row,1]]][[database[row,2]]][[index]][[8]]
      grandf1 <- population$breeding[[father[[1]]]][[father[[2]]]][[father[[3]]]][[7]]
      grandf1share <- population$breeding[[database[row,1]]][[database[row,2]]][[index]][[17]]
      grandm1 <- population$breeding[[father[[1]]]][[father[[2]]]][[father[[3]]]][[8]]
      grandm1share <- 1 - population$breeding[[database[row,1]]][[database[row,2]]][[index]][[17]]
      grandf2 <- population$breeding[[mother[[1]]]][[mother[[2]]]][[mother[[3]]]][[7]]
      grandf2share <- population$breeding[[database[row,1]]][[database[row,2]]][[index]][[18]]
      grandm2 <- population$breeding[[mother[[1]]]][[mother[[2]]]][[mother[[3]]]][[8]]
      grandm2share <- 1 - population$breeding[[database[row,1]]][[database[row,2]]][[index]][[18]]

      f1_t <- paste(if(father[2]==1) "M" else "F", father[3], "_", father[1], sep="")
      m1_t <- paste(if(mother[2]==1) "M" else "F", mother[3], "_", mother[1], sep="")
      grandf1_t <- paste(if(grandf1[2]==1) "M" else "F", grandf1[3], "_", grandf1[1], sep="")
      grandm1_t <- paste(if(grandm1[2]==1) "M" else "F", grandm1[3], "_", grandm1[1], sep="")
      grandf2_t <- paste(if(grandf2[2]==1) "M" else "F", grandf2[3], "_", grandf2[1], sep="")
      grandm2_t <- paste(if(grandm2[2]==1) "M" else "F", grandm2[3], "_", grandm2[1], sep="")
      child_t <- paste(if(database[row,2]==1) "M" else "F", index, "_", gen, sep="")

      father_t <- paste(if(father[2]==1) "M" else "F", father[3], "_", father[1], sep="")
      mother_t <- paste(if(mother[2]==1) "M" else "F", mother[3], "_", mother[1], sep="")
      child_t <- paste(if(database[row,2]==1) "M" else "F", index, "_", database[row,1], sep="")

      pedigree[rindex,] <- c(child_t, f1_t, m1_t, grandf1_t, grandm1_t, grandf2_t, grandm2_t)
      rindex <- rindex + 1
    }
  }
  if(founder.zero){
    set0 <- which(pedigree[,1]==pedigree[,2])
    if(length(set0)>0){
      pedigree[set0,2] <- "0"
    }
    set0 <- which(pedigree[,1]==pedigree[,3])
    if(length(set0)>0){
      pedigree[set0,3] <- "0"
    }
  }
  if(founder.zero){
    set0 <- which(pedigree[,2]==pedigree[,4])
    if(length(set0)>0){
      pedigree[set0,4] <- "0"
    }
    set0 <- which(pedigree[,2]==pedigree[,5])
    if(length(set0)>0){
      pedigree[set0,5] <- "0"
    }
  }
  if(founder.zero){
    set0 <- which(pedigree[,3]==pedigree[,6])
    if(length(set0)>0){
      pedigree[set0,6] <- "0"
    }
    set0 <- which(pedigree[,3]==pedigree[,7])
    if(length(set0)>0){
      pedigree[set0,7] <- "0"
    }
  }
  colnames(pedigree) <- c("offspring", "father", "mother", "grandfatherf", "grandmotherf", "grandfatherm", "grandmotherm")
  return(pedigree)
}
```
