## Supplementary material for "MoBPS - Modular Breeding Program Simulator": MoBPS R-package: get.pedmap.R

```
'#
  Authors
Torsten Pook,

Copyright (C) 2017 -- 2018  Torsten Pook

This program is free software; you can redistribute it and/or
modify it under the terms of the GNU General Public License
as published by the Free Software Foundation; either version 3
of the License, or (at your option) any later version.

This program is distributed in the hope that it will be useful,
but WITHOUT ANY WARRANTY; without even the implied warranty of
MERCHANTABILITY or FITNESS FOR A PARTICULAR PURPOSE.  See the
GNU General Public License for more details.

You should have received a copy of the GNU General Public License
along with this program; if not, write to the Free Software
Foundation, Inc., 59 Temple Place - Suite 330, Boston, MA  02111-1307, USA.

'#

#' Generate plink-file (pedmap)
#'
#' Generate a ped and map file (PLINK format) for selected groups and chromosome
#' @param population Population list
#' @param path Location to save pedmap-file
#' @param database Groups of individuals to consider for the export
#' @param gen Quick-insert for database (vector of all generations to export)
#' @param cohorts Quick-insert for database (vector of names of cohorts to export)
#' @param chromosomen Beschraenkung des Genotypen auf bestimmte Chromosomen (default: 1)
#' @export

get.pedmap <- function(population, path=NULL, database=NULL, gen=NULL, cohorts=NULL, chromosomen="all"){

  haplo <- get.haplo(population, database=database, gen=gen, cohorts=cohorts, chromosomen=chromosomen, export.alleles=FALSE)
  # haplo <- get.haplo(population, gen=1)
  if(length(path)==0){
    path <- "population"
  }
  chr.nr <- sum(population$info$snp)
  start <- 1
  for(index in 1:length(population$info$snp)){
    if(population$info$snp[index]>0){
      chr.nr[start:(start+population$info$snp[index]-1)] <- index
      start <- start + population$info$snp[index]
    }
  }
  mapfile <- cbind(chr.nr, population$info$snp.name, 0 ,population$info$bp)
  mapfile[is.na(mapfile)] <- 0
  haplo1 <- t(haplo[,(1:(ncol(haplo)/2))*2-1])
  haplo2 <- t(haplo[,(1:(ncol(haplo)/2))*2])
  ped <- cbind(haplo1, haplo2)
  ped <- ped[,c(0,ncol(haplo1))+ rep(1:ncol(haplo1), each=2)]
  ped[ped==0] <- "A"
  ped[ped==1] <- "C"
  pedfile <- cbind(1, 1:nrow(ped),0,0,0,0,ped)

  pedname <- paste0(path,".ped")
  utils::write.table(file=pedname, pedfile, col.names = FALSE, row.names = FALSE, quote=FALSE)
  mapname <- paste0(path,".map")
  utils::write.table(file=mapname, mapfile, col.names = FALSE, row.names = FALSE, quote=FALSE)
}
```
