## Supplementary material for "MoBPS - Modular Breeding Program Simulator": MoBPS R-package: get.vcf.R

```
'#
  Authors
Torsten Pook,

Copyright (C) 2017 -- 2018  Torsten Pook

This program is free software; you can redistribute it and/or
modify it under the terms of the GNU General Public License
as published by the Free Software Foundation; either version 3
of the License, or (at your option) any later version.

This program is distributed in the hope that it will be useful,
but WITHOUT ANY WARRANTY; without even the implied warranty of
MERCHANTABILITY or FITNESS FOR A PARTICULAR PURPOSE.  See the
GNU General Public License for more details.

You should have received a copy of the GNU General Public License
along with this program; if not, write to the Free Software
Foundation, Inc., 59 Temple Place - Suite 330, Boston, MA  02111-1307, USA.
'#

#' Generate vcf-file
#'
#' Generate a vcf-file for selected groups and chromosome
#' @param population Population list
#' @param path Location to save vcf-file
#' @param database Groups of individuals to consider for the export
#' @param gen Quick-insert for database (vector of all generations to export)
#' @param cohorts Quick-insert for database (vector of names of cohorts to export)
#' @param chromosomen Beschraenkung des Genotypen auf bestimmte Chromosomen (default: 1)
#' @export

get.vcf <- function(population, path=NULL, database=NULL, gen=NULL, cohorts=NULL, chromosomen="all"){

  haplo <- get.haplo(population, database=database, gen=gen, cohorts=cohorts, chromosomen=chromosomen, export.alleles=FALSE)
# haplo <- get.haplo(population, gen=1)
  if(length(path)==0){
    path <- "population.vcf"
  } else{
    path <- paste0(path,".vcf")
  }
  chr.nr <- sum(population$info$snp)
  start <- 1
  for(index in 1:length(population$info$snp)){
    if(population$info$snp[index]>0){
      chr.nr[start:(start+population$info$snp[index]-1)] <- index
      start <- start + population$info$snp[index]
    }
  }
  map <- data.frame(chr=chr.nr, pos=population$info$bp )
  rownames(map) <- rownames(haplo)
  geno <- haplo[,(1:(ncol(haplo)/2)*2)] + haplo[,(1:(ncol(haplo)/2)*2)-1]
  geno[geno==1] <- haplo[,(1:(ncol(haplo)/2)*2)][geno==1]*2-1

  if (requireNamespace("synbreed", quietly = TRUE)) {
    gp <- synbreed::create.gpData(geno=t(geno), map=map)
    synbreed::write.vcf(gp, file=path, unphased=FALSE)
    cat(paste0("Successfully generated vcf-file in ", getwd(), "/", path))
  } else{
    stop("writting of vcf currently uses synbreed-package. More efficient version coming soon!")
  }
}
```
