## Supplementary material for "MoBPS - Modular Breeding Program Simulator": MoBPS R-package: insert.bve.R

```
'#
  Authors
Torsten Pook,

Copyright (C) 2017 -- 2018  Torsten Pook

This program is free software; you can redistribute it and/or
modify it under the terms of the GNU General Public License
as published by the Free Software Foundation; either version 3
of the License, or (at your option) any later version.

This program is distributed in the hope that it will be useful,
but WITHOUT ANY WARRANTY; without even the implied warranty of
MERCHANTABILITY or FITNESS FOR A PARTICULAR PURPOSE.  See the
GNU General Public License for more details.

You should have received a copy of the GNU General Public License
along with this program; if not, write to the Free Software
Foundation, Inc., 59 Temple Place - Suite 330, Boston, MA  02111-1307, USA.
'#

#' Export estimated breeding values
#'
#' Function to export estimated breeding values
#' @param population Population list
#' @param bves Matrix of breeding values to enter (one row per individual with 1 element coding individual name)
#' @param type which time of values to input (default: "bve", alt: "bv", "pheno")
#' @param count Counting for economic cost calculation (default: 1 - (one observation (for "pheno"), one genotyping (for "bve")))
#' @export

insert.bve <- function(population, bves, type="bve", count=1){

  add <- 2
  if(type=="bv"){
    add <- 6
  } else if(type=="pheno"){
    add <- 8
  }

  for(index in 1:nrow(bves)){
    sex <- as.numeric(substr(bves[index,1], start=1, stop=1)=="F") + 1
    split <- strsplit(bves[index,1], split=c("_"))
    nr <- as.numeric(substr(split[[1]][1], start=2, stop=nchar(split[[1]][1])))
    gen <- as.numeric(split[[1]][2])
    population$breeding[[gen]][[sex+add]][,nr] <- as.numeric(bves[index,-1])
    if(add==2){
      population$breeding[[gen]][[sex]][[nr]][[16]] <- count
    } else if(add==8){
      population$breeding[[gen]][[sex]][[nr]][[15]] <- population$breeding[[gen]][[sex]][[nr]][[15]] + count
    }

  }

  return(population)
}
```
