## Supplementary material for "MoBPS - Modular Breeding Program Simulator": MoBPS R-package: json.simulation.R

```
'#
  Authors
Torsten Pook,

Copyright (C) 2017 -- 2018  Torsten Pook

This program is free software; you can redistribute it and/or
modify it under the terms of the GNU General Public License
as published by the Free Software Foundation; either version 3
of the License, or (at your option) any later version.

This program is distributed in the hope that it will be useful,
but WITHOUT ANY WARRANTY; without even the implied warranty of
MERCHANTABILITY or FITNESS FOR A PARTICULAR PURPOSE.  See the
GNU General Public License for more details.

You should have received a copy of the GNU General Public License
along with this program; if not, write to the Free Software
Foundation, Inc., 59 Temple Place - Suite 330, Boston, MA  02111-1307, USA.
'#

#' Generation of the starting population
#'
#' Generation of the starting population
#' @param file Path to a json-file generated by the user-interface
#' @param total Json-file imported via jsonlite::read_json
#' @param fast.mode Set to TRUE work on a small genome with few markers
#' @param progress.bars Set to TRUE to display progress bars
#' @param size.scaling Scale the size of nodes by this factor (especially for testing smaller examples)
#' @param rep.max Maximum number of repeats to use in fast.mode
#' @param verbose Set to FALSE to not display any prints
#' @export

json.simulation <- function(file=NULL, total=NULL, fast.mode=FALSE,
                            progress.bars=FALSE, size.scaling=NULL, rep.max=1,
                            verbose=TRUE){
  if(requireNamespace("miraculix", quietly = TRUE)){
    miraculix <- TRUE
    miraculix.dataset <- TRUE
  } else{
    miraculix <- FALSE
    miraculix.dataset <- FALSE
  }

  if(length(file)>0){
    total <- jsonlite::read_json(path=file)
  } else if(length(total)==0){
    cat("No dataset provided in file or total \n")
  }

  {
    {
      if(requireNamespace("RandomFieldsUtils", quietly = TRUE)){
        RandomFieldsUtils::RFoptions(helpinfo=FALSE)
      }

      nodes <- total$Nodes
      edges <- total$Edges
      geninfo <- total$`Genomic Info`
      traitinfo <- total$`Trait Info`
      cullinginfo <- total$Culling
      subpopulations <- total$Subpopulation$subpopulation_list
      major_table <- major <- list()
      n_traits <- length(traitinfo)
      map <- NULL
      bp <- NULL


      if(length(geninfo$`speed-mode`)>0 && geninfo$`speed-mode`=="Yes"){
        fast.mode <- TRUE
      }
      if(length(geninfo$`individual-scaling`)>0 && length(size.scaling)==0){
        size.scaling <- as.numeric(geninfo$`individual-scaling`)
      } else if(length(size.scaling)==0){
        size.scaling <- 1
      }


      fixed_costs <- as.numeric(total$Economy$`Fixed Cost`)
      fixed_annual_costs <- 0
      interest_rate <- 1+ as.numeric(total$Economy$`Interest Rate`)/100
      genotyping_costs <- as.numeric(total$Economy$`Genotyping Cost`)

      selection_index_raw <- total$`Selection Index`
      selection_index_scaling_raw <- total$`Selection Index Scaling`

      if(n_traits>0){
        selection_index_name <- numeric(length(selection_index_raw))
        selection_index <- matrix(0, nrow=length(selection_index_raw), ncol=n_traits)
        selection_index_miesenberger <- rep(FALSE,length(selection_index_raw))
        selection_index_miesenberger_wscaling <- rep(FALSE,length(selection_index_raw))
        for(index in 1:length(selection_index_raw)){
          selection_index[index,] <- as.numeric(unlist(selection_index_raw[[index]])[1:n_traits+1])
          selection_index_name[index] <- selection_index_raw[[index]]$'Name'
          if(length(selection_index_scaling_raw[[index]]$w_scaling)>0){
            if(length(selection_index_scaling_raw[[index]]$miesenberger)>0){
              selection_index_miesenberger[index] <- selection_index_scaling_raw[[index]]$miesenberger
            }
            if(selection_index_scaling_raw[[index]]$w_scaling=="Per Unit"){
              selection_index_miesenberger_wscaling[index] <- "unit"
            } else if(selection_index_scaling_raw[[index]]$w_scaling=="Per Phenotypic SD"){
              selection_index_miesenberger_wscaling[index] <- "pheno_sd"
            } else if(selection_index_scaling_raw[[index]]$w_scaling=="Per Breeding Value SD"){
              selection_index_miesenberger_wscaling[index] <- "bve_sd"
            }
          }

        }
      }
      if(sum(is.na(selection_index))>0){
        selection_index[is.na(selection_index)] <- 0
      }

      pheno_index_raw <- total$`Phenotyping Info`

      if(n_traits>0){
        pheno_index_name <- pheno_index_costs <- numeric(length(pheno_index_raw))
        pheno_index <- matrix(0, nrow=length(pheno_index_raw), ncol=n_traits)
        for(index in 1:length(pheno_index_raw)){
          pheno_index[index,] <- as.numeric(unlist(pheno_index_raw[[index]])[1:n_traits+2])
          pheno_index_costs[index] <- as.numeric(pheno_index_raw[[index]][[2]])
          pheno_index_name[index] <- pheno_index_raw[[index]]$'Name'
        }
      } else{
        pheno_index_name <- pheno_index_costs <- numeric(0)
        pheno_index <- NULL
      }

      culling_reason <- matrix(NA, nrow=length(cullinginfo$culling_reasons), ncol=8)
      if(length(culling_reason)>0){
        for(index in 1:nrow(culling_reason)){
          culling_reason[index,] <- unlist(cullinginfo$culling_reasons[[index]])
        }
        rearrange <- sort(as.numeric(culling_reason[,2]), index.return=TRUE)
        culling_reason <- culling_reason[rearrange$ix, ,drop=FALSE]
      }


      housing_index_raw <- total$Economy$`Animal Housing Costs`
      housing_index_name <- housing_index <- numeric(length(housing_index_raw))
      if(length(housing_index_raw)>0){
        for(index in 1:length(housing_index_raw)){
          housing_index[index] <- as.numeric(housing_index_raw[[index]][2])
          housing_index_name[index] <- housing_index_raw[[index]]$'Name'
        }
        housing_cost <- cbind(housing_index_name, housing_index)
      }


      if(n_traits>0){
        trait_matrix <- matrix(0, nrow=n_traits, ncol=8)
        for(index in 1:n_traits){
          trait_matrix[index,] <- unlist(traitinfo[[index]])[1:8]
        }
        # derive genetic variance based on total variance
        pheno_var <- as.numeric(trait_matrix[,4])
        trait_matrix[,4] <- as.numeric(trait_matrix[,4]) * sqrt(as.numeric(trait_matrix[,5]))


        #traitmean <- as.numeric(c(geninfo[[1]]$MilkYield_Mean, geninfo[[1]]$NonReturnRate_Mean, geninfo[[1]]$SomaticCellScore_Mean))
        traitmean <- as.numeric(trait_matrix[,3])
        trait_weigths <- as.numeric(trait_matrix[,8])

        #heritability <- as.numeric(c(geninfo[[1]]$MilkYield_Hertit, geninfo[[1]]$NonReturnRate_Hertit, geninfo[[1]]$SomaticCellScore_Hertit))
        heritability <- as.numeric(trait_matrix[,5])
        if(sum(is.na(heritability))>0){
          if(verbose) cat("No heritability entered - set heritability to 0.5 for all those traits\n")
          heritability[is.na(heritability)]==0
        }

      } else{
        heritability <- NULL
      }

      if(n_traits>0){


        cor_gen <- cor_pheno <- matrix(0, nrow=n_traits, ncol= n_traits)
        nr <- 1
        vector_pheno <- as.numeric(unlist(total$`Phenotypic Correlation`))
        vector_gen <- as.numeric(unlist(total$`Genetic Correlation`))
        for(index1 in 1:n_traits){
          for(index2 in 1:index1){
            cor_pheno[index2,index1] <- cor_pheno[index1,index2] <- vector_pheno[nr]
            cor_gen[index2,index1] <- cor_gen[index1,index2] <- vector_gen[nr]
            nr <- nr + 1
          }
        }

        eigen_gen <- eigen(cor_gen)
        if(sum(eigen_gen$values<0)>0){
          if(verbose) cat("Genetic covariance matrix is not positive definit.\n")
          if(verbose) cat("Generate projection on the set of positive definit matrices:")

          test <- eigen_gen

          test$values[test$values<0] <- 0
          M <- diag(test$values)

          S <- test$vectors

          newA <- S %*% M %*% solve(S)

          diag(newA) <- diag(newA) + 0.0000001 # Avoid numerical issues with inversion
          newA <- newA * matrix(1/sqrt(diag(newA)), nrow=nrow(newA), ncol=nrow(newA), byrow=TRUE) * matrix(1/sqrt(diag(newA)), nrow=nrow(newA), ncol=nrow(newA), byrow=FALSE)
          if(verbose) cat("new suggested residual covariance matrix:\n")
          cor_gen <- newA
          if(verbose) print(round(cor_gen, digits=3))
        }

        if(length(total$`PhenotypicResidual`)>0 && total$`PhenotypicResidual`){
          gen_var <- (diag(trait_matrix[,4])) %*% cor_gen %*% (diag(trait_matrix[,4]))
          pheno_var<-  (diag(pheno_var)) %*% cor_pheno %*% (diag(pheno_var))
          residual_var <- pheno_var - gen_var
          cor_pheno <- sqrt(diag(diag(1/residual_var))) %*% residual_var %*%  sqrt(diag(diag(1/residual_var)))
          if(verbose) cat("Used residual correlations:")
          if(verbose) print(cor_pheno)
        }
        eigen_pheno <- eigen(cor_pheno)
        if(sum(eigen_pheno$values<0)>0){
          if(verbose) cat("Residual covariance matrix is not positive definit.\n")
          if(verbose) cat("Generate projection on the set of positive definit matrices:")

          test <- eigen_pheno

          test$values[test$values<0] <- 0
          M <- diag(test$values)

          S <- test$vectors

          newA <- S %*% M %*% solve(S)

          diag(newA) <- diag(newA) + 0.0000001 # Avoid numerical issues with inversion
          newA <- newA * matrix(1/sqrt(diag(newA)), nrow=nrow(newA), ncol=nrow(newA), byrow=TRUE) * matrix(1/sqrt(diag(newA)), nrow=nrow(newA), ncol=nrow(newA), byrow=FALSE)
          if(verbose) cat("new suggested residual covariance matrix:\n")
          cor_pheno <- newA
          if(verbose) print(round(cor_pheno, digits=3))
        }


      }


      # Creating-types:
      # 0 - Founder
      # 1 - Selection
      # 2 - Reproduction
      # 3 - Recombination
      # 4 - Selfing
      # 5 - DH-Production
      # 6 - Cloning
      # 7 - Combine
      # 8 - Aging
      # 9 - Split


      #### Character - Numeric trafo ######
      for(index in 1:length(nodes)){
        nodes[[index]]$'Number of Individuals' <- as.numeric(nodes[[index]]$'Number of Individuals')
        nodes[[index]]$'Proportion of Male' <- as.numeric(nodes[[index]]$'Proportion of Male')
      }

    }
    #### MANUEL MODIFICATION:
    {

      if(fast.mode){
        if(verbose) cat("Reduce length of genome! Maximum number of Repeat set to ", rep.max, ". I like fast simulations!\n")
        geninfo$`Use Ensembl Map` <- "No"
        geninfo$`Use Own Map` <- "No"
        geninfo$`Chromosomes of Equal Length` <- "Yes"
        geninfo$'Chromosomes Info' <- list()
        geninfo$'Chromosomes Info'[[1]] <- list()
        geninfo$'Chromosomes Info'[[1]]$MD <- 10
        geninfo$'Chromosomes Info'[[1]]$Length <- 20
        geninfo$'Chromosomes Info'[[1]]$Recombination <- 1
        geninfo$'Number of Chromosomes' <- 5
        for(index in 1:length(nodes)){
          nodes[[index]]$'Path' <- NULL
        }
        for(index in 1:length(edges)){
          if(edges[[index]]$`Breeding Type`=="Repeat"){
            edges[[index]]$'Number of Repeat' <- min(as.numeric(edges[[index]]$'Number of Repeat'),rep.max)
          }
        }
      }
      if(size.scaling!=1){
        ids <- numeric(length(nodes))
        for(index in 1:length(nodes)){
          nodes[[index]]$`Number of Individuals` <- round(as.numeric(nodes[[index]]$`Number of Individuals`) * size.scaling, digits=0)
          if( nodes[[index]]$`Number of Individuals`==0){
            nodes[[index]]$`Number of Individuals` <- 1
          }
          ids[index] <- nodes[[index]]$id
        }
        to_node <- NULL
        to_number <- NULL

        for(index in 1:length(edges)){
          if(edges[[index]]$`Breeding Type`=="Combine"){
            to_node <- c(to_node, edges[[index]]$to)
            to_number <- c(to_number, nodes[[which(ids==edges[[index]]$from)]]$`Number of Individuals`)
          }
        }
        for(index in unique(to_node)){
          take <- which(to_node==index)
          if(nodes[[which(ids==index)]]$`Number of Individuals` != sum(to_number[take])){
            if(verbose) cat(paste0("Changed number of individuals in ", index, " from ",nodes[[which(ids==index)]]$`Number of Individuals`, " to ",
                                   sum(to_number[take]), " because of Combine rounding in size scaling.\n"))
            nodes[[which(ids==index)]]$`Number of Individuals` <- sum(to_number[take])
          }

        }

        to_node <- NULL
        to_number <- NULL

        for(index in 1:length(edges)){
          if(edges[[index]]$`Breeding Type`=="Split"){
            to_node <- c(to_node, edges[[index]]$from)
            to_number <- c(to_number, nodes[[which(ids==edges[[index]]$to)]]$`Number of Individuals`)
          }
        }
        for(index in unique(to_node)){
          take <- which(to_node==index)
          if(nodes[[which(ids==index)]]$`Number of Individuals` != sum(to_number[take])){
            if(verbose) cat(paste0("Changed number of individuals in ", index, " from ",nodes[[which(ids==index)]]$`Number of Individuals`, " to ",
                                   sum(to_number[take]), " because of Split rounding in size scaling.\n"))
            nodes[[which(ids==index)]]$`Number of Individuals` <- sum(to_number[take])
          }
        }
      }

      remove.effect.position <- FALSE

      for(index in 1:length(nodes)){
        if(length(nodes[[index]]$Path)>0 && nchar(nodes[[index]]$Path)>0 && nodes[[index]]$`Genotype generation`!="Upload Genotypes"){
          nodes[[index]]$Path <- ""
        }
      }
    }

    ######################## REALITY - CHECKS ############################
    {
      for(index in 1:length(edges)){
        if(length(edges[[index]]$`Time Needed`)>0){
          edges[[index]]$`Time Needed` <- as.numeric(gsub(",", ".", edges[[index]]$`Time Needed`))
        }
        if((edges[[index]]$`Breeding Type`=="Reproduction" || edges[[index]]$`Breeding Type`=="Aging" || edges[[index]]$`Breeding Type`=="Cloning" || edges[[index]]$`Breeding Type`=="Selfing" || edges[[index]]$`Breeding Type`=="DH-Production") && (is.na(as.numeric(edges[[index]]$`Time Needed`)) || as.numeric(edges[[index]]$`Time Needed`)==0 )){
          edges[[index]]$`Time Needed` <- 1
          if(verbose) cat("Edges to generate new individuals are not supposed to take 0 time. Automatically set to 1 time unit.\n")
        }
      }
      for(index in 1:length(nodes)){
        if(nodes[[index]]$Sex=="Both" && (nodes[[index]]$`Proportion of Male`==0 ||nodes[[index]]$`Proportion of Male`==1)){
          nodes[[index]]$`Proportion of Male` <- 0.5
          if(verbose) cat("Both node used with only one sex - set Proportion of male to 0.5. \n")
        }
      }
      for(index in 1:length(nodes)){
        if(length(nodes[[index]]$`Genotype generation`)==0 && nodes[[index]]$Founder=="Yes"){
          nodes[[index]]$`Genotype generation` <- "Random-sampling"
          if(length(nodes[[index]]$Path)>0 && nchar(nodes[[index]]$Path)>0){
            nodes[[index]]$`Genotype generation` <- "Upload Genotypes"
          }
          if(length(nodes[[index]]$`Genotype generation subpopulation`)==0){
            nodes[[index]]$`Genotype generation subpopulation` <- "Population 1"
          }

        } else if(length(nodes[[index]]$`Genotype generation subpopulation`)==0){
          nodes[[index]]$`Genotype generation subpopulation` <- "Population 1"
        }
      }
      for(index in 1:length(edges)){
        if(length(edges[[index]]$'Use Offspring for BVE')==0){
          if(verbose) cat("Manually entered Use offspring for BVE \n")
          edges[[index]]$'Use Offspring for BVE' <- "No" ## Ask Thuy if implemented XXX
        }
      }

      {
        for(index in 1:length(nodes)){
          if(length(nodes[[index]]$earliest_time)==0){
            # cat("Manually entered earliest time.\n")
            nodes[[index]]$earliest_time <- 0
          }
        }
      }
      {
        for(index in 1:length(edges)){
          if(length(edges[[index]]$'Time Needed')==0){
            if(verbose) cat("Manually entered time_needed\n")
            if(edges[[index]]$'Breeding Type'=="Selection" ||edges[[index]]$'Breeding Type'=="Combine" || edges[[index]]$'Breeding Type'=="Split"){
              edges[[index]]$'Time Needed' <- 0
            } else{
              edges[[index]]$'Time Needed' <- 1
            }
          }
        }
      }
      {
        for(index in 1:length(edges)){
          if(edges[[index]]$`Breeding Type`=="Selection" && (length(edges[[index]]$`Selection Index`)==0 || sum(selection_index_name==edges[[index]]$`Selection Index`)==0)){
            if(verbose) cat(paste0("Invalid Selection Index selected - use ", selection_index_name[1], "\n"))
            edges[[index]]$`Selection Index` <- selection_index_name[1]

          }
        }
      }
      # Diagonal of pheno/cor-matrix
      if(n_traits>0 && sum(diag(cor_gen)==1)!= nrow(cor_gen)){
        diag(cor_gen) <- 1
        if(verbose) cat("Diagonal of cor-matrix must be 1\n")
      }
      if(n_traits>0 && sum(diag(cor_pheno)==1)!= nrow(cor_pheno)){
        diag(cor_pheno) <- 1
        if(verbose) cat("Diagonal of cor-matrix must be 1\n")
      }


      # Correct nodes are Founders
      ids <- possible_founder <-  earliest_time <- numeric(length(nodes))
      for(index in 1:length(nodes)){
        ids[index] <- nodes[[index]]$id
        earliest_time[index] <- nodes[[index]]$earliest_time
      }
      for(index in 1:length(edges)){
        possible_founder[which(edges[[index]]$to==ids)] <- 1
      }

      for(index in which(possible_founder==0)){
        if(nodes[[index]]$Founder=="No"){
          nodes[[index]]$Founder <- "Yes"
          if(verbose) cat(paste0("Changed Note ", nodes[[index]]$id, " to Founder-Note! No incoming edge\n"))
        }
      }

      # All edges
      for(index in length(edges):1){
        if(sum(edges[[index]]$to==ids)==0 || sum(edges[[index]]$from==ids)==0){
          if(verbose) cat("Remove illegal edge. Connected Node not present\n")
          edges[[index]] <- NULL
        }
      }

      for(index in 1:length(edges)){
        if(edges[[index]]$'Breeding Type'==""){
          if(verbose) cat(paste0("Edge ", edges[[index]]$Name, " without breeding type. Invalid breeding program!\n"))

        }
      }

      # Selected cohort is not bigger than founder cohort
      for(index in 1:length(edges)){
        from <- which(edges[[index]]$from==ids)
        to <- which(edges[[index]]$to==ids)
        if((nodes[[from]]$'Number of Individuals' < nodes[[to]]$'Number of Individuals') && (edges[[index]]$'Breeding Type'=="Selection")){
          if(verbose) cat(paste0("more individuals selected than in founding note in ", edges[[index]]$Name, ". Automatically changed to selected all individuals\n"))
          nodes[[to]]$'Number of Individuals' <- nodes[[from]]$'Number of Individuals'
        }
      }

    }

    ################### Preparation ############################
    {
      # check for Ind
      same.sex.activ <- FALSE
      same.sex.sex <- 0.5
      for(index in 1:length(nodes)){
        if(nodes[[index]]$'Sex'=="Indefinit"){
          for(index2 in 1:length(nodes)){
            nodes[[index2]]$'Sex' <- "Male"
          }
          same.sex.activ <- TRUE
          same.sex.sex <- 0
        }
      }

      splits <- NULL
      n_male <- NULL
      for(index in 1:length(nodes)){
        if(nodes[[index]]$'Sex'=="Both"){
          splits <- c(splits, nodes[[index]]$id)
          node_id <- nodes[[index]]$id
          nodes[[length(nodes)+1]] <- nodes[[index]]
          nodes[[index]]$'Sex' <- "Male"
          nodes[[length(nodes)]]$'Sex' <- "Female"
          nodes[[index]]$id <- paste0(nodes[[index]]$id, "_M")
          nodes[[length(nodes)]]$id <- paste0(nodes[[length(nodes)]]$id, "_F")
          nodes[[index]]$label <- paste0(nodes[[index]]$label, "_M")
          nodes[[length(nodes)]]$label <- paste0(nodes[[length(nodes)]]$label, "_F")
          nodes[[index]]$'Number of Individuals' <- round(as.numeric(nodes[[index]]$'Number of Individuals') * as.numeric(nodes[[index]]$'Proportion of Male'))
          n_male <- c(n_male,nodes[[index]]$'Number of Individuals')
          nodes[[length(nodes)]]$'Number of Individuals' <- as.numeric(nodes[[length(nodes)]]$'Number of Individuals') - nodes[[index]]$'Number of Individuals'
          nodes[[index]]$'Proportion of Male' <- 1
          nodes[[length(nodes)]]$'Proportion of Male' <- 0

          for(index2 in 1:length(edges)){
            if(length(edges[[index2]]$`Manuel selected cohorts`)>0){
              for(index3 in 1:length(edges[[index2]]$`Manuel selected cohorts`)){
                split_up <- strsplit(edges[[index2]]$`Manuel selected cohorts`[[index3]],":")[[1]]
                if(split_up[1]==node_id){
                  edges[[index2]]$`Manuel selected cohorts`[[index3]] <- paste0(nodes[[index]]$id,":", split_up[2])
                  edges[[index2]]$`Manuel selected cohorts`[[length(edges[[index2]]$`Manuel selected cohorts`)+1]] <- paste0(nodes[[length(nodes)]]$id,":", split_up[2])
                }
              }
            }

          }
        }
      }

      for(index in 1:length(edges)){
        wfrom <- which(edges[[index]]$from==splits)
        wto <- which(edges[[index]]$to==splits)
        if(length(wfrom)>0 && length(wto)>0){
          edges[[length(edges)+1]] <- edges[[index]]
          edges[[index]]$to <- paste0(edges[[index]]$to, "_M")
          edges[[length(edges)]]$to <- paste0(edges[[length(edges)]]$to, "_F")
          edges[[index]]$from <- paste0(edges[[index]]$from, "_M")
          edges[[length(edges)]]$from <- paste0(edges[[length(edges)]]$from, "_F")
        } else if(length(wto)>0){
          edges[[length(edges)+1]] <- edges[[index]]
          edges[[index]]$to <- paste0(edges[[index]]$to, "_M")
          edges[[length(edges)]]$to <- paste0(edges[[length(edges)]]$to, "_F")
        } else if(length(wfrom)>0){
          edges[[length(edges)+1]] <- edges[[index]]
          edges[[index]]$from <- paste0(edges[[index]]$from, "_M")
          edges[[length(edges)]]$from <- paste0(edges[[length(edges)]]$from, "_F")
        }

      }


      ids <- earliest_time <- numeric(length(nodes))
      sex <- numeric(length(nodes))
      for(index in 1:length(nodes)){
        earliest_time[index] <- nodes[[index]]$earliest_time
        ids[index] <- nodes[[index]]$id
        sex[index] <- as.numeric(nodes[[index]]$'Sex'=="Female")
      }

      # Remove sex-switch
      for(index in length(edges):1){
        sex1 <- sex[which(edges[[index]]$from==ids)]
        sex2 <- sex[which(edges[[index]]$to==ids)]
        if(sex1!=sex2 && edges[[index]]$'Breeding Type'!="Reproduction"){
          edges[[index]] <- NULL
          #      if(sex1==2){
          #        cat("Invalid Edge to generate female from male individuals")
          #      } else{
          #        cat("Invalid Edge to generate male from female individuals")
          #      }

        }
      }

      # Identify founder cohorts
      # Add "Founder" as a breeding type - If selected allow for a dataset to be imported!
      founder <- NULL
      ids <- earliest_time <-  numeric(length(nodes))
      repeat_node <- numeric(length(nodes))
      incoming_repeat_node <- numeric(length(nodes))
      for(index in 1:length(nodes)){
        if(nodes[[index]]$Founder=="Yes"){
          founder <- c(founder,index)
        }
        ids[index] <- nodes[[index]]$id
        earliest_time[index] <- nodes[[index]]$earliest_time
      }
      for(index in 1:length(edges)){
        if(edges[[index]]$'Breeding Type'=="Repeat"){
          repeat_node[which(edges[[index]]$from==ids)] <- 1
          incoming_repeat_node[which(edges[[index]]$to==ids)] <- 1
        }
      }
    }
    ## REPEATS
    {
      repeat_links <- NULL
      for(index in 1:length(edges)){
        if(edges[[index]]$`Breeding Type`=="Repeat"){
          repeat_links <- rbind(repeat_links, c(edges[[index]]$from, edges[[index]]$to))
        }
      }

      while(sum(repeat_node)>0){
        n_nodes <- length(nodes)
        n_edges <- length(edges)
        link <- NULL
        link2 <- NULL
        start <- founder
        step <- start
        step_vali <- intersect(start, which(incoming_repeat_node==1))

        nodes_to_repeat <- NULL
        edges_to_repeat <- NULL
        prev <- NULL

        while(length(step)!=length(prev) || length(prev)==0){
          prev <- step
          for(index2 in 1:length(edges)){
            desti <- which(edges[[index2]]$to==ids)
            ori <- which(edges[[index2]]$from==ids)
            if(length(intersect(ori, step))>0){
              if(repeat_node[ori]==1){
                if(edges[[index2]]$`Breeding Type`=="Repeat"){
                  start <- unique(c(start, desti))
                  step <- unique(c(step, desti))
                  if(incoming_repeat_node[desti] || length(intersect(step_vali,ori))>0){
                    step_vali <- unique(c(step_vali, desti))
                  }
                  link <- rbind(link, c(ori, desti))
                  link2 <- unique(c(link2, index2))
                }
              } else{
                if(sum(ori==step)>0){
                  step <- unique(c(step, desti))

                  #if(incoming_repeat_node[desti]==1 && length(intersect(founder,desti))>0){
                  #  step_vali <- unique(c(step_vali, desti))
                  #} else
                  #if(incoming_repeat_node[desti]==1 || length(intersect(step_vali,ori))>0){
                  #  step_vali <- unique(c(step_vali, desti))
                  #  edges_to_repeat <- unique(c(edges_to_repeat, index2))
                  #}

                  if(incoming_repeat_node[desti]==1 || length(intersect(step_vali,ori))>0){
                    step_vali <- unique(c(step_vali, desti))
                  }

                }

              }
            }
          }
        }
        nodes_to_repeat <- step_vali[!duplicated(c(start, step_vali))[-(1:length(start))]]
        nodes_to_repeat <- sort(nodes_to_repeat)
        n_rep <- 0
        for(index in 1:length(edges)){
          if(sum(edges[[index]]$to==ids[nodes_to_repeat])>0 && edges[[index]]$`Breeding Type`!="Repeat"){
            edges_to_repeat <- c(edges_to_repeat, index)
          }
          if(sum(edges[[index]]$from==ids[nodes_to_repeat]) && edges[[index]]$`Breeding Type`=="Repeat"){
            n_rep <- as.numeric(edges[[index]]$`Number of Repeat`)
          }
        }


        #    edges_to_repeat <- sort(edges_to_repeat)
        link <- unique(link)
        #    for(index4 in start){
        #      edges[[index4]]$'Breeding Type' <- "PerformedRepeat"
        #    }
        for(index4 in 1:n_rep){
          for(nodrep in nodes_to_repeat){
            nodes[[length(nodes)+1]] <- nodes[[nodrep]]
            nodes[[length(nodes)]]$Founder <- "No"
            nodes[[length(nodes)]]$id <- paste0(nodes[[length(nodes)]]$id, "_",index4)
            nodes[[length(nodes)]]$label <- paste0(nodes[[length(nodes)]]$label, "_",index4)
          }
          for(edgesrep in edges_to_repeat){
            edges[[length(edges)+1]] <- edges[[edgesrep]]
            test <- ids[start]==edges[[length(edges)]]$from
            if(sum(edges[[edgesrep]]$from==ids[step_vali])==0){
              if(verbose) cat(paste0("Repeat from edge not included in the repeat: ", edges[[edgesrep]]$from, "\n"))
            } else if(sum(test)){
              take <- link[which(link[,2]==start[which(test)]),1]
              if(index4==1){
                edges[[length(edges)]]$from <- paste0(ids[take])
              } else{
                edges[[length(edges)]]$from <- paste0(ids[take],"_", index4-1)
              }

            } else{
              edges[[length(edges)]]$from <- paste0(edges[[length(edges)]]$from,"_", index4)
            }

            edges[[length(edges)]]$to <- paste0(edges[[length(edges)]]$to,"_", index4)
            edges[[length(edges)]]$id <- paste0(edges[[length(edges)]]$id,"_", index4)
          }
        }

        for(sonstcheck in (1:length(ids))[-step_vali]){
          for(edgeindex in 1:length(edges)){
            if(edges[[edgeindex]]$to==ids[sonstcheck] && sum(edges[[edgeindex]]$from==ids[nodes_to_repeat])>0){
              edges[[edgeindex]]$from <- paste0(edges[[edgeindex]]$from, "_", n_rep)
            }
          }
        }
        for(index in 1:length(edges)){
          if(edges[[index]]$`Breeding Type`=="Repeat" && sum(ids[step_vali]==edges[[index]]$to)>0){
            edges[[index]]$to <- paste0(edges[[index]]$to, "_", n_rep)
          }
        }


        for(index in 1:nrow(link)){
          if(verbose) cat(paste0("Successful repeat for ", ids[link[index,1]] ," to ",ids[link[index,2]],"\n" ))
        }
        if(verbose) cat(paste0(length(nodes)-n_nodes, " Nodes & ", length(edges) - n_edges, " Edges were added to generation script.\n"))


        ids <- earliest_time <- numeric(length(nodes))
        founder <- NULL
        for(index5 in 1:length(nodes)){
          if(nodes[[index5]]$Founder=="Yes"){
            founder <- c(founder,index5)
          }
          ids[index5] <- nodes[[index5]]$id
          earliest_time[index5] <- nodes[[index5]]$earliest_time
        }

        for(changeto in sort(link2, decreasing = TRUE)){
          edges[[changeto]] <- NULL
        }

        repeat_node <- incoming_repeat_node <- numeric(length(nodes))

        for(index in 1:length(edges)){
          if(edges[[index]]$'Breeding Type'=="Repeat"){

            repeat_node[which(edges[[index]]$from==ids)] <- 1
            incoming_repeat_node[which(edges[[index]]$to==ids)] <- 1
          }
        }

      }

      ## Change manuel selected according to repeats

      for(index in 1:length(edges)){
        if(length(edges[[index]]$`Manuel selected cohorts`)>0){
          for(index2 in 1:length(edges[[index]]$`Manuel selected cohorts`)){
            split_up <- strsplit(edges[[index]]$`Manuel selected cohorts`[[index2]], ":")[[1]]
            repeat_change <- which(split_up[1]==repeat_links[,2])
            if(length(repeat_change)==1 && split_up[2] != "X"){
              edges[[index]]$`Manuel selected cohorts`[[index2]] <- paste0(repeat_links[repeat_change,1], ":", as.numeric(split_up[2])-1)
            }
          }

        }
      }


      ## Determine genetic architecture

      if(geninfo$'Use Ensembl Map'=="Yes"){
        if (requireNamespace("MoBPSmaps", quietly = TRUE)) {
          if(geninfo$`Ensembl Dataset`=="Axiom Genotyping Array"){
            map <- MoBPSmaps::map_pig1
          } else if(geninfo$`Ensembl Dataset`=="GGP Porcine HD"){
            map <- MoBPSmaps::map_pig2
          } else if(geninfo$`Ensembl Dataset`=="GGP Porcine LD"){
            map <- MoBPSmaps::map_pig3
          } else if(geninfo$`Ensembl Dataset`=="Illumina_PorcineSNP60"){
            map <- MoBPSmaps::map_pig4
          } else if(geninfo$`Ensembl Dataset`=="Affymetrix Chicken600K Array"){
            map <- MoBPSmaps::map_chicken1
          } else if(geninfo$`Ensembl Dataset`=="Affymetrix Chicken600K Array_subset_diversity"){
            map <- MoBPSmaps::map_chicken2
          } else if(geninfo$`Ensembl Dataset`=="Affymetrix Chicken600K Array_subset_50k"){
            map <- MoBPSmaps::map_chicken3
          } else if(geninfo$`Ensembl Dataset`=="Illumina BovineSNP50 BeadChip"){
            map <- MoBPSmaps::map_cattle1
          } else if(geninfo$`Ensembl Dataset`=="Illumina BovineHD BeadChip"){
            map <- MoBPSmaps::map_cattle2
          } else if(geninfo$`Ensembl Dataset`=="Illumina BovineLD BeadChip"){
            map <- MoBPSmaps::map_cattle3
          }else if(geninfo$`Ensembl Dataset`=="Genotyping chip variations"){
            map <- MoBPSmaps::map_cattle4
          }else if(geninfo$`Ensembl Dataset`=="Illumina EquineSNP50 BeadChip"){
            map <- MoBPSmaps::map_horse1
          }else if(geninfo$`Ensembl Dataset`=="IlluminaOvineHDSNP"){
            map <- MoBPSmaps::map_sheep1
          }else if(geninfo$`Ensembl Dataset`=="IlluminaOvineSNP50"){
            map <- MoBPSmaps::map_sheep2
          } else if(geninfo$`Ensembl Dataset`=="Genotyping chip variants"){
            map <- MoBPSmaps::map_sheep3
          } else if(geninfo$`Ensembl Dataset`=="Illumina_GoatSNP50"){
            map <- MoBPSmaps::map_goat1
          } else if(geninfo$`Ensembl Dataset`=="Affy GeneChip500K"){
            map <- MoBPSmaps::map_human1
          } else if(geninfo$`Ensembl Dataset`=="Illumina_1M-duo"){
            map <- MoBPSmaps::map_human2
          } else if(geninfo$`Ensembl Dataset`== "Illumina_HumanHap550"){
            map <- MoBPSmaps::map_human3
          } else if(geninfo$`Ensembl Dataset`== "Affymetrix Axiom Maize Genotyping Array"){
            map <- MoBPSmaps::map_maize1
          } else if(geninfo$`Ensembl Dataset`== "3k_Sorghum_Bekele"){
            map <- MoBPSmaps::map_sorghum1
          } else if(geninfo$`Ensembl Dataset`== "12k_Wheat_Bekele"){
            map <- MoBPSmaps::map_wheat1
          } else if(geninfo$`Ensembl Dataset`== "29k_Wheat_Wen"){
            map <- MoBPSmaps::map_wheat2
          } else{
            map <- ensembl.map(dataset = geninfo$'Ensembl Dataset',
                               filter = geninfo$'Ensembl Filter',
                               filter.values = geninfo$'Ensembl Filter Values')
          }
        } else{
          stop("Use of Ensembl-Maps without MoBPSmaps R-package")
        }


        if(length(geninfo$'Max Number of SNPs')>0 && as.numeric(geninfo$'Max Number of SNPs')<nrow(map)){
          map <- map[sort(sample(1:nrow(map), as.numeric(geninfo$'Max Number of SNPs'))),]
        }


        chr.opt <- unique(map[,1])
        nchromo <- length(chr.opt)
        nsnp <- chromo.length <- numeric(nchromo)
        for(index in 1:nchromo){
          nsnp[index] <- sum(map[,1]==chr.opt[index])
          chromo.length[index] <- max(as.numeric(map[map[,1]==chr.opt[index],3])) / 100000000

        }
        if(verbose) cat("Assume 100.000.000 bp/M in Ensembl Map \n")

      } else if(geninfo$'Use Own Map'=="Yes" || length(map)>0){
        map_path <- geninfo$`Own Map Path`
        if(length(map_path)>0){
          map_type <- substr(map_path, start= nchar(map_path)-2, stop= nchar(map_path))
          if(map_type=="vcf"){
            if(verbose) cat("Map identified as vcf-file - extract map information")
            if(requireNamespace("vcfR", quietly = TRUE)){
              vcf_file <- vcfR::read.vcfR(map_path)
              map <- cbind(vcf_file@fix[,c(1,3,2)],NA,NA)
            } else{
              stop("Map-import failed! vcfR-package not available! \n")
            }
          } else if(map_type=="map"){
            if(verbose) cat("Map identified as Ped-map-file - extract map information")
            map_file <- utils::read.table(map_path)
            map <- cbind(map_file[,c(1,2,4,3)],NA)
            if(sum(map[,4])==0){
              map[,4] <- NA
            }
          } else if(map_type=="ata"){
            map_store <- load(map_path)
            if(length(map_store)>1){
              if(verbose) cat("More than one object contain in .Rdata - unless the map is named 'map' import will fail! \n")
            } else{
              map <- eval(parse(text=map_store[[1]]))
            }
          }
          map <- as.matrix(map)

        }

        chr.opt <- unique(map[,1])
        nchromo <- length(chr.opt)
        nsnp <- chromo.length <- numeric(nchromo)

        for(index in 1:nchromo){
          nsnp[index] <- sum(map[,1]==chr.opt[index])
          chromo.length[index] <- max(as.numeric(map[map[,1]==chr.opt[index],3])) / 100000000

        }
        if(verbose) cat("Assume 100.000.000 bp/M in Imported Map \n")

      } else if(geninfo$'Chromosomes of Equal Length'=="Yes"){
        nchromo <- as.numeric(geninfo$'Number of Chromosomes')
        nsnp <- ceiling(rep(as.numeric(geninfo$'Chromosomes Info'[[1]]$Length) * as.numeric(geninfo$'Chromosomes Info'[[1]]$MD), nchromo))
        chromo.length <- rep(as.numeric(geninfo$'Chromosomes Info'[[1]]$Length) * as.numeric(geninfo$'Chromosomes Info'[[1]]$Recombination, nchromo) /100, nchromo)
        bp <- ceiling((0.5:nsnp[1]) / nsnp[1] * as.numeric(geninfo$'Chromosomes Info'[[1]]$Length) * 1000000)

      } else{
        nchromo <- as.numeric(geninfo$'Number of Chromosomes')
        chromo.length <- nsnp <- numeric(nchromo)
        bp <- numeric(sum(nsnp))
        prev <- 0
        for(index in 1:nchromo){
          nsnp[index] <- ceiling(as.numeric(geninfo$'Chromosomes Info'[[index]]$Length) * as.numeric(geninfo$'Chromosomes Info'[[index]]$MD))
          chromo.length[index] <- as.numeric(geninfo$'Chromosomes Info'[[index]]$Length) * as.numeric(geninfo$'Chromosomes Info'[[index]]$Recombination, nchromo) /100
          bp[(1:nsnp[index])+prev] <- ceiling((0.5:nsnp[index]) / nsnp[index] * as.numeric(geninfo$'Chromosomes Info'[[index]]$Length) * 1000000)
          prev <- prev + nsnp[index]
        }

      }
      if(length(map)==0){
        for(index in 1:nchromo){
          map <- rbind(map, cbind(index, paste0("SNP", 1:nsnp[index]), NA, NA, NA))
        }
        if(length(bp)>0){
          map[,3] <- bp
        } else{
          if(verbose) cat("Why is there no base-pair in auto generated map?")
        }
      }


      groups <- length(nodes)


      # Species not needed, traitvariance not needed,
      position <- matrix(0, nrow=length(nodes), ncol=4)
      rownames(position) <- ids
      founding_a <- c(0,0)
      mig_m <- numeric(0)
      mig_f <- numeric(0)

      sex.s <- NULL
      genotyped.s <- NULL
      # CHECK FOR ME THAN 2 FOUNDERS (MORE THAN 1 of a sex- Migration level...)
      new_mig <- c(0,0)


      founder_data <- FALSE
      for(index in 1:length(founder)){

        sex <- as.numeric(nodes[[founder[index]]]$'Sex'=="Female")+1
        size <- as.numeric(nodes[[founder[index]]]$'Number of Individuals')
        founding_a[sex] <- founding_a[sex] + size
        sex.s <- c(sex.s, rep(sex, size))
        genotyped.s <- c(genotyped.s, stats::rbinom(as.numeric(nodes[[founder[index]]]$'Number of Individuals'), 1,
                                                    as.numeric(nodes[[founder[index]]]$`Proportion of genotyped individuals`)))
        if(sex==1){
          mig_m <- c(mig_m, rep(new_mig[1], size))
          mig <- new_mig[1]
          new_mig[1] <- new_mig[1] + 1
        }
        if(sex==2){
          mig_f <- c(mig_f, rep(new_mig[2], size))
          mig <- new_mig[2]
          new_mig[2] <- new_mig[2] + 1
        }
        position[founder[index],] <- c(1, sex, mig, size)
        if(nodes[[index]]$`Genotype generation` == "Upload Genotypes" && length(nodes[[founder[index]]]$Path)>0 && nchar(nodes[[founder[index]]]$Path)>0){
          founder_data <- TRUE
        }
      }


    }


    ################### Import genetic data  + major QTL import ####################
    {
      path_list <- NULL
      phasing_list <- NULL
      no_data <- numeric(length(founder))
      founder_pop <- numeric(length(founder))
      for(index in 1:length(founder)){
        path_list <- c(path_list, nodes[[founder[index]]]$Path )
        if(length(nodes[[founder[index]]]$'phasing_required') >0 && nodes[[founder[index]]]$'phasing_required'=="Yes"){
          phasing_list <- c(phasing_list, nodes[[founder[index]]]$Path )
        }
        founder_pop[index] <- nodes[[founder[index]]]$`Genotype generation subpopulation`
      }

      p_i <- list()
      major_sub <- list()
      major_table_sub <- list()
      subpopulation_info <- matrix(0, nrow=length(subpopulations), ncol=6)
      for(index in 1:length(subpopulations)){
        subpopulation_info[index,] <- unlist(subpopulations[[index]])[1:6]
      }

      if(length(subpopulations)>1 & miraculix.dataset){
        miraculix.dataset <- FALSE
        if(verbose) cat("miraculix dataset generation only for single subpopulation.\n")
      }

      if(length(unique(c("",path_list)))>1 && miraculix.dataset){
        miraculix.dataset <- FALSE
        if(verbose) cat("Detected data-path for founders. Deactive miraculix.dataset.\n")
      }
      if(!miraculix.dataset){
        dataset <- matrix(0L, nrow= sum(nsnp), ncol = sum(position[,4])*2)
      }

      starts <- cumsum(c(1, position[founder,4]))*2-1
      for(path in unique(c("",path_list))[-1]){
        data_path <- path
        if(length(data_path)>0){
          data_type <- substr(data_path, start= nchar(data_path)-2, stop= nchar(data_path))
          if(data_type=="vcf"|| data_type==".gz"){
            if(verbose) cat("Data input identified as vcf-file - extract genomic information \n")
            if(requireNamespace("vcfR", quietly = TRUE)){

              if(sum(path==phasing_list)>0){
                pedmap.to.phasedbeaglevcf(vcf_path = data_path)
                data_path <- "/home/nha/Plink/DB/temp_vcf_phased.vcf.gz"
              }

              vcf_file <- vcfR::read.vcfR(data_path)
              haplo1 <- substr(vcf_file@gt[,-1], start=1, stop=1)
              haplo2 <- substr(vcf_file@gt[,-1], start=3, stop=3)
              storage.mode(haplo1) <- storage.mode(haplo2) <- "integer"
              haplo_temp <- cbind(haplo1, haplo2)
              haplo_temp <- haplo_temp[,c(0,ncol(haplo1)) + rep(1:ncol(haplo1), each=2)]

              order <- numeric(nrow(map))
              prev <- 0
              it <- nrow(vcf_file@fix)
              for(index in 1:it){
                if(prev < it && map[prev+1,2]==vcf_file@fix[index,3]){
                  prev <- prev +1
                } else{
                  prev <- which(map[,2]==vcf_file@fix[index,3])
                }
                order[index] <- prev
              }
              haplo <- matrix(0,nrow=nrow(map), ncol=ncol(haplo_temp))
              haplo[order,] <- haplo_temp
            } else{
              stop("Data-import failed! vcfR-package not available! \n")
            }
          } else if(data_type=="ped"){
            if(verbose) cat("Data input identified as Ped-map-file - extract genomic information \n")
            if(sum(path==phasing_list)>0){
              pedmap.to.phasedbeaglevcf(ped_path=path, map_path=geninfo$`Own Map Path`)
              if(requireNamespace("vcfR", quietly = TRUE)){
                vcf_file <- vcfR::read.vcfR("/home/nha/Plink/DB/temp_vcf_phased.vcf.gz")
                haplo1 <- substr(vcf_file@gt[,-1], start=1, stop=1)
                haplo2 <- substr(vcf_file@gt[,-1], start=3, stop=3)
                storage.mode(haplo1) <- storage.mode(haplo2) <- "integer"
                haplo_temp <- cbind(haplo1, haplo2)
                haplo_temp <- haplo_temp[,c(0,ncol(haplo1)) + rep(1:ncol(haplo1), each=2)]

                order <- numeric(nrow(map))
                prev <- 0
                it <- nrow(vcf_file@fix)
                for(index in 1:it){
                  if(prev < it && map[prev+1,2]==vcf_file@fix[index,3]){
                    prev <- prev +1
                  } else{
                    prev <- which(map[,2]==vcf_file@fix[index,3])
                  }
                  order[index] <- prev
                }
                haplo <- matrix(0,nrow=nrow(map), ncol=ncol(haplo_temp))
                haplo[order,] <- haplo_temp
              } else{
                stop("Data-import failed! vcfR-package not available! \n")
              }
            } else{
              if(verbose) cat("Haplotype phase is assumed to be by colum - No internal phasing performed! \n")
              ped_file <- utils::read.table(data_path)
              haplo12 <- t(ped_file[,-(1:6)])
              haplo <- matrix(0, ncol = ncol(haplo12)*2, nrow=nrow(haplo12)/2)
              for(index1 in 1:ncol(haplo12)){
                haplo[,index1*2+c(-1,0)] <- matrix(haplo12[,index1], ncol=2, byrow=TRUE)
              }
            }


          } else if(data_type=="ata"){
            Map <- map ## Lisas subpos contain a element named map"
            data_store <- load(data_path)
            map <- Map
            if(length(data_store)>1){
              if(verbose) cat("More than one object contain in .Rdata - unless the data object is named 'haplo' import will fail! \n")
            } else{
              haplo <- eval(parse(text=map_store[[1]]))
            }
          }
        }

        takes <- which(path_list==path)
        no_data[takes] <- 1
        take <- NULL
        for(index in takes){
          take <- c(take, starts[index]:(starts[index+1]-1))
        }
        if(nrow(haplo)!=sum(nsnp)){
          if(data_type=="vcf" || data_type==".gz"){
            chr.opt <- unique(vcf_file@fix[,1])
            nsnp <- numeric(length(chr.opt))
            for(index in 1:length(chr.opt)){
              nsnp[index] <- sum(vcf_file@fix[,1]==chr.opt[index])
            }
          } else{
            nsnp <- floor(nsnp*nrow(haplo)/sum(nsnp))
            nsnp[1] <- nsnp[1] + nrow(haplo)-sum(nsnp)
          }

          dataset <- matrix(0L, nrow= sum(nsnp), ncol = sum(position[,4])*2)
          map <- NULL
          for(index in 1:nchromo){
            map <- rbind(map, cbind(index, paste0("SNP", 1:nsnp[index]), NA, NA, NA))
          }
        }
        if(length(haplo) != length(dataset[,take])){
          if(verbose) cat("Size of Founder-dataset not in concorance with node size!\n")
          haplo <- rep(haplo, length.out=length(dataset[,take]))
        }
        if(storage.mode(haplo)!="integer"){
          storage.mode(haplo) <- "integer"
        }
        dataset[,take] <- haplo
      }

      if((n_traits)>0){
        for(index in 1:n_traits){
          major[[index]] <- traitinfo[[index]]$'Trait QTL Info'

          if(length(major[[index]])>0 && length(major[[index]][[1]])>1){
            ncols <- length(major[[index]][[1]])
            mqtl <- matrix(unlist(major[[index]]), ncol=ncols, byrow=TRUE)
            to_enter <- mqtl[,c(1,4:8), drop=FALSE]
            to_enter_name <- mqtl[,c(2,3), drop=FALSE]
            storage.mode(to_enter) <- "numeric"
            if(sum(is.na(to_enter[,2]))>0 || sum(is.na(to_enter[,1]))>0){
              check_qtl <- unique(c(which(is.na(to_enter[,1])), which(is.na(to_enter[,2]))))

              for(sample_index in check_qtl){
                take_qtl <- which(to_enter_name[sample_index,1] == map[,2])
                if(length(take_qtl)>0){
                  to_enter[sample_index, 2] <- as.numeric(map[take_qtl[1], 1])
                  to_enter[sample_index, 1] <- sum(map[1:take_qtl[1], 1] == map[take_qtl[1], 1])
                  if(verbose) cat("SNP-position assigned via SNP-name\n")
                } else{
                  if(is.na(to_enter[sample_index,2])){
                    to_enter[sample_index,2] <- 1
                    if(verbose) cat("Illegal Chromosom-name. Simulate major SNP-effect on chromosome 1.\n")
                  }
                  if(!is.na(as.numeric(to_enter_name[sample_index,2]))){
                    diff_to <- - abs(as.numeric(map[,3]) - as.numeric(to_enter_name[sample_index,2]))
                    diff_to[map[,1]!=to_enter[sample_index,2]] <- -Inf
                    take_qtl <- which.max(diff_to)
                    to_enter[sample_index, 2] <- as.numeric(map[take_qtl[1], 1])
                    to_enter[sample_index, 1] <- sum(map[1:take_qtl[1], 1] == map[take_qtl[1], 1])
                    if(verbose) cat("SNP-position assigned via bp position\n")
                  } else{
                    to_enter[sample_index,1] <- sample(1:nsnp[to_enter[sample_index,2]],1)
                    if(verbose) cat(paste0("Illegal SNP-name/bp/position. SNP-effect generated on SNP ", to_enter[sample_index,1],"\n"))
                  }

                }

              }

            }

            to_enter[to_enter[,6]=="",6] <- NA
            major_table[[index]] <- to_enter
          }

        }
      }

      # Change Allele frequencies according to selection in major QTLs
      if(length(major_table)>0){
        for(index in 1:length(major_table)){
          if(length(major_table[[index]])>0){
            for(index2 in 1:nrow(major_table[[index]])){
              take <- which(cumsum((map[,1]==major_table[[index]][index2,2]))==major_table[[index]][index2,1])[1]
              map[take,5] <- major_table[[index]][index2,6]
            }
          }

        }
      }


      for(subpop in 1:length(subpopulations)){
        if(sum(no_data==0 & founder_pop==subpopulation_info[subpop,1])>0){
          takes <- which(no_data==0 & founder_pop==subpopulation_info[subpop,1])
          take <- NULL
          for(index in takes){
            take <- c(take, starts[index]:(starts[index+1]-1))
          }

          if(sum(no_data==1 & founder_pop==subpopulation_info[subpop,1])>0){

            takes2 <- which(no_data==1 & founder_pop==subpopulation_info[subpop,1])
            take2 <- NULL
            for(index in takes2){
              take2 <- c(take2, starts[index]:(starts[index+1]-1))
            }
            if(miraculix.dataset){
              p_i[[subpop]] <- miraculix::allele_freq(miraculix::genomicmatrix(dataset[,take2]))*2
            } else{
              p_i[[subpop]] <- rowMeans(dataset[,take2])
            }
          } else{
            p_i[[subpop]] <- stats::rbeta(sum(nsnp), shape1=as.numeric(subpopulation_info[subpop,2]),
                                          shape2=as.numeric(subpopulation_info[subpop,3]))
            set_zero <- sample(1:sum(nsnp), sum(nsnp) * as.numeric(subpopulation_info[subpop,4]))
            if(length(set_zero)>0){
              set_one <- sample((1:sum(nsnp))[-set_zero], sum(nsnp) * as.numeric(subpopulation_info[subpop,5]))
            } else{
              set_one <- sample((1:sum(nsnp)), sum(nsnp) * as.numeric(subpopulation_info[subpop,5]))
            }
            if(length(set_zero)>0){
              p_i[[subpop]][set_zero] <- 0
            }
            if(length(set_one)>0){
              p_i[[subpop]][set_one] <- 1
            }
          }

          if(sum(!is.na(map[,5]))){
            p_i[[subpop]][!is.na(map[,5])] <- as.numeric(map[!is.na(map[,5]),5])
          }
          ## Manually selected replaces!
          major_sub[[index]] <- subpopulations[[subpop]]$'QTL Info'
          if(length(major_sub[[index]])>0 && length(major_sub[[index]][[1]])>1){
            ncols <- length(major_sub[[index]][[1]])
            mqtl_sub <- matrix(unlist(major_sub[[index]]), ncol=ncols, byrow=TRUE)
            to_enter_sub <- mqtl_sub[,c(1,4:6), drop=FALSE]
            to_enter_name_sub <- mqtl[,c(2,3), drop=FALSE]
            storage.mode(to_enter_sub) <- "numeric"
            if(sum(is.na(to_enter_sub[,2]))>0 || sum(is.na(to_enter_sub[,1]))>0){
              check_qtl <- unique(c(which(is.na(to_enter_sub[,1])), which(is.na(to_enter_sub[,2]))))
              for(sample_index in check_qtl){
                take_qtl <- which(to_enter_name_sub[sample_index,1] == map[,2])
                if(length(take_qtl)>0){
                  to_enter_sub[sample_index, 2] <- as.numeric(map[take_qtl[1], 1])
                  to_enter_sub[sample_index, 1] <- sum(map[1:take_qtl[1], 1] == map[take_qtl[1], 1])
                  if(verbose) cat("SNP-position assigned via SNP-name\n")
                } else{
                  if(is.na(to_enter_sub[sample_index,2])){
                    to_enter_sub[sample_index,2] <- 1
                    if(verbose) cat("Illegal Chromosom-name. Simulate major SNP-effect on chromosome 1")
                  }
                  if(!is.na(as.numeric(to_enter_name_sub[sample_index,2]))){
                    diff_to <- - abs(as.numeric(map[,3]) - as.numeric(to_enter_name_sub[sample_index,2]))
                    diff_to[map[,1]!=to_enter_sub[sample_index,2]] <- -Inf
                    take_qtl <- which.max(diff_to)
                    to_enter_sub[sample_index, 2] <- as.numeric(map[take_qtl[1], 1])
                    to_enter_sub[sample_index, 1] <- sum(map[1:take_qtl[1], 1] == map[take_qtl[1], 1])
                    if(verbose) cat("SNP-position assigned via bp position\n")
                  } else{
                    to_enter_sub[sample_index,1] <- sample(1:nsnp[to_enter_sub[sample_index,2]],1)
                    if(verbose) cat(paste0("Illegal SNP-name/bp/position. SNP-effect generated on SNP ", to_enter_sub[sample_index,1],"\n"))
                  }
                }
              }

            }

            to_enter_sub[to_enter_sub[,4]=="",4] <- NA
            major_table_sub[[index]] <- to_enter_sub
            position_change <- cumsum(c(0, nsnp))[to_enter_sub[,2]] + to_enter_sub[,1]
            p_i[[subpop]][position_change] <- to_enter_sub[,3]
          }

        }
        if(miraculix.dataset){
          nindi <- (starts[max(takes)+1] - starts[min(takes)])/2
          chr.nr <- map[,1]
          chr.opt <- unique(chr.nr)
          dataset <- list()
          for(chr_index in 1:length(chr.opt)){
            dataset[[chr_index]] <- miraculix::rhaplo(p_i[[subpop]][which(chr.nr==chr.opt[chr_index])], indiv = nindi, loci = nsnp[chr_index])
          }
        } else{
          for(index in takes){
            # Integer * Integer > 2^32 --> NA! conversion to numerics
            take <- starts[index]:(starts[index+1]-1)
            input_type <- nodes[[founder[index]]]$`Genotype generation`
            if(input_type=="Random-sampling"){
              dataset[,take] <- stats::rbinom(as.numeric(nrow(dataset))*as.numeric(length(take)),1, prob=p_i[[subpop]])
            } else if(input_type=="Fully-homozygous"){
              dataset[,take[c(1:(length(take)/2)*2-1,1:(length(take)/2)*2)]] <- stats::rbinom(as.numeric(nrow(dataset))*as.numeric(length(take))/2,1, prob=p_i[[subpop]])
              # generate half as many alleles as spots to fill - rest is autocompleted
            } else if(input_type=="Fully-heterozygous"){
              dataset[,c(1:(length(take)/2)*2)-1] <- 0L # just for safety but not necessary
              dataset[,c(1:(length(take)/2)*2)] <- 1L
            } else if(input_type=="All-B-Allele"){
              dataset[,take] <- 1L
            } else if(input_type=="All-A-Allele "){
              dataset[,take] <- 0L
            } else{
              stop("Invalid input type for founder node!")
            }
          }
        }
      }


    }

    ############### Generate Base-Population
    {
      population <- NULL

      population <- creating.diploid(dataset = dataset, nindi=length(sex.s),
                                     sex.s = sex.s, genotyped.s = genotyped.s,
                                     chromosome.length = chromo.length,
                                     snps.equidistant = if(is.na(map[1,4])) {TRUE} else {FALSE}, miraculix = miraculix,
                                     miraculix.dataset = miraculix.dataset,
                                     chr.nr = map[,1], bp=map[,3], snp.name = map[,2],
                                     freq = map[,5], snp.position = if(is.na(map[1,4])) {NULL} else {map[,4]})

      # Cohort names
      gender_founder <- numeric(length(founder))
      nr <- 1
      for(index in 1:length(founder)){
        gender_founder[index] <- sex.s[nr]
        nr <- nr + position[founder[index],4]
      }
      founder_temp2 <- sort(gender_founder,index.return=TRUE)$ix
      founder_temp <- founder[sort(gender_founder,index.return=TRUE)$ix]
      cohort_info <- cbind(ids[founder_temp], 1, position[founder_temp,4], position[founder_temp,4], position[founder_temp,3], 1+cumsum(c(0,position[founder_temp,4]))[-(length(founder_temp)+1)], 1+cumsum(c(0,position[founder_temp,4]))[-(length(founder_temp)+1)] - sum(sex.s==1),
                           earliest_time[founder_temp], 0)


      cohort_info[(cohort_info[,7]>=1),c(3,6)] <- 0
      cohort_info[(cohort_info[,7]<1),c(4,7)] <- 0

      founder_pop <- founder_pop[founder_temp2]

      population$info$cohorts <- cohort_info

      colnames(population$info$cohorts) <- c("name","generation", "male individuals", "female individuals", "class", "position first male", "position first female",
                                             "time point", "creating.type")


      if(n_traits>0){
        population <- creating.trait(population, n.additive = as.numeric(trait_matrix[,6]),
                                     shuffle.cor = cor_gen, new.phenotype.correlation = cor_pheno,
                                     shuffle.traits=1:n_traits,
                                     trait.name = trait_matrix[,1])
        # Correct Scaling
        snp.before <- cumsum(c(0,population$info$snp))


        ## REASON?!
        if(TRUE){
          for(index in 1:n_traits){
            if(length(population$info$real.bv.add[[index]])>0){
              t <- population$info$real.bv.add[[index]]
              take <- sort(t[,1]+ snp.before[t[,2]], index.return=TRUE)
              t <- t[take$ix,,drop=FALSE]
              take <- sort(t[,1]+ t[,2] * 10^10)
              keep <- c(0,which(diff(take)!=0), length(take))
              if(length(keep) <= nrow(t)){
                for(index2 in 2:(length(keep))){
                  t[keep[index2],3:5] <- colSums(t[(keep[index2-1]+1):keep[index2],3:5, drop=FALSE])
                }
                population$info$real.bv.add[[index]] <- t[keep,]
              }
            }
          }
        }


        ## Variance Standardization
        for(index in 1:n_traits){
          new_var <- as.numeric(trait_matrix[index,4])^2

          if(population$info$bv.calculated==FALSE){
            population <- breeding.diploid(population, verbose=verbose)
          }
          active_sub <- subpopulation_info[1,1]
          standard_cohort <- population$info$cohorts[which(founder_pop==active_sub),1]
          var_test <- stats::var(get.bv(population, cohorts= standard_cohort)[index,])
          if(length(population$info$real.bv.add[[index]])>0){
            population$info$real.bv.add[[index]][,3:5] <- population$info$real.bv.add[[index]][,3:5] * sqrt(  new_var / var_test)
          } else{
            if(verbose) cat("You entered a trait without quantitative loci. Is this intentional?\n")
          }

        }
        population$info$bv.calculated <- FALSE

        ## Mean Standardization
        for(index in 1:n_traits){

          if(population$info$bv.calculated==FALSE){
            population <- breeding.diploid(population, verbose=verbose)
          }
          active_sub <- subpopulation_info[1,1]
          standard_cohort <- population$info$cohorts[which(founder_pop==active_sub),1]
          mean_test <- mean(get.bv(population, cohorts= standard_cohort)[index,])


          population$info$base.bv[index] <- traitmean[index] + population$info$base.bv[index] - mean_test

        }
        population$info$bv.calculated <- FALSE

        ## Add Major QTL
        for(index in 1:n_traits){
          if(length(major_table)>=index && length(major_table[[index]])>0){
            to_enter <- major_table[[index]][,1:5,drop=FALSE]
          } else{
            to_enter <- rbind(NULL)
          }
          if(length(to_enter)>0 || length(population$info$real.bv.add[[index]])>0){
            population$info$real.bv.add[[index]] <- rbind(to_enter, population$info$real.bv.add[[index]])

            if(population$info$real.bv.length[1]<index && length(to_enter)>0){
              population$info$real.bv.length[1] <- index
            }
          }

        }

        ## Subpopulation differences
        if(length(subpopulations)>1){
          for(subpop in 1:length(subpopulations)){
            if(population$info$bv.calculated==FALSE){
              population <- breeding.diploid(population, verbose=verbose)
            }
            if(length(subpopulations[[subpop]])>7){
              for(indexmod in 8:length(subpopulations[[subpop]])){
                active_trait <- as.numeric(substr(names(subpopulations[[subpop]])[[indexmod]], start=2, stop=nchar(names(subpopulations[[subpop]])[[indexmod]])))
                valid_markers0 <- valid_markers1 <- which(p_i[[subpop]]>0 & p_i[[subpop]]<1)
                for(reduction in (1:length(subpopulations))[-subpop]){
                  valid_markers0 <- intersect(valid_markers0, which(p_i[[reduction]]==0))
                }
                for(reduction in (1:length(subpopulations))[-subpop]){
                  valid_markers1 <- intersect(valid_markers1, which(p_i[[reduction]]==1))
                }
                valid_markers <- c(valid_markers0, valid_markers1)
                subpop_name <- subpopulation_info[subpop,1]
                sub_cohort <- population$info$cohorts[which(founder_pop==subpop_name),1]
                mean_ref <- mean(get.bv(population, cohorts= standard_cohort )[active_trait,])
                mean_sub <- mean(get.bv(population, cohorts= sub_cohort )[active_trait,])

                current_diff <- mean_sub - mean_ref
                target_diff <- as.numeric(subpopulations[[subpop]][[indexmod]])

                if(subpop==1){
                  change <- target_diff
                } else{
                  change <- target_diff - current_diff
                }

                diff_freq <- c(p_i[[subpop]][valid_markers0], - p_i[[subpop]][valid_markers1])


                effect_size <- change / sum(abs(diff_freq)) / 2
                direction <- diff_freq > 0

                snp_index <- chromo_index <- numeric(length(valid_markers))

                for(index2 in 1:length(valid_markers)){
                  chromo_index[index2] <- max(which(c(0,population$info$cumsnp)<=valid_markers[index2]))
                  snp_index[index2] <- valid_markers[index2] - c(0,population$info$cumsnp)[chromo_index[index2]]
                }
                add.effects <- cbind(snp_index, chromo_index, 2* effect_size, effect_size,   0)

                add.effects[direction==TRUE,3:5] <- cbind(rep(0, sum(direction)), rep(effect_size, sum(direction)), rep(2*effect_size, sum(direction)))


                population$info$real.bv.add[[active_trait]] <- rbind(population$info$real.bv.add[[active_trait]], add.effects)
                population$info$bv.calculated <- FALSE
                if(subpop==1){
                  population <- breeding.diploid(population, verbose=verbose)
                  population$info$bv.calculated <- FALSE
                }
              }
            }
          }
        }
      }
      population$breeding[[1]][[5]] <- mig_m
      population$breeding[[1]][[6]] <- mig_f

    }

    ################## Last preps ########################
    {
      # Add edges info to nodes
      for(index in 1:length(edges)){
        to_node <- which(edges[[index]]$to==ids)
        nodes[[to_node]]$'Breeding Type' <- edges[[index]]$'Breeding Type'
        if(nodes[[to_node]]$'Breeding Type'=="Selection" || nodes[[to_node]]$'Breeding Type'=="Split" || nodes[[to_node]]$'Breeding Type'=="Aging"  ){
          nodes[[to_node]]$'Selection Type' <- edges[[index]]$'Selection Type'
          #      nodes[[to_node]]$proportion <- edges[[index]]$proportion # not needed?
          nodes[[to_node]]$origin <- edges[[index]]$from
          nodes[[to_node]]$'Relationship Matrix' <- edges[[index]]$'Relationship Matrix'
          nodes[[to_node]]$'BVE Method' <- edges[[index]]$'BVE Method'
          nodes[[to_node]]$'Use Offspring for BVE' <- edges[[index]]$'Use Offspring for BVE'
          nodes[[to_node]]$edge.nr <- c(nodes[[to_node]]$edge.nr,index)
          nodes[[to_node]]$'Time Needed' <- c(nodes[[to_node]]$'Time Needed',edges[[index]]$'Time Needed')
          nodes[[to_node]]$'Selection Index' <- edges[[index]]$'Selection Index'
          nodes[[to_node]]$'Cohorts used in BVE' <- edges[[index]]$'Cohorts used in BVE'
          nodes[[to_node]]$'Selection Type Mean' <- edges[[index]]$'Selection Type Mean'
          if(nodes[[to_node]]$'Cohorts used in BVE'=="Manual select"){
            edges[[index]]$`Manuel selected cohorts`
            repeat_iter_test <- strsplit(nodes[[to_node]]$id, "_")[[1]]
            repeat_iter <- suppressWarnings(as.numeric(repeat_iter_test[length(repeat_iter_test)]))
            if(is.na(repeat_iter)){
              repeat_iter <- 0
            }
            id_bve <- NULL
            for(index3 in 1:length(edges[[index]]$`Manuel selected cohorts`)){
              cohort_split <- strsplit(edges[[index]]$`Manuel selected cohorts`[[index3]], ":")[[1]]
              if(length(cohort_split)==1){
                cohort_iter <- -(0:20) + repeat_iter
                cohort_iter <- cohort_iter[cohort_iter>=0]
              } else{
                if(!is.na(cohort_split[length(cohort_split)]) && cohort_split[length(cohort_split)]=="X"){
                  cohort_iter <- (- repeat_iter)
                } else{
                  cohort_iter <- as.numeric(cohort_split[length(cohort_split)])
                  if(is.na(cohort_iter)){
                    cohort_iter <- 0
                  }
                }
                cohort_iter <- repeat_iter + cohort_iter
              }

              for(index4 in cohort_iter){
                if(index4< (-1)){


                } else if(index4 == (-1)){
                  take_repeat <- which(repeat_links[,1]==cohort_split[1])
                  if(length(take_repeat)==1){
                    id_bve <- c(id_bve, repeat_links[take_repeat,2])
                  }

                } else if(index4==0){
                  id_bve <- c(id_bve, cohort_split[1])
                } else{
                  id_bve <- c(id_bve, paste0(cohort_split[1],"_", cohort_iter))
                }
              }

            }

            nodes[[to_node]]$'Manuel selected cohorts' <- id_bve
          }


          nodes[[to_node]]$'Depth of Pedigree' <- edges[[index]]$'Depth of Pedigree'
        }
        if(nodes[[to_node]]$'Breeding Type'=="Reproduction" || nodes[[to_node]]$'Breeding Type'=="Selfing" || nodes[[to_node]]$'Breeding Type'=="DH-Production" || nodes[[to_node]]$'Breeding Type'=="Cloning"){
          nodes[[to_node]]$origin <- c(nodes[[to_node]]$origin,edges[[index]]$from)
          nodes[[to_node]]$edge.nr <- c(nodes[[to_node]]$edge.nr,index)
          nodes[[to_node]]$'Time Needed' <- c(nodes[[to_node]]$'Time Needed',edges[[index]]$'Time Needed')
          if(length(edges[[index]]$OGC)>0 && edges[[index]]$OGC=="Yes"){
            nodes[[to_node]]$OGC <- TRUE
            if(length(edges[[index]]$'ogc_cAc')>0 || length(nodes[[to_node]]$ogc_cAc)==0){
              nodes[[to_node]]$ogc_cAc <- as.numeric(edges[[index]]$'ogc_cAc')
            } else{
              nodes[[to_node]]$ogc_cAc <- NA
            }

          } else if(length(nodes[[to_node]]$OGC)==0){
            nodes[[to_node]]$OGC <- FALSE
          }
          if(length(nodes[[to_node]]$selection_ratio)==0){
            nodes[[to_node]]$selection_ratio <- c(1,1)
          }
          if(length(nodes[[to_node]]$selection_ratio_index)==0){
            nodes[[to_node]]$selection_ratio_index <- c(selection_index_name[1], selection_index_name[1])
          }
          if(length(edges[[index]]$'selection_ratio')>0){
            sex <- as.numeric(nodes[[which(ids==edges[[index]]$from)]]$Sex=="Female") + 1
            if(length(edges[[index]]$'selection_ratio')==1){
              nodes[[to_node]]$selection_ratio[sex] <- as.numeric(edges[[index]]$'selection_ratio')
              if(is.na(nodes[[to_node]]$selection_ratio[sex])){
                nodes[[to_node]]$selection_ratio[sex] <- 1
              }
            }

          }
          if(length(edges[[index]]$'selection_ratio_index')>0){
            sex <- as.numeric(nodes[[which(ids==edges[[index]]$from)]]$Sex=="Female") + 1
            nodes[[to_node]]$selection_ratio_index[sex] <- (edges[[index]]$'selection_ratio_index'$Name)
          }

        }

        if(nodes[[to_node]]$'Breeding Type'=="Combine"){
          nodes[[to_node]]$origin <- c(nodes[[to_node]]$origin,edges[[index]]$from)
          nodes[[to_node]]$edge.nr <- c(nodes[[to_node]]$edge.nr,index)
          nodes[[to_node]]$'Time Needed' <- c(nodes[[to_node]]$'Time Needed',edges[[index]]$'Time Needed')
        }

        if(length(nodes[[to_node]]$max_offspring)!=2){
          nodes[[to_node]]$max_offspring <- c(Inf, Inf)
        }

        if(length(edges[[index]]$'Max Offspring')>0){
          from_node <- which(ids==edges[[index]]$from)
          switch <- as.numeric(nodes[[from_node]]$Sex=="Female") +1
          nodes[[to_node]]$max_offspring[switch] <- as.numeric(edges[[index]]$'Max Offspring')
        }

        if(length(edges[[index]]$repeat_mating)>0){
          nodes[[to_node]]$repeat_mating <- as.numeric(edges[[index]]$repeat_mating)
        } else{
          if(length(nodes[[to_node]]$repeat_mating)==0){
            nodes[[to_node]]$repeat_mating <- 1
          }
        }

      }

      phenotype_groups <- numeric(length(nodes))
      for(index in 1:length(edges)){
        phenotype_groups[which(ids==edges[[index]]$from)] <- 1
      }

      priority_breeding <- ids[(1-phenotype_groups)*1:length(ids)]


      n_tester <- n_tester_generated <- numeric(length(nodes))
      for(index in 1:length(edges)){
        if(length(intersect(priority_breeding, edges[[index]]$to))){
          n_tester[which(ids==edges[[index]]$from)] <- n_tester[which(ids==edges[[index]]$from)] +1
        }
      }

      for(index in 1:length(nodes)){
        exp_genotyped <- as.numeric(nodes[[index]]$`Proportion of genotyped individuals`)
        real_genotyped <- 0
        n_animals <- 0
        for(index2 in nodes[[index]]$origin){
          real_genotyped <- real_genotyped + as.numeric(nodes[[which(index2==ids)]]$`Proportion of genotyped individuals`) * as.numeric(nodes[[which(index2==ids)]]$`Number of Individuals`)
          n_animals <- n_animals + nodes[[which(index2==ids)]]$`Number of Individuals`
        }
        if(n_animals>0){
          nodes[[index]]$'Proportion of added genotypes' <- max(0, exp_genotyped - real_genotyped/n_animals)
        } else{
          nodes[[index]]$'Proportion of added genotypes' <- exp_genotyped
        }


      }


    }

    # Check for Split nodes
    {
      to_split <- NULL
      split_info <- list()
      split_part <- list()
      for(index in 1:length(edges)){
        if(edges[[index]]$'Breeding Type'=="Split"){
          to_split <- unique(c(to_split, edges[[index]]$from))
          split_nr <- which(to_split==edges[[index]]$from)
          nodes_nr <- which(edges[[index]]$from==ids)
          split_info[[split_nr]] <- 1:nodes[[nodes_nr]]$'Number of Individuals'
          if(length(split_part)>= split_nr){
            split_part[[split_nr]] <- c(split_part[[split_nr]], edges[[index]]$to)
          } else{
            split_part[[split_nr]] <- edges[[index]]$to
          }

        }
      }

    }
    ############## Founder Phenotypes ########################
    {
      if(n_traits>0){
        for(index in 1:nrow(population$info$cohorts)){
          population <- breeding.diploid(population, heritability = heritability,
                                         new.bv.observation.cohorts =  population$info$cohorts[index,1],
                                         sigma.e.database = cbind(1,(1:2)[population$info$size[1,]>0]),
                                         n.observation = pheno_index[which(pheno_index_name==nodes[[which(ids==population$info$cohorts[index,1])]]$'Phenotyping Class'),],
                                         verbose=verbose)

        }
      }
    }


    ### derive order of generation
    {
      {
        simulated <- founder
        left <- (1:groups)[-simulated]
        generation <- 1
        time.point.list <- numeric(length(ids))
        for(index in founder){
          time.point.list[index] <- nodes[[index]]$earliest_time
        }

        while(length(left)>0){
          generation <- generation + 1
          possible <- ids[left]
          stock <- ids[-left]
          for(index in 1:length(edges)){
            there <- which(edges[[index]]$to==possible)
            if(length(there)>0){


              if(sum(edges[[index]]$from==stock)==0){
                possible <- possible[-there]
                next
              }

              to_node <- which(ids==edges[[index]]$to)

              if(length(nodes[[to_node]]$'Manuel selected cohorts')>0){
                manu <- setdiff(nodes[[to_node]]$'Manuel selected cohorts', edges[[index]]$from)
                if(length(intersect(manu, stock))<length(manu)){
                  possible <- possible[-there]
                }
              }

            }
          }

          if(length(intersect(possible, priority_breeding))>0){
            possible <- intersect(possible, priority_breeding)
          } else{
            stock <- ids[-unique(c(left, (n_tester>n_tester_generated)*(1:length(n_tester))))]
            for(index in 1:length(edges)){
              there <- which(edges[[index]]$to==possible)
              if(length(there)>0){
                if(sum(edges[[index]]$from==stock)==0){
                  possible <- possible
                }
              }
            }
          }
          # Remove group for which not all testers are generated

          if(length(possible)==0){
            stop("invalite breeding program")
          }
          for(group in possible){
            groupnr <- which(ids==group)
            simulated <- c(simulated, groupnr)

            time.point <- 0
            origins <- nodes[[which(ids==group)]]$origin
            time_needed <- as.numeric(nodes[[which(ids==group)]]$'Time Needed')
            for(temp1 in 1:length(origins)){
              time.point <- max(time.point.list[simulated][which(origins[temp1]==ids[simulated])] + time_needed[temp1],time.point)
            }
            for(temp1 in nodes[[groupnr]]$'Manuel selected cohorts'){
              groupnr2 <- which(ids==temp1)
              time.point <- max(time.point.list[groupnr2],time.point)

            }

            time.point <- max(nodes[[which(ids==group)]]$earliest_time, time.point)

            time.point.list[groupnr] <- time.point
          }

          left <- (1:groups)[-simulated]
        }

        generation_times <- sort(unique(time.point.list))
        generation_group <- list()
        generation_bv_size <- list()
        for(index in 1:length(generation_times)){
          nrs <- setdiff(which(time.point.list==generation_times[[index]]), founder)
          btype <- numeric(length(nrs))
          if(length(nrs)>0){
            for(index2 in 1:length(nrs)){
              btype[index2] <- nodes[[nrs[index2]]]$'Breeding Type'
            }
          }

          prio <- which(btype=="Reproduction"| btype=="Selfing" | btype=="DH-Production" | btype=="Cloning" | btype=="Combine")
          if(length(prio)>0){
            nrs <- c(nrs[prio], nrs[-prio])
            btype <- c(btype[prio], btype[-prio])
          }
          if(sum(btype=="Split")>1){

            activ_type <- which(btype=="Split")
            activ_node <- nrs[activ_type]
            ns <- length(activ_node)
            activ_prior <- activ_time <- numeric(ns)
            for(index2 in 1:ns){
              activ_prior[index2] <- nodes[[activ_node[index2]]]$origin
              activ_time[index2] <- nodes[[activ_node[index2]]]$`Time Needed`
            }
            avail_prior <- unique(activ_prior)
            for(index2 in 1:length(avail_prior)){
              activ <- which(activ_prior==avail_prior[index2])
              new_order <- activ[sort(activ_time[activ], index.return=TRUE)$ix]
              nrs[activ_type][activ] <- nrs[activ_type][new_order]
              btype[activ_type][activ] <- btype[activ_type][new_order]
            }
          }

          generation_group[[index]] <- ids[nrs]
          changes <- 1
          while(changes>0){
            changes <- 0
            if(length(nrs)>0){
              for(index2 in 1:length(nrs)){
                if(length(intersect(c(nodes[[nrs[index2]]]$origin, nodes[[nrs[index2]]]$'Manuel selected cohorts'), generation_group[[index]][index2:length(nrs)]))>0){
                  back <- 0
                  for(i_origin in c(nodes[[nrs[index2]]]$origin, nodes[[nrs[index2]]]$'Manuel selected cohorts')){
                    back1 <- which(generation_group[[index]]==i_origin)
                    back <- max(back, back1[length(back1)])
                  }
                  store_temp <- nrs[index2]
                  nrs[index2:(back-1)] <- nrs[(index2+1):back]
                  nrs[back] <- store_temp
                  generation_group[[index]] <- ids[nrs]
                  changes <- 1
                }
              }
            }
          }

          btype <- numeric(length(nrs))
          if(length(nrs)>0){
            for(index2 in 1:length(nrs)){
              btype[index2] <- nodes[[nrs[index2]]]$'Breeding Type'
            }
          }

          database_add <- c(0,0)
          while(length(btype)>0 && (btype[1]=="Reproduction"| btype[1]=="Selfing" | btype[1]=="DH-Production" | btype[1]=="Cloning" | btype[1]=="Combine")){
            sex <- as.numeric(nodes[[nrs[1]]]$'Sex'=="Female") + 1
            database_add <- database_add + as.numeric(nodes[[nrs[1]]]$`Number of Individuals`) * c(sex==1, sex==2)
            btype <- btype[-1]
            nrs <- nrs[-1]
          }
          generation_bv_size[[index]] <- database_add

        }


      }
      {
        for(index in length(generation_group):1){
          if(length(generation_group[[index]])==0){
            generation_group[[index]] <- NULL
            generation_bv_size[[index]] <- NULL

          }
        }
      }
      #### Check if same breeding value estimation is done multiple times

      {
        bv_cohort <- NULL
        bv_cohort_ori <- NULL

        bv_included <- list()
        for(index in 1:length(nodes)){
          bve.breeding.type <- nodes[[index]]$`Breeding Type`=="Selection" || nodes[[index]]$`Breeding Type`=="Aging" || nodes[[index]]$`Breeding Type`=="Split"
          if(bve.breeding.type && nodes[[index]]$`Selection Type`=="BVE" && length(nodes[[index]]$'Cohorts used in BVE')>0 && nodes[[index]]$'Cohorts used in BVE'=="Manual select"){
            bv_cohort <- c(bv_cohort, nodes[[index]]$id)
            bv_cohort_ori <- c(bv_cohort_ori, nodes[[index]]$origin)
            bv_included[[length(bv_included)+1]] <- sort(unique( c(nodes[[index]]$origin, nodes[[index]]$'Manuel selected cohorts')))
          }
        }


        if(length(bv_included)>0){
          bv_class <- unique(bv_included)
          bv_cohort_class <- numeric(length(bv_cohort))
          bv_class_member <- list()
          for(index in 1:length(bv_cohort)){
            for(index2 in 1:length(bv_class)){
              if(length(bv_class[[index2]])==length(bv_included[[index]]) && prod(bv_class[[index2]] == bv_included[[index]])){
                bv_cohort_class[index] <- index2
                if(length(bv_class_member)<index2){
                  bv_class_member[[index2]] <- bv_cohort_ori[index]
                } else{
                  bv_class_member[[index2]] <- c(bv_class_member[[index2]], bv_cohort_ori[index])
                }

              }
            }
          }
          class_executed <- rep(FALSE,length(bv_class))
        }
      }
    }

    ############## Actual simulations ########################
    {
      # Derive Founders that are alive
      alive_cohorts <- population$info$cohorts[,1]
      alive_numbers <- as.numeric(population$info$cohorts[,3])
      alive_numbers[alive_numbers==0] <- as.numeric(population$info$cohorts[alive_numbers==0,4])
      death_to <- matrix(0, ncol=nrow(culling_reason), nrow=length(alive_cohorts))

      for(generation in 1:(length(generation_group)+1) +1){

        if(verbose) cat(paste0("Start simulation of generation:", generation," (time point: ", generation_times[generation-1], ")\n"))

        if(generation != (length(generation_group)+2)){
          for(group in generation_group[[generation-1]]){
            groupnr <- which(ids==group)
            sex <- as.numeric(nodes[[groupnr]]$'Sex'=="Female") + 1

            breeding.size <- as.numeric(nodes[[groupnr]]$'Number of Individuals') * c(sex==1, sex==2)
            involved_cohorts <- nodes[[groupnr]]$origin
            cohort_data <- population$info$cohorts[involved_cohorts,,drop=FALSE]
            sex_cohorts <- (as.numeric(cohort_data[,3])==0) +1
            selection.size <- c(sum(as.numeric(cohort_data[,3])), sum(as.numeric(cohort_data[,4])))

            share.genotyped <- as.numeric(nodes[[groupnr]]$`Proportion of genotyped individuals`)
            cohorts.m <- involved_cohorts[sex_cohorts==1]
            cohorts.f <- involved_cohorts[sex_cohorts==2]

            involved_groups <- cbind(as.numeric(cohort_data[,2]), sex_cohorts)
            # Derive time.point
            time.point <- 0
            origins <- nodes[[which(ids==group)]]$origin
            time_needed <- as.numeric(nodes[[which(ids==group)]]$'Time Needed')
            for(temp1 in 1:length(origins)){
              time.point <- as.numeric(population$info$cohorts[population$info$cohorts[,1]==origins[temp1],8]) + time_needed[temp1]
            }
            time.point <- max(nodes[[which(ids==group)]]$earliest_time, time.point)

            bve.database <- NULL
            bve_exe <- TRUE
            bve_insert_cohorts = c(cohorts.m, cohorts.f)


            bve.breeding.type <- nodes[[groupnr]]$`Breeding Type`=="Selection" || nodes[[groupnr]]$`Breeding Type`=="Aging" || nodes[[groupnr]]$`Breeding Type`=="Split"
            if(length(nodes[[groupnr]]$'Cohorts used in BVE') || bve.breeding.type){
              if(length(nodes[[groupnr]]$'Cohorts used in BVE')==0){
                bve.database <- involved_groups[,1:2, drop=FALSE]
              } else if(nodes[[groupnr]]$'Cohorts used in BVE'=="Last 2 Generations"){
                bve.database <- get.database(population, gen=max(1,generation-2):(generation-1))
                if(generation_bv_size[[generation-1]][1]>0){
                  bve.database <- rbind(bve.database, c(generation,1,1,generation_bv_size[[generation-1]][1]))
                }
                if(generation_bv_size[[generation-1]][2]>0){
                  bve.database <- rbind(bve.database, c(generation,2,1,generation_bv_size[[generation-1]][2]))
                }

              } else if(nodes[[groupnr]]$'Cohorts used in BVE'=="Last 3 Generations"){
                bve.database <- get.database(population, gen=max(1,generation-3):(generation-1))
                if(generation_bv_size[[generation-1]][1]>0){
                  bve.database <- rbind(bve.database, c(generation,1,1,generation_bv_size[[generation-1]][1]))
                }
                if(generation_bv_size[[generation-1]][2]>0){
                  bve.database <- rbind(bve.database, c(generation,2,1,generation_bv_size[[generation-1]][2]))
                }

              } else if(nodes[[groupnr]]$'Cohorts used in BVE'=="Last Generation"){
                bve.database <- get.database(population, gen=max(1,generation-1):(generation-1))
                if(generation_bv_size[[generation-1]][1]>0){
                  bve.database <- rbind(bve.database, c(generation,1,1,generation_bv_size[[generation-1]][1]))
                }
                if(generation_bv_size[[generation-1]][2]>0){
                  bve.database <- rbind(bve.database, c(generation,2,1,generation_bv_size[[generation-1]][2]))
                }

              } else if(nodes[[groupnr]]$'Cohorts used in BVE'=="All"){
                bve.database <- get.database(population, gen=1:(generation-1))
              } else if(nodes[[groupnr]]$'Cohorts used in BVE'=="Manual select") {
                bve.database <- get.database(population, cohorts=unique(nodes[[groupnr]]$'Manuel selected cohorts'))
                bve.database <- bve.database[!is.na(bve.database[,1]),, drop=FALSE]
                bv_cohort_index <- which(bv_cohort==group)
                bv_cohort_class_index <- bv_cohort_class[bv_cohort_index]
                bve_insert_cohorts <- bv_class_member[[bv_cohort_class_index]]
                if(class_executed[bv_cohort_class_index]){
                  bve_exe <- FALSE
                } else{
                  class_executed[bv_cohort_class_index] <- TRUE
                }

              } else if(nodes[[groupnr]]$'Cohorts used in BVE'=="Only this cohort"){
                bve.database <- involved_groups[,1:2, drop=FALSE]
              }
            }

            if(bve.breeding.type){
              activemmreml <-activesommer <- activemultisommer <- activbglr <- activerrblup <-FALSE
              singlestep.active <- FALSE
              depth <- 0
              parent_average <- FALSE
              grandparent_average <- FALSE
              mean_between <- NULL
              phenotype.bv <- FALSE
              computeA <- "vanRaden"
              if(length(nodes[[groupnr]]$'Selection Type')==0){
                if(verbose) cat("No selection type selected in some edges. Assume selection type 'Random'")
                bve <- FALSE
                selection <- "random"
              }else if(nodes[[groupnr]]$'Selection Type'=="Parent_Mean"){
                bve <- FALSE
                parent_average <- TRUE
                selection <- "function"
                if(nodes[[groupnr]]$'Selection Type Mean'=="BVE"){
                  mean_between <- "bve"
                } else if(nodes[[groupnr]]$'Selection Type Mean'=="Phenotype"){
                  mean_between <- "pheno"
                } else if(nodes[[groupnr]]$'Selection Type Mean'=="BV"){
                  mean_between <- "bv"
                } else {
                  mean_between <- "bvepheno"
                }
              } else if(nodes[[groupnr]]$'Selection Type'=="Grandparent_Mean"){
                bve <- FALSE
                grandparent_average <- TRUE
                selection <- "function"
                if(nodes[[groupnr]]$'Selection Type Mean'=="BVE"){
                  mean_between <- "bve"
                } else if(nodes[[groupnr]]$'Selection Type Mean'=="Phenotype"){
                  mean_between <- "pheno"
                } else if(nodes[[groupnr]]$'Selection Type Mean'=="BV"){
                  mean_between <- "bv"
                } else {
                  mean_between <- "bvepheno"
                }
              } else if(nodes[[groupnr]]$'Selection Type'=="Phenotypic"){
                bve <- FALSE
                selection <- "function"
                phenotype.bv <- TRUE
              } else if(nodes[[groupnr]]$'Selection Type'=="BVE"){
                bve <- TRUE
                selection <- "function"
                if(nodes[[groupnr]]$'Relationship Matrix'=="Pedigree"){
                  computeA <- "kinship"
                  depth <- as.numeric(nodes[[groupnr]]$'Depth of Pedigree')
                } else if(nodes[[groupnr]]$'Relationship Matrix'=="Single Step"){
                  depth <- as.numeric(nodes[[groupnr]]$'Depth of Pedigree')
                  singlestep.active <- TRUE
                }
                if(nodes[[groupnr]]$'BVE Method'=="REML-GBLUP" || nodes[[groupnr]]$'BVE Method'=="REML-GBLUP (EMMREML)"){
                  activemmreml <- TRUE
                } else if(nodes[[groupnr]]$'BVE Method'=="REML-GBLUP (sommer)") {
                  activesommer <- TRUE
                } else if(nodes[[groupnr]]$'BVE Method'=="Multi-trait REML-GBLUP (sommer)") {
                  activemultisommer <- TRUE
                } else if(nodes[[groupnr]]$'BVE Method'=="BayesA" || nodes[[groupnr]]$'BVE Method'=="REML-GBLUP (rrBLUP)"){
                  activerrblup <- TRUE
                } else if(nodes[[groupnr]]$'BVE Method'=="RKHS") {
                  activbglr <- TRUE
                }
              } else if(nodes[[groupnr]]$'Selection Type'=="Random"){
                bve <- FALSE
                selection <- "random"
              } else{
                if(verbose) cat("No selection type selected in some edges. Assume selection type 'Random'")
                bve <- FALSE
                selection <- "random"

              }
              if(length(involved_cohorts)>1){
                stop("Only one cohort to select from allowed in selection - check for error")
              } else{
                add.observation <- pheno_index[which(pheno_index_name==nodes[[groupnr]]$'Phenotyping Class'),] -
                  pheno_index[which(pheno_index_name==nodes[[which(ids==involved_cohorts[1])]]$'Phenotyping Class'),]
                add.observation[add.observation<0] <- 0
              }

              reduced.selection.panel.m <- NULL
              reduced.selection.panel.f <- NULL
              if(nodes[[groupnr]]$'Breeding Type'=="Selection"){
                creating.type <- 1
              } else if(nodes[[groupnr]]$'Breeding Type'=="Aging"){
                creating.type <- 8
              } else if(nodes[[groupnr]]$'Breeding Type'=="Split"){
                creating.type <- 9
                split_nr <- which(nodes[[groupnr]]$origin==to_split)
                reduced.selection.panel.m <- split_info[[split_nr]]
                reduced.selection.panel.f <- split_info[[split_nr]]
              }


              if(nodes[[groupnr]]$'Use Offspring for BVE'=="Yes"){
                offspring.bve.parents.database <- get.database(population, cohorts=c(cohorts.m, cohorts.f))
              } else{
                offspring.bve.parents.database <- NULL
              }

              population <- breeding.diploid(population, breeding.size=breeding.size,
                                             bve=(bve&bve_exe), computation.A = computeA,
                                             offspring.bve.parents.database=offspring.bve.parents.database,
                                             BGLR.bve = activbglr,
                                             emmreml.bve = activemmreml,
                                             rrblup.bve = activerrblup,
                                             sommer.bve = activesommer,
                                             sommer.multi.bve = activemultisommer,
                                             selection.size= breeding.size,
                                             copy.individual = TRUE,
                                             added.genotyped = nodes[[groupnr]]$`Proportion of added genotypes`,
                                             max.offspring = c(1,1),
                                             heritability = heritability,
                                             sigma.e.database = cbind(1,(1:2)[population$info$size[1,]>0]),
                                             new.bv.child="addobs",
                                             selection.m = selection,
                                             selection.f = selection,
                                             phenotype.bv = phenotype.bv,
                                             add.gen = generation,
                                             bve.database = bve.database,
                                             selfing.mating=TRUE,
                                             selfing.sex=(sex-1),
                                             selection.m.cohorts = cohorts.m,
                                             selection.f.cohorts = cohorts.f,
                                             new.class = new_mig[sex],
                                             selection.m.miesenberger = selection_index_miesenberger[which(selection_index_name==nodes[[groupnr]]$'Selection Index')],
                                             multiple.bve.scale.m = selection_index_miesenberger_wscaling[which(selection_index_name==nodes[[groupnr]]$'Selection Index')],
                                             n.observation = add.observation,
                                             remove.effect.position = remove.effect.position,
                                             name.cohort = nodes[[groupnr]]$id,
                                             time.point = time.point,
                                             creating.type = creating.type,
                                             depth.pedigree = depth,
                                             store.breeding.totals = TRUE,
                                             reduced.selection.panel.m = reduced.selection.panel.m,
                                             reduced.selection.panel.f = reduced.selection.panel.f,
                                             bve.cohorts = bve_insert_cohorts,
                                             bve.insert.cohorts = bve_insert_cohorts,
                                             display.progress=progress.bars,
                                             singlestep.active=singlestep.active,
                                             share.genotyped = share.genotyped,
                                             multiple.bve.weights.m = selection_index[which(selection_index_name==nodes[[groupnr]]$'Selection Index'),],
                                             verbose=verbose,
                                             bve.parent.mean=parent_average,
                                             bve.grandparent.mean=grandparent_average,
                                             bve.mean.between=mean_between
              )

              if(nodes[[groupnr]]$'Breeding Type'=="Split"){
                split_info[[split_nr]] <- sort(setdiff(split_info[[split_nr]], split_info[[split_nr]][population$info$breeding.totals[[length(population$info$breeding.totals)]][[7]][[sex]]]))
              }

            } else if(nodes[[groupnr]]$'Breeding Type'=="Reproduction"){

              generation.sex <- as.numeric(selection.size[2]>0)- 0.5 * as.numeric((selection.size[1]>0)*(selection.size[2]>0))
              if(generation.sex==0 || generation.sex==1){
                same.sex.activ <- TRUE
                same.sex.sex <- generation.sex
              } else{
                same.sex.activ <- FALSE
                same.sex.sex <- generation.sex
              }
              population <- breeding.diploid(population, breeding.size=breeding.size,
                                             selection.size= selection.size,
                                             heritability = heritability,
                                             sigma.e.database = cbind(1,(1:2)[population$info$size[1,]>0]),
                                             new.bv.child="obs",
                                             selection.m = if(sum(nodes[[groupnr]]$selection_ratio==1)==2){"random"} else{"function"},
                                             add.gen = generation,
                                             bve.database = bve.database,
                                             selection.m.cohorts = cohorts.m,
                                             selection.f.cohorts = cohorts.f,
                                             n.observation = pheno_index[which(pheno_index_name==nodes[[groupnr]]$'Phenotyping Class'),],
                                             new.class = new_mig[sex],
                                             same.sex.activ = same.sex.activ,
                                             same.sex.sex = same.sex.sex,
                                             same.sex.selfing = FALSE,
                                             name.cohort = nodes[[groupnr]]$id,
                                             time.point = time.point,
                                             creating.type = 2,
                                             store.breeding.totals = TRUE,
                                             display.progress=progress.bars,
                                             share.genotyped = share.genotyped,
                                             added.genotyped = nodes[[groupnr]]$`Proportion of added genotypes`,
                                             ogc = nodes[[groupnr]]$OGC,
                                             ogc.cAc = if(length(nodes[[groupnr]]$ogc_cAc)>0){nodes[[groupnr]]$ogc_cAc} else{NA},
                                             repeat.mating = nodes[[groupnr]]$repeat_mating,
                                             max.offspring = nodes[[groupnr]]$max_offspring,
                                             best.selection.ratio.m = nodes[[groupnr]]$selection_ratio[1],
                                             best.selection.ratio.f = nodes[[groupnr]]$selection_ratio[2],
                                             best.selection.criteria.m = "bv",
                                             best.selection.criteria.f = "bv",
                                             multiple.bve.weights.m = selection_index[max(1,which(selection_index_name==nodes[[groupnr]]$selection_ratio_index[1])),],
                                             multiple.bve.weights.f = selection_index[max(1,which(selection_index_name==nodes[[groupnr]]$selection_ratio_index[2])),],
                                             multiple.bve.scale.m = selection_index_miesenberger_wscaling[max(1,which(selection_index_name==nodes[[groupnr]]$selection_ratio_index[1]))],
                                             multiple.bve.scale.f = selection_index_miesenberger_wscaling[max(1,which(selection_index_name==nodes[[groupnr]]$selection_ratio_index[2]))],
                                             verbose=verbose
              )
            } else if(nodes[[groupnr]]$'Breeding Type'=="Selfing"){

              selfing.sex <- as.numeric(selection.size[2]>0)- 0.5 * as.numeric((selection.size[1]>0)*(selection.size[2]>0))
              population <- breeding.diploid(population, breeding.size=breeding.size,
                                             selection.size= selection.size,
                                             selfing.mating = TRUE,
                                             selfing.sex =  selfing.sex,
                                             heritability = heritability,
                                             sigma.e.database = cbind(1,(1:2)[population$info$size[1,]>0]),
                                             new.bv.child="obs",
                                             selection.m = "random",
                                             add.gen = generation,
                                             bve.database = bve.database,
                                             selection.m.cohorts = cohorts.m,
                                             selection.f.cohorts = cohorts.f,
                                             n.observation = pheno_index[which(pheno_index_name==nodes[[groupnr]]$'Phenotyping Class'),],
                                             new.class = new_mig[sex],
                                             name.cohort = nodes[[groupnr]]$id,
                                             time.point = time.point,
                                             creating.type = 4,
                                             store.breeding.totals = TRUE,
                                             display.progress=progress.bars,
                                             share.genotyped = share.genotyped,
                                             added.genotyped = nodes[[groupnr]]$`Proportion of added genotypes`,
                                             repeat.mating = nodes[[groupnr]]$repeat_mating,
                                             max.offspring = nodes[[groupnr]]$max_offspring,
                                             verbose=verbose)
            } else if(nodes[[groupnr]]$'Breeding Type'=="DH-Production"){

              dh.sex <- as.numeric(selection.size[2]>0)- 0.5 * as.numeric((selection.size[1]>0)*(selection.size[2]>0))
              population <- breeding.diploid(population, breeding.size=breeding.size,
                                             selection.size= selection.size,
                                             dh.mating = TRUE,
                                             dh.sex =  dh.sex,
                                             selfing.mating = TRUE,
                                             selfing.sex = dh.sex,
                                             heritability = heritability,
                                             sigma.e.database = cbind(1,(1:2)[population$info$size[1,]>0]),
                                             new.bv.child="obs",
                                             selection.m = "random",
                                             add.gen = generation,
                                             bve.database = bve.database,
                                             selection.m.cohorts = cohorts.m,
                                             selection.f.cohorts = cohorts.f,
                                             n.observation = pheno_index[which(pheno_index_name==nodes[[groupnr]]$'Phenotyping Class'),],
                                             new.class = new_mig[sex],
                                             name.cohort = nodes[[groupnr]]$id,
                                             time.point = time.point,
                                             creating.type = 5,
                                             store.breeding.totals = TRUE,
                                             display.progress=progress.bars,
                                             share.genotyped = share.genotyped,
                                             added.genotyped = nodes[[groupnr]]$`Proportion of added genotypes`,
                                             repeat.mating = nodes[[groupnr]]$repeat_mating,
                                             max.offspring = nodes[[groupnr]]$max_offspring,
                                             verbose=verbose)
            } else if(nodes[[groupnr]]$'Breeding Type'=="Recombination"){

              population <- breeding.diploid(population, breeding.size=breeding.size,
                                             mutation.rate = nodes[[groupnr]]$mutation,
                                             remutation.rate = nodes[[groupnr]]$remutation,
                                             recombination.rate = nodes[[groupnr]]$recom,
                                             selection.size= selection.size,
                                             heritability = heritability,
                                             sigma.e.database = cbind(1,(1:2)[population$info$size[1,]>0]),
                                             new.bv.child="obs",
                                             selection.m = "random",
                                             add.gen = generation,
                                             bve.database = bve.database,
                                             selection.m.cohorts = cohorts.m,
                                             selection.f.cohorts = cohorts.f,
                                             n.observation = pheno_index[which(pheno_index_name==nodes[[groupnr]]$'Phenotyping Class'),],
                                             new.class = new_mig[sex],
                                             name.cohort = nodes[[groupnr]]$id,
                                             time.point = time.point,
                                             creating.type = 3,
                                             store.breeding.totals = TRUE,
                                             display.progress=progress.bars,
                                             share.genotyped = share.genotyped,
                                             added.genotyped = nodes[[groupnr]]$`Proportion of added genotypes`,
                                             repeat.mating = nodes[[groupnr]]$repeat_mating,
                                             max.offspring = nodes[[groupnr]]$max_offspring,
                                             verbose=verbose)
            } else if(nodes[[groupnr]]$'Breeding Type'=="Cloning"){

              selfing.sex <- as.numeric(selection.size[2]>0)- 0.5 * as.numeric((selection.size[1]>0)*(selection.size[2]>0))
              if(length(involved_cohorts)>1){
                stop("Only one cohort to select from allowed in selection - check for error")
              } else{
                add.observation <- pheno_index[which(pheno_index_name==nodes[[groupnr]]$'Phenotyping Class'),] -
                  pheno_index[which(pheno_index_name==nodes[[which(ids==involved_cohorts[1])]]$'Phenotyping Class'),]
                add.observation[add.observation<0] <- 0
              }

              population <- breeding.diploid(population, breeding.size=breeding.size,
                                             selection.size= selection.size,
                                             copy.individual = TRUE,
                                             added.genotyped = nodes[[groupnr]]$`Proportion of added genotypes`,
                                             selfing.mating = TRUE,
                                             selfing.sex =  selfing.sex,
                                             heritability = heritability,
                                             sigma.e.database = cbind(1,(1:2)[population$info$size[1,]>0]),
                                             new.bv.child="addobs",
                                             selection.m = "random",
                                             add.gen = generation,
                                             bve.database = bve.database,
                                             selection.m.cohorts = cohorts.m,
                                             selection.f.cohorts = cohorts.f,
                                             n.observation = pheno_index[which(pheno_index_name==nodes[[groupnr]]$'Phenotyping Class'),],
                                             new.class = new_mig[sex],
                                             name.cohort = nodes[[groupnr]]$id,
                                             time.point = time.point,
                                             creating.type = 6,
                                             store.breeding.totals = TRUE,
                                             display.progress=progress.bars,
                                             share.genotyped = share.genotyped,
                                             repeat.mating = nodes[[groupnr]]$repeat_mating,
                                             max.offspring = nodes[[groupnr]]$max_offspring,
                                             verbose=verbose)

            } else if(nodes[[groupnr]]$'Breeding Type'=="Combine"){
              selfing.sex <- (as.numeric(selection.size[2])>0)- 0.5 * as.numeric((selection.size[1]>0)*(selection.size[2]>0))
              if(FALSE){
                stop("Only one cohort to select from allowed in selection - check for error")
              } else{
                add.observation <- pheno_index[which(pheno_index_name==nodes[[groupnr]]$'Phenotyping Class'),] -
                  pheno_index[which(pheno_index_name==nodes[[which(ids==involved_cohorts[1])]]$'Phenotyping Class'),]
                add.observation[add.observation<0] <- 0
                if(verbose) cat(paste0("Newly added phenotypes for combine note derived based on ", involved_cohorts[1],"\n"))
              }
              population <- breeding.diploid(population, breeding.size=breeding.size,
                                             selection.size= breeding.size,
                                             copy.individual = TRUE,
                                             added.genotyped = nodes[[groupnr]]$`Proportion of added genotypes`,
                                             max.offspring = c(1,1),
                                             new.bv.child="addobs",
                                             selection.m = "random",
                                             selfing.mating=TRUE,
                                             selfing.sex=selfing.sex,
                                             add.gen = generation,
                                             selection.m.cohorts = cohorts.m,
                                             selection.f.cohorts = cohorts.f,
                                             n.observation = add.observation,
                                             new.class = new_mig[sex],
                                             name.cohort = nodes[[groupnr]]$id,
                                             time.point = time.point,
                                             creating.type = 7,
                                             store.breeding.totals = TRUE,
                                             display.progress=progress.bars,
                                             share.genotyped = share.genotyped,
                                             verbose=verbose)
            }
            position[groupnr,] <- c(generation, sex, new_mig[sex], sum(breeding.size))
            new_mig[sex] <- new_mig[sex] + 1


            if(phenotype_groups[groupnr]==0){
              tested <- which(duplicated(c(nodes[[groupnr]]$origin, ids))[-(1:length(nodes[[groupnr]]$origin))])
              n_tester_generated[tested] <- n_tester_generated[tested] + 1
            }

          }
        }

        if(generation != (length(generation_group)+2)){
          if(verbose) cat("Generated groups: \n")
          if(verbose) cat(generation_group[[generation-1]])
          if(verbose) cat("\n")
          alive_cohorts <- c(alive_cohorts, generation_group[[generation-1]])

          new_cohorts <- population$info$cohorts[(nrow(population$info$cohorts)-length(generation_group[[generation-1]])+1):nrow(population$info$cohorts),,drop=FALSE]
          new_numbers <- as.numeric(new_cohorts[,3])
          new_numbers[new_numbers==0] <- as.numeric(new_cohorts[new_numbers==0,4])
          alive_numbers <- c(alive_numbers, new_numbers)
        }
        #      death_to <- rbind(death_to, matrix(0, nrow=length(generation_group[[generation-1]]), ncol=nrow(culling_reason)))

        from <- if(generation==2){0} else{generation_times[generation-2]}
        if(generation != (length(generation_group)+2)){
          to <- generation_times[generation-1]
        } else{
          to <- Inf
        }

        if(verbose) cat(paste0("Start culling modul for time span ", from ," to ", to,".\n"))


        for(cohort_index in alive_cohorts[alive_numbers>0]){
          age_start <- from - as.numeric(population$info$cohorts[cohort_index,8])
          age_end <- to - as.numeric(population$info$cohorts[cohort_index,8])
          sex <- if(population$info$cohorts[cohort_index,4]=="0"){"Male"} else{"Female"}
          # Time - period, sex
          active_culling <- as.numeric(culling_reason[,2]) > age_start & as.numeric(culling_reason[,2]) <= age_end & (culling_reason[,3] == sex | culling_reason[,3]=="Both")

          active_cohort <- which(population$info$cohorts[,1]==cohort_index)
          for(culling_index in which(active_culling)){
            population <- breeding.diploid(population,
                                           culling.cohort = cohort_index,
                                           culling.time = as.numeric(culling_reason[culling_index,2]),
                                           culling.name = culling_reason[culling_index,1],
                                           culling.bv1 = as.numeric(culling_reason[culling_index,6]),
                                           culling.bv2 = as.numeric(culling_reason[culling_index,8]),
                                           culling.share1 = as.numeric(culling_reason[culling_index,5]),
                                           culling.share2 = as.numeric(culling_reason[culling_index,7]),
                                           culling.index = selection_index[which(selection_index_name==culling_reason[culling_index,4]),],
                                           verbose=verbose
            )
            if(length(population$info$culling.stats)>=active_cohort){
              new_death <- as.numeric(population$info$culling.stats[[active_cohort]][population$info$culling.stats[[active_cohort]][,1]==culling_reason[culling_index,1],2])

            } else{
              new_death <- 0

            }
            if(length(new_death)>0){
              alive_numbers[active_cohort] <- alive_numbers[active_cohort] - new_death
            }


          }
          if(alive_numbers[active_cohort]==0){
            if(verbose) cat(paste0("All individuals in cohort ", cohort_index, " are death.\n"))
            if(verbose) cat("Reason for death:\n")
            for(index in 1:nrow(population$info$culling.stats[[active_cohort]])){
              if(verbose) cat(population$info$culling.stats[[active_cohort]][index,])
              if(verbose) cat("\n")
            }
          }

        }
      }

    }


    ############## Attach json-infos ########################
    housing <- list(housing_index, housing_index_name)
    phenotyping <- list(pheno_index_costs, pheno_index_name, pheno_index)

    population$info$json <- list(nodes, edges, geninfo, traitinfo, major, housing, phenotyping, ids)
  }

  return(population)

}
```
