## Supplementary material for "MoBPS - Modular Breeding Program Simulator": MoBPS R-package: kinship.development.R

```
'#
  Authors
Torsten Pook,

Copyright (C) 2017 -- 2018  Torsten Pook

This program is free software; you can redistribute it and/or
modify it under the terms of the GNU General Public License
as published by the Free Software Foundation; either version 3
of the License, or (at your option) any later version.

This program is distributed in the hope that it will be useful,
but WITHOUT ANY WARRANTY; without even the implied warranty of
MERCHANTABILITY or FITNESS FOR A PARTICULAR PURPOSE.  See the
GNU General Public License for more details.

You should have received a copy of the GNU General Public License
along with this program; if not, write to the Free Software
Foundation, Inc., 59 Temple Place - Suite 330, Boston, MA  02111-1307, USA.
'#

#' Devolopment of genetic/breeding value
#'
#' Function to plot genetic/breeding values for multiple generation/cohorts
#' @param population population list
#' @param database Groups of individuals to consider for the export
#' @param gen Quick-insert for database (vector of all generations to export)
#' @param cohorts Quick-insert for database (vector of names of cohorts to export)
#' @param json If TRUE extract which cohorts to plot according to the json-file used in json.simulation
#' @param ibd.obs Number of Individual pairs to sample for IBD estimation
#' @param hbd.obs Number of Individuals to sample for HBD estimation
#' @param display.cohort.name Set TRUE to display the name of the cohort in the x-axis
#' @param display.time.point Set TRUE to use time point of generated to sort groups
#' @param equal.spacing Equal distance between groups (independent of time.point)
#' @param time_reorder Set TRUE to order cohorts according to the time point of generation
#' @param display.hbd Set to TRUE to also display HBD in plot
#' @export

kinship.development <- function(population, database=NULL, gen=NULL, cohorts=NULL,
                           json=FALSE, ibd.obs=50, hbd.obs=10,
                           display.cohort.name=FALSE,
                           display.time.point=FALSE,
                           equal.spacing = FALSE,
                           time_reorder=FALSE,
                           display.hbd=FALSE){

  if(json){
    ids <- to_plot <- numeric(length(population$info$json[[1]]))
    for(index in 1:length(ids)){
      ids[index] <- population$info$json[[1]][[index]]$id
      if(length(population$info$json[[1]][[index]]$'BV Plot')>0){
        to_plot[index] <- population$info$json[[1]][[index]]$'BV Plot'
      }
      cohorts <- ids[which(to_plot=="Yes")]
      if(length(cohorts)==0){
        cohorts <- population$info$cohorts[,1]
      }
    }
  }
  inbred <- NULL
  time.point <- list()
  if(length(gen)>0){
    for(index in gen){
      inbred <- rbind(inbred, kinship.emp.fast(population=population, gen=index, ibd.obs=ibd.obs, hbd.obs=hbd.obs))
      time.point[[length(time.point)+1]] <- get.time.point(population, gen=index)
    }
  }
  if(length(database)>0){
    for(index in 1:nrow(database)){
      inbred <- rbind(inbred, kinship.emp.fast(population=population, database=database[index,1:2, drop=FALSE], ibd.obs=ibd.obs, hbd.obs=hbd.obs))
      time.point[[length(time.point)+1]] <- get.time.point(population, database=database[index,,drop=FALSE])

    }
  }
  if(length(cohorts)>0){
    for(index in cohorts){
      inbred <- rbind(inbred, kinship.emp.fast(population=population, cohorts=index, ibd.obs=ibd.obs, hbd.obs=hbd.obs))
      time.point[[length(time.point)+1]] <- get.time.point(population, cohorts=index)
    }
  }
  time_plot <-  1:nrow(inbred)

  if(display.time.point){
    for(index in 1:nrow(inbred)){
      time_plot[index] <- mean(time.point[[index]])
      if(length(unique(time.point[[index]]))>1){
        print("More than one time point in a plotted element")
      }
    }
  }

  if(time_reorder){
    reorder <- sort(time_plot, index.return=TRUE)$ix
  } else{
    reorder <- 1:length(time_plot)
  }

  if(equal.spacing){
    xlabel <- time_plot
    time_plot <- 1:length(time_plot)
  }

  if(display.cohort.name){
    graphics::par(mar=c(8.1,4.1,2.1,0.1))
  } else{
    graphics::par(mar=c(4.1,4.1,2.1,0.1))
  }

  graphics::plot(time_plot[reorder], inbred[reorder,1], main=paste("Kinship development"),
                 xaxt=if(display.cohort.name || equal.spacing){'n'}else {NULL},
                 xlab=if(display.cohort.name){""}else{"time"}, ylim=c(0,max(inbred[reorder,c(1,1+display.hbd)])), col="red", lwd=2, type="l", ylab="")
  if(display.hbd){
    graphics::lines(time_plot[reorder], inbred[reorder,2], main=paste("Kinship development"), lwd=2, col="blue")
  }

  if(equal.spacing && !display.cohort.name){
    graphics::axis(1, at=time_plot, labels=xlabel)
  }

  if(display.cohort.name){
    display.names <- NULL
    if(length(gen)>0){
      display.names <- c(display.names, paste("Generation", gen))
    }
    if(length(database)>0){
      display.names <- c(display.names, paste0(c("M", "F")[database[,2]], "_", database[,1]))
    }
    if(length(cohorts)>0){
      display.names <- c(display.names, cohorts)
    }
    display.names <- display.names
    graphics::axis(1, at=time_plot, labels=display.names,las=2)
    row.names(inbred) <- display.names
  }

  if(display.hbd){
    graphics::legend("topleft",c("IBD", "HBD"), lty=c(1,1), col=c("red","blue"), lwd=c(2,2))
  } else{
    graphics::legend("topleft",c("IBD"), lty=c(1), col=c("red"), lwd=c(2,2))
  }

  return(inbred)
}
```
