## Supplementary material for "MoBPS - Modular Breeding Program Simulator": MoBPS R-package: kinship.emp.R

```
'#
  Authors
Torsten Pook,

Copyright (C) 2017 -- 2018  Torsten Pook

This program is free software; you can redistribute it and/or
modify it under the terms of the GNU General Public License
as published by the Free Software Foundation; either version 3
of the License, or (at your option) any later version.

This program is distributed in the hope that it will be useful,
but WITHOUT ANY WARRANTY; without even the implied warranty of
MERCHANTABILITY or FITNESS FOR A PARTICULAR PURPOSE.  See the
GNU General Public License for more details.

You should have received a copy of the GNU General Public License
along with this program; if not, write to the Free Software
Foundation, Inc., 59 Temple Place - Suite 330, Boston, MA  02111-1307, USA.
'#

#' Empirical kinship
#'
#' Function to compute empirical kinship for a set of individuals)
#' @param animals List of animals to compute kinship for
#' @param population Population list
#' @param database Groups of individuals to consider for the export
#' @param gen Quick-insert for database (vector of all generations to export)
#' @param cohorts Quick-insert for database (vector of names of cohorts to export)
#' @param sym If True derive matrix entries below principle-diagonal
#' @export

kinship.emp <- function(animals=NULL, population=NULL, gen=NULL, database=NULL, cohorts=NULL, sym=FALSE){
  if(length(animals)==0 || length(gen)>0 || length(database)>0 || length(cohorts)>0){
    database <- get.database(population, gen, database, cohorts)
    animals <- list()
    for(index in 1:nrow(database)){
      if(diff(database[index,3:4])>=0){
        for(index2 in database[index,3]:database[index,4]){
          animals[[length(animals)+1]] <- population$breeding[[database[index,1]]][[database[[index,2]]]][[index2]]
        }
      }

    }
  }
  n <- length(animals)
  kinship <- matrix(0, nrow=n, ncol=n)
  chrom.length <- max(animals[[1]][[1]])
  for( i in 1:n){
    for( j in i:n){
        chr <- list()
        chr[[1]] <- animals[[i]][[1]][-1]
        chr[[2]] <- animals[[i]][[2]][-1]
        chr[[3]] <- animals[[j]][[1]][-1]
        chr[[4]] <- animals[[j]][[2]][-1]
        origin <- list()
        origin[[1]] <- animals[[i]][[5]]
        origin[[2]] <- animals[[i]][[6]]
        origin[[3]] <- animals[[j]][[5]]
        origin[[4]] <- animals[[j]][[6]]

        activ <- c(1,1,1,1)
        prev <- 0
        activ.recom <- c(chr[[1]][activ[1]], chr[[2]][activ[2]], chr[[3]][activ[3]], chr[[4]][activ[4]])
        activ.ursprung <- c(origin[[1]][activ[1]],origin[[2]][activ[2]],origin[[3]][activ[3]],origin[[4]][activ[4]])


        for(steps in 1:(length(c(chr[[1]], chr[[2]], chr[[3]], chr[[4]]))-3)){
          activ.min <- which.min(activ.recom)[1]
          activ.posi <- chr[[activ.min]][activ[activ.min]]
          ibd <- length(unique(activ.ursprung)) # Nur vergleich des Neuen mit bisherigen Rechenzeiteffizienter!

          ibd.factor <- 1-ibd*0.25 + 0.25*(ibd==1)
          kinship[i,j] <- kinship[i,j] + ibd.factor * (activ.posi - prev) / chrom.length
          prev <- activ.posi

          activ[activ.min] <- min(activ[activ.min] +1, length(chr[[activ.min]]))
          activ.recom[activ.min] <- chr[[activ.min]][activ[activ.min]]
          activ.ursprung[activ.min] <-  origin[[activ.min]][activ[activ.min]]

        }
      }

  }
  if(sym==TRUE){
    for(i in 1:n){
      for(j in 1:i){
        kinship[i,j] <- kinship[j,i]
      }
    }
  }
  return(kinship)
}

#' Empirical kinship
#'
#' Function to compute empirical kinship for a set of individuals)
#' @param animals List of animals to compute kinship for
#' @param population Population list
#' @param database Groups of individuals to consider for the export
#' @param gen Quick-insert for database (vector of all generations to export)
#' @param cohorts Quick-insert for database (vector of names of cohorts to export)
#' @param sym If True derive matrix entries below principle-diagonal
#' @param ibd.obs Number of Individual pairs to sample for IBD estimation
#' @param hbd.obs Number of Individuals to sample for HBD estimation
#' @export
#'
kinship.emp.fast <- function(animals=NULL, population=NULL, gen=NULL, database=NULL, cohorts=NULL, sym=FALSE, ibd.obs=50, hbd.obs=10){
  if(length(animals)==0 || length(gen)>0 || length(database)>0 || length(cohorts)>0){
    database <- get.database(population, gen=gen, database=database, cohorts=cohorts)
    animals <- list()
    for(index in 1:nrow(database)){
      if(diff(database[index,3:4])>=0){
        for(index2 in database[index,3]:database[index,4]){
          animals[[length(animals)+1]] <- population$breeding[[database[index,1]]][[database[[index,2]]]][[index2]]
        }
      }
    }
  }

  n <- length(animals)
  if(n==0){
    return(c(0,0))
  }

  kinship <- matrix(0, nrow=n, ncol=n)
  chrom.length <- max(animals[[1]][[1]])

  i1 <- sample(1:n, ibd.obs+ hbd.obs, replace=if((ibd.obs+hbd.obs)>n){TRUE} else{FALSE})
  j1 <- sample(1:n, ibd.obs+ hbd.obs, replace=if((ibd.obs+hbd.obs)>n){TRUE} else{FALSE})
  if(ibd.obs>0){
    while(sum(i1==j1)>0){
      j1[which(i1==j1)] <- sample(1:n, length(which(i1==j1)), replace=if((ibd.obs)>n){TRUE} else{FALSE})
    }
  }
  if(hbd.obs>0){
    j1[(ibd.obs+1):(ibd.obs+hbd.obs)] <- i1[(ibd.obs+1):(ibd.obs+hbd.obs)]
  }

  score <- numeric(length(i1))
  for(index in 1:length(i1)){
    i <- i1[index]
    j <- j1[index]
    chr <- list()
    chr[[1]] <- animals[[i]][[1]][-1]
    chr[[2]] <- animals[[i]][[2]][-1]
    chr[[3]] <- animals[[j]][[1]][-1]
    chr[[4]] <- animals[[j]][[2]][-1]
    origin <- list()
    origin[[1]] <- animals[[i]][[5]]
    origin[[2]] <- animals[[i]][[6]]
    origin[[3]] <- animals[[j]][[5]]
    origin[[4]] <- animals[[j]][[6]]

    activ <- c(1,1,1,1)
    prev <- 0
    activ.recom <- c(chr[[1]][activ[1]], chr[[2]][activ[2]], chr[[3]][activ[3]], chr[[4]][activ[4]])
    activ.ursprung <- c(origin[[1]][activ[1]],origin[[2]][activ[2]],origin[[3]][activ[3]],origin[[4]][activ[4]])


    for(steps in 1:(length(c(chr[[1]], chr[[2]], chr[[3]], chr[[4]]))-3)){
      activ.min <- which.min(activ.recom)[1]
      activ.posi <- chr[[activ.min]][activ[activ.min]]
      ibd <- length(unique(activ.ursprung)) # Nur vergleich des Neuen mit bisherigen Rechenzeiteffizienter!

      ibd.factor <- 1-ibd*0.25 + 0.25*(ibd==1)
      score[index] <- score[index] + ibd.factor * (activ.posi - prev) / chrom.length
      prev <- activ.posi

      activ[activ.min] <- min(activ[activ.min] +1, length(chr[[activ.min]]))
      activ.recom[activ.min] <- chr[[activ.min]][activ[activ.min]]
      activ.ursprung[activ.min] <-  origin[[activ.min]][activ[activ.min]]

    }
  }
  if(ibd.obs>0){
    ibd <- mean(score[1:ibd.obs])
  } else{
    ibd <- 0
  }
  if(hbd.obs>0){
    hbd <- mean(score[(ibd.obs+1):(ibd.obs+hbd.obs)])
  } else{
    hbd <- 0
  }

  return(c(ibd,hbd))
}
```
