## Supplementary material for "MoBPS - Modular Breeding Program Simulator": MoBPS R-package: kinship.exp.old.R

```
'#
  Authors
Torsten Pook,

Copyright (C) 2017 -- 2018  Torsten Pook

This program is free software; you can redistribute it and/or
modify it under the terms of the GNU General Public License
as published by the Free Software Foundation; either version 3
of the License, or (at your option) any later version.

This program is distributed in the hope that it will be useful,
but WITHOUT ANY WARRANTY; without even the implied warranty of
MERCHANTABILITY or FITNESS FOR A PARTICULAR PURPOSE.  See the
GNU General Public License for more details.

You should have received a copy of the GNU General Public License
along with this program; if not, write to the Free Software
Foundation, Inc., 59 Temple Place - Suite 330, Boston, MA  02111-1307, USA.
'#

#' Derive expected kinship
#'
#' Function to derive expected kinship
#' @param population Population list
#' @param prev.gen Maximum generatin gap between parent/offspring (default: Inf)
#' @param generation1.kinship Pedigree of the first generation (default: diag(1))
#' @param start.diagonal First X individuals are unrelated (default: 0)
#' @param calculate.averages Set to "generations" or "generationsex" to combine cohorts
#' @param ignore.diag Use kinship definition from Sitzenstock et al.
#' @param plot_grp Plot development of kinship per group
#' @param depth.pedigree Depth of the pedigree in generations


kinship.exp.old <- function(population, prev.gen=Inf, generation1.kinship=NULL, calculate.averages=FALSE, start.diagonal=0, ignore.diag=FALSE, plot_grp=FALSE,
                        depth.pedigree=Inf){

  generation.size <- cbind(population$info$size, rowSums(population$info$size))
  generations <- length(generation.size[,3])
  total <- sum(generation.size[,3])
  total.nr <- c(0,cumsum(generation.size[,3]))+1
  kinship <- matrix(0, ncol=total, nrow=total)
  if(length(generation1.kinship)>0){
    kinship <- generation1.kinship
  } else{
    # diag(1/2, ncol=nrow=generations.size[1,3])
    for(index in 1:generation.size[1,3]){
      kinship[index,index] <- 1/2
    }
  }
  if(start.diagonal>0){
    for(index in 1:start.diagonal){
      kinship[index,index] <- 1/2
    }
  }
  if(generations==1){
    return(kinship)
  }

  for(second in (total.nr[2]):total){
    gen.second <- sum(total.nr <= second)

    for(first in total.nr[max(gen.second - prev.gen,1)]:second){
      if(second>start.diagonal || first>start.diagonal){
        # Ermittle Vater/Mutter von second
        nr.second <- second - total.nr[gen.second] +1
        sex.second <- 2 - ((nr.second - generation.size[gen.second,1])<=0)
        if(sex.second==2)  nr.second <- nr.second - generation.size[gen.second,1]

        father <- population$breeding[[gen.second]][[sex.second]][[nr.second]][[7]]
        mother <- population$breeding[[gen.second]][[sex.second]][[nr.second]][[8]]

        nr.father <- total.nr[father[1]] -1 + father[3] + ((father[2]==2)*generation.size[mother[1],1]) # Selbstung!
        nr.mother <- total.nr[mother[1]] -1 + mother[3] + ((mother[2]==2)*generation.size[mother[1],1]) # add nr of male animals in the generation

        if(first!=second){
          kinship[first,second] <- 1/2 * (kinship[first, nr.father] + kinship[first, nr.mother])
          kinship[second,first] <- 1/2 * (kinship[first, nr.father] + kinship[first, nr.mother])
        } else{
          kinship[first,second] <- 1/2 + 1/2 * kinship[nr.father, nr.mother]
          kinship[second,first] <- 1/2 + 1/2 * kinship[nr.father, nr.mother]
        }

      }

    }
  }
  if(calculate.averages==FALSE){
    return(kinship)
  } else{
    groups <- calculate.averages
    if(groups=="generations"){
      groups <- total.nr
    }
    if(groups[1]=="generationsex"){
      groups <- sort(c(1, generation.size[,1]+total.nr[-length(total.nr)], generation.size[,3]+total.nr[-length(total.nr)]))
    }
    if(groups[1]=="generationsexmig"){
      groups <- 1
      for(index in 1:length(population$breeding)){
        for(sex in 1:2){
          migs <- unique(population$breeding[[index]][[4+sex]])
          for(index2 in migs){
            groups <- c(groups, groups[length(groups)] + sum(population$breeding[[index]][[4+sex]]==index2))
          }
        }
      }
    }
    n.groups <- (length(groups)-1)
    kinship.matrix <- matrix(0, nrow=n.groups, ncol = n.groups)
    hbd <- numeric(n.groups)

    for(i in 1:n.groups){
      for(j in 1:i){
        if((groups[i] <= (groups[i+1]-1) ) && groups[j] <= (groups[j+1]-1))
        kinship.matrix[i,j] <- mean(kinship[groups[i]:(groups[i+1]-1),groups[j]:(groups[j+1]-1)])
        kinship.matrix[j,i] <- kinship.matrix[i,j]
        if(ignore.diag==TRUE && i==j && (groups[i+1]-groups[i])>0){
          tot.diag <- sum( kinship[groups[i]:(groups[i+1]-1),groups[j]:(groups[j+1]-1)]) - sum(diag(kinship[groups[i]:(groups[i+1]-1),groups[j]:(groups[j+1]-1)]))
          kinship.matrix[i,j] <- tot.diag/ ((groups[i+1]-groups[i])) / ((groups[i+1]-1-groups[i]))
          if(length(groups[i]:(groups[i+1]-1))==1){
            hbd[i] <- -1 + 2*kinship[groups[i]:(groups[i+1]-1),groups[j]:(groups[j+1]-1)] / ((groups[i+1]-groups[i]))

          } else{
            hbd[i] <- -1 + 2*sum(diag(kinship[groups[i]:(groups[i+1]-1),groups[j]:(groups[j+1]-1)])) / ((groups[i+1]-groups[i]))

          }
        }


      }
    }
    if(plot_grp){
      graphics::plot(diag(kinship.matrix), xlab="Generation", ylab="Kinship", main="Entwicklung der mittleren Kinship")
    }
    return(list(kinship.matrix, hbd))
  }

}
```
