## Supplementary material for "MoBPS - Modular Breeding Program Simulator": MoBPS R-package: kinship.exp.R

```
'#
  Authors
Torsten Pook,

Copyright (C) 2017 -- 2018  Torsten Pook

This program is free software; you can redistribute it and/or
modify it under the terms of the GNU General Public License
as published by the Free Software Foundation; either version 3
of the License, or (at your option) any later version.

This program is distributed in the hope that it will be useful,
but WITHOUT ANY WARRANTY; without even the implied warranty of
MERCHANTABILITY or FITNESS FOR A PARTICULAR PURPOSE.  See the
GNU General Public License for more details.

You should have received a copy of the GNU General Public License
along with this program; if not, write to the Free Software
Foundation, Inc., 59 Temple Place - Suite 330, Boston, MA  02111-1307, USA.
'#

#' Derive expected kinship
#'
#' Function to derive expected kinship
#' @param population Population list
#' @param database Groups of individuals to consider for the export
#' @param gen Quick-insert for database (vector of all generations to export)
#' @param cohorts Quick-insert for database (vector of names of cohorts to export)
#' @param depth.pedigree Depth of the pedigree in generations
#' @param start.kinship Relationship matrix of the individuals in the first considered generation
#' @export

kinship.exp <- function(population, gen=NULL, database=NULL, cohorts=NULL, depth.pedigree=7,
                        start.kinship=NULL){

#                        prev.gen=Inf, generation1.kinship=NULL, calculate.averages=FALSE, start.diagonal=0, ignore.diag=FALSE, plot_grp=FALSE,

  database <- get.database(population, gen=gen, database=database, cohorts=cohorts)

  if(depth.pedigree==Inf){
    pedigree.database <- get.database(population, gen=1:max(database[,1]))
  } else{
    new.pedigree.database <- pedigree.database <- database
    remaining.depth <- depth.pedigree
    while(remaining.depth>0){
      parents <- get.pedigree(population, database = new.pedigree.database, raw=TRUE)
      m_parents <- rbind(parents[parents[,5]==1,4:6], parents[parents[,8]==1,7:9])
      f_parents <- rbind(parents[parents[,5]==2,4:6], parents[parents[,8]==2,7:9])
      if(nrow(m_parents)>0){
        m_gen <- unique(m_parents[,1])
        m_data <- cbind(m_gen, 1, 0,0)
        for(index in 1:length(m_gen)){
          m_data[index,3] <- min(m_parents[m_parents[,1]==m_gen[index],3])
          m_data[index,4] <- max(m_parents[m_parents[,1]==m_gen[index],3])
        }
      } else{
        m_data <- NULL
      }
      if(nrow(f_parents)>0){
        f_gen <- unique(f_parents[,1])
        f_data <- cbind(f_gen, 2, 0,0)
        for(index in 1:length(f_gen)){
          f_data[index,3] <- min(f_parents[f_parents[,1]==f_gen[index],3])
          f_data[index,4] <- max(f_parents[f_parents[,1]==f_gen[index],3])
        }

      } else{
        f_data <- NULL
      }

      new.pedigree.database <- get.database(population, database=rbind(m_data,f_data))
      new.pedigree.database <- unique(new.pedigree.database)
      remaining.depth <- remaining.depth - 1
      pedigree.database <- rbind(new.pedigree.database, pedigree.database)
    }

    pedigree.database <- get.database(population, database = pedigree.database)
  }
  n.animals <- sum(diff(t(database[,3:4, drop=FALSE]))+1)
  n.total <- sum(diff(t(pedigree.database[,3:4, drop=FALSE]))+1)
  position.pedigree <- numeric(n.animals)
  for(index in 1:nrow(database)){
    activ_ped <- which(pedigree.database[,1]==database[index,1] & pedigree.database[,2]==database[index,2] & pedigree.database[,3]<= database[index,3] & pedigree.database[,4]>= database[index,4])[1]
    if(activ_ped>1){
      prior <- sum(diff(t(pedigree.database[1:(activ_ped-1),3:4,drop=FALSE]))+1) - database[index,3] + pedigree.database[activ_ped,3]
    } else{
      prior <-  pedigree.database[activ_ped,3] - database[index,3]
    }
    if(index==1){
      prior2 <- 0
    } else{
      prior2 <- sum(diff(t(database[1:(index-1),3:4, drop=FALSE]))+1)
    }
    position.pedigree[1:(diff(database[index,3:4])+1) + prior2] <- 1:(diff(database[index,3:4])+1) + prior
  }

  kinship <- matrix(0, ncol=n.total, nrow=n.total)

  group.size <- pedigree.database[,4]-pedigree.database[,3] +1
  if(length(start.kinship)==0){
    size.firstgen <- sum(group.size[pedigree.database[,1]==pedigree.database[1,1]])
    kinship[1:size.firstgen, 1:size.firstgen] <- diag(1/2,size.firstgen)
  } else{
    kinship[1:nrow(start.kinship), 1:nrow(start.kinship)] <- start.kinship
    # Add reality check to validate size of start.kinship
  }
  first_new <- sum(pedigree.database[,1]==pedigree.database[1,1]) +1
  total <- sum(group.size)
  total.nr <- c(0,cumsum(group.size))+1

  ## Potential export individual id in the pedigree - more efficient for high number of copies!
  animal.nr <- get.id(population, database=pedigree.database)
  info.indi <- get.pedigree(population, database=pedigree.database)
  info.indi[info.indi=="0"] <- "M1_1" # Placeholder
  # necessary when using copy.individuals
  replaces <- which(duplicated(animal.nr))
#  for(replace in replaces){
#    new <- which(animal.nr==animal.nr[replace])[1]
#    info.indi[info.indi==info.indi[replace,1]] <- info.indi[new,1]
#  }

  if(length(replaces)>0){
    animal.nr.temp <- animal.nr[1:min(replaces)]
    for(replace in replaces){
      new <- which(animal.nr.temp==animal.nr[replace])[1]
      if(length(new)==0){
        animal.nr.temp <- animal.nr[1:replace]
        new <- which(animal.nr.temp==animal.nr[replace])[1]
      }
      info.indi[info.indi==info.indi[replace,1]] <- info.indi[new,1]
    }
  }

  sex.indi <- as.numeric(substr(info.indi[,1], start=1, stop=1)=="F") +1
  temp1 <- as.numeric(unlist(strsplit(substr(info.indi[,1], start=2, stop=nchar(info.indi[,1])), "\\_")))
  gen.indi <- temp1[1:(length(temp1)/2) *2]
  nr.indi <- temp1[1:(length(temp1)/2) *2 -1]

  sex.father <- as.numeric(substr(info.indi[,2], start=1, stop=1)=="F") +1
  temp1 <- as.numeric(unlist(strsplit(substr(info.indi[,2], start=2, stop=nchar(info.indi[,2])), "\\_")))
  if(length(temp1)>0){
    gen.father <- temp1[1:(length(temp1)/2) *2]
    nr.father <- temp1[1:(length(temp1)/2) *2 -1]
  }


  sex.mother <- as.numeric(substr(info.indi[,3], start=1, stop=1)=="F") +1
  temp1 <- as.numeric(unlist(strsplit(substr(info.indi[,3], start=2, stop=nchar(info.indi[,3])), "\\_")))
  if(length(temp1)>0){
    gen.mother <- temp1[1:(length(temp1)/2) *2]
    nr.mother <- temp1[1:(length(temp1)/2) *2 -1]
  }


  nr_father <- nr_mother <- numeric(total)
  for(index in (total.nr[first_new]):total){
    group_father <- which(pedigree.database[,1]==gen.father[index] & pedigree.database[,2] == sex.father[index] & pedigree.database[,3] <= nr.father[index] & pedigree.database[,4] >= nr.father[index])[1]
    group_mother <- which(pedigree.database[,1]==gen.mother[index] & pedigree.database[,2] == sex.mother[index] & pedigree.database[,3] <= nr.mother[index] & pedigree.database[,4] >= nr.mother[index])[1]
    nr_father[index] <- nr.father[index] - pedigree.database[group_father,3] + total.nr[group_father]
    nr_mother[index] <- nr.mother[index] - pedigree.database[group_mother,3] + total.nr[group_mother]

  }

#  if((total.nr[first_new]) <= total){
#    for(second in (total.nr[first_new]):total){
#      for(first in 1:second){
#        nr.father <- nr_father[second]
#        nr.mother <- nr_mother[second]
#        if(first!=second){
#          kinship[first,second] <- 1/2 * (kinship[first, nr.father] + kinship[first, nr.mother])
#          kinship[second,first] <- 1/2 * (kinship[first, nr.father] + kinship[first, nr.mother])
#        } else{
#         kinship[first,second] <- 1/2 + 1/2 * kinship[nr.father, nr.mother]
#          kinship[second,first] <- 1/2 + 1/2 * kinship[nr.father, nr.mother]
#        }
#      }
#    }
#  }


  animal_ids <- get.id(population, database = pedigree.database)
  if((total.nr[first_new]) <= total){
    for(second in (total.nr[first_new]):total){
      nr.father <- nr_father[second]
      nr.mother <- nr_mother[second]
      first <- 1:second
      if(is.na(nr.father) && is.na(nr.mother)){
        kinship[second,second] <- 1/2
        nr.mother <- nr.father <- 1
      }
      if(is.na(nr.father)){
        kinship[first,second] <- kinship[second,first] <-1/2 * (0 + kinship[first, nr.mother])
        nr.mother <- nr.father <- 1 # Founder-individual
      } else if(is.na(nr.mother)){
        kinship[first,second] <- kinship[second,first] <-1/2 * (kinship[first, nr.father] + 0)
        nr.mother <- nr.father <- 1 # Founder-individual
      } else{
        kinship[first,second] <- kinship[second,first] <-1/2 * (kinship[first, nr.father] + kinship[first, nr.mother])
      }

      if(nr.father==nr.mother && animal_ids[nr.father]==animal_ids[second]){
        kinship[second,second] <- 1/2
        # Individual is founder!
      } else{
        kinship[second,second] <- 1/2 + 1/2 * kinship[nr.father, nr.mother]
      }

    }
  }

#  for(replace in intersect(replaces, position.pedigree)){
#    new <- which(animal.nr==animal.nr[replace])[1]
#    kinship[replace,] <- kinship[new,]
#    kinship[,replace] <- kinship[,new]
#  }

  kinship.relevant <- kinship[position.pedigree,position.pedigree]

  return(kinship.relevant)

  if(FALSE){
    groups <- calculate.averages
    if(groups=="generations"){
      groups <- total.nr
    }
    if(groups[1]=="generationsex"){
      groups <- sort(c(1, generation.size[,1]+total.nr[-length(total.nr)], generation.size[,3]+total.nr[-length(total.nr)]))
    }
    if(groups[1]=="generationsexmig"){
      groups <- 1
      for(index in 1:length(population$breeding)){
        for(sex in 1:2){
          migs <- unique(population$breeding[[index]][[4+sex]])
          for(index2 in migs){
            groups <- c(groups, groups[length(groups)] + sum(population$breeding[[index]][[4+sex]]==index2))
          }
        }
      }
    }
    n.groups <- (length(groups)-1)
    kinship.matrix <- matrix(0, nrow=n.groups, ncol = n.groups)
    hbd <- numeric(n.groups)

    for(i in 1:n.groups){
      for(j in 1:i){
        if((groups[i] <= (groups[i+1]-1) ) && groups[j] <= (groups[j+1]-1))
          kinship.matrix[i,j] <- mean(kinship[groups[i]:(groups[i+1]-1),groups[j]:(groups[j+1]-1)])
        kinship.matrix[j,i] <- kinship.matrix[i,j]
        if(ignore.diag==TRUE && i==j && (groups[i+1]-groups[i])>0){
          tot.diag <- sum( kinship[groups[i]:(groups[i+1]-1),groups[j]:(groups[j+1]-1)]) - sum(diag(kinship[groups[i]:(groups[i+1]-1),groups[j]:(groups[j+1]-1)]))
          kinship.matrix[i,j] <- tot.diag/ ((groups[i+1]-groups[i])) / ((groups[i+1]-1-groups[i]))
          if(length(groups[i]:(groups[i+1]-1))==1){
            hbd[i] <- -1 + 2*kinship[groups[i]:(groups[i+1]-1),groups[j]:(groups[j+1]-1)] / ((groups[i+1]-groups[i]))

          } else{
            hbd[i] <- -1 + 2*sum(diag(kinship[groups[i]:(groups[i+1]-1),groups[j]:(groups[j+1]-1)])) / ((groups[i+1]-groups[i]))

          }
        }


      }
    }
    if(plot_grp){
      graphics::plot(diag(kinship.matrix), xlab="Generation", ylab="Kinship", main="Entwicklung der mittleren Kinship")
    }
    return(list(kinship.matrix, hbd))
  }

}
```
