## Supplementary material for "MoBPS - Modular Breeding Program Simulator": MoBPS R-package: ld.decay.R

```
'#
  Authors
Torsten Pook,

Copyright (C) 2017 -- 2018  Torsten Pook

This program is free software; you can redistribute it and/or
modify it under the terms of the GNU General Public License
as published by the Free Software Foundation; either version 3
of the License, or (at your option) any later version.

This program is distributed in the hope that it will be useful,
but WITHOUT ANY WARRANTY; without even the implied warranty of
MERCHANTABILITY or FITNESS FOR A PARTICULAR PURPOSE.  See the
GNU General Public License for more details.

You should have received a copy of the GNU General Public License
along with this program; if not, write to the Free Software
Foundation, Inc., 59 Temple Place - Suite 330, Boston, MA  02111-1307, USA.
'#

#' Generate LD plot
#'
#' Generate LD pot
#' @param population Population list
#' @param genotype.dataset Genotype dataset (default: NULL - just to save computation time when get.geno was already run)
#' @param step Stepsize to calculate LD
#' @param max Maximum distance between markers to consider for LD-plot
#' @param database Groups of individuals to consider for the export
#' @param gen Quick-insert for database (vector of all generations to export)
#' @param cohorts Quick-insert for database (vector of names of cohorts to export)
#' @param chromosomen Only consider a specific chromosome in calculations (default: 1)
#' @export

ld.decay <- function(population, genotype.dataset=NULL, chromosomen=1, step=5, max=500,database=NULL, gen=NULL, cohorts= NULL){
  max <- min(population$info$snp[chromosomen]-1, max)
  if(length(genotype.dataset)==0){
    dataset <- t(get.geno(population, chromosomen = chromosomen, gen=gen, database=database,cohorts=cohorts))
  } else{
    dataset <- t(genotype.dataset)
  }

  calc <- unique(c(1,seq(from=step, to=max, by=step)))
  ld <- numeric(length(calc))
  for(index in 1:length(calc)){
    lds <- numeric(population$info$snp[chromosomen]-calc[index])
    for(index2 in 1:(population$info$snp[chromosomen]-calc[index])){
      lds[index2] <- stats::cor(dataset[,index2], dataset[,index2+calc[index]])
    }
    ld[index] <- mean(lds^2, na.rm=TRUE)
  }
  graphics::plot(calc, ld, xlab="distance in SNP", ylab=expression(r^2))
  graphics::lines(calc, ld)
}
```
