## Supplementary material for "MoBPS - Modular Breeding Program Simulator": MoBPS R-package: miesenberger.index.R

```
#' gen/database/cohorts conversion
#'
#' Function to derive a database based on gen/database/cohorts
#' @param V1 Inverted phenotypic covarianz matrix
#' @param V Phenotypic covarianz matrix
#' @param G Genomic covarianz matrix
#' @param RG Genomic correlation matrix
#' @param r reliability for the breeding value estimation
#' @param w relative weighting of each trait (per genetic SD)
#' @param zw Estimated breeding value
#' @export

miesenberger.index <- function(V, G, V1=NULL, RG=NULL, r, w, zw=NULL){
  if(length(V1)==0){
    V1 <- chol2inv(chol(V))
  }
  if(length(RG)==0){
    RG <- sqrt(diag(1/diag(G))) %*% G %*% sqrt(diag(1/diag(G)))
  }

  d <- nrow(G)
  A <- RG * matrix(sqrt(diag(G)), nrow=d, ncol=d, byrow=TRUE) *
    matrix(sqrt(diag(V)), nrow=d, ncol=d, byrow=FALSE) *
    matrix(r, nrow=d, ncol=d, byrow=FALSE)
  bM <- as.numeric(V1 %*% A %*% w)
  if(length(zw)==0){
    return(bM)
  } else{
    IM <- sum( bM * zw )
    return(IM)
  }
}
```
