## Supplementary material for "MoBPS - Modular Breeding Program Simulator": MoBPS R-package: mutation.intro.R

```
'#
  Authors
Torsten Pook,

Copyright (C) 2017 -- 2018  Torsten Pook

This program is free software; you can redistribute it and/or
modify it under the terms of the GNU General Public License
as published by the Free Software Foundation; either version 3
of the License, or (at your option) any later version.

This program is distributed in the hope that it will be useful,
but WITHOUT ANY WARRANTY; without even the implied warranty of
MERCHANTABILITY or FITNESS FOR A PARTICULAR PURPOSE.  See the
GNU General Public License for more details.

You should have received a copy of the GNU General Public License
along with this program; if not, write to the Free Software
Foundation, Inc., 59 Temple Place - Suite 330, Boston, MA  02111-1307, USA.
'#

#' Mutation intro
#'
#' Function to change the base-pair in a specific loci
#' @param population Population list
#' @param gen Generation of the individual to introduce a mutation in
#' @param sex Sex of the individual to introduce a mutation in
#' @param individual.nr Individual Nr. of the individual to introduce a mutation in
#' @param qtl.posi Marker number to mutate
#' @param haplo.set Select chromosome set (default: 1 , alt: 2)
#' @export

mutation.intro <- function(population, gen, sex, individual.nr, qtl.posi, haplo.set=1) {

  if(sum(population$breeding[[gen]][[sex]][[individual.nr]][[2+ haplo.set]]==qtl.posi)==0){
    population$breeding[[gen]][[sex]][[individual.nr]][[2+ haplo.set]] <- sort(c(qtl.posi,population$breeding[[gen]][[sex]][[individual.nr]][[2+ haplo.set]]))
  } else{
    population$breeding[[gen]][[sex]][[individual.nr]][[2+ haplo.set]] <- unique(c(qtl.posi,population$breeding[[gen]][[sex]][[individual.nr]][[2+ haplo.set]]))[-1]
  }
  return(population)
}
```
