## Supplementary material for "MoBPS - Modular Breeding Program Simulator": MoBPS R-package: new.base.generation.R

```
'#
  Authors
Torsten Pook,

Copyright (C) 2017 -- 2018  Torsten Pook

This program is free software; you can redistribute it and/or
modify it under the terms of the GNU General Public License
as published by the Free Software Foundation; either version 3
of the License, or (at your option) any later version.

This program is distributed in the hope that it will be useful,
but WITHOUT ANY WARRANTY; without even the implied warranty of
MERCHANTABILITY or FITNESS FOR A PARTICULAR PURPOSE.  See the
GNU General Public License for more details.

You should have received a copy of the GNU General Public License
along with this program; if not, write to the Free Software
Foundation, Inc., 59 Temple Place - Suite 330, Boston, MA  02111-1307, USA.
'#

#' Set new base generation
#'
#' Function to set a new base generation for the population
#' @param population Population list
#' @param base.gen Vector containing all new base generations
#' @param delete.previous.gen Delete all data before base.gen (default: FALSE)
#' @param delete.breeding.totals Delete all breeding totals before base.gen (default: FALSE)
#' @param delete.bve.data Deleta all previous bve data (default: FALSE)
#' @param add.chromosome.ends Add chromosome ends as recombination points
#' @param miraculix If TRUE use miraculix package for data storage and computation time relevant computations
#' @export

new.base.generation <- function(population, base.gen=NULL, delete.previous.gen=FALSE, delete.breeding.totals=FALSE,
                                delete.bve.data=FALSE, add.chromosome.ends=TRUE, miraculix=FALSE){

  if (requireNamespace("miraculix", quietly = TRUE)) {
    codeOriginsU <- miraculix::codeOrigins
    decodeOriginsU <- miraculix::decodeOrigins
  } else{
    codeOriginsU <- codeOriginsR
    decodeOriginsU <- decodeOriginsR
  }
  if(length(population$info$miraculix)>0 && population$info$miraculix){
    miraculix <- TRUE
  }

  take <- which(population$info$origin.gen==base.gen)
  if(length(take)==1){
    origin_code <- population$info$origin.gen[take]
  } else{
    if(length(population$info$origin.gen)<64){
      population$info$origin.gen <- c(population$info$origin.gen, as.integer(base.gen))
      origin_code <- length(population$info$origin.gen)
    } else{
      print("To many origin generation!")
      print("Delete second lowest origin.gen")
      switch_gen <- sort(population$info$origin.gen, index.return=TRUE)[[2]]
      population$info$origin.gen[switch_gen] <- as.integer(base.gen)
      origin_code <- switch_gen
    }
  }


  if(length(base.gen)==0){
    base.gen <- length(population$breeding)
  }
  for(gen in base.gen){
    for(sex in 1:2){
      if(length(population$breeding[[gen]][[sex]])>0){
        for(nr in 1:length(population$breeding[[gen]][[sex]])){
          if(miraculix){
            population$breeding[[gen]][[sex]][[nr]][[9]] <- miraculix::computeSNPS(population, gen, sex, nr, what="haplo", output_compressed=TRUE)
          } else{
            snps <- compute.snps(population, gen, sex, nr, decodeOriginsU=decodeOriginsU)
            population$breeding[[gen]][[sex]][[nr]][[9]] <- snps[1,]
            population$breeding[[gen]][[sex]][[nr]][[10]] <- snps[2,]
          }

          population$breeding[[gen]][[sex]][[nr]][[1]] <- c(0, sum(population$info$length))
          population$breeding[[gen]][[sex]][[nr]][[2]] <- c(0, sum(population$info$length))
          if(add.chromosome.ends==TRUE){
            population$breeding[[gen]][[sex]][[nr]][[1]] <- population$info$length.total
            population$breeding[[gen]][[sex]][[nr]][[2]] <- population$info$length.total
          }
          population$breeding[[gen]][[sex]][[nr]][[3]] <- numeric(0)
          population$breeding[[gen]][[sex]][[nr]][[4]] <- numeric(0)
          population$breeding[[gen]][[sex]][[nr]][[5]] <- codeOriginsU(matrix(c(origin_code, sex, nr, 1),nrow=(length(population$breeding[[gen]][[sex]][[nr]][[1]])-1), ncol=4, byrow=TRUE))

          population$breeding[[gen]][[sex]][[nr]][[6]] <- codeOriginsU(matrix(c(origin_code, sex, nr, 2),nrow=(length(population$breeding[[gen]][[sex]][[nr]][[2]])-1), ncol=4, byrow=TRUE))
        }

      }

    }
  }
  if(delete.previous.gen){
    for(index in 1:(min(base.gen)-1)){
      population$breeding[[index]] <- "deleted"
    }
  }
  if(delete.breeding.totals){
    population$info$breeding.totals <- NULL
  }
  if(delete.bve.data){
    population$info$bve.data <- NULL
  }
  return(population)
}
```
