## Supplementary material for "MoBPS - Modular Breeding Program Simulator": MoBPS R-package: OGC.R

```
'#
  Authors
Torsten Pook,

Copyright (C) 2017 -- 2018  Torsten Pook

This program is free software; you can redistribute it and/or
modify it under the terms of the GNU General Public License
as published by the Free Software Foundation; either version 3
of the License, or (at your option) any later version.

This program is distributed in the hope that it will be useful,
but WITHOUT ANY WARRANTY; without even the implied warranty of
MERCHANTABILITY or FITNESS FOR A PARTICULAR PURPOSE.  See the
GNU General Public License for more details.

You should have received a copy of the GNU General Public License
along with this program; if not, write to the Free Software
Foundation, Inc., 59 Temple Place - Suite 330, Boston, MA  02111-1307, USA.
'#

#' THIS NEEDs SOME REWORK BEFORE USAGE
#'
#' THIS NEEDs SOME REWORK BEFORE USAGE
#' @param A A
#' @param u u
#' @param Q Q
#' @param cAc cAc
#' @param single single
#' @export

OGC <- function(A,u,Q,cAc=NA, single=TRUE){
  # bei NA wird cAc (Increase of average relationship) minimiert

  Opt.int <- function(A, u, Q, cAc=NA){

    A1 <- MASS::ginv(A)
    QAQ1 <- MASS::ginv(t(Q)%*%A1%*%Q)
    minA <- round(0.25*sum(QAQ1)+0.000005,5)
    if(minA<=0){
      minA <- 0.000005
    }
    if(!is.na(cAc)){
      if(minA>cAc){
        cat(paste0("Minimal increase possible is for cAc is ",minA,"\n"))
        cAc <- minA
        }
    }else{
      cAc <- minA
    }
    lambda0 <- as.numeric(sqrt((t(u)%*%(A1-A1%*%Q%*%QAQ1%*%t(Q)%*%A1)%*%u)/(4*cAc-sum(QAQ1))) )
    lambda <- QAQ1%*%(t(Q)%*%A1%*%u - lambda0)

    xopt <- A1%*%(u-Q%*%lambda)/(2*lambda0)
    return(list(xopt,cAc))
  }

  n_male <- sum(Q[,1])
  n_female <- sum(Q[,2])
  rr <- Opt.int(A,u,Q,cAc)
  xopt <- rr[[1]]
  minA <- minA1 <- rr[[2]]
  n <- length(u)
  names(u) <- 1:n
  for(i in 1:length(xopt)){
    if((min(xopt) >= 0)){break}
    neg <- which(xopt < 0)
    if(single && length(neg)>0){
      neg <- sample(neg, 1)
    }
    A <- A[-neg,-neg]
    u <- u[-neg]
    Q <- Q[-neg,]
    if(length(u)==2){xopt=rep(0.5,2);break}
    xopt1 <- Opt.int(A,u,Q,cAc)
    xopt <- xopt1[[1]]
    minA1 <- xopt1[[2]]
  }
  res <- numeric(n)
  res[as.numeric(names(u))] <- xopt
  cat(paste0("Realized gain in inbreeding via OGC: ", minA1 ,"\n"))
  cat(paste0(sum(Q[,1]), " of the ", n_male, " male individuals have positive contribution.\n"))
  cat(paste0(sum(Q[,2]), " of the ", n_female, " female individuals have positive contribution.\n"))

  return(list("Optimal c"=res, "c'u"=t(xopt)%*%u))
}
```
