## Supplementary material for "MoBPS - Modular Breeding Program Simulator": MoBPS R-package: panel.cor.R

```
'#
  Authors
Torsten Pook,

Copyright (C) 2017 -- 2018  Torsten Pook

This program is free software; you can redistribute it and/or
modify it under the terms of the GNU General Public License
as published by the Free Software Foundation; either version 3
of the License, or (at your option) any later version.

This program is distributed in the hope that it will be useful,
but WITHOUT ANY WARRANTY; without even the implied warranty of
MERCHANTABILITY or FITNESS FOR A PARTICULAR PURPOSE.  See the
GNU General Public License for more details.

You should have received a copy of the GNU General Public License
along with this program; if not, write to the Free Software
Foundation, Inc., 59 Temple Place - Suite 330, Boston, MA  02111-1307, USA.
'#

#' par function for plots
#'
#' par function for plots#'
#' @param x x
#' @param y y
#' @param digits digits
#' @param prefix prefix
#' @param cex.cor cex.cor

panel.cor <- function(x, y, digits=2, prefix="", cex.cor) {
  usr <- graphics::par("usr"); on.exit(graphics::par(usr))
  graphics::par(usr = c(0, 1, 0, 1))
  r <- stats::cor(x, y, use="complete.obs")
  txt <- format(c(r, 0.123456789), digits=digits)[1]
  txt <- paste(prefix, txt, sep="")
  if(missing(cex.cor)) cex <- 0.8/graphics::strwidth(txt)
  graphics::text(0.5, 0.5, txt, cex = cex * abs(r))
}
```
