## Supplementary material for "MoBPS - Modular Breeding Program Simulator": MoBPS R-package: parent.haeufig.R

```
'#
  Authors
Torsten Pook,

Copyright (C) 2017 -- 2018  Torsten Pook

This program is free software; you can redistribute it and/or
modify it under the terms of the GNU General Public License
as published by the Free Software Foundation; either version 3
of the License, or (at your option) any later version.

This program is distributed in the hope that it will be useful,
but WITHOUT ANY WARRANTY; without even the implied warranty of
MERCHANTABILITY or FITNESS FOR A PARTICULAR PURPOSE.  See the
GNU General Public License for more details.

You should have received a copy of the GNU General Public License
along with this program; if not, write to the Free Software
Foundation, Inc., 59 Temple Place - Suite 330, Boston, MA  02111-1307, USA.
'#
#' par function for plots
#'
#' par function for plots
#' @param population Population list
#' @param gen generation to consider

parent.haeufig <- function(population, gen){
  nanimals <- length(population$breeding[[gen]][[1]])
  previous <- numeric(population$info$size[(gen-1),1])
  for(index in 1:nanimals){
    father <- population$breeding[[gen]][[1]][[index]][[7]][3]
    mother <- population$breeding[[gen]][[1]][[index]][[8]][3]
    previous[father] <- previous[father]+1
    previous[mother] <- previous[mother]+1
  }
  return(previous)
}
```
