## Supplementary material for "MoBPS - Modular Breeding Program Simulator": MoBPS R-package: pedigree.emp.R

```
'#
  Authors
Torsten Pook,

Copyright (C) 2017 -- 2018  Torsten Pook

This program is free software; you can redistribute it and/or
modify it under the terms of the GNU General Public License
as published by the Free Software Foundation; either version 3
of the License, or (at your option) any later version.

This program is distributed in the hope that it will be useful,
but WITHOUT ANY WARRANTY; without even the implied warranty of
MERCHANTABILITY or FITNESS FOR A PARTICULAR PURPOSE.  See the
GNU General Public License for more details.

You should have received a copy of the GNU General Public License
along with this program; if not, write to the Free Software
Foundation, Inc., 59 Temple Place - Suite 330, Boston, MA  02111-1307, USA.
'#

#' Empirical pedigree
#'
#' Function to compute the empirical pedigree
#' @param animals List of individuals to consider
#' @param skip.diagonal Set to TRUE to ignore HBD
#' @export

pedigree.emp <- function(animals, skip.diagonal=FALSE){
  n <- length(animals)
  pedigree <- matrix(0, nrow=n, ncol=n)
  chrom.length <- max(animals[[1]][[1]])
  for( i in 1:n){
    for( j in i:n){
      if(i==j && skip.diagonal==TRUE){

      } else{
      chr <- list()
      chr[[1]] <- animals[[i]][[1]][-1]
      chr[[2]] <- animals[[i]][[2]][-1]
      chr[[3]] <- animals[[j]][[1]][-1]
      chr[[4]] <- animals[[j]][[2]][-1]
      origin <- list()
      origin[[1]] <- animals[[i]][[5]]
      origin[[2]] <- animals[[i]][[6]]
      origin[[3]] <- animals[[j]][[5]]
      origin[[4]] <- animals[[j]][[6]]

      activ <- c(1,1,1,1)
      prev <- 0
      activ.recom <- c(chr[[1]][activ[1]], chr[[2]][activ[2]], chr[[3]][activ[3]], chr[[4]][activ[4]])
      activ.ursprung <- rbind(origin[[1]][activ[1]],origin[[2]][activ[2]],origin[[3]][activ[3]],origin[[4]][activ[4]])


      for(steps in 1:(length(c(chr[[1]], chr[[2]], chr[[3]], chr[[4]]))-3)){
        activ.min <- which.min(activ.recom)[1]
        activ.posi <- chr[[activ.min]][activ[activ.min]]

        ibd.factor <- (prod(activ.ursprung[1,] == activ.ursprung[3,]) +
          prod(activ.ursprung[1,] == activ.ursprung[4,]) +
          prod(activ.ursprung[2,] == activ.ursprung[3,]) +
          prod(activ.ursprung[2,] == activ.ursprung[4,]))/4


        pedigree[i,j] <- pedigree[i,j] + ibd.factor * (activ.posi - prev) / chrom.length
        prev <- activ.posi

        activ[activ.min] <- activ[activ.min] +1
        if(steps<(length(c(chr[[1]], chr[[2]], chr[[3]], chr[[4]]))-3)){
          activ.recom[activ.min] <- chr[[activ.min]][activ[activ.min]]
          activ.ursprung[activ.min,] <-  origin[[activ.min]][activ[activ.min]]
        }


      }
      print(c(i,j))
      }
    }

  }

  return(pedigree)
}
```
