## Supplementary material for "MoBPS - Modular Breeding Program Simulator": MoBPS R-package: pedmap.to.phasedbeaglevcf.R

```
'#
  Authors
Torsten Pook,

Copyright (C) 2017 -- 2018  Torsten Pook

This program is free software; you can redistribute it and/or
modify it under the terms of the GNU General Public License
as published by the Free Software Foundation; either version 3
of the License, or (at your option) any later version.

This program is distributed in the hope that it will be useful,
but WITHOUT ANY WARRANTY; without even the implied warranty of
MERCHANTABILITY or FITNESS FOR A PARTICULAR PURPOSE.  See the
GNU General Public License for more details.

You should have received a copy of the GNU General Public License
along with this program; if not, write to the Free Software
Foundation, Inc., 59 Temple Place - Suite 330, Boston, MA  02111-1307, USA.
'#

#' Perform imputing/phasing
#'
#' Perform imputing/phasing (path chosen for the web-based application)
#' @param ped_path Directory of the ped-file
#' @param map_path Directory of the map-file
#' @param vcf_path Directory of the vcf-file (this will override any ped/map-file input)
#' @param beagle_jar Directory of BEAGLE
#' @param plink_dir Directory of Plink
#' @param db_dir Directory to save newly generated files (ped/map will be stored in the original folder)
#' @export

pedmap.to.phasedbeaglevcf <- function(ped_path=NULL, map_path=NULL, vcf_path=NULL, beagle_jar="/home/nha/beagle.03Jul18.40b.jar", plink_dir="/home/nha/Plink/plink", db_dir="/home/nha/Plink/DB/"){

  if(length(vcf_path)>0){

    plink_code <- paste0(plink_dir," --vcf ",vcf_path," --dog --recode --double-id --out ",db_dir,"temp_plink")
    system(plink_code)
    map_path <- paste0(db_dir,"temp_plink.map")
    ped_path <- paste0(db_dir,"temp_plink.ped")
  }

  map <- utils::read.table(map_path)

  ## Check if the dataset contains Markers with same ID
  remove_dup <- which(duplicated(map[,2]))
  if(length(remove_dup)>0){
    map <- map[-remove_dup,]
  }
  if(length(remove_dup)>0){
    cat(paste0("Removed ",changes, " markers from the set to avoid markers with the same name (duplicates?!).\n"))
  }


  ## Check if the dataset contains markers on same bp
  changes <- 0
  index <- 1
  change_posi <- which(duplicated(map[,c(1,4)]))

  while(index==1 || length(change_posi)>0){
    map[change_posi,4] <- map[change_posi,4] +1
    changes <- changes + length(change_posi)
    change_posi <- which(duplicated(map[,c(1,4)]))
    index <- index + 1
  }

  if(changes>0){
    cat(paste0("Increased bp of ",changes, "markers to avoid multiple Markers on same bp.\n"))
  }


  ## Write new map/ped files if necessary
  if(changes>0 || length(remove_dup)>0){
    new_map_name <- paste0(substr(map_path, start=1, stop=nchar(map_path)-4),"_reduced.map")
    utils::write.table(file=new_map_name, map, quote=FALSE, row.names = FALSE, col.names=FALSE)
  } else{
    new_map_name <- map_path
  }
  if(length(remove_dup)>0){
    ped <- utils::read.table(ped_path)
    ped <- ped[,-c((remove_dup*2)+5, remove_dup*2 +6)]

    new_ped_name <- paste0(substr(ped_path, start=1, stop=nchar(ped_path)-4),"_reduced.ped")
    utils::write.table(file=new_ped_name, ped, quote=FALSE, row.names = FALSE, col.names=FALSE)

  } else{
    new_ped_name <- ped_path
  }


  # Use Plink to generate a vcf file
  plink_code <- paste0(plink_dir," --noweb --dog --map ",new_map_name," --ped ",new_ped_name," --maf 0.0001 --recode vcf --out " ,db_dir, "temp_vcf")
  system(plink_code)

  # Use Beagle 5.0 to generated phased + imputed dataset
  beagle_code <- paste0("java -jar ",beagle_jar," gt=",db_dir,"temp_vcf.vcf out=",db_dir,"temp_vcf_phased")
  system(beagle_code)
}
```
