## Supplementary material for "MoBPS - Modular Breeding Program Simulator": MoBPS R-package: ssGBLUP.R

```
'#
  Authors
Torsten Pook,

Copyright (C) 2017 -- 2018  Torsten Pook

This program is free software; you can redistribute it and/or
modify it under the terms of the GNU General Public License
as published by the Free Software Foundation; either version 3
of the License, or (at your option) any later version.

This program is distributed in the hope that it will be useful,
but WITHOUT ANY WARRANTY; without even the implied warranty of
MERCHANTABILITY or FITNESS FOR A PARTICULAR PURPOSE.  See the
GNU General Public License for more details.

You should have received a copy of the GNU General Public License
along with this program; if not, write to the Free Software
Foundation, Inc., 59 Temple Place - Suite 330, Boston, MA  02111-1307, USA.
'#


#' ssGBLUP
#'
#' Function to perform single step GBLUP
#' @param A11 A11
#' @param A12 A12
#' @param A22 A22
#' @param G G
#' @export

ssGBLUP <- function(A11, A12, A22, G){


#  A <- cbind(rbind(A11,A21), rbind(A12,A22))
#  I <- diag(nrow(A22))
#  Front <- rbind(A12 %*% solve(A22) %*% I, I)
#  Back <- rbind(solve(A22) %*% A21 %*% I, I)
#  H <- A + Front %*% (G - A22) %*% Back # EXTREMELY INEFFICIENT + you should avoid MASS::ginv()

  # Legarra 2014 Single Step, A general Approach For Genomic Selection
  # A21 <- t(A12)
  # H11 <- A11 - A12 %*% A22inv %*% A21 + A12 %*% A22inv %*% G %*% A22inv %*% A21
  # H12 <- A12 %*% A22inv %*% G
  # H22 <- G
  # H1 <- rbind(cbind(H11,H12), cbind(t(H12), H22))

  A22inv <- solve(A22)
  A21 <- t(A12)
  A12A22inv <- A12 %*% A22inv
  H12 <- A12A22inv %*% G
  H11 <- A11 -  A12A22inv %*% A21 + H12 %*% t(A12A22inv)
  H22 <- G

  H <- rbind(cbind(H11,H12), cbind(t(H12), H22))

  return(H)
}
```
