## Supplementary material for "MoBPS - Modular Breeding Program Simulator": MoBPS R-package: summary.population.R

```
'#
  Authors
Torsten Pook,

Copyright (C) 2017 -- 2018  Torsten Pook

This program is free software; you can redistribute it and/or
modify it under the terms of the GNU General Public License
as published by the Free Software Foundation; either version 3
of the License, or (at your option) any later version.

This program is distributed in the hope that it will be useful,
but WITHOUT ANY WARRANTY; without even the implied warranty of
MERCHANTABILITY or FITNESS FOR A PARTICULAR PURPOSE.  See the
GNU General Public License for more details.

You should have received a copy of the GNU General Public License
along with this program; if not, write to the Free Software
Foundation, Inc., 59 Temple Place - Suite 330, Boston, MA  02111-1307, USA.
'#

#' Summary Population
#'
#' Summary of the population list
#' @param object Population-list
#' @param ... additional arguments affecting the summary produced
#' @return Summary of the population list including number of individuals, genone length and trait overview
#' @export


summary.population <- function(object, ...){
  population <- object
  cat("Population size:\n")
  nindi <- colSums(population$info$size)
  cat(paste0("Total: ", sum(nindi), " Individuals\n"))
  cat(paste0("Of which ", nindi[1], " are male and ", nindi[2], " are female.\n"))
  cat(paste0("There are ", nrow(population$info$size), " generations\n"))
  cat(paste0("and ", nrow(population$info$cohorts), " unique cohorts.\n \n"))

  cat("Genome Info:\n")
  if(population$info$chromosome==1){
    cat(paste0("There is ", population$info$chromosome, " unique chromosome.\n"))
  } else{
    cat(paste0("There are ", population$info$chromosome, " unique chromosomes.\n"))
  }

  cat(paste0("In total there are ", sum(population$info$snp), " SNPs.\n"))
  cat(paste0("The genome has a total length of ", sum(population$info$length), " Morgan.\n"))
  if(length(population$info$bp)==0 || prod(population$info$bp[population$info$cumsnp]==population$info$snp)==1){
    cat(paste0("No physical positions are stored.\n"))
  } else{
    cat(paste0("The genome has a physical size of about: ", round(sum(as.numeric(population$info$bp[population$info$cumsnp]))/1000000000, digits=4), " GB\n"))
  }

  if(population$info$bv.nr>1){
    cat("\nTrait Info:\n")
    cat(paste0("There are ", population$info$bv.nr, " modelled traits.\n"))
    cat(paste0("Of which ", population$info$bv.calc, " have underlying QTL.\n"))
    if(population$info$bv.nr>0){
      cat("Trait names are:")
      cat(population$info$trait.name)
      cat("\n")
      temp1 <- abs(population$info$bv.correlation)
      diag(temp1) <- 0
      if(sum(temp1)==0){
        cat("Genetics of traits are uncorrelated. \n")
      } else{
        cat(paste0("Highest correlation between genetics of traits is ", max(temp1)))
      }
      temp1 <- population$info$pheno.correlation %*% t(population$info$pheno.correlation)
      diag(temp1) <- 0
      if(sum(temp1)==0){
        cat("There are no interactions between enviromental effects.\n")
      } else{
        cat(paste0("Highest correlation between enviromental effects is ", max(temp1)))
      }

    }
  } else{
    cat("\nTrait Info:\n")
    cat(paste0("There is ", population$info$bv.nr, " modelled trait.\n"))
    if(population$info$bv.calc>0){
      cat(paste0("The trait has underlying QTL\n"))
    }

    if(population$info$bv.nr>0){
      cat("The trait is named: ")
      cat(population$info$trait.name)
      cat("\n")

    }
  }


}
```
