## Supplementary material for "MoBPS - Modular Breeding Program Simulator": MoBPS R-package: vlist.R

```
'#
  Authors
Torsten Pook,

Copyright (C) 2017 -- 2018  Torsten Pook

This program is free software; you can redistribute it and/or
modify it under the terms of the GNU General Public License
as published by the Free Software Foundation; either version 3
of the License, or (at your option) any later version.

This program is distributed in the hope that it will be useful,
but WITHOUT ANY WARRANTY; without even the implied warranty of
MERCHANTABILITY or FITNESS FOR A PARTICULAR PURPOSE.  See the
GNU General Public License for more details.

You should have received a copy of the GNU General Public License
along with this program; if not, write to the Free Software
Foundation, Inc., 59 Temple Place - Suite 330, Boston, MA  02111-1307, USA.
'#

#' Derive class
#'
#' Function to devide the class for each individual
#' @param list list you want to print details of
#' @param skip Skip first that many list-elements
#' @param first Only display first that many list-elements
#' @param select Display only selected list-elements
#' @export

vlist <- function(list, skip=NULL, first=NULL, select=NULL){
  if(length(skip)==0 && length(first)==0 && length(select)==0){
    skip <- 2
  }
  total <- list()
  if(length(skip)==1){
    for(index in (skip+1):length(list)){
      print(list[[index]])
      total[[length(total)+1]] <- list[[index]]
    }
  }
  if(length(first)==1){
    for(index in 1:first){
      print(list[[index]])
      total[[length(total)+1]] <- list[[index]]
    }
  }
  if(length(select)>0){
    for(index in select){
      print(list[[index]])
      total[[length(total)+1]] <- list[[index]]
    }
  }

}
```
