## Supplementary material for "MoBPS - Modular Breeding Program Simulator": miraculix R-package: configure

#! /bin/sh
# Guess values for system-dependent variables and create Makefiles.
# Generated by GNU Autoconf 2.69 for miraculix 0.9.
#
#
# Copyright (C) 1992-1996, 1998-2012 Free Software Foundation, Inc.
#
#
# This configure script is free software; the Free Software Foundation
# gives unlimited permission to copy, distribute and modify it.
## -------------------- ##
## M4sh Initialization. ##
## -------------------- ##

# Be more Bourne compatible
DUALCASE=1; export DUALCASE # for MKS sh
if test -n "${ZSH_VERSION+set}" && (emulate sh) >/dev/null 2>&1; then :
  emulate sh
  NULLCMD=:
  # Pre-4.2 versions of Zsh do word splitting on ${1+"$@"}, which
  # is contrary to our usage.  Disable this feature.
  alias -g '${1+"$@"}'='"$@"'
  setopt NO_GLOB_SUBST
else
  case `(set -o) 2>/dev/null` in #(
  *posix*) :
    set -o posix ;; #(
  *) :
     ;;
esac
fi


as_nl='
'
export as_nl
# Printing a long string crashes Solaris 7 /usr/bin/printf.
as_echo='\\\\\\\\\\\\\\\\\\\\\\\\\\\\\\\\\\\\\\\\\\\\\\\\\\\\\\\\\\\\\\\\\\\\\\\\\\\\\\\\\\\\\\\\\\\\\\\\\\\\\\\'
as_echo=$as_echo$as_echo$as_echo$as_echo$as_echo
as_echo=$as_echo$as_echo$as_echo$as_echo$as_echo$as_echo
# Prefer a ksh shell builtin over an external printf program on Solaris,
# but without wasting forks for bash or zsh.
if test -z "$BASH_VERSION$ZSH_VERSION" \
    && (test "X`print -r -- $as_echo`" = "X$as_echo") 2>/dev/null; then
  as_echo='print -r --'
  as_echo_n='print -rn --'
elif (test "X`printf %s $as_echo`" = "X$as_echo") 2>/dev/null; then
  as_echo='printf %s\n'
  as_echo_n='printf %s'
else
  if test "X`(/usr/ucb/echo -n -n $as_echo) 2>/dev/null`" = "X-n $as_echo"; then
    as_echo_body='eval /usr/ucb/echo -n "$1$as_nl"'
    as_echo_n='/usr/ucb/echo -n'
  else
    as_echo_body='eval expr "X$1" : "X\\(.*\\)"'
    as_echo_n_body='eval
      arg=$1;
      case $arg in #(
      *"$as_nl"*)
	expr "X$arg" : "X\\(.*\\)$as_nl";
	arg=`expr "X$arg" : ".*$as_nl\\(.*\\)"`;;
      esac;
      expr "X$arg" : "X\\(.*\\)" | tr -d "$as_nl"
    '
    export as_echo_n_body
    as_echo_n='sh -c $as_echo_n_body as_echo'
  fi
  export as_echo_body
  as_echo='sh -c $as_echo_body as_echo'
fi

# The user is always right.
if test "${PATH_SEPARATOR+set}" != set; then
  PATH_SEPARATOR=:
  (PATH='/bin;/bin'; FPATH=$PATH; sh -c :) >/dev/null 2>&1 && {
    (PATH='/bin:/bin'; FPATH=$PATH; sh -c :) >/dev/null 2>&1 ||
      PATH_SEPARATOR=';'
  }
fi


# IFS
# We need space, tab and new line, in precisely that order.  Quoting is
# there to prevent editors from complaining about space-tab.
# (If _AS_PATH_WALK were called with IFS unset, it would disable word
# splitting by setting IFS to empty value.)
IFS=" ""	$as_nl"

# Find who we are.  Look in the path if we contain no directory separator.
as_myself=
case $0 in #((
  *[\\/]* ) as_myself=$0 ;;
  *) as_save_IFS=$IFS; IFS=$PATH_SEPARATOR
for as_dir in $PATH
do
  IFS=$as_save_IFS
  test -z "$as_dir" && as_dir=.
    test -r "$as_dir/$0" && as_myself=$as_dir/$0 && break
  done
IFS=$as_save_IFS

     ;;
esac
# We did not find ourselves, most probably we were run as `sh COMMAND'
# in which case we are not to be found in the path.
if test "x$as_myself" = x; then
  as_myself=$0
fi
if test ! -f "$as_myself"; then
  $as_echo "$as_myself: error: cannot find myself; rerun with an absolute file name" >&2
  exit 1
fi

# Unset variables that we do not need and which cause bugs (e.g. in
# pre-3.0 UWIN ksh).  But do not cause bugs in bash 2.01; the "|| exit 1"
# suppresses any "Segmentation fault" message there.  '((' could
# trigger a bug in pdksh 5.2.14.
for as_var in BASH_ENV ENV MAIL MAILPATH
do eval test x\${$as_var+set} = xset \
  && ( (unset $as_var) || exit 1) >/dev/null 2>&1 && unset $as_var || :
done
PS1='$ '
PS2='> '
PS4='+ '

# NLS nuisances.
LC_ALL=C
export LC_ALL
LANGUAGE=C
export LANGUAGE

# CDPATH.
(unset CDPATH) >/dev/null 2>&1 && unset CDPATH

# Use a proper internal environment variable to ensure we don't fall
  # into an infinite loop, continuously re-executing ourselves.
  if test x"${_as_can_reexec}" != xno && test "x$CONFIG_SHELL" != x; then
    _as_can_reexec=no; export _as_can_reexec;
    # We cannot yet assume a decent shell, so we have to provide a
# neutralization value for shells without unset; and this also
# works around shells that cannot unset nonexistent variables.
# Preserve -v and -x to the replacement shell.
BASH_ENV=/dev/null
ENV=/dev/null
(unset BASH_ENV) >/dev/null 2>&1 && unset BASH_ENV ENV
case $- in # ((((
  *v*x* | *x*v* ) as_opts=-vx ;;
  *v* ) as_opts=-v ;;
  *x* ) as_opts=-x ;;
  * ) as_opts= ;;
esac
exec $CONFIG_SHELL $as_opts "$as_myself" ${1+"$@"}
# Admittedly, this is quite paranoid, since all the known shells bail
# out after a failed `exec'.
$as_echo "$0: could not re-execute with $CONFIG_SHELL" >&2
as_fn_exit 255
  fi
  # We don't want this to propagate to other subprocesses.
          { _as_can_reexec=; unset _as_can_reexec;}
if test "x$CONFIG_SHELL" = x; then
  as_bourne_compatible="if test -n \"\${ZSH_VERSION+set}\" && (emulate sh) >/dev/null 2>&1; then :
  emulate sh
  NULLCMD=:
  # Pre-4.2 versions of Zsh do word splitting on \${1+\"\$@\"}, which
  # is contrary to our usage.  Disable this feature.
  alias -g '\${1+\"\$@\"}'='\"\$@\"'
  setopt NO_GLOB_SUBST
else
  case \`(set -o) 2>/dev/null\` in #(
  *posix*) :
    set -o posix ;; #(
  *) :
     ;;
esac
fi
"
  as_required="as_fn_return () { (exit \$1); }
as_fn_success () { as_fn_return 0; }
as_fn_failure () { as_fn_return 1; }
as_fn_ret_success () { return 0; }
as_fn_ret_failure () { return 1; }

exitcode=0
as_fn_success || { exitcode=1; echo as_fn_success failed.; }
as_fn_failure && { exitcode=1; echo as_fn_failure succeeded.; }
as_fn_ret_success || { exitcode=1; echo as_fn_ret_success failed.; }
as_fn_ret_failure && { exitcode=1; echo as_fn_ret_failure succeeded.; }
if ( set x; as_fn_ret_success y && test x = \"\$1\" ); then :

else
  exitcode=1; echo positional parameters were not saved.
fi
test x\$exitcode = x0 || exit 1
test -x / || exit 1"
  as_suggested="  as_lineno_1=";as_suggested=$as_suggested$LINENO;as_suggested=$as_suggested" as_lineno_1a=\$LINENO
  as_lineno_2=";as_suggested=$as_suggested$LINENO;as_suggested=$as_suggested" as_lineno_2a=\$LINENO
  eval 'test \"x\$as_lineno_1'\$as_run'\" != \"x\$as_lineno_2'\$as_run'\" &&
  test \"x\`expr \$as_lineno_1'\$as_run' + 1\`\" = \"x\$as_lineno_2'\$as_run'\"' || exit 1"
  if (eval "$as_required") 2>/dev/null; then :
  as_have_required=yes
else
  as_have_required=no
fi
  if test x$as_have_required = xyes && (eval "$as_suggested") 2>/dev/null; then :

else
  as_save_IFS=$IFS; IFS=$PATH_SEPARATOR
as_found=false
for as_dir in /bin$PATH_SEPARATOR/usr/bin$PATH_SEPARATOR$PATH
do
  IFS=$as_save_IFS
  test -z "$as_dir" && as_dir=.
  as_found=:
  case $as_dir in #(
	 /*)
	   for as_base in sh bash ksh sh5; do
	     # Try only shells that exist, to save several forks.
	     as_shell=$as_dir/$as_base
	     if { test -f "$as_shell" || test -f "$as_shell.exe"; } &&
		    { $as_echo "$as_bourne_compatible""$as_required" | as_run=a "$as_shell"; } 2>/dev/null; then :
  CONFIG_SHELL=$as_shell as_have_required=yes
		   if { $as_echo "$as_bourne_compatible""$as_suggested" | as_run=a "$as_shell"; } 2>/dev/null; then :
  break 2
fi
fi
	   done;;
       esac
  as_found=false
done
$as_found || { if { test -f "$SHELL" || test -f "$SHELL.exe"; } &&
	      { $as_echo "$as_bourne_compatible""$as_required" | as_run=a "$SHELL"; } 2>/dev/null; then :
  CONFIG_SHELL=$SHELL as_have_required=yes
fi; }
IFS=$as_save_IFS


      if test "x$CONFIG_SHELL" != x; then :
  export CONFIG_SHELL
             # We cannot yet assume a decent shell, so we have to provide a
# neutralization value for shells without unset; and this also
# works around shells that cannot unset nonexistent variables.
# Preserve -v and -x to the replacement shell.
BASH_ENV=/dev/null
ENV=/dev/null
(unset BASH_ENV) >/dev/null 2>&1 && unset BASH_ENV ENV
case $- in # ((((
  *v*x* | *x*v* ) as_opts=-vx ;;
  *v* ) as_opts=-v ;;
  *x* ) as_opts=-x ;;
  * ) as_opts= ;;
esac
exec $CONFIG_SHELL $as_opts "$as_myself" ${1+"$@"}
# Admittedly, this is quite paranoid, since all the known shells bail
# out after a failed `exec'.
$as_echo "$0: could not re-execute with $CONFIG_SHELL" >&2
exit 255
fi

    if test x$as_have_required = xno; then :
  $as_echo "$0: This script requires a shell more modern than all"
  $as_echo "$0: the shells that I found on your system."
  if test x${ZSH_VERSION+set} = xset ; then
    $as_echo "$0: In particular, zsh $ZSH_VERSION has bugs and should"
    $as_echo "$0: be upgraded to zsh 4.3.4 or later."
  else
    $as_echo "$0: Please tell about your system,
$0: including any error possibly output before this
$0: message. Then install a modern shell, or manually run
$0: the script under such a shell if you do have one."
  fi
  exit 1
fi
fi
fi
SHELL=${CONFIG_SHELL-/bin/sh}
export SHELL
# Unset more variables known to interfere with behavior of common tools.
CLICOLOR_FORCE= GREP_OPTIONS=
unset CLICOLOR_FORCE GREP_OPTIONS

## --------------------- ##
## M4sh Shell Functions. ##
## --------------------- ##
# as_fn_unset VAR
# ---------------
# Portably unset VAR.
as_fn_unset ()
{
  { eval $1=; unset $1;}
}
as_unset=as_fn_unset

# as_fn_set_status STATUS
# -----------------------
# Set $? to STATUS, without forking.
as_fn_set_status ()
{
  return $1
} # as_fn_set_status

# as_fn_exit STATUS
# -----------------
# Exit the shell with STATUS, even in a "trap 0" or "set -e" context.
as_fn_exit ()
{
  set +e
  as_fn_set_status $1
  exit $1
} # as_fn_exit

# as_fn_mkdir_p
# -------------
# Create "$as_dir" as a directory, including parents if necessary.
as_fn_mkdir_p ()
{

  case $as_dir in #(
  -*) as_dir=./$as_dir;;
  esac
  test -d "$as_dir" || eval $as_mkdir_p || {
    as_dirs=
    while :; do
      case $as_dir in #(
      *\'*) as_qdir=`$as_echo "$as_dir" | sed "s/'/'\\\\\\\\''/g"`;; #'(
      *) as_qdir=$as_dir;;
      esac
      as_dirs="'$as_qdir' $as_dirs"
      as_dir=`$as_dirname -- "$as_dir" ||
$as_expr X"$as_dir" : 'X\(.*[^/]\)//*[^/][^/]*/*$' \| \
	 X"$as_dir" : 'X\(//\)[^/]' \| \
	 X"$as_dir" : 'X\(//\)$' \| \
	 X"$as_dir" : 'X\(/\)' \| . 2>/dev/null ||
$as_echo X"$as_dir" |
    sed '/^X\(.*[^/]\)\/\/*[^/][^/]*\/*$/{
	    s//\1/
	    q
	  }
	  /^X\(\/\/\)[^/].*/{
	    s//\1/
	    q
	  }
	  /^X\(\/\/\)$/{
	    s//\1/
	    q
	  }
	  /^X\(\/\).*/{
	    s//\1/
	    q
	  }
	  s/.*/./; q'`
      test -d "$as_dir" && break
    done
    test -z "$as_dirs" || eval "mkdir $as_dirs"
  } || test -d "$as_dir" || as_fn_error $? "cannot create directory $as_dir"


} # as_fn_mkdir_p

# as_fn_executable_p FILE
# -----------------------
# Test if FILE is an executable regular file.
as_fn_executable_p ()
{
  test -f "$1" && test -x "$1"
} # as_fn_executable_p
# as_fn_append VAR VALUE
# ----------------------
# Append the text in VALUE to the end of the definition contained in VAR. Take
# advantage of any shell optimizations that allow amortized linear growth over
# repeated appends, instead of the typical quadratic growth present in naive
# implementations.
if (eval "as_var=1; as_var+=2; test x\$as_var = x12") 2>/dev/null; then :
  eval 'as_fn_append ()
  {
    eval $1+=\$2
  }'
else
  as_fn_append ()
  {
    eval $1=\$$1\$2
  }
fi # as_fn_append

# as_fn_arith ARG...
# ------------------
# Perform arithmetic evaluation on the ARGs, and store the result in the
# global $as_val. Take advantage of shells that can avoid forks. The arguments
# must be portable across $(()) and expr.
if (eval "test \$(( 1 + 1 )) = 2") 2>/dev/null; then :
  eval 'as_fn_arith ()
  {
    as_val=$(( $* ))
  }'
else
  as_fn_arith ()
  {
    as_val=`expr "$@" || test $? -eq 1`
  }
fi # as_fn_arith


# as_fn_error STATUS ERROR [LINENO LOG_FD]
# ----------------------------------------
# Output "`basename $0`: error: ERROR" to stderr. If LINENO and LOG_FD are
# provided, also output the error to LOG_FD, referencing LINENO. Then exit the
# script with STATUS, using 1 if that was 0.
as_fn_error ()
{
  as_status=$1; test $as_status -eq 0 && as_status=1
  if test "$4"; then
    as_lineno=${as_lineno-"$3"} as_lineno_stack=as_lineno_stack=$as_lineno_stack
    $as_echo "$as_me:${as_lineno-$LINENO}: error: $2" >&$4
  fi
  $as_echo "$as_me: error: $2" >&2
  as_fn_exit $as_status
} # as_fn_error

if expr a : '\(a\)' >/dev/null 2>&1 &&
   test "X`expr 00001 : '.*\(...\)'`" = X001; then
  as_expr=expr
else
  as_expr=false
fi

if (basename -- /) >/dev/null 2>&1 && test "X`basename -- / 2>&1`" = "X/"; then
  as_basename=basename
else
  as_basename=false
fi

if (as_dir=`dirname -- /` && test "X$as_dir" = X/) >/dev/null 2>&1; then
  as_dirname=dirname
else
  as_dirname=false
fi

as_me=`$as_basename -- "$0" ||
$as_expr X/"$0" : '.*/\([^/][^/]*\)/*$' \| \
	 X"$0" : 'X\(//\)$' \| \
	 X"$0" : 'X\(/\)' \| . 2>/dev/null ||
$as_echo X/"$0" |
    sed '/^.*\/\([^/][^/]*\)\/*$/{
	    s//\1/
	    q
	  }
	  /^X\/\(\/\/\)$/{
	    s//\1/
	    q
	  }
	  /^X\/\(\/\).*/{
	    s//\1/
	    q
	  }
	  s/.*/./; q'`

# Avoid depending upon Character Ranges.
as_cr_letters='abcdefghijklmnopqrstuvwxyz'
as_cr_LETTERS='ABCDEFGHIJKLMNOPQRSTUVWXYZ'
as_cr_Letters=$as_cr_letters$as_cr_LETTERS
as_cr_digits='0123456789'
as_cr_alnum=$as_cr_Letters$as_cr_digits


  as_lineno_1=$LINENO as_lineno_1a=$LINENO
  as_lineno_2=$LINENO as_lineno_2a=$LINENO
  eval 'test "x$as_lineno_1'$as_run'" != "x$as_lineno_2'$as_run'" &&
  test "x`expr $as_lineno_1'$as_run' + 1`" = "x$as_lineno_2'$as_run'"' || {
  # Blame Lee E. McMahon (1931-1989) for sed's syntax.  :-)
  sed -n '
    p
    /[$]LINENO/=
  ' <$as_myself |
    sed '
      s/[$]LINENO.*/&-/
      t lineno
      b
      :lineno
      N
      :loop
      s/[$]LINENO\([^'$as_cr_alnum'_].*\n\)\(.*\)/\2\1\2/
      t loop
      s/-\n.*//
    ' >$as_me.lineno &&
  chmod +x "$as_me.lineno" ||
    { $as_echo "$as_me: error: cannot create $as_me.lineno; rerun with a POSIX shell" >&2; as_fn_exit 1; }

  # If we had to re-execute with $CONFIG_SHELL, we're ensured to have
  # already done that, so ensure we don't try to do so again and fall
  # in an infinite loop.  This has already happened in practice.
  _as_can_reexec=no; export _as_can_reexec
  # Don't try to exec as it changes $[0], causing all sort of problems
  # (the dirname of $[0] is not the place where we might find the
  # original and so on.  Autoconf is especially sensitive to this).
  . "./$as_me.lineno"
  # Exit status is that of the last command.
  exit
}

ECHO_C= ECHO_N= ECHO_T=
case `echo -n x` in #(((((
-n*)
  case `echo 'xy\c'` in
  *c*) ECHO_T='	';;	# ECHO_T is single tab character.
  xy)  ECHO_C='\c';;
  *)   echo `echo ksh88 bug on AIX 6.1` > /dev/null
       ECHO_T='	';;
  esac;;
*)
  ECHO_N='-n';;
esac

rm -f conf$$ conf$$.exe conf$$.file
if test -d conf$$.dir; then
  rm -f conf$$.dir/conf$$.file
else
  rm -f conf$$.dir
  mkdir conf$$.dir 2>/dev/null
fi
if (echo >conf$$.file) 2>/dev/null; then
  if ln -s conf$$.file conf$$ 2>/dev/null; then
    as_ln_s='ln -s'
    # ... but there are two gotchas:
    # 1) On MSYS, both `ln -s file dir' and `ln file dir' fail.
    # 2) DJGPP < 2.04 has no symlinks; `ln -s' creates a wrapper executable.
    # In both cases, we have to default to `cp -pR'.
    ln -s conf$$.file conf$$.dir 2>/dev/null && test ! -f conf$$.exe ||
      as_ln_s='cp -pR'
  elif ln conf$$.file conf$$ 2>/dev/null; then
    as_ln_s=ln
  else
    as_ln_s='cp -pR'
  fi
else
  as_ln_s='cp -pR'
fi
rm -f conf$$ conf$$.exe conf$$.dir/conf$$.file conf$$.file
rmdir conf$$.dir 2>/dev/null

if mkdir -p . 2>/dev/null; then
  as_mkdir_p='mkdir -p "$as_dir"'
else
  test -d ./-p && rmdir ./-p
  as_mkdir_p=false
fi

as_test_x='test -x'
as_executable_p=as_fn_executable_p

# Sed expression to map a string onto a valid CPP name.
as_tr_cpp="eval sed 'y%*$as_cr_letters%P$as_cr_LETTERS%;s%[^_$as_cr_alnum]%_%g'"

# Sed expression to map a string onto a valid variable name.
as_tr_sh="eval sed 'y%*+%pp%;s%[^_$as_cr_alnum]%_%g'"


test -n "$DJDIR" || exec 7<&0 </dev/null
exec 6>&1

# Name of the host.
# hostname on some systems (SVR3.2, old GNU/Linux) returns a bogus exit status,
# so uname gets run too.
ac_hostname=`(hostname || uname -n) 2>/dev/null | sed 1q`

#
# Initializations.
#
ac_default_prefix=/usr/local
ac_clean_files=
ac_config_libobj_dir=.
LIBOBJS=
cross_compiling=no
subdirs=
MFLAGS=
MAKEFLAGS=

# Identity of this package.
PACKAGE_NAME='miraculix'
PACKAGE_TARNAME='miraculix'
PACKAGE_VERSION='0.9'
PACKAGE_STRING='miraculix 0.9'
PACKAGE_BUGREPORT=''
PACKAGE_URL=''

ac_subst_vars='LTLIBOBJS
LIBOBJS
MIRACULIX_CXXFLAGS
OBJEXT
EXEEXT
ac_ct_CXX
CPPFLAGS
LDFLAGS
CXXFLAGS
CXX
target_alias
host_alias
build_alias
LIBS
ECHO_T
ECHO_N
ECHO_C
DEFS
mandir
localedir
libdir
psdir
pdfdir
dvidir
htmldir
infodir
docdir
oldincludedir
includedir
runstatedir
localstatedir
sharedstatedir
sysconfdir
datadir
datarootdir
libexecdir
sbindir
bindir
program_transform_name
prefix
exec_prefix
PACKAGE_URL
PACKAGE_BUGREPORT
PACKAGE_STRING
PACKAGE_VERSION
PACKAGE_TARNAME
PACKAGE_NAME
PATH_SEPARATOR
SHELL'
ac_subst_files=''
ac_user_opts='
enable_option_checking
'
      ac_precious_vars='build_alias
host_alias
target_alias
CXX
CXXFLAGS
LDFLAGS
LIBS
CPPFLAGS
CCC'


# Initialize some variables set by options.
ac_init_help=
ac_init_version=false
ac_unrecognized_opts=
ac_unrecognized_sep=
# The variables have the same names as the options, with
# dashes changed to underlines.
cache_file=/dev/null
exec_prefix=NONE
no_create=
no_recursion=
prefix=NONE
program_prefix=NONE
program_suffix=NONE
program_transform_name=s,x,x,
silent=
site=
srcdir=
verbose=
x_includes=NONE
x_libraries=NONE

# Installation directory options.
# These are left unexpanded so users can "make install exec_prefix=/foo"
# and all the variables that are supposed to be based on exec_prefix
# by default will actually change.
# Use braces instead of parens because sh, perl, etc. also accept them.
# (The list follows the same order as the GNU Coding Standards.)
bindir='${exec_prefix}/bin'
sbindir='${exec_prefix}/sbin'
libexecdir='${exec_prefix}/libexec'
datarootdir='${prefix}/share'
datadir='${datarootdir}'
sysconfdir='${prefix}/etc'
sharedstatedir='${prefix}/com'
localstatedir='${prefix}/var'
runstatedir='${localstatedir}/run'
includedir='${prefix}/include'
oldincludedir='/usr/include'
docdir='${datarootdir}/doc/${PACKAGE_TARNAME}'
infodir='${datarootdir}/info'
htmldir='${docdir}'
dvidir='${docdir}'
pdfdir='${docdir}'
psdir='${docdir}'
libdir='${exec_prefix}/lib'
localedir='${datarootdir}/locale'
mandir='${datarootdir}/man'

ac_prev=
ac_dashdash=
for ac_option
do
  # If the previous option needs an argument, assign it.
  if test -n "$ac_prev"; then
    eval $ac_prev=\$ac_option
    ac_prev=
    continue
  fi

  case $ac_option in
  *=?*) ac_optarg=`expr "X$ac_option" : '[^=]*=\(.*\)'` ;;
  *=)   ac_optarg= ;;
  *)    ac_optarg=yes ;;
  esac

  # Accept the important Cygnus configure options, so we can diagnose typos.

  case $ac_dashdash$ac_option in
  --)
    ac_dashdash=yes ;;

  -bindir | --bindir | --bindi | --bind | --bin | --bi)
    ac_prev=bindir ;;
  -bindir=* | --bindir=* | --bindi=* | --bind=* | --bin=* | --bi=*)
    bindir=$ac_optarg ;;

  -build | --build | --buil | --bui | --bu)
    ac_prev=build_alias ;;
  -build=* | --build=* | --buil=* | --bui=* | --bu=*)
    build_alias=$ac_optarg ;;

  -cache-file | --cache-file | --cache-fil | --cache-fi \
  | --cache-f | --cache- | --cache | --cach | --cac | --ca | --c)
    ac_prev=cache_file ;;
  -cache-file=* | --cache-file=* | --cache-fil=* | --cache-fi=* \
  | --cache-f=* | --cache-=* | --cache=* | --cach=* | --cac=* | --ca=* | --c=*)
    cache_file=$ac_optarg ;;

  --config-cache | -C)
    cache_file=config.cache ;;

  -datadir | --datadir | --datadi | --datad)
    ac_prev=datadir ;;
  -datadir=* | --datadir=* | --datadi=* | --datad=*)
    datadir=$ac_optarg ;;

  -datarootdir | --datarootdir | --datarootdi | --datarootd | --dataroot \
  | --dataroo | --dataro | --datar)
    ac_prev=datarootdir ;;
  -datarootdir=* | --datarootdir=* | --datarootdi=* | --datarootd=* \
  | --dataroot=* | --dataroo=* | --dataro=* | --datar=*)
    datarootdir=$ac_optarg ;;

  -disable-* | --disable-*)
    ac_useropt=`expr "x$ac_option" : 'x-*disable-\(.*\)'`
    # Reject names that are not valid shell variable names.
    expr "x$ac_useropt" : ".*[^-+._$as_cr_alnum]" >/dev/null &&
      as_fn_error $? "invalid feature name: $ac_useropt"
    ac_useropt_orig=$ac_useropt
    ac_useropt=`$as_echo "$ac_useropt" | sed 's/[-+.]/_/g'`
    case $ac_user_opts in
      *"
"enable_$ac_useropt"
"*) ;;
      *) ac_unrecognized_opts="$ac_unrecognized_opts$ac_unrecognized_sep--disable-$ac_useropt_orig"
	 ac_unrecognized_sep=', ';;
    esac
    eval enable_$ac_useropt=no ;;

  -docdir | --docdir | --docdi | --doc | --do)
    ac_prev=docdir ;;
  -docdir=* | --docdir=* | --docdi=* | --doc=* | --do=*)
    docdir=$ac_optarg ;;

  -dvidir | --dvidir | --dvidi | --dvid | --dvi | --dv)
    ac_prev=dvidir ;;
  -dvidir=* | --dvidir=* | --dvidi=* | --dvid=* | --dvi=* | --dv=*)
    dvidir=$ac_optarg ;;

  -enable-* | --enable-*)
    ac_useropt=`expr "x$ac_option" : 'x-*enable-\([^=]*\)'`
    # Reject names that are not valid shell variable names.
    expr "x$ac_useropt" : ".*[^-+._$as_cr_alnum]" >/dev/null &&
      as_fn_error $? "invalid feature name: $ac_useropt"
    ac_useropt_orig=$ac_useropt
    ac_useropt=`$as_echo "$ac_useropt" | sed 's/[-+.]/_/g'`
    case $ac_user_opts in
      *"
"enable_$ac_useropt"
"*) ;;
      *) ac_unrecognized_opts="$ac_unrecognized_opts$ac_unrecognized_sep--enable-$ac_useropt_orig"
	 ac_unrecognized_sep=', ';;
    esac
    eval enable_$ac_useropt=\$ac_optarg ;;

  -exec-prefix | --exec_prefix | --exec-prefix | --exec-prefi \
  | --exec-pref | --exec-pre | --exec-pr | --exec-p | --exec- \
  | --exec | --exe | --ex)
    ac_prev=exec_prefix ;;
  -exec-prefix=* | --exec_prefix=* | --exec-prefix=* | --exec-prefi=* \
  | --exec-pref=* | --exec-pre=* | --exec-pr=* | --exec-p=* | --exec-=* \
  | --exec=* | --exe=* | --ex=*)
    exec_prefix=$ac_optarg ;;

  -gas | --gas | --ga | --g)
    # Obsolete; use --with-gas.
    with_gas=yes ;;

  -help | --help | --hel | --he | -h)
    ac_init_help=long ;;
  -help=r* | --help=r* | --hel=r* | --he=r* | -hr*)
    ac_init_help=recursive ;;
  -help=s* | --help=s* | --hel=s* | --he=s* | -hs*)
    ac_init_help=short ;;

  -host | --host | --hos | --ho)
    ac_prev=host_alias ;;
  -host=* | --host=* | --hos=* | --ho=*)
    host_alias=$ac_optarg ;;

  -htmldir | --htmldir | --htmldi | --htmld | --html | --htm | --ht)
    ac_prev=htmldir ;;
  -htmldir=* | --htmldir=* | --htmldi=* | --htmld=* | --html=* | --htm=* \
  | --ht=*)
    htmldir=$ac_optarg ;;

  -includedir | --includedir | --includedi | --included | --include \
  | --includ | --inclu | --incl | --inc)
    ac_prev=includedir ;;
  -includedir=* | --includedir=* | --includedi=* | --included=* | --include=* \
  | --includ=* | --inclu=* | --incl=* | --inc=*)
    includedir=$ac_optarg ;;

  -infodir | --infodir | --infodi | --infod | --info | --inf)
    ac_prev=infodir ;;
  -infodir=* | --infodir=* | --infodi=* | --infod=* | --info=* | --inf=*)
    infodir=$ac_optarg ;;

  -libdir | --libdir | --libdi | --libd)
    ac_prev=libdir ;;
  -libdir=* | --libdir=* | --libdi=* | --libd=*)
    libdir=$ac_optarg ;;

  -libexecdir | --libexecdir | --libexecdi | --libexecd | --libexec \
  | --libexe | --libex | --libe)
    ac_prev=libexecdir ;;
  -libexecdir=* | --libexecdir=* | --libexecdi=* | --libexecd=* | --libexec=* \
  | --libexe=* | --libex=* | --libe=*)
    libexecdir=$ac_optarg ;;

  -localedir | --localedir | --localedi | --localed | --locale)
    ac_prev=localedir ;;
  -localedir=* | --localedir=* | --localedi=* | --localed=* | --locale=*)
    localedir=$ac_optarg ;;

  -localstatedir | --localstatedir | --localstatedi | --localstated \
  | --localstate | --localstat | --localsta | --localst | --locals)
    ac_prev=localstatedir ;;
  -localstatedir=* | --localstatedir=* | --localstatedi=* | --localstated=* \
  | --localstate=* | --localstat=* | --localsta=* | --localst=* | --locals=*)
    localstatedir=$ac_optarg ;;

  -mandir | --mandir | --mandi | --mand | --man | --ma | --m)
    ac_prev=mandir ;;
  -mandir=* | --mandir=* | --mandi=* | --mand=* | --man=* | --ma=* | --m=*)
    mandir=$ac_optarg ;;

  -nfp | --nfp | --nf)
    # Obsolete; use --without-fp.
    with_fp=no ;;

  -no-create | --no-create | --no-creat | --no-crea | --no-cre \
  | --no-cr | --no-c | -n)
    no_create=yes ;;

  -no-recursion | --no-recursion | --no-recursio | --no-recursi \
  | --no-recurs | --no-recur | --no-recu | --no-rec | --no-re | --no-r)
    no_recursion=yes ;;

  -oldincludedir | --oldincludedir | --oldincludedi | --oldincluded \
  | --oldinclude | --oldinclud | --oldinclu | --oldincl | --oldinc \
  | --oldin | --oldi | --old | --ol | --o)
    ac_prev=oldincludedir ;;
  -oldincludedir=* | --oldincludedir=* | --oldincludedi=* | --oldincluded=* \
  | --oldinclude=* | --oldinclud=* | --oldinclu=* | --oldincl=* | --oldinc=* \
  | --oldin=* | --oldi=* | --old=* | --ol=* | --o=*)
    oldincludedir=$ac_optarg ;;

  -prefix | --prefix | --prefi | --pref | --pre | --pr | --p)
    ac_prev=prefix ;;
  -prefix=* | --prefix=* | --prefi=* | --pref=* | --pre=* | --pr=* | --p=*)
    prefix=$ac_optarg ;;

  -program-prefix | --program-prefix | --program-prefi | --program-pref \
  | --program-pre | --program-pr | --program-p)
    ac_prev=program_prefix ;;
  -program-prefix=* | --program-prefix=* | --program-prefi=* \
  | --program-pref=* | --program-pre=* | --program-pr=* | --program-p=*)
    program_prefix=$ac_optarg ;;

  -program-suffix | --program-suffix | --program-suffi | --program-suff \
  | --program-suf | --program-su | --program-s)
    ac_prev=program_suffix ;;
  -program-suffix=* | --program-suffix=* | --program-suffi=* \
  | --program-suff=* | --program-suf=* | --program-su=* | --program-s=*)
    program_suffix=$ac_optarg ;;

  -program-transform-name | --program-transform-name \
  | --program-transform-nam | --program-transform-na \
  | --program-transform-n | --program-transform- \
  | --program-transform | --program-transfor \
  | --program-transfo | --program-transf \
  | --program-trans | --program-tran \
  | --progr-tra | --program-tr | --program-t)
    ac_prev=program_transform_name ;;
  -program-transform-name=* | --program-transform-name=* \
  | --program-transform-nam=* | --program-transform-na=* \
  | --program-transform-n=* | --program-transform-=* \
  | --program-transform=* | --program-transfor=* \
  | --program-transfo=* | --program-transf=* \
  | --program-trans=* | --program-tran=* \
  | --progr-tra=* | --program-tr=* | --program-t=*)
    program_transform_name=$ac_optarg ;;

  -pdfdir | --pdfdir | --pdfdi | --pdfd | --pdf | --pd)
    ac_prev=pdfdir ;;
  -pdfdir=* | --pdfdir=* | --pdfdi=* | --pdfd=* | --pdf=* | --pd=*)
    pdfdir=$ac_optarg ;;

  -psdir | --psdir | --psdi | --psd | --ps)
    ac_prev=psdir ;;
  -psdir=* | --psdir=* | --psdi=* | --psd=* | --ps=*)
    psdir=$ac_optarg ;;

  -q | -quiet | --quiet | --quie | --qui | --qu | --q \
  | -silent | --silent | --silen | --sile | --sil)
    silent=yes ;;

  -runstatedir | --runstatedir | --runstatedi | --runstated \
  | --runstate | --runstat | --runsta | --runst | --runs \
  | --run | --ru | --r)
    ac_prev=runstatedir ;;
  -runstatedir=* | --runstatedir=* | --runstatedi=* | --runstated=* \
  | --runstate=* | --runstat=* | --runsta=* | --runst=* | --runs=* \
  | --run=* | --ru=* | --r=*)
    runstatedir=$ac_optarg ;;

  -sbindir | --sbindir | --sbindi | --sbind | --sbin | --sbi | --sb)
    ac_prev=sbindir ;;
  -sbindir=* | --sbindir=* | --sbindi=* | --sbind=* | --sbin=* \
  | --sbi=* | --sb=*)
    sbindir=$ac_optarg ;;

  -sharedstatedir | --sharedstatedir | --sharedstatedi \
  | --sharedstated | --sharedstate | --sharedstat | --sharedsta \
  | --sharedst | --shareds | --shared | --share | --shar \
  | --sha | --sh)
    ac_prev=sharedstatedir ;;
  -sharedstatedir=* | --sharedstatedir=* | --sharedstatedi=* \
  | --sharedstated=* | --sharedstate=* | --sharedstat=* | --sharedsta=* \
  | --sharedst=* | --shareds=* | --shared=* | --share=* | --shar=* \
  | --sha=* | --sh=*)
    sharedstatedir=$ac_optarg ;;

  -site | --site | --sit)
    ac_prev=site ;;
  -site=* | --site=* | --sit=*)
    site=$ac_optarg ;;

  -srcdir | --srcdir | --srcdi | --srcd | --src | --sr)
    ac_prev=srcdir ;;
  -srcdir=* | --srcdir=* | --srcdi=* | --srcd=* | --src=* | --sr=*)
    srcdir=$ac_optarg ;;

  -sysconfdir | --sysconfdir | --sysconfdi | --sysconfd | --sysconf \
  | --syscon | --sysco | --sysc | --sys | --sy)
    ac_prev=sysconfdir ;;
  -sysconfdir=* | --sysconfdir=* | --sysconfdi=* | --sysconfd=* | --sysconf=* \
  | --syscon=* | --sysco=* | --sysc=* | --sys=* | --sy=*)
    sysconfdir=$ac_optarg ;;

  -target | --target | --targe | --targ | --tar | --ta | --t)
    ac_prev=target_alias ;;
  -target=* | --target=* | --targe=* | --targ=* | --tar=* | --ta=* | --t=*)
    target_alias=$ac_optarg ;;

  -v | -verbose | --verbose | --verbos | --verbo | --verb)
    verbose=yes ;;

  -version | --version | --versio | --versi | --vers | -V)
    ac_init_version=: ;;

  -with-* | --with-*)
    ac_useropt=`expr "x$ac_option" : 'x-*with-\([^=]*\)'`
    # Reject names that are not valid shell variable names.
    expr "x$ac_useropt" : ".*[^-+._$as_cr_alnum]" >/dev/null &&
      as_fn_error $? "invalid package name: $ac_useropt"
    ac_useropt_orig=$ac_useropt
    ac_useropt=`$as_echo "$ac_useropt" | sed 's/[-+.]/_/g'`
    case $ac_user_opts in
      *"
"with_$ac_useropt"
"*) ;;
      *) ac_unrecognized_opts="$ac_unrecognized_opts$ac_unrecognized_sep--with-$ac_useropt_orig"
	 ac_unrecognized_sep=', ';;
    esac
    eval with_$ac_useropt=\$ac_optarg ;;

  -without-* | --without-*)
    ac_useropt=`expr "x$ac_option" : 'x-*without-\(.*\)'`
    # Reject names that are not valid shell variable names.
    expr "x$ac_useropt" : ".*[^-+._$as_cr_alnum]" >/dev/null &&
      as_fn_error $? "invalid package name: $ac_useropt"
    ac_useropt_orig=$ac_useropt
    ac_useropt=`$as_echo "$ac_useropt" | sed 's/[-+.]/_/g'`
    case $ac_user_opts in
      *"
"with_$ac_useropt"
"*) ;;
      *) ac_unrecognized_opts="$ac_unrecognized_opts$ac_unrecognized_sep--without-$ac_useropt_orig"
	 ac_unrecognized_sep=', ';;
    esac
    eval with_$ac_useropt=no ;;

  --x)
    # Obsolete; use --with-x.
    with_x=yes ;;

  -x-includes | --x-includes | --x-include | --x-includ | --x-inclu \
  | --x-incl | --x-inc | --x-in | --x-i)
    ac_prev=x_includes ;;
  -x-includes=* | --x-includes=* | --x-include=* | --x-includ=* | --x-inclu=* \
  | --x-incl=* | --x-inc=* | --x-in=* | --x-i=*)
    x_includes=$ac_optarg ;;

  -x-libraries | --x-libraries | --x-librarie | --x-librari \
  | --x-librar | --x-libra | --x-libr | --x-lib | --x-li | --x-l)
    ac_prev=x_libraries ;;
  -x-libraries=* | --x-libraries=* | --x-librarie=* | --x-librari=* \
  | --x-librar=* | --x-libra=* | --x-libr=* | --x-lib=* | --x-li=* | --x-l=*)
    x_libraries=$ac_optarg ;;

  -*) as_fn_error $? "unrecognized option: \`$ac_option'
Try \`$0 --help' for more information"
    ;;

  *=*)
    ac_envvar=`expr "x$ac_option" : 'x\([^=]*\)='`
    # Reject names that are not valid shell variable names.
    case $ac_envvar in #(
      '' | [0-9]* | *[!_$as_cr_alnum]* )
      as_fn_error $? "invalid variable name: \`$ac_envvar'" ;;
    esac
    eval $ac_envvar=\$ac_optarg
    export $ac_envvar ;;

  *)
    # FIXME: should be removed in autoconf 3.0.
    $as_echo "$as_me: WARNING: you should use --build, --host, --target" >&2
    expr "x$ac_option" : ".*[^-._$as_cr_alnum]" >/dev/null &&
      $as_echo "$as_me: WARNING: invalid host type: $ac_option" >&2
    : "${build_alias=$ac_option} ${host_alias=$ac_option} ${target_alias=$ac_option}"
    ;;

  esac
done

if test -n "$ac_prev"; then
  ac_option=--`echo $ac_prev | sed 's/_/-/g'`
  as_fn_error $? "missing argument to $ac_option"
fi

if test -n "$ac_unrecognized_opts"; then
  case $enable_option_checking in
    no) ;;
    fatal) as_fn_error $? "unrecognized options: $ac_unrecognized_opts" ;;
    *)     $as_echo "$as_me: WARNING: unrecognized options: $ac_unrecognized_opts" >&2 ;;
  esac
fi

# Check all directory arguments for consistency.
for ac_var in	exec_prefix prefix bindir sbindir libexecdir datarootdir \
		datadir sysconfdir sharedstatedir localstatedir includedir \
		oldincludedir docdir infodir htmldir dvidir pdfdir psdir \
		libdir localedir mandir runstatedir
do
  eval ac_val=\$$ac_var
  # Remove trailing slashes.
  case $ac_val in
    */ )
      ac_val=`expr "X$ac_val" : 'X\(.*[^/]\)' \| "X$ac_val" : 'X\(.*\)'`
      eval $ac_var=\$ac_val;;
  esac
  # Be sure to have absolute directory names.
  case $ac_val in
    [\\/$]* | ?:[\\/]* )  continue;;
    NONE | '' ) case $ac_var in *prefix ) continue;; esac;;
  esac
  as_fn_error $? "expected an absolute directory name for --$ac_var: $ac_val"
done

# There might be people who depend on the old broken behavior: `$host'
# used to hold the argument of --host etc.
# FIXME: To remove some day.
build=$build_alias
host=$host_alias
target=$target_alias

# FIXME: To remove some day.
if test "x$host_alias" != x; then
  if test "x$build_alias" = x; then
    cross_compiling=maybe
  elif test "x$build_alias" != "x$host_alias"; then
    cross_compiling=yes
  fi
fi

ac_tool_prefix=
test -n "$host_alias" && ac_tool_prefix=$host_alias-

test "$silent" = yes && exec 6>/dev/null


ac_pwd=`pwd` && test -n "$ac_pwd" &&
ac_ls_di=`ls -di .` &&
ac_pwd_ls_di=`cd "$ac_pwd" && ls -di .` ||
  as_fn_error $? "working directory cannot be determined"
test "X$ac_ls_di" = "X$ac_pwd_ls_di" ||
  as_fn_error $? "pwd does not report name of working directory"


# Find the source files, if location was not specified.
if test -z "$srcdir"; then
  ac_srcdir_defaulted=yes
  # Try the directory containing this script, then the parent directory.
  ac_confdir=`$as_dirname -- "$as_myself" ||
$as_expr X"$as_myself" : 'X\(.*[^/]\)//*[^/][^/]*/*$' \| \
	 X"$as_myself" : 'X\(//\)[^/]' \| \
	 X"$as_myself" : 'X\(//\)$' \| \
	 X"$as_myself" : 'X\(/\)' \| . 2>/dev/null ||
$as_echo X"$as_myself" |
    sed '/^X\(.*[^/]\)\/\/*[^/][^/]*\/*$/{
	    s//\1/
	    q
	  }
	  /^X\(\/\/\)[^/].*/{
	    s//\1/
	    q
	  }
	  /^X\(\/\/\)$/{
	    s//\1/
	    q
	  }
	  /^X\(\/\).*/{
	    s//\1/
	    q
	  }
	  s/.*/./; q'`
  srcdir=$ac_confdir
  if test ! -r "$srcdir/$ac_unique_file"; then
    srcdir=..
  fi
else
  ac_srcdir_defaulted=no
fi
if test ! -r "$srcdir/$ac_unique_file"; then
  test "$ac_srcdir_defaulted" = yes && srcdir="$ac_confdir or .."
  as_fn_error $? "cannot find sources ($ac_unique_file) in $srcdir"
fi
ac_msg="sources are in $srcdir, but \`cd $srcdir' does not work"
ac_abs_confdir=`(
	cd "$srcdir" && test -r "./$ac_unique_file" || as_fn_error $? "$ac_msg"
	pwd)`
# When building in place, set srcdir=.
if test "$ac_abs_confdir" = "$ac_pwd"; then
  srcdir=.
fi
# Remove unnecessary trailing slashes from srcdir.
# Double slashes in file names in object file debugging info
# mess up M-x gdb in Emacs.
case $srcdir in
*/) srcdir=`expr "X$srcdir" : 'X\(.*[^/]\)' \| "X$srcdir" : 'X\(.*\)'`;;
esac
for ac_var in $ac_precious_vars; do
  eval ac_env_${ac_var}_set=\${${ac_var}+set}
  eval ac_env_${ac_var}_value=\$${ac_var}
  eval ac_cv_env_${ac_var}_set=\${${ac_var}+set}
  eval ac_cv_env_${ac_var}_value=\$${ac_var}
done

#
# Report the --help message.
#
if test "$ac_init_help" = "long"; then
  # Omit some internal or obsolete options to make the list less imposing.
  # This message is too long to be a string in the A/UX 3.1 sh.
  cat <<_ACEOF
\`configure' configures miraculix 0.9 to adapt to many kinds of systems.

Usage: $0 [OPTION]... [VAR=VALUE]...

To assign environment variables (e.g., CC, CFLAGS...), specify them as
VAR=VALUE.  See below for descriptions of some of the useful variables.

Defaults for the options are specified in brackets.

Configuration:
  -h, --help              display this help and exit
      --help=short        display options specific to this package
      --help=recursive    display the short help of all the included packages
  -V, --version           display version information and exit
  -q, --quiet, --silent   do not print \`checking ...' messages
      --cache-file=FILE   cache test results in FILE [disabled]
  -C, --config-cache      alias for \`--cache-file=config.cache'
  -n, --no-create         do not create output files
      --srcdir=DIR        find the sources in DIR [configure dir or \`..']

Installation directories:
  --prefix=PREFIX         install architecture-independent files in PREFIX
                          [$ac_default_prefix]
  --exec-prefix=EPREFIX   install architecture-dependent files in EPREFIX
                          [PREFIX]

By default, \`make install' will install all the files in
\`$ac_default_prefix/bin', \`$ac_default_prefix/lib' etc.  You can specify
an installation prefix other than \`$ac_default_prefix' using \`--prefix',
for instance \`--prefix=\$HOME'.

For better control, use the options below.

Fine tuning of the installation directories:
  --bindir=DIR            user executables [EPREFIX/bin]
  --sbindir=DIR           system admin executables [EPREFIX/sbin]
  --libexecdir=DIR        program executables [EPREFIX/libexec]
  --sysconfdir=DIR        read-only single-machine data [PREFIX/etc]
  --sharedstatedir=DIR    modifiable architecture-independent data [PREFIX/com]
  --localstatedir=DIR     modifiable single-machine data [PREFIX/var]
  --runstatedir=DIR       modifiable per-process data [LOCALSTATEDIR/run]
  --libdir=DIR            object code libraries [EPREFIX/lib]
  --includedir=DIR        C header files [PREFIX/include]
  --oldincludedir=DIR     C header files for non-gcc [/usr/include]
  --datarootdir=DIR       read-only arch.-independent data root [PREFIX/share]
  --datadir=DIR           read-only architecture-independent data [DATAROOTDIR]
  --infodir=DIR           info documentation [DATAROOTDIR/info]
  --localedir=DIR         locale-dependent data [DATAROOTDIR/locale]
  --mandir=DIR            man documentation [DATAROOTDIR/man]
  --docdir=DIR            documentation root [DATAROOTDIR/doc/miraculix]
  --htmldir=DIR           html documentation [DOCDIR]
  --dvidir=DIR            dvi documentation [DOCDIR]
  --pdfdir=DIR            pdf documentation [DOCDIR]
  --psdir=DIR             ps documentation [DOCDIR]
_ACEOF

  cat <<\_ACEOF
_ACEOF
fi

if test -n "$ac_init_help"; then
  case $ac_init_help in
     short | recursive ) echo "Configuration of miraculix 0.9:";;
   esac
  cat <<\_ACEOF

Some influential environment variables:
  CXX         C++ compiler command
  CXXFLAGS    C++ compiler flags
  LDFLAGS     linker flags, e.g. -L<lib dir> if you have libraries in a
              nonstandard directory <lib dir>
  LIBS        libraries to pass to the linker, e.g. -l<library>
  CPPFLAGS    (Objective) C/C++ preprocessor flags, e.g. -I<include dir> if
              you have headers in a nonstandard directory <include dir>

Use these variables to override the choices made by `configure' or to help
it to find libraries and programs with nonstandard names/locations.

Report bugs to the package provider.
_ACEOF
ac_status=$?
fi

if test "$ac_init_help" = "recursive"; then
  # If there are subdirs, report their specific --help.
  for ac_dir in : $ac_subdirs_all; do test "x$ac_dir" = x: && continue
    test -d "$ac_dir" ||
      { cd "$srcdir" && ac_pwd=`pwd` && srcdir=. && test -d "$ac_dir"; } ||
      continue
    ac_builddir=.

case "$ac_dir" in
.) ac_dir_suffix= ac_top_builddir_sub=. ac_top_build_prefix= ;;
*)
  ac_dir_suffix=/`$as_echo "$ac_dir" | sed 's|^\.[\\/]||'`
  # A ".." for each directory in $ac_dir_suffix.
  ac_top_builddir_sub=`$as_echo "$ac_dir_suffix" | sed 's|/[^\\/]*|/..|g;s|/||'`
  case $ac_top_builddir_sub in
  "") ac_top_builddir_sub=. ac_top_build_prefix= ;;
  *)  ac_top_build_prefix=$ac_top_builddir_sub/ ;;
  esac ;;
esac
ac_abs_top_builddir=$ac_pwd
ac_abs_builddir=$ac_pwd$ac_dir_suffix
# for backward compatibility:
ac_top_builddir=$ac_top_build_prefix

case $srcdir in
  .)  # We are building in place.
    ac_srcdir=.
    ac_top_srcdir=$ac_top_builddir_sub
    ac_abs_top_srcdir=$ac_pwd ;;
  [\\/]* | ?:[\\/]* )  # Absolute name.
    ac_srcdir=$srcdir$ac_dir_suffix;
    ac_top_srcdir=$srcdir
    ac_abs_top_srcdir=$srcdir ;;
  *) # Relative name.
    ac_srcdir=$ac_top_build_prefix$srcdir$ac_dir_suffix
    ac_top_srcdir=$ac_top_build_prefix$srcdir
    ac_abs_top_srcdir=$ac_pwd/$srcdir ;;
esac
ac_abs_srcdir=$ac_abs_top_srcdir$ac_dir_suffix

    cd "$ac_dir" || { ac_status=$?; continue; }
    # Check for guested configure.
    if test -f "$ac_srcdir/configure.gnu"; then
      echo &&
      $SHELL "$ac_srcdir/configure.gnu" --help=recursive
    elif test -f "$ac_srcdir/configure"; then
      echo &&
      $SHELL "$ac_srcdir/configure" --help=recursive
    else
      $as_echo "$as_me: WARNING: no configuration information is in $ac_dir" >&2
    fi || ac_status=$?
    cd "$ac_pwd" || { ac_status=$?; break; }
  done
fi

test -n "$ac_init_help" && exit $ac_status
if $ac_init_version; then
  cat <<\_ACEOF
miraculix configure 0.9
generated by GNU Autoconf 2.69

Copyright (C) 2012 Free Software Foundation, Inc.
This configure script is free software; the Free Software Foundation
gives unlimited permission to copy, distribute and modify it.
_ACEOF
  exit
fi

## ------------------------ ##
## Autoconf initialization. ##
## ------------------------ ##

# ac_fn_cxx_try_compile LINENO
# ----------------------------
# Try to compile conftest.$ac_ext, and return whether this succeeded.
ac_fn_cxx_try_compile ()
{
  as_lineno=${as_lineno-"$1"} as_lineno_stack=as_lineno_stack=$as_lineno_stack
  rm -f conftest.$ac_objext
  if { { ac_try="$ac_compile"
case "(($ac_try" in
  *\"* | *\`* | *\\*) ac_try_echo=\$ac_try;;
  *) ac_try_echo=$ac_try;;
esac
eval ac_try_echo="\"\$as_me:${as_lineno-$LINENO}: $ac_try_echo\""
$as_echo "$ac_try_echo"; } >&5
  (eval "$ac_compile") 2>conftest.err
  ac_status=$?
  if test -s conftest.err; then
    grep -v '^ *+' conftest.err >conftest.er1
    cat conftest.er1 >&5
    mv -f conftest.er1 conftest.err
  fi
  $as_echo "$as_me:${as_lineno-$LINENO}: \$? = $ac_status" >&5
  test $ac_status = 0; } && {
	 test -z "$ac_cxx_werror_flag" ||
	 test ! -s conftest.err
       } && test -s conftest.$ac_objext; then :
  ac_retval=0
else
  $as_echo "$as_me: failed program was:" >&5
sed 's/^/| /' conftest.$ac_ext >&5

	ac_retval=1
fi
  eval $as_lineno_stack; ${as_lineno_stack:+:} unset as_lineno
  as_fn_set_status $ac_retval

} # ac_fn_cxx_try_compile
cat >config.log <<_ACEOF
This file contains any messages produced by compilers while
running configure, to aid debugging if configure makes a mistake.

It was created by miraculix $as_me 0.9, which was
generated by GNU Autoconf 2.69.  Invocation command line was

  $ $0 $@

_ACEOF
exec 5>>config.log
{
cat <<_ASUNAME
## --------- ##
## Platform. ##
## --------- ##

hostname = `(hostname || uname -n) 2>/dev/null | sed 1q`
uname -m = `(uname -m) 2>/dev/null || echo unknown`
uname -r = `(uname -r) 2>/dev/null || echo unknown`
uname -s = `(uname -s) 2>/dev/null || echo unknown`
uname -v = `(uname -v) 2>/dev/null || echo unknown`

/usr/bin/uname -p = `(/usr/bin/uname -p) 2>/dev/null || echo unknown`
/bin/uname -X     = `(/bin/uname -X) 2>/dev/null     || echo unknown`

/bin/arch              = `(/bin/arch) 2>/dev/null              || echo unknown`
/usr/bin/arch -k       = `(/usr/bin/arch -k) 2>/dev/null       || echo unknown`
/usr/convex/getsysinfo = `(/usr/convex/getsysinfo) 2>/dev/null || echo unknown`
/usr/bin/hostinfo      = `(/usr/bin/hostinfo) 2>/dev/null      || echo unknown`
/bin/machine           = `(/bin/machine) 2>/dev/null           || echo unknown`
/usr/bin/oslevel       = `(/usr/bin/oslevel) 2>/dev/null       || echo unknown`
/bin/universe          = `(/bin/universe) 2>/dev/null          || echo unknown`

_ASUNAME

as_save_IFS=$IFS; IFS=$PATH_SEPARATOR
for as_dir in $PATH
do
  IFS=$as_save_IFS
  test -z "$as_dir" && as_dir=.
    $as_echo "PATH: $as_dir"
  done
IFS=$as_save_IFS

} >&5

cat >&5 <<_ACEOF


## ----------- ##
## Core tests. ##
## ----------- ##

_ACEOF


# Keep a trace of the command line.
# Strip out --no-create and --no-recursion so they do not pile up.
# Strip out --silent because we don't want to record it for future runs.
# Also quote any args containing shell meta-characters.
# Make two passes to allow for proper duplicate-argument suppression.
ac_configure_args=
ac_configure_args0=
ac_configure_args1=
ac_must_keep_next=false
for ac_pass in 1 2
do
  for ac_arg
  do
    case $ac_arg in
    -no-create | --no-c* | -n | -no-recursion | --no-r*) continue ;;
    -q | -quiet | --quiet | --quie | --qui | --qu | --q \
    | -silent | --silent | --silen | --sile | --sil)
      continue ;;
    *\'*)
      ac_arg=`$as_echo "$ac_arg" | sed "s/'/'\\\\\\\\''/g"` ;;
    esac
    case $ac_pass in
    1) as_fn_append ac_configure_args0 " '$ac_arg'" ;;
    2)
      as_fn_append ac_configure_args1 " '$ac_arg'"
      if test $ac_must_keep_next = true; then
	ac_must_keep_next=false # Got value, back to normal.
      else
	case $ac_arg in
	  *=* | --config-cache | -C | -disable-* | --disable-* \
	  | -enable-* | --enable-* | -gas | --g* | -nfp | --nf* \
	  | -q | -quiet | --q* | -silent | --sil* | -v | -verb* \
	  | -with-* | --with-* | -without-* | --without-* | --x)
	    case "$ac_configure_args0 " in
	      "$ac_configure_args1"*" '$ac_arg' "* ) continue ;;
	    esac
	    ;;
	  -* ) ac_must_keep_next=true ;;
	esac
      fi
      as_fn_append ac_configure_args " '$ac_arg'"
      ;;
    esac
  done
done
{ ac_configure_args0=; unset ac_configure_args0;}
{ ac_configure_args1=; unset ac_configure_args1;}

# When interrupted or exit'd, cleanup temporary files, and complete
# config.log.  We remove comments because anyway the quotes in there
# would cause problems or look ugly.
# WARNING: Use '\'' to represent an apostrophe within the trap.
# WARNING: Do not start the trap code with a newline, due to a FreeBSD 4.0 bug.
trap 'exit_status=$?
  # Save into config.log some information that might help in debugging.
  {
    echo

    $as_echo "## ---------------- ##
## Cache variables. ##
## ---------------- ##"
    echo
    # The following way of writing the cache mishandles newlines in values,
(
  for ac_var in `(set) 2>&1 | sed -n '\''s/^\([a-zA-Z_][a-zA-Z0-9_]*\)=.*/\1/p'\''`; do
    eval ac_val=\$$ac_var
    case $ac_val in #(
    *${as_nl}*)
      case $ac_var in #(
      *_cv_*) { $as_echo "$as_me:${as_lineno-$LINENO}: WARNING: cache variable $ac_var contains a newline" >&5
$as_echo "$as_me: WARNING: cache variable $ac_var contains a newline" >&2;} ;;
      esac
      case $ac_var in #(
      _ | IFS | as_nl) ;; #(
      BASH_ARGV | BASH_SOURCE) eval $ac_var= ;; #(
      *) { eval $ac_var=; unset $ac_var;} ;;
      esac ;;
    esac
  done
  (set) 2>&1 |
    case $as_nl`(ac_space='\'' '\''; set) 2>&1` in #(
    *${as_nl}ac_space=\ *)
      sed -n \
	"s/'\''/'\''\\\\'\'''\''/g;
	  s/^\\([_$as_cr_alnum]*_cv_[_$as_cr_alnum]*\\)=\\(.*\\)/\\1='\''\\2'\''/p"
      ;; #(
    *)
      sed -n "/^[_$as_cr_alnum]*_cv_[_$as_cr_alnum]*=/p"
      ;;
    esac |
    sort
)
    echo

    $as_echo "## ----------------- ##
## Output variables. ##
## ----------------- ##"
    echo
    for ac_var in $ac_subst_vars
    do
      eval ac_val=\$$ac_var
      case $ac_val in
      *\'\''*) ac_val=`$as_echo "$ac_val" | sed "s/'\''/'\''\\\\\\\\'\'''\''/g"`;;
      esac
      $as_echo "$ac_var='\''$ac_val'\''"
    done | sort
    echo

    if test -n "$ac_subst_files"; then
      $as_echo "## ------------------- ##
## File substitutions. ##
## ------------------- ##"
      echo
      for ac_var in $ac_subst_files
      do
	eval ac_val=\$$ac_var
	case $ac_val in
	*\'\''*) ac_val=`$as_echo "$ac_val" | sed "s/'\''/'\''\\\\\\\\'\'''\''/g"`;;
	esac
	$as_echo "$ac_var='\''$ac_val'\''"
      done | sort
      echo
    fi

    if test -s confdefs.h; then
      $as_echo "## ----------- ##
## confdefs.h. ##
## ----------- ##"
      echo
      cat confdefs.h
      echo
    fi
    test "$ac_signal" != 0 &&
      $as_echo "$as_me: caught signal $ac_signal"
    $as_echo "$as_me: exit $exit_status"
  } >&5
  rm -f core *.core core.conftest.* &&
    rm -f -r conftest* confdefs* conf$$* $ac_clean_files &&
    exit $exit_status
' 0
for ac_signal in 1 2 13 15; do
  trap 'ac_signal='$ac_signal'; as_fn_exit 1' $ac_signal
done
ac_signal=0

# confdefs.h avoids OS command line length limits that DEFS can exceed.
rm -f -r conftest* confdefs.h

$as_echo "/* confdefs.h */" > confdefs.h

# Predefined preprocessor variables.

cat >>confdefs.h <<_ACEOF
#define PACKAGE_NAME "$PACKAGE_NAME"
_ACEOF

cat >>confdefs.h <<_ACEOF
#define PACKAGE_TARNAME "$PACKAGE_TARNAME"
_ACEOF

cat >>confdefs.h <<_ACEOF
#define PACKAGE_VERSION "$PACKAGE_VERSION"
_ACEOF

cat >>confdefs.h <<_ACEOF
#define PACKAGE_STRING "$PACKAGE_STRING"
_ACEOF

cat >>confdefs.h <<_ACEOF
#define PACKAGE_BUGREPORT "$PACKAGE_BUGREPORT"
_ACEOF

cat >>confdefs.h <<_ACEOF
#define PACKAGE_URL "$PACKAGE_URL"
_ACEOF


# Let the site file select an alternate cache file if it wants to.
# Prefer an explicitly selected file to automatically selected ones.
ac_site_file1=NONE
ac_site_file2=NONE
if test -n "$CONFIG_SITE"; then
  # We do not want a PATH search for config.site.
  case $CONFIG_SITE in #((
    -*)  ac_site_file1=./$CONFIG_SITE;;
    */*) ac_site_file1=$CONFIG_SITE;;
    *)   ac_site_file1=./$CONFIG_SITE;;
  esac
elif test "x$prefix" != xNONE; then
  ac_site_file1=$prefix/share/config.site
  ac_site_file2=$prefix/etc/config.site
else
  ac_site_file1=$ac_default_prefix/share/config.site
  ac_site_file2=$ac_default_prefix/etc/config.site
fi
for ac_site_file in "$ac_site_file1" "$ac_site_file2"
do
  test "x$ac_site_file" = xNONE && continue
  if test /dev/null != "$ac_site_file" && test -r "$ac_site_file"; then
    { $as_echo "$as_me:${as_lineno-$LINENO}: loading site script $ac_site_file" >&5
$as_echo "$as_me: loading site script $ac_site_file" >&6;}
    sed 's/^/| /' "$ac_site_file" >&5
    . "$ac_site_file" \
      || { { $as_echo "$as_me:${as_lineno-$LINENO}: error: in \`$ac_pwd':" >&5
$as_echo "$as_me: error: in \`$ac_pwd':" >&2;}
as_fn_error $? "failed to load site script $ac_site_file
See \`config.log' for more details" "$LINENO" 5; }
  fi
done

if test -r "$cache_file"; then
  # Some versions of bash will fail to source /dev/null (special files
  # actually), so we avoid doing that.  DJGPP emulates it as a regular file.
  if test /dev/null != "$cache_file" && test -f "$cache_file"; then
    { $as_echo "$as_me:${as_lineno-$LINENO}: loading cache $cache_file" >&5
$as_echo "$as_me: loading cache $cache_file" >&6;}
    case $cache_file in
      [\\/]* | ?:[\\/]* ) . "$cache_file";;
      *)                      . "./$cache_file";;
    esac
  fi
else
  { $as_echo "$as_me:${as_lineno-$LINENO}: creating cache $cache_file" >&5
$as_echo "$as_me: creating cache $cache_file" >&6;}
  >$cache_file
fi

# Check that the precious variables saved in the cache have kept the same
# value.
ac_cache_corrupted=false
for ac_var in $ac_precious_vars; do
  eval ac_old_set=\$ac_cv_env_${ac_var}_set
  eval ac_new_set=\$ac_env_${ac_var}_set
  eval ac_old_val=\$ac_cv_env_${ac_var}_value
  eval ac_new_val=\$ac_env_${ac_var}_value
  case $ac_old_set,$ac_new_set in
    set,)
      { $as_echo "$as_me:${as_lineno-$LINENO}: error: \`$ac_var' was set to \`$ac_old_val' in the previous run" >&5
$as_echo "$as_me: error: \`$ac_var' was set to \`$ac_old_val' in the previous run" >&2;}
      ac_cache_corrupted=: ;;
    ,set)
      { $as_echo "$as_me:${as_lineno-$LINENO}: error: \`$ac_var' was not set in the previous run" >&5
$as_echo "$as_me: error: \`$ac_var' was not set in the previous run" >&2;}
      ac_cache_corrupted=: ;;
    ,);;
    *)
      if test "x$ac_old_val" != "x$ac_new_val"; then
	# differences in whitespace do not lead to failure.
	ac_old_val_w=`echo x $ac_old_val`
	ac_new_val_w=`echo x $ac_new_val`
	if test "$ac_old_val_w" != "$ac_new_val_w"; then
	  { $as_echo "$as_me:${as_lineno-$LINENO}: error: \`$ac_var' has changed since the previous run:" >&5
$as_echo "$as_me: error: \`$ac_var' has changed since the previous run:" >&2;}
	  ac_cache_corrupted=:
	else
	  { $as_echo "$as_me:${as_lineno-$LINENO}: warning: ignoring whitespace changes in \`$ac_var' since the previous run:" >&5
$as_echo "$as_me: warning: ignoring whitespace changes in \`$ac_var' since the previous run:" >&2;}
	  eval $ac_var=\$ac_old_val
	fi
	{ $as_echo "$as_me:${as_lineno-$LINENO}:   former value:  \`$ac_old_val'" >&5
$as_echo "$as_me:   former value:  \`$ac_old_val'" >&2;}
	{ $as_echo "$as_me:${as_lineno-$LINENO}:   current value: \`$ac_new_val'" >&5
$as_echo "$as_me:   current value: \`$ac_new_val'" >&2;}
      fi;;
  esac
  # Pass precious variables to config.status.
  if test "$ac_new_set" = set; then
    case $ac_new_val in
    *\'*) ac_arg=$ac_var=`$as_echo "$ac_new_val" | sed "s/'/'\\\\\\\\''/g"` ;;
    *) ac_arg=$ac_var=$ac_new_val ;;
    esac
    case " $ac_configure_args " in
      *" '$ac_arg' "*) ;; # Avoid dups.  Use of quotes ensures accuracy.
      *) as_fn_append ac_configure_args " '$ac_arg'" ;;
    esac
  fi
done
if $ac_cache_corrupted; then
  { $as_echo "$as_me:${as_lineno-$LINENO}: error: in \`$ac_pwd':" >&5
$as_echo "$as_me: error: in \`$ac_pwd':" >&2;}
  { $as_echo "$as_me:${as_lineno-$LINENO}: error: changes in the environment can compromise the build" >&5
$as_echo "$as_me: error: changes in the environment can compromise the build" >&2;}
  as_fn_error $? "run \`make distclean' and/or \`rm $cache_file' and start over" "$LINENO" 5
fi
## -------------------- ##
## Main body of script. ##
## -------------------- ##

ac_ext=c
ac_cpp='$CPP $CPPFLAGS'
ac_compile='$CC -c $CFLAGS $CPPFLAGS conftest.$ac_ext >&5'
ac_link='$CC -o conftest$ac_exeext $CFLAGS $CPPFLAGS $LDFLAGS conftest.$ac_ext $LIBS >&5'
ac_compiler_gnu=$ac_cv_c_compiler_gnu


CXX=`"${R_HOME}/bin/R" CMD config CXX`
ac_ext=cpp
ac_cpp='$CXXCPP $CPPFLAGS'
ac_compile='$CXX -c $CXXFLAGS $CPPFLAGS conftest.$ac_ext >&5'
ac_link='$CXX -o conftest$ac_exeext $CXXFLAGS $CPPFLAGS $LDFLAGS conftest.$ac_ext $LIBS >&5'
ac_compiler_gnu=$ac_cv_cxx_compiler_gnu
if test -z "$CXX"; then
  if test -n "$CCC"; then
    CXX=$CCC
  else
    if test -n "$ac_tool_prefix"; then
  for ac_prog in g++ c++ gpp aCC CC cxx cc++ cl.exe FCC KCC RCC xlC_r xlC
  do
    # Extract the first word of "$ac_tool_prefix$ac_prog", so it can be a program name with args.
set dummy $ac_tool_prefix$ac_prog; ac_word=$2
{ $as_echo "$as_me:${as_lineno-$LINENO}: checking for $ac_word" >&5
$as_echo_n "checking for $ac_word... " >&6; }
if ${ac_cv_prog_CXX+:} false; then :
  $as_echo_n "(cached) " >&6
else
  if test -n "$CXX"; then
  ac_cv_prog_CXX="$CXX" # Let the user override the test.
else
as_save_IFS=$IFS; IFS=$PATH_SEPARATOR
for as_dir in $PATH
do
  IFS=$as_save_IFS
  test -z "$as_dir" && as_dir=.
    for ac_exec_ext in '' $ac_executable_extensions; do
  if as_fn_executable_p "$as_dir/$ac_word$ac_exec_ext"; then
    ac_cv_prog_CXX="$ac_tool_prefix$ac_prog"
    $as_echo "$as_me:${as_lineno-$LINENO}: found $as_dir/$ac_word$ac_exec_ext" >&5
    break 2
  fi
done
  done
IFS=$as_save_IFS

fi
fi
CXX=$ac_cv_prog_CXX
if test -n "$CXX"; then
  { $as_echo "$as_me:${as_lineno-$LINENO}: result: $CXX" >&5
$as_echo "$CXX" >&6; }
else
  { $as_echo "$as_me:${as_lineno-$LINENO}: result: no" >&5
$as_echo "no" >&6; }
fi


    test -n "$CXX" && break
  done
fi
if test -z "$CXX"; then
  ac_ct_CXX=$CXX
  for ac_prog in g++ c++ gpp aCC CC cxx cc++ cl.exe FCC KCC RCC xlC_r xlC
do
  # Extract the first word of "$ac_prog", so it can be a program name with args.
set dummy $ac_prog; ac_word=$2
{ $as_echo "$as_me:${as_lineno-$LINENO}: checking for $ac_word" >&5
$as_echo_n "checking for $ac_word... " >&6; }
if ${ac_cv_prog_ac_ct_CXX+:} false; then :
  $as_echo_n "(cached) " >&6
else
  if test -n "$ac_ct_CXX"; then
  ac_cv_prog_ac_ct_CXX="$ac_ct_CXX" # Let the user override the test.
else
as_save_IFS=$IFS; IFS=$PATH_SEPARATOR
for as_dir in $PATH
do
  IFS=$as_save_IFS
  test -z "$as_dir" && as_dir=.
    for ac_exec_ext in '' $ac_executable_extensions; do
  if as_fn_executable_p "$as_dir/$ac_word$ac_exec_ext"; then
    ac_cv_prog_ac_ct_CXX="$ac_prog"
    $as_echo "$as_me:${as_lineno-$LINENO}: found $as_dir/$ac_word$ac_exec_ext" >&5
    break 2
  fi
done
  done
IFS=$as_save_IFS

fi
fi
ac_ct_CXX=$ac_cv_prog_ac_ct_CXX
if test -n "$ac_ct_CXX"; then
  { $as_echo "$as_me:${as_lineno-$LINENO}: result: $ac_ct_CXX" >&5
$as_echo "$ac_ct_CXX" >&6; }
else
  { $as_echo "$as_me:${as_lineno-$LINENO}: result: no" >&5
$as_echo "no" >&6; }
fi


  test -n "$ac_ct_CXX" && break
done

  if test "x$ac_ct_CXX" = x; then
    CXX="g++"
  else
    case $cross_compiling:$ac_tool_warned in
yes:)
{ $as_echo "$as_me:${as_lineno-$LINENO}: WARNING: using cross tools not prefixed with host triplet" >&5
$as_echo "$as_me: WARNING: using cross tools not prefixed with host triplet" >&2;}
ac_tool_warned=yes ;;
esac
    CXX=$ac_ct_CXX
  fi
fi

  fi
fi
# Provide some information about the compiler.
$as_echo "$as_me:${as_lineno-$LINENO}: checking for C++ compiler version" >&5
set X $ac_compile
ac_compiler=$2
for ac_option in --version -v -V -qversion; do
  { { ac_try="$ac_compiler $ac_option >&5"
case "(($ac_try" in
  *\"* | *\`* | *\\*) ac_try_echo=\$ac_try;;
  *) ac_try_echo=$ac_try;;
esac
eval ac_try_echo="\"\$as_me:${as_lineno-$LINENO}: $ac_try_echo\""
$as_echo "$ac_try_echo"; } >&5
  (eval "$ac_compiler $ac_option >&5") 2>conftest.err
  ac_status=$?
  if test -s conftest.err; then
    sed '10a\
... rest of stderr output deleted ...
         10q' conftest.err >conftest.er1
    cat conftest.er1 >&5
  fi
  rm -f conftest.er1 conftest.err
  $as_echo "$as_me:${as_lineno-$LINENO}: \$? = $ac_status" >&5
  test $ac_status = 0; }
done

cat confdefs.h - <<_ACEOF >conftest.$ac_ext
/* end confdefs.h.  */

int
main ()
{

  ;
  return 0;
}
_ACEOF
ac_clean_files_save=$ac_clean_files
ac_clean_files="$ac_clean_files a.out a.out.dSYM a.exe b.out"
# Try to create an executable without -o first, disregard a.out.
# It will help us diagnose broken compilers, and finding out an intuition
# of exeext.
{ $as_echo "$as_me:${as_lineno-$LINENO}: checking whether the C++ compiler works" >&5
$as_echo_n "checking whether the C++ compiler works... " >&6; }
ac_link_default=`$as_echo "$ac_link" | sed 's/ -o *conftest[^ ]*//'`

# The possible output files:
ac_files="a.out conftest.exe conftest a.exe a_out.exe b.out conftest.*"

ac_rmfiles=
for ac_file in $ac_files
do
  case $ac_file in
    *.$ac_ext | *.xcoff | *.tds | *.d | *.pdb | *.xSYM | *.bb | *.bbg | *.map | *.inf | *.dSYM | *.o | *.obj ) ;;
    * ) ac_rmfiles="$ac_rmfiles $ac_file";;
  esac
done
rm -f $ac_rmfiles

if { { ac_try="$ac_link_default"
case "(($ac_try" in
  *\"* | *\`* | *\\*) ac_try_echo=\$ac_try;;
  *) ac_try_echo=$ac_try;;
esac
eval ac_try_echo="\"\$as_me:${as_lineno-$LINENO}: $ac_try_echo\""
$as_echo "$ac_try_echo"; } >&5
  (eval "$ac_link_default") 2>&5
  ac_status=$?
  $as_echo "$as_me:${as_lineno-$LINENO}: \$? = $ac_status" >&5
  test $ac_status = 0; }; then :
  # Autoconf-2.13 could set the ac_cv_exeext variable to `no'.
# So ignore a value of `no', otherwise this would lead to `EXEEXT = no'
# in a Makefile.  We should not override ac_cv_exeext if it was cached,
# so that the user can short-circuit this test for compilers unknown to
# Autoconf.
for ac_file in $ac_files ''
do
  test -f "$ac_file" || continue
  case $ac_file in
    *.$ac_ext | *.xcoff | *.tds | *.d | *.pdb | *.xSYM | *.bb | *.bbg | *.map | *.inf | *.dSYM | *.o | *.obj )
	;;
    [ab].out )
	# We found the default executable, but exeext='' is most
	# certainly right.
	break;;
    *.* )
	if test "${ac_cv_exeext+set}" = set && test "$ac_cv_exeext" != no;
	then :; else
	   ac_cv_exeext=`expr "$ac_file" : '[^.]*\(\..*\)'`
	fi
	# We set ac_cv_exeext here because the later test for it is not
	# safe: cross compilers may not add the suffix if given an `-o'
	# argument, so we may need to know it at that point already.
	# Even if this section looks crufty: it has the advantage of
	# actually working.
	break;;
    * )
	break;;
  esac
done
test "$ac_cv_exeext" = no && ac_cv_exeext=

else
  ac_file=''
fi
if test -z "$ac_file"; then :
  { $as_echo "$as_me:${as_lineno-$LINENO}: result: no" >&5
$as_echo "no" >&6; }
$as_echo "$as_me: failed program was:" >&5
sed 's/^/| /' conftest.$ac_ext >&5

{ { $as_echo "$as_me:${as_lineno-$LINENO}: error: in \`$ac_pwd':" >&5
$as_echo "$as_me: error: in \`$ac_pwd':" >&2;}
as_fn_error 77 "C++ compiler cannot create executables
See \`config.log' for more details" "$LINENO" 5; }
else
  { $as_echo "$as_me:${as_lineno-$LINENO}: result: yes" >&5
$as_echo "yes" >&6; }
fi
{ $as_echo "$as_me:${as_lineno-$LINENO}: checking for C++ compiler default output file name" >&5
$as_echo_n "checking for C++ compiler default output file name... " >&6; }
{ $as_echo "$as_me:${as_lineno-$LINENO}: result: $ac_file" >&5
$as_echo "$ac_file" >&6; }
ac_exeext=$ac_cv_exeext

rm -f -r a.out a.out.dSYM a.exe conftest$ac_cv_exeext b.out
ac_clean_files=$ac_clean_files_save
{ $as_echo "$as_me:${as_lineno-$LINENO}: checking for suffix of executables" >&5
$as_echo_n "checking for suffix of executables... " >&6; }
if { { ac_try="$ac_link"
case "(($ac_try" in
  *\"* | *\`* | *\\*) ac_try_echo=\$ac_try;;
  *) ac_try_echo=$ac_try;;
esac
eval ac_try_echo="\"\$as_me:${as_lineno-$LINENO}: $ac_try_echo\""
$as_echo "$ac_try_echo"; } >&5
  (eval "$ac_link") 2>&5
  ac_status=$?
  $as_echo "$as_me:${as_lineno-$LINENO}: \$? = $ac_status" >&5
  test $ac_status = 0; }; then :
  # If both `conftest.exe' and `conftest' are `present' (well, observable)
# catch `conftest.exe'.  For instance with Cygwin, `ls conftest' will
# work properly (i.e., refer to `conftest.exe'), while it won't with
# `rm'.
for ac_file in conftest.exe conftest conftest.*; do
  test -f "$ac_file" || continue
  case $ac_file in
    *.$ac_ext | *.xcoff | *.tds | *.d | *.pdb | *.xSYM | *.bb | *.bbg | *.map | *.inf | *.dSYM | *.o | *.obj ) ;;
    *.* ) ac_cv_exeext=`expr "$ac_file" : '[^.]*\(\..*\)'`
	  break;;
    * ) break;;
  esac
done
else
  { { $as_echo "$as_me:${as_lineno-$LINENO}: error: in \`$ac_pwd':" >&5
$as_echo "$as_me: error: in \`$ac_pwd':" >&2;}
as_fn_error $? "cannot compute suffix of executables: cannot compile and link
See \`config.log' for more details" "$LINENO" 5; }
fi
rm -f conftest conftest$ac_cv_exeext
{ $as_echo "$as_me:${as_lineno-$LINENO}: result: $ac_cv_exeext" >&5
$as_echo "$ac_cv_exeext" >&6; }

rm -f conftest.$ac_ext
EXEEXT=$ac_cv_exeext
ac_exeext=$EXEEXT
cat confdefs.h - <<_ACEOF >conftest.$ac_ext
/* end confdefs.h.  */
#include <stdio.h>
int
main ()
{
FILE *f = fopen ("conftest.out", "w");
 return ferror (f) || fclose (f) != 0;

  ;
  return 0;
}
_ACEOF
ac_clean_files="$ac_clean_files conftest.out"
# Check that the compiler produces executables we can run.  If not, either
# the compiler is broken, or we cross compile.
{ $as_echo "$as_me:${as_lineno-$LINENO}: checking whether we are cross compiling" >&5
$as_echo_n "checking whether we are cross compiling... " >&6; }
if test "$cross_compiling" != yes; then
  { { ac_try="$ac_link"
case "(($ac_try" in
  *\"* | *\`* | *\\*) ac_try_echo=\$ac_try;;
  *) ac_try_echo=$ac_try;;
esac
eval ac_try_echo="\"\$as_me:${as_lineno-$LINENO}: $ac_try_echo\""
$as_echo "$ac_try_echo"; } >&5
  (eval "$ac_link") 2>&5
  ac_status=$?
  $as_echo "$as_me:${as_lineno-$LINENO}: \$? = $ac_status" >&5
  test $ac_status = 0; }
  if { ac_try='./conftest$ac_cv_exeext'
  { { case "(($ac_try" in
  *\"* | *\`* | *\\*) ac_try_echo=\$ac_try;;
  *) ac_try_echo=$ac_try;;
esac
eval ac_try_echo="\"\$as_me:${as_lineno-$LINENO}: $ac_try_echo\""
$as_echo "$ac_try_echo"; } >&5
  (eval "$ac_try") 2>&5
  ac_status=$?
  $as_echo "$as_me:${as_lineno-$LINENO}: \$? = $ac_status" >&5
  test $ac_status = 0; }; }; then
    cross_compiling=no
  else
    if test "$cross_compiling" = maybe; then
	cross_compiling=yes
    else
	{ { $as_echo "$as_me:${as_lineno-$LINENO}: error: in \`$ac_pwd':" >&5
$as_echo "$as_me: error: in \`$ac_pwd':" >&2;}
as_fn_error $? "cannot run C++ compiled programs.
If you meant to cross compile, use \`--host'.
See \`config.log' for more details" "$LINENO" 5; }
    fi
  fi
fi
{ $as_echo "$as_me:${as_lineno-$LINENO}: result: $cross_compiling" >&5
$as_echo "$cross_compiling" >&6; }

rm -f conftest.$ac_ext conftest$ac_cv_exeext conftest.out
ac_clean_files=$ac_clean_files_save
{ $as_echo "$as_me:${as_lineno-$LINENO}: checking for suffix of object files" >&5
$as_echo_n "checking for suffix of object files... " >&6; }
if ${ac_cv_objext+:} false; then :
  $as_echo_n "(cached) " >&6
else
  cat confdefs.h - <<_ACEOF >conftest.$ac_ext
/* end confdefs.h.  */

int
main ()
{

  ;
  return 0;
}
_ACEOF
rm -f conftest.o conftest.obj
if { { ac_try="$ac_compile"
case "(($ac_try" in
  *\"* | *\`* | *\\*) ac_try_echo=\$ac_try;;
  *) ac_try_echo=$ac_try;;
esac
eval ac_try_echo="\"\$as_me:${as_lineno-$LINENO}: $ac_try_echo\""
$as_echo "$ac_try_echo"; } >&5
  (eval "$ac_compile") 2>&5
  ac_status=$?
  $as_echo "$as_me:${as_lineno-$LINENO}: \$? = $ac_status" >&5
  test $ac_status = 0; }; then :
  for ac_file in conftest.o conftest.obj conftest.*; do
  test -f "$ac_file" || continue;
  case $ac_file in
    *.$ac_ext | *.xcoff | *.tds | *.d | *.pdb | *.xSYM | *.bb | *.bbg | *.map | *.inf | *.dSYM ) ;;
    *) ac_cv_objext=`expr "$ac_file" : '.*\.\(.*\)'`
       break;;
  esac
done
else
  $as_echo "$as_me: failed program was:" >&5
sed 's/^/| /' conftest.$ac_ext >&5

{ { $as_echo "$as_me:${as_lineno-$LINENO}: error: in \`$ac_pwd':" >&5
$as_echo "$as_me: error: in \`$ac_pwd':" >&2;}
as_fn_error $? "cannot compute suffix of object files: cannot compile
See \`config.log' for more details" "$LINENO" 5; }
fi
rm -f conftest.$ac_cv_objext conftest.$ac_ext
fi
{ $as_echo "$as_me:${as_lineno-$LINENO}: result: $ac_cv_objext" >&5
$as_echo "$ac_cv_objext" >&6; }
OBJEXT=$ac_cv_objext
ac_objext=$OBJEXT
{ $as_echo "$as_me:${as_lineno-$LINENO}: checking whether we are using the GNU C++ compiler" >&5
$as_echo_n "checking whether we are using the GNU C++ compiler... " >&6; }
if ${ac_cv_cxx_compiler_gnu+:} false; then :
  $as_echo_n "(cached) " >&6
else
  cat confdefs.h - <<_ACEOF >conftest.$ac_ext
/* end confdefs.h.  */

int
main ()
{
#ifndef __GNUC__
       choke me
#endif

  ;
  return 0;
}
_ACEOF
if ac_fn_cxx_try_compile "$LINENO"; then :
  ac_compiler_gnu=yes
else
  ac_compiler_gnu=no
fi
rm -f core conftest.err conftest.$ac_objext conftest.$ac_ext
ac_cv_cxx_compiler_gnu=$ac_compiler_gnu

fi
{ $as_echo "$as_me:${as_lineno-$LINENO}: result: $ac_cv_cxx_compiler_gnu" >&5
$as_echo "$ac_cv_cxx_compiler_gnu" >&6; }
if test $ac_compiler_gnu = yes; then
  GXX=yes
else
  GXX=
fi
ac_test_CXXFLAGS=${CXXFLAGS+set}
ac_save_CXXFLAGS=$CXXFLAGS
{ $as_echo "$as_me:${as_lineno-$LINENO}: checking whether $CXX accepts -g" >&5
$as_echo_n "checking whether $CXX accepts -g... " >&6; }
if ${ac_cv_prog_cxx_g+:} false; then :
  $as_echo_n "(cached) " >&6
else
  ac_save_cxx_werror_flag=$ac_cxx_werror_flag
   ac_cxx_werror_flag=yes
   ac_cv_prog_cxx_g=no
   CXXFLAGS="-g"
   cat confdefs.h - <<_ACEOF >conftest.$ac_ext
/* end confdefs.h.  */

int
main ()
{

  ;
  return 0;
}
_ACEOF
if ac_fn_cxx_try_compile "$LINENO"; then :
  ac_cv_prog_cxx_g=yes
else
  CXXFLAGS=""
      cat confdefs.h - <<_ACEOF >conftest.$ac_ext
/* end confdefs.h.  */

int
main ()
{

  ;
  return 0;
}
_ACEOF
if ac_fn_cxx_try_compile "$LINENO"; then :

else
  ac_cxx_werror_flag=$ac_save_cxx_werror_flag
	 CXXFLAGS="-g"
	 cat confdefs.h - <<_ACEOF >conftest.$ac_ext
/* end confdefs.h.  */

int
main ()
{

  ;
  return 0;
}
_ACEOF
if ac_fn_cxx_try_compile "$LINENO"; then :
  ac_cv_prog_cxx_g=yes
fi
rm -f core conftest.err conftest.$ac_objext conftest.$ac_ext
fi
rm -f core conftest.err conftest.$ac_objext conftest.$ac_ext
fi
rm -f core conftest.err conftest.$ac_objext conftest.$ac_ext
   ac_cxx_werror_flag=$ac_save_cxx_werror_flag
fi
{ $as_echo "$as_me:${as_lineno-$LINENO}: result: $ac_cv_prog_cxx_g" >&5
$as_echo "$ac_cv_prog_cxx_g" >&6; }
if test "$ac_test_CXXFLAGS" = set; then
  CXXFLAGS=$ac_save_CXXFLAGS
elif test $ac_cv_prog_cxx_g = yes; then
  if test "$GXX" = yes; then
    CXXFLAGS="-g -O2"
  else
    CXXFLAGS="-g"
  fi
else
  if test "$GXX" = yes; then
    CXXFLAGS="-O2"
  else
    CXXFLAGS=
  fi
fi
ac_ext=c
ac_cpp='$CPP $CPPFLAGS'
ac_compile='$CC -c $CFLAGS $CPPFLAGS conftest.$ac_ext >&5'
ac_link='$CC -o conftest$ac_exeext $CFLAGS $CPPFLAGS $LDFLAGS conftest.$ac_ext $LIBS >&5'
ac_compiler_gnu=$ac_cv_c_compiler_gnu

ac_ext=cpp
ac_cpp='$CXXCPP $CPPFLAGS'
ac_compile='$CXX -c $CXXFLAGS $CPPFLAGS conftest.$ac_ext >&5'
ac_link='$CXX -o conftest$ac_exeext $CXXFLAGS $CPPFLAGS $LDFLAGS conftest.$ac_ext $LIBS >&5'
ac_compiler_gnu=$ac_cv_cxx_compiler_gnu


CPPFLAGS="-march=native"
{ $as_echo "$as_me:${as_lineno-$LINENO}: checking which SIMD flags are available" >&5
$as_echo_n "checking which SIMD flags are available... " >&6; }

cat confdefs.h - <<_ACEOF >conftest.$ac_ext
/* end confdefs.h.  */

int
main ()
{

  ;
  return 0;
}
_ACEOF
if ac_fn_cxx_try_compile "$LINENO"; then :

else

  FX=""
  ## sysctl -a | grep "cpu.features:" # for OS X
  if test -a /proc/cpuinfo; then
    for SET in sse2 ssse3 avx2; do
      CPPFLAGS="-m$SET"
      info=`grep flags /proc/cpuinfo | grep " $SET "`
      if test "$info" != ""; then
        cat confdefs.h - <<_ACEOF >conftest.$ac_ext
/* end confdefs.h.  */

int
main ()
{

  ;
  return 0;
}
_ACEOF
if ac_fn_cxx_try_compile "$LINENO"; then :
   FX="$FX $CPPFLAGS"
fi
rm -f core conftest.err conftest.$ac_objext conftest.$ac_ext
      fi
    done
  fi
  CPPFLAGS="$FX"

fi
rm -f core conftest.err conftest.$ac_objext conftest.$ac_ext

##CPPFLAGS="" ## to be deleted if the above does not cause any problems
if test "$CPPFLAGS" != "" ; then
  { $as_echo "$as_me:${as_lineno-$LINENO}: result: $CPPFLAGS" >&5
$as_echo "$CPPFLAGS" >&6; }
else
  { $as_echo "$as_me:${as_lineno-$LINENO}: result: none" >&5
$as_echo "none" >&6; }
fi

## install.packages("miraculix_0.9.6.tar.gz", configure.args="CXX_FLAGS=-march=native", repos=NULL)
MIRACULIX_CXXFLAGS="$CXX_FLAGS $CPPFLAGS"

ac_config_files="$ac_config_files src/Makevars"

cat >confcache <<\_ACEOF
# This file is a shell script that caches the results of configure
# tests run on this system so they can be shared between configure
# scripts and configure runs, see configure's option --config-cache.
# It is not useful on other systems.  If it contains results you don't
# want to keep, you may remove or edit it.
#
# config.status only pays attention to the cache file if you give it
# the --recheck option to rerun configure.
#
# `ac_cv_env_foo' variables (set or unset) will be overridden when
# loading this file, other *unset* `ac_cv_foo' will be assigned the
# following values.

_ACEOF

# The following way of writing the cache mishandles newlines in values,
# but we know of no workaround that is simple, portable, and efficient.
# So, we kill variables containing newlines.
# Ultrix sh set writes to stderr and can't be redirected directly,
# and sets the high bit in the cache file unless we assign to the vars.
(
  for ac_var in `(set) 2>&1 | sed -n 's/^\([a-zA-Z_][a-zA-Z0-9_]*\)=.*/\1/p'`; do
    eval ac_val=\$$ac_var
    case $ac_val in #(
    *${as_nl}*)
      case $ac_var in #(
      *_cv_*) { $as_echo "$as_me:${as_lineno-$LINENO}: WARNING: cache variable $ac_var contains a newline" >&5
$as_echo "$as_me: WARNING: cache variable $ac_var contains a newline" >&2;} ;;
      esac
      case $ac_var in #(
      _ | IFS | as_nl) ;; #(
      BASH_ARGV | BASH_SOURCE) eval $ac_var= ;; #(
      *) { eval $ac_var=; unset $ac_var;} ;;
      esac ;;
    esac
  done

  (set) 2>&1 |
    case $as_nl`(ac_space=' '; set) 2>&1` in #(
    *${as_nl}ac_space=\ *)
      # `set' does not quote correctly, so add quotes: double-quote
      # substitution turns \\\\ into \\, and sed turns \\ into \.
      sed -n \
	"s/'/'\\\\''/g;
	  s/^\\([_$as_cr_alnum]*_cv_[_$as_cr_alnum]*\\)=\\(.*\\)/\\1='\\2'/p"
      ;; #(
    *)
      # `set' quotes correctly as required by POSIX, so do not add quotes.
      sed -n "/^[_$as_cr_alnum]*_cv_[_$as_cr_alnum]*=/p"
      ;;
    esac |
    sort
) |
  sed '
     /^ac_cv_env_/b end
     t clear
     :clear
     s/^\([^=]*\)=\(.*[{}].*\)$/test "${\1+set}" = set || &/
     t end
     s/^\([^=]*\)=\(.*\)$/\1=${\1=\2}/
     :end' >>confcache
if diff "$cache_file" confcache >/dev/null 2>&1; then :; else
  if test -w "$cache_file"; then
    if test "x$cache_file" != "x/dev/null"; then
      { $as_echo "$as_me:${as_lineno-$LINENO}: updating cache $cache_file" >&5
$as_echo "$as_me: updating cache $cache_file" >&6;}
      if test ! -f "$cache_file" || test -h "$cache_file"; then
	cat confcache >"$cache_file"
      else
        case $cache_file in #(
        */* | ?:*)
	  mv -f confcache "$cache_file"$$ &&
	  mv -f "$cache_file"$$ "$cache_file" ;; #(
        *)
	  mv -f confcache "$cache_file" ;;
	esac
      fi
    fi
  else
    { $as_echo "$as_me:${as_lineno-$LINENO}: not updating unwritable cache $cache_file" >&5
$as_echo "$as_me: not updating unwritable cache $cache_file" >&6;}
  fi
fi
rm -f confcache

test "x$prefix" = xNONE && prefix=$ac_default_prefix
# Let make expand exec_prefix.
test "x$exec_prefix" = xNONE && exec_prefix='${prefix}'

# Transform confdefs.h into DEFS.
# Protect against shell expansion while executing Makefile rules.
# Protect against Makefile macro expansion.
#
# If the first sed substitution is executed (which looks for macros that
# take arguments), then branch to the quote section.  Otherwise,
# look for a macro that doesn't take arguments.
ac_script='
:mline
/\\$/{
 N
 s,\\\n,,
 b mline
}
t clear
:clear
s/^[	 ]*#[	 ]*define[	 ][	 ]*\([^	 (][^	 (]*([^)]*)\)[	 ]*\(.*\)/-D\1=\2/g
t quote
s/^[	 ]*#[	 ]*define[	 ][	 ]*\([^	 ][^	 ]*\)[	 ]*\(.*\)/-D\1=\2/g
t quote
b any
:quote
s/[	 `~#$^&*(){}\\|;'\''"<>?]/\\&/g
s/\[/\\&/g
s/\]/\\&/g
s/\$/$$/g
H
:any
${
	g
	s/^\n//
	s/\n/ /g
	p
}
'
DEFS=`sed -n "$ac_script" confdefs.h`


ac_libobjs=
ac_ltlibobjs=
U=
for ac_i in : $LIBOBJS; do test "x$ac_i" = x: && continue
  # 1. Remove the extension, and $U if already installed.
  ac_script='s/\$U\././;s/\.o$//;s/\.obj$//'
  ac_i=`$as_echo "$ac_i" | sed "$ac_script"`
  # 2. Prepend LIBOBJDIR.  When used with automake>=1.10 LIBOBJDIR
  #    will be set to the directory where LIBOBJS objects are built.
  as_fn_append ac_libobjs " \${LIBOBJDIR}$ac_i\$U.$ac_objext"
  as_fn_append ac_ltlibobjs " \${LIBOBJDIR}$ac_i"'$U.lo'
done
LIBOBJS=$ac_libobjs

LTLIBOBJS=$ac_ltlibobjs


: "${CONFIG_STATUS=./config.status}"
ac_write_fail=0
ac_clean_files_save=$ac_clean_files
ac_clean_files="$ac_clean_files $CONFIG_STATUS"
{ $as_echo "$as_me:${as_lineno-$LINENO}: creating $CONFIG_STATUS" >&5
$as_echo "$as_me: creating $CONFIG_STATUS" >&6;}
as_write_fail=0
cat >$CONFIG_STATUS <<_ASEOF || as_write_fail=1
#! $SHELL
# Generated by $as_me.
# Run this file to recreate the current configuration.
# Compiler output produced by configure, useful for debugging
# configure, is in config.log if it exists.

debug=false
ac_cs_recheck=false
ac_cs_silent=false

SHELL=\${CONFIG_SHELL-$SHELL}
export SHELL
_ASEOF
cat >>$CONFIG_STATUS <<\_ASEOF || as_write_fail=1
## -------------------- ##
## M4sh Initialization. ##
## -------------------- ##

# Be more Bourne compatible
DUALCASE=1; export DUALCASE # for MKS sh
if test -n "${ZSH_VERSION+set}" && (emulate sh) >/dev/null 2>&1; then :
  emulate sh
  NULLCMD=:
  # Pre-4.2 versions of Zsh do word splitting on ${1+"$@"}, which
  # is contrary to our usage.  Disable this feature.
  alias -g '${1+"$@"}'='"$@"'
  setopt NO_GLOB_SUBST
else
  case `(set -o) 2>/dev/null` in #(
  *posix*) :
    set -o posix ;; #(
  *) :
     ;;
esac
fi


as_nl='
'
export as_nl
# Printing a long string crashes Solaris 7 /usr/bin/printf.
as_echo='\\\\\\\\\\\\\\\\\\\\\\\\\\\\\\\\\\\\\\\\\\\\\\\\\\\\\\\\\\\\\\\\\\\\\\\\\\\\\\\\\\\\\\\\\\\\\\\\\\\\\\\'
as_echo=$as_echo$as_echo$as_echo$as_echo$as_echo
as_echo=$as_echo$as_echo$as_echo$as_echo$as_echo$as_echo
# Prefer a ksh shell builtin over an external printf program on Solaris,
# but without wasting forks for bash or zsh.
if test -z "$BASH_VERSION$ZSH_VERSION" \
    && (test "X`print -r -- $as_echo`" = "X$as_echo") 2>/dev/null; then
  as_echo='print -r --'
  as_echo_n='print -rn --'
elif (test "X`printf %s $as_echo`" = "X$as_echo") 2>/dev/null; then
  as_echo='printf %s\n'
  as_echo_n='printf %s'
else
  if test "X`(/usr/ucb/echo -n -n $as_echo) 2>/dev/null`" = "X-n $as_echo"; then
    as_echo_body='eval /usr/ucb/echo -n "$1$as_nl"'
    as_echo_n='/usr/ucb/echo -n'
  else
    as_echo_body='eval expr "X$1" : "X\\(.*\\)"'
    as_echo_n_body='eval
      arg=$1;
      case $arg in #(
      *"$as_nl"*)
	expr "X$arg" : "X\\(.*\\)$as_nl";
	arg=`expr "X$arg" : ".*$as_nl\\(.*\\)"`;;
      esac;
      expr "X$arg" : "X\\(.*\\)" | tr -d "$as_nl"
    '
    export as_echo_n_body
    as_echo_n='sh -c $as_echo_n_body as_echo'
  fi
  export as_echo_body
  as_echo='sh -c $as_echo_body as_echo'
fi

# The user is always right.
if test "${PATH_SEPARATOR+set}" != set; then
  PATH_SEPARATOR=:
  (PATH='/bin;/bin'; FPATH=$PATH; sh -c :) >/dev/null 2>&1 && {
    (PATH='/bin:/bin'; FPATH=$PATH; sh -c :) >/dev/null 2>&1 ||
      PATH_SEPARATOR=';'
  }
fi


# IFS
# We need space, tab and new line, in precisely that order.  Quoting is
# there to prevent editors from complaining about space-tab.
# (If _AS_PATH_WALK were called with IFS unset, it would disable word
# splitting by setting IFS to empty value.)
IFS=" ""	$as_nl"

# Find who we are.  Look in the path if we contain no directory separator.
as_myself=
case $0 in #((
  *[\\/]* ) as_myself=$0 ;;
  *) as_save_IFS=$IFS; IFS=$PATH_SEPARATOR
for as_dir in $PATH
do
  IFS=$as_save_IFS
  test -z "$as_dir" && as_dir=.
    test -r "$as_dir/$0" && as_myself=$as_dir/$0 && break
  done
IFS=$as_save_IFS

     ;;
esac
# We did not find ourselves, most probably we were run as `sh COMMAND'
# in which case we are not to be found in the path.
if test "x$as_myself" = x; then
  as_myself=$0
fi
if test ! -f "$as_myself"; then
  $as_echo "$as_myself: error: cannot find myself; rerun with an absolute file name" >&2
  exit 1
fi

# Unset variables that we do not need and which cause bugs (e.g. in
# pre-3.0 UWIN ksh).  But do not cause bugs in bash 2.01; the "|| exit 1"
# suppresses any "Segmentation fault" message there.  '((' could
# trigger a bug in pdksh 5.2.14.
for as_var in BASH_ENV ENV MAIL MAILPATH
do eval test x\${$as_var+set} = xset \
  && ( (unset $as_var) || exit 1) >/dev/null 2>&1 && unset $as_var || :
done
PS1='$ '
PS2='> '
PS4='+ '

# NLS nuisances.
LC_ALL=C
export LC_ALL
LANGUAGE=C
export LANGUAGE

# CDPATH.
(unset CDPATH) >/dev/null 2>&1 && unset CDPATH


# as_fn_error STATUS ERROR [LINENO LOG_FD]
# ----------------------------------------
# Output "`basename $0`: error: ERROR" to stderr. If LINENO and LOG_FD are
# provided, also output the error to LOG_FD, referencing LINENO. Then exit the
# script with STATUS, using 1 if that was 0.
as_fn_error ()
{
  as_status=$1; test $as_status -eq 0 && as_status=1
  if test "$4"; then
    as_lineno=${as_lineno-"$3"} as_lineno_stack=as_lineno_stack=$as_lineno_stack
    $as_echo "$as_me:${as_lineno-$LINENO}: error: $2" >&$4
  fi
  $as_echo "$as_me: error: $2" >&2
  as_fn_exit $as_status
} # as_fn_error


# as_fn_set_status STATUS
# -----------------------
# Set $? to STATUS, without forking.
as_fn_set_status ()
{
  return $1
} # as_fn_set_status

# as_fn_exit STATUS
# -----------------
# Exit the shell with STATUS, even in a "trap 0" or "set -e" context.
as_fn_exit ()
{
  set +e
  as_fn_set_status $1
  exit $1
} # as_fn_exit

# as_fn_unset VAR
# ---------------
# Portably unset VAR.
as_fn_unset ()
{
  { eval $1=; unset $1;}
}
as_unset=as_fn_unset
# as_fn_append VAR VALUE
# ----------------------
# Append the text in VALUE to the end of the definition contained in VAR. Take
# advantage of any shell optimizations that allow amortized linear growth over
# repeated appends, instead of the typical quadratic growth present in naive
# implementations.
if (eval "as_var=1; as_var+=2; test x\$as_var = x12") 2>/dev/null; then :
  eval 'as_fn_append ()
  {
    eval $1+=\$2
  }'
else
  as_fn_append ()
  {
    eval $1=\$$1\$2
  }
fi # as_fn_append

# as_fn_arith ARG...
# ------------------
# Perform arithmetic evaluation on the ARGs, and store the result in the
# global $as_val. Take advantage of shells that can avoid forks. The arguments
# must be portable across $(()) and expr.
if (eval "test \$(( 1 + 1 )) = 2") 2>/dev/null; then :
  eval 'as_fn_arith ()
  {
    as_val=$(( $* ))
  }'
else
  as_fn_arith ()
  {
    as_val=`expr "$@" || test $? -eq 1`
  }
fi # as_fn_arith


if expr a : '\(a\)' >/dev/null 2>&1 &&
   test "X`expr 00001 : '.*\(...\)'`" = X001; then
  as_expr=expr
else
  as_expr=false
fi

if (basename -- /) >/dev/null 2>&1 && test "X`basename -- / 2>&1`" = "X/"; then
  as_basename=basename
else
  as_basename=false
fi

if (as_dir=`dirname -- /` && test "X$as_dir" = X/) >/dev/null 2>&1; then
  as_dirname=dirname
else
  as_dirname=false
fi

as_me=`$as_basename -- "$0" ||
$as_expr X/"$0" : '.*/\([^/][^/]*\)/*$' \| \
	 X"$0" : 'X\(//\)$' \| \
	 X"$0" : 'X\(/\)' \| . 2>/dev/null ||
$as_echo X/"$0" |
    sed '/^.*\/\([^/][^/]*\)\/*$/{
	    s//\1/
	    q
	  }
	  /^X\/\(\/\/\)$/{
	    s//\1/
	    q
	  }
	  /^X\/\(\/\).*/{
	    s//\1/
	    q
	  }
	  s/.*/./; q'`

# Avoid depending upon Character Ranges.
as_cr_letters='abcdefghijklmnopqrstuvwxyz'
as_cr_LETTERS='ABCDEFGHIJKLMNOPQRSTUVWXYZ'
as_cr_Letters=$as_cr_letters$as_cr_LETTERS
as_cr_digits='0123456789'
as_cr_alnum=$as_cr_Letters$as_cr_digits

ECHO_C= ECHO_N= ECHO_T=
case `echo -n x` in #(((((
-n*)
  case `echo 'xy\c'` in
  *c*) ECHO_T='	';;	# ECHO_T is single tab character.
  xy)  ECHO_C='\c';;
  *)   echo `echo ksh88 bug on AIX 6.1` > /dev/null
       ECHO_T='	';;
  esac;;
*)
  ECHO_N='-n';;
esac

rm -f conf$$ conf$$.exe conf$$.file
if test -d conf$$.dir; then
  rm -f conf$$.dir/conf$$.file
else
  rm -f conf$$.dir
  mkdir conf$$.dir 2>/dev/null
fi
if (echo >conf$$.file) 2>/dev/null; then
  if ln -s conf$$.file conf$$ 2>/dev/null; then
    as_ln_s='ln -s'
    # ... but there are two gotchas:
    # 1) On MSYS, both `ln -s file dir' and `ln file dir' fail.
    # 2) DJGPP < 2.04 has no symlinks; `ln -s' creates a wrapper executable.
    # In both cases, we have to default to `cp -pR'.
    ln -s conf$$.file conf$$.dir 2>/dev/null && test ! -f conf$$.exe ||
      as_ln_s='cp -pR'
  elif ln conf$$.file conf$$ 2>/dev/null; then
    as_ln_s=ln
  else
    as_ln_s='cp -pR'
  fi
else
  as_ln_s='cp -pR'
fi
rm -f conf$$ conf$$.exe conf$$.dir/conf$$.file conf$$.file
rmdir conf$$.dir 2>/dev/null


# as_fn_mkdir_p
# -------------
# Create "$as_dir" as a directory, including parents if necessary.
as_fn_mkdir_p ()
{

  case $as_dir in #(
  -*) as_dir=./$as_dir;;
  esac
  test -d "$as_dir" || eval $as_mkdir_p || {
    as_dirs=
    while :; do
      case $as_dir in #(
      *\'*) as_qdir=`$as_echo "$as_dir" | sed "s/'/'\\\\\\\\''/g"`;; #'(
      *) as_qdir=$as_dir;;
      esac
      as_dirs="'$as_qdir' $as_dirs"
      as_dir=`$as_dirname -- "$as_dir" ||
$as_expr X"$as_dir" : 'X\(.*[^/]\)//*[^/][^/]*/*$' \| \
	 X"$as_dir" : 'X\(//\)[^/]' \| \
	 X"$as_dir" : 'X\(//\)$' \| \
	 X"$as_dir" : 'X\(/\)' \| . 2>/dev/null ||
$as_echo X"$as_dir" |
    sed '/^X\(.*[^/]\)\/\/*[^/][^/]*\/*$/{
	    s//\1/
	    q
	  }
	  /^X\(\/\/\)[^/].*/{
	    s//\1/
	    q
	  }
	  /^X\(\/\/\)$/{
	    s//\1/
	    q
	  }
	  /^X\(\/\).*/{
	    s//\1/
	    q
	  }
	  s/.*/./; q'`
      test -d "$as_dir" && break
    done
    test -z "$as_dirs" || eval "mkdir $as_dirs"
  } || test -d "$as_dir" || as_fn_error $? "cannot create directory $as_dir"


} # as_fn_mkdir_p
if mkdir -p . 2>/dev/null; then
  as_mkdir_p='mkdir -p "$as_dir"'
else
  test -d ./-p && rmdir ./-p
  as_mkdir_p=false
fi


# as_fn_executable_p FILE
# -----------------------
# Test if FILE is an executable regular file.
as_fn_executable_p ()
{
  test -f "$1" && test -x "$1"
} # as_fn_executable_p
as_test_x='test -x'
as_executable_p=as_fn_executable_p

# Sed expression to map a string onto a valid CPP name.
as_tr_cpp="eval sed 'y%*$as_cr_letters%P$as_cr_LETTERS%;s%[^_$as_cr_alnum]%_%g'"

# Sed expression to map a string onto a valid variable name.
as_tr_sh="eval sed 'y%*+%pp%;s%[^_$as_cr_alnum]%_%g'"


exec 6>&1
## ----------------------------------- ##
## Main body of $CONFIG_STATUS script. ##
## ----------------------------------- ##
_ASEOF
test $as_write_fail = 0 && chmod +x $CONFIG_STATUS || ac_write_fail=1

cat >>$CONFIG_STATUS <<\_ACEOF || ac_write_fail=1
# Save the log message, to keep $0 and so on meaningful, and to
# report actual input values of CONFIG_FILES etc. instead of their
# values after options handling.
ac_log="
This file was extended by miraculix $as_me 0.9, which was
generated by GNU Autoconf 2.69.  Invocation command line was

  CONFIG_FILES    = $CONFIG_FILES
  CONFIG_HEADERS  = $CONFIG_HEADERS
  CONFIG_LINKS    = $CONFIG_LINKS
  CONFIG_COMMANDS = $CONFIG_COMMANDS
  $ $0 $@

on `(hostname || uname -n) 2>/dev/null | sed 1q`
"

_ACEOF

case $ac_config_files in *"
"*) set x $ac_config_files; shift; ac_config_files=$*;;
esac


cat >>$CONFIG_STATUS <<_ACEOF || ac_write_fail=1
# Files that config.status was made for.
config_files="$ac_config_files"

_ACEOF

cat >>$CONFIG_STATUS <<\_ACEOF || ac_write_fail=1
ac_cs_usage="\
\`$as_me' instantiates files and other configuration actions
from templates according to the current configuration.  Unless the files
and actions are specified as TAGs, all are instantiated by default.

Usage: $0 [OPTION]... [TAG]...

  -h, --help       print this help, then exit
  -V, --version    print version number and configuration settings, then exit
      --config     print configuration, then exit
  -q, --quiet, --silent
                   do not print progress messages
  -d, --debug      don't remove temporary files
      --recheck    update $as_me by reconfiguring in the same conditions
      --file=FILE[:TEMPLATE]
                   instantiate the configuration file FILE

Configuration files:
$config_files

Report bugs to the package provider."

_ACEOF
cat >>$CONFIG_STATUS <<_ACEOF || ac_write_fail=1
ac_cs_config="`$as_echo "$ac_configure_args" | sed 's/^ //; s/[\\""\`\$]/\\\\&/g'`"
ac_cs_version="\\
miraculix config.status 0.9
configured by $0, generated by GNU Autoconf 2.69,
  with options \\"\$ac_cs_config\\"

Copyright (C) 2012 Free Software Foundation, Inc.
This config.status script is free software; the Free Software Foundation
gives unlimited permission to copy, distribute and modify it."

ac_pwd='$ac_pwd'
srcdir='$srcdir'
test -n "\$AWK" || AWK=awk
_ACEOF

cat >>$CONFIG_STATUS <<\_ACEOF || ac_write_fail=1
# The default lists apply if the user does not specify any file.
ac_need_defaults=:
while test $# != 0
do
  case $1 in
  --*=?*)
    ac_option=`expr "X$1" : 'X\([^=]*\)='`
    ac_optarg=`expr "X$1" : 'X[^=]*=\(.*\)'`
    ac_shift=:
    ;;
  --*=)
    ac_option=`expr "X$1" : 'X\([^=]*\)='`
    ac_optarg=
    ac_shift=:
    ;;
  *)
    ac_option=$1
    ac_optarg=$2
    ac_shift=shift
    ;;
  esac

  case $ac_option in
  # Handling of the options.
  -recheck | --recheck | --rechec | --reche | --rech | --rec | --re | --r)
    ac_cs_recheck=: ;;
  --version | --versio | --versi | --vers | --ver | --ve | --v | -V )
    $as_echo "$ac_cs_version"; exit ;;
  --config | --confi | --conf | --con | --co | --c )
    $as_echo "$ac_cs_config"; exit ;;
  --debug | --debu | --deb | --de | --d | -d )
    debug=: ;;
  --file | --fil | --fi | --f )
    $ac_shift
    case $ac_optarg in
    *\'*) ac_optarg=`$as_echo "$ac_optarg" | sed "s/'/'\\\\\\\\''/g"` ;;
    '') as_fn_error $? "missing file argument" ;;
    esac
    as_fn_append CONFIG_FILES " '$ac_optarg'"
    ac_need_defaults=false;;
  --he | --h |  --help | --hel | -h )
    $as_echo "$ac_cs_usage"; exit ;;
  -q | -quiet | --quiet | --quie | --qui | --qu | --q \
  | -silent | --silent | --silen | --sile | --sil | --si | --s)
    ac_cs_silent=: ;;

  # This is an error.
  -*) as_fn_error $? "unrecognized option: \`$1'
Try \`$0 --help' for more information." ;;

  *) as_fn_append ac_config_targets " $1"
     ac_need_defaults=false ;;

  esac
  shift
done

ac_configure_extra_args=

if $ac_cs_silent; then
  exec 6>/dev/null
  ac_configure_extra_args="$ac_configure_extra_args --silent"
fi

_ACEOF
cat >>$CONFIG_STATUS <<_ACEOF || ac_write_fail=1
if \$ac_cs_recheck; then
  set X $SHELL '$0' $ac_configure_args \$ac_configure_extra_args --no-create --no-recursion
  shift
  \$as_echo "running CONFIG_SHELL=$SHELL \$*" >&6
  CONFIG_SHELL='$SHELL'
  export CONFIG_SHELL
  exec "\$@"
fi

_ACEOF
cat >>$CONFIG_STATUS <<\_ACEOF || ac_write_fail=1
exec 5>>config.log
{
  echo
  sed 'h;s/./-/g;s/^.../## /;s/...$/ ##/;p;x;p;x' <<_ASBOX
## Running $as_me. ##
_ASBOX
  $as_echo "$ac_log"
} >&5

_ACEOF
cat >>$CONFIG_STATUS <<_ACEOF || ac_write_fail=1
_ACEOF

cat >>$CONFIG_STATUS <<\_ACEOF || ac_write_fail=1

# Handling of arguments.
for ac_config_target in $ac_config_targets
do
  case $ac_config_target in
    "src/Makevars") CONFIG_FILES="$CONFIG_FILES src/Makevars" ;;

  *) as_fn_error $? "invalid argument: \`$ac_config_target'" "$LINENO" 5;;
  esac
done


# If the user did not use the arguments to specify the items to instantiate,
# then the envvar interface is used.  Set only those that are not.
# We use the long form for the default assignment because of an extremely
# bizarre bug on SunOS 4.1.3.
if $ac_need_defaults; then
  test "${CONFIG_FILES+set}" = set || CONFIG_FILES=$config_files
fi

# Have a temporary directory for convenience.  Make it in the build tree
# simply because there is no reason against having it here, and in addition,
# creating and moving files from /tmp can sometimes cause problems.
# Hook for its removal unless debugging.
# Note that there is a small window in which the directory will not be cleaned:
# after its creation but before its name has been assigned to `$tmp'.
$debug ||
{
  tmp= ac_tmp=
  trap 'exit_status=$?
  : "${ac_tmp:=$tmp}"
  { test ! -d "$ac_tmp" || rm -fr "$ac_tmp"; } && exit $exit_status
' 0
  trap 'as_fn_exit 1' 1 2 13 15
}
# Create a (secure) tmp directory for tmp files.

{
  tmp=`(umask 077 && mktemp -d "./confXXXXXX") 2>/dev/null` &&
  test -d "$tmp"
}  ||
{
  tmp=./conf$$-$RANDOM
  (umask 077 && mkdir "$tmp")
} || as_fn_error $? "cannot create a temporary directory in ." "$LINENO" 5
ac_tmp=$tmp

# Set up the scripts for CONFIG_FILES section.
# No need to generate them if there are no CONFIG_FILES.
# This happens for instance with `./config.status config.h'.
if test -n "$CONFIG_FILES"; then


ac_cr=`echo X | tr X '\015'`
# On cygwin, bash can eat \r inside `` if the user requested igncr.
# But we know of no other shell where ac_cr would be empty at this
# point, so we can use a bashism as a fallback.
if test "x$ac_cr" = x; then
  eval ac_cr=\$\'\\r\'
fi
ac_cs_awk_cr=`$AWK 'BEGIN { print "a\rb" }' </dev/null 2>/dev/null`
if test "$ac_cs_awk_cr" = "a${ac_cr}b"; then
  ac_cs_awk_cr='\\r'
else
  ac_cs_awk_cr=$ac_cr
fi

echo 'BEGIN {' >"$ac_tmp/subs1.awk" &&
_ACEOF


{
  echo "cat >conf$$subs.awk <<_ACEOF" &&
  echo "$ac_subst_vars" | sed 's/.*/&!$&$ac_delim/' &&
  echo "_ACEOF"
} >conf$$subs.sh ||
  as_fn_error $? "could not make $CONFIG_STATUS" "$LINENO" 5
ac_delim_num=`echo "$ac_subst_vars" | grep -c '^'`
ac_delim='%!_!# '
for ac_last_try in false false false false false :; do
  . ./conf$$subs.sh ||
    as_fn_error $? "could not make $CONFIG_STATUS" "$LINENO" 5

  ac_delim_n=`sed -n "s/.*$ac_delim\$/X/p" conf$$subs.awk | grep -c X`
  if test $ac_delim_n = $ac_delim_num; then
    break
  elif $ac_last_try; then
    as_fn_error $? "could not make $CONFIG_STATUS" "$LINENO" 5
  else
    ac_delim="$ac_delim!$ac_delim _$ac_delim!! "
  fi
done
rm -f conf$$subs.sh

cat >>$CONFIG_STATUS <<_ACEOF || ac_write_fail=1
cat >>"\$ac_tmp/subs1.awk" <<\\_ACAWK &&
_ACEOF
sed -n '
h
s/^/S["/; s/!.*/"]=/
p
g
s/^[^!]*!//
:repl
t repl
s/'"$ac_delim"'$//
t delim
:nl
h
s/\(.\{148\}\)..*/\1/
t more1
s/["\\]/\\&/g; s/^/"/; s/$/\\n"\\/
p
n
b repl
:more1
s/["\\]/\\&/g; s/^/"/; s/$/"\\/
p
g
s/.\{148\}//
t nl
:delim
h
s/\(.\{148\}\)..*/\1/
t more2
s/["\\]/\\&/g; s/^/"/; s/$/"/
p
b
:more2
s/["\\]/\\&/g; s/^/"/; s/$/"\\/
p
g
s/.\{148\}//
t delim
' <conf$$subs.awk | sed '
/^[^""]/{
  N
  s/\n//
}
' >>$CONFIG_STATUS || ac_write_fail=1
rm -f conf$$subs.awk
cat >>$CONFIG_STATUS <<_ACEOF || ac_write_fail=1
_ACAWK
cat >>"\$ac_tmp/subs1.awk" <<_ACAWK &&
  for (key in S) S_is_set[key] = 1
  FS = ""

}
{
  line = $ 0
  nfields = split(line, field, "@")
  substed = 0
  len = length(field[1])
  for (i = 2; i < nfields; i++) {
    key = field[i]
    keylen = length(key)
    if (S_is_set[key]) {
      value = S[key]
      line = substr(line, 1, len) "" value "" substr(line, len + keylen + 3)
      len += length(value) + length(field[++i])
      substed = 1
    } else
      len += 1 + keylen
  }

  print line
}

_ACAWK
_ACEOF
cat >>$CONFIG_STATUS <<\_ACEOF || ac_write_fail=1
if sed "s/$ac_cr//" < /dev/null > /dev/null 2>&1; then
  sed "s/$ac_cr\$//; s/$ac_cr/$ac_cs_awk_cr/g"
else
  cat
fi < "$ac_tmp/subs1.awk" > "$ac_tmp/subs.awk" \
  || as_fn_error $? "could not setup config files machinery" "$LINENO" 5
_ACEOF

# VPATH may cause trouble with some makes, so we remove sole $(srcdir),
# ${srcdir} and @srcdir@ entries from VPATH if srcdir is ".", strip leading and
# trailing colons and then remove the whole line if VPATH becomes empty
# (actually we leave an empty line to preserve line numbers).
if test "x$srcdir" = x.; then
  ac_vpsub='/^[	 ]*VPATH[	 ]*=[	 ]*/{
h
s///
s/^/:/
s/[	 ]*$/:/
s/:\$(srcdir):/:/g
s/:\${srcdir}:/:/g
s/:@srcdir@:/:/g
s/^:*//
s/:*$//
x
s/\(=[	 ]*\).*/\1/
G
s/\n//
s/^[^=]*=[	 ]*$//
}'
fi

cat >>$CONFIG_STATUS <<\_ACEOF || ac_write_fail=1
fi # test -n "$CONFIG_FILES"


eval set X "  :F $CONFIG_FILES      "
shift
for ac_tag
do
  case $ac_tag in
  :[FHLC]) ac_mode=$ac_tag; continue;;
  esac
  case $ac_mode$ac_tag in
  :[FHL]*:*);;
  :L* | :C*:*) as_fn_error $? "invalid tag \`$ac_tag'" "$LINENO" 5;;
  :[FH]-) ac_tag=-:-;;
  :[FH]*) ac_tag=$ac_tag:$ac_tag.in;;
  esac
  ac_save_IFS=$IFS
  IFS=:
  set x $ac_tag
  IFS=$ac_save_IFS
  shift
  ac_file=$1
  shift

  case $ac_mode in
  :L) ac_source=$1;;
  :[FH])
    ac_file_inputs=
    for ac_f
    do
      case $ac_f in
      -) ac_f="$ac_tmp/stdin";;
      *) # Look for the file first in the build tree, then in the source tree
	 # (if the path is not absolute).  The absolute path cannot be DOS-style,
	 # because $ac_f cannot contain `:'.
	 test -f "$ac_f" ||
	   case $ac_f in
	   [\\/$]*) false;;
	   *) test -f "$srcdir/$ac_f" && ac_f="$srcdir/$ac_f";;
	   esac ||
	   as_fn_error 1 "cannot find input file: \`$ac_f'" "$LINENO" 5;;
      esac
      case $ac_f in *\'*) ac_f=`$as_echo "$ac_f" | sed "s/'/'\\\\\\\\''/g"`;; esac
      as_fn_append ac_file_inputs " '$ac_f'"
    done

    # Let's still pretend it is `configure' which instantiates (i.e., don't
    # use $as_me), people would be surprised to read:
    #    /* config.h.  Generated by config.status.  */
    configure_input='Generated from '`
	  $as_echo "$*" | sed 's|^[^:]*/||;s|:[^:]*/|, |g'
	`' by configure.'
    if test x"$ac_file" != x-; then
      configure_input="$ac_file.  $configure_input"
      { $as_echo "$as_me:${as_lineno-$LINENO}: creating $ac_file" >&5
$as_echo "$as_me: creating $ac_file" >&6;}
    fi
    # Neutralize special characters interpreted by sed in replacement strings.
    case $configure_input in #(
    *\&* | *\|* | *\\* )
       ac_sed_conf_input=`$as_echo "$configure_input" |
       sed 's/[\\\\&|]/\\\\&/g'`;; #(
    *) ac_sed_conf_input=$configure_input;;
    esac

    case $ac_tag in
    *:-:* | *:-) cat >"$ac_tmp/stdin" \
      || as_fn_error $? "could not create $ac_file" "$LINENO" 5 ;;
    esac
    ;;
  esac

  ac_dir=`$as_dirname -- "$ac_file" ||
$as_expr X"$ac_file" : 'X\(.*[^/]\)//*[^/][^/]*/*$' \| \
	 X"$ac_file" : 'X\(//\)[^/]' \| \
	 X"$ac_file" : 'X\(//\)$' \| \
	 X"$ac_file" : 'X\(/\)' \| . 2>/dev/null ||
$as_echo X"$ac_file" |
    sed '/^X\(.*[^/]\)\/\/*[^/][^/]*\/*$/{
	    s//\1/
	    q
	  }
	  /^X\(\/\/\)[^/].*/{
	    s//\1/
	    q
	  }
	  /^X\(\/\/\)$/{
	    s//\1/
	    q
	  }
	  /^X\(\/\).*/{
	    s//\1/
	    q
	  }
	  s/.*/./; q'`
  as_dir="$ac_dir"; as_fn_mkdir_p
  ac_builddir=.

case "$ac_dir" in
.) ac_dir_suffix= ac_top_builddir_sub=. ac_top_build_prefix= ;;
*)
  ac_dir_suffix=/`$as_echo "$ac_dir" | sed 's|^\.[\\/]||'`
  # A ".." for each directory in $ac_dir_suffix.
  ac_top_builddir_sub=`$as_echo "$ac_dir_suffix" | sed 's|/[^\\/]*|/..|g;s|/||'`
  case $ac_top_builddir_sub in
  "") ac_top_builddir_sub=. ac_top_build_prefix= ;;
  *)  ac_top_build_prefix=$ac_top_builddir_sub/ ;;
  esac ;;
esac
ac_abs_top_builddir=$ac_pwd
ac_abs_builddir=$ac_pwd$ac_dir_suffix
# for backward compatibility:
ac_top_builddir=$ac_top_build_prefix

case $srcdir in
  .)  # We are building in place.
    ac_srcdir=.
    ac_top_srcdir=$ac_top_builddir_sub
    ac_abs_top_srcdir=$ac_pwd ;;
  [\\/]* | ?:[\\/]* )  # Absolute name.
    ac_srcdir=$srcdir$ac_dir_suffix;
    ac_top_srcdir=$srcdir
    ac_abs_top_srcdir=$srcdir ;;
  *) # Relative name.
    ac_srcdir=$ac_top_build_prefix$srcdir$ac_dir_suffix
    ac_top_srcdir=$ac_top_build_prefix$srcdir
    ac_abs_top_srcdir=$ac_pwd/$srcdir ;;
esac
ac_abs_srcdir=$ac_abs_top_srcdir$ac_dir_suffix


  case $ac_mode in
  :F)
  #
  # CONFIG_FILE
  #

_ACEOF

cat >>$CONFIG_STATUS <<\_ACEOF || ac_write_fail=1
# If the template does not know about datarootdir, expand it.
# FIXME: This hack should be removed a few years after 2.60.
ac_datarootdir_hack=; ac_datarootdir_seen=
ac_sed_dataroot='
/datarootdir/ {
  p
  q
}
/@datadir@/p
/@docdir@/p
/@infodir@/p
/@localedir@/p
/@mandir@/p'
case `eval "sed -n \"\$ac_sed_dataroot\" $ac_file_inputs"` in
*datarootdir*) ac_datarootdir_seen=yes;;
*@datadir@*|*@docdir@*|*@infodir@*|*@localedir@*|*@mandir@*)
  { $as_echo "$as_me:${as_lineno-$LINENO}: WARNING: $ac_file_inputs seems to ignore the --datarootdir setting" >&5
$as_echo "$as_me: WARNING: $ac_file_inputs seems to ignore the --datarootdir setting" >&2;}
_ACEOF
cat >>$CONFIG_STATUS <<_ACEOF || ac_write_fail=1
  ac_datarootdir_hack='
  s&@datadir@&$datadir&g
  s&@docdir@&$docdir&g
  s&@infodir@&$infodir&g
  s&@localedir@&$localedir&g
  s&@mandir@&$mandir&g
  s&\\\${datarootdir}&$datarootdir&g' ;;
esac
_ACEOF

# Neutralize VPATH when `$srcdir' = `.'.
# Shell code in configure.ac might set extrasub.
# FIXME: do we really want to maintain this feature?
cat >>$CONFIG_STATUS <<_ACEOF || ac_write_fail=1
ac_sed_extra="$ac_vpsub
$extrasub
_ACEOF
cat >>$CONFIG_STATUS <<\_ACEOF || ac_write_fail=1
:t
/@[a-zA-Z_][a-zA-Z_0-9]*@/!b
s|@configure_input@|$ac_sed_conf_input|;t t
s&@top_builddir@&$ac_top_builddir_sub&;t t
s&@top_build_prefix@&$ac_top_build_prefix&;t t
s&@srcdir@&$ac_srcdir&;t t
s&@abs_srcdir@&$ac_abs_srcdir&;t t
s&@top_srcdir@&$ac_top_srcdir&;t t
s&@abs_top_srcdir@&$ac_abs_top_srcdir&;t t
s&@builddir@&$ac_builddir&;t t
s&@abs_builddir@&$ac_abs_builddir&;t t
s&@abs_top_builddir@&$ac_abs_top_builddir&;t t
$ac_datarootdir_hack
"
eval sed \"\$ac_sed_extra\" "$ac_file_inputs" | $AWK -f "$ac_tmp/subs.awk" \
  >$ac_tmp/out || as_fn_error $? "could not create $ac_file" "$LINENO" 5

test -z "$ac_datarootdir_hack$ac_datarootdir_seen" &&
  { ac_out=`sed -n '/\${datarootdir}/p' "$ac_tmp/out"`; test -n "$ac_out"; } &&
  { ac_out=`sed -n '/^[	 ]*datarootdir[	 ]*:*=/p' \
      "$ac_tmp/out"`; test -z "$ac_out"; } &&
  { $as_echo "$as_me:${as_lineno-$LINENO}: WARNING: $ac_file contains a reference to the variable \`datarootdir'
which seems to be undefined.  Please make sure it is defined" >&5
$as_echo "$as_me: WARNING: $ac_file contains a reference to the variable \`datarootdir'
which seems to be undefined.  Please make sure it is defined" >&2;}

  rm -f "$ac_tmp/stdin"
  case $ac_file in
  -) cat "$ac_tmp/out" && rm -f "$ac_tmp/out";;
  *) rm -f "$ac_file" && mv "$ac_tmp/out" "$ac_file";;
  esac \
  || as_fn_error $? "could not create $ac_file" "$LINENO" 5
 ;;


  esac

done # for ac_tag


as_fn_exit 0
_ACEOF
ac_clean_files=$ac_clean_files_save

test $ac_write_fail = 0 ||
  as_fn_error $? "write failure creating $CONFIG_STATUS" "$LINENO" 5


# configure is writing to config.log, and then calls config.status.
# config.status does its own redirection, appending to config.log.
# Unfortunately, on DOS this fails, as config.log is still kept open
# by configure, so config.status won't be able to write to it; its
# output is simply discarded.  So we exec the FD to /dev/null,
# effectively closing config.log, so it can be properly (re)opened and
# appended to by config.status.  When coming back to configure, we
# need to make the FD available again.
if test "$no_create" != yes; then
  ac_cs_success=:
  ac_config_status_args=
  test "$silent" = yes &&
    ac_config_status_args="$ac_config_status_args --quiet"
  exec 5>/dev/null
  $SHELL $CONFIG_STATUS $ac_config_status_args || ac_cs_success=false
  exec 5>>config.log
  # Use ||, not &&, to avoid exiting from the if with $? = 1, which
  # would make configure fail if this is the last instruction.
  $ac_cs_success || as_fn_exit 1
fi
if test -n "$ac_unrecognized_opts" && test "$enable_option_checking" != no; then
  { $as_echo "$as_me:${as_lineno-$LINENO}: WARNING: unrecognized options: $ac_unrecognized_opts" >&5
$as_echo "$as_me: WARNING: unrecognized options: $ac_unrecognized_opts" >&2;}
fi
