## Supplementary material for "MoBPS - Modular Breeding Program Simulator": miraculix R-package: DESCRIPTION

Package: miraculix
Version: 0.9.7
Title: Algebraic and Statistical Functions for Genetics
Author: Martin Schlather [aut, cre], Malena Erbe [aut, cre], Florian Skene [aut], Alexander Freudenberg [ctr]
Description: This is a collection of fast tools for application in quantitative genetics. For instance, the SNP matrix can be stored in a minimum of memory  and the calculation of the genomic relationship matrix is based on a rapid algorithm. It also contains the window scanning approach by Kabluchko and Spodarev (2009), <doi:10.1239/aap/1240319575> to detect anomalous genomic areas <doi:10.1186/s12864-018-5009-y>. Furthermore, the package is used in the Modular Breeding Program Simulator (MoBPS, <https://github.com/tpook92/MoBPS>, <http://www.mobps.de/>). The tools are based on SIMD (Single Instruction Multiple Data, <https://en.wikipedia.org/wiki/SIMD>) and OMP (Open Multi-Processing, <https://de.wikipedia.org/wiki/OpenMP>).  
Maintainer: Martin Schlather <>
LinkingTo: RandomFieldsUtils
Depends: R (>= 3.0), RandomFieldsUtils (>= 0.5.9)
Imports: methods, graphics
Suggests:
License: GPL ( >= 3)
Biarch: true
URL: http://ms.math.uni-mannheim.de/de/publications/software/miraculix
NeedsCompilation: yes
Packaged: 2019-10-28 21:19:47 UTC; schlather
