## Supplementary material for "MoBPS - Modular Breeding Program Simulator": miraculix R-package: aaa_auto.R

```
# This file has been created automatically by 'rfGenerateConstants'


 ## from  ../../RandomFieldsUtils/RandomFieldsUtils/src/AutoRandomFieldsUtils.h

 MAXUNITS 	<- as.integer(4)
 MAXCHAR 	<- as.integer(18)
 RFOPTIONS 	<- "RFoptions"
 isGLOBAL 	<- as.integer(NA)

 WARN_UNKNOWN_OPTION_ALL 	<- as.integer(3)
 WARN_UNKNOWN_OPTION_SINGLE 	<- as.integer(2)
 WARN_UNKNOWN_OPTION_CAPITAL 	<- as.integer(1)
 WARN_UNKNOWN_OPTION_NONE 	<- as.integer(0)
 WARN_UNKNOWN_OPTION 	<- as.integer(10000)
 WARN_UNKNOWN_OPTION_CONDSINGLE 	<- as.integer((WARN_UNKNOWN_OPTION_SINGLE-WARN_UNKNOWN_OPTION))
 WARN_UNKNOWN_OPTION_DEFAULT 	<- as.integer(WARN_UNKNOWN_OPTION_ALL)


 ## from  src/AutoMiraculix.h

 AutoMiraculix_H 	<- as.integer(1)

 Shuffle 	<- as.integer(0)
 Shuffle256 	<- as.integer(1)
 TwoBit 	<- as.integer(2)
 ThreeBit 	<- as.integer(3)
 Hamming2 	<- as.integer(4)
 Hamming3 	<- as.integer(5)
 NoSNPcoding 	<- as.integer(6)
 NoSNPcodingR 	<- as.integer(7)
 AutoCoding 	<- as.integer(8)
 Haplo 	<- as.integer(9)


 last_usr_meth 	<- as.integer(AutoCoding)
 nr_snpcoding 	<- as.integer((Haplo+1))

 HAPLO 	<- as.integer(0)

 GENOMATRIX 	<- as.integer(1)
 LAST_WHAT 	<- as.integer(1)

 WHAT 	<- as.integer(0)
 SNPS 	<- as.integer(1)
 INDIVIDUALS 	<- as.integer(2)
 ADDR0 	<- as.integer(3)
 ADDR1 	<- as.integer(4)
 ALIGNADDR0 	<- as.integer(5)
 ALIGNADDR1 	<- as.integer(6)
 SUMGENO 	<- as.integer(7)
 SUMGENO_E9 	<- as.integer(8)

 SNPxIND 	<- as.integer(10)
 BITSPERCODE 	<- as.integer(12)
 BYTESPERBLOCK 	<- as.integer(13)
 CODESPERBLOCK 	<- as.integer(14)
 HEADER 	<- as.integer(15)
 DOUBLEINDIV 	<- as.integer(16)
 LEADINGCOL 	<- as.integer(17)
 MEMinUNITS0 	<- as.integer(18)

 MEMinUNITS1 	<- as.integer(19)
 ALIGNEDUNITS0 	<- as.integer(20)
 ALIGNEDUNITS1 	<- as.integer(21)

 INFO_LAST 	<- as.integer(ALIGNEDUNITS1)

 GENOMICMATRIX 	<- "genomicmatrix"
 HAPLOMATRIX 	<- "haplomatrix"
 ORIGINVECTOR 	<- "origindata"


 ## from  src/AutoMiraculix.cc

 SNPCODING_NAME <-
c( "Shuffle","Shuffle256","TwoBit","ThreeBit","Hamming2","Hamming3","NoSNPcoding","NoSNPcodingR","AutoCoding","Haplo" )


 WHAT_NAMES <-
c( "haplo vector","geno matrix" )
```
