## Supplementary material for "MoBPS - Modular Breeding Program Simulator": miraculix R-package: auxiliary.R

```
## Authors 
## Martin Schlather,
##
##
## Copyright (C) 2019 -- 2019 Martin Schlather
##
## This program is free software; you can redistribute it and/or
## modify it under the terms of the GNU General Public License
## as published by the Free Software Foundation; either version 3
## of the License, or (at your option) any later version.
##
## This program is distributed in the hope that it will be useful,
## but WITHOUT ANY WARRANTY; without even the implied warranty of
## MERCHANTABILITY or FITNESS FOR A PARTICULAR PURPOSE.  See the
## GNU General Public License for more details.
##
## You should have received a copy of the GNU General Public License
## along with this program; if not, write to the Free Software
## Foundation, Inc., 59 Temple Place - Suite 330, Boston, MA  02111-1307, USA.  

has.instruction.set <- function(which=c("SSE2", "SSSE3", "AVX", "AVX2")) {
  ans <- logical(length(which))
  for (i in 1:length(ans)) {
    ans[i] <-
      switch(which[i],
             ##  "MMX" = .Call(C_hasMMX),
             "SSE2" = .Call(C_hasSSE2),
             "SSSE3" = .Call(C_hasSSSE3),
             "AVX" = .Call(C_hasAVX),
             "AVX2" = .Call(C_hasAVX2),
             FALSE)
  }
  ans
}


dolocking <-function(do=TRUE) {
  if (length(do) > 0 && do) stop("sorry dolocking has not been programmed yet")
  ##.Call(C_dolocking, do)
  return(FALSE)
}


## RM, SAVE, PB, RETURN, DEBUG, unlock deleted 
```
