## Supplementary material for "MoBPS - Modular Breeding Program Simulator": miraculix R-package: debugging.R

```
## function for debugging only

SAVE <- function(name, ans) {
  print(name) ##
  variable.name <- "__Z"
  ENV <- .GlobalEnv
  name <- paste(variable.name, name, sep=".")
  name <- name[length(name)]
  if (!exists(name, envir=ENV)) assign(name, 0, envir=ENV)
  cur <- get(name, envir=ENV) + 1
  file <- paste(name, cur, "rda", sep=".")
  Print(file, ans) ##
  assign(name, cur, envir=ENV)
  save (file=file, ans) 
  return(ans)
}

RETURN <- function(name) {
  variable.name <- "__Z"
  ENV <- .GlobalEnv
  name <- paste(variable.name, name, sep=".")
  name <- name[length(name)]
  if (!exists(name, envir=ENV)) assign(name, 0, envir=ENV)
  cur <- get(name, envir=ENV) + 1
  file <- paste(name, cur, "rda", sep=".")  
  if (file.exists(file)) {
    cat("playback", file, "\n")
    load (file) 
    assign(name, cur, envir=ENV)
    return(name)
  }
  stop("hierher und nicht weiter")
}

PB <- function(name) {
  variable.name <- "__Z"
  ENV <- .GlobalEnv
  name <- paste(variable.name, name, sep=".")
  name <- name[length(name)]
  if (!exists(name, envir=ENV)) assign(name, 0, envir=ENV)
  cur <- get(name, envir=ENV) + 1
  file <- paste(name, cur, "rda", sep=".")  
  return(file.exists(file))
}


DEBUG <- function(name, population, gen, sex, nr) {
  variable.name <- "__Z"
  ENV <- .GlobalEnv
  name <- paste(variable.name, name, sep=".")
  name <- name[length(name)]
  if (!exists(name, envir=ENV)) assign(name, 0, envir=ENV)
  cur <- get(name, envir=ENV) + 1
  file <- paste(name, cur, "rda", sep=".")  
  if (cur == 2866) {
    .Call(C_Debug);
##    str(population$breeding[[1]][[1]][[318]])
##    str(population$breeding[[gen]][[sex]][[nr]])
##    print(population$breeding[[gen]][[sex]][[nr]][1:2])
  } # else .Call(C_StopDebug);
}


## only for development of rmCoded needed
## check: how to remove an object that has been defined on the upper level
## no further use
RM <- function(...) {
  env <- parent.env(environment())
  L <- list(...)
  if (is.character(L[[1]])) {
    if (!all(sapply(L, is.character)))
      stop("arguments must be all variables or all character vectors")
    L <- unlist(L)
  } else {
    L <- as.character(substitute(list(...)))[-1]
  }
  rm(list=L, envir=env)
}


#########################################################
## from below here are only temporary functions

unlock <- function(...,  envir = as.environment(-1)) {
  L <- list(...)
  if (!all(sapply(L, function(x) length(attr(x, "information")) > 0)))
    stop("not all arguments are coded objects (Note that 'codeOrigin' formally does return a coded object. So these objects cannot be unlock-ed.");
 if (is.character(L[[1]])) {
    if (!all(sapply(L, is.character)))
      stop("arguments must be all variables or all character vectors")
    L <- unlist(L)
  } else {
    L <- as.character(substitute(list(...)))[-1]
  }
  for (i in L)  .Call(C_unlock, get(i, envir=envir))
}
```
