## Supplementary material for "MoBPS - Modular Breeding Program Simulator": miraculix R-package: haplogeno.R

```
## Authors 
## Martin Schlather,
##
##
## Copyright (C) 2017 -- 2019 Martin Schlather
##
## This program is free software; you can redistribute it and/or
## modify it under the terms of the GNU General Public License
## as published by the Free Software Foundation; either version 3
## of the License, or (at your option) any later version.
##
## This program is distributed in the hope that it will be useful,
## but WITHOUT ANY WARRANTY; without even the implied warranty of
## MERCHANTABILITY or FITNESS FOR A PARTICULAR PURPOSE.  See the
## GNU General Public License for more details.
##
## You should have received a copy of the GNU General Public License
## along with this program; if not, write to the Free Software
## Foundation, Inc., 59 Temple Place - Suite 330, Boston, MA  02111-1307, USA.  


ENV <- environment()


decodeHaplo <- function(CM, indiv=integer(0),
                        sets=1:2,
                        IndividualsPerColumn=TRUE,
                        DoubledIndividuals=TRUE) {## method checked in C
 
  if (!is(CM, HAPLOMATRIX)) stop("matrix not a '", HAPLOMATRIX, "'")

  .Call(C_decodeHaplo, CM, as.integer(indiv), as.integer(sets),
        IndividualsPerColumn, DoubledIndividuals)
}


## code habpo-type efficiently (for original generation)
haplomatrix <- function(M,  # N,
                        IndividualsPerColumn=TRUE, DoubledIndividuals=TRUE) {
  ##%item{N}{vector of integers, which gives the selected rows. If missing all rows are selected.}
  ## if (!missing(N)) .Call(C_codeHaplo2, M, N, IndividualsPerColumn) else
  .Call(C_codeHaplo, M,  IndividualsPerColumn, DoubledIndividuals)
}


## geno functions
haplo2geno <- function(CM) {## method checked in C
  if (dolocking(NULL))
    stop("if 'dolocking' then implicite re-coding is not possible.")
  if (!is(CM, HAPLOMATRIX)) stop("matrix not a '", HAPLOMATRIX, "'")
  .Call(C_haplo2geno, CM)  # CM coded geno matrix
}


decodeGeno <- function(SNPxIndiv, snps, do.centering=FALSE) {
  if (!missing(snps))
    return(decodeNthGeno(SNPxIndiv, snps=snps, do.centering=do.centering))

  if (is(SNPxIndiv, GENOMICMATRIX)) {
     
    if (is.character(SNPxIndiv)) {
      snpXindiv <-.Call(C_file_get, SNPxIndiv)
      attr(snpXindiv, "filename") <- as.character(SNPxIndiv)
    } else {
      if (attr(SNPxIndiv, "method") == NoSNPcodingR) {
        snpXindiv <- SNPxIndiv
        storage.mode(snpXindiv) <- "integer"
      } else snpXindiv <-.Call(C_matrix_get, SNPxIndiv)
    }
  } else if (is.matrix(SNPxIndiv)) {
    snpXindiv <- SNPxIndiv
    d <- dim(snpXindiv)
    base::attributes(snpXindiv) <- NULL
    dim(snpXindiv) <- d
  } else stop("unknown format")

  if (do.centering) {
    centered <- RFoptions(GETOPTIONS="genetics")$centered      
    if (is.na(centered)) return(.Call(C_substract_centered, snpXindiv))
    else if (centered) return(snpXindiv - rowMeans(snpXindiv))
  }
  
  return(snpXindiv)
}

copyGeno <-function(SNPxIndiv) {
  if (is.matrix(SNPxIndiv)) return(SNPxIndiv + 0L)
  stopifnot(is(SNPxIndiv, GENOMICMATRIX))
  if (is.character(SNPxIndiv)) return(paste0("", SNPxIndiv))
  if (attr(SNPxIndiv, "method") == NoSNPcodingR) return(SNPxIndiv + 0L)
  .Call(C_copyGeno, SNPxIndiv)
}


Coding <-c(beagle="AB? ", plink="AB? ", plink2="12? ", plinkbinary="12345")
Header <-    c(beagle=1,    plink=0,    plink2=0,     plinkbinary=-3)
IndivPerCol <- c(beagle=TRUE, plink=TRUE, plink2=FALSE, plinkbinary=TRUE)
DoubledIndiv <-c(beagle=TRUE, plink=TRUE, plink2=FALSE, plinkbinary=FALSE)
LeadingCol <-c(beagle=2,    plink=4,    plink2=6,     plinkbinary=0)
Extension <- c(beagle="bgl", plink="phased", plink2="ped", plinkbinary="bed")


genomicmatrix <- function(snps, individuals, file.type,
                          coding, header, IndividualsPerColumn,
                          DoubledIndividuals, leadingcolumns,
                          loading = TRUE,
                          ...) {
  if (dolocking(NULL))
    stop("if 'dolocking' then implicite coding is not possible.")

  if (is(snps, GENOMICMATRIX)) {
    warning("'snps' is already of class '", GENOMICMATRIX,
            "'. Its original definition is kept.");
    if (length(list(match.call())) > 2)
      stop("further arguments may not be given")
    return(snps)
  }

  info <- NULL
  if (is(snps, HAPLOMATRIX)) {
    if (is.character(snps)) info <- attr(snps, "information") ## treat as if
    ##                                         file name + file.type were given
    else {
      if (length(list(match.call())) > 2)
        stop("further arguments may not be given")
      return(haplo2geno(snps))
    }
  }
  
  if (hasArg("individuals")) {
    if (length(as.list(match.call())) != 3)
      stop("if 'individuals' is given exactly 'snps' and 'individuals' must be given.")
     if (!is.numeric(snps) || length(snps) != 1)
      stop("'snps' and 'individuals' must be integers that give the size of the SNP matrix.")

    return (.Call(C_createSNPmatrix, as.integer(snps), as.integer(individuals)))
  }
 
  if (length(list(...)) > 0) {
    RFoptOld <- RFoptions(##LOCAL=1,
                          SAVEOPTIONS="genetics", ...)
    on.exit(RFoptions(LIST=RFoptOld))
  }
  RFopt <- RFoptions(GETOPTIONS="genetics")
  method <- RFopt$snpcoding
  if (method == AutoCoding) method <- .Call(C_getAutoCoding)

  if (is.numeric(snps) || is.logical(snps)) {
    if (method == NoSNPcodingR) {
      sumgeno <- sum(snps)
      mem <- length(snps)
      info <- c(what = GENOMATRIX,              ## 0 
                snps = nrow(snps),      ## 1
                individuals=ncol(snps),
                addr0=0,
                addr1=0,
                align=0,                ## 5
                align1=0,
                sumgeno= as.integer(sumgeno %% 1e9),
                sumgenoE9=as.integer(sumgeno / 1e9),
                unused = NA, ###
                unused2 = NA,
                isSNPxInd = TRUE,       ## 9
                bitspercode = 32,
                bytesperblock = 4,
                codesperblock = 1,
                header=NA,              ## 15
                DoubledIndividuals=NA,
                leadingcolumns=NA,
                memInUnits0 = as.integer(mem %% 1e9),
                meminUnits1 = as.integer(mem / 1e9),
                AlignedUnits0 = as.integer(mem %% 1e9),
                AlignedUnits1 = as.integer(mem / 1e9)
                )
      stopifnot(length(info) == INFO_LAST + 1)
      storage.mode(info) <- "integer"
      attr(snps, "information") <- info
      ##attr(snps, "coding") =  as.character(NULL)
      attr(snps, "method") <- method
      class(snps) <- GENOMICMATRIX
  } else snps <- .Call(C_matrix_coding, snps)
    return(snps)
  }

  if (!is.character(snps)) stop("'snps' is expected to be a file name.")
  object <- snps
  individuals <- snps <- NA
  if (missing(file.type)) {
    split <- strsplit(object, "\\.")[[1]]
    file.ext <-  c("txt", "bgl", "phased", "tped", "ped", "bed")
    s <- pmatch(split[length(split)], file.ext)
    if (!is.na(s)) { 
      file.type <- c(txt="beagle", bgl="beagle", phased="plink",
                     tped="plink2", ped="plink2",
                     bed="plinkbinary")[file.ext[s]]
    } else {
      message("Note: file.type could not be detected.")
    }
  }
  
  
  if (!missing(file.type)) {
    if (missing(coding)) coding <- Coding[file.type]
    if (missing(header)) header <- Header[file.type]
    if (missing(IndividualsPerColumn)) IndividualsPerColumn <- IndivPerCol[file.type]
    if (missing(DoubledIndividuals)) DoubledIndividuals <- DoubledIndiv[file.type]
    if (missing(leadingcolumns)) leadingcolumns <- LeadingCol[file.type]
    
    if (file.type == "plinkbinary") {
      s <- paste(split[-length(split)], collapse=".")
      
      file.lines <- function(object) {
        sys <- strsplit(system(paste("wc", object), intern=TRUE), " ")[[1]]
        sys <- sys[nchar(sys) > 0]
        as.integer(sys[1])
      }
      
      ##  #lines of the files give the snps or individuals
      individuals <- file.lines(paste(s, ".fam", sep=""))
      snps <- file.lines(paste(s, ".bim", sep=""))
    }
  }
  
  if (nchar(coding) != 4 && nchar(coding) != 5)
    stop("haplotype must be given as 4 characters, e.g. 'AB? '; genomic information 5 characters")
  if (nchar(coding) == 4) {
    ## never change this coding
    coding <- paste(substr(coding, 1, 2), substr(coding, 2, 4), sep="")
  }
  
  if (missing(coding) || missing(header) || missing(IndividualsPerColumn) ||
      missing(DoubledIndividuals) || missing(leadingcolumns)) {
    if (missing(coding) && missing(header) && missing(IndividualsPerColumn) &&
        missing(DoubledIndividuals) && missing(leadingcolumns))
      stop("unknown file.type. Please give the file.type")
    stopifnot(!missing(coding))
    stopifnot(!missing(header))
    stopifnot(!missing(IndividualsPerColumn))
    stopifnot(!missing(DoubledIndividuals))
    stopifnot(!missing(leadingcolumns))
  }

  object <- filename <- as.character(object)
  if (length(info) == 0)
    info <- c(
        what = NA,
        snps = as.integer(snps),# get rid of any attributes
        individuals=as.integer(individuals),# get rid of any attributes
        addr0=0,
        addr1=0,
        align=0,
        align1=0,
        sumgeno=0,
        sumgenoE9=0,
        unused = NA, 
        unused2 = NA,
        isSNPxInd = as.integer(IndividualsPerColumn), # get rid of name attribute
        bitspercode = NA, 
        bytesperblock = NA,
        codesperblock = NA,
        header=header,
        DoubledIndividuals=as.integer(DoubledIndividuals),#get rid of attributes
        leadingcolumns=as.integer(leadingcolumns),#get rid of attributes
        memInUnits0 = NA,
        meminUnits1 = NA,
        AlignedUnits0 = NA,
        AlignedUnits1 = NA
    )
  stopifnot(length(info) == INFO_LAST + 1)
  storage.mode(info) <- "integer"
  attr(object, "information") <- info
  attr(object, "coding") <-  as.character(coding)
  if (!loading) {
    attr(object, "method") <- method
    class(object) <- GENOMICMATRIX
    return(object)
  }

  object <- .Call(C_file_get, object);
  attr(object, "filename") <-  filename
  return(object)
}


fillGeno <- function(SNPxIndiv, indiv, values,
                     IndividualsPerColumn=TRUE,
                     DoubledIndividuals=TRUE) {## method checked in C
  ## SNPxIndiv[, indiv] <- values
  ## note: values can be anything: haplo, geno, coded or uncoded
  stopifnot(is(SNPxIndiv, GENOMICMATRIX))
  if (is.character(SNPxIndiv)) stop("function not available for files")
  if (!is(values, GENOMICMATRIX)) {    
    if (is.numeric(values)) {
      haplo <- is.matrix(values) &&
        (( IndividualsPerColumn && ncol(values) == 2 * length(indiv)) ||
         (!IndividualsPerColumn && nrow(values) == 2 * length(indiv)))
      
      if (!haplo) values <- .Call(C_matrix_coding, values)
      else values <- haplomatrix(values,
                                 IndividualsPerColumn=IndividualsPerColumn,
                                 DoubledIndividuals=DoubledIndividuals)
    }
    if (is(values, HAPLOMATRIX)) values <- haplo2geno(values)
  }
  .Call(C_fillSNPmatrix, SNPxIndiv, as.integer(indiv), values)
}


relationshipMatrix <- function(SNPxIndiv, ...) {
  if (length(list(...)) > 0) {
    RFoptOld <- RFoptions(## LOCAL=1,
                          SAVEOPTIONS="genetics", ...)
    on.exit(RFoptions(LIST=RFoptOld))
  }
  RFopt <- RFoptions(GETOPTIONS = c("basic", "genetics"))

  centered <- RFopt$genetics$centered
  opt <- RFopt$basic
  

  if (!is(SNPxIndiv, GENOMICMATRIX) || is.character(SNPxIndiv))
    SNPxIndiv <- genomicmatrix(SNPxIndiv)
  method <- attr(SNPxIndiv, "method")

  if (opt$verbose) cat("method =", SNPCODING_NAME[method + 1],
		       "using", opt$cores, "core(s).\n")
  
  if (method == NoSNPcodingR) {
    p <- rowMeans(SNPxIndiv)
    if (!is.na(centered) && centered) SNPxIndiv <- SNPxIndiv - p
    if (RFopt$genetics$normalized)
      SNPxIndiv <- SNPxIndiv / sqrt(sum(p * (1 - p / 2)))
    A <- base::crossprod(SNPxIndiv)
  }

  else A <- .Call(C_matrix_mult, SNPxIndiv)

  if (is.na(centered)) {
    p  <- .Call(C_get_centered)
    pA <- vectorGeno(V=p, SNPxIndiv=SNPxIndiv, do.centering=FALSE)
    A <- A - pA - base::'%*%'(rep(1, length(pA)), t(pA)) + sum(p^2)
  }
  attr(A, "centered") <- centered
  return(A)
}


crossprodx <- function(SNPxIndiv) relationshipMatrix(SNPxIndiv, centered=FALSE)


vectorGeno <- function(V, SNPxIndiv, do.centering=FALSE, decode=TRUE) {
  if (do.centering) stop("centered not programmed yet")
  ## much faster than to operate on packed matrix
  ## but packed might be of interest for very large matrices.

  if (is.matrix(SNPxIndiv)) return(base::'%*%'(V, SNPxIndiv))
  stopifnot(is(SNPxIndiv, GENOMICMATRIX))
  method <- attr(SNPxIndiv, "method")
  if (is.character(SNPxIndiv)) return(.Call(C_dot_file, SNPxIndiv,as.double(V)))
  else if (method %in% c(Shuffle, Shuffle256) && !decode)
    return(.Call(C_vectorGeno, V, SNPxIndiv))
  
  as.vector(base::'%*%'(V, as.matrix(SNPxIndiv)))
}


genoVector <- function(SNPxIndiv, V, do.centering=FALSE) {
  if (do.centering) stop("centering not programmed yet")
  if (is.matrix(SNPxIndiv)) return(as.vector(base::'%*%'(SNPxIndiv, V)))
  stopifnot(is(SNPxIndiv, GENOMICMATRIX))
  method <- attr(SNPxIndiv, "method")
  if (is.character(SNPxIndiv)) return(.Call(C_file_dot, SNPxIndiv,as.double(V)))
  else if (method %in% c(Shuffle, Shuffle256)) {
    return(.Call(C_genoVector, SNPxIndiv, V))
  }
  base::'%*%'(as.matrix(SNPxIndiv), V)
}


solveRelMat <- function(A, tau, vec, betahat, destroy_A=FALSE) {
  ## if not destroyed, more memory is consumed
  ## implements ( t(Z) Z + tau \1_{ind x ind} )^{-1} %*% vec
  stopifnot(length(tau) == 1 && length(betahat) ==1)
  .Call(C_solveRelMat, A, as.double(tau), as.double(vec), as.double(betahat),
	destroy_A)
}

                
SNPeffect <- function(SNPxIndiv, vec, centered=TRUE, tau=0) {
  snpXind <- SNPxIndiv
  if (!is(snpXind, GENOMICMATRIX)) snpXind <- genomicmatrix(snpXind)

  rM <- relationshipMatrix(snpXind, centered=centered, normalized=FALSE)
  if (tau != 0) rM <- rM + diag(nrow(rM)) * tau ## ehemals eps
  rM <- solvex(rM, vec)

  genoVector(SNPxIndiv, rM)
}


allele_freq <- function(SNPxIndiv) {
  # name <- as.character(as.list(match.call(envir=parent.frame(2L)))[[1]])
  if (is.character(SNPxIndiv)) SNPxIndiv <- genomicmatrix(SNPxIndiv)
  if (is.matrix(SNPxIndiv)) return(rowMeans(SNPxIndiv) * 0.5)
  stopifnot(is(SNPxIndiv, GENOMICMATRIX))
  .Call(C_allele_freq, SNPxIndiv)
}


 #  else if (any(!is.finite(snps) | snps < 1))
 ##   stop("all values of 'snps' must be positive integers")
zeroNthGeno <- function(SNPxIndiv, snps) {
  if (missing(snps)) snps <- 1 : attr(SNPxIndiv, "information")[1 + SNPS]
  if (is.matrix(SNPxIndiv)) {
    SNPxIndiv[snps, ] <- 0L
    return(SNPxIndiv)
  }
  
  stopifnot(is(SNPxIndiv, GENOMICMATRIX))
  snps <- as.integer(snps - 1)
  snpXindiv <- copyGeno(SNPxIndiv)
  
  .Call(C_zeroNthGeno, snpXindiv, snps)
  snpXindiv 
}


#  if (any(!is.finite(snps) | snps < 1))
#    stop("all values of 'snps' must be positive integers")
decodeNthGeno <- function(SNPxIndiv, snps, do.centering=FALSE) {
  if (missing(snps)) snps <- 1 : attr(SNPxIndiv, "information")[1 + SNPS]
 if (is.matrix(SNPxIndiv)) return(SNPxIndiv[snps, ])

  stopifnot(is(SNPxIndiv, GENOMICMATRIX)) 
  snps <- as.integer(snps - 1)
  snpXindiv <-.Call(C_get_matrix_N, SNPxIndiv, snps)

  if (do.centering) {
    centered <- RFoptions(GETOPTIONS="genetics")$centered      
    if (is.na(centered)) return(.Call(C_substract_centered, snpXindiv))
    else if (centered) return(snpXindiv - rowMeans(snpXindiv))
  }
  return(snpXindiv)
}
```
