## Supplementary material for "MoBPS - Modular Breeding Program Simulator": miraculix R-package: MoBPS.R

```
## Authors 
## Martin Schlather,
##
##
## Copyright (C) 2017 -- 2019 Martin Schlather
## functions designed for package MoBPS
##
## This program is free software; you can redistribute it and/or
## modify it under the terms of the GNU General Public License
## as published by the Free Software Foundation; either version 3
## of the License, or (at your option) any later version.
##
## This program is distributed in the hope that it will be useful,
## but WITHOUT ANY WARRANTY; without even the implied warranty of
## MERCHANTABILITY or FITNESS FOR A PARTICULAR PURPOSE.  See the
## GNU General Public License for more details.
##
## You should have received a copy of the GNU General Public License
## along with this program; if not, write to the Free Software
## Foundation, Inc., 59 Temple Place - Suite 330, Boston, MA  02111-1307, USA.  


## code "all" information of the animals effectively
codeOrigins <- function(M) .Call(C_codeOrigins, M)
 

decodeOrigins <- function(CM, row) .Call(C_decodeOrigins, CM, row)

computeSNPS <- function(population, gen, sex, nr,
			from_p = 1, to_p = Inf,
			output_compressed=FALSE,			
			select = NULL,
			what = c("geno", "haplo")
			) {
#  print("entering")
  ##  Print(gen, sex, nr)
##  str(population$breeding[[gen[1]]][[sex[1]]][[nr[1]]])
##  name <- as.character(as.list(match.call(envir=parent.frame(2L)))[[1]])
  ##  DEBUG(name, population, gen, sex, nr);
   
  if (is.matrix(gen)) {
    gen <- gen[, 1]
    sex <- gen[,2]
    nr <- gen[, 3]
  }
  geno <- what[1] != "haplo"

  ## Print(population[[1]])
 
  ans <-  .Call(C_computeSNPS, population,
		as.integer(gen),
		as.integer(sex),
		as.integer(nr), as.integer(from_p),
		as.integer(if (is.finite(to_p)) to_p else NA), select,
		geno)

  if (!output_compressed) {
    ##print(ans);    Print(output_compressed)    
    ans <- if (geno) decodeGeno(ans) else decodeHaplo(ans)
  }
  
  ##  if (PB(name)) RETURN(name) else SAVE(name, ans)
  ## genomicmatrix or haplomatrix
  return(ans)
}


compute <- function(population, gen, sex, nr, tau, vec, betahat,
		    select = NULL, matrix.return=FALSE) {
  ## implements ( t(Z) Z + tau \1_{ind x ind} )^{-1} %*% vec

    
  if (is.matrix(gen)) {
    gen <- gen[, 1]
    sex <- gen[,2]
    nr <- gen[, 3]
  }
  stopifnot(length(tau) == 1 && length(betahat) == 1)

  ## returns a list
   .Call(C_compute, population,
	as.integer(gen),
	as.integer(sex),
	as.integer(nr),
	as.double(tau),
	as.double(vec),
	as.double(betahat),
	as.integer(select),
	as.logical(matrix.return)
	)
}
```
