## Supplementary material for "MoBPS - Modular Breeding Program Simulator": miraculix R-package: rhaplo.R

```
## Authors 
## Martin Schlather,
##
##
## Copyright (C) 2019 -- 2019 Martin Schlather
##
## This program is free software; you can redistribute it and/or
## modify it under the terms of the GNU General Public License
## as published by the Free Software Foundation; either version 3
## of the License, or (at your option) any later version.
##
## This program is distributed in the hope that it will be useful,
## but WITHOUT ANY WARRANTY; without even the implied warranty of
## MERCHANTABILITY or FITNESS FOR A PARTICULAR PURPOSE.  See the
## GNU General Public License for more details.
##
## You should have received a copy of the GNU General Public License
## along with this program; if not, write to the Free Software
## Foundation, Inc., 59 Temple Place - Suite 330, Boston, MA  02111-1307, USA.  


rhaplo <- function(freq, indiv, loci, freq2, file,
                   file.type = c("beagle", "plink", "plink2"),
                   debugging = FALSE) {
  if (!hasArg("freq")) {
    if (!hasArg("loci")) stop("if 'freq' is not given, 'loci' must be given")
    freq <- 1/2
  } else freq <- as.double(freq)
  
  if (hasArg("loci")) {
    loci <- as.integer(loci)
    freq <- rep(freq, length.out=loci)
     if (hasArg("freq2")) freq2 <- rep(as.double(freq2), length.out=loci)
    else freq2 <- freq
  } else {
    loci <- length(freq)
    if (hasArg("freq2")) {
      freq2 <-
        if (length(freq2)==1) rep(as.double(freq2), length.out=loci)
        else as.double(freq2)
    } else freq2 <- freq
  }

  indiv <- as.integer(indiv)
 
  if (!hasArg("file")) {
    Code <-.Call(C_rhaplomatrix, freq, freq2, indiv)
    return(.Call(C_rhaplomatrixPart2, freq, freq2, indiv, Code))
  }

  file.type <- match.arg(file.type)
  file <- paste0(file, ".", Extension[file.type])
  header <- Header[file.type]
  leadingcol <- LeadingCol[file.type]
  doubledIndiv <- DoubledIndiv[file.type]
  Coding <- Coding[file.type]
  coding <- character(nchar(Coding))
  for (i in 1:length(coding)) coding[i] <- substring(Coding, i, i)

  codes <- coding[1 + (stats::runif( 2 * indiv * loci) < c(freq, freq2))]
 
  haplo <- length(coding) == 4
  M <- matrix(ncol = indiv * 2, codes)
  if (haplo) {
    if (!doubledIndiv) {
      M2 <- matrix(nrow = 2 * loci, ncol = indiv)
      M2[c(TRUE, FALSE), ] <- M[, c(TRUE, FALSE)]
      M2[c(FALSE, TRUE), ] <- M[, c(FALSE, TRUE)]
      M <- M2
    }
    if (!IndivPerCol[file.type]) M <- t(M)
    if (debugging) {
      codes <- codes == coding[2]
      m <- apply(array(codes, dim=c(loci, 2, indiv)), c(1,3), sum)
      attr(file, "M") <- m
    }
  } else M <- M[, c(TRUE, FALSE)] + M[, c(FALSE, TRUE)]

  if (file.type == "plinkbinary") {
    stop("not programmed yet")
  } else { ## ASCII file
    if (header == 0) file.create(file)
    else write(file=file, paste("This is header line no", 1:header))
    if (leadingcol > 0) M <- cbind(matrix("X", ncol=leadingcol, nrow=nrow(M)),M)
    sep <-coding[length(coding)]
    write.table(M, file=file, append=TRUE, quote=FALSE, sep = sep,
                row.names=FALSE, col.names=FALSE)
  }
  info <- c(
      what = HAPLO,
      snps = loci,# get rid of any attributes
      individuals=indiv,# get rid of any attributes
      addr0=0,
      addr1=0,
      align=0,
      align1=0,
      sumgeno=0,
      sumgenoE9=0,
      unused = NA, 
      unused2 = NA,
      isSNPxInd = as.integer(IndivPerCol[file.type]),# get rid of name attribute
      bitspercode = 1, 
      bytesperblock = NA,
      codesperblock = NA,
      header=header,
      DoubledIndividuals=as.integer(doubledIndiv),#get rid of attributes
      leadingcolumns=as.integer(leadingcol),#get rid of attributes
      memInUnits0 = NA,
      meminUnits1 = NA,
      AlignedUnits0 = NA,
      AlignedUnits1 = NA
   )
  attr(file, "information") <- info
  attr(file, "method") <- Haplo
  class(file) <- HAPLOMATRIX
  return (file)
}
```
