## Supplementary material for "MoBPS - Modular Breeding Program Simulator": miraculix R-package: S3.R

```
## Authors 
## Martin Schlather,
##
##
## Copyright (C) 2017 -- 2019 Martin Schlather
#               2014 Florian Skene
##
## This program is free software; you can redistribute it and/or
## modify it under the terms of the GNU General Public License
## as published by the Free Software Foundation; either version 3
## of the License, or (at your option) any later version.
##
## This program is distributed in the hope that it will be useful,
## but WITHOUT ANY WARRANTY; without even the implied warranty of
## MERCHANTABILITY or FITNESS FOR A PARTICULAR PURPOSE.  See the
## GNU General Public License for more details.
##
## You should have received a copy of the GNU General Public License
## along with this program; if not, write to the Free Software
## Foundation, Inc., 59 Temple Place - Suite 330, Boston, MA  02111-1307, USA.  


print_haplogeno <- function(x, name, ...) {
  info <- attr(x, "information")
  indiv <- info[INDIVIDUALS + 1]
  method <- attr(x, "method")
   if (indiv > 1) {
    cat(if (method == Haplo) "Loci" else "SNP",
        " x Indiv ", name, " matrix with ",
        SNPCODING_NAME[method + 1],
        " snpcoding: [1:", info[SNPS + 1], ", 1:", indiv,"]",
        sep="")
    if (method == Haplo) cat(" (twice)")
   } else
    cat(if (method == Haplo) "Haplotype " else "SNP",
        "information at ", info[SNPS + 1], " loci.", sep="")
    
  if (is.character(x))
    cat("\n<", x, "> (",
        if (length(attr(x, "M")) == 0) "unloaded" else "loaded", ")", sep="")
  else if (length(f <- attr(x, "filename")) > 0)
    cat("\n<", f, "> (loaded)", sep="")
  cat("\n")
}


as.matrix.haplomatrix <- function(x, ...) decodeHaplo(x, ...)


print.haplomatrix <- function(x, ...)  print_haplogeno(x, "haplo", ...)


str.haplomatrix <- function(object, ...) {
  print.haplomatrix(object, ...)
  cat("Attributes are a ")
  utils::str(base::attributes(object)) ##
}

as.haplomatrix <- function(object, ...) haplomatrix(object, ...)


as.matrix.genomicmatrix <- function(x, ...) decodeGeno(x, ...)
  
print.genomicmatrix <- function(x, ...) print_haplogeno(x, "genomic", ...)

str.genomicmatrix <- function(object, ...) {
  print.genomicmatrix(object, ...)
  cat("Attributes are a ")
  utils::str(base::attributes(object)) ##
  ##class(object) <- NULL;    str(object)
} 
  


as.genomicmatrix <- function(object, ...)
  genomicmatrix(object, ..., loading = FALSE)


if (FALSE) {
  setMethod("crossprod", signature = GENOMICMATRIX, definition=function(x) crossprodx(x))
  
  setMethod("%*%", signature = c(GENOMICMATRIX, "numeric"),
          definition=function(x,y) {
            stopifnot(is.vector(y))
            fdot (x, y)
          })
  
setMethod("%*%", signature = c("numeric", GENOMICMATRIX),
          definition=function(x,y) {
            stopifnot(is.vector(y))
            dotf
          })

setMethod("<-", signature = GENOMICMATRIX,
          definition=function(x) copyGenox())

}
```
