## Supplementary material for "MoBPS - Modular Breeding Program Simulator": miraculix R-package: scan.R

```
## Authors 
## Martin Schlather,
##
##
## Copyright (C) 2017 -- 2019 Martin Schlather
##
## This program is free software; you can redistribute it and/or
## modify it under the terms of the GNU General Public License
## as published by the Free Software Foundation; either version 3
## of the License, or (at your option) any later version.
##
## This program is distributed in the hope that it will be useful,
## but WITHOUT ANY WARRANTY; without even the implied warranty of
## MERCHANTABILITY or FITNESS FOR A PARTICULAR PURPOSE.  See the
## GNU General Public License for more details.
##
## You should have received a copy of the GNU General Public License
## along with this program; if not, write to the Free Software
## Foundation, Inc., 59 Temple Place - Suite 330, Boston, MA  02111-1307, USA.  

summary.scanning <- function(object, ...) {
  if (length(object$significant.areas) == 0 ||
       length(object$significant.areas) == 0 ) return(NULL)
  if (is.list(object$significant.areas)) {
    if (length(object$significant.areas) == 1) {
      return (object$significant.areas[[1]])
    } else {
      warning("not programmed yet") ## tabelle mit allen werten nebeneinander, sodass man sieht welche intervalle aufgesplittet oder verkleinert wurden 
      return(object$significant.areas)
    }
  } else {
    return (object$significant.areas)
  }
}

print.scanning <- function(x, ...) {
  str(x)#
}

plot.scanning <- function(x, ..., lwd=3, col=1:8) {
  graphics::plot(x$pos, x$freq, pch=".")
  if (length(x$pos) > 50000) {
    w <- windower(data=data.frame("-", x$pos, x$pos + 1, x$freq), length=5000)
    points(0.5*(w[,1] + w[,2]), w[,3], col="lightblue", pch=20, cex=0.5)
  }
  if (length(x$significant.areas) > 0) {
    if (is.list(x$significant.areas)) {
      for (i in 1:length(x$significant.areas)) {
        if (ncol(x$significant.areas[[i]]) > 0) {
          colour <- col[1 + (i) %% length(col)]
          for (j in 1:nrow(x$significant.areas[[i]]))
            lines(x$significant.areas[[i]][j, 1:2], c(0,0), col=colour, lwd=lwd)
        } else {
          message("no significant values for ",
                  if (x$was.null.threshold) "alpha" else "threshold",
                  "[",  i, "] = ",
                  if (x$was.null.threshold) x$alpha[i] else x$threshold[i])
        }
      }
    } else {      
      colour <- col[1 + (x$significant.areas[, 3]-1) %% length(col)]
      for (i in 1:nrow(x$significant.areas)) {
        lines(x$significant.areas[i, 1:2],
              c(0,0) + (x$significant.areas[i, 3] - 1) / 20,
              col=colour[i], lwd=lwd)
      }
    }
  } else {
    message("No significant area found at all.")
  }
  #points(x$pos, x$freq, pch=".")
}


scanning <- function(pos, freq, file, tuningUnits, alpha = 0.1,
                     coarsening = 1,
                     minscans = 0,
                     maxscans = 0,
                     sumscan = FALSE,  perSNP = TRUE,                     
                     colname, n,                 
                     threshold = NULL, collect=!old.def,
                     old.def = FALSE,
                     max.intervals = length(alpha) * 100000,
                     max.basepair.distance = 50000,
                     exclude.negative.at.boundary = TRUE,
                     maximum = TRUE,
                     mean.freq, sd.freq, mean.n) { 
  stopifnot(all(alpha > 0), all(alpha < 1)) ## Gleichheit nicht zulassen
  stopifnot(xor(missing(pos) && missing(freq), missing(file)))

  verbose <- RFoptions(GETOPTIONS="basic")$verbose
    
  if (missing(file)) {
    if (missing(colname)) colname <- "HeterAB"
  } else {
    ext <- strsplit(file, "\\.")[[1]]
    ext <- ext[length(ext)]
    if (ext == "rda") {
      load (file)
    } else {
      if (missing(colname)) {
        colname <- if (ext=="bed") c(pos=3, freq=4, n=5)
        else "HeterAB"
      }    
      if (is.character(colname)) {
        z <- read.table(file)[c("Pos", colname)]
        freq <- z[[colname]]
        pos <- z$Pos
      } else {
        stopifnot(is.numeric(colname))
        stopifnot(missing(n))
        z <- as.matrix(read.table(file)[colname])     
        pos <- z[, which(colname["pos"] == colname)]
        if (ext=="bed") {
          freq <- -z[, which(colname["freq"] == colname)]
        } else freq <- z[, which(colname["freq"] == colname)]
        n <- z[, which(colname["n"] == colname)]
       #
        if (FALSE) {
          idx <- pos > 3.5e7 & pos < 5e7
          freq <- freq[idx]
          pos <- pos[idx]
          n <- n[idx]
          save (file="data/genetics.rda", freq, pos, n)
        }
      }
      rm("z")
    }
    if (verbose) message(length(freq), " data read from file '", file, "'.")
  }

  coarsening <- as.integer(coarsening)
  if (coarsening > 1) {
    w <- windower(data=data.frame("-", pos, pos + 1, freq), length=coarsening)
    freq <- w[ , 3]
    idx <- !is.na(freq)
    if (!missing(n)) {
      w <- windower(data=data.frame("-", pos, pos + 1, n), length=coarsening)
      n <- w[idx, 3]
    }
    freq <- freq[idx]
    ##  pos <- w[idx, 1]
    pos <- 0.5 * (w[idx, 1] + w[idx, 2])
 }

  N <- length(pos)
  Nalpha <- if (is.null.threshold <- is.null(threshold))
              length(alpha) else length(threshold)
  
  stopifnot(all(abs(pos - round(pos)) < 1e-8))
  freq <- as.double(freq)
  pos <- as.integer(round(pos))
  perSNP <- as.integer(perSNP)
  minscans <- as.integer(minscans)
  maxscans <- as.integer(maxscans)

 
  if (missing(n)) n <- round(1 / min(freq[freq != 0])) ##
  if (length(n) == 1 && !all(abs(n * freq - round(n * freq)) < 1e-8)) {
    msg <- paste("Frequencies are not multiples of 1/n.",
                 if (length(n)==1) paste("Here, n=", n, ".", sep=""))
    stop(msg) # else warning(msg)
  }

  if (missing(mean.freq)) {
    mean.freq <- mean(freq)
    if (verbose) cat("mean freq:", mean.freq, "\n")
  }
  if (missing(sd.freq)) {
    sd.freq <- sd(freq)
    if (verbose) cat("sd freq:", sd.freq, "\n")
  }
  if (missing(mean.n)) {
    mean.n <- mean(n)
    if (verbose) cat("mean n:", mean.n, "\n")
  }
  freq <- (freq - mean.freq) / sd.freq
  freq <- freq - tuningUnits * mean.n / n
  mn <- mean(freq)
  eps <- 1e-16
  if (mn > -eps) {
    msg <- "The mean of the shifted frequencies should "
    deltatU <- sum(freq) / sum(1/n)
    deltatU <- deltatU +  10^{-log(deltatU) / log(10) - 2}
    if (mn > eps) stop(msg, "not be positive (but it is ",
                       formatC(mn, digits=4),
                       "; so 'tuningUnits' should be at least ",
                       format(deltatU, digits=4), " larger)")
    warning(msg, "be negative.")
  }

  if (perSNP) {
    v <- var(freq)
    nn <- N
  } else {
    stop("not programmed yet")
    v <- var(freq)
    nn <- diff(range(pos))
  }

 
  if (tuningUnits == 0.0) 
    KabluchkoSpodarev <- sqrt(nn * v) * qnorm(alpha, lower.tail=FALSE)
  else {
    KabluchkoSpodarev <- -999999 # not programmed yet
    
    if (FALSE) {
      u <- unique(freq)    
      f <- numeric(length(u))
      for (i in 1:length(u)) f[i] <- sum(freq == u[i])
      stopifnot(sum(f) == length(freq))
      logmoment <- function(theta) log(sum(f  * exp(theta * u)))
      theta.star <- uniroot(logmoment, interval=c(1/max(u), min(-log(f) / u)))
                                        # remark 4.1, (4.11)
      H.star <- log(H) - log(a.star * theta.star)
      KabluchkoSpodarev <- (log(nn) + H.star - log(-log(1-alpha))) / theta.star
    }
  }
  
  if (is.null(threshold))
    threshold <- as.double(KabluchkoSpodarev)
  else {
    threshold <- as.double(threshold)
   # if (FALSE && !is.null(alpha))
   #   cat("threshold by Kabluchko & Spodarev:", KabluchkoSpodarev, "\n")
  }

  above.threshold <- integer(if (sumscan) Nalpha * N else Nalpha)
  
  maximum <- double(1)
  if (old.def) {
    if (sumscan) 
      .Call(C_sumscan,
            pos, N, freq, minscans, maxscans, threshold, Nalpha, perSNP,
            above.threshold, maximum)
    else
      .Call(C_scan,
            pos, N, freq, minscans, maxscans, threshold, Nalpha, perSNP,
            above.threshold, maximum)
  } else {
    signif.areas <-
      .Call(C_collect_scan2,  pos, N, freq, minscans, maxscans,
            threshold, Nalpha,
            perSNP, as.integer(max.intervals),
            as.integer(max.basepair.distance),
            as.integer(exclude.negative.at.boundary),
            above.threshold, maximum)
  }
  
  if (any(above.threshold < 0))
    above.threshold <- rep(Inf, length(above.threshold))
  any.above.threshold <- any(above.threshold > 0)
   
  if (sumscan) {
    dim(above.threshold) <- c(N, Nalpha)
  }

  l <- list(if (!missing(file)) file=file,
            pos=pos, freq=freq,
            tuningUnits = tuningUnits,
            alpha=alpha,
            n = n,
            minscans=minscans,
            maxscans=maxscans,
            sumscan = sumscan,
            perSNP=perSNP,
            was.null.threshold = is.null.threshold,
            threshold=threshold,
            any.above.threshold =  any.above.threshold,
            max.basepair.distance = max.basepair.distance,
            exclude.negative.at.boundary = exclude.negative.at.boundary,
            # output
            above.threshold=above.threshold,
            maximum=maximum)
  if (!collect || sumscan) {
    if (!old.def) l$significant.areas <- t(signif.areas)
      class(l) <- "scanning"
    return(invisible(l))
  }  
  if (verbose) cat(if (old.def) "First scan done.\n" else "Scanning done.\n")
  
  significant.areas <- areas <- values <- NULL
  if (any.above.threshold) {
    ## Mengen vereinigen
    if (old.def) {
      n.areas <- max(above.threshold)
      if (n.areas > 1e8) stop(n.areas, " areas are above the threshold. This is too much.")
      areas <- integer(n.areas * 3) # start, end, thresholdlevel
      ## signif.areas <- integer(n.areas * 3) # start, end, thresholdlevel
      values <- double(n.areas)      # value
      
      signif.areas <-
        .Call(C_collect_scan,  pos, N, freq,
              minscans, maxscans, threshold, Nalpha,
              perSNP, areas, values)
      dim(areas) <- c(3, n.areas)
      signif.areas <- signif.areas[, signif.areas[3, ] != 0, drop=FALSE]   
      if (verbose) cat("Second scan done.\n")
    }
    
    significant.areas <- list()
    totallength <- numeric(Nalpha)
    START <- 1
    END <- 2
    for (i in 1:Nalpha) {
      s <- signif.areas[, signif.areas[3, ] >= i, drop=FALSE]
      j <- 1
      while (j <= ncol(s)) {
        k <- j + 1
        while (k <= ncol(s)) {
          if ((s[START, j] <= s[START, k] && s[START, k] <= s[END, j]) ||
              (s[START, k] <= s[START, j] && s[START, j] <= s[END, k])) {
            if (s[START, k] < s[START, j]) s[START, j] <- s[START, k]
            if (s[END, k] > s[END, j]) s[END, j] <- s[END, k]
            s <- s[, -k, drop=FALSE]
          } else k <- k + 1
        }
        j <- j + 1
      }
      s[3, ] <- i
      significant.areas[[i]] <- t(s)
      totallength[i] <- diff(colSums(significant.areas[[i]])[1:2])
    }
    Message <- paste("Null hypothesis is rejected at ",
                     if (is.null.threshold) paste("alpha =", alpha[1])
                     else  paste("threshold =",formatC(threshold[1], digits=4)),
                     " that the data are homogeneous", sep="")
  } else {
    totallength <- rep(0, Nalpha)
    p <- if (is.null(threshold))
      formatC(pnorm(maximum / sqrt(nn * v), lower.tail=FALSE), digits=4)
    else p <- "'unknown'"
     Message <- paste("Null hypothesis is accepted at ",
                     if (is.null.threshold) paste("alpha =", alpha[1])
                     else paste("threshold =", formatC(threshold[1], digits=4)),
                      " that the data are homogeneous (p=",p, ")", sep="")
  }

  stopifnot(length(totallength) == Nalpha)
  l <- c(l, list(areas = areas, values=values,
                 significant.areas = significant.areas,
                 totallength = totallength,
                 Message = Message))
  class(l) <- "scanning"
  return(invisible(l))
}


print.scan.statistics <- function(x, ...) {
  str(x) #
}

summary.scan.statistics <- function(object, ...) {
  cat(object$Message, "\n")  
}


plot.scan.statistics <- function(x, ..., pch="|", col=2:8) {
  txt <- ""
  alpha <- x$alpha
  Nalpha <- length(alpha)
  ylim <- if (hasArg("ylim")) list(...)$ylim else range(alpha, x$orig.freq)
  colname <- x$colname
  if (is.numeric(colname)) colname <- paste("column", colname["freq"])
  graphics::plot(x$pos, x$orig.freq, pch=pch, cex=0.75, ylim=ylim, main=x$file,
       xlab="positions", ylab=colname, col="black")
  if (x$sumscan) {
    above <- t(x$above.threshold)
    maxi <- apply(above, 1, max)
    txt <- paste("mx.cnts=", paste(maxi[1], collapse=","), "  ", sep="")
    above <- t(above / maxi)
    graphics::matplot(x$pos, above, type="l", add=TRUE, col=col);
    for (i in 1:Nalpha) {
      cur <- col[(i-1) %% length(col) + 1]
      points(x$pos, alpha[i] * (above[, i] > 0), col=cur, pch=20, cex=0.65) 
    }      
  } else { ### not sumscan
    for (i in 1:Nalpha) {
      xx <- x$significant.areas[[i]]
      if (length(xx) == 0) next;
      cur <- col[(i-1) %% length(col) + 1]
      for (j in 1:nrow(xx))
        if (xx[j, 1] == xx[j, 2]) points(xx[j, 1], alpha[i], col=cur, pch=20)
        else lines(c(xx[j, 1], xx[j, 2]), rep(alpha[i], 2), col=cur, lwd=3)
    }
  }
  
  if (x$minscans != 0) txt <- paste(txt, "minscans=", x$maxscans, "  ", sep="")
  if (x$maxscans != 0) txt <- paste(txt, "maxscans=", x$maxscans, sep="")

  if (txt!="") text(x=0, y=0.4, labels=txt, adj=c(0,0), col=6)
  title(sub=x$Message, col.sub="darkred")
  legend(x="topright",
         legend=alpha, col=col, text.col=col, lwd=2, adj=c(0, 0),
         title=expression(alpha))
}


scan.statistics <-
  function(file, tuningUnits, alpha=c(0.05, 0.01), repet=1000,
           coarsening = 1,
           minscans=0, maxscans=0,
           sumscan = FALSE,
           perSNP=TRUE, # otherwise per "basepair"
           colname, n, return.simu = FALSE,
           debug = FALSE,
           formula = FALSE, 
           old.def=FALSE,
           max.intervals = length(alpha) * 100000,
           max.basepair.distance = 50000,
           exclude.negative.at.boundary = TRUE,
           pos, freq) {
  if (!perSNP) stop("not programmed yet") ## simu muss auf pktprozessen beruhen
  if (formula) stop("Kabluchko-Spodarev formulae not programmed yet")
  if (1/min(alpha) > repet)
    stop("value of 'repet' too small to identify the 'alpha' level(s)")

  if (hasArg("file")) {
    if (hasArg("pos") || hasArg("freq") || hasArg("colname") || hasArg("n"))
      stop("either the file name or 'pos', 'freq', 'colname', 'n' must be given.")
    if (is.character(file)) {
      ext <- strsplit(file, "\\.")[[1]]
      ext <- ext[length(ext)]
      if (ext == "rda") {
        load (file)
      }
      else {
        if (missing(colname)) {
          colname <- if (ext=="bed")  c(pos=3, freq=4, n=5) else "HeterAB"
        }
        if (is.character(colname)) {
          z <- read.table(file)[c("Pos", colname)]
          freq <- z[[colname]]
          pos <- z$Pos
        } else {
          stopifnot(is.numeric(colname))
          stopifnot(missing(n))
          z <- as.matrix(read.table(file)[colname])
          pos <- z[, which(colname["pos"] == colname)]
          if (ext=="bed") {
            freq <- 1 - z[, which(colname["freq"] == colname)]
          } else freq <- z[, which(colname["freq"] == colname)]
          n <- z[, which(colname["n"] == colname)]  
        }
        rm("z")
      }
    } else {
      for (i in c('pos', 'freq', 'colname', 'n' ))
        assign(i, file[[i]])
    }
  }

  coarsening <- as.integer(coarsening)
  if (coarsening > 1) {
    w <- windower(data=data.frame("-", pos, pos + 1, freq), length=coarsening)
    freq <- w[ , 3]
    idx <- !is.na(freq)
    if (!missing(n)) {
      w <- windower(data=data.frame("-", pos, pos + 1, n), length=coarsening)
      n <- w[idx, 3]
    }
    freq <- freq[idx]
    ##  pos <- w[idx, 1]
    pos <- 0.5 * (w[idx, 1] + w[idx, 2])
  }

 
  N <- length(pos)
  Nalpha <- length(alpha)
  
  if (return.simu) ret.simu <- matrix(NA, nrow=N, ncol=repet)
  if (missing(n)) n <- round(1 / min(freq[freq != 0])) ##
  if (length(n) == 1) stopifnot(all(abs(n * freq - round(n * freq)) < 1e-8))
  
  ylim <- range(freq, alpha)
 
  maxima <- probab <- NULL
  if (debug && hasArg("file") &&
      file.exists(filename <- paste(file, ".max.rda", sep="")))
    load (filename)
  else if (is.null(formula) || !formula) {
    probab <- sum(freq) / N ## sum(n * freq) / (N * n) // counts
    maxima <- double(repet)
    verbose <- RFoptions(GETOPTIONS="basic")$verbose
     for (i in 1:repet) {
      if (verbose) cat(i, "")
      ## simufreq <- rbinom(N, n, probab) / n
      simufreq <- sample(freq, replace=TRUE)
      if (return.simu) {
        ret.simu[, i] <- simufreq
        stopifnot(!any(is.na(ret.simu[,i])))
      }
      maxima[i] <-
        scanning(pos, simufreq, tuningUnits=tuningUnits,
                 minscans=minscans, maxscans=maxscans,
                 sumscan=FALSE, perSNP=perSNP,
                 alpha=0.999, colname=colname, n=n,
                 old.def = old.def,
                 max.intervals = max.intervals,
                 max.basepair.distance = max.basepair.distance,
                 exclude.negative.at.boundary = exclude.negative.at.boundary
                 )$maximum      
      if (debug == 2) {
        graphics::plot(pos, simufreq, pch=".", ylim=ylim,
                       main= if (hasArg("file")) file,
                       sub=i)
        readline(paste(sum(simufreq), "press return"))
      }
    }
    maxima <- sort(maxima)
    threshold <- as.double(maxima[ceiling((1-alpha) * repet)])
    if (debug && hasArg("file")) {
      save (file=filename, maxima, threshold); return()
    }
  }

  res <-
    scanning(pos, freq, tuningUnits=tuningUnits,
             minscans=minscans, maxscans=maxscans,
             sumscan=sumscan, perSNP=perSNP,
             alpha=alpha,
             colname=colname, 
             threshold = if (is.null(formula) || !formula) threshold else NULL,#
             n=n,
             collect=TRUE,
             old.def = old.def,
             max.intervals = max.intervals,
             max.basepair.distance = max.basepair.distance,
             exclude.negative.at.boundary = exclude.negative.at.boundary
             )

  if ( (is.null(formula) || !formula) && !res$any.above.threshold)  {
     p <- mean(res$maximum >= maxima)
     res$Message <- paste("Null hypothesis is accepted at alpha=", alpha[1],
                          " that the data are homogeneous (p =",
                          formatC(p, digits=4), ")", sep="")
  }

  res <- c(res, list(file = if (hasArg("file")) file else NULL,
                     orig.freq = freq,
                     maxima = maxima, repet = repet,
                     colname = if (missing(colname)) NULL else colname ,
                     if (return.simu) ret.simu = ret.simu))
  class(res) <- "scan.statistics"

  return(res)
}
```
