## Supplementary material for "MoBPS - Modular Breeding Program Simulator": miraculix R-package: windower.R

```
## Authors 
## Martin Schlather,
##
##
## Copyright (C) 2017 -- 2019 Martin Schlather
##
## This program is free software; you can redistribute it and/or
## modify it under the terms of the GNU General Public License
## as published by the Free Software Foundation; either version 3
## of the License, or (at your option) any later version.
##
## This program is distributed in the hope that it will be useful,
## but WITHOUT ANY WARRANTY; without even the implied warranty of
## MERCHANTABILITY or FITNESS FOR A PARTICULAR PURPOSE.  See the
## GNU General Public License for more details.
##
## You should have received a copy of the GNU General Public License
## along with this program; if not, write to the Free Software
## Foundation, Inc., 59 Temple Place - Suite 330, Boston, MA  02111-1307, USA.  


windower <- function(data, length=20000, step=length/2, start=0,
                     n.min=0, na.rm=TRUE,
                     what=c("mean", "var", "sd", "min", "max", "median",
                         "sum")) {
  if (!na.rm) stop("'na.rm = FALSE' not programmed yet")
  if (any(isna <- is.na(data[,4]))) data <- data[!is.na, ]
  
  what <- match.arg(what)
  if (what == "var") return(windower.var(data=data, length=length, step=step,
        start=start, n.min=n.min, na.rm=na.rm))
  if (what == "sd") return(windower.sd(data=data, length=length, step=step,
        start=start, n.min=n.min, na.rm=na.rm))
  if (is.sum <- what == "sum") what <- "mean"
  ## +1 start notation for start
  if (!is.data.frame(data) || ncol(data) < 4) stop("not a 'bed' file")
  stopifnot(step > 0, length > 0)
  
  Start <- as.integer(data[ , 2])  ## alles 1er System
  Ende <-  as.integer(data[,  3])
  if (any(diff(Start) <= 0) || any(diff(Ende) <= 0))
    stop("intervals not disjoint or not ordered or file too long")
  stopifnot(!any(is.na(Ende)))

  n <- as.integer(ceiling((Ende[length(Ende)] - start) / step))

  whatnr <- which(what == as.character(as.list(args(windower))$what[-1]))
  
  win <- .Call(C_windower, whatnr,
               as.integer(start), as.integer(length), as.integer(step),
               Start, Ende, as.double(data[, 4]), nrow(data), n)

  if (is.sum) {
    win[3, is.na(win[3, ])] <- 0
    win[3, ] <- win[3, ] * win[4, ]
  }
  ##  dim(win) <- c(4, n)
  idx <- win[4, ] >= n.min
  return(t(win[, idx]))
}

windower.var <- function(data, length, step, start, n.min=2, na.rm) {
  if (n.min <= 1) stop("'n.min' must be at least 2")
  m <- windower(data=data, length=length, step=step, start=start,
                n.min=n.min, na.rm=na.rm)
  data$V4 <- data$V4^2
  m2 <- windower(data=data, length=length, step=step, start=start,
                 n.min=n.min, na.rm=na.rm)
  return(cbind(m[, 1:2], m[, 4] / (m[, 4] -1) * pmax(0, m2[, 3] - m[, 3]^2),
               m[, 4]))
}

windower.sd <- function(data, length, step, start, n.min=2, na.rm) {
 m <- windower.var(data=data, length=length, step=step, start=start,
                   n.min=n.min, na.rm=na.rm)
 m[, 3] <- sqrt(m[, 3])
 m
}
```
