## Supplementary material for "MoBPS - Modular Breeding Program Simulator": miraculix R-package: zzz.R

```
## Authors 
## Martin Schlather,
##
##
## Copyright (C) 2017 -- 2019 Martin Schlather
##
## This program is free software; you can redistribute it and/or
## modify it under the terms of the GNU General Public License
## as published by the Free Software Foundation; either version 3
## of the License, or (at your option) any later version.
##
## This program is distributed in the hope that it will be useful,
## but WITHOUT ANY WARRANTY; without even the implied warranty of
## MERCHANTABILITY or FITNESS FOR A PARTICULAR PURPOSE.  See the
## GNU General Public License for more details.
##
## You should have received a copy of the GNU General Public License
## along with this program; if not, write to the Free Software
## Foundation, Inc., 59 Temple Place - Suite 330, Boston, MA  02111-1307, USA.  


.onLoad <- function(lib, pkg) {
  ## DESCRIPTION MUSS RandomFieldsUtils EINGEFUEGT SEIN
  .Call("loadmiraculix");
  RFoptions(solve_method=c("cholesky", "cholesky"), max_chol=10^5)
}

.onAttach <- function (lib, pkg) {
  packageStartupMessage(.Call("attachmiraculix"));
}

.onDetach <- function(lib) {
}

.onUnload <- function(lib, pkg){
#   RFoptions(storing=FALSE) ## delete everything
  .C("detachmiraculix")
}
```
