## Supplementary material for "MoBPS - Modular Breeding Program Simulator": miraculix R-package: 2bit.cc

```
/*
 Authors 
 Martin Schlather,


 Copyright (C) 2018 -- 2019  Martin Schlather

This program is free software; you can redistribute it and/or
modify it under the terms of the GNU General Public License
as published by the Free Software Foundation; either version 3
of the License, or (at your option) any later version.

This program is distributed in the hope that it will be useful,
but WITHOUT ANY WARRANTY; without even the implied warranty of
MERCHANTABILITY or FITNESS FOR A PARTICULAR PURPOSE.  See the
GNU General Public License for more details.

You should have received a copy of the GNU General Public License
along with this program; if not, write to the Free Software
Foundation, Inc., 59 Temple Place - Suite 330, Boston, MA  02111-1307, USA.  
*/


// gcc -mavx -o hello_avx hello_avx.c

#define BitsPerCode 2L

#include <Basic_utils.h>
#include "Bit23.intern.h"
#include "haplogeno.h"

INLINER


#define TWO_GENUINEBITSPERMINIBLOCK 65536L // testing only

static double rev_geno_code[4] = {0, 1, RF_NA, 2}; 
static BlockType geno_code[nr_genotypes] = {0, 1, 3}, // 0, 1, 2
  result_code[nr_results] = {0, 1, 2, 3};
static Uint result_valueAND[nr_results] = {0, 1, 0 /* unused */, 4},
  result_valueXOR[nr_results] = {0, 0, 1, 0};

static table_type *TABLE2AND = NULL,
  *TABLE2XOR = NULL;

Uint CodesPerBlock2() { return CodesPerBlock; }
Uint UnitsPerIndiv2(Uint snps) { return Blocks(snps) * UnitsPerBlock; }


void initiate_table2() { // hash table 
  if (TABLE2AND != NULL || TABLE2XOR != NULL) BUG; 
  initiate_tableI(&TABLE2AND, TABLE_SIZE, CodesPerMiniblock, BitsPerCode,
		  result_code, result_valueAND, nr_results);
  initiate_tableI(&TABLE2XOR, TABLE_SIZE, CodesPerMiniblock, BitsPerCode,
		  result_code, result_valueXOR, nr_results);
}

/*
void static printbits(BlockType x) { //
  printbits(x, sizeof(BlockType) * BitsPerByte, BitsPerCode);//
}
*/

void Init2() {
  assert(BytesPerBlock == sizeof(BlockType0));
  if (TABLE2AND == NULL) initiate_table2();
}


// these defintions must match extactly as.genomicmatrix
// BODY-CODE IDENTICAL TO CODING3
SEXP matrix_start2 START23(Init2)
Ulong sumGeno2 sumGeno23

void matrix_coding2 Coding23Matrix;
Coding23(start_individual, end_individual, start_snp, end_snp, Mnrow, FROMINPUT);
} // see bit23intern.h


void haplo2geno2(Uint *X, Uint snps, Uint individuals, Uint unitsPerIndiv,
		 Uint *ans) {
  // note that X == ans is allowed !!
  Coding23(0, individuals, 0, snps, unitsPerIndiv, FROMHAPLO);
}


void ReUseAsTwoBit(SEXP Code) {
  Uint *info = GetInfo(Code),
    snps = info[SNPS],
    individuals = info[INDIVIDUALS];
  Ulong memInUnitsPerIndiv = Blocks(snps) * UnitsPerBlock;  
  InternReUseAs(Code, TwoBit, snps, individuals, memInUnitsPerIndiv, false);  
}
		   

void MmPsq2(Uint individuals, Uint min_row, Uint max_row,
	    Uint min_col, Uint max_col,
	    Uint compressed_start, Uint compressed_end, Uint blocks,
	    BlockType0 *M, double *ergb, bool Parallel) {  
  //  table_type *table = TABLE2AND, // auf lokalen array zu kopiern, scheint sich
   //                            nicht zu lohnen
    //  *tableXor = TABLE2XOR;

  //  printf("%d\n", TABLE2_SIZE); BUG;  
  assert(TABLE_SIZE == 65536);
  table_type tableAnd[TABLE_SIZE], tableXor[TABLE_SIZE];
  MEMCOPY(tableAnd, TABLE2AND, TABLE_SIZE * sizeof(table_type));
  MEMCOPY(tableXor, TABLE2XOR, TABLE_SIZE * sizeof(table_type));

  
  assert(//min_row >= 0 && min_col >= 0 &&
	 max_row <= individuals && max_col <= individuals);

#ifdef DO_PARALLEL
#pragma omp parallel for num_threads(CORES) if (Parallel)
#endif	  
  for (Uint c=min_col; c<max_col; c++) {
    Uint r  =  c > min_row ? c : min_row;
    BlockType * Mc = M + c * blocks,
      *Mr = M + r * blocks;
    double *ptr = ergb + c * individuals,
      *res_ptr_r = ptr + r;
    for ( ; r<max_row; r++, res_ptr_r++, Mr += blocks) {
      Uint sum = 0;
      BlockType *MR = Mr + compressed_start,
	*MC = Mc + compressed_start;
      for (Uint s=compressed_start; s<compressed_end; s++, MR++, MC++) {
	block_compressed x;
	x.x = *MR & *MC;
	sum+= tableAnd[x.b[0]] + tableAnd[x.b[1]] +
	  tableAnd[x.b[2]] + tableAnd[x.b[3]];

 	x.x = *MR xor *MC;
	sum += tableXor[x.b[0]] + tableXor[x.b[1]]
	  + tableXor[x.b[2]] + tableXor[x.b[3]];
     }
      ergb[c + r * individuals] = *res_ptr_r = (double) sum;
    }
  }
}


double getValue2(BlockType0 *Indiv, Uint s) {
  block_compressed x;
  x.x = Indiv[s / genuineCodesPerBlock];
  Uint idx = s % genuineCodesPerBlock,
    y = x.b[idx / CodesPerMiniblock] >> ( (idx % CodesPerMiniblock) * BitsPerCode); 
  //  printf("y=%d \n", (Uint) y);
  //  printf("%10e\n", rev_geno_code[y & CodeMask]);
  // printf("ok\n");

  return rev_geno_code[y & CodeMask];
}

SEXP get_matrix2(SEXP SNPxIndiv) {
  if (TABLE2AND == NULL) Init2();
  Uint 
    *info = GetInfo(SNPxIndiv),
    individuals = info[INDIVIDUALS],
    snps = info[SNPS],
    blocks = Blocks(snps),
    endfor = individuals * blocks;
  SEXP Ans;
  PROTECT(Ans = get_matrix23_start(individuals, snps, R_NilValue));
  Uint *ans = (Uint *) INTEGER(Ans);
  BlockType *M = (BlockType0 *) Align(SNPxIndiv, ALIGN_23);
  for (Uint a=0; a<endfor; a+=blocks) {
    BlockType *Ma = M + a;
    for (Uint s=0; s<snps; s++) *(ans++) = getValue2(Ma, s);
  }
  UNPROTECT(1);
  return Ans;
}


SEXP get_matrixN_2(SEXP SNPxIndiv, SEXP Snps) {
  if (TABLE2AND == NULL) Init2();
  GET_MATRIX_N_23(getValue2);
}


SEXP allele_freq2(SEXP SNPxIndiv) {
  if (TABLE2AND == NULL) Init2();
  ALLELE_FREQ23(getValue2);  
}


void matrix2_mult(Uint * SNPxIndiv, Uint snps, Uint individuals, double *A){
  Uint blocks = Blocks(snps);
  MmPsq2(individuals, 0, individuals, 0, individuals, 0, blocks, blocks,
	 (BlockType0 *) SNPxIndiv, A, true);
}

Uint *Align2(SEXP Code, Uint nr, bool test) {
  return AlignTest(Code, nr, test); }


void zeroNthGeno2 ZERONTH
```
