## Supplementary material for "MoBPS - Modular Breeding Program Simulator": miraculix R-package: 3bit.cc

```
/*
 Authors 
 Martin Schlather,


 Copyright (C) 2018 -- 2019  Martin Schlather

This program is free software; you can redistribute it and/or
modify it under the terms of the GNU General Public License
as published by the Free Software Foundation; either version 3
of the License, or (at your option) any later version.

This program is distributed in the hope that it will be useful,
but WITHOUT ANY WARRANTY; without even the implied warranty of
MERCHANTABILITY or FITNESS FOR A PARTICULAR PURPOSE.  See the
GNU General Public License for more details.

You should have received a copy of the GNU General Public License
along with this program; if not, write to the Free Software
Foundation, Inc., 59 Temple Place - Suite 330, Boston, MA  02111-1307, USA.  
*/


// gcc -mavx -o hello_avx hello_avx.c

// _MM_ALIGN16 Uint ccc;
#define BitsPerCode 3L

#include "Bit23.intern.h"
#include <Basic_utils.h>
#include "haplogeno.h"
#include "Haplo.h"

INLINER


#define TWO_GENUINEBITSPERMINIBLOCK 32768  // testing only

static double rev_geno_code[7] = {0, RF_NA, RF_NA, 1, RF_NA, RF_NA, 2}; 
static BlockType geno_code[nr_genotypes] = {0, 3, 6}, // 0, 1, 2
  result_code[nr_results] = {0, 3, 2, 6};
static Uint result_value[nr_results] = {0, 1, 2, 4};

// Codierung: 0->0; 1->3; 2->6
static table_type *TABLE3 = NULL;
void initiate_table3() { // hash table 
  if (TABLE3 != NULL) BUG; 
  initiate_tableI(&TABLE3, TABLE_SIZE, CodesPerMiniblock, BitsPerCode,
		  result_code, result_value, nr_results);
}

/*
void static printbits(BlockType x) { //
  printbits(x, sizeof(BlockType) * BitsPerByte, BitsPerCode);//
}
*/
Uint CodesPerBlock3() { return CodesPerBlock; }
Uint UnitsPerIndiv3(Uint snps) { return Blocks(snps) * UnitsPerBlock; }

void Init3() {
  assert(BytesPerBlock == sizeof(BlockType0));
  if (TABLE3 == NULL) initiate_table3();
}


// these defintions must match extactly as.genomicmatrix
SEXP matrix_start3 START23(Init3)
Ulong sumGeno3 sumGeno23


void haplo2geno3(Uint *X, Uint snps, Uint individuals, Uint unitsPerIndiv,
		 Uint *ans) {
  Coding23(0, individuals, 0, snps, unitsPerIndiv, FROMHAPLO);
}


void matrix_coding3 Coding23Matrix;
Coding23(start_individual, end_individual, start_snp, end_snp, Mnrow,
	 FROMINPUT);
} // see bit23intern.h

 
void MmPsq3(Uint individuals, Uint min_row, Uint max_row,
	    Uint min_col, Uint max_col,
	    Uint compressed_start, Uint compressed_end, Uint blocks,
	    BlockType0 *M, double *ergb, bool Parallel) {
  //   printf("i=%d r=%d %d c=%d %d comp=%d %d bl=%d\n",
  //	 individuals,  min_row, max_row, min_col, max_col,
  //	 compressed_start,  compressed_end, blocks);

   table_type *table = TABLE3;
  assert(//min_row >= 0 && min_col >= 0 &&
	 max_row <= individuals && max_col <= individuals);

#ifdef DO_PARALLEL
#pragma omp parallel for num_threads(CORES) if (Parallel)
#endif	  
  for (Uint c=min_col; c<max_col; c++) {
    Uint r  =  c > min_row ? c : min_row;
    BlockType * Mc = M + c * blocks,
      *Mr = M + r * blocks;
    double *ptr = ergb + c * individuals,
      *res_ptr_r = ptr + r;
    for ( ; r<max_row; r++, res_ptr_r++, Mr += blocks) {
      Uint sum = 0;
      BlockType *MR = Mr + compressed_start,
	*MC = Mc + compressed_start;
      for (Uint s=compressed_start; s<compressed_end; s++, MR++, MC++) {
	block_compressed x;
	x.x = *MR & *MC;
	sum+= table[x.b[0]] + table[x.b[1]] + table[x.b[2]] + table[x.b[3]];
      }
      ergb[c + r * individuals] = *res_ptr_r = (double) sum;
     }
  }
}


double getValue3(BlockType0 *Indiv, Uint s) {
  block_compressed x;
  x.x =  Indiv[s / genuineCodesPerBlock];
  Uint idx = s % genuineCodesPerBlock,
    y = x.b[idx / CodesPerMiniblock] >> ( (idx % CodesPerMiniblock) * BitsPerCode); 
  //  printf("y=%d \n", (Uint) y);
  // printf("%10e\n", rev_geno_code[y & CodeMask]);
  //  printf("ok\n");

  return rev_geno_code[y & CodeMask];
}


SEXP get_matrix3(SEXP SNPxIndiv) {
  if (TABLE3 == NULL) Init3();
  Uint 
    *info = GetInfo(SNPxIndiv),
    individuals = info[INDIVIDUALS],
    snps = info[SNPS],
    blocks = Blocks(snps),    
    endfor = individuals * blocks;
  SEXP Ans;
  PROTECT(Ans = get_matrix23_start(individuals, snps, R_NilValue));
  Uint *ans = (Uint *) INTEGER(Ans);
  BlockType *M = (BlockType0 *) Align(SNPxIndiv, ALIGN_23);
  for (Uint a=0; a<endfor; a+=blocks) {
    BlockType *Ma = M + a;
    for (Uint s=0; s<snps; s++) *(ans++) = getValue3(Ma, s);
  }
  UNPROTECT(1);
  return Ans;
}


SEXP get_matrixN_3(SEXP SNPxIndiv, SEXP Snps) {
  if (TABLE3 == NULL) Init3();
  GET_MATRIX_N_23(getValue3);
}


SEXP allele_freq3(SEXP SNPxIndiv) {
 if (TABLE3 == NULL) Init3();
 ALLELE_FREQ23(getValue3);
}


void matrix3_mult(Uint * SNPxIndiv, Uint snps, Uint individuals, double *A) {
  Uint blocks = Blocks(snps);
  MmPsq3(individuals, 0, individuals, 0, individuals, 0, blocks, blocks,
	 (BlockType0 *) SNPxIndiv, A, true);
}

Uint *Align3(SEXP Code, Uint nr, bool test) { return AlignTest(Code, nr, test); }

void zeroNthGeno3 ZERONTH
```
