## Supplementary material for "MoBPS - Modular Breeding Program Simulator": miraculix R-package: AutoMiraculix.cc

```
/*
 Authors 
 Martin Schlather,


 Copyright (C) 2018 -- 2019  Martin Schlather

This program is free software; you can redistribute it and/or
modify it under the terms of the GNU General Public License
as published by the Free Software Foundation; either version 3
of the License, or (at your option) any later version.

This program is distributed in the hope that it will be useful,
but WITHOUT ANY WARRANTY; without even the implied warranty of
MERCHANTABILITY or FITNESS FOR A PARTICULAR PURPOSE.  See the
GNU General Public License for more details.

You should have received a copy of the GNU General Public License
along with this program; if not, write to the Free Software
Foundation, Inc., 59 Temple Place - Suite 330, Boston, MA  02111-1307, USA.  
*/

#include "AutoMiraculix.h"

const char // muss in separater Zeile sein
// nachfolgende Zeile eingerueckt um 0 oder 2 Zeichen
*SNPCODING_NAME[nr_snpcoding] = {
  "Shuffle", "Shuffle256",
  "TwoBit", "ThreeBit", "Hamming2", "Hamming3", "NoSNPcoding",
  "NoSNPcodingR", "AutoCoding", "Haplo"},
  
*WHAT_NAMES[LAST_WHAT + 1] = {"haplo vector", "geno matrix"};
```
