## Supplementary material for "MoBPS - Modular Breeding Program Simulator": miraculix R-package: AutoMiraculix.h

```
/*
 Authors 
 Martin Schlather,

 Copyright (C) 2017 -- 2019 Martin Schlather

This program is free software; you can redistribute it and/or
modify it under the terms of the GNU General Public License
as published by the Free Software Foundation; either version 3
of the License, or (at your option) any later version.

This program is distributed in the hope that it will be useful,
but WITHOUT ANY WARRANTY; without even the implied warranty of
MERCHANTABILITY or FITNESS FOR A PARTICULAR PURPOSE.  See the
GNU General Public License for more details.

You should have received a copy of the GNU General Public License
along with this program; if not, write to the Free Software
Foundation, Inc., 59 Temple Place - Suite 330, Boston, MA  02111-1307, USA.  
*/

#ifndef AutoMiraculix_H
#define AutoMiraculix_H 1
#include <R.h>

typedef enum snpcoding {Shuffle,  // SSE2 0
			Shuffle256, // or AVX 1
			TwoBit,   // none needed 2
			ThreeBit, // none needed, 3
			Hamming2, // SSE2 4
			Hamming3, // SSSE3, 5
			NoSNPcoding, // none needed, 6
			NoSNPcodingR, // none needed, 7
			AutoCoding,  // 8
			Haplo
} snpcoding;

#define last_usr_meth AutoCoding // DO NEVER CHANGE
#define nr_snpcoding (Haplo + 1)


// WHAT just doubles the information that is also available through
// the class information, but easier to access on the C level for
// historical reasons. To be deleted maybe somewhen.
#define HAPLO 0
// #define GENO 1
#define GENOMATRIX 1
#define LAST_WHAT 1


// Coding of the attribute "information"
// !!!! ACHTUNG ZAHLEN MUESSEN DIE GLEICHEN BLEIBEN !!!!

#define WHAT 0
#define SNPS 1  // Wert darf auf keinen Fall geaendert werden
#define INDIVIDUALS 2 // Wert darf auf keinen Fall geaendert werden
#define ADDR0 3
#define ADDR1 4 // Achtung! Zweiter Teil von ADDR !!
#define ALIGNADDR0 5
#define ALIGNADDR1 6
#define SUMGENO 7
#define SUMGENO_E9 8
//#define MEMinUNITS1 9  //   unused !
// #define INFO_BLOCKS 11 // unsused !
#define SNPxIND 10 // INFO_INDIV_PER_COL, i.e. 'percolumn'
#define BITSPERCODE 12
#define BYTESPERBLOCK 13
#define CODESPERBLOCK 14
#define HEADER 15
#define DOUBLEINDIV 16
#define LEADINGCOL 17
#define MEMinUNITS0 18  // total genuine memory used to store all matrices --
// neither memory for allignment is not included, nor memory for info
#define MEMinUNITS1 19  
#define ALIGNEDUNITS0 20  // memory needed including alignment
#define ALIGNEDUNITS1 21
// ACHTUNG: gegebenenfalls haplogeno.cc:copyGeno aendern!!
#define INFO_LAST ALIGNEDUNITS1

#define GENOMICMATRIX "genomicmatrix"
#define HAPLOMATRIX "haplomatrix"
#define ORIGINVECTOR "origindata"


extern const char *SNPCODING_NAME[nr_snpcoding],
  *WHAT_NAMES[LAST_WHAT + 1];

#endif
```
