## Supplementary material for "MoBPS - Modular Breeding Program Simulator": miraculix R-package: Bit23.cc

```
/*
 Authors 
 Martin Schlather,


 Copyright (C) 2018 -- 2019  Martin Schlather

This program is free software; you can redistribute it and/or
modify it under the terms of the GNU General Public License
as published by the Free Software Foundation; either version 3
of the License, or (at your option) any later version.

This program is distributed in the hope that it will be useful,
but WITHOUT ANY WARRANTY; without even the implied warranty of
MERCHANTABILITY or FITNESS FOR A PARTICULAR PURPOSE.  See the
GNU General Public License for more details.

You should have received a copy of the GNU General Public License
along with this program; if not, write to the Free Software
Foundation, Inc., 59 Temple Place - Suite 330, Boston, MA  02111-1307, USA.  
*/


// gcc -mavx -o hello_avx hello_avx.c

#include "AutoMiraculix.h"
#include "Bit23.intern.h"
//#include "haplogeno.h"


void initiate_tableI(table_type **TABLE, Uint TABLESIZE,
		     Uint codesPerMiniblock, Uint bitsPerCode,
		     BlockType0 *result_code,
		     Uint *result_value, Uint NrResults) { // hash table

  Uint d,
    *nx;
  nx = (Uint *) CALLOC(codesPerMiniblock, sizeof(Uint));
  
  *TABLE = (table_type*) CALLOC(TABLESIZE, sizeof(table_type));
   table_type* table = *TABLE;

  while (true) {
    BlockType value = 0;
    Uint shift,
      sum = 0;
    
    for (d=shift=0; d<codesPerMiniblock; d++, shift+=bitsPerCode) {
      value |= result_code[nx[d]] << shift;
      sum += result_value[nx[d]];
    }
      
    assert(value < TABLESIZE && table[value] == 0);
    table[value] = sum;
    d = 0;
    nx[d]++;
    while (nx[d] >= NrResults) {
      nx[d] = 0;
      if (++d >= codesPerMiniblock) break;
      nx[d]++;
    }
    if (d >= codesPerMiniblock) break;
  }
  FREE(nx);
}


SEXP matrix_coding23(Uint *M, Uint snps, Uint individuals, snpcoding method){
  SEXP Code;
 
  if (method == TwoBit) {
    PROTECT(Code = matrix_start2(snps, individuals, R_NilValue));
    matrix_coding2(M, 0, individuals, 0, snps, snps, Code, NULL);
  } else if (method == ThreeBit) {
    PROTECT(Code = matrix_start3(snps, individuals, R_NilValue));
    matrix_coding3(M, 0, individuals, 0, snps, snps, Code, NULL);
  } else BUG;

  UNPROTECT(1);
  return Code;
}

SEXP get_matrix23_start(Uint individuals, Uint snps,
		      SEXP VARIABLE_IS_NOT_USED G) {
  SEXP Ans;
  PROTECT(Ans=allocMatrix(INTSXP, snps, individuals));
  UNPROTECT(1);
  return Ans;
}


void printbits(BlockType0 x, Uint size, Uint bitsPerCode) {
  BlockType mask = UINT64_C(1);  
  for (Uint d=0, zaehler=0; d<size; d++) {
    if (zaehler++ == bitsPerCode) {
      PRINTF(".");
      zaehler = 1;
    }
    PRINTF("%d", (x & mask) != 0);
    x >>= 1;
  }
  PRINTF(" ");
}
```
