## Supplementary material for "MoBPS - Modular Breeding Program Simulator": miraculix R-package: Bit23.h

```
/*
 Authors 
 Martin Schlather,


 Copyright (C) 2018 -- 2019  Martin Schlather

This program is free software; you can redistribute it and/or
modify it under the terms of the GNU General Public License
as published by the Free Software Foundation; either version 3
of the License, or (at your option) any later version.

This program is distributed in the hope that it will be useful,
but WITHOUT ANY WARRANTY; without even the implied warranty of
MERCHANTABILITY or FITNESS FOR A PARTICULAR PURPOSE.  See the
GNU General Public License for more details.

You should have received a copy of the GNU General Public License
along with this program; if not, write to the Free Software
Foundation, Inc., 59 Temple Place - Suite 330, Boston, MA  02111-1307, USA.  
*/


#ifndef miraculix_Bit23_H
#define miraculix_Bit23_H 1


#include "MX.h"
#include "AutoMiraculix.h"


SEXP matrix_coding23(Uint *M, Uint snps, Uint individuals, snpcoding method);


SEXP get_matrix2(SEXP SNPxIndiv);
SEXP matrix_start2(Uint snps, Uint individuals, SEXP file);
void matrix_coding2(Uint *SNPxIndiv, Uint start_individual, Uint end_individual,
		    Uint start_snp, Uint end_snp, Uint Mnrow,
		    SEXP Ans, double *G);
Uint *Align2(SEXP Code, Uint nr, bool test);
void Init2();
Uint CodesPerBlock2();
Uint UnitsPerIndiv2(Uint snps);
Ulong sumGeno2(Uint *S, Uint snps, Uint individuals);
void haplo2geno2(Uint * SNPxIndiv, Uint snps, Uint individuals, Uint, Uint *A);
Ulong sumGeno2(Uint *S, Uint snps, Uint individuals);
void matrix2_mult(Uint * SNPxIndiv, Uint snps, Uint individuals, double *A);
SEXP allele_freq2(SEXP SNPxIndiv);
SEXP get_matrixN_2(SEXP SNPxIndiv, SEXP Snps);
void zeroNthGeno2(SEXP SNPxIndiv, SEXP Snps);
void ReUseAsTwoBit(SEXP Code);


void matrix_coding3(Uint *SNPxIndiv, Uint start_individual, Uint end_individual,
		    Uint start_snp, Uint end_snp, Uint Mnrow,
		    SEXP Ans, double *G);
SEXP get_matrix3(SEXP SNPxIndiv);
SEXP matrix_start3(Uint snps, Uint individuals, SEXP file);

Uint *Align3(SEXP Code, Uint nr, bool test);

void Init3();
Uint CodesPerBlock3();
Uint UnitsPerIndiv3(Uint snps);
Ulong sumGeno3(Uint *S, Uint snps, Uint individuals);
void haplo2geno3(Uint * SNPxIndiv, Uint snps, Uint individuals, Uint, Uint *A);
Ulong sumGeno3(Uint *S, Uint snps, Uint individuals);
void matrix3_mult(Uint * SNPxIndiv, Uint snps, Uint individuals, double *A);
SEXP allele_freq3(SEXP SNPxIndiv);
SEXP get_matrixN_3(SEXP SNPxIndiv, SEXP Snps);
void zeroNthGeno3(SEXP SNPxIndiv, SEXP Snps);

#endif
```
