## Supplementary material for "MoBPS - Modular Breeding Program Simulator": miraculix R-package: Bit23.intern.h

```
/*
 Authors 
 Martin Schlather,


 Copyright (C) 2018 -- 2019  Martin Schlather

This program is free software; you can redistribute it and/or
modify it under the terms of the GNU General Public License
as published by the Free Software Foundation; either version 3
of the License, or (at your option) any later version.

This program is distributed in the hope that it will be useful,
but WITHOUT ANY WARRANTY; without even the implied warranty of
MERCHANTABILITY or FITNESS FOR A PARTICULAR PURPOSE.  See the
GNU General Public License for more details.

You should have received a copy of the GNU General Public License
along with this program; if not, write to the Free Software
Foundation, Inc., 59 Temple Place - Suite 330, Boston, MA  02111-1307, USA.  
*/


#ifndef miraculix_Bit23intern_H
#define miraculix_Bit23intern_H 1

#include <stdio.h>
#ifdef DO_PARALLEL
#include <omp.h>
#endif
#include "intrinsics.h"
//#include "haplogeno.h"
#include "options.h"
#include "error.h"
#include "MX.h"
#include "Haplo.h"

#define BytesPerMiniblock 2L
#define BitsPerMiniblock (BitsPerByte * BytesPerMiniblock)
#define MiniblocksPerBlock (BytesPerBlock / BytesPerMiniblock)
#define CodesPerMiniblock (BitsPerMiniblock / BitsPerCode)
#define genuineBitsPerMiniblock (CodesPerMiniblock * BitsPerCode)
#define two_genuineBitsPerMiniblock (1L << genuineBitsPerMiniblock)
#define MiniblocksPerBlock (BytesPerBlock / BytesPerMiniblock)
#define deltaMiniblockBits (BitsPerMiniblock - genuineBitsPerMiniblock)
#define genuineCodesPerBlock (MiniblocksPerBlock * CodesPerMiniblock)

#define TABLE_SIZE  two_genuineBitsPerMiniblock
 
#define nr_genotypes 3
#define nr_results 4

typedef uint16_t blockarray[MiniblocksPerBlock];
typedef union block_compressed {
  blockarray b;
  BlockType x;
} block_compressed;


void initiate_table2();
void initiate_table3();

typedef char table_type;
void initiate_tableI(table_type **TABLE, Uint TABLESIZE,
		     Uint codesPerMiniblock, Uint bits, BlockType0 *result_code,
		     Uint *result_value, Uint NrResults);

SEXP get_matrix23_start(Uint individuals, Uint snps, SEXP G);


void printbits(BlockType0 x, Uint size, Uint bits);

SEXP matrix_start2(Uint snps, Uint individuals, SEXP file);
void matrix_coding2(Uint *SNPxIndiv, Uint start_individual, Uint end_individual,
		    Uint start_snp, Uint end_snp, Uint Mnrow,
		    SEXP Ans, double *G);

SEXP matrix_start3(Uint snps, Uint individuals,  SEXP file);
void matrix_coding3(Uint *SNPxIndiv, Uint start_individual, Uint end_individual,
		    Uint start_snp, Uint end_snp, Uint Mnrow,
		    SEXP Ans, double *G);


/* matrix_start */
#define START23(Init23) (Uint snps, Uint individuals, SEXP file) {	\
  assert(genuineCodesPerBlock == CodesPerBlock);			\
  assert(TWO_GENUINEBITSPERMINIBLOCK == two_genuineBitsPerMiniblock);	\
  Init23();								\
  Ulong memPerIndiv = Blocks(snps) * UnitsPerBlock;			\
  SEXP Code =  CreateEmptyCodeVector(snps, individuals, memPerIndiv);	\
  PROTECT(Code);							\
  start_info(Code, file, UnitsPerBlock, genuineCodesPerBlock);		\
  UNPROTECT(1);								\
  return(Code);								\
  }


#define sumGeno23 (Uint *S, Uint snps, Uint individuals) {\
    Ulong						  \
      blocks = (Ulong) Blocks(snps) * individuals,	  \
      units = blocks * UnitsPerBlock,			  \
      sum = 0L;						  \
    Uint counter = 0;					  \
  for (Ulong i=0; i<units; i++) {			  \
    Uint s = S[i];					  \
    for (Uint u=0; u<CodesPerUnit; u++) {		  \
      sum += rev_geno_code[s & CodeMask];		  \
      s >>= BitsPerCode;				  \
      if (++counter >= CodesPerMiniblock) {		  \
	s >>= deltaMiniblockBits;			  \
	counter = 0;					  \
      }							  \
    }							  \
  }							  \
  return sum;						  \
  }


#define Coding23Matrix							\
  (Uint *X, Uint start_individual, Uint end_individual,		\
   Uint start_snp, Uint end_snp, Uint Mnrow,				\
   SEXP Ans, double VARIABLE_IS_NOT_USED * G) {	/* closing in .cc file */ \
  if (start_snp % genuineCodesPerBlock != 0) BUG;			\
  Uint *info = GetInfo(Ans),						\
    snps = info[SNPS];							\
  BlockType *ans = ((BlockType0*) Align(Ans, ALIGN_23)			\
		    ) + start_snp / genuineCodesPerBlock


#define Coding23(start_individual, end_individual, start_snp, end_snp, Mnrow, FROM) \
  Uint blocks = Blocks(snps);	 \
  for (Uint i=start_individual; i<end_individual; i++) {  \
    Uint *pX = X + (i - start_individual) * Mnrow,	\
      shift = 0,							\
      counter = 0;							\
    BlockType compressed = 0,						\
      *pAns = (BlockType0 *) ans + i * blocks;				\
    for (Uint s=start_snp; s<end_snp; s++) {				\
      compressed |= geno_code[FROM] << shift;				\
      shift += BitsPerCode;						\
      if (++counter >= CodesPerMiniblock) {				\
	shift += deltaMiniblockBits;					\
	counter = 0;							\
      }									\
      if (shift >= BitsPerBlock) {					\
	*pAns = compressed;						\
	pAns++;								\
	shift = counter = 0;						\
	compressed = 0;							\
      }									\
    }									\
    if (shift > 0) {							\
      *pAns = compressed;						\
      pAns++;								\
    }									\
  }


#define GET_MATRIX_N_23(GET)						\
  Uint									\
  *info = GetInfo(SNPxIndiv),						\
    individuals = info[INDIVIDUALS],					\
    len = length(Snps),							\
    *which = (Uint*) INTEGER(Snps),					\
    snps = info[SNPS],							\
    blocks = Blocks(snps),						\
    endfor = individuals * blocks;					\
  SEXP Ans;								\
  PROTECT(Ans = get_matrix23_start(individuals, len, R_NilValue));	\
  Uint *ans = (Uint *) INTEGER(Ans);					\
  BlockType *M = (BlockType0 *) Align(SNPxIndiv, ALIGN_23);		\
  for (Uint a=0; a<endfor; a+=blocks) {					\
    BlockType *Ma = M + a;						\
    for (Uint s=0; s<len; s++) {					\
      Rint S = which[s];						\
      *(ans++) = GET(Ma, S);						\
    }									\
  }									\
  UNPROTECT(1);								\
  return Ans

#define ALLELE_FREQ23(GET)						\
  Uint									\
  *info = GetInfo(SNPxIndiv),						\
    individuals = info[INDIVIDUALS],					\
    snps = info[SNPS],							\
    blocks = Blocks(snps),						\
    endfor = individuals * blocks;					\
  SEXP Ans;								\
  PROTECT(Ans=allocVector(REALSXP, snps));				\
  double *ans = REAL(Ans);						\
  for (Uint i=0; i<snps; ans[i++] = 0.0);				\
  BlockType *M = (BlockType0 *) Align(SNPxIndiv, ALIGN_23);		\
  for (Uint a=0; a<endfor; a+=blocks) {					\
    BlockType *Ma = M + a;						\
    for (Uint s=0; s<snps; s++) ans[s] += (double) GET(Ma, s);		\
  }									\
  double factor = 0.5 / (double) individuals;				\
  for (Uint i=0; i<snps; i++) ans[i] *= factor;				\
  UNPROTECT(1);								\
  return Ans


#define ZERONTH								\
  (SEXP SNPxIndiv, SEXP Snps) {						\
    Uint								\
      *info = GetInfo(SNPxIndiv),					\
      individuals = info[INDIVIDUALS],					\
      snps = info[SNPS],						\
      len = length(Snps),						\
      *which = (Uint*) INTEGER(Snps),					\
      blocks = Blocks(snps),						\
      endfor = individuals * blocks;					\
    BlockType *M = (BlockType0 *) Align(SNPxIndiv, ALIGN_23);		\
    for (Uint a=0; a<endfor; a+=blocks) {				\
      BlockType *Ma = M + a;						\
      for (Uint s=0; s<len; s++) {					\
	Rint S = which[s];						\
	Uint  j = S / genuineCodesPerBlock,				\
	  idx = S % genuineCodesPerBlock;				\
	block_compressed x;						\
	x.x = Ma[j];							\
	x.b[idx / CodesPerMiniblock] &=					\
	  ~(CodeMask << ( (idx % CodesPerMiniblock) * BitsPerCode));	\
	Ma[j] = x.x;							\
      }									\
    }									\
  }


#endif
```
