## Supplementary material for "MoBPS - Modular Breeding Program Simulator": miraculix R-package: error.h

```
/*
 Authors 
 Martin Schlather,


 Copyright (C) 2015 -- 2019 Martin Schlather

This program is free software; you can redistribute it and/or
modify it under the terms of the GNU General Public License
as published by the Free Software Foundation; either version 3
of the License, or (at your option) any later version.

This program is distributed in the hope that it will be useful,
but WITHOUT ANY WARRANTY; without even the implied warranty of
MERCHANTABILITY or FITNESS FOR A PARTICULAR PURPOSE.  See the
GNU General Public License for more details.

You should have received a copy of the GNU General Public License
along with this program; if not, write to the Free Software
Foundation, Inc., 59 Temple Place - Suite 330, Boston, MA  02111-1307, USA.  
*/


#ifndef miraculix_error_H
#define miraculix_error_H 1

#include <General_utils.h>

#ifdef DO_PARALLEL
#define LOCAL_MSG char MSG[LENERRMSG]

#else  // not DO_PARALLEL
#define LOCAL_MSG
extern char ERRMSG[LENERRMSG], MSG[LENERRMSG], BUG_MSG[LENERRMSG], MSG2[LENERRMSG];
extern errorloc_type ERROR_LOC;
extern errorstring_type ERRORSTRING;
#endif

#define STOP // assert(false)

#endif
```
