## Supplementary material for "MoBPS - Modular Breeding Program Simulator": miraculix R-package: hamming.cc

```
/*
 Authors 
 Martin Schlather,


Copyright (C) 2014 -- 2019 Martin Schlather
Copyright (C) 2014 -- 2015 Florian Skene: SSE2+SSSE3

This program is free software; you can redistribute it and/or
modify it under the terms of the GNU General Public License
as published by the Free Software Foundation; either version 3
of the License, or (at your option) any later version.

This program is distributed in the hope that it will be useful,
but WITHOUT ANY WARRANTY; without even the implied warranty of
MERCHANTABILITY or FITNESS FOR A PARTICULAR PURPOSE.  See the
GNU General Public License for more details.

You should have received a copy of the GNU General Public License
along with this program; if not, write to the Free Software
Foundation, Inc., 59 Temple Place - Suite 330, Boston, MA  02111-1307, USA.  
*/

#include "IntrinsicsBase.h"

#if defined SSSE3 || defined SSE2

#if defined SSSE3 && !defined SSE2
#define SSE2 1
#endif

#if defined AVX2
#undef AVX2
#endif

#if defined AVX256
#undef AVX256
#endif


#define BitsPerCode 4L
#include "intrinsics.h"
#include "miraculix.h"
#include "xport_import.h"
#include "options.h"
#include "AutoMiraculix.h"
#include <General_utils.h>
#include "MX.h"
#include "error.h"
#include "haplogeno.h"
#include "Haplo.h"
INLINER


#define repetSSE2 10L
#define atonceSSE2 (3L * repetSSE2)
#define atonce_SSSE3 4L

Uint CodesPerBlockH() { return CodesPerBlock; }
Ulong UnitsPerIndivH(Uint snps, Uint method) {
  Uint
    atonce = method == Hamming2 ? atonceSSE2 : atonce_SSSE3, // wechsel 64 bit auf 128; 3 bzw 4
    //                                       auf ein Mal; sse2: 10x wiederholt
    //minmem = (1L + (snps - 1L) / (2 * atonce)) * (2 * atonce),
    Xblocks = (1L + (snps - 1L) / (atonce * CodesPerBlock)) * atonce;
  return UnitsPerBlock * Xblocks;
}

static SEXP create_codevectorSSE(Uint snps, Uint individuals, snpcoding method){
  assert(BytesPerBlock == sizeof(BlockType0));
  Ulong mem2 = 2 * UnitsPerIndivH(snps, method); 
  SEXP Code = CreateEmptyCodeVector(snps, individuals, mem2);
  //if (PRESERVE) {BUG;  R_PreserveObject(C ode);}
  return Code;
}


static uint64_t coding1[] = {0, 5, 10},    //0, 1, 2
  coding2[] = {0, 3, 15},    //0, 1, 2
  coding3[] = {0, 6, 15},    //0, 1, 2
  coding4[] = {0, 3, 15};    //0, 1, 2
Uint rev_coding1[16] = {0, 0, 0, 0,  0, 1, 0, 0,  0, 0, 2, 0,  0, 0, 0, 0},
// rev_coding24[16] = {0, 0, 0, 1,  0, 0, 0, 0,  0, 0, 0, 0,  0, 0, 0, 2},
 rev_coding3[16] = {0, 0, 0, 0,  0, 0, 1, 0,  0, 0, 0, 0,  0, 0, 0, 2};


#define  MatrixCoding(start_snp, cur_snps, start_individual, cur_indiv, FROM) \
bool hamming2 = method == Hamming2;				   \
  Ulong memPerMatrix = UnitsPerIndivH(snps, method) * individuals, \
		 blocksPerMatrix =  memPerMatrix / UnitsPerBlock;	\
  Uint  blocks = blocksPerMatrix / individuals, /* NICHT Blocks(snps) !! */ \
		 mini = sizeof(BlockType0) / sizeof(uint64_t),		\
		 miniblocks = blocks * mini,				\
		 CodesPerMiniblock = CodesPerBlock / mini;		\
  uint64_t *code1 = hamming2 ? coding1 : coding3,			\
		 *code2 = hamming2 ? coding2 : coding4,			\
         *M = (uint64_t*) ((BlockType0 *) code + start_snp / CodesPerBlock + \
			   start_individual * blocks),			\
		 *M_T = (uint64_t*) ((BlockType0 *) M + blocksPerMatrix);	\
  for (Uint i = 0; i<cur_indiv; i++) {					\
    Uint count=0,							\
      *pX = X + i * Mnrow;						\
    uint64_t *Mptr = M + i * miniblocks,				\
      *M_Tptr = M_T + i * miniblocks;					\
    for (Uint s = 0; s<cur_snps; s++) {					\
      Uint snp = FROM;							\
      *Mptr = *Mptr << BitsPerCode;					\
      *Mptr |= code1[snp];						\
      *M_Tptr = *M_Tptr << BitsPerCode;					\
      *M_Tptr |= code2[snp];						\
      if (++count == CodesPerMiniblock) {				\
	count = 0;							\
	Mptr++;								\
	M_Tptr++;							\
      }									\
    }									\
    if (count > 0) {							\
      Uint shift = BitsPerCode * (CodesPerMiniblock - count);		\
      *Mptr = *Mptr << shift;						\
      *M_Tptr = *M_Tptr << shift;					\
    }									\
  }								       
 
   

void haplo2genoH(Uint *X, Uint snps, Uint individuals,
		 Uint Mnrow, snpcoding method, Uint *code) {
  MatrixCoding(0, snps, 0, individuals, FROMHAPLO);
}


void matrixH(Uint *X, Uint start_individual, Uint end_individual, 
	     Uint start_snp, Uint end_snp, Uint Mnrow,
	     snpcoding method,
	     SEXP Code) {
  if (start_snp % CodesPerBlock != 0) BUG;

  Uint
    *info = GetInfo(Code),
    individuals = info[INDIVIDUALS],
    snps = info[SNPS],
    cur_snps = end_snp - start_snp,
    cur_indiv = end_individual-start_individual,
    *code = (Uint *) Align(Code, ALIGN_SSE);

  MatrixCoding(start_snp, cur_snps, start_individual, cur_indiv, FROMINPUT);
}


void matrixH2(Uint *M, Uint start_individual, Uint end_individual, 
	     Uint start_snp, Uint end_snp, Uint Mnrow, 
	     SEXP Code,  double VARIABLE_IS_NOT_USED *G) { // from file_get
  matrixH(M, start_individual, end_individual, start_snp, end_snp, Mnrow,
	  Hamming2, Code);
}

void matrixH3(Uint *M, Uint start_individual, Uint end_individual, 
	     Uint start_snp, Uint end_snp, Uint Mnrow, 
	     SEXP Code,  double VARIABLE_IS_NOT_USED *G) { // from file_get
  matrixH(M, start_individual, end_individual, start_snp, end_snp, Mnrow,
	  Hamming3, Code);
}


#define DOGET(DO)							\
  Uint *rev = method == Hamming2 ? rev_coding1 : rev_coding3,		\
       memPerMatrix = UnitsPerIndivH(snps, method) * individuals, \
       blocksPerMatrix =  memPerMatrix / UnitsPerBlock,	\
       blocks = blocksPerMatrix / individuals,		\
       mini = sizeof(BlockType0) / sizeof(uint64_t),	\
       miniblocksPerMatrix = blocksPerMatrix * mini,	\
       miniblocks = blocks * mini,			\
       BitsPerMiniblock = BitsPerBlock / mini,		\
       CodesPerMiniblock = CodesPerBlock / mini;	\
  for (Uint a=0; a<miniblocksPerMatrix; a+=miniblocks) {		\
    uint64_t *Ma = ((uint64_t*) M) + a;					\
    for (Uint s=0; s<snps; s++) {					\
      DO rev[(Ma[s / CodesPerMiniblock]					\
	      >> (BitsPerMiniblock - BitsPerCode -			\
		  BitsPerCode  * (s % CodesPerMiniblock))) & CodeMask];	\
    }									\
  }


#define ANSPLUS *(ans++) =
SEXP get_matrixH(SEXP SNPxIndiv) {
  Uint 
     *info = GetInfo(SNPxIndiv),
    individuals = info[INDIVIDUALS],
    snps = info[SNPS];
  Uint method = MethodOf(SNPxIndiv);
  SEXP Ans;
  
  PROTECT(Ans=allocMatrix(INTSXP, snps, individuals));
  Uint *ans = (Uint *) INTEGER(Ans);
  uint64_t *M = (uint64_t*) Align(SNPxIndiv, ALIGN_SSE);

  DOGET(ANSPLUS);
 
  UNPROTECT(1);
  return Ans;
}

#define SUMUP sum +=
Ulong sumGenoH(Uint *M, Uint snps, Uint individuals, snpcoding method)  {
  Ulong sum = 0L;
  DOGET(SUMUP);
  return sum;
}


#define FREQ ans[s] += (double)

SEXP allele_freqH(SEXP SNPxIndiv) {
  Uint 
    *info = GetInfo(SNPxIndiv),
    individuals = info[INDIVIDUALS],
    snps = info[SNPS];
  Uint method = MethodOf(SNPxIndiv);
  SEXP Ans;
  uint64_t *M = (uint64_t*) Align(SNPxIndiv, ALIGN_SSE);
  
  PROTECT(Ans=allocVector(REALSXP, snps));
  double *ans = REAL(Ans); 
  for (Uint i=0; i<snps; ans[i++] = 0.0);
  

  DOGET(FREQ);
  double factor = 0.5 / (double) individuals;
  for (Uint i=0; i<snps; i++) ans[i] *= factor;
 
  UNPROTECT(1);
  return Ans;
}


#define DONGET(DO)							\
  Uint *rev = method == Hamming2 ? rev_coding1 : rev_coding3,		\
       memPerMatrix = UnitsPerIndivH(snps, method) * individuals, \
       blocksPerMatrix =  memPerMatrix / UnitsPerBlock,	\
       blocks = blocksPerMatrix / individuals,		\
       mini = sizeof(BlockType0) / sizeof(uint64_t),	\
       miniblocksPerMatrix = blocksPerMatrix * mini,	\
       miniblocks = blocks * mini,			\
       BitsPerMiniblock = BitsPerBlock / mini,		\
       CodesPerMiniblock = CodesPerBlock / mini;	\
  for (Uint a=0; a<miniblocksPerMatrix; a+=miniblocks) {		\
    uint64_t *Ma = ((uint64_t*) M) + a;					\
    for (Uint s=0; s<len; s++) {					\
      Rint S = which[s];						\
      DO rev[(Ma[S / CodesPerMiniblock]					\
	      >> (BitsPerMiniblock - BitsPerCode -			\
		  BitsPerCode  * (S % CodesPerMiniblock))) & CodeMask];	\
    }									\
  }

#define ANSPLUS *(ans++) =
SEXP get_matrixN_H(SEXP SNPxIndiv, SEXP Snps) {
  Uint 
     *info = GetInfo(SNPxIndiv),
    individuals = info[INDIVIDUALS],
    len = length(Snps),
    *which = (Uint*) INTEGER(Snps),
    snps = info[SNPS];
  Uint method = MethodOf(SNPxIndiv);
  SEXP Ans;
  
  PROTECT(Ans=allocMatrix(INTSXP, len, individuals));
  Uint *ans = (Uint *) INTEGER(Ans);
  uint64_t *M = (uint64_t*) Align(SNPxIndiv, ALIGN_SSE);

  DONGET(ANSPLUS);
 
  UNPROTECT(1);
  return Ans;
}


SEXP zeroNthGenoH(SEXP SNPxIndiv, SEXP Snps, snpcoding method) {
  Uint 
     *info = GetInfo(SNPxIndiv),
    individuals = info[INDIVIDUALS],
    len = length(Snps),
    *which = (Uint*) INTEGER(Snps),
    snps = info[SNPS];
  uint64_t *M = (uint64_t*) Align(SNPxIndiv, ALIGN_SSE);

  Uint 	
    memPerMatrix = UnitsPerIndivH(snps, method) * individuals, 
    blocksPerMatrix =  memPerMatrix / UnitsPerBlock,	
    blocks = blocksPerMatrix / individuals,		
    mini = sizeof(BlockType0) / sizeof(uint64_t),	
    miniblocksPerMatrix = blocksPerMatrix * mini,	
    miniblocks = blocks * mini,			
    BitsPerMiniblock = BitsPerBlock / mini,		
    CodesPerMiniblock = CodesPerBlock / mini;	
  for (Uint a=0; a<miniblocksPerMatrix; a+=miniblocks) {		
    uint64_t *Ma = ((uint64_t*) M) + a;					
    for (Uint s=0; s<len; s++) {					
      Rint S = which[s];						
      Ma[S / CodesPerMiniblock]	&=
	~ (CodeMask << (BitsPerMiniblock - BitsPerCode - 		
			BitsPerCode  * (S % CodesPerMiniblock)));  // <<
    }							
  }
 
  return SNPxIndiv;
}


static void matrix_multH2(BlockType0 *M,
		   BlockType0  *M_T,
		   Uint individuals,
		   Uint snps,
		   double *ergb,
		   Uint  blocks) {
  // M*M^T:
  // outer for-loop represents rows of M (only within partition, if > 1 cores
  // inner for-loop represents columns of M^T starting at diagonal element
  Uint
    Xblocks = UnitsPerIndivH(snps, Hamming2) / UnitsPerBlock,
    loops = Xblocks / atonceSSE2;
  BlockType m1, m2, m4, m8;
  uint64_t OOO1 = UINT64_C(0x0001000100010001);

  SET8(m1, 0x55);
  SET8(m2, 0x33);
  SET8(m4, 0x0f);
  SET16(m8,0x00ff);

#ifdef DO_PARALLEL
#pragma omp parallel for num_threads(CORES) schedule(dynamic, 20)
#endif 
  for (Uint i = 0; i<individuals; i++) {
    BlockType *mpp = M + i * blocks;
    for (Uint j = i; j<individuals; j++) {
      BlockType
	*m_tpp = M_T + j * blocks,
	*mptr = mpp; //mptr set back to first element of row in 	
      uint64_t tot = 0;
      for (Uint s=0; s<loops; s++) {
	uni128 acc;
	ZERO(acc VI);
	// do-while runs 10x before continuing
	for (Uint t=0; t<repetSSE2; t++) {
	  BlockType L1, L2, half1, half2;
	  LOADU(L1, mptr); mptr++;
	  LOADU(L2, m_tpp); m_tpp++;
	  AND(half1, L1, L2);
	  SHR64(L1, half1, 1);	  
	  AND(half2, L1, m1);
	  AND(half1, half1, m1);
	  OR(half1, half1, half2);
	  LOADU(L1, mptr); mptr++;
	  LOADU(L2, m_tpp); m_tpp++;
	    BlockType count1, count2;
	  AND(count1, L1, L2);
	  ADD64(count1, count1, half1);
	  LOADU(L1, mptr); mptr++;
	  LOADU(L2, m_tpp);m_tpp++;
	  AND(count2, L1, L2);
	  ADD64(count2, count2, half2);
#define dummy1 half1
#define dummy2 half2
	  AND(dummy1, count1, m2);
	  SHR64(L1, count1, 2);
	  AND(dummy2, L1, m2);
	  ADD64(count1,dummy1, dummy2);
	  AND(dummy1, count2, m2);
	  SHR64(L1, count2, 2);
	  AND(dummy2, L1, m2);
	  ADD64(dummy1, dummy1, dummy2);
	  ADD64(count1, count1, dummy1);
	  AND(dummy1, count1, m4);
	  SHR64(L1, count1, 4);
	  AND(dummy2, L1, m4);
	  ADD64(dummy1, dummy1, dummy2);
	  ADD64(acc VI, acc VI, dummy1);
	}
	BlockType L1, L2;
	AND(L1, acc VI, m8);
	SHR64(L2, acc VI, 8);
	AND(L2, L2, m8);
	ADD64(acc VI, L1, L2);
	tot += ((acc.u64[0] + acc.u64[1]) * OOO1) >> 48;
      }
      ergb[j + i * individuals] = ergb[i + j * individuals] = (double) tot;
    }
  }
}
 

static void matrix_multH3(BlockType0 *M,
		   BlockType0 *M_T,
		   Uint individuals,
		   Uint snps,
		   double *ergb,
		   Uint blocks) {   
#if !defined SSSE3
  ERR("'Hamming3' need the availablity of 'SSSE3'");
#endif

  Uint
    Xblocks =  UnitsPerIndivH(snps, Hamming3) / UnitsPerBlock,
    loops = Xblocks / atonce_SSSE3;
  
  const unsigned _LUT[] = {0x02010100, 0x03020201, 0x03020201, 0x04030302};
  BlockType LUT, mask, zero;


  LOADU(LUT, (BlockType0 *) _LUT); // _mm_set_epi32(,,,)
  SET8(mask, 0x0F);
  ZERO(zero);

#ifdef DO_PARALLEL
#pragma omp parallel for num_threads(CORES) schedule(dynamic, 20)
#endif
  for (Uint i = 0; i<individuals; i++) {
    BlockType
      *mpp = M + i * blocks;
     for (Uint j = i; j<individuals; j++) {
     uint64_t tot = 0;
      BlockType *mptr = mpp,
	*m_tpp = M_T + j * blocks;
      for (Uint k = 0; k < loops; k++) {    // '/4' hinzugefuegt
	uni128  intermed;
	BlockType v0, lo, count0, count1, L1, L2;
#define COUNT(count0)						\
	LOADU(L1, mptr);mptr++;					\
	LOADU(L2, m_tpp);m_tpp++;				\
	AND(v0, L1, L2);					\
	AND(lo, mask, v0);					\
	SHR16(L1, v0, 4);					\
	AND(v0, mask, L1); /* hi */				\
	SHUFFLE8(lo, LUT, lo);					\
	SHUFFLE8(v0, LUT, v0);					\
	ADD8(count0, lo, v0);
	
	COUNT(count0);
	COUNT(v0);
	ADD8(count0, count0, v0);
	COUNT(count1);
	COUNT(v0);
	ADD8(count1, count1, v0);
	ADD8(count0, count0, count1);
	
	//bitwise difference between 'tota' and all zeros
	SAD8(intermed VI, count0, zero);
	// adds the two bitcounts which are saved as unsigned 64bit integers
	// to continuous bitcount
	tot += (Uint) intermed.u64[0] + (Uint)intermed.u64[1];
      }
      ergb[j + i * individuals] = ergb[i + j * individuals] = (double) tot;
    }
  }
}

void matrixH_mult(Uint *M, Uint snps, Uint individuals,
		  snpcoding method, double *A) {
  Uint
    memPerMatrix = UnitsPerIndivH(snps, method) * individuals,
    blocksPerMatrix =  memPerMatrix / UnitsPerBlock,
    blocks = blocksPerMatrix / individuals;
  BlockType *M_T =  (BlockType0 *) M + blocksPerMatrix;

  if (method == Hamming2)
    matrix_multH2((BlockType0 *) M, M_T, individuals, snps, A, blocks);
  else
    matrix_multH3((BlockType0 *) M, M_T, individuals, snps, A, blocks);
 
}

SEXP matrix_startH( Uint snps,Uint individuals, snpcoding method, SEXP file) {  
  SEXP Code;
  PROTECT(Code = create_codevectorSSE(snps, individuals, method));
  start_info(Code, file, UnitsPerBlock, CodesPerBlock);
  UNPROTECT(1);
  return(Code);
}

SEXP matrix_startH2(Uint snps,Uint individuals,  SEXP file) {  
  return matrix_startH(snps, individuals, Hamming2, file);
}
 
SEXP matrix_startH3( Uint snps, Uint individuals,SEXP file) {  
  return matrix_startH(snps, individuals, Hamming3, file);
}

  
SEXP matrix_codingH(Uint *M, Uint snps, Uint individuals, snpcoding method){
  SEXP Code;
  PROTECT(Code = matrix_startH(snps, individuals, method, R_NilValue));
  matrixH(M, 0, individuals, 0, snps, snps, method,  Code);
  UNPROTECT(1);
  return Code;
}


Uint *AlignH(SEXP Code, Uint nr, bool test) {
  return AlignTest(Code, nr, test); }


#else // no sse2
#include "error.h"
#include "MX.h"
void SSEmissing() {
  ERR("'Hamming' needs at least the availablity of 'SSE2'");
}
#define Sm { SSEmissing(); return R_NilValue; }
#define Su { SSEmissing(); return 0; }
#define Sv { SSEmissing(); }
#if defined VARIABLE_IS_NOT_USED
#define V VARIABLE_IS_NOT_USED
#else
#define V
#endif


void matrixH_mult(Uint V *M, Uint V snps, Uint V individuals,
		  snpcoding method, double V *A) {
  if (method == Hamming2) {ERR("Hamming snp coding needs at least SSE2");}
  else ERR("Hamming3 snp coding needs at least SSSE3");
}

SEXP matrix_codingH(Uint V *M, Uint V snps, Uint V individuals,
		    snpcoding V method)Sm
SEXP get_matrixH(SEXP V SNPxIndiv)Sm
SEXP matrix_startH(Uint V snps,Uint V individuals,  snpcoding V method,
		   SEXP V file)Sm
SEXP matrix_startH2( Uint V snps, Uint V individuals,SEXP V file)Sm
SEXP matrix_startH3( Uint V snps, Uint V individuals, SEXP V file)Sm
void matrixH2(Uint V *M, Uint V start_individual, Uint V end_individual, 
	     Uint V start_snp, Uint V end_snp, Uint V Mnrow, SEXP V Ans,
	     double V *G)Sv
void matrixH3(Uint V *M, Uint V start_individual, Uint V end_individual, 
	     Uint V start_snp, Uint V end_snp, Uint V Mnrow, SEXP V Ans,
	     double V *G)Sv
Uint V *AlignH(SEXP V Code, Uint V nr, bool V test)Su
int CodesPerBlockH()Su
Ulong UnitsPerIndivH(Uint V snps, Uint V method)Su
Ulong sumGenoPlain(Uint V *S, Uint V snps, Uint V individuals,
		   snpcoding V method)Su
void haplo2genoH(Uint V * SNPxIndiv, Uint V snps, Uint V individuals, Uint V U,
		 snpcoding V method, Uint V *A)Sv
Ulong sumGenoH(Uint V *M, Uint V snps, Uint V individuals, snpcoding V method)Su
SEXP zeroNthGenoH(SEXP V SNPxIndiv, SEXP V Snps, snpcoding V method)Sm
SEXP get_matrixN_H(SEXP V SNPxIndiv, SEXP V Snps)Sm
SEXP allele_freqH(SEXP V SNPxIndiv)Sm


#endif // sse2
```
