## Supplementary material for "MoBPS - Modular Breeding Program Simulator": miraculix R-package: hamming.h

```
/*
 Authors 
 Martin Schlather,


 Copyright (C) 2018 -- 2019  Martin Schlather

This program is free software; you can redistribute it and/or
modify it under the terms of the GNU General Public License
as published by the Free Software Foundation; either version 3
of the License, or (at your option) any later version.

This program is distributed in the hope that it will be useful,
but WITHOUT ANY WARRANTY; without even the implied warranty of
MERCHANTABILITY or FITNESS FOR A PARTICULAR PURPOSE.  See the
GNU General Public License for more details.

You should have received a copy of the GNU General Public License
along with this program; if not, write to the Free Software
Foundation, Inc., 59 Temple Place - Suite 330, Boston, MA  02111-1307, USA.  
*/


#ifndef miraculix_sse_H
#define miraculix_sse_H 1

SEXP matrix_codingH(Uint *M, Uint snps, Uint individuals, snpcoding method);
SEXP get_matrixH(SEXP SNPxIndiv);

SEXP matrix_startH( Uint snps, Uint individuals,snpcoding method, SEXP file);
SEXP matrix_startH2( Uint snps,Uint individuals, SEXP file);
SEXP matrix_startH3(Uint snps, Uint individuals, SEXP file);
void matrixH2(Uint *M, Uint start_individual, Uint end_individual, 
	     Uint start_snp, Uint end_snp, Uint Mnrow, SEXP Ans,
	     double *G);
void matrixH3(Uint *M, Uint start_individual, Uint end_individual, 
	     Uint start_snp, Uint end_snp, Uint Mnrow, SEXP Ans,
	     double *G);

Uint *AlignH(SEXP Code, Uint nr, bool test);
int CodesPerBlockH();
Ulong UnitsPerIndivH(Uint snps, Uint Method);
Ulong sumGenoPlain(Uint *S, Uint snps, Uint individuals, snpcoding method);
void haplo2genoH(Uint * SNPxIndiv, Uint snps, Uint individuals, Uint U,
		 snpcoding method, Uint *A);
Ulong sumGenoH(Uint *M, Uint snps, Uint individuals, snpcoding method);
void matrixH_mult(Uint *SNPxIndiv, Uint snps, Uint individuals,
		    snpcoding method, double *A);

SEXP zeroNthGenoH(SEXP SNPxIndiv, SEXP Snps, snpcoding method);
SEXP get_matrixN_H(SEXP SNPxIndiv, SEXP Snps);
SEXP allele_freqH(SEXP SNPxIndiv);


#endif
```
