## Supplementary material for "MoBPS - Modular Breeding Program Simulator": miraculix R-package: Haplo.cc

```
/*
 Authors 
 Martin Schlather,


 Copyright (C) 2018 -- 2019  Martin Schlather

This program is free software; you can redistribute it and/or
modify it under the terms of the GNU General Public License
as published by the Free Software Foundation; either version 3
of the License, or (at your option) any later version.

This program is distributed in the hope that it will be useful,
but WITHOUT ANY WARRANTY; without even the implied warranty of
MERCHANTABILITY or FITNESS FOR A PARTICULAR PURPOSE.  See the
GNU General Public License for more details.

You should have received a copy of the GNU General Public License
along with this program; if not, write to the Free Software
Foundation, Inc., 59 Temple Place - Suite 330, Boston, MA  02111-1307, USA.  
*/


#define BitsPerCode 2L

#include "IntrinsicsBase.h"
#include "intrinsics.h"
#include <General_utils.h>
#include "xport_import.h"
#include "miraculix.h"
#include "MX.h"

#include "haplogeno.h"
INLINER

Uint CodesPerBlockHaplo() { return CodesPerBlock; }
Uint UnitsPerIndivHaplo(Uint snps) { return Blocks(snps) * UnitsPerBlock; }

void assert_haplo() {
}


Uint *AlignHaplo(SEXP Code, Uint nr, bool test) {
  return AlignTest(Code, nr, test); }


SEXP create_codevectorHaplo(Uint snps, Uint individuals) {
  assert_haplo();
  Uint memPerIndiv = Blocks(snps) * UnitsPerBlock;
  SEXP Code = CreateEmptyCodeVectorHaplo(snps, individuals, memPerIndiv);
  if (PRESERVE) {BUG; R_PreserveObject(Code);}
 return Code;
}


void InnerDetermHaplo(double *freq1, double *freq2,
		      Uint snps, Uint individuals, Uint *code) {
  Ulong units = Blocks(snps) * UnitsPerBlock,
    fullBlocks = snps / CodesPerBlock;
  //  printf("snps = %d\n", snps);
  for (Uint i=0; i<individuals; i++, code += units) {
    for (Uint j=0; j<fullBlocks; j++) {
      Uint *C = code + j * UnitsPerBlock,
	mm = j * CodesPerBlock;
      for (Uint u=0; u<CodesPerUnit; u++) {
	Uint shift = u * BitsPerCode;
	for (Uint k=0; k<UnitsPerBlock; k++, mm++) {
	  double f2 = freq2[mm];
	  Rint mm1 = freq1[mm] == 1.0,
	    mm2 = ISNA(f2) ? mm1 : (f2  == 1.0);
	  //	  printf("A mm=%d    %d %d     %f %f\n", mm, mm1, mm2, freq1[mm], freq2[mm]);
	  C[k] |= (mm1 << shift) | (mm2 << (shift + 1));
	}
      }
    }
    if (fullBlocks * CodesPerBlock < snps) {
      Uint 
	*C = code + fullBlocks * UnitsPerBlock,
	mm = CodesPerBlock * fullBlocks;
      for (Uint u=0; u<CodesPerUnit; u++) {     
	Uint shift = u * 2;
	for (Uint k=0; k<UnitsPerBlock && mm < snps; k++, mm++) {
	  double f2 = freq2[mm];
	  Rint mm1 = freq1[mm] == 1.0,
	    mm2 = ISNA(f2) ? mm1 : (f2  == 1.0);
	  // printf("B mm=%d    %d %d     %f %f\n", mm, mm1, mm2, freq1[mm], freq2[mm]);
	 
	  C[k] |= (mm1 << shift) | (mm2 << (shift + 1));
	}
	if (mm == snps) break;
      }
    }
  }
}


void InnerRandomHaplo(double *freq1, double *freq2,
		      Uint snps, Uint individuals, Uint *code) {
  Ulong units = Blocks(snps) * UnitsPerBlock;
  Uint 
    fullBlocks = snps / CodesPerBlock;
  //  printf("snps = %d\n", snps);
  for (Uint i=0; i<individuals; i++, code += units) {
    for (Uint j=0; j<fullBlocks; j++) {
      Uint *C = code + j * UnitsPerBlock,
	mm = j * CodesPerBlock;
      for (Uint u=0; u<CodesPerUnit; u++) {
	Uint shift = u * BitsPerCode;
	for (Uint k=0; k<UnitsPerBlock; k++, mm++) {
	  double f2 = freq2[mm];
	  Rint mm1 = UNIFORM_RANDOM <= freq1[mm],
	    mm2 = ISNA(f2) ? mm1 : UNIFORM_RANDOM <= f2;
	  //	  printf("C  mm=%d    %d %d     %f %f\n", mm, mm1, mm2, freq1[mm], freq2[mm]);
	  
	  C[k] |= (mm1 << shift) | (mm2 << (shift + 1));
	}
      }
    }
    if (fullBlocks * CodesPerBlock < snps) {
      Uint 
	*C = code + fullBlocks * UnitsPerBlock,
	mm = CodesPerBlock * fullBlocks;
      for (Uint u=0; u<CodesPerUnit; u++) {     
	Uint shift = u * 2;
	for (Uint k=0; k<UnitsPerBlock && mm < snps; k++, mm++) {
	  double f2 = freq2[mm];
	  Rint mm1 = UNIFORM_RANDOM <= freq1[mm],
	    mm2 = ISNA(f2) ? mm1 : UNIFORM_RANDOM <= f2;
	  //	  printf("D mm=%d    %d %d     %f %f %d\n", mm, mm1, mm2, freq1[mm], freq2[mm], ISNA(f2));
	  C[k] |= (mm1 << shift) | (mm2 << (shift + 1));
	}
	if (mm == snps) break;
      }
    }
    //  printf("\n");
  }
}


SEXP rhaplomatrix(SEXP Freq1, SEXP Freq2, SEXP Individuals) {
  Uint individuals = INTEGER(Individuals)[0],
    snps = length(Freq1);
  if (snps != (Uint) length(Freq2))
    ERR("'freq' and 'freq2' do not have the same length");
 return create_codevectorHaplo(snps, individuals);
}

  
SEXP rhaplomatrixPart2(SEXP Freq1, SEXP Freq2, SEXP Individuals, SEXP Code) {
  double *freq1 = REAL(Freq1),
    *freq2 = REAL(Freq2);
  Uint individuals = INTEGER(Individuals)[0],
    snps = length(Freq1),
    *C =  Align(Code, ALIGN_HAPLO);

  bool random = false;  
  for (Uint i=0; i < snps; i++) {
    double f = freq1[i];
    if (f < 0 || f > 1) ERR("frequencies not in [0,1]");
    random |= f != 0 && f != 1;
    f = freq2[i];
    if (!ISNA(f)) {
      if (f < 0 || f > 1) ERR("frequencies not in [0,1]");
      random |= f != 0 &&  f != 1;
    }
  }

  if (random) {
    GetRNGstate();
    // printf("snps=%d indivi=%d \n", snps, individuals);
    InnerRandomHaplo(freq1, freq2, snps, individuals, C);
    PutRNGstate();
  } else {
    InnerDetermHaplo(freq1, freq2, snps, individuals, C);
  }

  return Code;
}

void codeInnerHaplo2(Uint *M1, Uint *M2, Uint snps, Uint *code) {
  Uint j,
    fullBlocks = snps / CodesPerBlock;
#ifdef DO_PARALLEL
#pragma omp parallel for num_threads(CORES)
#endif
  for (j=0; j<fullBlocks; j++) {
    Uint *C = code + j * UnitsPerBlock,
      *mm1 = M1 + j * CodesPerBlock,
      *mm2 = M2 + j * CodesPerBlock;
    for (Uint u=0; u<CodesPerUnit; u++) {
      Uint shift = u * BitsPerCode;
      for (Uint k=0; k<UnitsPerBlock; k++, mm1++, mm2++) {
	assert(*mm1 == 0 || *mm1 == 1);
	assert(*mm2 == 0 || *mm2 == 1);
	C[k] |= (*mm1 << shift) | (*mm2 << (shift + 1));
      }
    } 
  }
  if (fullBlocks * CodesPerBlock < snps) {
    Uint *end = M1 + snps,
      *C = code + fullBlocks * UnitsPerBlock;
    M1 += CodesPerBlock * fullBlocks;
    M2 += CodesPerBlock * fullBlocks;
    for (Uint u=0; u<CodesPerUnit; u++) {
     
      Uint shift = u * 2;
      for (Uint k=0; k<UnitsPerBlock && M1 < end; k++, M1++, M2++) {
	assert(*M1 == 0 || *M1 == 1);
	assert(*M2 == 0 || *M2 == 1);
	C[k] |= (*M1 << shift) | (*M2 << (shift + 1));
      }
      if (M1 == end) break;
    }
  }
}


SEXP codeHaplo2(SEXP M1, SEXP M2) {// 2 x Haplo - Matrix, as vector gespeichert
  if (length(M1) == 0 || length(M1) != length(M2))
    ERR("'M' has length 0 or lengths are unequal.");
  if (!isVector(M1) || !isVector(M2))
    ERR("'M' or 'N' not a vector or of wrong size.");
  Uint snps = (Uint) length(M1);  
  SEXP Code;
  PROTECT(Code = create_codevectorHaplo(snps, 1));
  ToInt(M1);  
  ToInt(M2);  
  codeInnerHaplo2(M1int, M2int, snps, Align(Code, ALIGN_HAPLO));
  if (PRESERVE) R_PreserveObject(Code);
  FREEint(M1);
  FREEint(M2);
  UNPROTECT(1);
  return Code;
}


SEXP GetHaploSequence(Uint snps, Uint indiv,
		      bool indivpercol, bool doubledIndiv,
		      Uint currentBits, // BitsPerCode
		      Uint *blockIncr, Uint *bitIncr,
		      Uint *delta, Uint *indivIncr, bool create){
  SEXP Ans = R_NilValue;
  
  if (indivpercol) {
    *indivIncr = snps * currentBits;
    if (doubledIndiv) {
      *blockIncr = CodesPerBlock; // X
      *delta = currentBits == 1 ? 0L : snps ; // X; pointing to next bit
      *bitIncr = 1L; // X
      //printf("%d %d %d\n", snps, indiv, currentBits);
      if (create) Ans = allocMatrix(INTSXP, snps, indiv*currentBits);
   } else {
      *blockIncr = BitsPerBlock * currentBits / BitsPerCode;
      *delta = currentBits == 1 ? 0L : 1L;
      *bitIncr = currentBits;
      if (create) Ans = allocMatrix(INTSXP, (Ulong) currentBits * snps, indiv);
    }
  } else {
    *blockIncr = BitsPerBlock * indiv * currentBits / BitsPerCode;
    *bitIncr = currentBits * indiv;
    if (doubledIndiv) {
      *delta = currentBits == 1 ? 0L : 1L;
      *indivIncr = currentBits;
      if (create) Ans = allocMatrix(INTSXP, currentBits * indiv, snps);
    } else {
      *delta = currentBits == 1 ? 0L : indiv;
      *indivIncr = 1L;
      if (create) Ans=allocMatrix(INTSXP,  indiv, (Ulong) currentBits * snps);
    }
  }
  return Ans;
}


void codeInnerHaplo(Uint *MM, Uint snps, Uint indiv,
		    bool indivpercol, bool doubledIndiv, Uint *code) {  
  Uint j, blockIncr, bitIncr, delta, indivIncr,
    bits = snps * BitsPerCode,
    fullBlocks = bits / BitsPerBlock;
  Ulong units = Blocks(snps) * UnitsPerBlock;

  GetHaploSequence(snps, indiv, indivpercol, doubledIndiv, BitsPerCode,
		   &blockIncr, &bitIncr, &delta, &indivIncr, false);

#ifdef DO_PARALLEL
#pragma omp parallel for num_threads(CORES)
#endif


  
  for (Uint i=0; i<indiv; i++) {
    Uint *cm = code + i * units,
      *M = MM + i * indivIncr;
    for (j=0; j<fullBlocks; j++) {
      Uint *C = cm + j * UnitsPerBlock,
	*mm = M + j * blockIncr,
	*mm2 = mm + delta;
      for (Uint u=0; u<CodesPerUnit; u++) {
	Uint shift = u * BitsPerCode;
	for (Uint k=0; k<UnitsPerBlock; k++, mm+=bitIncr, mm2 += bitIncr) {
	  assert(*mm == 0 || *mm == 1);
	  assert(*mm2 == 0 || *mm2 == 1);
	  C[k] |= *mm2 << (shift + 1);
	  C[k] |= *mm << shift;	  
	}
      }
    }
    if (fullBlocks * CodesPerBlock < snps) {
      Uint 
	*C = cm  + fullBlocks * UnitsPerBlock,
	*mm = M + fullBlocks * blockIncr,
	*mm2 = mm + delta,
	*end = M + snps * bitIncr;
      for (Uint u=0; u<CodesPerUnit; u++) {
	Uint shift = u * BitsPerCode;
	for (Uint k=0; k<UnitsPerBlock && mm < end;
	     k++, mm+=bitIncr, mm2 += bitIncr) {
	  assert(*mm == 0 || *mm == 1);
	  assert(*mm2 == 0 || *mm2 == 1);
	  C[k] |= *mm2 << (shift + 1);
	  C[k] |= *mm << shift;
	}
	if (mm >= end) break;
      }
    }
  }
}

SEXP codeHaplo(SEXP M, SEXP IndivPerCol, SEXP DoubledIndiv) {// 2 x Haplo - Matrix, as vector gespeichert
  if (length(M) == 0) ERR("'M' has length 0.");
  Uint snps, indiv,
    nrow = (Uint) nrows(M),
    ncol = (Uint) ncols(M);
  bool doubledIndiv = LOGICAL(DoubledIndiv)[0],
    indivpercol = LOGICAL(IndivPerCol)[0];

  if (indivpercol) {
    snps = nrow;
    indiv = ncol;
  } else {
    snps = ncol;
    indiv = nrow;
  }
  if (doubledIndiv) {
    // printf("%d (%d %d) %d \n", indiv, nrow, ncol, BitsPerCode);
    if (indiv % BitsPerCode != 0)
      ERR("information on individuals not doubled");
    indiv /= BitsPerCode;
  } else {
    if (snps % BitsPerCode != 0) ERR("information on haplotype odd");
    snps /= BitsPerCode;
  }

  SEXP Code;
  PROTECT(Code = create_codevectorHaplo(snps, indiv));
  //  printf("? %d\n", TYPEOF(Code));
  ToInt(M);
  //  printf("! %d\n", TYPEOF(Code));
  codeInnerHaplo(Mint, snps, indiv, indivpercol, doubledIndiv,
		 Align(Code, ALIGN_HAPLO));
  //  printf("@\n");
  if (PRESERVE) R_PreserveObject(Code);
  FREEint(M);
  UNPROTECT(1);
  return Code;
}

void InitGetHaplo(SEXP CM, Uint **code, Uint *unitsPerIndiv) {
  assert_haplo();
  Uint 
    *info = GetInfo(CM),
    snps = info[SNPS];
  *unitsPerIndiv = (Ulong) Blocks(snps) * UnitsPerBlock;
  *code = Align(CM, ALIGN_HAPLO);
}

//*cm = code + unitsPerIndiv * i,
Uint rev_haplo_code[4] = {0L, 1L, 1L, 2L};
Uint GetHaplo(Uint *cm, Uint snp) {
  assert(BitsPerCode == 2);
  Uint 
    j = snp / CodesPerBlock,
    remainder = snp % CodesPerBlock,
    *C = cm + j * UnitsPerBlock,     
    u = remainder / UnitsPerBlock,
    k = remainder % UnitsPerBlock;
  return rev_haplo_code[(C[k] >> (u * BitsPerCode)) & CodeMask];
}

Uint GetHaploX(Uint *cm, Uint snp) { // to be deleted
  assert(BitsPerCode == 2);
  Uint 
    j = snp / CodesPerBlock,
    remainder = snp % CodesPerBlock,
    *C = cm + j * UnitsPerBlock,     
    u = remainder / UnitsPerBlock,
    k = remainder % UnitsPerBlock;

  //  printf("H=%d ", (C[k] >> (u * BitsPerCode)) & CodeMask);
  return (C[k] >> (u * BitsPerCode)) & CodeMask;
}


SEXP decodeHaplo(SEXP CM, SEXP Indiv,
		 SEXP Sets, SEXP IndivPerCol, SEXP DoubledIndiv) {
  assert_haplo();
  //printf("%d %d %d\n", length(CM), length(IndivPerCol), length(DoubledIndiv));
  bool IndivGiven = length(Indiv) > 0;
  Uint blockIncr, bitIncr, delta, indivIncr,
    *info = GetInfo(CM),
    currentBits = length(Sets),
    snps = info[SNPS], 
    indiv = info[INDIVIDUALS],
    currentIndiv = IndivGiven ? length(Indiv) : indiv,
    *I = IndivGiven ? (Uint*) INTEGER(Indiv) : NULL,
    bits = snps * BitsPerCode,
    fullBlocks = bits / BitsPerBlock,
    *code = Align(CM, ALIGN_HAPLO);
  Ulong units = Blocks(snps) * UnitsPerBlock;
  bool doubledIndiv = LOGICAL(DoubledIndiv)[0],
    indivpercol = LOGICAL(IndivPerCol)[0];
  if (info[WHAT] != HAPLO) ERR("not a haplo coding");
  //    printf("doubledindiv = %d\n", doubledIndiv);
  if (currentBits != 1 && currentBits != BitsPerCode)
    ERR("The number of chromosome sets can only be 1 or 2.");
  bool set1 = INTEGER(Sets)[0] == 1;
  if (currentBits == 2 && (!set1 || INTEGER(Sets)[1] != 2))
    ERR("The sets must be given in the order 1:2")
  else if (!set1 && INTEGER(Sets)[0] != 2)
    ERR("The value for a chromosome set can be 1 and 2 only.");
  
  SEXP Ans;
  PROTECT(Ans = GetHaploSequence(snps, currentIndiv, indivpercol, doubledIndiv,
				 currentBits, &blockIncr, &bitIncr, &delta,
				 &indivIncr, true));
  
  Uint *MM = (Uint*) INTEGER(Ans);
  Ulong total = (Ulong) currentBits * snps * currentIndiv;
  for (Ulong i=0; i<total; i++) MM[i] = 0; // noetig?
  assert(BitsPerCode == 2);

  assert(snps > 0);
  assert(indiv > 0);
  //printf("currentIndiv = %d\n", currentIndiv);
  
#ifdef DO_PARALLEL
#pragma omp parallel for num_threads(CORES)   
#endif
  for (Uint i=0; i<currentIndiv; i++) {
    Uint *cm = code + units * (IndivGiven ? I[i] - 1 : i),
      *M = MM + i * indivIncr;
#if defined SCHLATHERS_MACHINE
    Uint snp = 0;
#endif    
    for (Uint j=0; j<fullBlocks; j++) {
      Uint
	*C = cm + j * UnitsPerBlock,
	*mm = M + j * blockIncr,
	*mm2 = mm + delta;
      for (Uint u=0; u<CodesPerUnit; u++) {
	Uint shift = 1 << u * BitsPerCode,
	  shiftP1 = shift << 1;
	for (Uint k=0; k<UnitsPerBlock; k++, mm+=bitIncr, mm2 += bitIncr) {
	  // ordering important !!! as mm2 might be overwritten by mm!!
	  *mm2 = (C[k] & shiftP1) > 0;
	  if (set1) *mm = (C[k] & shift) > 0;
#if defined SCHLATHERS_MACHINE
	  assert(delta == 0 || GetHaploX(cm, snp++) == *mm + 2 * *mm2);
#endif    
	}
      }
    }
    if (fullBlocks * CodesPerBlock < snps) {
      Uint
	*C = cm + fullBlocks * UnitsPerBlock,
	*mm = M + fullBlocks * blockIncr,
	*mm2 = mm + delta,
	*end = M + snps * bitIncr;
      for (Uint u=0; u<CodesPerUnit; u++) {
	Uint shift = 1 << u * BitsPerCode,
	  shiftP1 = shift << 1;
	for (Uint k=0; k<UnitsPerBlock && mm < end;
	     k++, mm+=bitIncr, mm2 += bitIncr) {
	  *mm2= (C[k] & shiftP1) > 0;
	  if (set1) *mm = (C[k] & shift) > 0;
	  //	  printf("mm = %u %u %u; set1=%d delta=%d\n", *mm, *mm2, GetHaploX(cm,snp),set1, delta);
	  
	  assert(delta == 0 || GetHaploX(cm, snp++) == *mm  + 2 * *mm2);
	}
	if (mm >= end) break;
      }
    }
  }

  UNPROTECT(1);
  if (PRESERVE) R_PreserveObject(Ans);
  return Ans;
}


void zeroNthHaplo(SEXP CM, SEXP NN) { // 0.45 bei 25000 x 25000
  assert_haplo();
  Uint
    first2bits32 = 0x00000003,
    *N = (Uint*) INTEGER(NN),
    lenN = length(NN),
    *info = GetInfo(CM),
    snps = info[SNPS],
    individuals = info[INDIVIDUALS],
    blocks = Blocks(snps),
    *C0 = Align(CM, ALIGN_HAPLO);
  if (info[WHAT] != HAPLO) ERR("not a genomicmatrix coding");
  
  for (Ulong i=0; i<individuals; i++) {    
    BlockType *C1 = ((BlockType0 *) C0) + i * blocks;
    for (Uint ni=0; ni<lenN; ni++) {
      Uint j=N[ni] / CodesPerBlock,
	jj = N[ni] % CodesPerBlock,
	u = jj / UnitsPerBlock,
	k = jj % UnitsPerBlock;     
      BlockType *C = C1 + j;
      ((BlockUnitType0*) C)->u32[k] &= ~(first2bits32 << (BitsPerCode * u));
    }
  }
}
```
