## Supplementary material for "MoBPS - Modular Breeding Program Simulator": miraculix R-package: Haplo.h

```
#ifndef miraculix_Haplo_H
#define miraculix_Haplo_H 1

Uint *AlignHaplo(SEXP Code, Uint nr, bool test);
//SEXP codeHaplo(SEXP M1, SEXP M2);
Uint CodesPerBlockHaplo();
Uint UnitsPerIndivHaplo(Uint snps);
void InitGetHaplo(SEXP CM, Uint **code, Uint *units);
Uint GetHaplo(Uint *cm, Uint snp);
SEXP codeHaplo2(SEXP M1, SEXP M2);
SEXP create_codevectorHaplo(Uint snps, Uint individuals);
void zeroNthHaplo(SEXP CM, SEXP NN);


#endif
```
