## Supplementary material for "MoBPS - Modular Breeding Program Simulator": miraculix R-package: haplogeno.cc

```
/*
 Authors 
 Martin Schlather,


 Copyright (C) 2018 -- 2019  Martin Schlather

This program is free software; you can redistribute it and/or
modify it under the terms of the GNU General Public License
as published by the Free Software Foundation; either version 3
of the License, or (at your option) any later version.

This program is distributed in the hope that it will be useful,
but WITHOUT ANY WARRANTY; without even the implied warranty of
MERCHANTABILITY or FITNESS FOR A PARTICULAR PURPOSE.  See the
GNU General Public License for more details.

You should have received a copy of the GNU General Public License
along with this program; if not, write to the Free Software
Foundation, Inc., 59 Temple Place - Suite 330, Boston, MA  02111-1307, USA.  
*/

// _MM_ALIGN16 Uint ccc;

#include "AutoMiraculix.h"
#include <General_utils.h>
#include <zzz_RandomFieldsUtils.h>
#include "xport_import.h"
#include "error.h"
#include "Bit23.h"
#include "shuffle.h"
#include "hamming.h"
#include "nocoding.h"
#include "Haplo.h"
#include "options.h"
#include "miraculix.h"


typedef SEXP (*coding_start_t)(Uint , Uint, SEXP);
typedef void (*coding_main_t)(Uint *, Uint, Uint, Uint, Uint, Uint,
			      SEXP, double *);
typedef Uint (*codes_per_block_t)();

char EOL = '\n';


SEXP get_centered() {
  Uint len = GLOBAL.genetics.ncentered ;
  double *centered = GLOBAL.genetics.pcentered;
  assert (centered != NULL && GLOBAL.genetics.centered==Nan);
  SEXP Ans;
  PROTECT(Ans = allocVector(REALSXP, len));
  double *ans = REAL(Ans);
  MEMCOPY(ans, centered, sizeof(double) * len);
  UNPROTECT(1);
  return Ans;
}


			     


void DoCentering(double *ans, Uint individuals, bool centred, bool normalized,
		   Ulong sumGeno) {
  double
    nd = (double) individuals,
    nSq = nd * nd,
    factor = RF_NAN,
    sum_pi_qi = RF_NAN;
  if (centred || normalized) {
    double
      sum_P = RF_NAN,
      *sums = (double *) MALLOC(sizeof(double) * individuals),
      totalsum = 0.0;
    
    Uint k = 0,
      nP1 = individuals + 1;
    for (Uint i=0; i<individuals; i++) {
      double dummy = 0.0;
      for (Uint j=0; j<individuals; j++) dummy += ans[k++];
      sums[i] = nd * dummy;
      totalsum += dummy;
    }

    if (centred) {
      for (Uint i=0; i<individuals; i++) {
	Uint idx = i * nP1;
	for (Uint j=i; j<individuals; j++, idx++) {
	  ans[idx] = nSq * ans[idx] - sums[i] - sums[j] + totalsum;
	}
      }
    }
    
    if (normalized) {
      sum_P = nd *  (double) sumGeno; // ^= sigma^2 * nd^2
      factor = sum_pi_qi = sum_P - 0.5 * totalsum;
      //printf("f = %f %f %f\n", factor, sum_P, totalsum);
        if (!centred) factor /= nSq;
    } else factor = nSq;
    
    //    print("sumpq=%10g  %10g; sumGeno=%10g\n", sum_pi_qi, sum_pi_qi * nSq, sumGeno);
    
    //    print("sigma2 = %10g\n", sum_pi_qi);
    //    printf("factor = %f\n", factor);
    if (factor <= 0.0) ERR("strange input matrix?")
    
    for (Uint i=0; i<individuals; i++) {
      Uint idx = i * nP1;
      for (Uint j=i; j<individuals; j++) {
	ans[i + j * individuals] = (ans[idx++] /= factor);
	// printf("%f %f %d\n", ans[idx-1],ans[i + j * individuals], individuals);
      }
    }
    
    FREE(sums);
  }  
  // return sum_pi_qi /nSq ;
}


#define UPDATE								\
  if (isSNPxInd) {							\
    Uint jj, rP1 = r + 1;						\
    if (rP1 % nrow_matrix == 0) {					\
      /* always delivering matrix as snps x individuals */		\
      coding_main(matrix, 0, individuals, rP1-codesperblock, rP1,	\
		  nrow_matrix,  Ans, dG);				\
      rowidx = -plusrow;						\
      for (jj=0; jj<matrix_size; matrix[jj++] = 0);			\
    }									\
  } else {								\
    Uint jj;								\
    coding_main(matrix, r, r + 1, 0, snps, nrow_matrix, Ans, dG);	\
    rowidx = -plusrow;							\
    for (jj=0; jj<matrix_size; matrix[jj++] = 0);			\
  }


#define CLOSE								\
  if (isSNPxInd) {							\
    Uint remainder = r % nrow_matrix;					\
    if (remainder != 0) {						\
      coding_main(matrix, 0, individuals, (Uint)(r - remainder), r,	\
		  nrow_matrix,  Ans, dG);				\
    }									\
  } /* KEIN else  */							\
									\
  fclose(fp);								\
  FREE(matrix); 
  

SEXP file_binary_intern(char *file, 
			char *coding, Uint header, bool isSNPxInd, 
			bool doubledindividuals,
			// output
			Uint individuals,
			Uint snps,
			Uint codesperblock,
			coding_start_t coding_start,
			coding_main_t coding_main,
			SEXP G
			) {
  // for plinkbinary files

  char ch, 
    //   A = coding[0],
    B = coding[1],
    C = coding[2]
    //  NA = coding[3],
    //  SEP = coding[4]
    ;
  bool
    haplo = B == C;

  if (haplo) ERR("haplo not recognized for binary files");
   
  FILE *fp;
  Uint j, nrow_matrix, ncol_matrix, perbyte,
    *matrix, // snps x individuals
    rows, // rowsM1, 
    cols,  r, l, i, idx,
    plusrow, rowidx, hr, hc, haplorow, haplocol;
  
  if (snps <= 0 || individuals <= 0 || snps <= individuals) {
    char msg[200];
    SPRINTF(msg, "matrix is supposed to be a '%.20s' x '%.20s' matrix with %d individuals and %d SNPs, which looks odd.\n", 
	    isSNPxInd ? "SNPs" : "individuals",
	    isSNPxInd ? "individuals" : "SNPs",
	    individuals, snps);
    if (snps > 0 && individuals > 0) { warn(msg) } else ERR(msg);
  }
  
  if (isSNPxInd) {
    plusrow = 1;
    nrow_matrix = codesperblock; 
    rows = snps;
    cols = ncol_matrix = individuals;
  } else {
    // plusrow = snps;
    plusrow = codesperblock;
    rows = individuals;
    cols = ncol_matrix = snps;
    nrow_matrix = 1;
  }
  if (haplo) {
    if (isSNPxInd xor !doubledindividuals) {
      haplocol = 2;
      haplorow = 1;
    } else {
      haplocol = 1;
      haplorow = 2;    
    } 
  } else {
    haplocol = haplorow = 1;
  }

  SEXP Ans = coding_start(snps, individuals, G);
  Uint matrix_size = nrow_matrix * (ncol_matrix + 3);
  Ulong sumgeno = 0;
  double *dG = NULL;
  if (length(G) > 0) dG = REAL(G);

  if ((matrix = (Uint*) CALLOC(matrix_size, sizeof(Uint))) == NULL)
    ERR("memory space could not be acquired");
  cols = (Uint) CEIL(0.25 * cols);
  perbyte = 4 / haplocol;
  char twobitpattern = 3;

  //   printf("plusrow %d %d , %d %d c=%d %d haplorow/col %d %d\n", plusrow, nrow_matrix, individuals, haplo, cols, isSNPxInd, haplorow, haplocol);  //assert(false);

  if ((fp = fopen(file, "r")) == NULL)  {
    ERR1("file '%.50s' could not be opened", file);
  }
  for (i=0; i<header; i++) fgetc(fp);

  for (rowidx=r=0; r<rows; r++, rowidx+=plusrow) { // number of (multiple) rows 
    for (hr=0; hr<haplorow; hr++) { 
      // genuine loop only if haplo types are given rowwise
      idx = rowidx;

      //     printf("\n"); //assert(r < 10);

      for (l=0; l<cols; l++) { // number of (multiple cols 
	Uint k = 0;
	ch = fgetc(fp);
	for (j = 0; j < perbyte; j++, idx+=nrow_matrix) {
	  for (hc=0; hc<haplocol; hc++, k+=2) {
	    char value = (ch >> k) & twobitpattern;
	    //	  for (hc=0; hc<haplocol; hc++, ch >>= 2) {
	    // char value = ch & twobitpattern;
	    // genuine loop only if haplo types are given column wise
	    //  printf("%d", value==3 ? 2 : value==2 ? 1 : value == 0 ? 0 : value);

	    if (value != 0) { // == 0
	      // print("idx=%d %d (%d %d) %d\n", idx, matrix_size,  nrow_matrix, ncol_matrix, r);
	      assert(idx <= matrix_size);
	      if (value == 2) {
		matrix[idx]++;
		sumgeno++;
	      } else if (value == 3) {
		matrix[idx] += 2;
		sumgeno += 2;
	      } else if (value == 1) ERR("missing value detected.")
	      else {
		ERR8("unknown character detected (>%c<). Note that missings are not allowed here.\n Debugging information: row=%d of %d rows, col=%d of %d columns; plus=%d; current haplocol=%d of %d\n", ch, r, rows, l, cols, nrow_matrix, hc, haplocol);
	      }
	    //assert(matrix[idx] <=2);
	    } // ch != A
	  } //hc
	} // j
      } // cols l
    } // hr

    UPDATE;    

  } // rows r
   
  CLOSE;

  Uint *info = GetInfo(Ans);						
  StoreSumGeno(sumgeno);			       

  return Ans;
}


SEXP file_intern(SEXP file,
		 coding_start_t coding_start,
		 coding_main_t coding_main,
		 Uint codesperblock,
		 SEXP G) {

  Uint *info = GetInfo(file),
    leadingcols = info[LEADINGCOL],
    isSNPxInd = info[SNPxIND],
    snps = info[SNPS],
    individuals = info[INDIVIDUALS],
    doubledindividuals = info[DOUBLEINDIV];
  Rint header = info[HEADER]; // can be indeed negative !!
  info[CODESPERBLOCK] = codesperblock;

  // printf("individuals = %d doubledindiv=%d\n", individuals, doubledindividuals);
 
  char *coding = (char*) CHAR(STRING_ELT(getAttrib(file, Coding), 0));
  char *filename = (char*) CHAR(STRING_ELT(file, 0));

  if (header < 0) {
    // only for plinkbinary: snps & individuals already known
    return file_binary_intern(filename, coding,
			      -header, isSNPxInd, doubledindividuals,
			      individuals,
			      snps, codesperblock, 
		       // output
			      coding_start, coding_main, G
			      );
  } // else snps not known yet:

  // BEAGLE 3 
  // beides: ani x snps und snps x individ
  // A B A B ?
  // ? als voraussetzung nicht vorhanden
  // 1 header-Zeile
  // 2 erste Spalten weg
  //
  // plink: kein header
  // 4 spalten, die nicht brauchen
  char ch, oldch,
    A = coding[0],
    B = coding[1],
    C = coding[2],
    NA = coding[3],
    SEP = coding[4]
    ;
  bool 
    haplo = B == C;

  // printf("%.50s|\n", *coding);  assert(!haplo);
   
  FILE *fp;
  Uint nrow_matrix, ncol_matrix,
    *matrix = NULL, // snps x individuals
    rows, // rowsM1, 
    cols, colsM1,
    plusrow,  haplorow, haplocol;
  Ulong sumgeno = 0;
  if ((fp = fopen(filename, "r")) == NULL) {
    ERR1("file '%.50s' could not be opened", filename);
  }

  // determine number of rows and columns
  for (int i=0; i<header; i++) while (fgetc(fp) != EOL);

  // determine size of matrix:
  rows = 1;
  cols = 0;
  oldch = SEP;
  while ((ch  = fgetc(fp)) != EOL) {  
    if (ch == SEP && oldch != SEP) cols++;
    oldch = ch;
  }
  if (oldch != SEP) cols++;
  cols -= leadingcols;

  while ((ch  = fgetc(fp)) != EOF) { 
    if (ch == EOL) rows++;
    oldch = ch;
  }
  if (oldch != EOL) rows++;
  fclose(fp);

  // printf("isSNPxI=%d %d %d %d\n", isSNPxInd, cols, haplo, doubledindividuals);
  
  if (isSNPxInd) {
    individuals = cols;  
    snps = rows;
  } else {  
    individuals =  rows;
    snps = cols;    
  }
  
  if (haplo) {
    if (doubledindividuals) {
      Uint dummy = individuals / 2; 
      if (dummy * 2 != individuals) ERR("odd number of values (individuals)");
      individuals = dummy;
    } else {
      Uint dummy = snps / 2; 
      if (dummy * 2 != snps) ERR("odd number of values (SNPs)");
      snps = dummy;
    }
  }

  info[INDIVIDUALS] = individuals;
  info[SNPS] = snps;
 
  if (snps <= 0 || individuals <= 0 || snps <= individuals) {
    char msg[200];
    SPRINTF(msg, "matrix is supposed to be a '%.20s' x '%.20s' matrix with %d individuals and %d SNPs, which looks odd.\n", 
	    isSNPxInd ? "SNPs" : "individuals",
	    isSNPxInd ? "individuals" : "SNPs",
	    individuals, snps);
    if (snps > 0 && individuals > 0) { warn(msg) } else ERR(msg);
  }

  colsM1 = cols - 1;
  //rowsM1 = rows - 1;

  // read in table
  
  if (isSNPxInd) {
    plusrow = 1;
    nrow_matrix = codesperblock; 
    ncol_matrix = individuals;
  } else {
    // plusrow = snps;
    plusrow = codesperblock;
    ncol_matrix = snps;
    nrow_matrix = 1;
  }
  if (haplo) {
    if (isSNPxInd xor !doubledindividuals) {
      haplocol = 2;
      haplorow = 1;
    } else {
      haplocol = 1;
      haplorow = 2;    
    } 
  } else {
    haplocol = haplorow = 1;
  }

  // now, code matrix
  SEXP Ans = coding_start( snps, individuals,G);
  Uint matrix_size = nrow_matrix * ncol_matrix;
  if ((matrix = (Uint*) CALLOC(matrix_size, sizeof(Uint))) == NULL)
    ERR("memory space could not be acquired");
   double *dG = NULL;
  if (length(G) > 0) dG = REAL(G);

  rows /= haplorow;
  cols /= haplocol;

  //   printf("plusrow %d %d, %d %d c=%d %d haplorow/col %d %d\n", plusrow, nrow_matrix, individuals, haplo, cols, isSNPxInd, haplorow, haplocol);  //assert(false);
 
  fp = fopen(filename, "r");

  // jump header lines
  for (int i=0; i<header; i++) while ((ch=fgetc(fp)) != EOL);

  Uint r=0;
  for (Uint rowidx=0; r<rows; r++, rowidx+=plusrow) {//# of (multiple) rows
    for (Uint hr=0; hr<haplorow; hr++) { 
      // genuine loop only if haplo types are given rowwise
      Uint idx = rowidx;

      //   printf("0\n"); //assert(r < 10);
      
      // jump unnecessary leading stuff
      while ((ch = fgetc(fp)) == SEP);
      for (Uint l=0; l<leadingcols; l++) {
	while ((ch=fgetc(fp)) != SEP && ch!=EOF);
	if (ch == EOF) ERR("unexpected end of file");
	while ((ch = fgetc(fp)) == SEP);
      }

      for (Uint l=0; l<cols; l++, idx+=nrow_matrix) { // # of (multiple cols 
	for (Uint hc=0; hc<haplocol; hc++) {
	  // genuine loop only if haplo types are given column wise
	  
	  if (ch != A) { // == 0
	    //  if (idx>=nrow_matrix * ncol_matrix) 
	    // print("idx=%d %d, nrow_matrix=%d\n", idx, nrow_matrix * ncol_matrix, nrow_matrix);
	    if (ch == B) {
	      matrix[idx]++;
	      sumgeno++;
	    } else if (ch == C) {
	      matrix[idx] += 2;
	      sumgeno +=2;
	    } else if (ch == NA) ERR("missing value detected.")
	    else {
	      PRINTF(">%c< row=%d of %d rows, col=%d of %d columns; plus=%d hc=%d haplocol=%d\n", ch, r, rows, l, cols, nrow_matrix, hc, haplocol);
	      ERR("Unknown character detected. Note that missings are not allowed here.");
	    }
	    //assert(matrix[idx] <=2);
	  } 
	  if (l<colsM1 || hc!=haplocol-1)
	    while((ch = fgetc(fp)) == SEP); // printf("%c", ch);
	}
	
	//print("%d", matrix[idx]);
      } // cols l
      
      // search for end of line
      if (ch != EOL && ch != EOF) while ((ch=fgetc(fp)) != EOL && ch!=EOF);
    } // hr
   
    UPDATE;
    
    // printf("\n");
  } // rows r
  
  CLOSE;

  StoreSumGeno(sumgeno);			       

  return Ans;
}


Ulong sumGeno(Uint * SNPxIndiv, Uint snps, Uint individuals, snpcoding method) {
  switch (method) {
  case Shuffle : return sumGenoShuffle(SNPxIndiv, snps, individuals);
  case Shuffle256 : return sumGenoShuffle256(SNPxIndiv, snps, individuals);
  case TwoBit : return sumGeno2(SNPxIndiv, snps, individuals);
  case ThreeBit : return sumGeno3(SNPxIndiv, snps, individuals);
    
  case Hamming2 : 
  case Hamming3 : return sumGenoH(SNPxIndiv, snps, individuals, method);
    
  case NoSNPcoding:
  case NoSNPcodingR: return sumGenoPlain(SNPxIndiv, snps, individuals); 

  case AutoCoding :
  case Haplo : BUG;
      
  default : BUG;
  }

  return 0L;
}

 
  
Uint* DoAlign(SEXP CM, Uint nr, snpcoding method, bool test) {
  switch(method) {
  case Shuffle : return AlignShuffle(CM, nr, test);    
  case Shuffle256 : return AlignShuffle256(CM, nr, test);    
  case TwoBit : return Align2(CM, nr, test);
  case ThreeBit : return Align3(CM, nr, test);
  case Hamming2 :
  case Hamming3 : return AlignH(CM, nr, test);
  case NoSNPcoding:
  case NoSNPcodingR:  return AlignPlain(CM, nr, test);
  case Haplo : return AlignHaplo(CM, nr, test);    
  case AutoCoding : BUG;     
  default : ERR("Method is unknown!");
  }
  return NULL;
}

//Uint* DoAlignWithoutTest(SEXP CM, Uint nr, snpcoding method) {
//  // very dangerous command !!!
//  DoAlign(CM, nr, method, false);
//}

Uint* DoAlign(SEXP CM, Uint nr, snpcoding method) {
  return DoAlign(CM, nr, method, true);
}

void matrix_mult(Uint * SNPxIndiv, Uint snps, Uint individuals,
		 snpcoding method,  bool centred, bool normalized,
		 Ulong SumGeno, double *A) {
  assert(individuals > 0);
  double nSq = (double) individuals * (double) individuals;      
  if ((double) snps * nSq * 4.0 > 4.5035996e+15) // 2^{manissen-bits = 52}
    ERR("matrix too large to calculate the relationship matrix -- pls contact maintainer of 'miraculix'");
  switch (method) {
  case Shuffle : matrixshuffle_mult(SNPxIndiv, snps, individuals, A); break;
  case Shuffle256 : matrixshuffle256_mult(SNPxIndiv, snps, individuals, A); break;
  case TwoBit : matrix2_mult(SNPxIndiv, snps, individuals, A); break;
  case ThreeBit : matrix3_mult(SNPxIndiv, snps, individuals, A); break;
    
  case Hamming2 :
  case Hamming3 : matrixH_mult(SNPxIndiv, snps, individuals, method, A); break;
    
  case NoSNPcoding:
  case NoSNPcodingR:  matrixPlain_mult(SNPxIndiv, snps, individuals, A); break;
    
  case AutoCoding :
  case Haplo : BUG;
      
  default : BUG;
  }

  //  printf("\n\nmulti: \n"); for (int i=0; i<10; i++) printf("%f ", A[i]);
  

  assert(A[0] >= 0.0);
  if (centred || normalized) {
    //printf("\nSG=%ld %d\n", SumGeno, method);
    if (SumGeno == 0) // falls die Matrix tatsaechlich nur aus 0-en besteht,
      // wird eben doppelt gerechnet. Praktisch irrelevanter Fall
      SumGeno = sumGeno(SNPxIndiv, snps, individuals, method);
    //    printf("\nsumgeno = %lu %d %d %d \n", SumGeno, individuals ,centred, normalized);
    
   assert(individuals > 0);
   DoCentering(A, individuals, centred, normalized, SumGeno);
   assert(A[0] >= 0.0);
  }
  
  //printf("\n\nmultXXi: \n"); for (int i=0; i<10; i++) printf("%f ", A[i]);
}


double *matrix_mult(Uint * SNPxIndiv, Uint snps, Uint individuals,
		    snpcoding method,  bool centred, bool normalized,
		    Ulong SumGeno) {
  double *A = (double *) MALLOC(individuals * individuals * sizeof(double));
  if (A == NULL) ERR("mem allocation");

  // printf("method = %s\zn", SNPCODING_NAME[method]);
  
  matrix_mult(SNPxIndiv, snps, individuals, method, centred, normalized,
	      SumGeno, A);

  //  for (int i=0; i<10; i++) printf(">>%f ", A[i]);

 
  return A;
}
 

SEXP matrix_mult(SEXP SNPxIndiv) {
 snpcoding method = (snpcoding) MethodOf(SNPxIndiv);
   Uint
    *info = GetInfo(SNPxIndiv),
    individuals = info[INDIVIDUALS],
     snps = info[SNPS],
    *code = DoAlign(SNPxIndiv, ALIGN_RELATION, method);
  if (info[WHAT] != GENOMATRIX) ERR("not a coded Z matrix");
  
  SEXP Ans; 
  PROTECT(Ans = allocMatrix(REALSXP, individuals, individuals));
  matrix_mult(code, snps, individuals, method,
	      GLOBAL.genetics.centered == True,
	      GLOBAL.genetics.normalized == True,
	      SumGeno(info), REAL(Ans));
  
 UNPROTECT(1);
  if (PRESERVE) R_PreserveObject(Ans);
  return Ans;
}
    

int GetCodesPerBlock(snpcoding method) {
  switch (method) {
  case Shuffle : return CodesPerBlockShuffle();
  case Shuffle256 :  return CodesPerBlockShuffle256();
  case TwoBit : return CodesPerBlock2();
  case ThreeBit : return CodesPerBlock3();    
  case Hamming2 : case Hamming3 : return CodesPerBlockH();
  case NoSNPcoding: case NoSNPcodingR: return CodesPerBlockPlain();
  case AutoCoding : BUG;
  case Haplo : return CodesPerBlockHaplo();
  default : BUG;
  }
}

SEXP file_get(SEXP file) {
  switch (GLOBAL.genetics.method) {
  case Shuffle :  return file_intern(file, matrix_start_shuffle, matrix_shuffle,
				     CodesPerBlockShuffle(), R_NilValue);

  case Shuffle256 :  return file_intern(file, matrix_start_shuffle256,
					matrix_shuffle256,
					CodesPerBlockShuffle256(), R_NilValue);

  case TwoBit : return file_intern(file, matrix_start2, matrix_coding2,
				   CodesPerBlock2(), R_NilValue);
    
  case ThreeBit : return file_intern(file, matrix_start3, matrix_coding3,
				     CodesPerBlock3(), R_NilValue);
    
   case Hamming2 : return file_intern(file, matrix_startH2, matrixH2,
				     CodesPerBlockH(), R_NilValue);
    
  case Hamming3 : return file_intern(file, matrix_startH3, matrixH3,
				     CodesPerBlockH(), R_NilValue);

  case NoSNPcoding: 
  case NoSNPcodingR: return file_intern(file, matrix_start_plain, matrix_plain,
					CodesPerBlockPlain(), R_NilValue);

  case AutoCoding :// in R abgesichert
  case Haplo : BUG;    
    
  default : BUG;
    
  }
}


SEXP zeroNthGeno(SEXP SNPxIndiv, SEXP Snps) {
  Uint *info = GetInfo(SNPxIndiv),
    snps = info[SNPS],
    *which = (Uint*) INTEGER(Snps),
    len = length(Snps);  
  for (Uint i=0; i<len; i++) {
    if (which[i] >= snps) ERR("values of 'Snps' out of range.");
  }
    
  snpcoding method = (snpcoding)  MethodOf(SNPxIndiv);
  switch(method) {
  case Shuffle : zeroNthGenoShuffle(SNPxIndiv, Snps); break;
  case Shuffle256 : zeroNthGenoShuffle256(SNPxIndiv, Snps); break;
  case TwoBit :  zeroNthGeno2(SNPxIndiv, Snps);break;
  case ThreeBit : zeroNthGeno3(SNPxIndiv, Snps);break;
  case Hamming2 :
  case Hamming3 :  zeroNthGenoH(SNPxIndiv, Snps, method);break;
  case Haplo : zeroNthHaplo(SNPxIndiv, Snps);break;
    
  case NoSNPcoding: //return zeroNthGeno(SNPxIndiv, Snps);
  case NoSNPcodingR: 
  case AutoCoding :  BUG;
  default :  BUG;
  }

  return SNPxIndiv;
}


SEXP allele_freq(SEXP SNPxIndiv) {
  snpcoding method = (snpcoding) MethodOf(SNPxIndiv);
  switch (method) {
  case Shuffle : return allele_freqShuffle(SNPxIndiv);
  case Shuffle256 : return allele_freqShuffle256(SNPxIndiv);
    
  case TwoBit :return allele_freq2(SNPxIndiv);
  case ThreeBit : return allele_freq3(SNPxIndiv);
    
  case Hamming2 :
  case Hamming3 :return allele_freqH(SNPxIndiv);

  case Haplo : ERR("decoding of partial matrix not programmed yet"); 
    
  case NoSNPcoding:  //    return allele_freqPlain(SNPxIndiv); 
  case NoSNPcodingR:
  case AutoCoding : BUG;
      
  default : BUG;
  }
}


SEXP get_matrix_N(SEXP SNPxIndiv, SEXP Snps) {
  snpcoding method = (snpcoding) MethodOf(SNPxIndiv);
  switch (method) {
  case Shuffle : return get_matrixN_shuffle(SNPxIndiv, Snps);
  case Shuffle256 : return get_matrixN_shuffle256(SNPxIndiv, Snps);
    
  case TwoBit :return get_matrixN_2(SNPxIndiv, Snps);
  case ThreeBit : return get_matrixN_3(SNPxIndiv, Snps);
    
  case Hamming2 :
  case Hamming3 :return get_matrixN_H(SNPxIndiv, Snps);
    
  case Haplo : ERR("decoding of partial matrix not programmed yet"); 


  case NoSNPcoding:
  case NoSNPcodingR: // return get_matrixplain(SNPxIndiv, Snps);
  case AutoCoding :
  BUG;
      
  default : BUG;
  }
}


SEXP matrix_get(SEXP SNPxIndiv) {
  snpcoding method = (snpcoding) MethodOf(SNPxIndiv);
  //printf("snpcoding=%d\n", method);   BUG;
 switch (method) {
  case Shuffle : return get_matrixshuffle(SNPxIndiv);
  case Shuffle256 : return get_matrixshuffle256(SNPxIndiv);
    
  case TwoBit :return get_matrix2(SNPxIndiv);
  case ThreeBit : return get_matrix3(SNPxIndiv);
    
  case Hamming2 :
  case Hamming3 :return get_matrixH(SNPxIndiv);
    
  case NoSNPcoding:
  case NoSNPcodingR:  return get_matrixPlain(SNPxIndiv);
    
  case AutoCoding :
  case Haplo : BUG;
      
  default : BUG;
  }
}


SEXP matrix_coding(SEXP M) {
  if (PRESERVE) ERR("if 'dolocking' then implicite coding is impossible.");
  snpcoding method = (snpcoding) GLOBAL.genetics.method;
  if (length(M) == 0) ERR("'M' has length 0.");
  SEXP Code;
  Uint
    snps,
    indiv;
  if (isMatrix(M)) {
    snps = nrows(M);
    indiv = ncols(M);
  } else {
    snps = length(M);
    indiv = 1;
  }  
  
  ToInt(M);
  Ulong total = (Ulong) snps * indiv;
  for (Ulong i=0; i<total; i++) {
    if (Mint[i] > 2) ERR("SNP matrix has only the values 0,1,2");
  }
    
  switch (method) {
  case Shuffle : PROTECT(Code =  matrix_coding_shuffle(Mint, snps, indiv));
    break;
  case Shuffle256 : PROTECT(Code =  matrix_coding_shuffle256(Mint, snps,indiv));
    break;

  case TwoBit :
  case ThreeBit : PROTECT(Code = matrix_coding23(Mint, snps, indiv, method));
    break;
 
  case Hamming2 :
  case Hamming3 : PROTECT(Code = matrix_codingH(Mint, snps, indiv, method));
    break;

  case NoSNPcoding:
  case NoSNPcodingR: PROTECT(Code =  matrix_coding_plain(Mint, snps, indiv));
    break;

  case AutoCoding : // in R abgesichert
  case Haplo : BUG;
     
  default : BUG;
  }

   Ulong sumgeno = 0,
    endfor = (Ulong) snps * indiv;
  for (Ulong i=0; i<endfor; i++) sumgeno += (Ulong) Mint[i];

  Uint *info = GetInfo(Code);
  StoreSumGeno(sumgeno);


  FREEint(M);
  UNPROTECT(1);
  
  if (PRESERVE) R_PreserveObject(Code);
  return Code;
}


SEXP createSNPmatrix(Uint snps, Uint individuals, snpcoding method) {  
  SEXP Code;

  //printf("create snps=%d %d\n", snps, individuals);
  switch(method) {
  case Shuffle : PROTECT(Code = matrix_start_shuffle(snps, individuals,
						    R_NilValue));
    break;
  case Shuffle256 : PROTECT(Code = matrix_start_shuffle256(snps, individuals,
							   R_NilValue));
    break;

  case TwoBit : PROTECT(Code = matrix_start2(snps, individuals, 
						    R_NilValue));
    break;
    
  case ThreeBit : PROTECT(Code = matrix_start3(snps, individuals, 
						      R_NilValue));
    break;
 
  case Hamming2 :
  case Hamming3 : PROTECT(Code = matrix_startH(snps, individuals, method,
					       R_NilValue));
    break;

  case NoSNPcoding:
  case NoSNPcodingR:  PROTECT(Code = matrix_start_plain(snps, individuals,
							R_NilValue));
    break;
    
  case Haplo : PROTECT(Code = create_codevectorHaplo(snps, individuals));
    break;
 
  case AutoCoding :  BUG;
     
  default :  BUG;
  }

  if (PRESERVE) R_PreserveObject(Code);
  UNPROTECT(1);
  return Code;
}


SEXP createSNPmatrix(SEXP SNPs, SEXP Individuals) {
  Uint 
    n = Int0(Individuals),
    snps = Int0(SNPs);
  snpcoding method = (snpcoding) GLOBAL.genetics.method;
  if (method == AutoCoding) method = getAutoCodingIntern();
  return createSNPmatrix(snps, n, method);
}


SEXP fillSNPmatrix(SEXP Z, SEXP Idx, SEXP V) {
  ToInt(Idx);
  Uint upi,
    *infoZ = GetInfo(Z),
    *infoV = GetInfo(V),
    snpsZ = infoZ[SNPS],
    snpsV = infoV[SNPS],
    indiv = infoZ[INDIVIDUALS],
    methZ = MethodOf(Z),
    methV = MethodOf(V),
    *z = DoAlign(Z, ALIGN_RELATION, (snpcoding) methZ),
    *v = DoAlign(V, ALIGN_HAPLO, (snpcoding) methZ),
     len = length(Idx);
  if (infoZ[WHAT] != GENOMATRIX) ERR("not a geno matrix that is filled");
  if (snpsZ != snpsV) ERR("number of snps differ.");
  if (methZ != methV) ERR("methods of 'SNPxIndiv' and 'values' differ.");


  switch(methZ) {
  case Shuffle : upi = UnitsPerIndivShuffle(snpsZ); break;
  case Shuffle256 : upi = UnitsPerIndivShuffle256(snpsZ); break;
  case TwoBit : upi = UnitsPerIndiv2(snpsZ); break;
  case ThreeBit : upi = UnitsPerIndiv3(snpsZ); break;
  case Hamming2 :
  case Hamming3 : upi = (Uint) UnitsPerIndivH(snpsZ, methZ); break;
  case NoSNPcoding : upi = UnitsPerIndivPlain(snpsZ); break;
  case Haplo : upi = UnitsPerIndivHaplo(snpsZ); break;
  default : BUG;    
  }
  
  Ulong
    units = upi,
    bytes = BytesPerUnit * units;
  
  for (Ulong i=0; i<len; i++, v += units) {
    Uint idx = Idxint[i] - 1;
    if (idx >= indiv) ERR("Idx out of bound");
    MEMCOPY(z + idx * units, v, bytes);
  }
  FREEint(Idx);
  return R_NilValue;
}


SEXP copyGeno(SEXP CM) {
   Uint
    *info = GetInfo(CM),
    snps = info[SNPS],
    individuals = info[INDIVIDUALS],
    what = info[WHAT];
  
  if (what != GENOMATRIX) ERR("not a geno matrix");
  snpcoding method = (snpcoding) MethodOf(CM);
   
  //  printf("entering copy %d %d %d\n", snps, individuals, method);

  SEXP Code;
  PROTECT(Code = createSNPmatrix(snps, individuals, method));

  //  printf("created\n");
  
  Uint
    *from = DoAlign(CM, ALIGN_HAPLO, method),
    *to = DoAlign(Code, ALIGN_HAPLO, method),
    *info2 = GetInfo(Code),
    *addr1 = (Uint*) GetAddress(info, ALIGNADDR0),
    *addr2 = (Uint*) GetAddress(info2, ALIGNADDR0);
  Ulong mem = MemInUnits(info);

  assert(INFO_LAST== 21 && HEADER == 15); // minicheck ob sich die liste
  // geaendert hat, ohne hier zu korriegieren.
  info2[HEADER] = info[HEADER];
  info2[DOUBLEINDIV] = info[DOUBLEINDIV];
  info2[LEADINGCOL] = info[LEADINGCOL];
 
    
  //printf("lag1 = %ld %ld %lu %lu\n", lag1, lag2, mem, BytesPerUnit);

  if (mem != MemInUnits(info2)) BUG;

  intptr_t
    lag1 = (intptr_t) addr1 - (intptr_t) CM,
    lag2 = (intptr_t) addr2 - (intptr_t) Code;
  if (lag1 > 100 || lag1 < 0 || lag2 > 100 || lag2 < 0)
    ERR("Alignment problem -- pls contact maintainer.");
  //  if (lag1 - lag2 ) ERR("Memory was disclocated. Copying is not possible.");
  if (from != addr1 || to != addr2)
    ERR("Alignment problems. Pls contact maintainer.");
  
  MEMCOPY(to, from, mem * BytesPerUnit);
  //printf("leaving copy\n");
  UNPROTECT(1);
  return Code;
}


void ReUseAs(SEXP Code, snpcoding method) {
  if (method == Shuffle) ReUseAsShuffle(Code);
  else if (method == Shuffle256) ReUseAsShuffle256(Code);
  else if (method == TwoBit) ReUseAsTwoBit(Code);
  else BUG;  
}


Uint haplo2geno(Uint *H, Uint snps, Uint individuals,
		snpcoding method, Uint unitsPerIndivH, Uint *A) {
  //printf("******************** snps=%d %d %d\n", snps, individuals, method);
  switch (method) {
  case Shuffle :
    if (GetCodesPerBlock(Haplo) != GetCodesPerBlock(method))
      ERR("'Shuffle' is wrong. Choose 'Shuffle256' instead.");
    haplo2genoShuffle(H, snps, individuals, A);
    break;
    
  case Shuffle256 :
    if (GetCodesPerBlock(Haplo) != GetCodesPerBlock(method))
      ERR("'Shuffle256' is wrong. Choose 'Shuffle' instead.");
    haplo2genoShuffle256(H, snps, individuals, A);
    break;
    
  case TwoBit : haplo2geno2(H, snps, individuals, unitsPerIndivH, A);
    break;
    
  case ThreeBit :
    if (H == A) goto  ErrorHandling;
    haplo2geno3(H, snps, individuals, unitsPerIndivH, A);
    break;
    
  case Hamming2 :
  case Hamming3 :
    if (H == A) goto  ErrorHandling;
    haplo2genoH(H, snps, individuals, unitsPerIndivH, method, A);
    break;
    
  case NoSNPcoding:
  case NoSNPcodingR: 
    if (H == A) goto  ErrorHandling;
    haplo2genoPlain(H, snps, individuals, unitsPerIndivH, A);
    break;
    
  case AutoCoding :
  case Haplo : BUG;
      
  default : BUG;
  }

  return sumGeno(A, snps, individuals, method);
  
 ErrorHandling: 
  ERR("A memory effecient transformation of a haplotype matrix to a genomic matrix, e.g. from within 'MoBPS', relies on a two-bit-coding assuming for genomic matrices. So, only 'Shuffle', 'Shuffle256' and 'TwoBit' are allowed snp coding methods");
  return 0; // avoids compiler warning
}


SEXP haplo2geno(SEXP H) {
  snpcoding method = (snpcoding) GLOBAL.genetics.method;
  Uint
    *infoH = GetInfo(H),
    individuals = (Uint) infoH[INDIVIDUALS],
    snps =  (Uint) infoH[SNPS];
  if (infoH[WHAT] != HAPLO) ERR("not a haplo coding");

  SEXP Code;
  PROTECT(Code = createSNPmatrix(snps, individuals, method));
  Uint *haplo, unitsPerIndivH,
    *UintCode = (Uint*) INTEGER(Code),
    *info = GetInfo(Code),
    bytesperblock = info[BYTESPERBLOCK],
    *code = (Uint*) algn_general(UintCode, bytesperblock);

  // printf("H1\n");
  InitGetHaplo(H, &haplo, &unitsPerIndivH);
  //  printf("H2\n");
  
  Ulong sumgeno = haplo2geno(haplo, snps, // no PROTECT( neeeded
			     individuals, method, unitsPerIndivH, code);
  //  printf("H3\n");
  StoreSumGeno(sumgeno);
  //  printf("H4\n");
  
  UNPROTECT(1);
  if (PRESERVE) R_PreserveObject(Code);
  return Code;  
}

#define DotBlockSize 100 // arbitrary number


SEXP file_dot_start(Uint snps,Uint individuals,  SEXP G) {
  SEXP MTdot;
  if ((Uint) length(G) !=  individuals)
    ERR("vector must have length equal to number of individuals");
  if (PL > 1) {PRINTF("Data: %d individuals and %d SNPs\n", individuals, snps);}
  PROTECT(MTdot=allocVector(REALSXP, snps));
  for (Uint i=0; i<snps; REAL(MTdot)[i++] = 0.0);
  UNPROTECT(1);
  return MTdot;
}

void file_dot_do(Uint *M, Uint start_individual, Uint end_individual, 
		 Uint start_snp, Uint end_snp, Uint Mnrow,
		 SEXP Ans, double *G) {
  double *Fdot = REAL(Ans);
  for (Uint a=start_individual; a<end_individual; a++) {
    Uint *pM = M + (a - start_individual) * Mnrow;
    double dummy = G[a];
    for (Uint s=start_snp; s<end_snp; pM++) {
      Fdot[s++] += dummy * (double) *pM;
    }
  }
}

SEXP file_dot(SEXP file, SEXP G) {
  return file_intern(file, file_dot_start, file_dot_do,
			    DotBlockSize, G);
}


SEXP  dot_file_start(Uint snps, Uint individuals, SEXP G) {
  SEXP Ans;
  if ((Uint) length(G) != snps) ERR("vector must have length equal to number of snps");
  PROTECT(Ans=allocVector(REALSXP, individuals));
  for (Uint i=0; i<individuals; REAL(Ans)[i++] = 0.0);
  UNPROTECT(1);
  return Ans;
}


void dot_file_do(Uint *M, Uint start_individual, Uint end_individual, 
		 Uint start_snp, Uint end_snp, Uint Mnrow,
		 SEXP Ans, double *G) {
  double *dotF = REAL(Ans);
  for (Uint a=start_individual; a<end_individual; a++) {
    Uint *pM = M + (a - start_individual) * Mnrow;
    double dummy = 0;
    for (Uint s=start_snp; s<end_snp; pM++) {
      dummy += G[s++] * (double) *pM;
    }
    dotF[a] = dummy;
  }
}


SEXP dot_file(SEXP file, SEXP G) {
  return file_intern(file, dot_file_start, dot_file_do,
			    DotBlockSize, G);
}


SEXP substract_centered(SEXP SnpXindiv) {
  Uint indiv = ncols(SnpXindiv),
    len = GLOBAL.genetics.ncentered;
  Ulong snps = nrows(SnpXindiv);
  SEXP Ans;
  PROTECT(Ans = allocMatrix(REALSXP, snps, indiv));
  double *ans = REAL(Ans),
    *snpXindiv = REAL(SnpXindiv),
    *centered = GLOBAL.genetics.pcentered;
  assert(centered != NULL && GLOBAL.genetics.centered==Nan);
  if (snps != len) ERR("length of 'centered' must equal the number of SNPs.");

#ifdef DO_PARALLEL
#pragma omp parallel for num_threads(CORES)
#endif
  for (Uint i=0; i<indiv; i++) {
    Ulong i_snps = i * snps;
    double *s = snpXindiv + i_snps,
      *a = ans + i_snps;
    for (Uint j=0; j<snps; j++) a[j] = s[j] - centered[j];
  }
  UNPROTECT(1);
  return Ans;
}


bool do_warn_change = true, do_warn_changeOK = true;

Uint *Memory[ALIGN_LAST + 1] = { NULL },
  nMem[ALIGN_LAST + 1] = { 0 };

Uint *AlignBase(SEXP CM, Uint nr, Uint bytesperblock, bool test) {
  Uint *info = GetInfo(CM);

  //  printf("info=%d BPC=%d  bpb=%d meth=%d\n", info[BYTESPERBLOCK], info[BITSPERCODE], bytesperblock, MethodOf(CM));
  
  if (test && info[BYTESPERBLOCK] != bytesperblock) {
    // printf("info=%d BPC=%d  bpb=%d meth=%d\n", info[BYTESPERBLOCK], info[BITSPERCODE], bytesperblock, MethodOf(CM));
    // int *i;  printf("i=%d\n", i[9]);
    ERR("currently, data exchange between different kinds of machines (AVX2/SSE) is not possible");
  }
  Uint *address = (Uint *) INTEGER(CM),
    *algnaddress = (Uint *) algn_general(INTEGER(CM), bytesperblock),
    *infoaddress = (Uint*) GetAddress(info, ADDR0);
  
  if (infoaddress == address) {
    assert((uintptr_t) algnaddress % bytesperblock == 0);
    return algnaddress;
  } else if ((uintptr_t) infoaddress % BytesPerUnit ==
	     (uintptr_t) address % BytesPerUnit) {
#ifdef SCHLATHERS_MACHINE
    if (do_warn_changeOK) {
      PRINTF("Address has changed in a coded object (%d) by 'gc' or disk saving. But luckily got same modulus (%lu %lu)).\n", info[WHAT], (uintptr_t) infoaddress % (1024 * 1024), (uintptr_t) address % (1024 * 1024));
      do_warn_changeOK = debugging;
    }
#endif    
    assert((uintptr_t) algnaddress % bytesperblock == 0);
    return algnaddress;
  }
  
#ifdef SCHLATHERS_MACHINE
  if (do_warn_change) {
    PRINTF("Address has changed in a coded object (%d) by 'gc' or disk saving\n", info[WHAT]);
    do_warn_change = debugging;
  }
#endif    
  
  if (PRESERVE) ERR("severe error: has 'dolocking' be called in meanwhile? If 'dolocking' has not been called at all or only at the very beginning, please contaact maintainer.");

#ifdef DO_PARALLEL
  if (CORES > 1)
    ERR("Coded SNP matrix has been stored elsewhere or has been internally moved by R; in these cases set 'RFoptions(cores=1)'");
#endif  

  if ((uintptr_t) infoaddress % bytesperblock ==
      (uintptr_t) address % bytesperblock)
    return algnaddress;
  Ulong mem = MemInUnits(info),
    alignedmemU = AlignedInUnits(info);
  assert(mem <= alignedmemU);
  assert(alignedmemU == (Uint) length(CM));
  if (nMem[nr] < alignedmemU) {
    FREE(Memory[nr]);
    nMem[nr] = alignedmemU;
    Memory[nr] = (Uint*) CALLOC(nMem[nr], BytesPerUnit);
  }
  Uint *algnMem = (Uint *) algn_general(Memory[nr], bytesperblock);
  assert((uintptr_t) algnMem % bytesperblock == 0);
  
  MEMCOPY(algnMem, algnaddress, mem * BytesPerUnit);
  return algnMem;
}


SEXP dolocking(SEXP Do) {
  if (Do != R_NilValue) {
    PRESERVE = LOGICAL(Do)[0];
    if (PRESERVE) BUG;
    for (Uint i=0; i<= ALIGN_LAST; i++) {
      FREE(Memory[i]);
      nMem[i] = 0;
    }
  }

  SEXP Ans;
  PROTECT(Ans = allocVector(LGLSXP, 1));
  LOGICAL(Ans)[0] = PRESERVE;
  UNPROTECT(1);
  return Ans;
}

SEXP unlock(SEXP C) {
  BUG;
  //if (shuffleNotInit) InitShuffle();
  Uint 
    *info = GetInfo(C);
  if ((SEXP) info == R_NilValue) ERR("not a coded object");
  if (PRESERVE) {
#ifdef SCHLATHERS_MACHINE
   PRINTF("unlocking a %.50s at %lu\n", WHAT_NAMES[info[WHAT]],
	  (uintptr_t) INTEGER(C));
#endif  
  } else warning("'unlock' is called although 'dolocking()' has not been called");
  R_ReleaseObject(C);
  return R_NilValue;
}


SEXP Debug() { debugging = true; return R_NilValue; }
SEXP StopDebug() { debugging = false;  return R_NilValue; }


SEXP getAutoCoding() {
  SEXP dummy;
  PROTECT(dummy=allocVector(INTSXP, 1));
  INTEGER(dummy)[0] = GLOBAL.genetics.method = getAutoCodingIntern();
  switch(GLOBAL.genetics.method) {
  case Shuffle : InitShuffle(); break;
  case Shuffle256 : InitShuffle256(); break;
  case TwoBit : Init2(); break;
  case ThreeBit : Init3(); break;
  default: BUG;
  }    
  UNPROTECT(1);
  return dummy;
}


SEXP vectorGeno(SEXP V, SEXP Z) {
  Uint
    *info = GetInfo(Z),
    individuals = info[INDIVIDUALS];
  SEXP Ans;

  PROTECT(Ans = allocVector(REALSXP, individuals));
  double *ans = REAL(Ans);
  for (Uint i=0, endfor= individuals; i<endfor; i++) ans[i] = 0.0; // noetig?

  switch(GLOBAL.genetics.method) {
  case Shuffle : vectorGenoShuffle(V, Z, ans); break;
  case Shuffle256 : vectorGenoShuffle256(V, Z, ans); break;
  default:
    UNPROTECT(1);
    BUG;
  }
  
  UNPROTECT(1);
  if (PRESERVE) R_PreserveObject(Ans);
  return(Ans);
}

SEXP genoVector(SEXP Z, SEXP V) {
   Uint
    *info = GetInfo(Z),
    snps = info[SNPS];
  SEXP Ans;
 
  PROTECT(Ans = allocVector(REALSXP, snps));
  double *ans = REAL(Ans);
 
  switch(GLOBAL.genetics.method) {
  case Shuffle :  genoVectorShuffle(Z, V, ans); break;
  case Shuffle256 : genoVectorShuffle256(Z, V, ans); break;
  default:
    UNPROTECT(1);
    BUG;
  }
  
  UNPROTECT(1);
  if (PRESERVE) R_PreserveObject(Ans);
  return(Ans);
}
```
