## Supplementary material for "MoBPS - Modular Breeding Program Simulator": miraculix R-package: haplogeno.h

```
/*
 Authors 
 Martin Schlather,


 Copyright (C) 2016 -- 2019 Martin Schlather

This program is free software; you can redistribute it and/or
modify it under the terms of the GNU General Public License
as published by the Free Software Foundation; either version 3
of the License, or (at your option) any later version.

This program is distributed in the hope that it will be useful,
but WITHOUT ANY WARRANTY; without even the implied warranty of
MERCHANTABILITY or FITNESS FOR A PARTICULAR PURPOSE.  See the
GNU General Public License for more details.

You should have received a copy of the GNU General Public License
along with this program; if not, write to the Free Software
Foundation, Inc., 59 Temple Place - Suite 330, Boston, MA  02111-1307, USA.  
*/


#ifndef miraculix_haplogeno_H
#define miraculix_haplogeno_H 1

#include <inttypes.h>
#include "miraculix.h"
#include "AutoMiraculix.h"
// #include "IntrinsicsBase.h" --- darf nicht rein !!!
#include "intrinsics.h"


#define FROMINPUT  *(pX++)
#define FROMHAPLO GetHaplo(pX, s)


#define INLINER								\
  Uint static inline *algn(int *X) {assert(algn_general(X, BytesPerBlock)>=(uintptr_t)X); return (Uint *) algn_general(X, BytesPerBlock); } \
									\
  Uint static inline Blocks(Uint X) { return 1L + (X - 1L) / CodesPerBlock; } 

#define Units(snps) (Blocks(snps) * UnitsPerBlock)

//  (Ulong) Blocks(S) * I * UnitsPerBlock;


#define ALL_INLINER INLINER						\
  real static inline *algnrl(real *X) {assert(algn_general(X, BytesPerBlock)>=(uintptr_t)X); return (real *)  algn_general(X, BytesPerBlock); } \
  \
Uint static inline RealAlign(Uint X) { return BytesPerBlock / sizeof(real) + (1L +  (X - 1L) / CodesPerBlock) * CodesPerBlock; }


Uint *AlignBase(SEXP CM, Uint nr, Uint bytesperblock, bool test);
#define Align(CM, nr) AlignBase(CM, nr, BytesPerBlock, true)
#define AlignTest(CM, nr, test) AlignBase(CM, nr, BytesPerBlock, test)

Uint* DoAlign(SEXP SNPxIndiv, Uint nr, snpcoding method);
//Uint* DoAlignWithoutTest(SEXP CM, Uint nr, snpcoding method);
SEXP createSNPmatrix(Uint snps, Uint individuals, snpcoding method);
Ulong sumGeno(Uint * SNPxIndiv, Uint snps, Uint individuals, snpcoding method);

void ReUseAs(SEXP Code, snpcoding method);
Uint haplo2geno(Uint * SNPxIndiv, Uint snps, Uint individuals,
		snpcoding method, Uint unitsPerIndiv, Uint *A);

double *matrix_mult(Uint * SNPxIndiv, Uint snps, Uint individuals,
		    snpcoding method,  bool centred, bool normalized,
		    Ulong SumGeno);

int GetCodesPerBlock(snpcoding method);

#endif
```
