## Supplementary material for "MoBPS - Modular Breeding Program Simulator": miraculix R-package: initNerror.cc

```
/* 
 Authors
 Martin Schlather,

 library for simulation of random fields -- init part and error messages

 Copyright (C) 2001 -- 2019 Martin Schlather

This program is free software; you can redistribute it and/or
modify it under the terms of the GNU General Public License
as published by the Free Software Foundation; either version 3
of the License, or (at your option) any later version.

This program is distributed in the hope that it will be useful,
but WITHOUT ANY WARRANTY; without even the implied warranty of
MERCHANTABILITY or FITNESS FOR A PARTICULAR PURPOSE.  See the
GNU General Public License for more details.

You should have received a copy of the GNU General Public License
along with this program; if not, write to the Free Software
Foundation, Inc., 59 Temple Place - Suite 330, Boston, MA  02111-1307, USA.
*/


#include <stdio.h>  
//#include <stdlib.h>
//#include <sys/timeb.h> 
 
#include <string.h>
#include "error.h"


char ERRMSG[LENERRMSG], MSG[LENERRMSG], BUG_MSG[LENMSG], MSG2[LENERRMSG];
errorloc_type ERROR_LOC;
errorstring_type ERRORSTRING;
```
