## Supplementary material for "MoBPS - Modular Breeding Program Simulator": miraculix R-package: intrinsics.h

```
/*
 Authors 
 Martin Schlather,


 Copyright (C) 2018 -- 2019  Martin Schlather

This program is free software; you can redistribute it and/or
modify it under the terms of the GNU General Public License
as published by the Free Software Foundation; either version 3
of the License, or (at your option) any later version.

This program is distributed in the hope that it will be useful,
but WITHOUT ANY WARRANTY; without even the implied warranty of
MERCHANTABILITY or FITNESS FOR A PARTICULAR PURPOSE.  See the
GNU General Public License for more details.

You should have received a copy of the GNU General Public License
along with this program; if not, write to the Free Software
Foundation, Inc., 59 Temple Place - Suite 330, Boston, MA  02111-1307, USA.  
*/


#ifndef miraculix_initrinsics_H
#define miraculix_initrinsics_H 1

#include <inttypes.h>
#include "MX.h"

// PKG_CXXFLAGS =  $(SHLIB_OPENMP_CXXFLAGS) -mavx -msse -msse2 -msse3 -msse4 -msse4a -march=core-avx2


#if defined SSE
//#include <xmmintrin.h>
#endif

#if defined SSE2
#include <emmintrin.h>
#endif

#if defined SSE3
//#include <pmmintrin.h>
#endif

#if defined SSSE3
#include <tmmintrin.h>
#endif

#if defined SSE4A
//#include <ammintrin.h>
#endif

#if defined SSE412
//#include <smmintrin.h>
#endif

#if defined AVX || defined AVX2
#include <x86intrin.h>
#endif

#if defined AVX512
//#include <immintrin.h>
#endif


#ifndef UINT64_C
#define UINT64_C(X) X
#endif


typedef union uni64{
  uint64_t u64;
  double d8;
  float f4[2];
  uint32_t u32[2];
  uint16_t u16[4];
  uint8_t u8[8];
} uni64;

typedef union uni128 {
#if defined SSE || defined SSE2
  __m128i vi; // hamming
  __m128d d;
  __m128d d128[1];
  __m64 m64[2];
#endif
  uint64_t u64[2];// hamming
  uint32_t u32[4];
  uint8_t u8[16]; // hamming, shuffle
} uni128;


typedef union uni256 {
#if defined AVX || defined AVX2
  __m256i vi; // hamming
  __m256d d;
  __m128d d128[2];
  __m64 m64[4];
#endif
  uint64_t u64[4];// hamming
  uint32_t u32[8];
  uint8_t u8[32]; // hamming, shuffle
} uni256;


#if defined AVX2
#define BlockType0 __m256i
#define BytesPerBlock 32L
#define Double __m256d
#define Float __m256
#define BlockUnitType0 uni256

#elif defined SSE2
#define BlockType0 __m128i 
#define BytesPerBlock 16
#define Double __m128d
#define Float __m128
#define BlockUnitType0 uni128

#elif defined PseudoSSE // particularly shuffle (but too slow && with errors)
//_CRT_ALIGN(16) 
#define uni uni128
#define __m128i uni // ok
#define __m128d uni // ok
#define __m128 uni  // ok
#define BlockType0 uni
#define BytesPerBlock 16L
#define Double uni
#define Float uni
#define BlockUnitType0 uni
#define VI

#elif defined UNCOMPRESSED
#define BlockType0 Uint
#define BytesPerBlock 4L

#else  // particularly Bit23
#define uni uni64
#define BlockType0 uint64_t
#define BytesPerBlock 8L
#define BlockUnitType0 uni

#endif


#define UnitsPerBlock (BytesPerBlock / BytesPerUnit) 
#define BitsPerUnit (BytesPerUnit * BitsPerByte) 
#define BitsPerBlock (BytesPerBlock * BitsPerByte) 
#define CodesPerUnit (BitsPerUnit / BitsPerCode)
#define CodesPerBlock (CodesPerUnit * UnitsPerBlock)
#define CodesPerByte (BitsPerByte / BitsPerCode)
#define DoublesPerBlock (BytesPerBlock / BytesPerDouble)
#define CodeMask ((1L << BitsPerCode) - 1L)


 // Achtung! bei 16-bit-Verfahren wie TwoBits oder ThreeBits kann
  // BitsPerUnit != 2 * BitsPer16Bit sein.
  // dies wuerde alle definitionen hier in 'intrinsic.h' zerstoeren !!
//#if defined BitsPerCode && defined SCHLATHERS_MACHINE
//s-tatic_assert(BitsPerCode == 2L || BitsPerCode == 3L || BitsPerCode == 4L ||
//	      BitsPerCode == BitsPerUnit, "'BitsPerCode' not of correct size");
//#endif


#define ALIGNED __attribute__ ((aligned (BytesPerBlock)))
#define BlockType BlockType0 ALIGNED
#define BlockUnitType BlockUnitType0 ALIGNED


// #if defined AVX || defined SSE2 || defined AVX2 || defined AVX256
#if defined SSE2 || defined AVX2
#define VI .vi
#define U128(X) .u128[X]
#endif


// needed for immitations of sse/ssse3 functions
#define X128F64none(Z,A) Z(A,0); Z(A,1); 
#define X128F64(Z,A,B) Z(A,B,0); Z(A,B,1); 
#define X128F32(Z,A,B) X128F64(Z,A,B); Z(A,B,2); Z(A,B,3); 
#define X128F16(Z,A,B) X128F32(Z,A,B); Z(A,B,4); Z(A,B,5); Z(A,B,6); Z(A,B,7);
#define X128F8(Z,A,B) X128F16(Z,A,B); Z(A,B,8); Z(A,B,9);Z(A,B,10); Z(A,B,11);Z(A,B,12); Z(A,B,13); Z(A,B,14); Z(A,B,15);
#define X128F64bi(Z,A,B,C) Z(A,B,C,0); Z(A,B,C,1); 
#define X128F32bi(Z,A,B,C) X128F64bi(Z,A,B,C); Z(A,B,C,2); Z(A,B,C,3);
#define X128F16bi(Z,A,B,C) X128F32bi(Z,A,B,C); Z(A,B,C,4); Z(A,B,C,5); Z(A,B,C,6); Z(A,B,C,7);
#define X128F8bi(Z,A,B,C) X128F16bi(Z,A,B,C); Z(A,B,C,8); Z(A,B,C,9);Z(A,B,C,10); Z(A,B,C,11);Z(A,B,C,12); Z(A,B,C,13); Z(A,B,C,14); Z(A,B,C,15);


#if defined AVX2
#define AND(A,B,C) A = _mm256_and_si256(B,C)
#define OR(A,B,C) A = _mm256_or_si256(B,C)
#define XOR(A,B,C) A = _mm256_xor_si256(B,C)
#define SHR32(A,B,C) A = _mm256_srli_epi32(B,C) // see also _mm256512_rol_epi64,
#define SHL32(A,B,C) A = _mm256_slli_epi32(B,C)
#define SHR16(A,B,C) A = _mm256_srli_epi16(B,C)
#define SHR64(A,B,C) A = _mm256_srli_epi64(B,C)
#define SHUFFLE8(A,B,C) A = _mm256_shuffle_epi8(B,C)

#define SET8(A,B) A = _mm256_set1_epi8(B)
#define SETREV8(A,B0,B1,B2,B3,B4,B5,B6,B7,B8,B9,B10,B11,B12,B13,B14,B15) A = _mm256_setr_epi8(B0,B1,B2,B3,B4,B5,B6,B7,B8,B9,B10,B11,B12,B13,B14,B15,B0,B1,B2,B3,B4,B5,B6,B7,B8,B9,B10,B11,B12,B13,B14,B15)
#define SET16(A,B) A = _mm256_set1_epi16(B)
BlockType set_epi32(Uint X);
#define SET32(A,X) A = set_epi32(X) // _mm256_set1_epi32(X) funktioniert nicht!?
#define ZERO(A)  A = _mm256_setzero_si256()
#define LOAD(A,B) A = _mm256_load_si256(B)
#define LOADU(A,B) A = _mm256_loadu_si256(B)
//#define LOAD_DOUBLE(A,B) A =  _mm256_load_pd(B)
//#define LOAD1_DOUBLE(A,B) A =  _mm256_load1_pd(B)
#define STORE_DOUBLE(A,B) _mm256_store_pd(A,B)
#define EXTRACT16(A, B, C) A = _mm256_extract_epi16(B, C)

#define ADD8(A,B,C) A= _mm256_add_epi8(B,C)
#define ADD32(A,B,C) A = _mm256_add_epi32(B,C)
#define ADD64(A,B,C) A = _mm256_add_epi64(B,C)
#define SAD8(A,B,C) A =  _mm256_sad_epu8(B,C)
#define INT2FLOAT(A,B) A = _mm_cvtepi32_ps(B) // 128 bit only !!
#define MULTFLOAT(A,B,C) A = _mm256_mul_ps (B,C)
#define ADDFLOAT(A,B,C) A = _mm256_add_ps (B,C)
#define SUBFLOAT(A,B,C) A =  _mm256_sub_ps (B,C)
#define ZEROFLOAT(A,B) A =_mm256_setzero_ps()
#define INT2DOUBLE(A,B) A =_mm_cvtpi32_pd(B) // 128 bit only !!
#define MULTDOUBLE(A,B,C) A = _mm256_mul_pd(B,C)
#define ADDDOUBLE(A,B,C) A =  _mm256_add_pd(B,C)
#define SUBDOUBLE(A,B,C) A =  _mm256_sub_pd(B,C)
#define ZERODOUBLE(A) A = _mm256_setzero_pd()


#elif defined SSE2 

#define AND(A,B,C) A = _mm_and_si128(B,C)
#define OR(A,B,C) A = _mm_or_si128(B,C)
#define XOR(A,B,C) A = _mm_xor_si128(B,C)
#define SHR32(A,B,C) A = _mm_srli_epi32(B,C) // see also _mm512_rol_epi64,
#define SHL32(A,B,C) A = _mm_slli_epi32(B,C)
#define SHR16(A,B,C) A = _mm_srli_epi16(B,C)
#define SHR64(A,B,C) A = _mm_srli_epi64(B,C)

#define SET8(A,B) A = _mm_set1_epi8(B)
#define SETREV8(A,B0,B1,B2,B3,B4,B5,B6,B7,B8,B9,B10,B11,B12,B13,B14,B15) A = _mm_setr_epi8(B0,B1,B2,B3,B4,B5,B6,B7,B8,B9,B10,B11,B12,B13,B14,B15)
#define SET16(A,B) A = _mm_set1_epi16(B)
#define SET32(A,B) A = _mm_set1_epi32(B)
//#define SET_EPI64(A,B) A = _mm_set1_epi64(B,C)
//#define SET_EPI32(X) _mm_set_epi32(X,X,X,X)
#define ZERO(A)  A = _mm_setzero_si128()
#define LOAD(A,B) A = _mm_load_si128(B)
#define LOADU(A,B) A = _mm_loadu_si128(B)
//#define LOAD_DOUBLE(A,B) A =  _mm_load_pd(B)
//#define LOAD1_DOUBLE(A,B) A =  _mm_load1_pd(B)
#define STORE_DOUBLE(A,B) _mm_store_pd(A,B)
#define EXTRACT16(A, B, C) A = _mm_extract_epi16(B, C)

#define ADD8(A,B,C) A= _mm_add_epi8(B,C)
#define ADD32(A,B,C) A = _mm_add_epi32(B,C)
#define ADD64(A,B,C) A = _mm_add_epi64(B,C)
#define SAD8(A,B,C) A =  _mm_sad_epu8(B,C)
#define INT2FLOAT(A,B) A = _mm_cvtepi32_ps(B)
#define MULTFLOAT(A,B,C) A = _mm_mul_ps (B,C)
#define ADDFLOAT(A,B,C) A = _mm_add_ps (B,C)
#define SUBFLOAT(A,B,C) A =  _mm_sub_ps (B,C)
#define ZEROFLOAT(A,B) A =_mm_setzero_ps()
#define INT2DOUBLE(A,B) A =_mm_cvtpi32_pd(B) // very expensive
#define MULTDOUBLE(A,B,C) A = _mm_mul_pd(B,C)
#define ADDDOUBLE(A,B,C) A =  _mm_add_pd(B,C)
#define SUBDOUBLE(A,B,C) A =  _mm_sub_pd(B,C)
#define ZERODOUBLE(A) A = _mm_setzero_pd()
//#define MOVEMASK _mm_movemask_ps
//#define BLEND _mm_blend_pd //see also _mm512_mask_inserti64x4_mm_insert_epi64

#if defined SSSE3
#define SHUFFLE8(A,B,C) A = _mm_shuffle_epi8(B,C)
#else
#define SHUFFLE8X(A,B,C,X) ((uni128*) &A)->u8[X] = ((uni128*) &B)->u8[((uni128*) &C)->u8[X]] // ACHTUNG A != B !!
#define SHUFFLE8(A,B,C) { X128F8bi(SHUFFLE8X,A,B,C) }
#endif

// NONE
#elif defined PseudoSSE

#define ANDX(A,B,C,X) (A).u64[X] = (B).u64[X] & (C).u64[X] 
#define AND(A,B,C) { X128F64bi(ANDX,A,B,C) }
#define ORX(A,B,C,X) (A).u64[X] = (B).u64[X] | (C).u64[X] 
#define OR(A,B,C) { X128F64bi(ORX,A,B,C) }
#define XORX(A,B,C,X) \
  (A).u64[X] = (B).u64[X] xor (C).u64[X] 
#define XOR(A,B,C) { X128F64bi(XORX,A,B,C) }

#define SHR32X(A,B,C,X) (A).u32[X] = (B).u32[X] >> C; 
#define SHR32(A,B,C) { X128F32bi(SHR32X,A,B,C) }
#define SHL32X(A,B,C,X) (A).u32[X] = (B).u32[X] << C; 
#define SHL32(A,B,C) { X128F32bi(SHL32X,A,B,C) }
#define SHR16X(A,B,C,X) (A).u16[X] = (B).u16[X] >> C;
#define SHR16(A,B,C) { X128F16bi(SHR16X,A,B,C) }
#define SHR64X(A,B,C,X) (A).u64[X] = (B).u64[X] >> C;
#define SHR64(A,B,C) { X128F64bi(SHR64X,A,B,C) }
#define SHUFFLE8X(A,B,C,X)  (A).u8[X] = (B).u8[(C).u8[X]] // ACHTUNG A != B !!
#define SHUFFLE8(A,B,C) { assert((A) != (B)); X128F8bi(SHUFFLE8X,A,B,C) }

#define SET8X(A,B,X) (A).u8[X] = B
#define SET8(A,B) { X128F8(SET8X,A,B) }
#define SETREV8(A,B0,B1,B2,B3,B4,B5,B6,B7,B8,B9,B10,B11,B12,B13,B14,B15) { \
  (A).u8[0] = B0; (A).u8[1] = B1; (A).u8[2] = B2; (A).u8[3] = B3;	\
  (A).u8[4] = B4; (A).u8[5] = B5; (A).u8[6] = B6; (A).u8[7] = B7;	\
  (A).u8[8] = B8; (A).u8[9] = B9; (A).u8[10] = B10; (A).u8[11] = B11;	\
  (A).u8[12] = B12; (A).u8[13] = B13; (A).u8[14] = B14; (A).u8[15] = B15; \
  }
#define SET16X(A,B,X) (A).u16[X] = B
#define SET16(A,B) { X128F16(SET16X,A,B) }
#define SET32X(A,B,X) (A).u32[X] = B
#define SET32(A,B) { X128F32(SET32X,A,B) }

#define ZEROX(A,X) (A).u64[X] = 0
#define ZERO(A) { X128F64none(ZEROX,A) }
#define LOADX(A,B,X) (A).u64[X] = (B)->u64[X]
#define LOAD(A,B) { X128F64(LOADX,A,B) }
#define LOADU LOAD
#define STORE_DOUBLEX(A,B,X) (A).d8[X] = (B)->d8[X]
#define STORE_DOUBLE(A,B) { X128F64(STORE_DOUBLEX,A,B) }
#define EXTRACT16(A, B, C) A = (int) ((B).u32[C/2])

#define ADD8X(A,B,C,X) (A).u8[X] = (B).u8[X] + (C).u8[X];
#define ADD8(A,B,C) { X128F8bi(ADD8X,A,B,C) }
#define ADD32X(A,B,C,X) (A).u32[X] = (B).u32[X] + (C).u32[X];
#define ADD32(A,B,C) { X128F32bi(ADD32X,A,B,C) }
#define ADD64X(A,B,C,X) (A).u64[X] = (B).u64[X]+(C).u64[X];
#define ADD64(A,B,C) { X128F64bi(ADD64X,A,B,C) }
#define SAD8X(B,C,X) (int) ((B).u8[X]>(C).u8[X] ? (B).u8[X]-(C).u8[X] : (C).u8[X]-(B).u8[X])
#define SAD8(A,B,C) {						\
    (A).u64[0] = SAD8X(B,C,0) + SAD8X(B,C,1) + SAD8X(B,C,2) + SAD8X(B,C,3) + SAD8X(B,C,4) + SAD8X(B,C,5) + SAD8X(B,C,6) + SAD8X(B,C,7); \
    (A).u64[1] = SAD8X(B,C,0) + SAD8X(B,C,1) + SAD8X(B,C,2) + SAD8X(B,C,3) + SAD8X(B,C,4) + SAD8X(B,C,5) + SAD8X(B,C,6) + SAD8X(B,C,7); \
}


#define INT2FLOATX(A,B,X) (A).f4[X] = (float) ((B).u32[X])
#define INT2FLOAT(A,B) { uni BB=B; X128F32(INT2FLOATX,A,B) }
#define MULTFLOATX(A,B,C,X) (A).f4[X] = (B).f4[X] * (C).f4[X]
#define MULTFLOAT(A,B,C) { X128F32bi(MULTFLOATX,A,B,C) }
#define ADDFLOATX(A,B,C,X) (A).f4[X] = (B).f4[X] + (C).f4[X]
#define ADDFLOAT(A,B,C) { X128F32bi(ADDFLOATX,A,B,C) }
#define SUBFLOATX(A,B,C,X) (A).f4[X] = (B).f4[X] - (C).f4[X]
#define SUBFLOAT(A,B,C) { X128F32bi(SUBFLOATX,A,B,C) }
#define ZEROFLOATX(A,X) (A).f4[X] = 0.0;
#define ZEROFLOAT(A) { X128F32none(ZEROFLOATX,A,B,C) }

#define INT2DOUBLEX(A,B,X)  (A).d8[X] = (double) ((B).u32[X])
// #define INT2DOUBLE(A,B) { assert({uni64  BB=B;}) X128F64(INT2DOUBLEX,A,B) }
#define INT2DOUBLE(A,B) { X128F64(INT2DOUBLEX,A,B) }
#define MULTDOUBLEX(A,B,C,X) (A).d8[X] = (B).d8[X] * (C).d8[X]
#define MULTDOUBLE(A,B,C) { X128F64bi(MULTDOUBLEX,A,B,C) }
#define ADDDOUBLEX(A,B,C,X) (A).d8[X] = (B).d8[X] + (C).d8[X]
#define ADDDOUBLE(A,B,C) { X128F64bi(ADDDOUBLEX,A,B,C) }
#define SUBDOUBLEX(A,B,C,X) (A).d8[X] = (B).d8[X] - (C).d8[X]
#define SUBDOUBLE(A,B,C) { X128F64bi(SUBDOUBLEX,A,B,C) }
#define ZERODOUBLEX(A,X) (A).d8[X] = 0.0;
#define ZERODOUBLE(A) { X128F64none(ZERODOUBLEX,A) }
//#define MOVEMASK _mm_movemask_ps
//#define BLEND _mm_blend_pd //see also _mm512_mask_inserti64x4_mm_insert_epi64

#elif defined UNCOMPRESSED
// nothing to do

#else 
// do not do anything

#endif


 
// #define Uint uint32 // Window

#endif
```
