## Supplementary material for "MoBPS - Modular Breeding Program Simulator": miraculix R-package: IntrinsicsBase.h

```
/*
 Authors 
 Martin Schlather,


 Copyright (C) 2018 -- 2019  Martin Schlather

This program is free software; you can redistribute it and/or
modify it under the terms of the GNU General Public License
as published by the Free Software Foundation; either version 3
of the License, or (at your option) any later version.

This program is distributed in the hope that it will be useful,
but WITHOUT ANY WARRANTY; without even the implied warranty of
MERCHANTABILITY or FITNESS FOR A PARTICULAR PURPOSE.  See the
GNU General Public License for more details.

You should have received a copy of the GNU General Public License
along with this program; if not, write to the Free Software
Foundation, Inc., 59 Temple Place - Suite 330, Boston, MA  02111-1307, USA.  
*/


#ifndef miraculix_IntrinsicsBase_H
#define miraculix_IntrinsicsBase_H 1

//#if !defined _WIN32

#if defined __AVX2__
#define AVX2 1
#endif
#if defined __AVX__ 
#define AVX 1
#endif
#if defined __SSE41__ 
#define SSE41 1
#endif
#if defined  __SSE4A__
#define SSE4A 1
#endif
#if defined  __SSSE3__  // needs SSE2 as well
#define SSSE3 1
#endif
#if defined  __SSE3__ 
#define SSE3 1
#endif
#if defined  __SSE2__ 
#define SSE2 1 
#endif
#if defined __SSE__ 
#define SSE 1
#endif

// #endif // not WIN32

#endif
```
