## Supplementary material for "MoBPS - Modular Breeding Program Simulator": miraculix R-package: kleinkram.cc

```
/*
 Authors 
 Martin Schlather,

 Copyright (C) 2015 -- 2017 Martin Schlather

This program is free software; you can redistribute it and/or
modify it under the terms of the GNU General Public License
as published by the Free Software Foundation; either version 3
of the License, or (at your option) any later version.

This program is distributed in the hope that it will be useful,
but WITHOUT ANY WARRANTY; without even the implied warranty of
MERCHANTABILITY or FITNESS FOR A PARTICULAR PURPOSE.  See the
GNU General Public License for more details.

You should have received a copy of the GNU General Public License
along with this program; if not, write to the Free Software
Foundation, Inc., 59 Temple Place - Suite 330, Boston, MA  02111-1307, USA.  
*/

#include <R_ext/Lapack.h>
//#include "def.h" // never change this line
#include <General_utils.h> //#include <General_utils.h>
#include "xport_import.h"


#define SCALAR(A,B,C) Ext_scalarX(A,B,C, SCALAR_AVX)
//#define SCALARINT(A,B,C) Ext_scalarInt(A,B,C, SCALAR_BASE)
#define LINEAR(A,B,C,D) Ext_linearX(A,B,C,D,6)

// void xxx() {  solvePosDefSp(NULL, 0, 0, NULL, 0, NULL, NULL, NULL); }

void strcopyN(char *dest, const char *src, int n) {
  if (n > 1) {
    n--; 
    strncpy(dest, src, n);
  }
  dest[n] = '\0';
}

void AtA(double *a, int nrow, int ncol, double *C) {
  // C =  A^T %*% A
#ifdef DO_PARALLEL
#pragma omp parallel for num_threads(CORES) if (MULTIMINSIZE(ncol)) schedule(dynamic, 20)
#endif  
  for (int i=0; i<ncol; i++) {
    double 
      *A = a + i * nrow,
      *B = A;
    for (int j=i; j<ncol; j++, B+=nrow) {
      C[i * ncol + j] = C[i + ncol * j] = SCALAR(A, B, nrow);
    }
  }
}
 

void xA_noomp(double *x, double*A, int nrow, int ncol, double *y) {
  if (A == NULL) {
    if (nrow != ncol || nrow <= 0) BUG;
    MEMCOPY(y, x, sizeof(double) * nrow);
  } else {
    for (int i=0; i<ncol; i++) {
      y[i] = SCALAR(x, A + i * nrow, nrow);
    }
  }
}

void xA(double *x, double*A, int nrow, int ncol, double *y) {
  if (A == NULL) {
    if (nrow != ncol || nrow <= 0) BUG;
    MEMCOPY(y, x, sizeof(double) * nrow);
  } else {
#ifdef DO_PARALLEL
#pragma omp parallel for num_threads(CORES) if (MULTIMINSIZE(ncol) && MULTIMINSIZE(nrow))
#endif  
   for (int i=0; i<ncol; i++) y[i] = SCALAR(x, A + i * nrow, nrow);
  } 
} 
  
void xA(double *x1, double *x2,  double*A, int nrow, int ncol, double *y1,
	double *y2) {
  if (A == NULL) {
    if (nrow != ncol || nrow <= 0) BUG;
    MEMCOPY(y1, x1, sizeof(double) * nrow);
    MEMCOPY(y2, x2, sizeof(double) * nrow);
  } else {
    double *a = A;
    for (int i=0; i<ncol; i++, a += nrow) {
      y1[i] = SCALAR(x1, a, nrow);
      y2[i] = SCALAR(x2, a, nrow);
    }
  }	
}

void xAx(double *x, double*A, int nrow,  double *y) {
  double sum = 0.0;
#ifdef DO_PARALLEL
#pragma omp parallel for num_threads(CORES) reduction(+:sum) if (MULTIMINSIZE(nrow) && MULTIMINSIZE(nrow))
#endif  
  for (int i=0; i<nrow; i++) sum += x[i] * SCALAR(x, A + i * nrow, nrow);
  *y = sum;
}

void Ax(double *A, double*x, int nrow, int ncol, double *y) {
  if (A == NULL) {
    if (nrow != ncol || nrow <= 0) BUG;
    MEMCOPY(y, x, sizeof(double) * nrow);
  } else {
#ifdef DO_PARALLEL
#pragma omp parallel for num_threads(CORES) if (MULTIMINSIZE(ncol) && MULTIMINSIZE(nrow))
    for (int j=0; j<nrow; j++) {
      double dummy = 0.0;
      int k = j;
      for (int i=0; i<ncol; i++, k+=nrow) { 
	dummy += A[k] * x[i];
      }
      y[j] = dummy;
    }
#else
    for (int i=0; i<nrow; i++) y[i]=0.0;
    for (int k=0, i=0; i<ncol; i++) { 
      for (int j=0; j<nrow; j++) {
	y[j] += A[k++] * x[i];
      }
    }
#endif  
  }
}


void Ax(double *A, double*x1, double*x2, int nrow, int ncol, double *y1,
	double *y2) {
  if (A == NULL) {
    if (nrow != ncol || nrow <= 0) BUG;
    MEMCOPY(y1, x1, sizeof(double) * nrow);
    MEMCOPY(y2, x2, sizeof(double) * nrow);
  } else {
    for (int i=0; i<nrow; i++) y1[i]=y2[i]=0.0;
    for (int k=0, i=0; i<ncol; i++) { 
      for (int j=0; j<nrow; j++) {
	y1[j] += A[k] * x1[i];
	y2[j] += A[k++] * x2[i];
      }
    }
  }
}


double XkCXtl(double *X, double *C, int nrow, int dim, int k, int l) {
  // (k-th row of X) * C * (l-th row of X)
  // X is nrow x dim matrix
  // C is dim x dim matrix
  double
    *pX = X + k, 
    *pY = X + l, 
    result = 0.0;
  int size = nrow * dim;
  
#ifdef DO_PARALLEL
#pragma omp parallel for num_threads(CORES) reduction(+:result)
#endif
  for (int j=0; j<size; j+=nrow) {
    double scalar = 0.0;
    int ci = j * dim;
    for (int i=0; i<size; i+=nrow) scalar += pX[i] * C[ci++];
    result += scalar * pY[j];
  }
  return result;
}


void XCXt(double *X, double *C, double *V, int nrow, int dim /* dim of C */) {
  int size = nrow * dim;
  double  
    *endpX = X + nrow,
    *dummy = (double*) MALLOC(sizeof(double) * size); // dummy = XC
  if (dummy == NULL) RFERROR("XCXt: memory allocation error in XCXt");
 
#ifdef DO_PARALLEL
#pragma omp parallel for num_threads(CORES) 
#endif
  for (double *pX = X; pX < endpX; pX++) {
    double *pdummy = dummy + (pX - X);
    for (int ci=0, cd=0; cd<size; cd+=nrow) {
      double scalar = 0.0;
      for (int i=0; i<size; i+=nrow) {
        scalar += pX[i] * C[ci++];
      }
      pdummy[cd] = scalar;
    }
  }

  // V = dummy X^t
#ifdef DO_PARALLEL
#pragma omp parallel for num_threads(CORES) 
#endif
  for (int rv=0; rv<nrow; rv++) {
    for (int cv=rv; cv<nrow; cv++) {
      double scalar=0.0;
      for (int i=0; i<size; i+=nrow) {
	scalar += dummy[rv + i] * X[cv + i];
     }
      V[rv + cv * nrow] = V[cv + rv * nrow] = scalar;
    }
  }

  UNCONDFREE(dummy);
}


double xUy(double *x, double *U, double *y, int dim) {
  // U a symmetric matrix given by its upper triangular part
  double xVy = 0.0;
  int    dimM1 = dim - 1;
#ifdef DO_PARALLEL
#pragma omp parallel for num_threads(CORES) if (MULTIMINSIZE(dim)) reduction(+:xVy) 
#endif  
  for (int d=0; d<dim; d++) {
    int i, 
      j = dim * d;
    double dummy = 0.0;
    for (i=0; i<=d; i++) dummy += x[i] * U[j++];
    for (j += dimM1; i<dim; i++, j+=dim) dummy += x[i] * U[j];
    xVy += dummy * y[d];
  }
  return xVy;
}

/*

  // U a symmetric matrix given by its upper triangular part
  assert(z != NULL);
  int   dimM1 = dim - 1;
#ifdef DO_PARALLEL
#pragma omp parallel for num_threads(CORES) if (MULTIMINSIZE(dim))
#endif  
  for (int d=0; d<dim; d++) {
    double dummy;
    int i,
      j = dim * d;
    for (dummy = 0.0, i=0; i<=d; i++) dummy += x[i] * U[j++];
    for (j += dimM1; i<dim; i++, j+=dim) dummy += x[i] * U[j];
    if (z!=NULL) z[d] = dummy;
  }
  double xVx;
  SCALAR_PROD(z, x, dim, xVx);
  return xVx;

 */

double xUxz(double *x, double *U, int dim, double *z) {
 double xVx = 0.0;
  int dimM1 = dim - 1;
#ifdef DO_PARALLEL
#pragma omp parallel for num_threads(CORES) reduction(+:xVx)
#endif
  for (int d=0; d<dim; d++) {
    int i, 
      j = dim * d;
    double dummy = 0.0;
    for (dummy = 0.0, i=0; i<=d; i++) dummy += x[i] * U[j++];
    for (j += dimM1; i<dim; i++, j+=dim) dummy += x[i] * U[j];
    if (z != NULL) z[d] = dummy;
    xVx += dummy * x[d];
  }
  return xVx;
}

double xUx(double *x, double *U, int dim) {
    return xUxz(x, U, dim, NULL);
}

double x_UxPz(double *x, double *U, double *z, int dim) {
// x^t (Ux + z); U dreieckmatrix
  double xVx = 0.0;
  int    dimM1 = dim - 1;
#ifdef DO_PARALLEL
#pragma omp parallel for num_threads(CORES) reduction(+:xVx)
#endif
  for (int d=0; d<dim; d++) {
    int i,
      j = dim * d;
    double dummy = z[d];
    for (i=0; i<=d; i++) dummy += x[i] * U[j++];
    for (j += dimM1; i<dim; i++, j+=dim) dummy += x[i] * U[j];
    xVx += dummy * x[d];
  }
  return xVx;
}


void matmult(double *a, double *b, double *c, int l, int m, int n) {
// multiplying an lxm- and an mxn-matrix, saving result in C
#ifdef DO_PARALLEL
#pragma omp parallel for num_threads(CORES) 
#endif
   for (int i=0; i<l; i++) {
     double *A = a + i,
       *C = c + i;
     for (int j=0; j<n; j++) {
       double dummy = 0.0,
	 *B = b + j * m;
       for (int k=0; k<m; k++) dummy += A[k*l] * B[k];
       C[j * l] = dummy;
     }
   }
}


void matmulttransposed(double *A, double *B, double *c, int m, int l, int n) {
// multiplying t(A) and B with dim(A)=(m,l) and dim(B)=(m,n),
// saving result in C
#ifdef DO_PARALLEL
#pragma omp parallel for num_threads(CORES) 
#endif
  for (int i=0; i<l; i++) {    
    double *C = c + i,
      *Aim = A + i * m;
    for (int j=0; j<n; j++) C[j * l] = SCALAR(Aim, B + j * m, m);
  }
}


/*
void matmulttransposedInt(int *A, int *B, int *c, int m, int l, int n) {
// multiplying t(A) and B with dim(A)=(m,l) and dim(B)=(m,n),
// saving result in C
#ifdef DO_PARALLEL
#pragma omp parallel for num_threads(CORES) 
#endif
  for (int i=0; i<l; i++) {    
    int *C = c + i,
      *Aim = A + i * m;
    for (int j=0; j<n; j++) C[j * l] = SCALARINT(Aim, B + j * m, m);
  }
}

*/


void matmult_2ndtransp(double *a, double *B, double *c, int l, int m, int n) {
// multiplying A and t(B) with dim(A)=(l, m) and dim(B)=(n, m),
// saving result in C
#ifdef DO_PARALLEL
#pragma omp parallel for num_threads(CORES) if (l * m * n > 1000)
#endif
  for (int i=0; i<l; i++) {
    double *C = c + i,
      *A = a + i;
    for (int j=0; j<n; j++) {
       double dummy = 0.0,
	 *Bj = B + j;
       for (int k=0; k<m; k++) dummy += A[k * l] * Bj[k * n];
       C[j*l] = dummy;
    }
  }
}


void matmult_2ndtransp(double *a, double *B, double *c, int l, int m) {
// multiplying A and t(B) with dim(A)=(l, m) and dim(B)=(n, m),
// saving result in C
  int lm = l  * m;
#ifdef DO_PARALLEL
#pragma omp parallel for num_threads(CORES) if (l * m * l > 1000)
#endif
  for (int i=0; i<l; i++) {
    double *C = c + i,
      *A = a + i;
    for (int j=0; j<l; j++) {
       double dummy = 0.0,
	 *Bj = B + j;
       for (int k=0; k<lm; k+=l) dummy += A[k] * Bj[k];
       C[j*l] = dummy;
    }
  }
}


void matmult_tt(double *a, double *B, double *c, int m, int l, int n) {
// calculating t(A B) with dim(A)=(m,l) and dim(B)=(m,n),
// saving result in C
#ifdef DO_PARALLEL
#pragma omp parallel for num_threads(CORES) 
#endif
  for (int i=0; i<l; i++) {
    double *A = a + i,
      *C = c + i * l;
    for (int j=0; j<n; j++) {
      double dummy = 0.0,
	*Bjm = B + j * m;
      for (int k=0; k<m; k++) dummy += A[k * l] * Bjm[k];
      C[j] = dummy;
    }
  }
}


void Xmatmult(double *A, double *B, double *C, int l, int m, int n) {
// multiplying an lxm- and an mxn-matrix, saving result in C
#ifdef DO_PARALLEL
#pragma omp parallel for num_threads(CORES) 
#endif
  for (int i=0; i<l; i++) {
    for (int jl=i, jm=0, j=0; j<n; j++, jl+=l, jm+=m) {
      double dummy = 0.0;
      int endfor = jm + m;
      for (int kl=i, k=jm; k<endfor; k++, kl+=l) dummy += A[kl] * B[k]; 
      C[jl] = dummy;
    }
  }
}

void Xmatmulttransposed(double *A, double *B, double *C, int m, int l, int n) {
// multiplying t(A) and B with dim(A)=(m,l) and dim(B)=(m,n),
// saving result in C
#ifdef DO_PARALLEL
#pragma omp parallel for num_threads(CORES) 
#endif
  for (int i=0; i<l; i++) {
    int im = i * m;
    for (int jl=i, jm=0, j=0; j<n; j++, jl+=l, jm+=m) {
      double dummy = 0.0;
      int endfor = im + m;
      for (int jmk=jm, k=im; k<endfor; k++) dummy += A[k] * B[jmk++]; 
      C[jl] = dummy;
    }
  }
}


double *matrixmult(double *m1, double *m2, int dim1, int dim2, int dim3) {
  double *m0 = (double*) MALLOC(sizeof(double) * dim1 * dim3);
  matmult(m1, m2, m0, dim1, dim2, dim3);
  return m0;
}


SEXP TooLarge(int *n, int l){
#define nTooLarge 2 // mit op
  const char *tooLarge[nTooLarge] = {"size", "msg"};
  SEXP namevec, info;
  PROTECT(info=allocVector(VECSXP, nTooLarge));
  PROTECT(namevec = allocVector(STRSXP, nTooLarge));
  for (int i=0; i<nTooLarge; i++)
    SET_STRING_ELT(namevec, i, mkChar(tooLarge[i]));
  setAttrib(info, R_NamesSymbol, namevec);
  int i=0;
  SET_VECTOR_ELT(info, i++, Int(n, l, l));
  SET_VECTOR_ELT(info, i,
		 mkString("too many elements - increase max.elements"));
  UNPROTECT(2);
  return info;
}

SEXP TooSmall(){
  SEXP namevec;
  const char *msg = "value has not been initialized";
  PROTECT(namevec = allocVector(STRSXP, 1));
  SET_STRING_ELT(namevec, 0, mkChar(msg));
  UNPROTECT(1);
  return namevec;
}


SEXP Int(int *V, int n, int max) {
  SEXP dummy;
  if (V==NULL) return allocVector(INTSXP, 0);
  if (n>max) return TooLarge(&n, 1);
   if (n<0) return TooSmall();
   PROTECT(dummy=allocVector(INTSXP, n));
  for (int i=0; i<n; i++) INTEGER(dummy)[i] = V[i];
  UNPROTECT(1);
  return dummy;
}

SEXP Int(int* V, int n) {
  return Int(V, n, MAXINT);
}


SEXP Logic(bool* V, int n, int max) {
  SEXP dummy;
  if (V==NULL) return allocVector(VECSXP, 0);
  if (n>max) return TooLarge(&n, 1);
  if (n<0) return TooSmall();
  PROTECT(dummy=allocVector(LGLSXP, n));
  for (int i=0; i<n; i++) LOGICAL(dummy)[i] = V[i];
  UNPROTECT(1);
  return dummy;
}
SEXP Logic(bool* V, int n) {
  return Logic(V, n, MAXINT);
}

SEXP Num(double* V, int n, int max) {
  SEXP dummy;
  if (V==NULL) return allocVector(REALSXP, 0);
  if (n>max) return TooLarge(&n, 1);
   if (n<0) return TooSmall();
  PROTECT(dummy=allocVector(REALSXP, n));
  for (int i=0; i<n; i++) REAL(dummy)[i] = V[i];
  UNPROTECT(1);
  return dummy;
}
SEXP Num(double* V, int n) {
  return Num(V, n, MAXINT);
}

SEXP Result(double* V, int n, int max) {
  SEXP dummy;
  if (V==NULL) return allocVector(REALSXP, 0);
  if (n>max) return TooLarge(&n, 1);
   if (n<0) return TooSmall();
 PROTECT(dummy=allocVector(REALSXP, n));
  for (int i=0; i<n; i++) REAL(dummy)[i] = (double) V[i];
  UNPROTECT(1);
  return dummy;
}

SEXP Result(double* V, int n) {
  return Result(V, n, MAXINT);
}

SEXP Char(const char **V, int n, int max) {
  SEXP dummy;
  if (V==NULL) return allocVector(STRSXP, 0);
  if (n>max) return TooLarge(&n, 1);
   if (n<0) return TooSmall();
   PROTECT(dummy=allocVector(STRSXP, n));
   for (int i=0; i<n; i++){
     SET_STRING_ELT(dummy, i, mkChar(V[i]));  
   }
  UNPROTECT(1);
  return dummy;
}

SEXP Char(const char **V, int n) {
  return Char(V, n, MAXINT);
}

SEXP Mat(double* V, int row, int col, int max) {
  if (V==NULL) return allocMatrix(REALSXP, 0, 0);
  int n = row * col;
  if (n>max) {
    int nn[2];
    nn[0] = row;
    nn[1] = col;
    return TooLarge(nn, 2);
  }
  SEXP dummy;
  PROTECT(dummy=allocMatrix(REALSXP, row, col));
  for (int i=0; i<n; i++) REAL(dummy)[i] = V[i];
  UNPROTECT(1);
  return dummy;
}

SEXP Mat(double* V, int row, int col) {
  return Mat(V, row, col, MAXINT);
}


SEXP Mat_t(double* V, int row, int col, int max) {
  if (V==NULL) return allocMatrix(REALSXP, 0, 0);
  int n = row * col;
  if (n>max) {
    int nn[2];
    nn[0] = row;
    nn[1] = col;
    return TooLarge(nn, 2);
  }
  SEXP dummy;
  PROTECT(dummy=allocMatrix(REALSXP, row, col));
  for (int k=0, j=0; j<col; j++) {
     for (int i=0; i<row; i++) {
      REAL(dummy)[k++] = V[j + col * i];
    }
  }
  UNPROTECT(1);
  return dummy;
}

SEXP Mat_t(double* V, int row, int col) {
  return Mat_t(V, row, col, MAXINT);
}


SEXP MatString(char **V, int row, int col, int max) {
  if (V==NULL) return allocMatrix(STRSXP, 0, 0);
  int n = row * col;
  if (n>max) {
    int nn[2];
    nn[0] = row;
    nn[1] = col;
    return TooLarge(nn, 2);
  }
  SEXP dummy;
  PROTECT(dummy=allocMatrix(STRSXP, row, col));
  for (int k=0; k<n; k++)
    SET_STRING_ELT(dummy, k, mkChar(V[k]));
  UNPROTECT(1);
  return dummy;
}

SEXP MatString(char** V, int row, int col) {
  return MatString(V, row, col, MAXINT);
}

SEXP MatInt(int* V, int row, int col, int max) {
  if (V==NULL) return allocMatrix(INTSXP, 0, 0);
  int n = row * col;
  if (n>max) {
    int nn[2];
    nn[0] = row;
    nn[1] = col;
    return TooLarge(nn, 2);
  }
  SEXP dummy;
  PROTECT(dummy=allocMatrix(INTSXP, row, col));
  for (int i=0; i<n; i++) INTEGER(dummy)[i] = V[i];
  UNPROTECT(1);
  return dummy;
}

SEXP MatInt(int* V, int row, int col) {
  return MatInt(V, row, col, MAXINT);
}

SEXP Array3D(double** V, int depth, int row, int col, int max) {
  if (V==NULL) return alloc3DArray(REALSXP, 0, 0, 0);
  int
    m = row * col,
    n = row * col * depth;
  if (n>max) {
    int nn[3];
    nn[0] = row;
    nn[1] = col;
    nn[2] = depth;
    return TooLarge(nn, 3);
  }
  SEXP dummy;
  PROTECT(dummy=alloc3DArray(REALSXP, depth, row, col));
  for (int j=0; j<depth; j++) {
    for (int i=0; i<m; i++) {
      REAL(dummy)[j*m+i] = V[j][i];
    }
  }
  UNPROTECT(1);
  return dummy;
}

SEXP Array3D(double** V, int depth, int row, int col) {
  return Array3D(V, depth, row, col, MAXINT);
}


usr_bool UsrBoolRelaxed(SEXP p, char *name, int idx) {
  double dummy = Real(p, name, idx);
  if (!R_finite(dummy)) return Nan;
  return dummy==0.0 ? False : True ;
}


usr_bool UsrBool(SEXP p, char *name, int idx) {
  double dummy = Real(p, name, idx);
  if (dummy == 0.0) return False;
  else if (dummy == 1.0) return True;
  else if (ISNAN(dummy)) return Nan;
  RFERROR2("invalid value (%d) for boolean variable '%.50s'.", (int) dummy, name);
}


SEXP String(char *V) {
  SEXP str;
  PROTECT(str = allocVector(STRSXP, 1)); 
  SET_STRING_ELT(str, 1, mkChar(V));
  UNPROTECT(1);
  return str;
}

SEXP String(char V[][MAXCHAR], int n, int max) {
  SEXP str;
  if (V==NULL) return allocVector(STRSXP, 0);
  if (n>max) return TooLarge(&n, 1);
  if (n<0) return TooSmall();
  PROTECT(str = allocVector(STRSXP, n)); 
  for (int i=0; i<n; i++) {
    SET_STRING_ELT(str, i, mkChar(V[i]));
  }
  UNPROTECT(1);
  return str;
}


SEXP String(int *V, const char * List[], int n, int endvalue) {
  SEXP str;
  if (V==NULL || n <= 0) return allocVector(STRSXP, 0);
  int k;
  for (k=0; k<n; k++) {
    if (V[k] == endvalue) break;
  }
  //  printf("k=%d, n=%d\n", k, n);
  PROTECT(str = allocVector(STRSXP, k)); 
  for (int i=0; i<k; i++) {
    //  printf("V[%d]=%d\n", i, V[i]);
    //    printf("%.50s\n", List[V[i]]);
    SET_STRING_ELT(str, i, mkChar(List[V[i]]));
  }
  UNPROTECT(1);
  return str;
}

double Real(SEXP p, char *name, int idx) {
  //  {printf("%.50s type=%d \n", name,TYPEOF(p));}
  if (p != R_NilValue) {
    assert(idx < length(p));
    switch (TYPEOF(p)) {
    case REALSXP :  return REAL(p)[idx];
    case INTSXP :
      if (INTEGER(p)[idx]==NA_INTEGER) return RF_NA;
      else return((double) INTEGER(p)[idx]);
    case LGLSXP :
      if (LOGICAL(p)[idx]==NA_LOGICAL) return(RF_NA);
      else return((double) LOGICAL(p)[idx]);
    default : {}
    }
  }

 RFERROR2("'%.50s' can not be transformed to double! (type=%d)\n", name, TYPEOF(p));  
  return RF_NA;  // to avoid warning from compiler
}


void Real(SEXP el,  char *name, double *vec, int maxn) {
  if (el == R_NilValue) {
    RFERROR1("'%.50s' cannot be transformed to double.\n", name);
  }
  int n = length(el);
  for (int j=0, i=0; i<maxn; i++) {
    vec[i] = Real(el, name, j);
    if (++j >= n) j=0;
  }
  return;
}


int Integer(SEXP p, char *name, int idx, bool nulltoNA) {
  if (p != R_NilValue) {
    assert(idx < length(p));
    // printf("typeof = %d %d idx=%d len=%d\n", TYPEOF(p), REALSXP, idx, length(p));
    switch(TYPEOF(p)) {
    case INTSXP : 
      return INTEGER(p)[idx]; 
    case REALSXP : 
      double value;
      value = REAL(p)[idx];
      if (ISNAN(value)) {
	return NA_INTEGER;
      }
      int intvalue;
      intvalue = (int) value;
      //      printf("%10g %d %10e\n ", value, intvalue, value-intvalue);
      if (value == intvalue) return intvalue;      
      else {
	RFERROR2("%.50s: integer value expected. Got %10e.", name, value);
      }
    case LGLSXP :
      if (LOGICAL(p)[idx]==NA_LOGICAL) return(NA_INTEGER);
      else return((int) LOGICAL(p)[idx]);
    default : {}
    }
  } else if (nulltoNA) return NA_INTEGER;
  RFERROR2("%.50s: unmatched type of parameter [type=%d]", name, TYPEOF(p));
  return NA_INTEGER; // compiler warning vermeiden
}

int Integer(SEXP p, char *name, int idx) {
  return Integer(p, name, idx, false);
}


void Integer(SEXP el, char *name, int *vec, int maxn) {
  if (el == R_NilValue) {
    RFERROR1("'%.50s' cannot be transformed to integer.\n",name);
  }
  int n = length(el);
  for (int j=0, i=0; i<maxn; i++) {
    vec[i] = Integer(el, name, j);
    if (++j >= n) j=0;
  }
}


void Integer2(SEXP el, char *name, int *vec) {
  int n;
  if (el == R_NilValue || (n = length(el))==0) {
      RFERROR1("'%.50s' cannot be transformed to integer.\n",name);
  }
 
  vec[0] = Integer(el, name, 0);
  if (vec[0] == NA_INTEGER || vec[0] < 1)
    RFERROR1("first component of '%.50s' must be at least 1", name);
  if (n==1) vec[1] = vec[0];
  else {
    vec[1] = Integer(el, name, n-1);    
    if ( vec[1] != NA_INTEGER && vec[1] < vec[0])
      RFERROR1("'%.50s' must be increasing", name);
    if (n > 2) {
      int v = vec[0] + 1;
      for (int i = 1; i<n; i++, v++)
	if (Integer(el, name, i) != v)
	  RFERROR1("'%.50s' is not a sequence of numbers",name); 

    }
  }
}


bool Logical(SEXP p, char *name, int idx) {
   if (p != R_NilValue)
    assert(idx < length(p));
    switch (TYPEOF(p)) {
    case REALSXP:
      if (ISNAN(REAL(p)[idx])) return NA_LOGICAL ;
      else return (bool) REAL(p)[idx];
    case INTSXP :
      if (INTEGER(p)[idx]==NA_INTEGER) return NA_LOGICAL;
      else return (bool) INTEGER(p)[idx];
    case LGLSXP : return LOGICAL(p)[idx];
    default : {}
    }
  RFERROR1("'%.50s' cannot be transformed to logical.\n", name);  
  return NA_LOGICAL;  // to avoid warning from compiler
}


char Char(SEXP el, char *name) {
  SEXPTYPE type;
  if (el == R_NilValue) goto ErrorHandling;
  type = TYPEOF(el);
  if (type == CHARSXP) return CHAR(el)[0];
  if (type == STRSXP) {
    if (length(el)==1) {
      if (STRLEN(CHAR(STRING_ELT(el,0))) == 1)
	return (CHAR(STRING_ELT(el,0)))[0];
      else if (STRLEN(CHAR(STRING_ELT(el,0))) == 0)
	return '\0';
    }
  }
 
 ErrorHandling:
  RFERROR1("'%.50s' cannot be transformed to character.\n",  name);  
  return 0; // to avoid warning from compiler
}


void String(SEXP el, char *name, char names[][MAXCHAR], int maxlen) {
  int l = length(el);
  SEXPTYPE type;  
  if (el == R_NilValue) goto ErrorHandling;
  if (l > maxlen)  {
    RFERROR1("number of variable names exceeds %d. Take abbreviations?", maxlen);
  }
  type = TYPEOF(el);
  //  printf("type=%d %d %d %d\n", TYPEOF(el), INTSXP, REALSXP, LGLSXP);
  if (type == CHARSXP) {
    for (int i=0; i<l; i++) {
      names[i][0] = CHAR(el)[i];
      names[i][1] = '\0';
    }
  } else if (type == STRSXP) {
    for (int i=0; i<l; i++) {
      //print("%d %d\n", i, l);
      strcopyN(names[i], CHAR(STRING_ELT(el, i)), MAXCHAR);
    }
  } else goto ErrorHandling;
  return;
 
 ErrorHandling:
  RFERROR1("'%.50s' cannot be transformed to character.\n",  name);  
}


double NonNegInteger(SEXP el, char *name) {
  int num;

  num = INT;
  if (num<0) {
    num=0; 
    WARN1("'%.50s', which has been negative, is set 0.\n",name);
  }
  return num;
}

double NonNegReal(SEXP el, char *name) {
  double num;
  num = NUM;
  if (num<0.0) {
    num=0.0; 
    WARN1("%.50s, which has been negative, is set 0.\n",name);
   }
  return num;
}

double NonPosReal(SEXP el, char *name) {
  double num;
  num = NUM;
  if (num>0.0) {
    num=0.0; 
    WARN1("%.50s, which has been positive, is set 0.\n",name);
  }
  return num;
}

double PositiveInteger(SEXP el, char *name) {
  int num;
  num = INT;
   if (num<=0) {
    WARN2("'%.50s', which has been %.50s, is set 1.\n",
	  name, num==0L ? "0" : "negative");
    num=1L;
  }
   return num;
}

double PositiveReal(SEXP el, char *name) {
  double num;
  num = NUM;
  if (num<=0.0) {
     WARN2("'%.50s', which has been %.50s, is set 1.\n",
	   name, num==0.0 ? "0" : "negative");
     num=1.0; 
   }
  return num;
}


SEXP ExtendedInteger(double x) {
  return ScalarInteger(R_FINITE(x) ? x : NA_INTEGER);
}

SEXP ExtendedBooleanUsr(usr_bool x) {
  return ScalarLogical((int) x);
}


int Match(char *name, name_type List, int n) {
  // == NOMATCHING, -1, if no matching function is found
  // == MULTIPLEMATCHING,-2, if multiple matching fctns are found,  
  // if more than one match exactly, the last one is taken (enables overwriting 
  // standard functions)
  unsigned int ln;
  int Nr;
  Nr=0;
  ln=STRLEN(name);
  //  print("Match %d %d %.50s %.50s %d\n", Nr, n, name, List[Nr], ln);

  while ( Nr < n  && STRNCMP(name, List[Nr], ln)) {
    Nr++;
  }
  if (Nr < n) { 
    if (ln==STRLEN(List[Nr])) // exactmatching -- take first -- changed 1/7/07
      return Nr;
    // a matching function is found. Are there other functions that match?
    int j; 
    bool multiplematching=false;
    j=Nr+1; // if two or more covariance functions have the same name 
    //            the last one is taken 
    while (j<n) {
      while ( (j<n) && STRNCMP(name, List[j], ln)) {j++;}
      if (j<n) {
	if (ln==STRLEN(List[j])) { // exactmatching -- take first 
	  return j;
	}
	else {multiplematching=true;}
      }
      j++;
    }
    if (multiplematching) {return MULTIPLEMATCHING;}
  } else return NOMATCHING;
  return Nr;
}

int Match(char *name, const char * List[], int n) {
  // printf("Matching\n");
   // == -1 if no matching name is found
  // == -2 if multiple matching names are found, without one matching exactly
  unsigned int ln;
  int Nr;
  Nr=0;
  ln=STRLEN(name);
  //    print("Matchx %d %d %.50s %.50s %d\n", Nr, n, name, List[Nr], ln);

  while ( Nr < n  && STRNCMP(name, List[Nr], ln)) {
    //     print("       %d %d %.50s %.50s %d\n", Nr, n, name, List[Nr], ln);
    //   printf("%.50s\n", List[Nr]);
    Nr++;
  }
  if (Nr < n) { 
    if (ln==STRLEN(List[Nr])) {// exactmatching -- take first -- changed 1/7/07
      //      print(" found  X    %d %d %.50s %.50s %d\n", Nr, n, name, List[Nr], ln);
      return Nr;
    }
    // a matching function is found. Are there other functions that match?
    int j; 
    bool multiplematching=false;
    j=Nr+1; // if two or more covariance functions have the same name 
    //            the last one is taken 
    while (j<n) {
      while ( (j<n) && STRNCMP(name, List[j], ln)) {j++;}
      if (j<n) {
	if (ln==STRLEN(List[j])) { // exactmatching -- take first 
	  return j;
	}
	else {multiplematching=true;}
      }
      j++;
    }
    if (multiplematching) {return MULTIPLEMATCHING;}
  } else return NOMATCHING;

  //    print(" found      %d %d %.50s %.50s %d\n", Nr, n, name, List[Nr], ln);
 
  return Nr;
}


void GetName(SEXP el, char *name, const char * List[], int n,
	     int defaultvalue, int endvalue, int *ans, int maxlen_ans) {
  char dummy[1000];
  int 
    k = 0,
    len_el = length(el);

  if (TYPEOF(el) == NILSXP) goto ErrorHandling;
  if (len_el > maxlen_ans) 
    RFERROR2("option '%.50s' is too lengthy. Maximum length is %d.", name, maxlen_ans);

  if (TYPEOF(el) == STRSXP) {    
    for (k=0; k<len_el; k++) {
      ans[k] = Match((char*) CHAR(STRING_ELT(el, k)), List, n);
      if (ans[k] < 0) {
	if (STRCMP((char*) CHAR(STRING_ELT(el, k)), " ") == 0 ||
	    STRCMP((char*) CHAR(STRING_ELT(el, k)), "") == 0) {
	  goto ErrorHandling;
	}
	goto ErrorHandling0;
      }
    }
    for (; k<maxlen_ans; k++) ans[k] = endvalue;
    return;
  }

ErrorHandling0:
  SPRINTF(dummy, "'%.50s': unknown value '%.50s'. Possible values are:", 
	  name, CHAR(STRING_ELT(el, k)));
  int i;
  for (i=0; i<n-1; i++) {
    char info[1000];
    SPRINTF(info, "%.50s '%.50s',", dummy, List[i]);    
    STRCPY(dummy, info);
  }
  RFERROR2("%.50s and '%.50s'.", dummy, List[i]);  
 
 ErrorHandling:
  if (defaultvalue >= 0) {
    ans[0] = defaultvalue;
    for (k=1; k<maxlen_ans; k++) ans[k] = endvalue;
    return;
  }
  
  RFERROR1("'%.50s': no value given.", name);
}

int GetName(SEXP el, char *name, const char * List[], int n,
	    int defaultvalue) {
  int i;
  GetName(el, name, List, n, defaultvalue, defaultvalue, &i, 1);
  return i;
}


int GetName(SEXP el, char *name, const char * List[], int n) {
 return GetName(el, name, List, n, -1);
}


double ownround(double x) { return TRUNC((x + SIGN(x) * 0.5)); }


double lonmod(double x, double modulus) {  
  double 
    halfmodulus = 0.5 * modulus,
    y = x + modulus + halfmodulus;
  return Mod(y, modulus) - halfmodulus;
}

/*


double intpow(double x, int p) {
  //int p0 = p;
  // double x0 = x;

  double res = 1.0;
  if (p < 0) {
    p = -p;
    x = 1.0 / x;
  } 
  while (p != 0) {
    //    printf("  ... %10e %d : %10e\n" , x, p, res);
  if (p % 2 == 1) res *= x;
    x *= x;
    p /= 2;
  }
  return res;
}


void distInt(int *X, int*N, int *Genes, double *dist) {
    int i,j, k, di, diff, *x, *y, ve, ho, endfor,
	n = *N,
	nP1 = n + 1,
	genes = *Genes;
 
  x = y = X;
  for (j=0, i=0;  j<n;  i += nP1, j++, y += genes) {
    dist[i] = 0.0;
    endfor = i + (n - j);
    for (ve = i + 1, ho = i + n, x = y + genes; 
         ve < endfor; 
	 ve++, ho += n) {
      for (di=0.0, k=0; k<genes; k++, x++) {
	diff = *x - y[k];
	di += diff * diff;
      }
      dist[ve] = dist[ho] = SQRT((double) di);
    }
  }
}


void vectordist(double *v, int *Dim, double *Dist, int *diag){
  int d, dim, dr;
  double *v1, *v2, *end;
  bool notdiag = (*diag==0);
  dim = Dim[0];
  end = v + Dim[1] * dim; 

//  print("%d %d %10g %10g\n", dim , Dim[0], v, end);

  for (dr=0, v1=v; v1<end; v1+=dim) { // loop is one to large??
    v2 = v1;
    if (notdiag) {
       v2 += dim;
    }
    for (; v2<end; ) {
      for (d=0; d<dim; v2++) {
	Dist[dr++] = v1[d++] - *v2;
      }
    }
  }
} 


int addressbits(void VARIABLE_IS_NOT_USED *addr) {
#ifndef RANDOMFIELDS_DEBUGGING  
  return 0;
#else
  double x = (Long) addr,
    cut = 1e9;
  x = x - TRUNC(x / cut) * cut;
  return (int) x;
#endif

}


 */
```
