## Supplementary material for "MoBPS - Modular Breeding Program Simulator": miraculix R-package: kleinkram.h

```
/*
 Authors 
 Martin Schlather,


 Copyright (C) 2015 -- 2017 Martin Schlather

This program is free software; you can redistribute it and/or
modify it under the terms of the GNU General Public License
as published by the Free Software Foundation; either version 3
of the License, or (at your option) any later version.

This program is distributed in the hope that it will be useful,
but WITHOUT ANY WARRANTY; without even the implied warranty of
MERCHANTABILITY or FITNESS FOR A PARTICULAR PURPOSE.  See the
GNU General Public License for more details.

You should have received a copy of the GNU General Public License
along with this program; if not, write to the Free Software
Foundation, Inc., 59 Temple Place - Suite 330, Boston, MA  02111-1307, USA.  
*/


#ifndef kleinkram_rfutils_h
#define kleinkram_rfutils_h 1

#include <R.h>
#include <Rinternals.h>
#include <Basic_utils.h> //#include <Basic_utils.h>

typedef char name_type[][MAXCHAR];

void strcopyN(char *dest, const char *src, int n);

usr_bool UsrBool(SEXP p, char *name, int idx);
usr_bool UsrBoolRelaxed(SEXP p, char *name, int idx);

#define INT Integer(el, name, 0)
#define LOGI Logical(el, name, 0)
#define NUM Real(el, name, 0)
#define USRLOG UsrBool(el, name, 0)
#define USRLOGRELAXED UsrBoolRelaxed(el, name, 0)
#define CHR Char(el, name)
#define STR(X, N)  strcopyN(X, CHAR(STRING_ELT(el, 0)), N);
#define POS0INT NonNegInteger(el, name) /* better: non-negative */
#define POS0NUM NonNegReal(el, name)
#define NEG0NUM NonPosReal(el, name)
#define POSINT PositiveInteger(el, name) /* better: non-negative */
#define POSNUM PositiveReal(el, name)


SEXP Logic(bool* V, int n, int max) ;
SEXP Num(double* V, int n, int max) ;
SEXP Int(int *V, int n, int max) ;
SEXP Char(const char **V, int n, int max) ;
SEXP Mat(double* V, int row, int col, int max);
SEXP Mat_t(double* V, int row, int col, int max);
SEXP MatInt(int* V, int row, int col, int max) ;
SEXP MatString(char **V, int row, int col, int max);
SEXP Array3D(int** V, int depth, int row, int col, int max) ;
SEXP String(char *V);

SEXP Logic(bool* V, int n) ;
SEXP Num(double* V, int n) ;
SEXP Int(int *V, int n) ;
SEXP Char(const char **V, int n) ;
SEXP Mat(double* V, int row, int col);
SEXP Mat_t(double* V, int row, int col);
SEXP MatInt(int* V, int row, int col) ;
SEXP MatString(char** V, int row, int col);
SEXP Array3D(int** V, int depth, int row, int col) ;
SEXP String(char V[][MAXCHAR], int n, int max);
SEXP String(int *V, const char * List[], int n, int endvalue);


SEXP TooLarge(int *n, int l);
SEXP TooSmall();


double Real(SEXP p, char *name, int idx);
void Real(SEXP el,  char *name, double *vec, int maxn) ;
int Integer(SEXP p, char *name, int idx, bool nulltoNA) ;
int Integer(SEXP p, char *name, int idx);
void Integer(SEXP el, char *name, int *vec, int maxn) ;
void Integer2(SEXP el, char *name, int *vec) ;
bool Logical(SEXP p, char *name, int idx);
char Char(SEXP el, char *name) ;
double NonNegInteger(SEXP el, char *name) ;
double NonNegReal(SEXP el, char *name) ;
double NonPosReal(SEXP el, char *name) ;
double PositiveInteger(SEXP el, char *name) ;
double PositiveReal(SEXP el, char *name) ;
void String(SEXP el, char *name, char names[][MAXCHAR], int maxlen);


#define MULTIPLEMATCHING -2
#define NOMATCHING -1
#define MATCHESINTERNAL -3
int Match(char *name, const char * List[], int n);
int Match(char *name, name_type List, int n);


SEXP ExtendedInteger(double x);
SEXP ExtendedBooleanUsr(usr_bool x);
 


double XkCXtl(double *X, double *C, int nrow, int dim, int k, int l);
void XCXt(double *X, double *C, double *V, int nrow, int dim);
void AtA(double *a, int nrow, int ncol, double *A);
void xA(double *x, double*A, int nrow, int ncol, double *y);
void xA_noomp(double *x, double*A, int nrow, int ncol, double *y);
void xA(double *x1, double *x2,  double*A, int nrow, int ncol, double *y1,
	double *y2);
void xAx(double *x, double*A, int nrow,  double *y);
void Ax(double *A, double*x, int nrow, int ncol, double *y);
void Ax(double *A, double*x1, double*x2, int nrow, int ncol, double *y1,
	double *y2);
double xUy(double *x, double *U, double *y, int dim);
double xUxz(double *x, double *U, int dim, double *z);
double x_UxPz(double *x, double *U, double *z, int dim);
double xUx(double *x, double *U, int dim);
void matmult(double *A, double *B, double *C, int l, int m, int n);
void matmulttransposed(double *A, double *B, double *C, int m, int l, int n);
//void matmulttransposedInt(int *A, int *B, int *c, int m, int l, int n); 
void matmult_2ndtransp(double *A, double *B, double *C, int m, int l, int n);
void matmult_2ndtransp(double *A, double *B, double *C, int m, int l);
void matmult_tt(double *A, double *B, double *C, int m, int l, int n);
double *matrixmult(double *m1, double *m2, int dim1, int dim2, int dim3);


void GetName(SEXP el, char *name, const char * List[], int n,
	     int defaultvalue, int endvalue, int *ans, int maxlen_ans);
int GetName(SEXP el, char *name, const char * List[], int n) ;
int GetName(SEXP el, char *name, const char * List[], int n,
	    int defaultvalue) ;


#define SCALAR_PROD(A, B, N, ANS) {			\
    int  k_ =0,				\
    end_ = N - 4;				\
  ANS = 0.0;					\
  for (; k_<end_; k_+=4) {				\
    ANS += A[k_] * B[k_]				\
      + A[k_ + 1] * B[k_ + 1]				\
      + A[k_ + 2] * B[k_ + 2]				\
      + A[k_ + 3] * B[k_ + 3];				\
  }							\
  for (; k_<N; k_++) ANS += A[k_] * B[k_];		\
  }


// unused :
#define FILL_IN(A, N, VALUE) {				\
    int end_ = N;					\
    for (int k_=0; k_<end_; (A)[k_++]=VALUE);		\
}

double scalar(double *A, double *B, int N);
double ownround(double x);

#define Mod(Z, modulus) ((Z) - FLOOR((Z) / (modulus)) * (modulus))
double lonmod(double x, double modulus); 

/*
extern "C" void vectordist(double *v, int *dim, double *dist, int *diag); 
bool is_diag(double *aniso, int dim);
*/ 

#endif
```
