## Supplementary material for "MoBPS - Modular Breeding Program Simulator": miraculix R-package: miraculix.h

```
/*
 Authors 
 Martin Schlather,


 Copyright (C) 2018 -- 2019  Martin Schlather

This program is free software; you can redistribute it and/or
modify it under the terms of the GNU General Public License
as published by the Free Software Foundation; either version 3
of the License, or (at your option) any later version.

This program is distributed in the hope that it will be useful,
but WITHOUT ANY WARRANTY; without even the implied warranty of
MERCHANTABILITY or FITNESS FOR A PARTICULAR PURPOSE.  See the
GNU General Public License for more details.

You should have received a copy of the GNU General Public License
along with this program; if not, write to the Free Software
Foundation, Inc., 59 Temple Place - Suite 330, Boston, MA  02111-1307, USA.  
*/


#ifndef miraculix_main_H
#define miraculix_main_H 1

//
#include <R.h>
#include <Rinternals.h>
#include <Rdefines.h>

#ifdef __cplusplus
extern "C" {
#endif 

 // general
  SEXP RFoptions(SEXP options);

  SEXP loadmiraculix();
  SEXP attachmiraculix();
  void detachmiraculix();
 
  // scan
  SEXP scan(SEXP positions, SEXP length, SEXP freq, 
	    SEXP minscan,  SEXP maxscan, SEXP threshold, SEXP nthres,
	    SEXP PER_SNP,
	    SEXP above_threshold, SEXP maximum);
  
  SEXP sumscan(SEXP positions, SEXP length, SEXP freq, 
	       SEXP minscan,  SEXP maxscan, SEXP threshold, SEXP nthres,
	       SEXP PER_SNP,
	       SEXP above_threshold, SEXP maximum);

  SEXP collect_scan(SEXP positions, SEXP length, SEXP freq, 
		    SEXP minscan,  SEXP maxscan, SEXP threshold, SEXP nthres,
		    SEXP PER_SNP, 
		    // additional return values:
		    SEXP areas,  SEXP value
		    );
  SEXP collect_scan2(SEXP positions, SEXP length, SEXP freq, 
		     SEXP minscan,  SEXP maxscan, SEXP threshold, SEXP nthres,
		     SEXP PER_SNP, SEXP max_intervals,
		     SEXP max_max_basepair_dist, SEXP exclude_negative,
		     // additional return values:
		     SEXP above_threshold, SEXP maximum		     
		     );


  // windower
  SEXP windower(SEXP Init, SEXP, SEXP Length, SEXP Step, SEXP start, SEXP ende,
		     SEXP data, SEXP Lendata, SEXP N);


  // Verwandtschaft
  SEXP vector012matrix(SEXP vector, SEXP matrix);
  SEXP matrixvector012(SEXP vector, SEXP matrix);

  SEXP getAutoCoding();
  SEXP hasSSE2();
  SEXP hasSSSE3();
  SEXP hasAVX2();
  SEXP hasAVX();
  
  SEXP matrix_coding(SEXP M) ;  //
  SEXP matrix_get(SEXP M);  
  SEXP matrix_mult(SEXP M);

  SEXP file_get(SEXP file);
  SEXP file_dot(SEXP file, SEXP g);
  SEXP dot_file(SEXP file, SEXP g);
  
  SEXP codeOrigins(SEXP M);
  SEXP decodeOrigins(SEXP M, SEXP line);

  // shot table coding
  SEXP fillSNPmatrix(SEXP Z, SEXP Idx, SEXP V) ;
  SEXP createSNPmatrix(SEXP SNPSs, SEXP Individuals) ;  
  SEXP get_matrix_N(SEXP CM, SEXP NN);
  SEXP decodeHaplo(SEXP CM, SEXP, SEXP, SEXP, SEXP) ;
  SEXP haplo2geno(SEXP Haplo);
  SEXP zeroNthGeno(SEXP CM, SEXP NN);
  SEXP copyGeno(SEXP CM);
  SEXP codeHaplo(SEXP M, SEXP IndivPerCol, SEXP DoubledIndiv) ;// 2 x Haplo - Matrix, as vector gespeichert
  //  SEXP codeHaplo2(SEXP M1, SEXP M2, SEXP);
  SEXP vectorGeno(SEXP V, SEXP Z);
  SEXP genoVector(SEXP Z, SEXP V);
  SEXP unlock(SEXP X);
  SEXP dolocking(SEXP Do);
  SEXP computeSNPS(SEXP population, SEXP Generation, SEXP Sex, SEXP Nr,
		   SEXP From_SNP, SEXP To_SNP, SEXP Select, SEXP Geno);
  SEXP allele_freq(SEXP CGM);
  SEXP Debug();
  SEXP StopDebug();
  SEXP compute(SEXP population, SEXP Generation, SEXP Sex, SEXP Nr,
	       SEXP tau, SEXP vec, SEXP b, SEXP Select, SEXP matrix_return);
  SEXP solveRelMat(SEXP R, SEXP tau, SEXP vec, SEXP b, SEXP destroy);
  SEXP substract_centered(SEXP SnpXindiv);
  SEXP get_centered();
  SEXP rhaplomatrix(SEXP freq, SEXP freq2,  SEXP indiv);
  SEXP rhaplomatrixPart2(SEXP freq, SEXP freq2, SEXP indiv, SEXP Code);
  

#ifdef __cplusplus
}
#endif

#endif /* RF_simu_PUBLIC_H*/
```
