## Supplementary material for "MoBPS - Modular Breeding Program Simulator": miraculix R-package: MoBPS.cc

```
/*
 Authors 
 Martin Schlather,


 Copyright (C) 2018 -- 2019  Martin Schlather

This program is free software; you can redistribute it and/or
modify it under the terms of the GNU General Public License
as published by the Free Software Foundation; either version 3
of the License, or (at your option) any later version.

This program is distributed in the hope that it will be useful,
but WITHOUT ANY WARRANTY; without even the implied warranty of
MERCHANTABILITY or FITNESS FOR A PARTICULAR PURPOSE.  See the
GNU General Public License for more details.

You should have received a copy of the GNU General Public License
along with this program; if not, write to the Free Software
Foundation, Inc., 59 Temple Place - Suite 330, Boston, MA  02111-1307, USA.  
*/


#define BitsPerCode 2L
#define PseudoSSE 1 // fallback option

#include "IntrinsicsBase.h"

#include "intrinsics.h"
#include "miraculix.h"
#include <General_utils.h>

#include <inttypes.h> 
#include "Vector.matrix.h"
#include "AutoMiraculix.h"
#include "options.h"
#include "Haplo.h"
#include "haplogeno.h"
#include "xport_import.h"
#include "shuffle.h"

//#define DO_FLOAT 1  // must be completely re-checked if defined

// #define DO_PARALLEL 1 // done by basic_utils.h

// #define INLINE
#define INLINE static inline


Uint zaehler = 0;

//#define show printf
//
#define show(X)
// void show(const char *s) {  if (debugging) PRINTF("%.50s", s); }
				   
INLINER

BlockType BitMaskStart[CodesPerBlock], BitMaskEnd[CodesPerBlock];
bool MoBPSNotInit = true;
void InitMoBPS() {
  if (!MoBPSNotInit) BUG;
  MoBPSNotInit = false;
  snpcoding method = (snpcoding) GLOBAL.genetics.method;
  if (method != Shuffle && method != Shuffle256) {
#if defined SSE2
    PRINTF("MoBPS will run much faster and with a minimum of memory if 'Shuffle' or 'Shuffle256' is used as 'snpcoding'\n");
#else
    if (method != TwoBit) {
      GLOBAL.genetics.method = method = TwoBit;
      PRINTF("The default value of 'RFoption(snpcoding=...)' in miraculix, which is 'ThreeBit' or 'Hamming2' in case SSSE3 is not available, is overwritten by the default value of 'MoBPS', which is 'TwoBit'.\nNote that 'MoBPS' will run much faster if 'miraculix' is compiled with SSSE3.\n");
    } else {
      PRINTF("MoBPS will run with a minimum of memory if 'TwoBit' is used as 'snpcoding'\n");
    }
#endif
  }
  
  assert(BytesPerBlock == sizeof(BlockType0));
 
  BlockType Einsen;
  SET32(Einsen, 0xFFFFFFFF);
  BlockUnitType *BME = (BlockUnitType*) BitMaskEnd;
  BME->u32[0] = 0x03;

  for (Uint k=1; k<CodesPerBlock; k++) {
    XOR(BitMaskStart[k], BitMaskEnd[k-1], Einsen); // not
    BitMaskEnd[k] = BitMaskEnd[k-1];
    BME = (BlockUnitType*) BitMaskEnd + k;
    BME->u32[k % UnitsPerBlock] |= 0x03 << (BitsPerCode * (k / UnitsPerBlock));
  }
  BitMaskStart[0] = Einsen;
}


void assert_MoBPS() {
  if (MoBPSNotInit) InitMoBPS();
}


#define ORIGIN_GENERATION 0
#define ORIGIN_SEX 1
#define ORIGIN_NR 2
#define ORIGIN_HAPLO 3

#define BITS_GENE_INPUT 6
#define BITS_SEX 1
#define BITS_INDIVIDUALS 22
#define BITS_HAPLO 3

#define MAX_GENE_INPUT (1 << BITS_GENE_INPUT)
#define MAX_SEX (1 << BITS_SEX)
#define MAX_INDIVIDUALS (1 << BITS_INDIVIDUALS)
#define MAX_HAPLO (1 << BITS_HAPLO)

#define PATTERN_GENE_INPUT  (MAX_GENE_INPUT - 1)
#define PATTERN_SEX (MAX_SEX - 1)
#define PATTERN_INDIVIDUALS (MAX_INDIVIDUALS - 1)
#define PATTERN_HAPLO (MAX_HAPLO - 1)


void INLINE decodeOrigin(Uint x, Uint *ans) {
  ans[ORIGIN_HAPLO] = (x & PATTERN_HAPLO);
  x >>= BITS_HAPLO;
  ans[ORIGIN_NR] = (x & PATTERN_INDIVIDUALS);
  x >>= BITS_INDIVIDUALS;  
  ans[ORIGIN_SEX] = (x & PATTERN_SEX);
  x >>= BITS_SEX;
  ans[ORIGIN_GENERATION] = x;
}


SEXP decodeOrigins(SEXP M, SEXP Line) {  
  assert_MoBPS();
  Uint line = Int0(Line);
  assert(line > 0); // R index
  SEXP Ans;
  PROTECT(Ans = allocVector(INTSXP, 4));
  Uint *ans = (Uint*) INTEGER(Ans);
  decodeOrigin(Inti(M, line - 1), ans);
  for (Uint i=0; i<4; ans[i++]++);
  UNPROTECT(1);
  if (PRESERVE) R_PreserveObject(Ans);
  return Ans;
}


SEXP codeOrigins(SEXP M) {
  assert_MoBPS();
  Uint nrow = nrows(M);
  assert(ncols(M) == 4);
  
  SEXP Ans;
  PROTECT(Ans = allocVector(INTSXP, nrow));
 
  Uint *ans = (Uint*) INTEGER(Ans);
  for (Uint i=0; i<nrow; i++) ans[i] = 0;  // unnoetig?!

#define DO1(TYPE, RTYPE)			\
  TYPE *gen = RTYPE(M),				\
    *sex = gen + nrow,				\
    *nr = sex + nrow,				\
    *haplo = nr + nrow					       
    
#define DO2								\
  for (Uint i=0; i<nrow; i++) {						\
    Uint g = (Uint) gen[i],						\
      s = (Uint) sex[i],						\
      n = (Uint) nr[i],							\
      H = (Uint) haplo[i];						\
    if (g < 1 || g > MAX_GENE_INPUT|| s < 1 || s > MAX_SEX ||		\
	n < 1 || n > MAX_INDIVIDUALS ||	H < 1 || H > MAX_HAPLO)		\
      ERR1("some value in row %d out of bound", i + 1);			\
    ans[i] = ((((((g - 1) << BITS_SEX) +				\
		 s - 1) << BITS_INDIVIDUALS) +				\
	       n - 1) << BITS_HAPLO) +					\
      H - 1;								\
      									\
    Uint control[4];							\
    decodeOrigin(ans[i], control);					\
    if (control[0]+1 != g || control[1]+1 != s || control[2]+1 != n ||	\
	control[3]+1 != H) {						\
      /*	p rintf("i=%d: generation:%d->%d sex:%d->%d nr:%d->%d haplo:%d->%d\n", i, g, control[0], s, control[1], n, control[2], H, control[3]); */ \
      BUG;								\
    }									\
  }

  
  if (TYPEOF(M) == REALSXP) {
    DO1(double, REAL);
#ifdef DO_PARALLEL
#pragma omp parallel for num_threads(CORES)   
#endif
    DO2   
      } else if (TYPEOF(M) == INTSXP) {
    DO1(int, INTEGER);
#ifdef DO_PARALLEL
#pragma omp parallel for num_threads(CORES)  
#endif
    DO2   
      } else BUG;
  
  UNPROTECT(1);
  if (PRESERVE) R_PreserveObject(Ans);
  return Ans;
}


// population
#define INFO 0
#define BREEDING 1

// info
#define INFO_DUMMY 0
#define NSNPS 2
#define POSITION 3
#define SNP_POSITION 5
#define LENGTH_CHROM 6
#define LENGTH_TOTAL 7
#define SNPS_EQUIDIST 17
#define GENERATION_REFTABLE 18

// individual
#define RECOMBI1 0
#define RECOMBI2 1
#define MUTAT1 2
#define MUTAT2 3
#define ORIGIN1 4
#define ORIGIN2 5
//#define FATHER 6
//#define MOTHER 7
#define HAPLOTYP1 8 // in case of founder, otherwise empty
#define HAPLOTYP2 9
#define CODEDHAPLO HAPLOTYP1

#define MAX_CHROM 100


Uint INLINE GetGenoCode(BlockType0 *code, Uint pos) {
  Uint block = pos / CodesPerBlock,
    unit = pos % UnitsPerBlock,
    elmt = (pos % CodesPerBlock) / UnitsPerBlock;
  BlockUnitType *c = (BlockUnitType*) code + block;
  return (c[0].u32[unit] >> (2 * elmt)) & 0x00000003;
}


void INLINE PutGenoCode(Uint value, Uint pos, BlockType0 *code) {
  Uint 
    unit = pos % UnitsPerBlock,
    elmt = (pos % CodesPerBlock) / UnitsPerBlock,
    shift = 2 * elmt;
  BlockUnitType *c = ((BlockUnitType*) code) + pos / CodesPerBlock;

  c[0].u32[unit] =
    (c[0].u32[unit] & (0xFFFFFFFF xor (0x00000003 << shift)))//clear
    | (value << shift);
}


Uint INLINE GetHaploCode(BlockType0 *code, Uint pos, Uint haplo) {
  Uint unit = pos % UnitsPerBlock,
    elmt = (pos % CodesPerBlock) / UnitsPerBlock; 
  BlockUnitType *c = (BlockUnitType*) code + pos / CodesPerBlock; 
  assert(haplo <= 1);
  assert(elmt < CodesPerUnit);
  return (c[0].u32[unit] >> (2 * elmt + haplo)) & 0x00000001;
}


void INLINE PutHaploCode(Uint value, Uint pos, Uint haplo, BlockType0 *code) {
  Uint 
    unit = pos % UnitsPerBlock,
    elmt = (pos % CodesPerBlock) / UnitsPerBlock,
    shift = 2 * elmt + haplo;
  BlockUnitType *c = (BlockUnitType*) code + pos / CodesPerBlock; 
  if (value) c[0].u32[unit] |= (0x00000001 << shift);
  else c[0].u32[unit] &= 0xFFFFFFFF xor (0x00000001 << shift);
}


void INLINE PutHaploOne(Uint pos, Uint haplo, BlockType0 *code) {
  Uint 
    unit = pos % UnitsPerBlock,
    elmt = (pos % CodesPerBlock) / UnitsPerBlock,
    shift = 2 * elmt + haplo;
  BlockUnitType *c = (BlockUnitType*) code + pos / CodesPerBlock; 
  c[0].u32[unit] |= (0x00000001 << shift);
}


  
Uint getNrSNPposition(double cur_recomb, double *position, Uint Min, Uint Max) {
  Uint min = Min,
    max = Max;
  //  print("%d %d\n", min, max);
  while (min < max) {
    Uint inbetween = (min + max + 1) / 2;
    //printf("min=%d,max=%d %10g cr=%10g, pm=%10g\n", min, max,position[inbetween],
    //	   cur_recomb, position[max] );
    if (position[inbetween] <= cur_recomb) min = inbetween;
    if (position[inbetween] > cur_recomb) max = inbetween - 1;
  }
  if (min > max) BUG;
  //  printf("%10g %10g min=%d max=%d (start:%d %d) %10g %10g %10e\n", position[max], cur_recomb, min, max, Min, Max, position[Max-1], position[Max], cur_recomb -  position[Max]);
  if (position[max] > cur_recomb) {
    if (max == Min) return max - 1; // links des ersten Markers auf dem
    //                                 Chromosom; somit wird letzter marker 
    //                                 vom Vor-Chromoosom zugeordnet
    BUG;
  }
  if (max < Max && position[max+1] <= cur_recomb) BUG;
  return max;
}


#define ASSERT(WHERE,WHAT,NAME)						\
  assert(STRCMP(CHAR(STRING_ELT(getAttrib(WHERE, R_NamesSymbol), WHAT)), \
		NAME) == 0);
int IcomputeSNPS(SEXP population, SEXP Generation, SEXP Sex, SEXP Nr,
		 Rint From_SNP, Rint To_SNP, SEXP Select, BlockType0 *ans){
  
  //  zaehler ++; printf("%d\n", zaehler);
  Uint
    cur_select = NA_INTEGER,
    individuals = length(Generation),
    len_sel = length(Select);
  bool do_select = len_sel > 0;
  if (individuals != (Uint) length(Sex) || individuals != (Uint) length(Nr))
    ERR("'gen', 'gender' and 'nr' do not have same length");
  if (From_SNP <= 0) ERR("'from_p' not valid.");

  CondToInt(do_select, Select);
   
  ASSERT(population, BREEDING, "breeding");
  SEXP breeding = VECTOR_ELT(population, BREEDING);

  ASSERT(population, INFO, "info")
    SEXP info = VECTOR_ELT(population, INFO);
  Uint len_info = length(info);

  static Uint gen_ref[MAX_GENE_INPUT] = { 0 };
  if (GENERATION_REFTABLE >= len_info ||
      length(VECTOR_ELT(info, GENERATION_REFTABLE)) == 0) {
    if (gen_ref[MAX_GENE_INPUT - 1] == 0) { // should never happen except uninit
      for (Uint i=0; i<MAX_GENE_INPUT; i++) gen_ref[i] = i;
      PRINTF("Generation reference table now initialized.\n"); 
    }
  } else {
    ASSERT(info, GENERATION_REFTABLE, "origin.gen");
    Uint *p = (Uint*) INTEGER(VECTOR_ELT(info, GENERATION_REFTABLE)); 
    for (Uint i=0; i<MAX_GENE_INPUT; i++) gen_ref[i] = (Uint) (p[i] - 1);
  }

  ASSERT(info, LENGTH_TOTAL, "length.total");
  Uint n_chrom = length(VECTOR_ELT(info, LENGTH_TOTAL)) - 1;
  if (n_chrom >= MAX_CHROM) ERR1("maximum chromosoms is %d", MAX_CHROM);
  double *length_total = REAL(VECTOR_ELT(info, LENGTH_TOTAL));
  
  ASSERT(info, LENGTH_CHROM, "length");
  double *LengthChrom = REAL(VECTOR_ELT(info, LENGTH_CHROM));
  
  ASSERT(info, NSNPS, "snp");
  Uint n_snps;
  SEXP Dummy = VECTOR_ELT(info, INFO_DUMMY);

  //   if (zaehler >= 11424) printf("AB\n");
#ifdef DOCH_AUF_INFO_ABSPEICHERN  
  if (TYPEOF(Dummy) != INTSXP || length(Dummy) == 0){
    Uint mem = n_chromP1 + n_chromP1;
    SEXP dummy;
    PROTECT(dummy = allocVector(INTSXP, mem));    
    Uint *snps_perchrom = (Uint*) INTEGER(VECTOR_ELT(info, NSNPS)),
      *length_prec_chroms = (Uint*) INTEGER(dummy) + 0,
      *totalsnps_prec_chroms = (Uint*) INTEGER(dummy) + n_chromP1;
    length_prec_chroms[0] = 0;
    totalsnps_prec_chroms[0] = 0;
    for (Uint c=1; c<=n_chrom; c++) {
      length_prec_chroms[c] =  length_prec_chroms[c-1] + LengthChrom[c-1];
      totalsnps_prec_chroms[c] = totalsnps_prec_chroms[c-1] +
	snps_perchrom[c-1];
    }
    SET_VECTOR_ELT(info, INFO_DUMMY, dummy);
    UNPROTECT(1);
  }
  Uint
    *length_prec_chroms = (Uint*) INTEGER(VECTOR_ELT(info, INFO_DUMMY)) + 0,
    *totalsnps_prec_chroms = length_prec_chroms + n_chromP1;
#else
  if (TYPEOF(Dummy) != LGLSXP) {
    SEXP dummy;
    PROTECT(dummy = allocVector(LGLSXP, 1));
    LOGICAL(dummy)[0] = true;
    SET_VECTOR_ELT(info, INFO_DUMMY, dummy);
    UNPROTECT(1);  
  }
   
  double length_prec_chroms[MAX_CHROM + 1];
  SEXP snps_perchrom = VECTOR_ELT(info, NSNPS);
  ToInt(snps_perchrom);
  Uint
    totalsnps_prec_chroms[MAX_CHROM + 1];
  length_prec_chroms[0] = 0;
  totalsnps_prec_chroms[0] = 0;
  for (Uint c=1; c<=n_chrom; c++) {
    length_prec_chroms[c] =  length_prec_chroms[c-1] + LengthChrom[c-1];
    totalsnps_prec_chroms[c] = totalsnps_prec_chroms[c-1] +
      snps_perchromint[c-1];
  }
#endif

  // if (zaehler >= 11424)  printf("ABC\n");
  n_snps = totalsnps_prec_chroms[n_chrom];
  assert(n_snps > 0);
  Rint fromSNP = From_SNP - 1,
    toSNP = To_SNP==NA_INTEGER || To_SNP >= (int) n_snps ? MAXINT : To_SNP -1;
  if (toSNP < fromSNP) ERR("'to_p' smaller than 'from_p'");
  if (fromSNP % CodesPerBlock != 0)
    ERR1("(from_p-1) must be devisible by %d", CodesPerBlock);
  if (toSNP < MAXINT && (toSNP + 1)  % CodesPerBlock != 0)
    ERR1("'to_p' must be devisible by %d or 'NA'", CodesPerBlock);


  Uint cur_nsnps = 0;
  if (do_select) {
    cur_select = Selectint[0] - 1;
    Rint toSNP_P1 = toSNP + 1;
    for (Uint k=0; k<len_sel; k++)
      cur_nsnps +=
	Selectint[k] > (Uint) fromSNP && Selectint[k] <= (Uint) toSNP_P1;
  } else cur_nsnps = 1L + MIN(n_snps - 1L, (Uint) toSNP) - fromSNP;
  if (ans == NULL) return cur_nsnps;
  Uint blocks = Blocks(cur_nsnps);

  ASSERT(info, SNPS_EQUIDIST, "snps.equidistant");
  bool equidist = (bool) LOGICAL(VECTOR_ELT(info, SNPS_EQUIDIST))[0];
  
  ASSERT(info, POSITION, "position");
  SEXP position = VECTOR_ELT(info, POSITION);

  ASSERT(info, SNP_POSITION, "snp.position");
  double *snp_positions = REAL(VECTOR_ELT(info, SNP_POSITION)),
    last_snp_position = snp_positions[n_snps - 1],
    first_snp_position = snp_positions[0];

  ToInt(Generation);
  ToInt(Sex);
  ToInt(Nr);

  //  if (zaehler >= 11424)  printf("ABD\n");
  for (Uint i=0; i<individuals; i++) {
    //    print("i=%d %d\n", i, individuals);
    //    printf("%d %d\n", i, individuals);
    if (length(breeding) < (int) Generationint[i]) ERR("value of 'generation' too large.");
    SEXP
      g = VECTOR_ELT(breeding, Generationint[i]-1);
    if (length(g) < (int) Sexint[i]) ERR("'sex' out of range.");
    SEXP 
      s = VECTOR_ELT(g, (int) Sexint[i]-1);
    if (length(s) < (int) Nrint[i]) ERR("value of 'nr' too large..");  
    SEXP
      Indiv = VECTOR_ELT(s, Nrint[i]-1);
    BlockType *haplo = ans + i * blocks,
      NullEins[2];
    SET32(NullEins[0], 0x55555555);
    SET32(NullEins[1], 0xAAAAAAAA);
    for (Uint hapnr=0; hapnr<=1; hapnr++) {
      //printf("h=%d \n",  hapnr);
      //      if (zaehler >= 11424) printf("haplo=%u\n", hapnr);      
      if (TYPEOF(VECTOR_ELT(Indiv, RECOMBI1 + hapnr)) != REALSXP)
	ERR("real valued vector for recombination information expected");
      double *recombi = REAL(VECTOR_ELT(Indiv, RECOMBI1 + hapnr));
      Uint
	n_recomb = length(VECTOR_ELT(Indiv, RECOMBI1 + hapnr)) - 1;
      SEXP origin = VECTOR_ELT(Indiv, ORIGIN1 + hapnr);
      ToInt(origin);
      Uint
	n_mutat = length(VECTOR_ELT(Indiv, MUTAT1 + hapnr)),
	// separation of the following 3 allows for selection but also
	// for mutation error such as insertion and deletion
	p = 0, // position what is written if there is no selection
	q = 0, // position what is written actually
	psel = 0, pmutat = 0,
	till = 0,
	from = 0; // current recombination starting point
      bool mutations = n_mutat > 0;
      
      SEXP mutat = mutations ? VECTOR_ELT(Indiv, MUTAT1 + hapnr) : NULL;
      CondToInt(mutations, mutat);
      
      for (Uint index=0; index <= n_recomb; index++) {
	// printf("n_rec=%d %d\n", n_recomb, index);
	
	double cur_recomb = recombi[index];
	Uint cur_chrom; // starting with number 0

	//	if (zaehler >= 11424) printf("index=%d %d\n", index, n_recomb);

	if (cur_recomb <= first_snp_position) {
	  from = 0;
	  continue;
	}
	
	for (cur_chrom=0; cur_chrom<n_chrom; cur_chrom++)
	  if (length_total[cur_chrom + 1] >= cur_recomb) break;
	if (equidist) {
	  if (cur_recomb > last_snp_position) till = n_snps - 1;
	  else {
	    double halfbetween = REAL(VECTOR_ELT(position, cur_chrom))[0];
	    till = totalsnps_prec_chroms[cur_chrom] +
	      CEIL((cur_recomb - length_prec_chroms[cur_chrom] - halfbetween) /
		   (2.0 * halfbetween)) - 1;
	  }
	} else {

	  //	  printf("cur_chrom =%d\n", cur_chrom);
	  Uint min = cur_chrom == 0 ? 0 : totalsnps_prec_chroms[cur_chrom],
	    max = cur_chrom == n_chrom ? n_chrom - 1
	    : totalsnps_prec_chroms[cur_chrom + 1] - 1;
	  till = getNrSNPposition(cur_recomb, snp_positions, min, max);
	  // printf("%d %d %d last=%d till=%d\n", cur_recomb, min, max, last_snp_position, till);
	}

	if (index == 0) {
	  from = till + 1;
	  assert(from == 0);
	  continue;
	}
	
	assert(toSNP != NA_INTEGER && toSNP <= MAXINT);
	       //printf("%d %d %d %d\n", till, from, toSNP, fromSNP);
	if (till >= from && from <= (Uint) toSNP && till >= (Uint) fromSNP) {
	  Uint von = MAX((Uint) fromSNP, from),
	    bis = MIN((Uint) toSNP, till),
	    o_info[4],
	    n_protect = 0;
 	  decodeOrigin(originint[index-1], o_info);
	  //
	  assert(bis < n_snps);
	  assert(von <= bis);

	  //  SEXP Orig_gen = VECTOR_ELT(breeding, o_info[ORIGIN_GENERATION]), 
	  SEXP Orig_gen = VECTOR_ELT(breeding,
				     gen_ref[o_info[ORIGIN_GENERATION]]), 
	    O2 = VECTOR_ELT(Orig_gen, o_info[ORIGIN_SEX]),
	    Orig = VECTOR_ELT(O2, o_info[ORIGIN_NR]);
	  Uint o_haplotype = o_info[ORIGIN_HAPLO];
	  SEXP coded_haplo = VECTOR_ELT(Orig, CODEDHAPLO);
	  //	  if (n_snps <= CodesPerBlock*3)//Coding selbst + hinten & vorne Platz
	  //   ERR2("number of snps is %d, but should be more than %d so that the programme can be used.", n_snps, CodesPerBlock * 3);
	  if ((Uint) length(coded_haplo) == n_snps) {
	    PROTECT(coded_haplo = codeHaplo2(VECTOR_ELT(Orig, HAPLOTYP1),
					     VECTOR_ELT(Orig, HAPLOTYP2)));
	    SET_VECTOR_ELT(Orig, CODEDHAPLO, coded_haplo);
	  }
 
	  //if (zaehler >= 11424) printf("ok\n");
	  BlockType
	    *o_haplo = (BlockType0*) DoAlign(coded_haplo, ALIGN_CODED, Haplo);
	 
	  //	  if (zaehler >= 11424 || zaehler >= 7220) printf("ok %d!\n", zaehler);
	  //	  printf("her rre\n");


	  for (Uint op=von; op<=bis; ) {
	    //
	    
	    //  printf("haplo=%d index=%d von=%d %d %d\n", hapnr, index, von, bis, op);
	    // printf("%d %d %.50s\n", (op % CodesPerBlock), (q % CodesPerBlock),  do_select);
	    if ( (op % CodesPerBlock) == (q % CodesPerBlock) && !do_select) {
	      // if (zaehler >= 11000) printf(".");
	      // blockwise copying	      
	      Uint 
		bnr = q / CodesPerBlock, 
		start = q % CodesPerBlock,
		o_rest = 1 + bis - op,
		rest = CodesPerBlock - start;
	      if (rest > o_rest) rest = o_rest;
	      BlockType L1,
		o_code = o_haplo[op / CodesPerBlock];
	      ////	      printf("rest = %d\n", rest);

	      //  if (zaehler >= 11424)   printf("a\n");
	      
	      if (mutations) {
		//if (zaehler >= 11424) printf("mutat = %d\n", n_mutat);
		Uint end = op + rest; // excluded, R-Coding !!
		if (mutatint[pmutat] <= end) { // mutat hat R codierung !!
		  // insbesondere bei select koennen Mutation uebersprungen
		  // werden. Somit muss zunaechst pmutat gegebenfalls erhoeht
		  // werden:
		  while (pmutat < n_mutat && mutatint[pmutat] <= op) pmutat++;
		  while (pmutat < n_mutat && mutatint[pmutat] <= end) {
		    //		    printf("%d %d\n", pmutat, n_mutat);
		    Uint idx = (mutatint[pmutat] - 1) % CodesPerBlock;
		    AND(L1 , BitMaskStart[idx], BitMaskEnd[idx]);
		    XOR(o_code, o_code, L1);
		    pmutat++;
		  }
		  mutations = pmutat < n_mutat;
		}
	      }
	      //  if (zaehler >= 11424) printf("xxxx\n");
	      AND(L1, BitMaskStart[start], BitMaskEnd[start + rest - 1]);
	      AND(L1, L1, NullEins[o_haplotype]);
	      AND(o_code, o_code, L1);

	      if (o_haplotype != hapnr) {
		if (o_haplotype > hapnr) { SHR32(o_code, o_code, 1); }
		else SHL32(o_code, o_code, 1);
	      }
	      
	      //   if (zaehler >= 11424) printf("fast ende %d\n", zaehler);
	      OR(haplo[bnr], haplo[bnr], o_code);
	      // printf("%d %ld\n", bnr, haplo[bnr]);
	      //	      showblock((Uint*) haplo + bnr, 1);
	      //  if (zaehler >= 11424) printf("ende\n");
	      op += rest;
	      p += rest; // p is now endpoint + 1 !
	      q += rest;
	    } else {
	      BUG;
	      // if (op % 1000 == 0) printf(";");
	      if (false)	      
		printf("von %u %u %ld %ld z=%d %d %d %d %d\n",//
		       von, bis, 
		       op % CodesPerBlock, q % CodesPerBlock, zaehler,
		       from, till, index, n_recomb);
	      // BUG;
	      // bitwise copying
	      if (do_select) {
		if (psel >= len_sel || cur_select != p) {
		  op++;
		  p++;
		  continue;
		}
		cur_select = Selectint[psel++];
	      }
	      Uint H = GetHaploCode((BlockType0 *) o_haplo, op, o_haplotype);
	      if (mutations) {
		// note mutat:R coding; op:C-Coding
		while (pmutat < n_mutat && mutatint[pmutat] <= op) pmutat++;
		if (mutatint[pmutat] == op + 1) {
		  H = 1- H;
		  pmutat++;
		}
		mutations = pmutat < n_mutat;
	      }
	      if (H) PutHaploOne(q, hapnr, haplo);
	      //printf("H = %d\n", H);
	      op++;
	      p++;
	      q++;
	    }
	  }// for loop through all snps between two recombination points
	  if (n_protect > 0) UNPROTECT(n_protect);
	} // if space between two recombination points is not empty
	from = till + 1;
      } // for n_recomb
      FREEint(mutat);
      FREEint(origin);
    } // hapnr 
  }

  FREEint(snps_perchrom);
  FREEint(Select);
  FREEint(Generation);
  FREEint(Sex); 
  FREEint(Nr); 

  
  return 0;
}

    
SEXP computeSNPS(SEXP population, SEXP Generation, SEXP Sex, SEXP Nr,
		 SEXP From_SNP, SEXP To_SNP, SEXP Select, SEXP Geno){
  // printf("A\n");
  assert_MoBPS();
  snpcoding method = (snpcoding) GLOBAL.genetics.method;
  bool geno = LOGICAL(Geno)[0];
  Uint individuals = length(Generation);
  //  printf("B\n");
  Uint snps = IcomputeSNPS(population, Generation, Sex, Nr,
			   INTEGER(From_SNP)[0], INTEGER(To_SNP)[0],
			   Select, NULL);
  // blocks = Blocks(snps);
  //unitsPerIndiv = blocks * UnitsPerBlock;
  //  printf("C\n");
  SEXP Ans;
  PROTECT(Ans = createSNPmatrix(snps, individuals, Haplo));// Shuffle==Haplo braucht
  //                einen Tick mehr Speicher als TwoBit, da letzteres auf 64 Bit
  //                laeuft und Shuffle/Haplo auf 128 Bit.
  
  //   printf("D\n");
  
 Uint *ans = (Uint*) DoAlign(Ans, ALIGN_COMPUTE, Haplo); 
 // printf("E\n");
  IcomputeSNPS(population, Generation, Sex, Nr, INTEGER(From_SNP)[0],
	       INTEGER(To_SNP)[0], Select, (BlockType0*) ans);
  //  printf("F m=%d\n", method);
 
  if (geno) {
    //  printf("geno\n");
   if ( (method == Shuffle || method == Shuffle256)
	  &&  GetCodesPerBlock(Haplo) == GetCodesPerBlock(method)
	  ){
     //printf("ok %d %d \n", GetCodesPerBlock(Haplo), GetCodesPerBlock(method));
      Uint *ansG = (Uint*) DoAlign(Ans, ALIGN_COMPUTE, method);
      assert(ansG == ans);
      if (method == Shuffle) haplo2genoShuffle(ans, snps, individuals, ansG);
      else haplo2genoShuffle256(ans, snps, individuals, ansG);
      ReUseAs(Ans, method);
      Uint *info = GetInfo(Ans);
      Ulong sumgeno = sumGeno(ans, snps, individuals, method);
      StoreSumGeno(sumgeno);
    } else {
      //    HELPINFO("Not using 'Shuffle' as snp coding needs more memory and time.");
     //     printf("haplo\n");
     SEXP h = haplo2geno(Ans); // no PROTECT( needed
     //printf(" ende\n");
     UNPROTECT(1);
     return h;
    }
  }
  // printf("F m done =%d\n", method);

  if (PRESERVE) R_PreserveObject(Ans);
  //  printf("END\n");
 
  UNPROTECT(1);
  return Ans;
 }


SEXP compute(SEXP popul, SEXP Generation, SEXP Sex, SEXP Nr,
	     SEXP Tau, SEXP Vec, SEXP Betahat,
	     SEXP Select, SEXP Matrix_return) {
  assert_MoBPS();
  snpcoding method = (snpcoding) GLOBAL.genetics.method;
  Uint snps = IcomputeSNPS(popul, Generation, Sex, Nr, 1, NA_INTEGER,
			   Select, NULL),
    blocks = Blocks(snps),
    individuals = length(Generation),
    unitsPerIndiv = blocks * UnitsPerBlock;
  Ulong mem = blocks * individuals * UnitsPerBlock + UnitsPerBlock - 1;
  Rint *G = (Rint *) CALLOC(mem, BytesPerUnit);
  if (G == NULL) ERR1("mem allocation (%lu bytes)", mem * BytesPerUnit);
  Uint *g = (Uint*) algn(G),
    *g0 = g,
    n_protect = 0;
  if ((Uint) length(Vec) != individuals) ERR("'vec' not of correct length");

  IcomputeSNPS(popul, Generation, Sex, Nr,
	       1, NA_INTEGER, // R coded indices !!
	       Select, (BlockType0 *) g);


  if ( (method != Shuffle && method != Shuffle256)) {
    //    printf("method = %d\n", method);
    //  HELPINFO("Not using 'Shuffle' or 'TwoBit' coding needs more memory.");
    SEXP G0;
    PROTECT(G0 = createSNPmatrix(snps, individuals, method));
    n_protect++;
    g0 = (Uint*) DoAlign(G0, ALIGN_COMPUTE, method); 
  }
  haplo2geno(g, snps, individuals, method, unitsPerIndiv, g0);
  // printf("g0 ");for (int i=0; i<10; i++) printf("%d ", g0[i]); printf("\n\n\n");
  
  double *A = matrix_mult(g0, // no PROTECT( needed;
			  snps, individuals, method, true, true, 0);
  
  FREE(G);

  SEXP Ans = IsolveRelMat(individuals, A, REAL(Tau)[0], // no PROTECT( needed;
			  REAL(Vec),
			  REAL(Betahat)[0], LOGICAL(Matrix_return)[0]);
  FREE(A);
  if (n_protect > 0) UNPROTECT(n_protect);

  return Ans;
}
```
