## Supplementary material for "MoBPS - Modular Breeding Program Simulator": miraculix R-package: MX.cc

```
/*
 Authors 
 Martin Schlather,


 Copyright (C) 2018 -- 2019  Martin Schlather

This program is free software; you can redistribute it and/or
modify it under the terms of the GNU General Public License
as published by the Free Software Foundation; either version 3
of the License, or (at your option) any later version.

This program is distributed in the hope that it will be useful,
but WITHOUT ANY WARRANTY; without even the implied warranty of
MERCHANTABILITY or FITNESS FOR A PARTICULAR PURPOSE.  See the
GNU General Public License for more details.

You should have received a copy of the GNU General Public License
along with this program; if not, write to the Free Software
Foundation, Inc., 59 Temple Place - Suite 330, Boston, MA  02111-1307, USA.  
*/


#include <R.h>
#include <Rinternals.h>
#include <inttypes.h> 
//#include <Rdefines.h>
#include "AutoMiraculix.h"
#include "MX.h"
#include <General_utils.h>
#include <zzz_RandomFieldsUtils.h>
#include "options.h"

Rint PL=C_PRINTLEVEL,
  CORES=INITCORES; // INITI
bool PRESERVE = false;
SEXP Information = R_NilValue,
  Coding = R_NilValue,
  Method = R_NilValue;

bool debugging = false;


void ReUseAsX(SEXP Code, snpcoding method,
	     Uint snps, Uint individuals,
	     Uint memInUnitsPerIndiv, 
	     Uint bytesPerBlock, Uint bitsPerCode,
	     Uint codesperblock, bool haplo) {
  //  printf(" X=%d ", method);
  // int *i;  if (method == 0) printf("i=%d\n", i[100]);
  
  SEXP Class;
  Uint *info = GetInfo(Code);
  Ulong 
    memInUnits = (Ulong) individuals * memInUnitsPerIndiv,
    alignedMem = memInUnits + bytesPerBlock / BytesPerUnit - 1,
    origaligned = AlignedInUnits(info);

  if (origaligned != alignedMem) {
    // printf("mem %lu -> %lu\n", origaligned, alignedMem);
    if (origaligned < alignedMem) ERR("BUG: Reuse of memory impossible");
    Uint *code = (Uint *) INTEGER(Code);
    for (Uint i=alignedMem; i<origaligned; code[i++] = 0L);
  }
 
  MethodOf(Code) = haplo ? Haplo : method;
  info[WHAT] = haplo ? HAPLO : GENOMATRIX;
  info[SNPS] = snps;
  info[INDIVIDUALS] = individuals;  
  info[SNPxIND] = true;  
  info[BITSPERCODE] = bitsPerCode;
  info[CODESPERBLOCK] = codesperblock;
  info[BYTESPERBLOCK] = bytesPerBlock;
  info[SUMGENO] = info[SUMGENO_E9] = 0;
  StoreMemInUnits(memInUnits);
  ADDADDRESS((Rint*) algn_general(INTEGER(Code), bytesPerBlock), ALIGNADDR0); 

  PROTECT(Class = allocVector(STRSXP, 1));
  SET_STRING_ELT(Class, 0, mkChar(haplo ? HAPLOMATRIX : GENOMICMATRIX));
  SET_CLASS(Code, Class);
  UNPROTECT(1);
}


SEXP CreateEmptyCodeVectorALL(Uint snps, Uint individuals,
			      Uint memInUnitsPerIndiv, 
			      Uint bytesPerBlock, Uint bitsPerCode,
			      Uint codesperblock, bool haplo) {
  

  //  uint64_t OOO1 = UINT64_C(0x0001000100010001);
  //  printf("%ld %ld\n", OOO1, 0x0001000100010001);
  //uint64_t OOO1 = 281479271743489LL;
 

   SEXP Code, Info, method;
  Ulong 
    memInUnits = (Ulong) individuals * memInUnitsPerIndiv,
    alignedMem = memInUnits + bytesPerBlock / BytesPerUnit - 1;
 
  //  printf("create: snps = %u %u %lu %lu\n", snps, bitsPerCode, memInUnits, alignedMem);

   if (bitsPerCode == 32) { 
    if (snps * (Ulong) individuals != memInUnits) BUG;
    PROTECT(Code = allocMatrix(INTSXP, snps, individuals));
  } else {
    //printf("snps = %d %d prod=%d %d, %s\n", snps, individuals,snps * individuals, memInUnits, SNPCODING_NAME[GLOBAL.genetics.method]);
    
    if (snps * (Ulong) individuals <= memInUnits && snps > 1000) {
      //  printf("snps = %d %d prod=%d %d, %s\n", snps, individuals,snps * individuals, memInUnits, SNPCODING_NAME[GLOBAL.genetics.method]);
      BUG;
    }
    PROTECT(Code = allocVector(INTSXP, alignedMem)); 
    Uint* A = (Uint*) INTEGER(Code);
    for (Ulong k=0; k<alignedMem; A[k++] = 0);
   }
 
  
  PROTECT(method = allocVector(INTSXP, 1));
  PROTECT(Info = allocVector(INTSXP, INFO_LAST + 1));
  Uint *info = (Uint*) INTEGER(Info);
  for (Uint i=0; i<=INFO_LAST; i++) info[i] = NA_INTEGER;
  
  StoreAlignedInUnits(alignedMem);
  ADDADDRESS(INTEGER(Code), ADDR0);
  setAttrib(Code, Information, Info); // install("information")
  setAttrib(Code, Method, method); // install("method")

  ReUseAsX(Code, (snpcoding) GLOBAL.genetics.method,
	  snps, individuals, memInUnitsPerIndiv, bytesPerBlock, bitsPerCode,
	  codesperblock, haplo);
  
  UNPROTECT(3);

  return Code;
}


void start_info(SEXP Code, SEXP file, Uint unitsPerBlock, Uint codesperblock) {
   Uint *info = GetInfo(Code);
 
  if (file != R_NilValue) {
    Uint *fileinfo = GetInfo(file);
    info[SNPxIND] = fileinfo[SNPxIND];					
  }

  if (PL > 1) {								
    Uint
      mem = MemInUnits(info),
      individuals = info[INDIVIDUALS];
    bool inMB = mem > 5000000;
    PRINTF("Data: %d individuals and %d SNPs\nStorage mode: %d block(s) of %d codes\nSize of M: %d %sB.", 
	   individuals, info[SNPS], mem / unitsPerBlock, codesperblock,
	   1 + (mem - 1) / (inMB ? 1048576 : 1024),
	   inMB ? "M" : "k");	       
  }
}


Uint *GetInfo(SEXP M) { // has negative integers included!!
#define INFO_INFO 0
  SEXP Infos = getAttrib(M, Information);
  if (TYPEOF(Infos) == INTSXP) {
    return (Uint *) INTEGER(Infos);
  } else {
    //    PRINTF("obsolete storing of information");
    return (Uint *) INTEGER(VECTOR_ELT(Infos, INFO_INFO));
  }
}


Uint Inti(SEXP X, Uint i) {
  switch(TYPEOF(X)) {
  case INTSXP : return (Uint) INTEGER(X)[i];
  case LGLSXP : return (Uint) LOGICAL(X)[i];
  case REALSXP : return (Uint) REAL(X)[i];
  default : ERR("not of numerical type");
  }
}


Rint * GetAddress(Uint *info, Uint where) {
  addr_Uint addalign;
  for (Uint addr=0; addr<UnitsPerAddress; addr++)      
    addalign.u[addr] = info[where + addr];
  return addalign.a;
}
```
