## Supplementary material for "MoBPS - Modular Breeding Program Simulator": miraculix R-package: MX.h

```
/*
 Authors 
 Martin Schlather,


 Copyright (C) 2018 -- 2019  Martin Schlather

This program is free software; you can redistribute it and/or
modify it under the terms of the GNU General Public License
as published by the Free Software Foundation; either version 3
of the License, or (at your option) any later version.

This program is distributed in the hope that it will be useful,
but WITHOUT ANY WARRANTY; without even the implied warranty of
MERCHANTABILITY or FITNESS FOR A PARTICULAR PURPOSE.  See the
GNU General Public License for more details.

You should have received a copy of the GNU General Public License
along with this program; if not, write to the Free Software
Foundation, Inc., 59 Temple Place - Suite 330, Boston, MA  02111-1307, USA.  
*/


#ifndef miraculix_MX_H
#define miraculix_MX_H 1


// 1

//#include "miraculix.h"
//#include "AutoMiraculix.h"
//#include "intrinsics.h"
// include liste nicht erweitern, sondern stattdessen haplogeno.h verwenden

#include <R.h>
#include <Rinternals.h>
#include <inttypes.h>
#include "AutoMiraculix.h"


typedef int Rint;
typedef unsigned int Uint;
//typedef unsigned long long Ulong;
typedef uint64_t Ulong;
typedef int64_t Long;

extern bool debugging, PRESERVE;
extern Rint PL, CORES;
extern SEXP Information, Method, Coding; // READONLY !!!!

SEXP CreateEmptyCodeVectorALL(Uint snps, Uint individuals,
			      Uint memInUnitsPerIndiv, 
			      Uint bytesPerBlock, Uint bitsPerCode,
			      Uint codesperblock, bool haplo);

void ReUseAsX(SEXP Code, snpcoding method,
	      Uint snps, Uint individuals,
	      Uint memInUnitsPerIndiv, 
	      Uint bytesPerBlock, Uint bitsPerCode,
	      Uint codesperblock, bool haplo);
 
#define CreateEmptyCodeVector(snps, individuals, memInUnitsPerIndiv)	\
  CreateEmptyCodeVectorALL(snps, individuals, memInUnitsPerIndiv,	\
			     BytesPerBlock, BitsPerCode, CodesPerBlock, false);
#define CreateEmptyCodeVectorHaplo(snps, individuals, memInUnitsPerIndiv) \
  CreateEmptyCodeVectorALL(snps, individuals, memInUnitsPerIndiv, 	\
 			     BytesPerBlock, BitsPerCode, CodesPerBlock, true);

#define InternReUseAs(Code,method,snps,individuals,memInUnitsPerIndiv, haplo) \
  ReUseAsX(Code, method, snps, individuals, memInUnitsPerIndiv,		\
	   BytesPerBlock, BitsPerCode, CodesPerBlock, haplo);


void start_info(SEXP Code, SEXP file, Uint unitsperblock, Uint codesperblock);

// #define GetInfo(M) (Uint *) INTEGER(getAttrib(M, Information))
// OBIGES WAERE DAS RICHTIGE, ABER ALTES FORMAT EXISTIERT NOCH:
Uint *GetInfo(SEXP M);
#define MethodOf(M) INTEGER(getAttrib(M, Method))[0]
#define ClassOf(M) INTEGER(getAttrib(M, Class))[0]

#define SumGeno(info)							\
    ((Ulong) info[SUMGENO] +(uint64_t) info[SUMGENO_E9] * (Ulong) 1000000000)
#define StoreSumGeno(X) {					\
    Ulong GenoSum = X;					\
    info[SUMGENO] = GenoSum % (uint64_t) 1000000000;	\
    info[SUMGENO_E9] = GenoSum / (uint64_t) 1000000000;	\
  }


#define MemInUnits(info)						\
  ((Ulong) info[MEMinUNITS0] +(uint64_t) info[MEMinUNITS1] * (Ulong) 1000000000)
#define StoreMemInUnits(X) {					\
    Ulong memX = X;					\
    info[MEMinUNITS0] = memX % (uint64_t) 1000000000;	\
    info[MEMinUNITS1] = memX / (uint64_t) 1000000000;	\
  }


#define AlignedInUnits(info)						\
  ((Ulong) info[ALIGNEDUNITS0] +(uint64_t) info[ALIGNEDUNITS1] * (Ulong) 1000000000)
#define StoreAlignedInUnits(X) {					\
    Ulong memX = X;					\
    info[ALIGNEDUNITS0] = memX % (uint64_t) 1000000000;	\
    info[ALIGNEDUNITS1] = memX / (uint64_t) 1000000000;	\
  }


#define BitsPerByte 8L
#define BytesPerDouble 8L
#define BytesPerUnit 4L // R unit (int) !
#define MaxUnitsPerAddress 2L
#define UnitsPerAddress (1L + (sizeof(Uint *) - 1L) / BytesPerUnit)

typedef union addr_Uint{
  Rint * a;
  Uint u[UnitsPerAddress];
  uint8_t bbbb_CXXb[2 * sizeof(Uint*)]; // nur zum testen
} addr_Uint;


#define ALIGN_HAPLO 0
#define ALIGN_HAPLOGENO 1
#define ALIGN_VECTOR 2
#define ALIGN_RELATION 3
#define ALIGN_ALLELE 4
#define ALIGN_COMPUTE 5
#define ALIGN_CODED 6
#define ALIGN_LAST ALIGN_CODED

#define ALIGN_23 ALIGN_CODED
#define ALIGN_SSE ALIGN_CODED

#define algn_general(X, bytesperblock)  ((1L + (uintptr_t) (((uintptr_t) X - 1L) / bytesperblock)) * bytesperblock) // ok!!


	  
#define ADDADDRESS(WHICH, WHERE)	{				\
    addr_Uint addalign;							\
    addalign.a = WHICH;							\
    for (Uint addr=0; addr<UnitsPerAddress; addr++)			\
      info[WHERE + addr] = addalign.u[addr];				\
  }


Rint* GetAddress(Uint *info, Uint where);


Uint Inti(SEXP X, Uint i);
#define Int0(X) Inti(X, 0L)


#define HELPINFO(M) if (GLOBAL_UTILS->basic.helpinfo) PRINTF("%s\n(Note that you can unable this information by 'RFoptions(helpinfo=FALSE)'.)\n", M) //


#endif
```
