## Supplementary material for "MoBPS - Modular Breeding Program Simulator": miraculix R-package: nocoding.cc

```
/*
 Authors 
 Martin Schlather,


Copyright (C) 2019 -- 2019  Martin Schlather

This program is free software; you can redistribute it and/or
modify it under the terms of the GNU General Public License
as published by the Free Software Foundation; either version 3
of the License, or (at your option) any later version.

This program is distributed in the hope that it will be useful,
but WITHOUT ANY WARRANTY; without even the implied warranty of
MERCHANTABILITY or FITNESS FOR A PARTICULAR PURPOSE.  See the
GNU General Public License for more details.

You should have received a copy of the GNU General Public License
along with this program; if not, write to the Free Software
Foundation, Inc., 59 Temple Place - Suite 330, Boston, MA  02111-1307, USA.  
*/

#define BitsPerCode 32L
#define UNCOMPRESSED 1

//#include <stdio.h>
//#include "options.h"
#include "AutoMiraculix.h"
#include "intrinsics.h"
#include <General_utils.h>
// #include "error.h"
#include "haplogeno.h"
#include "Haplo.h"
#include "Scalar.h"

INLINER

Uint CodesPerBlockPlain() { return CodesPerBlock; }
Uint UnitsPerIndivPlain(Uint snps) { return snps; }

SEXP matrix_start_plain(Uint individuals, Uint snps,
			SEXP VARIABLE_IS_NOT_USED  file) {
  SEXP Code = CreateEmptyCodeVector(snps, individuals, snps);
  return Code;
}


void matrix_plain(Uint *M, Uint start_individual, Uint end_individual, 
		  Uint start_snp, Uint end_snp, Uint Mnrow,
		  SEXP Ans, double VARIABLE_IS_NOT_USED *G) {
  Uint
    *info = GetInfo(Ans),
    snps = info[SNPS],
    *ans = (Uint *) INTEGER(Ans);
  for (Ulong i=start_individual; i<end_individual; i++) {
    Uint *Mptr = ans + i * snps,
      *pM = M + (i - start_individual) * Mnrow;
    for (Uint s=start_snp; s<end_snp; pM++) {
      Mptr[s++] = *pM;
    }
  }
}


SEXP matrix_coding_plain(Uint *M, Uint snps, Uint individuals){
  SEXP Code;
  PROTECT(Code = matrix_start_plain(individuals, snps, R_NilValue));
  matrix_plain(M, 0, individuals, 0, snps, snps, Code, NULL);
  UNPROTECT(1);
  return Code;
}

SEXP get_matrixPlain(SEXP SNPxIndiv) {
   Uint 
     *info = GetInfo(SNPxIndiv),
      individuals = info[INDIVIDUALS],
     snps = info[SNPS];
   Uint
     *M = (Uint*) Align(SNPxIndiv, ALIGN_SSE);
   SEXP Ans;
   PROTECT(Ans=allocMatrix(INTSXP, snps, individuals));
   MEMCOPY(INTEGER(Ans), M, (Ulong) snps * individuals * sizeof(Uint));
   UNPROTECT(1);
   return Ans;
}


Uint *AlignPlain(SEXP Code, Uint nr, bool test) {return AlignTest(Code, nr, test); }


Ulong sumGenoPlain(Uint *S, Uint snps, Uint individuals) {
  Ulong units = (Ulong) snps * individuals;  
  Ulong sum = 0L;					  
  for (Ulong i=0; i<units; i++) sum += S[i];
  return sum;						  
}

void haplo2genoPlain(Uint *code, Uint snps, Uint individuals,
		    Uint unitsPerIndiv, Uint *MM) {
  Ulong total = (Ulong) snps * individuals;
  for (Ulong i=0; i<total; i++) MM[i] = 0; // noetig?
  assert(BitsPerCode == 32);

#ifdef DO_PARALLEL
#pragma omp parallel for num_threads(CORES)   
#endif
  for (Ulong i=0; i<individuals; i++) {
    Uint
      *cm = code + unitsPerIndiv * i,
      *M = MM + i * snps;
    for (Uint s = 0; s < snps; s++) M[s] = GetHaplo(cm, s);
  }
}

void matrixPlain_mult(Uint * SNPxIndiv, Uint snps, Uint individuals, double *A){
  matmulttransposedUint(SNPxIndiv, SNPxIndiv, A, snps, individuals, individuals);
}
```
