## Supplementary material for "MoBPS - Modular Breeding Program Simulator": miraculix R-package: nocoding.h

```
/*
 Authors 
 Martin Schlather,


 Copyright (C) 2018 -- 2019  Martin Schlather

This program is free software; you can redistribute it and/or
modify it under the terms of the GNU General Public License
as published by the Free Software Foundation; either version 3
of the License, or (at your option) any later version.

This program is distributed in the hope that it will be useful,
but WITHOUT ANY WARRANTY; without even the implied warranty of
MERCHANTABILITY or FITNESS FOR A PARTICULAR PURPOSE.  See the
GNU General Public License for more details.

You should have received a copy of the GNU General Public License
along with this program; if not, write to the Free Software
Foundation, Inc., 59 Temple Place - Suite 330, Boston, MA  02111-1307, USA.  
*/


#ifndef miraculix_auto_H
#define miraculix_auto_H 1

SEXP matrix_start_plain(Uint individuals, Uint snps, SEXP G);
void matrix_plain(Uint *M, Uint start_individual, Uint end_individual, 
		  Uint start_snp, Uint end_snp,  Uint Mnrow, SEXP Ans,
		  double *G);
SEXP matrix_coding_plain(Uint *M, Uint snps, Uint individuals);
SEXP get_matrixPlain(SEXP SNPxIndiv);
Uint *AlignPlain(SEXP Code, Uint nr, bool test);
Uint CodesPerBlockPlain();
Uint UnitsPerIndivPlain(Uint snps);
Ulong sumGenoPlain(Uint *S, Uint snps, Uint individuals);
void haplo2genoPlain(Uint * SNPxIndiv, Uint snps, Uint individuals,
		     Uint unitsPerIndiv, Uint *A);
Ulong sumGenoPlain(Uint *S, Uint snps, Uint individuals);
void matrixPlain_mult(Uint * SNPxIndiv, Uint snps, Uint individuals, double *A);


#endif
```
