## Supplementary material for "MoBPS - Modular Breeding Program Simulator": miraculix R-package: options.cc

```
/* 
 Authors
 Martin Schlather,

 library for simulation of random fields 

 Copyright (C) 2018 -- 2019 Martin Schlather, 

This program is free software; you can redistribute it and/or
modify it under the terms of the GNU General Public License
as published by the Free Software Foundation; either version 3
of the License, or (at your option) any later version.
RO
This program is distributed in the hope that it will be useful,
but WITHOUT ANY WARRANTY; without even the implied warranty of
MERCHANTABILITY or FITNESS FOR A PARTICULAR PURPOSE.  See the
GNU General Public License for more details.

You should have received a copy of the GNU General Public License
along with this program; if not, write to the Free Software
Foundation, Inc., 59 Temple Place - Suite 330, Boston, MA  02111-1307, USA.
*/


#include <R.h>
#include <Rinternals.h>
#include <Rdefines.h>
#include <R_ext/Linpack.h>
#include <stdio.h>  
//#include <stdlib.h>
#include <unistd.h>
#include <string.h>

// ACHTUNG: Reihenfolge nicht aendern!
#include "IntrinsicsBase.h"
#include "intrinsics.h"
#include <Basic_utils.h>
#include "error.h"
#include <zzz_RandomFieldsUtils.h>
#include "miraculix.h"
#include "MX.h"
#include "options.h"
//#include "def.h"
#include "AutoMiraculix.h"
#include "kleinkram.h"
#include "xport_import.h" // important! must be very last!
//                           (HAS_XXX in zzz_RandomFieldsUtils.h)


snpcoding getAutoCodingIntern() {
  return
#if defined AVX2
    Shuffle256
#elif defined SSSE3
    Shuffle
#elif defined SSE2
    Hamming2
#else
    ThreeBit
#endif
    ;  
}


#define AVX2_USED true
#define AVX_USED false
#define SSSE3_USED true
#define SSE2_USED true
#define SSE_USED false

void PrintSystem() {
#ifdef AttachMessage
  PRINTF(AttachMessage(miraculix,false));
  PRINTF("\n");
#endif
}

SEXP attachmiraculix() {
#ifdef SCHLATHERS_MACHINE
  PRINTF("floating point double precision: %s\n",
#if defined DO_FLOAT
	 "no"
#else
	 "yes"
	 );
#endif
#endif

#ifdef ReturnAttachMessage
  ReturnAttachMessage(miraculix, true);
#else
  return R_NilValue;
#endif
}

void detachmiraculix() {
  Ext_detachRFoptions(prefixlist, prefixN);
}


 
CALL1(void, getErrorString, errorstring_type, errorstring)
CALL1(void, setErrorLoc, errorloc_type, errorloc)

const char * prefixlist[prefixN] = {"genetics"};


// IMPORTANT: all names of general must have at least 3 letters !!!
const char *genetics[geneticsN] = 
  {"digits", "snpcoding", "centered", "normalized", "returnsigma"};

  

globalparam GLOBAL = {
genetics_START
};
utilsparam *GLOBAL_UTILS;


const char **all[prefixN] = {genetics};
int allN[prefixN] = {geneticsN};
 
void setparameter(Rint i, Rint j, SEXP el, char name[200], 
		  bool VARIABLE_IS_NOT_USED isList, Rint local) {
#ifdef DO_PARALLEL
  if (local != isGLOBAL) ERR1("Options specific to RandomFieldsUtils, here '%.50s', can be set only via 'RFoptions' outside any parallel code.", name);
#endif  
  globalparam *options = &GLOBAL; 
  switch(i) {
  case 0: {// genetics
    genetics_param *gp;
    gp = &(options->genetics);
    switch(j) {
    case 0: gp->digits = NUM; break;
    case 1: {
      Rint m = TYPEOF(el) != STRSXP ? POS0NUM
	: GetName(el, name, SNPCODING_NAME, last_usr_meth + 1, gp->method);
#if !defined SSE2
      if (m == Hamming2) {
	PrintSystem();
 	ERR("'Hamming2', which needs 'SSE2', is not available under the current compilation. See the starting message for a remedy.");
      }
#endif
#if !defined SSSE3 
      if (m == Hamming3 || m == Shuffle) {
	PrintSystem();
	ERR1("'%10s', which needs 'SSSE3' is not available under the current compilation. See the starting message for a remedy.",
	     m == Hamming3 ? "Hamming3" : "Shuffle");
      }
#endif
#if !defined AVX2
      if (m == Shuffle256) {
	PrintSystem();
	ERR("'Shuffle256', which needs 'AVX2', is not available under the current compilation. See the starting message for a remedy.");
      }
#endif
      if (m > last_usr_meth) ERR("given snp coding not allowed");
      gp->method = m;
      break;
    }
    case 2:
      if (TYPEOF(el) == LGLSXP) {
	gp->normalized = (gp->centered = USRLOG);
	if (gp->centered == Nan) {
	  if (gp->pcentered == NULL) {
	    WARN1("'%.50s' set to TRUE", name); // OK
	    gp->normalized = (gp->centered = True);
	  }
	} else {
	  FREE(gp->pcentered);
	  gp->ncentered = 0;
	}
      } else {
	Uint len = length(el);
	gp->ncentered = len;
	FREE(gp->pcentered);
	gp->pcentered = (double*) MALLOC(len * sizeof(double));
	Real(el, name, gp->pcentered, len);
	gp->centered = Nan;
	gp->normalized = False;
      }
      break;      
    case 3:
      gp->normalized = LOGI;
      if (gp->normalized && gp->centered != True) { 
	warn("'normalized=TRUE' only allowed with 'centered=TRUE'.\n'normalized=FALSE' is kept"); // OK
	gp->normalized = false;
      }
      break;
    case 4 : gp->returnsigma = LOGI; break;
    default: BUG; 
    }
  }
    break;
  default: BUG;
  }
}


#define PLoffset -10
  void finalparameter(Rint VARIABLE_IS_NOT_USED local) {
  PL = GLOBAL_UTILS->basic.Cprintlevel - PLoffset;
  CORES = GLOBAL_UTILS->basic.cores;
}


void getparameter(SEXP sublist, Rint i, Rint VARIABLE_IS_NOT_USED local) {
  Uint k;
#ifdef DO_PARALLEL
  //  if (local != isGLOBAL) ERR("Options specific to RandomFieldsUtils can be obtained only on a global level and outside any parallel code.");
#endif  
  globalparam *options = &GLOBAL; 
 switch(i) {
  case 0 : {
    k = 0;
    genetics_param *p = &(options->genetics);
    ADD(ScalarReal(p->digits));
    //    ADD(ScalarString(mkChar(RELSHIP_METH_NAME[p->method])));
    ADD(ScalarInteger(p->method));
    ADD(ExtendedBooleanUsr(p->centered));    
    ADD(ScalarLogical(p->normalized));
    ADD(ScalarLogical(p->returnsigma));
  }
    break;
  default : BUG;
  }
}


SEXP loadmiraculix() {
  includeXport();
  Ext_getUtilsParam(&GLOBAL_UTILS);
  GLOBAL_UTILS->solve.max_chol = 8192;
  GLOBAL_UTILS->solve.max_svd = 6555;  

  finalparameter(isGLOBAL);
  Ext_attachRFoptions(prefixlist, prefixN, all, allN,
		      setparameter, finalparameter, getparameter,
		      NULL, -10, false);

  finalparameter(isGLOBAL);
  
  Information = install("information");
  Method = install("method");
  Coding = install("coding");

  if (BytesPerUnit != sizeof(Uint))
    ERR1("The programme relies on the assumption that an unsigned integer has %d Bytes.", BytesPerUnit);
  
  if (sizeof(int) != sizeof(Uint))
    ERR2("The programme relies on the assumption that a signed integer the same lenth than an unsigned integer. Found %d and %d bytes respectively.", sizeof(int), sizeof(Uint));
  
  if (sizeof(uint64_t) != sizeof(double))
    ERR2("The programme relies on the assumption that an 'uint64_t' has the size of a 'double'. Found %d and %d bytes respectively.", sizeof(uint64_t), sizeof(double));
  
  if (sizeof(uint64_t) != 8) ERR1("The programme relies on the assumption that an 'uint64_t' has 8 Bytes. Found %d bytes.", sizeof(uint64_t));
  
  if (sizeof(void*) > MaxUnitsPerAddress * BytesPerUnit)
    ERR2("The programme relies on the assumption that an 'void*' has at most %d Bytes. Found %d bytes.", MaxUnitsPerAddress * BytesPerUnit, sizeof(void*));
  return R_NilValue;
}


#ifndef HAS_PARALLEL
#ifdef DO_PARALLEL
#define HAS_PARALLEL true
#else
#define HAS_PARALLEL false
#endif
#if defined AVX2
#define HAS_AVX2 true
#else
#define HAS_AVX2 false
#endif
#if defined AVX
#define HAS_AVX true
#else
#define HAS_AVX false
#endif
#if defined SSSE3
#define HAS_SSSE3 true
#else
#define HAS_SSSE3 false
#endif
#if defined SSE2
#define HAS_SSE2 true
#else
#define HAS_SSE2 false
#endif
#if defined SSE
#define HAS_SSE true
#else
#define HAS_SSE false
#endif
#endif


SEXP hasSSE2() {
  SEXP Ans;
  PROTECT(Ans = allocVector(LGLSXP, 1));
  LOGICAL(Ans)[0] = HAS_SSE2;
  UNPROTECT(1);
  return Ans;
}


SEXP hasSSSE3() {
  SEXP Ans;
  PROTECT(Ans = allocVector(LGLSXP, 1));
  LOGICAL(Ans)[0] = HAS_SSSE3;
  UNPROTECT(1);
  return Ans;
}


SEXP hasAVX() {
  SEXP Ans;
  PROTECT(Ans = allocVector(LGLSXP, 1));
  LOGICAL(Ans)[0] = HAS_AVX;
  UNPROTECT(1);
  return Ans;
}


SEXP hasAVX2() {
  SEXP Ans;
  PROTECT(Ans = allocVector(LGLSXP, 1));
  LOGICAL(Ans)[0] = HAS_AVX2;
  UNPROTECT(1);
  return Ans;
}
```
