## Supplementary material for "MoBPS - Modular Breeding Program Simulator": miraculix R-package: options.h

```
/*
 Authors 
 Martin Schlather,


 Copyright (C) 2016 -- 2019 Martin Schlather

This program is free software; you can redistribute it and/or
modify it under the terms of the GNU General Public License
as published by the Free Software Foundation; either version 3
of the License, or (at your option) any later version.

This program is distributed in the hope that it will be useful,
but WITHOUT ANY WARRANTY; without even the implied warranty of
MERCHANTABILITY or FITNESS FOR A PARTICULAR PURPOSE.  See the
GNU General Public License for more details.

You should have received a copy of the GNU General Public License
along with this program; if not, write to the Free Software
Foundation, Inc., 59 Temple Place - Suite 330, Boston, MA  02111-1307, USA.  
*/


#ifndef miraculix_options_H
#define miraculix_options_H 1

#include <Rinternals.h>
#include <Basic_utils.h>
#include "AutoMiraculix.h"

#define geneticsN 5
typedef struct genetics_param {
  bool normalized, returnsigma;
  usr_bool centered;
  double digits;
  Rint method;
  // internal:
  double *pcentered;
  Rint ncentered;
} genetics_param;

#define genetics_START {			\
    true, false,				\
      True,					\
      3.0,					\
      (int) getAutoCodingIntern(),			\
      NULL, 0 /* internal */			\
}


typedef struct globalparam {
  genetics_param genetics;
} globalparam;
extern globalparam GLOBAL;

#define prefixN 1
extern const char * prefixlist[prefixN], **all[prefixN];
extern Rint allN[prefixN];
void setparameter(Rint i, Rint j, SEXP el, char name[200], bool isList, Rint local);
void getRFoptions(SEXP *sublist);
void finalparameter(Rint local);

snpcoding getAutoCodingIntern();
  
#endif
```
