## Supplementary material for "MoBPS - Modular Breeding Program Simulator": miraculix R-package: Scalar.cc

```
/* 
 Authors
 Martin Schlather,

 Collection of system specific auxiliary functions

 Copyright (C) 2019 -- 2019 Martin Schlather, 

This program is free software; you can redistribute it and/or
modify it under the terms of the GNU General Public License
as published by the Free Software Foundation; either version 3
of the License, or (at your option) any later version.

This program is distributed in the hope that it will be useful,
but WITHOUT ANY WARRANTY; without even the implied warranty of
MERCHANTABILITY or FITNESS FOR A PARTICULAR PURSE.  See the
GNU General Public License for more details.

You should have received a copy of the GNU General Public License
along with this program; if not, write to the Free Software
Foundation, Inc., 59 Temple Place - Suite 330, Boston, MA  02111-1307, USA.
*/

/*

Makefile must be:

PKG_LIBS =  $(SHLIB_OPENMP_CXXFLAGS) $(LAPACK_LIBS) $(BLAS_LIBS) $(FLIBS)  -march=native  -mssse3 
PKG_CXXFLAGS =  $(SHLIB_OPENMP_CXXFLAGS)  -march=native -mssse3 

 */


//#include <assert.h>
//#include "kleinkram.h"
#include "Scalar.h"
#include <General_utils.h>
//#include "intrinsics.h"
//#include "Basic_utils.h"
//#include "errors_messages.h"
//#include "zzz_RandomFieldsUtils.h"


//#define Nmodi 9
//name_type modi = { "1x1", "2x2", "4x4", "8x8", "near", "simple", "precise", "kahan", "1x1p"};


#define size 8
#define vectorlen (256 / (size * 8))
#define repet 8
#define atonce (vectorlen * repet)

#if (8 != repet)
  wrong repet length
#endif
#if (4 != vectorlen)
  wrong vector length
#endif


void matmulttransposedUint(Uint *A, Uint *B, double *c, Uint m, Uint l, Uint n) {
// multiplying t(A) and B with dim(A)=(m,l) and dim(B)=(m,n),
// saving result in C
#ifdef DO_PARALLEL
#pragma omp parallel for num_threads(CORES) 
#endif
  for (Uint i=0; i<l; i++) {    
    double *C = c + i;
    Uint *Aim = A + i * m;
    for (Uint j=0; j<n; j++) C[j * l] = (double) SCALARUINT(Aim, B + j * m, m);
  }
}


 
Uint scalarUint2by2( Uint * v1, Uint * v2, Uint N){
  Uint *endv1 = v1 + (N / 2) * 2,
    *end = v1 + N,
    sum = 0;
  for(; v1 < endv1; v1 += 2, v2 += 2) sum += v2[0] * v1[0] + v2[1] * v1[1];
  if (v1 < end) sum += v2[0] * v1[0]; 
  return sum;
}
 
 
Uint scalarUint4by4( Uint * v1, Uint * v2, Uint N){
  Uint*endv1 = v1 + (N / 4) * 4,
    *end = v1 + N,
    sum = 0;
  for(; v1 < endv1; v1 += 4, v2 += 4)
    sum += v2[0] * v1[0] + v2[1] * v1[1] + v2[2] * v1[2]+ v2[3] * v1[3];
  for(; v1 < end; v1++, v2++) sum += v2[0] * v1[0];        
  return sum;
}

 
Uint scalarUint8by8( Uint * v1, Uint * v2, Uint N){
  Uint
    *endv1 = v1 + (N / 8L) * 8L,
    *end = v1 + N,
    sum = 0.0;
  for(; v1 < endv1; v1 += 8L, v2 += 8L)
    sum += v2[0] * v1[0] + v2[1] * v1[1]+ v2[2] * v1[2] + v2[3] * v1[3] +
      v2[4] * v1[4] + v2[5] * v1[5]+ v2[6] * v1[6]+ v2[7] * v1[7];
  for(; v1 < end; v1++, v2++) sum +=  v2[0] * v1[0];        
  return sum;
}


Uint scalarUintPX( Uint * V1, Uint * V2, Uint N){
#define AtOnce 16
  Uint
    *endv1 = V1 + (N / AtOnce) * AtOnce,
    *end = V1 + N,
    sum = 0;
#ifdef DO_PARALLEL
#pragma omp parallel for num_threads(CORES) if (N > 200) reduction(+:sum) schedule(dynamic, 50) 
#endif
  for(Uint *v1=V1; v1 < endv1; v1 += AtOnce) {
    Uint *v2 = V2 + (V1 - v1);
    sum +=  v2[0] * v1[0] + v2[1] * v1[1]+ v2[2] * v1[2] + v2[3] * v1[3] +
      v2[4] * v1[4] + v2[5] * v1[5]+ v2[6] * v1[6]+ v2[7] * v1[7] +
      v2[8] * v1[8] + v2[9] * v1[9]+ v2[10] * v1[10] + v2[11] * v1[11] +
      v2[12] * v1[12] + v2[13] * v1[13]+ v2[14] * v1[14]+ v2[15] * v1[15];
  }
  Uint
    *v1 = V1 + (N / AtOnce),
    *v2 = V2 + (V1 - v1);
  for(; v1 < end; v1++, v2++) sum += v2[0] * v1[0];        
  return sum;
}


//bool pr = true;
Uint scalarUint(Uint *x, Uint *y, Uint len, Uint n) {
 //  if (pr) { prUintf("mode = %d\n", n); pr = false; }
 // 0 : 7.9
// 1:  7.55
// 2: 7.8
// 3:7.58
//4: 7.5 
// 5: 7.4!
//6:7.4
//7: 7.9
// 8: "ewige" schleife

  switch(n) {
  case 0 : //return scalarUint(x, y, len);
  case SCALAR_BASE : return scalarUint8by8(x, y, len); 
  case 2 : return scalarUint4by4(x, y, len); 
  case 3 : return scalarUint2by2(x, y, len);
#ifdef FMA_AVAILABLE_X_X_X_X_X_X
  case 4 : //return avx_scalarUintDfma(x, y, len);
#endif    
#ifdef AVX_X_X_X_X_X_X
  case 5 : //return avx_scalarUintDnearfma(x, y, len); 
  case SCALAR_AVX : return avx_scalarUintD(x, y, len); // best one kernel
  case 7 : return avx_scalarUintDP(x, y, len);  //best
    //  case SCALAR_KAHAN : return avx_scalarUintDK(x, y, len);  -- macht keinen Sinn bei Uint
#else
  case 4: case 5: case 6: case 7: case 8 : return scalarUint8by8(x, y, len);
#endif


    
#ifndef DO_PARALLEL // so does not work
    
#ifdef DO_PARALLEL
  case SCALAR_AVX_PARALLEL :
#if defined AVX && defined OpenMP
    return avx_scalarUintDparallel(x, y, len);
#endif    
  case SCALAR_BASE_PARALLEL : return scalarUintP(x, y, len);// parallel, nicht-vectoriell
#else
  case SCALAR_AVX_PARALLEL :
#ifdef AVX
  return avx_scalarUintD(x, y, len);
#endif     
   case SCALAR_BASE_PARALLEL : return scalarUint2by2(x, y, len); 
#endif
     
#endif
  default : ERR("method not available"); 
  }
  return RF_NAN;
}
  
```
