## Supplementary material for "MoBPS - Modular Breeding Program Simulator": miraculix R-package: Scalar.h

```
/*
 Authors 
 Martin Schlather,


 Copyright (C) 2019 -- 2019 Martin Schlather

This program is free software; you can redistribute it and/or
modify it under the terms of the GNU General Public License
as published by the Free Software Foundation; either version 3
of the License, or (at your option) any later version.

This program is distributed in the hope that it will be useful,
but WITHOUT ANY WARRANTY; without even the implied warranty of
MERCHANTABILITY or FITNESS FOR A PARTICULAR PURPOSE.  See the
GNU General Public License for more details.

You should have received a copy of the GNU General Public License
along with this program; if not, write to the Free Software
Foundation, Inc., 59 Temple Place - Suite 330, Boston, MA  02111-1307, USA.  
*/


#ifndef miraculix_ScalarInt_H
#define miraculix_ScalarInt_H 1

#include "MX.h"

#define SCALAR_AVX 6
#define SCALAR_KAHAN 8
#define SCALAR_BASE 1
#define SCALAR_AVX_PARALLEL 9
#define SCALAR_BASE_PARALLEL 10


#define SCALARUINT(x, y, len) scalarUint(x, y, len, SCALAR_BASE);
Uint scalarUint(Uint *x, Uint *y, Uint len, Uint n);
void matmulttransposedUint(Uint *A, Uint *B, double *c, Uint m, Uint l, Uint n);

#endif
```
