## Supplementary material for "MoBPS - Modular Breeding Program Simulator": miraculix R-package: scan.cc

```
/*
 Authors 
 Martin Schlather,


 Copyright (C) 2018 -- 2019  Martin Schlather

This program is free software; you can redistribute it and/or
modify it under the terms of the GNU General Public License
as published by the Free Software Foundation; either version 3
of the License, or (at your option) any later version.

This program is distributed in the hope that it will be useful,
but WITHOUT ANY WARRANTY; without even the implied warranty of
MERCHANTABILITY or FITNESS FOR A PARTICULAR PURPOSE.  See the
GNU General Public License for more details.

You should have received a copy of the GNU General Public License
along with this program; if not, write to the Free Software
Foundation, Inc., 59 Temple Place - Suite 330, Boston, MA  02111-1307, USA.  
*/


#include <R.h>
#include "miraculix.h"
#include "MX.h"
#include <Basic_utils.h>
#include "error.h"
#include <zzz_RandomFieldsUtils.h>


bool debug = false;

#define START 0
#define END 1
#define LEVEL 2
#define IDX(N) (3 * (N))
#define DELETED 0

#define COLLECT(exclude_negative, DO, Check)				\
  int i, j,								\
    nthr = *nthres,							\
    len = *length,							\
    *pos = positions,							\
    min = *minscan,							\
    max = *maxscan,							\
    perSNP = *PER_SNP							\
    ;									\
  double mass;								\
									\
 /* // printf("Len=%d %d %d perSNP=%d\n", len, min, max, perSNP);*/	\
  for (i = 0 ; i<len; i++) {						\
    if (exclude_negative) for ( ; i<len; i++) if (freq[i] >= 0) break;  \
    if (i >= len) break;						\
    int pos_i = pos[i];							\
    mass = 0.0;								\
    for (j=i; j<len; j++) {						\
       double freq_j = freq[j];						\
      mass += freq_j;							\
      Long laenge = perSNP ? j-i+1 : pos[j] - pos_i + 1;		\
      if (laenge < min) {						\
	continue;							\
      }									\
      if (max > 0 && laenge > max) {			\
        if (debug) { PRINTF("break %d %d\n", max, (int) laenge);}	\
	break;								\
      }									\
      DO;								\
    }									\
    Check;								\
    for ( ; i<len; i++) if (freq[i] < 0) break;				\
  }					

  
void scanC(int *positions, int *length, double  *freq, int *minscan,
	   int *maxscan, double *threshold, int *nthres, int *PER_SNP,
	   int *above_threshold,  double *maximum) {
  double maxi = -1e-40;
  int k;
  bool toomanyintervals = false;
  for (k=0; k<*nthres; above_threshold[k++] = 0);

  COLLECT(false,							
	  if (mass >= threshold[0]) {					\
	    int d=0;	/* // printf("%10g %d %10g\n", mass, d, threshold[d]);*/ \
	    while (nthr > d && mass >= threshold[d]) above_threshold[d++]++; \
	  }								\
	  if (mass > maxi) maxi = mass;					\
	  , if (above_threshold[0] < 0) toomanyintervals = true;);
  *maximum = maxi;
  if (toomanyintervals) above_threshold[0] = -1;
}

		

SEXP scan(SEXP positions, SEXP length, SEXP freq, 
	  SEXP minscan,  SEXP maxscan, SEXP threshold, SEXP nthres,
	  SEXP PER_SNP,
	  SEXP above_threshold, SEXP Maximum) {
  scanC(INTEGER(positions), INTEGER(length), REAL(freq), 
	INTEGER(minscan),  INTEGER(maxscan), REAL(threshold),
	INTEGER(nthres),
	INTEGER(PER_SNP),
	INTEGER(above_threshold), REAL(Maximum));
  return R_NilValue;
}

SEXP collect_scan(int *positions, int *length, double  *freq, int *minscan,
		  int *maxscan, double *threshold, int *nthres,
		  int *PER_SNP, 
		  // additional return values:
		  int *areas,  double *value) {
  int n = 0,
    *a = areas;
  SEXP Res;
  
  COLLECT(false,					  \
      if (mass >= threshold[0]) {			  \
	int d=1;					  \
	a[START] = pos[i];				  \
	a[END] = pos[j];				  \
	while (nthr>d && mass >= threshold[d]) d++;	  \
	a[LEVEL] = d; /* R referencing starting with 1 */ \
	value[n] = mass;				  \
	n++;						  \
	a += 3;						  \
      }							  \
      , {});
  
  PROTECT(Res = allocMatrix(INTSXP, 3, n));
  int *res = INTEGER(Res);
  MEMCOPY(res, areas, 3 * sizeof(int) * n);

  for (i=0; i<n; i++) {
    int level = res[IDX(i) + LEVEL];
    if (res[IDX(i) + LEVEL] == DELETED) continue; 
    for (j=i+1; j<n; j++) {
      if (res[IDX(j) + START] == DELETED || level > res[IDX(j) + LEVEL])
	continue;  
      if (res[IDX(i) + END] < res[IDX(j) + START]) break;
      // NOTE that value of res[IDX(i) + END] is changing
      if (res[IDX(i) + END] < res[IDX(j) + END])
	res[IDX(i) + END] = res[IDX(j) + END];
      if (res[IDX(i) + LEVEL] == res[IDX(j) + LEVEL]) {
	res[IDX(j) + START] = res[IDX(j) + END] = res[IDX(j) + LEVEL] = DELETED;
      }
    }
  }

  UNPROTECT(1);
  return Res;
}


  
SEXP collect_scan(SEXP positions, SEXP length, SEXP freq, 
		  SEXP minscan,  SEXP maxscan, SEXP threshold, SEXP nthres,
		  SEXP PER_SNP, SEXP areas,  SEXP value
		  ) {
   return collect_scan(INTEGER(positions), INTEGER(length), REAL(freq), 
	   INTEGER(minscan), INTEGER(maxscan), REAL(threshold), INTEGER(nthres),
	   INTEGER(PER_SNP), INTEGER(areas), REAL(value));
 }


  

SEXP collect_scan2(int *positions, int *length, double  *freq, int *minscan,
		   int *maxscan, double *threshold, int *nthres,
		   int *PER_SNP, 
		   int max_intervals, int max_basepair_dist,
		   bool exclude_negative,
		   // additonal return values
		   int *above_threshold, double *maximum) {
  int n = 0, k,
    threemaxI = 3 * max_intervals,
    *res = (int*) MALLOC(sizeof(int) * threemaxI),
    *a = res;
  double maxi = -1e-40;
  bool not_exclude = !exclude_negative;

  for (k=0; k<*nthres; above_threshold[k++] = 0);

  COLLECT(exclude_negative,						\
	  if (j > i && pos[j] > pos[j-1] + max_basepair_dist) break;	\
	  if (mass >= threshold[0] && (not_exclude || freq_j>=0.0)) {	\
	    if (mass > maxi) maxi = mass;				\
	    int m;							\
	    int d=1;							\
	    a[START] = pos_i;						\
	    a[END] = pos[j];						\
	    while (nthr>d && mass >= threshold[d]) d++;			\
	    a[LEVEL] = d; /* R referencing starting with 1 */		\
	    bool del = false;						\
	    for (m=0; m<n; ) {						\
	      int idxm = IDX(m++);					\
	      assert(idxm + LEVEL < threemaxI && idxm + END < threemaxI); \
	      if (res[idxm + LEVEL] > a[LEVEL] ||			\
		  res[idxm + END] <  pos_i) continue;			\
	      if (res[idxm + END] < a[END]) res[idxm + END] = a[END];	\
	      if (res[idxm + LEVEL] == a[LEVEL]) {del=true; break;}	\
	    }								\
	    if (!del) {							\
	      for (k=0; k<d; above_threshold[k++]++);			\
	      n++;							\
	      if (n >= max_intervals) {					\
		ERR("too many intervals found; analysis stopped.");	\
	      }								\
	      a += 3;							\
	    }								\
	  }								\
	  , {});
  
  *maximum = maxi;
  SEXP Res;
  PROTECT(Res = allocMatrix(INTSXP, 3, n));

  if (n>0) MEMCOPY(INTEGER(Res), res, 3 * n * sizeof(int));
  FREE(res);
 
  // int m; for(m=0; m<n; m++) print("n=%d %d %d %d\n", n, INTEGER(Res)[3 *m ], INTEGER(Res)[3 *m +1], INTEGER(Res)[3 *m + 2]);
  UNPROTECT(1);

  return Res;
}


SEXP collect_scan2(SEXP positions, SEXP length, SEXP freq, 
		   SEXP minscan,  SEXP maxscan, SEXP threshold, SEXP nthres,
		   SEXP PER_SNP, SEXP max_intervals, SEXP max_basepair_dist, 
		   SEXP exclude_negative, 
		   SEXP above_threshold, SEXP maximum
		  ) {
   return collect_scan2(INTEGER(positions), INTEGER(length), REAL(freq), 
			INTEGER(minscan), INTEGER(maxscan), REAL(threshold), 
			INTEGER(nthres),
			INTEGER(PER_SNP), INTEGER(max_intervals)[0],
			INTEGER(max_basepair_dist)[0],
			(bool) INTEGER(exclude_negative)[0],
			INTEGER(above_threshold),
			REAL(maximum));
 }

void sumscanC(int *positions, int *length, double  *freq, 
	     int *minscan,  int *maxscan, double *threshold, int *nthres,
	     int *PER_SNP,
	     int *above_threshold,  double *maximum) {
  double res = -1e-40;
  int k;
  for (k=0; k<*nthres; above_threshold[k++] = 0);
  if (!*PER_SNP) ERR("sumscan only for 'perSNP=TRUE'");

  COLLECT(false,							\
     if (mass >= threshold[0]) {					\
       int *cp = above_threshold;					\
       for (int d=0; d<nthr; d++) {					\
	   if (mass < threshold[d]) break;				\
	   for (k=i; k<=j; cp[k++]++);					\
	   cp+=len;							\
       }								\
     }									\
     if (mass > res) res = mass;					\
     , {});
    *maximum = res;
}
							
  
SEXP sumscan(SEXP positions, SEXP length, SEXP freq, 
	     SEXP minscan,  SEXP maxscan, SEXP threshold, SEXP nthres,
	     SEXP PER_SNP,
	     SEXP above_threshold, SEXP Maximum) {
  sumscanC(INTEGER(positions), INTEGER(length), REAL(freq), 
	  INTEGER(minscan), INTEGER(maxscan), REAL(threshold), INTEGER(nthres),
	  INTEGER(PER_SNP),
	  INTEGER(above_threshold), REAL(Maximum));
  return R_NilValue;
}
```
