## Supplementary material for "MoBPS - Modular Breeding Program Simulator": miraculix R-package: shuffle256.cc

```
/*
 Authors 
 Martin Schlather,


 Copyright (C) 2018 -- 2019  Martin Schlather

This program is free software; you can redistribute it and/or
modify it under the terms of the GNU General Public License
as published by the Free Software Foundation; either version 3
of the License, or (at your option) any later version.

This program is distributed in the hope that it will be useful,
but WITHOUT ANY WARRANTY; without even the implied warranty of
MERCHANTABILITY or FITNESS FOR A PARTICULAR PURPOSE.  See the
GNU General Public License for more details.

You should have received a copy of the GNU General Public License
along with this program; if not, write to the Free Software
Foundation, Inc., 59 Temple Place - Suite 330, Boston, MA  02111-1307, USA.  
*/


#include "IntrinsicsBase.h"

#if defined AVX2

#define BitsPerCode 2L
#include "intrinsics.h"
#include <General_utils.h>
#include "xport_import.h"
#include "options.h"
#include "MX.h"
#include "shuffleDef.h"


// [1 + (X - 1) / UnitsPerBlock] * UnitsPerBlock + (UnitsPerBlock - 1)
// last part: to make finally sure that first used element of R-vector
// is aligned (maximum distance to 0-elemtn: UnitsPerBlock - 1)
// first part: then memory is read in blocks; so next number of Bytes that is
// is divisible by Unitsperblock * Bytesperunit is taken:
// 1 + (X - 1) / UnitsPerBlock]


static void vectorGenoKahan(Real *V,  BlockType0 *CM, Uint snps,
		     Uint individuals, double *ans) {
  Uint blocks = Blocks(snps);
  BlockType first2bits32;
  // NullEins = SET32(0x55555555),
  SET32(first2bits32, 0x00000003);


#ifdef DO_PARALLEL
#pragma omp parallel for num_threads(CORES)   
#endif
  for (Uint i=0; i<individuals; i++) {
    BlockType
      *cm = CM + i * blocks;
    Real sum, corr,
      *v = V;
    ZEROREAL(sum);
    ZEROREAL(corr);
    for (Uint b=0; b<blocks; b++, cm++) {
      BlockUnitType d,f;
      BlockType c = *cm;
      //   d VI = SHR32(c, 1);
      // d VI = AND(NullEins, d VI);      
      // c = XOR(c, d VI);      
      for (Uint j=0; j<CodesPerUnit; j++) {
	AND(d VI, c, first2bits32);
	Real  dummy, fcorr, roughsum;
	
	IR(0,0);
	IR(1,1);
	VGK;

#if !defined DO_FLOAT
	IR(0,2); IR(1,3);
	v++;
	VGK;
#endif	
	v++;
	SHR32(c, c, 2L);
      }
    }
    real *s = (real*) &sum;
    ans[i] = (double) (s[0] + s[1] + s[2] + s[3]
#if  defined DO_FLOAT    
		       + s[4] + s[5] + s[6] + s[7]
#endif		     
		       );
  }

}


static void vectorGenoIntern(Real *V, BlockType0 *CM, Uint snps,
		      Uint individuals, double *ans) {

  if (GLOBAL_UTILS->basic.kahanCorrection) {
    vectorGenoKahan(V, CM, snps, individuals, ans);
    return;
  }
  Uint blocks = Blocks(snps);
  BlockType    first2bits32;
  // NullEins = SET32(0x55555555),
  SET32(first2bits32, 0x00000003);

#ifdef DO_PARALLEL
#pragma omp parallel for num_threads(CORES)   
#endif
  for (Uint i=0; i<individuals; i++) {
    BlockType
      *cm = CM + i * blocks;
    Real sum,
      *v = V;
    ZEROREAL(sum);
    for (Uint b=0; b<blocks; b++, cm++) {
      BlockUnitType d,f;
      BlockType c = *cm;
      //      d VI = SHR32(c, 1);
      //d VI = AND(NullEins, d VI);      
      // c = XOR(c, d VI);      
      for (Uint j=0; j<CodesPerUnit; j++) {
	AND(d VI, c, first2bits32);
	IR(0, 0);
	IR(1, 1);
	VGI;

#if !defined DO_FLOAT
	IR(0, 2); IR(1, 3);
	v++;
	VGI;
#endif
	
	v++;
	SHR32(c, c, 2L);
      }
    }
    real *s = (real*) &sum;
    ans[i] = (double) (s[0] + s[1] + s[2] + s[3]
#if defined DO_FLOAT    
		       + s[4] + s[5] + s[6] + s[7]
#endif		     
		       );
  }
}


Uint INLINE ADDUPVECTOR(BlockType0 x) {
  BlockType zero;
  ZERO(zero);
  BlockUnitType vsum;
  SAD8(vsum VI, x, zero);  
  return vsum.u32[0]+ vsum.u32[2] + vsum.u32[4] + vsum.u32[6];
}


#include "shuffleIntern.h"


void InitShuffle256() { InitIntern();}
Uint CodesPerBlockShuffle256() { return CodesPerBlock; }
Uint UnitsPerIndivShuffle256(Uint snps) { return Blocks(snps) * UnitsPerBlock; }

static void assert_intern() { ASSERT_INTERN(Shuffle256); } 
static void assert_intern(SEXP M) { ASSERT_INTERN_M(Shuffle256); }

Ulong sumGenoShuffle256(Uint *S, Uint snps, Uint individuals) {
  return sumGenoIntern(S, snps, individuals);
}

void ReUseAsShuffle256(SEXP Code) { ReUseAsIntern(Code, Shuffle256); }

void haplo2genoShuffle256(Uint *X, Uint snps, Uint individuals, Uint *Ans) {
  // Achtung!! Nach Aufruf muss sumGeno() berechnet werden!!
  haplo2genoIntern(X, snps, individuals, Ans);
}

     
void zeroNthGenoShuffle256(SEXP CM, SEXP NN) {
  assert_intern(CM);
  zeroNthGenoIntern(CM, NN);
}

SEXP get_matrixN_shuffle256(SEXP CM, SEXP NN) {
  assert_intern(CM);
  return get_matrixN_intern(CM, NN);
}


void vectorGenoShuffle256(SEXP V, SEXP Z, double *ans) {
  assert_intern(Z);
  vGIntern(V, Z, ans);
}


static void genoVectorIntern(BlockType0 *CM, double *V, Uint snps,
		      Uint individuals, double *ans) {
  if (GLOBAL_UTILS->basic.kahanCorrection) {
    //    genoVectorKahan(V, CM, snps, individuals, ans);
    // return;
  }
  Uint blocks = Blocks(snps),
    blocksM1 = blocks - 1;  
  double *end_ans = ans + snps;
  BlockType c,first2bits32;
  SET32(first2bits32, 0x00000003);
  for (Uint i=0; i<snps; ans[i++] = 0.0);

  for (Uint i=0; i<individuals; i++) {
    BlockType *cm = CM + i * blocks;
    double
      *a = ans,
      f0 = V[i],
      factor[4] = {0.0, f0, 2.0 * f0, 0}; // nur fuer geno mit 00, 01, 10 Code

    for (Uint b=0; b<=blocksM1; b++) {
       c = cm[b];
     
      Uint endfor = b < blocksM1 ? CodesPerUnit
		: (snps - blocksM1 * CodesPerBlock) / UnitsPerBlock;

      for (Uint j=0; j<endfor; j++) {
	BlockUnitType d;
	AND(d VI, c, first2bits32);
	a[0] += factor[d.u32[0]]; 
	a[1] += factor[d.u32[1]];
	a[2] += factor[d.u32[2]];
	a[3] += factor[d.u32[3]];
	a[4] += factor[d.u32[4]]; 
	a[5] += factor[d.u32[5]];
	a[6] += factor[d.u32[6]];
	a[7] += factor[d.u32[7]];
	a += 8;

	/* todo speed might be improved using
mm256_bitshuffle256_epi64_mask
_mm_blendv_epi8 (__m128i a, __m128i b, __m128i mask)
Rint _mm_cmpestra (__m128i a, Rint la, __m128i b, Rint lb, const Rint imm8)
__m128i _mm_hadd_epi32 (__m128i a, __m128i b)
__m64 _mm_maddubs_pi16 (__m64 a, __m64 b)
void _mm_maskmove_si64 (__m64 a, __m64 mask, char* mem_addr)
Rint _mm_movemask_ps (__m128 a)
Rint _m_pmovmskb (__m64 a)
Rint _mm256_movemask_pd (__m256d a)
_mm256_mask_bitshuffle256_epi64_mask (__mmask32 k2, __m256i b, __m256i c)
__m128d _mm_mask_permute_pd (__m128d src, __mmask8 k, __m128d a, const Rint imm8)
__m256d _mm256_permute_pd (__m256d a, Rint imm8)
__m256d _mm256_permute2f128_pd (__m256d a, __m256d b, Rint imm8)
__m256d _mm256_permute4x64_pd (__m256d a, const Rint imm8)
	*/

	SHR32(c, c, BitsPerCode);
      }
    }

    BlockUnitType d;
    AND(d VI, c, first2bits32);
    
    assert(end_ans >= a && end_ans - a < UnitsPerBlock);
    for(Uint k=0; a < end_ans; a++) {
      *a += factor[d.u32[k++]];
    }
  }
}


void genoVectorShuffle256(SEXP Z, SEXP V, double *ans) {
  assert_intern(Z);
  Uint
    *info = GetInfo(Z),
    snps = info[SNPS],
    individuals = info[INDIVIDUALS],
    len = length(V);
  if (len != individuals) ERR("vector 'V' not of correct length");
  genoVectorIntern((BlockType0 *) Align(Z, ALIGN_VECTOR), REAL(V), 
 		   snps, individuals, ans);
}


SEXP get_matrixshuffle256(SEXP CM) { // 0.45 bei 25000 x 25000
  assert_intern(CM);
  return get_matrixIntern(CM);
}


SEXP matrix_start_shuffle256( Uint snps,Uint individuals, SEXP file) {  
  assert_intern();  
  return matrix_start_Intern(snps, individuals, file);
}
 

void matrix_shuffle256(Uint *M, Uint start_individual, Uint end_individual, 
		       Uint start_snp, Uint end_snp, Uint Mnrow, SEXP Ans,
		       double VARIABLE_IS_NOT_USED *G) {
  matrix_Intern(M, start_individual, end_individual, 
		       start_snp, end_snp, Mnrow, Ans, G);
}


SEXP matrix_coding_shuffle256(Uint *M, Uint snps, Uint individuals) {  
  SEXP Code;
  PROTECT(Code = matrix_start_shuffle256(snps, individuals, R_NilValue));
  matrix_shuffle256(M, 0, individuals, 0, snps, snps, Code, NULL);
  UNPROTECT(1);
  return Code;
}


void matrixshuffle256_mult(Uint *CGM, Uint snps, Uint individuals, double *ans){
  matrix_multIntern(CGM, snps, individuals, ans);
}


Uint *AlignShuffle256(SEXP Code, Uint nr, bool test) {
  return AlignTest(Code, nr, test);
}

      
SEXP allele_freqShuffle256(SEXP GM) {
  assert_intern(GM);
  return allele_freqIntern(GM);
}


#else // !defined AVX2
#include "error.h"
#include "MX.h"
void AVX2missing() { ERR("'shuffle256' needs the availablity of 'AVX2'"); }
#define Sm { AVX2missing(); return R_NilValue; }
#define Su { AVX2missing(); return 0; }
#define Sv { AVX2missing(); }
#if defined VARIABLE_IS_NOT_USED
#define V VARIABLE_IS_NOT_USED
#else
#define V
#endif
SEXP allele_freqShuffle256(SEXP V GM) Sm
void matrixshuffle256_mult(Uint V* CGM, Uint V snps, Uint V individuals,
			double V *ans) Sv
SEXP matrix_start_shuffle256(Uint V snps, Uint V individuals, SEXP V G) Sm
void matrix_shuffle256(Uint V *M, Uint V start_individual,
		       Uint V end_individual, 
		       Uint V start_snp, Uint V end_snp, Uint V Mnrow,
		       SEXP V Ans, double V * G) Sv

SEXP get_matrixshuffle256(SEXP V SNPxIndiv) Sm
SEXP matrix_coding_shuffle256(Uint V *M, Uint V snps, Uint V individuals) Sm
Uint *AlignShuffle256(SEXP V Code, Uint V nr, bool V test) Su
void InitShuffle256() Sv
Uint CodesPerBlockShuffle256() Su
Uint UnitsPerIndivShuffle256(Uint V snps) Su
void haplo2genoShuffle256(Uint V * SNPxIndiv, Uint V snps, Uint V individuals,
		       Uint V *A) Sv
Ulong sumGenoShuffle256(Uint V *S, Uint V snps, Uint V individuals) Su
void zeroNthGenoShuffle256(SEXP V CM, SEXP V NN) Sv
SEXP get_matrixN_shuffle256(SEXP V CM, SEXP V NN) Sm
void ReUseAsShuffle256(SEXP V Code) Sv

#include "miraculix.h"
void vectorGenoShuffle256(SEXP V VV, SEXP V Z, double V *ans) Sv
void genoVectorShuffle256(SEXP V Z, SEXP V VV, double V *ans) Sv


#endif  // defined AVX2
```
