## Supplementary material for "MoBPS - Modular Breeding Program Simulator": miraculix R-package: shuffle.h

```
/*
 Authors 
 Martin Schlather,


 Copyright (C) 2019 -- 2019  Martin Schlather

This program is free software; you can redistribute it and/or
modify it under the terms of the GNU General Public License
as published by the Free Software Foundation; either version 3
of the License, or (at your option) any later version.

This program is distributed in the hope that it will be useful,
but WITHOUT ANY WARRANTY; without even the implied warranty of
MERCHANTABILITY or FITNESS FOR A PARTICULAR PURPOSE.  See the
GNU General Public License for more details.

You should have received a copy of the GNU General Public License
along with this program; if not, writne to the Free Software
Foundation, Inc., 59 Temple Place - Suite 330, Boston, MA  02111-1307, USA.  
*/


#ifndef miraculix_shuffle_H
#define miraculix_shuffle_H 1

#include "MX.h"


void matrixshuffle_mult(Uint* CGM, Uint snps, Uint individuals, double *ans);
SEXP matrix_start_shuffle(Uint snps,Uint individuals,  SEXP G);
void matrix_shuffle(Uint *M, Uint start_individual, Uint end_individual, 
		    Uint start_snp, Uint end_snp, Uint Mnrow,
		    SEXP Ans, double * G);

SEXP get_matrixshuffle(SEXP SNPxIndiv);
SEXP matrix_coding_shuffle(Uint *M, Uint snps, Uint individuals);

Uint *AlignShuffle(SEXP Code, Uint nr, bool test);
void InitShuffle();
Uint CodesPerBlockShuffle();
Uint UnitsPerIndivShuffle(Uint snps);
void haplo2genoShuffle(Uint * SNPxIndiv, Uint snps, Uint individuals, Uint *A);
Ulong sumGenoShuffle(Uint *S, Uint snps, Uint individuals);

void zeroNthGenoShuffle(SEXP CM, SEXP NN);
SEXP allele_freqShuffle(SEXP GM);
SEXP get_matrixN_shuffle(SEXP CM, SEXP NN);
void ReUseAsShuffle(SEXP Code);
void genoVectorShuffle(SEXP Z, SEXP V, double *ans);
void vectorGenoShuffle(SEXP Z, SEXP V, double *ans);


void matrixshuffle256_mult(Uint* CGM, Uint snps, Uint individuals, double *ans);
SEXP matrix_start_shuffle256(Uint snps, Uint individuals, SEXP G);
void matrix_shuffle256(Uint *M, Uint start_individual, Uint end_individual, 
		    Uint start_snp, Uint end_snp, Uint Mnrow,
		    SEXP Ans, double * G);

SEXP get_matrixshuffle256(SEXP SNPxIndiv);
SEXP matrix_coding_shuffle256(Uint *M, Uint snps, Uint individuals);

Uint *AlignShuffle256(SEXP Code, Uint nr, bool test);
void InitShuffle256();
Uint CodesPerBlockShuffle256();
Uint UnitsPerIndivShuffle256(Uint snps);
void haplo2genoShuffle256(Uint * SNPxIndiv, Uint snps, Uint individuals,
			  Uint *A);
Ulong sumGenoShuffle256(Uint *S, Uint snps, Uint individuals);

void zeroNthGenoShuffle256(SEXP CM, SEXP NN);
SEXP allele_freqShuffle256(SEXP GM);
SEXP get_matrixN_shuffle256(SEXP CM, SEXP NN);
void ReUseAsShuffle256(SEXP Code);
void genoVectorShuffle256(SEXP Z, SEXP V, double *ans);
void vectorGenoShuffle256(SEXP Z, SEXP V, double *ans);


#endif
```
