## Supplementary material for "MoBPS - Modular Breeding Program Simulator": miraculix R-package: shuffleDef.h

```
/*
 Authors 
 Martin Schlather,


 Copyright (C) 2019 -- 2019  Martin Schlather

This program is free software; you can redistribute it and/or
modify it under the terms of the GNU General Public License
as published by the Free Software Foundation; either version 3
of the License, or (at your option) any later version.

This program is distributed in the hope that it will be useful,
but WITHOUT ANY WARRANTY; without even the implied warranty of
MERCHANTABILITY or FITNESS FOR A PARTICULAR PURPOSE.  See the
GNU General Public License for more details.

You should have received a copy of the GNU General Public License
along with this program; if not, writne to the Free Software
Foundation, Inc., 59 Temple Place - Suite 330, Boston, MA  02111-1307, USA.  
*/


#ifndef miraculix_shuffleDef_H
#define miraculix_shuffleDef_H 1


#include "haplogeno.h"
#include "Haplo.h"


// #define INLINE
#define INLINE static inline

#if defined DO_FLOAT
#define real float
#define Real Float
#define ZEROREAL ZEROFLOAT 
//#define INT2REAL INT2FLOAT 
#define MULTREAL MULTFLOAT 
#define ADDREAL ADDFLOAT  
#define SUBREAL SUBFLOAT
#define REALVALUE .f
#define IR(X, Y) INT2FLOAT(f.u128[X], d.u128[Y]);
#else // ! DO_FLOAT
#define real double
#define Real Double
#define ZEROREAL ZERODOUBLE 
//#define INT2REAL INT2DOUBLE 
#define MULTREAL MULTDOUBLE 
#define ADDREAL ADDDOUBLE  
#define SUBREAL SUBDOUBLE  
#define REALVALUE .d
#define IR(X, Y) INT2DOUBLE(f.d128[X], d.m64[Y]);
#endif // not DO_FLOAT

ALL_INLINER

#define VGK					\
	MULTREAL(f REALVALUE, *v, f REALVALUE);			\
	SUBREAL(fcorr, f REALVALUE, corr);		\
	ADDREAL(roughsum, sum, fcorr);		\
	SUBREAL(dummy, roughsum, sum);		\
	SUBREAL(corr, dummy, fcorr);		\
	sum = roughsum

#define VGI							\
	MULTREAL(f REALVALUE, *v, f REALVALUE);			\
	ADDREAL(sum, sum, f REALVALUE);				\


#define ASSERT_INTERN(Shuffle) {					\
    Uint method = GLOBAL.genetics.method;				\
    if (method != Shuffle) { /* will work if aligned: (&& method != TwoBit) */ \
      PRINTF("NOTE: 'RFoptions(snpcoding=%s)' is necessary for this function.\nSo, 'RFoptions' is set accordingly by the package.\n", #Shuffle); \
      GLOBAL.genetics.method = Shuffle;					\
      /* ERR("Set 'RFoptions(snpcoding=Shuffle)' first."); */		\
    }									\
    if (shuffleNotInit) InitIntern();					\
  }

#define ASSERT_INTERN_M(Shuffle) {				\
    Uint method = MethodOf(M);					\
  if (method != Shuffle && method != Haplo)			\
    ERR1("Function only available for method '%10s'\n", #Shuffle);	\
  if (shuffleNotInit) InitIntern();			       \
}


static BlockType and2scalar, xor2scalar, LH;
static real *Pd0 = NULL; 					   


#endif // shuffleDef.h
```
