## Supplementary material for "MoBPS - Modular Breeding Program Simulator": miraculix R-package: shuffleIntern.h

```
/*
 Authors 
 Martin Schlather,


 Copyright (C) 2019 -- 2019  Martin Schlather

This program is free software; you can redistribute it and/or
modify it under the terms of the GNU General Public License
as published by the Free Software Foundation; either version 3
of the License, or (at your option) any later version.

This program is distributed in the hope that it will be useful,
but WITHOUT ANY WARRANTY; without even the implied warranty of
MERCHANTABILITY or FITNESS FOR A PARTICULAR PURPOSE.  See the
GNU General Public License for more details.

You should have received a copy of the GNU General Public License
along with this program; if not, writne to the Free Software
Foundation, Inc., 59 Temple Place - Suite 330, Boston, MA  02111-1307, USA.  
*/


#ifndef miraculix_shuffleIntern_H
#define miraculix_shuffleIntern_H 1

/*
static void showblock(Uint *X, Uint snps) { 
  Uint *x = X,
    bits = snps * BitsPerCode,
    endfor = 1L + (Uint) ((bits - 1L) / BitsPerBlock);
  
  for (Uint d=0; d<endfor; d++, x+=UnitsPerBlock) {
    for (Uint b=0; b<UnitsPerBlock; b++, x++) {
      Uint pattern = (Uint) 0x80000000;
      for (Uint s=0; s<BitsPerUnit; s++) {
        PRINTF("%d", (*x & pattern) != 0);
	pattern >>= 1;
	if (s==15) { PRINTF("."); }
      }
      PRINTF(" ");
    }
  }

  PRINTF("\n");
}


static void showblock(BlockType0 X) { showblock((Uint *) &X, CodesPerBlock); }
static void showblock(BlockType0 X, Uint blocks) {
  showblock((Uint *) &X, blocks * CodesPerBlock);
}

static void showblock(Uint *X, Uint snps, Uint individuals) { 
  for (Uint i=0; i<individuals; i++) {
#ifdef SCHLATHERS_MACHINE    
    // PRINTF("%ld\n", (uint ptr_t) (X + i * Blocks(snps) * UnitsPerBlock));
#endif 
    showblock(X + i * Blocks(snps) * UnitsPerBlock, snps);
  }
}
*/

static Uint permut_finv_g_h[CodesPerBlock];
static bool shuffleNotInit = true;


static void InitIntern() {
  assert(BytesPerBlock == sizeof(BlockType0));
  if (!shuffleNotInit ) BUG;
 
//                   0  1  2  3  4  5  6  7   8  9 10 11 12 13 14 15
//#define andshuffle 0, 1, 4, 0, 1, 2, 5, 0,  4, 5, 8, 0, 0, 0, 0, 0
//#define xorshuffle 0, 0, 0, 2, 0, 0, 0, 2,  0, 0, 0, 2, 2, 2, 2, 4
//    SETREV8(as[i], andshuffle);
//    SETREV8(xs[i], xorshuffle);
    SETREV8(and2scalar, 0, 1, 4, 0, 1, 2, 5, 0,  4, 5, 8, 0, 0, 0, 0, 0);
    SETREV8(xor2scalar, 0, 0, 0, 2, 0, 0, 0, 2,  0, 0, 0, 2, 2, 2, 2, 4);

  SET32(LH, 0x0F0F0F0F);

  // see permutation.tex !
  Uint f[CodesPerBlock], g[CodesPerBlock], h[CodesPerBlock],
    finv[CodesPerBlock];
    
  for (Uint i=0, z=0; i<CodesPerUnit; i++) {
    for (Uint j=0; j<UnitsPerBlock; j++) {
      f[z++] = j * CodesPerUnit + i;
    }
  }
  Ext_orderingInt((int *) f, CodesPerBlock, 1, (int *) finv);

  for (Uint i=0, z=0; i<CodesPerByte; i++) {
    for (Uint j=0; j<BytesPerBlock; j++) {
      g[z++] = j * CodesPerByte + i;
    }
  }
  
  for (Uint k=0, z=0; k<CodesPerByte; k++) {
    for (Uint j=0; j<BytesPerUnit; j++) {
      for (Uint i=0; i<UnitsPerBlock; i++) {
	h[z++] = i * BytesPerUnit + j + k * BytesPerBlock;
      }
    }
  }

  for (Uint k=0; k<CodesPerBlock; k++) {
    permut_finv_g_h[k] = finv[g[h[k]]];
  }
}


static Ulong sumGenoIntern(Uint *S, Uint snps, Uint individuals) { 
 Ulong ans = 0;
 Ulong blocks = (Ulong) Blocks(snps) * individuals;

#define x31  (255L / 8L) // 2 : max{0,1,2}
  //                         4 : 2-bit-coding in 1 Byte
#define BytesPer64  (64 / BitsPerByte)
 //#define intgroup_size (2147483648L / (x31 * 8L * BytesPerUnit)) // 2^31 / x31
#define intgroup_size 3 // 2^31 / x31
#define intgroup_sizeM1 (intgroup_size - 1L)
  Ulong
    intgroupsM1 = (blocks - 1L) / (x31 * intgroup_size),
    sizex31 = intgroup_size * x31;
  //    groupsM1 = (blocks - 1L) / x31;
  BlockType first2bits8, first8bits;
    // NullEins = SET32(0x55555555),
  SET32(first2bits8, 0x03030303);
  SET32(first8bits, 0x000000FF);
  BlockUnitType sum32;
 
  for (Ulong X=0; X<=intgroupsM1; X++) { // this loop needed only for huge
    // data sizes of more than 50000 individuals and 10^5 SNPS each
    ZERO(sum32 VI);
    BlockType *S_int = (BlockType0 *) S + X * sizex31 ;
    Ulong groupsM1 = X < intgroupsM1 ? intgroup_sizeM1
			: (blocks - 1L) / x31 - intgroupsM1 * intgroup_size;
    for (Ulong i=0; i<=groupsM1; i++) {
      BlockType sum,
	*cgm = S_int + i * x31;
      ZERO(sum);
      Uint endx = i < groupsM1 || X < intgroupsM1 ? x31 :
	blocks - intgroupsM1 * sizex31 - x31 * groupsM1;
      for (Uint x=0; x<endx; x++, cgm++) {
	BlockType d,
	  c = *cgm;
	//	d = SHR32(c, 1);
	//	d = AND(NullEins, d);
	//	c = XOR(c, d);      
	for (Uint j=0; j<CodesPerByte; j++) { // for-schleife aufloesen ?
	  AND(d, c, first2bits8);
	  ADD8(sum, sum, d);
	  SHR32(c, c, 2);
	}
      }
      for (Uint j=0; j<BytesPerUnit; j++) {
	BlockType L1;
	AND(L1, sum, first8bits);
	ADD32(sum32 VI, sum32 VI, L1); 
	SHR32(sum, sum, BitsPerByte);
      }
    } // for groups

    for (Uint i=0; i<UnitsPerBlock; i++) {
      ans += (Ulong) sum32.u32[i];
    }
  }

  return ans;
}

static void ReUseAsIntern(SEXP Code, snpcoding method) {
  Uint *info = GetInfo(Code),
    snps = info[SNPS],
    individuals = info[INDIVIDUALS];
  Ulong memInUnitsPerIndiv = Blocks(snps) * UnitsPerBlock;  
  InternReUseAs(Code, method, snps, individuals, memInUnitsPerIndiv, false);  
}

	
static void haplo2genoIntern(Uint *X, Uint snps, Uint individuals, Uint *Ans) {
  // Achtung!! Nach Aufruf muss sumGeno() berechnet werden!!
  Uint blocks = Blocks(snps);
  BlockType EN, NE;
  SET32(EN, 0xAAAAAAAA);
  SET32(NE, 0x55555555); 

  for (Uint j=0; j<individuals; j++) {
    Uint jb = j * blocks;
    BlockType *x = ((BlockType0 *) X)  + jb,
      *a = ((BlockType0 *) Ans)  + jb;
    for (Uint b=0; b<blocks; b++) {
      BlockType y,z,dummy;
      AND(y, x[b], EN);
      AND(z, x[b], NE);
      SHL32(dummy, z, 1L);
      AND(dummy, y, dummy);
      SHR32(y, y, 1);
      XOR(y, y, z);
      
      OR(a[b], dummy, y); 
      // a[b] = OR(SHR32(y, 1L), OR(z, AND(y, SHL32(z, 1L))));
    }
  }
}

static void zeroNthGenoIntern(SEXP CM, SEXP NN) {
  Uint
    first2bits32 = 0x00000003,
    *N = (Uint*) INTEGER(NN),
    lenN = length(NN),
    *info = GetInfo(CM),
    snps = info[SNPS],
    individuals = info[INDIVIDUALS],
    blocks = Blocks(snps),
    *C0 = Align(CM, ALIGN_HAPLO);
  if (info[WHAT] != GENOMATRIX) ERR("not a genomicmatrix coding");
  
  for (Ulong i=0; i<individuals; i++) {    
    BlockType *C1 = ((BlockType0 *) C0) + i * blocks;
    for (Uint ni=0; ni<lenN; ni++) {
      Uint j=N[ni] / CodesPerBlock,
	jj = N[ni] % CodesPerBlock,
	u = jj / UnitsPerBlock,
	k = jj % UnitsPerBlock;     
      BlockType *C = C1 + j;
      ((BlockUnitType*) C)->u32[k] &= ~(first2bits32 << (BitsPerCode * u));
    }
  }
}


static SEXP get_matrixN_intern(SEXP CM, SEXP NN) {
  SEXP Ans = R_NilValue;
  //  BlockType first2bits32 = SET32(0x00000003);
  Uint
    first2bits32 = 0x00000003,
    *N = (Uint*) INTEGER(NN),
    lenN = length(NN),
    *info = GetInfo(CM),
    snps = info[SNPS],
    individuals = info[INDIVIDUALS],
    blocks = Blocks(snps),
    *C0 = Align(CM, ALIGN_HAPLO);
  if (info[WHAT] != GENOMATRIX) ERR("not a genoimatrix coding");
  
  if (lenN ==1) PROTECT(Ans = allocVector(INTSXP, individuals));
  else PROTECT(Ans = allocMatrix(INTSXP, lenN, individuals));
  Uint *M = (Uint*) INTEGER(Ans);

  for (Ulong i=0; i<individuals; i++) {    
    Uint *mm = M + i * lenN;
    for (Uint ni=0; ni<lenN; ni++, mm++) {
      Uint nni = N[ni];
      Uint j=nni / CodesPerBlock,
	jj = nni % CodesPerBlock,
	u = jj / UnitsPerBlock,
	k = jj % UnitsPerBlock;     
      BlockType c = ((BlockType0 *) C0)[i * blocks + j];
      BlockUnitType d;
      d VI = c;
      // d VI = AND(SHR32(c, BitsPerCode * u), first2bits32);  *mm = d.u32[k];
      *mm = (d.u32[k] >> (BitsPerCode * u)) & first2bits32;
    }
  }
 
  UNPROTECT(1);  
  return Ans;
}

#define VGK					\
	MULTREAL(f REALVALUE, *v, f REALVALUE);			\
	SUBREAL(fcorr, f REALVALUE, corr);		\
	ADDREAL(roughsum, sum, fcorr);		\
	SUBREAL(dummy, roughsum, sum);		\
	SUBREAL(corr, dummy, fcorr);		\
	sum = roughsum

#define VGI							\
	MULTREAL(f REALVALUE, *v, f REALVALUE);			\
	ADDREAL(sum, sum, f REALVALUE);				\


static void vGIntern(SEXP V, SEXP Z, double *ans) {
  Uint
    *info = GetInfo(Z),
    snps = info[SNPS],
    individuals = info[INDIVIDUALS],
    len = length(V);
  if (len != snps) ERR("vector 'V' not of correct length");

  real *v0 = (real *) CALLOC(RealAlign(len), sizeof(real)),
    *v = (real*) algnrl(v0);    
  switch(TYPEOF(V)) {
  case INTSXP : {
    ToInt(V);
    for (Uint i=0; i<len; i++) v[i] = (real) Vint[i];
    FREEint(V);
    break;
  }
  case REALSXP : {
    double *vp = REAL(V);
#ifdef DO_FLOAT
    for (Uint i=0; i<len; i++) v[i] = (real) vp[i];
#else
    MEMCOPY(v, vp, len * sizeof(double));
#endif
    break;
  }
  case LGLSXP : {
    Rint *vp = LOGICAL(V);
    for (Uint i=0; i<len; i++) v[i] = (real) vp[i];
    break;
  }
  default : ERR("'V' of unknown type.");
  }

  vectorGenoIntern((Real *) v, (BlockType0 *) Align(Z, ALIGN_VECTOR),
		   snps, individuals, ans);

  FREE(v0);
}


#if defined AVX2 || defined SSSE3

BlockType INLINE VECTORADD(BlockType0 xx, BlockType0 yy) {
  BlockType z, sum, y;  
  AND(z, xx, yy);
  AND(y, z, LH); // 0.25 von 4
  SHUFFLE8(sum, and2scalar, y); // 0.5 von 4

  SHR32(y, z, 4L);
  AND(y, y, LH); // 
  SHUFFLE8(y, and2scalar, y);// 
  ADD8(sum, sum, y); // 
  //   showblock(sum, CodesPerBlock);

#ifndef OLD_VECTORADD
  XOR(z, xx, yy);
  AND(y, z, LH);
  SHUFFLE8(y, xor2scalar, y);
  ADD8(sum, sum, y);

  SHR32(y, z, 4L);
  AND(y, y, LH);
  SHUFFLE8(y, xor2scalar, y);
#else  // FOLGENDES IST NICHT SCHNELLER !!!
  XOR(z, xx, yy);
  SHR32(y, z, 1L);
  AND(z, y, z);
  SHR32(y, z, 3L);
  OR(z, z, y);
  AND(y, z, LH);
  SHUFFLE8(y, xor2scalar, y);
#endif

  ADD8(sum, sum, y);
  return sum;
}


static Uint scalar012(BlockType0 *x, BlockType0 *y, Uint blocks) {
  Uint ans = 0;
#define x15  (255L / 16L) // 4 : max=2^2
  //                         4 : 2-bit-coding in 1 Byte                    
  Uint
    bundles = blocks / x15;

  for (Uint i=0; i<bundles; i++) { 
    BlockType L1, L2, sum,
      *xx = x + i * x15,
      *yy = y + i * x15;
    ZERO(sum);
#if x15 == 15L
    LOAD(L1, xx); xx++; LOAD(L2, yy); yy++; ADD8(sum, sum, VECTORADD(L1, L2));
    LOAD(L1, xx); xx++; LOAD(L2, yy); yy++; ADD8(sum, sum, VECTORADD(L1, L2));
    LOAD(L1, xx); xx++; LOAD(L2, yy); yy++; ADD8(sum, sum, VECTORADD(L1, L2));
    LOAD(L1, xx); xx++; LOAD(L2, yy); yy++; ADD8(sum, sum, VECTORADD(L1, L2));
    LOAD(L1, xx); xx++; LOAD(L2, yy); yy++; ADD8(sum, sum, VECTORADD(L1, L2));

    LOAD(L1, xx); xx++; LOAD(L2, yy); yy++; ADD8(sum, sum, VECTORADD(L1, L2));
    LOAD(L1, xx); xx++; LOAD(L2, yy); yy++; ADD8(sum, sum, VECTORADD(L1, L2));
    LOAD(L1, xx); xx++; LOAD(L2, yy); yy++; ADD8(sum, sum, VECTORADD(L1, L2));
    LOAD(L1, xx); xx++; LOAD(L2, yy); yy++; ADD8(sum, sum, VECTORADD(L1, L2));
    LOAD(L1, xx); xx++; LOAD(L2, yy); yy++; ADD8(sum, sum, VECTORADD(L1, L2));

    LOAD(L1, xx); xx++; LOAD(L2, yy); yy++; ADD8(sum, sum, VECTORADD(L1, L2));
    LOAD(L1, xx); xx++; LOAD(L2, yy); yy++; ADD8(sum, sum, VECTORADD(L1, L2));
    LOAD(L1, xx); xx++; LOAD(L2, yy); yy++; ADD8(sum, sum, VECTORADD(L1, L2));
    LOAD(L1, xx); xx++; LOAD(L2, yy); yy++; ADD8(sum, sum, VECTORADD(L1, L2));
    LOAD(L1, xx); xx++; LOAD(L2, yy); yy++; ADD8(sum, sum, VECTORADD(L1, L2));

#else
    for (Uint j=0; j<x15; j++) {
      LOAD(L1, xx); xx++; LOAD(L2, yy); yy++; ADD8(sum, sum, VECTORADD(L1, L2));
    }
#endif
    ans += ADDUPVECTOR(sum); 
  }

   
  BlockType sum ,
    *endx = x + blocks,
    *xx = x + bundles * x15,
    *yy = y + bundles * x15;
  ZERO(sum);
  for (; xx < endx; xx++, yy++) ADD8(sum, sum, VECTORADD(*xx, *yy));
  ans += ADDUPVECTOR(sum);
  return ans;
}


static void matrix_multIntern(Uint *CGM, Uint snps, Uint individuals,
			      double *ans) {
  Uint blocks = Blocks(snps);
#ifdef DO_PARALLEL
#pragma omp parallel for num_threads(CORES) schedule(dynamic, 16)  
#endif
  for (Uint i=0; i<individuals; i++) {
    for (Uint j=i; j<individuals; j++) {
      Uint Mij = scalar012(((BlockType0 *) CGM) + i * blocks,
			    ((BlockType0 *) CGM) + j * blocks, blocks);
      ans[i + j * individuals] = ans[i * individuals + j] = (double) Mij;
    }
  }
}

#endif // AVX2, SSSE3


static SEXP get_matrixIntern(SEXP CM) { // 0.45 bei 25000 x 25000
  SEXP Ans = R_NilValue;
  BlockType first2bits32;
  SET32(first2bits32, 0x00000003);
  Uint 
    *info = GetInfo(CM),
    snps = info[SNPS],
    individuals = info[INDIVIDUALS],
    bits = snps * BitsPerCode,
    fullBlocks = bits / BitsPerBlock,
    blocks = Blocks(snps),    
    *C0 = Align(CM, ALIGN_HAPLO);
  if (info[WHAT] != GENOMATRIX) ERR("not a geno coding");
   
  PROTECT(Ans = allocMatrix(INTSXP, snps, individuals));
  Uint *M = (Uint*) INTEGER(Ans);
  for (Uint i=0, endfor= snps; i<endfor; i++) M[i] = 0; // noetig?

#ifdef DO_PARALLEL
#pragma omp parallel for num_threads(CORES)   
#endif
  for (Ulong i=0; i<individuals; i++) {
    Uint *mm = M + i * snps;
    BlockType *C = (BlockType0 *) C0 + i * blocks;
    for (Uint j=0; j<fullBlocks; j++, C++) {
      BlockType c = *C;
      for (Uint u=0; u<CodesPerUnit; u++, mm+=UnitsPerBlock) {	  
	BlockUnitType d;
	AND(d VI, c, first2bits32);
#define MD(i) mm[i] = d.u32[i];
#if UnitsPerBlock == 4
	MD(0); MD(1); MD(2); MD(3);
#elif UnitsPerBlock == 8
	MD(0); MD(1); MD(2); MD(3); MD(4); MD(5); MD(6); MD(7);
#else 
	for (Uint k=0; k<UnitsPerBlock; k++) MD(k);
#endif	    
	SHR32(c, c, BitsPerCode);
      }
    }
    if (fullBlocks < blocks) {
      Uint *end = M + (i + 1L) * snps;
      BlockType c = ((BlockType0 *) C0)[i * blocks + fullBlocks];
      for (Uint u=0; u<CodesPerUnit && mm < end; u++) {
	BlockUnitType d;
	AND(d VI, c, first2bits32);
	for (Uint k=0; k<UnitsPerBlock && mm < end; k++, mm++)  *mm = d.u32[k];
	SHR32(c, c, BitsPerCode);
      } 
    }
  }
 
  UNPROTECT(1);
  if (PRESERVE) R_PreserveObject(Ans);

  return Ans;
}

    
static SEXP matrix_start_Intern(Uint snps, Uint individuals, SEXP file){ 
  Uint  memPerIndiv = Blocks(snps) * UnitsPerBlock;
  SEXP Code = CreateEmptyCodeVector(snps, individuals, memPerIndiv);
  PROTECT(Code);

  start_info(Code, file, UnitsPerBlock, CodesPerBlock);
  if (PRESERVE) {BUG; R_PreserveObject(Code);}
  UNPROTECT(1);
  return Code;
}
 


static void codeInnerGeno(Uint *M,
		   Uint snps,
		   Uint *code) {
  Uint j,
    bits = snps * BitsPerCode,
    //fullBlocks = (bits - 1)  / BitsPerBlock;
    fullBlocks =  bits / BitsPerBlock; // 30.5.2019

 
  if (false) {
    for (j=0; j<fullBlocks; j++) {
      Uint *C = code + j * UnitsPerBlock,
	*mm = M + j * CodesPerBlock;
      for (Uint u=0; u<CodesPerUnit; u++) {
	Uint shift = u * 2;
	for (Uint k=0; k<UnitsPerBlock; k++, mm++) {
	  Uint m = *mm;
	  C[k] |= m << shift; 
	}
      }
    }
  } else {
    BlockType0 *Code = (BlockType0 *) code;
    for (j=0; j<fullBlocks; j++) {
      BlockType0 C;
      ZERO(C);
      Uint *mm =  (M + j * CodesPerBlock);
      for (Uint u=0; u<CodesPerUnit; u++, mm+=UnitsPerBlock) {
	Uint shift = u * 2;
	BlockType0 m;
	LOADU(m, (BlockType0 *) mm);
	SHL32(m, m, shift);
	OR(C, C, m);
      }
      Code[j] = C;
    }
  }
  
  Uint snps_done = fullBlocks * CodesPerBlock;
  if (snps_done < snps) {
    Uint *end = M + snps,
      *C = code + UnitsPerBlock * fullBlocks;
    M += snps_done;
    for (Uint u=0; u<CodesPerUnit; u++) {
      Uint shift = u * 2;
      for (Uint k=0; k<UnitsPerBlock && M < end; k++, M++) {
	Uint m = *M;
	//	if (checking && (m < 0 || m > 2)) ERR("wrong values for SNP matrix");
	C[k] |= m << shift;
      }
      if (M == end) break;
    }
  }  
}


static void matrix_Intern(Uint *M, Uint start_individual, Uint end_individual, 
		    Uint start_snp, Uint end_snp, Uint Mnrow, SEXP Ans,
		    double VARIABLE_IS_NOT_USED *G) {
  if (start_snp % CodesPerBlock != 0) BUG;
  Uint
    *info = GetInfo(Ans),
    //    individuals = info[INDIVIDUALS],
    snps = info[SNPS],
    cur_snps = end_snp - start_snp,
    *pM = M,
    *endM = M + Mnrow * (end_individual - start_individual),
    blocks = Blocks(snps);
  Ulong units = (Ulong) blocks * UnitsPerBlock;
  Uint *ans = ((Uint*) Align(Ans, ALIGN_HAPLOGENO)
	       ) + start_snp / CodesPerUnit + start_individual * units;    

  for (; pM<endM; pM += Mnrow, ans += units) {
    codeInnerGeno(pM, cur_snps,ans);
  }
}


static void allele_freq2(BlockType0 *CGM, Uint snps, Uint individuals,
			      double *sumP, double *sumP2) {
  // NOTE! twice allel frequency (indicated by the 2 at the end) 
  double n =  (double) individuals;
  if (snps > 536870912) ERR("too many SNPs");
  Uint blocks = Blocks(snps);

  Uint
    byteblocks = (snps - 1L) / BytesPerBlock + CodesPerByte,
    alignedByteBlocks = byteblocks + 1L,
    units = byteblocks * BytesPerBlock,
    alignedUnits = units + UnitsPerBlock,
    // here: P= 2(p_i), not 2 p_i - 1 == expected values == means
    groupsM1 = (individuals - 1L) / x31; 
  Rint
    *Sums0 = (Rint*) MALLOC(alignedUnits * BytesPerUnit),      
    *sum0 = (Rint*) MALLOC(alignedByteBlocks * BytesPerBlock);// snps + zuschlag
  BlockType first8bits, first2bits8,
    *sum = (BlockType0*) algn(sum0),
    *Sums  = (BlockType0*) algn(Sums0);
  // NullEins = SET32(0x55555555),
  SET32(first8bits, 0x000000FF);
  SET32(first2bits8, 0x03030303);
  if (Sums == NULL || sum == NULL) ERR("allocation error");
  { BlockType zero;
    ZERO(zero);
    Uint endfor = units / UnitsPerBlock;
    for (Uint k=0; k<endfor; Sums[k++] = zero); }
  // parallel funktioniert nicht gut!
  for (Uint i=0; i<=groupsM1; i++) {
    BlockType *cgm = CGM + i * blocks * x31;
    Uint endx = i < groupsM1 ? x31 : individuals - x31 * groupsM1;
    {
      BlockType zero;
      ZERO(zero);
      for (Uint k=0; k < byteblocks; sum[k++] = zero);
    }
    for (Uint x=0; x<endx; x++) {
      BlockType *s = sum;
      for (Uint k=0; k<blocks; k++, cgm++) { // see permutation.tex (*)
	BlockType d,
	  c = *cgm;
       	//	d = SHR32(c, 1);
	//	d = AND(NullEins, d);
	//	c = XOR(c, d);      
	for (Uint j=0; j<CodesPerByte; j++, s++) { // for-schleife aufloesen?
	  AND(d, c, first2bits8);
	  ADD8(*s, *s, d);
	  SHR32(c, c, 2);
	}
      }
    }
    
    //    R < Simianer_sims_23_01.R --vanilla
	 
    BlockType *pS = (BlockType0*) Sums;      
    for (Uint k=0; k<byteblocks; k++) { // see permutation.tex (*)
      BlockType s0 = sum[k];
      for (Uint j=0; j<BytesPerUnit; j++, pS++) {
	BlockType L1;
	AND(L1, s0, first8bits);
	ADD32(*pS, *pS, L1); 
	SHR32(s0, s0, BitsPerByte);
      }
    }
  } // for groups
  FREE(sum0);
  
  Uint *S = (Uint*) Sums,
    alignedreal = byteblocks * BytesPerBlock * sizeof(real) + BytesPerBlock;
  Pd0 = (real*) MALLOC(alignedreal);
  real
    *Pd = (real *) algnrl(Pd0),
    *pPd = Pd;
  double
    sum_Psq = 0.0,
    sum_P = 0.0;
  if (GLOBAL_UTILS->basic.kahanCorrection) {
    double corr = 0.0,
      corrSq = 0.0;
    for (Uint b=0,i=0; b<blocks; b++, pPd += CodesPerBlock) {
      for (Uint c=0; c<CodesPerBlock; c++, i++) {
	double Pi = (double) S[i] / n;
	pPd[permut_finv_g_h[c]] = (real) Pi;

	double fcorr = Pi - corr,
	  roughsum = sum_P + fcorr;
	corr = (roughsum - sum_P) - fcorr;	
	sum_P = roughsum;
	
	fcorr = Pi * Pi - corrSq;
	roughsum = sum_Psq + fcorr;
	corrSq = (roughsum - sum_Psq) - fcorr;	
	sum_Psq = roughsum;
      }
    }
  } else {
    for (Uint b=0,i=0; b<blocks; b++, pPd += CodesPerBlock) {
      for (Uint c=0; c<CodesPerBlock; c++, i++) {
	double Pi = (double) S[i] / n;
	pPd[permut_finv_g_h[c]] = (real) Pi;
	sum_P += Pi;
	sum_Psq += Pi * Pi;
      }
    }
  }
    
  FREE(Sums0);
  if (sumP != NULL) *sumP = sum_P;
  if (sumP2 != NULL) *sumP2 = sum_Psq;
}


static SEXP allele_freqIntern(SEXP GM) {
  SEXP Ans = R_NilValue; 
  Uint 
    *info = GetInfo(GM),
   individuals = info[INDIVIDUALS],
    snps = info[SNPS],
    *cgm = Align(GM, ALIGN_ALLELE);
  if (info[WHAT] != GENOMATRIX) ERR("not a coded Z matrix");
  PROTECT(Ans = allocVector(REALSXP, snps));
  allele_freq2((BlockType0 *) cgm, snps, individuals, NULL,
	       NULL); // Pd0 set as side effect
  real *Pd = (real*) algnrl(Pd0);
  double *ans = REAL(Ans);
  for (Uint i=0; i<snps; i++) ans[i] = 0.5 * (double) Pd[i];
  FREE(Pd0);
  UNPROTECT(1);
  if (PRESERVE) R_PreserveObject(Ans);
  return Ans;
}

#endif // shuffleIntern.h
```
