## Supplementary material for "MoBPS - Modular Breeding Program Simulator": miraculix R-package: thuy.cc

```
/* 
 Authors
 Martin Schlather,

 library for simulation of random fields 

 Copyright (C) 2012 -- 2019 Martin Schlather, 

This program is free software; you can redistribute it and/or
modify it under the terms of the GNU General Public License
as published by the Free Software Foundation; either version 3
of the License, or (at your option) any later version.

This program is distributed in the hope that it will be useful,
but WITHOUT ANY WARRANTY; without even the implied warranty of
MERCHANTABILITY or FITNESS FOR A PARTICULAR PURPOSE.  See the
GNU General Public License for more details.

You should have received a copy of the GNU General Public License
along with this program; if not, write to the Free Software
Foundation, Inc., 59 Temple Place - Suite 330, Boston, MA  02111-1307, USA.
*/

#include <Basic_utils.h>


void intEV(int* x, int* z, int* Len, int *K, int *sumsq, int *n,
	   int* pos) {
  #define every 10000
  int h,diff,
    k = *K,
    len = *Len;

  for (int i=0; i<k; i++) sumsq[i] = n[i] =0;
  for (int i=0; i<len; pos[i++]=0);
  
  for (int i=0; i<len; i++) {
    if ((i % every) == 0) { PRINTF("%d (%d)\n", i / every, len / every); }
    //  print("%d %d\n", i, len);
    for (int j=i+1; j<len; j++) {      
      h = x[j] - x[i];
      if (h >= k) break;    
      diff = z[j] - z[i];
      diff *= diff;
      if (diff > 0) {
	(pos[i])++;
	(pos[j])++;
      }
      sumsq[h] += diff;
      n[h] ++;
    }
  }
  // printf("ok\n");
}
```
