## Supplementary material for "MoBPS - Modular Breeding Program Simulator": miraculix R-package: Vector.matrix.cc

```
/*
 Authors 
 Martin Schlather,


 Copyright (C) 2018 -- 2019  Martin Schlather

This program is free software; you can redistribute it and/or
modify it under the terms of the GNU General Public License
as published by the Free Software Foundation; either version 3
of the License, or (at your option) any later version.

This program is distributed in the hope that it will be useful,
but WITHOUT ANY WARRANTY; without even the implied warranty of
MERCHANTABILITY or FITNESS FOR A PARTICULAR PURPOSE.  See the
GNU General Public License for more details.

You should have received a copy of the GNU General Public License
along with this program; if not, write to the Free Software
Foundation, Inc., 59 Temple Place - Suite 330, Boston, MA  02111-1307, USA.  
*/


#include <R.h>
#include <Rinternals.h>
#include "kleinkram.h"
#include "miraculix.h"
#include <General_utils.h>
#include "MX.h"
#include "xport_import.h"


#define MULTIPLY				\
  for (int j=0; j<c; m += r) {			\
  double sum;					\
  SCALAR_PROD(v, m, r, sum);			\
  a[j++] = sum;					\
}


SEXP vector012matrixXX(SEXP vector, SEXP matrix) {
  int err = NOERROR,
    n = length(vector),
    r = nrows(matrix),
    c = ncols(matrix)
    ;
  if (r != n) ERR("vector and matrix do not match");
  SEXP ans;
  PROTECT(ans = allocVector(REALSXP, c));
  double *m = REAL(matrix),
    *a = REAL(ans);


#define FOR					\
  for(int i=0; i<n; i++)			\
    if (v[i] == 1) idx1[n1++] = i;		\
    else if (v[i] == 2) idx2[n2++] = i;		\
  //  else if (v[i] != 0) {err=1; goto ErrorHandling;}
      
  if (c < 999999) {
    switch (TYPEOF(vector)) {
    case REALSXP : { double *v = REAL(vector);MULTIPLY } break;
    case INTSXP : { int *v = INTEGER(vector); MULTIPLY } break;
    case LGLSXP : { int *v = LOGICAL(vector); MULTIPLY } break;
    default : ERR("vector type incompatible");  
    } 
  } else {
    int
      n1 = 0,
      n2 = 0;
    unsigned int
      *idx1 = (unsigned int*) MALLOC(sizeof(int) * n),
      *idx2 = (unsigned int *) MALLOC(sizeof(int) * n);
    switch (TYPEOF(vector)) {
    case REALSXP : { double *v = REAL(vector); FOR } break;
    case INTSXP  : { int *v = INTEGER(vector); FOR } break;
    case LGLSXP : {
      int *v = LOGICAL(vector);
      for(int i=0; i<n; i++)
	if (v[i] == 1) idx1[n1++] = i;
	else if (v[i] != 0) { err = 1; goto ErrorHandling; }
    }
      break;
    default : err = 2; goto ErrorHandling;
    }
    
    // printf("    martin todo : parallelisieren");
    for (int j=0; j<c; j++) {  
    double *mm = m + j * r,
      sum = 0.0;
      for (int i=0; i<n2; sum += mm[idx2[i++]]);
      sum *= 2;
      for (int i=0; i<n1; sum += mm[idx1[i++]]);
      a[j] = sum;
    }
    
 ErrorHandling:
    FREE(idx1);
    FREE(idx2); 
    
    if (err == 1) { ERR("only 012 allowed for the vector") }
    else if (err == 2) { ERR("unknown type of the vector") }
  }
  UNPROTECT(1);
  return ans;
}


SEXP matrixvector012(SEXP matrix, SEXP vector) {
  int
    n = length(vector),
    r = nrows(matrix),
    c = ncols(matrix)
    ;
  if (c != n) ERR("vector and matrix do not match");
  SEXP ans;
  PROTECT(ans = allocVector(REALSXP, r));
  double *m = REAL(matrix),
    *a = REAL(ans),
    *b = (double *) CALLOC(r, sizeof(double));

  // Martin : todo : replace by SIMD
#define MULTIPLYv							\
  for (int j=0; j<c; j++) {						\
    if (v[j] == 1) for (int i=0; i < r;  m++) a[i++] += *m;		\
    else if (v[j] == 2) for (int i=0; i < r; m++) b[i++] += *m;		\
    else m += r;							\
  }

  for (int j=0; j<r; a[j++] = 0);
  switch (TYPEOF(vector)) {
  case REALSXP : { double *v = REAL(vector);MULTIPLYv } break;
  case INTSXP : { int *v = INTEGER(vector); MULTIPLYv } break;
  case LGLSXP : { int *v = LOGICAL(vector); MULTIPLYv } break;
  default : ERR("vector type incompatible");  
  }

  for (int i=0; i < r; i++) a[i] += 2.0 * b[i];
  FREE(b);
    
  UNPROTECT(1);
  return ans;
}  


SEXP vector012matrix(SEXP vector, SEXP matrix) {
  int err = NOERROR,
    n = length(vector),
    r = nrows(matrix),
    c = ncols(matrix)
    ;
  if (r != n) ERR("vector and matrix do not match");
  SEXP ans;
  PROTECT(ans = allocVector(REALSXP, c));
  double *m = REAL(matrix),
    *a = REAL(ans);


#define FOR2					\
  for(int i=0; i<n; i++)			\
    if (v[i] == 1) idx1[n1++] = i;		\
    else if (v[i] == 2) idx2[n2++] = i;		\
    else if (v[i] != 0) {err=1; goto ErrorHandling;}	\
  int old = idx1[0];					\
  for(int i=1; i<n1; ) {					\
    int dummy = idx1[i];					\
    idx1[i++] -= old;						\
    old = dummy;						\
  }							\
  old = idx2[0];					\
  for(int i=1; i<n2; ) {				\
     int dummy = idx2[i];				\
    idx2[i++] -= old;					\
     old = dummy;					\
  }							\

      
  if (c < 9) {
    switch (TYPEOF(vector)) {
    case REALSXP : { double *v = REAL(vector);MULTIPLY } break;
    case INTSXP : { int *v = INTEGER(vector); MULTIPLY } break;
    case LGLSXP : { int *v = LOGICAL(vector); MULTIPLY } break;
    default : ERR("vector type incompatible");  
    } 
  } else {
    int
      n1 = 0,
      n2 = 0;
    unsigned int
      *idx1 = (unsigned int*) MALLOC(sizeof(int) * n),
      *idx2 = (unsigned int *) MALLOC(sizeof(int) * n);
    switch (TYPEOF(vector)) {
    case REALSXP : { double *v = REAL(vector); FOR2 } break;
    case INTSXP  : { int *v = INTEGER(vector); FOR2 } break;
    case LGLSXP : {
      int *v = LOGICAL(vector);
      for(int i=0; i<n; i++)
	if (v[i] == 1) idx1[n1] = n1 > 0 ? i - idx1[n1-1] : i;
	else if (v[i] != 0) { err = 1; goto ErrorHandling; }
    }
      break;
    default : err = 2; goto ErrorHandling;
    }
    
    // martin todo : SIMD / parallel
    for (int j=0; j<c; j++) {  
      double *mm = m + j * r,
	sum = 0.0;
      for (int i=0; i<n2; ) {
	//	print("%d 2: %d %d %d \n", j, i, idx1[i], mm - m);
	mm += idx2[i++];
	sum += *mm;
      }
      sum *= 2.0;
      mm = m + j * r;
      for (int i=0; i<n1; ) {
	//	print("%d 1: %d %d %d \n", j, i, idx1[i], mm - m);
	mm += idx1[i++];
	sum += *mm;
      }
      a[j] = sum;
    }
    
 ErrorHandling:
    FREE(idx1);
    FREE(idx2); 
    
    if (err == 1) { ERR("only 012 allowed for the vector") }
    else if (err == 2) { ERR("unknown type of the vector") }
  }
  UNPROTECT(1);
  return ans;
}


SEXP IsolveRelMat(Uint individuals, double *Atau, double tau,
		  double *Vec, double beta, bool matrix) {
  const char *info[3] = {"rest", "yhat", "rel.matrix"};
  double
     *A = NULL,
    *pA = NULL;

  if (tau <= 0) ERR("'tau' must be positive");

  Uint elmts = 2 + matrix,
    iP1 = individuals + 1,
    i2 = individuals * individuals;
  SEXP Ans, namevec;
  PROTECT(Ans = allocVector(VECSXP, elmts));
  PROTECT(namevec = allocVector(STRSXP, elmts));
  setAttrib(Ans, R_NamesSymbol, namevec);
  for (Uint k=0; k<elmts; k++) SET_STRING_ELT(namevec, k, mkChar(info[k]));
 
  if (matrix) {
    SEXP RA;
    PROTECT(RA = allocMatrix(REALSXP, individuals, individuals));
    SET_VECTOR_ELT(Ans, 2, RA);
    MEMCOPY(REAL(RA), Atau, sizeof(double) * i2);
    pA = REAL(RA);
  } else {
    A =(double *) MALLOC(sizeof(double) * i2),
    MEMCOPY(A, Atau, sizeof(double) * i2);
    pA = A;
  }
  for (Uint i=0; i<i2; i+=iP1) Atau[i] += tau;

  SEXP rest, yhat;
  PROTECT(rest = allocVector(REALSXP, individuals));
  SET_VECTOR_ELT(Ans, 0, rest);
  double *r = REAL(rest);
  MEMCOPY(r, Vec, sizeof(double) * individuals);
  Rint err = Ext_solvePosDef(Atau, individuals, true, r, 1, NULL, NULL);
  if (err != NOERROR) {
    errorstring_type errorstring;
    Ext_getErrorString(errorstring);
    ERR1("error occurred when solving the system (%.50s)", errorstring);
  }
  PROTECT(yhat = allocVector(REALSXP, individuals));
  SET_VECTOR_ELT(Ans, 1, yhat);
  double *y = REAL(yhat);
  xA(r, pA, individuals, individuals, y);
  FREE(A);
  for (Uint i=0; i<individuals; y[i++] += beta);
  UNPROTECT(4 + matrix);
  if (PRESERVE) R_PreserveObject(Ans);
   
  return(Ans);
}

SEXP solveRelMat(SEXP A, SEXP Tau, SEXP Vec, SEXP Beta, SEXP Destroy) {
  Uint individuals = nrows(A),
      bytes = individuals * individuals * sizeof(double);
  bool destroy = LOGICAL(Destroy)[0];
  double *a = NULL;

  if (!destroy) {
    a = (double *) MALLOC(bytes);
    MEMCOPY(a, REAL(A), bytes);
  }
  SEXP Ans = IsolveRelMat(individuals, destroy ? REAL(A) : a,//no PROTECT( needd
			  REAL(Tau)[0], REAL(Vec), REAL(Beta)[0], false);
  FREE(a);

  return Ans;
}
```
