## Supplementary material for "MoBPS - Modular Breeding Program Simulator": miraculix R-package: windower.cc

```
/*
 Authors 
 Martin Schlather,


 Copyright (C) 2018 -- 2019  Martin Schlather

This program is free software; you can redistribute it and/or
modify it under the terms of the GNU General Public License
as published by the Free Software Foundation; either version 3
of the License, or (at your option) any later version.

This program is distributed in the hope that it will be useful,
but WITHOUT ANY WARRANTY; without even the implied warranty of
MERCHANTABILITY or FITNESS FOR A PARTICULAR PURPOSE.  See the
GNU General Public License for more details.

You should have received a copy of the GNU General Public License
along with this program; if not, write to the Free Software
Foundation, Inc., 59 Temple Place - Suite 330, Boston, MA  02111-1307, USA.  
*/


#include <R.h>
#include <General_utils.h>
#include "miraculix.h"
#include "xport_import.h"
#include "MX.h"


void windower_meanC(int *Init, int *Length, int* Step, int *start, int *ende,
		   double *data, int *Lendata, double *res, int *N){
  Long is, ie, j, win_n, k,
    win_ende = *Init + *Length,
    win_start = *Init,
    step = *Step, 
    lendata = *Lendata,
    n = *N
    ;									
  double mass = 0.0;
  
  for (win_n = is = ie = k = j = 0; j < n; j++) {
    while(ie < lendata && ende[ie] <= win_start) {
      mass -= data[ie++];
      win_n--;
    }
    while(is < lendata && start[is] < win_ende) { // <= win_ende??
      mass += data[is++];
      win_n++;
    }
    // printf("mass %10g %d \n", mass, win_n);
    res[k++] = win_start;
    res[k++] = win_ende;
    res[k++] = win_n == 0 ?  NA_REAL : mass / (double) win_n;
    res[k++] = win_n;
 
    win_ende += step;
    win_start += step;
  }
}
  

SEXP windower_mean(SEXP Init, SEXP Length, SEXP Step, SEXP start, SEXP ende,
		   SEXP data, SEXP Lendata, SEXP N) {
  SEXP res;
  PROTECT(res=allocMatrix(REALSXP, 4,  INTEGER(N)[0]));
  windower_meanC(INTEGER(Init), INTEGER(Length), INTEGER(Step),
		INTEGER(start), INTEGER(ende), REAL(data), INTEGER(Lendata),
		REAL(res), INTEGER(N));
  UNPROTECT(1);
  return res;
}


void windower_minC(int *Init, int *Length, int* Step, int *start, int *ende,
		   double *data, int *Lendata, double *res, int *N){
  Long is, ie, j, win_n, k,
    win_ende = *Init + *Length,
    win_start = *Init,
    step = *Step, 
    lendata = *Lendata,
    n = *N
    ;									
  double min = RF_INF;
  
  for (win_n = is = ie = k = j = 0; j < n; j++) {
    while(is < lendata && start[is] < win_ende) { // <= win_ende??
      double dummy = data[is++];
      if (dummy < min) min = dummy;
      win_n++;
    }
    bool redo = false;
    while(ie < lendata && ende[ie] <= win_start) {
      if (data[ie++] == min) redo = true;
      win_n--;
    }
    if (redo) {
      // 'ie' erste Position; 'is' nach letzter Position
      int i = ie;
      min = i < is ? data[i++] : RF_INF;
      for (; i < is; i++) if (data[i] < min) min = data[i];
    }
    res[k++] = win_start;
    res[k++] = win_ende;
    res[k++] = min;
    res[k++] = win_n;
 
    win_ende += step;
    win_start += step;
  }
}
  
SEXP windower_min(SEXP Init, SEXP Length, SEXP Step, SEXP start, SEXP ende,
		   SEXP data, SEXP Lendata, SEXP N) {
  SEXP res;
  PROTECT(res=allocMatrix(REALSXP, 4,  INTEGER(N)[0]));
  windower_minC(INTEGER(Init), INTEGER(Length), INTEGER(Step),
		INTEGER(start), INTEGER(ende), REAL(data), INTEGER(Lendata),
		REAL(res), INTEGER(N));
  UNPROTECT(1);
  return res;
}

void windower_maxC(int *Init, int *Length, int* Step, int *start, int *ende,
		   double *data, int *Lendata, double *res, int *N){
  Long is, ie, j, win_n, k,
    win_ende = *Init + *Length,
    win_start = *Init,
    step = *Step, 
    lendata = *Lendata,
    n = *N
    ;									
  double max = RF_NEGINF;
  
  for (win_n = is = ie = k = j = 0; j < n; j++) {
    while(is < lendata && start[is] < win_ende) { // <= win_ende??
      double dummy = data[is++];
      if (dummy > max) max = dummy;
      win_n++;
    }
    bool redo = false;
    while(ie < lendata && ende[ie] <= win_start) {
      if (data[ie++] == max) redo = true;
      win_n--;
    }
    if (redo) {
      // 'ie' erste Position; 'is' nach letzter Position
      int i = ie;
      max = i < is ? data[i++] : RF_NEGINF;
      for (; i < is; i++) if (data[i] > max) max = data[i];
    }
    res[k++] = win_start;
    res[k++] = win_ende;
    res[k++] = max;
    res[k++] = win_n;
 
    win_ende += step;
    win_start += step;
  }
}


SEXP windower_max(SEXP Init, SEXP Length, SEXP Step, SEXP start, SEXP ende,
		   SEXP data, SEXP Lendata, SEXP N) {
  SEXP res;
  PROTECT(res=allocMatrix(REALSXP, 4,  INTEGER(N)[0]));
  windower_maxC(INTEGER(Init), INTEGER(Length), INTEGER(Step),
		INTEGER(start), INTEGER(ende), REAL(data), INTEGER(Lendata),
		REAL(res), INTEGER(N));
  UNPROTECT(1);
  return res;
}

void windower_medianC(int *Init, int *Length, int* Step, int *start, int *ende,
		   double *data, int *Lendata, double *res, int *N){
  Long is, ie, j, win_n, k,
    win_ende = *Init + *Length,
    win_start = *Init,
    step = *Step, 
    lendata = *Lendata,
    n = *N
    ;									
  int *pos = (int *) MALLOC(sizeof(int) * *Length);
  if (pos == NULL) ERR("memory allocation error");
  
  for (win_n = is = ie = k = j = 0; j < n; j++) {
    while(ie < lendata && ende[ie] <= win_start) {
      ie++;
      win_n--;
    }
    while(is < lendata && start[is] < win_ende) { // <= win_ende??
      is++;
      win_n++;
    }
    Ext_ordering(data + ie, win_n, 1, pos);
    res[k++] = win_start;
    res[k++] = win_ende;
    res[k++] = win_n % 2 == 1 ? data[ie + pos[(win_n - 1) / 2]] :
      0.5 * (data[ie + pos[win_n / 2 - 1]] + data[ie + pos[win_n / 2]]);
    res[k++] = win_n;
 
    win_ende += step;
    win_start += step;
  }
  FREE(pos);
}

SEXP windower_median(SEXP Init, SEXP Length, SEXP Step, SEXP start, SEXP ende,
		   SEXP data, SEXP Lendata, SEXP N) {
  SEXP res;
  PROTECT(res=allocMatrix(REALSXP, 4,  INTEGER(N)[0]));
  windower_medianC(INTEGER(Init), INTEGER(Length), INTEGER(Step),
		INTEGER(start), INTEGER(ende), REAL(data), INTEGER(Lendata),
		REAL(res), INTEGER(N));
  UNPROTECT(1);
  return res;
}

SEXP windower(SEXP what, SEXP Init, SEXP Length, SEXP Step, SEXP start, SEXP ende,
	      SEXP data, SEXP Lendata, SEXP N) {
  switch(INTEGER(what)[0]) {
  case 1 : return windower_mean(Init, Length, Step, start, ende, data, Lendata, N);
  case 4 : return windower_min(Init, Length, Step, start, ende, data, Lendata, N);
  case 5 : return windower_max(Init, Length, Step, start, ende, data, Lendata, N);    
  case 6 : return windower_median(Init, Length, Step, start, ende, data, Lendata, N);
  default : BUG;
  }
  return R_NilValue;
}
```
