## Supplementary material for "MoBPS - Modular Breeding Program Simulator": miraculix R-package: xport_import.cc

```
/*
 Authors
 Martin Schlather,

 Copyright (C) 2015 -- 2019 Martin Schlather, 

This program is free software; you can redistribute it and/or
modify it under the terms of the GNU General Public License
as published by the Free Software Foundation; either version 3
of the License, or (at your option) any later version.

This program is distributed in the hope that it will be useful,
but WITHOUT ANY WARRANTY; without even the implied warranty of
MERCHANTABILITY or FITNESS FOR A PARTICULAR PURPOSE.  See the
GNU General Public License for more details.

You should have received a copy of the GNU General Public License
along with this program; if not, write to the Free Software
Foundation, Inc., 59 Temple Place - Suite 330, Boston, MA  02111-1307, USA.
*/

#include <R_ext/Rdynload.h>
//#include "RF.h"
#include "xport_import.h"

#define pkg "RandomFieldsUtils"

#ifdef CALL
#undef CALL
#endif
#define CALL(what) what##_type Ext_##what = NULL
UTILSCALLS;

#undef CALL
#define CALL(what) Ext_##what = (what##_type) R_GetCCallable(pkg, #what)
void includeXport() {
  UTILSCALLS;
} // export C

bool ToFalse[1] = { false };
```
