## Supplementary material for "MoBPS - Modular Breeding Program Simulator": miraculix R-package: xport_import.h

```
/*
 Authors 
 Martin Schlather,


 Copyright (C) 2015 -- 2019 Martin Schlather

This program is free software; you can redistribute it and/or
modify it under the terms of the GNU General Public License
as published by the Free Software Foundation; either version 3
of the License, or (at your option) any later version.

This program is distributed in the hope that it will be useful,
but WITHOUT ANY WARRANTY; without even the implied warranty of
MERCHANTABILITY or FITNESS FOR A PARTICULAR PURPOSE.  See the
GNU General Public License for more details.

You should have received a copy of the GNU General Public License
along with this program; if not, write to the Free Software
Foundation, Inc., 59 Temple Place - Suite 330, Boston, MA  02111-1307, USA.  
*/


#ifndef RFxport_H
#define RFxport_H 1

#include <zzz_RandomFieldsUtils.h>

#define UTILSCALLS \
  CALL(getErrorString);				\
  CALL(setErrorLoc);				\
  CALL(getUtilsParam);				\
  CALL(attachRFoptions);			\
  CALL(detachRFoptions);			\
  CALL(ordering);				\
  CALL(orderingInt);				\
  CALL(solvePosDef);				\
  CALL(scalarX);				\
  CALL(ToIntI)				
  //  CALL(scalarInt);			       
  
#ifdef CALL
#undef CALL
#endif
#define CALL(what) extern what##_type Ext_##what
UTILSCALLS;

void includeXport();
extern utilsparam* GLOBAL_UTILS;

// #undef ToReal
// #define ToReal(X) (TYPEOF(X) == REALSXP ? REAL(X) : Ext_ToRealI(X, ToFalse))
//#define ToRealX(X, ) (TYPEOF(X) == REALSXP ? REAL(X) : Ext_ToRealI(X, ToFalse))

// extern bool ToFalse[1];
// #define ToInt(X) Ext_ToIntI(X, ToFalse, false) -- nicht verwenden, da
// parallele Aufrufe !!
//#define ToIntX(X,C) Ext_ToIntI(X, C, false)


#define ToInt(X)				\
  bool create##X = true;			\
  Uint *X##int = (Uint*) Ext_ToIntI(X, &create##X, false)

#define CondToInt(C,X)							\
  bool create##X = C;							\
  Uint *X##int = create##X ? (Uint*) Ext_ToIntI(X, &create##X, false) : NULL;


#define FREEint(X) if (create##X) FREE(X##int)

/*

int *XToIntI(SEXP X, bool *create, bool round);

#define ToInt(X)				\
  bool create##X = true;			\
  Uint *X##int = (Uint*) XToIntI(X, &create##X, false)

#define CondToInt(C,X)							\
  bool create##X = C;							\
  Uint *X##int = create##X ? (Uint*) XToIntI(X, &create##X, false) : NULL;


#define FREEint(X) if (create##X) {BUG; FREE(X##int)}

*/

#endif
```
