## Supplementary material for "MoBPS - Modular Breeding Program Simulator": miraculix R-package: zzz.c

```
/*
 Authors 
 Martin Schlather,

 Copyright (C) 2017 -- 2019 Martin Schlather

This program is free software; you can redistribute it and/or
modify it under the terms of the GNU General Public License
as published by the Free Software Foundation; either version 3
of the License, or (at your option) any later version.

This program is distributed in the hope that it will be useful,
but WITHOUT ANY WARRANTY; without even the implied warranty of
MERCHANTABILITY or FITNESS FOR A PARTICULAR PURPOSE.  See the
GNU General Public License for more details.

You should have received a copy of the GNU General Public License
along with this program; if not, write to the Free Software
Foundation, Inc., 59 Temple Place - Suite 330, Boston, MA  02111-1307, USA.  
*/

#include <R_ext/Rdynload.h> 
#include "miraculix.h"
#include <Basic_utils.h>

#define none 0

void F77_SUB(specific_lines)(int *ntotal, int* nlines, int *whichlines,
			     int *datacols, int *idcols, int *datamat);
void F77_SUB(gmatrix)(double *x, double *y, int * nsnps, int *nind);
void F77_SUB(gmatrix_data)(int *nsnps, int *nind, int *nn, int *runnr,
			   int *ndec, double *y, double *nenner,
			   char *paramfile);
void F77_SUB(gmatrix_data_recoded)(int *nsnps, int *nind, int *nn, int *runnr,
				   int *ndec, double *y, double *nenner,
				   char *paramfile);
				  
#define FDEF(name, n, type) {#name,  (DL_FUNC) &F77_SUB(name), n, type}
/*
static R_NativePrimitiveArgType
  real2int2[] = {REALSXP, REALSXP, INTSXP, INTSXP},
  int5real2char[] = {INTSXP, INTSXP, INTSXP, INTSXP, INTSXP, REALSXP, REALSXP,
		     STRSXP},
  int6[] = {INTSXP, INTSXP, INTSXP, INTSXP, INTSXP, INTSXP}
  ;
*/
static R_FortranMethodDef fortranMethods[] = {
  /* 
  FDEF(gmatrix, 4, real2int2),
  FDEF(gmatrix_data, 8, int5real2char),
  FDEF(gmatrix_data_recoded, 8, int5real2char),
  FDEF(specific_lines, 6, int6),
  */
  {NULL, NULL, 0, none}
};
			    
#define CDEF(name, n, type) {#name, (DL_FUNC) &name, n, type}
//static R_NativePrimitiveArgType
//int1[] = { INTSXP };
//char2int7[] = { CHARSXP, CHARSXP, INTSXP, INTSXP, INTSXP, INTSXP, INTSXP,
//	      INTSXP, INTSXP}; 

static const R_CMethodDef cMethods[]  = {
  CDEF(detachmiraculix, 0, none),
  {NULL, NULL, 0, none}
};

#define CALLDEF(name, n) {#name, (DL_FUNC) &name, n}
static R_CallMethodDef callMethods[]  = {
  // in die respectiven C-Dateien muss adoption.h eingebunden sein
  CALLDEF(loadmiraculix, 0),
  CALLDEF(attachmiraculix, 0),
  
  CALLDEF(scan, 10),
  CALLDEF(sumscan, 10),
  CALLDEF(collect_scan, 10),
  CALLDEF(collect_scan2, 13),
  
  CALLDEF(windower, 9),

  CALLDEF(vector012matrix, 2),
  CALLDEF(matrixvector012, 2),
  CALLDEF(matrix_coding, 1),
  CALLDEF(matrix_get, 1),
  CALLDEF(matrix_mult, 1),
  

  CALLDEF(getAutoCoding, 0),
  CALLDEF(hasSSE2, 0),
  CALLDEF(hasSSSE3, 0),
  CALLDEF(hasAVX, 0),
  CALLDEF(hasAVX2, 0),

  CALLDEF(file_get, 1),
  CALLDEF(file_dot, 2),
  CALLDEF(dot_file, 2),
  
  CALLDEF(codeOrigins, 1),
  CALLDEF(decodeOrigins, 2),
  CALLDEF(codeHaplo, 3),
  CALLDEF(decodeHaplo, 5),
  CALLDEF(haplo2geno, 1),
  CALLDEF(copyGeno, 1),
  CALLDEF(zeroNthGeno, 2),
  CALLDEF(get_matrix_N, 2),
  CALLDEF(rhaplomatrix, 3),
  CALLDEF(rhaplomatrixPart2, 4),

  CALLDEF(createSNPmatrix, 2),
  CALLDEF(fillSNPmatrix, 3),
  CALLDEF(vectorGeno, 2),
  CALLDEF(genoVector, 2),
  CALLDEF(unlock, 1),
  CALLDEF(dolocking, 1),
  CALLDEF(computeSNPS, 8),
  CALLDEF(compute, 9),
  CALLDEF(allele_freq, 1),
  CALLDEF(Debug, 0),
  CALLDEF(StopDebug, 0),
  CALLDEF(solveRelMat, 5),
  CALLDEF(substract_centered, 1),
  CALLDEF(get_centered, 0),
  
  //  CALLDEF(),
 //  CALLDEF(codeSNPs, 2),
  {NULL, NULL, 0}
};


#define CALLABLE(FCTN) R_RegisterCCallable("CHaploBlocker",#FCTN,(DL_FUNC) FCTN)
void R_init_miraculix(DllInfo  *dll) {
  R_registerRoutines(dll, cMethods, // .C
		     callMethods,
		     fortranMethods, // .Fortran
		     NULL); // ext
  R_useDynamicSymbols(dll, FALSE); // OK
  R_forceSymbols(dll, TRUE); // OK
}


void R_unload_miraculix(DllInfo *info) {
  // just to avoid warning from compiler on my computer
#ifdef SCHLATHERS_MACHINE
  if (info == NULL) {};
#endif  
  /* Release resources. */
}
```
