## Supplementary material for "MoBPS - Modular Breeding Program Simulator": RandomFieldsUtils R-package: DESCRIPTION

Package: RandomFieldsUtils
Version: 0.5.9
Title: Utilities for the Simulation and Analysis of Random Fields
Author: Martin Schlather [aut, cre], Reinhard Furrer [ctb], Martin Kroll [ctb], Brian D. Ripley [ctb]
Maintainer: Martin Schlather <>
Depends: R (>= 3.0)
Imports: utils, methods
Description: Various utilities are provided that might be used in spatial statistics and elsewhere. It delivers a method for solving linear equations that checks the sparsity of the matrix before any algorithm is used. In particular the following algorithms are implemented: the pivotised Cholesky decomposition by Harbrecht, Peter and Schneider (2012) <doi:10.1016/j.apnum.2011.10.001>, the Struve functions by MacLeod (1993) <doi:10.1090/S0025-5718-1993-1176713-8>, and a precise algorithm for evaluating the Matern covariance function.
License: GPL ( >= 3)
URL:
        http://ms.math.uni-mannheim.de/de/publications/software/randomfieldsutils
Packaged: 2019-10-28 21:09:49 UTC; schlather
NeedsCompilation: yes
