## Supplementary material for "MoBPS - Modular Breeding Program Simulator": RandomFieldsUtils R-package: AutoRandomFieldsUtils.h

```
#ifndef auto_rfutils_h
#define auto_rfutils_h 1

#include "AutoRandomFieldsUtilsLocal.h"

#define MAXUNITS 4
#define MAXCHAR 18 // max number of characters for (covariance) names  
#define RFOPTIONS "RFoptions"
#define isGLOBAL NA_INTEGER


#define WARN_UNKNOWN_OPTION_ALL 3
#define WARN_UNKNOWN_OPTION_SINGLE 2
#define WARN_UNKNOWN_OPTION_CAPITAL 1
#define WARN_UNKNOWN_OPTION_NONE 0
#define WARN_UNKNOWN_OPTION 10000
#define WARN_UNKNOWN_OPTION_CONDSINGLE ( WARN_UNKNOWN_OPTION_SINGLE - WARN_UNKNOWN_OPTION)
#define WARN_UNKNOWN_OPTION_DEFAULT WARN_UNKNOWN_OPTION_ALL


#endif
```
