## Supplementary material for "MoBPS - Modular Breeding Program Simulator": RandomFieldsUtils R-package: AutoRandomFieldsUtilsLocal.h

```
#ifndef auto_rfutils_local_h
#define auto_rfutils_local_h 1

#define PIVOT_NONE 0  
#define PIVOT_AUTO 1  
#define PIVOT_DO 2    
#define PIVOT_IDX 3    // IDX isn't returned by RFoptions
#define PIVOT_UNDEFINED 4
#define PIVOTLAST PIVOT_UNDEFINED

#define PIVOTSPARSE_MMD 1 // for spam
#define PIVOTSPARSE_RCM 2 // for spam


#endif
```
