## Supplementary material for "MoBPS - Modular Breeding Program Simulator": RandomFieldsUtils R-package: Basic_utils.h

```
/*
 Authors 
 Martin Schlather,


 Copyright (C) 2015 -- 2019  Martin Schlather

This program is free software; you can redistribute it and/or
modify it under the terms of the GNU General Public License
as published by the Free Software Foundation; either version 3
of the License, or (at your option) any later version.

This program is distributed in the hope that it will be useful,
but WITHOUT ANY WARRANTY; without even the implied warranty of
MERCHANTABILITY or FITNESS FOR A PARTICULAR PURPOSE.  See the
GNU General Public License for more details.

You should have received a copy of the GNU General Public License
along with this program; if not, write to the Free Software
Foundation, Inc., 59 Temple Place - Suite 330, Boston, MA  02111-1307, USA.  
*/


#ifndef basic_rfutils_h
#define basic_rfutils_h 1
#ifndef __cplusplus
#include <stdbool.h>
#endif
#include <inttypes.h>
#include <R.h>
#include <Rmath.h>
#include "AutoRandomFieldsUtils.h"


#ifndef DO_PARALLEL_ALREADY_CONSIDERED

#ifdef _OPENMP
#define DO_PARALLEL 1
#else
#ifdef DO_PARALLEL
#undef DO_PARALLEL
#endif
#endif

#ifdef DO_PARALLEL
//#undef DO_PARALLEL
#endif


#endif // DO_PARALLEL_ALREADY_CONSIDERED


//#ifdef WIN32
//#ifdef DO_PARALLEL
//#undef DO_PARALLEL // make a comment to get parallel (part 1, see also part 2)
//#endif
//#endif


#define MULTIMINSIZE(S) ((S) > 20)
// #define MULTIMINSIZE(S) false
// #define MULTIMINSIZE(S) true


#ifndef showfree
#define showfree !true 
#endif


#define DOPRINT true
// 1


//g++ -std=gnu++11 -I"/usr/local/lib/R/include" -DNDEBUG  -I"/home/schlather/R/x86_64-pc-linux-gnu-library/3.6/RandomFieldsUtils/include" -I/usr/local/include  -mavx  -fpic  -I/usr/lib/jvm/java-8-openjdk-amd64/include -I/usr/lib/jvm/java-8-openjdk-amd64/include/linux/ -O2 -fopenmp -pthread -g -fPIC -DNDEBUG -pthread  -c operator.cc -o operator.o
//g++ -std=gnu++11 -I"/usr/local/lib/R/include" -DNDEBUG  -I"/home/schlather/R/x86_64-pc-linux-gnu-library/3.6/RandomFieldsUtils/include" -I/usr/local/include  -mavx  -fpic  -I/usr/lib/jvm/java-8-openjdk-amd64/include -I/usr/lib/jvm/java-8-openjdk-amd64/include/linux/ -O2 -fopenmp -pthread -g -fPIC -DNDEBUG -pthread  -c PoissonPolygon.cc -o PoissonPolygon.o
//


// // 1


#ifdef __cplusplus
extern "C" {
#endif
  void spamcsrdns_(int*, double *, int *, int*, double*);
  void spamdnscsr_(int*, int*, double *, int*, double*, int*, int*, double*);
  void cholstepwise_(int*, int*, double*, int*, int*, int*, int*, int*,
		    int*, int*, int*, int*, int*, int*, double*, int*,
		    int*, int*, int*, int*);
  void backsolves_(int*, int*, int*, int*, int*, double*, int*, int*, int*,
		  int*, double*, double*);
  void calcja_(int*, int*, int*, int*, int*, int*, int*);
  void amuxmat_(int*, int*, int*, double*, double*, double*, int*, int*);
  //  void transpose_(int *, int *, double *, int * int *, double*, int*, int*);
  //  void spamback_();
  //  void spamforward();
#ifdef __cplusplus
}
#endif


typedef enum usr_bool {
  // NOTE: if more options are included, change ExtendedBoolean in
  // userinterface.cc of RandomFields
  False=false, 
  True=true, 
  //Exception=2, // for internal use only
  Nan=INT_MIN
} usr_bool;


#define RF_NA NA_REAL 
#define RF_NAN R_NaN
#define RF_NEGINF R_NegInf
#define RF_INF R_PosInf
#define T_PI M_2_PI

#define OBSOLETENAME "obsolete" 

#define MAXINT 2147483647
#define MININT -2147483647
#define MAXUNSIGNED (MAXINT * 2) + 1
#define INFDIM MAXINT
#define INFTY INFDIM

#define LENGTH length // to avoid the unvoluntiered use of LENGTH defined by R
#define complex Rcomplex
#define DOT "."
#define GAUSS_RANDOM(SIGMA) rnorm(0.0, SIGMA)
#define UNIFORM_RANDOM unif_rand()
#define POISSON_RANDOM(x) rpois(x)
#define SQRT2 M_SQRT2
#define SQRTPI M_SQRT_PI
#define INVPI M_1_PI
#define PIHALF M_PI_2 
#define ONETHIRD 0.333333333333333333333333
#define TWOTHIRD 0.6666666666666666666666667
#define TWOPI 6.283185307179586476925286766559
#define INVLOG2 1.442695040888963
#define INVSQRTTWO 0.70710678118654752440084436210
#define INVSQRTTWOPI 0.39894228040143270286
#define SQRTTWOPI 2.5066282746310002416
#define SQRTINVLOG005 0.5777613700268771079749
//#define LOG05 -0.69314718055994528623
#define LOG3 1.0986122886681096913952452369225257046474905578227
#define LOG2 M_LN2


#define EPSILON     0.00000000001
#define EPSILON1000 0.000000001

#define MIN(A,B) ((A) < (B) ? (A) : (B))
#define MAX(A,B) ((A) > (B) ? (A) : (B))


#define ACOS std::acos
#define ASIN std::asin
#define ATAN std::atan
#define CEIL(X) std::ceil((double) X) // keine Klammern um X!
#define COS std::cos
#define EXP std::exp
#define FABS(X) std::fabs((double) X) // keine Klammern um X!
#define FLOOR std::floor
#define LOG std::log
#define POW(X, Y) R_pow((double) X, (double) Y) // keine Klammern um X!
#define SIGN(X) sign((double) X)
#define SIN std::sin
#define SQRT(X) std::sqrt((double) X)
#define STRCMP(A, B) std::strcmp(A, B)
#define STRCPY(A, B) std::strcpy(A, B)
#define STRLEN std::strlen
#define STRNCMP(A, B, C) std::strncmp(A, B, C)
#define STRNCPY(A, B, N) strcopyN(A, B, N)
#define TAN std::tan
#define MEMCOPYX std::memcpy
#define MEMSET std::memset  
#define AALLOC std::aligned_alloc
#define CALLOCX std::calloc
#define MALLOCX std::malloc
#define FREEX std::free
#define SPRINTF std::sprintf //
#define ROUND(X) ownround((double) X)
#define TRUNC(X) ftrunc((double) X) // keine Klammern um X!
#define QSORT std::qsort


typedef unsigned int Uint;
typedef uint64_t  Ulong;
typedef int64_t  Long;

#define PRINTF Rprintf //
#ifdef SCHLATHERS_MACHINE
#ifdef DO_PARALLEL
#include <omp.h>
#undef PRINTF
#define PRINTF if (omp_get_num_threads() > 1) { error("\n\nnever use Rprintf/PRINTF within parallel constructions!!\n\n"); } else Rprintf // OK
#endif
#endif

#define DOPRINTF if (!DOPRINT) {} else PRINTF
#define print NEVER_USE_print_or_PRINTF_WITHIN_PARALLEL /* // */


#endif
```
