## Supplementary material for "MoBPS - Modular Breeding Program Simulator": RandomFieldsUtils R-package: errors_messages.h

```
/*
 Authors 
 Martin Schlather,


 Copyright (C) 2015 -- 2017 Martin Schlather

This program is free software; you can redistribute it and/or
modify it under the terms of the GNU General Public License
as published by the Free Software Foundation; either version 3
of the License, or (at your option) any later version.

This program is distributed in the hope that it will be useful,
but WITHOUT ANY WARRANTY; without even the implied warranty of
MERCHANTABILITY or FITNESS FOR A PARTICULAR PURPOSE.  See the
GNU General Public License for more details.

You should have received a copy of the GNU General Public License
along with this program; if not, write to the Free Software
Foundation, Inc., 59 Temple Place - Suite 330, Boston, MA  02111-1307, USA.  
*/


// Datei wi

#ifndef rfutils_error_H
#define rfutils_error_H 1


#define NOERROR 0                 
#define ERRORMEMORYALLOCATION 1 
#define ERRORFAILED 2      /* method didn't work for the specified parameters */
#define ERRORNOTPROGRAMMEDYET 3
#define ERRORM 4          /* a single error message */
#define ERRORMEND 12      /* a single error message -- und alles dazwischen */
 


#define LENMSG 1000
#define MAXERRORSTRING 1000
#define nErrorLoc 1000
#define LENERRMSG 1000
typedef char errorstring_type[MAXERRORSTRING];
typedef char errorloc_type[nErrorLoc];


#ifdef DO_PARALLEL
#define LOCAL_ERRMSG2 char MSG2[LENERRMSG]
#ifndef LOCAL_ERRLOC_MSG
#define LOCAL_ERRLOC_MSG errorloc_type ERROR_LOC=""; char ERRMSG[LENERRMSG];
#endif
#ifndef LOCAL_ERRORSTRING
#define LOCAL_ERRORSTRING errorstring_type ERRORSTRING
#endif

#else  // not DO_PARALLEL

#define LOCAL_ERRMSG2
#ifndef LOCAL_ERRLOC_MSG
#define LOCAL_ERRLOC_MSG
#endif
#ifndef LOCAL_ERRORSTRING
#define LOCAL_ERRORSTRING
#endif
extern char ERRMSG[LENERRMSG], // used by Error_utils.h. Never use elsewhere
  MSG2[LENERRMSG];// used at the same time with MSG and ERR()
extern errorstring_type ERRORSTRING; // used by ERRORM in RandomFields

#ifndef ERROR_LOC
extern errorloc_type ERROR_LOC;
#endif
//  extern char   MSG[LENERRMSG]; // used by RandomFields in intermediate steps
#endif

#endif

#ifndef WHICH_ERRORSTRING 
#define WHICH_ERRORSTRING ERRORSTRING
#endif

#ifndef LOCAL_ERROR
#define LOCAL_ERROR(N) {}
#endif


// #define ERRMSG(X) if (PL>=PL_ERRORS){errorMSG(X,MSG); PRINTF("error: %s%s\n",ERROR_LOC,MSG);}


#ifdef SCHLATHERS_MACHINE
#define ERRLINE0 PRINTF("(ERROR in %s, line %d)\n", __FILE__, __LINE__); LOCAL_ERRLOC_MSG
//#define ERRLINE ERRLINE0; LOCAL_ERRMSG2
#else
#define ERRLINE0 LOCAL_ERRLOC_MSG
#endif
#define ERRLINE ERRLINE0; LOCAL_ERRMSG2


#define W_ERRLINE0 char W_ERRMSG[LENERRMSG]
#define W_ERRLINE  char W_MSG2[LENERRMSG]


#define RFERROR error
#define ERR(X) {ERRLINE0;SPRINTF(ERRMSG, "%.90s %.790s",ERROR_LOC,X);RFERROR(ERRMSG);}
#define ERR00(X) ERRLINE;SPRINTF(ERRMSG, "%.90s %.790s", ERROR_LOC, X)
#define ERR1(X, Y) {ERR00(X); SPRINTF(MSG2, ERRMSG, Y); RFERROR(MSG2);}
#define ERR2(X, Y, Z) {ERR00(X); SPRINTF(MSG2, ERRMSG, Y, Z); RFERROR(MSG2);}
#define ERR3(X, Y, Z, A) {ERR00(X); SPRINTF(MSG2, ERRMSG,Y,Z,A); RFERROR(MSG2);}
#define ERR4(X, Y, Z, A, B) {ERR00(X); SPRINTF(MSG2,ERRMSG,Y,Z,A,B);	\
    RFERROR(MSG2);}
#define ERR5(X, Y, Z, A, B, C) {ERR00(X); SPRINTF(MSG2, ERRMSG,Y,Z,A,B,C); \
    RFERROR(MSG2);}
#define ERR6(X, Y, Z, A, B,C,D) {ERR00(X); SPRINTF(MSG2, ERRMSG,Y,Z,A,B,C,D); \
    RFERROR(MSG2);}
#define ERR7(X, Y, Z,A,B,C,D,E) {ERR00(X); SPRINTF(MSG2,ERRMSG,Y,Z,A,B,C,D,E); \
    RFERROR(MSG2);}
#define ERR8(X,Y,Z,A,B,C,D,E,F){ERR00(X);SPRINTF(MSG2,ERRMSG,Y,Z,A,B,C,D,E,F); \
    RFERROR(MSG2);}

#define FERR(X) LOCAL_ERRORSTRING; STRCPY(WHICH_ERRORSTRING, X); DEBUGINFOERR
#define FERR1(X,Y) LOCAL_ERRORSTRING; \
  SPRINTF(WHICH_ERRORSTRING, X, Y); DEBUGINFOERR
#define FERR2(X,Y,Z) LOCAL_ERRORSTRING; \
  SPRINTF(WHICH_ERRORSTRING, X, Y, Z); DEBUGINFOERR
#define FERR3(X,Y,Z,A) LOCAL_ERRORSTRING; \
  SPRINTF(WHICH_ERRORSTRING, X, Y, Z, A); DEBUGINFOERR
#define FERR4(X,Y,Z,A,B) LOCAL_ERRORSTRING; \
  SPRINTF(WHICH_ERRORSTRING,X,Y,Z,A,B); DEBUGINFOERR
#define FERR5(X,Y,Z,A,B,C) LOCAL_ERRORSTRING; \
  SPRINTF(WHICH_ERRORSTRING,X,Y,Z,A,B,C); DEBUGINFOERR 
#define FERR6(X,Y,Z,A,B,C,D) LOCAL_ERRORSTRING; \
  SPRINTF(WHICH_ERRORSTRING,X,Y,Z,A,B,C,D); DEBUGINFOERR 
#define FERR7(X,Y,Z,A,B,C,D,E) LOCAL_ERRORSTRING; \
  SPRINTF(WHICH_ERRORSTRING,X,Y,Z,A,B,C,D,E); DEBUGINFOERR

#define NERR00(N) LOCAL_ERROR(N); return N;
#define NERR(N,X) {FERR(X); NERR00(N)}
#define NERR1(N,X,Y) { FERR1(X, Y); NERR00(N)}
#define NERR2(N,X, Y, Z) { FERR2(X, Y, Z); NERR00(N)}
#define NERR3(N,X, Y, Z, A) { FERR3(X, Y, Z, A); NERR00(N)}
#define NERR4(N,X, Y, Z, A, B) { FERR4(X, Y, Z, A, B); NERR00(N)}
#define NERR5(N,X, Y, Z, A, B, C) { FERR5(X, Y, Z, A, B, C); NERR00(N)}
#define NERR6(N,X, Y, Z, A, B, C, D) { FERR6(X, Y, Z, A,B,C,D); NERR00(N)}
#define NERR7(N,X,Y,Z, A, B, C, D, E) { FERR7(X,Y,Z,A,B,C,D,E); NERR00(N)}

#define SERR(X) NERR(ERRORM, X)
#define SERR1(X,Y) NERR1(ERRORM, X, Y)
#define SERR2(X,Y,Z) NERR2(ERRORM, X, Y, Z)
#define SERR3(X,Y,Z, A) NERR3(ERRORM, X, Y, Z, A)
#define SERR4(X,Y,Z, A, B) NERR4(ERRORM, X, Y, Z, A, B)
#define SERR5(X,Y,Z, A, B, C) NERR5(ERRORM, X, Y, Z, A, B, C)
#define SERR6(X,Y,Z, A, B, C, D) NERR6(ERRORM, X, Y, Z, A, B, C, D)
#define SERR7(X,Y,Z, A, B, C, D, E) NERR7(ERRORM, X, Y, Z, A, B, C, D, E)

#define CERR00 err=ERRORM; continue;
#define CERR(X) { FERR(X); CERR00}
#define CERR1(X,Y) { FERR1(X, Y); CERR00}
#define CERR2(X, Y, Z) { FERR2(X, Y, Z);  CERR00}
#define CERR3(X, Y, Z, A) { FERR3(X, Y, Z, A); CERR00}


#define GERR00 LOCAL_ERROR(ERRORM); err = ERRORM; goto ErrorHandling;
#define GERR(X) {FERR(X); GERR00}
#define GERR1(X,Y) {FERR1(X,Y); GERR00}
#define GERR2(X,Y,Z) {FERR2(X,Y,Z); GERR00}
#define GERR3(X,Y,Z,A) {FERR3(X,Y,Z,A); GERR00}
#define GERR4(X,Y,Z,A,B) {FERR4(X,Y,Z,A,B); GERR00}
#define GERR5(X,Y,Z,A,B,C) {FERR5(X,Y,Z,A,B,C); GERR00}
#define GERR6(X,Y,Z,A,B,C,D) {FERR6(X,Y,Z,A,B,C,D); GERR00}

#define GNERR00(N) err = N; goto ErrorHandling;
#define GNERR(N,X) {FERR(X); GNERR00(N)}
#define GNERR1(N,X,Y) {FERR1(X,Y);GNERR00(N)}
#define GNERR2(N,X,Y,Z) {FERR2(X,Y,Z); GNERR00(N)}
#define GNERR3(N,X,Y,Z,A) {FERR3(X,Y,Z,A); GNERR00(N)}
#define GNERR4(N,X,Y,Z,A,B) {FERR4(X,Y,Z,A,B); GNERR00(N)}
#define GNERR5(N,X,Y,Z,A,B,C) {FERR5(X,Y,Z,A,B,C); GNERR00(N)}
#define GNERR6(N,X,Y,Z,A,B,C,D) {FERR6(X,Y,Z,A,B,C,D); GNERR00(N)}

#define RFWARNING warning
#define warn(X) {RFWARNING(X);}
#define WARN0 warn
#define WARN1(X, Y) {W_ERRLINE; \
    SPRINTF(W_MSG2, X, Y); RFWARNING(W_MSG2);}
#define WARN2(X, Y, Z) {W_ERRLINE; \
    SPRINTF(W_MSG2, X, Y, Z); RFWARNING(W_MSG2);}
#define WARN3(X, Y, Z, A) {W_ERRLINE;\
    SPRINTF(W_MSG2, X, Y, Z, A); RFWARNING(W_MSG2);}
#define WARN4(X, Y, Z, A, B) {W_ERRLINE;	\
    SPRINTF(W_MSG2, X, Y, Z, A, B); RFWARNING(W_MSG2);}
#define WARN5(X, Y, Z, A, B, C) {W_ERRLINE;	\
    SPRINTF(W_MSG2, X, Y, Z, A, B, C); RFWARNING(W_MSG2);}
#define WARN6(X, Y, Z, A, B,C,D) {W_ERRLINE;	\
    SPRINTF(W_MSG2, X, Y, Z, A, B, C, D); RFWARNING(W_MSG2);}
#define WARN7(X, Y, Z,A,B,C,D,E) {W_ERRLINE;	\
    SPRINTF(W_MSG2, X, Y, Z, A, B, C, D, E); RFWARNING(W_MSG2);}


#define RFERROR1(M,A) {errorstring_type ERR_STR; \
    SPRINTF(ERR_STR, M, A); RFERROR(ERR_STR);}
#define RFERROR2(M,A,B) {errorstring_type ERR_STR; \
    SPRINTF(ERR_STR, M, A,B); RFERROR(ERR_STR);}
#define RFERROR3(M,A,B,C) {errorstring_type ERR_STR;\
    SPRINTF(ERR_STR, M, A,B,C); RFERROR(ERR_STR);}
#define RFERROR4(M,A,B,C,D) {errorstring_type ERR_STR; \
    SPRINTF(ERR_STR, M, A,B,C,D); RFERROR(ERR_STR);}
#define RFERROR5(M,A,B,C,D,E) {errorstring_type ERR_STR; \
    SPRINTF(ERR_STR, M, A,B,C,D,E); RFERROR(ERR_STR);}
#define RFERROR6(M,A,B,C,D,E,F) {errorstring_type ERR_STR;\
    SPRINTF(ERR_STR, M, A,B,C,D,E,F); RFERROR(ERR_STR);}
#define RFERROR7(M,A,B,C,D,E,F,G) {errorstring_type ERR_STR;\
    SPRINTF(ERR_STR, M, A,B,C,D,E,F,G); RFERROR(ERR_STR);}
```
