## Supplementary material for "MoBPS - Modular Breeding Program Simulator": RandomFieldsUtils R-package: General_utils.h

```
/*
 Authors 
 Martin Schlather,


 Copyright (C) 2015 -- 2017 Martin Schlather

This program is free software; you can redistribute it and/or
modify it under the terms of the GNU General Public License
as published by the Free Software Foundation; either version 3
of the License, or (at your option) any later version.

This program is distributed in the hope that it will be useful,
but WITHOUT ANY WARRANTY; without even the implied warranty of
MERCHANTABILITY or FITNESS FOR A PARTICULAR PURPOSE.  See the
GNU General Public License for more details.
 
You should have received a copy of the GNU General Public License
along with this program; if not, write to the Free Software
Foundation, Inc., 59 Temple Place - Suite 330, Boston, MA  02111-1307, USA.  
*/


#ifndef rfutils_H
#define rfutils_H 1

#include <R.h>
#include <Rinternals.h>
#include <Rmath.h>
#include <string.h>
#include "Basic_utils.h"
#include "errors_messages.h"
#include "kleinkram.h"
#include "Solve.h"
#include "scalar.h"


#ifdef HIDE_UNUSED_VARIABLE
#define VARIABLE_IS_NOT_USED __attribute__ ((unused))
#else
#ifdef __GNUC__
#define VARIABLE_IS_NOT_USED __attribute__ ((unused))
#else
#define VARIABLE_IS_NOT_USED
#endif
#endif


// not SCHLATHERS_MACHINE
#ifndef SCHLATHERS_MACHINE
#define INTERNAL SERR("Sorry. This functionality doesn't exist currently. There is work in progress at the moment by the maintainer.")
#define assert(X) {}
#define BUG {								\
    RFERROR3("Severe error occured in function '%.50s' (file '%.50s', line %d). Please contact maintainer .", \
	    __FUNCTION__, __FILE__, __LINE__);				\
  }

#define DO_TESTS false
//#define MEMCOPY(A,B,C) {MEMCPY(A,B,C); printf("memcpy %.50s %d\n", __FILE__, __LINE__);}
#define MEMCOPY(A,B,C) MEMCOPYX(A,B,C)
#define AMALLOC(ELEMENTS, SIZE) AALLOC(SIZE, (SIZE) * (ELEMENTS))
#define MALLOC MALLOCX
#define CALLOC CALLOCX
#define XCALLOC CALLOCX
//
#define FREE(X) if ((X) != NULL) {FREEX(X); (X)=NULL;}
//#define FREE(X) if ((X) != NULL) {printf("utils free %.50s %ld Line %d %s\n", #X, (Long) X, __LINE__, __FILE__); FREEX(X); (X)=NULL;}
#define UNCONDFREE(X) {FREEX(X); (X)=NULL;}
#endif // not SCHLATHERS_MACHINE


// SCHLATHERS_MACHINE
#ifdef SCHLATHERS_MACHINE 
#define MAXALLOC 1000000000L

// __extension__ unterdrueckt Fehlermeldung wegen geklammerter Argumente
#define INTERNAL  {		\
    RFERROR3("made to be an internal function '%.50s' ('%.50s', line %d).", \
	     __FUNCTION__, __FILE__, __LINE__);				\
  }

#define assert(X) if (!__extension__ (X)) {	     \
    RFERROR3("'assert' failed in function '%.50s' (%.50s, line %d).", \
	    __FUNCTION__, __FILE__, __LINE__);				\
  }
#define SHOW_ADDRESSES 1
#define BUG { RFERROR2("BUG in '%.50s' line %d.\n",  __FUNCTION__, __LINE__);}
#define DO_TESTS true

#define MEMCOPY(A,B,C) __extension__ ({ assert((A)!=NULL && (B)!=NULL && (C)>0 && (C)<=MAXALLOC); MEMCOPYX(A,B,C); })
//#define MEMCOPY(A,B,C) memory_copy(A, B, C)
#define MALLOC(X) __extension__ ({assert((X)>0 && (X)<=MAXALLOC); MALLOCX(X);})
#define CALLOC(X, Y) __extension__({assert((X)>0 && (X)<=MAXALLOC && (Y)>0 && (Y)<=64); CALLOCX(X,Y);})
#define XCALLOC(X, Y) __extension__({assert((X)>0 && (X)<=MAXALLOC && (Y)>0); CALLOCX(X,Y);})
#define FREE(X) { if ((X) != NULL) {if (showfree) { DOPRINTF("(free in %s, line %d)\n", __FILE__, __LINE__);} FREEX(X); (X)=NULL;}}
#define UNCONDFREE(X) { if (showfree) {DOPRINTF("(free in %s, line %d)\n", __FILE__, __LINE__);} FREEX(X); (X)=NULL;}
#endif // SCHLATHERS_MACHINE


#ifdef RANDOMFIELDS_DEBUGGING
#undef MALLOC
#define MALLOC(X) __extension__({DOPRINTF("(MLLC %s, line %d)\n", __FILE__, __LINE__);assert((X)>0 && (X)<=3e9); MALLOCX(X);})
//
#undef CALLOC
#undef XCALLOC
#define CALLOC(X, Y) __extension__({DOPRINTF("(CLLC %s, line %d)\n",__FILE__, __LINE__);assert((X)>0 && (X)<MAXALLOC && (Y)>0 && (Y)<=216); CALLOCX(X,Y);})
#define XCALLOC(X, Y) __extension__({DOPRINTF("(CLLC %s, line %d)\n",__FILE__, __LINE__);assert((X)>0 && (X)<MAXALLOC && (Y)>0); CALLOCX(X,Y);})
//#define MALLOC malloc
//#define CALLOC calloc


// note that DEBUGINDOERR is redefined in MachineDebugging.h
#define DEBUGINFOERR {						\
    errorstring_type dummy_; STRCPY(dummy_, WHICH_ERRORSTRING);		\
    SPRINTF(WHICH_ERRORSTRING, "%.50s (%.50s, line %d)\n", dummy_, __FILE__, __LINE__); \
  }
#define DEBUGINFO DOPRINTF("(currently at  %s, line %d)\n", __FILE__, __LINE__)

#else
#define DEBUGINFO
#define DEBUGINFOERR if (PL >= PL_ERRORS) {PRINTF("error: %s\n", WHICH_ERRORSTRING);}
#endif


extern int PLoffset;
#define PL_IMPORTANT 1 
#define PL_BRANCHING 2
#define PL_DETAILSUSER 3
#define PL_RECURSIVE 4
#define PL_STRUCTURE 5 // see also initNerror.ERROROUTOFMETHOD
#define PL_ERRORS  6 // only those that are caught internally

#define PL_FCTN_DETAILS 7  // R
#define PL_FCTN_SUBDETAILS 8

#define PL_COV_STRUCTURE 7 // C
#define PL_DIRECT_SEQU 8
#define PL_DETAILS 9
#define PL_SUBDETAILS 10

#define MATERN_NU_THRES 100
#define BESSEL_NU_THRES 100
#define LOW_MATERN 1e-20
#define LOW_BESSEL 1e-20

#ifdef SCHLATHERS_MACHINE 
#define ZZ printf("%s, %s, %d\n",__FUNCTION__,__FILE__,__LINE__) //
#else
#define ZZ
#endif


#endif
```
