## Supplementary material for "MoBPS - Modular Breeding Program Simulator": RandomFieldsUtils R-package: intrinsics.h

```
#ifndef miraculix_initrinsics_H
#define miraculix_initrinsics_H 1

#include <inttypes.h> // uint ptr_t


// PKG_CXXFLAGS =  $(SHLIB_OPENMP_CXXFLAGS) -mavx ODER -march=native 
#ifdef __SSE__
#define SSE __SSE__
#endif
#ifdef  __SSE2__
#define SSE2 __SSE2__
#endif
#ifdef  __SSE3__
#define SSE3 __SSE3__
#endif
#ifdef  __SSSE3__
#define SSSE3 __SSSE3__
#endif
#ifdef  __SSE4A__
#define SSE4A __SSE4A__
#endif
#if defined __SSE41__ || defined __SS42__
#define SSE412 1
#endif
//
#ifdef __AVX__
#define AVX 1
#endif
#ifdef __AVX2__
#define AVX2 1
#endif


#if defined (AVX512)
#define SSEBITS 512
#define SSEMODE 30
#elif defined (SSE)
#define SSEBITS 256
#define SSEMODE 20
#elif defined (SSE)
#define SSEBITS 128
#define SSEMODE 10
#else
#define SSEBITS 64
#define SSEMODE 0
#endif

#ifndef WIN32
// #define FMA_AVAILABLE __FMA__
#endif


#if __GNUC__ > 4 ||							\
  (__GNUC__ == 4 && (__GNUC_MINOR__ > 9 ||				\
		     (__GNUC_MINOR__ == 9 &&  __GNUC_PATCHLEVEL__ >= 1)))
//#define OpenMP4 1
#endif

#ifdef SSE
#include <xmmintrin.h>
#endif

#ifdef SSE2
//#include <emmintrin.h>
#endif

#ifdef SSE3
//#include <pmmintrin.h>
#endif

#ifdef SSSE3
//#include <tmmintrin.h>
#endif

#ifdef SSE4A
//#include <ammintrin.h>
#endif

#ifdef SSE412
//#include <smmintrin.h>
#endif

#if defined AVX || defined AVX2
#include <x86intrin.h>
#endif

#ifdef AVX512
//#include <immintrin.h>
#endif


#if defined AVX
#define BytesPerBlock 32
#define UBlockType __m256i 
#define BlockType __m256i ALIGNED
#define Double __m256d
#define MAXDOUBLE _mm256_max_pd
#define MAXINTEGER _mm256_max_epi32
#define LOAD _mm256_load_si256
// #define EXPDOUBLE mm256_exp_pd // only on intel compiler
#define ADDDOUBLE  _mm256_add_pd
#define SUBDOUBLE  _mm256_sub_pd
#define MULTDOUBLE _mm256_mul_pd 
#define LOADuDOUBLE _mm256_loadu_pd
#define LOADDOUBLE _mm256_load_pd
#define STOREuDOUBLE _mm256_storeu_pd
#define ZERODOUBLE _mm256_setzero_pd()

#elif defined SSE2
#define BytesPerBlock 16
#define UBlockType __m128i
#define BlockType __m128i ALIGNED
#define Double __m128d
#define MAXDOUBLE _mm_max_pd
#define MAXINTEGER _mm_max_epi32
#define LOAD _mm_load_si128
// #define EXPDOUBLE _mm_exp_pd  // only on intel compiler
#define ADDDOUBLE  _mm_add_pd
#define SUBDOUBLE  _mm_sub_pd
#define MULTDOUBLE _mm_mul_pd 
#define LOADuDOUBLE _mm_loadu_pd
#define LOADDOUBLE _mm_load_pd
#define STOREuDOUBLE _mm_storeu_pd
#define ZERODOUBLE _mm_setzero_pd()

#else
#define BytesPerBlock 8
#endif

#define ALIGNED __attribute__ ((aligned (BytesPerBlock)))
#define doubles (BytesPerBlock / 8)
#define integers (BytesPerBlock / 8)

#endif
```
