## Supplementary material for "MoBPS - Modular Breeding Program Simulator": RandomFieldsUtils R-package: linear.h

```
#ifndef LINEAR_RU_H
#define LINEAR_RU_H 1


#define LINEAR_AVX 6
#define LINEAR_BASE 1
#define LINEAR_AVX_PARALLEL 9
#define LINEAR_BASE_PARALLEL 10

void linearX(double *x, double y, int len, double *out, int n);


#endif
```
