## Supplementary material for "MoBPS - Modular Breeding Program Simulator": RandomFieldsUtils R-package: Options_utils.h

```
/*
 Authors 
 Martin Schlather,


 Copyright (C) 2015 -- 2017 Martin Schlather

This program is free software; you can redistribute it and/or
modify it under the terms of the GNU General Public License
as published by the Free Software Foundation; either version 3
of the License, or (at your option) any later version.

This program is distributed in the hope that it will be useful,
but WITHOUT ANY WARRANTY; without even the implied warranty of
MERCHANTABILITY or FITNESS FOR A PARTICULAR PURPOSE.  See the
GNU General Public License for more details.
 
You should have received a copy of the GNU General Public License
along with this program; if not, write to the Free Software
Foundation, Inc., 59 Temple Place - Suite 330, Boston, MA  02111-1307, USA.  
*/


#ifndef rfutils_options_H
#define rfutils_options_H 1

#include <R.h>
#include <Rdefines.h>
#include "Basic_utils.h"
#include "Solve.h"


#define R_PRINTLEVEL 1
#define C_PRINTLEVEL 1
#ifdef SCHLATHERS_MACHINE
#define INITCORES 4
#else
#define INITCORES 1
#endif

extern int PL, CORES;


#define LEN_OPTIONNAME 201

#define basicN 10
// IMPORTANT: all names of basic must be have least 3 letters !!!
extern const char *basic[basicN];
#define BASIC_WARN_OPTION 9
typedef struct basic_param {
  int 
  Rprintlevel,
    Cprintlevel,
    seed, cores, warn_unknown_option;
  bool skipchecks, asList /* hidden:verbose */, kahanCorrection, helpinfo;
} basic_param;
#define basic_START \
  { R_PRINTLEVEL, C_PRINTLEVEL, NA_INTEGER, INITCORES,	 \
      WARN_UNKNOWN_OPTION_DEFAULT,			 \
      false, true, false, true				 \
      }


extern const char * InversionNames[nr_InversionMethods];

#define SOLVE_SVD_TOL 3
#define solveN 20
typedef struct solve_param {
  usr_bool sparse, pivot_check;
  bool det_as_log;
  double spam_tol, spam_min_p, svd_tol, eigen2zero, pivot_relerror,
    max_deviation, max_reldeviation;
  InversionMethod Methods[SOLVE_METHODS];
  int spam_min_n, spam_sample_n, spam_factor, pivotsparse, max_chol,
    max_svd, pivot,
    actual_pivot, actual_size,
    *pivot_idx, pivot_idx_n;//permutation; phys+logi laenge
  //  bool tmp_delete;
} solve_param;
#ifdef SCHLATHERS_MACHINE
#define svd_tol_start 1e-08
#else
#define svd_tol_start 0
#endif
#define solve_START							\
  { Nan, False, true, 							\
    DBL_EPSILON, 0.8, svd_tol_start, 1e-12, 1e-11,			\
    1e-10, 1e-10,							\
  {NoInversionMethod,  NoFurtherInversionMethod},			\
      400, 500, 4294967, PIVOTSPARSE_MMD, 16384,			\
	10000, PIVOT_NONE, /* never change -- see RFoptions.Rd */	\
        PIVOT_UNDEFINED, 0, NULL, 0}
extern const char * solve[solveN];


typedef struct utilsparam{
  basic_param basic;
  solve_param solve;
} utilsparam;


typedef void (*setparameterfct) (int, int, SEXP, char[200], bool, int);
typedef void (*getparameterfct) (SEXP, int, int);
typedef void (*finalsetparameterfct) (int);
typedef void (*deleteparameterfct) (int);
#define ADD(ELT) SET_VECTOR_ELT(sublist, k++, ELT);
#define ADDCHAR(ELT) x[0] = ELT; ADD(ScalarString(mkChar(x)));


#endif
```
