## Supplementary material for "MoBPS - Modular Breeding Program Simulator": RandomFieldsUtils R-package: scalar.h

```
#ifndef SCALAR_RU_H
#define SCALAR_RU_H 1


#define SCALAR_AVX 6
#define SCALAR_KAHAN 8
#define SCALAR_BASE 1
#define SCALAR_AVX_PARALLEL 9
#define SCALAR_BASE_PARALLEL 10

// double scalarX(double *x, double *y, int len, int n);


#endif
```
