## Supplementary material for "MoBPS - Modular Breeding Program Simulator": RandomFieldsUtils R-package: Solve.h

```
/*
 Authors 
 Martin Schlather,


 Copyright (C) 2015 -- 2017 Martin Schlather

This program is free software; you can redistribute it and/or
modify it under the terms of the GNU General Public License
as published by the Free Software Foundation; either version 3
of the License, or (at your option) any later version.

This program is distributed in the hope that it will be useful,
but WITHOUT ANY WARRANTY; without even the implied warranty of
MERCHANTABILITY or FITNESS FOR A PARTICULAR PURPOSE.  See the
GNU General Public License for more details.

You should have received a copy of the GNU General Public License
along with this program; if not, write to the Free Software
Foundation, Inc., 59 Temple Place - Suite 330, Boston, MA  02111-1307, USA.  
*/

#include "errors_messages.h"

#ifndef rfutils_solve_H
#define rfutils_solve_H 1


typedef enum InversionMethod { 
  Cholesky, SVD, Eigen, Sparse,
  NoInversionMethod, // last user available method
  QR, LU, // currently not propagated
  NoFurtherInversionMethod, // local values
  direct_formula, 
  Diagonal // always last one!
} InversionMethod;
#define nr_InversionMethods ((int) Diagonal + 1)
#define nr_user_InversionMethods ((int) NoFurtherInversionMethod + 1)


#define SOLVE_METHODS 3
typedef struct solve_storage {
  errorstring_type err_msg;
  InversionMethod method, newMethods[SOLVE_METHODS];
  usr_bool sparse;
  int SICH_n, MM_n, workspaceD_n, workspaceU_n, VT_n, U_n, D_n, 
    iwork_n, work_n, w2_n, ipiv_n, workLU_n, pivotsparse_n,
    xlnz_n, snode_n, xsuper_n, xlindx_n, invp_n, 
    cols_n, rows_n, DD_n, lindx_n, xja_n,
    diagonal_n,
     lnz_n, w3_n, result_n,
     nsuper, nnzlindx, size, actual_size, actual_pivot,
    *pivot_idx, pivot_idx_n, 
    *iwork, *ipiv,
    *pivotsparse, *xlnz, *snode, *xsuper, *xlindx, 
    *invp, *cols, *rows, *lindx, *xja; //*t_cols, *t_rows;
  double *SICH, *MM, *workspaceD, *workspaceU,
    *VT, *work, *w2, *U, *D, *workLU, *diagonal,
    *lnz, *DD, *w3, *result,
    *to_be_deleted; //, *t_DD;
} solve_storage;

#define SOLVE 0
#define MATRIXSQRT 1
#define DETERMINANT 2

#endif
```
