## Supplementary material for "MoBPS - Modular Breeding Program Simulator": RandomFieldsUtils R-package: Utils.h

```
/*
 Authors 
 Martin Schlather,


 Copyright (C) 2018 -- 2018 Martin Schlather

This program is free software; you can redistribute it and/or
modify it under the terms of the GNU General Public License
as published by the Free Software Foundation; either version 3
of the License, or (at your option) any later version.

This program is distributed in the hope that it will be useful,
but WITHOUT ANY WARRANTY; without even the implied warranty of
MERCHANTABILITY or FITNESS FOR A PARTICULAR PURPOSE.  See the
GNU General Public License for more details.

You should have received a copy of the GNU General Public License
along with this program; if not, write to the Free Software
Foundation, Inc., 59 Temple Place - Suite 330, Boston, MA  02111-1307, USA.  
*/


#ifndef rfutils_utils_H
#define rfutils_utils_H 1

double *ToReal(SEXP X);
int *ToInt(SEXP X);
//double *ToRealI(SEXP X, bool *create);


#ifdef __cplusplus
extern "C" {
#endif 
  int *ToIntI(SEXP X, bool *create, bool round);
 void freeGlobals();
  //  double *ToRealI(SEXP X, bool *create);
  // int *ToIntI(SEXP X, bool *create, bool round);
#ifdef __cplusplus
}
#endif


#endif
```
