## Supplementary material for "MoBPS - Modular Breeding Program Simulator": RandomFieldsUtils R-package: win_linux_aux.h

```
/*
 Authors 
 Martin Schlather,


 Copyright (C) 2015 -- 2017 Martin Schlather

This program is free software; you can redistribute it and/or
modify it under the terms of the GNU General Public License
as published by the Free Software Foundation; either version 3
of the License, or (at your option) any later version.

This program is distributed in the hope that it will be useful,
but WITHOUT ANY WARRANTY; without even the implied warranty of
MERCHANTABILITY or FITNESS FOR A PARTICULAR PURPOSE.  See the
GNU General Public License for more details.

You should have received a copy of the GNU General Public License
along with this program; if not, write to the Free Software
Foundation, Inc., 59 Temple Place - Suite 330, Boston, MA  02111-1307, USA.  
*/


#ifndef WIN_LINUX_AUX_H
#define WIN_LINUX_AUX_H 1

extern "C" void sleepMilli(int *milli);
extern "C" void sleepMicro(int *milli);
extern "C" void pid(int *i);
extern "C" void hostname(char **h, int *i);


#endif /* WIN_LINUX_AUX_H */
```
