## Supplementary material for "MoBPS - Modular Breeding Program Simulator": RandomFieldsUtils R-package: zzz_calls.h

```
/*
 Authors 
 Martin Schlather,


 Copyright (C) 2015 -- 2017 Martin Schlather

This program is free software; you can redistribute it and/or
modify it under the terms of the GNU General Public License
as published by the Free Software Foundation; either version 3
of the License, or (at your option) any later version.

This program is distributed in the hope that it will be useful,
but WITHOUT ANY WARRANTY; without even the implied warranty of
MERCHANTABILITY or FITNESS FOR A PARTICULAR PURPOSE.  See the
GNU General Public License for more details.

You should have received a copy of the GNU General Public License
along with this program; if not, write to the Free Software
Foundation, Inc., 59 Temple Place - Suite 330, Boston, MA  02111-1307, USA.  
*/

#ifndef rfutils_calls_H
#define rfutils_calls_H 1
#include <R_ext/Rdynload.h>

  
#define CALL0(V, N)							\
  attribute_hidden V RU_##N() {						\
    static V(*fun)(AV) = NULL;						\
    if (fun == NULL) fun = (V (*) ()) R_GetCCallable(MY_PACKAGE, #N);	\
    return fun(); }
#define DECLARE0(V, N)							\
  typedef V (*N##_type)();						\
  /* extern N##_type Ext_##N; */					\
  attribute_hidden V RU_##N();						\
  V N();

#define CALL1(V, N, AV, AN)						\
  /* N##_type Ext_##N = NULL; */					\
  attribute_hidden V RU_##N(AV AN) {					\
  static N##_type fun = NULL;						\
  if (fun == NULL) fun = (N##_type) R_GetCCallable(MY_PACKAGE, #N);	\
  return fun(AN); }						
#define DECLARE1(V, N, AV, AN)						\
  typedef V (*N##_type)(AV AN);						\
  /* extern N##_type Ext_##N; */					\
  attribute_hidden V RU_##N(AV AN);					\
  V N(AV AN);

#define CALL2(V, N, AV, AN, BV, BN)					\
  /* N##_type Ext_##N = NULL; */					\
  attribute_hidden V RU_##N(AV AN, BV BN) {			\
  static N##_type fun = NULL;						\
  if (fun == NULL) fun = (N##_type) R_GetCCallable(MY_PACKAGE, #N);	\
  return fun(AN, BN); }					       
#define DECLARE2(V, N, AV, AN, BV, BN)		\
  typedef V (*N##_type)(AV AN, BV BN);	\
  /* extern N##_type Ext_##N; */		\
  attribute_hidden V RU_##N(AV AN, BV BN);	\
  V N(AV AN, BV BN);
  
#define CALL3(V, N, AV, AN, BV, BN, CV, CN)				\
  /* N##_type Ext_##N = NULL; */					\
  attribute_hidden V RU_##N(AV AN, BV BN, CV CN) {		\
  static N##_type fun = NULL;						\
  if (fun == NULL) fun = (N##_type) R_GetCCallable(MY_PACKAGE, #N);	\
  return fun(AN, BN, CN); }						
#define DECLARE3(V, N, AV, AN, BV, BN, CV, CN)				\
  typedef V (*N##_type)(AV AN, BV BN, CV CN);				\
  /* extern N##_type Ext_##N; */					\
  attribute_hidden V RU_##N(AV AN, BV BN, CV CN);			\
  V N(AV AN, BV BN, CV CN);
  
#define CALL4(V, N, AV, AN, BV, BN, CV, CN, DV, DN)			\
  /* N##_type Ext_##N = NULL; */					\
  attribute_hidden V RU_##N(AV AN, BV BN, CV CN, DV DN) {	\
  static N##_type fun = NULL;						\
  if (fun == NULL) fun = (N##_type) R_GetCCallable(MY_PACKAGE, #N);	\
  return fun(AN, BN, CN, DN); }					
#define DECLARE4(V, N, AV, AN, BV, BN, CV, CN, DV, DN)			\
  typedef V (*N##_type)(AV AN, BV BN, CV CN, DV DN);			\
  /* extern N##_type Ext_##N; */					\
  attribute_hidden V RU_##N(AV AN, BV BN, CV CN, DV DN);		\
  V N(AV AN, BV BN, CV CN, DV DN);
  
#define CALL5(V, N, AV, AN, BV, BN, CV, CN, DV, DN, EV, EN)		\
  /* N##_type Ext_##N = NULL; */					\
  attribute_hidden V RU_##N(AV AN, BV BN, CV CN, DV DN, EV EN) {	\
  static N##_type fun = NULL;						\
  if (fun == NULL) fun = (N##_type) R_GetCCallable(MY_PACKAGE, #N);	\
  return fun(AN, BN, CN, DN, EN); }					
#define DECLARE5(V, N, AV, AN, BV, BN, CV, CN, DV, DN, EV, EN)		\
  typedef V (*N##_type)(AV AN, BV BN, CV CN, DV DN, EV EN);		\
  /* extern N##_type Ext_##N; */					\
  attribute_hidden V RU_##N(AV AN, BV BN, CV CN, DV DN, EV EN);		\
  V N(AV AN, BV BN, CV CN, DV DN, EV EN);
  
#define CALL6(V, N, AV, AN, BV, BN, CV, CN, DV, DN, EV, EN, FV, FN)	\
  /* N##_type Ext_##N = NULL; */					\
  attribute_hidden V RU_##N(AV AN, BV BN, CV CN, DV DN, EV EN, FV FN) { \
    static N##_type fun = NULL;						\
      if (fun == NULL) fun = (N##_type) R_GetCCallable(MY_PACKAGE, #N);	\
      return fun(AN, BN, CN, DN, EN, FN); }				
#define DECLARE6(V, N, AV, AN, BV, BN, CV, CN, DV, DN, EV, EN, FV, FN)	\
  typedef V (*N##_type)(AV AN, BV BN, CV CN, DV DN, EV EN, FV FN);	\
  /* extern N##_type Ext_##N; */					\
  attribute_hidden V RU_##N(AV AN, BV BN, CV CN, DV DN, EV EN, FV FN);	\
  V N(AV AN, BV BN, CV CN, DV DN, EV EN, FV FN);
  
#define CALL7(V, N, AV, AN, BV, BN, CV, CN, DV, DN, EV, EN, FV, FN, GV, GN) \
  /* N##_type Ext_##N = NULL; */					\
  attribute_hidden V RU_##N(AV AN, BV BN, CV CN, DV DN, EV EN, FV FN, GV GN) { \
    static N##_type fun = NULL;						\
      if (fun == NULL) fun = (N##_type) R_GetCCallable(MY_PACKAGE, #N);	\
      return fun(AN, BN, CN, DN, EN, FN, GN); }			       
#define DECLARE7(V, N, AV, AN, BV, BN, CV, CN, DV, DN, EV, EN, FV, FN, GV, GN) \
  typedef V (*N##_type)(AV AN, BV BN, CV CN, DV DN, EV EN, FV FN, GV GN); \
  /* extern N##_type Ext_##N; */					\
  attribute_hidden V RU_##N(AV AN, BV BN, CV CN, DV DN, EV EN, FV FN, GV GN); \
  V N(AV AN, BV BN, CV CN, DV DN, EV EN, FV FN, GV GN);
  
#define CALL8(V, N, AV, AN, BV, BN, CV, CN, DV, DN, EV, EN, FV, FN, GV, GN, HV, HN) \
  /* N##_type Ext_##N = NULL; */					\
  attribute_hidden V RU_##N(AV AN, BV BN, CV CN, DV DN, EV EN, FV FN, GV GN, HV HN) { \
  static N##_type fun = NULL;						\
  if (fun == NULL) fun = (N##_type) R_GetCCallable(MY_PACKAGE, #N);	\
  return fun(AN, BN, CN, DN, EN, FN, GN, HN); }		      
#define DECLARE8(V, N, AV, AN, BV, BN, CV, CN, DV, DN, EV, EN, FV, FN, GV, GN, HV, HN) \
  typedef V (*N##_type)(AV AN, BV BN, CV CN, DV DN, EV EN, FV FN, GV GN, HV HN); \
  /* extern N##_type Ext_##N; */					\
  attribute_hidden V RU_##N(AV AN, BV BN, CV CN, DV DN, EV EN, FV FN, GV GN, HV HN); \
  V N(AV AN, BV BN, CV CN, DV DN, EV EN, FV FN, GV GN, HV HN);

#define CALL9(V, N, AV, AN, BV, BN, CV, CN, DV, DN, EV, EN, FV, FN, GV, GN, HV, HN, IV, IN) \
  /* N##_type Ext_##N = NULL; */					\
  attribute_hidden V RU_##N(AV AN, BV BN, CV CN, DV DN, EV EN, FV FN, GV GN, HV HN, IV IN) { \
  static N##_type fun = NULL;						\
  if (fun == NULL) fun = (N##_type) R_GetCCallable(MY_PACKAGE, #N);	\
  return fun(AN, BN, CN, DN, EN, FN, GN, HN, IN); }		      
#define DECLARE9(V, N, AV, AN, BV, BN, CV, CN, DV, DN, EV, EN, FV, FN, GV, GN, HV, HN, IV, IN) \
  typedef V (*N##_type)(AV AN, BV BN, CV CN, DV DN, EV EN, FV FN, GV GN, HV HN, IV IN); \
  /* extern N##_type Ext_##N; */					\
  attribute_hidden V RU_##N(AV AN, BV BN, CV CN, DV DN, EV EN, FV FN, GV GN, HV HN, IV IN); \
  V N(AV AN, BV BN, CV CN, DV DN, EV EN, FV FN, GV GN, HV HN, IV IN);


#define CALL10(V, N, AV, AN, BV, BN, CV, CN, DV, DN, EV, EN, FV, FN, GV, GN, HV, HN, IV, IN, JV, JN) \
  /* N##_type Ext_##N = NULL; */					\
  attribute_hidden V RU_##N(AV AN, BV BN, CV CN, DV DN, EV EN, FV FN, GV GN, HV HN, IV IN, JV JN) { \
  static N##_type fun = NULL;						\
  if (fun == NULL) fun = (N##_type) R_GetCCallable(MY_PACKAGE, #N);	\
  return fun(AN, BN, CN, DN, EN, FN, GN, HN, IN, JN); }		      
#define DECLARE10(V, N, AV, AN, BV, BN, CV, CN, DV, DN, EV, EN, FV, FN, GV, GN, HV, HN, IV, IN, JV, JN) \
  typedef V (*N##_type)(AV AN, BV BN, CV CN, DV DN, EV EN, FV FN, GV GN, HV HN, IV IN, JV JN); \
  /* extern N##_type Ext_##N; */					\
  attribute_hidden V RU_##N(AV AN, BV BN, CV CN, DV DN, EV EN, FV FN, GV GN, HV HN, IV IN, JV JN); \
  V N(AV AN, BV BN, CV CN, DV DN, EV EN, FV FN, GV GN, HV HN, IV IN, JV JN);


#endif
```
