## Supplementary material for "MoBPS - Modular Breeding Program Simulator": RandomFieldsUtils R-package: zzz_RandomFieldsUtils.h

```
/*
 Authors 
 Martin Schlather,


 Copyright (C) 2015 -- 2017 Martin Schlather

This program is free software; you can redistribute it and/or
modify it under the terms of the GNU General Public License
as published by the Free Software Foundation; either version 3
of the License, or (at your option) any later version.

This program is distributed in the hope that it will be useful,
but WITHOUT ANY WARRANTY; without even the implied warranty of
MERCHANTABILITY or FITNESS FOR A PARTICULAR PURPOSE.  See the
GNU General Public License for more details.

You should have received a copy of the GNU General Public License
along with this program; if not, write to the Free Software
Foundation, Inc., 59 Temple Place - Suite 330, Boston, MA  02111-1307, USA.  
*/

#ifndef rfutils_init_H
#define rfutils_init_H 1


 /* 
!!!!! HIER NIE EIN SEXP OBJEKT ZURUECKGEBEN  !!!!  
  */

#include "Options_utils.h"
#include "errors_messages.h"
#include "scalar.h"
#include "Utils.h"


#ifdef HAVE_VISIBILITY_ATTRIBUTE
  # define attribute_hidden __attribute__ ((visibility ("hidden")))
#else
  # define attribute_hidden
#endif

#ifdef __cplusplus
extern "C" {
#endif

#define MY_PACKAGE "RandomFieldsUtils"
#define MY_ACRONYM XX
#include "zzz_calls.h"

  /* 
!!!!! HIER NIE EIN SEXP OBJEKT ZURUECKGEBEN  !!!!  
  */

  
  DECLARE1(void, solve_DELETE, solve_storage**, S)
  DECLARE1(void, solve_NULL, solve_storage*, x)
  DECLARE7(int, solvePosDef, double*, M, int, size, bool, posdef, 
	   double *, rhs, int, rhs_cols, double *, logdet, solve_storage *, PT)
  DECLARE8(int, solvePosDefSp, double *, M, int, size, bool, posdef,
	   double *, rhs, int, rhs_cols, double *,logdet,
	   solve_storage *, Pt, solve_param *,sp)
  DECLARE8(int, solvePosDefResult, double*, M, int, size, bool, posdef, 
	   double *, rhs, int, rhs_cols, double *, result, double*, logdet, 
	   solve_storage*, PT)
  DECLARE4(int, sqrtPosDefFree, double *, M, int, size, solve_storage *, pt,
	   solve_param *, sp)
  DECLARE3(int, sqrtRHS, solve_storage *, pt, double*, RHS, double *, res)
  DECLARE2(int, invertMatrix, double *, M, int, size)
  DECLARE2(double, StruveH, double, x, double, nu)
  DECLARE3(double, StruveL, double, x, double, nu, bool, expScale1d)
  DECLARE1(double, I0mL0, double, x)
  DECLARE3(double, WM, double, x, double, nu, double, factor)
  DECLARE3(double, DWM, double, x, double, nu, double, factor)
  DECLARE3(double, DDWM, double, x, double, nu, double, factor)
  DECLARE3(double, D3WM, double, x, double, nu, double, factor)
  DECLARE3(double, D4WM, double, x, double, nu, double, factor)
  DECLARE4(double, logWM, double, x, double, nu1, double, nu2, double, factor)
  DECLARE1(double, Gauss, double, x)
  DECLARE1(double, DGauss, double, x)
  DECLARE1(double, DDGauss, double, x)
  DECLARE1(double, D3Gauss, double, x)
  DECLARE1(double, D4Gauss, double, x)
  DECLARE1(double, logGauss, double, x)
  
  DECLARE1(void, getErrorString, errorstring_type, errorstring)
  DECLARE1(void, setErrorLoc, errorloc_type, errorloc)
  DECLARE1(void, getUtilsParam, utilsparam **, up)
  DECLARE10(void, attachRFoptions, const char **, prefixlist, int, N, 
	   const char ***, all, int *, allN, setparameterfct, set, 
	   finalsetparameterfct, final, getparameterfct, get,
	    deleteparameterfct, del,
	   int, PLoffset,
	   bool, basicopt)
  DECLARE2(void, detachRFoptions, const char **, prefixlist, int, N)

  DECLARE3(void, sorting, double*, data, int, len, usr_bool, NAlast)
  DECLARE3(void, sortingInt, int*, data, int, len, usr_bool, NAlast)
  DECLARE4(void, ordering, double*, data, int, len, int, dim, int *, pos)
  DECLARE4(void, orderingInt, int*, data, int, len, int, dim, int *, pos)
  DECLARE4(double, scalarX, double *, x, double *, y, int, len, int, n)
  //  DECLARE4(int, scalarInt, int *, x, int *, y, int, len, int, n)
  DECLARE2(double, detPosDef, double *, M, int, size)
  DECLARE8(int, XCinvXdet,double, *M, int, size, double *,X, int, X_cols,
	  double *, XCinvX, double *, det, bool, log, solve_storage, *PT)
  DECLARE10(int, XCinvYdet,double, *M, int, size, bool, posdef,
	    double *, X, double *, Y, int, cols,
	    double *, XCinvY, double *, det, bool, log, solve_storage, *PT)
  //  DECLARE5(double, XCinvXlogdet, double *, M, int, size, double *, X,
  //	   int, X_cols, solve_storage *, PT)
  DECLARE2(bool, is_positive_definite, double *, C, int, dim)
  DECLARE2(void, chol2inv, double *, MPT, int, size)
  DECLARE2(int, chol, double *, MPT, int, size)
  // DECLARE2(double *, ToRealI, SEXP, X, bool *, create)
  DECLARE3(int *, ToIntI, SEXP, X, bool *, create, bool, round)
  DECLARE1(void, pid, int *, i)
  DECLARE1(void, sleepMicro, int *, i)

 /* 
!!!!! HIER NIE EIN SEXP OBJEKT ZURUECKGEBEN  !!!!  
  */
 
  /*

    See in R package RandomFields, /src/userinterfaces.cc 
          CALL#(...)
    at the beginning for how to make the functions available
    in a calling package

  */
#ifdef __cplusplus
}
#endif

#ifdef DO_PARALLEL
#define HAS_PARALLEL true
#else
#define HAS_PARALLEL false
#endif
#if defined AVX2
#define HAS_AVX2 true
#else
#define HAS_AVX2 false
#endif
#if defined AVX
#define HAS_AVX true
#else
#define HAS_AVX false
#endif
#if defined SSSE3
#define HAS_SSSE3 true
#else
#define HAS_SSSE3 false
#endif
#if defined SSE2
#define HAS_SSE2 true
#else
#define HAS_SSE2 false
#endif
#if defined SSE
#define HAS_SSE true
#else
#define HAS_SSE false
#endif


#define AttachMessageN 1000
#define HAS_ONE_RELEVANT (HAS_PARALLEL || (HAS_AVX2 && AVX2_USED) || (HAS_AVX && AVX_USED) || (HAS_SSSE3 && SSSE3_USED) ||  HAS_SSE2 || HAS_SSE)
#define HAS_ALL_RELEVANT (HAS_PARALLEL && (HAS_AVX2 || !AVX2_USED) && (HAS_AVX || !AVX_USED) && (HAS_SSSE3 || !SSSE3_USED) &&  HAS_SSE2 && HAS_SSE)

#define AttachMessage(PKG, HINT)					\
  "'"#PKG"' %.20s %.10s%.10s%.10s%.10s%.10s%.10s%.10s%.10s%.10s%.10s%.10s%.10s%.10s.%.320s%.120s%.120s", \
    HAS_ONE_RELEVANT ? "sees" : "does not see any of",			\
    HAS_PARALLEL ? "OMP, " : "",					\
    HAS_AVX2 && AVX2_USED ? "AVX2, " : "",				\
    HAS_AVX && AVX_USED ? "AVX, " : "",					\
    HAS_SSSE3 && SSSE3_USED ? "SSSE3, " : "",				\
    HAS_SSE2 && SSE2_USED ? "SSE2, " : "",				\
    HAS_SSE && SSE_USED ? "SSE, " : "",					\
    !HAS_ONE_RELEVANT || HAS_ALL_RELEVANT ? "" : "but not ",		\
    !HAS_PARALLEL ? "OMP, " : "",					\
    !HAS_AVX2 && AVX2_USED ? "AVX2" : "",				\
    !HAS_AVX && AVX_USED ? ", AVX" : "",				\
    !HAS_SSSE3 && SSSE3_USED ? ", SSSE3" : "",				\
    !HAS_SSE2 && SSE2_USED ? ", SSE2" : "",				\
    !HAS_SSE && SSE_USED ? ", SSE" : "",				\
    HINT && ((!HAS_AVX2 && AVX2_USED) || (!HAS_AVX && AVX_USED) || !HAS_SSE2) ? "\nNote that without appropriate SIMD the calculations become slow.\nConsider recompiling '"#PKG"' and 'RandomFieldsUtils' with flags e.g.,\n  install.packages(\""#PKG"\", configure.args=\"CXX_FLAGS=-march=native\")" : "", \
    HINT && (!HAS_AVX2 && AVX2_USED) ?					\
    "\n  install.packages(\""#PKG"\", configure.args=\"CXX_FLAGS=-mavx2\")" \
    : "",								\
    HINT && (!HAS_AVX && AVX_USED) ?					\
    "\n  install.packages(\""#PKG"\" , configure.args=\"CXX_FLAGS=-mavx\")"\
    : ""							

#define ReturnAttachMessage(PKG,HINT) 	\
  SEXP Ans = PROTECT(allocVector(STRSXP, 1));	\
  char simd_msg[AttachMessageN];			\
  SPRINTF(simd_msg, AttachMessage(PKG,HINT)); 	\
  SET_STRING_ELT(Ans, 0, mkChar(simd_msg));		\
  UNPROTECT(1);						\
  return Ans;

#endif

      
```
