## Supplementary material for "MoBPS - Modular Breeding Program Simulator": RandomFieldsUtils R-package: aaa_auto.R

```
# This file has been created automatically by 'rfGenerateConstants'


 ## from  src/AutoRandomFieldsUtils.h

 MAXUNITS 	<- as.integer(4)
 MAXCHAR 	<- as.integer(18)
 RFOPTIONS 	<- "RFoptions"
 isGLOBAL 	<- as.integer(NA)

 WARN_UNKNOWN_OPTION_ALL 	<- as.integer(3)
 WARN_UNKNOWN_OPTION_SINGLE 	<- as.integer(2)
 WARN_UNKNOWN_OPTION_CAPITAL 	<- as.integer(1)
 WARN_UNKNOWN_OPTION_NONE 	<- as.integer(0)
 WARN_UNKNOWN_OPTION 	<- as.integer(10000)
 WARN_UNKNOWN_OPTION_CONDSINGLE 	<- as.integer((WARN_UNKNOWN_OPTION_SINGLE-WARN_UNKNOWN_OPTION))
 WARN_UNKNOWN_OPTION_DEFAULT 	<- as.integer(WARN_UNKNOWN_OPTION_ALL)


 ## from  src/AutoRandomFieldsUtilsLocal.h

 PIVOT_NONE 	<- as.integer(0)
 PIVOT_AUTO 	<- as.integer(1)
 PIVOT_DO 	<- as.integer(2)
 PIVOT_IDX 	<- as.integer(3)
 PIVOT_UNDEFINED 	<- as.integer(4)
 PIVOTLAST 	<- as.integer(PIVOT_UNDEFINED)

 PIVOTSPARSE_MMD 	<- as.integer(1)
 PIVOTSPARSE_RCM 	<- as.integer(2)
```
