## Supplementary material for "MoBPS - Modular Breeding Program Simulator": RandomFieldsUtils R-package: internal_use.R

```
checkExamples <- function(exclude=NULL, include=1:length(.fct.list),
                          ask=FALSE, echo=TRUE, halt=FALSE, ignore.all=FALSE,
                          path=package, package="RandomFields",
                          read.rd.files=TRUE, local = TRUE,
                          libpath = NULL, single.runs = FALSE,
                          reset) {
  .exclude <- exclude
  .ask <- ask
  .echo <- echo
  .halt <- halt
  .ignore.all <- ignore.all
  .package <- package
  .path <- path
  .local <- local
  useDynLib <- importClassesFrom <- import <-
  importFrom <- exportClasses <-
  importMethodsFrom <- exportMethods <- S3method <- function(...) NULL
  .env <- new.env()
  stopifnot(is.na(RFoptions()$basic$seed))

  exportPattern <- function(p) { ## necessary to read NAMESPACE??!!
    if (p == "^R\\.") p <- "^R."
    all.pattern <- p %in% c("^[^\\.]", "^[^.]", ".") | get("all.pattern", .env)
    if (!.ignore.all) assign("all.pattern", all.pattern, .env)
    if (all.pattern) return(NULL)
    stopifnot(nchar(p)==3, substr(p,1,1)=="^")
    assign("p", c(get("p", .env), substring(p, 2)), .env)
  }

  export <- function(...) {
    ## code from 'rm'
    dots <- match.call(expand.dots = FALSE)$...
    z <-deparse(substitute(...))
    if (length(dots) && !all(sapply(dots, function(x) is.symbol(x) || 
                                    is.character(x)))) 
      stop("... must contain names or character strings")
    z <- sapply(dots, as.character)
    cat("export", z, "\n")
    assign("export", c(get("export", .env), z), .env)
  }
  import <- importClassesFrom <- importMethodsFrom <- importFrom <- useDynLib <-
    exportClasses <- S3method <- exportMethods <-
      function(...) {
        dots <- match.call(expand.dots = FALSE)$...
       # cat("other:", sapply(dots, as.character), "\n")
      }
  assign("export", NULL, .env)
  assign("all.pattern", FALSE, .env)
  assign("p", NULL, .env)
  
  cat("'source' causes problems in valgrind")
  .content <- readLines(paste(.path, "NAMESPACE", sep="/"), -1)
  eval(parse(text = .content))      
  cat("\tend source\n")
  if (is.logical(read.rd.files) && !read.rd.files) {
    .package.env <- parent.env(.GlobalEnv)
    while (attr(.package.env, "name") != paste("package:", .package, sep="")) {
      .package.env <- parent.env(.package.env)
    }
    .orig.fct.list <- ls(envir=.package.env)

    .ok <- (get("all.pattern", .env) |
            substr(.orig.fct.list, 1, 1) %in% get("p", .env) | 
            .orig.fct.list %in% get("export", .env))
    .fct.list <- .orig.fct.list[.ok]
  } else {
    if (is.logical(read.rd.files))
      .path <- paste("./", .path, "/man", sep="")
    else .path <- read.rd.files
    .files <- dir(.path, pattern="d$")
    .fct.list <- character(length(.files))
    for (i in 1:length(.files)) {
                                        #cat(i, .path, .files[i], "\n")
                                        #if (i == 152) {cat("jumped\n"); next} 
                                        #Print(.path, .files[i])
      #.content <- scan(paste(.path, .files[i], sep="/") , what=character(),
      #                 quiet=TRUE)
      .fn <- paste(.path, .files[i], sep="/") 
      .content <- readLines(.fn, n = 2)
      if (substr(.content[1], 1, 5) != "\\name" &&
          (substr(.content[1], 1, 1) != "%" || substr(.content[2], 1, 5) != "\\name"))
        stop(.files[i], " does not start with '\\name' -- what at least in 2018 has caused problems in valgrind")
      
      .content <- scan(.fn, what=character(), quiet=TRUE)
      .content <- strsplit(.content, "alias\\{")
      .content <- .content[which(lapply(.content, length) > 1)][[1]][2]
      .fct.list[i] <-
        strsplit(strsplit(.content,"\\}")[[1]][1], ",")[[1]][1]
    }
  }

  .include <- if (is.numeric(include)) include else 1:99999
  .include.name <- include
  if (exists("RMexp")) RFoptions(graphics.close_screen = TRUE, graphics.split_screen = TRUE)
  .RFopt <- RFoptions()
  .not_working_no <- .not_working <- NULL
  .included.fl <- .fct.list[.include]
  .not.isna <- !is.na(.included.fl)
  .include <- .include[.not.isna]
  .included.fl <- .included.fl[.not.isna]
  .max.fct.list <- max(.included.fl)
  if (single.runs) {
    file.in <- "example..R"
    file.out <- "example..Rout"
    if (file.exists(file.out)) file.remove(file.out)
  }

  if (is.character(.include.name)) {
    .include.name <- sapply(strsplit(.include.name, ".Rd"), function(x) x[1])
  }

 
  ENVIR <- new.env()
  .allwarnings <- list()
  for (.idx in .include) {
    if (is.character(.include.name) && !(.fct.list[.idx] %in% .include.name))
      next
    try(repeat dev.off(), silent=TRUE)
    if (.idx %in% .exclude) next
    cat("\n\n\n\n\n", .idx, " ", .package, ":", .fct.list[.idx],
        " (total=", length(.fct.list), ") \n", sep="")
     RFoptions(LIST=.RFopt)
    if (!missing(reset)) do.call(reset)
    if (.echo) cat(.idx, "")
    .tryok <- TRUE
    if (single.runs) {
      txt <- paste("library(", package,", ", libpath, "); example(",
		   .fct.list[.idx],
		   ", ask =", .ask,
		   ", echo =", .echo,
		   ")", sep="")
      write(file=file.in,  txt)
      command <- paste("R < ", file.in, ">>", file.out)
    } else {
      ##stopifnot(RFoptions()$basic$print <=2 )
      ##
      .time <- system.time(.res <- try(do.call(utils::example, 
                                               list(.fct.list[.idx], ask=.ask,
					       echo=.echo, local=.local))))
      w <- warnings()
      .allwarnings <- c(.allwarnings, list(c("Help page ", .idx)), w)
      print(w) ## ok
      if (is(.res, "try-error")) {
	if (.halt) {
	  stop("\n\n\t***** ",.fct.list[.idx], " (", .idx,
	       " out of ", max(.include), "). has failed. *****\n\n")
	} else {
	  .not_working_no <- c(.not_working_no, .idx)
	  .not_working <- c(.not_working, .fct.list[.idx])
	  .tryok <- FALSE
	}
      }
      if (exists("RMexp")) RFoptions(storing = FALSE)
      cat("****** '", .fct.list[.idx], "' (", .idx, ") done. ******\n")
      print(.time) #
      if (.tryok && !is.na(RFoptions()$basic$seed)) {
	Print(.not_working, paste(.not_working_no, collapse=", "), #
	      RFoptions()$basic$seed)
	stop("seed not NA: ", .fct.list[.idx])
      }
    }
  }
  Print(.not_working, paste(.not_working_no, collapse=", ")) #
  .ret <- list(.not_working, .not_working_no)
  names(.ret) <- c(.package, "")
  return(.ret)
}


reverse_dependencies_with_maintainers <-
  function(packages, which = c("Depends", "Imports", "LinkingTo"),
           recursive = FALSE) {
    ## function taken from CRAN developer website. 
    repos <- getOption("repos")["CRAN"]
    ## if (substr(repos, 1, 1) == "@") repos <- "http://cran.r-project.org"
    Print(repos) #
    contrib.url(repos, "source") # trigger chooseCRANmirror() if required
    description <- sprintf("%s/web/packages/packages.rds", repos)
    con <- if(substring(description, 1L, 7L) == "file://")
      file(description, "rb")
    else
      url(description, "rb")
    on.exit(close(con))
    db <- readRDS(gzcon(con))
    rownames(db) <- NULL
    
    rdepends <- tools::package_dependencies(packages, db, which,
                                            recursive = recursive,
                                            reverse = TRUE)
    rdepends <- sort(unique(unlist(rdepends)))
    pos <- match(rdepends, db[, "Package"], nomatch = 0L)
    
    db <- db[pos, c("Package", "Version", "Maintainer")]
    if (is.vector(db)) dim(db) <- c(1, length(db))
    db
  }

ShowInstallErrors <-
  function(dir=".", pkgs=unlist(strsplit( dir(pattern="*.Rcheck"), ".Rcheck")))
    for (i in 1:length(pkgs)) {
      cat("\n\n", pkgs[i], "\n")
      for (f in c("00install.out", "00check.log")) {
	system(paste("grep [eE][rR][rR][oO][rR] ", dir, "/",  pkgs[i],
		     ".Rcheck/", f, sep=""))
 	system(paste("grep \"user system elapsed\" -A 2 ", dir, "/",  pkgs[i],
		     ".Rcheck/", f, sep=""))
 ##	system(paste("grep \"Warning messages\" -A 4 ", dir, "/",  pkgs[i],
        ##		     ".Rcheck/", f, sep=""))
### find -type f -name "00*" -exec grep Warning {} \; -print
### find -type f -name "00*" -exec grep "user system elapse" -A 3 {} \; -print

        
        }
    }
  
    

Dependencies <- function(pkgs = all.pkgs, dir = "Dependencies",
                         install = FALSE, check=TRUE, reverse=FALSE,
  			 package="RandomFields") {
  Print(utils::packageDescription(package)) #
  all <- reverse_dependencies_with_maintainers(package #, which="Suggests")
                                               , which="all")
  all.pkgs <- all[, 1]
  PKGS <- paste(all[,1], "_", all[,2], ".tar.gz", sep="")   
  
  ## getOption("repos")["CRAN"]
  URL <- "http://cran.r-project.org/src/contrib/"

  if (install) {
    system(paste("mkdir ", dir))
    system(paste("rm ", dir, "/*tar.gz*", sep=""))
    for (i in 1:length(pkgs)) {
      cat("PACKAGE:", PKGS[i], ":", i, "out of ", length(pkgs),"\n")
      x <- system(paste("(cd ", dir, "; wget ", URL, PKGS[i], ")", sep=""))
      if (x != 0) stop(PKGS[i], "not downloadable")
    ## extended version see RandomFields V 3.0.51 or earlier     
    }
  }
  if (!hasArg("pkgs")) {
    if (check) {
      reverse <- if (reverse) list(repos = getOption("repos")["CRAN"]) else NULL
      tools::check_packages_in_dir(dir=dir, check_args = c("--as-cran", ""),
                                   reverse=reverse)
    }
    ShowInstallErrors(dir, pkgs)
    return(NULL)
  } else { ### old:
    if (check) {
      for (j in 1:length(pkgs)) {
	i <- pmatch(pkgs[j], PKGS)
	if (is.na(i)) next
	command <- paste("(cd ", dir, "; time R CMD check --as-cran", PKGS[i],")")
	Print(command) #
	x <- system(command)
	ShowInstallErrors(dir, pkgs)
	if (x != 0) stop(PKGS[i], "failed")
      }
    }
  }

}
# R Under development (unstable) (2014-12-09 r67142) -- "Unsuffered Consequences"


#  Dependencies(check=FALSE)
```
