## Supplementary material for "MoBPS - Modular Breeding Program Simulator": RandomFieldsUtils R-package: maths.R

```
## Authors 
## Martin Schlather,
##
##
## Copyright (C) 2015 Martin Schlather
##
## This program is free software; you can redistribute it and/or
## modify it under the terms of the GNU General Public License
## as published by the Free Software Foundation; either version 3
## of the License, or (at your option) any later version.
##
## This program is distributed in the hope that it will be useful,
## but WITHOUT ANY WARRANTY; without even the implied warranty of
## MERCHANTABILITY or FITNESS FOR A PARTICULAR PURPOSE.  See the
## GNU General Public License for more details.
##
## You should have received a copy of the GNU General Public License
## along with this program; if not, write to the Free Software
## Foundation, Inc., 59 Temple Place - Suite 330, Boston, MA  02111-1307, USA.  


I0L0 <- function(x) {
  storage.mode(x) <- "double"
#  res <- double(length(x))
  .Call(C_I0ML0, x)
}


.struve <- function(x, nu, sign, expon.scaled) {
  storage.mode(x) <- "double"
  storage.mode(nu) <- "double"
  storage.mode(expon.scaled) <- "logical"
  storage.mode(sign) <- "double"
#  res <- double(max(length(x), length(nu)))
 .Call(C_struve, x, nu, sign, expon.scaled)
}
struveH <- function(x, nu)  .struve(x, nu, -1, FALSE)
struveL <- function(x, nu, expon.scaled=FALSE)  .struve(x, nu, 1, expon.scaled)

wmscale <- function(scaling) {
  switch(scaling,
         whittle = 0.0,
         matern = sqrt(2),
         handcockwallis = 2.0
         )
}

whittle <- function(x, nu, derivative=0,
                   scaling=c("whittle", "matern", "handcockwallis")) {
  .Call(C_WMr, as.double(x), as.double(nu), as.integer(derivative),
        if (is.character(scaling)) wmscale(match.arg(scaling))
        else as.double(scaling))
}

matern <- function(x, nu, derivative=0,
                   scaling=c("matern", "whittle", "handcockwallis")) {
  whittle(x, nu, derivative,
          if (is.character(scaling)) wmscale(match.arg(scaling)) else scaling)
}

 
nonstwm <- function(x, y, nu, log=FALSE,
                    scaling=c("whittle", "matern", "handcockwallis")) {
  if (is.function(nu)) {
    nu1 <- nu(x)
    nu2 <- nu(y)
  } else nu1 <- nu2 <- nu
  L <- .Call(C_logWMr, sqrt(sum((x - y)^2)), as.double(nu1), as.double(nu2),
             if (is.character(scaling)) wmscale(match.arg(scaling))
             else as.double(scaling))
  if (log) L else exp(L)
}
  

gauss <- function(x, derivative=0) {
  .Call(C_gaussr, as.double(x), as.integer(derivative))
}
```
