## Supplementary material for "MoBPS - Modular Breeding Program Simulator": RandomFieldsUtils R-package: RFoptions.R

```
summary.RFopt <- function(object, ...) {  
  object <- lapply(object, function(z) z[order(names(z))])
  object <- object[c(1, 1 + order(names(object[-1])))]
  class(object) <- "summary.RFopt"
  object
}


print.summary.RFopt <- function(x, ...) {
  str(x, give.attr=FALSE, ...) #
  invisible(x)
}

print.RFopt <- function(x, ...) {
  print.summary.RFopt(summary.RFopt(x, ...)) #
  invisible(x)
}

summary.RFoptElmnt <- function(object, ...) {
  object <- object[order(names(object))]
  class(object) <- "summary.RFoptElmt"
  object
}

print.summary.RFoptElmnt <- function(x, ...) {
  str(x, give.attr=FALSE, ...) #
  invisible(x)
}

print.RFoptElmnt <- function(x, ...) {
  print.summary.RFoptElmnt(summary.RFoptElmnt(x, ...)) #
  invisible(x)
}

RFoptions <- function(..., no.readonly=TRUE) {
  opt <- .External(C_RFoptions, ...)
  if (length(opt) == 0) return(invisible(NULL))
  if (is.list(opt[[1]])) {
    opt <- lapply(opt,
		  function(x) {
		    class(x) <- "RFoptElmnt"
		    x
		})
    class(opt) <-  "RFopt"
  } else class(opt) <-  "RFoptElmnt"
  if (!no.readonly) {
    opt$readonly <- list()
  }
  opt
}
```
