## Supplementary material for "MoBPS - Modular Breeding Program Simulator": RandomFieldsUtils R-package: utils.R

```
## Authors 
## Martin Schlather,
##
##
## Copyright (C) 2015 Martin Schlather
##
## This program is free software; you can redistribute it and/or
## modify it under the terms of the GNU General Public License
## as published by the Free Software Foundation; either version 3
## of the License, or (at your option) any later version.
##
## This program is distributed in the hope that it will be useful,
## but WITHOUT ANY WARRANTY; without even the implied warranty of
## MERCHANTABILITY or FITNESS FOR A PARTICULAR PURPOSE.  See the
## GNU General Public License for more details.
##
## You should have received a copy of the GNU General Public License
## along with this program; if not, write to the Free Software
## Foundation, Inc., 59 Temple Place - Suite 330, Boston, MA  02111-1307, USA.  


sleep.milli <- function(n) {
  .C(C_sleepMilli, as.integer(n))
  invisible(NULL)
}

sleep.micro <- function(n) {
  .C(C_sleepMicro, as.integer(n))
  invisible(NULL)
}
	
hostname<-function(){.C(C_hostname, h=paste(seq(0,0,l=100), collapse=""),
                        as.integer(100))$h}

pid <- function() {.C(C_pid, i=integer(1))$i}

LockFile <- function(file, printlevel=RFoptions()$basic$printlevel) {
  PL_ERRORS <- 6
  lock.ext <- ".lock"
  LockFile <- paste(file, lock.ext, sep="")
  if (file.exists(LockFile)) { #2.
    if (printlevel>=PL_ERRORS ) cat("'",file,"' is locked.\n");
    return(2);
  }
  PID <- pid();
  write(file=LockFile,c(PID,hostname()),ncolumns=2,append=TRUE); #3.a.
  Pid <- matrix(scan(LockFile,what=character(0), quiet=TRUE),nrow=2)
  if ((sum(Pid[1,]==PID)!=1) || (sum(Pid[1,]>PID)>0)){ #3.b.
    if (printlevel>PL_ERRORS )
      cat("Lock file of '", file, "' is knocked out.\n");
    return(3);
  }
  return(0);
}

FileExists <- function(file, printlevel=RFoptions()$basic$printlevel) {
    ## for parallel simulation studies: the same data output file should not
  ## be created twice. So:
  ## 1. if file exists then assume another process has done the work already
  ## 2. if file.lock existss then assume another process is doing the work
  ## 3.a. otherwise create file.lock to show other processes that the process
  ##      will do the work
  ## 3.b. check if another process has started with the same work at the same
  ##      time it may happen that in case of simulatenous creation of file.lock
  ##      no process will do the work...(then the lock file will rest.)
  PL_ERRORS <- 6
  if (file.exists(file)) { #1.
    if (printlevel>=PL_ERRORS ) cat("'", file, "' already exists.\n");
    return(1)
  } else {
    return(LockFile(file, printlevel=printlevel))
  }
}

LockRemove <- function(file) {
  ## removes auxiliary files created by FileExists
  lock.ext <- ".lock";
  file.remove(paste0(file, lock.ext))
}


Print <- function(..., digits=6, empty.lines=2) { #
  ## ?"..1"
#  print(..1)
#  print(substitute(..1))
#   print(missing(..100))
   
  max.elements <- 99
  l <- list(...)
  n <- as.character(match.call())[-1]
  cat(paste(rep("\n", empty.lines), collapse="")) #
  for (i in 1:length(l)) {
    cat(n[i]) #
    if (!is.list(l[[i]]) && is.vector(l[[i]])) {
      L <- length(l[[i]])
      if (L==0) cat(" = <zero>")#
      else {
        cat(" [", L, "] = ", sep="")
        cat(if (is.numeric(l[[i]]))
            round(l[[i]][1:min(L , max.elements)], digits=digits)#
            else l[[i]][1:min(L , max.elements)]) #
        if (max.elements < L) cat(" ...")
      }
    } else {
       if (is.list(l[[i]])) {
        cat(" =  ") #
        str(l[[i]], digits.d=digits) #
      } else {
        cat(" =")
        if (length(l[[i]]) <= 100 && FALSE) {
          print(if (is.numeric(l[[i]])) round(l[[i]], digits=digits)#
                else l[[i]])
        } else {
          if (length(l[[i]]) > 1 && !is.vector(l[[i]]) && !is.matrix(l[[i]])
              && !is.array(l[[i]])) cat("\n")
          str(l[[i]]) #
        }
      }
    }
    cat("\n")
  }
}


cholx <- function(a) .Call(C_Chol, a)

cholPosDef <- function() stop("please use 'cholx' instead of 'cholPosDef'.")

solvePosDef <- function(a, b=NULL, logdeterminant=FALSE) {
  stop("please use 'solvex' instead of 'solvePosDef'.")
}

solvex <- function(a, b=NULL, logdeterminant=FALSE) {
  if (logdeterminant) {
    logdet <- double(1)
    res <- .Call(C_SolvePosDef, a, b, logdet)
    return(list(inv=res, logdet=logdet))
  } else {
    .Call(C_SolvePosDef, a, b, double(0))
  }
}

sortx <- function(x, from=1, to=length(x),
                        decreasing=FALSE, na.last = NA) {
  n <- length(x)
 if (n <= 4000 || (to - from) < (0.35 + is.double(x) * 0.15) * n) {   
    if (decreasing) {
      x <- -x
      if (!is.na(na.last)) na.last <- !na.last
    }
    ans <- .Call(C_sortX, x, as.integer(from), as.integer(to),
                 as.logical(na.last))
    return(if (decreasing) -ans else ans)
 } else {
    return(if (from==1 && to==n)
           sort(x, decreasing=decreasing, na.last=na.last) else
           sort(x, decreasing=decreasing, na.last=na.last)[from:to])
  }
}


orderx <- function(x, from=1, to=length(x),
                         decreasing=FALSE, na.last = NA) {
#  cat((to - from) *  (0.35 + 0.14 * log(length(x)))^2, "", length(x), "\n")
  if ((to - from) * (0.35 + 0.14 * log(length(x)))^2 > length(x)) { #10^2:1, 10^3:1.5, 10^4:3 10^5:5 10^6:5, 10^7: 8, 10^8:10,
    #    cat("old", from, to ,"\n");
    ans <- order(x, decreasing=decreasing, na.last=na.last)
    return(if (from==1 && to==length(x)) ans else ans[from:to])
  }
  if (decreasing) {
    x <- -x
    if (!is.na(na.last)) na.last <- !na.last
  }

  .Call(C_orderX, x, as.integer(from), as.integer(to),
        as.logical(na.last))
}


# scalar <- function(x, y, mode="1x1") .Call(C_scalarX, x, y, mode)


confirm <- function(x, y, ...) {
  e <- all.equal(x, y, ...)
  if (is.logical(e) && e) {
    cat("'", deparse(substitute(x)) , "' and '", deparse(substitute(y)),
        "' are the same.\n", sep="")
  } else {
    if (R.Version()$os=="linux-gnu") stop(e)
    else {
      message(x)
      cat("(under linux systems they are the same.)")
      return(FALSE)
    }
  }
}

chol2mv <- function(C, n) .Call(C_chol2mv, C, as.integer(n))
tcholRHS <- function(C, RHS) {
  if (!is.double(RHS)) storage.mode(RHS) <- "double"
  .Call(C_tcholRHS, C, RHS)
}
colMax <- function(x) .Call(C_colMaxs, x)
rowMeansx <- function(x, weight=NULL) .Call(C_rowMeansX, x, weight)
rowProd <- function(x) .Call(C_rowProd, x)
SelfDivByRow <- function(x, v) .Call(C_DivByRow, x, v)
quadratic <- function(x, v) .Call(C_quadratic, v, x)
dotXV <- function(x, w) .Call(C_dotXV, x, w)

dbinorm <- function(x, S) .Call(C_dbinorm, x, S)
```
