## Supplementary material for "MoBPS - Modular Breeding Program Simulator": RandomFieldsUtils R-package: zzz.R

```
.onLoad <- function(lib, pkg) {
  .Call("loadRandomFieldsUtils")
}

.onAttach <- function (lib, pkg) {
  packageStartupMessage(.Call("attachRandomFieldsUtils"))
}

.onDetach <- function(lib) {
# .Call("detachRanodmFieldsUtils")
}

.onUnload <- function(lib, pkg){
  .Call("detachRandomFieldsUtils")
}
```
