## Supplementary material for "MoBPS - Modular Breeding Program Simulator": RandomFieldsUtils R-package: bckslvmodified.f

```
c 
c     Authors:
c     Reinhard Furrer
c       
c  Copyright (C) 2017 -- 2017 Reinhard Furrer
c 
c This program is free software; you can redistribute it and/or
c modify it under the terms of the GNU General Public License
c as published by the Free Software Foundation; either version 3
c of the License, or (at your option) any later version.
c 
c This program is distributed in the hope that it will be useful,
c but WITHOUT ANY WARRANTY; without even the implied warranty of
c MERCHANTABILITY or FITNESS FOR A PARTICULAR PURPOSE.  See the
c GNU General Public License for more details.
c 
c You should have received a copy of the GNU General Public License
c along with this program; if not, write to the Free Software
c Foundation, Inc., 59 Temple Place - Suite 330, Boston, MA  02111-1307, USA.  


      subroutine backsolve(m,nsuper,nrhs,lindx,xlindx,lnz,
     &                   xlnz,xsuper,b)
c see below...
      implicit none

      integer m,nsuper,nrhs,lindx(*),xlindx(m+1),
     &        xlnz(m+1),xsuper(m+1)
      double precision lnz(*),b(m,nrhs)
      integer j
      do j = 1,nrhs
         call blkslb(nsuper,xsuper,xlindx,lindx,xlnz,lnz,b(1,j))
      enddo
      return
      end


      subroutine forwardsolve(m,nsuper,nrhs,lindx,xlindx,
     &  lnz,xlnz,xsuper,b)
c INPUT:
c     m -- the number of column in the matrix
c     lindx -- an nsub-vector of interger which contains, in 
c           column major oder, the row subscripts of the nonzero
c           entries in L in a compressed storage format
c     xlindx -- an nsuper-vector of integer of pointers for lindx
c     lnz -- First contains the non-zero entries of d; later
c            contains the entries of the Cholesky factor
c     xlnz -- column pointer for L stored in lnz
c     xsuper -- array of length m+1 containing the supernode
c               partitioning
c     b -- the rhs of the equality constraint
c OUTPUT:
c     b -- the solution
      
      implicit none

      integer m,nsuper,nrhs,lindx(*),xlindx(m+1),
     &        xlnz(m+1),xsuper(m+1)
      double precision lnz(*),b(m,nrhs)
      integer j
c
      do j = 1,nrhs
         call blkslf(nsuper,xsuper,xlindx,lindx,xlnz,lnz,b(1,j))
      enddo
      return
      end
C***********************************************************************
C***********************************************************************
C
C   Version:        0.4
C   Last modified:  December 27, 1994
C   Authors:        Esmond G. Ng and Barry W. Peyton
C                   Slight modification by Reinhard Furrer 
C
C   Mathematical Sciences Section, Oak Ridge National Laboratory
C
C***********************************************************************
C***********************************************************************
C
C***********************************************************************
      subroutine pivotforwardsolve(m,nsuper,nrhs,lindx,xlindx,lnz,
     &                   xlnz,invp,perm,xsuper,newrhs,sol,b)
c Sparse least squares solver via Ng-Peyton's sparse Cholesky 
c    factorization for sparse symmetric positive definite
c INPUT:
c     m -- the number of column in the design matrix X
c     nsubmax -- upper bound of the dimension of lindx
c     lindx -- an nsub-vector of interger which contains, in 
c           column major oder, the row subscripts of the nonzero
c           entries in L in a compressed storage format
c     xlindx -- an nsuper-vector of integer of pointers for lindx
c     lnz -- First contains the non-zero entries of d; later
c            contains the entries of the Cholesky factor
c     xlnz -- column pointer for L stored in lnz
c     invp -- an m-vector of integer of inverse permutation
c             vector
c     perm -- an m-vector of integer of permutation vector
c     xsuper -- array of length m+1 containing the supernode
c               partitioning
c     newrhs -- extra work vector for right-hand side and
c               solution
c     sol -- the least squares solution
c     b -- an m-vector, usualy the rhs of the equality constraint
c          X'a = (1-tau)X'e in the rq setting
c OUTPUT:
c     y -- an m-vector of least squares solution
c WORK ARRAYS:
c     b -- an m-vector, usually the rhs of the equality constraint
c          X'a = (1-tau)X'e in the rq setting
      implicit none

      integer m,nsuper,nrhs,lindx(*),xlindx(m+1),
     &        invp(m),perm(m),xlnz(m+1), xsuper(m+1)
      integer i,j
      double precision lnz(*),b(m,nrhs),newrhs(m),sol(m,nrhs)

      do j = 1,nrhs
         do i = 1,m
            newrhs(i) = b(perm(i),j)
         enddo
         call blkslf(nsuper,xsuper,xlindx,lindx,xlnz,lnz,newrhs)
         do i = 1,m
            sol(i,j) = newrhs(invp(i))
         enddo
      enddo
      return
      end
C***********************************************************************
      subroutine pivotbacksolve(m,nsuper,nrhs,lindx,xlindx,lnz,
     &                   xlnz,invp,perm,xsuper,newrhs,sol,b)
c     see above
      implicit none

      integer m, nsuper,nrhs,lindx(*),xlindx(m+1),
     &        invp(m),perm(m),xlnz(m+1), xsuper(m+1)
      double precision lnz(*),b(m,nrhs),newrhs(m),sol(m,nrhs)
      integer i,j
      do j = 1,nrhs
         do i = 1,m
            newrhs(i) = b(perm(i),j)
         enddo
         call blkslb(nsuper,xsuper,xlindx,lindx,xlnz,lnz,newrhs)
         do i = 1,m
            sol(i,j) = newrhs(invp(i))
         enddo
      enddo
      return
      end
C***********************************************************************
      subroutine backsolves(m,nsuper,nrhs,lindx,xlindx,lnz,
     &                   xlnz,invp,perm,xsuper,newrhs,b)
c MODIFIED, overwritting b now!! (M.Schlather, 4.4.15)
c
c Sparse least squares solver via Ng-Peyton's sparse Cholesky 
c    factorization for sparse symmetric positive definite
c INPUT:
c     m -- the number of column in the design matrix X
c     nsubmax -- upper bound of the dimension of lindx
c     lindx -- an nsub-vector of interger which contains, in 
c           column major oder, the row subscripts of the nonzero
c           entries in L in a compressed storage format
c     xlindx -- an nsuper-vector of integer of pointers for lindx
c     nnzlmax -- the upper bound of the non-zero entries in
c                L stored in lnz, including the diagonal entries
c     lnz -- First contains the non-zero entries of d; later
c            contains the entries of the Cholesky factor
c     xlnz -- column pointer for L stored in lnz
c     invp -- an m-vector of integer of inverse permutation
c             vector
c     perm -- an m-vector of integer of permutation vector
c     xsuper -- array of length m+1 containing the supernode
c               partitioning
c     newrhs -- extra work vector for right-hand side and
c               solution
c    ( sol -- the least squares solution)
c     b -- an m-vector, usualy the rhs of the equality constraint
c          X'a = (1-tau)X'e in the rq setting
c OUTPUT:
c     y -- an m-vector of least squares solution
c WORK ARRAYS:
c     b -- an m-vector, usually the rhs of the equality constraint
c          X'a = (1-tau)X'e in the rq setting
      implicit none

      integer m,nsuper,nrhs,lindx(*),xlindx(m+1),
     &        invp(m),perm(m),xlnz(m+1), xsuper(m+1)
      double precision lnz(*),b(m,nrhs),newrhs(m)

      integer i,j
      do j = 1,nrhs
         do i = 1,m
           newrhs(i) = b(perm(i),j) 
         enddo
         call blkslv(nsuper,xsuper,xlindx,lindx,xlnz,lnz,newrhs)
         do i = 1,m
            b(i,j) = newrhs(invp(i))
         enddo
      enddo
      return
      end
```
