## Supplementary material for "MoBPS - Modular Breeding Program Simulator": RandomFieldsUtils R-package: brdomain.cc

```
/* 
 Authors
 Martin Schlather,

 Collection of system specific auxiliary functions

 Copyright (C) 2001 -- 2017 Martin Schlather, 

This program is free software; you can redistribute it and/or
modify it under the terms of the GNU General Public License
as published by the Free Software Foundation; either version 3
of the License, or (at your option) any later version.

This program is distributed in the hope that it will be useful,
but WITHOUT ANY WARRANTY; without even the implied warranty of
MERCHANTABILITY or FITNESS FOR A PARTICULAR PURSE.  See the
GNU General Public License for more details.

You should have received a copy of the GNU General Public License
along with this program; if not, write to the Free Software
Foundation, Inc., 59 Temple Place - Suite 330, Boston, MA  02111-1307, USA.
*/


// #define SIMD_AVAIALBLE 1


#include <Rmath.h>
#include <R.h>
#include "RandomFieldsUtils.h"
#include "General_utils.h"


//
#ifdef SCHLATHERS_MACHINE
//#include "kleinkram.h"


SEXP brdomain(SEXP Srf, SEXP Ssigma2, SEXP Sinstances, SEXP Smaxn) {
  double
    *rf = REAL(Srf),
    *sigma2 = REAL(Ssigma2);
  int
    instances = INTEGER(Sinstances)[0],
    maxn = INTEGER(Smaxn)[0],
    nloc = nrows(Ssigma2),
    total = instances * nloc,
    N = ncols(Srf);
  SEXP Ans;
  PROTECT(Ans = allocVector(REALSXP, total));
  double *ans = REAL(Ans);
  
  for (int i=0; i<total; ans[i++] = RF_NEGINF);
  GetRNGstate();
  for (int t=0; t<instances; t++, ans += nloc) {
    PRINTF("\nt=%d ", t);
    double pois = 0;
    for (int m=0; m<maxn; m++) {
      double e = rexp(1.0);
      pois += e;
      int
	x0 = (int) (UNIFORM_RANDOM * nloc);
      if (t==1) { PRINTF(" pois=%10g e=%10g %d : ", pois, e, x0);}
      double
	*field = rf + nloc * (int) (UNIFORM_RANDOM * N),
	*s2 = sigma2 + x0 * nloc,
	uz0 = - LOG(pois) - field[x0];
      
      for (int j=0; j<nloc; j++) {
        if (j == x0 && field[j] != field[x0]) BUG;
	double w = uz0 + field[j] - s2[j];
	if (t==1 && j == 97-68 - 1 && (m <= 10)) {
	  PRINTF("j=%d %10g %10g u=%10g\n", j, w, ans[j], - LOG(pois));
	}
	if (t==1 && j == 97-68 - 1 &&  x0==j) {
	  PRINTF("\nGREAT j=%d %10g %10g u=%10g\n", j, w, ans[j], - LOG(pois));
	}
	if (w > ans[j]) ans[j] = w;
      }
    }
  }
  PutRNGstate();
  UNPROTECT(1);
  return Ans;
}


//
#endif // SCHLATHERS_MACHINE
```
