## Supplementary material for "MoBPS - Modular Breeding Program Simulator": RandomFieldsUtils R-package: cholmodified.f

```
c 
c     Authors:
c     Reinhard Furrer
c       
c  Copyright (C) 2017 -- 2017 Reinhard Furrer
c 
c This program is free software; you can redistribute it and/or
c modify it under the terms of the GNU General Public License
c as published by the Free Software Foundation; either version 3
c of the License, or (at your option) any later version.
c 
c This program is distributed in the hope that it will be useful,
c but WITHOUT ANY WARRANTY; without even the implied warranty of
c MERCHANTABILITY or FITNESS FOR A PARTICULAR PURPOSE.  See the
c GNU General Public License for more details.
c 
c You should have received a copy of the GNU General Public License
c along with this program; if not, write to the Free Software
c Foundation, Inc., 59 Temple Place - Suite 330, Boston, MA  02111-1307, USA.  


      subroutine updatefactor( m,nnzd,
     &     d,jd,id, invp,perm, 
     &                lindx,xlindx, nsuper,lnz,xlnz,
     &                snode, xsuper,
     &                cachesize,ierr)

      implicit none
      integer m,nnzd
      integer nsuper,tmpsiz,
     &        ierr,
     &        jd(nnzd),cachesize,
     &        id(m+1),lindx(*),xlindx(*),
     &        invp(m),perm(m),xlnz(m+1),
     &        snode(m),xsuper(m+1)
      double precision d(nnzd),lnz(*)

c temp and working stuff, loops, etc
      integer iwork(7*m+3)
      integer split(m)
c
c Clean L
      call cleanlnz(nsuper,xsuper,xlnz,lnz)
c
c Input numerical values into data structures of L
      call inpnv(id,jd,d,perm,invp,nsuper,xsuper,xlindx,lindx,
     &           xlnz,lnz,iwork)
c
c Initialization for block factorization
      call bfinit(m,nsuper,xsuper,snode,xlindx,lindx,cachesize,tmpsiz,
     &            split)
c
c Numerical factorization
      call blkfc2(nsuper,xsuper,snode,split,xlindx,lindx,xlnz,
     &     lnz,iwork(1),iwork(nsuper+1),iwork(2*nsuper+1),
     &     iwork(2*nsuper+m+1),tmpsiz,ierr)
      if (ierr .eq. -1) then
         ierr = 1
         go to 100
      elseif (ierr .eq. -2) then
         ierr = 3
         go to 100
      endif
  100 continue
      return
      end


      

      subroutine cholstepwise(m,nnzd,
     &     d,jd,id,    doperm,invp,perm,
     &                nsub,nsubmax,
     &                lindx,xlindx,nsuper,nnzlmax,lnz,xlnz,
     &                snode,xsuper,
     &                cachsz,ierr)
c     Modified chol routine  
c
c
c Sparse least squares solver via Ng-Peyton's sparse Cholesky 
c    factorization for sparse symmetric positive definite
c INPUT:
c     m -- the number of column in the matrix A
c     d -- an nnzd-vector of non-zero values of A
c     jd -- an nnzd-vector of indices in d
c     id -- an (m+1)-vector of pointers to the begining of each
c           row in d and jd
c     nsubmax -- upper bound of the dimension of lindx
c     lindx -- an nsub-vector of integer which contains, in 
c           column major oder, the row subscripts of the nonzero
c           entries in L in a compressed storage format
c     xlindx -- an nsuper-vector of integer of pointers for lindx
c     nsuper -- the length of xlindx   ???
c     nnzlmax -- the upper bound of the non-zero entries in
c                L stored in lnz, including the diagonal entries
c     lnz -- First contains the non-zero entries of d; later
c            contains the entries of the Cholesky factor
c     xlnz -- column pointer for L stored in lnz
c     invp -- an n-vector of integer of inverse permutation
c             vector
c     perm -- an n-vector of integer of permutation vector
c     colcnt -- array of length m, containing the number of
c               non-zeros in each column of the factor, including
c               the diagonal entries
c     snode -- array of length m for recording supernode
c              membership
c     xsuper -- array of length m+1 containing the supernode
c               partitioning
c     split -- an m-vector with splitting of supernodes so that
c              they fit into cache
c     tmpmax -- upper bound of the dimension of tmpvec
c     tmpvec -- a tmpmax-vector of temporary vector
c     cachsz -- size of the cache (in kilobytes) on the target
c               machine
c     ierr -- error flag
c       1 -- insufficient work space in call to extract
c       2 -- insufficient storage in iwork when calling ordmmd;
c       3 -- insufficient storage in iwork when calling sfinit;
c       4 -- nnzl > nnzlmax when calling sfinit
c       5 -- nsub > nsubmax when calling sfinit
c       6 -- insufficient work space in iwork when calling symfct
c       7 -- inconsistancy in input when calling symfct
c       8 -- tmpsiz > tmpmax when calling symfct; increase tmpmax
c       9 -- nonpositive diagonal encountered when calling
c            blkfct
c       10 -- insufficient work storage in tmpvec when calling
c            blkfct
c       11 -- insufficient work storage in iwork when calling
c            blkfct
c OUTPUT:
c     y -- an m-vector of least squares solution
c     nsub -- number of subscripts in lindx
c WORK ARRAYS:
c     adjncy -- the indices of non diag elements
c     iwsiz -- set at 7*m+3
c     iwork -- an iwsiz-vector of integer as work space
c
c
      implicit none
      integer m,nnzd,doperm
      integer nsub,nsuper,nnzl,iwsiz,tmpsiz,
     &        nnzlmax,nsubmax,cachsz,ierr,
     &        adj(m+1),adjncy(nnzd-m+1),jd(nnzd),
c  fix introduced in 29-3
c     &        adj(m+1),adjncy(nnzd-m),jd(nnzd),
     &        id(m+1),lindx(nsubmax),xlindx(m+1),
     &        invp(m),perm(m),xlnz(m+1),
     &        colcnt(m),snode(m),xsuper(m+1),split(m)
      double precision d(nnzd),lnz(nnzlmax)

c temp and working stuff, loops, etc
      integer i,j,k,  nnzadj, jtmp
      integer iwork(7*m+3)


c iwsiz is used temporalily
      iwsiz=0

c Create the adjacency matrix: eliminate the diagonal elements from
c    (d,id,jd) and make two copies: (*,xlindx,lindx),(*,adj,adjncy)
c    Also to lindx and xlindx, because the matrix structure is destroyed 
c    by the minimum degree ordering routine.

      nsub = 0

c the adj matrix has m elements less than d
      nnzadj = nnzd - m

      k=1
      do i=1,m
c copy id, but ajust for the missing diagonal.
         xlindx(i) = id(i)-i+1
         adj(i) = xlindx(i)

c now cycle over all rows
         do j=id(i),id(i+1)-1
            jtmp=jd(j)
            if (jtmp.ne.i) then
               lindx(k) = jtmp               
               adjncy(k) = jtmp               
               k=k+1
            else
               if ( d(j) .le. 0) then
                  ierr = 1
                  return
               endif
               iwsiz = iwsiz + 1
            endif
         enddo
      enddo
      jtmp=m+1
      xlindx(jtmp) = id(jtmp)-m
      adj(jtmp) = xlindx(jtmp)

c check if we actually had m elements on the diagonal...
      if ( iwsiz .lt. m) then
         ierr = 1
         return
      endif

c initialize iwsiz to the later used value...        
      iwsiz=7*m+3

c
c
c reorder the matrix using minimum degree ordering routine.
c we call the genmmd function directly (do not pass via ordmmd).


      if (doperm.eq.1) then

c       delta  - tolerance value for multiple elimination.
c                set to 0 below
c       maxint - maximum machine representable (short) integer
c                (any smaller estimate will do) for marking
c                nodes.
c                set to 32767 below
         call genmmd  (  m, xlindx,lindx, invp,perm,0,
     1        iwork(1),  iwork(m+1), iwork(2*m+1), iwork(3*m+1) ,
     1        32767, nsub   )
      endif
      if (doperm.eq.2) then
         call genrcm ( m, nnzadj, xlindx,lindx, perm )
         do i=1,m
            invp(perm(i))=i
         enddo
      endif
      if (doperm.eq.0) then
         do i=1,m
            invp(perm(i))=i
         enddo
      endif
         
c
c Call sfinit: Symbolic factorization initialization
c   to compute supernode partition and storage requirements
c   for symbolic factorization. New ordering is a postordering 
c   of the nodal elimination tree
c
      call sfinit(m,nnzadj,adj(1),adjncy(1),perm,
     &            invp,colcnt,nnzl,nsub,nsuper,snode,xsuper,iwsiz,
     &            iwork,ierr)
c we do not have to test ierr, as we have hardwired iwsiz to 7*m+3
      if (nnzl .gt. nnzlmax) then
         ierr = 4
         go to 100
      endif
      if (nsub .gt. nsubmax) then
         ierr = 5
         go to 100
      endif
c
c Call symfct: Perform supernodal symbolic factorization
c
      iwsiz = nsuper + 2 * m + 1

      call symfc2(m,nnzadj,adj(1),adjncy(1),perm,invp,
     &            colcnt,nsuper,xsuper,snode,nsub,xlindx,lindx,
     &            xlnz,
     &     iwork(1), iwork(nsuper+1), iwork(nsuper+m+2) ,ierr)
c ierr = -2 "inconsistency in the input"
      if (ierr .eq. -2) then
         ierr = 6
         go to 100
      endif


c
c Input numerical values into data structures of L
      call inpnv(id,jd,d,perm,invp,nsuper,xsuper,xlindx,lindx,
     &           xlnz,lnz,iwork)
c
c Initialization for block factorization
      call bfinit(m,nsuper,xsuper,snode,xlindx,lindx,cachsz,tmpsiz,
     &            split)
c
c Numerical factorization
      call blkfc2(nsuper,xsuper,snode,split,xlindx,lindx,xlnz,
     &     lnz,iwork(1),iwork(nsuper+1),iwork(2*nsuper+1),
     &     iwork(2*nsuper+m+1),tmpsiz,ierr)
      if (ierr .eq. -1) then
         ierr = 1
         go to 100
      elseif (ierr .eq. -2) then
         ierr = 3
         go to 100
      endif
  100 continue
c      WRITE(6,699) nnzd
c699    FORMAT(1X,' FOUND ',I6,' RETURNING!')
      return
      end


C***********************************************************************
C***********************************************************************
C
C   Authors:        Reinhard Furrer, based on inpnv
C
C
C***********************************************************************
C***********************************************************************
C
C     ------------------------------------------------------
C     Clean the array lnz
C     ------------------------------------------------------
C
      SUBROUTINE  CLEANLNZ (NSUPER, XSUPER, XLNZ, LNZ)
C
      IMPLICIT NONE

      INTEGER             NSUPER
      INTEGER             XSUPER(*), XLNZ(*)
      DOUBLE PRECISION    LNZ(*)
C     
      INTEGER             II, J, JSUPER
C
      DO  500  JSUPER = 1, NSUPER
         DO  400  J = XSUPER(JSUPER), XSUPER(JSUPER+1)-1
            DO  200  II = XLNZ(J), XLNZ(J+1)-1
               LNZ(II) = 0.0
 200        CONTINUE
 400     CONTINUE
C     
 500  CONTINUE
      RETURN
      END


C***********************************************************************
C***********************************************************************
C
C   Version:        0.4
C   Last modified:  December 27, 1994
C   Authors:        Esmond G. Ng and Barry W. Peyton
C
C   Mathematical Sciences Section, Oak Ridge National Laboratory
C
C***********************************************************************
C***********************************************************************
C************     ASSMB .... INDEXED ASSEMBLY OPERATION     ************
C***********************************************************************
C***********************************************************************
C
C   PURPOSE:
C       THIS ROUTINE PERFORMS AN INDEXED ASSEMBLY (I.E., SCATTER-ADD)
C       OPERATION, ASSUMING DATA STRUCTURES USED IN SOME OF OUR SPARSE
C       CHOLESKY CODES.
C
C   INPUT PARAMETERS:
C       M               -   NUMBER OF ROWS IN Y.
C       Q               -   NUMBER OF COLUMNS IN Y.
C       Y               -   BLOCK UPDATE TO BE INCORPORATED INTO FACTOR
C                           STORAGE.
C       RELIND          -   RELATIVE INDICES FOR MAPPING THE UPDATES
C                           ONTO THE TARGET COLUMNS.
C       XLNZ            -   POINTERS TO THE START OF EACH COLUMN IN THE
C                           TARGET MATRIX.
C
C   OUTPUT PARAMETERS:
C       LNZ             -   CONTAINS COLUMNS MODIFIED BY THE UPDATE
C                           MATRIX.
C
C***********************************************************************
C
      SUBROUTINE  ASSMB  (  M     , Q     , Y     , RELIND, XLNZ  ,
     &                      LNZ   , LDA                             )
C
C***********************************************************************
C
C       -----------
C       PARAMETERS.
C       -----------
C
        INTEGER             LDA   , M     , Q
        INTEGER             XLNZ(*)
        INTEGER             RELIND(*)
        DOUBLE PRECISION    LNZ(*)        , Y(*)
C
C       ----------------
C       LOCAL VARIABLES.
C       ----------------
C
        INTEGER             ICOL  , IL1   , IR    , IY1   , LBOT1 ,
     &                      YCOL  , YOFF1
C
C***********************************************************************
C
C
        YOFF1 = 0
        IY1 = 0
        DO  200  ICOL = 1, Q
            YCOL = LDA - RELIND(ICOL)
            LBOT1 = XLNZ(YCOL+1) - 1
CDIR$ IVDEP
            DO  100  IR = ICOL, M
                IL1 = LBOT1 - RELIND(IR)
                IY1 = YOFF1 + IR
                LNZ(IL1) = LNZ(IL1) + Y(IY1)
                Y(IY1) = 0.0D0
  100       CONTINUE
            YOFF1 = IY1 - ICOL
  200   CONTINUE
C
      RETURN
      END
C***********************************************************************
C***********************************************************************
C
C   Version:        0.4
C   Last modified:  December 27, 1994
C   Authors:        Joseph W.H. Liu
C
C   Mathematical Sciences Section, Oak Ridge National Laboratory
C
C***********************************************************************
C***********************************************************************
C******     BETREE ..... BINARY TREE REPRESENTATION OF ETREE     *******
C***********************************************************************
C***********************************************************************
C
C   WRITTEN BY JOSEPH LIU (JUL 17, 1985)
C
C   PURPOSE:
C       TO DETERMINE THE BINARY TREE REPRESENTATION OF THE ELIMINATION
C       TREE GIVEN BY THE PARENT VECTOR.  THE RETURNED REPRESENTATION
C       WILL BE GIVEN BY THE FIRST-SON AND BROTHER VECTORS.  THE ROOT
C       OF THE BINARY TREE IS ALWAYS NEQNS.
C
C   INPUT PARAMETERS:
C       NEQNS           -   NUMBER OF EQUATIONS.
C       PARENT          -   THE PARENT VECTOR OF THE ELIMINATION TREE.
C                           IT IS ASSUMED THAT PARENT(I) > I EXCEPT OF
C                           THE ROOTS.
C
C   OUTPUT PARAMETERS:
C       FSON            -   THE FIRST SON VECTOR.
C       BROTHR          -   THE BROTHER VECTOR.
C
C***********************************************************************
C
      SUBROUTINE  BETREE (  NEQNS , PARENT, FSON  , BROTHR          )
C
C***********************************************************************
C
        INTEGER(4)           BROTHR(*)     , FSON(*)       ,
     &                      PARENT(*)
C
        INTEGER(4)           NEQNS
C
C***********************************************************************
C
        INTEGER(4)           LROOT , NODE  , NDPAR
C
C***********************************************************************
C
        IF  ( NEQNS .LE. 0 )  RETURN
C
        DO  100  NODE = 1, NEQNS
            FSON(NODE) = 0
            BROTHR(NODE) = 0
  100   CONTINUE
        LROOT = NEQNS
C       ------------------------------------------------------------
C       FOR EACH NODE := NEQNS-1 STEP -1 DOWNTO 1, DO THE FOLLOWING.
C       ------------------------------------------------------------
        IF  ( NEQNS .LE. 1 )  RETURN
        DO  300  NODE = NEQNS-1, 1, -1
            NDPAR = PARENT(NODE)
            IF  ( NDPAR .LE. 0  .OR.  NDPAR .EQ. NODE )  THEN
C               -------------------------------------------------
C               NODE HAS NO PARENT.  GIVEN STRUCTURE IS A FOREST.
C               SET NODE TO BE ONE OF THE ROOTS OF THE TREES.
C               -------------------------------------------------
                BROTHR(LROOT) = NODE
                LROOT = NODE
            ELSE
C               -------------------------------------------
C               OTHERWISE, BECOMES FIRST SON OF ITS PARENT.
C               -------------------------------------------
                BROTHR(NODE) = FSON(NDPAR)
                FSON(NDPAR) = NODE
            ENDIF
  300   CONTINUE
        BROTHR(LROOT) = 0
C
        RETURN
      END
C***********************************************************************
C***********************************************************************
C
C   Version:        0.4
C   Last modified:  December 27, 1994
C   Authors:        Esmond G. Ng and Barry W. Peyton
C
C   Mathematical Sciences Section, Oak Ridge National Laboratory
C
C***********************************************************************
C***********************************************************************
C******     BFINIT ..... INITIALIZATION FOR BLOCK FACTORIZATION   ******
C***********************************************************************
C***********************************************************************
C
C   PURPOSE:
C       THIS SUBROUTINE COMPUTES ITEMS NEEDED BY THE LEFT-LOOKING
C       BLOCK-TO-BLOCK CHOLESKY FACTORITZATION ROUTINE BLKFCT.
C
C   INPUT PARAMETERS:
C       NEQNS           -   NUMBER OF EQUATIONS.
C       NSUPER          -   NUMBER OF SUPERNODES.
C       XSUPER          -   INTEGER ARRAY OF SIZE (NSUPER+1) CONTAINING
C                           THE SUPERNODE PARTITIONING.
C       SNODE           -   SUPERNODE MEMBERSHIP.
C       (XLINDX,LINDX)  -   ARRAYS DESCRIBING THE SUPERNODAL STRUCTURE.
C       CACHSZ          -   CACHE SIZE (IN KBYTES).
C
C   OUTPUT PARAMETERS:
C       TMPSIZ          -   SIZE OF WORKING STORAGE REQUIRED BY BLKFCT.
C       SPLIT           -   SPLITTING OF SUPERNODES SO THAT THEY FIT
C                           INTO CACHE.
C
C***********************************************************************
C
      SUBROUTINE  BFINIT  ( NEQNS , NSUPER, XSUPER, SNODE , XLINDX, 
     &                      LINDX , CACHSZ, TMPSIZ, SPLIT           )
C
C***********************************************************************
C
        INTEGER     CACHSZ, NEQNS , NSUPER, TMPSIZ
        INTEGER     XLINDX(*)       , XSUPER(*)
        INTEGER     LINDX (*)       , SNODE (*)   ,
     &              SPLIT(*)
C
C***********************************************************************
C
C       ---------------------------------------------------
C       DETERMINE FLOATING POINT WORKING SPACE REQUIREMENT.
C       ---------------------------------------------------
        CALL  FNTSIZ (  NSUPER, XSUPER, SNODE , XLINDX, LINDX ,
     &                  TMPSIZ                                  )
C
C       -------------------------------
C       PARTITION SUPERNODES FOR CACHE.
C       -------------------------------
        CALL  FNSPLT (  NEQNS , NSUPER, XSUPER, XLINDX, CACHSZ,
     &                  SPLIT                                   )
C
        RETURN
      END
C***********************************************************************
C***********************************************************************
C
C   Version:        0.3
C   Last modified:  March 6, 1995
C   Authors:        Esmond G. Ng and Barry W. Peyton
C                   RF eliminated dependence on SMXPY and MMPY
C
C   Mathematical Sciences Section, Oak Ridge National Laboratoy
C
C***********************************************************************
C***********************************************************************
C*********     BLKFC2 .....  BLOCK GENERAL SPARSE CHOLESKY     *********
C***********************************************************************
C***********************************************************************
C
C   PURPOSE:
C       THIS SUBROUTINE FACTORS A SPARSE POSITIVE DEFINITE MATRIX.
C       THE COMPUTATION IS ORGANIZED AROUND KERNELS THAT PERFORM
C       SUPERNODE-TO-SUPERNODE UPDATES, I.E., BLOCK-TO-BLOCK UPDATES.
C
C   INPUT PARAMETERS:
C       NSUPER          -   NUMBER OF SUPERNODES.
C       XSUPER          -   SUPERNODE PARTITION.
C       SNODE           -   MAPS EACH COLUMN TO THE SUPERNODE CONTAINING
C                           IT.
C       SPLIT           -   SPLITTING OF SUPERNODES SO THAT THEY FIT
C                           INTO CACHE.
C       (XLINDX,LINDX)  -   ROW INDICES FOR EACH SUPERNODE (INCLUDING
C                           THE DIAGONAL ELEMENTS).
C       (XLNZ,LNZ)      -   ON INPUT, CONTAINS MATRIX TO BE FACTORED.
C       TMPSIZ          -   SIZE OF TEMPORARY WORKING STORAGE.
C
C   OUTPUT PARAMETERS:
C       LNZ             -   ON OUTPUT, CONTAINS CHOLESKY FACTOR.
C       IFLAG           -   ERROR FLAG.
C                               0: SUCCESSFUL FACTORIZATION.
C                              -1: NONPOSITIVE DIAGONAL ENCOUNTERED,
C                                  MATRIX IS NOT POSITIVE DEFINITE.
C                              -2: INSUFFICIENT WORKING STORAGE 
C                                  [TEMP(*)].
C
C   WORKING PARAMETERS:
C       LINK            -   LINKS TOGETHER THE SUPERNODES IN A SUPERNODE
C                           ROW.
C       LENGTH          -   LENGTH OF THE ACTIVE PORTION OF EACH 
C                           SUPERNODE.
C       INDMAP          -   VECTOR OF SIZE NEQNS INTO WHICH THE GLOBAL
C                           INDICES ARE SCATTERED.
C       RELIND          -   MAPS LOCATIONS IN THE UPDATING COLUMNS TO 
C                           THE CORRESPONDING LOCATIONS IN THE UPDATED 
C                           COLUMNS.  (RELIND IS GATHERED FROM INDMAP).
C       TEMP            -   REAL VECTOR FOR ACCUMULATING UPDATES.  MUST
C                           ACCOMODATE ALL COLUMNS OF A SUPERNODE. 
C       
C***********************************************************************
C
      SUBROUTINE  BLKFC2 (  NSUPER, XSUPER, SNODE , SPLIT , XLINDX,
     &                      LINDX , XLNZ  , LNZ   , LINK  , LENGTH,
     &                      INDMAP, RELIND, TMPSIZ, IFLAG )
C
C*********************************************************************
C
C       -----------
C       PARAMETERS.
C       -----------
C
        INTEGER             XLINDX(*)     , XLNZ(*)
        INTEGER             INDMAP(*)     , LENGTH(*)     ,
     &                      LINDX(*)      , LINK(*)       ,
     &                      RELIND(*)     , SNODE(*)      ,
     &                      SPLIT(*)      , XSUPER(*)
        INTEGER             IFLAG , NSUPER, TMPSIZ
        DOUBLE PRECISION    LNZ(*)
C
C       ----------------
C       LOCAL VARIABLES.
C       ----------------
C
        INTEGER             FJCOL , FKCOL , I     , ILEN  , ILPNT ,
     &                      INDDIF, JLEN  , JLPNT , JSUP  , JXPNT ,
     &                      KFIRST, KLAST , KLEN  , KLPNT , KSUP  ,
     &                      KXPNT , LJCOL , NCOLUP, NJCOLS, NKCOLS,
     &                      NXKSUP, NXTCOL, NXTSUP, STORE
        DOUBLE PRECISION    TEMP(TMPSIZ) 
C     RF: put TEMP(*) into a local variable

        DOUBLE PRECISION MXDIAG
        INTEGER NTINY
C*********************************************************************
C
        IFLAG = 0
        NTINY = 0
C
C       -----------------------------------------------------------
C       INITIALIZE EMPTY ROW LISTS IN LINK(*) AND ZERO OUT TEMP(*).
C       -----------------------------------------------------------
        DO  100  JSUP = 1, NSUPER
            LINK(JSUP) = 0
  100   CONTINUE
        DO  200  I = 1, TMPSIZ
            TEMP(I) = 0.0D+00
  200   CONTINUE

C       COMPUTE MAXIMUM DIAGONAL ELEMENT IN INPUT MATRIX
        MXDIAG = 0.D0
        DO 201 I = 1, XSUPER(NSUPER+1)-1
          FJCOL = XLNZ(I)
          MXDIAG = MAX(MXDIAG, LNZ(FJCOL))
 201    CONTINUE
C
C       ---------------------------
C       FOR EACH SUPERNODE JSUP ...
C       ---------------------------
        DO  600  JSUP = 1, NSUPER
C
C           ------------------------------------------------
C           FJCOL  ...  FIRST COLUMN OF SUPERNODE JSUP.
C           LJCOL  ...  LAST COLUMN OF SUPERNODE JSUP.
C           NJCOLS ...  NUMBER OF COLUMNS IN SUPERNODE JSUP.
C           JLEN   ...  LENGTH OF COLUMN FJCOL.
C           JXPNT  ...  POINTER TO INDEX OF FIRST
C                       NONZERO IN COLUMN FJCOL.
C           ------------------------------------------------
            FJCOL  = XSUPER(JSUP)
            NJCOLS = XSUPER(JSUP+1) - FJCOL
            LJCOL  = FJCOL + NJCOLS - 1
            JLEN   = XLNZ(FJCOL+1) - XLNZ(FJCOL)
            JXPNT  = XLINDX(JSUP)

C            print *, 'Super Node: ', JSUP, ' first: ', FJCOL, 
C     .           ' last: ', LJCOL

            
C
C
C           -----------------------------------------------------
C           SET UP INDMAP(*) TO MAP THE ENTRIES IN UPDATE COLUMNS
C           TO THEIR CORRESPONDING POSITIONS IN UPDATED COLUMNS, 
C           RELATIVE THE THE BOTTOM OF EACH UPDATED COLUMN.
C           -----------------------------------------------------
            CALL  LDINDX ( JLEN, LINDX(JXPNT), INDMAP )
C
C           -----------------------------------------
C           FOR EVERY SUPERNODE KSUP IN ROW(JSUP) ...
C           -----------------------------------------
            KSUP = LINK(JSUP)
  300       IF  ( KSUP .GT. 0 )  THEN
                NXKSUP = LINK(KSUP)
C
C               -------------------------------------------------------
C               GET INFO ABOUT THE CMOD(JSUP,KSUP) UPDATE.
C
C               FKCOL  ...  FIRST COLUMN OF SUPERNODE KSUP.
C               NKCOLS ...  NUMBER OF COLUMNS IN SUPERNODE KSUP.
C               KLEN   ...  LENGTH OF ACTIVE PORTION OF COLUMN FKCOL.
C               KXPNT  ...  POINTER TO INDEX OF FIRST NONZERO IN ACTIVE
C                           PORTION OF COLUMN FJCOL.
C               -------------------------------------------------------
                FKCOL = XSUPER(KSUP)
                NKCOLS = XSUPER(KSUP+1) - FKCOL
                KLEN = LENGTH(KSUP)
                KXPNT = XLINDX(KSUP+1) - KLEN
C
C               -------------------------------------------
C               PERFORM CMOD(JSUP,KSUP), WITH SPECIAL CASES
C               HANDLED DIFFERENTLY.
C               -------------------------------------------
C
                IF  ( KLEN .NE. JLEN )  THEN
C
C                   -------------------------------------------
C                   SPARSE CMOD(JSUP,KSUP).
C
C                   NCOLUP ... NUMBER OF COLUMNS TO BE UPDATED.
C                   -------------------------------------------
C
                    DO  400  I = 0, KLEN-1
                        NXTCOL = LINDX(KXPNT+I)
                        IF  ( NXTCOL .GT. LJCOL )  GO TO 500
  400               CONTINUE
                    I = KLEN
  500               CONTINUE
                    NCOLUP = I
C
                    IF  ( NKCOLS .EQ. 1 )  THEN
C
C                       ----------------------------------------------
C                       UPDATING TARGET SUPERNODE BY TRIVIAL
C                       SUPERNODE (WITH ONE COLUMN).
C
C                       KLPNT  ...  POINTER TO FIRST NONZERO IN ACTIVE
C                                   PORTION OF COLUMN FKCOL.
C                       ----------------------------------------------
                        KLPNT = XLNZ(FKCOL+1) - KLEN
                        CALL  MMPYI ( KLEN, NCOLUP, LINDX(KXPNT),
     &                                LNZ(KLPNT), XLNZ, LNZ, INDMAP )
C
                    ELSE
C
C                       --------------------------------------------
C                       KFIRST ...  FIRST INDEX OF ACTIVE PORTION OF
C                                   SUPERNODE KSUP (FIRST COLUMN TO
C                                   BE UPDATED).
C                       KLAST  ...  LAST INDEX OF ACTIVE PORTION OF
C                                   SUPERNODE KSUP.
C                       --------------------------------------------
C
                        KFIRST = LINDX(KXPNT)
                        KLAST  = LINDX(KXPNT+KLEN-1)
                        INDDIF = INDMAP(KFIRST) - INDMAP(KLAST)
C
                        IF  ( INDDIF .LT. KLEN )  THEN
C
C                           ---------------------------------------
C                           DENSE CMOD(JSUP,KSUP).
C
C                           ILPNT  ...  POINTER TO FIRST NONZERO IN
C                                       COLUMN KFIRST.
C                           ILEN   ...  LENGTH OF COLUMN KFIRST.
C                           ---------------------------------------
                            ILPNT = XLNZ(KFIRST)
                            ILEN = XLNZ(KFIRST+1) - ILPNT
                            CALL  MMPY ( KLEN, NKCOLS, NCOLUP,
     &                                   SPLIT(FKCOL), XLNZ(FKCOL),
     &                                   LNZ, LNZ(ILPNT), ILEN  )
C
                        ELSE
C
C                           -------------------------------
C                           GENERAL SPARSE CMOD(JSUP,KSUP).
C                           COMPUTE CMOD(JSUP,KSUP) UPDATE
C                           IN WORK STORAGE.
C                           -------------------------------
                            STORE = KLEN * NCOLUP - NCOLUP * 
     &                              (NCOLUP-1) / 2
                            IF  ( STORE .GT. TMPSIZ )  THEN
                                IFLAG = -2
                                RETURN
                            ENDIF
                            CALL  MMPY ( KLEN, NKCOLS, NCOLUP,
     &                                   SPLIT(FKCOL), XLNZ(FKCOL),
     &                                   LNZ, TEMP, KLEN  )
C                           ----------------------------------------
C                           GATHER INDICES OF KSUP RELATIVE TO JSUP.
C                           ----------------------------------------
                            CALL  IGATHR ( KLEN, LINDX(KXPNT),
     &                                     INDMAP, RELIND )
C                           --------------------------------------
C                           INCORPORATE THE CMOD(JSUP,KSUP) BLOCK
C                           UPDATE INTO THE TO APPROPRIATE COLUMNS
C                           OF L.
C                           --------------------------------------
                            CALL  ASSMB ( KLEN, NCOLUP, TEMP, RELIND,
     &                                    XLNZ(FJCOL), LNZ, JLEN )
C
                        ENDIF
C
                    ENDIF
C
                ELSE
C
C                   ----------------------------------------------
C                   DENSE CMOD(JSUP,KSUP).
C                   JSUP AND KSUP HAVE IDENTICAL STRUCTURE.
C
C                   JLPNT  ...  POINTER TO FIRST NONZERO IN COLUMN
C                               FJCOL.
C                   ----------------------------------------------
                    JLPNT = XLNZ(FJCOL)
                    CALL  MMPY ( KLEN, NKCOLS, NJCOLS, SPLIT(FKCOL),
     &                           XLNZ(FKCOL), LNZ, LNZ(JLPNT), JLEN)
                    NCOLUP = NJCOLS
                    IF  ( KLEN .GT. NJCOLS )  THEN
                        NXTCOL = LINDX(JXPNT+NJCOLS)
                    ENDIF
C
                ENDIF
C
C               ------------------------------------------------
C               LINK KSUP INTO LINKED LIST OF THE NEXT SUPERNODE
C               IT WILL UPDATE AND DECREMENT KSUP'S ACTIVE
C               LENGTH.
C               ------------------------------------------------
                IF  ( KLEN .GT. NCOLUP )  THEN
                    NXTSUP = SNODE(NXTCOL)
                    LINK(KSUP) = LINK(NXTSUP)
                    LINK(NXTSUP) = KSUP
                    LENGTH(KSUP) = KLEN - NCOLUP
                ELSE
                    LENGTH(KSUP) = 0
                ENDIF
C
C               -------------------------------
C               NEXT UPDATING SUPERNODE (KSUP).
C               -------------------------------
                KSUP = NXKSUP
                GO TO 300
C
            ENDIF
C
C           ----------------------------------------------
C           APPLY PARTIAL CHOLESKY TO THE COLUMNS OF JSUP.
C           ----------------------------------------------
CxPCxPCxPCxPCxPCxPCxPCxPCxPCxPCxPCxPCxPCxPCxPCxPCxPCxPCxPCxPCxPCxPCxPC
            CALL CHLSUP ( JLEN, NJCOLS, SPLIT(FJCOL), XLNZ(FJCOL), LNZ,
     &                    MXDIAG, NTINY )
            IF  ( IFLAG .NE. 0 )  THEN
                IFLAG = -1
                RETURN
            ENDIF
C
C           -----------------------------------------------
C           INSERT JSUP INTO LINKED LIST OF FIRST SUPERNODE
C           IT WILL UPDATE.
C           -----------------------------------------------
            IF  ( JLEN .GT. NJCOLS )  THEN
                NXTCOL = LINDX(JXPNT+NJCOLS)
                NXTSUP = SNODE(NXTCOL)
                LINK(JSUP) = LINK(NXTSUP)
                LINK(NXTSUP) = JSUP
                LENGTH(JSUP) = JLEN - NJCOLS
            ELSE
                LENGTH(JSUP) = 0
            ENDIF
C
  600   CONTINUE
C
CxPCxPCxPCxPCxPCxPCxPCxPCxPCxPCxPCxPCxPCxPCxPCxPCxPCxPCxPCxPCxPCxPCxPC
C        IF(NTINY .NE. 0) WRITE(6,699) NTINY
C 699    FORMAT(1X,' FOUND ',I6,' TINY DIAGONALS; REPLACED WITH INF')
C
C SET IFLAG TO -1 TO INDICATE PRESENCE OF TINY DIAGONALS
C
        IF(NTINY .NE. 0) IFLAG = -1
CxPCxPCxPCxPCxPCxPCxPCxPCxPCxPCxPCxPCxPCxPCxPCxPCxPCxPCxPCxPCxPCxPCxPC
        RETURN
      END
C***********************************************************************
C***********************************************************************
C
C   Written:        October 6, 1996 by SJW. Based on routine BLKSLV of
C                   Esmond G. Ng and Barry W. Peyton.
C
C   Modified:       Sept 30, 1999 to improve efficiency in the case
C                   in which the right-hand side and solution are both
C                   expected to be sparse. Happens a lot in "dense"
C                   column handling.
C
C***********************************************************************
C***********************************************************************
C*********     BLKSLB ... BACK TRIANGULAR SUBSTITUTION        **********
C***********************************************************************
C***********************************************************************
C
C   PURPOSE:
C       GIVEN THE CHOLESKY FACTORIZATION OF A SPARSE SYMMETRIC
C       POSITIVE DEFINITE MATRIX, THIS SUBROUTINE PERFORMS THE
C       BACKWARD TRIANGULAR SUBSTITUTION.  IT USES OUTPUT FROM BLKFCT.
C
C   INPUT PARAMETERS:
C       NSUPER          -   NUMBER OF SUPERNODES.
C       XSUPER          -   SUPERNODE PARTITION.
C       (XLINDX,LINDX)  -   ROW INDICES FOR EACH SUPERNODE.
C       (XLNZ,LNZ)      -   CHOLESKY FACTOR.
C
C   UPDATED PARAMETERS:
C       RHS             -   ON INPUT, CONTAINS THE RIGHT HAND SIDE.  ON
C                           OUTPUT, CONTAINS THE SOLUTION.
C
C***********************************************************************
C
      SUBROUTINE  BLKSLB (  NSUPER, XSUPER, XLINDX, LINDX , XLNZ  ,
     &                      LNZ   , RHS                             )
C
C***********************************************************************
C
        INTEGER             NSUPER
        INTEGER             LINDX(*)      , XSUPER(*)
        INTEGER             XLINDX(*)     , XLNZ(*)
        DOUBLE PRECISION    LNZ(*)        , RHS(*)
C
C***********************************************************************
C
        INTEGER             FJCOL , I     , IPNT  , IX    , IXSTOP,
     &                      IXSTRT, JCOL  , JPNT  , JSUP  , LJCOL
        DOUBLE PRECISION    T
C
C***********************************************************************
C
        IF  ( NSUPER .LE. 0 )  RETURN
C       -------------------------
C       BACKWARD SUBSTITUTION ...
C       -------------------------
        LJCOL = XSUPER(NSUPER+1) - 1
        DO  600  JSUP = NSUPER, 1, -1
            FJCOL  = XSUPER(JSUP)
            IXSTOP = XLNZ(LJCOL+1) - 1
            JPNT   = XLINDX(JSUP) + (LJCOL - FJCOL)
            DO  500  JCOL = LJCOL, FJCOL, -1
                IXSTRT = XLNZ(JCOL)
                IPNT   = JPNT + 1
                T      = RHS(JCOL)
CDIR$           IVDEP
                DO  400  IX = IXSTRT+1, IXSTOP
                   I = LINDX(IPNT)
                   IF(RHS(I) .NE. 0.D0) T = T - LNZ(IX)*RHS(I)
                   IPNT = IPNT + 1
 400            CONTINUE

                IF(T .NE. 0.D0) THEN
                   RHS(JCOL) = T/LNZ(IXSTRT)
                ELSE
                   RHS(JCOL) = 0.D0
                ENDIF

                IXSTOP    = IXSTRT - 1
                JPNT      = JPNT - 1
 500            CONTINUE

            LJCOL = FJCOL - 1
  600   CONTINUE
C
        RETURN
      END
C***********************************************************************
C***********************************************************************
C
C   Written:        October 6, 1996 by SJW. Based on routine BLKSLV of
C                   Esmond G. Ng and Barry W. Peyton.
C
C   Modified:       Sept 30, 1999 to improve efficiency in the case
C                   in which the right-hand side and solution are both
C                   expected to be sparse. Happens a lot in "dense"
C                   column handling.
C
C***********************************************************************
C***********************************************************************
C*********     BLKSLF ... FORWARD TRIANGULAR SUBSTITUTION     **********
C***********************************************************************
C***********************************************************************
C
C   PURPOSE:
C       GIVEN THE CHOLESKY FACTORIZATION OF A SPARSE SYMMETRIC
C       POSITIVE DEFINITE MATRIX, THIS SUBROUTINE PERFORMS THE
C       FORWARD TRIANGULAR SUBSTITUTIOn.  IT USES OUTPUT FROM BLKFCT.
C
C   INPUT PARAMETERS:
C       NSUPER          -   NUMBER OF SUPERNODES.
C       XSUPER          -   SUPERNODE PARTITION.
C       (XLINDX,LINDX)  -   ROW INDICES FOR EACH SUPERNODE.
C       (XLNZ,LNZ)      -   CHOLESKY FACTOR.
C
C   UPDATED PARAMETERS:
C       RHS             -   ON INPUT, CONTAINS THE RIGHT HAND SIDE.  ON
C                           OUTPUT, CONTAINS THE SOLUTION.
C
C***********************************************************************
C
      SUBROUTINE  BLKSLF (  NSUPER, XSUPER, XLINDX, LINDX , XLNZ  ,
     &                      LNZ   , RHS                             )
C
C***********************************************************************
C
        INTEGER             NSUPER
        INTEGER             LINDX(*)      , XSUPER(*)
        INTEGER             XLINDX(*)     , XLNZ(*)
        DOUBLE PRECISION    LNZ(*)        , RHS(*)
C
C***********************************************************************
C
        INTEGER             FJCOL , I     , IPNT  , IX    , IXSTOP,
     &                      IXSTRT, JCOL  , JPNT  , JSUP  , LJCOL
        DOUBLE PRECISION    T
C
C***********************************************************************
C
        IF  ( NSUPER .LE. 0 )  RETURN
C
C       ------------------------
C       FORWARD SUBSTITUTION ...
C       ------------------------
        FJCOL = XSUPER(1)
        DO  300  JSUP = 1, NSUPER
            LJCOL  = XSUPER(JSUP+1) - 1
            IXSTRT = XLNZ(FJCOL)
            JPNT   = XLINDX(JSUP)
            DO  200  JCOL = FJCOL, LJCOL
                IXSTOP    = XLNZ(JCOL+1) - 1

                IF(RHS(JCOL) .NE. 0.D0) THEN
                   T         = RHS(JCOL)/LNZ(IXSTRT)
                   RHS(JCOL) = T
                   IPNT      = JPNT + 1
CDIR$           IVDEP
                   DO  100  IX = IXSTRT+1, IXSTOP
                      I      = LINDX(IPNT)
                      RHS(I) = RHS(I) - T*LNZ(IX)
                      IPNT   = IPNT + 1
 100               CONTINUE
                ENDIF

                IXSTRT = IXSTOP + 1
                JPNT   = JPNT + 1
  200       CONTINUE
            FJCOL = LJCOL + 1
  300   CONTINUE
C
        RETURN
      END
C***********************************************************************
C***********************************************************************
C
C   Version:        0.4
C   Last modified:  December 27, 1994
C   Authors:        Esmond G. Ng and Barry W. Peyton
C
C   Mathematical Sciences Section, Oak Ridge National Laboratory
C
C   Modified:       Sept 30, 1999 to improve efficiency in the case
C                   in which the right-hand side and solution are both
C                   expected to be sparse. Happens a lot in "dense"
C                   column handling.
C
C***********************************************************************
C***********************************************************************
C*********     BLKSLV ... BLOCK TRIANGULAR SOLUTIONS          **********
C***********************************************************************
C***********************************************************************
C
C   PURPOSE:
C       GIVEN THE CHOLESKY FACTORIZATION OF A SPARSE SYMMETRIC
C       POSITIVE DEFINITE MATRIX, THIS SUBROUTINE PERFORMS THE
C       TRIANGULAR SOLUTION.  IT USES OUTPUT FROM BLKFCT.
C
C   INPUT PARAMETERS:
C       NSUPER          -   NUMBER OF SUPERNODES.
C       XSUPER          -   SUPERNODE PARTITION.
C       (XLINDX,LINDX)  -   ROW INDICES FOR EACH SUPERNODE.
C       (XLNZ,LNZ)      -   CHOLESKY FACTOR.
C
C   UPDATED PARAMETERS:
C       RHS             -   ON INPUT, CONTAINS THE RIGHT HAND SIDE.  ON
C                           OUTPUT, CONTAINS THE SOLUTION.
C
C***********************************************************************
C
      SUBROUTINE  BLKSLV (  NSUPER, XSUPER, XLINDX, LINDX , XLNZ  ,
     &                      LNZ   , RHS                             )
C
C***********************************************************************
C
        INTEGER             NSUPER
        INTEGER             LINDX(*)      , XSUPER(*)
        INTEGER             XLINDX(*)     , XLNZ(*)
        DOUBLE PRECISION    LNZ(*)        , RHS(*)
C
C***********************************************************************
C
        INTEGER             FJCOL , I     , IPNT  , IX    , IXSTOP,
     &                      IXSTRT, JCOL  , JPNT  , JSUP  , LJCOL
        DOUBLE PRECISION    T
C
C***********************************************************************
C
        IF  ( NSUPER .LE. 0 )  RETURN
C
C       ------------------------
C       FORWARD SUBSTITUTION ...
C       ------------------------
        FJCOL = XSUPER(1)
        DO  300  JSUP = 1, NSUPER
            LJCOL  = XSUPER(JSUP+1) - 1
            IXSTRT = XLNZ(FJCOL)
            JPNT   = XLINDX(JSUP)
C            print *, "JSUP", JSUP, FJCOL, LJCOL
            DO  200  JCOL = FJCOL, LJCOL
                IXSTOP    = XLNZ(JCOL+1) - 1

C                print *, JSUP, JCOL
C                print *, RHS(JCOL)

                IF(RHS(JCOL) .NE. 0.D0) THEN
                   T         = RHS(JCOL)/LNZ(IXSTRT)
                   RHS(JCOL) = T
                   IPNT      = JPNT + 1
CDIR$           IVDEP
                   DO  100  IX = IXSTRT+1, IXSTOP
                      I      = LINDX(IPNT)
                      RHS(I) = RHS(I) - T*LNZ(IX)
                      IPNT   = IPNT + 1
 100               CONTINUE
                ENDIF
                
                IXSTRT = IXSTOP + 1
                JPNT   = JPNT + 1
  200       CONTINUE
            FJCOL = LJCOL + 1
  300   CONTINUE
C
C       -------------------------
C       BACKWARD SUBSTITUTION ...
C       -------------------------
        LJCOL = XSUPER(NSUPER+1) - 1
        DO  600  JSUP = NSUPER, 1, -1
            FJCOL  = XSUPER(JSUP)
            IXSTOP = XLNZ(LJCOL+1) - 1
            JPNT   = XLINDX(JSUP) + (LJCOL - FJCOL)
            DO  500  JCOL = LJCOL, FJCOL, -1
                IXSTRT = XLNZ(JCOL)
                IPNT   = JPNT + 1
                T      = RHS(JCOL)
CDIR$           IVDEP
                DO  400  IX = IXSTRT+1, IXSTOP
                    I    = LINDX(IPNT)
                    IF(RHS(I) .NE. 0.D0) T = T - LNZ(IX)*RHS(I)
                    IPNT = IPNT + 1
  400           CONTINUE

                IF(T .NE. 0.D0) THEN
                   RHS(JCOL) = T/LNZ(IXSTRT)
                ELSE
                   RHS(JCOL) = 0.D0
                ENDIF

                IXSTOP    = IXSTRT - 1
                JPNT      = JPNT - 1
  500       CONTINUE
            LJCOL = FJCOL - 1
  600   CONTINUE
C
        RETURN
      END


C***********************************************************************
C***********************************************************************
C
C   Version:        0.4
C   Last modified:  January 12, 1995
C   Authors:        Esmond G. Ng and Barry W. Peyton
C
C   Mathematical Sciences Section, Oak Ridge National Laboratory
C
C***********************************************************************
C***********************************************************************
C******     BTREE2 ..... BINARY TREE REPRESENTATION OF ETREE     *******
C***********************************************************************
C***********************************************************************
C
C   PURPOSE:
C       TO DETERMINE A BINARY TREE REPRESENTATION OF THE ELIMINATION 
C       TREE, FOR WHICH EVERY "LAST CHILD" HAS THE MAXIMUM POSSIBLE
C       COLUMN NONZERO COUNT IN THE FACTOR.  THE RETURNED REPRESENTATION 
C       WILL BE GIVEN BY THE FIRST-SON AND BROTHER VECTORS.  THE ROOT OF 
C       THE BINARY TREE IS ALWAYS NEQNS.
C 
C   INPUT PARAMETERS:
C       NEQNS           -   NUMBER OF EQUATIONS.
C       PARENT          -   THE PARENT VECTOR OF THE ELIMINATION TREE.
C                           IT IS ASSUMED THAT PARENT(I) > I EXCEPT OF
C                           THE ROOTS.
C       COLCNT          -   COLUMN NONZERO COUNTS OF THE FACTOR.
C
C   OUTPUT PARAMETERS:
C       FSON            -   THE FIRST SON VECTOR.
C       BROTHR          -   THE BROTHER VECTOR.
C
C   WORKING PARAMETERS:
C       LSON            -   LAST SON VECTOR.
C
C***********************************************************************
C
      SUBROUTINE  BTREE2 (  NEQNS , PARENT, COLCNT, FSON  , BROTHR,
     &                      LSON    )
C
C***********************************************************************
C
        INTEGER             BROTHR(*)     , COLCNT(*)     ,
     &                      FSON(*)       , LSON(*)       ,
     &                      PARENT(*)
C
        INTEGER             NEQNS
C
C***********************************************************************
C
        INTEGER(4)           LROOT , NODE  , NDLSON, NDPAR
C
C***********************************************************************
C
        IF  ( NEQNS .LE. 0 )  RETURN
C
        DO  100  NODE = 1, NEQNS
            FSON(NODE) = 0
            BROTHR(NODE) = 0
            LSON(NODE) = 0
  100   CONTINUE
        LROOT = NEQNS
C       ------------------------------------------------------------
C       FOR EACH NODE := NEQNS-1 STEP -1 DOWNTO 1, DO THE FOLLOWING.
C       ------------------------------------------------------------
        IF  ( NEQNS .LE. 1 )  RETURN
        DO  300  NODE = NEQNS-1, 1, -1
            NDPAR = PARENT(NODE)
            IF  ( NDPAR .LE. 0  .OR.  NDPAR .EQ. NODE )  THEN
C               -------------------------------------------------
C               NODE HAS NO PARENT.  GIVEN STRUCTURE IS A FOREST.
C               SET NODE TO BE ONE OF THE ROOTS OF THE TREES.
C               -------------------------------------------------
                BROTHR(LROOT) = NODE
                LROOT = NODE
            ELSE
C               -------------------------------------------
C               OTHERWISE, BECOMES FIRST SON OF ITS PARENT.
C               -------------------------------------------
                NDLSON = LSON(NDPAR)
                IF  ( NDLSON .NE. 0 )  THEN
                    IF  ( COLCNT(NODE) .GE. COLCNT(NDLSON) )  THEN
                        BROTHR(NODE) = FSON(NDPAR)
                        FSON(NDPAR) = NODE
                    ELSE
                        BROTHR(NDLSON) = NODE
                        LSON(NDPAR) = NODE
                    ENDIF
                ELSE
                    FSON(NDPAR) = NODE
                    LSON(NDPAR) = NODE
                ENDIF
            ENDIF
  300   CONTINUE
        BROTHR(LROOT) = 0
C
        RETURN
      END
C***********************************************************************
C***********************************************************************
C
C   Version:        0.3
C   Last modified:  December 27, 1994
C   Authors:        Esmond G. Ng and Barry W. Peyton
C                   
C   Modified by RF:   Eliminated the  MMPYN, SMXPY as arguments
C
C   Mathematical Sciences Section, Oak Ridge National Laboratoy
C
C***********************************************************************
C***********************************************************************
C******     CHLSUP .... DENSE CHOLESKY WITHIN SUPERNODE   **************
C***********************************************************************
C***********************************************************************
C
C     PURPOSE - THIS ROUTINE PERFORMS CHOLESKY
C               FACTORIZATION ON THE COLUMNS OF A SUPERNODE
C               THAT HAVE RECEIVED ALL UPDATES FROM COLUMNS
C               EXTERNAL TO THE SUPERNODE.
C
C     INPUT PARAMETERS -
C        M      - NUMBER OF ROWS (LENGTH OF THE FIRST COLUMN).
C        N      - NUMBER OF COLUMNS IN THE SUPERNODE.
C        XPNT   - XPNT(J+1) POINTS ONE LOCATION BEYOND THE END
C                 OF THE J-TH COLUMN OF THE SUPERNODE.
C        X(*)   - CONTAINS THE COLUMNS OF OF THE SUPERNODE TO
C                 BE FACTORED.
C
C     EXTERNAL ROUTINES -
C        MMPY8  -  MATRIX-MATRIX MULTIPLY WITH 8 LOOP UNROLLING.
C
C     OUTPUT PARAMETERS -
C        X(*)   - ON OUTPUT, CONTAINS THE FACTORED COLUMNS OF
C                 THE SUPERNODE.
C        IFLAG  - UNCHANGED IF THERE IS NO ERROR.
C                 =1 IF NONPOSITIVE DIAGONAL ENTRY IS ENCOUNTERED.
C
C***********************************************************************
C
      SUBROUTINE  CHLSUP  ( M, N, SPLIT, XPNT, X, MXDIAG, NTINY
     &                      )
C
C***********************************************************************
C
C     -----------
C     PARAMETERS.
C     -----------
C
      EXTERNAL            MMPY8
C
      INTEGER             M, N
C
      INTEGER             XPNT(*), SPLIT(*)
C
      DOUBLE PRECISION    X(*), MXDIAG
      INTEGER             NTINY
C
C     ----------------
C     LOCAL VARIABLES.
C     ----------------
C
      INTEGER             FSTCOL, JBLK  , JPNT  , MM    , NN    ,
     &                    NXTCOL, Q
C
C***********************************************************************
C
        
        JBLK = 0
        FSTCOL = 1
        MM = M
        JPNT = XPNT(FSTCOL)
C
C       ----------------------------------------
C       FOR EACH BLOCK JBLK IN THE SUPERNODE ...
C       ----------------------------------------
  100   CONTINUE
        IF  ( FSTCOL .LE. N )  THEN
            JBLK = JBLK + 1
            NN = SPLIT(JBLK)
C           ------------------------------------------
C           ... PERFORM PARTIAL CHOLESKY FACTORIZATION
C               ON THE BLOCK.
C           ------------------------------------------
            CALL PCHOL ( MM, NN, XPNT(FSTCOL), X, MXDIAG, NTINY)
C           ----------------------------------------------
C           ... APPLY THE COLUMNS IN JBLK TO ANY COLUMNS
C               OF THE SUPERNODE REMAINING TO BE COMPUTED.
C           ----------------------------------------------
            NXTCOL = FSTCOL + NN
            Q = N - NXTCOL + 1
            MM = MM - NN
            JPNT = XPNT(NXTCOL)
            IF  ( Q .GT. 0 )  THEN
                CALL  MMPY8( MM, NN, Q, XPNT(FSTCOL), X, X(JPNT), MM )
            ENDIF
            FSTCOL = NXTCOL
            GO TO 100
        ENDIF
C
        RETURN
        END
C***********************************************************************
C***********************************************************************
C
C   Version:        0.4
C   Last modified:  December 27, 1994
C   Authors:        Esmond G. Ng and Barry W. Peyton
C
C   Mathematical Sciences Section, Oak Ridge National Laboratory
C
C***********************************************************************
C***********************************************************************
C**********     CHORDR ..... CHILD REORDERING                ***********
C***********************************************************************
C***********************************************************************
C
C   PURPOSE:
C       REARRANGE THE CHILDREN OF EACH VERTEX SO THAT THE LAST ONE 
C       MAXIMIZES (AMONG THE CHILDREN) THE NUMBER OF NONZEROS IN THE 
C       CORRESPONDING COLUMN OF L.  ALSO DETERMINE AN NEW POSTORDERING 
C       BASED ON THE STRUCTURE OF THE MODIFIED ELIMINATION TREE.
C
C   INPUT PARAMETERS:
C       NEQNS           -   NUMBER OF EQUATIONS.
C
C   UPDATED PARAMETERS:
C       (PERM,INVP)     -   ON INPUT, THE GIVEN PERM AND INVERSE PERM
C                           VECTORS.  ON OUTPUT, THE NEW PERM AND
C                           INVERSE PERM VECTORS OF THE NEW
C                           POSTORDERING.
C       COLCNT          -   COLUMN COUNTS IN L UNDER INITIAL ORDERING;
C                           MODIFIED TO REFLECT THE NEW ORDERING.
C
C   OUTPUT PARAMETERS:
C       PARENT          -   THE PARENT VECTOR OF THE ELIMINATION TREE
C                           ASSOCIATED WITH THE NEW ORDERING.
C
C   WORKING PARAMETERS:
C       FSON            -   THE FIRST SON VECTOR.
C       BROTHR          -   THE BROTHER VECTOR.
C       INVPOS          -   THE INVERSE PERM VECTOR FOR THE
C                           POSTORDERING.
C
C   PROGRAM SUBROUTINES:
C       BTREE2, EPOST2, INVINV.
C
C***********************************************************************
C
      SUBROUTINE  CHORDR (  NEQNS , PERM  , INVP  ,
     &                      COLCNT, PARENT, FSON  , BROTHR, INVPOS  )
C
C***********************************************************************
C
        INTEGER             BROTHR(*)     ,
     &                      COLCNT(*)     , FSON(*)       , 
     &                      INVP(*)       , INVPOS(*)     , 
     &                      PARENT(*)     , PERM(*)
C
        INTEGER             NEQNS
C
C***********************************************************************
C
C       ----------------------------------------------------------
C       COMPUTE A BINARY REPRESENTATION OF THE ELIMINATION TREE, 
C       SO THAT EACH "LAST CHILD" MAXIMIZES AMONG ITS SIBLINGS THE 
C       NUMBER OF NONZEROS IN THE CORRESPONDING COLUMNS OF L.
C       ----------------------------------------------------------
        CALL  BTREE2  ( NEQNS , PARENT, COLCNT, FSON  , BROTHR, 
     &                  INVPOS                                  )
C
C       ----------------------------------------------------
C       POSTORDER THE ELIMINATION TREE (USING THE NEW BINARY  
C       REPRESENTATION.  
C       ----------------------------------------------------
        CALL  EPOST2  ( NEQNS , FSON  , BROTHR, INVPOS, PARENT, 
     &                  COLCNT, PERM                            ) 
C
C       --------------------------------------------------------
C       COMPOSE THE ORIGINAL ORDERING WITH THE NEW POSTORDERING.
C       --------------------------------------------------------
        CALL  INVINV  ( NEQNS , INVP  , INVPOS, PERM    )
C
        RETURN
      END
C***********************************************************************
C***********************************************************************
C
C   Version:        0.4
C   Last modified:  December 27, 1994
C   Authors:        Esmond G. Ng and Barry W. Peyton
C
C   Mathematical Sciences Section, Oak Ridge National Laboratory
C
C***********************************************************************
C***********************************************************************
C******     DSCAL1 .... SCALE A VECTOR                     **************
C***********************************************************************
C***********************************************************************
C
C     PURPOSE - THIS ROUTINE COMPUTES A <-- AX, WHERE A IS A
C               SCALAR AND X IS A VECTOR.
C
C     INPUT PARAMETERS -
C        N - LENGTH OF THE VECTOR X.
C        A - SCALAR MULIPLIER.
C        X - VECTOR TO BE SCALED.
C
C     OUTPUT PARAMETERS - 
C        X - REPLACED BY THE SCALED VECTOR, AX.
C
C***********************************************************************
C
      SUBROUTINE  DSCAL1 ( N, A, X )
C
C***********************************************************************
C
C     -----------
C     PARAMETERS.
C     -----------
      INTEGER             N
      DOUBLE PRECISION    A, X(N)
C
C     ----------------
C     LOCAL VARIABLES.
C     ----------------
      INTEGER             I
C
C***********************************************************************
C
      DO  100  I = 1, N
          X(I) = A * X(I)
  100 CONTINUE
      RETURN
      END
C***********************************************************************
C***********************************************************************
C
C   Version:        0.4
C   Last modified:  December 27, 1994
C   Authors:        Esmond G. Ng and Barry W. Peyton
C
C   Mathematical Sciences Section, Oak Ridge National Laboratory
C
C***********************************************************************
C***********************************************************************
C***************     EPOST2 ..... ETREE POSTORDERING #2  ***************
C***********************************************************************
C***********************************************************************
C
C   PURPOSE:
C       BASED ON THE BINARY REPRESENTATION (FIRST-SON,BROTHER) OF THE 
C       ELIMINATION TREE, A POSTORDERING IS DETERMINED. THE
C       CORRESPONDING PARENT AND COLCNT VECTORS ARE ALSO MODIFIED TO 
C       REFLECT THE REORDERING.
C
C   INPUT PARAMETERS:
C       ROOT            -   ROOT OF THE ELIMINATION TREE (USUALLY IT
C                           IS NEQNS).
C       FSON            -   THE FIRST SON VECTOR.
C       BROTHR          -   THE BROTHR VECTOR.
C
C   UPDATED PARAMETERS:
C       PARENT          -   THE PARENT VECTOR.
C       COLCNT          -   COLUMN NONZERO COUNTS OF THE FACTOR.
C
C   OUTPUT PARAMETERS:
C       INVPOS          -   INVERSE PERMUTATION FOR THE POSTORDERING.
C
C   WORKING PARAMETERS:
C       STACK           -   THE STACK FOR POSTORDER TRAVERSAL OF THE
C                           TREE.
C
C***********************************************************************
C
      SUBROUTINE  EPOST2 (  ROOT  , FSON  , BROTHR, INVPOS, PARENT,
     &                      COLCNT, STACK                           )
C
C***********************************************************************
C
        INTEGER(4)           BROTHR(*)     , COLCNT(*)     , 
     &                      FSON(*)       , INVPOS(*)     , 
     &                      PARENT(*)     , STACK(*)
C
        INTEGER(4)           ROOT
C
C***********************************************************************
C
        INTEGER(4)           ITOP  , NDPAR , NODE  , NUM   , NUNODE
C
C***********************************************************************
C
        NUM = 0
        ITOP = 0
        NODE = ROOT
C       -------------------------------------------------------------
C       TRAVERSE ALONG THE FIRST SONS POINTER AND PUSH THE TREE NODES
C       ALONG THE TRAVERSAL INTO THE STACK.
C       -------------------------------------------------------------
  100   CONTINUE
            ITOP = ITOP + 1
            STACK(ITOP) = NODE
            NODE = FSON(NODE)
            IF  ( NODE .GT. 0 )  GO TO 100
C           ----------------------------------------------------------
C           IF POSSIBLE, POP A TREE NODE FROM THE STACK AND NUMBER IT.
C           ----------------------------------------------------------
  200       CONTINUE
                IF  ( ITOP .LE. 0 )  GO TO 300
                NODE = STACK(ITOP)
                ITOP = ITOP - 1
                NUM = NUM + 1
                INVPOS(NODE) = NUM
C               ----------------------------------------------------
C               THEN, TRAVERSE TO ITS YOUNGER BROTHER IF IT HAS ONE.
C               ----------------------------------------------------
                NODE = BROTHR(NODE)
                IF  ( NODE .LE. 0 )  GO TO 200
            GO TO 100
C
  300   CONTINUE
C       ------------------------------------------------------------
C       DETERMINE THE NEW PARENT VECTOR OF THE POSTORDERING.  BROTHR
C       IS USED TEMPORARILY FOR THE NEW PARENT VECTOR.
C       ------------------------------------------------------------
        DO  400  NODE = 1, NUM
            NUNODE = INVPOS(NODE)
            NDPAR = PARENT(NODE)
            IF  ( NDPAR .GT. 0 )  NDPAR = INVPOS(NDPAR)
            BROTHR(NUNODE) = NDPAR
  400   CONTINUE
C
        DO  500  NUNODE = 1, NUM
            PARENT(NUNODE) = BROTHR(NUNODE)
  500   CONTINUE
C
C       ----------------------------------------------
C       PERMUTE COLCNT(*) TO REFLECT THE NEW ORDERING.
C       ----------------------------------------------
        DO  600  NODE = 1, NUM
            NUNODE = INVPOS(NODE)
            STACK(NUNODE) = COLCNT(NODE)
  600   CONTINUE
C
        DO  700  NODE = 1, NUM
            COLCNT(NODE) = STACK(NODE)
  700   CONTINUE
C
        RETURN
      END
C***********************************************************************
C***********************************************************************
C
C   Version:        0.4
C   Last modified:  December 27, 1994
C   Authors:        Joseph W.H. Liu
C
C   Mathematical Sciences Section, Oak Ridge National Laboratory
C
C***********************************************************************
C***********************************************************************
C**********     ETORDR ..... ELIMINATION TREE REORDERING     ***********
C***********************************************************************
C***********************************************************************
C
C   WRITTEN BY JOSEPH LIU (JUL 17, 1985)
C
C   PURPOSE:
C       TO DETERMINE AN EQUIVALENT REORDERING BASED ON THE STRUCTURE OF
C       THE ELIMINATION TREE.  A POSTORDERING OF THE GIVEN ELIMINATION
C       TREE IS RETURNED.
C
C   INPUT PARAMETERS:
C       NEQNS           -   NUMBER OF EQUATIONS.
C       (XADJ,ADJNCY)   -   THE ADJACENCY STRUCTURE.
C
C   UPDATED PARAMETERS:
C       (PERM,INVP)     -   ON INPUT, THE GIVEN PERM AND INVERSE PERM
C                           VECTORS.  ON OUTPUT, THE NEW PERM AND
C                           INVERSE PERM VECTORS OF THE EQUIVALENT
C                           ORDERING.
C
C   OUTPUT PARAMETERS:
C       PARENT          -   THE PARENT VECTOR OF THE ELIMINATION TREE
C                           ASSOCIATED WITH THE NEW ORDERING.
C
C   WORKING PARAMETERS:
C       FSON            -   THE FIRST SON VECTOR.
C       BROTHR          -   THE BROTHER VECTOR.
C       INVPOS          -   THE INVERSE PERM VECTOR FOR THE
C                           POSTORDERING.
C
C   PROGRAM SUBROUTINES:
C       BETREE, ETPOST, ETREE , INVINV.
C
C***********************************************************************
C
      SUBROUTINE  ETORDR (  NEQNS , XADJ  , ADJNCY, PERM  , INVP  ,
     &                      PARENT, FSON  , BROTHR, INVPOS          )
C
C***********************************************************************
C
        INTEGER(4)           ADJNCY(*)     , BROTHR(*)     ,
     &                      FSON(*)       , INVP(*)       ,
     &                      INVPOS(*)     , PARENT(*)     ,
     &                      PERM(*)
C
        INTEGER(4)           XADJ(*)
        INTEGER(4)           NEQNS
C
C***********************************************************************
C
C       -----------------------------
C       COMPUTE THE ELIMINATION TREE.
C       -----------------------------
        CALL  ETREE ( NEQNS, XADJ, ADJNCY, PERM, INVP, PARENT, INVPOS )
C
C       --------------------------------------------------------
C       COMPUTE A BINARY REPRESENTATION OF THE ELIMINATION TREE.
C       --------------------------------------------------------
        CALL  BETREE ( NEQNS, PARENT, FSON, BROTHR )
C
C       -------------------------------
C       POSTORDER THE ELIMINATION TREE.
C       -------------------------------
        CALL  ETPOST ( NEQNS, FSON, BROTHR, INVPOS, PARENT, PERM )
C
C       --------------------------------------------------------
C       COMPOSE THE ORIGINAL ORDERING WITH THE NEW POSTORDERING.
C       --------------------------------------------------------
        CALL  INVINV ( NEQNS, INVP, INVPOS, PERM )
C
        RETURN
      END
C***********************************************************************
C***********************************************************************
C
C   Version:        0.4
C   Last modified:  December 27, 1994
C   Authors:        Joseph W.H. Liu
C
C   Mathematical Sciences Section, Oak Ridge National Laboratory
C
C***********************************************************************
C***********************************************************************
C***************     ETPOST ..... ETREE POSTORDERING     ***************
C***********************************************************************
C***********************************************************************
C
C   WRITTEN BY JOSEPH LIU (SEPT 17, 1986)
C
C   PURPOSE:
C       BASED ON THE BINARY REPRESENTATION (FIRST-SON,BROTHER) OF
C       THE ELIMINATION TREE, A POSTORDERING IS DETERMINED. THE
C       CORRESPONDING PARENT VECTOR IS ALSO MODIFIED TO REFLECT
C       THE REORDERING.
C
C   INPUT PARAMETERS:
C       ROOT            -   ROOT OF THE ELIMINATION TREE (USUALLY IT
C                           IS NEQNS).
C       FSON            -   THE FIRST SON VECTOR.
C       BROTHR          -   THE BROTHR VECTOR.
C
C   UPDATED PARAMETERS:
C       PARENT          -   THE PARENT VECTOR.
C
C   OUTPUT PARAMETERS:
C       INVPOS          -   INVERSE PERMUTATION FOR THE POSTORDERING.
C
C   WORKING PARAMETERS:
C       STACK           -   THE STACK FOR POSTORDER TRAVERSAL OF THE
C                           TREE.
C
C***********************************************************************
C
      SUBROUTINE  ETPOST (  ROOT  , FSON  , BROTHR, INVPOS, PARENT,
     &                      STACK                                   )
C
C***********************************************************************
C
        INTEGER(4)           BROTHR(*)     , FSON(*)       ,
     &                      INVPOS(*)     , PARENT(*)     ,
     &                      STACK(*)
C
        INTEGER(4)           ROOT
C
C***********************************************************************
C
        INTEGER(4)           ITOP  , NDPAR , NODE  , NUM   , NUNODE
C
C***********************************************************************
C
        NUM = 0
        ITOP = 0
        NODE = ROOT
C       -------------------------------------------------------------
C       TRAVERSE ALONG THE FIRST SONS POINTER AND PUSH THE TREE NODES
C       ALONG THE TRAVERSAL INTO THE STACK.
C       -------------------------------------------------------------
  100   CONTINUE
            ITOP = ITOP + 1
            STACK(ITOP) = NODE
            NODE = FSON(NODE)
            IF  ( NODE .GT. 0 )  GO TO 100
C           ----------------------------------------------------------
C           IF POSSIBLE, POP A TREE NODE FROM THE STACK AND NUMBER IT.
C           ----------------------------------------------------------
  200       CONTINUE
                IF  ( ITOP .LE. 0 )  GO TO 300
                NODE = STACK(ITOP)
                ITOP = ITOP - 1
                NUM = NUM + 1
                INVPOS(NODE) = NUM
C               ----------------------------------------------------
C               THEN, TRAVERSE TO ITS YOUNGER BROTHER IF IT HAS ONE.
C               ----------------------------------------------------
                NODE = BROTHR(NODE)
                IF  ( NODE .LE. 0 )  GO TO 200
            GO TO 100
C
  300   CONTINUE
C       ------------------------------------------------------------
C       DETERMINE THE NEW PARENT VECTOR OF THE POSTORDERING.  BROTHR
C       IS USED TEMPORARILY FOR THE NEW PARENT VECTOR.
C       ------------------------------------------------------------
        DO  400  NODE = 1, NUM
            NUNODE = INVPOS(NODE)
            NDPAR = PARENT(NODE)
            IF  ( NDPAR .GT. 0 )  NDPAR = INVPOS(NDPAR)
            BROTHR(NUNODE) = NDPAR
  400   CONTINUE
C
        DO  500  NUNODE = 1, NUM
            PARENT(NUNODE) = BROTHR(NUNODE)
  500   CONTINUE
C
        RETURN
      END
C***********************************************************************
C***********************************************************************
C
C   Version:        0.4
C   Last modified:  December 27, 1994
C   Authors:        Joseph W.H. Liu
C
C   Mathematical Sciences Section, Oak Ridge National Laboratory
C
C***********************************************************************
C***********************************************************************
C****************     ETREE ..... ELIMINATION TREE     *****************
C***********************************************************************
C***********************************************************************
C
C   WRITTEN BY JOSEPH LIU (JUL 17, 1985)
C
C   PURPOSE:
C       TO DETERMINE THE ELIMINATION TREE FROM A GIVEN ORDERING AND
C       THE ADJACENCY STRUCTURE.  THE PARENT VECTOR IS RETURNED.
C
C   INPUT PARAMETERS:
C       NEQNS           -   NUMBER OF EQUATIONS.
C       (XADJ,ADJNCY)   -   THE ADJACENCY STRUCTURE.
C       (PERM,INVP)     -   PERMUTATION AND INVERSE PERMUTATION VECTORS
C
C   OUTPUT PARAMETERS:
C       PARENT          -   THE PARENT VECTOR OF THE ELIMINATION TREE.
C
C   WORKING PARAMETERS:
C       ANCSTR          -   THE ANCESTOR VECTOR.
C
C***********************************************************************
C
      SUBROUTINE  ETREE (   NEQNS , XADJ  , ADJNCY, PERM  , INVP  ,
     &                      PARENT, ANCSTR                          )
C
C***********************************************************************
C
        INTEGER(4)           ADJNCY(*)     , ANCSTR(*)     ,
     &                      INVP(*)       , PARENT(*)     ,
     &                      PERM(*)
C
        INTEGER(4)           NEQNS
        INTEGER(4)           XADJ(*)
C
C***********************************************************************
C
        INTEGER(4)           I     , J     , JSTOP , JSTRT , NBR   ,
     &                      NEXT  , NODE
C
C***********************************************************************
C
        IF  ( NEQNS .LE. 0 )  RETURN
C
        DO  400  I = 1, NEQNS
            PARENT(I) = 0
            ANCSTR(I) = 0
            NODE = PERM(I)
C
            JSTRT = XADJ(NODE)
            JSTOP = XADJ(NODE+1) - 1
            IF  ( JSTRT .LE. JSTOP )  THEN
                DO  300  J = JSTRT, JSTOP
                    NBR = ADJNCY(J)
                    NBR = INVP(NBR)
                    IF  ( NBR .LT. I )  THEN
C                       -------------------------------------------
C                       FOR EACH NBR, FIND THE ROOT OF ITS CURRENT
C                       ELIMINATION TREE.  PERFORM PATH COMPRESSION
C                       AS THE SUBTREE IS TRAVERSED.
C                       -------------------------------------------
  100                   CONTINUE
                            IF  ( ANCSTR(NBR) .EQ. I )  GO TO 300
                            IF  ( ANCSTR(NBR) .GT. 0 )  THEN
                                NEXT = ANCSTR(NBR)
                                ANCSTR(NBR) = I
                                NBR = NEXT
                                GO TO 100
                            ENDIF
C                       --------------------------------------------
C                       NOW, NBR IS THE ROOT OF THE SUBTREE.  MAKE I
C                       THE PARENT NODE OF THIS ROOT.
C                       --------------------------------------------
                        PARENT(NBR) = I
                        ANCSTR(NBR) = I
                    ENDIF
  300           CONTINUE
            ENDIF
  400   CONTINUE
C
        RETURN
      END
C***********************************************************************
C***********************************************************************
C
C   Version:        0.4
C   Last modified:  January 12, 1995
C   Authors:        Esmond G. Ng and Barry W. Peyton
C
C   Mathematical Sciences Section, Oak Ridge National Laboratory
C
C***********************************************************************
C***********************************************************************
C**************     FCNTHN  ..... FIND NONZERO COUNTS    ***************
C***********************************************************************
C***********************************************************************
C
C   PURPOSE:
C       THIS SUBROUTINE DETERMINES THE ROW COUNTS AND COLUMN COUNTS IN
C       THE CHOLESKY FACTOR.  IT USES A DISJOINT SET UNION ALGORITHM.
C
C       TECHNIQUES:
C       1) SUPERNODE DETECTION.
C       2) PATH HALVING.
C       3) NO UNION BY RANK.
C
C   NOTES:
C       1) ASSUMES A POSTORDERING OF THE ELIMINATION TREE.
C
C   INPUT PARAMETERS:
C       (I) NEQNS       -   NUMBER OF EQUATIONS.
C       (I) ADJLEN      -   LENGTH OF ADJACENCY STRUCTURE.
C       (I) XADJ(*)     -   ARRAY OF LENGTH NEQNS+1, CONTAINING POINTERS
C                           TO THE ADJACENCY STRUCTURE.
C       (I) ADJNCY(*)   -   ARRAY OF LENGTH XADJ(NEQNS+1)-1, CONTAINING
C                           THE ADJACENCY STRUCTURE.
C       (I) PERM(*)     -   ARRAY OF LENGTH NEQNS, CONTAINING THE
C                           POSTORDERING.
C       (I) INVP(*)     -   ARRAY OF LENGTH NEQNS, CONTAINING THE
C                           INVERSE OF THE POSTORDERING.
C       (I) ETPAR(*)    -   ARRAY OF LENGTH NEQNS, CONTAINING THE
C                           ELIMINATION TREE OF THE POSTORDERED MATRIX.
C
C   OUTPUT PARAMETERS:
C       (I) ROWCNT(*)   -   ARRAY OF LENGTH NEQNS, CONTAINING THE NUMBER
C                           OF NONZEROS IN EACH ROW OF THE FACTOR,
C                           INCLUDING THE DIAGONAL ENTRY.
C       (I) COLCNT(*)   -   ARRAY OF LENGTH NEQNS, CONTAINING THE NUMBER
C                           OF NONZEROS IN EACH COLUMN OF THE FACTOR,
C                           INCLUDING THE DIAGONAL ENTRY.
C       (I) NLNZ        -   NUMBER OF NONZEROS IN THE FACTOR, INCLUDING
C                           THE DIAGONAL ENTRIES.
C
C   WORK PARAMETERS:
C       (I) SET(*)      -   ARRAY OF LENGTH NEQNS USED TO MAINTAIN THE
C                           DISJOINT SETS (I.E., SUBTREES).
C       (I) PRVLF(*)    -   ARRAY OF LENGTH NEQNS USED TO RECORD THE
C                           PREVIOUS LEAF OF EACH ROW SUBTREE.
C       (I) LEVEL(*)    -   ARRAY OF LENGTH NEQNS+1 CONTAINING THE LEVEL
C                           (DISTANCE FROM THE ROOT).
C       (I) WEIGHT(*)   -   ARRAY OF LENGTH NEQNS+1 CONTAINING WEIGHTS
C                           USED TO COMPUTE COLUMN COUNTS.
C       (I) FDESC(*)    -   ARRAY OF LENGTH NEQNS+1 CONTAINING THE
C                           FIRST (I.E., LOWEST-NUMBERED) DESCENDANT.
C       (I) NCHILD(*)   -   ARRAY OF LENGTH NEQNS+1 CONTAINING THE
C                           NUMBER OF CHILDREN.
C       (I) PRVNBR(*)   -   ARRAY OF LENGTH NEQNS USED TO RECORD THE
C                           PREVIOUS ``LOWER NEIGHBOR'' OF EACH NODE.
C
C   FIRST CREATED ON    APRIL 12, 1990.
C   LAST UPDATED ON     JANUARY 12, 1995.
C
C***********************************************************************
C
      SUBROUTINE FCNTHN  (  NEQNS , ADJLEN, XADJ  , ADJNCY, PERM  ,
     &                      INVP  , ETPAR , ROWCNT, COLCNT, NLNZ  ,
     &                      SET   , PRVLF , LEVEL , WEIGHT, FDESC ,
     &                      NCHILD, PRVNBR                          )
C
C       -----------
C       PARAMETERS.
C       -----------
        INTEGER             ADJLEN, NEQNS , NLNZ
        INTEGER             ADJNCY(ADJLEN)  , COLCNT(NEQNS) ,
     &                      ETPAR(NEQNS)    , FDESC(0:NEQNS),
     &                      INVP(NEQNS)     , LEVEL(0:NEQNS),
     &                      NCHILD(0:NEQNS) , PERM(NEQNS)   ,
     &                      PRVLF(NEQNS)    , PRVNBR(NEQNS) ,
     &                      ROWCNT(NEQNS)   , SET(NEQNS)    ,
     &                      WEIGHT(0:NEQNS)
        INTEGER             XADJ(*)
C
C       ----------------
C       LOCAL VARIABLES.
C       ----------------
        INTEGER             HINBR , IFDESC, J     , JSTOP , JSTRT , 
     &                      K     , LAST1 , LAST2 , LCA   , LFLAG , 
     &                      LOWNBR, OLDNBR, PARENT, PLEAF , TEMP  , 
     &                      XSUP
C
C***********************************************************************
C
C       --------------------------------------------------
C       COMPUTE LEVEL(*), FDESC(*), NCHILD(*).
C       INITIALIZE XSUP, ROWCNT(*), COLCNT(*),
C                  SET(*), PRVLF(*), WEIGHT(*), PRVNBR(*).
C       --------------------------------------------------
        XSUP = 1
        LEVEL(0) = 0
        DO  100  K = NEQNS, 1, -1
            ROWCNT(K) = 1
            COLCNT(K) = 0
            SET(K) = K
            PRVLF(K) = 0
            LEVEL(K) = LEVEL(ETPAR(K)) + 1
            WEIGHT(K) = 1
            FDESC(K) = K
            NCHILD(K) = 0
            PRVNBR(K) = 0
  100   CONTINUE
        NCHILD(0) = 0
        FDESC(0) = 0
        DO  200  K = 1, NEQNS
            PARENT = ETPAR(K)
            WEIGHT(PARENT) = 0
            NCHILD(PARENT) = NCHILD(PARENT) + 1
            IFDESC = FDESC(K)
            IF  ( IFDESC .LT. FDESC(PARENT) )  THEN
                FDESC(PARENT) = IFDESC
            ENDIF
  200   CONTINUE
C       ------------------------------------
C       FOR EACH ``LOW NEIGHBOR'' LOWNBR ...
C       ------------------------------------
        DO  600  LOWNBR = 1, NEQNS
            LFLAG = 0
            IFDESC = FDESC(LOWNBR)
            OLDNBR = PERM(LOWNBR)
            JSTRT = XADJ(OLDNBR)
            JSTOP = XADJ(OLDNBR+1) - 1
C           -----------------------------------------------
C           FOR EACH ``HIGH NEIGHBOR'', HINBR OF LOWNBR ...
C           -----------------------------------------------
            DO  500  J = JSTRT, JSTOP
                HINBR = INVP(ADJNCY(J))
                IF  ( HINBR .GT. LOWNBR )  THEN
                    IF  ( IFDESC .GT. PRVNBR(HINBR) )  THEN
C                       -------------------------
C                       INCREMENT WEIGHT(LOWNBR).
C                       -------------------------
                        WEIGHT(LOWNBR) = WEIGHT(LOWNBR) + 1
                        PLEAF = PRVLF(HINBR)
C                       -----------------------------------------
C                       IF HINBR HAS NO PREVIOUS ``LOW NEIGHBOR'' 
C                       THEN ...
C                       -----------------------------------------
                        IF  ( PLEAF .EQ. 0 )  THEN
C                           -----------------------------------------
C                           ... ACCUMULATE LOWNBR-->HINBR PATH LENGTH 
C                               IN ROWCNT(HINBR).
C                           -----------------------------------------
                            ROWCNT(HINBR) = ROWCNT(HINBR) +
     &                                  LEVEL(LOWNBR) - LEVEL(HINBR)
                        ELSE
C                           -----------------------------------------
C                           ... OTHERWISE, LCA <-- FIND(PLEAF), WHICH 
C                               IS THE LEAST COMMON ANCESTOR OF PLEAF 
C                               AND LOWNBR.
C                               (PATH HALVING.)
C                           -----------------------------------------
                            LAST1 = PLEAF
                            LAST2 = SET(LAST1)
                            LCA = SET(LAST2)
  300                       CONTINUE
                                IF  ( LCA .NE. LAST2 )  THEN
                                    SET(LAST1) = LCA
                                    LAST1 = LCA
                                    LAST2 = SET(LAST1)
                                    LCA = SET(LAST2)
                                    GO TO 300
                                ENDIF
C                           -------------------------------------
C                           ACCUMULATE PLEAF-->LCA PATH LENGTH IN 
C                           ROWCNT(HINBR).
C                           DECREMENT WEIGHT(LCA).
C                           -------------------------------------
                            ROWCNT(HINBR) = ROWCNT(HINBR)
     &                                  + LEVEL(LOWNBR) - LEVEL(LCA)
                            WEIGHT(LCA) = WEIGHT(LCA) - 1
                        ENDIF
C                       ----------------------------------------------
C                       LOWNBR NOW BECOMES ``PREVIOUS LEAF'' OF HINBR.
C                       ----------------------------------------------
                        PRVLF(HINBR) = LOWNBR
                        LFLAG = 1
                    ENDIF
C                   --------------------------------------------------
C                   LOWNBR NOW BECOMES ``PREVIOUS NEIGHBOR'' OF HINBR.
C                   --------------------------------------------------
                    PRVNBR(HINBR) = LOWNBR
                ENDIF
  500       CONTINUE
C           ----------------------------------------------------
C           DECREMENT WEIGHT ( PARENT(LOWNBR) ).
C           SET ( P(LOWNBR) ) <-- SET ( P(LOWNBR) ) + SET(XSUP).
C           ----------------------------------------------------
            PARENT = ETPAR(LOWNBR)
            WEIGHT(PARENT) = WEIGHT(PARENT) - 1
            IF  ( LFLAG .EQ. 1     .OR.
     &            NCHILD(LOWNBR) .GE. 2 )  THEN
                XSUP = LOWNBR
            ENDIF
            SET(XSUP) = PARENT
  600   CONTINUE
C       ---------------------------------------------------------
C       USE WEIGHTS TO COMPUTE COLUMN (AND TOTAL) NONZERO COUNTS.
C       ---------------------------------------------------------
        NLNZ = 0
        DO  700  K = 1, NEQNS
            TEMP = COLCNT(K) + WEIGHT(K)
            COLCNT(K) = TEMP
            NLNZ = NLNZ + TEMP
            PARENT = ETPAR(K)
            IF  ( PARENT .NE. 0 )  THEN
                COLCNT(PARENT) = COLCNT(PARENT) + TEMP
            ENDIF
  700   CONTINUE
C
        RETURN
      END
C***********************************************************************
C***********************************************************************
C
C   Version:        0.4
C   Last modified:  May 26, 1995
C   Authors:        Esmond G. Ng and Barry W. Peyton
C
C   Mathematical Sciences Section, Oak Ridge National Laboratory
C
C***********************************************************************
C***********************************************************************
C****     FNSPLT ..... COMPUTE FINE PARTITIONING OF SUPERNODES     *****
C***********************************************************************
C***********************************************************************
C
C   PURPOSE:
C       THIS SUBROUTINE DETERMINES A FINE PARTITIONING OF SUPERNODES
C       WHEN THERE IS A CACHE AVAILABLE ON THE MACHINE.  THE FINE
C       PARTITIONING IS CHOSEN SO THAT DATA RE-USE IS MAXIMIZED.
C       
C   INPUT PARAMETERS:
C       NEQNS           -   NUMBER OF EQUATIONS.
C       NSUPER          -   NUMBER OF SUPERNODES.
C       XSUPER          -   INTEGER ARRAY OF SIZE (NSUPER+1) CONTAINING
C                           THE SUPERNODE PARTITIONING.
C       XLINDX          -   INTEGER ARRAY OF SIZE (NSUPER+1) CONTAINING
C                           POINTERS IN THE SUPERNODE INDICES.
C       CACHSZ          -   CACHE SIZE IN KILO BYTES.
C                           IF THERE IS NO CACHE, SET CACHSZ = 0.
C
C   OUTPUT PARAMETERS:
C       SPLIT           -   INTEGER ARRAY OF SIZE NEQNS CONTAINING THE
C                           FINE PARTITIONING.
C
C***********************************************************************
C
        SUBROUTINE  FNSPLT ( NEQNS , NSUPER, XSUPER, XLINDX,
     &                       CACHSZ, SPLIT )
C
C***********************************************************************
C
C       -----------
C       PARAMETERS.
C       -----------
        INTEGER         CACHSZ, NEQNS , NSUPER
        INTEGER         XSUPER(*), SPLIT(*)
        INTEGER         XLINDX(*)
C
C       ----------------
C       LOCAL VARIABLES.
C       ----------------
        INTEGER         CACHE , CURCOL, FSTCOL, HEIGHT, KCOL  , 
     1                  KSUP  , LSTCOL, NCOLS , NXTBLK, USED  , 
     1                  WIDTH
C
C *******************************************************************
C
C       --------------------------------------------
C       COMPUTE THE NUMBER OF 8-BYTE WORDS IN CACHE.
C       --------------------------------------------
        IF  ( CACHSZ .LE. 0 )  THEN
            CACHE = 2 000 000 000
        ELSE
           CACHE = INT(( FLOAT(CACHSZ) * 1024. / 8. ) * 0.9)
        ENDIF
C
C       ---------------
C       INITIALIZATION.
C       ---------------
        DO  100  KCOL = 1, NEQNS
            SPLIT(KCOL) = 0
  100   CONTINUE
C
C       ---------------------------
C       FOR EACH SUPERNODE KSUP ...
C       ---------------------------
        DO  1000  KSUP = 1, NSUPER
C           -----------------------
C           ... GET SUPERNODE INFO.
C           -----------------------
            HEIGHT = XLINDX(KSUP+1) - XLINDX(KSUP)
            FSTCOL = XSUPER(KSUP)
            LSTCOL = XSUPER(KSUP+1) - 1
            WIDTH = LSTCOL - FSTCOL + 1
            NXTBLK = FSTCOL
C           --------------------------------------
C           ... UNTIL ALL COLUMNS OF THE SUPERNODE 
C               HAVE BEEN PROCESSED ...
C           --------------------------------------
            CURCOL = FSTCOL - 1
  200       CONTINUE
C               -------------------------------------------
C               ... PLACE THE FIRST COLUMN(S) IN THE CACHE.
C               -------------------------------------------
                CURCOL = CURCOL + 1
                IF  ( CURCOL .LT. LSTCOL )  THEN
                    CURCOL = CURCOL + 1
                    NCOLS = 2
                    USED = 4 * HEIGHT - 1
                    HEIGHT = HEIGHT - 2
                ELSE
                    NCOLS = 1
                    USED = 3 * HEIGHT
                    HEIGHT = HEIGHT - 1
                ENDIF
C
C               --------------------------------------
C               ... WHILE THE CACHE IS NOT FILLED AND
C                   THERE ARE COLUMNS OF THE SUPERNODE 
C                   REMAINING TO BE PROCESSED ...
C               --------------------------------------
  300           CONTINUE
                IF  ( USED+HEIGHT .LT. CACHE  .AND.
     &                CURCOL      .LT. LSTCOL       )  THEN
C                   --------------------------------
C                   ... ADD ANOTHER COLUMN TO CACHE.
C                   --------------------------------
                    CURCOL = CURCOL + 1
                    NCOLS = NCOLS + 1
                    USED = USED + HEIGHT
                    HEIGHT = HEIGHT - 1
                    GO TO 300
                ENDIF
C               -------------------------------------
C               ... RECORD THE NUMBER OF COLUMNS THAT 
C                   FILLED THE CACHE.
C               -------------------------------------
                SPLIT(NXTBLK) = NCOLS
                NXTBLK = NXTBLK + 1
C               --------------------------
C               ... GO PROCESS NEXT BLOCK.
C               --------------------------
                IF  ( CURCOL .LT. LSTCOL )  GO TO 200
 1000   CONTINUE
C
        RETURN
        END
C***********************************************************************
C***********************************************************************
C
C   Version:        0.4
C   Last modified:  December 27, 1994
C   Authors:        Esmond G. Ng and Barry W. Peyton
C
C   Mathematical Sciences Section, Oak Ridge National Laboratory
C
C***********************************************************************
C***********************************************************************
C******     FNTSIZ ..... COMPUTE WORK STORAGE SIZE FOR BLKFCT     ******
C***********************************************************************
C***********************************************************************
C
C   PURPOSE:
C       THIS SUBROUTINE DETERMINES THE SIZE OF THE WORKING STORAGE
C       REQUIRED BY BLKFCT.
C
C   INPUT PARAMETERS:
C       NSUPER          -   NUMBER OF SUPERNODES.
C       XSUPER          -   INTEGER ARRAY OF SIZE (NSUPER+1) CONTAINING
C                           THE SUPERNODE PARTITIONING.
C       SNODE           -   SUPERNODE MEMBERSHIP.
C       (XLINDX,LINDX)  -   ARRAYS DESCRIBING THE SUPERNODAL STRUCTURE.
C
C   OUTPUT PARAMETERS:
C       TMPSIZ          -   SIZE OF WORKING STORAGE REQUIRED BY BLKFCT.
C
C***********************************************************************
C
        SUBROUTINE  FNTSIZ ( NSUPER, XSUPER, SNODE , XLINDX, 
     &                       LINDX , TMPSIZ  )
C
C***********************************************************************
C
        INTEGER     NSUPER, TMPSIZ
        INTEGER     XLINDX(*)       , XSUPER(*)
        INTEGER     LINDX (*)       , SNODE (*)
C
        INTEGER     BOUND , CLEN  , CURSUP, I     , IBEGIN, IEND  , 
     &              KSUP  , LENGTH, NCOLS , NXTSUP, 
     &              TSIZE , WIDTH
C
C***********************************************************************
C
C       RETURNS SIZE OF TEMP ARRAY USED BY BLKFCT FACTORIZATION ROUTINE.
C       NOTE THAT THE VALUE RETURNED IS AN ESTIMATE, THOUGH IT IS USUALLY
C       TIGHT.
C
C       ----------------------------------------
C       COMPUTE SIZE OF TEMPORARY STORAGE VECTOR
C       NEEDED BY BLKFCT.
C       ----------------------------------------
        TMPSIZ = 0
        DO  500  KSUP = NSUPER, 1, -1
            NCOLS = XSUPER(KSUP+1) - XSUPER(KSUP)
            IBEGIN = XLINDX(KSUP) + NCOLS
            IEND = XLINDX(KSUP+1) - 1
            LENGTH = IEND - IBEGIN + 1
            BOUND = LENGTH * (LENGTH + 1) / 2
            IF  ( BOUND .GT. TMPSIZ )  THEN
                CURSUP = SNODE(LINDX(IBEGIN))
                CLEN = XLINDX(CURSUP+1) - XLINDX(CURSUP)
                WIDTH = 0
                DO  400  I = IBEGIN, IEND
                    NXTSUP = SNODE(LINDX(I))
                    IF  ( NXTSUP .EQ. CURSUP )  THEN
                        WIDTH = WIDTH + 1
                        IF  ( I .EQ. IEND )  THEN
                            IF  ( CLEN .GT. LENGTH )  THEN
                                TSIZE = LENGTH * WIDTH - 
     &                                  (WIDTH - 1) * WIDTH / 2
                                TMPSIZ = MAX ( TSIZE , TMPSIZ )
                            ENDIF
                        ENDIF
                    ELSE
                        IF  ( CLEN .GT. LENGTH )  THEN
                            TSIZE = LENGTH * WIDTH - 
     &                              (WIDTH - 1) * WIDTH / 2
                            TMPSIZ = MAX ( TSIZE , TMPSIZ )
                        ENDIF
                        LENGTH = LENGTH - WIDTH
                        BOUND = LENGTH * (LENGTH + 1) / 2
                        IF  ( BOUND .LE. TMPSIZ )  GO TO 500
                        WIDTH = 1
                        CURSUP = NXTSUP
                        CLEN = XLINDX(CURSUP+1) - XLINDX(CURSUP)
                    ENDIF
  400           CONTINUE
            ENDIF
  500   CONTINUE
C
        RETURN
        END
C***********************************************************************
C***********************************************************************
C
C   Version:        0.4
C   Last modified:  December 27, 1994
C   Authors:        Esmond G. Ng and Barry W. Peyton
C
C   Mathematical Sciences Section, Oak Ridge National Laboratory
C
C***********************************************************************
C***********************************************************************
C****************    FSUP1 ..... FIND SUPERNODES #1    *****************
C***********************************************************************
C***********************************************************************
C
C   PURPOSE:
C       THIS SUBROUTINE IS THE FIRST OF TWO ROUTINES FOR FINDING A
C       MAXIMAL SUPERNODE PARTITION.  IT RETURNS ONLY THE NUMBER OF
C       SUPERNODES NSUPER AND THE SUPERNODE MEMBERSHIP VECTOR SNODE(*), 
C       WHICH IS OF LENGTH NEQNS.  THE VECTORS OF LENGTH NSUPER ARE 
C       COMPUTED SUBSEQUENTLY BY THE COMPANION ROUTINE FSUP2.
C
C   METHOD AND ASSUMPTIONS:
C       THIS ROUTINE USES THE ELIMINATION TREE AND THE FACTOR COLUMN 
C       COUNTS TO COMPUTE THE SUPERNODE PARTITION; IT ALSO ASSUMES A 
C       POSTORDERING OF THE ELIMINATION TREE.
C
C   INPUT PARAMETERS:
C       (I) NEQNS       -   NUMBER OF EQUATIONS.
C       (I) ETPAR(*)    -   ARRAY OF LENGTH NEQNS, CONTAINING THE
C                           ELIMINATION TREE OF THE POSTORDERED MATRIX.
C       (I) COLCNT(*)   -   ARRAY OF LENGTH NEQNS, CONTAINING THE
C                           FACTOR COLUMN COUNTS: I.E., THE NUMBER OF 
C                           NONZERO ENTRIES IN EACH COLUMN OF L
C                           (INCLUDING THE DIAGONAL ENTRY).
C
C   OUTPUT PARAMETERS:
C       (I) NOFSUB      -   NUMBER OF SUBSCRIPTS.
C       (I) NSUPER      -   NUMBER OF SUPERNODES (<= NEQNS).
C       (I) SNODE(*)    -   ARRAY OF LENGTH NEQNS FOR RECORDING
C                           SUPERNODE MEMBERSHIP.
C
C   FIRST CREATED ON    JANUARY 18, 1992.
C   LAST UPDATED ON     NOVEMBER 11, 1994.
C
C***********************************************************************
C
      SUBROUTINE  FSUP1  (  NEQNS , ETPAR , COLCNT, NOFSUB, NSUPER,
     &                      SNODE                                   )
C
C***********************************************************************
C
C       -----------
C       PARAMETERS.
C       -----------
        INTEGER             NEQNS , NOFSUB, NSUPER
        INTEGER             COLCNT(*)     , ETPAR(*)      ,
     &                      SNODE(*)
C
C       ----------------
C       LOCAL VARIABLES.
C       ----------------
        INTEGER             KCOL
C
C***********************************************************************
C
C       --------------------------------------------
C       COMPUTE THE FUNDAMENTAL SUPERNODE PARTITION.
C       --------------------------------------------
        NSUPER = 1
        SNODE(1) = 1
        NOFSUB = COLCNT(1)
        DO  300  KCOL = 2, NEQNS
            IF  ( ETPAR(KCOL-1) .EQ. KCOL )  THEN
                IF  ( COLCNT(KCOL-1) .EQ. COLCNT(KCOL)+1 )  THEN
                    SNODE(KCOL) = NSUPER
                    GO TO 300
                ENDIF
            ENDIF
            NSUPER = NSUPER + 1
            SNODE(KCOL) = NSUPER
            NOFSUB = NOFSUB + COLCNT(KCOL)
  300   CONTINUE
C
      RETURN
      END
C***********************************************************************
C***********************************************************************
C
C   Version:        0.4
C   Last modified:  December 27, 1994
C   Authors:        Esmond G. Ng and Barry W. Peyton
C
C   Mathematical Sciences Section, Oak Ridge National Laboratory
C
C***********************************************************************
C***********************************************************************
C****************    FSUP2  ..... FIND SUPERNODES #2   *****************
C***********************************************************************
C***********************************************************************
C
C   PURPOSE:
C       THIS SUBROUTINE IS THE SECOND OF TWO ROUTINES FOR FINDING A
C       MAXIMAL SUPERNODE PARTITION.  IT'S SOLE PURPOSE IS TO 
C       CONSTRUCT THE NEEDED VECTOR OF LENGTH NSUPER: XSUPER(*).  THE
C       FIRST ROUTINE FSUP1 COMPUTES THE NUMBER OF SUPERNODES AND THE 
C       SUPERNODE MEMBERSHIP VECTOR SNODE(*), WHICH IS OF LENGTH NEQNS.
C
C
C   ASSUMPTIONS:
C       THIS ROUTINE ASSUMES A POSTORDERING OF THE ELIMINATION TREE.  IT
C       ALSO ASSUMES THAT THE OUTPUT FROM FSUP1 IS AVAILABLE.
C
C   INPUT PARAMETERS:
C       (I) NEQNS       -   NUMBER OF EQUATIONS.
C       (I) NSUPER      -   NUMBER OF SUPERNODES (<= NEQNS).
C       (I) SNODE(*)    -   ARRAY OF LENGTH NEQNS FOR RECORDING
C                           SUPERNODE MEMBERSHIP.
C
C   OUTPUT PARAMETERS:
C       (I) XSUPER(*)   -   ARRAY OF LENGTH NSUPER+1, CONTAINING THE
C                           SUPERNODE PARTITIONING.
C
C   FIRST CREATED ON    JANUARY 18, 1992.
C   LAST UPDATED ON     NOVEMEBER 22, 1994.
C
C***********************************************************************
C
      SUBROUTINE  FSUP2  (  NEQNS , NSUPER, SNODE , XSUPER  )
C
C***********************************************************************
C
C       -----------
C       PARAMETERS.
C       -----------
        INTEGER             NEQNS , NSUPER
        INTEGER             SNODE(*)      , 
     &                      XSUPER(*)
C
C       ----------------
C       LOCAL VARIABLES.
C       ----------------
        INTEGER             KCOL  , KSUP  , LSTSUP
C
C***********************************************************************
C
C       -------------------------------------------------
C       COMPUTE THE SUPERNODE PARTITION VECTOR XSUPER(*).
C       -------------------------------------------------
        LSTSUP = NSUPER + 1
        DO  100  KCOL = NEQNS, 1, -1
            KSUP = SNODE(KCOL)
            IF  ( KSUP .NE. LSTSUP )  THEN
                XSUPER(LSTSUP) = KCOL + 1
            ENDIF
            LSTSUP = KSUP
  100   CONTINUE
        XSUPER(1) = 1
C
      RETURN
      END
C***********************************************************************
C***********************************************************************
C
C   Version:        0.4
C   Last modified:  December 27, 1994
C   Authors:        Joseph W.H. Liu
C
C   Mathematical Sciences Section, Oak Ridge National Laboratory
C
C***********************************************************************
C***********************************************************************
C--- SPARSPAK-A (ANSI FORTRAN) RELEASE III --- NAME = GENMMD
C  (C)  UNIVERSITY OF WATERLOO   JANUARY 1984
C***********************************************************************
C***********************************************************************
C****     GENMMD ..... MULTIPLE MINIMUM EXTERNAL DEGREE     ************
C***********************************************************************
C***********************************************************************
C
C     PURPOSE - THIS ROUTINE IMPLEMENTS THE MINIMUM DEGREE
C        ALGORITHM.  IT MAKES USE OF THE IMPLICIT REPRESENTATION
C        OF ELIMINATION GRAPHS BY QUOTIENT GRAPHS, AND THE
C        NOTION OF INDISTINGUISHABLE NODES.  IT ALSO IMPLEMENTS
C        THE MODIFICATIONS BY MULTIPLE ELIMINATION AND MINIMUM
C        EXTERNAL DEGREE.
C        ---------------------------------------------
C        CAUTION - THE ADJACENCY VECTOR ADJNCY WILL BE
C        DESTROYED.
C        ---------------------------------------------
C
C     INPUT PARAMETERS -
C        NEQNS  - NUMBER OF EQUATIONS.
C        (XADJ,ADJNCY) - THE ADJACENCY STRUCTURE.
C        DELTA  - TOLERANCE VALUE FOR MULTIPLE ELIMINATION.
C        MAXINT - MAXIMUM MACHINE REPRESENTABLE (SHORT) INTEGER
C                 (ANY SMALLER ESTIMATE WILL DO) FOR MARKING
C                 NODES.
C
C     OUTPUT PARAMETERS -
C        PERM   - THE MINIMUM DEGREE ORDERING.
C        INVP   - THE INVERSE OF PERM.
C        NOFSUB - AN UPPER BOUND ON THE NUMBER OF NONZERO
C                 SUBSCRIPTS FOR THE COMPRESSED STORAGE SCHEME.
C
C     WORKING PARAMETERS -
C        DHEAD  - VECTOR FOR HEAD OF DEGREE LISTS.
C        INVP   - USED TEMPORARILY FOR DEGREE FORWARD LINK.
C        PERM   - USED TEMPORARILY FOR DEGREE BACKWARD LINK.
C        QSIZE  - VECTOR FOR SIZE OF SUPERNODES.
C        LLIST  - VECTOR FOR TEMPORARY LINKED LISTS.
C        MARKER - A TEMPORARY MARKER VECTOR.
C
C     PROGRAM SUBROUTINES -
C        MMDELM, MMDINT, MMDNUM, MMDUPD.
C
C***********************************************************************
C
      SUBROUTINE  GENMMD ( NEQNS, XADJ, ADJNCY, INVP, PERM,
     1                     DELTA, DHEAD, QSIZE, LLIST, MARKER,
     1                     MAXINT, NOFSUB )
C
C***********************************************************************
C
         INTEGER    ADJNCY(*), DHEAD(*) , INVP(*)  , LLIST(*) ,
     1              MARKER(*), PERM(*)  , QSIZE(*)
         INTEGER    XADJ(*)
         INTEGER    DELTA , EHEAD , I     , MAXINT, MDEG  ,
     1              MDLMT , MDNODE, NEQNS , NEXTMD, NOFSUB,
     1              NUM, TAG
C
C***********************************************************************
C
         IF  ( NEQNS .LE. 0 )  RETURN
C
C        ------------------------------------------------
C        INITIALIZATION FOR THE MINIMUM DEGREE ALGORITHM.
C        ------------------------------------------------
         NOFSUB = 0
         CALL  MMDINT ( NEQNS, XADJ, DHEAD, INVP, PERM,
     1                  QSIZE, LLIST, MARKER )
C
C        ----------------------------------------------
C        NUM COUNTS THE NUMBER OF ORDERED NODES PLUS 1.
C        ----------------------------------------------
         NUM = 1
C
C        -----------------------------
C        ELIMINATE ALL ISOLATED NODES.
C        -----------------------------
         NEXTMD = DHEAD(1)
  100    CONTINUE
             IF  ( NEXTMD .LE. 0 )  GO TO 200
                 MDNODE = NEXTMD
                 NEXTMD = INVP(MDNODE)
                 MARKER(MDNODE) = MAXINT
                 INVP(MDNODE) = - NUM
                 NUM = NUM + 1
                 GO TO 100
C
  200    CONTINUE
C        ----------------------------------------
C        SEARCH FOR NODE OF THE MINIMUM DEGREE.
C        MDEG IS THE CURRENT MINIMUM DEGREE;
C        TAG IS USED TO FACILITATE MARKING NODES.
C        ----------------------------------------
         IF  ( NUM .GT. NEQNS )  GO TO 1000
         TAG = 1
         DHEAD(1) = 0
         MDEG = 2
  300    CONTINUE
             IF  ( DHEAD(MDEG) .GT. 0 )  GO TO 400
                 MDEG = MDEG + 1
                 GO TO 300
  400        CONTINUE
C            -------------------------------------------------
C            USE VALUE OF DELTA TO SET UP MDLMT, WHICH GOVERNS
C            WHEN A DEGREE UPDATE IS TO BE PERFORMED.
C            -------------------------------------------------
             MDLMT = MDEG + DELTA
             EHEAD = 0
C
  500        CONTINUE
                 MDNODE = DHEAD(MDEG)
                 IF  ( MDNODE .GT. 0 )  GO TO 600
                     MDEG = MDEG + 1
                     IF  ( MDEG .GT. MDLMT )  GO TO 900
                         GO TO 500
  600            CONTINUE
C                ----------------------------------------
C                REMOVE MDNODE FROM THE DEGREE STRUCTURE.
C                ----------------------------------------
                 NEXTMD = INVP(MDNODE)
                 DHEAD(MDEG) = NEXTMD
                 IF  ( NEXTMD .GT. 0 )  PERM(NEXTMD) = - MDEG
                 INVP(MDNODE) = - NUM
                 NOFSUB = NOFSUB + MDEG + QSIZE(MDNODE) - 2
                 IF  ( NUM+QSIZE(MDNODE) .GT. NEQNS )  GO TO 1000
C                ----------------------------------------------
C                ELIMINATE MDNODE AND PERFORM QUOTIENT GRAPH
C                TRANSFORMATION.  RESET TAG VALUE IF NECESSARY.
C                ----------------------------------------------
                 TAG = TAG + 1
                 IF  ( TAG .LT. MAXINT )  GO TO 800
                     TAG = 1
                     DO  700  I = 1, NEQNS
                         IF  ( MARKER(I) .LT. MAXINT )  MARKER(I) = 0
  700                CONTINUE
  800            CONTINUE
                 CALL  MMDELM ( MDNODE, XADJ, ADJNCY, DHEAD, INVP,
     1                          PERM, QSIZE, LLIST, MARKER, MAXINT,
     1                          TAG )
                 NUM = NUM + QSIZE(MDNODE)
                 LLIST(MDNODE) = EHEAD
                 EHEAD = MDNODE
                 IF  ( DELTA .GE. 0 )  GO TO 500
  900        CONTINUE
C            -------------------------------------------
C            UPDATE DEGREES OF THE NODES INVOLVED IN THE
C            MINIMUM DEGREE NODES ELIMINATION.
C            -------------------------------------------
             IF  ( NUM .GT. NEQNS )  GO TO 1000
             CALL  MMDUPD ( EHEAD, NEQNS, XADJ, ADJNCY, DELTA, MDEG,
     1                      DHEAD, INVP, PERM, QSIZE, LLIST, MARKER,
     1                      MAXINT, TAG )
             GO TO 300
C
 1000    CONTINUE
         CALL  MMDNUM ( NEQNS, PERM, INVP, QSIZE )
         RETURN
C
      END
C***********************************************************************
C***********************************************************************
C
C   Version:        0.4
C   Last modified:  December 27, 1994
C   Authors:        Esmond G. Ng and Barry W. Peyton
C
C   Mathematical Sciences Section, Oak Ridge National Laboratory
C
C***********************************************************************
C***********************************************************************
C******         IGATHR .... INTEGER GATHER OPERATION      **************
C***********************************************************************
C***********************************************************************
C
C     PURPOSE - THIS ROUTINE PERFORMS A STANDARD INTEGER GATHER
C               OPERATION.
C
C     INPUT PARAMETERS -
C        KLEN   - LENGTH OF THE LIST OF GLOBAL INDICES.
C        LINDX  - LIST OF GLOBAL INDICES.
C        INDMAP - INDEXED BY GLOBAL INDICES, IT CONTAINS THE
C                 REQUIRED RELATIVE INDICES.
C
C     OUTPUT PARAMETERS - 
C        RELIND - LIST RELATIVE INDICES.
C
C***********************************************************************
C
      SUBROUTINE  IGATHR ( KLEN  , LINDX, INDMAP, RELIND )
C
C***********************************************************************
C
C     -----------
C     PARAMETERS.
C     -----------
      INTEGER             KLEN  
      INTEGER             INDMAP(*), LINDX (*), RELIND(*)
C
C     ----------------
C     LOCAL VARIABLES.
C     ----------------
      INTEGER             I
C
C***********************************************************************
C
CDIR$ IVDEP
      DO  100  I = 1, KLEN  
          RELIND(I) = INDMAP(LINDX(I))
  100 CONTINUE
      RETURN
      END
C***********************************************************************
C***********************************************************************
C
C   Version:        0.4
C   Last modified:  December 27, 1994
C   Authors:        Esmond G. Ng and Barry W. Peyton
C
C   Mathematical Sciences Section, Oak Ridge National Laboratory
C
C***********************************************************************
C***********************************************************************
C
C     ------------------------------------------------------
C     INPUT NUMERICAL VALUES INTO SPARSE DATA STRUCTURES ...
C     ------------------------------------------------------
C
      SUBROUTINE  INPNV  (  XADJF, ADJF, ANZF, PERM, INVP,
     &                      NSUPER, XSUPER, XLINDX, LINDX,
     &                      XLNZ, LNZ, OFFSET )
C
        INTEGER             XADJF(*), ADJF(*)
        DOUBLE PRECISION    ANZF(*)
        INTEGER             PERM(*), INVP(*)
        INTEGER             NSUPER
        INTEGER             XSUPER(*), XLINDX(*), LINDX(*)
        INTEGER             XLNZ(*)
        DOUBLE PRECISION    LNZ(*)
        INTEGER             OFFSET(*)
C
        INTEGER             I, II, J, JLEN, JSUPER, LAST, OLDJ
C
        DO  500  JSUPER = 1, NSUPER
C
C           ----------------------------------------
C           FOR EACH SUPERNODE, DO THE FOLLOWING ...
C           ----------------------------------------
C
C           -----------------------------------------------
C           FIRST GET OFFSET TO FACILITATE NUMERICAL INPUT.
C           -----------------------------------------------
            JLEN = XLINDX(JSUPER+1) - XLINDX(JSUPER)
            DO  100  II = XLINDX(JSUPER), XLINDX(JSUPER+1)-1
                I = LINDX(II)
                JLEN = JLEN - 1
                OFFSET(I) = JLEN
  100       CONTINUE
C
            DO  400  J = XSUPER(JSUPER), XSUPER(JSUPER+1)-1
C               -----------------------------------------
C               FOR EACH COLUMN IN THE CURRENT SUPERNODE,
C               FIRST INITIALIZE THE DATA STRUCTURE.
C               -----------------------------------------
c                DO  200  II = XLNZ(J), XLNZ(J+1)-1
c                    LNZ(II) = 0.0
c  200           CONTINUE
c     The previous lines are not required as R initializes the arrays
c     Reinhard Furrer, Nov 19, 2007
C
C               -----------------------------------
C               NEXT INPUT THE INDIVIDUAL NONZEROS.
C               -----------------------------------
                OLDJ = PERM(J)
                LAST = XLNZ(J+1) - 1
                DO  300  II = XADJF(OLDJ), XADJF(OLDJ+1)-1
                    I = INVP(ADJF(II))
                    IF  ( I .GE. J )  THEN
                        LNZ(LAST-OFFSET(I)) = ANZF(II)
                    ENDIF
  300           CONTINUE
  400       CONTINUE
C
  500   CONTINUE
        RETURN
      END
C***********************************************************************
C***********************************************************************
C
C   Version:        0.4
C   Last modified:  December 27, 1994
C   Authors:        Joseph W.H. Liu
C
C   Mathematical Sciences Section, Oak Ridge National Laboratory
C
C***********************************************************************
C***********************************************************************
C***********     INVINV ..... CONCATENATION OF TWO INVP     ************
C***********************************************************************
C***********************************************************************
C
C   WRITTEN BY JOSEPH LIU (JUL 17, 1985)
C
C   PURPOSE:
C       TO PERFORM THE MAPPING OF
C           ORIGINAL-INVP --> INTERMEDIATE-INVP --> NEW INVP
C       AND THE RESULTING ORDERING REPLACES INVP.  THE NEW PERMUTATION
C       VECTOR PERM IS ALSO COMPUTED.
C
C   INPUT PARAMETERS:
C       NEQNS           -   NUMBER OF EQUATIONS.
C       INVP2           -   THE SECOND INVERSE PERMUTATION VECTOR.
C
C   UPDATED PARAMETERS:
C       INVP            -   THE FIRST INVERSE PERMUTATION VECTOR.  ON
C                           OUTPUT, IT CONTAINS THE NEW INVERSE
C                           PERMUTATION.
C
C   OUTPUT PARAMETER:
C       PERM            -   NEW PERMUTATION VECTOR (CAN BE THE SAME AS
C                           INVP2).
C
C***********************************************************************
C
      SUBROUTINE  INVINV (  NEQNS , INVP  , INVP2 , PERM            )
C
C***********************************************************************
C
        INTEGER(4)           INVP(*)       , INVP2(*)      ,
     &                      PERM(*)
C
        INTEGER(4)           NEQNS
C
C***********************************************************************
C
        INTEGER(4)           I     , INTERM, NODE
C
C***********************************************************************
C
        DO  100  I = 1, NEQNS
            INTERM = INVP(I)
            INVP(I) = INVP2(INTERM)
  100   CONTINUE
C
        DO  200  I = 1, NEQNS
            NODE = INVP(I)
            PERM(NODE) = I
  200   CONTINUE
C
        RETURN
      END
C***********************************************************************
C***********************************************************************
C
C   Version:        0.4
C   Last modified:  December 27, 1994
C   Authors:        Esmond G. Ng and Barry W. Peyton
C
C   Mathematical Sciences Section, Oak Ridge National Laboratory
C
C***********************************************************************
C***********************************************************************
C******         LDINDX .... LOAD INDEX VECTOR             **************
C***********************************************************************
C***********************************************************************
C
C     PURPOSE - THIS ROUTINE COMPUTES THE SECOND INDEX VECTOR
C               USED TO IMPLEMENT THE DOUBLY-INDIRECT SAXPY-LIKE
C               LOOPS THAT ALLOW US TO ACCUMULATE UPDATE 
C               COLUMNS DIRECTLY INTO FACTOR STORAGE.
C
C     INPUT PARAMETERS -
C        JLEN   - LENGTH OF THE FIRST COLUMN OF THE SUPERNODE,
C                 INCLUDING THE DIAGONAL ENTRY.
C        LINDX  - THE OFF-DIAGONAL ROW INDICES OF THE SUPERNODE, 
C                 I.E., THE ROW INDICES OF THE NONZERO ENTRIES
C                 LYING BELOW THE DIAGONAL ENTRY OF THE FIRST
C                 COLUMN OF THE SUPERNODE.
C
C     OUTPUT PARAMETERS - 
C        INDMAP - THIS INDEX VECTOR MAPS EVERY GLOBAL ROW INDEX
C                 OF NONZERO ENTRIES IN THE FIRST COLUMN OF THE 
C                 SUPERNODE TO ITS POSITION IN THE INDEX LIST 
C                 RELATIVE TO THE LAST INDEX IN THE LIST.  MORE
C                 PRECISELY, IT GIVES THE DISTANCE OF EACH INDEX
C                 FROM THE LAST INDEX IN THE LIST.
C
C***********************************************************************
C
      SUBROUTINE  LDINDX ( JLEN, LINDX, INDMAP )
C
C***********************************************************************
C
C     -----------
C     PARAMETERS.
C     -----------
      INTEGER             JLEN
      INTEGER             LINDX(*), INDMAP(*)
C
C     ----------------
C     LOCAL VARIABLES.
C     ----------------
      INTEGER             CURLEN, J, JSUB
C
C***********************************************************************
C

      CURLEN = JLEN

      DO  200  J = 1, JLEN
          JSUB = LINDX(J)
          CURLEN = CURLEN - 1
          INDMAP(JSUB) = CURLEN
  200 CONTINUE
      RETURN
      END
C***********************************************************************
C***********************************************************************
C
C   Version:        0.4
C   Last modified:  December 27, 1994
C   Authors:        Joseph W.H. Liu
C
C   Mathematical Sciences Section, Oak Ridge National Laboratory
C
C***********************************************************************
C***********************************************************************
C--- SPARSPAK-A (ANSI FORTRAN) RELEASE III --- NAME = MMDELM
C  (C)  UNIVERSITY OF WATERLOO   JANUARY 1984
C***********************************************************************
C***********************************************************************
C**     MMDELM ..... MULTIPLE MINIMUM DEGREE ELIMINATION     ***********
C***********************************************************************
C***********************************************************************
C
C     PURPOSE - THIS ROUTINE ELIMINATES THE NODE MDNODE OF
C        MINIMUM DEGREE FROM THE ADJACENCY STRUCTURE, WHICH
C        IS STORED IN THE QUOTIENT GRAPH FORMAT.  IT ALSO
C        TRANSFORMS THE QUOTIENT GRAPH REPRESENTATION OF THE
C        ELIMINATION GRAPH.
C
C     INPUT PARAMETERS -
C        MDNODE - NODE OF MINIMUM DEGREE.
C        MAXINT - ESTIMATE OF MAXIMUM REPRESENTABLE (SHORT)
C                 INTEGER.
C        TAG    - TAG VALUE.
C
C     UPDATED PARAMETERS -
C        (XADJ,ADJNCY) - UPDATED ADJACENCY STRUCTURE.
C        (DHEAD,DFORW,DBAKW) - DEGREE DOUBLY LINKED STRUCTURE.
C        QSIZE  - SIZE OF SUPERNODE.
C        MARKER - MARKER VECTOR.
C        LLIST  - TEMPORARY LINKED LIST OF ELIMINATED NABORS.
C
C***********************************************************************
C
      SUBROUTINE  MMDELM ( MDNODE, XADJ, ADJNCY, DHEAD, DFORW,
     1                     DBAKW, QSIZE, LLIST, MARKER, MAXINT,
     1                     TAG )
C
C***********************************************************************
C
         INTEGER    ADJNCY(*), DBAKW(*) , DFORW(*) , DHEAD(*) ,
     1              LLIST(*) , MARKER(*), QSIZE(*)
         INTEGER    XADJ(*)
         INTEGER    ELMNT , I     , ISTOP , ISTRT , J     ,
     1              JSTOP , JSTRT , LINK  , MAXINT, MDNODE,
     1              NABOR , NODE  , NPV   , NQNBRS, NXNODE,
     1              PVNODE, RLMT  , RLOC  , RNODE , TAG   ,
     1              XQNBR
C
C***********************************************************************
C
C        -----------------------------------------------
C        FIND REACHABLE SET AND PLACE IN DATA STRUCTURE.
C        -----------------------------------------------
         MARKER(MDNODE) = TAG
         ISTRT = XADJ(MDNODE)
         ISTOP = XADJ(MDNODE+1) - 1
C        -------------------------------------------------------
C        ELMNT POINTS TO THE BEGINNING OF THE LIST OF ELIMINATED
C        NABORS OF MDNODE, AND RLOC GIVES THE STORAGE LOCATION
C        FOR THE NEXT REACHABLE NODE.
C        -------------------------------------------------------
         ELMNT = 0
         RLOC = ISTRT
         RLMT = ISTOP
         DO  200  I = ISTRT, ISTOP
             NABOR = ADJNCY(I)
             IF  ( NABOR .EQ. 0 )  GO TO 300
                 IF  ( MARKER(NABOR) .GE. TAG )  GO TO 200
                     MARKER(NABOR) = TAG
                     IF  ( DFORW(NABOR) .LT. 0 )  GO TO 100
                         ADJNCY(RLOC) = NABOR
                         RLOC = RLOC + 1
                         GO TO 200
  100                CONTINUE
                     LLIST(NABOR) = ELMNT
                     ELMNT = NABOR
  200    CONTINUE
  300    CONTINUE
C            -----------------------------------------------------
C            MERGE WITH REACHABLE NODES FROM GENERALIZED ELEMENTS.
C            -----------------------------------------------------
             IF  ( ELMNT .LE. 0 )  GO TO 1000
                 ADJNCY(RLMT) = - ELMNT
                 LINK = ELMNT
  400            CONTINUE
                     JSTRT = XADJ(LINK)
                     JSTOP = XADJ(LINK+1) - 1
                     DO  800  J = JSTRT, JSTOP
                         NODE = ADJNCY(J)
                         LINK = - NODE
C                         IF  ( NODE )  400, 900, 500
                         if ( NODE .LT. 0) GO TO 400
                         if ( NODE .EQ. 0) GO TO 900
C  500                    CONTINUE
                         IF  ( MARKER(NODE) .GE. TAG  .OR.
     1                         DFORW(NODE) .LT. 0 )  GO TO 800
                             MARKER(NODE) = TAG
C                            ---------------------------------
C                            USE STORAGE FROM ELIMINATED NODES
C                            IF NECESSARY.
C                            ---------------------------------
  600                        CONTINUE
                                 IF  ( RLOC .LT. RLMT )  GO TO 700
                                     LINK = - ADJNCY(RLMT)
                                     RLOC = XADJ(LINK)
                                     RLMT = XADJ(LINK+1) - 1
                                     GO TO 600
  700                        CONTINUE
                             ADJNCY(RLOC) = NODE
                             RLOC = RLOC + 1
  800                CONTINUE
  900            CONTINUE
                 ELMNT = LLIST(ELMNT)
                 GO TO 300
 1000    CONTINUE
         IF  ( RLOC .LE. RLMT )  ADJNCY(RLOC) = 0
C        --------------------------------------------------------
C        FOR EACH NODE IN THE REACHABLE SET, DO THE FOLLOWING ...
C        --------------------------------------------------------
         LINK = MDNODE
 1100    CONTINUE
             ISTRT = XADJ(LINK)
             ISTOP = XADJ(LINK+1) - 1
             DO  1700  I = ISTRT, ISTOP
                 RNODE = ADJNCY(I)
                 LINK = - RNODE
C                 IF  ( RNODE )  1100, 1800, 1200
                 if ( RNODE .LT. 0) GO TO 1100
                 if ( RNODE .EQ. 0) GO TO 1800
C 1200            CONTINUE
C                --------------------------------------------
C                IF RNODE IS IN THE DEGREE LIST STRUCTURE ...
C                --------------------------------------------
                 PVNODE = DBAKW(RNODE)
                 IF  ( PVNODE .EQ. 0  .OR.
     1                 PVNODE .EQ. (-MAXINT) )  GO TO 1300
C                    -------------------------------------
C                    THEN REMOVE RNODE FROM THE STRUCTURE.
C                    -------------------------------------
                     NXNODE = DFORW(RNODE)
                     IF  ( NXNODE .GT. 0 )  DBAKW(NXNODE) = PVNODE
                     IF  ( PVNODE .GT. 0 )  DFORW(PVNODE) = NXNODE
                     NPV = - PVNODE
                     IF  ( PVNODE .LT. 0 )  DHEAD(NPV) = NXNODE
 1300            CONTINUE
C                ----------------------------------------
C                PURGE INACTIVE QUOTIENT NABORS OF RNODE.
C                ----------------------------------------
                 JSTRT = XADJ(RNODE)
                 JSTOP = XADJ(RNODE+1) - 1
                 XQNBR = JSTRT
                 DO  1400  J = JSTRT, JSTOP
                     NABOR = ADJNCY(J)
                     IF  ( NABOR .EQ. 0 )  GO TO 1500
                         IF  ( MARKER(NABOR) .GE. TAG )  GO TO 1400
                             ADJNCY(XQNBR) = NABOR
                             XQNBR = XQNBR + 1
 1400            CONTINUE
 1500            CONTINUE
C                ----------------------------------------
C                IF NO ACTIVE NABOR AFTER THE PURGING ...
C                ----------------------------------------
                 NQNBRS = XQNBR - JSTRT
                 IF  ( NQNBRS .GT. 0 )  GO TO 1600
C                    -----------------------------
C                    THEN MERGE RNODE WITH MDNODE.
C                    -----------------------------
                     QSIZE(MDNODE) = QSIZE(MDNODE) + QSIZE(RNODE)
                     QSIZE(RNODE) = 0
                     MARKER(RNODE) = MAXINT
                     DFORW(RNODE) = - MDNODE
                     DBAKW(RNODE) = - MAXINT
                     GO TO 1700
 1600            CONTINUE
C                --------------------------------------
C                ELSE FLAG RNODE FOR DEGREE UPDATE, AND
C                ADD MDNODE AS A NABOR OF RNODE.
C                --------------------------------------
                 DFORW(RNODE) = NQNBRS + 1
                 DBAKW(RNODE) = 0
                 ADJNCY(XQNBR) = MDNODE
                 XQNBR = XQNBR + 1
                 IF  ( XQNBR .LE. JSTOP )  ADJNCY(XQNBR) = 0
C
 1700        CONTINUE
 1800    CONTINUE
         RETURN
C
      END
C***********************************************************************
C***********************************************************************
C
C   Version:        0.4
C   Last modified:  December 27, 1994
C   Authors:        Joseph W.H. Liu
C
C   Mathematical Sciences Section, Oak Ridge National Laboratory
C
C***********************************************************************
C***********************************************************************
C--- SPARSPAK-A (ANSI FORTRAN) RELEASE III --- NAME = MMDINT
C  (C)  UNIVERSITY OF WATERLOO   JANUARY 1984
C***********************************************************************
C***********************************************************************
C***     MMDINT ..... MULT MINIMUM DEGREE INITIALIZATION     ***********
C***********************************************************************
C***********************************************************************
C
C     PURPOSE - THIS ROUTINE PERFORMS INITIALIZATION FOR THE
C        MULTIPLE ELIMINATION VERSION OF THE MINIMUM DEGREE
C        ALGORITHM.
C
C     INPUT PARAMETERS -
C        NEQNS  - NUMBER OF EQUATIONS.
C        (XADJ,ADJNCY) - ADJACENCY STRUCTURE.
C
C     OUTPUT PARAMETERS -
C        (DHEAD,DFORW,DBAKW) - DEGREE DOUBLY LINKED STRUCTURE.
C        QSIZE  - SIZE OF SUPERNODE (INITIALIZED TO ONE).
C        LLIST  - LINKED LIST.
C        MARKER - MARKER VECTOR.
C
C***********************************************************************
C
      SUBROUTINE  MMDINT ( NEQNS, XADJ, DHEAD, DFORW,
     1                     DBAKW, QSIZE, LLIST, MARKER )
C
C***********************************************************************
C
         INTEGER    DBAKW(*) , DFORW(*) , DHEAD(*) ,
     1              LLIST(*) , MARKER(*), QSIZE(*)
         INTEGER    XADJ(*)
         INTEGER    FNODE , NDEG  , NEQNS , NODE
C
C***********************************************************************
C
         DO  100  NODE = 1, NEQNS
             DHEAD(NODE) = 0
             QSIZE(NODE) = 1
             MARKER(NODE) = 0
             LLIST(NODE) = 0
  100    CONTINUE
C        ------------------------------------------
C        INITIALIZE THE DEGREE DOUBLY LINKED LISTS.
C        ------------------------------------------
         DO  200  NODE = 1, NEQNS
             NDEG = XADJ(NODE+1) - XADJ(NODE) + 1
             FNODE = DHEAD(NDEG)
             DFORW(NODE) = FNODE
             DHEAD(NDEG) = NODE
             IF  ( FNODE .GT. 0 )  DBAKW(FNODE) = NODE
             DBAKW(NODE) = - NDEG
  200    CONTINUE
         RETURN
C
      END
C***********************************************************************
C***********************************************************************
C
C   Version:        0.4
C   Last modified:  December 27, 1994
C   Authors:        Joseph W.H. Liu
C
C   Mathematical Sciences Section, Oak Ridge National Laboratory
C
C***********************************************************************
C***********************************************************************
C--- SPARSPAK-A (ANSI FORTRAN) RELEASE III --- NAME = MMDNUM
C  (C)  UNIVERSITY OF WATERLOO   JANUARY 1984
C***********************************************************************
C***********************************************************************
C*****     MMDNUM ..... MULTI MINIMUM DEGREE NUMBERING     *************
C***********************************************************************
C***********************************************************************
C
C     PURPOSE - THIS ROUTINE PERFORMS THE FINAL STEP IN
C        PRODUCING THE PERMUTATION AND INVERSE PERMUTATION
C        VECTORS IN THE MULTIPLE ELIMINATION VERSION OF THE
C        MINIMUM DEGREE ORDERING ALGORITHM.
C
C     INPUT PARAMETERS -
C        NEQNS  - NUMBER OF EQUATIONS.
C        QSIZE  - SIZE OF SUPERNODES AT ELIMINATION.
C
C     UPDATED PARAMETERS -
C        INVP   - INVERSE PERMUTATION VECTOR.  ON INPUT,
C                 IF QSIZE(NODE)=0, THEN NODE HAS BEEN MERGED
C                 INTO THE NODE -INVP(NODE); OTHERWISE,
C                 -INVP(NODE) IS ITS INVERSE LABELLING.
C
C     OUTPUT PARAMETERS -
C        PERM   - THE PERMUTATION VECTOR.
C
C***********************************************************************
C
      SUBROUTINE  MMDNUM ( NEQNS, PERM, INVP, QSIZE )
C
C***********************************************************************
C
         INTEGER    INVP(*)  , PERM(*)  , QSIZE(*)
         INTEGER    FATHER, NEQNS , NEXTF , NODE  , NQSIZE,
     1              NUM   , ROOT
C
C***********************************************************************
C
         DO  100  NODE = 1, NEQNS
             NQSIZE = QSIZE(NODE)
             IF  ( NQSIZE .LE. 0 )  PERM(NODE) = INVP(NODE)
             IF  ( NQSIZE .GT. 0 )  PERM(NODE) = - INVP(NODE)
  100    CONTINUE
C        ------------------------------------------------------
C        FOR EACH NODE WHICH HAS BEEN MERGED, DO THE FOLLOWING.
C        ------------------------------------------------------
         DO  500  NODE = 1, NEQNS
             IF  ( PERM(NODE) .GT. 0 )  GO TO 500
C                -----------------------------------------
C                TRACE THE MERGED TREE UNTIL ONE WHICH HAS
C                NOT BEEN MERGED, CALL IT ROOT.
C                -----------------------------------------
                 FATHER = NODE
  200            CONTINUE
                     IF  ( PERM(FATHER) .GT. 0 )  GO TO 300
                         FATHER = - PERM(FATHER)
                         GO TO 200
  300            CONTINUE
C                -----------------------
C                NUMBER NODE AFTER ROOT.
C                -----------------------
                 ROOT = FATHER
                 NUM = PERM(ROOT) + 1
                 INVP(NODE) = - NUM
                 PERM(ROOT) = NUM
C                ------------------------
C                SHORTEN THE MERGED TREE.
C                ------------------------
                 FATHER = NODE
  400            CONTINUE
                     NEXTF = - PERM(FATHER)
                     IF  ( NEXTF .LE. 0 )  GO TO 500
                         PERM(FATHER) = - ROOT
                         FATHER = NEXTF
                         GO TO 400
  500    CONTINUE
C        ----------------------
C        READY TO COMPUTE PERM.
C        ----------------------
         DO  600  NODE = 1, NEQNS
             NUM = - INVP(NODE)
             INVP(NODE) = NUM
             PERM(NUM) = NODE
  600    CONTINUE
         RETURN
C
      END
C***********************************************************************
C***********************************************************************
C
C   Version:        0.4
C   Last modified:  December 27, 1994
C   Authors:        Joseph W.H. Liu
C
C   Mathematical Sciences Section, Oak Ridge National Laboratory
C
C***********************************************************************
C***********************************************************************
C--- SPARSPAK-A (ANSI FORTRAN) RELEASE III --- NAME = MMDUPD
C  (C)  UNIVERSITY OF WATERLOO   JANUARY 1984
C***********************************************************************
C***********************************************************************
C*****     MMDUPD ..... MULTIPLE MINIMUM DEGREE UPDATE     *************
C***********************************************************************
C***********************************************************************
C
C     PURPOSE - THIS ROUTINE UPDATES THE DEGREES OF NODES
C        AFTER A MULTIPLE ELIMINATION STEP.
C
C     INPUT PARAMETERS -
C        EHEAD  - THE BEGINNING OF THE LIST OF ELIMINATED
C                 NODES (I.E., NEWLY FORMED ELEMENTS).
C        NEQNS  - NUMBER OF EQUATIONS.
C        (XADJ,ADJNCY) - ADJACENCY STRUCTURE.
C        DELTA  - TOLERANCE VALUE FOR MULTIPLE ELIMINATION.
C        MAXINT - MAXIMUM MACHINE REPRESENTABLE (SHORT)
C                 INTEGER.
C
C     UPDATED PARAMETERS -
C        MDEG   - NEW MINIMUM DEGREE AFTER DEGREE UPDATE.
C        (DHEAD,DFORW,DBAKW) - DEGREE DOUBLY LINKED STRUCTURE.
C        QSIZE  - SIZE OF SUPERNODE.
C        LLIST  - WORKING LINKED LIST.
C        MARKER - MARKER VECTOR FOR DEGREE UPDATE.
C        TAG    - TAG VALUE.
C
C***********************************************************************
C
      SUBROUTINE  MMDUPD ( EHEAD, NEQNS, XADJ, ADJNCY, DELTA,
     1                     MDEG, DHEAD, DFORW, DBAKW, QSIZE,
     1                     LLIST, MARKER, MAXINT, TAG )
C
C***********************************************************************
C
         INTEGER    ADJNCY(*), DBAKW(*) , DFORW(*) , DHEAD(*) ,
     1              LLIST(*) , MARKER(*), QSIZE(*)
         INTEGER    XADJ(*)
         INTEGER    DEG   , DEG0  , DELTA , EHEAD , ELMNT ,
     1              ENODE , FNODE , I     , IQ2   , ISTOP ,
     1              ISTRT , J     , JSTOP , JSTRT , LINK  ,
     1              MAXINT, MDEG  , MDEG0 , MTAG  , NABOR ,
     1              NEQNS , NODE  , Q2HEAD, QXHEAD, TAG
C
C***********************************************************************
C
         MDEG0 = MDEG + DELTA
         ELMNT = EHEAD
  100    CONTINUE
C            -------------------------------------------------------
C            FOR EACH OF THE NEWLY FORMED ELEMENT, DO THE FOLLOWING.
C            (RESET TAG VALUE IF NECESSARY.)
C            -------------------------------------------------------
             IF  ( ELMNT .LE. 0 )  RETURN
             MTAG = TAG + MDEG0
             IF  ( MTAG .LT. MAXINT )  GO TO 300
                 TAG = 1
                 DO  200  I = 1, NEQNS
                     IF  ( MARKER(I) .LT. MAXINT )  MARKER(I) = 0
  200            CONTINUE
                 MTAG = TAG + MDEG0
  300        CONTINUE
C            ---------------------------------------------
C            CREATE TWO LINKED LISTS FROM NODES ASSOCIATED
C            WITH ELMNT: ONE WITH TWO NABORS (Q2HEAD) IN
C            ADJACENCY STRUCTURE, AND THE OTHER WITH MORE
C            THAN TWO NABORS (QXHEAD).  ALSO COMPUTE DEG0,
C            NUMBER OF NODES IN THIS ELEMENT.
C            ---------------------------------------------
             Q2HEAD = 0
             QXHEAD = 0
             DEG0 = 0
             LINK = ELMNT
  400        CONTINUE
                 ISTRT = XADJ(LINK)
                 ISTOP = XADJ(LINK+1) - 1
                 DO  700  I = ISTRT, ISTOP
                     ENODE = ADJNCY(I)
                     LINK = - ENODE
C                     IF  ( ENODE )  400, 800, 500
                         if ( ENODE .LT. 0) GO TO 400
                         if ( ENODE .EQ. 0) GO TO 800
C
C 500                 CONTINUE
                     IF  ( QSIZE(ENODE) .EQ. 0 )  GO TO 700
                         DEG0 = DEG0 + QSIZE(ENODE)
                         MARKER(ENODE) = MTAG
C                        ----------------------------------
C                        IF ENODE REQUIRES A DEGREE UPDATE,
C                        THEN DO THE FOLLOWING.
C                        ----------------------------------
                         IF  ( DBAKW(ENODE) .NE. 0 )  GO TO 700
C                            ---------------------------------------
C                            PLACE EITHER IN QXHEAD OR Q2HEAD LISTS.
C                            ---------------------------------------
                             IF  ( DFORW(ENODE) .EQ. 2 )  GO TO 600
                                 LLIST(ENODE) = QXHEAD
                                 QXHEAD = ENODE
                                 GO TO 700
  600                        CONTINUE
                             LLIST(ENODE) = Q2HEAD
                             Q2HEAD = ENODE
  700            CONTINUE
  800        CONTINUE
C            --------------------------------------------
C            FOR EACH ENODE IN Q2 LIST, DO THE FOLLOWING.
C            --------------------------------------------
             ENODE = Q2HEAD
             IQ2 = 1
  900        CONTINUE
                 IF  ( ENODE .LE. 0 )  GO TO 1500
                 IF  ( DBAKW(ENODE) .NE. 0 )  GO TO 2200
                     TAG = TAG + 1
                     DEG = DEG0
C                    ------------------------------------------
C                    IDENTIFY THE OTHER ADJACENT ELEMENT NABOR.
C                    ------------------------------------------
                     ISTRT = XADJ(ENODE)
                     NABOR = ADJNCY(ISTRT)
                     IF  ( NABOR .EQ. ELMNT )  NABOR = ADJNCY(ISTRT+1)
C                    ------------------------------------------------
C                    IF NABOR IS UNELIMINATED, INCREASE DEGREE COUNT.
C                    ------------------------------------------------
                     LINK = NABOR
                     IF  ( DFORW(NABOR) .LT. 0 )  GO TO 1000
                         DEG = DEG + QSIZE(NABOR)
                         GO TO 2100
 1000                CONTINUE
C                        --------------------------------------------
C                        OTHERWISE, FOR EACH NODE IN THE 2ND ELEMENT,
C                        DO THE FOLLOWING.
C                        --------------------------------------------
                         ISTRT = XADJ(LINK)
                         ISTOP = XADJ(LINK+1) - 1
                         DO  1400  I = ISTRT, ISTOP
                             NODE = ADJNCY(I)
                             LINK = - NODE
                             IF  ( NODE .EQ. ENODE )  GO TO 1400
C                             IF  ( NODE )  1000, 2100, 1100
                             if ( NODE .LT. 0 ) GO TO 1000
                             if ( NODE .EQ. 0 ) GO TO 2100

C
C 1100                        CONTINUE
                             IF  ( QSIZE(NODE) .EQ. 0 )  GO TO 1400
                             IF  ( MARKER(NODE) .GE. TAG )  GO TO 1200
C                                -------------------------------------
C                                CASE WHEN NODE IS NOT YET CONSIDERED.
C                                -------------------------------------
                                 MARKER(NODE) = TAG
                                 DEG = DEG + QSIZE(NODE)
                                 GO TO 1400
 1200                        CONTINUE
C                            ----------------------------------------
C                            CASE WHEN NODE IS INDISTINGUISHABLE FROM
C                            ENODE.  MERGE THEM INTO A NEW SUPERNODE.
C                            ----------------------------------------
                             IF  ( DBAKW(NODE) .NE. 0 )  GO TO 1400
                             IF  ( DFORW(NODE) .NE. 2 )  GO TO 1300
                                 QSIZE(ENODE) = QSIZE(ENODE) +
     1                                          QSIZE(NODE)
                                 QSIZE(NODE) = 0
                                 MARKER(NODE) = MAXINT
                                 DFORW(NODE) = - ENODE
                                 DBAKW(NODE) = - MAXINT
                                 GO TO 1400
 1300                        CONTINUE
C                            --------------------------------------
C                            CASE WHEN NODE IS OUTMATCHED BY ENODE.
C                            --------------------------------------
                             IF  ( DBAKW(NODE) .EQ.0 )
     1                             DBAKW(NODE) = - MAXINT
 1400                    CONTINUE
                         GO TO 2100
 1500            CONTINUE
C                ------------------------------------------------
C                FOR EACH ENODE IN THE QX LIST, DO THE FOLLOWING.
C                ------------------------------------------------
                 ENODE = QXHEAD
                 IQ2 = 0
 1600            CONTINUE
                     IF  ( ENODE .LE. 0 )  GO TO 2300
                     IF  ( DBAKW(ENODE) .NE. 0 )  GO TO 2200
                         TAG = TAG + 1
                         DEG = DEG0
C                        ---------------------------------
C                        FOR EACH UNMARKED NABOR OF ENODE,
C                        DO THE FOLLOWING.
C                        ---------------------------------
                         ISTRT = XADJ(ENODE)
                         ISTOP = XADJ(ENODE+1) - 1
                         DO  2000  I = ISTRT, ISTOP
                             NABOR = ADJNCY(I)
                             IF  ( NABOR .EQ. 0 )  GO TO 2100
                             IF  ( MARKER(NABOR) .GE. TAG )  GO TO 2000
                                 MARKER(NABOR) = TAG
                                 LINK = NABOR
C                                ------------------------------
C                                IF UNELIMINATED, INCLUDE IT IN
C                                DEG COUNT.
C                                ------------------------------
                                 IF  ( DFORW(NABOR) .LT. 0 )  GO TO 1700
                                     DEG = DEG + QSIZE(NABOR)
                                     GO TO 2000
 1700                            CONTINUE
C                                    -------------------------------
C                                    IF ELIMINATED, INCLUDE UNMARKED
C                                    NODES IN THIS ELEMENT INTO THE
C                                    DEGREE COUNT.
C                                    -------------------------------
                                     JSTRT = XADJ(LINK)
                                     JSTOP = XADJ(LINK+1) - 1
                                     DO  1900  J = JSTRT, JSTOP
                                         NODE = ADJNCY(J)
                                         LINK = - NODE
C                                         IF  ( NODE )  1700, 2000, 1800
                                         if ( NODE .LT. 0) GO TO 1700
                                         if ( NODE .EQ. 0) GO TO 2000
C
C 1800                                    CONTINUE
                                         IF  ( MARKER(NODE) .GE. TAG )
     1                                         GO TO 1900
                                             MARKER(NODE) = TAG
                                             DEG = DEG + QSIZE(NODE)
 1900                                CONTINUE
 2000                    CONTINUE
 2100                CONTINUE
C                    -------------------------------------------
C                    UPDATE EXTERNAL DEGREE OF ENODE IN DEGREE
C                    STRUCTURE, AND MDEG (MIN DEG) IF NECESSARY.
C                    -------------------------------------------
                     DEG = DEG - QSIZE(ENODE) + 1
                     FNODE = DHEAD(DEG)
                     DFORW(ENODE) = FNODE
                     DBAKW(ENODE) = - DEG
                     IF  ( FNODE .GT. 0 )  DBAKW(FNODE) = ENODE
                     DHEAD(DEG) = ENODE
                     IF  ( DEG .LT. MDEG )  MDEG = DEG
 2200                CONTINUE
C                    ----------------------------------
C                    GET NEXT ENODE IN CURRENT ELEMENT.
C                    ----------------------------------
                     ENODE = LLIST(ENODE)
                     IF  ( IQ2 .EQ. 1 )  GO TO 900
                         GO TO 1600
 2300        CONTINUE
C            -----------------------------
C            GET NEXT ELEMENT IN THE LIST.
C            -----------------------------
             TAG = MTAG
             ELMNT = LLIST(ELMNT)
             GO TO 100
C
      END
C***********************************************************************
C***********************************************************************
C
C   Version:        0.4
C   Last modified:  December 27, 1994
C   Authors:        Esmond G. Ng and Barry W. Peyton
C                   RF: modified mmpy8 dependence
C
C   Mathematical Sciences Section, Oak Ridge National Laboratory
C
C***********************************************************************
C***********************************************************************
C**************     MMPY  .... MATRIX-MATRIX MULTIPLY     **************
C***********************************************************************
C***********************************************************************
C
C   PURPOSE -
C       THIS ROUTINE PERFORMS A MATRIX-MATRIX MULTIPLY, Y = Y + XA,
C       ASSUMING DATA STRUCTURES USED IN SOME OF OUR SPARSE CHOLESKY
C       CODES.
C
C   INPUT PARAMETERS -
C       M               -   NUMBER OF ROWS IN X AND IN Y.
C       N               -   NUMBER OF COLUMNS IN X AND NUMBER OF ROWS
C                           IN A.
C       Q               -   NUMBER OF COLUMNS IN A AND Y.
C       SPLIT(*)        -   BLOCK PARTITIONING OF X.
C       XPNT(*)         -   XPNT(J+1) POINTS ONE LOCATION BEYOND THE
C                           END OF THE J-TH COLUMN OF X.  XPNT IS ALSO
C                           USED TO ACCESS THE ROWS OF A.
C       X(*)            -   CONTAINS THE COLUMNS OF X AND THE ROWS OF A.
C       LDY             -   LENGTH OF FIRST COLUMN OF Y.
C
C   EXTERNAL ROUTINES:
C       MMPYN           -   MATRIX-MATRIX MULTIPLY,
C                           WITH LEVEL 8 LOOP UNROLLING.
C
C   UPDATED PARAMETERS -
C       Y(*)            -   ON OUTPUT, Y = Y + AX.
C
C***********************************************************************
C
      SUBROUTINE  MMPY   (  M     , N     , Q     , SPLIT , XPNT  ,
     &                      X     , Y     , LDY    )
C
C***********************************************************************
C
C       -----------
C       PARAMETERS.
C       -----------
C
        EXTERNAL            MMPY8
        INTEGER             LDY   , M     , N     , Q
        INTEGER             SPLIT(*)      , XPNT(*)
        DOUBLE PRECISION    X(*)          , Y(*)
C
C       ----------------
C       LOCAL VARIABLES.
C       ----------------
C
        INTEGER             BLK   , FSTCOL, NN
C
C***********************************************************************
C
        BLK = 1
        FSTCOL = 1
  100   CONTINUE
        IF  ( FSTCOL .LE. N )  THEN
            NN = SPLIT(BLK)
            CALL  MMPY8 ( M, NN, Q, XPNT(FSTCOL), X, Y, LDY )
            FSTCOL = FSTCOL + NN
            BLK = BLK + 1
            GO TO 100
        ENDIF
        RETURN
C
      END
C***********************************************************************
C***********************************************************************
C
C   Version:        0.4
C   Last modified:  May 26, 1995
C   Authors:        Esmond G. Ng, Barry W. Peyton, and Guodong Zhang
C
C   Mathematical Sciences Section, Oak Ridge National Laboratory
C
C***********************************************************************
C***********************************************************************
C*************     MMPY8  .... MATRIX-MATRIX MULTIPLY     **************
C***********************************************************************
C***********************************************************************
C
C   PURPOSE -
C       THIS ROUTINE PERFORMS A MATRIX-MATRIX MULTIPLY, Y = Y + XA,
C       ASSUMING DATA STRUCTURES USED IN SOME OF OUR SPARSE CHOLESKY
C       CODES.
C
C       LOOP UNROLLING: LEVEL 8 UPDATING TWO COLUMNS AT A TIME
C
C   INPUT PARAMETERS -
C       M               -   NUMBER OF ROWS IN X AND IN Y.
C       N               -   NUMBER OF COLUMNS IN X AND NUMBER OF ROWS
C                           IN A.
C       Q               -   NUMBER OF COLUMNS IN A AND Y.
C       XPNT(*)         -   XPNT(J+1) POINTS ONE LOCATION BEYOND THE
C                           END OF THE J-TH COLUMN OF X.  XPNT IS ALSO
C                           USED TO ACCESS THE ROWS OF A.
C       X(*)            -   CONTAINS THE COLUMNS OF X AND THE ROWS OF A.
C       LDY             -   LENGTH OF FIRST COLUMN OF Y.
C
C   UPDATED PARAMETERS -
C       Y(*)            -   ON OUTPUT, Y = Y + AX.
C
C***********************************************************************
C
        SUBROUTINE  MMPY8  (  M     , N     , Q     , XPNT  , X     ,
     &                        Y     , LDY                             )
C
C***********************************************************************
C
C       -----------
C       PARAMETERS.
C       -----------
C
        INTEGER               LDY   , M     , N     , Q
        INTEGER               XPNT(*)
        DOUBLE PRECISION      X(*)          , Y(*)
C
C       ----------------
C       LOCAL VARIABLES.
C       ----------------
C
        INTEGER               I     , J     , K     , QQ
        INTEGER               I1    , I2    , I3    , I4    , I5    ,
     &                        I6    , I7    , I8
        INTEGER               IYBEG , IYBEG1, IYBEG2, LENY  , MM    
        DOUBLE PRECISION      A1    , A2    , A3    , A4    , A5    ,
     &                        A6    , A7    , A8    , A9    , A10   ,
     &                        A11   , A12   , A13   , A14   , A15   ,
     &                        A16
        DOUBLE PRECISION      B1    , B2    , B3    , B4    , B5    ,
     &                        B6    , B7    , B8    , Y1    , Y2
C
C***********************************************************************
C
C       ----------------------------------------------------
C       COMPUTE EACH DIAGONAL ENTRY OF THE ODD COLUMNS OF Y.
C       ----------------------------------------------------
C
        MM = M
        QQ = MIN(M,Q)
        IYBEG = 1
        LENY = LDY - 1
        DO  200 J = 1, QQ-1 , 2
CDIR$   IVDEP
            DO  100  I = 1, N
                I1 = XPNT(I+1) - MM
                A1 = X(I1)
                Y(IYBEG) = Y(IYBEG) - A1*A1
  100       CONTINUE
            IYBEG = IYBEG + 2*LENY + 1
            LENY = LENY - 2
            MM = MM - 2
  200   CONTINUE
C       
C       -------------------------------------------------------
C       UPDATE TWO COLUMNS OF Y AT A TIME,  EXCEPT THE DIAGONAL 
C       ELEMENT.
C       NOTE: THE DIAGONAL ELEMENT OF THE ODD COLUMN HAS
C             BEEN COMPUTED, SO WE COMPUTE THE SAME NUMBER OF
C             ELEMENTS FOR THE TWO COLUMNS.
C       -------------------------------------------------------
C
        MM = M
        IYBEG = 1
        LENY = LDY - 1 
C
        DO  3000  J = 1, QQ-1, 2
C
            IYBEG1 = IYBEG 
            IYBEG2 = IYBEG + LENY
C
            DO  400  K = 1, N-7, 8
C
C               -----------------------------------
C               EIGHT COLUMNS UPDATING TWO COLUMNS.
C               -----------------------------------
C
                I1 = XPNT(K+1) - MM
                I2 = XPNT(K+2) - MM
                I3 = XPNT(K+3) - MM
                I4 = XPNT(K+4) - MM
                I5 = XPNT(K+5) - MM
                I6 = XPNT(K+6) - MM
                I7 = XPNT(K+7) - MM
                I8 = XPNT(K+8) - MM
                A1 = X(I1)
                A2 = X(I2)
                A3 = X(I3)
                A4 = X(I4)
                A5 = X(I5)
                A6 = X(I6)
                A7 = X(I7)
                A8 = X(I8)
                A9  = X(I1+1)
                A10 = X(I2+1)
                A11 = X(I3+1)
                A12 = X(I4+1)
                A13 = X(I5+1)
                A14 = X(I6+1)
                A15 = X(I7+1)
                A16 = X(I8+1)
C
                Y(IYBEG1+1) =  Y(IYBEG1+1) -
     &              A1*A9 - A2*A10 - A3*A11 - A4*A12 - A5*A13 - 
     &              A6*A14 - A7*A15 - A8*A16
C
                Y(IYBEG2+1) =  Y(IYBEG2+1) -
     &              A9*A9 - A10*A10 - A11*A11 - A12*A12 - A13*A13 - 
     &              A14*A14 - A15*A15 - A16*A16
C
                DO  300  I = 2, MM-1
                    Y1 = Y(IYBEG1+I)
                    B1 = X(I1+I)
                    Y1 =  Y1 - B1 * A1
                    Y2 = Y(IYBEG2+I)
                    B2 = X(I2+I)
                    Y2 =  Y2 - B1 * A9
                    Y1 =  Y1 - B2 * A2
                    B3 = X(I3+I)
                    Y2 =  Y2 - B2 * A10
                    Y1 =  Y1 - B3 * A3
                    B4 = X(I4+I)
                    Y2 =  Y2 - B3 * A11
                    Y1 =  Y1 - B4 * A4
                    B5 = X(I5+I)
                    Y2 =  Y2 - B4 * A12
                    Y1 =  Y1 - B5 * A5
                    B6 = X(I6+I)
                    Y2 =  Y2 - B5 * A13
                    Y1 =  Y1 - B6 * A6
                    B7 = X(I7+I)
                    Y2 =  Y2 - B6 * A14
                    Y1 = Y1 - B7 * A7
                    B8 = X(I8+I)
                    Y2 = Y2 - B7 * A15
                    Y1 = Y1 - B8 * A8       
                    Y(IYBEG1+I) = Y1
                    Y2 =  Y2 - B8 * A16
                    Y(IYBEG2+I) = Y2               
  300           CONTINUE
C
  400       CONTINUE
C
C           -----------------------------
C           BOUNDARY CODE FOR THE K LOOP.
C           -----------------------------
C
C             GO TO ( 2000, 1700, 1500, 1300,
C     &              1100,  900,  700,  500  ), N-K+2
             if (N .LT. K) go to 2000
             if (N .EQ. K) go to 1700
             if (N .EQ. K + 1) go to 1500
             if (N .EQ. K + 2) go to 1300
             if (N .EQ. K + 3) go to 1100
             if (N .EQ. K + 4) go to 900
             if (N .EQ. K + 5) go to 700
C
C  500       CONTINUE
C
C               -----------------------------------
C               SEVEN COLUMNS UPDATING TWO COLUMNS.
C               -----------------------------------
C
                I1 = XPNT(K+1) - MM
                I2 = XPNT(K+2) - MM
                I3 = XPNT(K+3) - MM
                I4 = XPNT(K+4) - MM
                I5 = XPNT(K+5) - MM
                I6 = XPNT(K+6) - MM
                I7 = XPNT(K+7) - MM
                A1 = X(I1) 
                A2 = X(I2) 
                A3 = X(I3) 
                A4 = X(I4) 
                A5 = X(I5) 
                A6 = X(I6) 
                A7 = X(I7) 
                A9  = X(I1+1) 
                A10 = X(I2+1)
                A11 = X(I3+1)
                A12 = X(I4+1)
                A13 = X(I5+1)
                A14 = X(I6+1)
                A15 = X(I7+1)
C
                Y(IYBEG1+1) =  Y(IYBEG1+1) -
     &              A1*A9 - A2*A10 - A3*A11 - A4*A12 - A5*A13 - 
     &              A6*A14 - A7*A15
C
                Y(IYBEG2+1) =  Y(IYBEG2+1) -
     &              A9*A9 - A10*A10 - A11*A11 - A12*A12 - A13*A13 - 
     &              A14*A14 - A15*A15
C
                DO  600  I = 2, MM-1
                    Y1 = Y(IYBEG1+I)
                    B1 = X(I1+I)
                    Y1 =  Y1 - B1 * A1
                    Y2 = Y(IYBEG2+I)
                    B2 = X(I2+I)
                    Y2 =  Y2 - B1 * A9
                    Y1 =  Y1 - B2 * A2
                    B3 = X(I3+I)
                    Y2 =  Y2 - B2 * A10
                    Y1 =  Y1 - B3 * A3
                    B4 = X(I4+I)
                    Y2 =  Y2 - B3 * A11
                    Y1 =  Y1 - B4 * A4
                    B5 = X(I5+I)
                    Y2 =  Y2 - B4 * A12
                    Y1 =  Y1 - B5 * A5
                    B6 = X(I6+I)
                    Y2 =  Y2 - B5 * A13
                    Y1 =  Y1 - B6 * A6
                    B7 = X(I7+I)
                    Y2 =  Y2 - B6 * A14
                    Y1 = Y1 - B7 * A7
                    Y(IYBEG1+I) = Y1
                    Y2 = Y2 - B7 * A15
                    Y(IYBEG2+I) = Y2               
  600           CONTINUE
C
                GO TO 2000
C
  700       CONTINUE
C
C               ---------------------------------
C               SIX COLUMNS UPDATING TWO COLUMNS.
C               ---------------------------------
C
                I1 = XPNT(K+1) - MM
                I2 = XPNT(K+2) - MM
                I3 = XPNT(K+3) - MM
                I4 = XPNT(K+4) - MM
                I5 = XPNT(K+5) - MM
                I6 = XPNT(K+6) - MM
                A1 = X(I1)
                A2 = X(I2)
                A3 = X(I3)
                A4 = X(I4)
                A5 = X(I5)
                A6 = X(I6)
                A9  = X(I1+1)
                A10 = X(I2+1)
                A11 = X(I3+1)
                A12 = X(I4+1)
                A13 = X(I5+1)
                A14 = X(I6+1)
C
                Y(IYBEG1+1) =  Y(IYBEG1+1) -
     &              A1*A9 - A2*A10 - A3*A11 - A4*A12 - A5*A13 - 
     &              A6*A14
C
                Y(IYBEG2+1) =  Y(IYBEG2+1) -
     &              A9*A9 - A10*A10 - A11*A11 - A12*A12 - A13*A13 - 
     &              A14*A14
C
                DO  800  I = 2, MM-1
                    Y1 = Y(IYBEG1+I)
                    B1 = X(I1+I)
                    Y1 =  Y1 - B1 * A1
                    Y2 = Y(IYBEG2+I)
                    B2 = X(I2+I)
                    Y2 =  Y2 - B1 * A9
                    Y1 =  Y1 - B2 * A2
                    B3 = X(I3+I)
                    Y2 =  Y2 - B2 * A10
                    Y1 =  Y1 - B3 * A3
                    B4 = X(I4+I)
                    Y2 =  Y2 - B3 * A11
                    Y1 =  Y1 - B4 * A4
                    B5 = X(I5+I)
                    Y2 =  Y2 - B4 * A12
                    Y1 =  Y1 - B5 * A5
                    B6 = X(I6+I)
                    Y2 =  Y2 - B5 * A13
                    Y1 =  Y1 - B6 * A6
                    Y(IYBEG1+I) = Y1
                    Y2 =  Y2 - B6 * A14
                    Y(IYBEG2+I) = Y2               
  800           CONTINUE
C
                GO TO 2000
C
  900       CONTINUE
C
C               ----------------------------------
C               FIVE COLUMNS UPDATING TWO COLUMNS.
C               ----------------------------------
C
                I1 = XPNT(K+1) - MM
                I2 = XPNT(K+2) - MM
                I3 = XPNT(K+3) - MM
                I4 = XPNT(K+4) - MM
                I5 = XPNT(K+5) - MM
                A1 = X(I1)
                A2 = X(I2)
                A3 = X(I3)
                A4 = X(I4)
                A5 = X(I5)
                A9  = X(I1+1)
                A10 = X(I2+1)
                A11 = X(I3+1)
                A12 = X(I4+1)
                A13 = X(I5+1)
C
                Y(IYBEG1+1) =  Y(IYBEG1+1) -
     &              A1*A9 - A2*A10 - A3*A11 - A4*A12 - A5*A13
C
                Y(IYBEG2+1) =  Y(IYBEG2+1) -
     &              A9*A9 - A10*A10 - A11*A11 - A12*A12 - A13*A13
C
                DO  1000  I = 2, MM-1
                    Y1 = Y(IYBEG1+I)
                    B1 = X(I1+I)
                    Y1 =  Y1 - B1 * A1
                    Y2 = Y(IYBEG2+I)
                    B2 = X(I2+I)
                    Y2 =  Y2 - B1 * A9
                    Y1 =  Y1 - B2 * A2
                    B3 = X(I3+I)
                    Y2 =  Y2 - B2 * A10
                    Y1 =  Y1 - B3 * A3
                    B4 = X(I4+I)
                    Y2 =  Y2 - B3 * A11
                    Y1 =  Y1 - B4 * A4
                    B5 = X(I5+I)
                    Y2 =  Y2 - B4 * A12
                    Y1 =  Y1 - B5 * A5
                    Y(IYBEG1+I) = Y1
                    Y2 =  Y2 - B5 * A13
                    Y(IYBEG2+I) = Y2               
 1000           CONTINUE
C
                GO TO 2000
C
 1100       CONTINUE
C
C               ----------------------------------
C               FOUR COLUMNS UPDATING TWO COLUMNS.
C               ----------------------------------
C
                I1 = XPNT(K+1) - MM
                I2 = XPNT(K+2) - MM
                I3 = XPNT(K+3) - MM
                I4 = XPNT(K+4) - MM
                A1 = X(I1)
                A2 = X(I2)
                A3 = X(I3)
                A4 = X(I4)
                A9  = X(I1+1)
                A10 = X(I2+1)
                A11 = X(I3+1)
                A12 = X(I4+1)
C
                Y(IYBEG1+1) =  Y(IYBEG1+1) -
     &              A1*A9 - A2*A10 - A3*A11 - A4*A12
C
                Y(IYBEG2+1) =  Y(IYBEG2+1) -
     &              A9*A9 - A10*A10 - A11*A11 - A12*A12
C
                DO  1200  I = 2, MM-1
                    Y1 = Y(IYBEG1+I)
                    B1 = X(I1+I)
                    Y1 =  Y1 - B1 * A1
                    Y2 = Y(IYBEG2+I)
                    B2 = X(I2+I)
                    Y2 =  Y2 - B1 * A9
                    Y1 =  Y1 - B2 * A2
                    B3 = X(I3+I)
                    Y2 =  Y2 - B2 * A10
                    Y1 =  Y1 - B3 * A3
                    B4 = X(I4+I)
                    Y2 =  Y2 - B3 * A11
                    Y1 =  Y1 - B4 * A4
                    Y(IYBEG1+I) = Y1
                    Y2 =  Y2 - B4 * A12
                    Y(IYBEG2+I) = Y2               
 1200           CONTINUE
C
                GO TO 2000
C
 1300       CONTINUE
C
C               -----------------------------------
C               THREE COLUMNS UPDATING TWO COLUMNS.
C               -----------------------------------
C
                I1 = XPNT(K+1) - MM
                I2 = XPNT(K+2) - MM
                I3 = XPNT(K+3) - MM
                A1 = X(I1)
                A2 = X(I2)
                A3 = X(I3)
                A9  = X(I1+1)
                A10 = X(I2+1)
                A11 = X(I3+1)
C
                Y(IYBEG1+1) =  Y(IYBEG1+1) -
     &              A1*A9 - A2*A10 - A3*A11
C
                Y(IYBEG2+1) =  Y(IYBEG2+1) -
     &              A9*A9 - A10*A10 - A11*A11
C
                DO  1400  I = 2, MM-1
                    Y1 = Y(IYBEG1+I)
                    B1 = X(I1+I)
                    Y1 =  Y1 - B1 * A1
                    Y2 = Y(IYBEG2+I)
                    B2 = X(I2+I)
                    Y2 =  Y2 - B1 * A9
                    Y1 =  Y1 - B2 * A2
                    B3 = X(I3+I)
                    Y2 =  Y2 - B2 * A10
                    Y1 =  Y1 - B3 * A3
                    Y(IYBEG1+I) = Y1
                    Y2 =  Y2 - B3 * A11
                    Y(IYBEG2+I) = Y2               
 1400           CONTINUE
C
                GO TO 2000
C
 1500       CONTINUE
C
C               ---------------------------------
C               TWO COLUMNS UPDATING TWO COLUMNS.
C               ---------------------------------
C
                I1 = XPNT(K+1) - MM
                I2 = XPNT(K+2) - MM
                A1 = X(I1)
                A2 = X(I2)
                A9  = X(I1+1)
                A10 = X(I2+1)
C
                Y(IYBEG1+1) =  Y(IYBEG1+1) -
     &              A1*A9 - A2*A10
C
                Y(IYBEG2+1) =  Y(IYBEG2+1) -
     &              A9*A9 - A10*A10
C
                DO  1600  I = 2, MM-1
                    Y1 = Y(IYBEG1+I)
                    B1 = X(I1+I)
                    Y1 =  Y1 - B1 * A1
                    Y2 = Y(IYBEG2+I)
                    B2 = X(I2+I)
                    Y2 =  Y2 - B1 * A9
                    Y1 =  Y1 - B2 * A2
                    Y(IYBEG1+I) = Y1
                    Y2 =  Y2 - B2 * A10
                    Y(IYBEG2+I) = Y2               
 1600           CONTINUE
C
                GO TO 2000
C
 1700       CONTINUE
C
C               --------------------------------
C               ONE COLUMN UPDATING TWO COLUMNS.
C               --------------------------------
C
                I1 = XPNT(K+1) - MM
                A1 = X(I1)
                A9  = X(I1+1)
C
                Y(IYBEG1+1) =  Y(IYBEG1+1) -
     &              A1*A9
C
                Y(IYBEG2+1) =  Y(IYBEG2+1) -
     &              A9*A9
C
                DO  1800  I = 2, MM-1
                    Y1 = Y(IYBEG1+I)
                    B1 = X(I1+I)
                    Y1 =  Y1 - B1 * A1
                    Y2 = Y(IYBEG2+I)
                    Y(IYBEG1+I) = Y1
                    Y2 =  Y2 - B1 * A9
                    Y(IYBEG2+I) = Y2               
 1800           CONTINUE
C
                GO TO 2000
C
C           -----------------------------------------------
C           PREPARE FOR NEXT PAIR OF COLUMNS TO BE UPDATED.
C           -----------------------------------------------
C
 2000       CONTINUE
            MM = MM - 2
            IYBEG = IYBEG2 + LENY + 1
            LENY = LENY - 2
C
 3000   CONTINUE
C
C       -----------------------------------------------------
C       BOUNDARY CODE FOR J LOOP:  EXECUTED WHENVER Q IS ODD.
C       -----------------------------------------------------
C
        IF  ( J .EQ. QQ )  THEN
            CALL  SMXPY8  ( MM, N, Y(IYBEG), XPNT, X )
        ENDIF
C
        RETURN
        END
C***********************************************************************
C***********************************************************************
C
C   Version:        0.4
C   Last modified:  December 27, 1994
C   Authors:        Esmond G. Ng and Barry W. Peyton
C
C   Mathematical Sciences Section, Oak Ridge National Laboratory
C
C***********************************************************************
C***********************************************************************
C*************     MMPYI  .... MATRIX-MATRIX MULTIPLY     **************
C***********************************************************************
C***********************************************************************
C
C   PURPOSE -
C       THIS ROUTINE PERFORMS A MATRIX-MATRIX MULTIPLY, Y = Y + XA,
C       ASSUMING DATA STRUCTURES USED IN SOME OF OUR SPARSE CHOLESKY
C       CODES.
C
C       MATRIX X HAS ONLY 1 COLUMN.
C
C   INPUT PARAMETERS -
C       M               -   NUMBER OF ROWS IN X AND IN Y.
C       Q               -   NUMBER OF COLUMNS IN A AND Y.
C       XPNT(*)         -   XPNT(J+1) POINTS ONE LOCATION BEYOND THE
C                           END OF THE J-TH COLUMN OF X.  XPNT IS ALSO
C                           USED TO ACCESS THE ROWS OF A.
C       X(*)            -   CONTAINS THE COLUMNS OF X AND THE ROWS OF A.
C       IY(*)           -   IY(COL) POINTS TO THE BEGINNING OF COLUMN
C       RELIND(*)       -   RELATIVE INDICES.
C
C   UPDATED PARAMETERS -
C       Y(*)            -   ON OUTPUT, Y = Y + AX.
C
C***********************************************************************
C
      SUBROUTINE  MMPYI  (  M     , Q     , XPNT  , X     , IY    ,
     &                      Y     , RELIND                          )
C
C***********************************************************************
C
C       -----------
C       PARAMETERS.
C       -----------
C
        INTEGER             M     , Q
        INTEGER             IY(*)         , RELIND(*)     ,
     &                      XPNT(*)
        DOUBLE PRECISION    X(*)      , Y(*)
C
C       ----------------
C       LOCAL VARIABLES.
C       ----------------
C
        INTEGER             COL   , I     , ISUB  , K     , YLAST
        DOUBLE PRECISION    A
C
C***********************************************************************
C
        DO  200  K = 1, Q
            COL = XPNT(K)
            YLAST = IY(COL+1) - 1
            A = - X(K)
CDIR$   IVDEP
            DO  100  I = K, M
                ISUB = XPNT(I)
                ISUB = YLAST - RELIND(ISUB)
                Y(ISUB) = Y(ISUB) + A*X(I)
  100       CONTINUE
  200   CONTINUE
        RETURN
C
      END
C***********************************************************************
C***********************************************************************
C
C   Version:        0.4
C   Last modified:  December 27, 1994
C   Authors:        Esmond G. Ng and Barry W. Peyton
C
C   Mathematical Sciences Section, Oak Ridge National Laboratory
C
C***********************************************************************
C***********************************************************************
C****     ORDMMD ..... MULTIPLE MINIMUM EXTERNAL DEGREE     ************
C***********************************************************************
C***********************************************************************
C
C     PURPOSE - THIS ROUTINE CALLS LIU'S MULTIPLE MINIMUM DEGREE
C               ROUTINE.
C
C     INPUT PARAMETERS -
C        NEQNS  - NUMBER OF EQUATIONS.
C        (XADJ,ADJNCY) - THE ADJACENCY STRUCTURE.
C        IWSIZ  - SIZE OF INTEGER WORKING STORAGE.
C
C     OUTPUT PARAMETERS -
C        PERM   - THE MINIMUM DEGREE ORDERING.
C        INVP   - THE INVERSE OF PERM.
C        NOFSUB - AN UPPER BOUND ON THE NUMBER OF NONZERO
C                 SUBSCRIPTS FOR THE COMPRESSED STORAGE SCHEME.
C        IFLAG  - ERROR FLAG.
C                   0: SUCCESSFUL ORDERING
C                  -1: INSUFFICIENT WORKING STORAGE
C                      [IWORK(*)].
C
C     WORKING PARAMETERS -
C        IWORK  - INTEGER WORKSPACE OF LENGTH 4*NEQNS.
C
C***********************************************************************
C
      SUBROUTINE ORDMMD  (  NEQNS , XADJ  , ADJNCY, INVP  , PERM  ,
     1                      IWSIZ , IWORK , NOFSUB, IFLAG           )
C
C***********************************************************************
C
         INTEGER    ADJNCY(*), INVP(*)  , IWORK(*) , PERM(*)
         INTEGER    XADJ(*)
         INTEGER    DELTA , IFLAG , IWSIZ , MAXINT, NEQNS , 
     &              NOFSUB
C
C*********************************************************************
C
        IFLAG = 0
        IF  ( IWSIZ .LT. 4*NEQNS )  THEN
            IFLAG = -1
            RETURN
        ENDIF
C
C       DELTA  - TOLERANCE VALUE FOR MULTIPLE ELIMINATION.
C       MAXINT - MAXIMUM MACHINE REPRESENTABLE (SHORT) INTEGER
C                (ANY SMALLER ESTIMATE WILL DO) FOR MARKING
C                NODES.
C
        DELTA  = 0
        MAXINT = 32767
        CALL GENMMD  (  NEQNS , XADJ  , ADJNCY, INVP  , PERM  ,
     1                  DELTA , 
     1                  IWORK(1)              ,
     1                  IWORK(NEQNS+1)        ,
     1                  IWORK(2*NEQNS+1)      ,
     1                  IWORK(3*NEQNS+1)      ,
     1                  MAXINT, NOFSUB          )
         RETURN
C
      END
C***********************************************************************
C***********************************************************************
C
C   Version:        0.3
C   Last modified:  December 27, 1994
C   Authors:        Esmond G. Ng and Barry W. Peyton
C
C   Mathematical Sciences Section, Oak Ridge National Laboratoy
C
C***********************************************************************
C***********************************************************************
C******     PCHOL .... DENSE PARTIAL CHOLESKY             **************
C***********************************************************************
C***********************************************************************
C
C     PURPOSE - THIS ROUTINE PERFORMS CHOLESKY
C               FACTORIZATION ON THE COLUMNS OF A SUPERNODE
C               THAT HAVE RECEIVED ALL UPDATES FROM COLUMNS
C               EXTERNAL TO THE SUPERNODE.
C
C     INPUT PARAMETERS -
C        M      - NUMBER OF ROWS (LENGTH OF THE FIRST COLUMN).
C        N      - NUMBER OF COLUMNS IN THE SUPERNODE.
C        XPNT   - XPNT(J+1) POINTS ONE LOCATION BEYOND THE END
C                 OF THE J-TH COLUMN OF THE SUPERNODE.
C        X(*)   - CONTAINS THE COLUMNS OF OF THE SUPERNODE TO
C                 BE FACTORED.
C     
C     EXTERNAL ROUTINE:
C        SMXPY8 -  MATRIX-VECTOR MULTIPLY WITH 8 LOOP UNROLLING.
C
C     OUTPUT PARAMETERS -
C        X(*)   - ON OUTPUT, CONTAINS THE FACTORED COLUMNS OF
C                 THE SUPERNODE.
C
C***********************************************************************
C
      SUBROUTINE  PCHOL  ( M, N, XPNT, X, MXDIAG, NTINY )
C
C***********************************************************************
C
C     -----------
C     PARAMETERS.
C     -----------
C
      EXTERNAL            SMXPY8
C
      INTEGER             M, N
C
      INTEGER             XPNT(*)
C
CxPCxPCxPCxPCxPCxPCxPCxPCxPCxPCxPCxPCxPCxPCxPCxPCxPCxPCxPCxPCxPCxPCxPC
      DOUBLE PRECISION    X(*), MXDIAG
      INTEGER             NTINY
C
C     ----------------
C     LOCAL VARIABLES.
C     ----------------
C
      INTEGER             JPNT  , JCOL  , MM
C
      DOUBLE PRECISION    DIAG
C
C***********************************************************************
C
C       ------------------------------------------
C       FOR EVERY COLUMN JCOL IN THE SUPERNODE ...
C       ------------------------------------------
        MM     = M
        JPNT = XPNT(1)
        DO  100  JCOL = 1, N
C
C           ----------------------------------
C           UPDATE JCOL WITH PREVIOUS COLUMNS.
C           ----------------------------------
            IF  ( JCOL .GT. 1 )  THEN
                CALL SMXPY8 ( MM, JCOL-1, X(JPNT), XPNT, X )
            ENDIF
C
C           ---------------------------
C           COMPUTE THE DIAGONAL ENTRY.
C           ---------------------------
            DIAG = X(JPNT)
CxPCxPCxPCxPCxPCxPCxPCxPCxPCxPCxPCxPCxPCxPCxPCxPCxPCxPCxPCxPCxPCxPCxPC
            IF (DIAG .LE. 1.0D-30*MXDIAG) THEN
               DIAG = 1.0D+128
               NTINY = NTINY+1
            ENDIF
CxPCxPCxPCxPCxPCxPCxPCxPCxPCxPCxPCxPCxPCxPCxPCxPCxPCxPCxPCxPCxPCxPCxPC

            DIAG = SQRT ( DIAG )
            X(JPNT) = DIAG
            DIAG = 1.0D+00 / DIAG
C
C           ----------------------------------------------------
C           SCALE COLUMN JCOL WITH RECIPROCAL OF DIAGONAL ENTRY.
C           ----------------------------------------------------
            MM = MM - 1
            JPNT = JPNT + 1
            CALL DSCAL1 ( MM, DIAG, X(JPNT) )
            JPNT = JPNT + MM
C
  100   CONTINUE
C
      RETURN
      END
C***********************************************************************
C***********************************************************************
C
C   Version:        0.4
C   Last modified:  January 12, 1995
C   Authors:        Esmond G. Ng and Barry W. Peyton
C
C   Mathematical Sciences Section, Oak Ridge National Laboratory
C
C***********************************************************************
C***********************************************************************
C**************    SFINIT  ..... SET UP FOR SYMB. FACT.     ************
C***********************************************************************
C***********************************************************************
C
C   PURPOSE:
C       THIS SUBROUTINE COMPUTES THE STORAGE REQUIREMENTS AND SETS UP 
C       PRELIMINARY DATA STRUCTURES FOR THE SYMBOLIC FACTORIZATION.
C
C   NOTE:
C       THIS VERSION PRODUCES THE MAXIMAL SUPERNODE PARTITION (I.E.,
C       THE ONE WITH THE FEWEST POSSIBLE SUPERNODES).
C
C   INPUT PARAMETERS:
C       NEQNS       -   NUMBER OF EQUATIONS.
C       NNZA        -   LENGTH OF ADJACENCY STRUCTURE.
C       XADJ(*)     -   ARRAY OF LENGTH NEQNS+1, CONTAINING POINTERS
C                       TO THE ADJACENCY STRUCTURE.
C       ADJNCY(*)   -   ARRAY OF LENGTH XADJ(NEQNS+1)-1, CONTAINING
C                       THE ADJACENCY STRUCTURE.
C       PERM(*)     -   ARRAY OF LENGTH NEQNS, CONTAINING THE
C                       POSTORDERING.
C       INVP(*)     -   ARRAY OF LENGTH NEQNS, CONTAINING THE
C                       INVERSE OF THE POSTORDERING.
C       IWSIZ       -   SIZE OF INTEGER WORKING STORAGE.
C
C   OUTPUT PARAMETERS:
C       COLCNT(*)   -   ARRAY OF LENGTH NEQNS, CONTAINING THE NUMBER
C                       OF NONZEROS IN EACH COLUMN OF THE FACTOR,
C                       INCLUDING THE DIAGONAL ENTRY.
C       NNZL        -   NUMBER OF NONZEROS IN THE FACTOR, INCLUDING
C                       THE DIAGONAL ENTRIES.
C       NSUB        -   NUMBER OF SUBSCRIPTS.
C       NSUPER      -   NUMBER OF SUPERNODES (<= NEQNS).
C       SNODE(*)    -   ARRAY OF LENGTH NEQNS FOR RECORDING
C                       SUPERNODE MEMBERSHIP.
C       XSUPER(*)   -   ARRAY OF LENGTH NEQNS+1, CONTAINING THE
C                       SUPERNODE PARTITIONING.
C       IFLAG(*)    -   ERROR FLAG.
C                          0: SUCCESSFUL SF INITIALIZATION.
C                         -1: INSUFFICENT WORKING STORAGE
C                             [IWORK(*)].
C
C   WORK PARAMETERS:
C       IWORK(*)    -   INTEGER WORK ARRAY OF LENGTH 7*NEQNS+3.
C
C   FIRST CREATED ON    NOVEMEBER 14, 1994.
C   LAST UPDATED ON     January 12, 1995.
C
C***********************************************************************
C
      SUBROUTINE  SFINIT (  NEQNS , NNZA  , XADJ  , ADJNCY, PERM  ,
     &                      INVP  , COLCNT, NNZL  , NSUB  , NSUPER,
     &                      SNODE , XSUPER, IWSIZ , IWORK , IFLAG   )
C
C       ----------- 
C       PARAMETERS.  
C       -----------
        INTEGER             IFLAG , IWSIZ , NNZA  , NEQNS , NNZL  , 
     &                      NSUB  , NSUPER
        INTEGER             ADJNCY(NNZA)    , COLCNT(NEQNS)   ,
     &                      INVP(NEQNS)     , IWORK(7*NEQNS+3),
     &                      PERM(NEQNS)     , SNODE(NEQNS)    , 
     &                      XADJ(NEQNS+1)   , XSUPER(NEQNS+1)
C
C***********************************************************************
C
C       --------------------------------------------------------
C       RETURN IF THERE IS INSUFFICIENT INTEGER WORKING STORAGE.
C       --------------------------------------------------------
        IFLAG = 0
        IF  ( IWSIZ .LT. 7*NEQNS+3 )  THEN
            IFLAG = -1
            RETURN
        ENDIF
C
C       ------------------------------------------
C       COMPUTE ELIMINATION TREE AND POSTORDERING.
C       ------------------------------------------
        CALL  ETORDR (  NEQNS , XADJ  , ADJNCY, PERM  , INVP  ,
     &                  IWORK(1)              ,
     &                  IWORK(NEQNS+1)        ,
     &                  IWORK(2*NEQNS+1)      ,
     &                  IWORK(3*NEQNS+1)        ) 
C
C       ---------------------------------------------
C       COMPUTE ROW AND COLUMN FACTOR NONZERO COUNTS.
C       ---------------------------------------------
        CALL  FCNTHN (  NEQNS , NNZA  , XADJ  , ADJNCY, PERM  , 
     &                  INVP  , IWORK(1)      , SNODE , COLCNT, 
     &                  NNZL  ,
     &                  IWORK(NEQNS+1)        ,
     &                  IWORK(2*NEQNS+1)      ,
     &                  XSUPER                ,
     &                  IWORK(3*NEQNS+1)      ,
     &                  IWORK(4*NEQNS+2)      ,
     &                  IWORK(5*NEQNS+3)      ,
     &                  IWORK(6*NEQNS+4)        )
C
C       ---------------------------------------------------------
C       REARRANGE CHILDREN SO THAT THE LAST CHILD HAS THE MAXIMUM 
C       NUMBER OF NONZEROS IN ITS COLUMN OF L.
C       ---------------------------------------------------------
        CALL  CHORDR (  NEQNS , PERM  , INVP  ,
     &                  COLCNT, 
     &                  IWORK(1)              ,
     &                  IWORK(NEQNS+1)        ,
     &                  IWORK(2*NEQNS+1)      ,
     &                  IWORK(3*NEQNS+1)        )
C
C       ----------------
C       FIND SUPERNODES.
C       ----------------
        CALL  FSUP1  (  NEQNS , IWORK(1)      , COLCNT, NSUB  , 
     &                  NSUPER, SNODE                           )
        CALL  FSUP2  (  NEQNS , NSUPER,  SNODE,     XSUPER      )
C
        RETURN
      END
C***********************************************************************
C***********************************************************************
C
C   Version:        0.4
C   Last modified:  December 27, 1994
C   Authors:        Esmond G. Ng and Barry W. Peyton
C
C   Mathematical Sciences Section, Oak Ridge National Laboratory
C
C***********************************************************************
C***********************************************************************
C******     SMXPY8 .... MATRIX-VECTOR MULTIPLY            **************
C***********************************************************************
C***********************************************************************
C
C     PURPOSE - THIS ROUTINE PERFORMS A MATRIX-VECTOR MULTIPLY,
C               Y = Y + AX, ASSUMING DATA STRUCTURES USED IN
C               RECENTLY DEVELOPED SPARSE CHOLESKY CODES.  THE 
C               '8' SIGNIFIES LEVEL 8 LOOP UNROLLING.
C
C     INPUT PARAMETERS -
C        M      - NUMBER OF ROWS.
C        N      - NUMBER OF COLUMNS.
C        Y      - M-VECTOR TO WHICH AX WILL BE ADDED.
C        APNT   - INDEX VECTOR FOR A.  APNT(I) POINTS TO THE
C                 FIRST NONZERO IN COLUMN I OF A.
C        Y      - ON OUTPUT, CONTAINS Y = Y + AX.
C
C***********************************************************************
C
      SUBROUTINE  SMXPY8 ( M, N, Y, APNT, A )
C
C***********************************************************************
C
C     -----------
C     PARAMETERS.
C     -----------
C
      INTEGER             M, N, LEVEL
C
      INTEGER             APNT(*)
C
      DOUBLE PRECISION    Y(*), A(*)
C
      PARAMETER           ( LEVEL = 8 )
C
C     ----------------
C     LOCAL VARIABLES.
C     ----------------
C
      INTEGER             I, I1, I2, I3, I4, I5, I6, I7, I8,
     &                    J, REMAIN
C
      DOUBLE PRECISION    A1, A2, A3, A4, A5, A6, A7, A8
C
C***********************************************************************
C
      REMAIN = MOD ( N, LEVEL )
C
C      GO TO ( 2000, 100, 200, 300,
C     &         400, 500, 600, 700  ), REMAIN+1
      if (REMAIN .eq. 0) go to 2000
C      if (REMAIN .eq. 1) go to 100
      if (REMAIN .eq. 2) go to 200
      if (REMAIN .eq. 3) go to 300
      if (REMAIN .eq. 4) go to 400
      if (REMAIN .eq. 5) go to 500
      if (REMAIN .eq. 6) go to 600
      if (REMAIN .eq. 7) go to 700
C
C  100 CONTINUE
      I1 = APNT(1+1) - M
      A1 = - A(I1)
      DO  150  I = 1, M
          Y(I) = Y(I) + A1*A(I1)
          I1 = I1 + 1
  150 CONTINUE
      GO TO 2000
C
  200 CONTINUE
      I1 = APNT(1+1) - M
      I2 = APNT(1+2) - M
      A1 = - A(I1)
      A2 = - A(I2)
      DO  250  I = 1, M
          Y(I) = ( (Y(I))
     &           + A1*A(I1)) + A2*A(I2)
          I1 = I1 + 1
          I2 = I2 + 1
  250 CONTINUE
      GO TO 2000
C
  300 CONTINUE
      I1 = APNT(1+1) - M
      I2 = APNT(1+2) - M
      I3 = APNT(1+3) - M
      A1 = - A(I1)
      A2 = - A(I2)
      A3 = - A(I3)
      DO  350  I = 1, M
          Y(I) = (( (Y(I))
     &           + A1*A(I1)) + A2*A(I2))
     &           + A3*A(I3)
          I1 = I1 + 1
          I2 = I2 + 1
          I3 = I3 + 1
  350 CONTINUE
      GO TO 2000
C
  400 CONTINUE
      I1 = APNT(1+1) - M
      I2 = APNT(1+2) - M
      I3 = APNT(1+3) - M
      I4 = APNT(1+4) - M
      A1 = - A(I1)
      A2 = - A(I2)
      A3 = - A(I3)
      A4 = - A(I4)
      DO  450  I = 1, M
          Y(I) = ((( (Y(I))
     &           + A1*A(I1)) + A2*A(I2))
     &           + A3*A(I3)) + A4*A(I4)
          I1 = I1 + 1
          I2 = I2 + 1
          I3 = I3 + 1
          I4 = I4 + 1
  450 CONTINUE
      GO TO 2000
C
  500 CONTINUE
      I1 = APNT(1+1) - M
      I2 = APNT(1+2) - M
      I3 = APNT(1+3) - M
      I4 = APNT(1+4) - M
      I5 = APNT(1+5) - M
      A1 = - A(I1)
      A2 = - A(I2)
      A3 = - A(I3)
      A4 = - A(I4)
      A5 = - A(I5)
      DO  550  I = 1, M
          Y(I) = (((( (Y(I))
     &           + A1*A(I1)) + A2*A(I2))
     &           + A3*A(I3)) + A4*A(I4))
     &           + A5*A(I5)
          I1 = I1 + 1
          I2 = I2 + 1
          I3 = I3 + 1
          I4 = I4 + 1
          I5 = I5 + 1
  550 CONTINUE
      GO TO 2000
C
  600 CONTINUE
      I1 = APNT(1+1) - M
      I2 = APNT(1+2) - M
      I3 = APNT(1+3) - M
      I4 = APNT(1+4) - M
      I5 = APNT(1+5) - M
      I6 = APNT(1+6) - M
      A1 = - A(I1)
      A2 = - A(I2)
      A3 = - A(I3)
      A4 = - A(I4)
      A5 = - A(I5)
      A6 = - A(I6)
      DO  650  I = 1, M
          Y(I) = ((((( (Y(I))
     &           + A1*A(I1)) + A2*A(I2))
     &           + A3*A(I3)) + A4*A(I4))
     &           + A5*A(I5)) + A6*A(I6)
          I1 = I1 + 1
          I2 = I2 + 1
          I3 = I3 + 1
          I4 = I4 + 1
          I5 = I5 + 1
          I6 = I6 + 1
  650 CONTINUE
      GO TO 2000
C
  700 CONTINUE
      I1 = APNT(1+1) - M
      I2 = APNT(1+2) - M
      I3 = APNT(1+3) - M
      I4 = APNT(1+4) - M
      I5 = APNT(1+5) - M
      I6 = APNT(1+6) - M
      I7 = APNT(1+7) - M
      A1 = - A(I1)
      A2 = - A(I2)
      A3 = - A(I3)
      A4 = - A(I4)
      A5 = - A(I5)
      A6 = - A(I6)
      A7 = - A(I7)
      DO  750  I = 1, M
          Y(I) = (((((( (Y(I))
     &           + A1*A(I1)) + A2*A(I2))
     &           + A3*A(I3)) + A4*A(I4))
     &           + A5*A(I5)) + A6*A(I6))
     &           + A7*A(I7)
          I1 = I1 + 1
          I2 = I2 + 1
          I3 = I3 + 1
          I4 = I4 + 1
          I5 = I5 + 1
          I6 = I6 + 1
          I7 = I7 + 1
  750 CONTINUE
      GO TO 2000
C
 2000 CONTINUE
      DO  4000  J = REMAIN+1, N, LEVEL
          I1 = APNT(J+1) - M
          I2 = APNT(J+2) - M
          I3 = APNT(J+3) - M
          I4 = APNT(J+4) - M
          I5 = APNT(J+5) - M
          I6 = APNT(J+6) - M
          I7 = APNT(J+7) - M
          I8 = APNT(J+8) - M
          A1 = - A(I1)
          A2 = - A(I2)
          A3 = - A(I3)
          A4 = - A(I4)
          A5 = - A(I5)
          A6 = - A(I6)
          A7 = - A(I7)
          A8 = - A(I8)
          DO  3000  I = 1, M
              Y(I) = ((((((( (Y(I))
     &               + A1*A(I1)) + A2*A(I2))
     &               + A3*A(I3)) + A4*A(I4))
     &               + A5*A(I5)) + A6*A(I6))
     &               + A7*A(I7)) + A8*A(I8)
              I1 = I1 + 1
              I2 = I2 + 1
              I3 = I3 + 1
              I4 = I4 + 1
              I5 = I5 + 1
              I6 = I6 + 1
              I7 = I7 + 1
              I8 = I8 + 1
 3000     CONTINUE
 4000 CONTINUE
C
      RETURN
      END
C***********************************************************************
C***********************************************************************
C
C   Version:        0.4
C   Last modified:  February 13, 1995
C   Authors:        Esmond G. Ng and Barry W. Peyton
C
C   Mathematical Sciences Section, Oak Ridge National Laboratory
C
C***********************************************************************
C***********************************************************************
C*************     SYMFC2 ..... SYMBOLIC FACTORIZATION    **************
C***********************************************************************
C***********************************************************************
C
C   PURPOSE: 
C       THIS ROUTINE PERFORMS SUPERNODAL SYMBOLIC FACTORIZATION ON A 
C       REORDERED LINEAR SYSTEM.  IT ASSUMES ACCESS TO THE COLUMNS 
C       COUNTS, SUPERNODE PARTITION, AND SUPERNODAL ELIMINATION TREE
C       ASSOCIATED WITH THE FACTOR MATRIX L.
C
C   INPUT PARAMETERS:
C       (I) NEQNS       -   NUMBER OF EQUATIONS
C       (I) ADJLEN      -   LENGTH OF THE ADJACENCY LIST.
C       (I) XADJ(*)     -   ARRAY OF LENGTH NEQNS+1 CONTAINING POINTERS
C                           TO THE ADJACENCY STRUCTURE.
C       (I) ADJNCY(*)   -   ARRAY OF LENGTH XADJ(NEQNS+1)-1 CONTAINING
C                           THE ADJACENCY STRUCTURE.
C       (I) PERM(*)     -   ARRAY OF LENGTH NEQNS CONTAINING THE
C                           POSTORDERING.
C       (I) INVP(*)     -   ARRAY OF LENGTH NEQNS CONTAINING THE
C                           INVERSE OF THE POSTORDERING.
C       (I) COLCNT(*)   -   ARRAY OF LENGTH NEQNS, CONTAINING THE NUMBER
C                           OF NONZEROS IN EACH COLUMN OF THE FACTOR,
C                           INCLUDING THE DIAGONAL ENTRY.
C       (I) NSUPER      -   NUMBER OF SUPERNODES.
C       (I) XSUPER(*)   -   ARRAY OF LENGTH NSUPER+1, CONTAINING THE
C                           FIRST COLUMN OF EACH SUPERNODE.
C       (I) SNODE(*)    -   ARRAY OF LENGTH NEQNS FOR RECORDING
C                           SUPERNODE MEMBERSHIP.
C       (I) NOFSUB      -   NUMBER OF SUBSCRIPTS TO BE STORED IN
C                           LINDX(*).
C
C   OUTPUT PARAMETERS:
C       (I) XLINDX      -   ARRAY OF LENGTH NEQNS+1, CONTAINING POINTERS 
C                           INTO THE SUBSCRIPT VECTOR.
C       (I) LINDX       -   ARRAY OF LENGTH MAXSUB, CONTAINING THE
C                           COMPRESSED SUBSCRIPTS.
C       (I) XLNZ        -   COLUMN POINTERS FOR L.
C       (I) FLAG        -   ERROR FLAG:
C                               0 - NO ERROR.
C                               1 - INCONSISTANCY IN THE INPUT.
C       
C   WORKING PARAMETERS:
C       (I) MRGLNK      -   ARRAY OF LENGTH NSUPER, CONTAINING THE 
C                           CHILDREN OF EACH SUPERNODE AS A LINKED LIST.
C       (I) RCHLNK      -   ARRAY OF LENGTH NEQNS+1, CONTAINING THE 
C                           CURRENT LINKED LIST OF MERGED INDICES (THE 
C                           "REACH" SET).
C       (I) MARKER      -   ARRAY OF LENGTH NEQNS USED TO MARK INDICES
C                           AS THEY ARE INTRODUCED INTO EACH SUPERNODE'S
C                           INDEX SET.
C
C***********************************************************************
C
      SUBROUTINE  SYMFC2 (  NEQNS , ADJLEN, XADJ  , ADJNCY, PERM  , 
     &                      INVP  , COLCNT, NSUPER, XSUPER, SNODE ,
     &                      NOFSUB, XLINDX, LINDX , XLNZ  , MRGLNK,
     &                      RCHLNK, MARKER, FLAG    )
C
C***********************************************************************
C
C       -----------
C       PARAMETERS.
C       -----------
        INTEGER             ADJLEN, FLAG  , NEQNS , NOFSUB, NSUPER
        INTEGER             ADJNCY(ADJLEN), COLCNT(NEQNS) ,
     &                      INVP(NEQNS)   , MARKER(NEQNS) ,
     &                      MRGLNK(NSUPER), LINDX(NOFSUB) , 
     &                      PERM(NEQNS)   , RCHLNK(0:NEQNS), 
     &                      SNODE(NEQNS)  , XSUPER(NSUPER+1)
        INTEGER             XADJ(NEQNS+1) , XLINDX(NSUPER+1),
     &                      XLNZ(NEQNS+1)
C
C       ----------------
C       LOCAL VARIABLES.
C       ----------------
        INTEGER             FSTCOL, HEAD  , I     , JNZBEG, JNZEND,
     &                      JPTR  , JSUP  , JWIDTH, KNZ   , KNZBEG,
     &                      KNZEND, KPTR  , KSUP  , LENGTH, LSTCOL,
     &                      NEWI  , NEXTI , NODE  , NZBEG , NZEND ,
     &                      PCOL  , PSUP  , POINT , TAIL  , WIDTH
C
C***********************************************************************
C
        FLAG = 0
        IF  ( NEQNS .LE. 0 )  RETURN
C
C       ---------------------------------------------------
C       INITIALIZATIONS ...
C           NZEND  : POINTS TO THE LAST USED SLOT IN LINDX.
C           TAIL   : END OF LIST INDICATOR 
C                    (IN RCHLNK(*), NOT MRGLNK(*)).
C           MRGLNK : CREATE EMPTY LISTS.
C           MARKER : "UNMARK" THE INDICES.
C       ---------------------------------------------------
        NZEND = 0
        HEAD = 0
        TAIL = NEQNS + 1
        POINT = 1
        DO  50  I = 1, NEQNS
            MARKER(I) = 0
            XLNZ(I) = POINT
            POINT = POINT + COLCNT(I)
   50   CONTINUE
        XLNZ(NEQNS+1) = POINT
        POINT = 1
        DO  100  KSUP = 1, NSUPER
            MRGLNK(KSUP) = 0
            FSTCOL = XSUPER(KSUP)
            XLINDX(KSUP) = POINT
            POINT = POINT + COLCNT(FSTCOL)
  100   CONTINUE
        XLINDX(NSUPER+1) = POINT
C
C       ---------------------------
C       FOR EACH SUPERNODE KSUP ... 
C       ---------------------------
        DO  1000  KSUP = 1, NSUPER
C
C           ---------------------------------------------------------
C           INITIALIZATIONS ...
C               FSTCOL : FIRST COLUMN OF SUPERNODE KSUP.
C               LSTCOL : LAST COLUMN OF SUPERNODE KSUP.
C               KNZ    : WILL COUNT THE NONZEROS OF L IN COLUMN KCOL.
C               RCHLNK : INITIALIZE EMPTY INDEX LIST FOR KCOL.
C           ---------------------------------------------------------
            FSTCOL = XSUPER(KSUP)
            LSTCOL = XSUPER(KSUP+1) - 1
            WIDTH  = LSTCOL - FSTCOL + 1
            LENGTH = COLCNT(FSTCOL)
            KNZ = 0
            RCHLNK(HEAD) = TAIL
            JSUP = MRGLNK(KSUP)
C
C           -------------------------------------------------
C           IF KSUP HAS CHILDREN IN THE SUPERNODAL E-TREE ...
C           -------------------------------------------------
            IF  ( JSUP .GT. 0 )  THEN
C               ---------------------------------------------
C               COPY THE INDICES OF THE FIRST CHILD JSUP INTO 
C               THE LINKED LIST, AND MARK EACH WITH THE VALUE 
C               KSUP.
C               ---------------------------------------------
                JWIDTH = XSUPER(JSUP+1) - XSUPER(JSUP)
                JNZBEG = XLINDX(JSUP) + JWIDTH
                JNZEND = XLINDX(JSUP+1) - 1
                DO  200  JPTR = JNZEND, JNZBEG, -1
                    NEWI = LINDX(JPTR)
                    KNZ = KNZ+1
                    MARKER(NEWI) = KSUP
                    RCHLNK(NEWI) = RCHLNK(HEAD)
                    RCHLNK(HEAD) = NEWI
  200           CONTINUE
C               ------------------------------------------
C               FOR EACH SUBSEQUENT CHILD JSUP OF KSUP ...
C               ------------------------------------------
                JSUP = MRGLNK(JSUP)
  300           CONTINUE
                IF  ( JSUP .NE. 0  .AND.  KNZ .LT. LENGTH )  THEN
C                   ----------------------------------------
C                   MERGE THE INDICES OF JSUP INTO THE LIST,
C                   AND MARK NEW INDICES WITH VALUE KSUP.
C                   ----------------------------------------
                    JWIDTH = XSUPER(JSUP+1) - XSUPER(JSUP)
                    JNZBEG = XLINDX(JSUP) + JWIDTH
                    JNZEND = XLINDX(JSUP+1) - 1
                    NEXTI = HEAD
                    DO  500  JPTR = JNZBEG, JNZEND
                        NEWI = LINDX(JPTR)
  400                   CONTINUE
                            I = NEXTI
                            NEXTI = RCHLNK(I)
                            IF  ( NEWI .GT. NEXTI )  GO TO 400
                        IF  ( NEWI .LT. NEXTI )  THEN
                            KNZ = KNZ+1
                            RCHLNK(I) = NEWI
                            RCHLNK(NEWI) = NEXTI
                            MARKER(NEWI) = KSUP
                            NEXTI = NEWI
                        ENDIF
  500               CONTINUE
                    JSUP = MRGLNK(JSUP)
                    GO TO 300
                ENDIF
            ENDIF
C           ---------------------------------------------------
C           STRUCTURE OF A(*,FSTCOL) HAS NOT BEEN EXAMINED YET.  
C           "SORT" ITS STRUCTURE INTO THE LINKED LIST,
C           INSERTING ONLY THOSE INDICES NOT ALREADY IN THE
C           LIST.
C           ---------------------------------------------------
            IF  ( KNZ .LT. LENGTH )  THEN
                NODE = PERM(FSTCOL)
                KNZBEG = XADJ(NODE)
                KNZEND = XADJ(NODE+1) - 1
                DO  700  KPTR = KNZBEG, KNZEND
                    NEWI = ADJNCY(KPTR)
                    NEWI = INVP(NEWI)
                    IF  ( NEWI .GT. FSTCOL  .AND.
     &                    MARKER(NEWI) .NE. KSUP )  THEN
C                       --------------------------------
C                       POSITION AND INSERT NEWI IN LIST
C                       AND MARK IT WITH KCOL.
C                       --------------------------------
                        NEXTI = HEAD
  600                   CONTINUE
                            I = NEXTI
                            NEXTI = RCHLNK(I)
                            IF  ( NEWI .GT. NEXTI )  GO TO 600
                        KNZ = KNZ + 1
                        RCHLNK(I) = NEWI
                        RCHLNK(NEWI) = NEXTI
                        MARKER(NEWI) = KSUP
                    ENDIF
  700           CONTINUE
            ENDIF
C           ------------------------------------------------------------
C           IF KSUP HAS NO CHILDREN, INSERT FSTCOL INTO THE LINKED LIST.
C           ------------------------------------------------------------
            IF  ( RCHLNK(HEAD) .NE. FSTCOL )  THEN
                RCHLNK(FSTCOL) = RCHLNK(HEAD)
                RCHLNK(HEAD) = FSTCOL
                KNZ = KNZ + 1
            ENDIF
C
C           --------------------------------------------
C           COPY INDICES FROM LINKED LIST INTO LINDX(*).
C           --------------------------------------------
            NZBEG = NZEND + 1
            NZEND = NZEND + KNZ
            IF  ( NZEND+1 .NE. XLINDX(KSUP+1) )  GO TO 8000
            I = HEAD
            DO  800  KPTR = NZBEG, NZEND
                I = RCHLNK(I)
                LINDX(KPTR) = I
  800       CONTINUE
C
C           ---------------------------------------------------
C           IF KSUP HAS A PARENT, INSERT KSUP INTO ITS PARENT'S 
C           "MERGE" LIST.
C           ---------------------------------------------------
            IF  ( LENGTH .GT. WIDTH )  THEN
                PCOL = LINDX ( XLINDX(KSUP) + WIDTH )
                PSUP = SNODE(PCOL)
                MRGLNK(KSUP) = MRGLNK(PSUP)
                MRGLNK(PSUP) = KSUP
            ENDIF
C
 1000   CONTINUE
C
        RETURN
C
C       -----------------------------------------------
C       INCONSISTENCY IN DATA STRUCTURE WAS DISCOVERED.
C       -----------------------------------------------
 8000   CONTINUE
        FLAG = -2
        RETURN
C
      END


      subroutine genrcm ( node_num, adj_num, adj_row, adj, perm )

!*****************************************************************************80
!
!! GENRCM finds the reverse Cuthill-Mckee ordering for a general graph.
!
!  Discussion:
!
!    For each connected component in the graph, the routine obtains
!    an ordering by calling RCM.
!
!  Modified:
!
!    04 January 2003
!
!  Author:
!
!    Alan George, Joseph Liu
!    FORTRAN90 version by John Burkardt
!
!  Reference:
!
!    Alan George, Joseph Liu,
!    Computer Solution of Large Sparse Positive Definite Systems,
!    Prentice Hall, 1981.
!
!  Parameters:
!
!    Input, integer NODE_NUM, the number of nodes.
!
!    Input, integer ADJ_NUM, the number of adjacency entries.
!
!    Input, integer ADJ_ROW(NODE_NUM+1).  Information about row I is stored
!    in entries ADJ_ROW(I) through ADJ_ROW(I+1)-1 of ADJ.
!
!    Input, integer ADJ(ADJ_NUM), the adjacency structure.
!    For each row, it contains the column indices of the nonzero entries.
!
!    Output, integer PERM(NODE_NUM), the RCM ordering.
!
!  Local Parameters:
!
!    Local, integer LEVEL_ROW(NODE_NUM+1), the index vector for a level
!    structure.  The level structure is stored in the currently unused 
!    spaces in the permutation vector PERM.
!
!    Local, integer MASK(NODE_NUM), marks variables that have been numbered.
!
      implicit none

      integer adj_num,node_num

      integer adj(adj_num)
      integer adj_row(node_num+1)
      integer i
      integer iccsze
      integer mask(node_num)
      integer level_num
      integer level_row(node_num+1)
      integer num
      integer perm(node_num)
      integer root
      

      do i=1,node_num
         mask(i) = 1
      enddo
      num = 1
      
      do i = 1, node_num
!
!  For each masked connected component...
!
         if ( mask(i).ne. 0 ) then

            root = i

!     
!  Find a pseudo-peripheral node ROOT.  The level structure found by
!  ROOT_FIND is stored starting at PERM(NUM).
!
            call root_find ( root, adj_num, adj_row, adj, mask,  
     &       level_num,    level_row, perm(num), node_num )
!     
!  RCM orders the component using ROOT as the starting node.
!     
            call rcm ( root, adj_num, adj_row, adj, mask, perm(num), 
     &        iccsze,    node_num )
            
            num = num + iccsze
!
!  We can stop once every node is in one of the connected components.
!
            if ( node_num .lt. num ) then
               return
            endif
            
         endif

      enddo

      return
      end

      subroutine rcm ( root, adj_num, adj_row, adj, mask, perm, iccsze, 
     &   node_num )

!*****************************************************************************80
!
!! RCM renumbers a connected component by the reverse Cuthill McKee algorithm.
!
!  Discussion:
!
!    The connected component is specified by a node ROOT and a mask.
!    The numbering starts at the root node.
!
!    An outline of the algorithm is as follows:
!
!    X(1) = ROOT.
!
!    for ( I = 1 to N-1)
!      Find all unlabeled neighbors of X(I),
!      assign them the next available labels, in order of increasing degree.
!
!    When done, reverse the ordering.
!
!  Modified:
!
!    02 January 2007
!
!  Author:
!
!    Alan George, Joseph Liu
!    FORTRAN90 version by John Burkardt
!
!  Reference:
!
!    Alan George, Joseph Liu,
!    Computer Solution of Large Sparse Positive Definite Systems,
!    Prentice Hall, 1981.
!
!  Parameters:
!
!    Input, integer ROOT, the node that defines the connected component.
!    It is used as the starting point for the RCM ordering.
!
!    Input, integer ADJ_NUM, the number of adjacency entries.
!
!    Input, integer ADJ_ROW(NODE_NUM+1).  Information about row I is stored
!    in entries ADJ_ROW(I) through ADJ_ROW(I+1)-1 of ADJ.
!
!    Input, integer ADJ(ADJ_NUM), the adjacency structure.
!    For each row, it contains the column indices of the nonzero entries.
!
!    Input/output, integer MASK(NODE_NUM), a mask for the nodes.  Only 
!    those nodes with nonzero input mask values are considered by the 
!    routine.  The nodes numbered by RCM will have their mask values 
!    set to zero.
!
!    Output, integer PERM(NODE_NUM), the RCM ordering.
!
!    Output, integer ICCSZE, the size of the connected component
!    that has been numbered.
!
!    Input, integer NODE_NUM, the number of nodes.
!
!  Local Parameters:
!
!    Workspace, integer DEG(NODE_NUM), a temporary vector used to hold 
!    the degree of the nodes in the section graph specified by mask and root.
!
      implicit none
      
      integer adj_num
      integer node_num
      
      integer adj(adj_num)
      integer adj_row(node_num+1)
      integer deg(node_num)
      integer fnbr
      integer i
      integer iccsze
      integer j
      integer jstop
      integer jstrt
      integer k
      integer l
      integer lbegin
      integer lnbr
      integer lperm
      integer lvlend
      integer mask(node_num)
      integer nbr
      integer node
      integer perm(node_num)
      integer root
!     
!  Find the degrees of the nodes in the component specified by MASK and ROOT.
!
      call degree ( root, adj_num, adj_row, adj, mask, deg, iccsze, 
     &     perm, node_num )

      mask(root) = 0

      if ( iccsze .le. 1 ) then
         return
      end if

      lvlend = 0
      lnbr = 1
!     
!  LBEGIN and LVLEND point to the beginning and
!  the end of the current level respectively.
!
      do while ( lvlend .lt. lnbr )

         lbegin = lvlend + 1
         lvlend = lnbr
         
         do i = lbegin, lvlend
!
!  For each node in the current level...
!
            node = perm(i)
            jstrt = adj_row(node)
            jstop = adj_row(node+1) - 1
!     
!  Find the unnumbered neighbors of NODE.
!
!  FNBR and LNBR point to the first and last neighbors
!  of the current node in PERM.
!
            fnbr = lnbr + 1

            do j = jstrt, jstop

               nbr = adj(j)

               if ( mask(nbr) .ne. 0 ) then
                  lnbr = lnbr + 1
                  mask(nbr) = 0
                  perm(lnbr) = nbr
               end if

            end do
!     
!  If no neighbors, skip to next node in this level.
!
cc            if ( lnbr .le. fnbr ) then
cc               cycle
cc            end if
            if ( lnbr .gt. fnbr ) then
!
!  Sort the neighbors of NODE in increasing order by degree.
!  Linear insertion is used.
!
               k = fnbr

               do while ( k .lt. lnbr )

                  l = k
                  k = k + 1
                  nbr = perm(k)
                  
                  do while ( fnbr .lt. l )

                     lperm = perm(l)

                     if ( deg(lperm) .le. deg(nbr) ) then
                        exit
                     end if

                     perm(l+1) = lperm
                     l = l - 1
                  
                  end do
                  
                  perm(l+1) = nbr

               end do
            end if
            
         end do
         
      end do
!
!  We now have the Cuthill-McKee ordering.  Reverse it.
!
      k=iccsze/2
      l=iccsze
      do i=1,k
         lperm=perm(l)
         perm(l)=perm(i)
         perm(i)=lperm
         l=l-1
      enddo

      return
      end
      subroutine root_find ( root, adj_num, adj_row, adj, mask, 
     & level_num,   level_row, level, node_num )

!*****************************************************************************80
!
!! ROOT_FIND finds a pseudo-peripheral node.
!
!  Discussion:
!
!    The diameter of a graph is the maximum distance (number of edges)
!    between any two nodes of the graph.
!
!    The eccentricity of a node is the maximum distance between that
!    node and any other node of the graph.
!
!    A peripheral node is a node whose eccentricity equals the
!    diameter of the graph.
!
!    A pseudo-peripheral node is an approximation to a peripheral node;
!    it may be a peripheral node, but all we know is that we tried our
!    best.
!
!    The routine is given a graph, and seeks pseudo-peripheral nodes,
!    using a modified version of the scheme of Gibbs, Poole and
!    Stockmeyer.  It determines such a node for the section subgraph
!    specified by MASK and ROOT.
!
!    The routine also determines the level structure associated with
!    the given pseudo-peripheral node; that is, how far each node
!    is from the pseudo-peripheral node.  The level structure is
!    returned as a list of nodes LS, and pointers to the beginning
!    of the list of nodes that are at a distance of 0, 1, 2, ...,
!    NODE_NUM-1 from the pseudo-peripheral node.
!
!  Modified:
!
!    28 October 2003
!
!  Author:
!
!    Alan George, Joseph Liu
!    FORTRAN90 version by John Burkardt
!
!  Reference:
!
!    Alan George, Joseph Liu,
!    Computer Solution of Large Sparse Positive Definite Systems,
!    Prentice Hall, 1981.
!
!    Norman Gibbs, William Poole, Paul Stockmeyer,
!    An Algorithm for Reducing the Bandwidth and Profile of a Sparse Matrix,
!    SIAM Journal on Numerical Analysis,
!    Volume 13, pages 236-250, 1976.
!
!    Norman Gibbs,
!    Algorithm 509: A Hybrid Profile Reduction Algorithm,
!    ACM Transactions on Mathematical Software,
!    Volume 2, pages 378-387, 1976.
!
!  Parameters:
!
!    Input/output, integer ROOT.  On input, ROOT is a node in the
!    the component of the graph for which a pseudo-peripheral node is
!    sought.  On output, ROOT is the pseudo-peripheral node obtained.
!
!    Input, integer ADJ_NUM, the number of adjacency entries.
!
!    Input, integer ADJ_ROW(NODE_NUM+1).  Information about row I is stored
!    in entries ADJ_ROW(I) through ADJ_ROW(I+1)-1 of ADJ.
!
!    Input, integer ADJ(ADJ_NUM), the adjacency structure.
!    For each row, it contains the column indices of the nonzero entries.
!
!    Input, integer MASK(NODE_NUM), specifies a section subgraph.  Nodes 
!    for which MASK is zero are ignored by FNROOT.
!
!    Output, integer LEVEL_NUM, is the number of levels in the level structure
!    rooted at the node ROOT.
!
!    Output, integer LEVEL_ROW(NODE_NUM+1), LEVEL(NODE_NUM), the 
!    level structure array pair containing the level structure found.
!
!    Input, integer NODE_NUM, the number of nodes.
!
      implicit none

      integer adj_num
      integer node_num
      
      integer adj(adj_num)
      integer adj_row(node_num+1)
      integer iccsze
      integer j
      integer jstrt
      integer k
      integer kstop
      integer kstrt
      integer level(node_num)
      integer level_num
      integer level_num2
      integer level_row(node_num+1)
      integer mask(node_num)
      integer mindeg
      integer nabor
      integer ndeg
      integer node
      integer root
!
!  Determine the level structure rooted at ROOT.
!
      call level_set ( root, adj_num, adj_row, adj, mask, level_num, 
     &     level_row, level, node_num )
!
!  Count the number of nodes in this level structure.
!
      iccsze = level_row(level_num+1) - 1
!
!  Extreme case:
!    A complete graph has a level set of only a single level.
!    Every node is equally good (or bad).
!
      if ( level_num .eq. 1 ) then
         return
      end if
!     
!  Extreme case:
!    A "line graph" 0--0--0--0--0 has every node in its only level.
!    By chance, we've stumbled on the ideal root.
!
      if ( level_num .eq. iccsze ) then
         return
      end if
!
!  Pick any node from the last level that has minimum degree
!  as the starting point to generate a new level set.
!
      do

         mindeg = iccsze
         
         jstrt = level_row(level_num)
         root = level(jstrt)

         if ( jstrt .lt. iccsze ) then

            do j = jstrt, iccsze

               node = level(j)
               ndeg = 0
               kstrt = adj_row(node)
               kstop = adj_row(node+1) - 1
               
               do k = kstrt, kstop
                  nabor = adj(k)
                  if ( 0 .lt. mask(nabor) ) then
                     ndeg = ndeg + 1
                  end if
               end do

               if ( ndeg .lt. mindeg ) then
                  root = node
                  mindeg = ndeg
               end if

            end do
            
         end if
!
!  Generate the rooted level structure associated with this node.
!
         call level_set ( root, adj_num, adj_row, adj, mask, 
     &        level_num2,   level_row, level, node_num )
!
!  If the number of levels did not increase, accept the new ROOT.
!
        if ( level_num2 .le. level_num ) then
           exit
        end if

        level_num = level_num2
!
!  In the unlikely case that ROOT is one endpoint of a line graph,
!  we can exit now.
!
        if ( iccsze .le. level_num ) then
           exit
        end if
        
      end do
      
      return
      end

      subroutine level_set ( root, adj_num, adj_row, adj, mask, 
     & level_num,   level_row, level, node_num )

!*****************************************************************************80
!
!! LEVEL_SET generates the connected level structure rooted at a given node.
!
!  Discussion:
!
!    Only nodes for which MASK is nonzero will be considered.
!
!    The root node chosen by the user is assigned level 1, and masked.
!    All (unmasked) nodes reachable from a node in level 1 are
!    assigned level 2 and masked.  The process continues until there
!    are no unmasked nodes adjacent to any node in the current level.
!    The number of levels may vary between 2 and NODE_NUM.
!
!  Modified:
!
!    28 October 2003
!
!  Author:
!
!    Alan George, Joseph Liu
!    FORTRAN90 version by John Burkardt
!
!  Reference:
!
!    Alan George, Joseph Liu,
!    Computer Solution of Large Sparse Positive Definite Systems,
!    Prentice Hall, 1981.
!
!  Parameters:
!
!    Input, integer ROOT, the node at which the level structure
!    is to be rooted.
!
!    Input, integer ADJ_NUM, the number of adjacency entries.
!
!    Input, integer ADJ_ROW(NODE_NUM+1).  Information about row I is stored
!    in entries ADJ_ROW(I) through ADJ_ROW(I+1)-1 of ADJ.
!
!    Input, integer ADJ(ADJ_NUM), the adjacency structure.
!    For each row, it contains the column indices of the nonzero entries.
!
!    Input/output, integer MASK(NODE_NUM).  On input, only nodes with nonzero
!    MASK are to be processed.  On output, those nodes which were included
!    in the level set have MASK set to 1.
!
!    Output, integer LEVEL_NUM, the number of levels in the level
!    structure.  ROOT is in level 1.  The neighbors of ROOT
!    are in level 2, and so on.
!
!    Output, integer LEVEL_ROW(NODE_NUM+1), LEVEL(NODE_NUM), the rooted 
!    level structure.
!
!    Input, integer NODE_NUM, the number of nodes.
!
      implicit none

      integer adj_num
      integer node_num
      
      integer adj(adj_num)
      integer adj_row(node_num+1)
      integer i
      integer iccsze
      integer j
      integer jstop
      integer jstrt
      integer lbegin
      integer level_num
      integer level_row(node_num+1)
      integer level(node_num)
      integer lvlend
      integer lvsize
      integer mask(node_num)
      integer nbr
      integer node
      integer root
      
      mask(root) = 0
      level(1) = root
      level_num = 0
      lvlend = 0
      iccsze = 1
!
!  LBEGIN is the pointer to the beginning of the current level, and
!  LVLEND points to the end of this level.
!
      do

         lbegin = lvlend + 1
         lvlend = iccsze
         level_num = level_num + 1
         level_row(level_num) = lbegin
!     
!  Generate the next level by finding all the masked neighbors of nodes
!  in the current level.
!
         do i = lbegin, lvlend

            node = level(i)
            jstrt = adj_row(node)
            jstop = adj_row(node+1) - 1
            
            do j = jstrt, jstop

               nbr = adj(j)

               if ( mask(nbr) .ne. 0 ) then
                  iccsze = iccsze + 1
                  level(iccsze) = nbr
                  mask(nbr) = 0
               end if
               
            end do
            
         end do
!
!  Compute the current level width (the number of nodes encountered.)
!  If it is positive, generate the next level.
!
         lvsize = iccsze - lvlend

         if ( lvsize .le. 0 ) then
            exit
         end if
         
      end do
      
      level_row(level_num+1) = lvlend + 1
!
!  Reset MASK to 1 for the nodes in the level structure.
!
      do i =1 ,iccsze
         mask(level(i)) = 1
      enddo
      return
      end


      subroutine degree ( root, adj_num, adj_row, adj, mask, deg, 
     & iccsze, ls,  node_num )

!*****************************************************************************80
!
!! DEGREE computes the degrees of the nodes in the connected component.
!
!  Discussion:
!
!    The connected component is specified by MASK and ROOT.
!    Nodes for which MASK is zero are ignored.
!
!  Modified:
!
!    05 January 2003
!
!  Author:
!
!    Alan George, Joseph Liu
!    FORTRAN90 version by John Burkardt
!
!  Reference:
!
!    Alan George, Joseph Liu,
!    Computer Solution of Large Sparse Positive Definite Systems,
!    Prentice Hall, 1981.
!
!  Parameters:
!
!    Input, integer ROOT, the node that defines the connected component.
!
!    Input, integer ADJ_NUM, the number of adjacency entries.
!
!    Input, integer ADJ_ROW(NODE_NUM+1).  Information about row I is stored
!    in entries ADJ_ROW(I) through ADJ_ROW(I+1)-1 of ADJ.
!
!    Input, integer ADJ(ADJ_NUM), the adjacency structure.
!    For each row, it contains the column indices of the nonzero entries.
!
!    Input, integer MASK(NODE_NUM), is nonzero for those nodes which are
!    to be considered.
!
!    Output, integer DEG(NODE_NUM), contains, for each  node in the connected
!    component, its degree.
!
!    Output, integer ICCSIZE, the number of nodes in the connected component.
!
!    Output, integer LS(NODE_NUM), stores in entries 1 through ICCSIZE the nodes
!    in the connected component, starting with ROOT, and proceeding 
!    by levels.
!
!    Input, integer NODE_NUM, the number of nodes.
!
      implicit none
      
      integer adj_num
      integer node_num

      integer adj(adj_num)
      integer adj_row(node_num+1)
      integer deg(node_num)
      integer i
      integer iccsze
      integer ideg
      integer j
      integer jstop
      integer jstrt
      integer lbegin
      integer ls(node_num)
      integer lvlend
      integer lvsize
      integer mask(node_num)
      integer nbr
      integer node
      integer root
!     
!  The sign of ADJ_ROW(I) is used to indicate if node I has been considered.
      ls(1) = root
      adj_row(root) = -adj_row(root)
      lvlend = 0
      iccsze = 1
!
!  LBEGIN is the pointer to the beginning of the current level, and
!  LVLEND points to the end of this level.
      do

         lbegin = lvlend + 1
         lvlend = iccsze
!
!  Find the degrees of nodes in the current level,
!  and at the same time, generate the next level.
         do i = lbegin, lvlend

            node = ls(i)
            jstrt = -adj_row(node)
            jstop = abs ( adj_row(node+1) ) - 1
            ideg = 0

            do j = jstrt, jstop

               nbr = adj(j)

               if ( mask(nbr) .ne. 0 ) then

                  ideg = ideg + 1

                  if ( 0 .le. adj_row(nbr) ) then
                     adj_row(nbr) = -adj_row(nbr)
                     iccsze = iccsze + 1
                     ls(iccsze) = nbr
                  end if
                  
               end if

            end do

            deg(node) = ideg
            
         end do
!
!  Compute the current level width.    
         lvsize = iccsze - lvlend
!
!  If the current level width is nonzero, generate another level.
         if ( lvsize .eq. 0 ) then
            exit
         end if

      end do
!
!  Reset ADJ_ROW to its correct sign and return.
      do i = 1, iccsze
         node = ls(i)
         adj_row(node) = -adj_row(node)
      end do
      
      return
      end
```
