## Supplementary material for "MoBPS - Modular Breeding Program Simulator": RandomFieldsUtils R-package: linear.cc

```
/* 
 Authors
 Martin Schlather,

 Collection of system specific auxiliary functions

 Copyright (C) 2001 -- 2017 Martin Schlather, 

This program is free software; you can redistribute it and/or
modify it under the terms of the GNU General Public License
as published by the Free Software Foundation; either version 3
of the License, or (at your option) any later version.

This program is distributed in the hope that it will be useful,
but WITHOUT ANY WARRANTY; without even the implied warranty of
MERCHANTABILITY or FITNESS FOR A PARTICULAR PURSE.  See the
GNU General Public License for more details.

You should have received a copy of the GNU General Public License
along with this program; if not, write to the Free Software
Foundation, Inc., 59 Temple Place - Suite 330, Boston, MA  02111-1307, USA.
*/

/*

Makefile must be:

PKG_LIBS =  $(SHLIB_OPENMP_CXXFLAGS) $(LAPACK_LIBS) $(BLAS_LIBS) $(FLIBS)  -march=native  -mssse3 
PKG_CXXFLAGS =  $(SHLIB_OPENMP_CXXFLAGS)  -march=native -mssse3 

 */


#define BUG assert(false);

#include <assert.h>
#include "kleinkram.h"
#include "linear.h"
#include "intrinsics.h"
#include "Basic_utils.h"
#include "errors_messages.h"

#define Nlinmodi 9
name_type linmodi = { "1x1", "2x2", "4x4", "8x8", "near", "simple", "precise",
		      "kahan", "1x1p"};


typedef unsigned int uint32;


#define size 8
#define vectorlen (256 / (size * 8))
#define repet 8
#define atonce (vectorlen * repet)
#define VECTOR _mm256_loadu_pd
#define SET_0(NR) sum##NR = _mm256_setzero_pd()
#define P_0(NR) prod##NR = _mm256_setzero_pd()
#define SUMUP(NR, nr) sum##NR = _mm256_add_pd(sum##NR, sum##nr)
#define ADDF(NR) \
  sum##NR = _mm256_fmadd_pd(VECTOR(x + i + NR * vectorlen),\
			    VECTOR(y + i + NR * vectorlen), sum##NR)
#define ADDN(NR)							\
  prod##NR = _mm256_mul_pd(VECTOR(x + i + NR * vectorlen),		\
			   VECTOR(y + i + NR * vectorlen));		\
  sum##NR = _mm256_add_pd(sum##NR, prod##NR) 


#if (7 != repet - 1)
  wrong repet length
#endif
#if (3 != vectorlen - 1)
  wrong vector length
#endif

  
#ifdef AVX
  /*
void avx_linearprodDnearfma(double * x, double y, int len) {
  // deutlich genauer zum 0 tarif
  int i = 0,
     lenM = len - (atonce - 1);  
  __m256d SET_0(0), SET_0(1), SET_0(2), SET_0(3), SET_0(4), SET_0(5), SET_0(6),SET_0(7),
    P_0(0), P_0(1), P_0(2), P_0(3), P_0(4), P_0(5), P_0(6),P_0(7);

   double *D  = (double *) &sum0;

  if ( len >= atonce) {
    for (; i < lenM; i += atonce) {
      //
      ADDN(0); ADDN(1); ADDN(2); ADDN(3); ADDN(4); ADDN(5); ADDN(6); ADDN(7); 
    }
    SUMUP(0, 1); SUMUP(2, 3); SUMUP(4, 5); SUMUP(6, 7);
    SUMUP(0, 2); SUMUP(4, 6); SUMUP(0, 4);
  }
  lenM = len - vectorlen + 1;
  for (; i < lenM; i += vectorlen) { 
    ADDN(0);
  }

  double sum = D[0] + D[1] + D[2] + D[3];

  for (; i < len; i++) sum += x[i] * y[i];
}
 
  */

#define MUL(NR)								\
  _mm256_storeu_pd(inout + i + NR * vectorlen,				\
		   _mm256_add_pd(VECTOR(inout + i + NR * vectorlen),	\
				 _mm256_mul_pd(VECTOR(x + i + NR * vectorlen), \
					       y)))
			  
void avx_linearprodD(double * x, double  Y, int len, double *inout) {
  int i = 0,
    lenM = len - (atonce - 1);  
  __m256d y = _mm256_set1_pd(Y);
 
  for (; i < lenM; i += atonce) {
    MUL(0); MUL(1); MUL(2); MUL(3); MUL(4); MUL(5); MUL(6); MUL(7);
    // for (int k=0; k<atonce; k++) printf("k=%d %10g %10g %10g\n", i+k, inout[i+k], Y, x[i+k]);
  }
 

  lenM = len - vectorlen + 1;
  for (; i < lenM; i += vectorlen) {
    MUL(0);
  }
 
  for (; i < len; i++) inout[i] += x[i] * Y;
 }


/*

void avx_linearprodDparallel(double * x, double y, int len) {
   int i = 0,
    lenM = len - (atonce - 1);  
  __m256d SET_0(0), P_0(0);
    //zero =_mm256_setzero_pd();
   double *D  = (double *) &sum0;


  if ( len >= atonce) {
#ifdef DO_PARALLEL
#pragma omp declare reduction(addpd:  __m256d:				\
			      omp_out = _mm256_add_pd(omp_out, omp_in))	\
			      initializer (omp_priv = _mm256_setzero_pd())
#endif
#ifdef DO_PARALLEL
#pragma omp parallel for num_threads(CORES) reduction(addpd:sum0)
#endif
  for (i=0; i < lenM; i += atonce) {
    //
    ADD(0); ADD(1); ADD(2); ADD(3); ADD(4); ADD(5); ADD(6); ADD(7); 
    }
  }

  lenM = len - vectorlen + 1;
  for (; i < lenM; i += vectorlen) { 
    ADD(0);
  }
  
  double sum = D[0] + D[1] + D[2] + D[3];

  for (; i < len; i++) sum += x[i] * y[i];
}


void avx_linearprodDP(double * x, double y, int len) {
  int i = 0,
     lenM = len - (atonce - 1);  
  __m256d SET_0(0), SET_0(1), P_0(0);
   double *D  = (double *) &sum1;

  if ( len >= atonce) {
    
    for (; i < lenM; ) {
      int lenMM = i + vectorlen * (repet * 10 + 1);
      if (lenMM > lenM) lenMM = lenM;
      sum0 = _mm256_mul_pd(VECTOR(x + i), VECTOR(y + i));
      i += vectorlen;
      for (; i < lenMM; i += atonce) {
	ADD(0); ADD(1); ADD(2); ADD(3); ADD(4); ADD(5); ADD(6); ADD(7); 
	  }
      sum1 = _mm256_add_pd(sum0, sum1);
    }
  }
  
 lenM = len - vectorlen + 1;
 for (; i < lenM; i += vectorlen) { 
    prod0 = _mm256_mul_pd(VECTOR(x + i), VECTOR(y + i));
    sum1 = _mm256_add_pd(sum1, prod0);
  }
  
  double sum = D[0] + D[1] + D[2] + D[3];

  for (; i < len; i++) {
    // printf("final %d\n", i);
    sum += x[i] * y[i];
  }
}


#define ADDK(NR)								\
  prod0 = _mm256_mul_pd(VECTOR(x + i + NR * vectorlen),		\
			   VECTOR(y + i + NR * vectorlen));		\
  sum2 = _mm256_sub_pd(prod0, sum1);\
  sum3 = _mm256_add_pd(sum0, sum2);		\
  sum1 = _mm256_sub_pd(sum3, sum0);		\
  sum0 = sum3;					\
  sum1 = _mm256_sub_pd(sum1, sum2);
void avx_linearprodDK(double * x, double  y, int len) {
   // Kahan enhanced
  int i = 0,
     lenM = len - (atonce - 1);  
  __m256d SET_0(0), // sum
    SET_0(1),  // c
    SET_0(2), // y
    SET_0(3),  // t
    P_0(0),
    P_0(1);
   double *D  = (double *) &sum0;

  if ( len >= atonce) {
  for (; i < lenM; i += atonce) {
    ADDK(0); ADDK(1); ADDK(2); ADDK(3); ADDK(4); ADDK(5); ADDK(6); ADDK(7);
  }
  }
 lenM = len - vectorlen + 1;
 for (; i < lenM; i += vectorlen) { 
    ADDK(0);
 }
 sum0 = _mm256_add_pd(sum0, prod1); 
  
  double sum = D[0] + D[1] + D[2] + D[3];

  for (; i < len; i++) sum += x[i] * y[i];
}

  */
  
#endif
 
/*
void linearprod( double * v1,  double v2, int N){
  double *endv1 = v1 + N,
    sum = 0;
    for(; v1!= endv1; v1++) sum+= v2 * v1[0];
}
 
 
 
void linearprodP( double * v1,  double v2, int N){
  double // *endv1 = v1 + N,
    sum = 0;
#ifdef DO_PARALLEL
#pragma omp parallel for num_threads(CORES) reduction(+:sum)
#endif
  for(int i=0; i<=N; i++) sum += v2 * v1[i];
}
*/
 


void linearprod2by2( double * v1,  double v2, int N, double *inout){
  double *endv1 = v1 + (N / 2) * 2,
    *end = v1 + N;
  for(; v1 < endv1; v1+=2, inout+=2) {
      inout[0] += v2 * v1[0];
      inout[1] += v2 * v1[1];
  }
  if (v1 < end) inout[0] += v2 * v1[0];
}
 
 /*
void linearprod4by4( double * v1,  double v2, int N){
  double*endv1 = v1 + (N / 4) * 4,
    *end = v1 + N,
      sum = 0;
    for(; v1 < endv1; v1+=4) {
      sum+= v2 * v1[0] + v2 * v1[1] + v2 * v1[2]+ v2 * v1[3];
    }
    for(; v1 < end; v1++) sum += v2 * v1[0];        
}

 
void linearprod8by8( double * v1,  double v2, int N){
  double
    *endv1 = v1 + (N / 8) * 8,
    *end = v1 + N,
      sum = 0;
    for(; v1 < endv1; v1+=8) {
      sum+= v2 * v1[0] + v2 * v1[1]+ v2 * v1[2] + v2 * v1[3] +
	v2 * v1[4] + v2 * v1[5]+ v2 * v1[6]+ v2 * v1[7];
    }
    for(; v1 < end; v1++) sum +=  v2 * v1[0];        
}
*/

//bool pr = true;
void linearX(double *x, double y, int len, double *inout, int n) {
  //  if (n < 0) {  }
//  if (pr) { printf("mode = %d\n", n); pr = false; }
 // 0 : 7.9
// 1:  7.55
// 2: 7.8
// 3:7.58
//4: 7.5 
// 5: 7.4!
//6:7.4
//7: 7.9
// 8: "ewige" schleife

  switch(n) {
  case 0 : BUG; // linearprod(x, y, len); break;
  case LINEAR_BASE :  linearprod2by2(x, y, len, inout); break; 
  case 2 : BUG;  //linearprod4by4(x, y, len); break; 
  case 3 : BUG; // linearprod8by8(x, y, len); break; 
#ifdef FMA_AVAILABLE
  case 4 : BUG;  //avx_linearprodDfma(x, y, len); break;
#endif    
#ifdef AVX
  case 5 : BUG;  //avx_linearprodDnearfma(x, y, len); break; 
  case LINEAR_AVX :  avx_linearprodD(x, y, len, inout); break; // best one kernel
  case 7 : BUG; // avx_linearprodDP(x, y, len); break;  //best
  case 8 : BUG  //avx_linearprodDK(x, y, len); break; // kahan
#else
  case 4: case 5: case 6: case 7: case 8 :  linearprod2by2(x, y, len, inout); break;
#endif
    
#ifdef DO_PARALLEL
  case LINEAR_AVX_PARALLEL :
#ifdef AVX
    BUG; // avx_linearprodDparallel(x, y, len); break;
#endif    
  case LINEAR_BASE_PARALLEL :  BUG; //linearprodP(x, y, len); break;// parallel, nicht-vectoriell
#else
  case LINEAR_AVX_PARALLEL :
#ifdef AVX
    BUG; // avx_linearprodD(x, y, len); break;
#endif    
  case LINEAR_BASE_PARALLEL :  BUG; // linearprod2by2(x, y, len); break; 
#endif  
  default : {{ ERR("method not available"); }} //
  }
}
  
```
