## Supplementary material for "MoBPS - Modular Breeding Program Simulator": RandomFieldsUtils R-package: maths.cc

```
/*
 Authors 
 Martin Schlather,

 Copyright (C) 2015 -- 2017 Martin Schlather

This program is free software; you can redistribute it and/or
modify it under the terms of the GNU General Public License
as published by the Free Software Foundation; either version 3
of the License, or (at your option) any later version.

This program is distributed in the hope that it will be useful,
but WITHOUT ANY WARRANTY; without even the implied warranty of
MERCHANTABILITY or FITNESS FOR A PARTICULAR PURPOSE.  See the
GNU General Public License for more details.

You should have received a copy of the GNU General Public License
along with this program; if not, write to the Free Software
Foundation, Inc., 59 Temple Place - Suite 330, Boston, MA  02111-1307, USA.  
*/
#include <R_ext/Lapack.h>
#include "RandomFieldsUtils.h"
#include "zzz_RandomFieldsUtils.h"
#include "General_utils.h"


double struve_intern(double x, double nu, double factor_Sign, bool expscaled)
{ 
 if ((x == 0.0) && (nu>-1.0)) return 0.0;
  if (x <= 0.0) return RF_NA; // not programmed yet
  double exp_dummy,
     dummy = 0.0, 
     logx = 2.0 * LOG(0.5 * x), 
     x1 = 1.5, 
     x2 = nu + 1.5,
     value = 1.0, 
     fsign = factor_Sign,
    epsilon=1e-20;

  
   do {
     dummy += logx - LOG(x1) - LOG(FABS(x2));
     exp_dummy = EXP(dummy);
     value += (1 - 2 * (x2 < 0))  * fsign * exp_dummy;
     //  printf("%10g %10g %10g %10g\n", value, fsign, x1, x2);
     x1 += 1.0;
     x2 += 1.0;
     fsign = factor_Sign * fsign; 
   } while (exp_dummy > FABS(value) * epsilon);

 

   x1 = 1.5;
   x2 = nu + 1.5;
   if (x2 > 0.0) { 
     dummy = (nu + 1.0) * 0.5 * logx - lgammafn(x1) - lgammafn(x2);
     if (expscaled) dummy -= x;
     value *= EXP(dummy);
   } else {
     //if ( (double) ((int) (x1-0.5)) != x1-0.5 ) return RF_NA;
     value *= POW(0.5 * x, nu + 1.0) / (gammafn(x1) * gammafn(x2));
     if (expscaled) value *= EXP(-x);
   }

  return value;
}

//void Struv eH(double *x, double *nu) {*x=struv e(*x, *nu, -1.0, false);}
//void Struv eL(double *x, double *nu, int * expScaled) {
//  *x=struv e(*x, *nu, 1.0, (bool) *expScaled);
//}
double StruveH(double x, double nu) {return struve_intern(x, nu, -1.0, false);}
double StruveL(double x, double nu, bool expScaled) {
 return struve_intern(x, nu, 1.0, expScaled);
}


SEXP struve(SEXP X, SEXP Nu, SEXP Factor_Sign, SEXP Expscaled) {
  int i,    
    lenx = length(X),
    lennu = length(Nu),
    len = lenx;  
  if (len < lennu) len = lennu;
  SEXP Result;
  PROTECT(Result = allocVector(REALSXP, len));
  double *x = REAL(X),
    *nu  = REAL(Nu),
    factor_sign = REAL(Factor_Sign)[0],
    *result = REAL(Result);
  bool expscaled = LOGICAL(Expscaled)[0];
  for (i=0; i<len; i++)
    result[i]=struve_intern(x[i % lenx], nu[i % lennu], factor_sign, expscaled);
 
  UNPROTECT(1);
  return Result;
}


//void I0ML0(double *x, int *n) {
//  int i;
//  for (i=0; i<*n; i++) x[i] = I0mL0(x[i]);
//} 


double I0mL0(double x){
  /* Bessel I_0 - Struve L_0 for non-negative arguments x */
  /* see J. MacLeod, Chebyshev expansions for modified {S}truve and 
     related functions, Mathematics of Computation, 60, 735-747, 1993 */
  static double g2[24] = {
	0.52468736791485599138e0,
	-0.35612460699650586196e0,
	0.20487202864009927687e0,
	-0.10418640520402693629e0,
	0.4634211095548429228e-1,
	-0.1790587192403498630e-1,
	0.597968695481143177e-2,
	-0.171777547693565429e-2,
	0.42204654469171422e-3,
	-0.8796178522094125e-4,
	0.1535434234869223e-4,
	-0.219780769584743e-5,
	0.24820683936666e-6,
	-0.2032706035607e-7,
	0.90984198421e-9,
	0.2561793929e-10,
	-0.710609790e-11,
	0.32716960e-12,
	0.2300215e-13,
	-0.292109e-14,
	-0.3566e-16,
	0.1832e-16,
	-0.10e-18,
	-0.11e-18
  };
  static double g3[24] = {
	2.00326510241160643125e0,
	0.195206851576492081e-2,
	0.38239523569908328e-3,
	0.7534280817054436e-4,
	0.1495957655897078e-4,
	0.299940531210557e-5,
	0.60769604822459e-6,
	0.12399495544506e-6,
	0.2523262552649e-7,
	0.504634857332e-8,
	0.97913236230e-9,
	0.18389115241e-9,
	0.3376309278e-10,
	0.611179703e-11,
	0.108472972e-11,
	0.18861271e-12,
	0.3280345e-13,
	0.565647e-14,
	0.93300e-15,
	0.15881e-15,
	0.2791e-16,
	0.389e-17,
	0.70e-18,
	0.16e-18
  };
  double r, x2, ac;
  int i;
  
  if (x < 0.0) {return RF_NA;}
  if (x < 16.0) {
    r = 0.5 * g2[0];
    ac = ACOS((6.0 * x - 40.0) / (x + 40.0));
    for (i=1; i<24; i++) {
      r += g2[i] * cos(i * ac);
    }
  } else {
    r = 0.5 * g3[0];
    x2 = x * x;
    ac = ACOS((800.0 - x2) / (288.0 + x2));
    for (i=1; i<24; i++) {
      r += g3[i] * cos(i * ac);
    }
    r *= T_PI /* 2/pi */ / x;
  }
  return r;
}


SEXP I0ML0(SEXP X) {
  SEXP Result;
  PROTECT(Result = allocVector(REALSXP, length(X)));
  double *x = REAL(X),
    *result = REAL(Result);
  int i,    
    lenx = length(X);  
  for (i=0; i<lenx; i++) result[i] = I0mL0(x[i]);

  UNPROTECT(1);
  return Result;
}


/* Gausian model */
double Gauss(double x) {
  return EXP(- x * x);
  //  printf("%10g %10g\n", *x, *v);
}
double logGauss(double x) {
  return - x * x;
}
double DGauss(double y) {
  return -2.0 * y * EXP(- y * y);
}
double DDGauss(double x) {
  double y = x * x; 
  return (4.0 * y - 2.0)* EXP(- y);
}
double D3Gauss(double x) {
  double y = x * x; 
  return x * (12 - 8 * y) * EXP(- y);
}
double D4Gauss(double x) {
  double y = x * x; 
  return ((16 * y - 48) * y + 12) * EXP(- y);
}


/* Whittle-Matern or Whittle or Besset ---- rescaled form of Whittle-Matern,
    see also there */ 

#define LOW_MATERN 1e-20
double logWM(double x, double nu1, double nu2, double factor) {
  // check calling functions, like hyperbolic and gneiting if any changings !!

  //  printf("%10g %10g %10g %10g\n", x, nu1, nu2, factor);

#ifdef DO_PARALLEL
  double loggamma;
#else 
  static double loggamma, loggamma1old, loggamma2old, loggamma_old, 
    nuOld=-RF_INF,
    nu1old=-RF_INF,
    nu2old=-RF_INF;
#endif  
  double v, y, 
    nu = 0.5 * (nu1 + nu2),
    nuThres = nu < MATERN_NU_THRES ? nu : MATERN_NU_THRES,
		   scale = 1.0;
  if (factor!=0.0) scale = factor * SQRT(nuThres);
  bool simple = nu1 == nu2 || nu > MATERN_NU_THRES;
  double bk[MATERN_NU_THRES + 1L];

  if (x > LOW_MATERN) {
#ifdef DO_PARALLEL
    if (simple) loggamma = lgammafn(nuThres);
    else loggamma = 0.5*(lgammafn(nu1) + lgammafn(nu2));
#else 
    if (simple) {
      if (nuThres != nuOld) {
	nuOld = nuThres;
	loggamma_old = lgammafn(nuThres);
      } 
      loggamma = loggamma_old;      
    } else {
      if (nu1 != nu1old) {
	nu1old = nu1;
	loggamma1old = lgammafn(nu1);
      }
      if (nu2 != nu2old) {
	nu2old = nu2;
	loggamma2old = lgammafn(nu2);
      }
      loggamma = 0.5 * (loggamma1old + loggamma2old);
    }
#endif  
    
    y = x  * scale;
    v = LOG2 + nuThres * LOG(0.5 * y) - loggamma + 
      LOG(bessel_k_ex(y, nuThres, 2.0, bk)) - y;
  } else v = 0.0;
    
  if (nu > MATERN_NU_THRES) { // factor!=0.0 && 
    double w, 
      g = MATERN_NU_THRES / nu;
    y = x * factor / 2;
    w = logGauss(y);

    //if (nu>100) printf("nu=%10g %10e %10e %10e\n", nu, v, g, w);

    v = v * g + (1.0 - g) * w;
    if (nu1 != nu2) { // consistenz zw. nu1, nu2 und nuThres wiederherstellen
      v += lgammafn(nu)- 0.5 * (lgammafn(nu1) + lgammafn(nu2)); // !nuThres
    }
    
    // if (!R_FINITE(v)) ERR("non-finite value in the whittle-matern model -- value of 'nu' is much too large");

    //if (nu>100) printf("v=%10g \n", v);
  }

  return v;
}


double WM(double x, double nu, double factor) {
  // check calling functions, like hyperbolic and gneiting if any changings !!
  return EXP(logWM(x, nu, nu, factor));
}

double DWM(double x, double nu, double factor) { 
#ifdef DO_PARALLEL
  double loggamma;
#else 
  static double nuOld=RF_INF;
  static double loggamma;
#endif  
  double   y, v,
    nuThres = nu < MATERN_NU_THRES ? nu : MATERN_NU_THRES,
		   scale = 1.0;
  if (factor != 0.0) scale = factor * SQRT(nuThres);
  double bk[MATERN_NU_THRES + 1L];
  
  if (x > LOW_MATERN) {
#ifdef DO_PARALLEL
    loggamma = lgammafn(nuThres);
#else 
   if (nuThres!=nuOld) {
      nuOld = nuThres;
      loggamma = lgammafn(nuThres);
    }
#endif    
    y = x * scale;  
    v = - 2.0 * EXP(nuThres * LOG(0.5 * y) - loggamma + 
			     LOG(bessel_k_ex(y, nuThres - 1.0, 2.0, bk)) - y);
  } else {
    v = (nuThres > 0.5) ? 0.0 : (nuThres < 0.5) ? INFTY : 1.253314137;
  }
  v *= scale;

  if (nu > MATERN_NU_THRES) {
    double w, 
      g = MATERN_NU_THRES / nu;
    scale = factor / 2.0;
    y = x * scale;
    w = DGauss(y) * scale;
    v = v * g + (1.0 - g) * w;
  }
  return v;
}

double DDWM(double x, double nu, double factor) { 
#ifdef DO_PARALLEL
  double gamma;
#else 
  static double nuOld=RF_INF;
  static double gamma;
#endif  
  double  y, v,
    nuThres = nu < MATERN_NU_THRES ? nu : MATERN_NU_THRES,
		   scale = 1.0;
  if (factor != 0.0) scale = factor * SQRT(nuThres);
  double scaleSq  = scale * scale,
		   bk[MATERN_NU_THRES + 1L];
  
  if (x > LOW_MATERN) {
#ifdef DO_PARALLEL
    gamma = gammafn(nuThres);
#else 
    if (nuThres!=nuOld) {
      nuOld = nuThres;
      gamma = gammafn(nuThres);
    }
#endif    
    y = x * scale;
    v = POW(0.5 * y , nuThres - 1.0) / gamma *
      (- bessel_k_ex(y, nuThres - 1.0, 1.0, bk)
       + y * bessel_k_ex(y, nuThres - 2.0, 1.0, bk));
  } else {
    v = (nu > 1.0) ? -0.5 / (nu - 1.0) : INFTY;
  }
  v *= scaleSq;

  if (nu > MATERN_NU_THRES) {
    double w, 
      g = MATERN_NU_THRES / nu;
    scale = factor / 2.0;
    scaleSq = scale * scale;
    y = x * scale;
    w = DDGauss(y) * scaleSq;
    v = v * g + (1.0 - g) * w;
  }
  return v;
}

double D3WM(double x, double nu, double factor) { 
#ifdef DO_PARALLEL
  double gamma;
#else 
  static double nuOld=RF_INF;
  static double gamma;
#endif  
  double y, v,
    nuThres = nu < MATERN_NU_THRES ? nu : MATERN_NU_THRES,
    scale = (factor != 0.0) ? factor * SQRT(nuThres) : 1.0,
    scaleSq  = scale * scale;
   double bk[MATERN_NU_THRES + 1L];
 
  if (x > LOW_MATERN) {
#ifdef DO_PARALLEL
     gamma = gammafn(nuThres);
 #else 
    if (nuThres!=nuOld) {
      nuOld = nuThres;
      gamma = gammafn(nuThres);
    }
#endif    
    y = x * scale;
    v = POW(0.5 * y , nuThres - 1.0) / gamma *
      ( 3.0 * bessel_k_ex(y, nuThres - 2.0, 1.0, bk) 
	-y * bessel_k_ex(y, nuThres - 3.0, 1.0, bk)); 
  } else {
    v = 0.0;
  }
  v *= scaleSq * scale;
 
  if (nu > MATERN_NU_THRES) {
    double w, 
      g = MATERN_NU_THRES / nu;
    scale = factor / 2.0;
    scaleSq = scale * scale;
    y = x * scale;
    w = D3Gauss(y) * scaleSq * scale;
    v = v * g + (1.0 - g) * w;
  }
  return v;
}

double D4WM(double x,  double nu, double factor) { 
  double y, v,
    nuThres = nu < MATERN_NU_THRES ? nu : MATERN_NU_THRES,
    scale = (factor != 0.0) ? factor * SQRT(nuThres) : 1.0,
    scaleSq  = scale * scale;
  double bk[MATERN_NU_THRES + 1L];

  //  printf("x=%10g nu=%10g\n", x, nuThres);
  
  if (x > LOW_MATERN) {
#ifdef DO_PARALLEL
    double gamma = gammafn(nuThres);
#else 
    static double nuOld=RF_INF;
    static double gamma;
    if (nuThres!=nuOld) {
      nuOld = nuThres;
      gamma = gammafn(nuThres);
    }
#endif    
    y = x * scale;
    v = 0.25 * POW(0.5 * y , nuThres - 3.0) / gamma *
      (+ 6.0 * (nuThres - 3.0 - y * y) * bessel_k_ex(y, nuThres - 3.0, 1.0, bk)
       + y * (3.0  + y * y) * bessel_k_ex(y, nuThres - 4.0, 1.0, bk)); 
  } else {
    v = INFTY;
    if (nuThres > 2.0) v = 0.75 / ((nuThres - 1.0) * (nuThres - 2.0));
  }
  v *= scaleSq * scaleSq;

  if (nu > MATERN_NU_THRES) {
    double w, 
      g = MATERN_NU_THRES / nu;
    scale = factor / 2.0;
    scaleSq = scale * scale;
    y = x * scale;
    w = D4Gauss(y) * scaleSq * scaleSq;
    v = v * g + (1.0 - g) * w;
  }

  // printf("v=%10g\n", v);
  
  return v;
}


typedef double (*primfct1)(double);
typedef double (*primfct3)(double, double, double);
#define CALCULATE(PRIMFCTN)			\
  double *x = REAL(X);				\
  int n = length(X),				\
    deriv = INTEGER(Derivative)[0];					\
  if (deriv < 0 || deriv > 4) ERR("value of 'derivative' out of range"); \
  PRIMFCTN F = fctns[deriv];						\
									\
  SEXP Ans;								\
  PROTECT(Ans=allocVector(REALSXP, n));					\
  double *ans = REAL(Ans);						\
  for (int i=0; i<n; i++) ans[i] = F

#define RETURN					\
  UNPROTECT(1);					\
  return(Ans);


SEXP gaussr(SEXP X, SEXP Derivative) {  
  static primfct1 fctns[] = {Gauss, DGauss, DDGauss, D3Gauss, D4Gauss};
  CALCULATE(primfct1)(FABS(x[i]));
  RETURN;
}

SEXP WMr(SEXP X, SEXP Nu, SEXP Derivative, SEXP Factor) {  
  static primfct3 fctns[] = {WM, DWM, DDWM, D3WM, D4WM };
  double 
    *nu = REAL(Nu),
    *factor = REAL(Factor);
  int 
    nnu = length(Nu),
    nfactor = length(Factor);  
  CALCULATE(primfct3)(FABS(x[i]), nu[i % nnu], factor[i % nfactor]);
  RETURN;
}
 

SEXP logWMr(SEXP X, SEXP Nu1, SEXP Nu2, SEXP Factor) {  
  double 
    nu1 = REAL(Nu1)[0],
    nu2 = REAL(Nu2)[0],
    factor = REAL(Factor)[0];
  double *x = REAL(X);				
  //  int n = length(X);	
  if (nu1 <= 0.0 || nu2 <= 0.0) ERR("'nu' must be positive");
  if (factor < 0.0) ERR("'factor' must be positive");
 									
  SEXP Ans;								
  PROTECT(Ans=allocVector(REALSXP, 1));					
  double *ans = REAL(Ans);						
  ans[0] = logWM(FABS(x[0]), nu1, nu2, factor);
  UNPROTECT(1);					
  return(Ans);
}


/////////////////////////////////////
// not documente yet:


double incomplete_gamma(double start, double end, double s) {
  // int_start^end t^{s-1} e^{-t} \D t

  // print("incomplete IN s=%10g e=%10g s=%10g\n", start, end, s);

  double
    v = 0.0, 
    w = 0.0;

  if (s <= 1.0) {
    if (start == 0.0) return RF_NAN;
  }
  
  double 
    e_start = EXP(-start),
    e_end = EXP(-end),
    power_start = POW(start, s),      
    power_end = end < RF_INF ? POW(end, s) : 0,
    factor = 1.0; 
  
  
  while (s < 0.0) {
    factor /= s;    
    v +=  factor * (power_end * e_end - power_start * e_start);
    power_start *= start;
    if (end < RF_INF) power_end *= end;
    s += 1.0;
  }
  
  w = pgamma(start, s, 1.0, 0, 0);  // q, shape, scale, lower, log
  if (R_FINITE(end)) w -= pgamma(end, s, 1.0, 0, 0);

  //  print("incomplete s=%10g e=%10g s=%10g v=%10g g=%10g w=%10g\n", start, end, s, v, gammafn(s), w);

  return v + gammafn(s) * w * factor;
}
```
