## Supplementary material for "MoBPS - Modular Breeding Program Simulator": RandomFieldsUtils R-package: options.cc

```
/*
 Authors 
 Martin Schlather,


 Copyright (C) 2016 -- 2017  Martin Schlather

This program is free software; you can redistribute it and/or
modify it under the terms of the GNU General Public License
as published by the Free Software Foundation; either version 3
of the License, or (at your option) any later version.

This program is distributed in the hope that it will be useful,
but WITHOUT ANY WARRANTY; without even the implied warranty of
MERCHANTABILITY or FITNESS FOR A PARTICULAR PURPOSE.  See the
GNU General Public License for more details.
 
You should have received a copy of the GNU General Public License
along with this program; if not, write to the Free Software
Foundation, Inc., 59 Temple Place - Suite 330, Boston, MA  02111-1307, USA.  
*/

#include "Basic_utils.h" // must be before anything else
#ifdef DO_PARALLEL
#include <omp.h>
#endif
#include <unistd.h>
#include "General_utils.h"
#include "kleinkram.h"
#include "zzz_RandomFieldsUtils.h"

#define PLverbose 2

// IMPORTANT: all names of general must have at least 3 letters !!!
const char *basic[basicN] =
  { "printlevel","cPrintlevel", "seed", "cores", "skipchecks",
    "asList", "verbose", "kahanCorrection", "helpinfo", "warn_unknown_option"};

const char * solve[solveN] = 
  { "use_spam", "spam_tol", "spam_min_p", "svdtol", "eigen2zero",
    "solve_method", "spam_min_n", "spam_sample_n", "spam_factor", "spam_pivot",
    "max_chol", "max_svd", "pivot",
    "pivot_idx", // dynamic parameter
    "pivot_relerror", "pivot_max_deviation", "pivot_max_reldeviation",
    "det_as_log", "pivot_actual_size", "pivot_check"
    //, "tmp_delete"
  };


#define ownprefixN 2
const char * ownprefixlist[ownprefixN] = {"basic", "solve"};
const char **ownall[ownprefixN] = {basic, solve};
int ownallN[ownprefixN] = {basicN, solveN};


int PL = C_PRINTLEVEL,
  CORES = 1;

utilsparam GLOBAL = {
  basic_START,
  solve_START
};


//#if defined(unix) || defined(__unix__) || defined(__unix)
#if defined (__unix__) || (defined (__APPLE__) && defined (__MACH__))
int numCPU = sysconf(_SC_NPROCESSORS_ONLN);
#else
int numCPU = MAXINT;
#endif


void setparameterUtils(int i, int j, SEXP el, char name[LEN_OPTIONNAME], 
		       bool isList, int VARIABLE_IS_NOT_USED local) {
#ifdef DO_PARALLEL
  if (local != isGLOBAL) ERR1("Options specific to RandomFieldsUtils, here '%.50s', can be set only via 'RFoptions' outside any parallel code.", name);
#endif
  //printf("i=%d j=%d\n", i, j);
  utilsparam *options = &GLOBAL; 
  switch(i) {
  case 0: {// general
    basic_param *gp = &(options->basic);
    switch(j) {
    case 0: {  // general options
      int threshold = 1000; //PL_ERRORS;
      gp->Rprintlevel = INT;
      if (gp->Rprintlevel > threshold) gp->Rprintlevel = threshold;
      PL = gp->Cprintlevel = gp->Rprintlevel + PLoffset;
    }
      break;
    case 1: PL = gp->Cprintlevel = INT + PLoffset ;
      break;
    case 2: gp->seed = Integer(el, name, 0, true); break;
    case 3: gp->cores = POSINT; 
      if (gp->cores > numCPU) {
	WARN1("number of 'cores' is set to %d", numCPU);
	gp->cores = numCPU;
      }
#ifndef DO_PARALLEL      
      if (gp->cores != 1) {
	gp->cores = 1;
	PRINTF("The system does not allow for OpenMP: the value 1 is kept for 'cores'.");
      }										       
#endif
      break;
    case 4: gp->skipchecks = LOGI;    break;
    case 5: gp->asList = LOGI; break;
    case 6 : if (!isList) {
	PL = gp->Cprintlevel = gp->Rprintlevel = 1 + (LOGI * (PLverbose - 1));
      }
      break;
    case 7: gp->kahanCorrection = LOGI; break;
    case 8: gp->helpinfo = LOGI; break;
    case BASIC_WARN_OPTION: {
	int v = INT;
	if (v > -3) gp->warn_unknown_option = MIN(v, 3);
	else if (gp->warn_unknown_option == WARN_UNKNOWN_OPTION_DEFAULT) {
	  v += WARN_UNKNOWN_OPTION;
	  gp->warn_unknown_option = MIN(v, 3);
	} // else idle (respect the changing of the user)
      }
	break;

   default: BUG;
    }}
    break;
 
 case 1: {
   //   printf("name = %.50s %d\n", name, j);
   
    solve_param *so = &(options->solve);
    switch(j) {
    case 0: so->sparse = USRLOG; break; // USRLOGRELAXED??
    case 1: so->spam_tol = POS0NUM; break;      
    case 2: so->spam_min_p = POS0NUM; break;      
    case SOLVE_SVD_TOL: so->svd_tol = POS0NUM; break;        
    case 4: so->eigen2zero = POS0NUM; break;        
    case 5:
      GetName(el, name, InversionNames, nr_user_InversionMethods,
		    (int) NoInversionMethod, (int) NoFurtherInversionMethod, 
		    (int *)so->Methods, SOLVE_METHODS);
      break;
    case 6: so->spam_min_n = POSINT; break;      
    case 7: so->spam_sample_n = POSINT; break;      
    case 8: so->spam_factor = POSINT; break;      
    case 9: so->pivotsparse = POSINT; 
      if (so->pivotsparse > 2) so->pivotsparse = PIVOT_NONE;
      break;    
    case 10:
      //      printf("max chol = %d\n",  so->max_chol);
      so->max_chol = POSINT;
      //      printf("X max chol = %d\n",  so->max_chol);
       break;      
    case 11: so->max_svd = POS0INT; break;    
      //    case 11: so->tmp_delete = LOGI; break;    
    case 12: so->pivot = POS0INT;
      if (so->pivot > PIVOTLAST) so->pivot = PIVOT_UNDEFINED;
      break;    
    case 13: if (!isList) {
      int len = length(el);
      if (len == 0) {
	if (so->pivot_idx_n > 0) FREE(so->pivot_idx);
      } else {
	if (so->pivot_idx_n != len) {
	  FREE(so->pivot_idx);
	  so->pivot_idx = (int*) MALLOC(len * sizeof(int));
	}
	for (int L=0; L<len; L++) so->pivot_idx[L] = Integer(el, name, L);
      }
      so->pivot_idx_n = len;     
    }
      break; 
    case 14: so->pivot_relerror = POS0NUM; break;    
    case 15: so->max_deviation = POSNUM; break;    
    case 16: so->max_reldeviation = POS0NUM; break;    
    case 17: so->det_as_log = LOGI; break;    
    case 18: so->actual_size = POS0NUM; break;    
    case 19: so->pivot_check = USRLOG; break;    
   default: BUG;
    }}
    break;
    
  default: BUG;
  }

    }


void getparameterUtils(SEXP sublist, int i,
		       int VARIABLE_IS_NOT_USED local) {
  //  printf("hier %d\n", i);
  int  k = 0;
    //#define ADD(ELT) {printf(#ELT"\n");SET_VECTOR_ELT(sublist[i], k++, ELT);}
#ifdef DO_PARALLEL
  //  if (local != isGLOBAL) ERR("Options specific to RandomFieldsUtils can be obtained only on a global level and outside any parallel code.");
#endif  
  utilsparam *options = &GLOBAL; 
  switch(i) {
  case 0 : {
    //    printf("OK %d\n", i);
    basic_param *p = &(options->basic);
    ADD(ScalarInteger(p->Rprintlevel));    
    ADD(ScalarInteger(p->Cprintlevel - PLoffset));
    ADD(ScalarInteger(p->seed));    
    ADD(ScalarInteger(p->cores));    
    ADD(ScalarLogical(p->skipchecks));
    ADD(ScalarLogical(p->asList));   
    ADD(ScalarLogical(p->Rprintlevel >= PLverbose));
    ADD(ScalarLogical(p->kahanCorrection));   
    ADD(ScalarLogical(p->helpinfo));    
    ADD(ScalarInteger(p->warn_unknown_option));    
   }
    break;
  
  case 1 : {
    solve_param *p = &(options->solve);
    //    printf("sparse user interface %d %d %d\n", p->sparse, NA_LOGICAL, NA_INTEGER);
    ADD(ExtendedBooleanUsr(p->sparse));
    //    printf("A\n");
    ADD(ScalarReal(p->spam_tol));    
    ADD(ScalarReal(p->spam_min_p));    
    ADD(ScalarReal(p->svd_tol)); 
    ADD(ScalarReal(p->eigen2zero));    
    SET_VECTOR_ELT(sublist, k++,
		   String((int*) p->Methods, InversionNames, SOLVE_METHODS,
			  (int) NoFurtherInversionMethod));	
    ADD(ScalarInteger(p->spam_min_n));    
    ADD(ScalarInteger(p->spam_sample_n));    
    ADD(ScalarInteger(p->spam_factor));    
    ADD(ScalarInteger(p->pivotsparse));    
    ADD(ScalarInteger(p->max_chol));    
    ADD(ScalarInteger(p->max_svd)); 
    ADD(ScalarInteger(p->pivot));
    //if (true)
    SET_VECTOR_ELT(sublist, k++,
			     Int(p->pivot_idx, p->pivot_idx_n, p->pivot_idx_n));
    //  else ADD(ScalarInteger(NA_INTEGER));
    //    ADD(ScalarLogical(p->tmp_delete));
    ADD(ScalarReal(p->pivot_relerror));    
    ADD(ScalarReal(p->max_deviation));    
    ADD(ScalarReal(p->max_reldeviation));    
    ADD(ScalarLogical(p->det_as_log));
    ADD(ScalarInteger(p->actual_size));
    ADD(ExtendedBooleanUsr(p->pivot_check));
  }
    break;
  default : BUG;
  }
  //printf("EE A\n");
}

void delparameterUtils(int VARIABLE_IS_NOT_USED local) {
#ifdef DO_PARALLEL
  if (local != isGLOBAL) RFERROR("'pivot_idx' cannot be freed on a local level");
#endif  
  utilsparam *options = &GLOBAL;
  FREE(options->solve.pivot_idx);
}

void getErrorString(errorstring_type errorstring){
#ifdef DO_PARALLEL
  STRCPY(errorstring, "error occurred in package RandomFieldsUtils");
#else
  strcopyN(errorstring, ERRORSTRING, MAXERRORSTRING);
#endif  
}

void setErrorLoc(errorloc_type VARIABLE_IS_NOT_USED errorloc){
#ifndef DO_PARALLEL
  strcopyN(ERROR_LOC, errorloc, nErrorLoc);
#endif  
}
```
