## Supplementary material for "MoBPS - Modular Breeding Program Simulator": RandomFieldsUtils R-package: own.cc

```
/*
 Authors 
 Martin Schlather,

 Copyright (C) 2015 -- 2017 Martin Schlather

This program is free software; you can redistribute it and/or
modify it under the terms of the GNU General Public License
as published by the Free Software Foundation; either version 3
of the License, or (at your option) any later version.

This program is distributed in the hope that it will be useful,
but WITHOUT ANY WARRANTY; without even the implied warranty of
MERCHANTABILITY or FITNESS FOR A PARTICULAR PURPOSE.  See the
GNU General Public License for more details.

You should have received a copy of the GNU General Public License
along with this program; if not, write to the Free Software
Foundation, Inc., 59 Temple Place - Suite 330, Boston, MA  02111-1307, USA.  
*/

//#include "Basic_utils.h"  // must be before anything else
#include "RandomFieldsUtils.h"  // must be before anything else
#ifdef DO_PARALLEL
#include <omp.h>
#endif
#include <R.h>
#include <Rinternals.h>
#include "General_utils.h"
#include "own.h"
#include "Utils.h"
#include "intrinsics.h"
#include "zzz_RandomFieldsUtils.h" // always last


// local
#ifdef DO_PARALLEL
#else
char ERRMSG[LENERRMSG], MSG[LENERRMSG], MSG2[LENERRMSG];
errorloc_type ERROR_LOC="";
errorstring_type ERRORSTRING;
#endif

SEXP loadRandomFieldsUtils() {
  attachRFoptions(ownprefixlist, ownprefixN,
		  ownall, ownallN,
  		  setparameterUtils, NULL,
		  getparameterUtils,
		  delparameterUtils,
		  0, true);
  return R_NilValue;
}


SEXP attachRandomFieldsUtils() {
#define AVX2_USED true
#define AVX_USED true
  //#define SSE4_USED true
#define SSSE3_USED false
#define SSE2_USED true
#define SSE_USED false
  
  ReturnAttachMessage(RandomFieldsUtils, false);  
}

SEXP detachRandomFieldsUtils(){
  detachRFoptions(ownprefixlist, ownprefixN);
  freeGlobals();
  return R_NilValue;
}
```
