## Supplementary material for "MoBPS - Modular Breeding Program Simulator": RandomFieldsUtils R-package: own.h

```
/*
 Authors 
 Martin Schlather,


 Copyright (C) 2015 -- 2017 Martin Schlather

This program is free software; you can redistribute it and/or
modify it under the terms of the GNU General Public License
as published by the Free Software Foundation; either version 3
of the License, or (at your option) any later version.

This program is distributed in the hope that it will be useful,
but WITHOUT ANY WARRANTY; without even the implied warranty of
MERCHANTABILITY or FITNESS FOR A PARTICULAR PURPOSE.  See the
GNU General Public License for more details.
 
You should have received a copy of the GNU General Public License
along with this program; if not, write to the Free Software
Foundation, Inc., 59 Temple Place - Suite 330, Boston, MA  02111-1307, USA.  
*/


#ifndef rfutils_own_H
#define rfutil_own_H 1
#include "Options_utils.h"

void setparameterUtils(int i, int j, SEXP el, char name[200], bool isList, int local);
void getparameterUtils(SEXP sublist, int i, int local);
void delparameterUtils(int local);
void set_num_threads();


extern utilsparam GLOBAL;
#define ownprefixN 2
extern const char * ownprefixlist[ownprefixN],
  **ownall[ownprefixN];
extern int ownallN[ownprefixN];

#define HELPINFO(M) if (GLOBAL.basic.helpinfo) { PRINTF("%s\n(Note that you can unable this information by 'RFoptions(helpinfo=FALSE)'.)\n", M); } //

#endif
```
