## Supplementary material for "MoBPS - Modular Breeding Program Simulator": RandomFieldsUtils R-package: RandomFieldsUtils.h

```
/*
 Authors 
 Martin Schlather,


 Copyright (C) 2015 -- 2017 Martin Schlather

This program is free software; you can redistribute it and/or
modify it under the terms of the GNU General Public License
as published by the Free Software Foundation; either version 3
of the License, or (at your option) any later version.

This program is distributed in the hope that it will be useful,
but WITHOUT ANY WARRANTY; without even the implied warranty of
MERCHANTABILITY or FITNESS FOR A PARTICULAR PURPOSE.  See the
GNU General Public License for more details.

You should have received a copy of the GNU General Public License
along with this program; if not, write to the Free Software
Foundation, Inc., 59 Temple Place - Suite 330, Boston, MA  02111-1307, USA.  
*/


#ifndef RFutils_public_H
#define RFutils_public_H 1

#include <R.h>
#include <Rinternals.h>
#include <R_ext/Rdynload.h>
#include "Basic_utils.h"

#ifdef __cplusplus
extern "C" {
#endif 
  void R_init_RandomFieldsUtils(DllInfo *info);
  void R_unload_RandomFieldsUtils(DllInfo *info);

  SEXP struve(SEXP X, SEXP Nu, SEXP Factor_Sign, SEXP Expscaled);
  SEXP I0ML0(SEXP X);
  SEXP gaussr(SEXP X, SEXP Derivative); 
  SEXP WMr(SEXP X, SEXP Nu, SEXP Derivative, SEXP Factor);
  SEXP logWMr(SEXP X, SEXP Nu1, SEXP Nu2, SEXP Factor);

  SEXP SolvePosDef(SEXP M, SEXP rhs, SEXP logdet);
  SEXP Chol(SEXP M);
  
  SEXP RFoptions(SEXP options);

  SEXP loadRandomFieldsUtils();
  SEXP attachRandomFieldsUtils();
  SEXP detachRandomFieldsUtils();

  SEXP sortX(SEXP Data, SEXP From, SEXP To, SEXP NAlast);
  SEXP orderX(SEXP Data, SEXP From, SEXP To, SEXP NAlast);

  void sleepMicro(int *micro);
  void sleepMilli(int *milli);
  void hostname(char **h, int *i);
  void pid(int *i);
  SEXP getChar();
  SEXP colMaxs(SEXP M);
  SEXP rowMeansX(SEXP M, SEXP Factor);
  SEXP rowProd(SEXP M);
  SEXP chol2mv(SEXP Chol, SEXP N);
  SEXP tcholRHS(SEXP C, SEXP RHS);
  SEXP DivByRow(SEXP M, SEXP V);
  SEXP quadratic(SEXP x, SEXP A);
  SEXP dbinorm(SEXP X, SEXP Sigma);
  SEXP dotXV(SEXP M, SEXP V);
  void Ordering(double *d, int *len, int *dim, int *pos);


  
#ifdef SCHLATHERS_MACHINE
  //  SEXP scalarX(SEXP x, SEXP y, SEXP mode);
  //  SEXP brdomain(SEXP Srf, SEXP Sgamma, SEXP Sinstances, SEXP Smaxn);
#endif

#ifdef __cplusplus
}
#endif


#endif 
 
```
