## Supplementary material for "MoBPS - Modular Breeding Program Simulator": RandomFieldsUtils R-package: RFoptions.cc

```
/*
 Authors 
 Martin Schlather,


 Copyright (C) 2016 -- 2017 Martin Schlather

This program is free software; you can redistribute it and/or
modify it under the terms of the GNU General Public License
as published by the Free Software Foundation; either version 3
of the License, or (at your option) any later version.

This program is distributed in the hope that it will be useful,
but WITHOUT ANY WARRANTY; without even the implied warranty of
MERCHANTABILITY or FITNESS FOR A PARTICULAR PURPOSE.  See the
GNU General Public License for more details.
 
You should have received a copy of the GNU General Public License
along with this program; if not, write to the Free Software
Foundation, Inc., 59 Temple Place - Suite 330, Boston, MA  02111-1307, USA.  
*/

#include "RandomFieldsUtils.h"
#include "General_utils.h"
#include "own.h"
#include "zzz_RandomFieldsUtils.h"


typedef struct {
  int ListNr, i;
} getlist_type;


void setpDef(int VARIABLE_IS_NOT_USED  i, 
	     int VARIABLE_IS_NOT_USED  j, 
	     SEXP VARIABLE_IS_NOT_USED  el,
	     char VARIABLE_IS_NOT_USED  name[LEN_OPTIONNAME], 
	     bool VARIABLE_IS_NOT_USED  isList,
	     int VARIABLE_IS_NOT_USED local) {
  BUG;
}
void getpDef(SEXP VARIABLE_IS_NOT_USED  sublist, int VARIABLE_IS_NOT_USED i,
	     int VARIABLE_IS_NOT_USED local) {
  BUG;
}


#define MAXNLIST 5
int NList = 0; // originally 1
int nbasic_options = 0,
  AllprefixN[MAXNLIST] = {ownprefixN, 0, 0, 0, 0},
  *AllallN[MAXNLIST] = {ownallN, NULL, NULL, NULL, NULL};
const char  *basic_options[MAXNLIST] = {ownprefixlist[1], NULL, NULL, NULL},
  **Allprefix[MAXNLIST] = {ownprefixlist, NULL, NULL, NULL, NULL},
  ***Allall[MAXNLIST] = { ownall, NULL, NULL, NULL, NULL};
setparameterfct setparam[MAXNLIST] = 
  {setparameterUtils, setpDef, setpDef, setpDef, setpDef};
getparameterfct getparam[MAXNLIST] = 
  {getparameterUtils, getpDef, getpDef, getpDef, getpDef};
finalsetparameterfct finalparam[MAXNLIST] = { NULL, NULL, NULL, NULL, NULL };
deleteparameterfct delparam[MAXNLIST] = { NULL, NULL, NULL, NULL, NULL };


void hintVariable(char *name) {
  if (GLOBAL.basic.warn_unknown_option > 0) {
    PRINTF("'%s' is considered as a variable (not as an option).\n", name);
    if (GLOBAL.basic.helpinfo) {
      PRINTF("(This hint can be turned off by 'RFoptions(%s=-2)",
	     basic[BASIC_WARN_OPTION]);
    }
  }
}

void setparameter(SEXP el, char *prefix, char *mainname, bool isList,
		  getlist_type *getlist, int local) {  
  int 
    j = NOMATCHING,
    i = NOMATCHING,
    ListNr = NOMATCHING;
  char name[LEN_OPTIONNAME];

  SPRINTF(name, "%.50s%.50s%.50s", prefix, STRLEN(prefix)==0 ? "" : ".", mainname);

  if (STRCMP(prefix, "")) {
    for (ListNr=0; ListNr<NList; ListNr++) {
      i = Match(prefix, Allprefix[ListNr], AllprefixN[ListNr]);
      if (i != NOMATCHING) break;
    }
    if (i == NOMATCHING) ERR1("option prefix name '%.50s' not found.", prefix); 
    if (i < 0 || STRCMP(prefix, Allprefix[ListNr][i])) {
      for (int k=ListNr + 1; k < NList; k++) {
	int ii = Match(prefix, Allprefix[ListNr], AllprefixN[ListNr]);
	if (ii == NOMATCHING) continue;
	i = MULTIPLEMATCHING;
	if (ii >= 0 && STRCMP(prefix, Allprefix[k][ii])==0) {
	  ListNr = k;
	  i = ii;
	  break;
	} // ii >0
      } // for k
      if (i == MULTIPLEMATCHING) 
	ERR1("option prefix name '%.50s' is ambiguous.", prefix);   
    } // prefix == List

    j = Match(mainname, Allall[ListNr][i], AllallN[ListNr][i]);
      
  } else { // (i==0), no prefix given
#define MinNameLength 3
    for (ListNr=0; ListNr<NList; ListNr++) {
      int trueprefixN = AllprefixN[ListNr];
      for (i=0; i < trueprefixN; i++) {	
	j = Match(mainname, Allall[ListNr][i], AllallN[ListNr][i]);
	if (j != NOMATCHING) break;     
      }
      if (j != NOMATCHING) break;
    }
    if (j==NOMATCHING) {
      switch(GLOBAL.basic.warn_unknown_option) {
      case WARN_UNKNOWN_OPTION_NONE : return;
      case WARN_UNKNOWN_OPTION_CAPITAL : case -WARN_UNKNOWN_OPTION_CAPITAL :
	if (name[0] >= 'A' && name[1] <= 'Z') {
	  hintVariable(name);
	  return;
	}
	break;
      case WARN_UNKNOWN_OPTION_SINGLE : case -WARN_UNKNOWN_OPTION_SINGLE :
	if (STRLEN(name) == 1 && name[0] >= 'A' && name[1] <= 'Z') {
	  hintVariable(name);
	  return;
	}
	break;
      case WARN_UNKNOWN_OPTION_ALL :
      default :
	ERR1("Unknown option '%.50s'.", name);
      }
      return;
    }
    
    // printf("j=%d  %.50s\n", j,  j >=0 ? Allall[ListNr][i][j] : "multi");
    
    if (j < 0  || STRCMP(mainname, Allall[ListNr][i][j])) {
      int starti = i + 1;
      for (int k = ListNr; k<NList; k++, starti=0) {
	int tpN = AllprefixN[k];
	for (int ii=starti; ii < tpN; ii++) {	
	  int jj = Match(mainname, Allall[k][ii], AllallN[k][ii]);
	  if (jj == NOMATCHING) continue;
	  
	  //  printf("listnr=%d %.50s jj=%d %.50s\n", ListNr, Allall[ListNr][i][j], 
	  //	 jj, jj < 0 ? "none" : Allall[k][ii][jj]);
	  j = MULTIPLEMATCHING;
	  if (jj >= 0 && STRCMP(mainname, Allall[k][ii][jj])==0) {
	    ListNr = k;
	    i = ii;
	    j = jj;
	    break;
	  } // jj
	} // for ii
	if (j >= 0) break;
      } // for k
    } // if j < 0 || !=
  } // no prefix given

  if (j<0) ERR1("Multiple partial matching for option '%.50s'.", name); 
 
  if (getlist != NULL) {
    int k=0;
    while((getlist[k].ListNr != ListNr || getlist[k].i != i)
	  && getlist[k].ListNr >= 0) k++;
    if (getlist[k].ListNr < 0)
      ERR2("Option '%.50s' not allowed for this call.\n   In case you really need this option, use the command 'RFoption(%.50s=...)'", mainname, mainname);
  }
  
  setparam[ListNr](i, j, el, name, isList, local); 
}


SEXP getRFoptions(int ListNr, int i, int local) {
  SEXP sublist, subnames;
  int elmts = AllallN[ListNr][i];
  PROTECT(sublist = allocVector(VECSXP, elmts));
  PROTECT(subnames = allocVector(STRSXP, elmts));
  for (int k=0; k<elmts; k++) {
    //    printf("getopt %d k=%d elmnts=%d %d %.50s\n", i, k, elmts, ListNr, Allall[ListNr][i][k]);
    SET_STRING_ELT(subnames, k, mkChar(Allall[ListNr][i][k])); 
  }
  getparam[ListNr](sublist, i, local);
  setAttrib(sublist, R_NamesSymbol, subnames);
  UNPROTECT(2);
  return sublist;
}

SEXP getRFoptions(int local) {
  SEXP list, names;
 
  int  prefixN, totalN, i, ListNr,
    itot = 0;
  for (totalN=ListNr=0; ListNr<NList; ListNr++) {
    prefixN = AllprefixN[ListNr];
    for (i=0; i<prefixN; i++) {
      totalN += STRCMP(Allprefix[ListNr][i], OBSOLETENAME) != 0;
    }
  }

  PROTECT(list = allocVector(VECSXP, totalN));
  PROTECT(names = allocVector(STRSXP, totalN));

  for (ListNr =0; ListNr<NList; ListNr++) {
    //    printf("ListNr %d\n", ListNr);
    prefixN = AllprefixN[ListNr];    
    for (i=0; i<prefixN; i++) {
      if (STRCMP(Allprefix[ListNr][i], OBSOLETENAME) == 0) continue;
      SET_VECTOR_ELT(list, itot, getRFoptions(ListNr, i, local));
      SET_STRING_ELT(names, itot, mkChar(Allprefix[ListNr][i]));
      itot ++;
    } 
  }

  setAttrib(list, R_NamesSymbol, names);   
  UNPROTECT(2);

  return list;
}


void getListNr(bool save, int t, int actual_nbasic, SEXP which,
	      getlist_type *getlist,
	      int *Nr, int *idx // output
	      ){
  int i, ListNr;
  const char *z;
  if (save && t < nbasic_options) z = basic_options[t];
  else z = (char*) CHAR(STRING_ELT(which, t - actual_nbasic));
  for (ListNr=0; ListNr<NList; ListNr++) {
    int prefixN = AllprefixN[ListNr];
    for (i=0; i<prefixN; i++)
      if (STRCMP(Allprefix[ListNr][i], z) == 0) break;
    if (i < prefixN) break;
  }
  if (ListNr >= NList) ERR("unknown value for 'GETOPTIONS'");
  if (getlist != NULL) {
    getlist[t].ListNr = ListNr;
    getlist[t].i = i;
  }
  *Nr = ListNr;
  *idx = i;
}

  
SEXP getRFoptions(SEXP which, getlist_type *getlist, bool save, int local) {
  
  int ListNr, idx,
    actual_nbasic = nbasic_options * save,
    totalN = length(which) + actual_nbasic;  

  if (totalN == 0) return R_NilValue;
  if (totalN == 1) {
    getListNr(save, 0, actual_nbasic, which, getlist, &ListNr, &idx);
    return getRFoptions(ListNr, idx, local);
  }

  SEXP list, names;
  PROTECT(list = allocVector(VECSXP, totalN));
  PROTECT(names = allocVector(STRSXP, totalN));
  for (int t=0; t<totalN; t++) {
    getListNr(save, t, actual_nbasic,  which, getlist, &ListNr, &idx);
    SET_VECTOR_ELT(list, t, getRFoptions(ListNr, idx, local));
    SET_STRING_ELT(names, t, mkChar(Allprefix[ListNr][idx]));
  }
  setAttrib(list, R_NamesSymbol, names);     
  UNPROTECT(2);
  return list;
}


void splitAndSet(SEXP el, char *name, bool isList, getlist_type *getlist,
		 int local) {
  int i, len;
  char prefix[LEN_OPTIONNAME / 2], mainname[LEN_OPTIONNAME / 2];   
  //  printf("splitandset\n");
  len = STRLEN(name);
  for (i=0; i < len && name[i]!='.'; i++);
  if (i==0) { ERR1("argument '%.50s' not valid\n", name); }
  if (i==len) {
    STRCPY(prefix, "");
    strcopyN(mainname, name, LEN_OPTIONNAME / 2);
  } else {
    strcopyN(prefix, name, MIN(i + 1, LEN_OPTIONNAME / 2));
    strcopyN(mainname, name+i+1, MIN(STRLEN(name) - i, LEN_OPTIONNAME / 2) );
  }

  // 
  //  printf("i=%d %d %.50s %.50s\n", i, len, prefix, mainname);
  setparameter(el, prefix, mainname, isList && GLOBAL.basic.asList, getlist,
	       local);
  //   printf("ende\n");
}


SEXP RFoptions(SEXP options) {
  int i, j, lenlist, lensub;
  SEXP el, list, sublist, names, subnames,
     ans = R_NilValue;
  char *name, *pref;
  bool isList = false;
  int
    n_protect = 0,
    local = isGLOBAL;
  

    /* 
     In case of strange values of a parameter, undelete
     the comment for PRINTF
  */

  
  // printf("start %d %d\n", GLOBAL.basic.warn_unknown_option, options == R_NilValue);
  options = CDR(options); /* skip 'name' */

  // printf(" %d %d\n", GLOBAL.basic.warn_unknown_option, options == R_NilValue);
  if (options == R_NilValue) return getRFoptions(local); 

  
  //  printf("hierA %d\n", isNull(TAG(options)));
 
  if (isNull(TAG(options))) name = (char*) ""; // (char*)
  else name = (char*) CHAR(PRINTNAME(TAG(options)));
  
  //printf("hierB, %d\n", STRCMP(name, "LOCAL"));
  if (STRCMP(name, "LOCAL")==0) {
    el = CAR(options);
    local = INT;
    options = CDR(options); /* skip 'name' */
    if (isNull(TAG(options))) name = (char*) "";
    else name = (char*) CHAR(PRINTNAME(TAG(options)));
  }

  //printf("hier\n");
  
  if ((isList = STRCMP(name, "LIST")==0)) {   
    //   printf("isList\n");
    list = CAR(options);
    if (TYPEOF(list) != VECSXP)
      ERR1("'LIST' needs as argument the output of '%.50s'", RFOPTIONS);
    PROTECT(names = getAttrib(list, R_NamesSymbol));
    lenlist = length(list);
    for (i=0; i<lenlist; i++) {
      int len;
      pref = (char*) CHAR(STRING_ELT(names, i));  

      //     printf("%d %.50s\n", i, pref);

      sublist = VECTOR_ELT(list, i);
      len = STRLEN(pref);
      for (j=0; j < len && pref[j]!='.'; j++);
      if (TYPEOF(sublist) == VECSXP && j==len) { // no "."
	// so, general parameters may not be lists,
	// others yes
	lensub = length(sublist);
	PROTECT(subnames = getAttrib(sublist, R_NamesSymbol));
	for (j=0; j<lensub; j++) {
	  name = (char*) CHAR(STRING_ELT(subnames, j));


	  //    print("%d %d %.50s : %10g %10g\n", i, j, name,
	  //	     GLOBAL.gauss.exactness, GLOBAL.TBM.linesimustep);	  
	  //
	  //	  printf("  %d %d pref=%.50s name=%.50s\n", i, j, pref, name);


	  setparameter(VECTOR_ELT(sublist, j), pref, name, 
		       isList & GLOBAL.basic.asList, NULL, local);
	}
	UNPROTECT(1);
      } else {   
	splitAndSet(sublist, pref, isList, NULL, local);
      }
    }
    UNPROTECT(1);
    //    print("end1 %10g\n", GLOBAL.TBM.linesimufactor);
  } else {    
    getlist_type *getlist = NULL;
    bool save;
    if ((save = STRCMP(name, "SAVEOPTIONS") ==0 ) ||
	STRCMP(name, "GETOPTIONS")==0) {
      SEXP getoptions = CAR(options);
      options = CDR(options);
      if (options != R_NilValue) {
	//	printf("hier\n");
	int len = length(getoptions) + nbasic_options * save;
	getlist = (getlist_type *) MALLOC(sizeof(getlist_type) * (len + 1));
	getlist[len].ListNr = -1;
      }
      //     printf("l=%d\n", local);
      PROTECT(ans = getRFoptions(getoptions, getlist, save, local));
      n_protect++;
    }
    //    printf("iok %d\n", length(CAR(options)));
    for(i = 0; options != R_NilValue; i++, options = CDR(options)) {     
      //       printf("set opt i=%d\n", i);
      el = CAR(options);
      if (isNull(TAG(options))) name = (char*) "";
      else name = (char*) CHAR(PRINTNAME(TAG(options)));
      //      printf("AAxx %s %d %ld\n", name, isList);
      //      printf("%ld\n", REAL(el));
      splitAndSet(el, name, isList, getlist, local);
      //printf("yy %s %d\n", name, isList);
    }
    FREE(getlist);
    //    printf("end2\n");
  }


  //  printf("Nlist = %d %ld %ld\n", NList, ans, R_NilValue);
  for (i=0; i<NList; i++)
    if (finalparam[i] != NULL) {
      finalparam[i](local);
    }

  if (n_protect > 0) UNPROTECT(n_protect);
  
  GLOBAL.basic.asList = true;

  // printf("Ende\n");
  
  return(ans);
} 
 

int PLoffset = 0;
void attachRFoptions(const char **prefixlist, int N, 
		     const char ***all, int *allN,
		     setparameterfct set, finalsetparameterfct final,
		     getparameterfct get,
		     deleteparameterfct del,
		     int pl_offset, bool basicopt) {
  for (int ListNr=0; ListNr<NList; ListNr++) {    
    if (AllprefixN[ListNr] == N && 
	STRCMP(Allprefix[ListNr][0], prefixlist[0]) == 0) {
      if (PL > 0) {
	PRINTF("options starting with prefix '%s' have been already attached.",
		prefixlist[0]);
      }
      return;    
    }
  }
  if (basicopt) basic_options[nbasic_options++] = prefixlist[0];
  if (NList >= MAXNLIST) BUG;
  Allprefix[NList] = prefixlist;
  AllprefixN[NList] = N;
  Allall[NList] = all;
  AllallN[NList] = allN;
  setparam[NList] = set;
  finalparam[NList] = final;
  getparam[NList] = get;
  delparam[NList] = del;
  NList++;
  PLoffset = pl_offset;
  basic_param *gp = &(GLOBAL.basic);
  PL = gp->Cprintlevel = gp->Rprintlevel + PLoffset;
  CORES = gp->cores;
}


void detachRFoptions(const char **prefixlist, int N) {
  int ListNr;
  for (ListNr=0; ListNr<NList; ListNr++) {    
    if (AllprefixN[ListNr] == N && 
	STRCMP(Allprefix[ListNr][0], prefixlist[0]) == 0) break;
  }  
  if (ListNr >= NList) {
    ERR1("options starting with prefix '%.50s' have been already detached.",
	prefixlist[0]);
  }

  if (delparam[ListNr] != NULL) delparam[ListNr](isGLOBAL);
  
  int i;
  for (i=0; i<nbasic_options ; i++)
    if (STRCMP(basic_options[i], prefixlist[0]) == 0) break;
  for (i++ ; i < nbasic_options; i++) basic_options[i - 1] = basic_options[i];
  
  for (ListNr++; ListNr<NList; ListNr++) {    
    Allprefix[ListNr - 1] = Allprefix[ListNr];
    AllprefixN[ListNr - 1] =  AllprefixN[ListNr];
    Allall[ListNr - 1] = Allall[ListNr];
    AllallN[ListNr - 1] = AllallN[ListNr];
    setparam[ListNr - 1] = setparam[ListNr];
    finalparam[ListNr - 1] = finalparam[ListNr];
    getparam[ListNr - 1] = getparam[ListNr];
  }

  NList--;
  if (NList <= 1) PLoffset = 0;
}

void getUtilsParam(utilsparam **global) { 
  // printf("GLO %ld\n", &GLOBAL);
  *global = &GLOBAL; 
}
```
