## Supplementary material for "MoBPS - Modular Breeding Program Simulator": RandomFieldsUtils R-package: scalar.cc

```
/* 
 Authors
 Martin Schlather,

 Collection of system specific auxiliary functions

 Copyright (C) 2001 -- 2017 Martin Schlather, 

This program is free software; you can redistribute it and/or
modify it under the terms of the GNU General Public License
as published by the Free Software Foundation; either version 3
of the License, or (at your option) any later version.

This program is distributed in the hope that it will be useful,
but WITHOUT ANY WARRANTY; without even the implied warranty of
MERCHANTABILITY or FITNESS FOR A PARTICULAR PURSE.  See the
GNU General Public License for more details.

You should have received a copy of the GNU General Public License
along with this program; if not, write to the Free Software
Foundation, Inc., 59 Temple Place - Suite 330, Boston, MA  02111-1307, USA.
*/

/*

Makefile must be:

PKG_LIBS =  $(SHLIB_OPENMP_CXXFLAGS) $(LAPACK_LIBS) $(BLAS_LIBS) $(FLIBS)  -march=native  -mssse3 
PKG_CXXFLAGS =  $(SHLIB_OPENMP_CXXFLAGS)  -march=native -mssse3 

 */

#define BUG assert(false);

#include <assert.h>
#include "kleinkram.h"
#include "scalar.h"
#include "intrinsics.h"
#include "Basic_utils.h"
#include "errors_messages.h"
#include "zzz_RandomFieldsUtils.h"


#define Nmodi 9
name_type modi = { "1x1", "2x2", "4x4", "8x8", "near", "simple", "precise", "kahan", "1x1p"};


typedef unsigned int uint32;


#define size 8
#define vectorlen (256 / (size * 8))
#define repet 8
#define atonce (vectorlen * repet)
#define SET_0(NR) sum##NR = ZERODOUBLE
#define P_0(NR) prod##NR = ZERODOUBLE
#define SUMUP(NR, nr) sum##NR = ADDDOUBLE(sum##NR, sum##nr)
#define ADDN(NR)							\
  prod##NR = MULTDOUBLE(LOADuDOUBLE(x + i + NR * vectorlen),		\
			LOADuDOUBLE(y + i + NR * vectorlen));		\
  sum##NR = ADDDOUBLE(sum##NR, prod##NR) 


#if (7 != repet - 1)
  wrong repet length
#endif
#if (3 != vectorlen - 1)
  wrong vector length
#endif

  
#ifdef AVX

#ifdef FMA_AVAILABLE
#define ADDF(NR) \
  sum##NR = _mm256_fmadd_pd(LOADuDOUBLE(x + i + NR * vectorlen),\
			    LOADuDOUBLE(y + i + NR * vectorlen), sum##NR)
double avx_scalarprodDfma(double * x, double * y, int len) {
 int i = 0,
    lenM = len - (atonce - 1);  
   __m256d SET_0(0);
   double *D  = (double *) &sum0;

  if (len >= atonce) {
    __m256d SET_0(1), SET_0(2), SET_0(3), SET_0(4), SET_0(5), SET_0(6),SET_0(7);
   for (; i < lenM; i += atonce) { 
     ADDF(0); ADDF(1); ADDF(2); ADDF(3); ADDF(4); ADDF(5); ADDF(6); ADDF(7); 
   }
   SUMUP(0, 1); SUMUP(2, 3); SUMUP(4, 5); SUMUP(6, 7);
   SUMUP(0, 2); SUMUP(4, 6); SUMUP(0, 4);
  }
  lenM = len - vectorlen + 1;
  for (; i < lenM; i += vectorlen) { ADDF(0);  }
  double sum = D[0] + D[1] + D[2] + D[3];
  for (; i < len; i++) sum += x[i] * y[i];
  return sum;
}
#endif


double avx_scalarprodDnearfma(double * x, double * y, int len) {
  // deutlich genauer zum 0 tarif
  int i = 0,
     lenM = len - (atonce - 1);  
  __m256d SET_0(0), SET_0(1), SET_0(2), SET_0(3), SET_0(4), SET_0(5),
    SET_0(6),SET_0(7),
    P_0(0), P_0(1), P_0(2), P_0(3), P_0(4), P_0(5), P_0(6), P_0(7);
  double *D  = (double *) &sum0;

  if ( len >= atonce) {
    for (; i < lenM; i += atonce) {
      ADDN(0); ADDN(1); ADDN(2); ADDN(3); ADDN(4); ADDN(5); ADDN(6); ADDN(7); 
    }
    SUMUP(0, 1); SUMUP(2, 3); SUMUP(4, 5); SUMUP(6, 7);
    SUMUP(0, 2); SUMUP(4, 6); SUMUP(0, 4);
  }
  lenM = len - vectorlen + 1;
  for (; i < lenM; i += vectorlen) {  ADDN(0);  }
  double sum = D[0] + D[1] + D[2] + D[3];
  for (; i < len; i++) sum += x[i] * y[i];
  
  return sum;
}
 

#define ADDM(NR)								\
  prod0 = MULTDOUBLE(LOADuDOUBLE(x + i + NR * vectorlen),		\
			   LOADuDOUBLE(y + i + NR * vectorlen));		\
  sum0 = ADDDOUBLE(sum0, prod0)
double avx_scalarprodD(double * x, double * y, int len) {
  int i = 0,
    lenM = len - (atonce - 1);  
  __m256d SET_0(0), P_0(0);
   double *D  = (double *) &sum0;

  if ( len >= atonce) {
  for (; i < lenM; i += atonce) {
    ADDM(0); ADDM(1); ADDM(2); ADDM(3); ADDM(4); ADDM(5); ADDM(6); ADDM(7); 
    }
  }
  lenM = len - vectorlen + 1;
  for (; i < lenM; i += vectorlen) { ADDM(0); } 
  double sum = D[0] + D[1] + D[2] + D[3];
  for (; i < len; i++) sum += x[i] * y[i];
  return sum;
}

//#p r a g m a   o m p declare reduction(minabs : int :  omp_out = a bs(omp_in) > omp_out ? omp_out : a bs(omp_in)   initializer (omp_priv=LARGENUM)

#if defined OpenMP4
double avx_scalarprodDparallel(double * x, double * y, int len) {
   int i = 0,
    lenM = len - (atonce - 1);  
   __m256d SET_0(0), P_0(0);
   double *D  = (double *) &sum0;

#ifdef DO_PARALLEL
#pragma omp declare reduction(addpd:  __m256d:				\
			      omp_out = ADDDOUBLE(omp_out, omp_in))	\
			      initializer (omp_priv = ZERODOUBLE)
#endif
 
  if ( len >= atonce) {
#ifdef DO_PARALLEL
#pragma omp parallel for num_threads(CORES) reduction(addpd:sum0) schedule(dynamic, 100)
#endif
    for (i=0; i < lenM; i += atonce) {
      ADDM(0); ADDM(1); ADDM(2); ADDM(3); ADDM(4); ADDM(5); ADDM(6); ADDM(7); 
    }
  }
  lenM = len - vectorlen + 1;
  for (; i < lenM; i += vectorlen) { ADDM(0); }
  double sum = D[0] + D[1] + D[2] + D[3];
  for (; i < len; i++) sum += x[i] * y[i];
  return sum;
}
#endif


double avx_scalarprodDP(double * x, double * y, int len) {
  int i = 0,
     lenM = len - (atonce - 1);  
  __m256d SET_0(0), SET_0(1), P_0(0);
   double *D  = (double *) &sum1;
  if ( len >= atonce) {
    for (; i < lenM; ) {
      int lenMM = i + vectorlen * (repet * 10 + 1);
      if (lenMM > lenM) lenMM = lenM;
      sum0 = MULTDOUBLE(LOADuDOUBLE(x + i), LOADuDOUBLE(y + i));
      i += vectorlen;
      for (; i < lenMM; i += atonce) {
	ADDM(0); ADDM(1); ADDM(2); ADDM(3); ADDM(4); ADDM(5); ADDM(6); ADDM(7); 
      }
      sum1 = ADDDOUBLE(sum0, sum1);
    }
  }
  
 lenM = len - vectorlen + 1;
 for (; i < lenM; i += vectorlen) { 
    prod0 = MULTDOUBLE(LOADuDOUBLE(x + i), LOADuDOUBLE(y + i));
    sum1 = ADDDOUBLE(sum1, prod0);
  }
  double sum = D[0] + D[1] + D[2] + D[3];
  for (; i < len; i++) sum += x[i] * y[i];
  return sum;
}


#define ADDK(NR)							\
  prod0 = MULTDOUBLE(LOADuDOUBLE(x + i + NR * vectorlen),		\
		     LOADuDOUBLE(y + i + NR * vectorlen));		\
  sum2 = SUBDOUBLE(prod0, sum1);					\
  sum3 = ADDDOUBLE(sum0, sum2);						\
  sum1 = SUBDOUBLE(sum3, sum0);						\
  sum0 = sum3;								\
  sum1 = SUBDOUBLE(sum1, sum2);
double avx_scalarprodDK(double * x, double * y, int len) {
  // Kahan
  int i = 0,
    lenM = len - (atonce - 1);  
  __m256d SET_0(0), // sum
    SET_0(1), 
    SET_0(2), // y
    SET_0(3), // t
    P_0(0),
    P_0(1);
  double *D  = (double *) &sum0;  
  if ( len >= atonce) {
    for (; i < lenM; i += atonce) {
      ADDK(0); ADDK(1); ADDK(2); ADDK(3); ADDK(4); ADDK(5); ADDK(6); ADDK(7);
    }
  }
  lenM = len - vectorlen + 1;
  for (; i < lenM; i += vectorlen) { ADDK(0); }
  sum0 = ADDDOUBLE(sum0, prod1);
  double sum = D[0] + D[1] + D[2] + D[3];
  
  for (; i < len; i++) sum += x[i] * y[i];
  return sum;
}

#endif
 
 
double scalarprod( double * v1, double * v2, int N){
  double *endv1 = v1 + N,
    sum = 0;
  for(; v1!= endv1; v1++, v2++) sum +=  v2[0] * v1[0];
  return sum;
}
 
 
double scalarprodP( double * v1, double * v2, int N){
  double 
    sum = 0.0;
#ifdef DO_PARALLEL
#pragma omp parallel for num_threads(CORES) if (N > 200) reduction(+:sum) schedule(dynamic, 100) 
#endif
  for(int i=0; i<=N; i++) sum += v2[i] * v1[i];
  return sum;
}
 
 
double scalarprod2by2( double * v1, double * v2, int N){
  double *endv1 = v1 + (N / 2) * 2,
    *end = v1 + N,
    sum = 0;
  for(; v1 < endv1; v1 += 2, v2 += 2) sum += v2[0] * v1[0] + v2[1] * v1[1];
  if (v1 < end) sum += v2[0] * v1[0]; 
  return sum;
}
 
 
double scalarprod4by4( double * v1, double * v2, int N){
  double*endv1 = v1 + (N / 4) * 4,
    *end = v1 + N,
    sum = 0;
  for(; v1 < endv1; v1 += 4, v2 += 4)
    sum += v2[0] * v1[0] + v2[1] * v1[1] + v2[2] * v1[2]+ v2[3] * v1[3];
  for(; v1 < end; v1++, v2++) sum += v2[0] * v1[0];        
  return sum;
}

 
double scalarprod8by8( double * v1, double * v2, int N){
  double
    *endv1 = v1 + (N / 8) * 8,
    *end = v1 + N,
    sum = 0.0;
  for(; v1 < endv1; v1 += 8, v2 += 8)
    sum += v2[0] * v1[0] + v2[1] * v1[1]+ v2[2] * v1[2] + v2[3] * v1[3] +
      v2[4] * v1[4] + v2[5] * v1[5]+ v2[6] * v1[6]+ v2[7] * v1[7];
  for(; v1 < end; v1++, v2++) sum +=  v2[0] * v1[0];        
  return sum;
}


double scalarprodPX( double * V1, double * V2, int N){
#define AtOnce 16
  double
    *endv1 = V1 + (N / AtOnce) * AtOnce,
    *end = V1 + N,
    sum = 0;
#ifdef DO_PARALLEL
#pragma omp parallel for num_threads(CORES) if (N > 200) reduction(+:sum) schedule(dynamic, 50) 
#endif
  for(double *v1=V1; v1 < endv1; v1 += AtOnce) {
    double *v2 = V2 + (V1 - v1);
    sum +=  v2[0] * v1[0] + v2[1] * v1[1]+ v2[2] * v1[2] + v2[3] * v1[3] +
      v2[4] * v1[4] + v2[5] * v1[5]+ v2[6] * v1[6]+ v2[7] * v1[7] +
      v2[8] * v1[8] + v2[9] * v1[9]+ v2[10] * v1[10] + v2[11] * v1[11] +
      v2[12] * v1[12] + v2[13] * v1[13]+ v2[14] * v1[14]+ v2[15] * v1[15];
  }
  double
    *v1 = V1 + (N / AtOnce),
    *v2 = V2 + (V1 - v1);
  for(; v1 < end; v1++, v2++) sum += v2[0] * v1[0];        
  return sum;
}


//bool pr = true;
double scalarX(double *x, double *y, int len, int n) {
  if (n < 0) {
  }
//  if (pr) { printf("mode = %d\n", n); pr = false; }
 // 0 : 7.9
// 1:  7.55
// 2: 7.8
// 3:7.58
//4: 7.5 
// 5: 7.4!
//6:7.4
//7: 7.9
// 8: "ewige" schleife
  //  printf("n=%d ", n);
  
  switch(n) {
  case 0 : return scalarprod(x, y, len);
  case SCALAR_BASE : return scalarprod2by2(x, y, len); 
  case 2 : return scalarprod4by4(x, y, len); 
  case 3 : return scalarprod8by8(x, y, len); 
#ifdef FMA_AVAILABLE
  case 4 : return avx_scalarprodDfma(x, y, len);
#endif    
#ifdef AVX
  case 5 : return avx_scalarprodDnearfma(x, y, len); 
  case SCALAR_AVX : return avx_scalarprodD(x, y, len); // best one kernel
  case 7 : return avx_scalarprodDP(x, y, len);  //best
  case SCALAR_KAHAN : return avx_scalarprodDK(x, y, len); // kahan
#else
  case 4: case 5: case 6: case 7: case 8 : return scalarprod2by2(x, y, len);
#endif
    
#ifdef DO_PARALLEL
  case SCALAR_AVX_PARALLEL :
#if defined AVX && defined OpenMP
    return avx_scalarprodDparallel(x, y, len);
#endif    
  case SCALAR_BASE_PARALLEL : return scalarprodP(x, y, len);// parallel, nicht-vectoriell
#else
  case SCALAR_AVX_PARALLEL :
#ifdef AVX
  return avx_scalarprodD(x, y, len);
#endif     
   case SCALAR_BASE_PARALLEL : return scalarprod2by2(x, y, len); 
#endif  
  default : ERR("method not available"); 
  }
  return RF_NAN;
}
  


SEXP scalarX(SEXP x, SEXP y, SEXP mode) {
  int len = length(x);
  if (length(y) != len) ERR("x and y differ in length");
  int n;
  if (length(mode) == 0) n = -1;
  else {
    n = Match((char*) CHAR(STRING_ELT(mode, 0)), modi, Nmodi);
    if (n < 0) ERR("unknown modus");
  }
  SEXP Ans;
  PROTECT(Ans = allocVector(REALSXP, 1));
  double *ans = REAL(Ans);
  *ans = scalarX(REAL(x), REAL(y), len, n); // no PROTECT( needed
  UNPROTECT(1);
  return Ans;
}
```
