## Supplementary material for "MoBPS - Modular Breeding Program Simulator": RandomFieldsUtils R-package: solve.cc

```
/*
 Authors 
 Martin Schlather,

 Copyright (C) 2015 -- 2017 Martin Schlather, Reinhard Furrer, Martin Kroll

This program is free software; you can redistribute it and/or
modify it under the terms of the GNU General Public License
as published by the Free Software Foundation; either version 3
of the License, or (at your option) any later version.

This program is distributed in the hope that it will be useful,
but WITHOUT ANY WARRANTY; without even the implied warranty of
MERCHANTABILITY or FITNESS FOR A PARTICULAR PURPOSE.  See the
GNU General Public License for more details.

You should have received a copy of the GNU General Public License
along with this program; if not, write to the Free Software
Foundation, Inc., 59 Temple Place - Suite 330, Boston, MA  02111-1307, USA.  
*/
 
#include "Basic_utils.h"  // must be before anything else
#ifdef DO_PARALLEL
#include <omp.h>
#endif
#include <R_ext/Lapack.h>

#define LOCAL_ERRORSTRING
#define WHICH_ERRORSTRING pt->err_msg
#include "RandomFieldsUtils.h"
#include "own.h"
#include "zzz_RandomFieldsUtils.h"
#include "General_utils.h"
#include "kleinkram.h" 
#include "linear.h"


// 0 : 7.9
// 1:  7.55
// 2: 7.8
// 3:7.58
//4: 7.5
// 5: 7.4!
//6:7.4
//7: 7.9
// 8:

#define SCALAR(A,B,C) scalarX(A,B,C,NR)
#define LINEAR(A,B,C,D) linearX(A,B,C,D,6)

/* three trials n=2500 | 2 crs usr/sys/elap      |  new with linear (1;2crs)
0 : 8.140 8.076 8.036
1 : 6.288 6.296 6.284  |  6.988   0.012   5.665  | 6.032; 6.672   0.004   5.395
2 : 6.164 6.288 6.244 
3 : 6.596 6.696 6.636 
4 : ---
5 : 6.112 6.092 6.080  |  6.736   0.008   5.464
6 : 6.084 6.112 6.020  |  6.652   0.008   5.383  | 5.600; 6.756   0.008   3.413 

7 : 6.300 6.208 6.176 
8 : 7.828 7.836 7.792
9 : 10.24 10.10 10.00
10: >1min
2+10: >1min
10+2:9.14  9.24  9.23
2+9: 6.18  6.13  6.08  |  6.968   0.020   5.640
9+2:10.2  10.11 10.00   
*/


const char * InversionNames[nr_InversionMethods] = {
  "cholesky", "svd", "eigen", "sparse",
  "method undefined",
  "qr", "lu", 
  "no method left",
  "direct formula",
  "diagonal"};


    //  double *A_= A, *B_= B;				
     // i_ = N,					

#define KAHAN GLOBAL.basic.kahanCorrection

#define CMALLOC(WHICH, N, TYPE)	{				 \
    int _N_ = N;						 \
    if (pt->WHICH##_n < _N_) {					 \
      if (pt->WHICH##_n < 0) BUG; 				 \
      FREE(pt->WHICH);						 \
      pt->WHICH##_n = _N_;						\
	if ((pt->WHICH = (TYPE *) CALLOC(_N_, sizeof(TYPE))) == NULL)	\
	  return ERRORMEMORYALLOCATION;					\
    } else {								\
      assert( (_N_ > 0 && pt->WHICH != NULL) || _N_ == 0);		\
      for (int iii=0; iii<_N_; pt->WHICH[iii++] = 0);			\
    }									\
  }									\
    TYPE VARIABLE_IS_NOT_USED *WHICH = pt->WHICH


//  sqrtPosDef nutzt pt->U fuer das Ergebnis		
#define FREEING(WHICH)						\
  assert(int VARIABLE_IS_UNUSED *_i = WHICH);	\
  if (pt->WHICH != NULL && pt->WHICH != result) {	\
    UNCONDFREE(pt->WHICH);						\
    pt->WHICH##_n = 0;						\
  }						
 					       
#define FREEING_INT(WHICH)			\
  assert(int VARIABLE_IS_UNUSED *_i = WHICH);	\
  if (pt->WHICH != NULL) {			\
    UNCONDFREE(pt->WHICH);				\
    pt->WHICH##_n = 0;				\
  }						


double Determinant(double *M, int size, bool log) {
  int sizeP1 = size + 1,
    sizeSq = size * size;
  if (log) {
    double tmp = 0.0;
    for (int i=0; i<sizeSq; i+=sizeP1) tmp += LOG(M[i]);
    return tmp;
  } 
  double tmp = 1.0;
  for (int i=0; i<sizeSq; i+=sizeP1) tmp *= M[i];
  return tmp;
}

double cumProd(double *D, int size, bool log) {
  if (log) {
    double dummy = 0.0;
    for (int i = 0; i < size; dummy += LOG(D[i++]));
    return dummy;
  }
  double dummy = 1.0;
  for (int i = 0; i < size; dummy *= D[i++]);
  return dummy;
}


void solve_DELETE0(solve_storage *x) {
    FREE(x->iwork);
    FREE(x->ipiv);

    FREE(x->pivotsparse);
    FREE(x->pivot_idx);
    FREE(x->xlnz);
    FREE(x->snode);
    FREE(x->xsuper);
    FREE(x->xlindx);
    FREE(x->invp);
    FREE(x->cols);
    FREE(x->rows);
    FREE(x->lindx);
    FREE(x->xja);
    //    FREE(x->);
   
    // double *
    FREE(x->SICH);
    FREE(x->MM);
    FREE(x->workspaceD);
    FREE(x->workspaceU);

    FREE(x->VT);
    FREE(x->work);
    FREE(x->w2);
    FREE(x->U);
    FREE(x->D);
    
    FREE(x->workLU);
    FREE(x->diagonal);
  
    FREE(x->lnz); 
    FREE(x->DD);
    FREE(x->w3);
    FREE(x->result);
    FREE(x->to_be_deleted);
}

void solve_DELETE(solve_storage **S) {
  solve_storage *x = *S;
  if (x!=NULL) {
    solve_DELETE0(*S);
    UNCONDFREE(*S);
  }
}
void solve_NULL(solve_storage* x) {
  if (x == NULL) return;
  MEMSET(x, 0, sizeof(solve_storage));  
  x->nsuper = x->nnzlindx = x->size = -1;
  x->method = NoInversionMethod;
  for (int i=0; i<SOLVE_METHODS; x->newMethods[i++] = NoInversionMethod);
  x->actual_pivot = PIVOT_UNDEFINED;
}

double static inline det3(double *M, int size) {
  double det;
  switch(size){ // Abfrage nach Groesse der Matrix M + Berechnung der Determinante per Hand
  case 1: det = M[0];
    break;
  case 2: det = M[0] * M[3] - M[1] * M[2];
    break;
  case 3: det = 
      M[0] * (M[4] * M[8] - M[5] * M[7]) 
      - M[1] * (M[3] * M[8] - M[5] * M[6]) 
      + M[2] * (M[3] * M[7] - M[4] * M[6]); // Entwicklung nach 1. Spalte
    break;
  default : BUG;
    break;
  }
  return det;
}

int logdet3(double det, bool posdef, double *logdet, bool log) {
  if (posdef && det < 0) return ERRORFAILED;
  if (logdet != NULL) {
    if (log) {
      if (det <= 0) return ERRORFAILED;
      *logdet = LOG(det);
    } else *logdet = det;
  }
  return NOERROR;
}

int solve3(double *M, int size, bool posdef,
	   double *rhs, int rhs_cols,
	   double *result, double *logdet, bool log,
	   solve_storage *pt	
	   ){
  
  assert(size <= 3);
  if (size <= 0) SERR("matrix in 'solvePosDef' of non-positive size.");

  double det = det3(M, size);
  if (logdet3(det, posdef, logdet, log) != NOERROR) return ERRORFAILED;

  double detinv = 1.0 / det; // determinant of inverse of M
  
  switch(size){
  case 1 : {// size of matrix == 1
    if (rhs_cols == 0) result[0] = detinv;   
    else for (int i=0; i<rhs_cols; i++) result[i] = rhs[i] * detinv;
  }
    break;
  case 2 : { // size of matrix == 2
    double a = M[0] * detinv,
      d = M[3] * detinv;
    if (rhs_cols == 0) {
      result[0] = d;
      result[1] = -M[1] * detinv;
      result[2] = -M[2] * detinv;
      result[3] = a;
    } else { // rhs_cols != 0
      double *p = rhs, *q = result;
      if (M[1] != 0.0 || M[2] != 0.0) {
	double 
	  b = M[1] * detinv,
	  c = M[2] * detinv;
	for (int i=0; i<rhs_cols; i++, p+=2, q+=2) {
	  double swap = d * p[0] - c * p[1];
	  q[1] = a * p[1] - b * p[0];
	  q[0] = swap;
	}
      } else {
	for (int i=0; i<rhs_cols; i++, p+=2, q+=2) {
	  double swap = d * p[0];
	  q[1] = a * p[1];
	  q[0] = swap;
	}
      }
    }
  }
    break;
  case 3 : {// size of matrix == 3   
    double swap0 = detinv * (M[4] * M[8] - M[5] * M[7]),
      swap1 = detinv * (M[5] * M[6] - M[3] * M[8]),
      swap2 = detinv * (M[3] * M[7] - M[4] * M[6]),
      swap3 = detinv * (M[2] * M[7] - M[1] * M[8]),
      swap4 = detinv * (M[0] * M[8] - M[2] * M[6]),
      swap5 = detinv * (M[1] * M[6] - M[0] * M[7]),
      swap6 = detinv * (M[1] * M[5] - M[2] * M[4]),
      swap7 = detinv * (M[2] * M[3] - M[0] * M[5]),
      swap8 = detinv * (M[0] * M[4] - M[1] * M[3]);
    if(rhs_cols == 0){ // invert matrix
      result[0] = swap0;
      result[1] = swap1;
      result[2] = swap2;
      result[3] = swap3;
      result[4] = swap4;
      result[5] = swap5;
      result[6] = swap6;
      result[7] = swap7;
      result[8] = swap8;
    } else { // solve system given by M and rhs
      double *p = rhs, *q = result;
      for (int i=0; i<rhs_cols; i++, p+=3, q+=3) {
	double swapA = p[0] * swap0 + p[1] * swap3 + p[2] * swap6;
	double swapB = p[0] * swap1 + p[1] * swap4 + p[2] * swap7;
	q[2] = p[0] * swap2 + p[1] * swap5 + p[2] * swap8;
	q[0] = swapA;
	q[1] = swapB;
      }
    }
  }
    break;
  default: BUG;
  }
  
  return NOERROR;
}

int chol3(double *M, int size, double *res, solve_storage *pt){
  // UNBEDINGT in sqrtRHS.cc auch aendern
  assert(size <= 3);
  if (size <= 0) SERR("matrix in 'solvePosDef' not of positive size.");
  //  if (M[0] < 0) return ERRORFAILED;
  res[0] = SQRT(M[0]);
  if (size == 1) return NOERROR;
  res[1] = 0.0;
  res[size] = res[0] > 0.0 ? M[size] / res[0] : 0.0;
  double dummy = M[size + 1] - res[size] * res[size];
  res[size + 1] = SQRT(MAX(0.0, dummy));
  if (size == 2) return NOERROR;
  res[2] = res[5] = 0.0;
  res[6] = res[0] > 0.0 ? M[6] / res[0] : 0.0;
  res[7] = res[4] > 0.0 ? (M[7] - res[3] * res[6]) / res[4] : 0.0;
  dummy = M[8] - res[6] * res[6] - res[7] * res[7];
  res[8] = SQRT(MAX(0.0, dummy));
  return NOERROR;
}


void Sort(double *RESULT, int size, int rhs_cols, int *pi, int *rank,
	  double *dummy) {
  orderingInt(pi, size, 1, rank);
  int i=0,
    totalRHS = size * rhs_cols;
  while(i < size && i == rank[i]) i++;
  while (i < size) {
    int stored_i = i,
      read = i;
    double *p_write = NULL,
      *p_read = RESULT + read;
    for (int k=0; k<rhs_cols; k++) dummy[k] = p_read[k * size];
    while (true) {
      int write = read;
      p_write = RESULT + write;
      read = rank[write];
      rank[write] = write;
      if (read == stored_i) {
	for (int k=0; k<rhs_cols; k++) p_write[k*size]=dummy[k];
	break;
      }
      p_read = RESULT + read;
      for (int k=0; k<totalRHS; k+=size) p_write[k] = p_read[k];
    }
    while(i < size && i == rank[i]) i++;
  }
}


// to do: fehlt: chol2solve(chol, x)


void chol2inv(double *MPT, int size) {
  int sizeP1 = size + 1,
    sizeSq = size * size,
    NR = KAHAN ? SCALAR_KAHAN : SCALAR_AVX;

  double *diagonal = (double *) MALLOC(sizeof(double) * size);
#ifdef DO_PARALLEL
#pragma omp parallel for num_threads(CORES) if (size > 60) schedule(dynamic, 20)
#endif	    
  for (int k=0; k<size; k++) {
    double *p_RESULT = MPT + k * sizeP1,
      diagK = diagonal[k] = 1.0 / p_RESULT[0];
    for (int i=1; i<size - k; i++) {
      double *pM = p_RESULT + i * size;
      p_RESULT[i] = (-diagK * pM[0] - SCALAR(pM + 1, p_RESULT + 1, i -1))
	/ pM[i];
    }
    // i == k
  }
  
#ifdef DO_PARALLEL
#pragma omp parallel for num_threads(CORES) if (size > 60) schedule(dynamic, 20)
#endif	    
  for (int k=0; k<size; k++) {	      
    double *p_RESULT = MPT + k * size;
    for (int i=size-1; i>k; i--) {
      double *pM = MPT + i * size,
	r = (p_RESULT[i] /= pM[i]);
      diagonal[k] -= r *pM[k];
      LINEAR(pM + k + 1, -r, i-k-1, p_RESULT + k + 1);
      // for (int j=k+1; j<i; j++) p_RESULT[j] -= r * pM[j]; 
    }
    // i == k
  }
  
  for (int k=0; k<size; k++) {
    double *pM = MPT + k * size;	         
    pM[k] = diagonal[k] / pM[k];
  }
  
  for (int i2=0,i=0; i<size; i++, i2+=size + 1) {
    int i3 = i2 + 1;
    for (int j = i2 + size; j<sizeSq; j+=size) 
      MPT[j] = MPT[i3++];
  }
  FREE(diagonal);
}


int doPosDef(double *M, int size, bool posdef,
	     double *rhs, int rhs_cols, double *result,
	     double *logdet, int calculate, solve_storage *Pt,
	     solve_param *sp
	     ){
  // it is ok to have
 // ==15170== Syscall param sched_setaffinity(mask) points to unaddressable byte(s)
  // caused by gcc stuff

  //  printf("doPosDef %ld %d %d rhs=%ld %d %ld %ld calc=%d %ld %ld\n",
  //	 M, size, posdef, rhs, rhs_cols, result, logdet, calculate, Pt,sp);
/*
    M: (in/out) a square matrix (symmetry is not checked) of size x size;
       NOTE THAT THE CONTENTS OF M IS DESTROYED IFF NO RHS IS GIVEN
       AND result IS NOT GIVEN.
       In case rhs is not given, the inverse of M is returned here
       In case of sqrtonly M is expected to be a positive definite matrix
    posdef (in): whether or not the matrix is positiv (semi)definite --
            to some extend doPosDef can deal with non-positiv definite
            functions
    rhs (in/out) : right hand side of the equality with rhs_cols columns
          NOTE THAT THE CONTENTS OF rhs WILL BE DESTROYED IF rhs IS GIVEN 
 
          the solution of the equality is returned in rhs
    rhs_cols : number of colums of the matrix on the right hand side
    result (out) : NULL or matrix of the size of the result (inverse matrix or 
           of size of the matrix on the right hand side); see also 'M' and 'rhs'
    logdet (out): if not NULL the logarithm of the determinant is returned
    pt (in/out) : working space. If NULL, internal working spaces are used.
 
          A non-NULL value gives an advantage only if doPosDef is called
          repeatedly. Then 
            solve_storage *pt = (solve_storage*) m alloc(sizeof(solve_storage);
            solve_NULL(pt);
          prepares pt. Deletion is done only at the very end by
            solve_DELETE(pt);
          In meanwhile the working space is managed by doPosDef;
     Sp (in): parameters. If NULL, the behaviour is as described in
          the R manual for doPosDef.
          The parameters are described in the package 'RandomFields'
          under ?RFoptions
	  
  */

  
  // http://www.nag.com/numeric/fl/nagdoc_fl23/xhtml/F01/f01intro.xml#
  assert(NA_LOGICAL == INT_MIN && NA_LOGICAL == NA_INTEGER); // nur zur sicherheit, wegen usr_bool
  //          eigentlich sollte usr_bool unabhaengig davon funktionieren
  assert(calculate != DETERMINANT ||
	 (logdet != NULL && result == NULL && rhs == NULL));
  assert(calculate != MATRIXSQRT || (rhs == NULL && posdef));
  assert(rhs_cols == 0 || rhs != NULL);

  double *RESULT = result != NULL ? result : rhs_cols > 0 ? rhs : M;
  assert(sp != NULL);

  if (size <= 3 && sp->pivot == PIVOT_AUTO) {
    if (Pt != NULL) {
      Pt->method = direct_formula;
      Pt->size = size;
    }
    if (calculate == DETERMINANT)
      return logdet3(det3(M, size), posdef, logdet, sp->det_as_log);
    else if (calculate  == MATRIXSQRT) return chol3(M, size, RESULT, Pt);
    else return solve3(M, size, posdef, rhs, rhs_cols, RESULT, logdet,
		       sp->det_as_log, Pt);
  }

  assert(SOLVE_METHODS >= 2);
 
  solve_storage *pt;
  if (Pt != NULL) {
    pt = Pt; 
  } else {
    pt = (solve_storage*) MALLOC(sizeof(solve_storage));
    solve_NULL(pt);    
  }
  int  
    err = NOERROR,
    spam_zaehler = 0,
    nnzA = 0,
    sizeSq = size * size,
    sizeP1 = size + 1;
  usr_bool
    sparse = sp->sparse;
  double 
     spam_tol = sp->spam_tol;
  bool diag  = false;
 
  pt->method = NoFurtherInversionMethod;
  pt->size = size;
 
  if (sparse == Nan && (sparse = (usr_bool) (size > sp->spam_min_n))) {
    double mean_diag = 0.0;
    for (int i=0; i<sizeSq; i += sizeP1) mean_diag += M[i];
    mean_diag /= (double) size;
    spam_tol *= mean_diag;

    bool random_sample = sizeSq >= sp->spam_sample_n * 3;
    if (random_sample) {
      double 
	thr = sp->spam_sample_n * (1.0 - sp->spam_min_p);
      int	
	threshold = (int) (thr + SQRT(thr) * 3),
	notZero = 0;
      for (int i=0; i<sp->spam_sample_n; i++) {
	if ((notZero += FABS(M[(i * sp->spam_factor) % sizeSq]) > 
	     spam_tol) >= threshold){
	  sparse = False;
	  break;
	}
      }
      if (PL >= PL_FCTN_DETAILS) {
	PRINTF("random sampling: sparse=%d\n",
	       sparse == Nan ? NA_INTEGER : (int) sparse);
      }
    }
    if (!random_sample || sparse == True) {
      int diag_nnzA = 0;
      //#ifdef DO_PARALLEL
      //#pragma omp parallel for num_threads(CORES) schedule(dynamic,10) reduction(+:nnzA,diag_nnzA)
      //#endif
      for (int i=0; i<size; i++) {
	int end = i * sizeP1;
	Long j;
	for (j=i * size; j<end; nnzA += FABS(M[j++]) >= spam_tol);
	diag_nnzA += FABS(M[j++]) > spam_tol;
	end = (i+1) * size;
	if (!posdef) for (; j<end; j++) nnzA += FABS(M[j++]) >= spam_tol;
      }
      diag = (nnzA == 0);
      if (posdef) nnzA *= 2;
      nnzA += diag_nnzA;
      sparse = (usr_bool) (nnzA <= sizeSq * (1.0 - sp->spam_min_p));
      spam_zaehler = nnzA + 1;
      if (PL >= PL_DETAILSUSER) {
	if (diag) { PRINTF("diagonal matrix detected\n"); }
	else if (sparse == True) {
	  PRINTF("sparse matrix detected (%3.2f%% zeros)\n", 
		 100.0 * (1.0 - nnzA / (double) sizeSq));
	}
	else {  PRINTF("full matrix detected (%3.2f%% nonzeros)\n", 
		       100.0 * nnzA / (double) sizeSq); }
      }
    }
  } else {
    diag = true;
    for (int i=0; i<size && diag; i++) {
      int end = i * sizeP1;
      Long j;
      for (j=i * size; j<end; j++) {
	//	printf("(%d %d %10g %d)\n", i, j, M[j], size);
	if (FABS(M[j]) > spam_tol) {
	  diag = false;
	  break;
	}
      }
      if (!diag) break;
      j++;
      end = (i+1) * size;
      if (!posdef) {
	for (; j<end; j++) {
	  if (FABS(M[j]) > spam_tol) {
	    diag = false;
	    break;
	  }
	}
      }
    }
  }

  if (diag) {
    pt->method = Diagonal;
    if (PL>=PL_STRUCTURE) { PRINTF("dealing with diagonal matrix\n"); }
    if (logdet != NULL) {
      *logdet = Determinant(M, size, sp->det_as_log);
      if (calculate == DETERMINANT) return NOERROR;
    }
    if (rhs_cols == 0) {
      MEMCOPY(RESULT, M, sizeSq * sizeof(double));
      if (calculate == MATRIXSQRT) {
	for (int i=0; i<sizeSq; i += sizeP1) {
	  RESULT[i] = M[i] > 0.0 ? SQRT(M[i]) : 0.0;	
	}
      } else {
	for (int i=0; i<sizeSq; i += sizeP1) 
	  RESULT[i] = M[i] <= 0.0 ? 0.0 : 1.0 / M[i];
      }
    } else {
      CMALLOC(MM, size, double);
      for (int i=0; i<size; i++) {
	int idx = i * sizeP1;
	MM[i] = M[idx] == 0.0 ? 0.0 : 1.0 / M[idx];
      }
      int j;
      for (int k=j=0; j<rhs_cols; j++)
	for (int i=0; i<size; i++, k++) RESULT[k] = rhs[k] * MM[i];
    }
    
    err = NOERROR;
    goto ErrorHandling;
  }


  // size of matrix at least 4 x 4, and not diagonal
  InversionMethod *Meth;
  if (sparse == True || sp->Methods[0] == NoFurtherInversionMethod ||
      sp->Methods[0] == NoInversionMethod) {
    Meth = pt->newMethods;
    if (sparse == True) {
      Meth[0] = Sparse;
      bool given0 =  sp->Methods[0] != NoFurtherInversionMethod &&
	sp->Methods[0] != NoInversionMethod;
      Meth[1] = given0 && sp->Methods[0] != Sparse
	? sp->Methods[0] 
	: posdef ? Cholesky : LU;  
      if (SOLVE_METHODS > 2) {
	bool given1 = sp->Methods[1] != NoFurtherInversionMethod &&
	  sp->Methods[1] != NoInversionMethod;
	Meth[2] = given0 && sp->Methods[0] != Sparse &&
	  given1 && sp->Methods[1] != Sparse
	  ? sp->Methods[1] 
	  : posdef ? Eigen : LU;
      }
      // pt->newMethods[1] = Sparse;
    } else {
      Meth[0] = posdef ? Cholesky : LU;  
      Meth[1] =  posdef ? Eigen : LU;
      if (SOLVE_METHODS > 2) Meth[2] = Eigen;
    }
    for (int i=3; i<SOLVE_METHODS; Meth[i++]=NoFurtherInversionMethod);   
  } else Meth = sp->Methods;

  if (!posdef && Meth[0] != Eigen && Meth[0] != Eigen) {
    err = ERRORNOTPROGRAMMEDYET;
    goto ErrorHandling;
  }

  // cholesky, QR, SVD, Eigen, LU always destroy original matrix M
  bool gesichert;
  if ((gesichert = rhs_cols==0 && result == NULL)) {
    if ((gesichert = (SOLVE_METHODS > sparse + 1 &&
		      Meth[sparse + 1] != Meth[sparse] &&
		      Meth[sparse + 1] != NoFurtherInversionMethod)
	 || (Meth[sparse] == SVD && sp->svd_tol > 0.0 && calculate != SOLVE)
	 )) { // at least two different Methods in the list
      CMALLOC(SICH, sizeSq, double);
      MEMCOPY(SICH, M, sizeSq * sizeof(double));
    }
  }
  double *SICH, *MPT;
  SICH = pt->SICH;

  MPT = M; // also for sparse result
  if (rhs_cols > 0) {
    CMALLOC(MM, sizeSq, double);
    MPT = MM;
  } else if (result != NULL) MPT = result;


  errorstring_type ErrStr;
  STRCPY(ErrStr, "");
 
  for (int m=0; m<SOLVE_METHODS && (m==0 || Meth[m] != Meth[m-1]); m++) {    
    pt->method = Meth[m];
    if (pt->method<0) break;
    if (calculate != SOLVE) {
      if (pt->method == NoInversionMethod && m<=sparse) BUG;
       if (pt->method == NoFurtherInversionMethod) break;
     if (PL>=PL_STRUCTURE) { 
	PRINTF("method to calculate the square root : %s\n", 
	       InversionNames[pt->method]);
      }
    } else {
	if (PL>=PL_STRUCTURE) { 
	  PRINTF("method to calculate the inverse : %s\n",
		 InversionNames[pt->method]);
      }
    }
     
    if (rhs_cols == 0 && result == NULL) {
      if (m > sparse) {      
	MEMCOPY(MPT, SICH, sizeSq * sizeof(double));
      }
    } else if (pt->method != Sparse && MPT != M) {
      MEMCOPY(MPT, M, sizeSq * sizeof(double));
    }
    
    switch(pt->method) {
    case Cholesky : {
#define C_GERR(X,G) {STRCPY(ErrStr, X); FERR(X); err = ERRORM; goto G;}
#define C_GERR1(X,Y,G) {SPRINTF(ErrStr,X,Y); FERR(ErrStr);err = ERRORM; goto G;}
#define C_GERR2(X,Y,Z,G){SPRINTF(ErrStr,X,Y,Z);FERR(ErrStr);err=ERRORM; goto G;}
#define C_GERR3(X,Y,Z,A,G) {SPRINTF(ErrStr,X,Y,Z,A); FERR(ErrStr);err = ERRORM; goto G;}
      int NR = KAHAN ? SCALAR_KAHAN : SCALAR_AVX;
      if (!posdef) CERR("Cholesky needs positive definite matrix");
      if (size > sp->max_chol)
	CERR2("Matrix  is too large for Cholesky decomposition. Maximum ist currently a %d x %d matrix. Increase 'max_chol' in 'RFoption' if necessary.",
	      sp->max_chol, sp->max_chol);

      
      CMALLOC(diagonal, size > rhs_cols ? size : rhs_cols, double);
      for (int i=0; i<size; i++) diagonal[i] = MPT[sizeP1 * i];
      
      pt->actual_pivot = PIVOT_UNDEFINED;

      //      printf("sp->pivot = %d\n", sp->pivot);
      
      if (sp->pivot == PIVOT_NONE || sp->pivot == PIVOT_AUTO) {// cholesky

	// cmp for instance http://stackoverflow.com/questions/22479258/cholesky-decomposition-with-openmp

	// obere und untere dreiecksmatrix wird gelesen und obere geschrieben
	err = NOERROR;
	pt->actual_pivot = PIVOT_NONE;
	{
	  double *A = MPT;
	  for (int i=0; i<size; i++, A += size) {
	    double sclr = SCALAR(A, A, i);
	    if (A[i] <= sclr) {
	      if (sp->pivot == PIVOT_NONE)
		C_GERR3("Got %10e as %d-th eigenvalue.%.50s)'.",
			A[i] - sclr, i, sp->pivot == PIVOT_NONE
			? " Try with 'RFoptions(pivot=PIVOT_DO" : "",
			Pivot_Cholesky)
		else C_GERR(".", Pivot_Cholesky);
	      break;
	    }
	    A[i] = SQRT(A[i] - sclr);
	    
	    //	  double invsum = 1.0 / A[i];
	    double sum = A[i];
#ifdef DO_PARALLEL
#pragma omp parallel for num_threads(CORES) if (MULTIMINSIZE(size - i)) schedule(dynamic, 8) 
#endif
	    //	    for (double *B=MPT + (i+1) * size; B<endB; B+=size) {
	    for (int j = i + 1; j < size; j++) {
	      double *B = MPT + j * size;
	      B[i] = (B[i] - SCALAR(A, B, i)) / sum;
	    }
	  }
	}
	
	if (err == NOERROR) {
	  if (calculate == MATRIXSQRT) {
	    int deltaend = size - 1;
	    double *end = MPT + sizeSq;
	    for (double *p=MPT + 1; p<end; p+=sizeP1, deltaend--)
	      FILL_IN(p, deltaend, 0.0);	    
	  } else {
	    if (logdet != NULL) {
	      *logdet = Determinant(MPT, size, sp->det_as_log);
	      if (sp->det_as_log) *logdet *=2; else *logdet *= *logdet;
	      if (calculate == DETERMINANT) return NOERROR;
	    }

	    if (rhs_cols == 0) chol2inv(MPT, size);
	    else { // rhs_cols > 0
	      //int totalRHS = size * rhs_cols;
	      //if (result!=NULL) MEMCOPY(RESULT, rhs, sizeof(double)*totalRHS);
#ifdef DO_PARALLEL
#pragma omp parallel for num_threads(CORES) if (rhs_cols > 30)
#endif	      
	      for (int k=0; k<rhs_cols; k++) {
		double *p_RESULT = RESULT + k * size,
		  *p_rhs = rhs + k * size;
		for (int i=0; i<size; i++) {
		  double *pM = MPT + i * size;
		  p_RESULT[i] = (p_rhs[i] - SCALAR(pM, p_RESULT, i)) / pM[i];
		}
	      }
	      
#ifdef DO_PARALLEL
#pragma omp parallel for num_threads(CORES) if (rhs_cols > 30)
#endif	      
	      for (int k=0; k<rhs_cols; k++) {	      
		double *p_RESULT = RESULT + k * size;
		for (int i=size-1; i>=0; i--) {
		  double *pM = MPT + i * size,
		    r = (p_RESULT[i] /= pM[i]);
		  LINEAR(pM, -r, i, p_RESULT);
		  // for (int j=0; j<i; j++)  p_RESULT[j] -= r * pM[j];
		}
	      }
	    }
	  } // not sqrt only
	} // err == NOERROR
      } else err = ERRORFAILED;
    
      Pivot_Cholesky:
      if (err != NOERROR && sp->pivot != PIVOT_NONE) {
	if (PL > PL_DETAILS) { PRINTF("trying pivoting\n"); }
	int actual_size = NA_INTEGER;
	// code according to Helmut Harbrecht,Michael Peters,Reinhold Schneider
	// talk: The pivoted Cholesky decomposition and its application to
	//       stochastic PDEs
	// ebenso: untere dreiecksmatrix wird gelesen; obere geschrieben
	if (pt->actual_pivot == PIVOT_NONE) {
	  // wiederherstellung der Diagonalen und der unteren dreiecksmatrix
	  for (int i=0; i<size; i++) MPT[sizeP1 * i] = diagonal[i];
	}
	int *pi;
	if (sp->pivot == PIVOT_DO || sp->pivot == PIVOT_AUTO) {
 	  FREE(pt->pivot_idx); // ALWAYS FREE IT!!! cp Chol(SEXP M)
	  pt->pivot_idx = (int*) MALLOC(size * sizeof(int));
	  pt->pivot_idx_n = size;
	  pt->actual_pivot = PIVOT_DO;
	  for (int i=0; i<size; i++) pt->pivot_idx[i] = i;
	  pt->actual_size = actual_size = size;
	  pi = pt->pivot_idx;
	} else { // PIVOT_IDX 
	  if (sp->pivot_idx_n < size || sp->actual_size > size) {
	    //printf("XA, %d %d %d\n", sp->pivot_idx_n , size, sp->actual_size);
	    CERR("pivot_idx does not have the correct length.\nSet 'RFoption(pivot_idx=, pivot_actual_size=)' to the attributes of a\npivoted Cholesky decomposition.");
	  }
	  actual_size = pt->actual_size = sp->actual_size;
	  if (actual_size > size) BUG;
	  FREE(pt->pivot_idx);
	  int bytes = size * sizeof(int);
	  pt->pivot_idx = (int*) MALLOC(bytes);
	  MEMCOPY(pt->pivot_idx, sp->pivot_idx, bytes);
	  pt->actual_pivot = PIVOT_IDX;
	  pt->pivot_idx_n = sp->pivot_idx_n;
	  pi = sp->pivot_idx;
	}

	err = NOERROR;
	double
	  rel_thres = 0,
	  max_deviation = sp->max_deviation, //  1e-10,
	  max_reldeviation = sp->max_reldeviation, //  1e-10,
	  *Morig = gesichert ? SICH : M;
	if (MPT == Morig || (rhs_cols > 0 && rhs == RESULT))
	  CERR("Pivoted cholesky cannot be performed on place! Either you are a programmer or you should contact the maintainer.");
	for (int q=0; q<actual_size; q++) {
	  if (pt->actual_pivot == PIVOT_DO) {
	    double 
	      max = RF_NEGINF,
	      deviation = 0.0;
	    int k,
	      argmax = NA_INTEGER;	    
	    for (k=q; k<size; k++) {
	      double dummy = diagonal[pi[k]];
	      // if (diagonal[pi[k]] < 0)

	      //printf("k=%d %10e %10e\n", k, dummy, -1e-15 * size * size);
	      if (dummy < -1e-4 * sp->pivot_relerror* size * size){
		C_GERR1("matrix not positive definite or increase 'pivot_relerror' by at least factor %10g.", dummy * -1e4 / (size * size), ERR_CHOL);
	      }
	      deviation += dummy;
	      if (max < dummy) {
		max = dummy;
		argmax = k;		    
	      }
	    }
	    double dev = rel_thres * max_reldeviation;
	    if (deviation <= max_deviation || (q > 0 && deviation <= dev) ) {
	      actual_size = pt->actual_size = q;
	      if (sp->pivot_check != False) {
		double largest = 0;
		//printf("q=%d %d\n", q, size);
		for (int i=q; i<size; i++) {
		  double *mpt = MPT + pi[i] * size;
		  for (int j=q; j<=i; j++) {
		    double absm = FABS(mpt[j]);
		    largest = absm > largest ? absm : largest;
		    // if(absm == largest || absm > 5) printf("%10e %d %d; %d\n", absm, i, j, size);
		  }
		}
		if (largest > max_deviation || (q > 0 && largest > dev)) {
		  char msg[500];
		  SPRINTF(msg, "Matrix has a numerically zero, leading value at the %d-th pivoted line, but the largest deviation %10e from zero in the rest of the matrix is greater than the tolerance %10e. %.50s.",
			  q,
			  largest,
			  MAX(max_deviation, dev),
			  sp->pivot_check == True
			  ? "If you are sure that the matrix is semi-definite, set 'RFoptions(pivot_check=NA)' or 'RFoptions(pivot_check=True)'"
			  : "The result can be imprecise");
		  if (sp->pivot_check == True) C_GERR(msg, ERR_CHOL)
		    else warn(msg);
		}
	      }
	      break;
	    }
	    rel_thres += diagonal[pi[q]];
	    int dummy = pi[q];
	    pi[q] = pi[argmax];
	    pi[argmax] = dummy;
	  }
	  
	  
	  int pqq = pi[q],
	    col_q = pqq * size;
	  
	  if (diagonal[pqq] < 0) {
	    C_GERR1("Negative leading value found at the %d-th pivoted line.",
		    q, ERR_CHOL);
	  }
	  double lqpq = MPT[q + col_q] = SQRT(diagonal[pqq]);	    
#ifdef DO_PARALLEL
#pragma omp parallel for num_threads(CORES) if (MULTIMINSIZE(size - q)) schedule(dynamic, 8) 
#endif
	  for (int i=q+1; i<size; i++) {
	    int
	      pii = pi[i],
	      col_i = pii * size;
	    assert(pii != pqq);
	    double scalar = SCALAR(MPT + col_q, MPT + col_i, q);
	    MPT[q + col_i] = (Morig[pqq + col_i] - scalar) / lqpq;
	    diagonal[pii] -=  MPT[q + col_i] *  MPT[q + col_i];
	    // in Harbrecht: lqpq * MPT[q + col_i];
	  }
	  
	  /*	  if (!true) {
		  for (int k=0; k<size; k++) {  
		  for (int j=0; j<size; j++)
		  p rintf("%10g ", MPT[size * j + k]);
		  p rintf("\n");
			  } p rintf("\n");
			  }
	  */
	  
	} // for q

	
	if (err == NOERROR) {
	  if (calculate == MATRIXSQRT) {
	    int i = 0;
	    for ( ; i<actual_size; i++) {
	      FILL_IN(MPT + i + 1 + size * pi[i], size - 1 - i, 0.0);
	    }
	    for ( ; i<size; i++) {
	      FILL_IN(MPT + actual_size + size*pi[i], size - actual_size, 0.0);
	    }
	    
	  } else {
	    if (logdet != NULL) {
	      if (sp->det_as_log) {
		double logD = 0.0;
		for (int i=0; i < size; i++) logD += LOG(MPT[i + pi[i] * size]);
		*logdet = logD * 2;
	      } else {
		double logD = 1.0;
		for (int i=0; i < size; i++) logD *= MPT[i + pi[i] * size];
		*logdet = logD * logD;
	      }
	      if (calculate == DETERMINANT) return NOERROR;
	    }
	    
	    //////////////////////////////////////////////////
	    //////////////////////////////////////////////////

	    if (rhs_cols == 0) {
 	      if (actual_size < size) 
		GERR("Matrix not definite. Try 'solve(M , diag(nrow(M)))' instead or add a small positive constant to the diagonal.")
	      
#ifdef DO_PARALLEL
#pragma omp parallel for num_threads(CORES) if (size > 60) schedule(dynamic, 20)
#endif
		  for (int k=0 ; k<actual_size; k++) {
		    double *p_RESULT = MPT + pi[k] * size + k,
		      diagK = diagonal[k] = 1.0 / p_RESULT[0];
		    for (int i=1; i<size - k; i++) {
		      double *pM = MPT + k + pi[k + i] * size;
		      p_RESULT[i] = (-diagK * pM[0]
				     -SCALAR(pM + 1, p_RESULT + 1, i -1)) / pM[i];
		    }
		  }     
	      
	      
#ifdef DO_PARALLEL
#pragma omp parallel for num_threads(CORES) if (size > 60) schedule(dynamic, 20)
#endif	    
	      for (int k=0; k<size; k++) {	      
		double *p_RESULT = MPT + pi[k] * size;
		for (int i=actual_size-1; i>k; i--) {
		  double *pM = MPT + pi[i] * size,		    
		    r = (p_RESULT[i] /= pM[i]);
		  diagonal[k] -= r * pM[k];
		  LINEAR(pM + k + 1, -r, i-k-1, p_RESULT + k + 1);
		  // for (int j=k+1; j<i; j++) p_RESULT[j] -= r * pM[j]; 
		}
		// i == k
	      }
	      
	      for (int k=0; k<size; k++) {	      
		double *p_RESULT = MPT + pi[k] * size;
		for (int q=0; q<k; p_RESULT[q++] = RF_INF);
		//	for (int q=actual_size; q<size; p_RESULT[q++] = RF_NA);
	      }
	      
	      for (int k=0; k<actual_size; k++) {
		double *pM = MPT + pi[k] * size;	         
		pM[k] = diagonal[k] / pM[k];
	      }
	      
	      CMALLOC(xja, size, int);
	      Sort(RESULT, size, size, pi, xja, diagonal);
	      for (int i=0; i<size; i++) {
		for (int j=i+1; j<size; j++) {
		  int idx = i + j * size;
		  if (MPT[idx] == RF_INF) MPT[idx] = MPT[j + i * size];
		  else MPT[j + i * size] = MPT[idx];
		}
	      }
	      
	      
	      //////////////////////////////////////////////////
	      //////////////////////////////////////////////////
	      
	    } else { // rhs_cols > 0
	      assert(rhs != RESULT);
	      double eps = diagonal[0] * sp->pivot_relerror;
			      
#ifdef DO_PARALLEL
#pragma omp parallel for num_threads(CORES) if (rhs_cols > 30)
#endif
	      for (int k=0; k<rhs_cols; k++) {
		double *p_RESULT = RESULT + k * size,
		  *p_rhs = rhs + k * size;
		int i=0;
		for ( ; i<actual_size; i++) {
		  int pii = pi[i];
		  double *pM = MPT + pii * size;
		  p_RESULT[i] = (p_rhs[pii] - SCALAR(pM, p_RESULT, i)) / pM[i];
		}		
	 	for ( ; i<size; i++) {
		  int pii = pi[i];
		  double *pM = MPT + pii * size;
		  p_RESULT[i] = 0.0;
		  if (FABS((p_rhs[pii] - SCALAR(pM, p_RESULT, i))) > eps) {
		    if (Pt == NULL) solve_DELETE(&pt);		   
		    ERR1("Equation system not solvable (difference %10e). Try increasing 'pivot_relerror' in 'RFoptions' to get an approximate solution.",
			 p_rhs[pii] - SCALAR(pM, p_RESULT, i));
		  }
		}
	      }

#ifdef DO_PARALLEL
#pragma omp parallel for num_threads(CORES) if (rhs_cols > 30)
#endif 
	      for (int k=0; k<rhs_cols; k++) {	      
		double *p_RESULT = RESULT + k * size;
		for (int i=actual_size - 1; i>=0; i--) {
		  int pii = pi[i];
		  double *pM = MPT + pii * size,
		    r = (p_RESULT[i] /= pM[i]);
		  LINEAR(pM, -r, i, p_RESULT);
		  // for (int j=0; j<i; j++) p_RESULT[j] -= r * pM[j];
		}
	      }
	      CMALLOC(xja, size, int);
	      Sort(RESULT, size, rhs_cols, pi, xja, diagonal);
	    }// rhs_cols > 0
	  } // not sqrt only

	} // err == NOERROR	
      }

      ERR_CHOL:
      
      if (err != NOERROR) {	
	if (pt->actual_pivot == PIVOT_NONE)
	  CERR1("Probably matrix not positive semi definite: %.50s\nTry more flexible options in 'RFoptions' ('pivot = PIVOT_AUTO or 'pivot = PIVOT_DO'').\n", ErrStr)
	else // pt->actual_pivot == PIVOT_DO  or  PIVOT_IDX
	  CERR1("Likely, the matrix is not positive semi definite: %.50s\n", ErrStr)
      }

      if (PL >=  PL_DETAILSUSER) {
	PRINTF("Cholesky decomposition successful\n");
      }
      
      break;
    }
    case QR : {// QR returns transposed of the inverse !!
      if (rhs_cols > 0 || logdet != NULL || calculate != SOLVE) {
	err = ERRORFAILED;
	continue;
      }

      err = ERRORNOTPROGRAMMEDYET; /// to do: clarify transposed !
      continue;

      CMALLOC(workspaceD, size, double);
      CMALLOC(workspaceU, size, double);

      F77_CALL(dgeqrf)(&size, &size, // QR
		       MPT, &size, // aijmax, &irank, inc, workspaceD, 
		       workspaceU, workspaceD, &size, &err);     
      
      if (err != NOERROR) {	
	CERR1("'dgeqrf' failed with err=%d.", err);
      }
      if (PL >=  PL_DETAILSUSER) { PRINTF("QR successful\n"); }
      break;
    }
      
    case Eigen : { //  M = U D UT
      int max_eigen = sp->max_svd;
      double eigen2zero = sp->eigen2zero;
      if (size > max_eigen) CERR("matrix too large for Cholesky or eigen value decomposition. Increase 'max_chol' and 'max_svd' in 'RFoption' if necessary.");
      
      double 
	optimal_work,
	*pt_work = &optimal_work;
      int k=0,
	optimal_intwork,
	*pt_iwork = &optimal_intwork, 
	lwork = -1,
	lintwork = -1;
      
      CMALLOC(U, sizeSq, double);
      CMALLOC(D, size, double); 
      CMALLOC(xja, 2 * size, int);
      
      for (int i=0; i<=1; i++) {
	double dummy = 0.0,
	  abstol = 0.0;
	int dummy_nr;

	F77_CALL(dsyevr)("V", "A", "U", &size, // Eigen
			 MPT, &size, &dummy, &dummy, &k, &k, 
			 &abstol,// or DLAMCH
			 &dummy_nr, D, U, &size, 
			 xja, // 2 * size * sizeof(integer); nonzeros_idx
			 pt_work, &lwork,
			 pt_iwork, &lintwork,
			 &err
			 );
	if (i==1 || err != NOERROR || ISNAN(D[0])) break;
	lwork = (int) optimal_work;
	lintwork = (int) optimal_intwork;
	CMALLOC(work, lwork, double);
	CMALLOC(iwork, lintwork, int);
	pt_iwork = iwork;
	pt_work = work; 
      }

      
      if (err != NOERROR) {
	if (PL>PL_ERRORS) { PRINTF("Error code F77_CALL(dsyevr) = %d\n", err);}
	CERR1("'dsyevr' failed with err=%d.", err);
	break;
      }
      
      for (int i=0; i<size; i++) if (D[i] < -eigen2zero) {
	  const char *advice[2]={"",
				 " Consider increasing the value of 'eigen2value'."};
	  double min  = D[i];
	  for (int j=i+1; j<size; j++) if (D[j] < min) min = D[j];
	  //print("negative eigen values!!!! %10g\n", D[i]);
	  GERR3("Negative eigenvalues found (less than -eigen2zero=%10e). Smallest one equals %10e. %.50s", -eigen2zero, min, advice[min > -eigen2zero * 100]);
	    
	} //else print("%10g ", D[i]);
      
      if (calculate == MATRIXSQRT) {
	for (int j=0; j<size; j++) {
	  double dummy;
	  dummy = 0.0;
	  if (D[j] >= eigen2zero) dummy = SQRT(D[j]);
	  for (int i=0; i<size; i++, k++) RESULT[k] = U[k] * dummy;
	}
      } else {
	// calculate determinant 
	if (logdet != NULL) {
	  *logdet = cumProd(D, size, sp->det_as_log);
	  if (calculate == DETERMINANT) return NOERROR;
	}
	for (int j=0; j<size; j++) D[j] = D[j] < eigen2zero ? 0.0 : 1.0 / D[j];
	if (rhs_cols > 0) {
	  int tot = size * rhs_cols;
	  CMALLOC(w2, tot, double);	
	  matmulttransposed(U, rhs, w2, size, size, rhs_cols);
	  for (k=0; k<tot; )
	    for (int i=0; i<size; i++) w2[k++] *= D[i];
	  matmult(U, w2, RESULT, size, size, rhs_cols);
	} else {
	  int j;
	  CMALLOC(w2, sizeSq, double);
	  for (k=0, j=0; j<size; j++) {
	    double dummy = D[j];
	    for (int i=0; i<size; i++, k++) w2[k] = U[k] * dummy;
	  }
	  matmult_2ndtransp(w2, U, RESULT, size, size, size); // V * U^T
	}
      }
      if (PL >=  PL_DETAILSUSER) {
	PRINTF("eigen value decomposition successful\n");
      }
      break;
    }
	
    case SVD : {// SVD : M = U D VT
      if (size > sp->max_svd) CERR("matrix too large for SVD decomposition.");
      int k = 0,  
	lwork = -1,
	size8 = size * 8;
      double  optim_lwork,
	eigen2zero = sp->eigen2zero,
	*pt_work = &optim_lwork;

      CMALLOC(VT, sizeSq, double);
      CMALLOC(U, sizeSq, double);
      CMALLOC(D, size, double); 
      CMALLOC(iwork, size8, int);

      for (int i=0; i<=1; i++) {
	F77_CALL(dgesdd)("A", &size, &size, // SVD
			 MPT, &size, D, U, &size, VT, &size, 
			 pt_work, &lwork, iwork, &err);
	if (i==1 || err != NOERROR || ISNAN(D[0])) break;
	lwork = (int) optim_lwork;
	CMALLOC(work, lwork, double);
	pt_work = work;
      }
      if (err != NOERROR) {
	if (PL>PL_ERRORS) {
	  PRINTF("Error code F77_CALL(dgesdd) = %d\n", err);
	}
	CERR1("'dgesdd' failed with err=%d.", err);
	break;
      }

      if (calculate == MATRIXSQRT) {
	double svdtol = sp->svd_tol;
	/* calculate SQRT of covariance matrix */
	for (int j=0; j<size; j++) {
	  double dummy;
	  if (D[j] < -eigen2zero) CERR("negative eigenvalues found.");
	  dummy = 0.0;
	  if (D[j] >= eigen2zero) dummy = SQRT(D[j]);
	  for (int i=0; i<size; i++, k++) RESULT[k] = U[k] * dummy;
	}
 
	/* check SVD */
 	if (svdtol > 0.0) {
	  double *Morig = gesichert ? SICH : M;
	  for (int i=0; i<size; i++) {
	    double *Ui = RESULT + i;
	    for (k=i; k<size; k++) {
	      double *Uk = RESULT + k,
		sum = 0.0;
	      for (int j=0; j<sizeSq; j+=size) {
		sum += Ui[j] * Uk[j];
	      }
	      
	      if (FABS(Morig[i * size + k] - sum) > svdtol) {
		if (PL > PL_ERRORS) {
		  PRINTF("difference %10e at (%d,%d) between the value (%10e) of the covariance matrix and the square of its root (%10e).\n", 
			 Morig[i * size +k] - sum, i, k, Morig[i*size+k], sum);
		}
		FERR3("required precision not attained  (%10e > %10e): probably invalid model. See also '%.50s'.", FABS(Morig[i * size + k] - sum), svdtol,
		      solve[SOLVE_SVD_TOL]);

		err=ERRORM;
		break;
	      } //else printf("ok (%d,%d) %10g %10g\n", i, k, Morig[i*size+k],sum);
	    }
	    if (err != NOERROR) break;		
	  }
	  if (err != NOERROR) break;		
	} // end if svdtol > 0

      } else {
        // calculate determinant 
        if (logdet != NULL) {
	  *logdet = cumProd(D, size, sp->det_as_log);
	  if (calculate == DETERMINANT) return NOERROR;
	}
 	
        for (int j=0; j<size; j++) 
	  D[j] = FABS(D[j]) < eigen2zero ? 0.0 : 1.0 / D[j];

	if (rhs_cols > 0) {
	  int tot = size * rhs_cols;
	  CMALLOC(w2, tot, double);	
	  matmulttransposed(U, rhs, w2, size, size, rhs_cols);
	  for (k=0; k<tot; )
	    for (int i=0; i<size; i++) w2[k++] *= D[i];
	  matmulttransposed(VT, w2, RESULT, size, size, rhs_cols);
	} else {
	  // calculate inverse of covariance matrix
	  int j;	  
	  for (k=0, j=0; j<size; j++) {
	    double dummy = D[j];
	    for (int i=0; i<size; i++) U[k++] *= dummy;
	  }
	  matmult_tt(U, VT, RESULT, size, size, size); // V * U^T
	}
      }

      if (PL >=  PL_DETAILSUSER) { PRINTF("svd successful\n"); }
      break;
    }

    case LU : {// LU
      if (calculate == MATRIXSQRT) {
	err = ERRORFAILED;
	continue;
      }
      
      CMALLOC(ipiv, size, int);		    
      F77_CALL(dgetrf)(&size, &size, // LU
		       MPT, &size, ipiv, &err);
      if (err != NOERROR) {
	CERR1("'dgetrf' (LU) failed with err=%d.", err);
      }
 
      if (logdet != NULL) {
	CERR("logdet cannot be determined for 'LU'.");
	*logdet = Determinant(MPT, size, sp->det_as_log);
        if (calculate == DETERMINANT) return NOERROR;
      }

      if (rhs_cols > 0) {
	int totalRHS = size * rhs_cols;
	if (result != NULL) MEMCOPY(RESULT, rhs, sizeof(double) * totalRHS);
	F77_CALL(dgetrs)("N", &size, // LU rhs
			 &rhs_cols, MPT, &size, ipiv, 
			 RESULT, &size, &err);
	if (err != NOERROR) {	
	  CERR1("'dgetrs' (LU) failed with err=%d.", err);
	}
      } else {
	int lwork = -1;
	double dummy,
	  *p = &dummy;
	for (int i=0; i<=1; i++) { 
	  F77_CALL(dgetri)(&size, MPT, // LU solve
			   &size, ipiv, p, &lwork, &err);	
	  if (err != NOERROR) break;
	  lwork = (int) dummy;
	  CMALLOC(workLU, lwork, double);
	  p = workLU;
	}
      }      
      if (PL >=  PL_DETAILSUSER) { PRINTF("LU decomposition successful\n"); }
      break;
    }
	
    case Sparse : {// sparse matrix
      int nnzlindx, 
	doperm = sp->pivotsparse,
	halfsq = size * (size + 1) / 2,
	nnzcolindices = 0,
	nnzR = 0,
	cache = 512, // to do: CPU cache size
	nnzcfact[3] = { 5, 1, 5 }, 
	nnzRfact[3] = { 5, 1, 2 };
      double
	cholincrease_nnzcol = 1.25,
	cholincrease_nnzR = 1.25;

      if (!posdef) CERR("'spam' needs a positive definite matrix.");
      CMALLOC(pivotsparse, size, int);
      if (!doperm) for (int i=0; i<size; i++) pivotsparse[i] = i + 1;

      if (spam_zaehler == 0) {
	for (int i=0; i<sizeSq; i++) nnzA += FABS(M[i]) >= spam_tol;
	spam_zaehler = nnzA + 1; // falls nur aus Nullen bestehend
      }
      
      CMALLOC(xlnz, sizeP1, int);
      CMALLOC(snode, size, int);
      CMALLOC(xsuper, sizeP1, int);
      CMALLOC(xlindx, sizeP1, int);
      CMALLOC(invp, size, int);
      CMALLOC(w3, size, double);

      CMALLOC(cols, spam_zaehler, int);
      CMALLOC(rows, sizeP1, int);
   
      int nDD = spam_zaehler;
      if (nDD < size) nDD = size;
      CMALLOC(DD, nDD, double);
      // prepare spam

      F77_CALL(spamdnscsr)(&size, &size, M, &size, DD,
			   cols, // ja
			   rows, // ia
			   &spam_tol); // create spam object   
      pt->nsuper = 0;
      // calculate spam_cholesky
      err = 4; // to get into the while loop
      while (err == 4 || err == 5) {
	if (nnzcolindices == 0) {
	  double rel = nnzA / (double) size;
	  if (rel < 5) {
	    nnzcolindices = (int) CEIL(nnzA * (1.05 * rel - 3.8));
	    if (nnzcolindices < 1000) nnzcolindices = 1000;
	  } else {
	    nnzcolindices = nnzA;
	  }
	  nnzcolindices *= nnzcfact[doperm];
	  if (nnzcolindices < nnzA) nnzcolindices = nnzA;
	} else if (err == 5) {
	  int tmp = (int) CEIL(nnzcolindices * cholincrease_nnzcol);
	  if (PL > PL_RECURSIVE) {
	    PRINTF("Increased 'nnzcolindices' with 'NgPeyton' method\n(currently set to %d from %d)", tmp, nnzR);
	  }
	  nnzcolindices = tmp;
	}
	if (nnzcolindices < pt->lindx_n) nnzcolindices = pt->lindx_n;
	
	if (nnzR == 0) {
	  double u = FLOOR(.4 * POW(nnzA, 1.2));
	  u = u < 4 * nnzA ? 4 * nnzA : CEIL(u);
	  nnzR = (int) u * nnzRfact[doperm];
	} else if (err == 4) {
	  int tmp = (int) CEIL(nnzR * cholincrease_nnzR);
	  if (PL > PL_RECURSIVE) {
	    PRINTF("Increased 'nnzR' with 'NgPeyton' method\n(currently set to %d from %d)", tmp, nnzR);
	  }
	  nnzR = tmp;
	}
	if (nnzR < pt->lnz_n) nnzR = pt->lnz_n;
	else if (nnzR > halfsq) nnzR = halfsq;	
	
	CMALLOC(lindx, nnzcolindices, int);	
	CMALLOC(lnz, nnzR, double);
	 	
	F77_CALL(cholstepwise)(&size, &nnzA, DD, cols, rows, &doperm,
			       invp, pivotsparse, 
			       &nnzlindx, &nnzcolindices, 
			       lindx, // 
			       xlindx,// 
			       &(pt->nsuper), // length of lindx
			       &nnzR,  // physical length of lindx
			       lnz,   // output:result
			       xlnz,  // cols of lnz "ja"
			       snode,  // supernode membership ??
			       xsuper, // supernode partioning
			       &cache, // cache size of the CPU
			       &err
			       );       
	
	if (err != NOERROR) {
	  CERR1("'cholstepwise' failed with err=%d.", err);
	  break;
	}	 
      } // while
      
      if (err != NOERROR) CERR("'spam' failed.");
      if (PL >=  PL_DETAILSUSER) { PRINTF("'spam' successful\n"); }
      
      // spam solve
      
      if (calculate == MATRIXSQRT) {
	
	//BUG; // unexpected behaviour in spam
	
	nnzR = xlnz[size] - 1;
	CMALLOC(xja, nnzR, int);
	F77_CALL(calcja)(&size, &(pt->nsuper), pt->xsuper, 
			 pt->lindx, pt->xlindx, pt->xlnz, xja);
	for (int i=0; i<size; invp[i++]--); 
	F77_CALL(spamcsrdns)(&size, pt->lnz, xja, pt->xlnz, RESULT);
	for (int i=0; i<size; i++) {
	  int endfor = (i + 1) * size;
	  for (int j = i * (size + 1) + 1; j<endfor; RESULT[j++]=0.0);
	}
      } else {     
	double *lnz = pt->lnz;
	int RHS_COLS, 	
	  *lindx = pt->lindx;
	
	// spam determinant
	if (logdet != NULL) {
	  if (sp->det_as_log) {
	    double tmp = 0.0;
	    for (int i=0; i<size; i++) tmp += LOG(lnz[xlnz[i] - 1]);	  
	    *logdet = 2.0 * tmp;
	  } else {
	    double tmp = 1.0;
	    for (int i=0; i<size; i++) tmp *= lnz[xlnz[i] - 1] ;
	    *logdet = tmp * tmp;
	  }
	  if (calculate == DETERMINANT) return NOERROR;
	}
	
	/*   z = .Fortran("backsolves", m = nrow,
	     nsuper, p, a@colindices,
	     a@colpointers, as.double(a@entries),
	     a@rowpointers, a@invpivot, a@pivot,
	     a@supernodes, vector("double",nrow),
	     sol = vector("double",nrow*p),
	     as.vector(b,"double"),
	     NAOK = .Spam$NAOK,PACKAGE = "spam")$sol
	*/
	if (rhs_cols <= 0) { // UNBEDINGT VOR double *RHS;
	  RHS_COLS = size;	
	  FILL_IN(RESULT, sizeSq, 0.0);
	  for (int i=0; i<sizeSq; i += sizeP1) RESULT[i] = 1.0; 
	  
	} else {
	  RHS_COLS = rhs_cols;	
	  if (result != NULL) 
	    MEMCOPY(RESULT, rhs, size * rhs_cols * sizeof(double));
	}
	
	//printf("nsuper=%d\n", pt->nsuper);
	//	  for (int ii=0; ii<size; ii++) 
	//printf("%d %d %d %d %10e\n", ii, pt->nsuper, sizeP1, xsuper[ii],
	//	   w3[ii]);
	
	//	  if (false)
	//	  for (int jsub=0; jsub<=pt->nsuper; jsub++) {
	//	    int fj = xsuper[1 - 1],
	//	      Lj = xsuper[jsub + 1 - 1] -1;
	//	    printf("%d %d %d\n", jsub, fj, Lj);
	//	    for (int jcol=fj; jcol <= Lj; jcol++) {
	//	      printf("%d,%10g  ", jcol, w3[jcol -  1]);
	//	    }	    
	//	  }
	
	//	  for (int jcol=1; jcol <= 600; jcol++) {
	//	    w3[jcol - 1] = jcol;
	//   printf("%d,%10g  ", jcol, w3[jcol -  1]);
	//  }	    
	
	
	//	  printf("%ld %ld %d\n", RESULT, rhs, rhs_cols);
	//	  for (int ii=0; ii<size; ii++) printf("%d %10e\n", ii, RESULT[ii]);
	//	  BUG;
	
	F77_CALL(backsolves)(&size, &(pt->nsuper), &RHS_COLS, 
			     lindx, // colindices
			     xlindx, //colpointers
			     lnz, 
			     xlnz, //  rowpointers
			     invp, pivotsparse,
			     xsuper, // supernodes
			     w3, RESULT);	
	if (PL >=  PL_DETAILSUSER) { PRINTF("'spam' successful\n"); }
      }         
      break;
    } // Sparse

    case NoInversionMethod: GERR("no inversion method given.");
    case NoFurtherInversionMethod:
#ifdef DO_PARALLEL
       GERR("All specified matrix inversion methods have failed.");     
#else
      STRCPY(ErrStr, ERRORSTRING);
      GERR1("%.50s (All specified matrix inversion methods have failed.)",
	    ErrStr);
#endif
    case direct_formula: case Diagonal: 
      GERR("strange method appeared: please contact author.");
    default : GERR1("unknown method (%d) in 'RandomFieldsUtils'.", pt->method);
    } // switch
    
    if (err==NOERROR) break;
  } // for m

 
 ErrorHandling:
  if (Pt == NULL) {
    solve_DELETE(&pt);      
  } else {
    Pt->sparse = sparse;
  }
  

  return err; //  -method;
}
  
 
SEXP doPosDef(SEXP M, SEXP rhs, SEXP logdet, int calculate,
	      solve_storage *Pt, solve_param *Sp){
  int 
    rhs_rows, rhs_cols,
    err = NOERROR,
    size = ncols(M), 
    rows = nrows(M);
  bool deleteMM = false,
    deleteRHS = false;
  SEXP res;
  solve_storage Pt0, *pt = Pt;
  if (pt == NULL) {
    solve_NULL(&Pt0);
    pt = &Pt0;
  }
  

  if (rhs == R_NilValue) {
    rhs_rows = rhs_cols = 0;
  } else if (isMatrix(rhs)) {
    rhs_rows = nrows(rhs);
    rhs_cols = ncols(rhs);
  } else if ((rhs_rows = length(rhs)) == 0) {
    rhs_cols = 0;
  } else {
    rhs_cols = 1;
  }
  if (rows != size) ERR("not a square matrix");
  if (rhs_rows > 0 && rhs_rows != size)
    ERR("vector size does not match the matrix size");
  
  int 
    new_cols = rhs_cols == 0 ? size : rhs_cols,
    total = size * new_cols;

  //  res =  PROTECT(isReal(M) ? duplicate(M): coerceVector(M, REALSXP)); UNPROTECT(1); return res;

  if (rhs_cols==0 || isMatrix(rhs)) {
    res = PROTECT(allocMatrix(REALSXP, size, new_cols));
  } else {
    res =  PROTECT(allocVector(REALSXP, total));
  }


  double *MM=NULL, 
    *RHS = NULL;
  if (TYPEOF(M) != REALSXP) {
    if (TYPEOF(M) != INTSXP && TYPEOF(M) != LGLSXP) 
      GERR("numerical matrix expected");
    if ((deleteMM = rhs_cols != 0))
      MM = (double*) MALLOC(total * sizeof(double));
    else MM = REAL(res);
    if (TYPEOF(M) == INTSXP) {
      for (int i=0; i<total; i++) 
	MM[i] = INTEGER(M)[i] == NA_INTEGER ? RF_NA : (double) INTEGER(M)[i];
    } else {
      for (int i=0; i<total; i++) 
	MM[i] = LOGICAL(M)[i] == NA_LOGICAL ? RF_NA : (double) LOGICAL(M)[i];
    } 
  } else MM = REAL(M); 

  if (rhs_cols > 0) {
    if ((deleteRHS = TYPEOF(rhs) != REALSXP)) {
      if (TYPEOF(res) != INTSXP && TYPEOF(rhs) != LGLSXP) 
	GERR("numerical matrix expected");
      int totalRHS = rhs_cols * rhs_rows; 
      RHS = (double*) MALLOC(totalRHS * sizeof(double));
      if (TYPEOF(rhs) == INTSXP) {
	for (int i=0; i<totalRHS; i++) 
	  RHS[i] = INTEGER(rhs)[i] == NA_INTEGER 
	    ? RF_NA : (double) INTEGER(rhs)[i];
      } else if (TYPEOF(rhs) == LGLSXP) {
	for (int i=0; i<totalRHS; i++) 
	  RHS[i] = LOGICAL(rhs)[i] == NA_LOGICAL
	    ? RF_NA : (double) LOGICAL(rhs)[i];
      } 
    } else RHS = REAL(rhs);
  }
  
  err = doPosDef(MM, size, true, rhs_cols == 0 ? NULL : RHS,  rhs_cols, // no PROTECT( needed
		 (rhs_cols == 0 && TYPEOF(M) == REALSXP) ||
		 (rhs_cols > 0 && TYPEOF(rhs) == REALSXP) ? REAL(res) : NULL, 
		 length(logdet) == 0 ? NULL : REAL(logdet),
		 calculate, pt, Sp);

 ErrorHandling:
  if (deleteMM) FREE(MM);
  if (deleteRHS) FREE(RHS);
  if (pt != Pt) solve_DELETE0(pt);
  
  UNPROTECT(1);
  if (err != NOERROR) {
    const char *methname[] = {"solvePosDef", "cholesky", "determinant"};
    errorstring_type msg;
    switch (err) {
    case ERRORMEMORYALLOCATION : STRCPY(msg, "memory allocation error"); break;
    case ERRORNOTPROGRAMMEDYET : STRCPY(msg, "not programmed yet"); break;
    case ERRORFAILED : STRCPY(msg, "algorithm has failed"); break;
    case ERRORM : STRCPY(msg, pt->err_msg);
      break;
    default:  STRCPY(msg, "<unknown error>");
    }
    RFERROR2("'%.50s': %.50s.\n", methname[calculate], msg);
  }

  return res;
}


SEXP SolvePosDef(SEXP M, SEXP rhs, SEXP logdet){
  return doPosDef(M, rhs, logdet, SOLVE, NULL, &(GLOBAL.solve));
}


int solvePosDefResult(double *M, int size, bool posdef, 
		      double *rhs, int rhs_cols, double *result,
		      double *logdet, solve_storage *PT) {
  return doPosDef(M, size, posdef, rhs, rhs_cols, result, logdet, SOLVE,
		  PT, &(GLOBAL.solve));
}


int solvePosDef(double *M, int size, bool posdef, 
		double *rhs, int rhs_cols, double *logdet, 
		solve_storage *PT) {
  return doPosDef(M, size, posdef, rhs, rhs_cols,  NULL,  logdet, SOLVE,
		  PT, &(GLOBAL.solve));
}


int solvePosDefSp(double *M, int size, bool posdef, 
		  double *rhs, int rhs_cols,  double *logdet, 
		  solve_storage *PT,  solve_param * sp) {
  return doPosDef(M, size, posdef, rhs, rhs_cols, NULL, logdet, SOLVE,
		  PT, sp);
}


int XCinvYdet(double *M, int size, bool posdef, double *X, double *Y, int cols, 
	      double *XCinvY, double *det, bool log, solve_storage *PT) {
  int NR = KAHAN ? SCALAR_KAHAN : SCALAR_AVX;
  bool pt = PT != NULL && PT->result != NULL;
  double *result;
  if (pt) result=PT->result;
  else result= (double *) MALLOC(sizeof(double) * size * cols);  
  if (result == NULL) return ERRORMEMORYALLOCATION;
  double *res = result;
  solve_param sp = GLOBAL.solve;
  sp.det_as_log = log;
  int err =  doPosDef(M, size, posdef, Y, cols, result, det, // no PROTECT( needed
		      SOLVE, PT, &sp);
  for (int i=0; i<cols; i++, res += size, X += size)
    XCinvY[i] = SCALAR(res, X, size);
  if (!pt) FREE(result);
  return err;
}

int XCinvXdet(double *M, int size, double *X, int X_cols,
	      double *XCinvX, double *det, bool log, solve_storage *PT) {
  return XCinvYdet(M, size, true, X, X, X_cols, XCinvX, det, log, PT);
}


double XCinvXlogdet(double *M, int size, double *X, int X_cols,
		    solve_storage *PT) {
  int NR = KAHAN ? SCALAR_KAHAN : SCALAR_AVX;
  bool pt = PT != NULL && PT->result != NULL;
  double ans, *result;
  if (pt) result = PT->result;
  else result = (double *) MALLOC(sizeof(double) * size * X_cols);
  if (result == NULL) ERR("memory allocation error in 'xcxlogdet'");  
  double *res = result;
  solve_param sp = GLOBAL.solve;
  sp.det_as_log = true;
  int err =  doPosDef(M, size, true, X, X_cols, result, &ans,// no PROTECT( needed
		      SOLVE, PT, &sp);
  ans *= X_cols; 
  for (int i=0; i<X_cols; i++, res += size, X += size) {
    ans += SCALAR(res, X, size);
  }
  if (!pt) FREE(result);
  if (err != NOERROR)
    ERR("error occurred when calculating determinant of a pos def matrix.");
  return ans;
}


double detPosDef(double *M, int size) {//never log of det --always small sizes!!
  solve_param sp = GLOBAL.solve;
  sp.det_as_log = false;
  double det;
  int err= doPosDef(M, size, true, NULL, 0, NULL, &det, DETERMINANT, // no PROTECT( needed
		    NULL, &sp);
  if (err != NOERROR)
    ERR("error occurred when calculating determinant of a pos def matrix.");
  return det;
}


int invertMatrix(double *M, int size) {
  return doPosDef(M, size, false, NULL, 0, NULL, NULL, SOLVE, NULL, NULL);
}


SEXP Chol(SEXP M) {
  solve_param sp = GLOBAL.solve;
  sp.Methods[0] = sp.Methods[1] = Cholesky;
  sp.sparse = False; // currently does not work, waiting for Reinhard
  // sp.pivot = PIVOT_NONE;
  solve_storage Pt;
  solve_NULL(&Pt);
  SEXP Ans;
  PROTECT(Ans = doPosDef(M, R_NilValue, R_NilValue, MATRIXSQRT, &Pt, &sp));

  if (Pt.actual_pivot == PIVOT_DO || Pt.actual_pivot ==  PIVOT_IDX) {    
    // NEVER: FREE(GLOBAL.solve.pivot_idx); See Pivot_Cholesky:
    SEXP Idx, Info1, Info3;
    PROTECT(Idx = allocVector(INTSXP, Pt.pivot_idx_n));
    MEMCOPY(INTEGER(Idx), Pt.pivot_idx, sizeof(int) * Pt.pivot_idx_n);
    setAttrib(Ans, install("pivot_idx"), Idx);
    
    PROTECT(Info1 = allocVector(INTSXP, 1));
    INTEGER(Info1)[0] = Pt.actual_size;
    setAttrib(Ans, install("pivot_actual_size"), Info1);
  
    PROTECT(Info3 = allocVector(INTSXP, 1));
    INTEGER(Info3)[0] = PIVOT_DO;
    setAttrib(Ans, install("actual_pivot"), Info3);
   
    UNPROTECT(3);
    assert(Pt.pivot_idx_n == ncols(M));
  }
  
  solve_DELETE0(&Pt);
  UNPROTECT(1);
  return Ans;
}


int chol(double *M, int size) {
  solve_param sp = GLOBAL.solve;
  sp.Methods[0] = sp.Methods[1] = Cholesky;
  sp.sparse = False; // currently does not work, waiting for Reinhard
  sp.pivot = PIVOT_NONE;
  return doPosDef(M, size, true, NULL, 0, NULL, NULL, MATRIXSQRT, NULL, &sp);   
}


bool is_positive_definite(double *C, int dim) { // bool not allowed in C
  int err,
    bytes = sizeof(double) * dim * dim;
  double *test;
  test = (double*) MALLOC(bytes);
  MEMCOPY(test, C, bytes);
  err = chol(test, dim);
  UNCONDFREE(test);
  return(err == NOERROR);
}


/*

  ## extrem wichter check -- folgendes funktioniert bislang bei spam nicht:
  library(RandomFields, lib="~/TMP")
  RFoptions(printlevel = 3, pch="", seed=999, use_spam = TRUE) #//
  z = RFsimulate(RMspheric(), x, max_variab=10000, n=10000, spC=F ALSE)
  C = cov(t(z))
  c = RFcovmatrix(RMspheric(), x) #//
  p rint(summary(as.double(c - C))) ##//
  stopifnot(max(a b s(c-C)) < 0.05)

*/


int sqrtPosDefFree(double *M,  // in out
		   int size,  
		   solve_storage *pt,     // in out
		   solve_param *sp
		   ){
  int err, sizeSq = size * size;
  if (sp == NULL) sp = &(GLOBAL.solve);
  InversionMethod *Meth = sp->Methods;
  double *res = NULL;
  bool extra_alloc = Meth[0] == NoInversionMethod ||
    Meth[0] == NoFurtherInversionMethod ||
    (Meth[1] != NoInversionMethod && Meth[1] != NoFurtherInversionMethod &&
     Meth[1] != Meth[0]) ||
    (Meth[0] != Cholesky && Meth[0] != Eigen && Meth[0] != SVD);
  assert(pt != NULL);

  if (sp->sparse == True) 
    warning("package 'spam' is currently not used for simulation");
  sp->sparse = False;

  if (extra_alloc) {
    CMALLOC(result, sizeSq, double);
    res = result;
  } else {
     FREE(pt->result);
    pt->result = M;
    pt->result_n = sizeSq;
  }


  // printf("%ld %d %ld %d %ld %ld\n", M, size, res, MATRIXSQRT, pt, sp);
  
  // it is ok to have
  // ==15170== Syscall param sched_setaffinity(mask) points to unaddressable byte(s)
  // caused by gcc stuff
  err = doPosDef(M, size, true, NULL, 0, res, NULL, MATRIXSQRT, pt, sp);// no PROTECT( needed
  if (extra_alloc) {  
#ifdef WIN32
    pt->to_be_deleted = M;
#else
    FREE(M);
#endif
  }
  return err;
}


void sqrtRHS_Chol(double *U, int size, double* RHS, int RHS_size, int n,
		  double *result,
		  bool pivot, int act_size, int *pi) {
  //  printf("n=%d,rhss=%d si=%d pivot=%d, act=%d U=%d  RHS=%d %d pi=%d\n",
  //	 n,  RHS_size, size,pivot,  act_size, U!=NULL, RHS!=NULL,   result!=NULL, pi!=NULL );
//  for (int i=0; i<size; i++) printf("%d ", pi[i]); printf("\n");

  int
    nsize = n * size,
    NR = KAHAN ? SCALAR_KAHAN : SCALAR_AVX;
  assert(U != NULL);
  if (pivot){
    int n_act_size = n * act_size,
      diff = size - act_size,
      n_diff = nsize - n_act_size;
#ifdef DO_PARALLEL
#pragma omp parallel for num_threads(CORES) if (MULTIMINSIZE(n_act_size)) schedule(dynamic, 8) 
#endif
    for (int i=0 ; i<n_act_size; i++) {
      int ii = i % act_size,
	j = i / act_size;
      result[pi[ii] + j * size] = SCALAR(RHS + j * RHS_size, U + pi[ii] * size,
					 ii + 1);
    }
    pi += act_size;
#ifdef DO_PARALLEL
#pragma omp parallel for num_threads(CORES) if (MULTIMINSIZE(n_diff))
#endif
    for (int i=0; i<n_diff; i++) {
      int ii = pi[i % diff],
	j = i / diff;
      result[ii + j * size] = SCALAR(RHS + j * RHS_size, U + ii*size, act_size);
    }
  } else {
#ifdef DO_PARALLEL
#pragma omp parallel for num_threads(CORES) if (MULTIMINSIZE(nsize)) schedule(dynamic, 8) 
#endif
    for (int i=0; i<nsize; i++) {
      int ii = i % size,
	j = (i / size) * size;
      result[ii + j] = SCALAR(RHS + j, U + ii * size, ii + 1);
    }
  }
}

SEXP tcholRHS(SEXP C, SEXP RHS) {  
  int n_protect = 2;
  SEXP Ans, Idx;
  PROTECT(Idx = getAttrib(C, install("pivot_idx")));
  bool pivot = length(Idx) > 0;
  int 
    n = isMatrix(RHS) ? ncols(RHS) : 1,
    rows = isMatrix(RHS) ? nrows(RHS) : length(RHS),
    size = ncols(C),
    act_size =size;
  if (pivot) {
    SEXP dummy;
    PROTECT(dummy = getAttrib(C, install("pivot_actual_size")));
    act_size=INTEGER(dummy)[0];
    n_protect++;
  }
  int *pi = pivot ?  (int *) INTEGER(Idx) : NULL;
    if (isMatrix(RHS)) PROTECT(Ans = allocMatrix(REALSXP, size, n));
    else PROTECT(Ans = allocVector(REALSXP, size));
  if (rows < act_size) ERR("too few rows of RHS");
  sqrtRHS_Chol(REAL(C), size, REAL(RHS), rows, n, REAL(Ans),
	       pivot, act_size, pi);
  UNPROTECT(n_protect);
  return Ans;
}


SEXP chol2mv(SEXP C, SEXP N) {  
  int n_protect = 2;
  SEXP Ans, Idx;
  PROTECT(Idx= getAttrib(C, install("pivot_idx")));
  bool pivot = length(Idx) > 0;
  int
    n = INTEGER(N)[0],
    size = ncols(C),
    act_size = size;
  if (pivot) {
    SEXP dummy;
    PROTECT(dummy = getAttrib(C, install("pivot_actual_size")));
    act_size = INTEGER(dummy)[0];
    n_protect++;
  }
  int
    n_act_size = n * act_size,
    *pi = pivot ? INTEGER(Idx) : NULL;
  if (n == 1) PROTECT(Ans = allocVector(REALSXP, size));
  else PROTECT(Ans = allocMatrix(REALSXP, size, n));
  double *gauss = (double *) MALLOC(sizeof(double) * n_act_size);
  if (gauss == NULL) ERR("memory allocation error");
  GetRNGstate();
  for (int i=0; i<n_act_size; gauss[i++] = GAUSS_RANDOM(1.0));
  PutRNGstate();
  sqrtRHS_Chol(REAL(C), size, gauss, act_size, n, REAL(Ans),
	       pivot, act_size, pi);
  FREE(gauss);
  UNPROTECT(n_protect);
  return Ans;
}


int sqrtRHS(solve_storage *pt, double* RHS, double *result){
  assert(pt != NULL);
  int 
    size = pt->size;
  switch (pt->method) { 
  case direct_formula : 
  case Cholesky : {
    bool pivot = (pt->actual_pivot == PIVOT_DO ||
		  pt->actual_pivot == PIVOT_IDX) &&
      pt->method != direct_formula;
    if (pivot && pt->pivot_idx_n != size) BUG;
    
    sqrtRHS_Chol(pt->result, size, RHS, size, 1, result, pivot,
		 pivot ? pt->actual_size : NA_INTEGER,
		 pt->pivot_idx);
    return NOERROR;
  }
  case SVD : case Eigen : {  
    double *U = pt->result;
    assert(U != NULL);
#ifdef DO_PARALLEL
#pragma omp parallel for num_threads(CORES) if (MULTIMINSIZE(size))
#endif
    for (int i=0; i<size; i++){
      double dummy = 0.0;
      int k = i;
      for (int j=0; j<size; j++, k+=size) dummy += U[k] * RHS[j];
      result[i] = (double) dummy; 
    }
  }
    break;

  case Sparse : {
    BUG; // SEE ALSO solve, calculate, tmp_delete !!
    int one = 1;
    assert(pt->DD != NULL);
    F77_CALL(amuxmat)(&size, &size, &one, RHS, pt->DD, pt->lnz, 
		      pt->xja, pt->xlnz);
    for (int i=0; i<size; i++) result[i] = pt->DD[pt->invp[i]];
  }
    break;

  case Diagonal : {  
    int  i, j,
      sizeP1 = size + 1;
    double *D = pt->result;
    assert(D != NULL);
    for (i=j=0; j<size; j++, i+=sizeP1) result[j] = RHS[j] * D[i];
  }
    break;
    
  default : 
    BUG;
  }
  
  return NOERROR;
}
```
