## Supplementary material for "MoBPS - Modular Breeding Program Simulator": RandomFieldsUtils R-package: sort.cc

```
/*
 Authors 
 Martin Schlather,

 Copyright (C) 2017 -- 2017 Martin Schlather

This program is free software; you can redistribute it and/or
modify it under the terms of the GNU General Public License
as published by the Free Software Foundation; either version 3
of the License, or (at your option) any later version.

This program is distributed in the hope that it will be useful,
but WITHOUT ANY WARRANTY; without even the implied warranty of
MERCHANTABILITY or FITNESS FOR A PARTICULAR PURPOSE.  See the
GNU General Public License for more details.

You should have received a copy of the GNU General Public License
along with this program; if not, write to the Free Software
Foundation, Inc., 59 Temple Place - Suite 330, Boston, MA  02111-1307, USA.  
*/

#include "RandomFieldsUtils.h"
#include "General_utils.h"
#include "zzz_RandomFieldsUtils.h"


typedef bool (*vergleich)(int, int, void *O);

bool smaller1(int i, int j, void *ORDERD) {
  return ((double *) ORDERD)[i] < ((double *) ORDERD)[j];
}

bool greater1(int i, int j, void *ORDERD) {
  return ((double *) ORDERD)[i] > ((double *) ORDERD)[j];
}

bool smallerInt1(int i, int j, void *ORDERDINT) {
  return ((int *) ORDERDINT)[i] < ((int *)ORDERDINT)[j];
}

bool greaterInt1(int i, int j, void *ORDERDINT) {
  return ((int *)ORDERDINT)[i] > ((int *)ORDERDINT)[j];
}

typedef bool (*vergleichX)(int, int, int, void *O);
vergleichX SMALLERX=NULL, GREATERX=NULL;

bool smaller(int i, int j, int ORDERDIM, void *O)
{
  double *x, *y, *ORDERD = (double*) O;
  int d;
  x = ORDERD + i * ORDERDIM;
  y = ORDERD + j * ORDERDIM;
  for(d=0; d<ORDERDIM; d++)
     if (x[d] != y[d]) return x[d] < y[d];
  return false;
}

bool greater(int i, int j, int ORDERDIM, void *O)
{
  double *x, *y, *ORDERD = (double*) O;
  int d;
  x = ORDERD + i * ORDERDIM;
  y = ORDERD + j * ORDERDIM;
  for(d=0; d<ORDERDIM; d++)
    if (x[d] != y[d]) return x[d] > y[d];
  return false;
}

bool smallerInt(int i, int j, int ORDERDIM, void *O)
{
  int *x, *y, *ORDERDINT = (int*) O;
  int d;
  x = ORDERDINT + i * ORDERDIM;
  y = ORDERDINT + j * ORDERDIM;
  for(d=0; d<ORDERDIM; d++) {
    if (x[d] != y[d]) {
     return x[d] < y[d];
    }
  }
  return false;
}

bool greaterInt(int i, int j, int ORDERDIM, void *O)
{
  int *x, *y, *ORDERDINT = (int*) O;
  int d;
  x = ORDERDINT + i * ORDERDIM;
  y = ORDERDINT + j * ORDERDIM;
  for(d=0; d<ORDERDIM; d++)
    if (x[d] != y[d]) return x[d] > y[d];
  return false;
}


void order(int *pos, int start, int end, vergleich SMALLER, vergleich GREATER,
	  void * ORDERD, int order_from, int order_to) {
  int randpos, pivot, left, right, pivotpos, swap;
  
  if( start < end ) {   
    //Get RNGstate();randpos = start + (int) (UNIFORM_RANDOM * (end-start+1)); PutRNGstate(); // use Get/Put RNGstate with great care !!
    randpos = (int) (0.5 * (start + end));
    pivot = pos[randpos];
    pos[randpos] = pos[start];
    pos[start] = pivot;
    
    pivotpos=start; 
    left = start;
    right=end+1;   
    while (left < right) {      
      //printf("order > %ld start=%d %d left=%d %d %d pivot=%d\n", pos, start, end, left, right, pos[left], pivot);
      while (++left < right && SMALLER(pos[left], pivot, ORDERD)) pivotpos++;
      while (--right > left && GREATER(pos[right], pivot, ORDERD));      
      if (left < right) {
	swap=pos[left]; pos[left]=pos[right]; pos[right]=swap;
	pivotpos++;
      }
    }
    pos[start] = pos[pivotpos];
    pos[pivotpos] = pivot;
    if (start <= order_to && pivotpos > order_from)
      order(pos, start, pivotpos-1, SMALLER, GREATER,
	    ORDERD,  order_from,  order_to);
    if (pivotpos < order_to && end >= order_from)
      order(pos, pivotpos + 1, end, SMALLER, GREATER,
	    ORDERD, order_from, order_to);
  }
}


void Xorder(int *pos, int start, int end, vergleichX SMALLER,vergleichX GREATER,
	    int D, void * ORDERD, int order_from, int order_to ) {
  int randpos, pivot, left, right, pivotpos, swap;
  
  if( start < end ) {   
    //Get RNGstate();randpos = start + (int) (UNIFORM_RANDOM * (end-start+1)); PutRNGstate(); // use Get/Put RNGstate with great care !!
    randpos = (int) (0.5 * (start + end));
    pivot = pos[randpos];
    pos[randpos] = pos[start];
    pos[start] = pivot;
    
    pivotpos=start; 
    left = start;
    right=end+1;   
    while (left < right) {      
      //printf("order > %ld start=%d %d left=%d %d %d pivot=%d\n", pos, start, end, left, right, pos[left], pivot);
      while (++left < right && SMALLER(pos[left], pivot, D, ORDERD)) pivotpos++;
      while (--right > left && GREATER(pos[right], pivot, D, ORDERD));
      if (left < right) {
	swap=pos[left]; pos[left]=pos[right]; pos[right]=swap;
	pivotpos++;
      }
    }
    pos[start] = pos[pivotpos];
    pos[pivotpos] = pivot;
    if (start <= order_to && pivotpos > order_from)
      Xorder(pos, start, pivotpos-1, SMALLER, GREATER,
	     D, ORDERD, order_from, order_to);
    if (pivotpos < order_to && end >= order_from)
      Xorder(pos, pivotpos + 1, end, SMALLER, GREATER,
	     D, ORDERD, order_from, order_to);
  }
}


void orderingFromTo(double *d, int len, int dim, int *pos, int from, int to,
	      usr_bool NAlast) 
{
  int start, end;
  if (NAlast == Nan) {
    for (int i=0; i<len; i++) pos[i]=i;
    end = len-1;
    start = 0;
  } else {
    if (dim != 1) ERR("NAs only allowed for scalars");
    if (NAlast == True) {
       start = 0;
       end = -1;
       int NAend = len;
       for (int i=0; i<len; i++) 
	 if (ISNA(d[i]) || ISNAN(d[i])) pos[--NAend] = i;
	 else pos[++end] = i;
       assert(NAend - 1 == end);
    } else { // if (NAlast == False) {
      start = len;
      end = len -1;
      int NAstart = -1;
      for (int i=0; i<len; i++) 
	if (ISNA(d[i]) || ISNAN(d[i])) pos[++NAstart] = i;
	else pos[--start] = i;
      assert(NAstart + 1 == start);
    }
  }
  if (dim == 1) {
    order(pos, start, end, smaller1, greater1, (void *) d, from - 1, to - 1);
  } else {
    Xorder(pos, start, end, smaller, greater, dim, (void*) d, from - 1, to - 1);
  }
}

void Ordering(double *d, int *len, int *dim, int *pos) {
  orderingFromTo(d, *len, *dim, pos, 1, *len, Nan);
}

 void ordering(double *d, int len, int dim, int *pos) {
  orderingFromTo(d, len, dim, pos, 1, len, Nan);
}


void orderingIntFromTo(int *d, int len, int dim, int *pos, int from, int to, 
		 usr_bool NAlast) {
  /* quicksort algorithm, slightly modified:
     does not sort the data, but d[pos] will be ordered 
     NOTE: pos must have the values 0,1,2,...,start-end !
     (orderdouble is a kind of sorting pos according to
     the variable d)
  */  
    int start, end;
  if (NAlast == Nan) {
    for (int i=0; i<len; i++) pos[i]=i;
    end = len-1;
    start = 0;
  } else {
    if (dim != 1) ERR("NAs only allowed for scalars");
    if (NAlast == True) {
       start = 0;
       end = -1;
       int NAend = len;
       for (int i=0; i<len; i++) 
	 if (d[i] == NA_INTEGER) pos[--NAend] = i;
	 else pos[++end]=i;
       if (NAend - 1 != end) BUG;
    } else { // if (NAlast == False) {
      start = len;
      end = len -1;
      int NAstart = -1;
      for (int i=0; i<len; i++) 
	if (d[i] == NA_INTEGER) pos[++NAstart] = i;
	else pos[--start]=i;
      if (NAstart + 1 != start) BUG;
    }
  }
  if (dim == 1) {
    order(pos, start, end, smallerInt1, greaterInt1, (void *) d, from-1, to-1);
  } else {
    Xorder(pos, start, end, smallerInt, greaterInt, dim, (void*) d, from-1, to-1);
  }  
}
 
void orderingInt(int *d, int len, int dim, int *pos)  {
  orderingIntFromTo(d, len, dim, pos, 1, len, Nan);
}


void quicksort(int start, int end, double *ORDERD, int order_from, int order_to)
{
  //  printf("start %d %d\n", start, end);

  int left, right, pivotpos;
  
  if( start < end ) {   
    //Get RNGstate();randpos = start + (int) (UNIFORM_RANDOM * (end-start+1)); PutRNGstate(); // use Get/Put RNGstate with great care !!
    int randpos = (int) (0.5 * (start + end));
    double pivot = ORDERD[randpos];
    ORDERD[randpos] = ORDERD[start];
    ORDERD[start] = pivot;
      
    pivotpos=start; 
    left = start;
    right = end+1;   
    
    while (left < right) {      
      //printf("order > start=%d %d left=%d %d %10g pivot=%10g\n", start, end, left, right, ORDERD[left], pivot);
      while (++left < right && ORDERD[left] < pivot) pivotpos++;
      while (--right > left && ORDERD[right] > pivot);      
      if (left < right) {
	double swap = ORDERD[left];
	ORDERD[left]=ORDERD[right]; 
	ORDERD[right]=swap;
	pivotpos++;
      }
    }
    ORDERD[start] = ORDERD[pivotpos];
    ORDERD[pivotpos] = pivot;
    if (start <= order_to && pivotpos > order_from)
      quicksort(start, pivotpos-1, ORDERD, order_from, order_to);
    if (pivotpos < order_to && end >= order_from)
      quicksort(pivotpos + 1, end, ORDERD, order_from, order_to);
  }
}


void sortingFromTo(double *d, int len, int from, int to, usr_bool NAlast) {
   int start, end;
  if (NAlast == Nan) {
    end = len-1;
    start = 0;
  } if (NAlast == True) {
    start = end = 0;
    int NAend = len - 1;
    while (end < NAend) {
      while (NAend >= 0 && (ISNA(d[NAend]) || ISNAN(d[NAend]))) NAend--;
      while (end < NAend && !ISNA(d[end]) && !ISNAN(d[end])) end++;
      if (end < NAend) {
	double swap = d[end];
	d[end] = d[NAend]; 
	d[NAend--] = swap;
      }
    }
     assert(NAend == end && false);
  } else { // if (NAlast == False) {
    start = end = len - 1;
    int NAstart = 0;
    while (start > NAstart) {
      while(NAstart < len && (ISNA(d[NAstart]) || ISNAN(d[NAstart]))) NAstart++;
      while (start > NAstart && !ISNA(d[start]) && !ISNAN(d[start])) start--;
      // printf("s = %d\n", start);
      if (start > NAstart) {
	double swap = d[start];
	d[start] = d[NAstart]; 
	d[NAstart++] = swap;
      }
    }   
    //   print("Rstart %d %d %d\n", start, end, NAstart);
   assert(NAstart == start);
  }
  // print("Xstart %d %d\n", start, end);
  quicksort(start, end, d, from - 1, to - 1);
  // for (int i=0; i<len; i++) printf("%10g\n", d[i]); BUG;
}

void sorting(double *d, int len, usr_bool NAlast) {
  sortingFromTo(d, len, 1, len, NAlast);
}

void sortInt(int start, int end, int *ORDERDINT, int order_from, int order_to) {
  //  printf("start %d %d\n", start, end);

  int left, right, pivotpos;
  
  if( start < end ) {   
    //Get RNGstate();randpos = start + (int) (UNIFORM_RANDOM * (end-start+1)); PutRNGstate(); // use Get/Put RNGstate with great care !!
    int randpos = (int) (0.5 * (start + end));
    int pivot = ORDERDINT[randpos];
    ORDERDINT[randpos] = ORDERDINT[start];
    ORDERDINT[start] = pivot;
      
    pivotpos=start; 
    left = start;
    right = end+1;   
    
    while (left < right) {      
      while (++left < right && ORDERDINT[left] < pivot) pivotpos++;
      while (--right > left && ORDERDINT[right] > pivot);      
      if (left < right) {
	int swap = ORDERDINT[left];
	ORDERDINT[left]=ORDERDINT[right]; 
	ORDERDINT[right]=swap;
	pivotpos++;
      }
    }
    ORDERDINT[start] = ORDERDINT[pivotpos];
    ORDERDINT[pivotpos] = pivot;
    if (start <= order_to && pivotpos > order_from)
      sortInt(start, pivotpos-1, ORDERDINT, order_from, order_to);
    if (pivotpos < order_to && end >= order_from)
      sortInt(pivotpos + 1, end, ORDERDINT, order_from, order_to);
  }
}


void sortingIntFromTo(int *d, int len, int from, int to,  usr_bool NAlast) {
  /* quicksort algorithm, slightly modified:
     does not sort the data, but d[pos] will be ordered 
     NOTE: pos must have the values 0,1,2,...,start-end !
     (orderdouble is a kind of sorting pos according to
     the variable d)
  */  

   int start, end;
  if (NAlast == Nan) {
    end = len-1;
    start = 0;
  } if (NAlast == True) {
    start = end = 0;
    int NAend = len - 1;
    while (end < NAend) {
      while (NAend >= 0 && d[NAend] == NA_INTEGER) NAend--;
      while (end < NAend && d[end] != NA_INTEGER) end++;
      if (end < NAend) {
	int swap = d[end];
	d[end] = d[NAend]; 
	d[NAend--] = swap;
      }
    }
     assert(NAend == end && false);
  } else { // if (NAlast == False) {
    start = end = len - 1;
    int NAstart = 0;
    while (start > NAstart) {
      while(NAstart < len && d[NAstart] == NA_INTEGER) NAstart++;
      while (start > NAstart && d[start] != NA_INTEGER) start--;
      if (start > NAstart) {
	double swap = d[start];
	d[start] = d[NAstart]; 
	d[NAstart++] = swap;
      }
    }   
   assert(NAstart == start);
  }
  sortInt(start, end, d, from - 1, to - 1);
}
 
void sortingInt(int *d, int len, usr_bool NAlast) {
  sortingIntFromTo(d, len, 1, len, NAlast);
}


SEXP sortX(SEXP Data, SEXP From, SEXP To, SEXP NAlast) {
  int 
    err = NOERROR,
    len = length(Data),
    from = MAX(1, INTEGER(From)[0]),
    to = MIN(INTEGER(To)[0], len);
  if (from > to) return R_NilValue; 

  usr_bool nalast;
  if (LOGICAL(NAlast)[0] == NA_LOGICAL) nalast = Nan;
  else nalast = LOGICAL(NAlast)[0] ? True : False;
  SEXP Ans;

  if (TYPEOF(Data) == REALSXP) {
    //     printf("%d %d %d %d\n",  from, to, INTEGER(To)[0], len);
    PROTECT(Ans=allocVector(REALSXP, to - from + 1));
    int bytes = len * sizeof(double);
    double *data;
    if ((data = (double*) MALLOC(bytes)) == NULL) { 
      err = ERRORMEMORYALLOCATION; goto ErrorHandling; 
    }
    MEMCOPY(data, REAL(Data), bytes);
    sortingFromTo(data, len, from, to, nalast);
    from--;
    double *ans;
    ans = REAL(Ans);
    for (int i=from; i<to; i++) ans[i - from] = data[i];
    FREE(data);
  } else if  (TYPEOF(Data) == INTSXP) {
    PROTECT(Ans=allocVector(INTSXP, to - from + 1));
    int *data,
      bytes = len * sizeof(int);
    if ((data = (int*) MALLOC(bytes)) == NULL) { 
      err = ERRORMEMORYALLOCATION; goto ErrorHandling; 
    }
    MEMCOPY(data, INTEGER(Data), bytes);
    sortingIntFromTo(data, len, from, to, nalast);
    from--;
    int *ans;
    ans = INTEGER(Ans);
    for (int i=from ; i<to; i++) ans[i - from] = data[i];
    FREE(data);
  } else ERR("Data must be real valued or integer valued.");

  
 ErrorHandling :
  UNPROTECT(1);

  switch(err) {
  case ERRORMEMORYALLOCATION : ERR("not enough memory");
  default:;
  }

  return Ans;
}
 

SEXP orderX(SEXP Data, SEXP From, SEXP To, SEXP NAlast) {
  int 
    err = NOERROR,
    len = length(Data),
    from = MAX(1, INTEGER(From)[0]),
    to = MIN(INTEGER(To)[0], len);
  if (from > to) return R_NilValue; 

  SEXP Ans;
  PROTECT(Ans=allocVector(INTSXP, to - from + 1));

  usr_bool nalast;
  if ( LOGICAL(NAlast)[0] == NA_LOGICAL) nalast = Nan;
  else nalast = LOGICAL(NAlast)[0] ? True : False;
  int
    bytes = len * sizeof(int),
    *pos = (int*) MALLOC(bytes);
  if (pos == NULL) {err = ERRORMEMORYALLOCATION; goto ErrorHandling;}

  if (TYPEOF(Data) == REALSXP) {
    //     printf("%d %d %d %d\n",  from, to, INTEGER(To)[0], len);
    orderingFromTo(REAL(Data), len, 1, pos, from, to, nalast);
  } else if  (TYPEOF(Data) == INTSXP) {
    orderingIntFromTo(INTEGER(Data), len, 1, pos, from, to, nalast);
  } else {
    err = ERRORFAILED;
    goto ErrorHandling;
  }

  from--;
  
  int *ans;
  ans = INTEGER(Ans);
  for (int i=from; i<to; i++) ans[i - from] = pos[i] + 1;

 ErrorHandling :
  FREE(pos);
  UNPROTECT(1);
  
  switch(err) {
  case ERRORFAILED : ERR("Data must be real valued or integer valued.");
  case ERRORMEMORYALLOCATION : ERR("not enough memory");
  default:;
  }

  return Ans;
}
 

/* 
   extendable to higher dim :

  if (from > to) return R_NilValue; 

  int
    *pos = (int*) MALLOC(len * sizeof(int));
  usr_bool nalast = LOGICAL(NAlast)[0] == NA_LOGICAL ? Nan :
    LOGICAL(NAlast)[0] ? True : False;
  SEXP Ans;


  if (TYPEOF(Data) == REALSXP) {
    // printf("%d %d %d %d\n",  from, to, INTEGER(To)[0], len);
    PROTECT(Ans=allocVector(REALSXP, to - from + 1));
    double *ans = REAL(Ans),
      *data = REAL(Data);
    ordering(data, len, dim, pos, from, to, nalast);
    from--;
    for (int i=from; i<to; i++) {
      //printf("%d %d %d %10g     ", i, from, pos[i], data[pos[i]]);
      ans[i - from] = data[pos[i]];
    }
  } else if  (TYPEOF(Data) == INTSXP) {
    PROTECT(Ans=allocVector(INTSXP, to - from + 1));
    int *ans = INTEGER(Ans),
      *data = INTEGER(Data);
    orderingInt(data, len, dim, pos, from, to, nalast);
    from--;
    for (int i=from ; i<to; i++) ans[i - from] = data[pos[i]];
  } else ERR("Data must be real valued or integer valued.");
  UNPROTECT(1);
  f ree(pos);
  return Ans;
}
 
*/
```
