## Supplementary material for "MoBPS - Modular Breeding Program Simulator": RandomFieldsUtils R-package: spamown.f

```
c 
c     Authors:
c     Reinhard Furrer
c       
c  Copyright (C) 2017 -- 2017 Reinhard Furrer
c 
c This program is free software; you can redistribute it and/or
c modify it under the terms of the GNU General Public License
c as published by the Free Software Foundation; either version 3
c of the License, or (at your option) any later version.
c 
c This program is distributed in the hope that it will be useful,
c but WITHOUT ANY WARRANTY; without even the implied warranty of
c MERCHANTABILITY or FITNESS FOR A PARTICULAR PURPOSE.  See the
c GNU General Public License for more details.
c 
c You should have received a copy of the GNU General Public License
c along with this program; if not, write to the Free Software
c Foundation, Inc., 59 Temple Place - Suite 330, Boston, MA  02111-1307, USA.  


      subroutine amuxmat (n,m,p, x, y, a,ja,ia)
      implicit none
      integer n, m, p, ja(*), ia(*)
      double precision  x(m,p), y(n,p), a(*)
c-----------------------------------------------------------------------
c Multiplies a sparse matrix by a full matrix using consecutive dot 
c products, cf. subroutine amux from sparse kit. 
c Matrix A is stored in compressed sparse row storage.
c
c on entry:
c----------
c n     = row dimension of A
c p     = column dimension of x
c x     = array of dimension mxp, m column dimension of A.
c a, ja,
c    ia = input matrix in compressed sparse row format.
c
c on return:
c-----------
c y     = array of dimension nxp, containing the product y=Ax
c
c Reinhard Furrer
c-----------------------------------------------------------------------
c local variables
c
      double precision t
      integer j, i, k
c-----------------------------------------------------------------------
      do j = 1,p
         do i = 1,n
c
c     compute the inner product of row i with vector x
c
            t = 0.0d0
            do  k=ia(i), ia(i+1)-1
               t = t + a(k)*x(ja(k),j)
            enddo
c
            y(i,j) = t
         enddo
      enddo
c
      return
c---------end-of-amuxmat------------------------------------------------
c-----------------------------------------------------------------------
      end
c
 
      subroutine notzero (ja,ia,nrow,ncol,nnz,nz,jao,iao)
c Return the structure of the zero entries in ra,ja,ia, in 
c  compressed sparse row format via rao, jao, iao.
c INPUT:
c     ja, ia -- sparse structure of the matrix A
c     nrow -- number of rows in `a'
c     ncol -- number of columns in `a'
c     nnz -- number of non-zero elements
c     nz -- number of zero elements
c OUTPUT:
c     jao, iao --  sparse structure of the zero entries
c WORK ARRAY:
c     colmn -- logical vector of length ncol

      implicit none
      integer  nrow,ncol,nnz,nz,inz,
     &       ja(nnz),ia(nrow+1),jao(nz),iao(nrow+1)
      logical colmn(ncol)
      integer i,j,k
      inz = 0
      iao(1) = 1
      do i = 1,nrow
         iao(i+1) = iao(i)
         do k = 1,ncol
            colmn(k) = .true.
         enddo
         do j = ia(i),ia(i+1)-1
            colmn(ja(j)) = .false.
         enddo
         do k = 1,ncol
            if(colmn(k)) then
               inz = inz + 1
               jao(inz) = k
               iao(i+1) = iao(i+1) + 1
            endif
         enddo
      enddo
      return
      end


      subroutine setdiagmat (nrow, n, a, ja, ia, diag, iw) 
      implicit none
      integer nrow, n
      double precision a(*),  diag(n) 
      integer ja(*), ia(nrow+1), iw(nrow)
c-----------------------------------------------------------------------
c Sets the diagonal entries of a sparse matrix
c-----------------------------------------------------------------------
c on entry:
c ---------
c nrow	= integer. The row dimension of A
c n = integer. Smallest dimension of A
c
c a, ja, ia   = Matrix A in compressed sparse row format. Sorted.
c diag = diagonal matrix stored as a vector diag(1:n)
c iw   = n vector of zeros.
c
c on return:
c----------
c updated matrix A
c iw    = iw contains the positions of the diagonal entries in the 
c         output matrix. (i.e., a(iw(k)), ja(iw(k)), k=1,...n,
c         are the values/column indices of the diagonal elements 
c         of the output matrix. ). 
c
c Reinhard Furrer
c-----------------------------------------------------------------
      logical insert
      integer i,j, k, k1, k2, icount
     
c
c     get positions of diagonal elements in data structure.
c     
      do  11 i=1,n
         do 21 j= ia(i),ia(i+1)-1
            if (ja(j) .ge. i) then
               if (ja(j) .eq. i) then
                  iw(i) = j
               endif
               goto 11
            endif
 21      continue
 11   continue
c     
c     count number of holes in diagonal and add diag(*) elements to
c     valid diagonal entries.
c     
      icount = 0
      do 31 i=1, n
         if (iw(i) .eq. 0) then
            icount = icount+1
         else
            a(iw(i)) = diag(i) 
         endif
 31      continue
c     
c     if no diagonal elements to insert return
c     
      if (icount .eq. 0) return
c     
c     shift the nonzero elements if needed, to allow for created 
c     diagonal elements. 
c     
c     
c     copy rows backward
c     
      do 5 i=nrow, 1, -1 
c     
c     go through  row ii
c     
         k1 = ia(i)
         k2 = ia(i+1)-1 

         ia(i+1) = ia(i+1)+icount

         if ((i .gt. n) .or. (iw(i) .gt. 0)) then
c     iw(ii) equal to 0, means no diagonal element in a, we need to insert it
c     test is thus true.

c     no fill-in, only copying
            do 4 k = k2,k1,-1 
               ja(k+icount)=ja(k)
               a(k+icount)=a(k)
 4          continue  
            iw(i)=-i
         else
            insert=.TRUE.
            if (k2.lt.k1) then
               ja(k2+icount)=i
               a(k2+icount)=diag(i)
               iw(i)=k2+icount
               icount=icount-1
               insert = .FALSE.
               if (icount .eq. 0) return
            else
               do 6 k = k2,k1,-1
                  if (ja(k).gt. i) then
                     ja(k+icount)=ja(k)
                     a(k+icount)=a(k)
                  else  if  (insert) then
                     ja(k+icount)=i
                     a(k+icount)=diag(i)
                     iw(i)=k+icount
                     icount=icount-1
                     insert = .FALSE.
                     if (icount .eq. 0) return
                  endif
                  if (ja(k).lt. i) then
                     ja(k+icount)=ja(k)
                     a(k+icount)=a(k)
                  endif
 6             continue
c     in case there is only one element, larger than i, we still need to 
c     add the diagonal element
               if  (insert) then
                   ja(k+icount)=i
                   a(k+icount)=diag(i)
                   iw(i)=k+icount
                   icount=icount-1
                   insert = .FALSE.
                   if (icount .eq. 0) return
                endif
            endif
         endif 
 5    continue
      return
c-----------------------------------------------------------------------
c------------end-of-diagaddmat------------------------------------------
      end
      subroutine diagaddmat (nrow, n, a, ja, ia, diag, iw) 
      implicit none
      integer nrow, n
      double precision a(*),  diag(n) 
      integer ja(*), ia(nrow+1), iw(nrow)
c-----------------------------------------------------------------------
c Adds a diagonal matrix to a sparse matrix:  A = Diag + A 
c-----------------------------------------------------------------------
c on entry:
c ---------
c nrow	= integer. The row dimension of A
c n = integer. Smallest dimension of A
c
c a, ja, ia   = Matrix A in compressed sparse row format. Sorted.
c diag = diagonal matrix stored as a vector diag(1:n)
c iw   = n vector of zeros.
c
c on return:
c----------
c updated matrix A
c iw    = iw contains the positions of the diagonal entries in the 
c         output matrix. (i.e., a(iw(k)), ja(iw(k)), k=1,...n,
c         are the values/column indices of the diagonal elements 
c         of the output matrix. ). 
c
c Reinhard Furrer
c-----------------------------------------------------------------
      logical insert
      integer i,j, k, k1, k2, icount
     
c
c     get positions of diagonal elements in data structure.
c     
      do  11 i=1,n
         do 21 j= ia(i),ia(i+1)-1
            if (ja(j) .ge. i) then
               if (ja(j) .eq. i) then
                  iw(i) = j
               endif
               goto 11
            endif
 21      continue
 11   continue
c     
c     count number of holes in diagonal and add diag(*) elements to
c     valid diagonal entries.
c     
      icount = 0
      do 31 i=1, n
         if (iw(i) .eq. 0) then
            icount = icount+1
         else
            a(iw(i)) = a(iw(i)) + diag(i) 
         endif
 31      continue
c     
c     if no diagonal elements to insert return
c     
      if (icount .eq. 0) return
c     
c     shift the nonzero elements if needed, to allow for created 
c     diagonal elements. 
c     
c     
c     copy rows backward
c     
      do 5 i=nrow, 1, -1 
c     
c     go through  row ii
c     
         k1 = ia(i)
         k2 = ia(i+1)-1 

         ia(i+1) = ia(i+1)+icount

         if ((i .gt. n) .or. (iw(i) .gt. 0)) then
c     iw(ii) equal to 0, means no diagonal element in a, we need to insert it
c     test is thus true.

c     no fill-in, only copying
            do 4 k = k2,k1,-1 
               ja(k+icount)=ja(k)
               a(k+icount)=a(k)
 4          continue  
            iw(i)=-i
         else
            insert=.TRUE.
            if (k2.lt.k1) then
               ja(k2+icount)=i
               a(k2+icount)=diag(i)
               iw(i)=k2+icount
               icount=icount-1
               insert = .FALSE.
               if (icount .eq. 0) return
            else
               do 6 k = k2,k1,-1
                  if (ja(k).gt. i) then
                     ja(k+icount)=ja(k)
                     a(k+icount)=a(k)
                  else  if  (insert) then
                     ja(k+icount)=i
                     a(k+icount)=diag(i)
                     iw(i)=k+icount
                     icount=icount-1
                     insert = .FALSE.
                     if (icount .eq. 0) return
                  endif
                  if (ja(k).lt. i) then
                     ja(k+icount)=ja(k)
                     a(k+icount)=a(k)
                  endif
 6             continue
c     in case there is only one element, larger than i, we still need to 
c     add the diagonal element
               if  (insert) then
                   ja(k+icount)=i
                   a(k+icount)=diag(i)
                   iw(i)=k+icount
                   icount=icount-1
                   insert = .FALSE.
                   if (icount .eq. 0) return
                endif
            endif
         endif 
 5    continue
      return
c-----------------------------------------------------------------------
c------------end-of-setdiagmat------------------------------------------
      end
c-----------------------------------------------------------------------
      subroutine diagmua (nrow, a, ia, diag)
      implicit none
      integer          nrow, ia(nrow+1)
      double precision a(*),  diag(nrow), scal
c-----------------------------------------------------------------------
c performs the matrix by matrix product A = Diag * A  (in place) 
c (diamua from sparsekit provides more functionality)
c-----------------------------------------------------------------------
c on entry:
c ---------
c nrow	= integer. The row dimension of A
c a, ia   = Matrix A in compressed sparse row format.
c           (ja is not needed) 
c 
c diag = diagonal matrix stored as a vector diag(1:n)
c
c on return:
c----------
c a, 	= resulting matrix A in compressed sparse row sparse format.
c 
c Notes: 
c-------
c     Reinhard Furrer 2007-06-21
c	    
c-----------------------------------------------------------------
c     local variables
      integer          ii, k, k1, k2

      do 1 ii=1,nrow
c     
c     normalize each row 
c     
         k1 = ia(ii)
         k2 = ia(ii+1)-1
         scal = diag(ii) 
         do 2 k=k1, k2
            a(k) = a(k)*scal
 2       continue
 1    continue
c     
      return
c----------end-of-diagmua------------------------------------------------
c-----------------------------------------------------------------------
      end 
c----------------------------------------------------------------------- 

c-----------------------------------------------------------------------
      subroutine getdiag (a,ja,ia,len,diag)

      implicit none
      double precision diag(*),a(*)
      integer len, ia(*), ja(*)
c-----------------------------------------------------------------------
c This subroutine extracts the main diagonal.
c (getdia from sparsekit provides more functionality)
c----------------------------------------------------------------------- 
c 
c on entry:
c---------- 
c
c len= min(nrow, ncol) = min dimension of the matrix a.
c a,ja,ia = matrix stored in sorted compressed sparse row a,ja,ia,format
c diag  = array of zeros.
c
c on return:
c----------- 
c diag  = array of length containing the wanted diagonal.
c
c Notes: 
c-------
c     Reinhard Furrer 2006-11-02
c----------------------------------------------------------------------c
c     local variables
      integer i, k
c     
c     extract  diagonal elements
c     
      do 1 i=1, len
         do k= ia(i),ia(i+1) -1
            if (ja(k) .ge. i) then
c     we are at or beyond the diagonal. 
               if (ja(k) .eq. i) then
                  diag(i)= a(k)
               endif
               goto 1
            endif
         enddo
 1    continue
      return
c------------end-of-getdiag----------------------------------------------
c-----------------------------------------------------------------------
      end


c Functions that are new or  modified.

      subroutine subsparsefull(nrow,a,ja,ia,b)
c
c     subtracts a sparse matrix from a full one
c     algorithm is in-place, i.e. b is changed
c
c
c Notes: 
c-------
c     Reinhard Furrer 2006-09-21
c-----------------------------------------------------------------------

      implicit none
      integer nrow,ja(*),ia(nrow+1)
      double precision a(*), b(nrow,*)

      integer i,k


      do i=1,nrow
         do k=ia(i),ia(i+1)-1
            b(i,ja(k)) = b(i,ja(k))-a(k)
         enddo
      enddo
      return
      end

      subroutine subfullsparse(nrow,ncol,a,ja,ia,b)
c
c     subtracts a full matrix from a sparse one
c     algorithm is in-place, i.e. b is changed
c
c
c Notes: 
c-------
c     Reinhard Furrer 2006-09-21
c-----------------------------------------------------------------------

      implicit none
      integer nrow,ncol,ja(*),ia(nrow+1)
      double precision a(*), b(nrow,*)

      integer i,j,k


      do i=1,nrow
         do j=1,ncol
            b(i,j) = -b(i,j)
         enddo
         do k=ia(i),ia(i+1)-1
            b(i,ja(k)) = b(i,ja(k))+a(k)
         enddo
      enddo
      return
      end

      subroutine addsparsefull(nrow,a,ja,ia,b)
c
c     adds a sparse matrix to a full one
c     algorithm is in-place, i.e. b is changed
c
c
c Notes: 
c-------
c     Reinhard Furrer 2006-09-21
c-----------------------------------------------------------------------

      implicit none
      integer nrow,ja(*),ia(nrow+1)
      double precision a(*), b(nrow,*)

      integer i,k


      do i=1,nrow
         do k=ia(i),ia(i+1)-1
            b(i,ja(k)) = b(i,ja(k))+a(k)
         enddo
      enddo
      return
      end

      subroutine constructia(nrow,nir,ia,ir)
c     
c     constructs from a regular row index vector a sparse ia vector.
c     note that a regular column index vector corresponds to the 
c     sparse ja vector. for example:
c         A[ir,jc] =>  A@ja = jc, A@ia = constructia(nrow,nir,ia,ir)$ia
c
c     nrow: row dimension of A
c     nir:  length of ir
c     ir:   array of length nir+1!!!
c
c Notes: 
c-------
c     _*Row indices have to be ordered!*_
c
c     Reinhard Furrer 2006-09-13
c-----------------------------------------------------------------------

      implicit none
      integer nrow,nir
      integer ia(nrow+1),ir(*)

      integer i,k

      k=1
      ia(1)=1
      do i=1,nrow
 5       continue
         if (ir(k) .eq. i) then
            k=k+1
            if (k .le. nir) goto 5
         endif
         ia(i+1)=k
      enddo

      ia(nrow+1)=nir+1
      
      return
      end   


 


      subroutine disttospam(nrow,x,a,ja,ia,eps)
      
      implicit none
      integer nrow, ia(nrow+1), ja(*)
      double precision x(*), a(*), eps
c
c     Convertion of an R dist object (removes zero entries as well). 
c
c On entry:
c----------
c     nrow    -- row dimension of the matrix
c     x       -- elements of the dist object (is lower diagonal) 
c                        n*(i-1) - i*(i-1)/2 + j-i  for i < j
c
c     a,ja,ia -- input matrix in CSR format
c
c On return:
c-----------
c     a,ja,ia -- cleaned matrix
c
c Notes: 
c-------
c     Reinhard Furrer 2008-08-13
c-----------------------------------------------------------------------
c
c     Local
      integer i,j,k, tmp

      
      ia(1) = 1 
      k = 1
      do i = 2, nrow
         ia(i) = k
         do j=1 , i-1 
            tmp = nrow*(j-1)-j*(j-1)/2+i-j
            if (.not.(dabs(x(tmp)) .le. eps)) then
               ja(k) = j
               a(k) = x(tmp)
               k = k + 1
            endif
            
         enddo
      enddo

      ia(nrow+1) = k
      return

c---- end of disttospam -------------------------------------------------
c-----------------------------------------------------------------------
      end

      subroutine setdiaold (nrow,ncol,a,ja,ia,c,jc,ic,cmax,diag,eps)

      implicit none

      double precision  a(*),c(*),diag(*),eps
      integer nrow, ncol, ia(*), ja(*), ic(*), jc(*), cmax
c
c     this routine sets the diagonal entries of a matix, provided they
c     are non-zero.
c
c On entry:
c----------
c     nrow,ncol    --  dimensions of the matrix
c     a,ja,ia -- input matrix in CSR format
c     c,jc,ic -- input matrix in CSR format with enough space, see below
c     diag -- diagonal values to set
c     eps  -- what is smaller than zero?
c
c On return:
c-----------
c     c,jc,ic -- matrix with modified diag in CSR format
c
c Notes: 
c-------
c     Reinhard Furrer 2006-10-30
c-----------------------------------------------------------------------
c
c     Local
      double precision b(nrow)
      integer i,k, len, ib(nrow+1), jb(nrow)

c     
      len=0
      ib(1)=1
      do i=1,nrow
         jb(i)=0
      enddo


      do 10 i=1,nrow
         do 15 k= ia(i),ia(i+1) -1
            if (ja(k) .eq. i) then
               a(k)=diag(i)
               c(k)=diag(i)
               ib(i+1)=ib(i)
               goto 10
            endif
            if (ja(k) .gt. i) then
               if (diag(i).gt.eps) then
                  len=len+1
                  jb(len)=i
                  ib(i+1)=ib(i)+1
                  b(len)=diag(i)
               else
                  ib(i+1)=ib(i)
               endif
               goto 10
            endif
 15      continue
 10   continue
      
      if (len .eq. 0) return
c     
c     set nonexisiting elements.
c     

      call subass(nrow,ncol,a,ja,ia,b,jb,ib,c,jc,ic,cmax)
 
      return
c------------end of setdia----------------------------------------------
c-----------------------------------------------------------------------
      end
c-----------------------------------------------------------------------


c
c-----------------------------------------------------------------------
      subroutine subass(nrow,ncol,a,ja,ia,b,jb,ib,c,jc,ic,nzmax)
      implicit none
      integer nrow,ncol,nzmax
      integer ja(*),jb(*),jc(*),ia(*),ib(*),ic(*)
      double precision a(*), b(*), c(*) 

c-----------------------------------------------------------------------
c replaces the elements of A with those of B for matrices in sorted CSR 
c format. we assume that each row is sorted with increasing column 
c indices.
c-----------------------------------------------------------------------
c on entry:
c ---------
c nrow	= integer. The row dimension of A and B
c ncol  = integer. The column dimension of A and B.
c
c a,ja,ia,
c b,jb,ib = Matrices A and B in compressed sparse row format with column
c           entries sorted ascendly in each row   
c
c nzmax	= integer. The max length of the arrays c and jc.
c 
c on return:
c----------
c c,jc,ic = resulting matrix C in compressed sparse row sparse format
c           with entries sorted ascendly in each row. 
c	    
c Notes: 
c-------
c     Reinhard Furrer 2006-09-13, based on sparsekit2 subroutine aplb1
c-----------------------------------------------------------------------

c     local variables
      integer i,j1,j2,ka,kb,kc,kamax,kbmax

      kc = 1
      ic(1) = kc 
c
c     looping over the rows:
      do 6 i=1, nrow
         ka = ia(i)
         kb = ib(i)
         kamax = ia(i+1)-1
         kbmax = ib(i+1)-1 
 5       continue 

c     If we have one or more entries then ka <= kamax
c     If we do not have any entries in both A and B
c     we will not enter the if clause. In which case 
c     we repeatedly copy ic(i+1) <- ic(i).
         if (ka .le. kamax .or. kb .le. kbmax) then 

c     j1 and j2 are left hand pointers of the first entry
c     of A and B. If no entry, they are set to ncol+1
            if (ka .le. kamax) then
               j1 = ja(ka)
            else
               j1 = ncol+1
            endif
            if (kb .le. kbmax) then 
               j2 = jb(kb)         
            else 
               j2 = ncol+1
            endif
c     
c     Three cases:
c       j1=j2: copy element of b in c, incr. all three pointers
c       j1<j2: copy element of a in c, incr. a and c pointers
c       j1>j2: copy element of b in c, incr. b and c pointers
            if (j1 .eq. j2) then 
               c(kc) = b(kb)
               jc(kc) = j1
               ka = ka+1
               kb = kb+1
               kc = kc+1
            else if (j1 .lt. j2) then
               jc(kc) = j1
               c(kc) = a(ka)
               ka = ka+1
               kc = kc+1
            else if (j1 .gt. j2) then
               jc(kc) = j2
               c(kc) = b(kb)
               kb = kb+1
               kc = kc+1
            endif

C     the next four lines should not be required...
            if (kc .gt. nzmax+1) then
c               write (*,*) "exceeding array capacities...",i,nzmax,
c     & ka,kb,kc,j1,j2,kamax,kbmax,ncol,jb(kb)
               return
            endif
            goto 5
         endif
         
         ic(i+1) = kc
 6    continue
      return
c------------end-of-subass---------------------------------------------- 
c-----------------------------------------------------------------------
      end

      subroutine spamcsrdns(nrow,a,ja,ia,dns)

      implicit none
      integer i,k
      integer nrow,ja(*),ia(*)
      double precision dns(nrow,*),a(*)

c-----------------------------------------------------------------------
c Compressed Sparse Row    to    Dense
c-----------------------------------------------------------------------
c
c converts a row-stored sparse matrix into a densely stored one
c
c On entry:
c----------
c
c nrow  = row-dimension of a
c a,
c ja,
c ia    = input matrix in compressed sparse row format.
c         (a=value array, ja=column array, ia=pointer array)
c dns   = array where to store dense matrix
c
c on return:
c-----------
c dns   = the sparse matrix a, ja, ia has been stored in dns(nrow,*)
c
c changes:
c---------
c eliminated the ierr 
c eliminated the filling of zeros: all done with 
c-----------------------------------------------------------------------
      do i=1,nrow
         do  k=ia(i),ia(i+1)-1
            dns(i,ja(k)) = a(k)
         enddo
      enddo
      return
c---- end of csrdns ----------------------------------------------------
c-----------------------------------------------------------------------
      end


c-----------------------------------------------------------------------
      subroutine spamdnscsr(nrow,ncol,dns,ndns,a,ja,ia,eps)

      implicit none
      integer i,j,next
      integer nrow,ncol,ndns,ia(*),ja(*)
      double precision dns(ndns,*),a(*),eps
c-----------------------------------------------------------------------
c Converts a densely stored matrix into a CSR  sparse matrix.
c-----------------------------------------------------------------------
c on entry:
c---------
c
c nrow  = row-dimension of a
c ncol  = column dimension of a
c nzmax = maximum number of nonzero elements allowed. This
c         should be set to be the lengths of the arrays a and ja.
c dns   = input nrow x ncol (dense) matrix.
c ndns  = first dimension of dns.
c
c on return:
c----------
c
c a, ja, ia = value, column, pointer  arrays for output matrix
c
c changes:
c---------
c eliminated the ierr 
c introduced epsilon
c-----------------------------------------------------------------------
      next = 1
      ia(1) = 1
      do  i=1,nrow
         do  j=1, ncol
            if (.not.(dabs(dns(i,j)) .le. eps)) then 
               ja(next) = j
c            write(*,*) next,dns(i,j), eps, i, j
               a(next) = dns(i,j)
               next = next+1
            endif
         enddo
         ia(i+1) = next
      enddo
      return
c---- end of dnscsr ----------------------------------------------------
c-----------------------------------------------------------------------
      end

c----------------------------------------------------------------------- 
      subroutine getmask(nrow,nnz,ir,jc,jao,iao)
c----------------------------------------------------------------------- 
      implicit none
      integer nrow,nnz,ir(*),jc(*),jao(*),iao(*)
      integer k,k0,j,i,iad
c-----------------------------------------------------------------------
c  Gets Compressed Sparse Row indices from Coordinate ones
c----------------------------------------------------------------------- 
c  Loosely based on coocsr from Sparsekit.
c
c on entry:
c--------- 
c nrow	= dimension of the matrix 
c nnz	= number of nonzero elements in matrix
c ir, 
c jc    = matrix in coordinate format. ir(k), jc(k) store the nnz
c         nonzero index. The order of the elements is arbitrary. 
c iao   = vector of 0 of size nrow+1
c
c on return:
c----------- 
c ir 	is destroyed
c
c jao, iao = matrix index in general sparse matrix format with 
c       jao containing the column indices, 
c	and iao being the pointer to the beginning of the row
c
c------------------------------------------------------------------------

c determine row-lengths.
      do 2 k=1, nnz
         iao(ir(k)) = iao(ir(k))+1
 2    continue
c starting position of each row..
      k = 1
      do 3 j=1,nrow+1
         k0 = iao(j)
         iao(j) = k
         k = k+k0
 3    continue
c go through the structure  once more. Fill in output matrix.
      do 4 k=1, nnz
         i = ir(k)
         j = jc(k)
         iad = iao(i)
         jao(iad) = j
         iao(i) = iad+1
 4    continue
c shift back iao
      do 5 j=nrow,1,-1
         iao(j+1) = iao(j)
 5    continue
      iao(1) = 1
      return
c----------------------------------------------------------------------- 
      end


c----------------------------------------------------------------------- 
      subroutine getblock(a,ja,ia, nrw, rw, ncl, cl, bnz, b,jb,ib)
c-----------------------------------------------------------------------
c     purpose:
c     -------- 
c     this function returns the elements a(rw,cl) of a matrix a, 
c     for any index vector rw and cl. the matrix is assumed to be stored 
c     in compressed sparse row (csr) format. 
c
c
c     Reinhard Furrer 2006-09-12
c-----------------------------------------------------------------------
c     parameters:
c     ----------- 
c on entry: 
c---------- 
c     a,ja,ia = the matrix a in compressed sparse row format (input).
c     nrw,rw
c     ncl,cl  = length of and the vector containing the rows and columns
c               to extract
c
c on return:
c----------- 
c     bnz     = nonzero elements of b
c     b,jb,ib = the matrix a(rw,cl) in compressed sparse row format.
c
c note:
c------
c     no error testing is done. It is assumed that b has enough space
c     allocated.
c-----------------------------------------------------------------------
      implicit none

      integer nrw,rw(*),  ncl, cl(*)
      integer bnz, ia(*),ja(*), ib(*),jb(*)
      double precision a(*),b(*)
c
c     local variables.
c
      integer irw, jcl, jja
c
c      write(*,*) cl(1),cl(2)
      bnz = 1
      ib(1) = 1
      do irw = 1,nrw
         do jcl = 1,ncl

            do jja = ia(rw(irw)),ia(rw(irw)+1)-1

               if (cl(jcl) .eq. ja(jja)) then
c     we've found one...
                  b(bnz)  = a(jja)
                  jb(bnz) = jcl
                  bnz = bnz + 1
               endif
            enddo
         enddo
         ib(irw+1) = bnz
c     end irw, we've cycled over all lines 
      enddo 
      bnz = bnz - 1
c      write(*,*) cl(1),cl(2)

      return
c--------end-of-getblock------------------------------------------------
c-----------------------------------------------------------------------
      end 


c----------------------------------------------------------------------- 
      subroutine getlines(a,ja,ia, nrw, rw, bnz, b,jb,ib)
c-----------------------------------------------------------------------
c     purpose:
c     -------- 
c     this function returns the lines rw of a matrix a. 
c     the matrix is assumed to be stored 
c     in compressed sparse row (csr) format. 
c
c
c     Reinhard Furrer 2012-04-04
c-----------------------------------------------------------------------
c     parameters:
c     ----------- 
c on entry: 
c---------- 
c     a,ja,ia = the matrix a in compressed sparse row format (input).
c     nrw,rw  = length of and the vector containing the rows and columns
c               to extract
c
c on return:
c----------- 
c     bnz     = nonzero elements of b
c     b,jb,ib = the matrix a(rw,cl) in compressed sparse row format.
c
c note:
c------
c     no error testing is done. It is assumed that b has enough space
c     allocated.
c-----------------------------------------------------------------------
      implicit none

      integer nrw,rw(*)
      integer bnz, ia(*),ja(*), ib(*),jb(*)
      double precision a(*),b(*)
c
c     local variables.
c
      integer irw, jja
c
      bnz = 1
      ib(1) = 1
      do irw = 1,nrw
         do jja = ia(rw(irw)),ia(rw(irw)+1)-1

            b(bnz)  = a(jja)
            jb(bnz) = ja(jja)
            bnz = bnz + 1
         enddo
         ib(irw+1) = bnz
c     end irw, we've cycled over all lines 
      enddo 
      bnz = bnz - 1

      return
c--------end-of-getlines------------------------------------------------
c-----------------------------------------------------------------------
      end 


c----------------------------------------------------------------------- 
      subroutine getelem(i,j,a,ja,ia,iadd,elem) 
c-----------------------------------------------------------------------
c     purpose:
c     -------- 
c     this function returns the element a(i,j) of a matrix a, 
c     for any pair (i,j).  the matrix is assumed to be stored 
c     in compressed sparse row (csr) format. getelem performs a
c     binary search. 
c     also returns (in iadd) the address of the element a(i,j) in 
c     arrays a and ja when the search is successsful (zero if not).
c-----------------------------------------------------------------------
c     parameters:
c     ----------- 
c on entry: 
c---------- 
c     i      = the row index of the element sought (input).
c     j      = the column index of the element sought (input).
c     a      = the matrix a in compressed sparse row format (input).
c     ja     = the array of column indices (input).
c     ia     = the array of pointers to the rows' data (input).
c on return:
c----------- 
c     elem = value of a(i,j). 
c     iadd   = address of element a(i,j) in arrays a, ja if found,
c              zero if not found. (output) 
c
c     note: the inputs i and j are not checked for validity. 
c-----------------------------------------------------------------------
c     noel m. nachtigal october 28, 1990 -- youcef saad jan 20, 1991.
c
c     Reinhard Furrer: converted to subroutine and eliminated sorted
c         many manipulations... last for 0.31; Sept 13
c----------------------------------------------------------------------- 
      implicit none

      integer i, ia(*), iadd, j, ja(*)
      double precision a(*),elem
c
c     local variables.
c
      integer ibeg, iend, imid
c
c     initialization 
c
      iadd = 0 
      ibeg = ia(i)
      iend = ia(i+1)-1

c     empty line! test at beginning
 10   if (iend .lt. ibeg) return

c     
c     begin binary search: 
c     test of bounds
      if (ja(ibeg).gt.j) return
      if (ja(iend).lt.j) return

      if (ja(ibeg).eq.j) then
         iadd = ibeg 
         goto 20
      endif
      if (ja(iend).eq.j) then
         iadd = iend 
         goto 20
      endif
     
c     compute the middle index and test if found
      imid = ( ibeg + iend ) / 2
      if (ja(imid).eq.j) then
         iadd = imid 
         goto 20
      endif

c     update the interval bounds. 
      if (ja(imid).gt.j) then
         iend = imid -1
      else 
         ibeg = imid +1
      endif

      goto 10  
c     
c     set iadd and elem before returning
 20   elem = a(iadd) 
      return
c--------end-of-getelem-------------------------------------------------
c-----------------------------------------------------------------------
      end 

      subroutine getallelem(nir,ir,jr,a,ja,ia,alliadd,allelem)
c-----------------------------------------------------------------------
c     purpose:
c     -------- 
c     wrapper to getelem to retrieve several elements.
c----------------------------------------------------------------------- 
c     Reinhard Furrer 2006-09-12
c----------------------------------------------------------------------- 
      implicit none
      
      integer nir,ir(nir),jr(nir),ja(*),ia(*),alliadd(nir)
      double precision a(*),allelem(nir)
c     local vars
      integer i
      do i = 1,nir
         call getelem(ir(i),jr(i),a,ja,ia,alliadd(i),allelem(i))
      enddo
      return
c--------end-of-allgetelem----------------------------------------------
c-----------------------------------------------------------------------
      end


c-----------------------------------------------------------------------
c-----------------------------------------------------------------------
c-
c- Modified by P. T. Ng from sparsekit
c-----------------------------------------------------------------------
c-----------------------------------------------------------------------
      subroutine aemub (nrow,ncol,a,ja,ia,amask,jmask,imask,
     *                  c,jc,ic,iw,aw,nzmax,ierr)
c---------------------------------------------------------------------
      real(8) a(*),c(*),amask(*),aw(ncol)
      integer ia(nrow+1),ja(*),jc(*),ic(nrow+1),jmask(*),imask(nrow+1)
      logical iw(ncol)
c-----------------------------------------------------------------------
c Modified from amask by Pin T. Ng on 2/27/03 to perform 
c element-wise multiplication
c-----------------------------------------------------------------------
c On entry:
c---------
c nrow  = integer. row dimension of input matrix
c ncol  = integer. Column dimension of input matrix.
c
c a,
c ja,
c ia    = the A matrix in Compressed Sparse Row format
c
c amask,
c jmask,
c imask = matrix defining mask stored in compressed
c         sparse row format. (This is the B matrix)
c
c nzmax = length of arrays c and jc. see ierr.
c
c On return:
c-----------
c
c a, ja, ia and amask, jmask, imask are unchanged.
c
c c
c jc,
c ic    = the output matrix in Compressed Sparse Row format.
c
c ierr  = integer. serving as error message.c
c         ierr = 1  means normal return
c         ierr .gt. 1 means that amask stopped when processing
c         row number ierr, because there was not enough space in
c         c, jc according to the value of nzmax.
c
c work arrays:
c-------------
c iw    = logical work array of length ncol.
c aw    = real work array of length ncol.
c
c note:
c------ the  algorithm is in place: c, jc, ic can be the same as
c a, ja, ia in which cas the code will overwrite the matrix c
c on a, ja, ia
c
c-----------------------------------------------------------------------
      ierr = 0
      len = 0
      do 1 j=1, ncol
         iw(j) = .false.
         aw(j) = 0.0
 1    continue
c     unpack the mask for row ii in iw
      do 100 ii=1, nrow
c     save pointer and value in order to be able to do things in place
         do 2 k=imask(ii), imask(ii+1)-1
            iw(jmask(k)) = .true.
            aw(jmask(k)) = amask(k)
 2       continue
c     add umasked elemnts of row ii
         k1 = ia(ii)
         k2 = ia(ii+1)-1
         ic(ii) = len+1
         do 200 k=k1,k2
            j = ja(k)
            if (iw(j)) then
               len = len+1
               if (len .gt. nzmax) then
                  ierr = ii
                  return
               endif
               jc(len) = j
               c(len) = a(k)*aw(j)
            endif
 200     continue
c
         do 3 k=imask(ii), imask(ii+1)-1
            iw(jmask(k)) = .false.
            aw(jmask(k)) = 0.0
 3       continue
 100  continue
      ic(nrow+1)=len+1
c
      return
c-----end-of-aemub -----------------------------------------------------
c-----------------------------------------------------------------------
      end
c-----------------------------------------------------------------------
      subroutine aemub1 (nrow,ncol,a,ja,ia,b,jb,ib,c,jc,ic,
     *     nzmax,ierr)
      real(8) a(*), b(*), c(*)
      integer ja(*),jb(*),jc(*),ia(nrow+1),ib(nrow+1),ic(nrow+1)
c-----------------------------------------------------------------------
c A modification of aplsb by Pin Ng on 6/12/02 to
c perform the element-wise operation C = A*B for matrices in 
c sorted CSR format.
c the difference with aplsb is that the resulting matrix is such that
c the elements of each row are sorted with increasing column indices in
c each row, provided the original matrices are sorted in the same way. 
c-----------------------------------------------------------------------
c on entry:
c ---------
c nrow	= integer. The row dimension of A and B
c ncol  = integer. The column dimension of A and B.
c
c a,
c ja,
c ia   = Matrix A in compressed sparse row format with entries sorted
c
c b, 
c jb, 
c ib	=  Matrix B in compressed sparse row format with entries sorted
c        ascendly in each row   
c
c nzmax	= integer. The  length of the arrays c and jc.
c         amub will stop if the result matrix C  has a number 
c         of elements that exceeds exceeds nzmax. See ierr.
c 
c on return:
c----------
c c, 
c jc, 
c ic	= resulting matrix C in compressed sparse row sparse format
c         with entries sorted ascendly in each row. 
c	    
c ierr	= integer. serving as error message. 
c         ierr = 0 means normal return,
c         ierr .gt. 0 means that amub stopped while computing the
c         i-th row  of C with i=ierr, because the number 
c         of elements in C exceeds nzmax.
c
c Notes: 
c-------
c     this will not work if any of the two input matrices is not sorted
c-----------------------------------------------------------------------
      ierr = 0
      kc = 1
      ic(1) = kc 
c
c     the following loop does a merge of two sparse rows and 
c     multiplies  them.
c 
      do 6 i=1, nrow
         ka = ia(i)
         kb = ib(i)
         kamax = ia(i+1)-1
         kbmax = ib(i+1)-1 
 5       continue 
c
c     this is a while  -- do loop -- 
c 
         if (ka .le. kamax .or. kb .le. kbmax) then 
c     
            if (ka .le. kamax) then
               j1 = ja(ka)
            else
c     take j1 large enough  that always j2 .lt. j1
               j1 = ncol+1
            endif
            if (kb .le. kbmax) then 
               j2 = jb(kb)         
            else 
c     similarly take j2 large enough  that always j1 .lt. j2 
               j2 = ncol+1
            endif
c     
c     three cases
c     
            if (j1 .eq. j2) then 
               c(kc) = a(ka)*b(kb)
               jc(kc) = j1
               ka = ka+1
               kb = kb+1
               kc = kc+1
            else if (j1 .lt. j2) then
               ka = ka+1
            else if (j1 .gt. j2) then
               kb = kb+1
            endif
            if (kc .gt. nzmax) goto 999
            goto 5
c
c     end while loop
c
         endif
         ic(i+1) = kc
 6    continue
      return
 999  ierr = i 
      return
c------------end-of-aemub1 --------------------------------------------- 
c-----------------------------------------------------------------------
      end
c-----------------------------------------------------------------------
      subroutine aedib (nrow,ncol,job,a,ja,ia,b,jb,ib,
     *     c,jc,ic,nzmax,iw,aw,ierr)
      real(8) a(*), b(*), c(*), aw(ncol) 
      integer ja(*),jb(*),jc(*),ia(nrow+1),ib(nrow+1),ic(nrow+1),
     *     iw(ncol)
c-----------------------------------------------------------------------
c performs the element-wise matrix division  C = A/B. 
c Modified from aplsb by Pin Ng on 2/27/03
c-----------------------------------------------------------------------
c on entry:
c ---------
c nrow	= integer. The row dimension of A and B
c ncol  = integer. The column dimension of A and B.
c job   = integer. Job indicator. When job = 0, only the structure
c                  (i.e. the arrays jc, ic) is computed and the
c                  real values are ignored.
c
c a,
c ja,
c ia   = Matrix A in compressed sparse row format.
c 
c b, 
c jb, 
c ib	=  Matrix B in compressed sparse row format.
c
c nzmax	= integer. The  length of the arrays c and jc.
c         amub will stop if the result matrix C  has a number 
c         of elements that exceeds exceeds nzmax. See ierr.
c 
c on return:
c----------
c c, 
c jc, 
c ic	= resulting matrix C in compressed sparse row sparse format.
c	    
c ierr	= integer. serving as error message. 
c         ierr = 0 means normal return,
c         ierr .gt. 0 means that amub stopped while computing the
c         i-th row  of C with i=ierr, because the number 
c         of elements in C exceeds nzmax.
c
c work arrays:
c------------
c iw	= integer work array of length equal to the number of
c         columns in A.
c aw	= real work array of length equal to the number of
c         columns in A.
c
c-----------------------------------------------------------------------
      logical values
      values = (job .ne. 0) 
      ierr = 0
      len = 0
      ic(1) = 1 
      do 1 j=1, ncol
         iw(j) = 0
 1    continue
c     
      do 500 ii=1, nrow
c     row i 
         do 200 ka=ia(ii), ia(ii+1)-1 
            len = len+1
            jcol    = ja(ka)
            if (len .gt. nzmax) goto 999
            jc(len) = jcol 
            if (values) c(len)  = a(ka)/0.0 
            iw(jcol)= len
            aw(jcol) = a(ka)
 200     continue
c     
         do 300 kb=ib(ii),ib(ii+1)-1
            jcol = jb(kb)
            jpos = iw(jcol)
            if (jpos .eq. 0) then
               len = len+1
               if (len .gt. nzmax) goto 999
               jc(len) = jcol
               if (values) c(len)  = 0.0
               iw(jcol)= len
            else
               if (values) c(jpos) = aw(jcol)/b(kb)
            endif
 300     continue
         do 301 k=ic(ii), len
            iw(jc(k)) = 0
 301     continue
         ic(ii+1) = len+1
 500  continue
      return
 999  ierr = ii
      return
c------------end of aedib ----------------------------------------------- 
c-----------------------------------------------------------------------
      end
c-----------------------------------------------------------------------
      subroutine aeexpb (nrow,ncol,job,a,ja,ia,b,jb,ib,
     *     c,jc,ic,nzmax,iw,aw,ierr)
      real(8) a(*), b(*), c(*), aw(ncol) 
      integer ja(*),jb(*),jc(*),ia(nrow+1),ib(nrow+1),ic(nrow+1),
     *     iw(ncol)
c-----------------------------------------------------------------------
c performs the element-wise matrix division  C = A/B. 
c Modified from aplsb by Pin Ng on 2/27/03
c-----------------------------------------------------------------------
c on entry:
c ---------
c nrow	= integer. The row dimension of A and B
c ncol  = integer. The column dimension of A and B.
c job   = integer. Job indicator. When job = 0, only the structure
c                  (i.e. the arrays jc, ic) is computed and the
c                  real values are ignored.
c
c a,
c ja,
c ia   = Matrix A in compressed sparse row format.
c 
c b, 
c jb, 
c ib	=  Matrix B in compressed sparse row format.
c
c nzmax	= integer. The  length of the arrays c and jc.
c         amub will stop if the result matrix C  has a number 
c         of elements that exceeds exceeds nzmax. See ierr.
c 
c on return:
c----------
c c, 
c jc, 
c ic	= resulting matrix C in compressed sparse row sparse format.
c	    
c ierr	= integer. serving as error message. 
c         ierr = 0 means normal return,
c         ierr .gt. 0 means that amub stopped while computing the
c         i-th row  of C with i=ierr, because the number 
c         of elements in C exceeds nzmax.
c
c work arrays:
c------------
c iw	= integer work array of length equal to the number of
c         columns in A.
c aw	= real work array of length equal to the number of
c         columns in A.
c
c-----------------------------------------------------------------------
      logical values
      values = (job .ne. 0) 
      ierr = 0
      len = 0
      ic(1) = 1 
      do 1 j=1, ncol
         iw(j) = 0
 1    continue
c     
      do 500 ii=1, nrow
c     row i 
         do 200 ka=ia(ii), ia(ii+1)-1 
            len = len+1
            jcol    = ja(ka)
            if (len .gt. nzmax) goto 999
            jc(len) = jcol 
            if (values) c(len)  = 1.0
            iw(jcol)= len
            aw(jcol) = a(ka)
 200     continue
c     
         do 300 kb=ib(ii),ib(ii+1)-1
            jcol = jb(kb)
            jpos = iw(jcol)
            if (jpos .eq. 0) then
               len = len+1
               if (len .gt. nzmax) goto 999
               jc(len) = jcol
               if (values) c(len)  = 0.0**b(kb)
               iw(jcol)= len
            else
               if (values) c(jpos) = aw(jcol)**b(kb)
            endif
 300     continue
         do 301 k=ic(ii), len
            iw(jc(k)) = 0
 301     continue
         ic(ii+1) = len+1
 500  continue
      return
 999  ierr = ii
      return
c------------end of aeexpb ----------------------------------------------- 
c-----------------------------------------------------------------------
      end


      SUBROUTINE CALCJA(nrow,nsuper,
     %     xsuper,lindx,xlindx,xlnz,
     %     cholcja)
c     small function to calculate ja for the cholesky factor
c     as they use a condensed format. GRATULIERU LIT!


c     INPUT:
c     nrow (integer)           number of rows
c     nsuper (integer)         number of supernodes
c     xsuper (integer)         supernode partition
c     xlindx,lindx  (integer)  row indices for each supernode
c     xlnz (integer)           ia for cholesky factor
c
c     OUTPUT:
c     cholcja (integer)        ja for cholesky factor

      IMPLICIT NONE

      INTEGER nrow,nsuper
      INTEGER xsuper(nrow),lindx(*),xlindx(nrow+1),xlnz(nrow+1)
      INTEGER cholcja(*)

      INTEGER k, i, j, m, n

      k=1
      m=1
      DO i=1,nsuper
         DO j=1,( xsuper(i+1)-xsuper(i))
            DO n=1,(xlnz(k+1)-xlnz(k))
               cholcja(m)=lindx( xlindx(i)+j-2  + n)
               m=m+1
            ENDDO
            k=k+1
         ENDDO
      ENDDO

      RETURN
      END


      subroutine transpose(n,m,a,ja,ia,ao,jao,iao)

      implicit none
      integer n,m,ia(n+1),iao(m+1),ja(*),jao(*)
      double precision  a(*),ao(*)


      integer i,j,k,next
c-----------------------------------------------------------------------
c     Transposition
c     similar to csrcsc from sparsekit
c----------------------------------------------------------------------- 
c on entry:
c----------
c n	= number of rows of CSR matrix.
c m    = number of columns of CSC matrix.
c a	= real array of length nnz (nnz=number of nonzero elements in input 
c         matrix) containing the nonzero elements.
c ja	= integer array of length nnz containing the column positions
c 	  of the corresponding elements in a.
c ia	= integer of size n+1. ia(k) contains the position in a, ja of
c	  the beginning of the k-th row.
c
c on return:
c ---------- 
c ao	= real array of size nzz containing the "a" part of the transpose
c jao	= integer array of size nnz containing the column indices.
c iao	= integer array of size n+1 containing the "ia" index array of
c	  the transpose. 
c
c----------------------------------------------------------------------- 
c----------------- compute lengths of rows of transp(A) ----------------
      do  i=1, n
         do  k=ia(i), ia(i+1)-1 
            j = ja(k)+1
            iao(j) = iao(j)+1
         enddo
      enddo
c---------- compute pointers from lengths ------------------------------
      iao(1) = 1
      do  i=1,m
         iao(i+1) = iao(i) + iao(i+1)
      enddo
c--------------- now do the actual copying ----------------------------- 
      do  i=1,n
         do  k=ia(i),ia(i+1)-1 
            j = ja(k) 
            next = iao(j)
            ao(next) = a(k)
            jao(next) = i
            iao(j) = next+1
         enddo
      enddo
c-------------------------- reshift iao and leave ---------------------- 
      do  i=m,1,-1
         iao(i+1) = iao(i)
      enddo
      iao(1) = 1
c-----------------------------------------------------------------------
      end
c-----------------------------------------------------------------------
 
c-----------------------------------------------------------------------


      subroutine reducedim(a,ja,ia,eps,bnrow,bncol,k,b,jb,ib)

      implicit none
      double precision a(*),b(*),eps
      integer bnrow, bncol,k
      integer ia(*),ja(*),ib(*),jb(*)

      integer   i, j, jaj

c-----------------------------------------------------------------------
c  Reduces the dimension of A to (,bnrow,bncol) by copying it to B.
c  (Hence not in place - for R purposes).
c  Only elements smaller than eps are copied.
c-----------------------------------------------------------------------
c on entry:
c---------
c
c------------------------------------------------------------------------

      k=1
      do i=1,bnrow
         ib(i)=k
         do j=ia(i), ia(i+1)-1
            jaj=ja(j)
            if (jaj .le.bncol) then
               if (abs( a(j)) .gt. eps) then
                  b(k)=a(j)
                  jb(k)=jaj
                  k=k+1
               endif
            endif
         enddo
      enddo
      ib(bnrow+1)=k
      return
c-----------------------------------------------------------------------
      end

c-----------------------------------------------------------------------
c Currently not used...
      subroutine reducediminplace(eps,nrow,ncol,k,a,ja,ia)

      implicit none
      double precision a(*),eps
      integer nrow, ncol,k
      integer ia(*),ja(*)

      integer   i, j, jj, itmp

c-----------------------------------------------------------------------
c  Reduces the dimension of A to (nrow,ncol) _in place_
c  Only elements smaller than eps are copied.
c-----------------------------------------------------------------------
c  Reinhard Furrer, June 2008
c------------------------------------------------------------------------

      k=1
      do i=1,nrow
         itmp = ia(i)
         ia(i)=k
         do j=itmp, ia(i+1)-1
            jj=ja(j)
            if (jj .le. ncol) then
               if (abs( a(jj)) .gt. eps) then
                  a(k)=a(jj)
                  ja(k)=jj
                  k=k+1
               endif
            endif
         enddo
      enddo
      ia(nrow+1)=k
      return
c-----------------------------------------------------------------------
      end
c-----------------------------------------------------------------------


c----------------------------------------------------------------------c
c      T R I A N G U L A R    S Y S T E M    S O L U T I O N S         c
c                                                                      c
c      spamforward and spamback                                        c
c----------------------------------------------------------------------c

      subroutine spamforward (n,p,x,b,l,jl,il)
      implicit none

      integer n, p, jl(*),il(n+1)
      double precision  x(n,p), b(n,p), l(*)

      integer i, k, j
      double precision  t

c-----------------------------------------------------------------------
c   solves    L x = y ; L = lower triang. /  CSR format
c                        sequential forward elimination 
c-----------------------------------------------------------------------
c
c On entry:
c----------
c n,p      = integer. dimensions of problem.
c b      = real array containg the right side.
c
c l, jl, il,    = Lower triangular matrix stored in CSR format.
c
c On return:
c-----------
c       x  = The solution of  L x  = b.
c--------------------------------------------------------------------
c     Reinhard Furrer June 2008, April 2012
      
c     if first diagonal element is zero, break
      if (l(1) .eq. 0.0 ) goto 5

c     cycle over all columns of b
      do i=1,p 
         
c     first row has one element then cycle over all rows
         x(1,i) = b(1,i) / l(1)
         do 3 k = 2, n
            t = b(k,i)
            do 1 j = il(k), il(k+1)-1
               if (jl(j) .lt. k) then
                  t = t-l(j)*x(jl(j),i)
               else
                  if (jl(j) .eq. k) then
                     if (l(j) .eq. 0.0) goto 5 
c     diagonal element is not zero, hence we divide and leave the loop
                     x(k,i) = t / l(j)
                     goto 3
                  endif 
               endif
 1          continue
 3       continue      
      enddo

      return
 5    n = -k
      return
      end
c-----------------------------------------------------------------------
      subroutine spamback (n,p,x,b,r,jr,ir)
      implicit none

      integer n, p, jr(*),ir(n+1)
      double precision  x(n,p), b(n,p), r(*)

      integer l, k, j
      double precision  t
c-----------------------------------------------------------------------
c             Solves   R x = b    R = upper triangular.
c-----------------------------------------------------------------------
c
c On entry:
c----------
c n,p      = integers. dimension of problem.
c b        = real array containg the right side.
c
c r, jr, ir,    = Upper triangular matrix stored in CSR format.
c
c On return:
c-----------
c       x = The solution of  R x = b .
c--------------------------------------------------------------------
c     Reinhard Furrer June 2008, April 2012
      
      k = 0
      if (r(ir(n+1)-1) .eq. 0.0 ) goto 5
      do l=1,p
         x(n,l) = b(n,l) / r(ir(n+1)-1)
         do 3 k = n-1,1,-1
            t = b(k,l)
            do 1  j = ir(k+1)-1,ir(k),-1
               if (jr(j) .gt. k) then
                  t = t - r(j)*x(jr(j),l)
               else
                  if (jr(j) .eq. k) then
                     if (r(j) .eq. 0.0) goto 5 
c     diagonal element is not zero, hence we divide and leave the loop
                     x(k,l) = t / r(j)
                     goto 3
                  endif 
               endif
 1          continue
 3       continue      
      enddo
      return
 5    n = -k
      return
      end
c-----------------------------------------------------------------------
```
