## Supplementary material for "MoBPS - Modular Breeding Program Simulator": RandomFieldsUtils R-package: utils.cc

```
/* 
 Authors
 Martin Schlather,

 Collection of system specific auxiliary functions

 Copyright (C) 2001 -- 2017 Martin Schlather, 

This program is free software; you can redistribute it and/or
modify it under the terms of the GNU General Public License
as published by the Free Software Foundation; either version 3
of the License, or (at your option) any later version.

This program is distributed in the hope that it will be useful,
but WITHOUT ANY WARRANTY; without even the implied warranty of
MERCHANTABILITY or FITNESS FOR A PARTICULAR PURSE.  See the
GNU General Public License for more details.

You should have received a copy of the GNU General Public License
along with this program; if not, write to the Free Software
Foundation, Inc., 59 Temple Place - Suite 330, Boston, MA  02111-1307, USA.
*/

//#include <Rmath.h>
//#include <unistd.h>
#include "RandomFieldsUtils.h"
//#include "win_linux_aux.h"
#include "General_utils.h"
#include "intrinsics.h"
//#include "Solve.h"
#include "Utils.h"
#include "own.h"


SEXP getChar() {  ERR("does not work");
#ifdef WIN32
  ERR("input limitations on windows");
#endif
#define maxGetChar 255
  //typedef char intchar[sizeof(int) / sizeof(char)];
  //typedef char intchar[sizeof(int) / sizeof(char)];
  SEXP str;
  int //g,
    i = -1;
  char // c, *t = NULL,
    *s = NULL
    ;
  s = (char*) MALLOC(sizeof(char) * maxGetChar);
  // initscr();
  //  fflush(stdin);
  //  nocbreak();
  while (++i < maxGetChar) {
    // g = getchar();       
    //s[i] = ((intchar*) &g)[0][0];
    // g = scanf("%c\n", s); s[1] = '\0'; break;
    //t = fgets(s, 2, stdin); break;
    //    s[i] = getch();
    //fflush(stdin);
    if (false) {
      s[i+1] = '\0';
      // printf("%d i=%d  '%c' '%c' '%c' '%c' '%c'\n", g, i,  s[i],
      // ((intchar*) &g)[0][0],
      //  ((intchar*) &g)[0][1],
      // ((intchar*) &g)[0][2],
      // ((intchar*) &g)[0][3]
      // );
    }
    if (s[i] == '\n') {
      s[i] = '\0';
      break;
    }
  }
  //endwin();
//printf(">%.50s<\n", s);
  PROTECT(str=allocVector(STRSXP, 1));
  SET_STRING_ELT(str, 0, mkChar(s));  
  UNPROTECT(1);
  FREE(s);
  return str;
}


SEXP DivByRow(SEXP M, SEXP V) {
  int
    l = length(V),
    r = nrows(M),
    c = ncols(M);

  double *m = REAL(M),
    *v = REAL(V);
  
  if (l != c) ERR("vector does not match matrix");
  for (int j=0; j<c; j++) {
    double vj = v[j];
    for (int i=0; i<r; i++) {
      *(m++) /= vj;
    }
  }

  return M;
}

#define algn_general(X)  ((1L + (uintptr_t) (((uintptr_t) X - 1L) / BytesPerBlock)) * BytesPerBlock)

double static inline *algn(double *X) {assert(algn_general(X)>=(uintptr_t)X); return (double *) algn_general(X); }

int static inline *algnInt(int *X) {assert(algn_general(X)>=(uintptr_t)X); return (int *) algn_general(X); }


void colMaxsIint(int *M, int r, int c, int *ans) {
  if (r < 32) {
    for (int i=0; i<c; i++) {
      int *m = M + r * i,
	dummy = m[0];    
      for (int j=1; j<r; j++) dummy = MAX(dummy, m[j]);
      ans[i] = dummy;
    }
    return;
  }
#ifdef DO_PARALLEL
#pragma omp parallel for num_threads(CORES)
#endif  
  for (int i=0; i<c; i++) {
     int dummy,
      *m = M + r * i;
#if defined SSE4 || defined AVX2
     int *start = algnInt(m),
       *end = m + r;
    uintptr_t End = (uintptr_t) (end - integers);
    if ((uintptr_t) start < End) {
      BlockType *m0 = (BlockType*) start,
	Dummy = LOAD((BlockType*) m0);
      for (m0++ ; (uintptr_t) m0 < End; m0++) {
	Dummy = MAXINTEGER(Dummy, LOAD(m0));
      }
      int *d = (int *) &Dummy;
      dummy = d[0];
      dummy = MAX(dummy, d[1]);
      dummy = MAX(dummy, d[2]);
      dummy = MAX(dummy, d[3]);
#if defined AVX2
      dummy = MAX(dummy, d[4]);
      dummy = MAX(dummy, d[5]);
      dummy = MAX(dummy, d[6]);
      dummy = MAX(dummy, d[7]);
#endif // AVX
      for ( ; m<start; m++) dummy = MAX(dummy, *m);
      m = (int *) m0;
      for ( ; m<end; m++) dummy = MAX(dummy, *m);
    } else {
      dummy = m[0];    
      for (int j=1; j<r; j++) dummy = MAX(dummy, m[j]);
    }
#else // not SSE4
    dummy = m[0];    
    for (int j=1; j<r; j++) dummy = MAX(dummy, m[j]);
#endif    
    ans[i] = dummy;
  }
}


void colMaxsI(double *M, int r, int c, double *ans) {
  if (r < 16) {
    for (int i=0; i<c; i++) {
      double *m = M + r * i,
	dummy = m[0];    
      for (int j=1; j<r; j++) dummy = MAX(dummy, m[j]);
      ans[i] = dummy;
    }
    return;
  }
#ifdef DO_PARALLEL
#pragma omp parallel for num_threads(CORES)
#endif  
  for (int i=0; i<c; i++) {
    double dummy,
      *m = M + r * i;
#if defined SSE2
    double *start = algn(m),
      *end = m + r;
    uintptr_t End = (uintptr_t) (end - doubles);
    if ((uintptr_t) start < End) {
      Double * m0 = (Double*) start,
	Dummy = (Double) LOAD((UBlockType*) m0);
      for (m0++ ; (uintptr_t) m0 < End; m0++) {
	Dummy = MAXDOUBLE(Dummy, (Double) LOAD((UBlockType*) m0));
      }
      double *d = (double *) &Dummy;
      dummy = d[0];
      dummy = MAX(dummy, d[1]);
#if defined AVX
      dummy = MAX(dummy, d[2]);
      dummy = MAX(dummy, d[3]);
#endif
      for ( ; m<start; m++) dummy = MAX(dummy, *m);
      m = (double *) m0;
      for ( ; m<end; m++) dummy = MAX(dummy, *m);
    } else {
      dummy = m[0];    
      for (int j=1; j<r; j++) dummy = MAX(dummy, m[j]);
    }
#else
    dummy = m[0];    
    for (int j=1; j<r; j++) dummy = MAX(dummy, m[j]);
#endif    
    ans[i] = dummy;
  }
}


SEXP colMaxs(SEXP M) {
  int
    r = nrows(M),
    c = ncols(M);
  if (r == 0) return R_NilValue;
  SEXP Ans;
  if (TYPEOF(M) == REALSXP) {
    PROTECT(Ans = allocVector(REALSXP, c));
    colMaxsI(REAL(M), r, c, REAL(Ans));
  } else {
    bool i = TYPEOF(M) == INTSXP;
    PROTECT(Ans = allocVector(i ? INTSXP : LGLSXP, c));
    int *m, *a;
    if (i) {
      m = INTEGER(M);
      a = INTEGER(Ans);
    } else {
      m = LOGICAL(M);
      a = LOGICAL(Ans);
    }
    colMaxsIint(m, r, c, a);
  }
  UNPROTECT(1);
  return Ans;
}


SEXP rowProd(SEXP M) {
  int
    r = nrows(M),
    r4 = r / 4,
    c = ncols(M);
  if (r == 0) return R_NilValue;
  SEXP Ans;
  if (TYPEOF(M) == REALSXP) {
    PROTECT(Ans = allocVector(REALSXP, r));
    double *ans = REAL(Ans),
      *m = REAL(M);
    MEMCOPY(ans, m, sizeof(double) * r);
    m += r;
    for (int ic=1; ic<c; ic++) {
      double *a = ans;
      for (int ir=0; ir<r4; ir++) {
	*(a++) *= *(m++);
	*(a++) *= *(m++);
	*(a++) *= *(m++);
	*(a++) *= *(m++);
      }
      for (int ir=r4 * 4; ir<r; ir++) *(a++) *= *(m++);
    }
  } else {
    // printf("type = %d", TYPEOF(M));
    RFERROR("transform to double first") ;
  }
  UNPROTECT(1);
  return Ans;
}

SEXP rowMeansX(SEXP M, SEXP Weight) {
  // todo : SSE2 / AVX
  int
    r = nrows(M),
    c = ncols(M);
  if (r == 0 || c == 0) return R_NilValue;
  if (length(Weight) != c && length(Weight) != 0)
    ERR("Length of 'weight' must equal number of columns of 'x'.");
  SEXP Ans;
  PROTECT(Ans = allocVector(REALSXP, r));
  double *ans = REAL(Ans);
  for (int j=0; j<r; j++) ans[j] = 0.0;
  if (length(Weight) == 0) {    
#define for1					\
    for (int i=0; i<c; i++, m+=r) {			\
      for (int j=0; j<r; j++) ans[j] += (double) m[j];	\
    }
  
    if (TYPEOF(M) == REALSXP) { double *m = REAL(M); for1; }
    else {
      int *m;
      if (TYPEOF(M) == INTSXP) m = INTEGER(M); else m = LOGICAL(M);
      for1;
    }
    
  } else {    
    double *weight = ToReal(Weight);
#define for2							\
    for (int i=0; i<c; i++, m+=r) {				\
      double dummy = weight[i]; /* load1(weight); MULTDOUBLE */ \
      for (int j=0; j<r; j++) ans[j] += (double) m[j] * dummy;	\
    }

    if (TYPEOF(M) == REALSXP) { double *m = REAL(M); for2; }
    else {
      int *m;
      if (TYPEOF(M) == INTSXP) m = INTEGER(M); else m = LOGICAL(M);
      for2;
    }
    
    if (TYPEOF(Weight) != REALSXP) FREE(weight);
  }
  double invc = 1.0 / (double) c;
  for (int j=0; j<r; j++) ans[j] *= invc;
  UNPROTECT(1);
  return Ans;
}

SEXP dbinorm(SEXP X, SEXP Sigma) { // 12'41
  int nrow,
    ncol = 2;
  double *x, *y;
  if (TYPEOF(X) == VECSXP) {
    if (length(X) != ncol) BUG;
    SEXP xx = VECTOR_ELT(X, 0);
    nrow = length(xx);
    x = REAL(xx);
    y = REAL(VECTOR_ELT(X, 1));
  } else {
    if (isMatrix(X)) {
      if (ncols(X) != ncol) BUG;
      nrow = nrows(X);
    } else if (isVector(X)) {
      if (length(X) != ncol) BUG;
      nrow = 1;
    } else BUG;
    x = REAL(X);
    y = x + nrow;
  }

  
  SEXP Ans;
  PROTECT(Ans = allocVector(REALSXP, nrow));
  double *ans = REAL(Ans);
  //  int nrow4 = nrow - 4;
  if (length(Sigma) == 0) {
    double invtwopi = 1.0 / TWOPI;
    /*
      minushalfX[4] ={-0.5, -0.5, -0.5, -0.5},
      invtwopiX [4] = {invtwopi, invtwopi, invtwopi, invtwopi};
    int i=0;

#define atonce 4
    __m256d minushalf4 = LOADuDOUBLE(minushalfX),
       invtwopi4 = LOADuDOUBLE(invtwopiX);
      
    for (; i<nrow4; i+=atonce) {
      __m256d x4 = LOADuDOUBLE(x + i);
      double *xx4 = (double *) &x4;
      x4 = MULTDOUBLE(x4, x4);
      {
	__m256d y4 = LOADuDOUBLE(y + i);
	y4 = MULTDOUBLE(y4, y4);
	x4 = ADDDOUBLE(x4, y4);
      }
      x4 = MULTDOUBLE(minushalf4, x4);
      xx4[0] = EXP(xx4[0]);
      xx4[1] = EXP(xx4[1]);
      xx4[2] = EXP(xx4[2]);
      xx4[3] = EXP(xx4[3]);
      x4 = MULTDOUBLE(x4, invtwopi4);
      STOREuDOUBLE(ans + i, x4);
    }
    */
    for (int i=0; i<nrow; i++) 
      ans[i] = EXP(-0.5 * (x[i] * x[i] + y[i] * y[i])) * invtwopi;
    } else {
    double *sigma=REAL(Sigma),
      sigma1 = sigma[0],
      sigma4 = sigma[3],
      inv2piSrtS = 1.0 / (TWOPI * SQRT(sigma1 * sigma4)),
      invS1half = 0.5 / sigma1,
      invS4half = 0.5 / sigma4;
    if (sigma[1] == 0.0 && sigma[2] == 0.0) {
      for (int i=0 ; i<nrow; i++)
	ans[i] = EXP(- (x[i] * x[i] * invS1half + y[i] * y[i] * invS4half) )
	  * inv2piSrtS;
    } else BUG;
  }
  UNPROTECT(1);
  return Ans;
}

double *ToRealDummy = NULL;
int  ToRealN = 0;
double *ToRealI(SEXP X, bool *create) {
  if (TYPEOF(X) == REALSXP) { 
    *create = false;
    return REAL(X);
  }
  HELPINFO("Better use 'double' as storage mode (for one of the arguments).");
  int len = length(X); 
  double *y;
  if (create || ToRealN < len) {
    y = (double *) MALLOC(sizeof(double) * len);
    if (y == NULL) ERR1("not enough memory for an %d vector of doubles", len);
    if (!create) {
      FREE(ToRealDummy);
      ToRealDummy = y;
      ToRealN = len;
    }
  } else y = ToRealDummy;
  int *x;
  if (TYPEOF(X)==INTSXP) x=INTEGER(X); else x=LOGICAL(X);
  for (int i=0; i<len; i++) y[i] = (double) x[i];
  return y;
}

double *ToReal(SEXP X) {
  bool ToFalse[1] = { false };
 if (TYPEOF(X) == REALSXP) return REAL(X);
  return ToRealI(X, ToFalse);
}


int *ToIntDummy = NULL;
int  ToIntN = 0;
int *ToIntI(SEXP X, bool *create, bool round) {
  if (TYPEOF(X) == INTSXP) {
    *create = false;
    return INTEGER(X);
  }
  if (TYPEOF(X) == LGLSXP) {
    *create = false;
    return LOGICAL(X);
  }
  int len = length(X);
  if (len > 100 || PL > 1)
    HELPINFO("Better use 'integer' as storage mode (for one of the arguments).");
  int *y;
  if (*create || ToIntN < len) {
    y = (int *) MALLOC(sizeof(int) * len);    
    if (y == NULL) ERR1("not enough memory for an %d vector of integers", len);
    if (!*create) {
      FREE(ToIntDummy);
      ToIntDummy = y;
      ToIntN = len;
    }
  } else y = ToIntDummy;
  double *x = (double *) REAL(X);
  if (round) for (int i=0; i<len; i++) y[i] = (int) ROUND(x[i]);
  else for (int i=0; i<len; i++) y[i] = (int) x[i];
  return y;
}

int *ToInt(SEXP X) {
  bool ToFalse[1] = { false };
  return ToIntI(X, ToFalse, false);
}

void freeGlobals() {
  FREE(ToRealDummy);
  FREE(ToIntDummy);
}


SEXP quadratic(SEXP x, SEXP A) {
  SEXP ans;
  int len = length(x);
  if (len != nrows(A) || len != ncols(A)) ERR("'x' and 'A' do not match.");
  PROTECT(ans = allocVector(REALSXP, 1));
  xAx(REAL(x), REAL(A), len, REAL(ans));
  UNPROTECT(1);
  return ans;
}

SEXP dotXV(SEXP M, SEXP V) {
  int
    r = nrows(M),
    c = ncols(M),
    l = length(V)
    ;
  if (l != r) ERR("X and v do not match");
  if (r == 0) return R_NilValue;
  SEXP Ans;
  PROTECT(Ans = allocMatrix(REALSXP, r, c));

  // bringt nix
  //#ifdef DO_PARALLEL
  //#pragma omp parallel for num_threads(CORES)
  //#endif  
  for (int i=0; i<c; i++) {
    //  printf("i=%d\n", i);
#if defined SSE2_DONOTUSE_AS_SLOWER
    double 
      *ans = REAL(Ans) + r * i,
      *v = REAL(V),
      *m = REAL(M) + r * i,
      *end = m + r - doubles;
    for ( ; m < end; m += doubles, ans += doubles, v += doubles)
      STOREuDOUBLE(ans, MULTDOUBLE(LOADuDOUBLE(m), LOADuDOUBLE(v)));
    end += doubles;
    for (; m < end; m++) *ans = *m * *v;
#else
    double
      *ans = REAL(Ans) + r * i,
      *v = REAL(V),
      *m = REAL(M) + r * i;
    for (int j=0; j<r; j++) {
      ans[j] = m[j] * v[j];
    }
     
#endif    
  }

  UNPROTECT(1);
  return Ans;
}

 
```
