## Supplementary material for "MoBPS - Modular Breeding Program Simulator": RandomFieldsUtils R-package: win_linux_aux.cc

```
/* 
 Authors
 Martin Schlather,

 Collection of system specific auxiliary functions

 Copyright (C) 2001 -- 2017 Martin Schlather, 

This program is free software; you can redistribute it and/or
modify it under the terms of the GNU General Public License
as published by the Free Software Foundation; either version 3
of the License, or (at your option) any later version.

This program is distributed in the hope that it will be useful,
but WITHOUT ANY WARRANTY; without even the implied warranty of
MERCHANTABILITY or FITNESS FOR A PARTICULAR PURSE.  See the
GNU General Public License for more details.

You should have received a copy of the GNU General Public License
along with this program; if not, write to the Free Software
Foundation, Inc., 59 Temple Place - Suite 330, Boston, MA  02111-1307, USA.
*/

#ifdef WIN32
// #define WIN32_LEAN_AND_MEAN
#define VC_EXTRALEAN
#include <windows.h>
#endif


// achtung! windows.h zusammen mit <Rmath.h oder R.graphics>
// gibt warnung, da ERROR mehrfach definiert !
// deshalb auch in auxiliary.h nicht basic.h einbinden // obsolette ?!!
#include <unistd.h>
#include <Rinternals.h>
#include "win_linux_aux.h"
/*

#include <unistd.h>
 
#include <R_ext/Lapack.h>
#include <R_ext/Linpack.h>
#include "RF.h"
#include <Rdefines.h>
*/


 
void sleepMilli(int *milli) {
#ifdef WIN32
  Sleep((long) *milli); // along
#else 
  usleep((useconds_t) (1000 * (unsigned long) *milli));// along
#endif
}

void sleepMicro(int *micro) {
#ifdef WIN32
  Sleep((long) ((*micro + 500) / 1000));// along
#else
  usleep((useconds_t) *micro);
#endif
}

void pid(int *i)  {
#ifdef WIN32 
  *i = _getpid();
#else
  *i = getpid(); 
#endif
}

void hostname(char **h, int *i){
#ifdef WIN32
  *h[0]=0;
#else
  gethostname(*h, *i);
#endif
}  
```
