## Supplementary material for "MoBPS - Modular Breeding Program Simulator": RandomFieldsUtils R-package: zzz.c

```
/*
 Authors 
 Martin Schlather,

 Copyright (C) 2015 -- 2017 Martin Schlather

This program is free software; you can redistribute it and/or
modify it under the terms of the GNU General Public License
as published by the Free Software Foundation; either version 3
of the License, or (at your option) any later version.

This program is distributed in the hope that it will be useful,
but WITHOUT ANY WARRANTY; without even the implied warranty of
MERCHANTABILITY or FITNESS FOR A PARTICULAR PURPOSE.  See the
GNU General Public License for more details.

You should have received a copy of the GNU General Public License
along with this program; if not, write to the Free Software
Foundation, Inc., 59 Temple Place - Suite 330, Boston, MA  02111-1307, USA.  
*/

//#include "Basic_utils.h" // must be before anything else

#include "RandomFieldsUtils.h"
#include "zzz_RandomFieldsUtils.h"
#include "Utils.h"

 
static R_NativePrimitiveArgType 
    int_arg[] = { INTSXP },
    host_arg[] = { STRSXP, INTSXP};
  //  static R_NativeArgStyle argin[] = {R_ARG_IN},
  //    argout[] = {R_ARG_OUT},
  //   hostarg[] = {R_ARG_OUT, R_ARG_OUT};

#define CDEF(name, n, type) {#name, (DL_FUNC) & name, n, type}
static const R_CMethodDef cMethods[]  = {
  CDEF(sleepMilli,  1, int_arg),
  CDEF(sleepMicro, 1, int_arg),
  CDEF(pid, 1, int_arg),
  CDEF(hostname, 2, host_arg),
  // {"attachRFoptionsUtils", (DL_FUNC) &attachRFoptionsUtils, 0, NULL, NULL},
  // {"detachRFoptionsUtils", (DL_FUNC) &detachRFoptionsUtils, 0, NULL, NULL},
  {NULL, NULL, 0, NULL}
};


#define CALLDEF_DO(name, n) {#name, (DL_FUNC) &name, n}
static R_CallMethodDef callMethods[]  = {
  // in die respectiven C-Dateien muss RandomFieldsUtils.h eingebunden sein
  CALLDEF_DO(Chol, 1),
  CALLDEF_DO(SolvePosDef, 3),
  CALLDEF_DO(struve, 4),
  CALLDEF_DO(I0ML0, 1),
  CALLDEF_DO(gaussr, 2),
  CALLDEF_DO(WMr, 4),
  CALLDEF_DO(logWMr, 4),
  CALLDEF_DO(loadRandomFieldsUtils, 0),
  CALLDEF_DO(attachRandomFieldsUtils, 0),
  CALLDEF_DO(detachRandomFieldsUtils, 0),
  CALLDEF_DO(sortX, 4),
  CALLDEF_DO(orderX, 4), 
  CALLDEF_DO(getChar, 0),
  CALLDEF_DO(DivByRow, 2),
  CALLDEF_DO(colMaxs, 1),
  CALLDEF_DO(quadratic, 2),
  CALLDEF_DO(dotXV, 2),
  CALLDEF_DO(rowMeansX, 2),
  CALLDEF_DO(rowProd, 1),
  CALLDEF_DO(dbinorm, 2),
  CALLDEF_DO(chol2mv, 2),
  CALLDEF_DO(tcholRHS, 2),
  //  CALLDEF_DO(),
  {NULL, NULL, 0}
};


 
#define EXTDEF_DO(name, n)  {#name, (DL_FUNC) &name, n}
static const R_ExternalMethodDef extMethods[] = {
  // in die respectiven C-Dateien muss RandomFieldsUtils.h eingebunden sein
  EXTDEF_DO(RFoptions, -1), 
  {NULL, NULL, 0} 
};


#define CALLABLE(FCTN)  R_RegisterCCallable("RandomFieldsUtils", #FCTN, (DL_FUNC)  FCTN)
void R_init_RandomFieldsUtils(DllInfo  *dll) {
  CALLABLE(solve_DELETE);
  CALLABLE(solve_NULL);
  CALLABLE(solvePosDef);
  CALLABLE(invertMatrix);
  
  CALLABLE(solvePosDefSp);
  CALLABLE(sqrtPosDefFree); 
  CALLABLE(sqrtRHS);
  
  CALLABLE(detPosDef);
  CALLABLE(XCinvXdet);
  CALLABLE(XCinvYdet);
  CALLABLE(is_positive_definite);
  CALLABLE(chol2inv);
  CALLABLE(chol);

  CALLABLE(StruveH);
  CALLABLE(StruveL);
  CALLABLE(I0mL0);

  CALLABLE(WM);
  CALLABLE(DWM);
  CALLABLE(DDWM);
  CALLABLE(D3WM);
  CALLABLE(D4WM);
  CALLABLE(logWM);
  
  CALLABLE(Gauss);
  CALLABLE(DGauss);
  CALLABLE(DDGauss);
  CALLABLE(D3Gauss);
  CALLABLE(D4Gauss);
  CALLABLE(logGauss);
  
  CALLABLE(getErrorString);
  CALLABLE(setErrorLoc);
  CALLABLE(getUtilsParam);
  CALLABLE(attachRFoptions);
  CALLABLE(detachRFoptions);

  CALLABLE(ordering);
  CALLABLE(orderingInt);
  CALLABLE(sorting);
  CALLABLE(sortingInt);
  CALLABLE(scalarX);
  //  CALLABLE(scalarInt);

  CALLABLE(ToIntI);
  //  CALLABLE(ToRealI);

  CALLABLE(pid);
  CALLABLE(sleepMicro);
 

  R_registerRoutines(dll, cMethods, callMethods, NULL, // .Fortran
		     extMethods);
  R_useDynamicSymbols(dll, FALSE); //
}


void R_unload_RandomFieldsUtils(DllInfo *info) {
  // just to avoid warning from compiler on my computer
#ifdef SCHLATHERS_MACHINE  
  if (0) Rprintf("%ld\n", (unsigned long) info);
#endif  
  /* Release resources. */
}

 
```
